## Supplementary figures and images for "Large, three-generation CEPH families reveal post-zygotic mosaicism and variability in germline mutation accumulation"

### 1.pdf

# 1

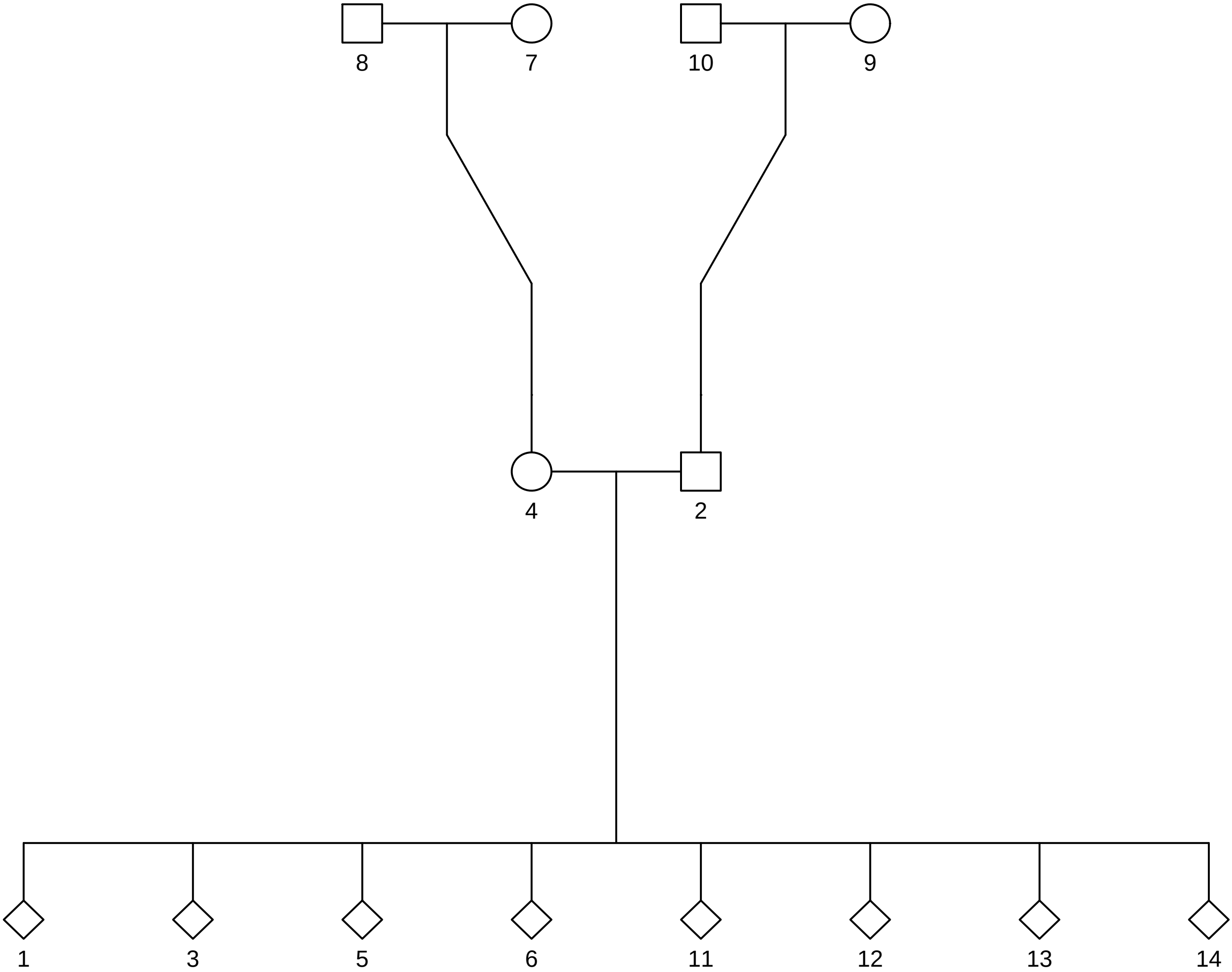

### 2.pdf

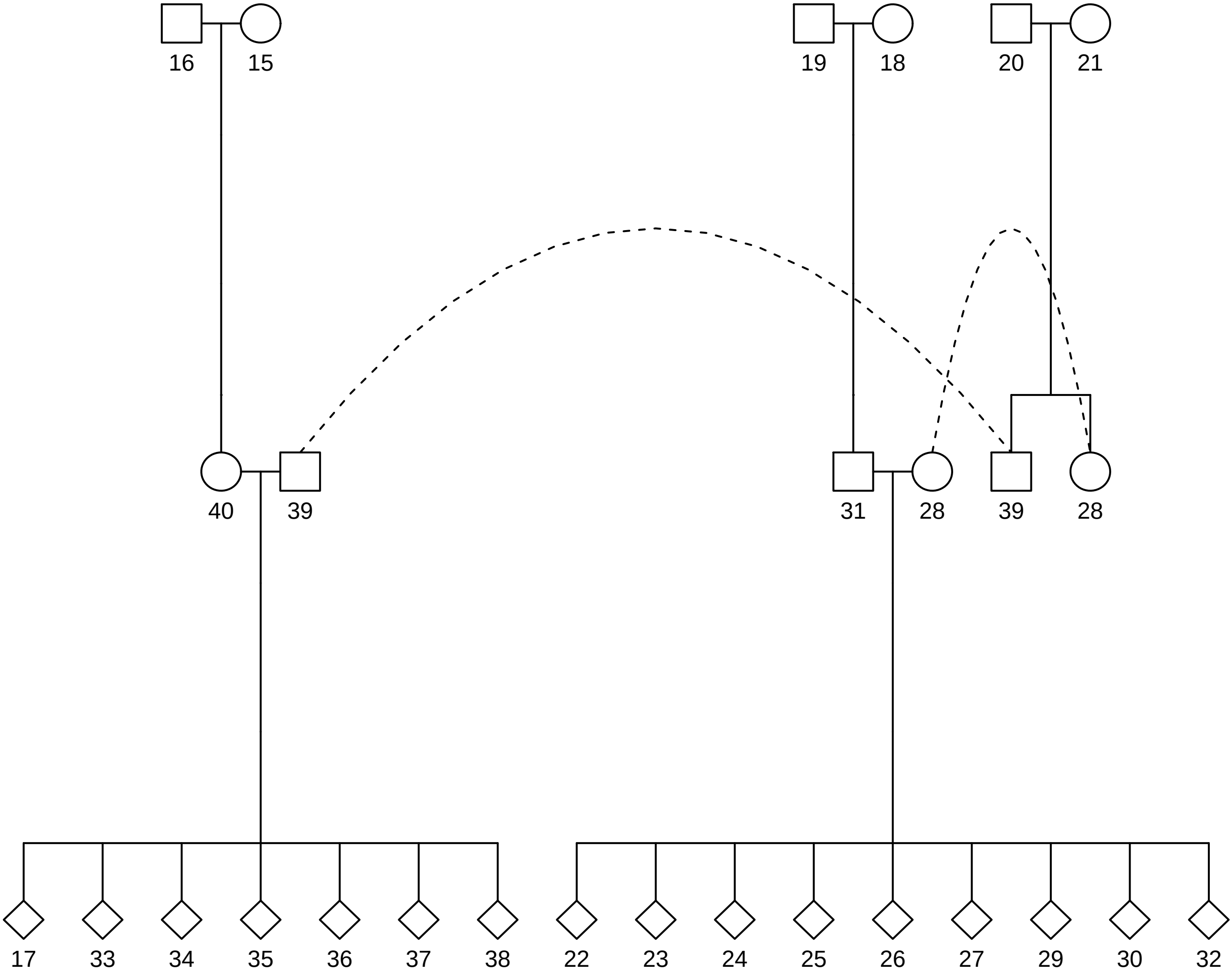

### 3.pdf

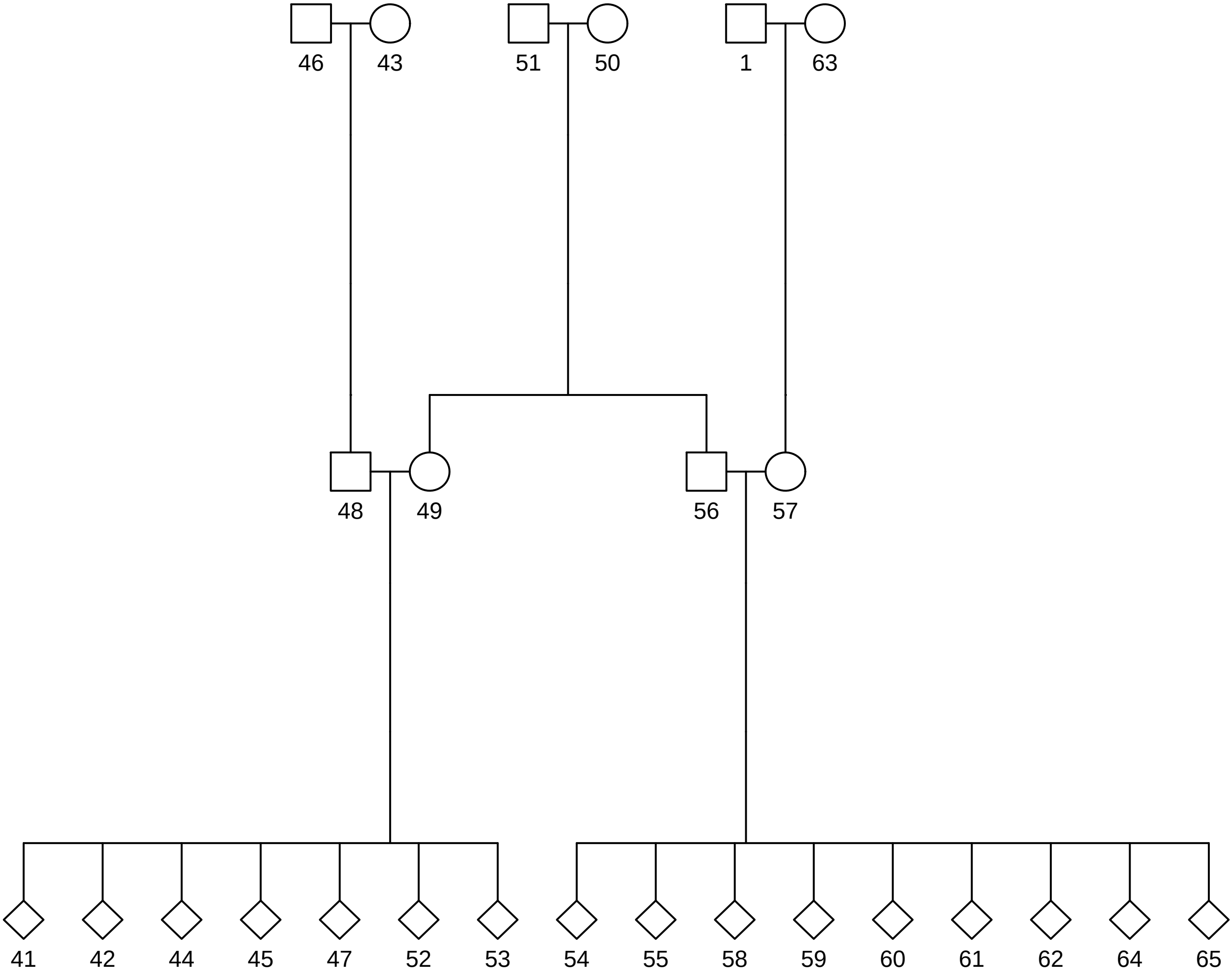

### 4.pdf

**4**

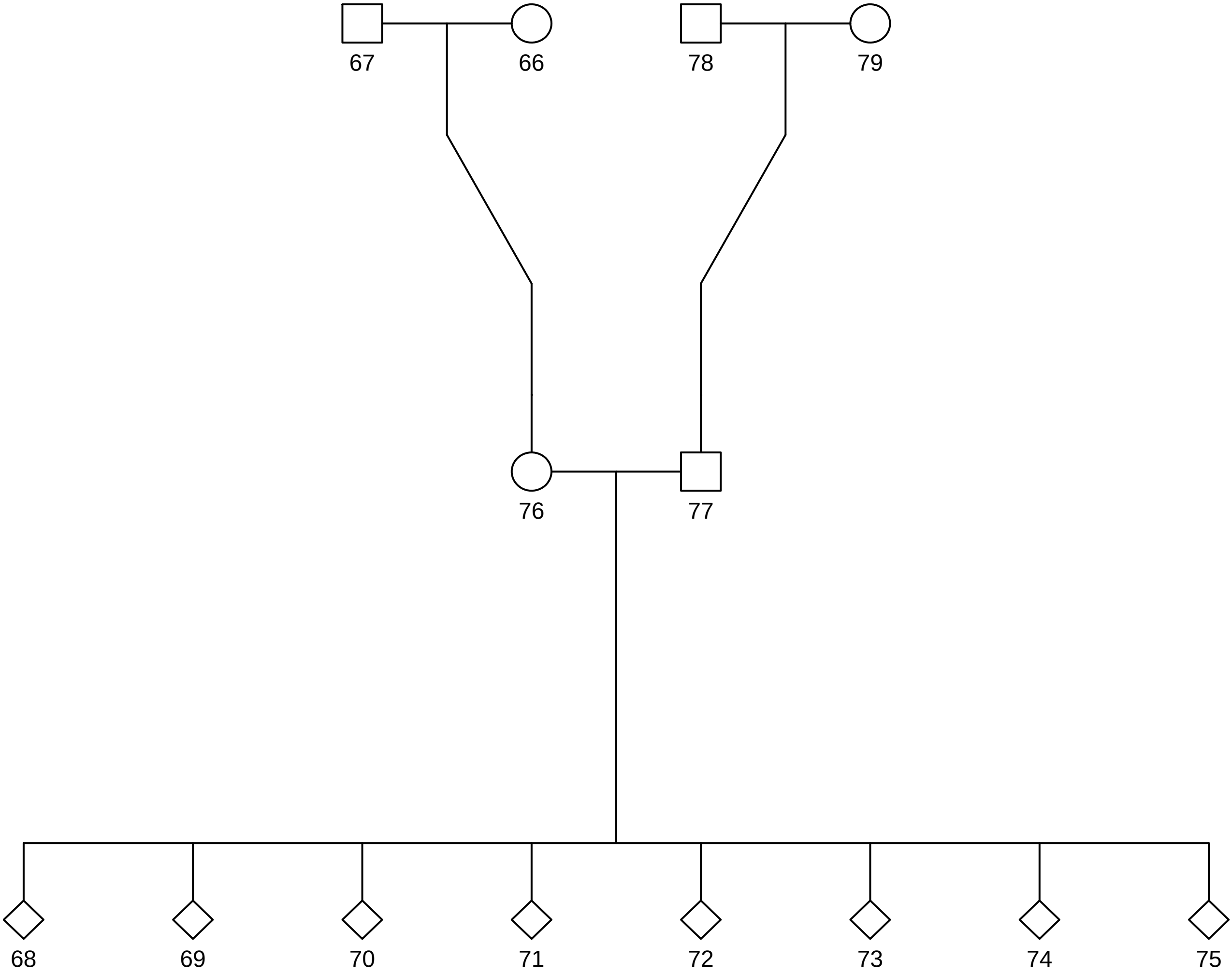

### 5.pdf

# 5

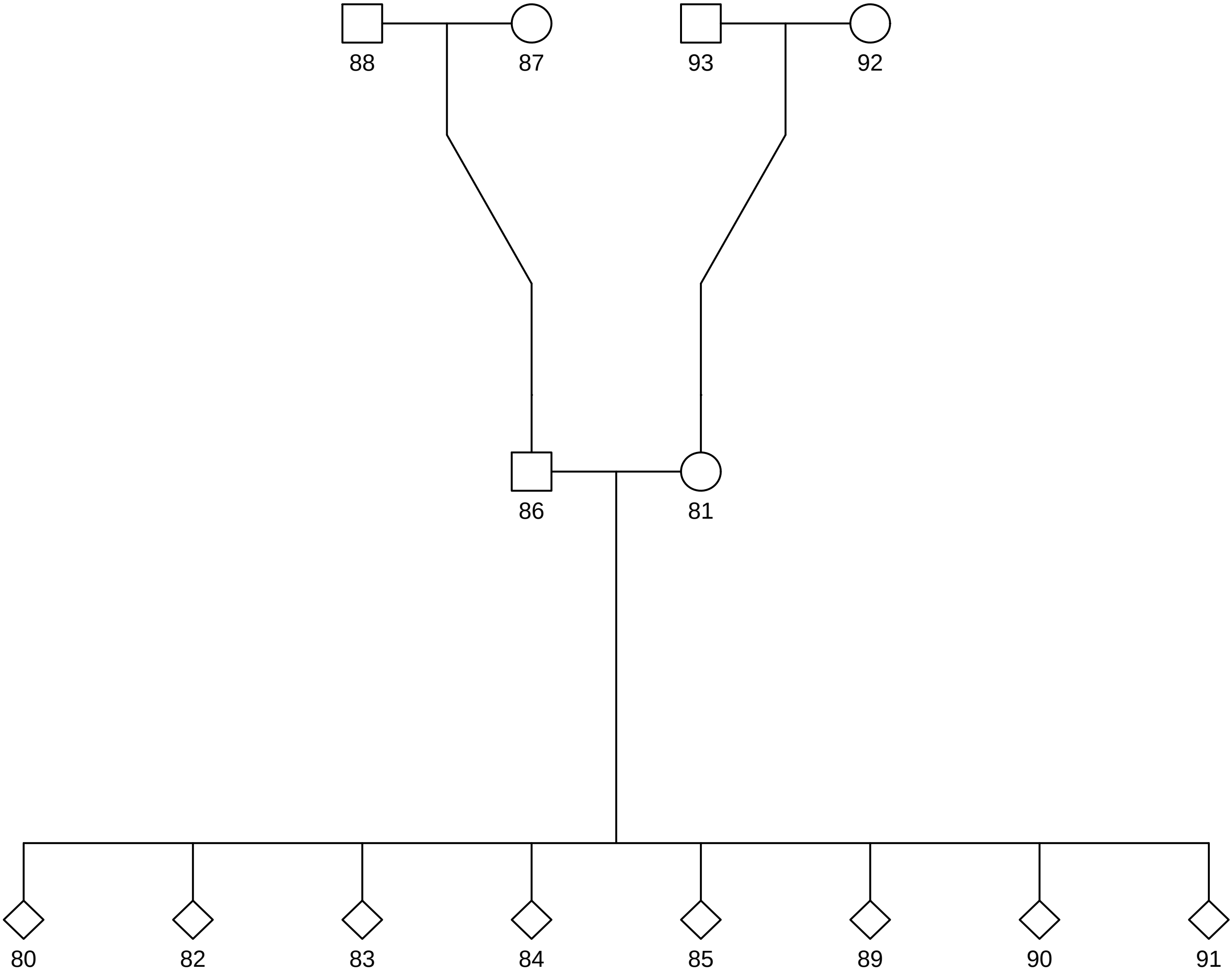

### 6.pdf

# 6

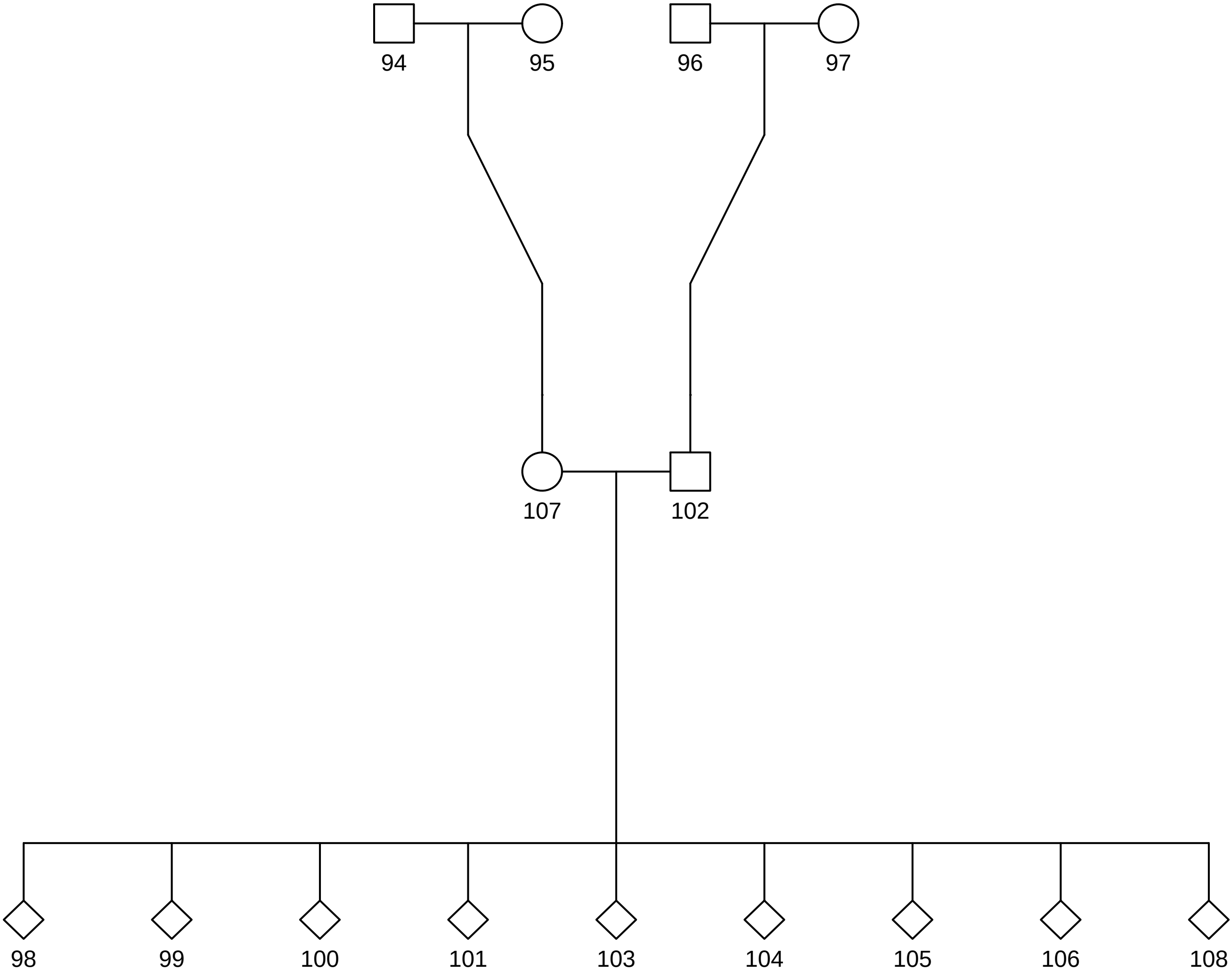

### 7.pdf

# 7

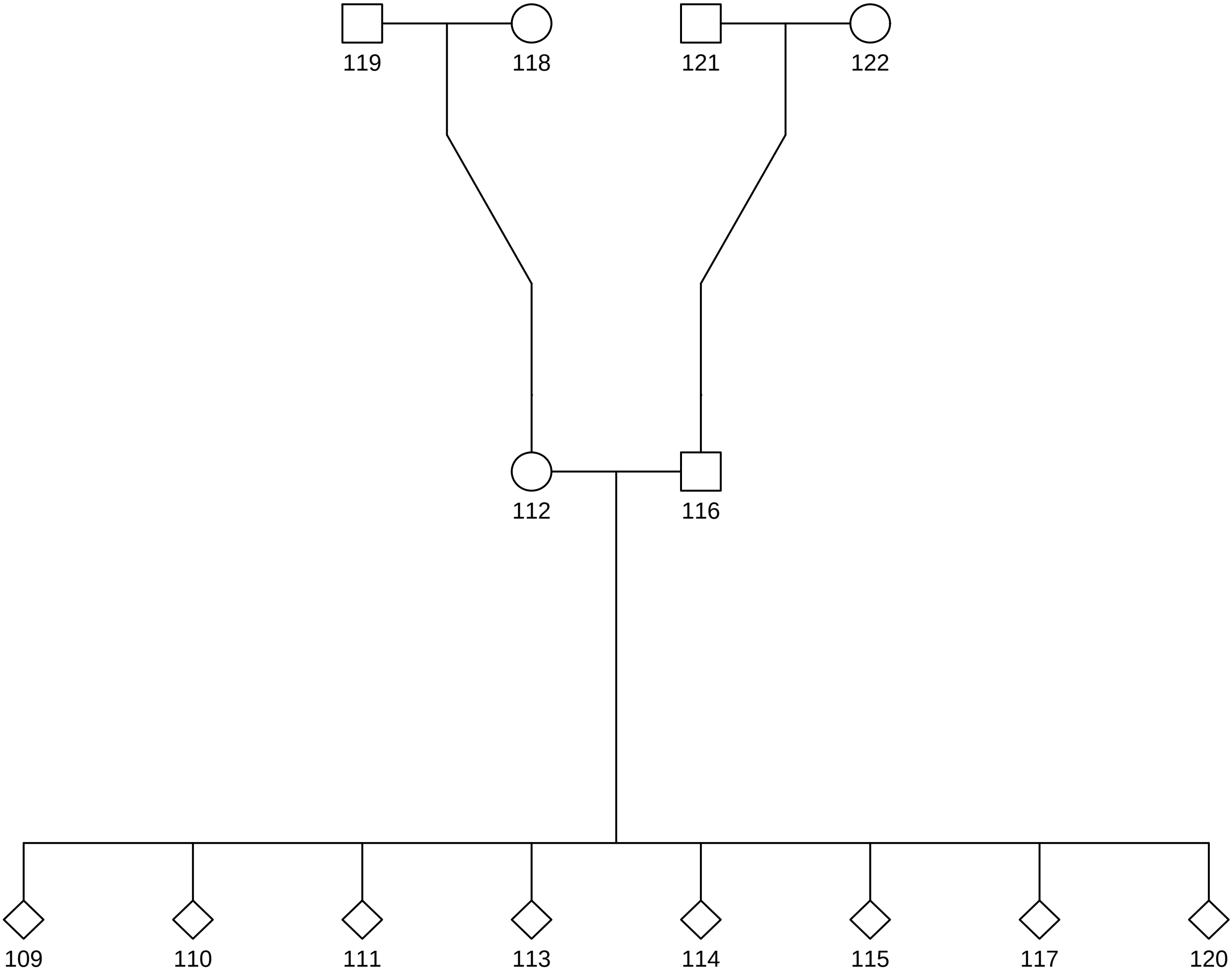

### 8.pdf

# 8

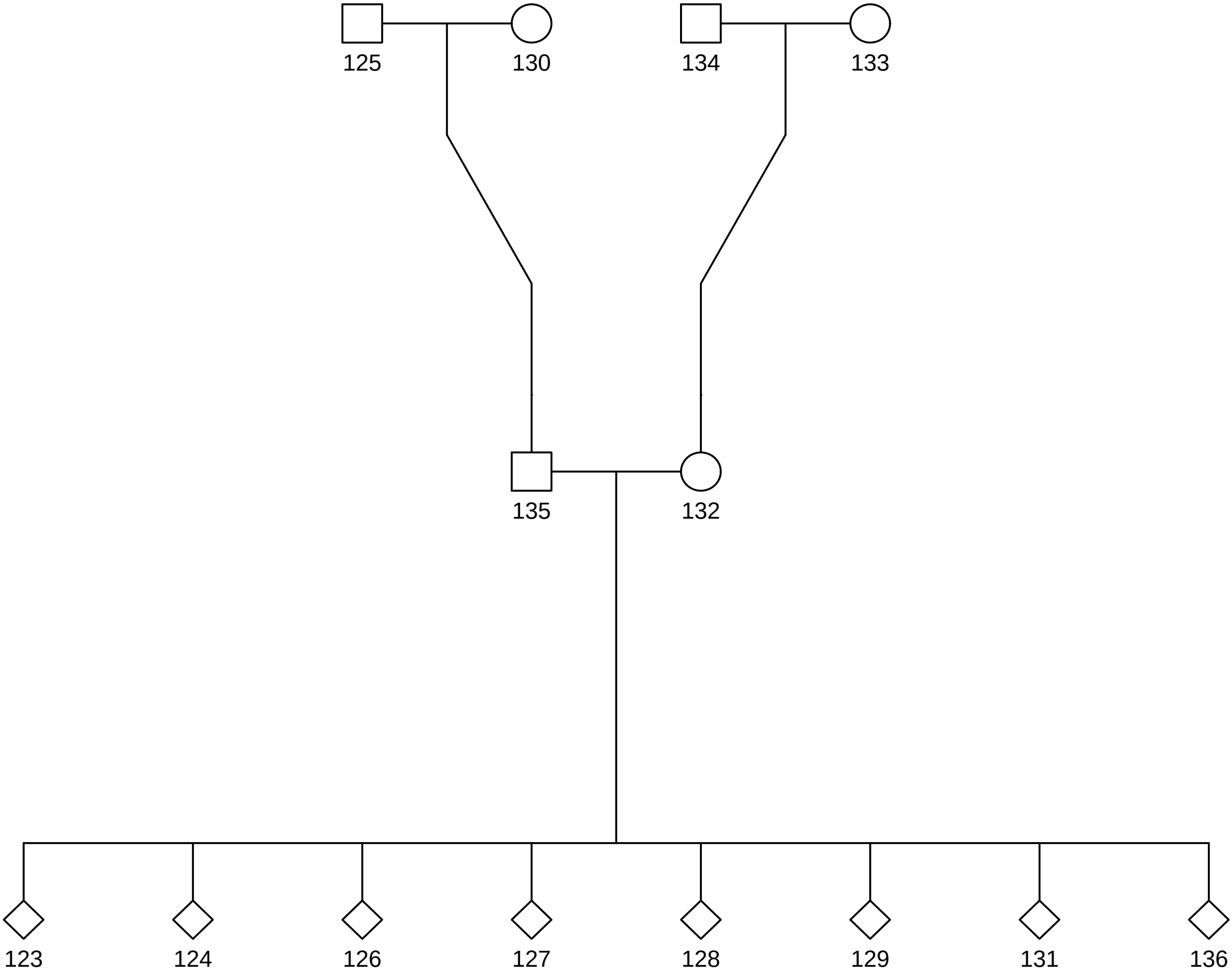

### 9.pdf

# 9

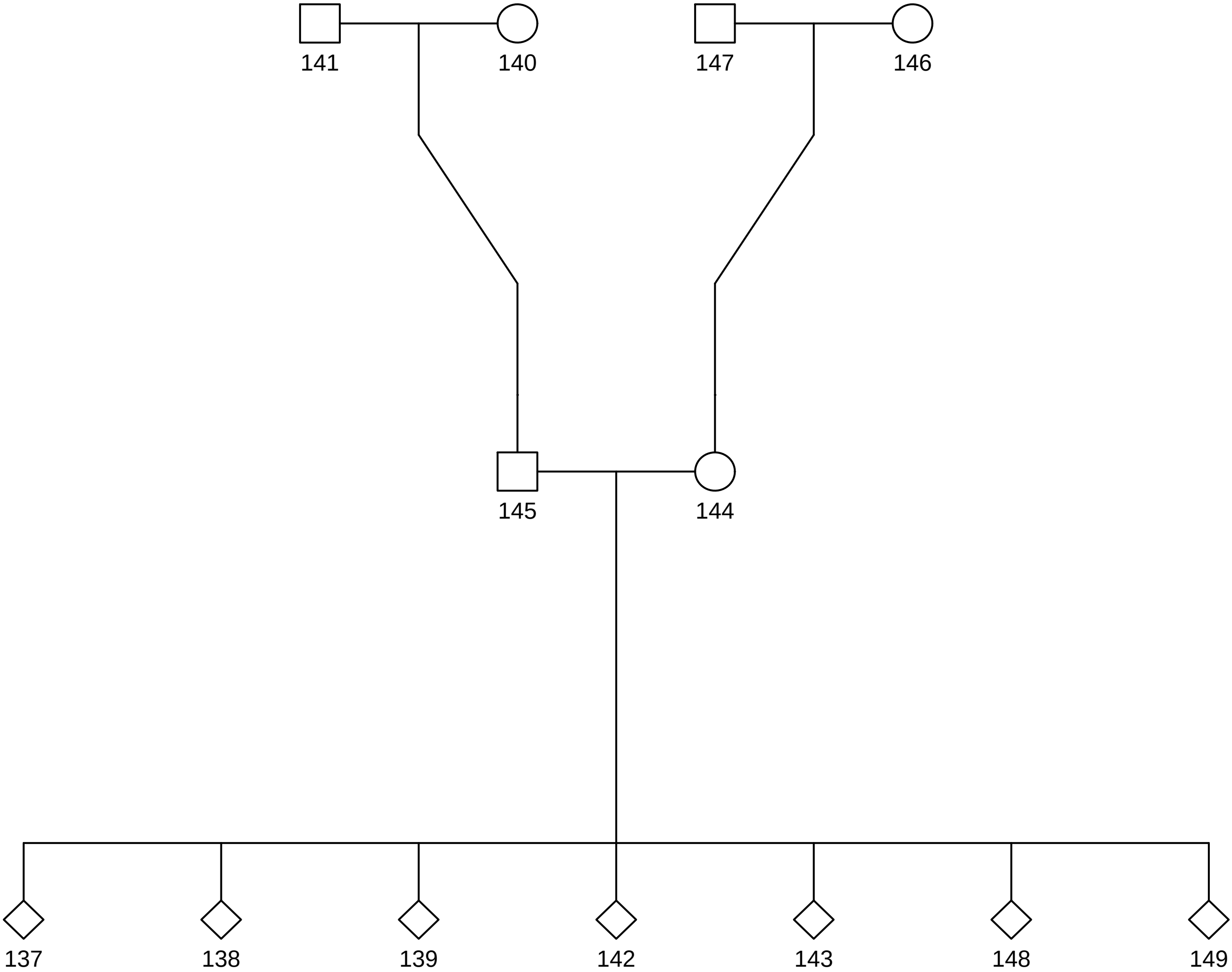

### 10.pdf

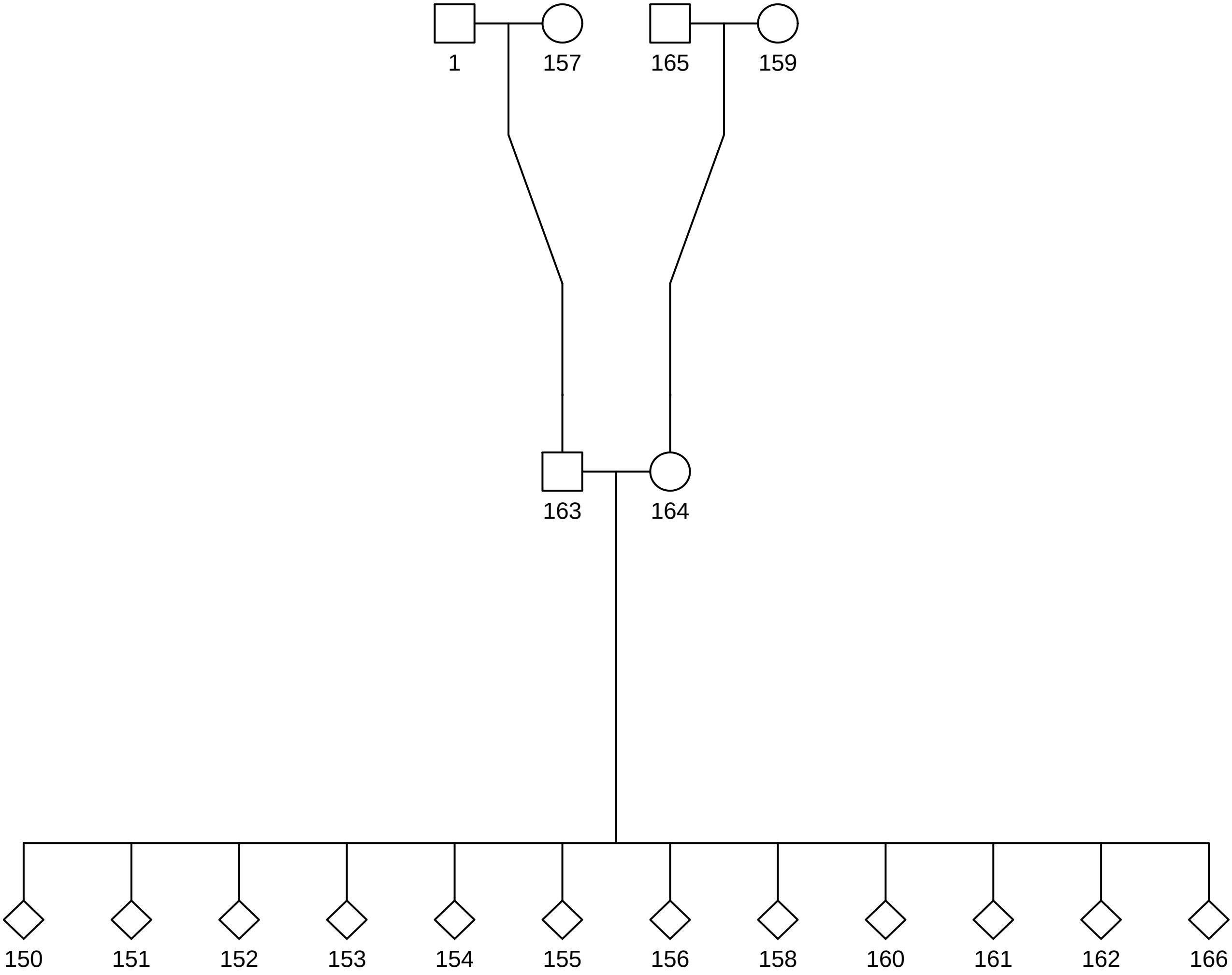

### 11.pdf

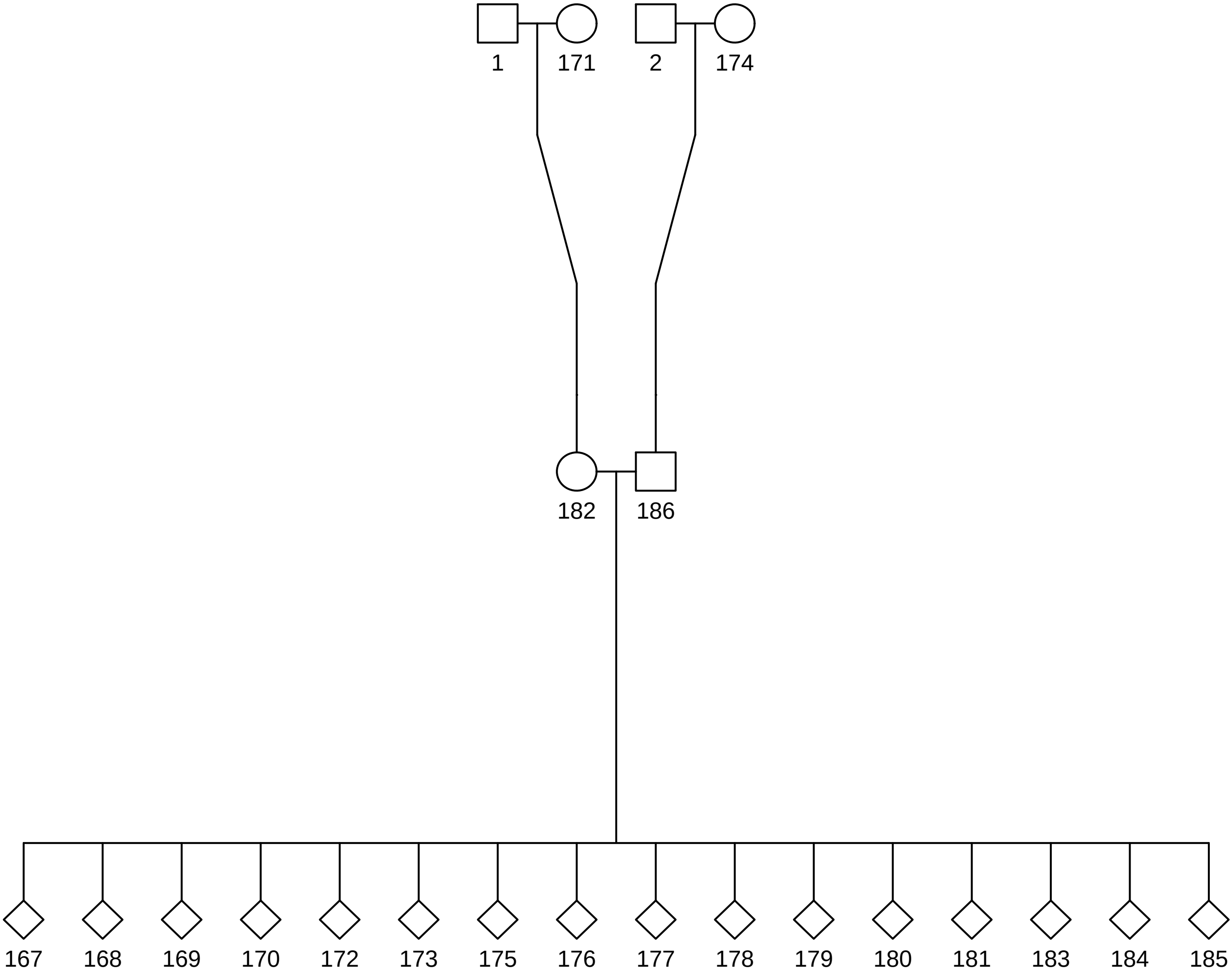

### 12.pdf

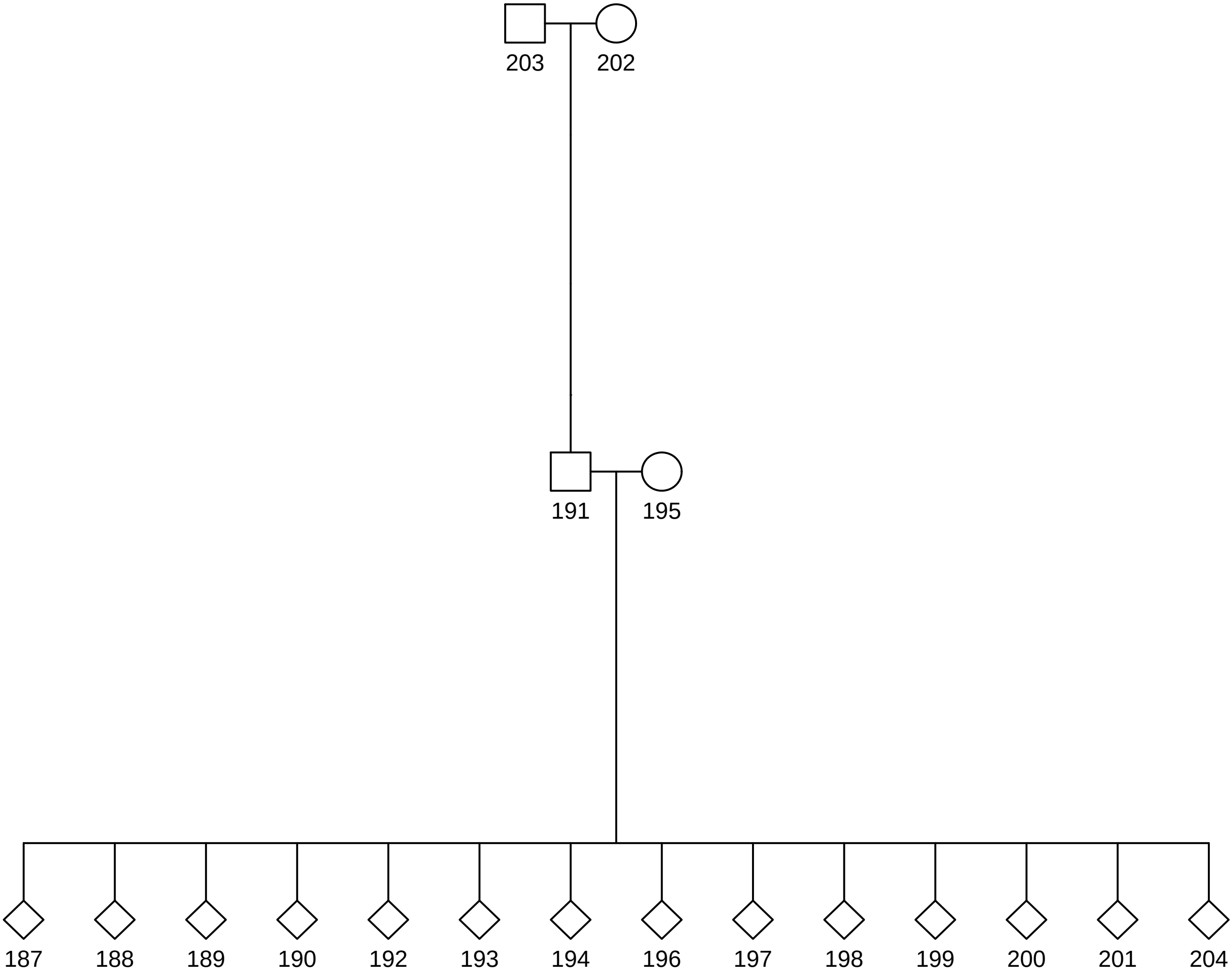

### 13.pdf

# 13

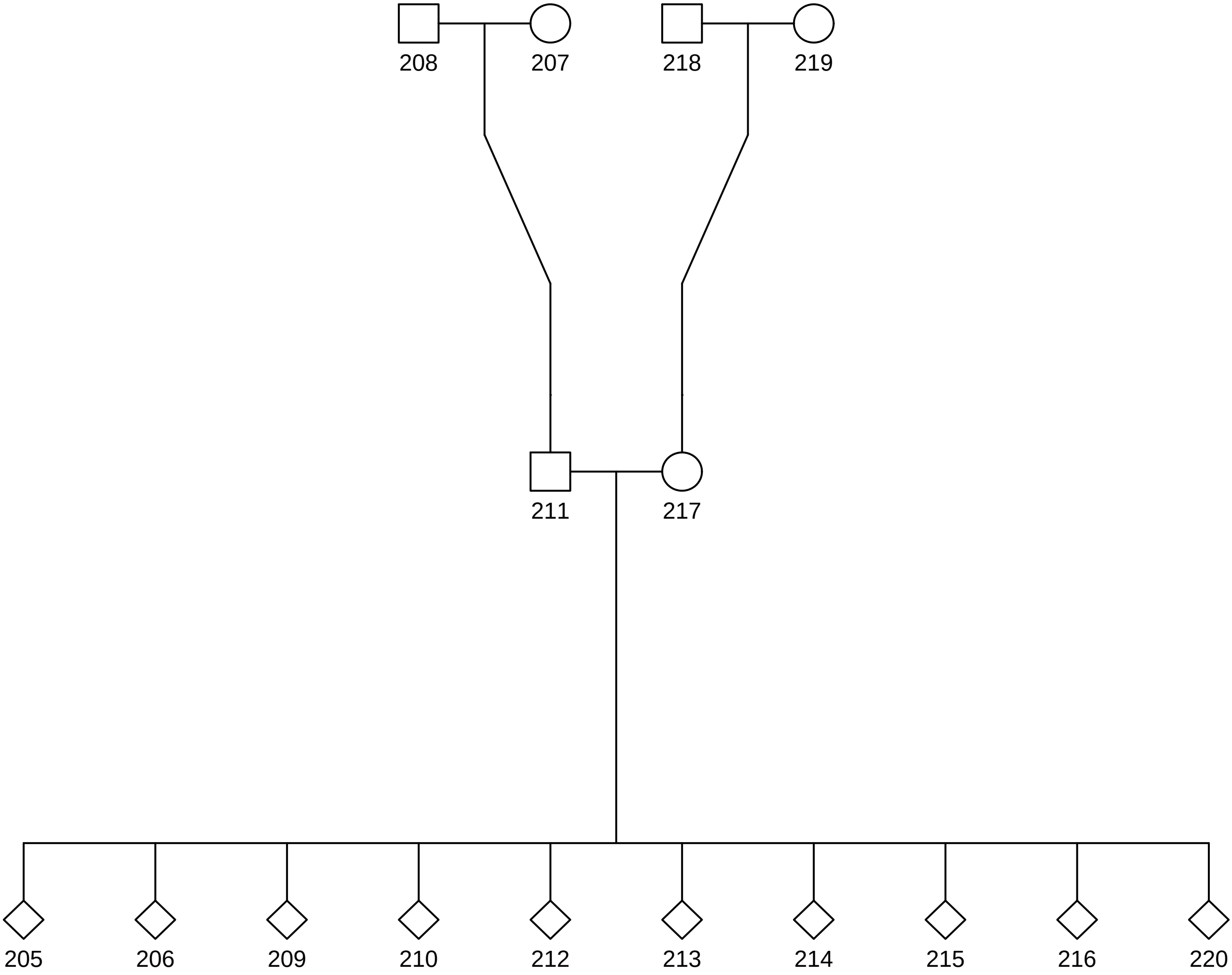

### 14.pdf

# 14

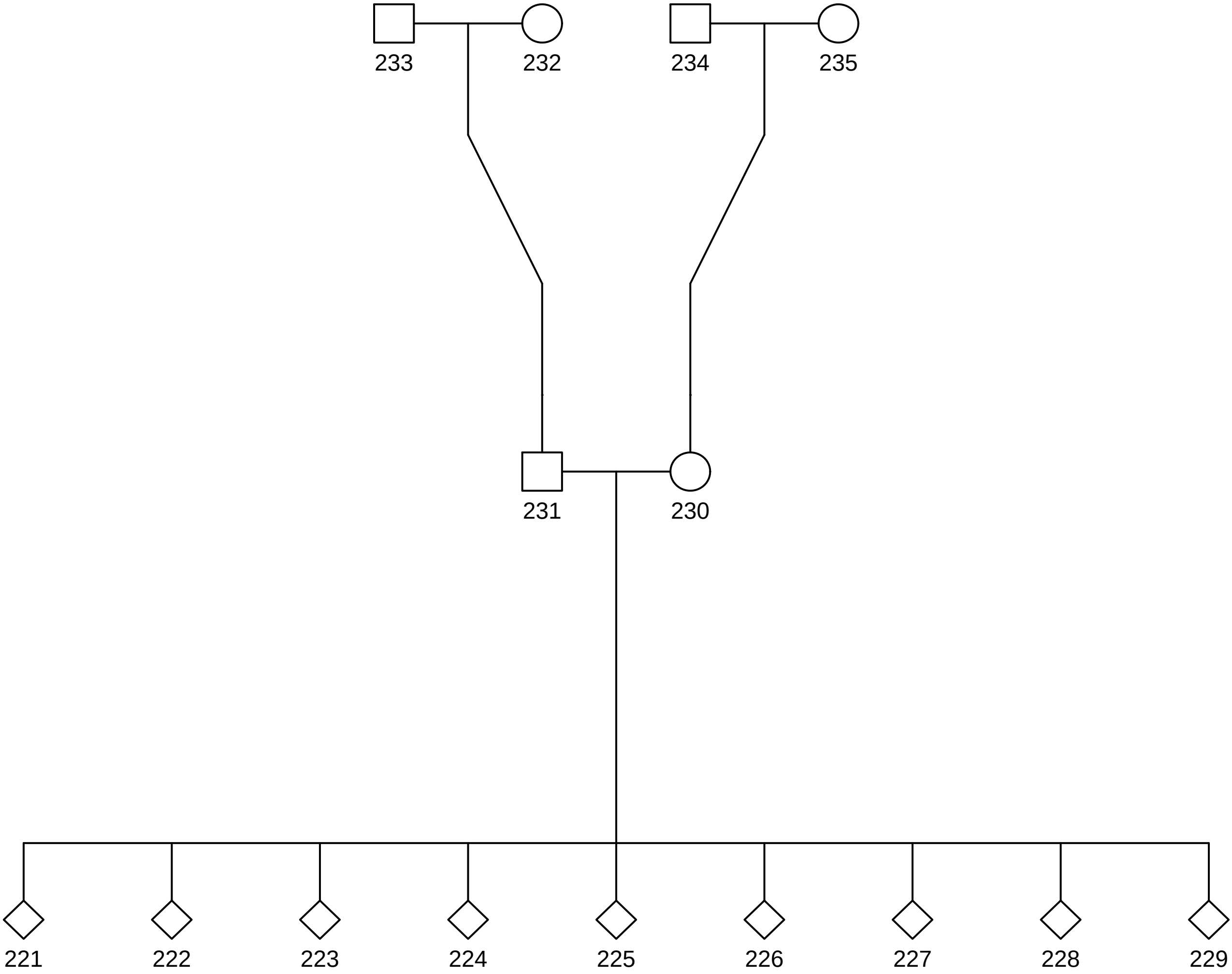

### 15.pdf

# 15

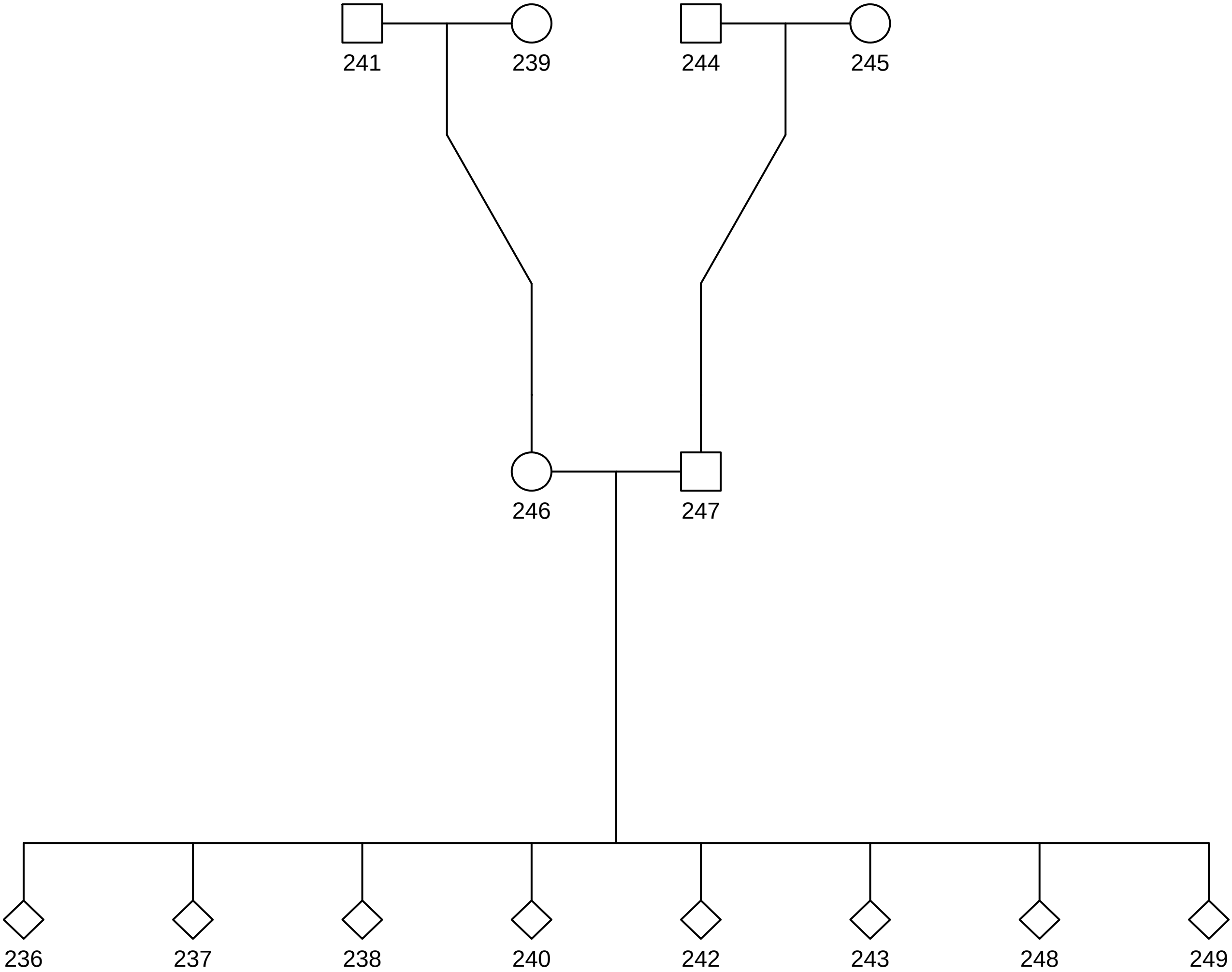

### 16.pdf

# 16

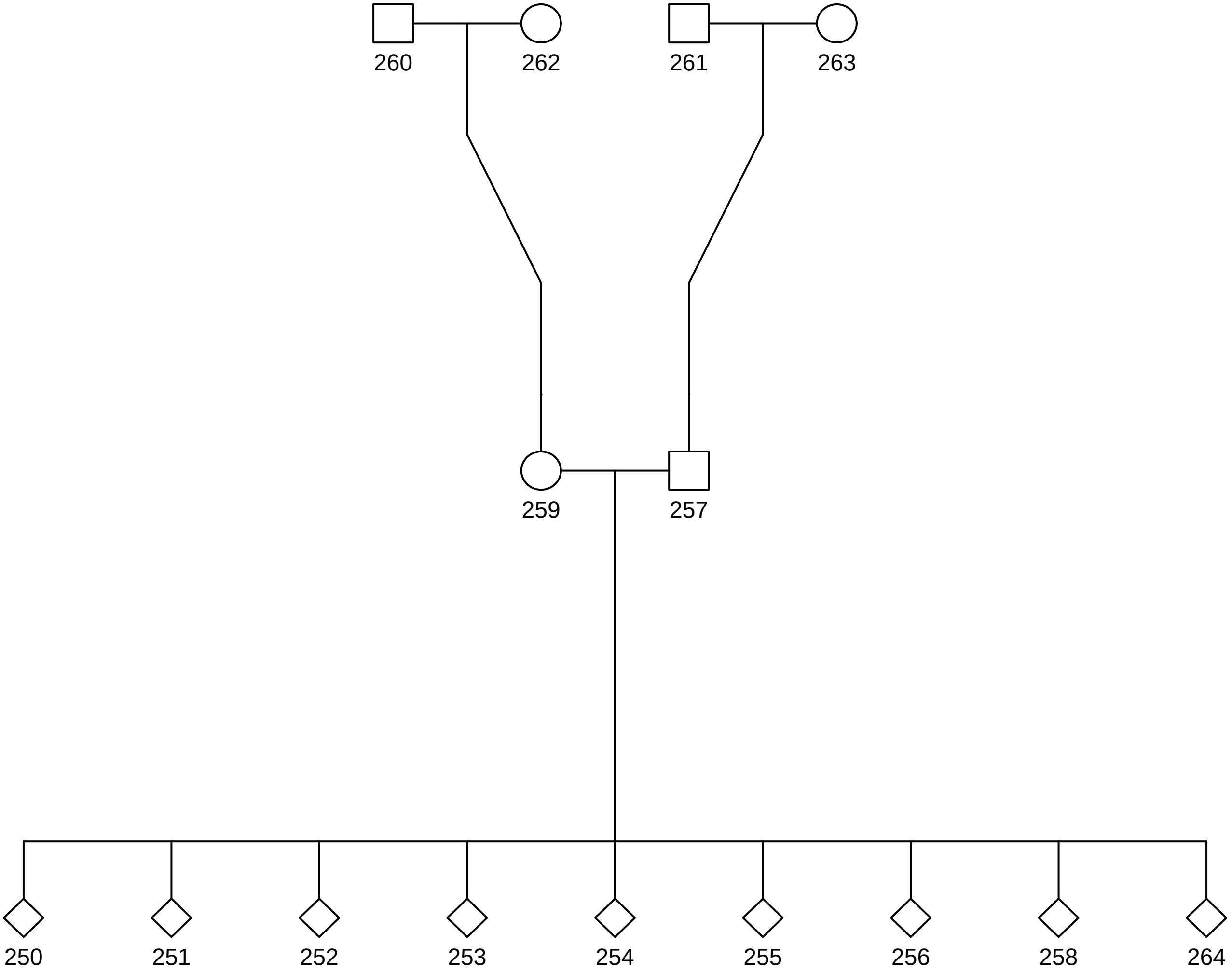

### 17.pdf

# 17

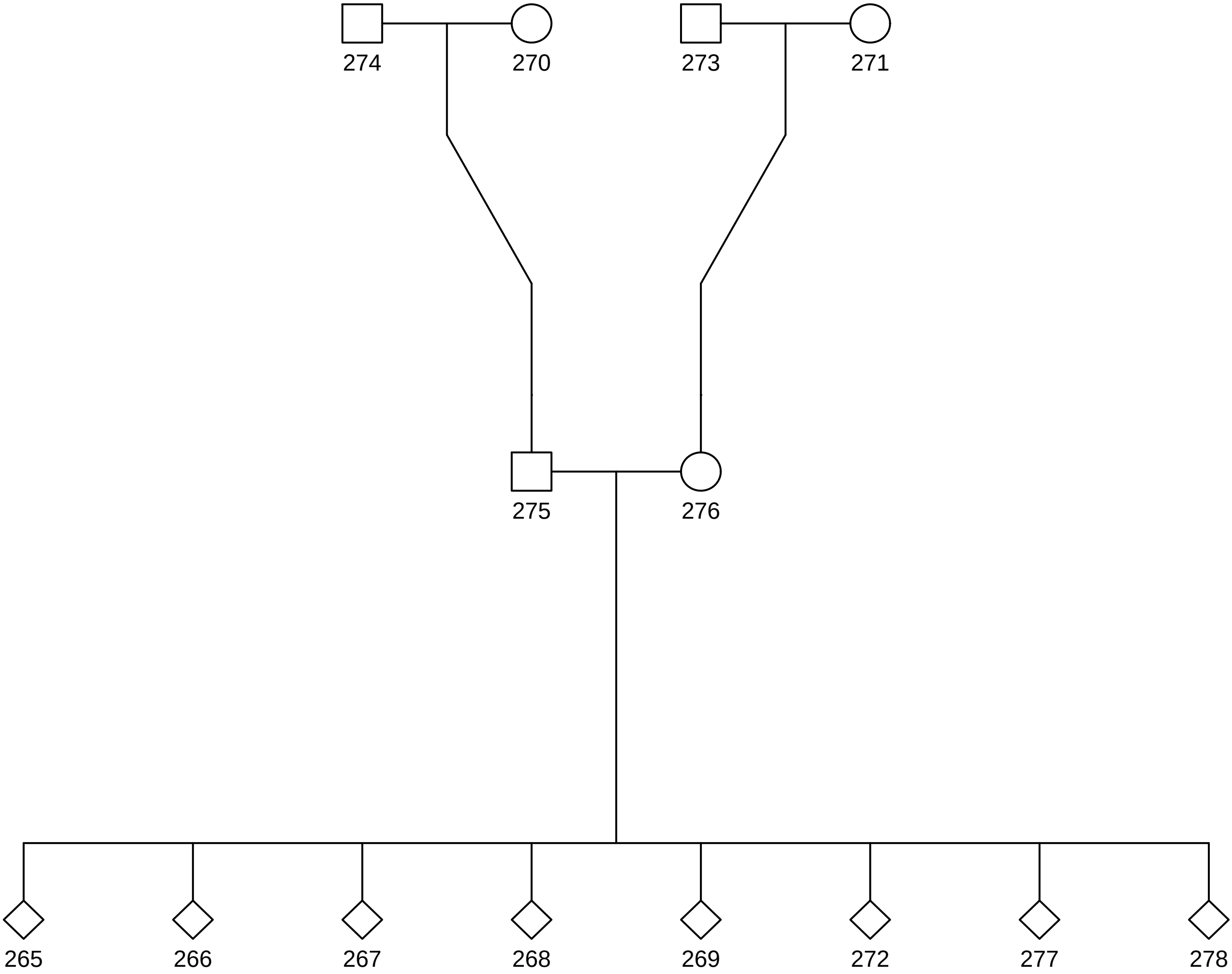

### 18.pdf

# 18

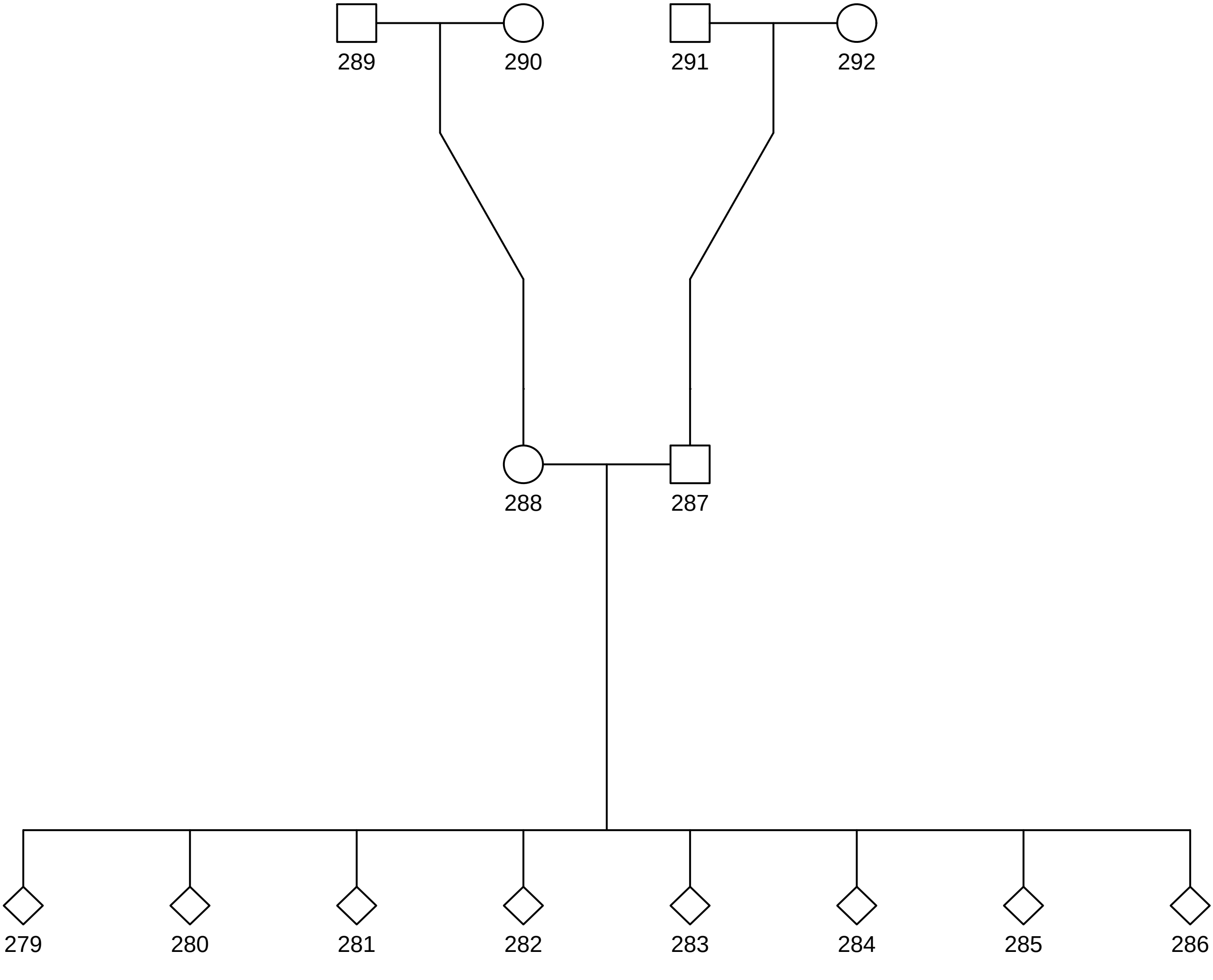

### 19.pdf

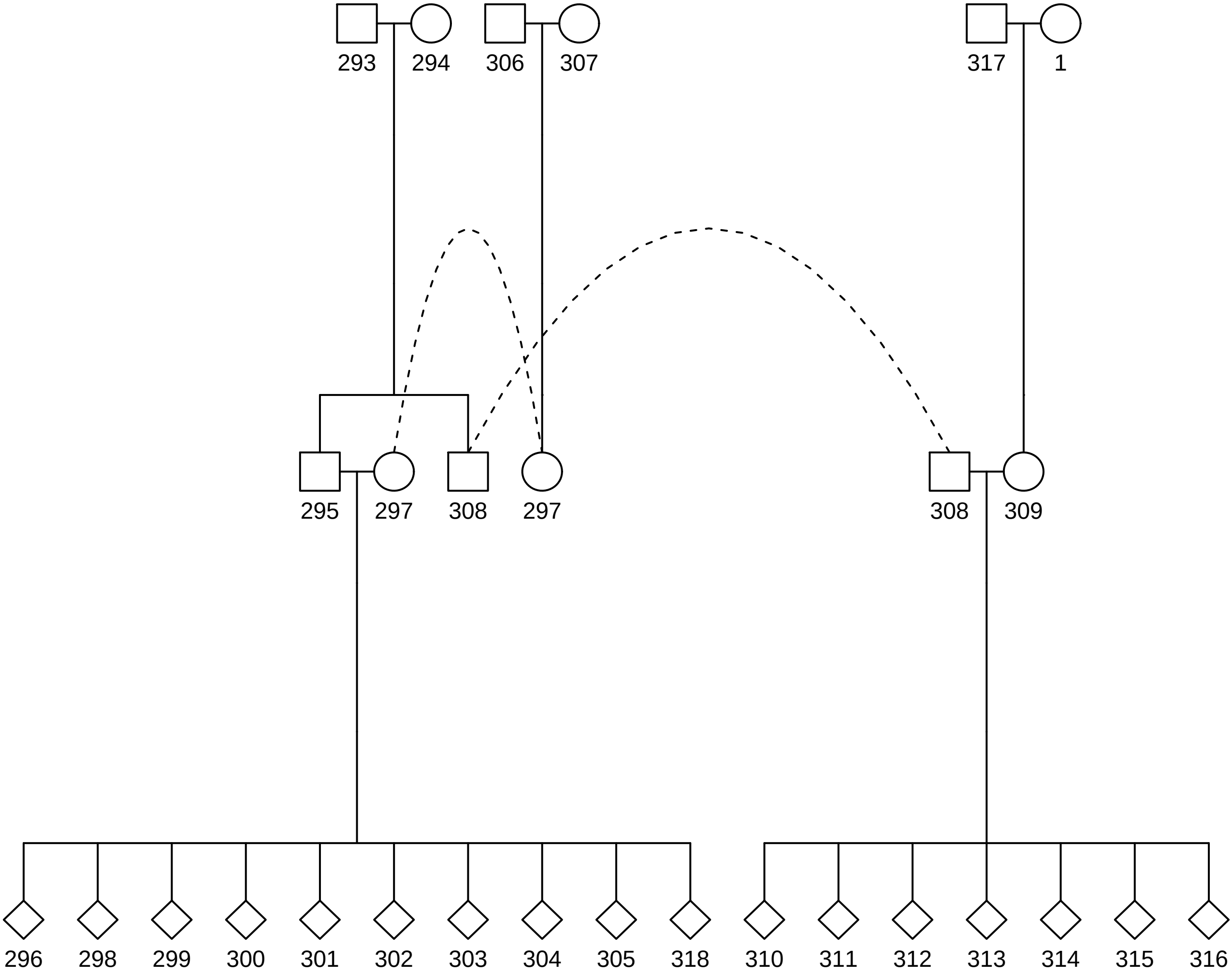

### 20.pdf

# 20

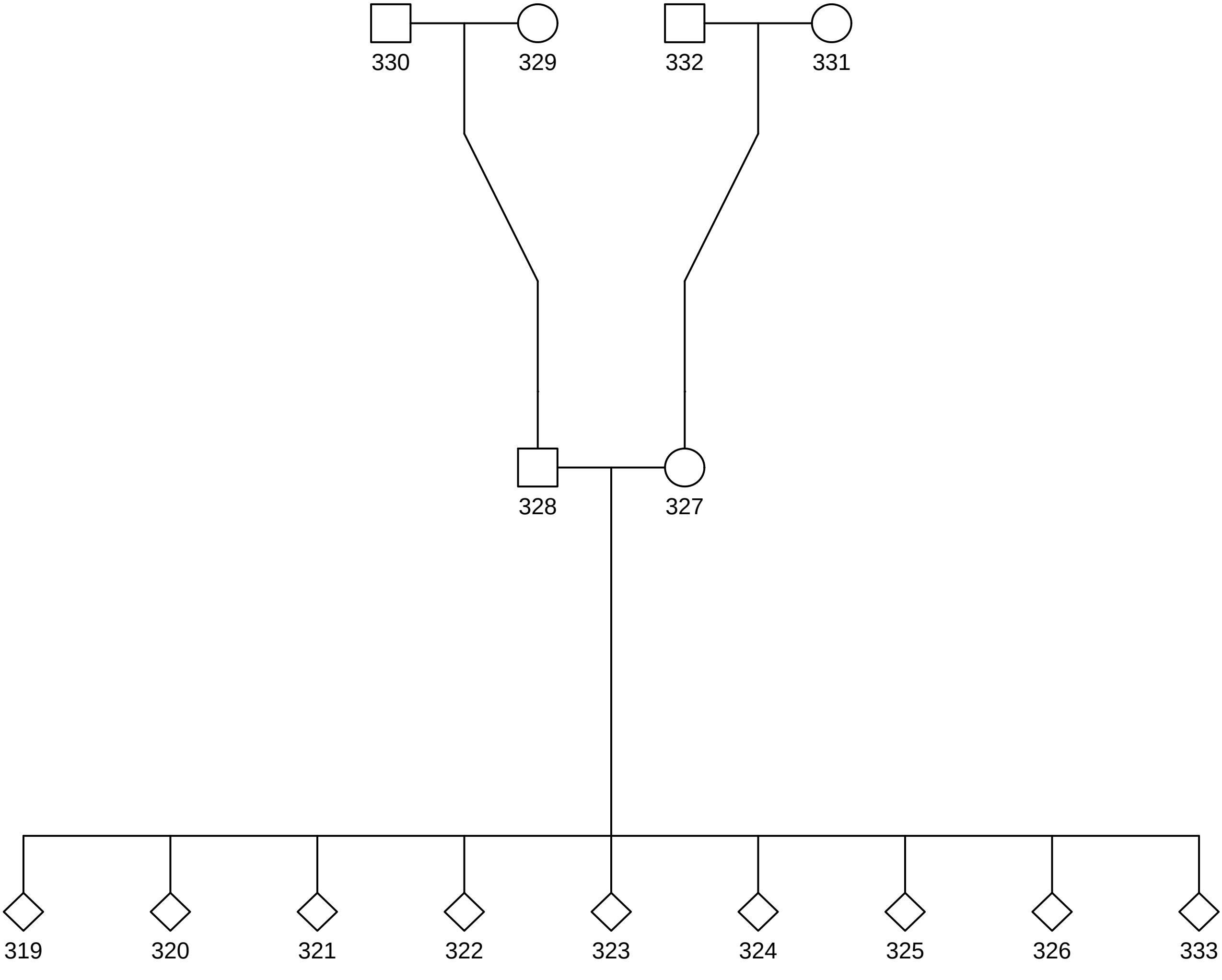

### 21.pdf

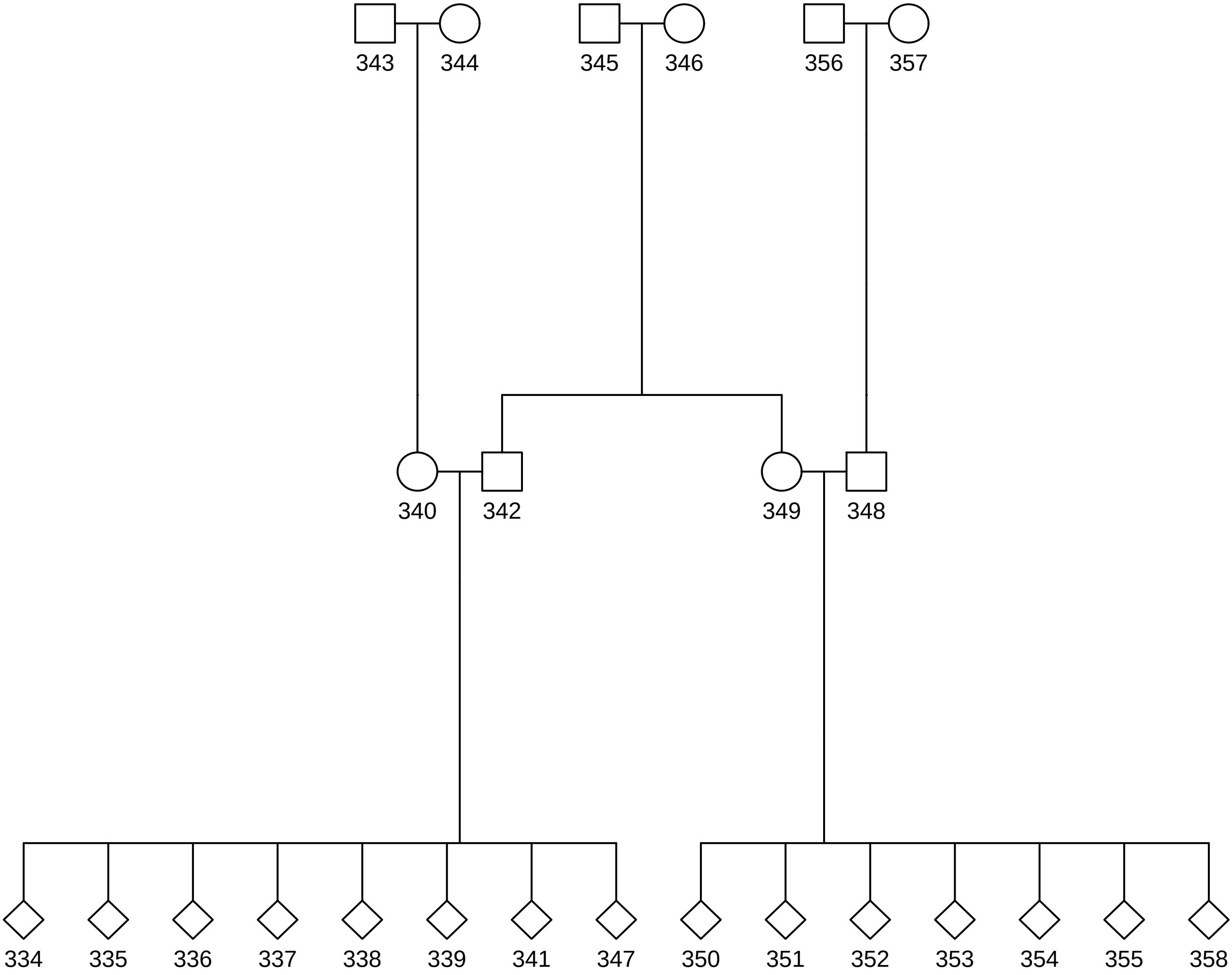

### 22.pdf

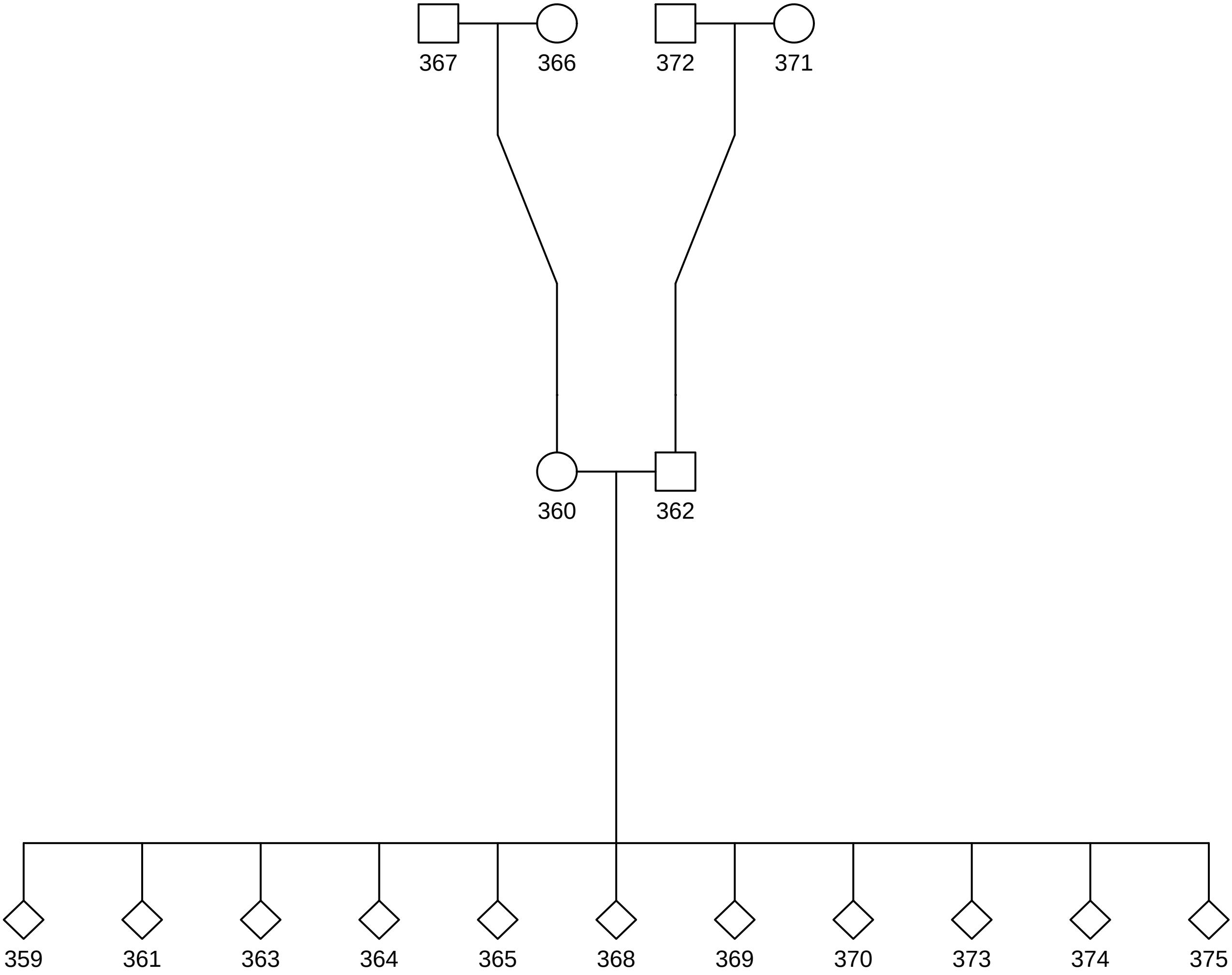

### 23.pdf

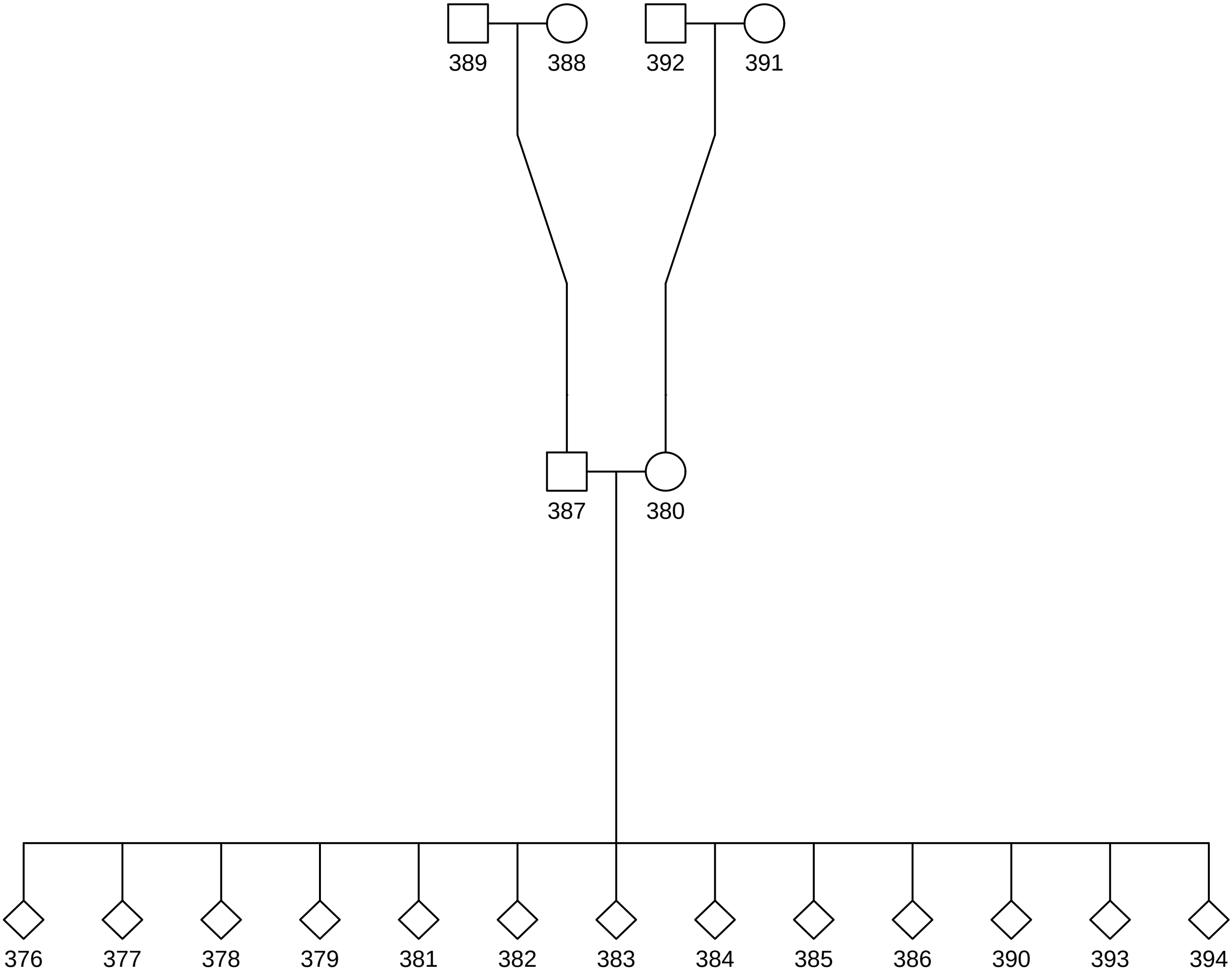

### 24.pdf

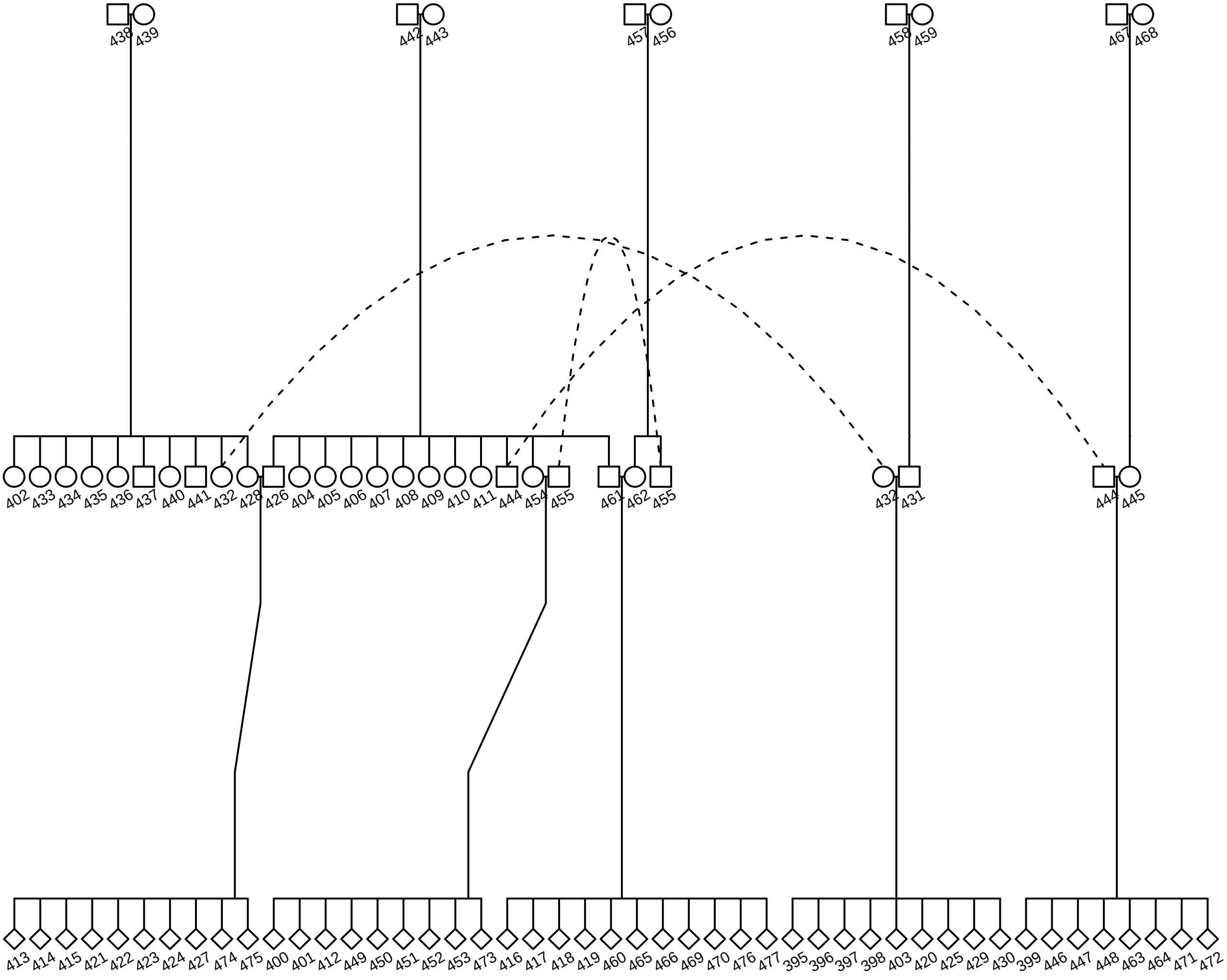

### 25.pdf

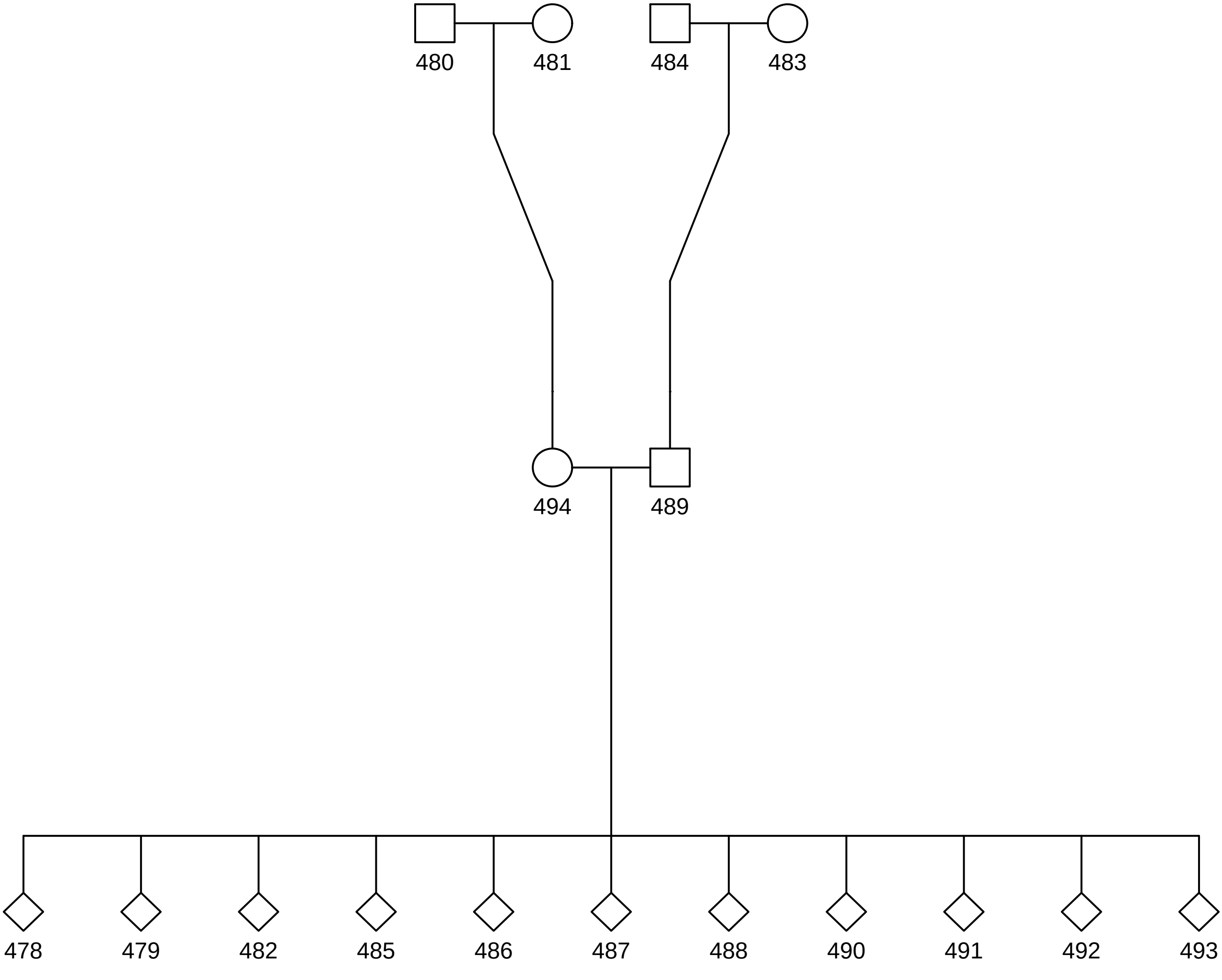

### 26.pdf

# 26

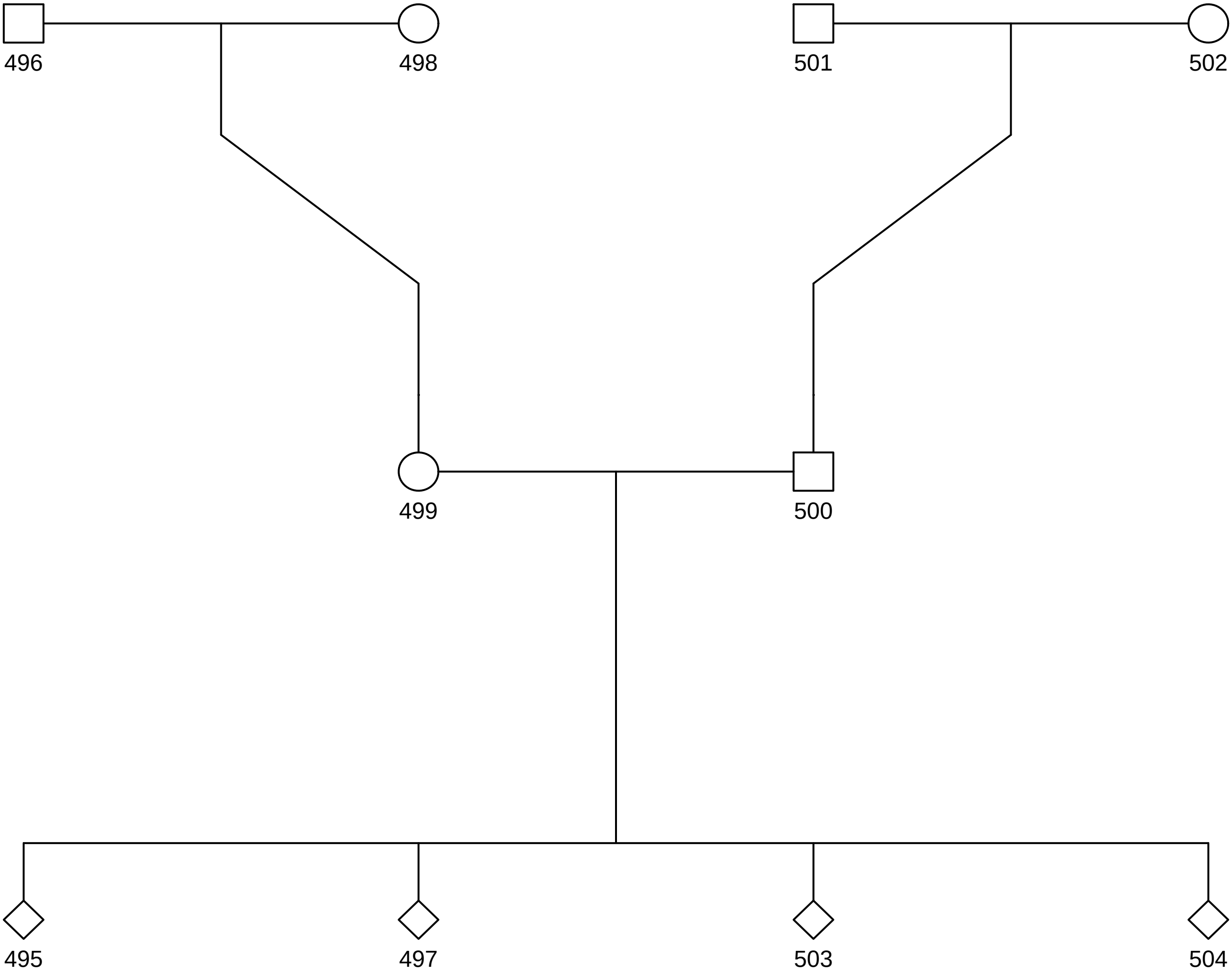

### 27.pdf

# 27

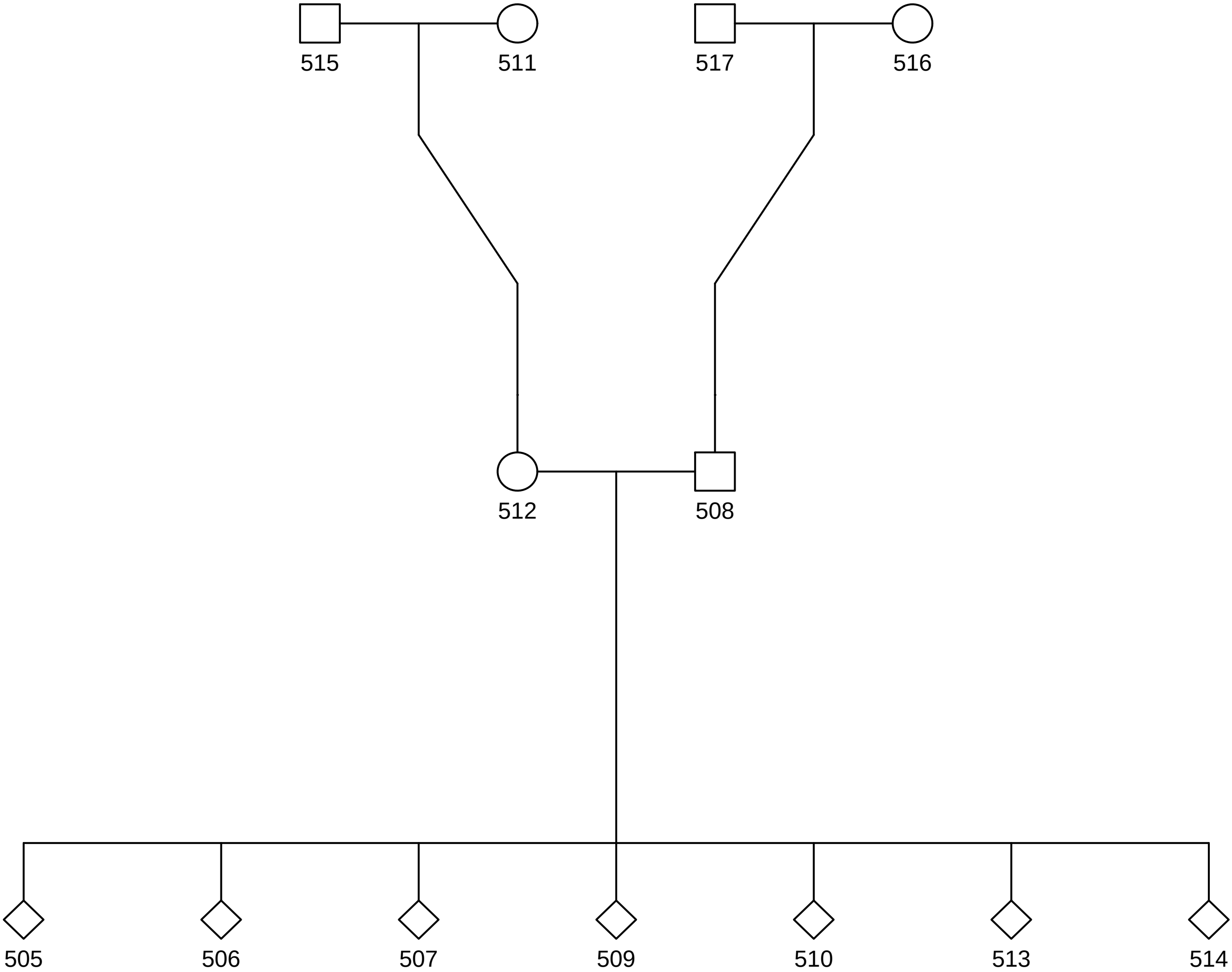

### 28.pdf

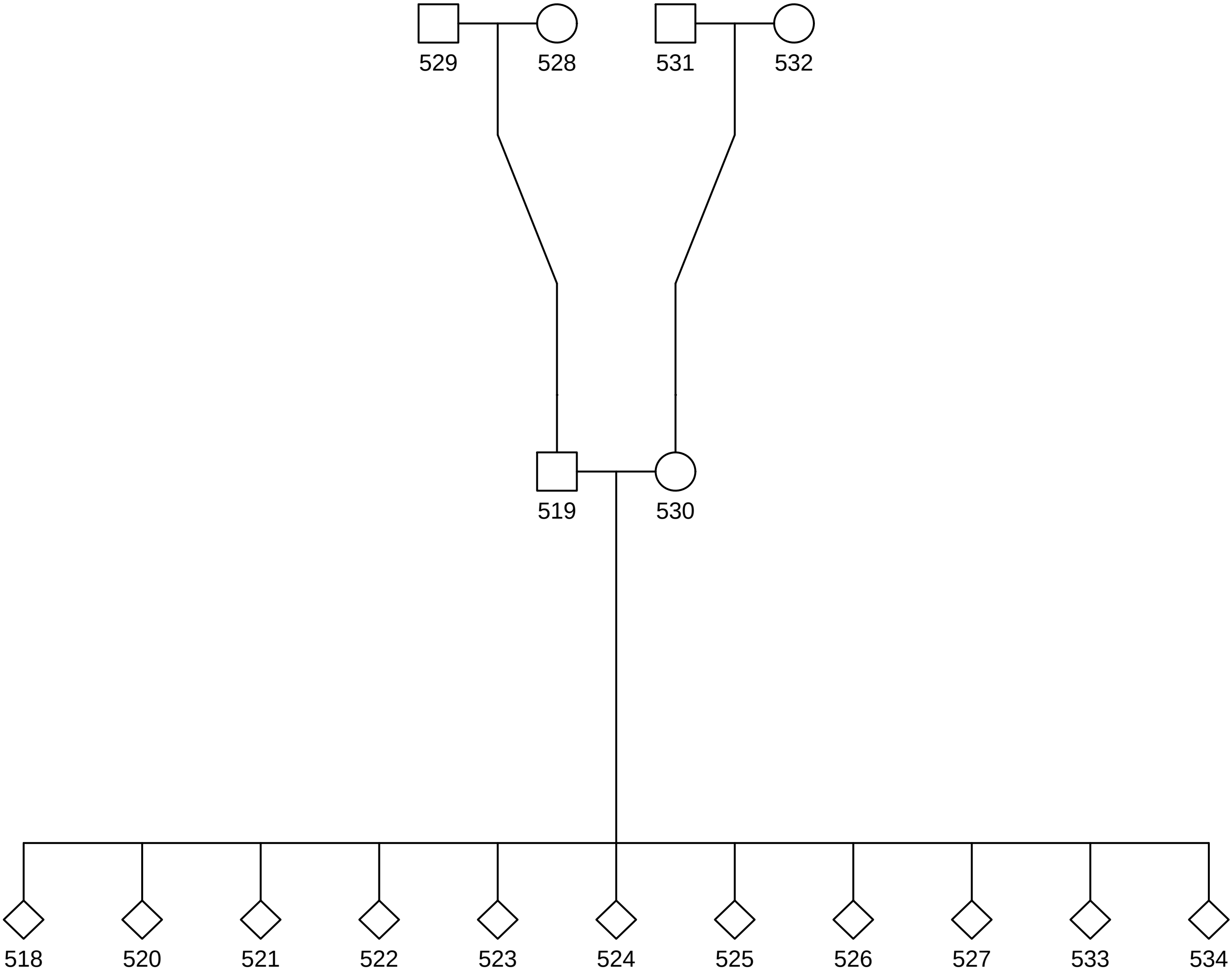

### 29.pdf

# 29

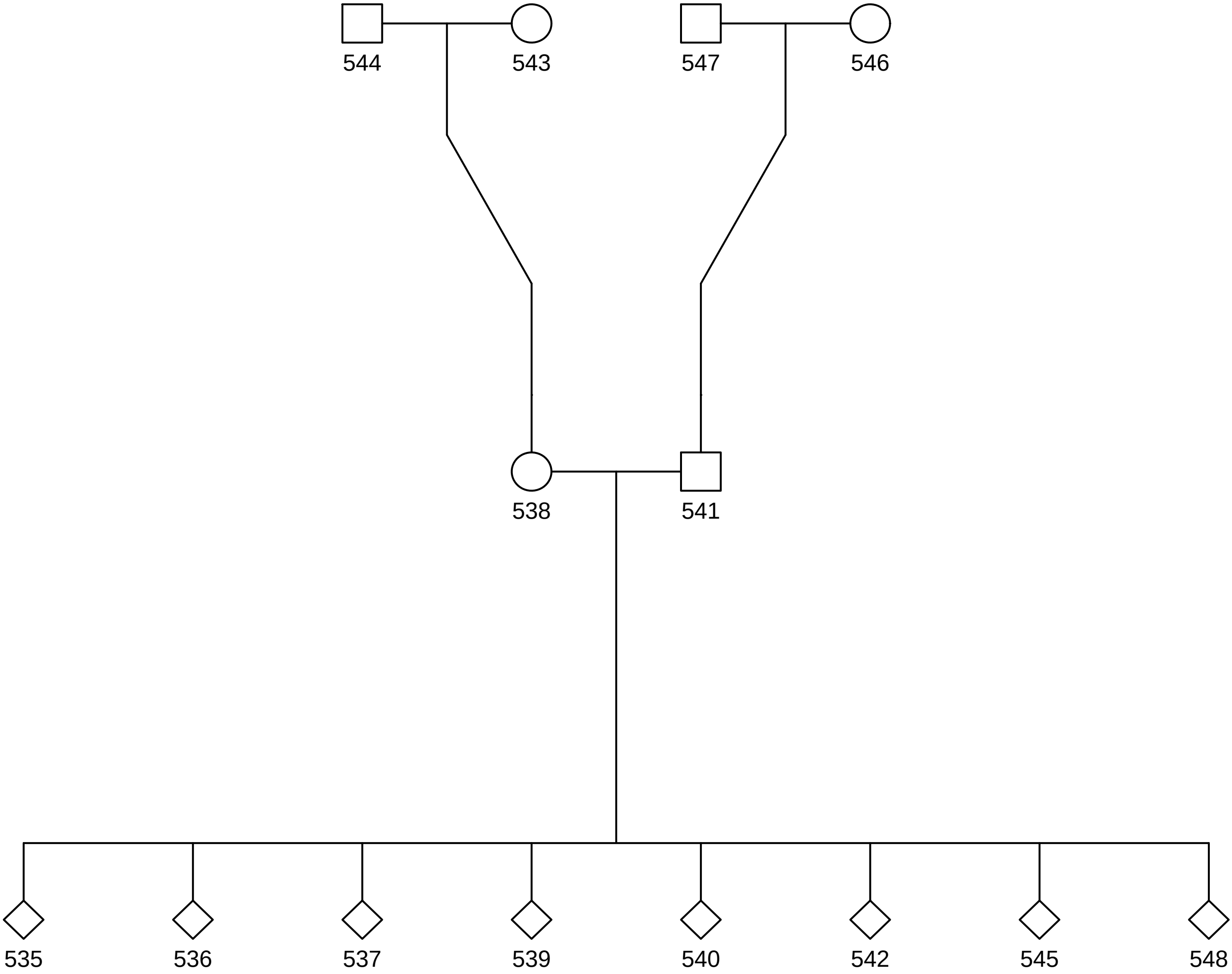

### 30.pdf

# 30

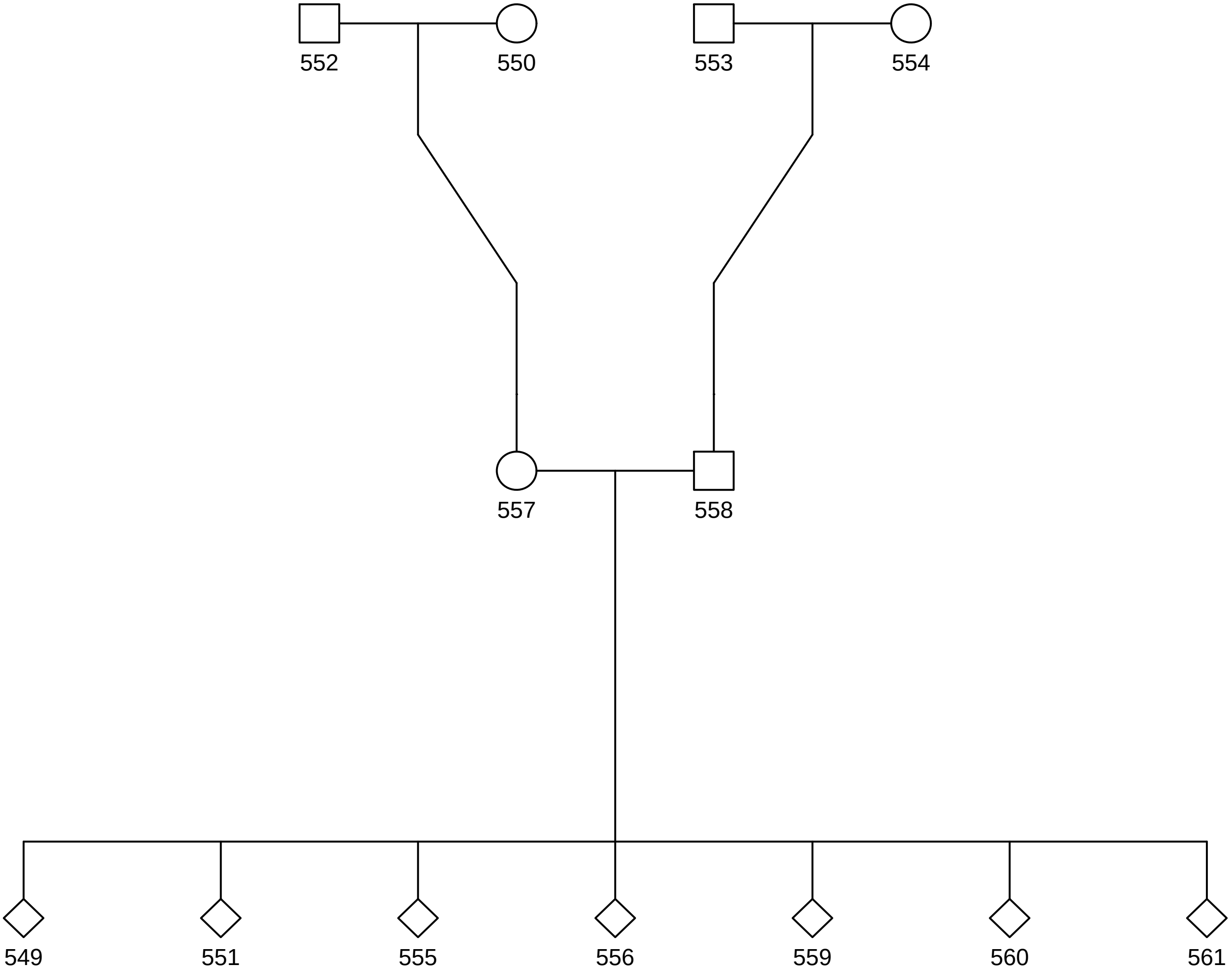

### 31.pdf

# 31

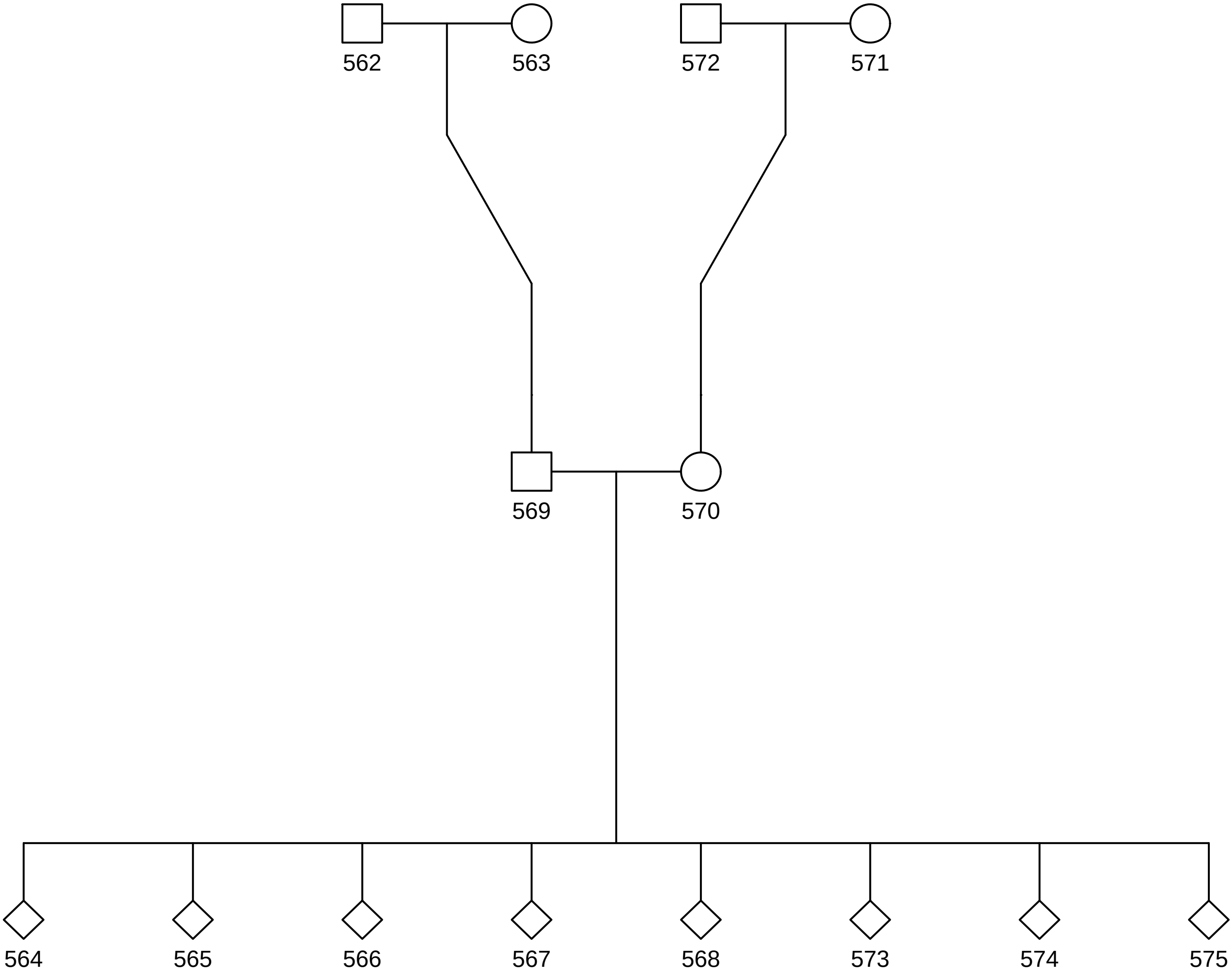

### 32.pdf

# 32

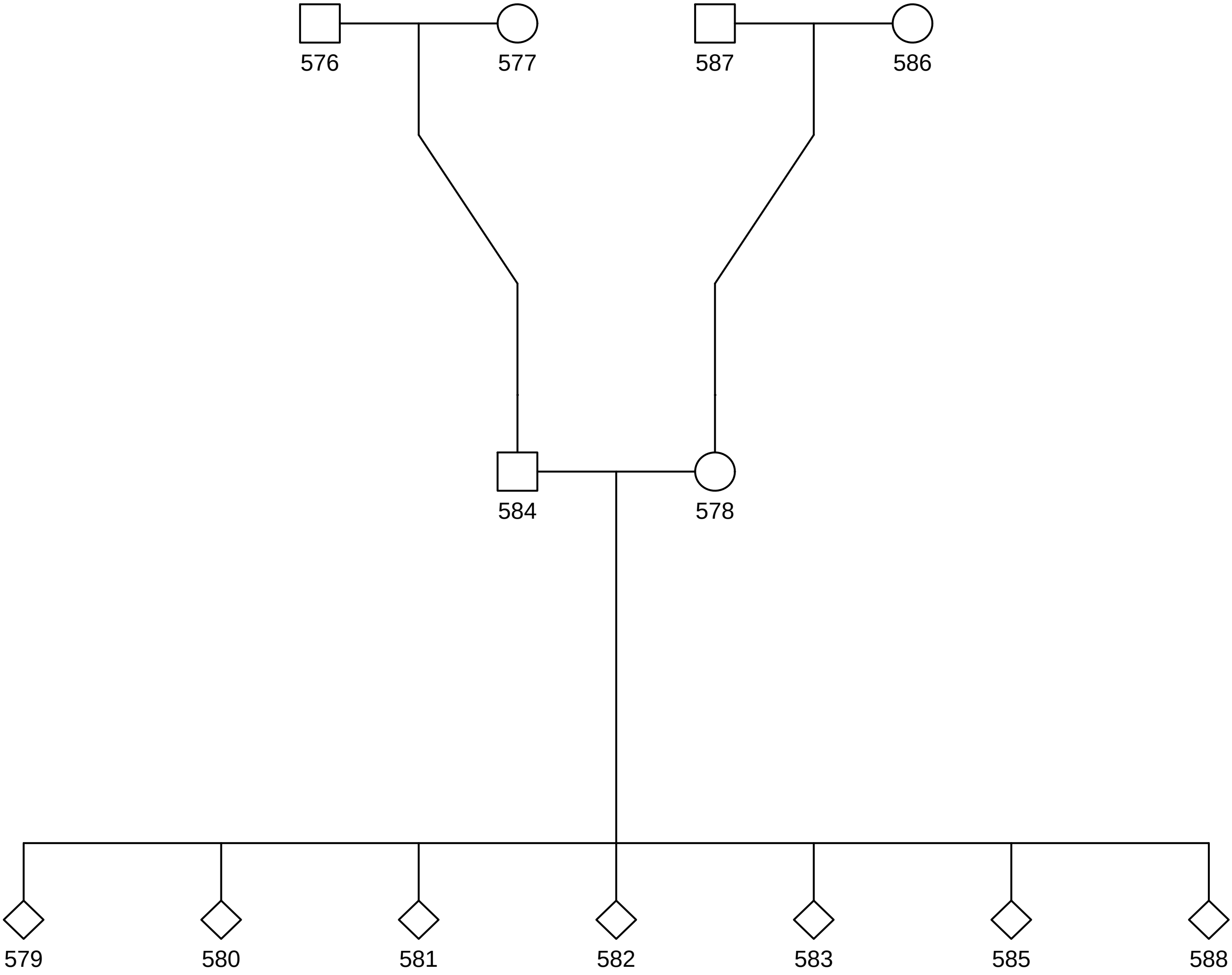

### 33.pdf

# 33

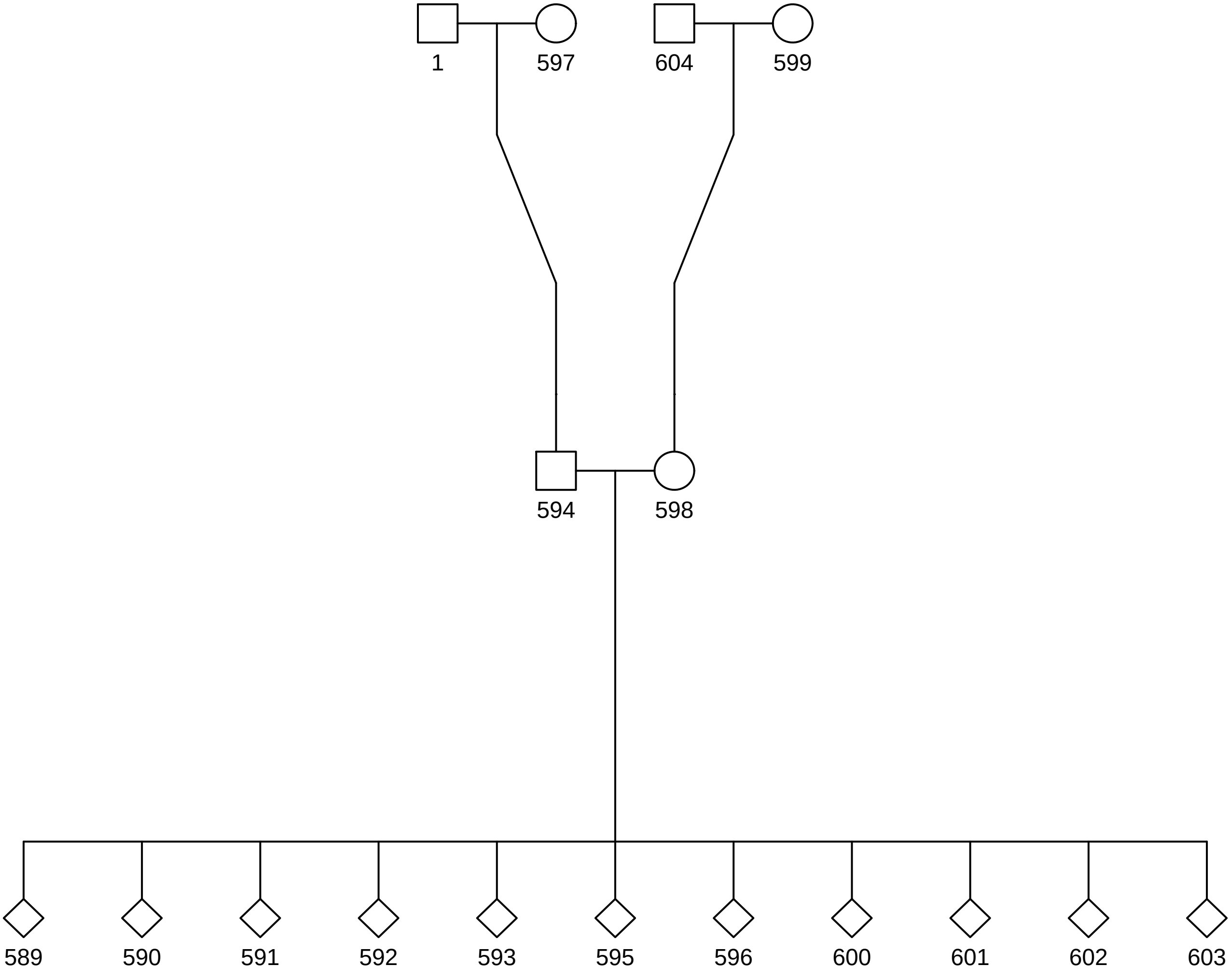

### chr1_1,142,234_1,142,274.png

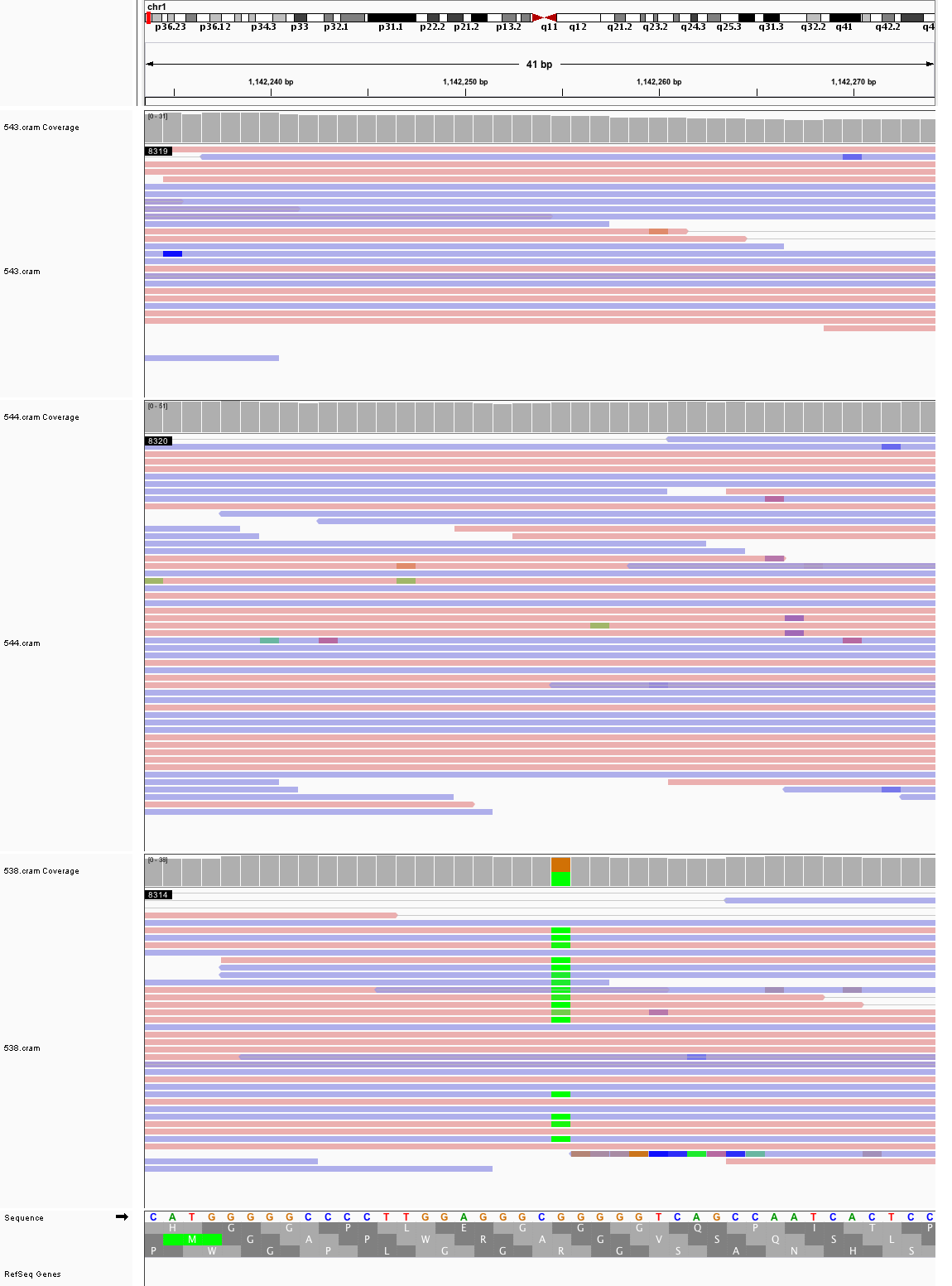

### chr1_8,217,448_8,217,488.png

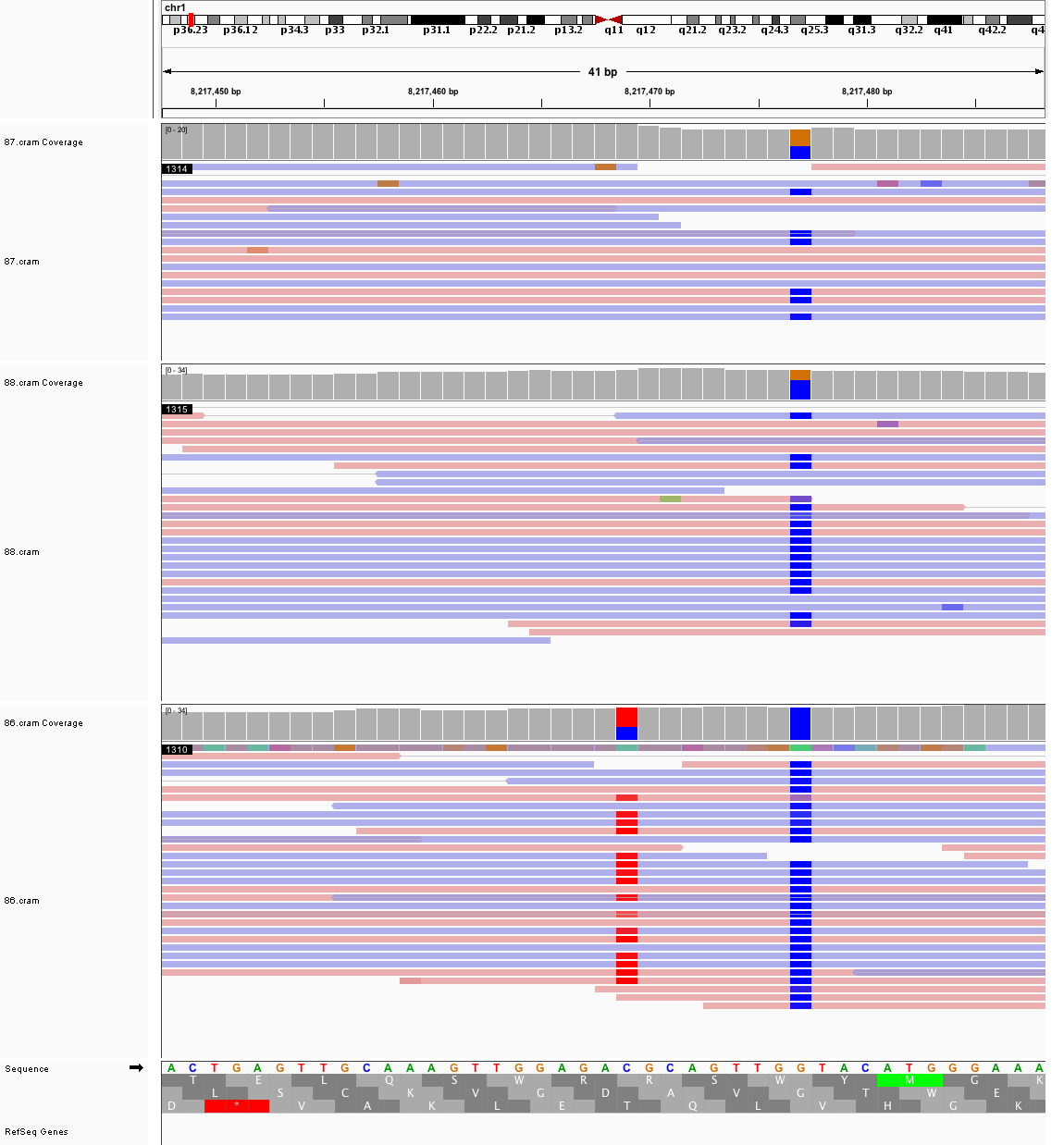

### chr1_75,351,486_75,351,526.png

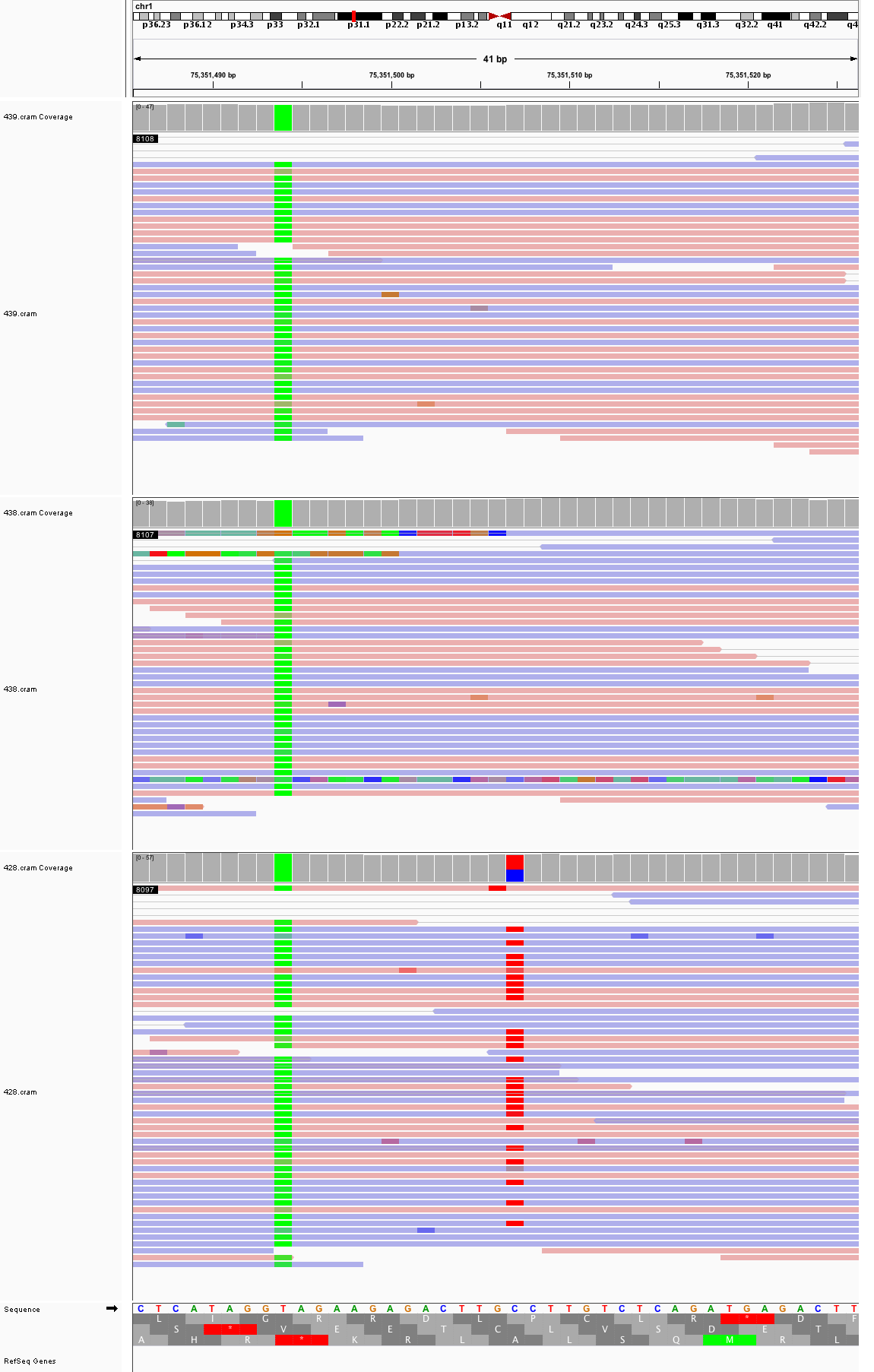

### chr1_85,664,265_85,664,305.png

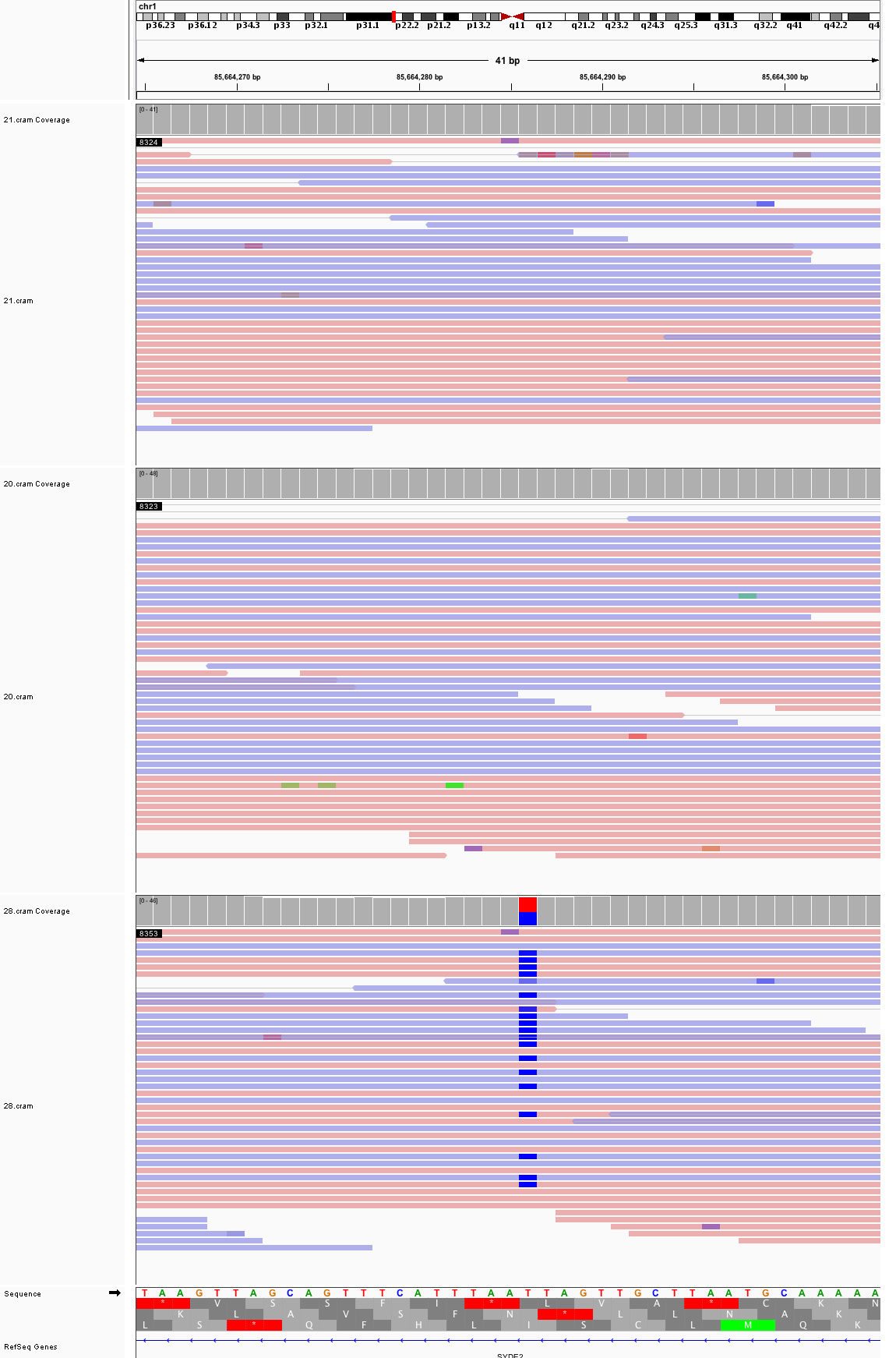

### chr1_143,877,217_143,877,257.png

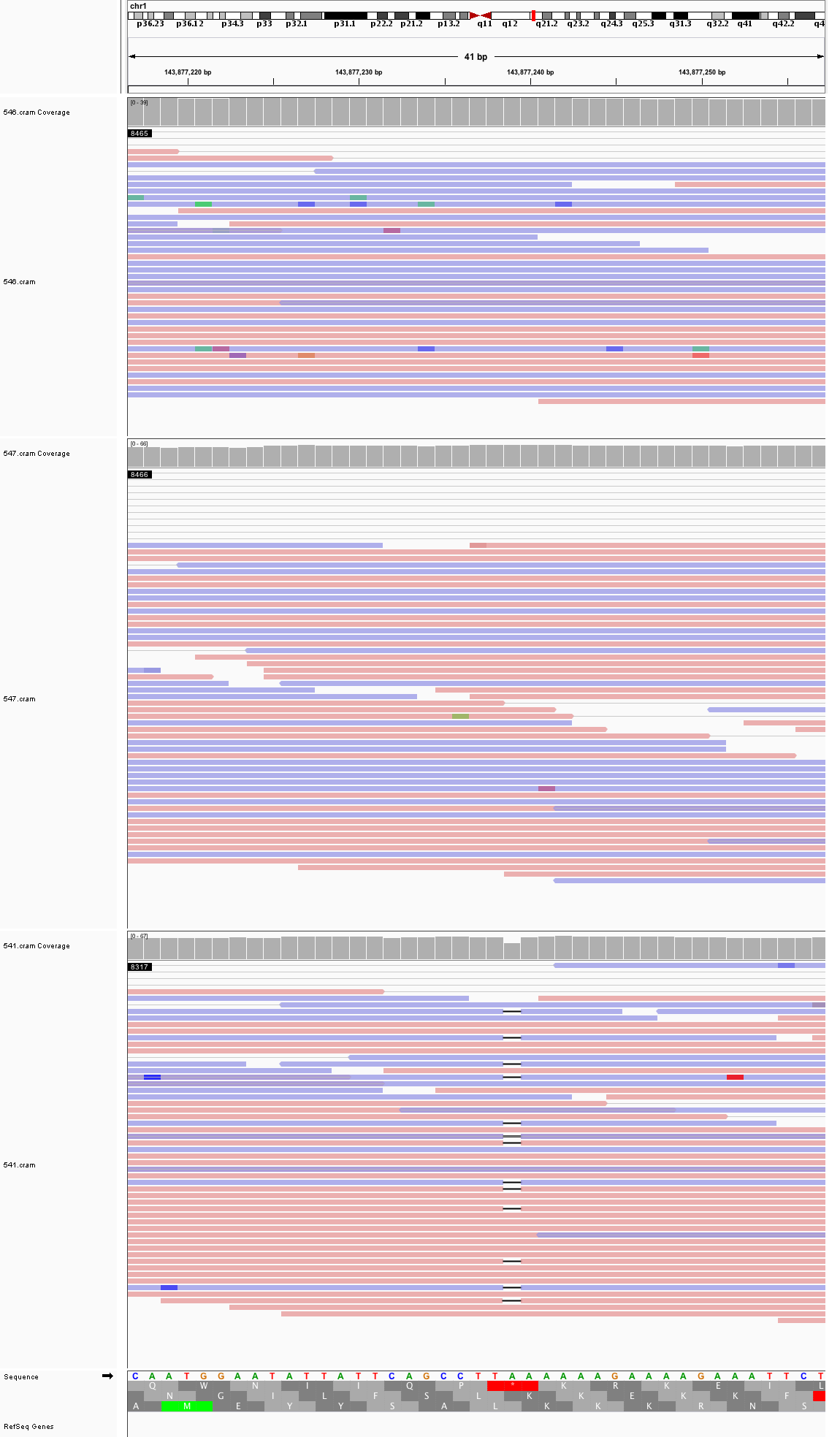

### chr1_191,935,677_191,935,717.png

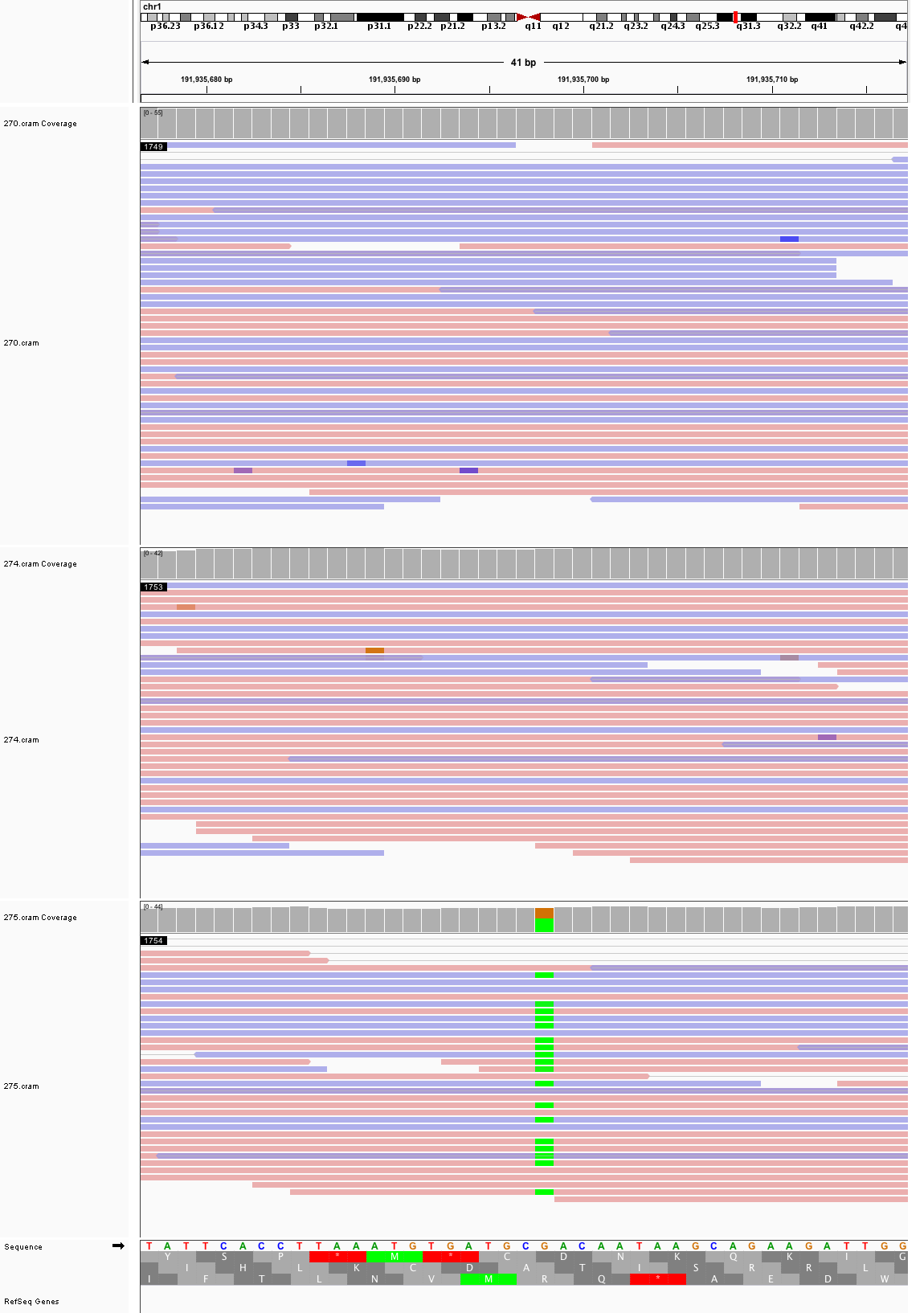

### chr1_234,581,349_234,581,389.png

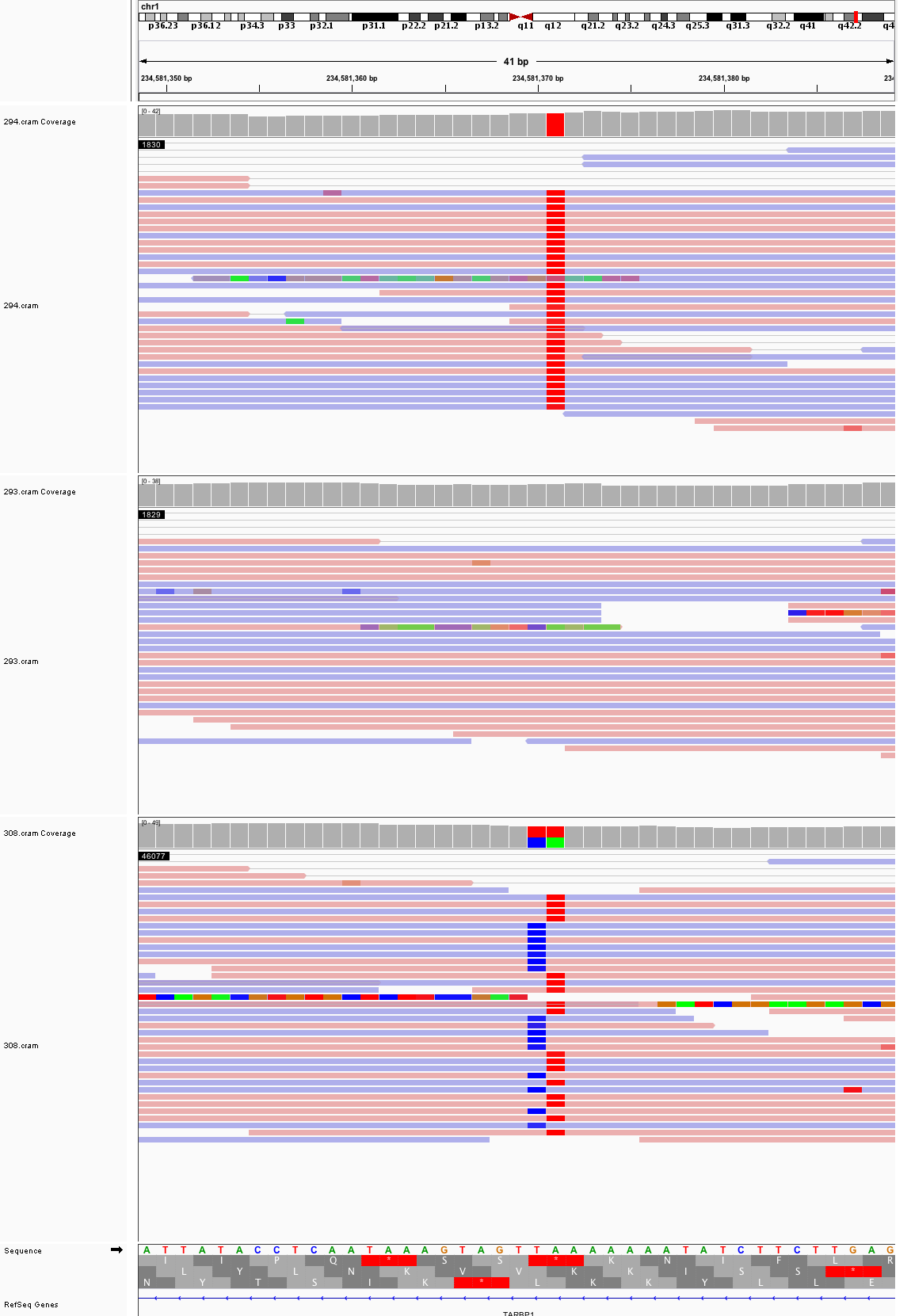

### chr2_9,729,471_9,729,511.png

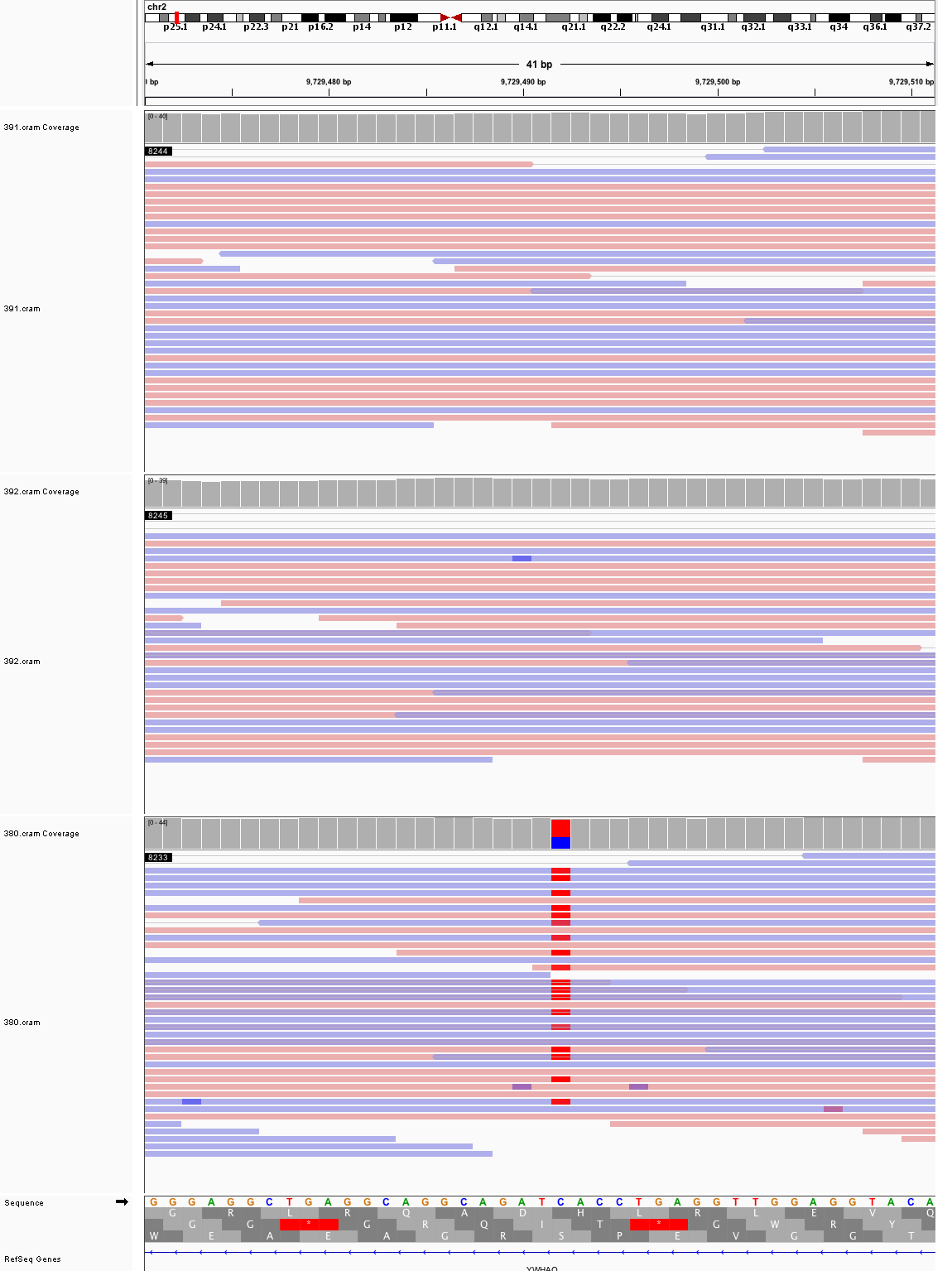

### chr2_39,865,281_39,865,321.png

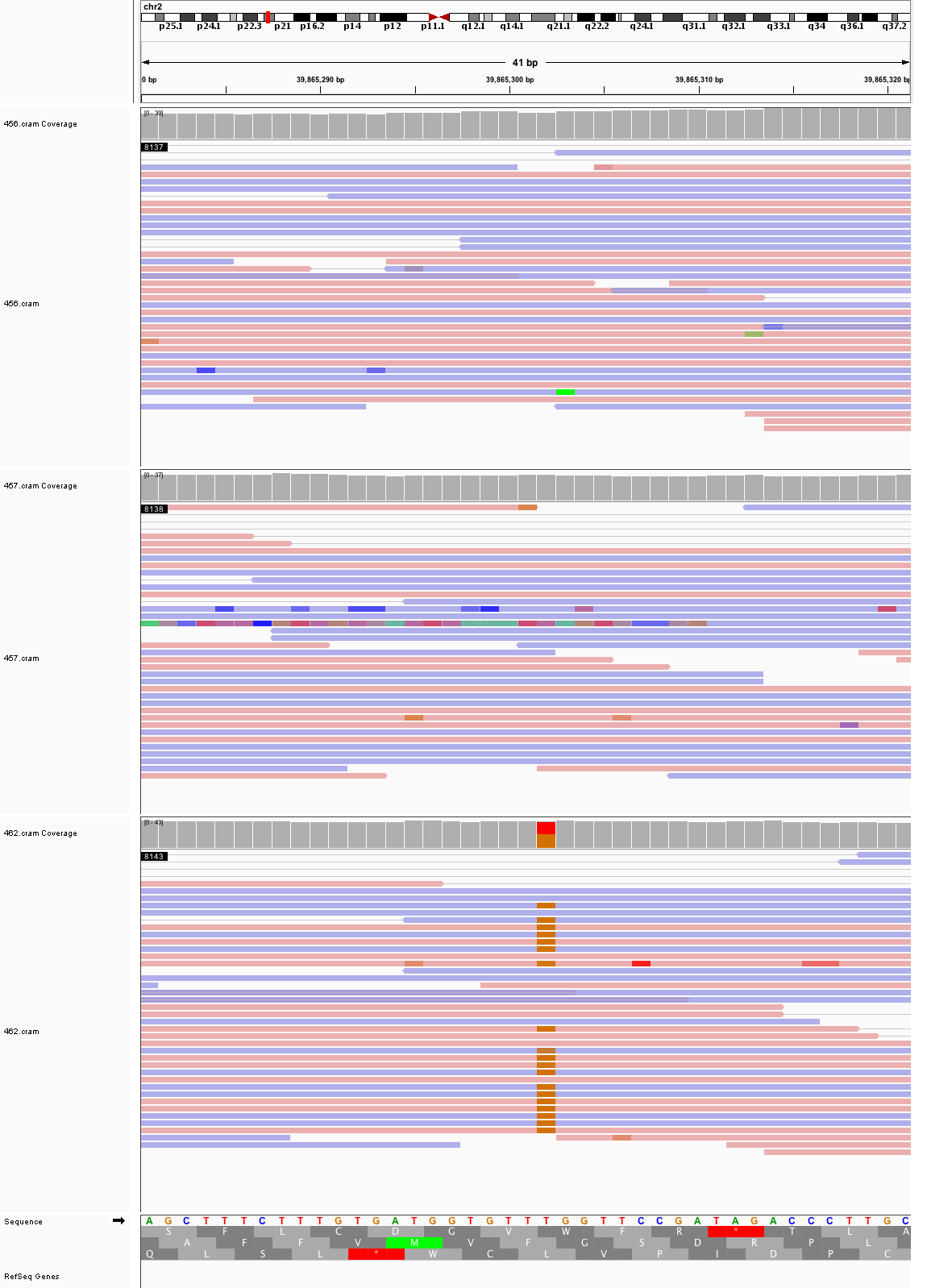

### chr2_117,430,371_117,430,411.png

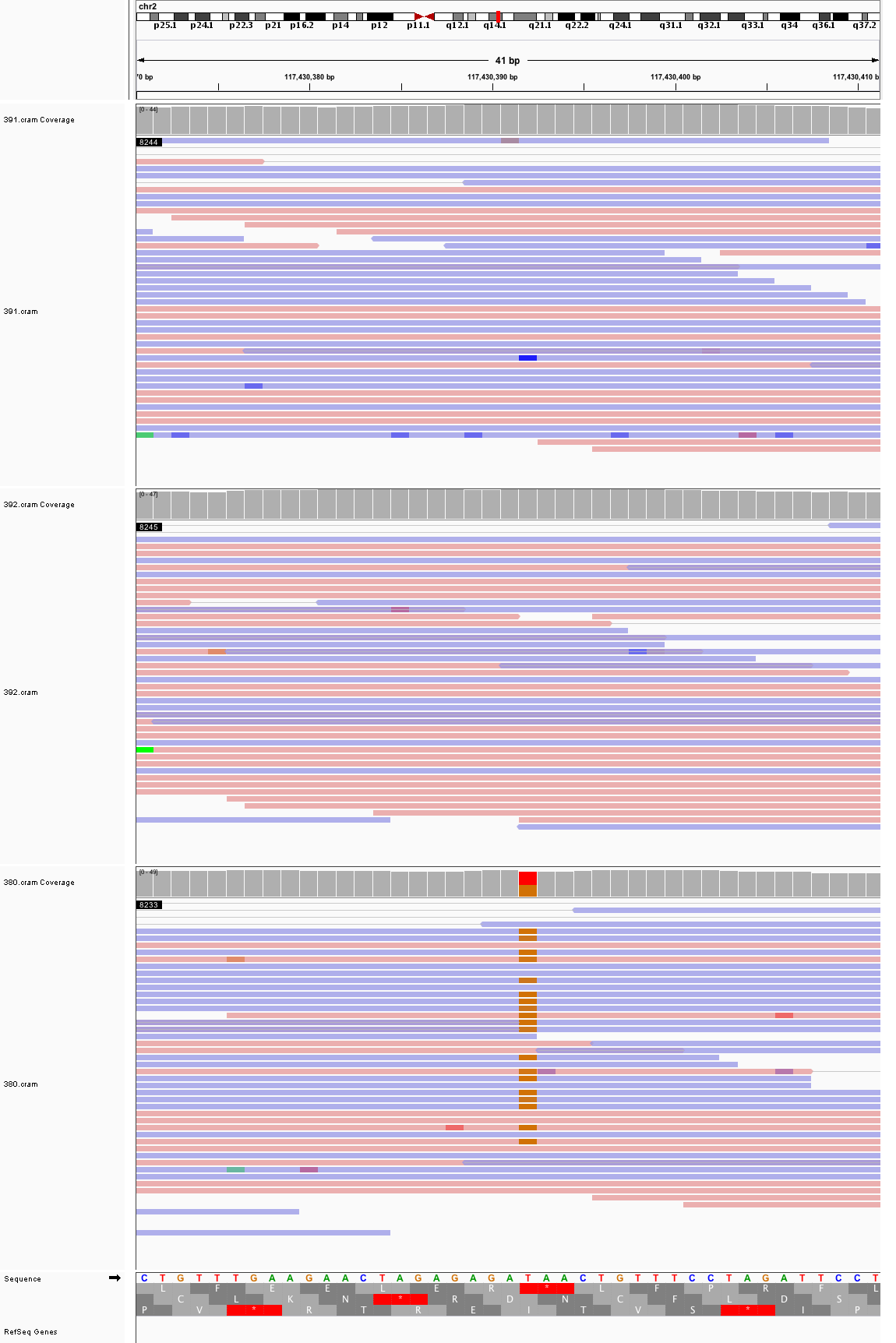

### chr2_118,776,384_118,776,424.png

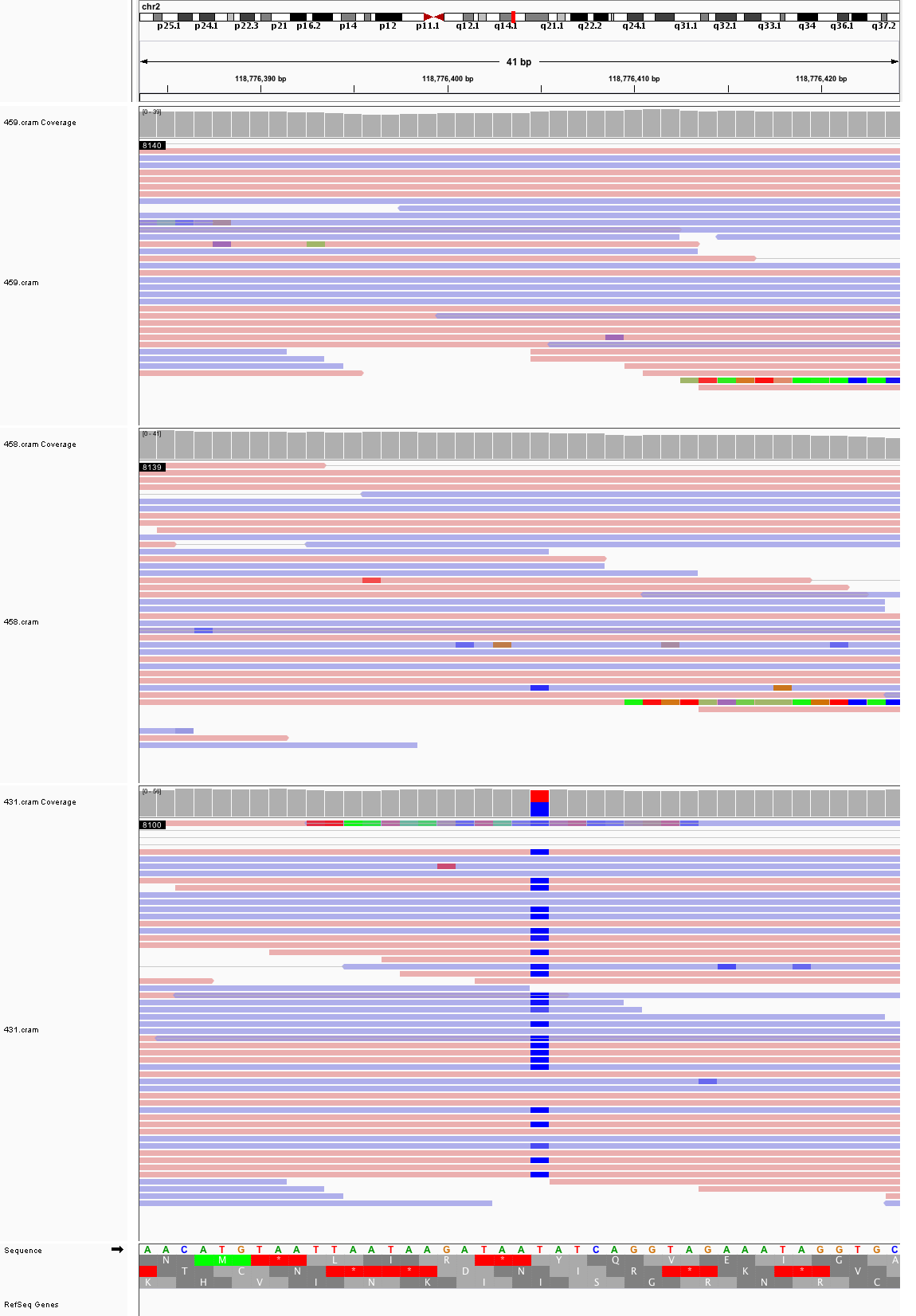

### chr2_190,266,379_190,266,419.png

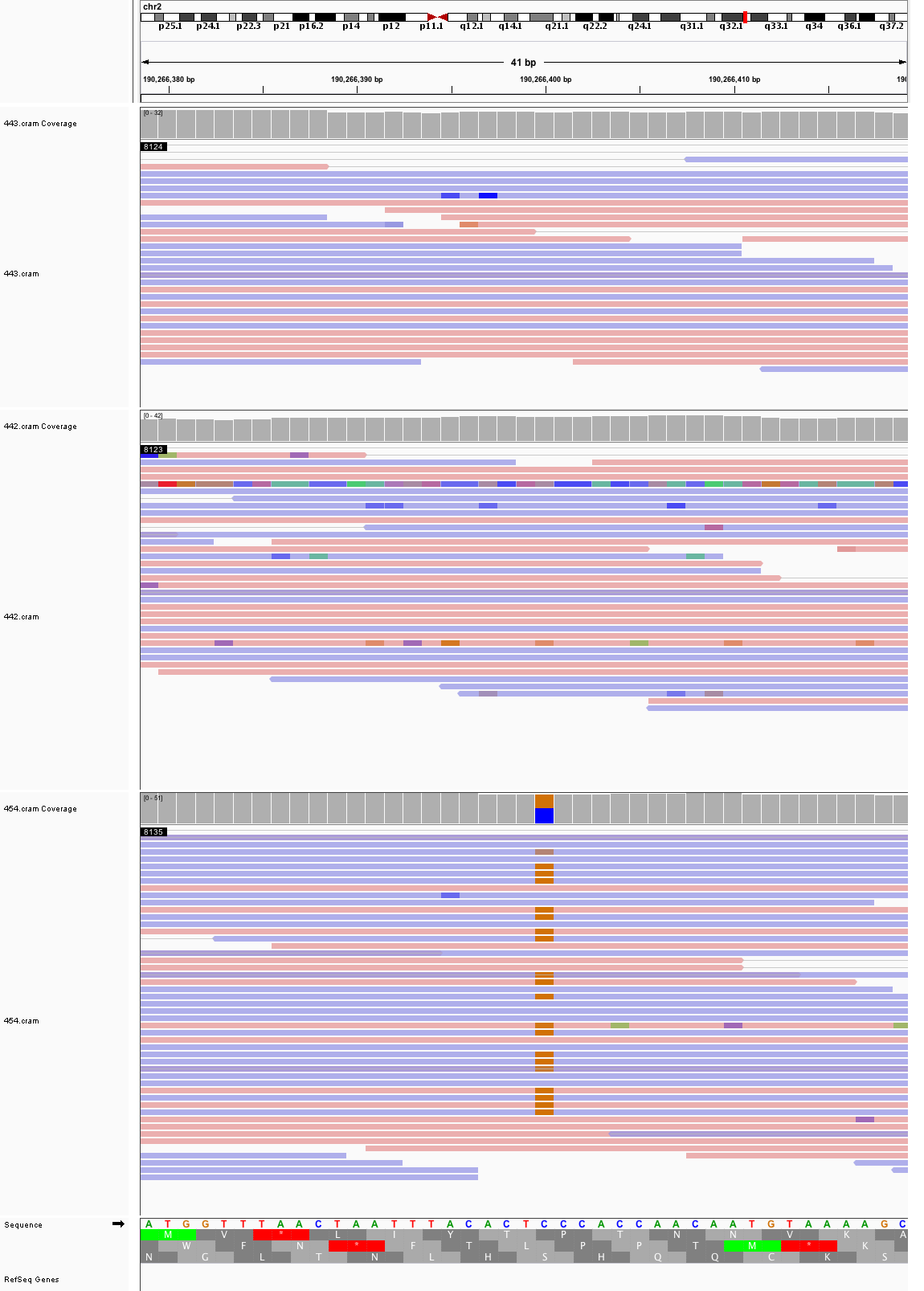

### chr2_207,078,587_207,078,627.png

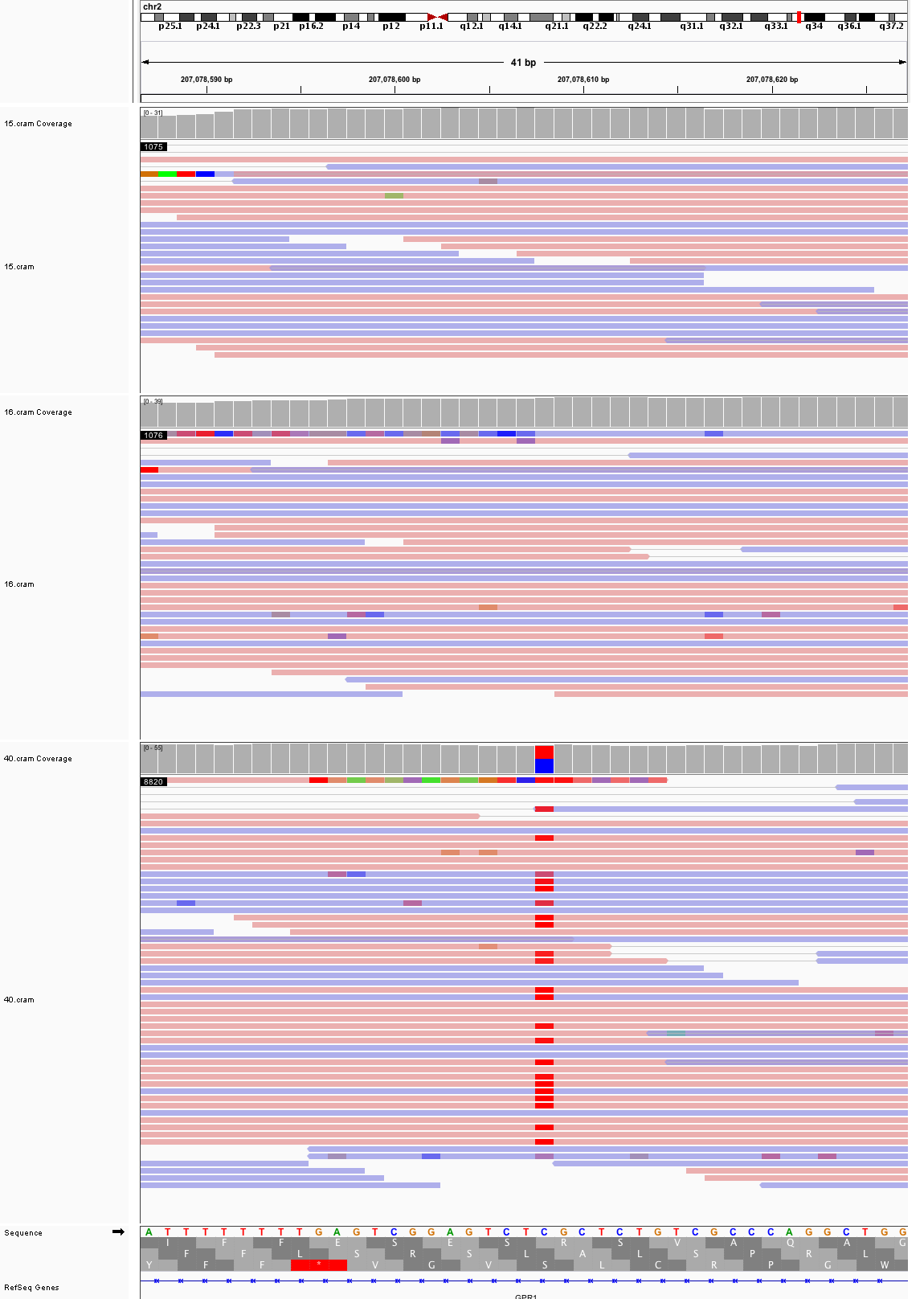

### chr2_220,502,391_220,502,431.png

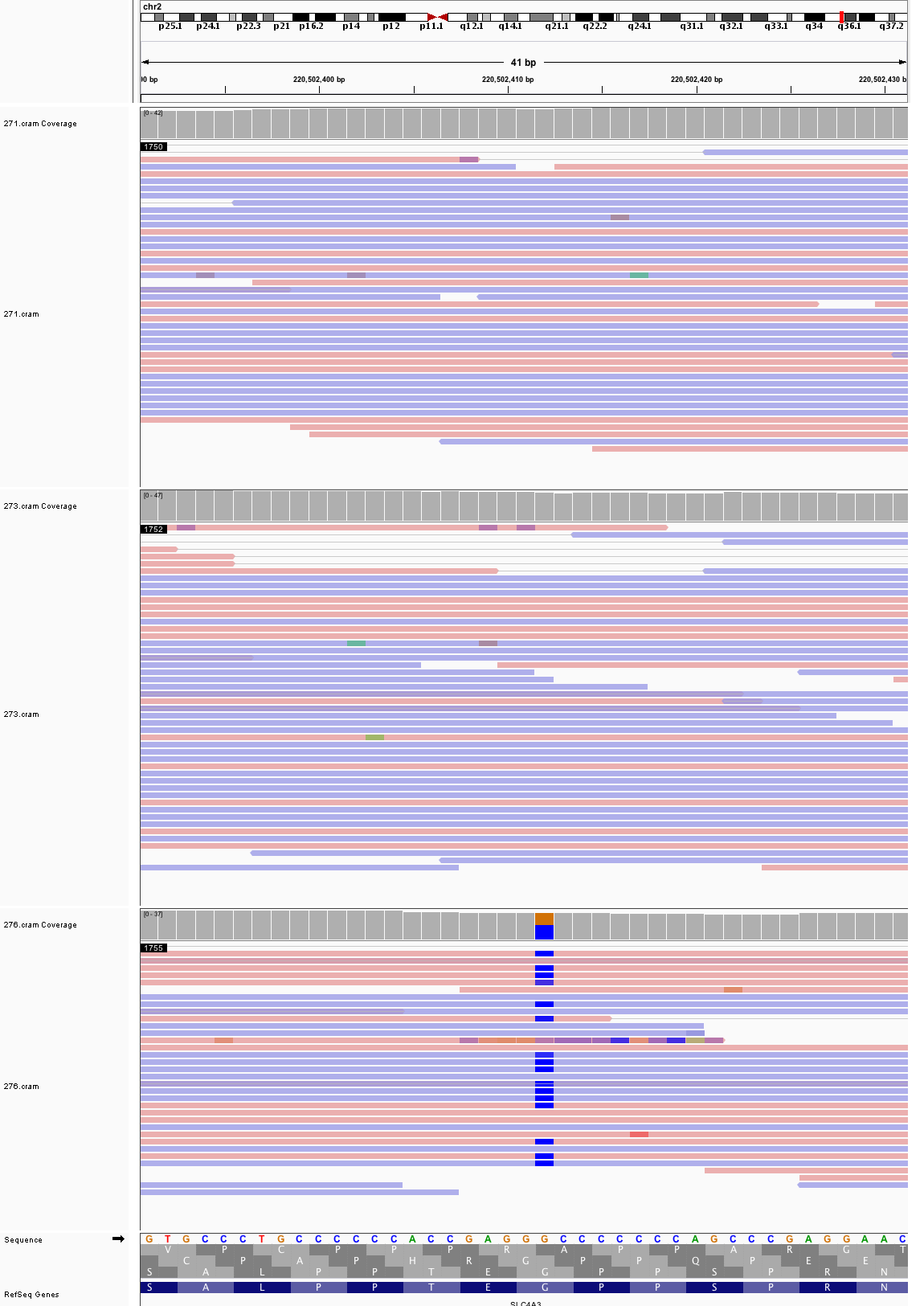

### chr3_34,394,482_34,394,522.png

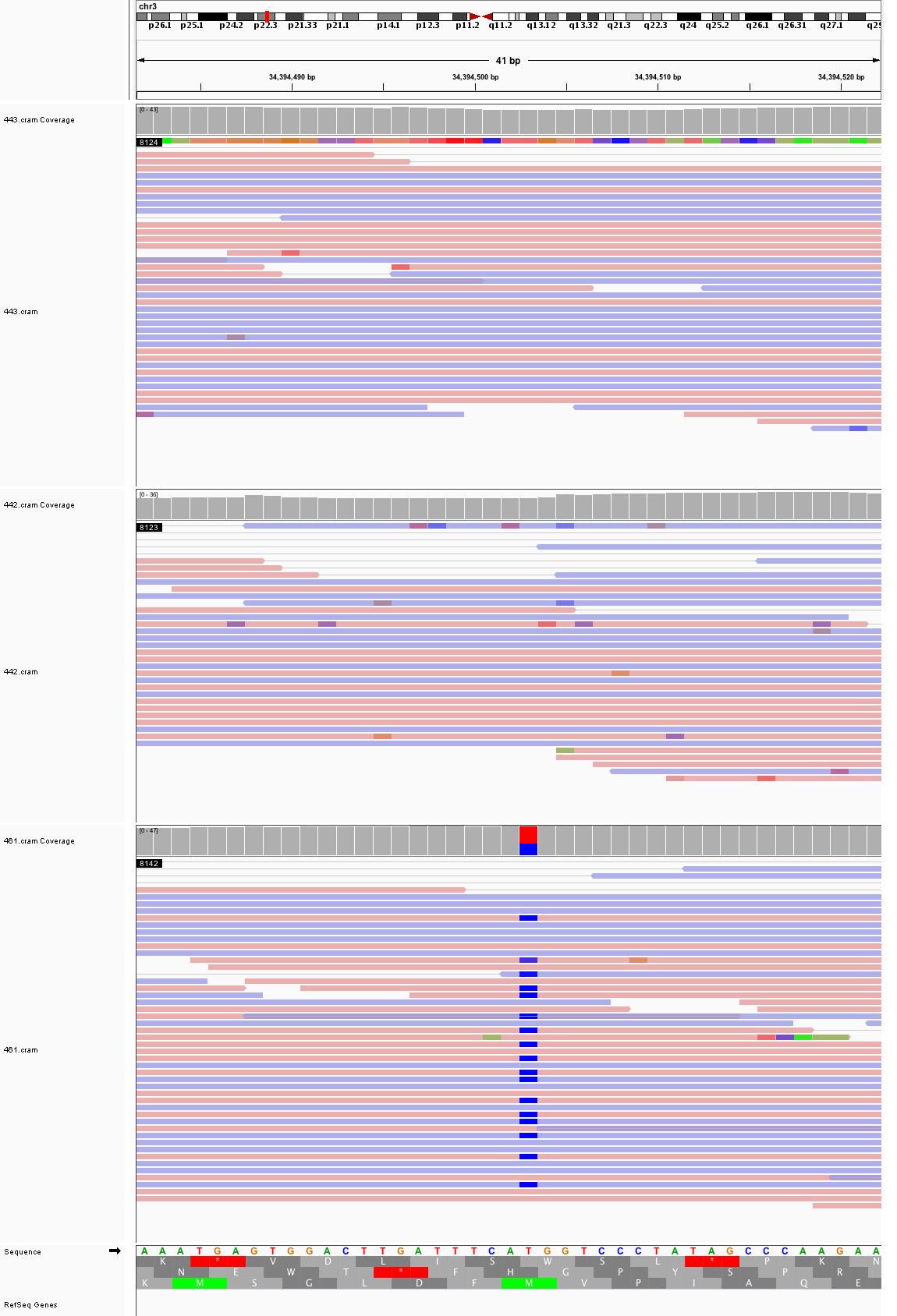

### chr3_40,972,849_40,972,889.png

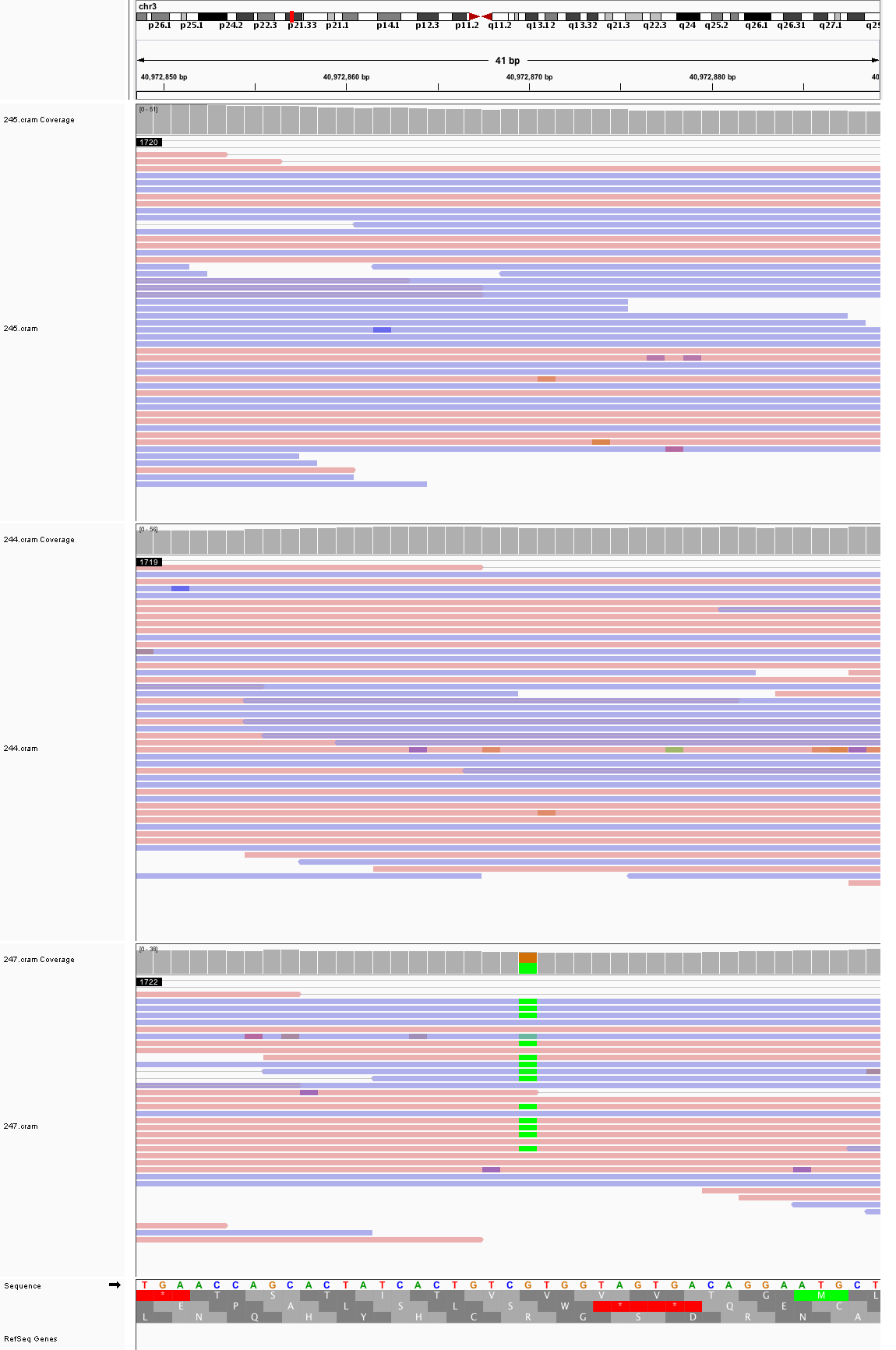

### chr3_42,918,874_42,918,914.png

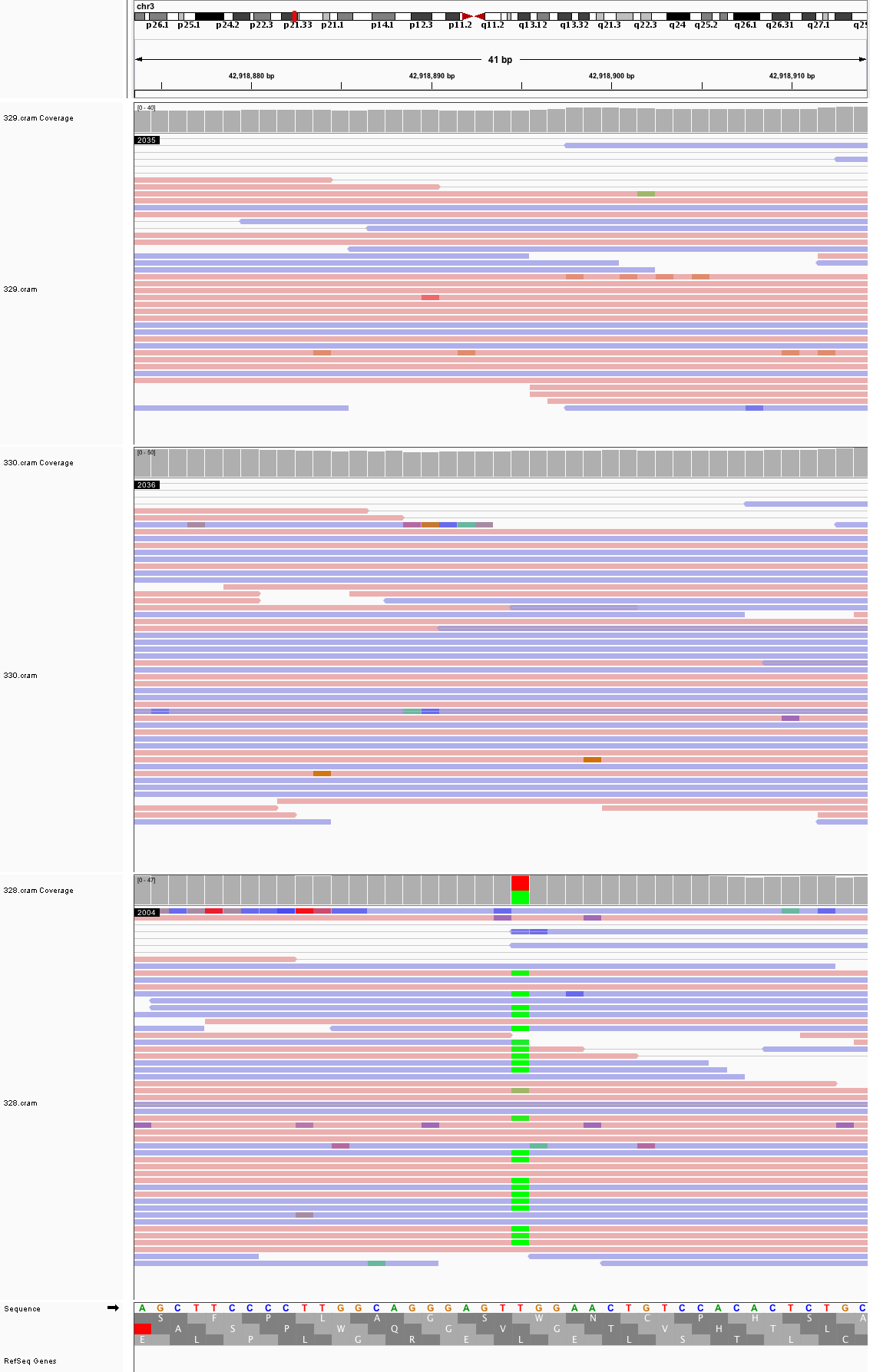

### chr3_80,428,845_80,428,885.png

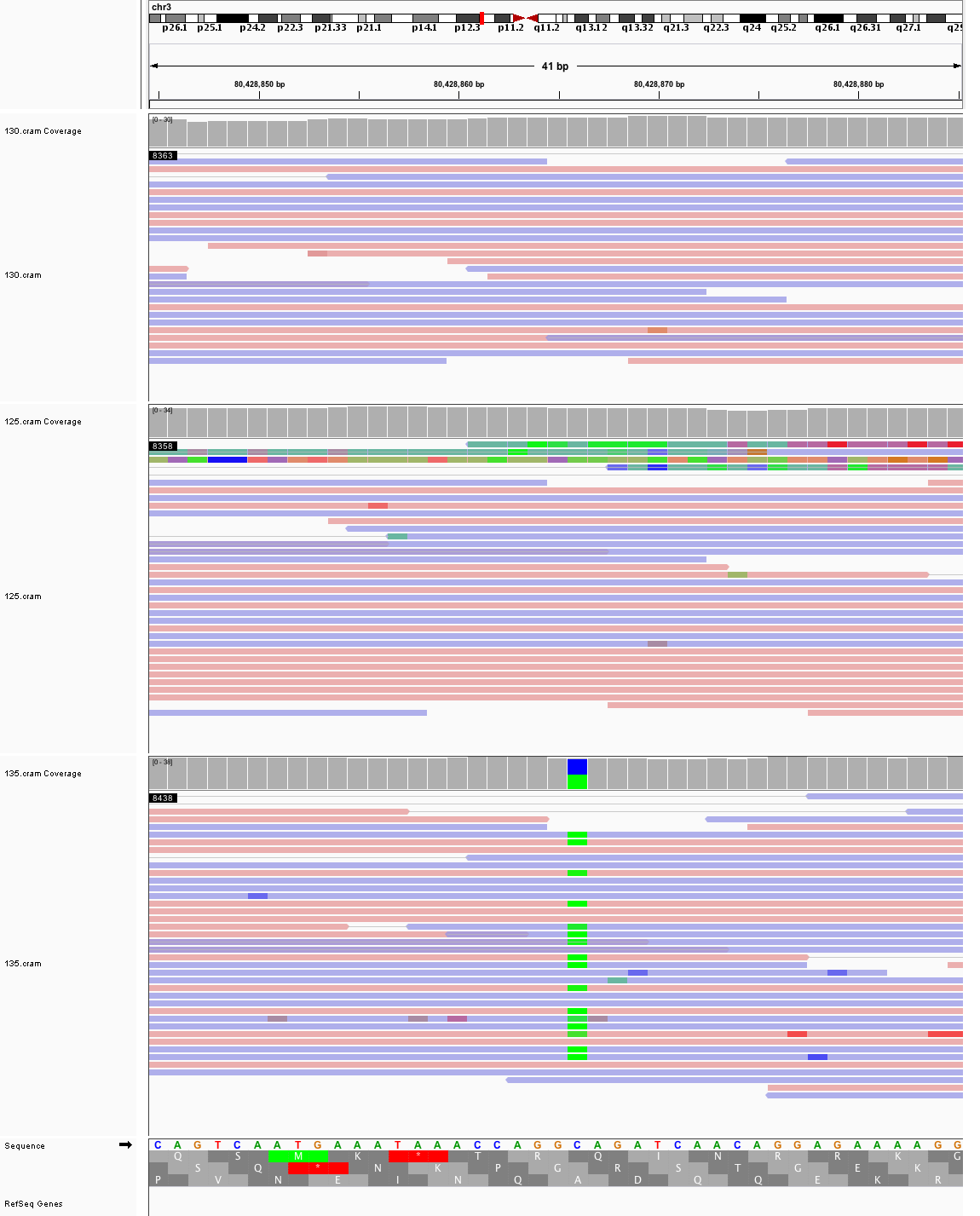

### chr3_97,442,181_97,442,221.png

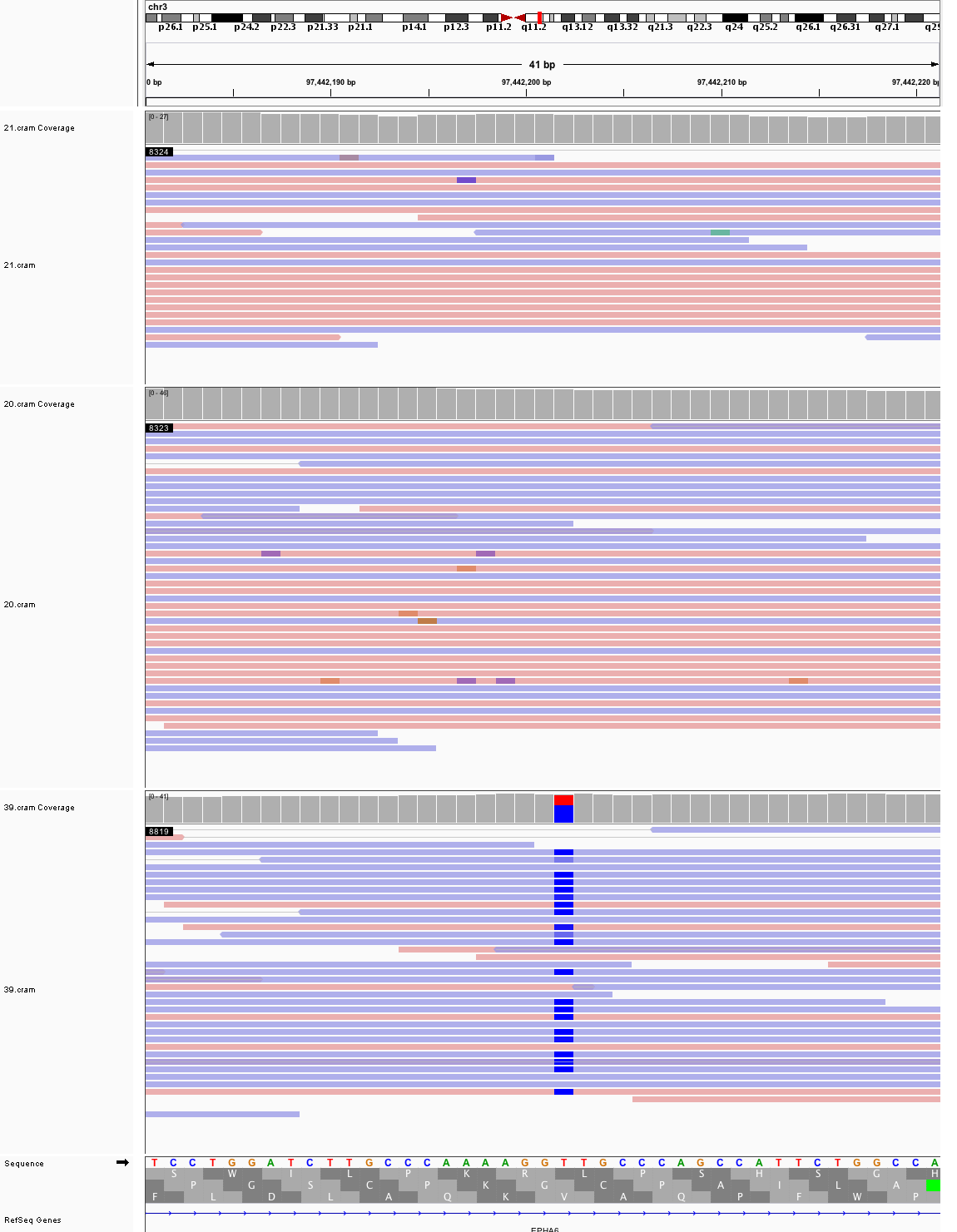

### chr3_175,698,258_175,698,298.png

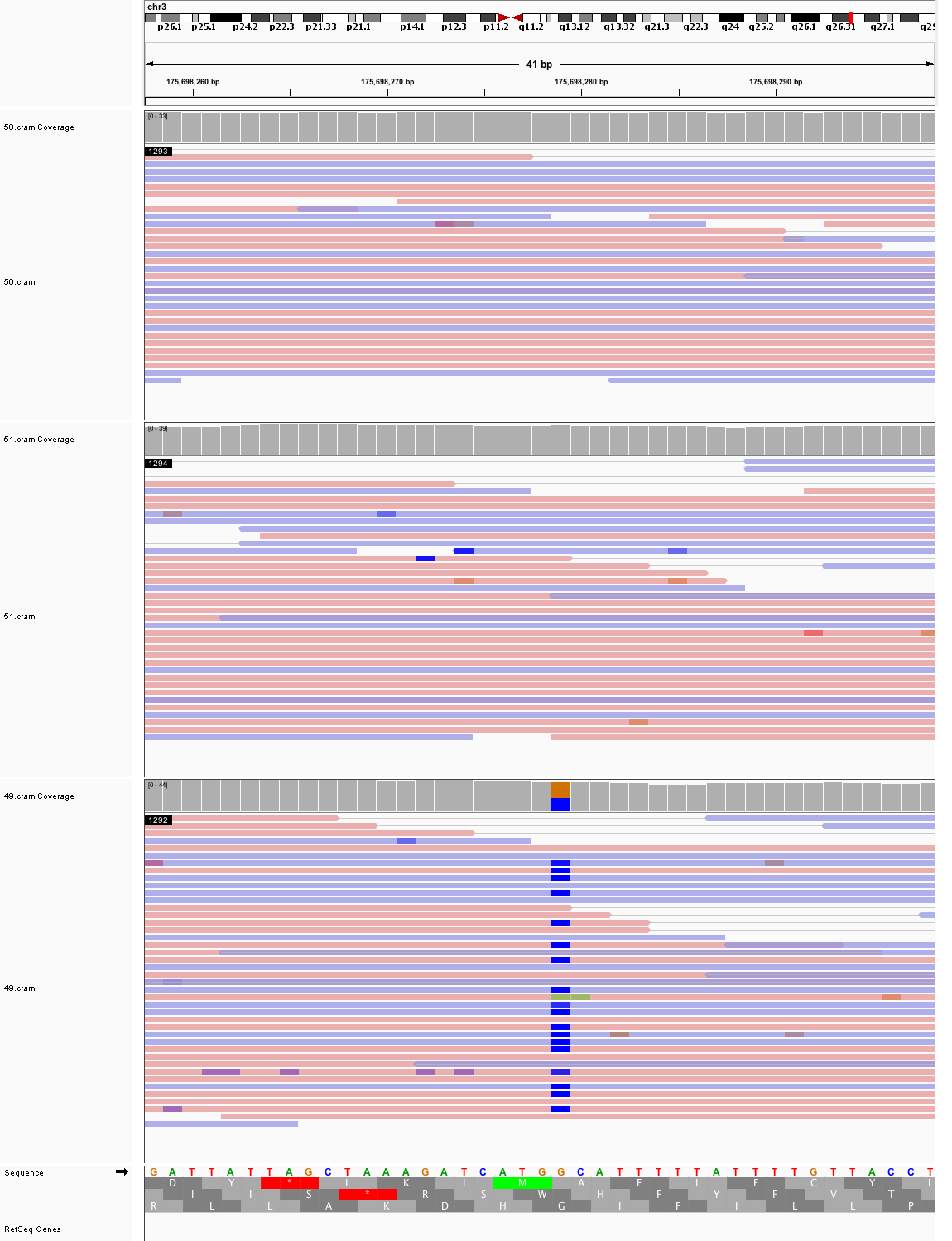

### chr3_195,829,161_195,829,201.png

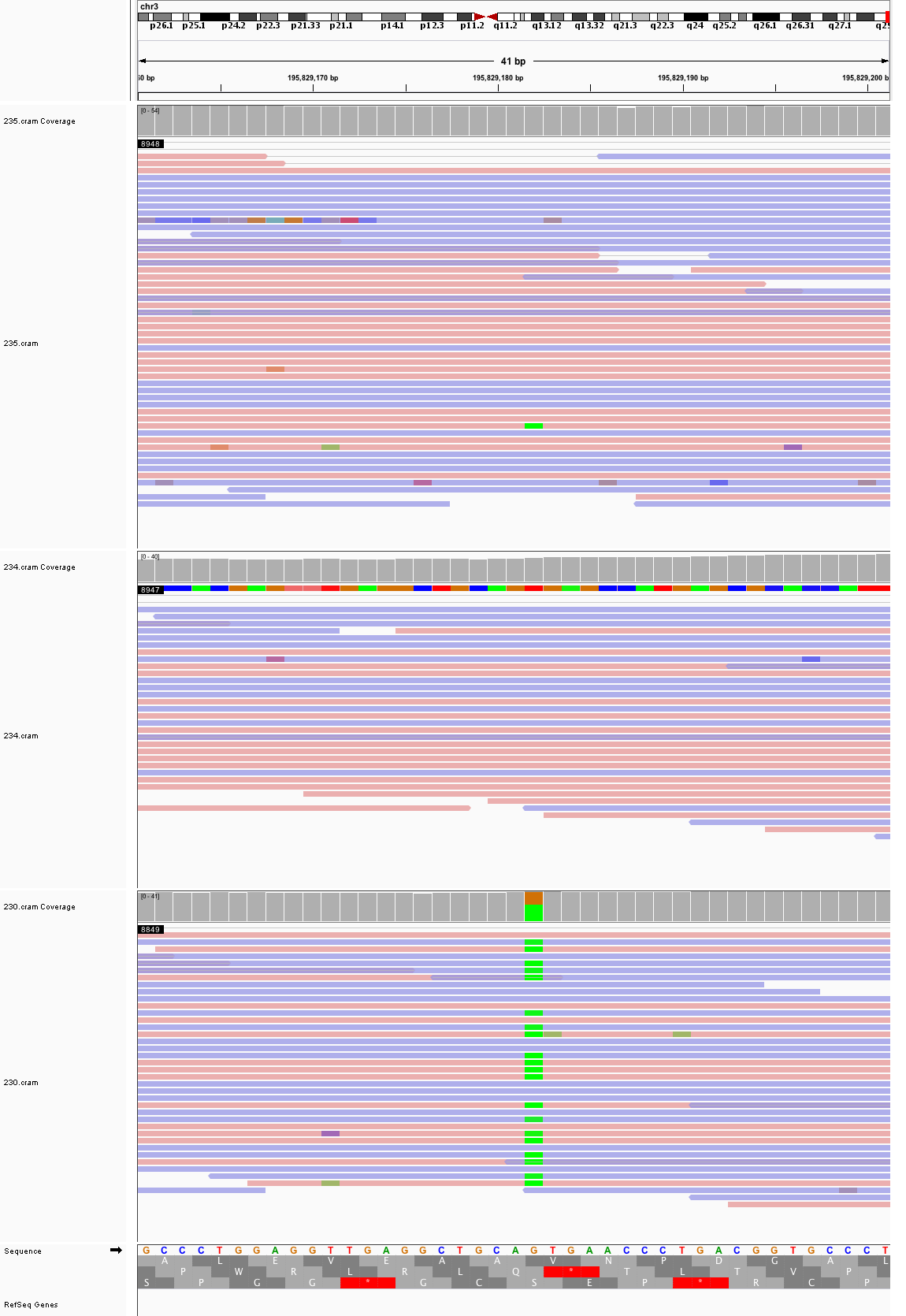

### chr4_1,937,913_1,937,953.png

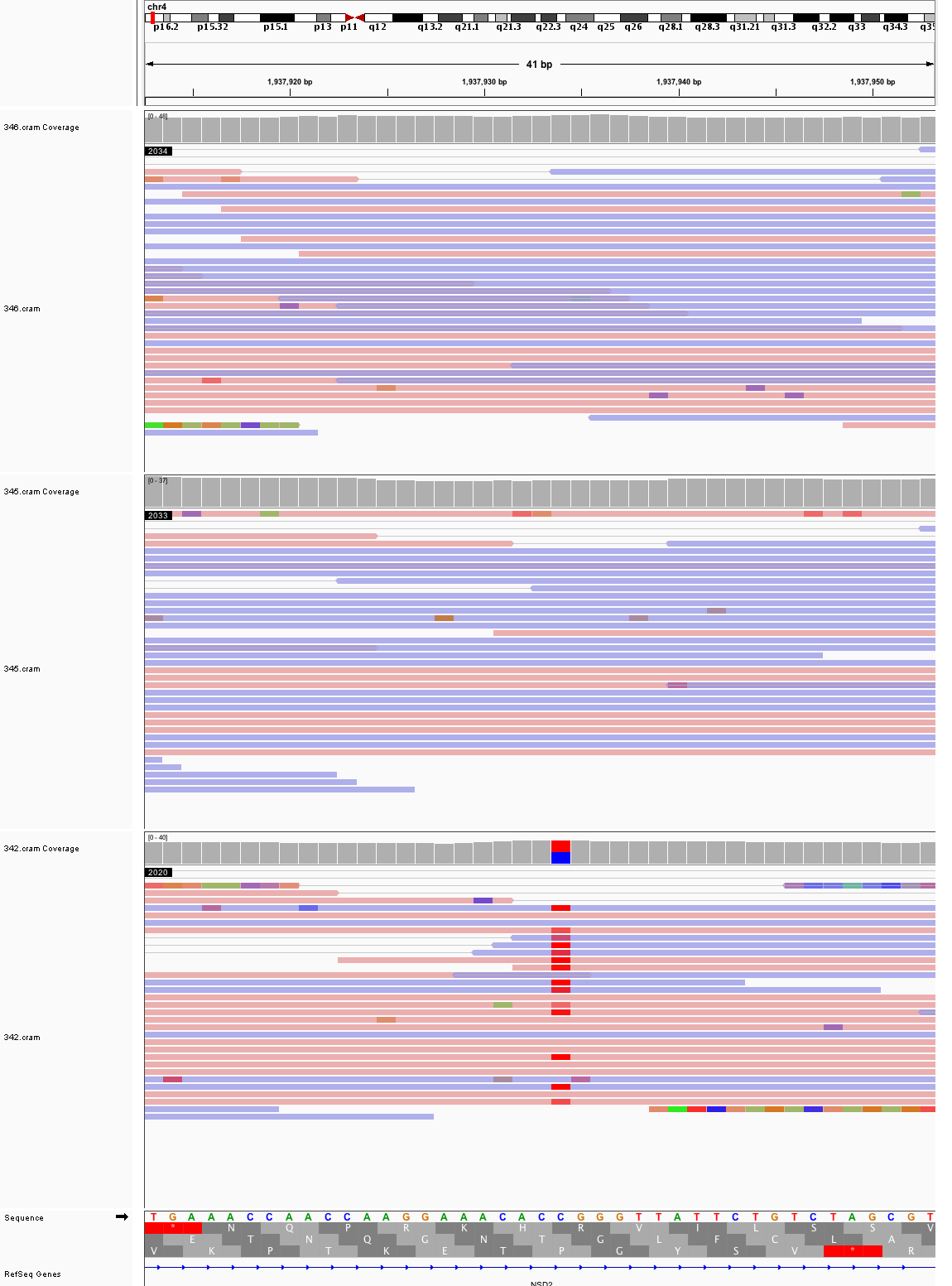

### chr4_7,710,300_7,710,340.png

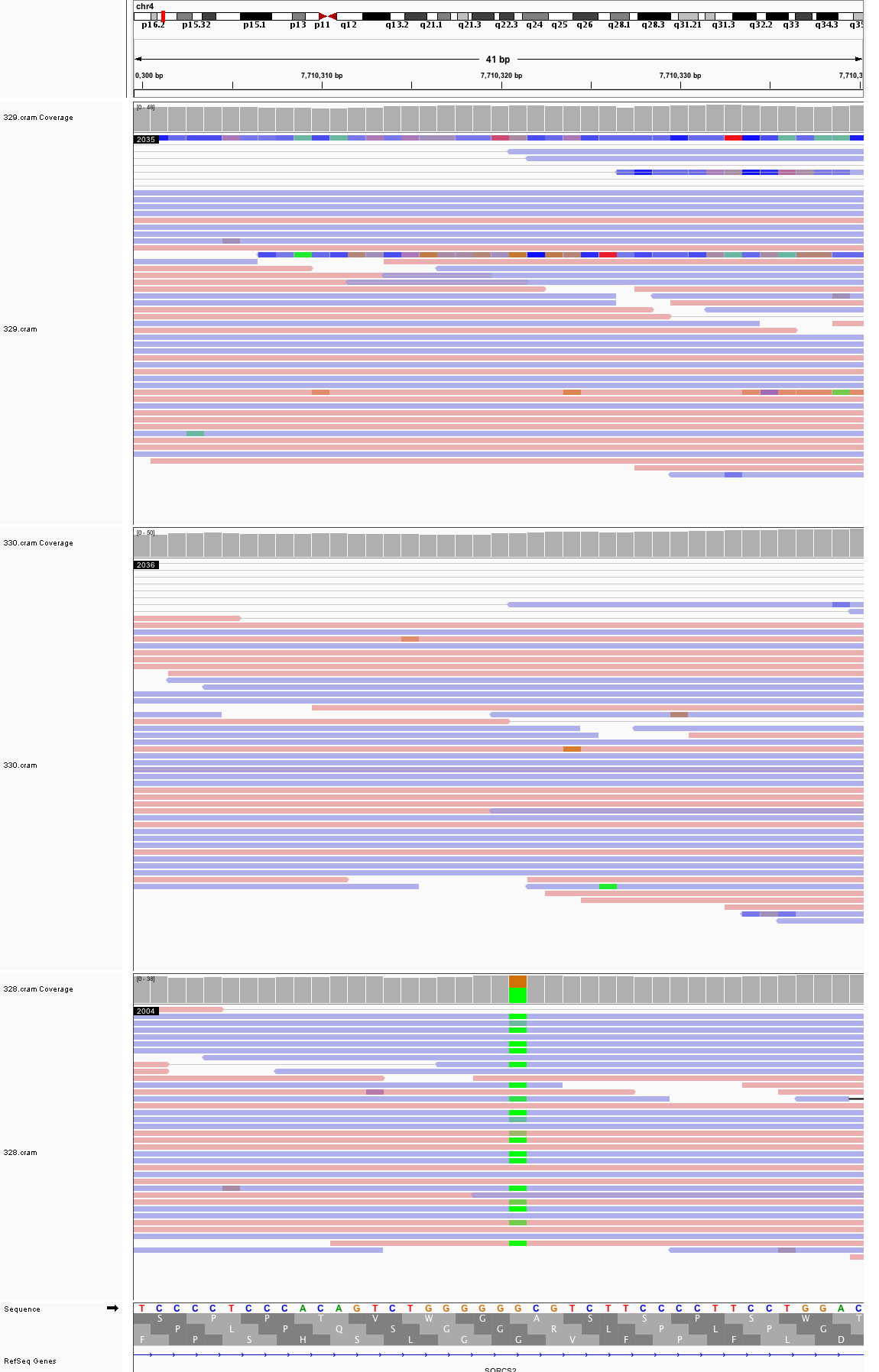

### chr4_22,961,833_22,961,873.png

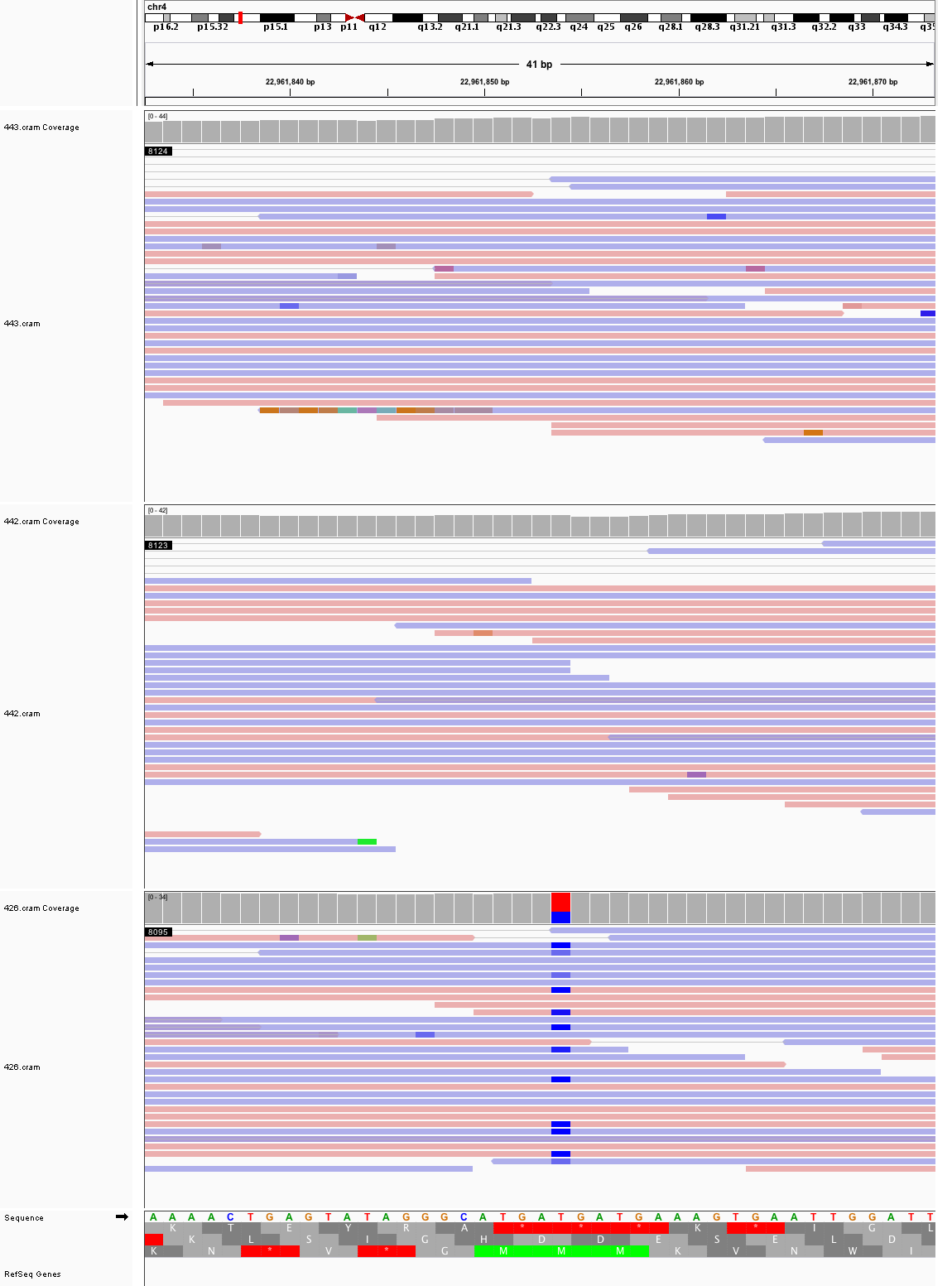

### chr4_23,189,014_23,189,054.png

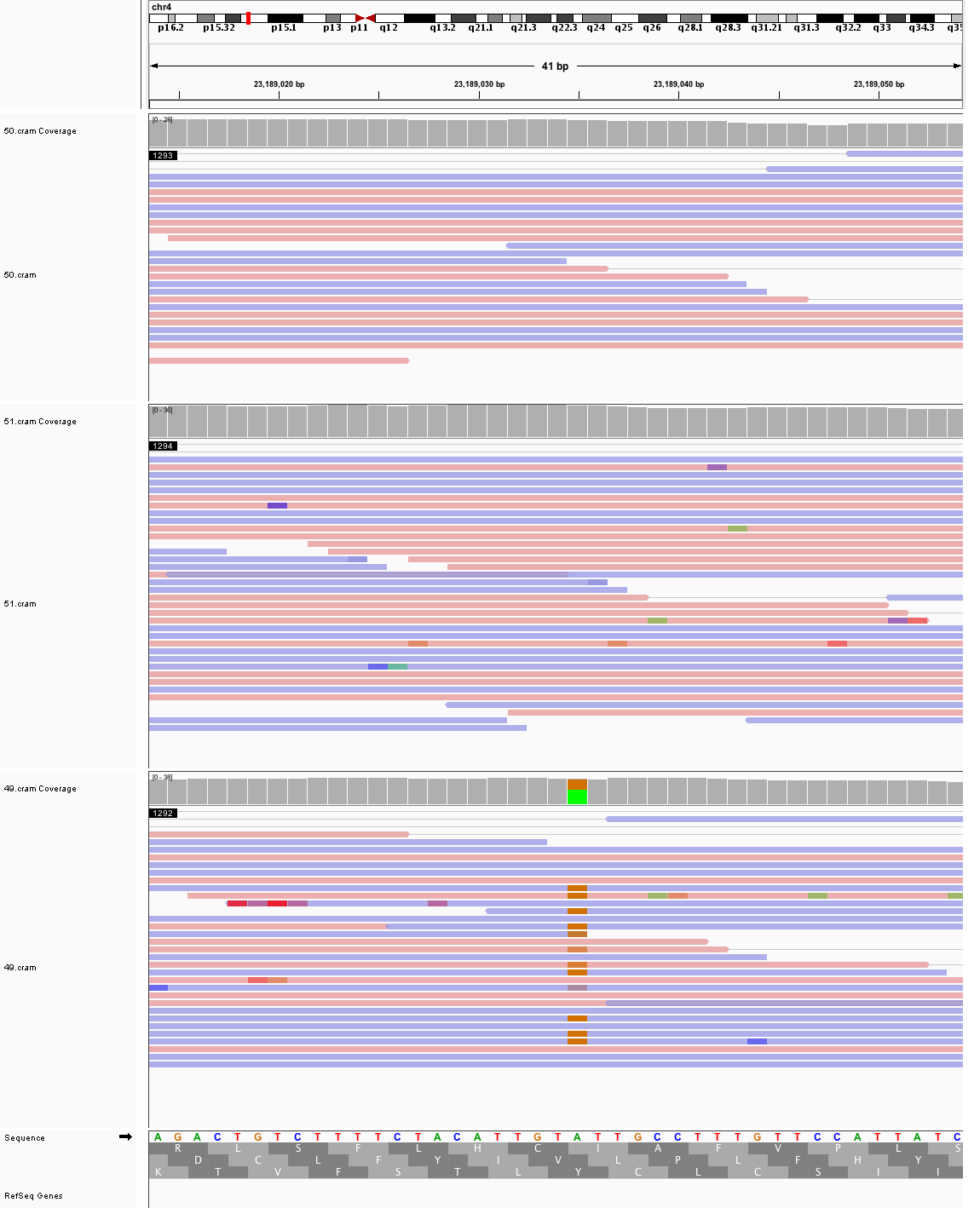

### chr4_57,950,071_57,950,111.png

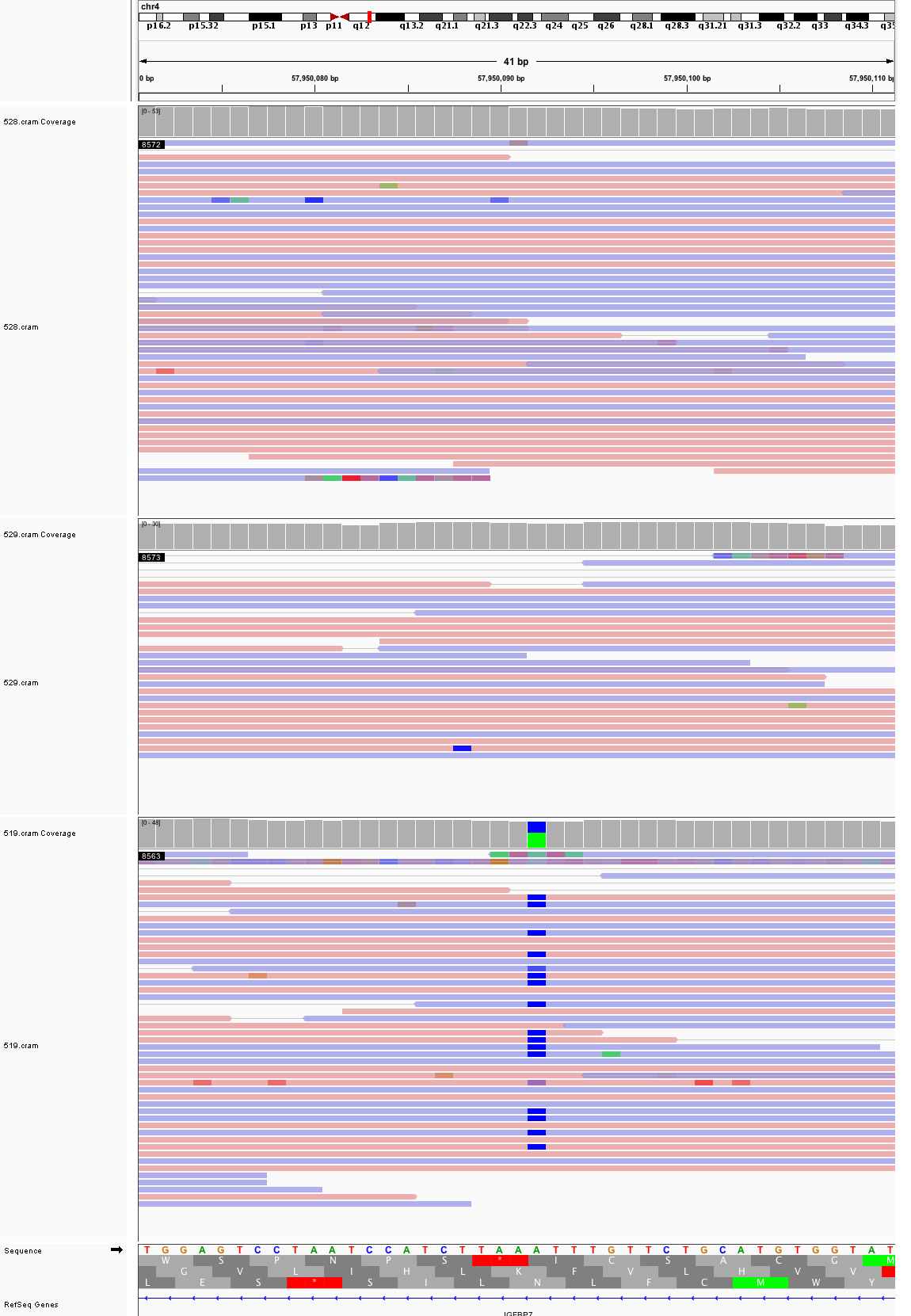

### chr5_24,249,187_24,249,227.png

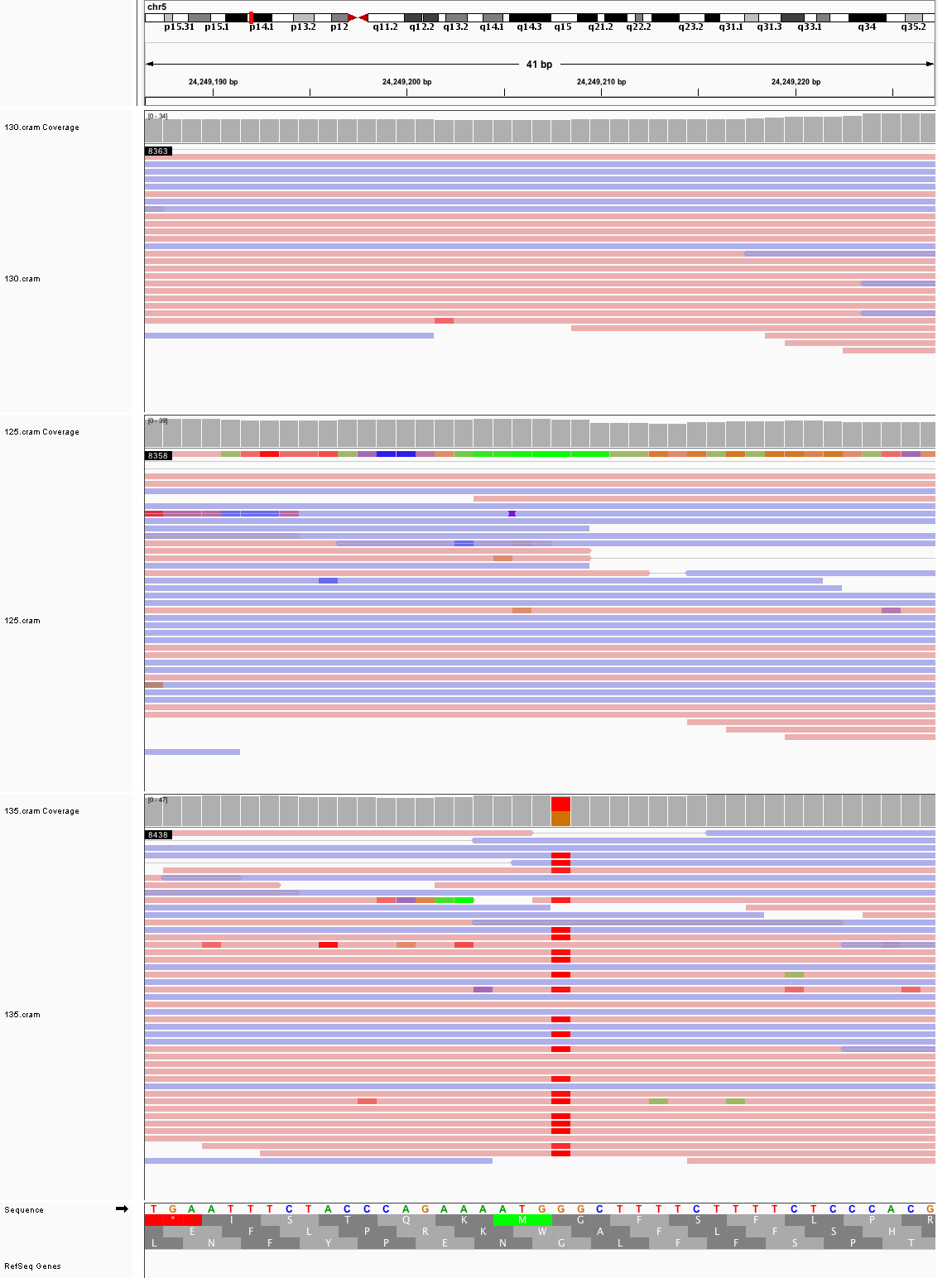

### chr5_25,565,330_25,565,370.png

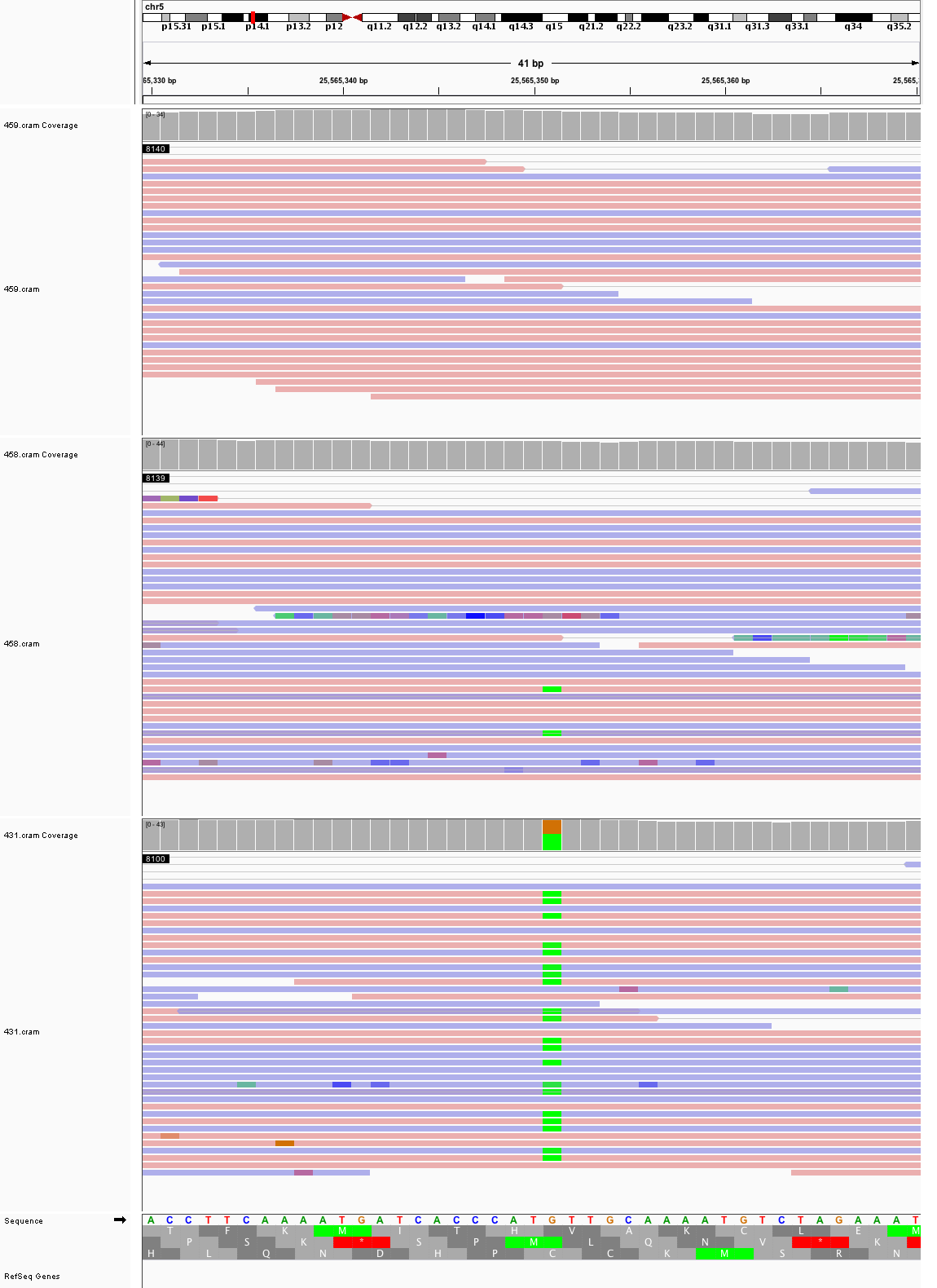

### chr5_103,367,331_103,367,371.png

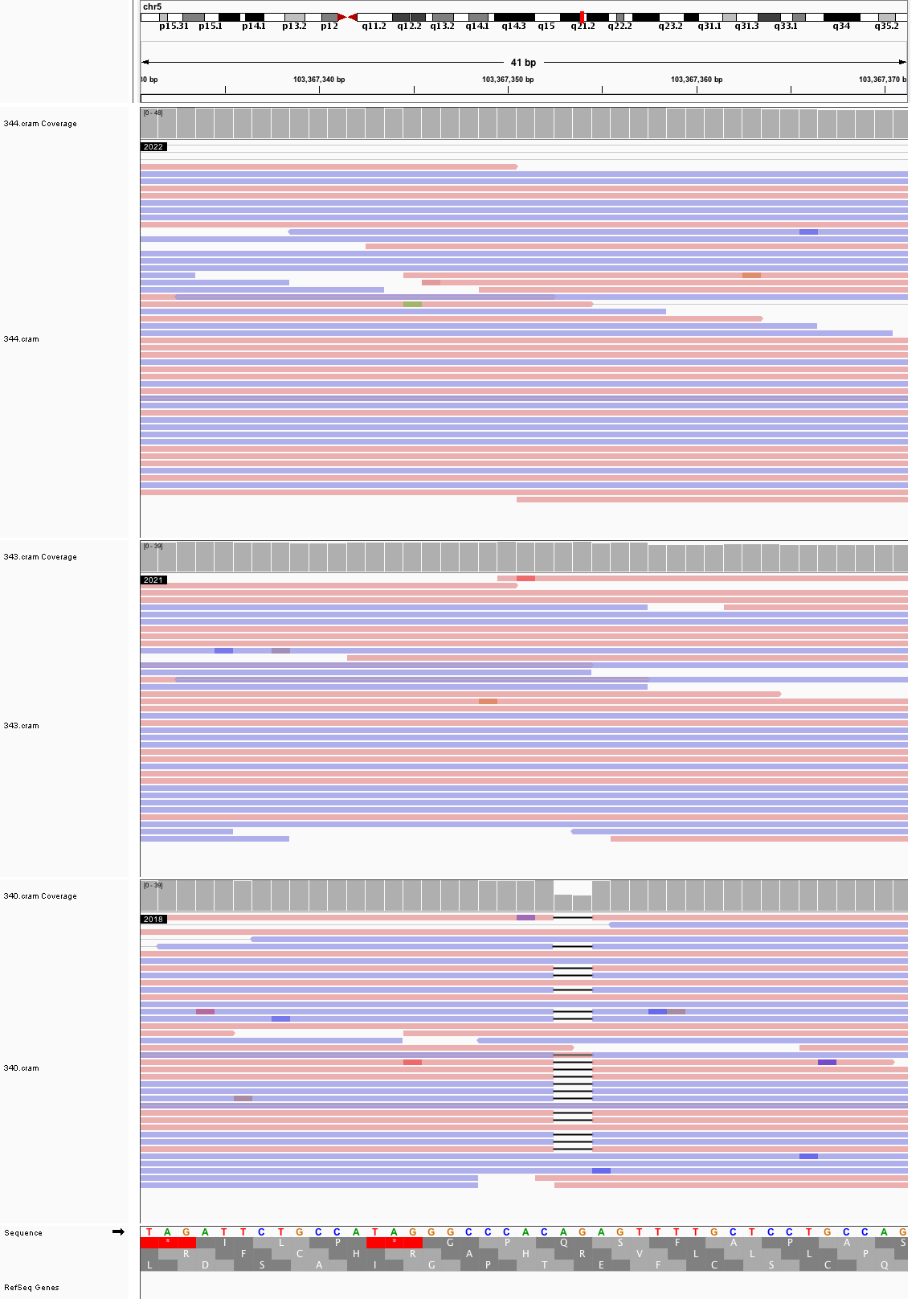

### chr6_3,483,409_3,483,449.png

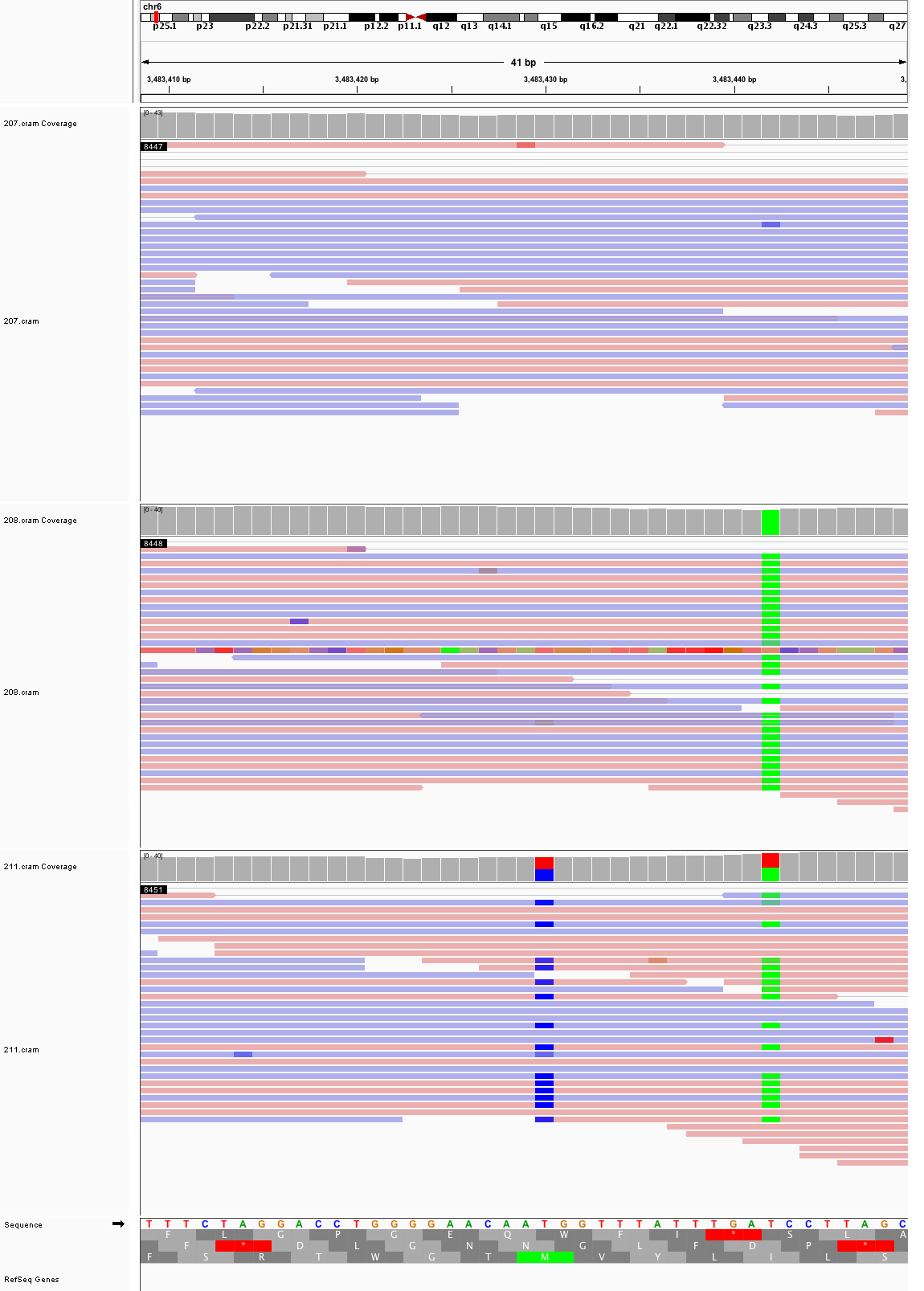

### chr6_81,158,408_81,158,448.png

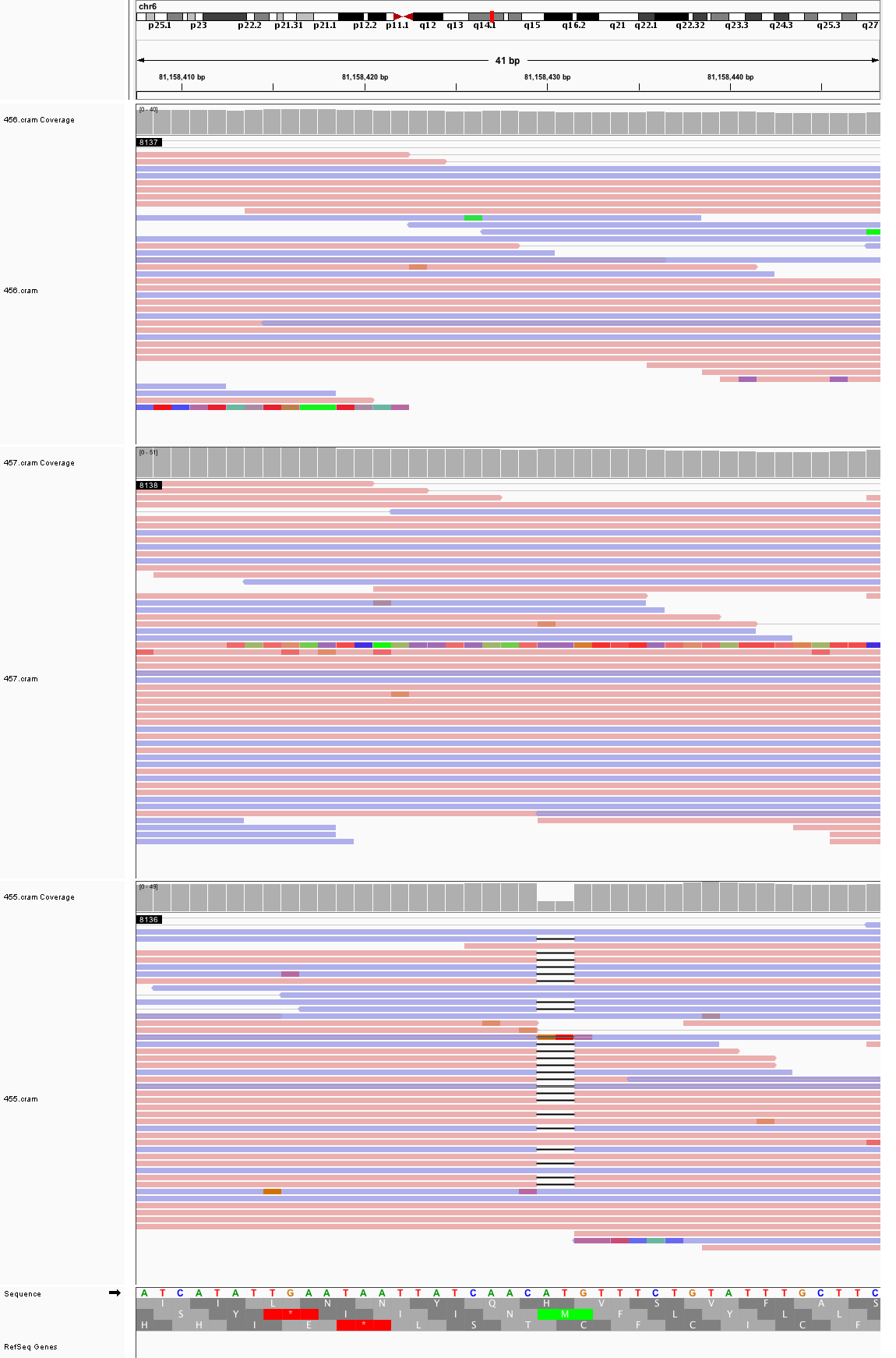

### chr6_124,951,766_124,951,806.png

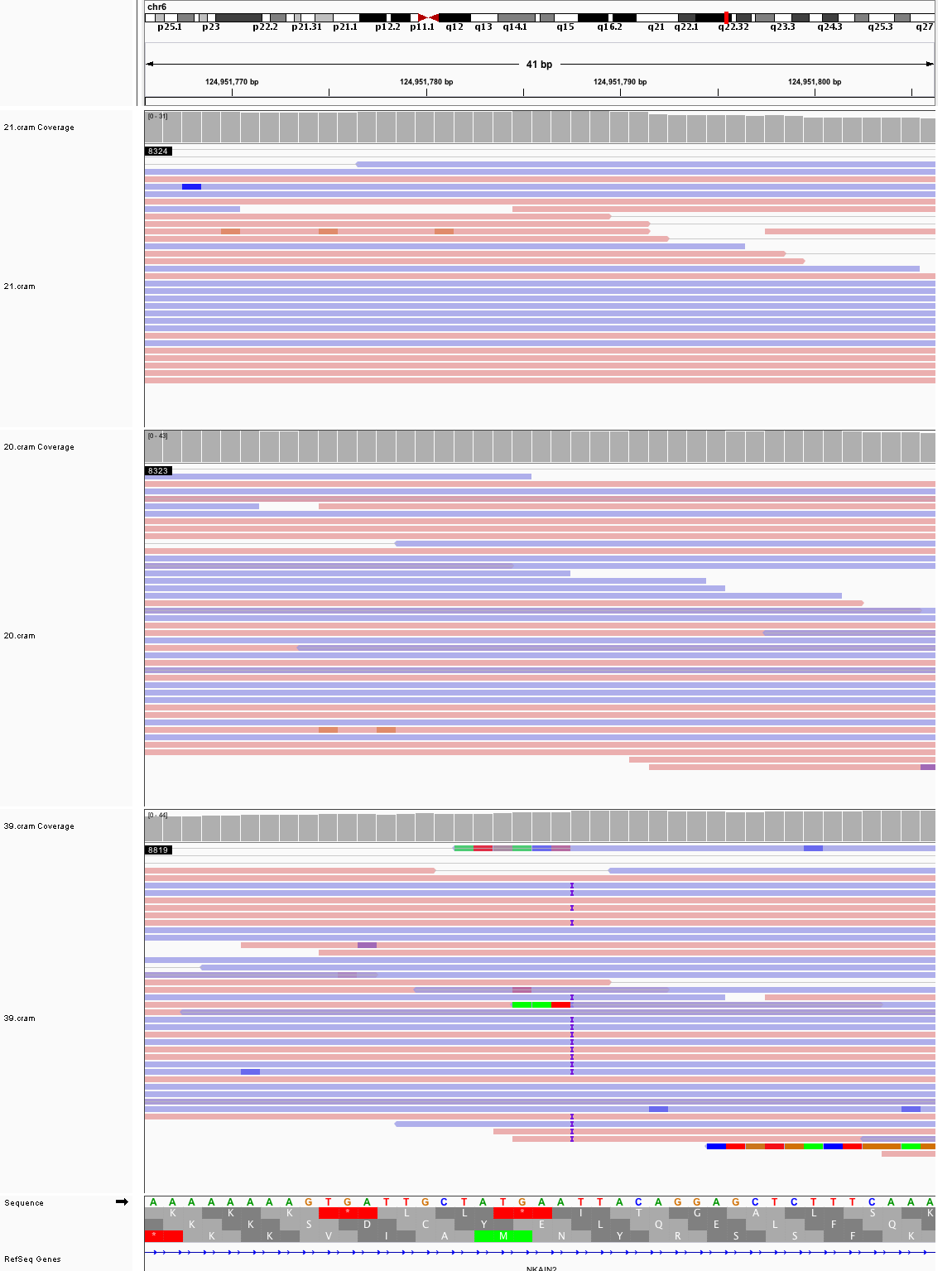

### chr7_24,970,787_24,970,827.png

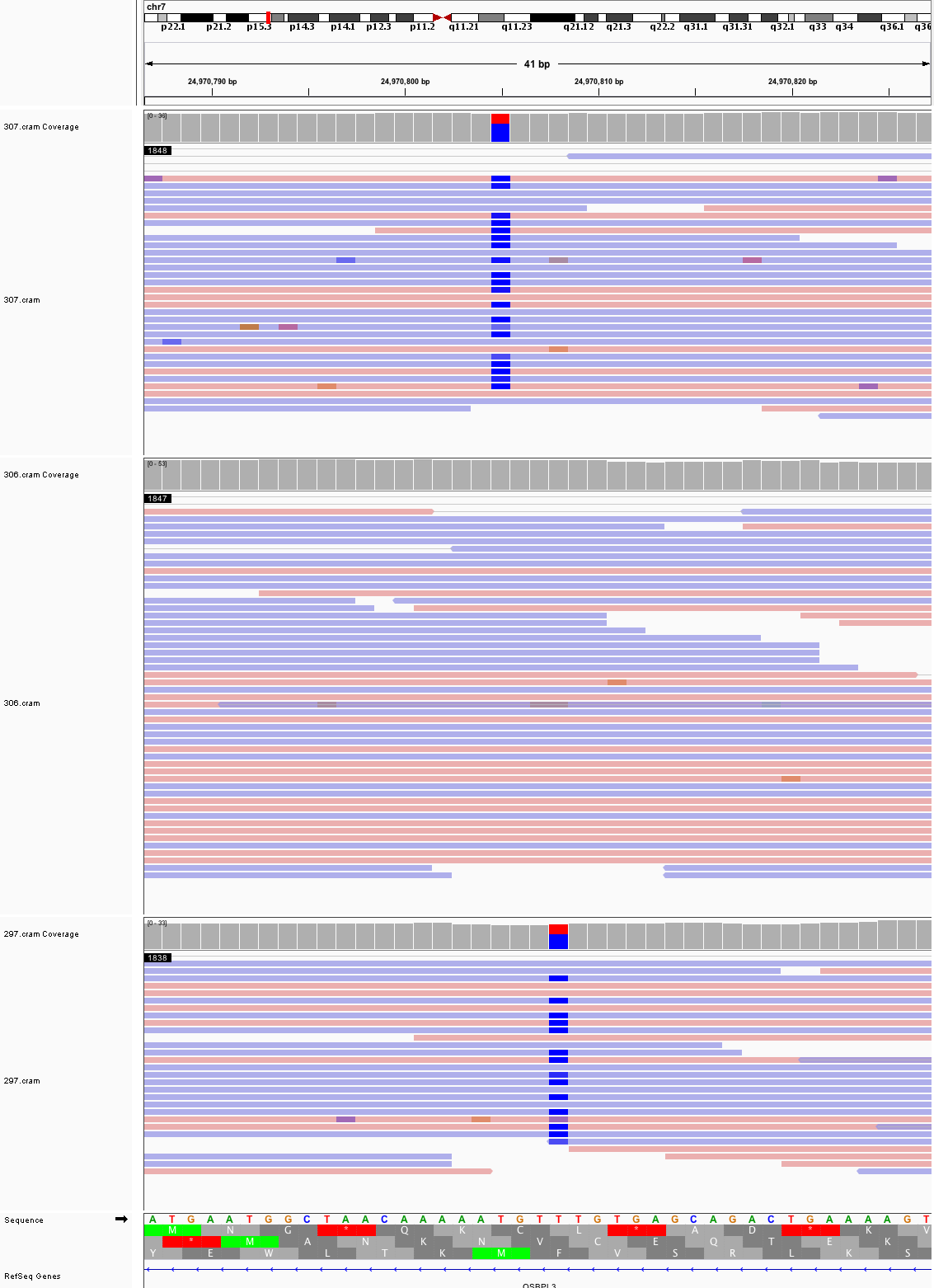

### chr7_45,294,177_45,294,217.png

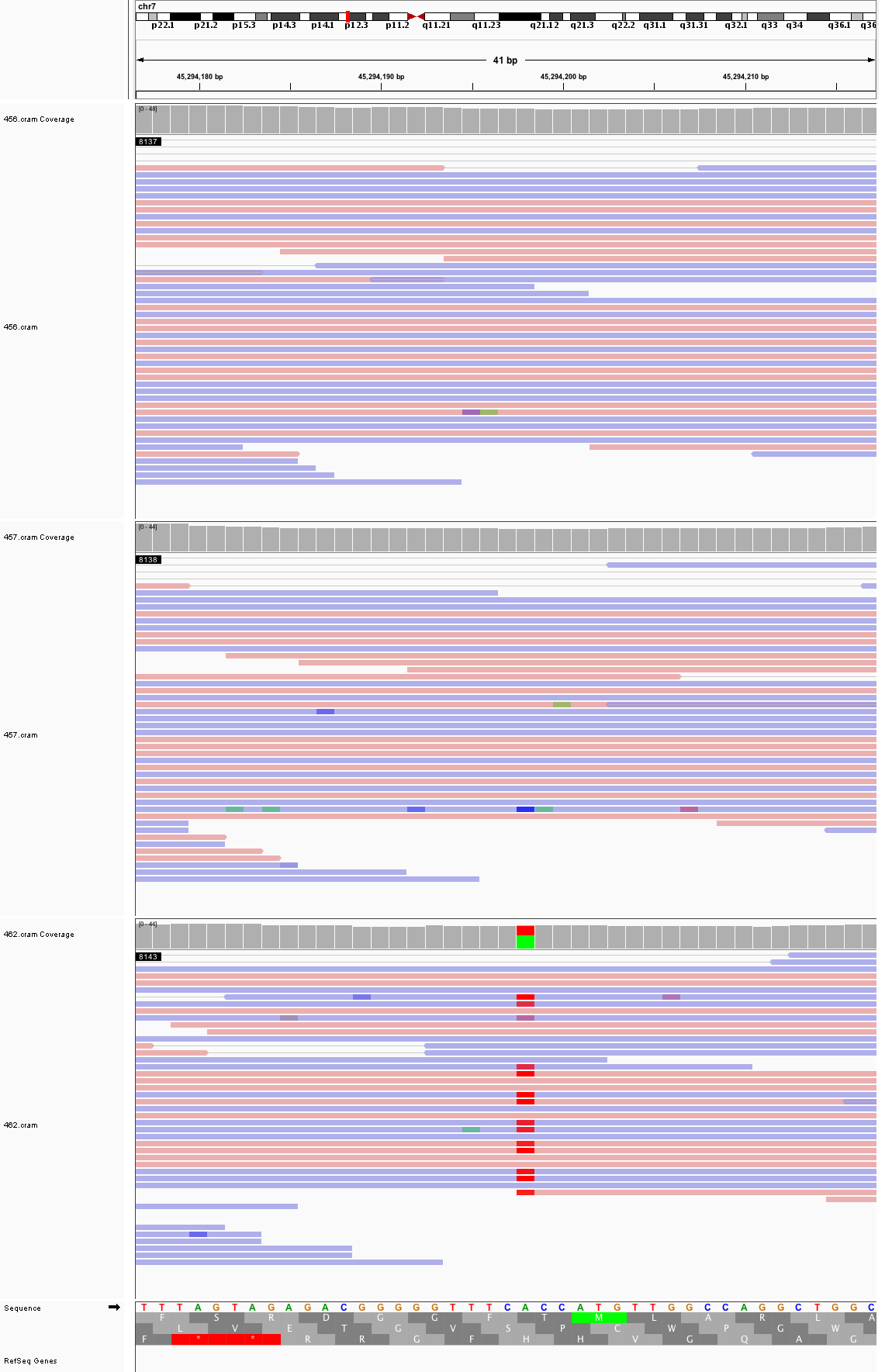

### chr7_53,269,122_53,269,162.png

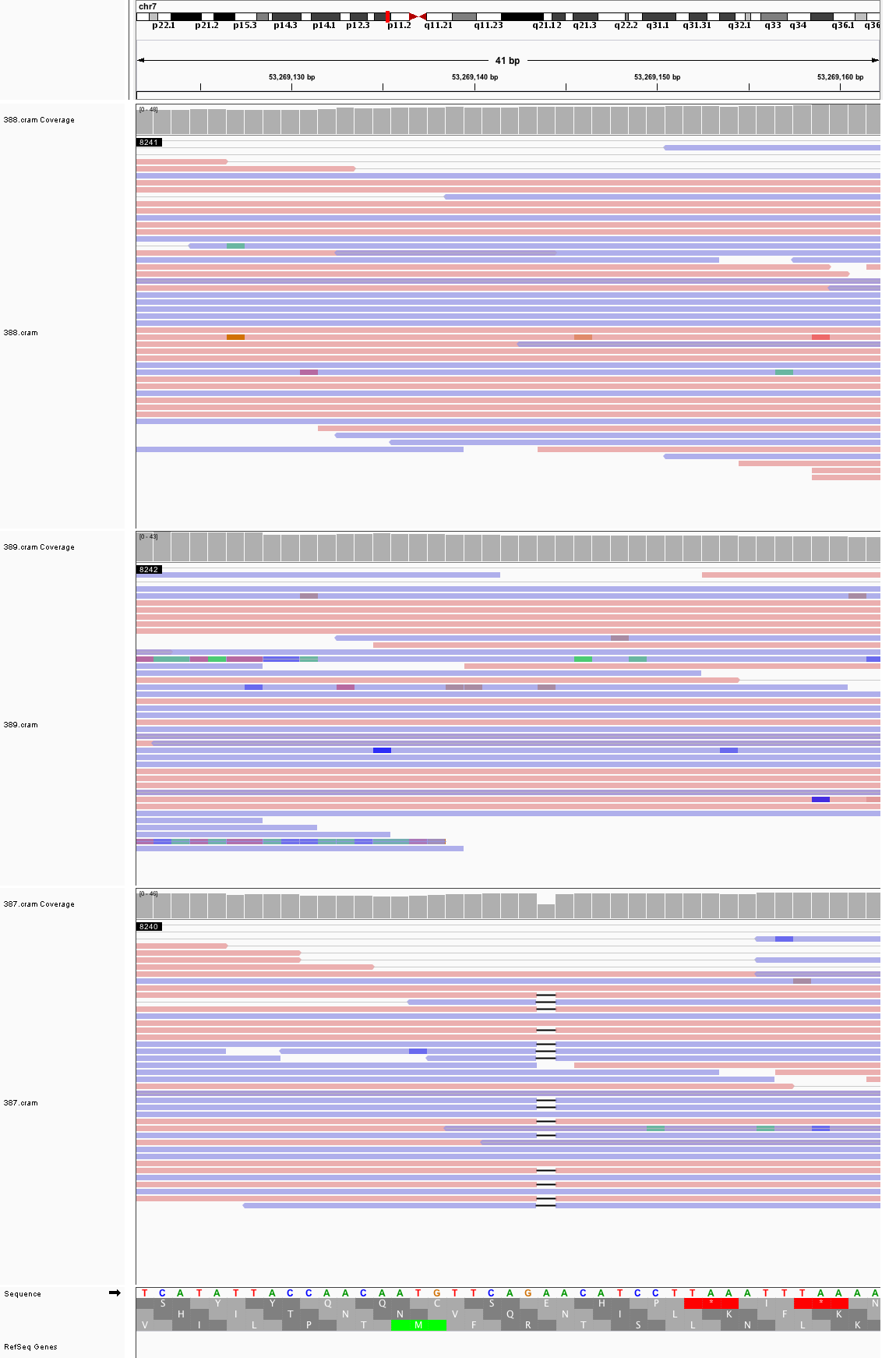

### chr7_71,545,005_71,545,045.png

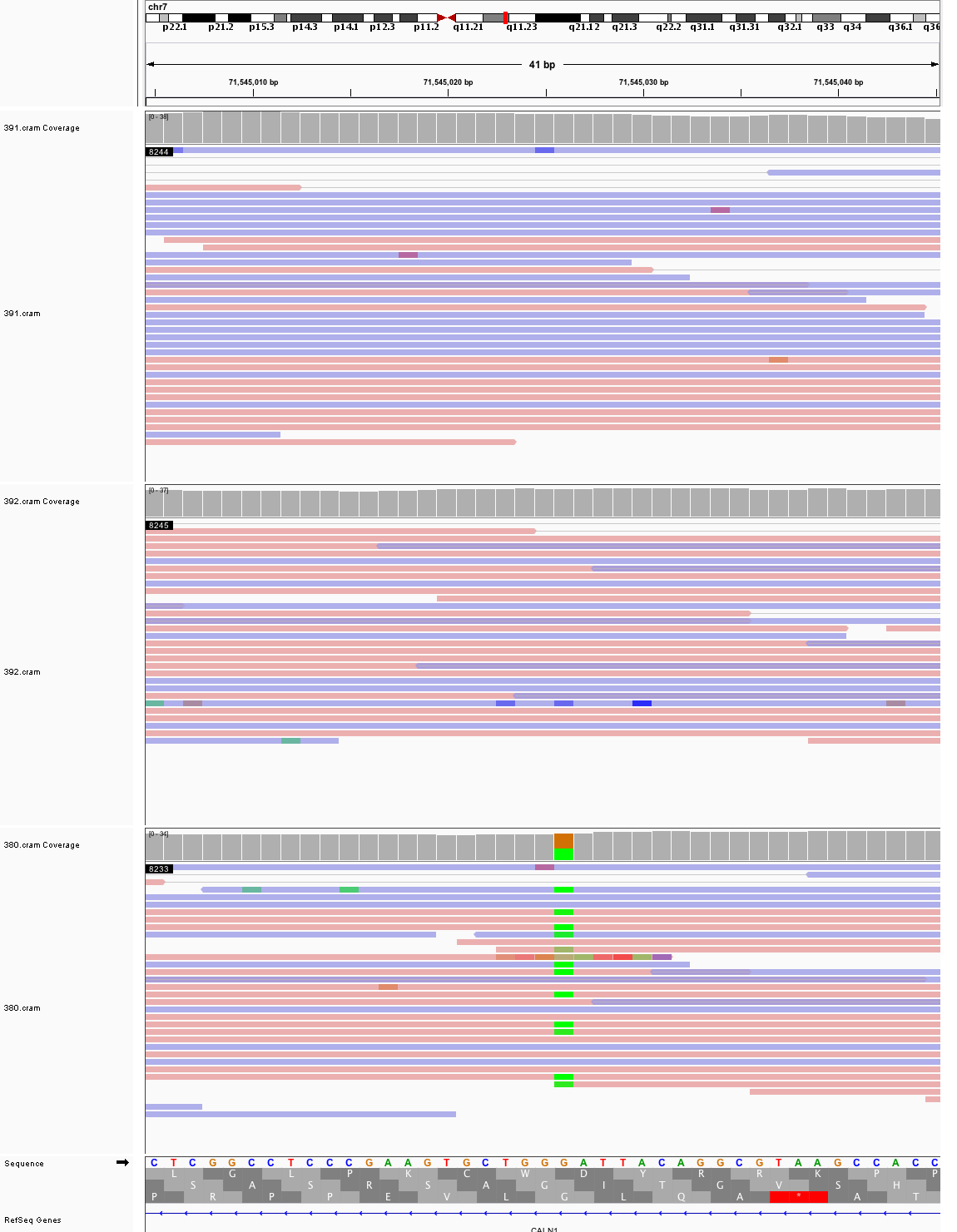

### chr7_98,507,616_98,507,656.png

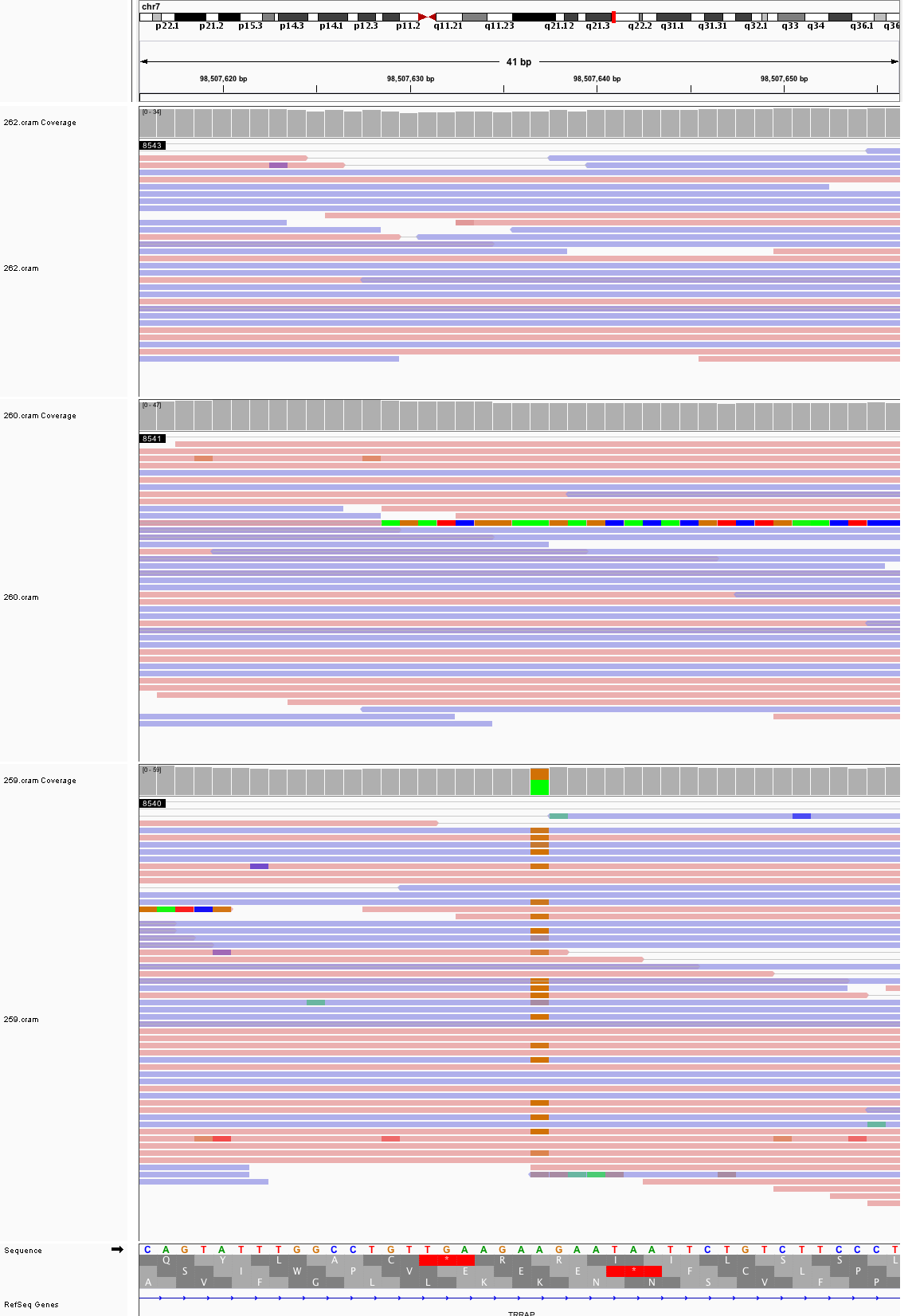

### chr7_103,386,139_103,386,179.png

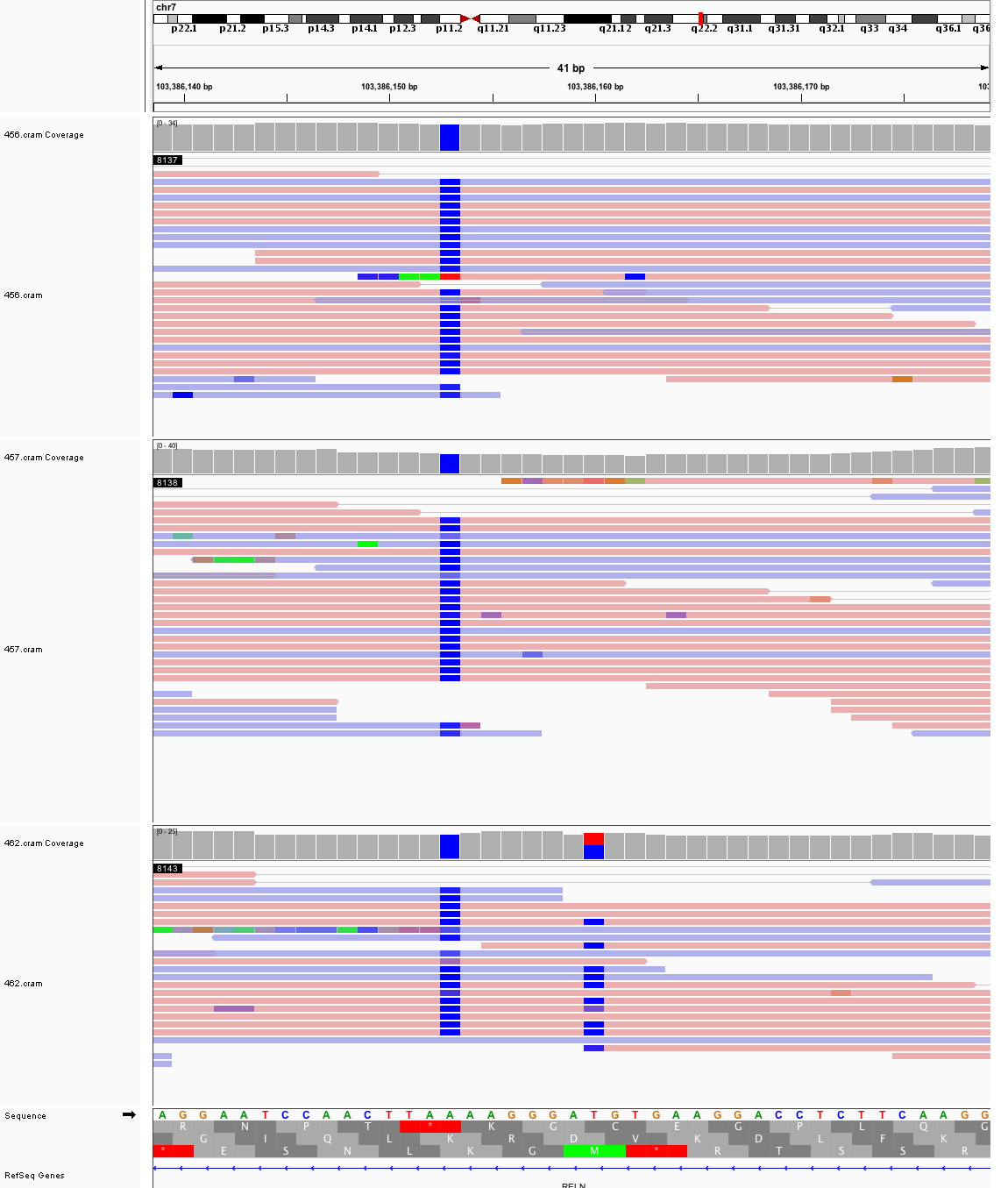

### chr7_156,931,355_156,931,395.png

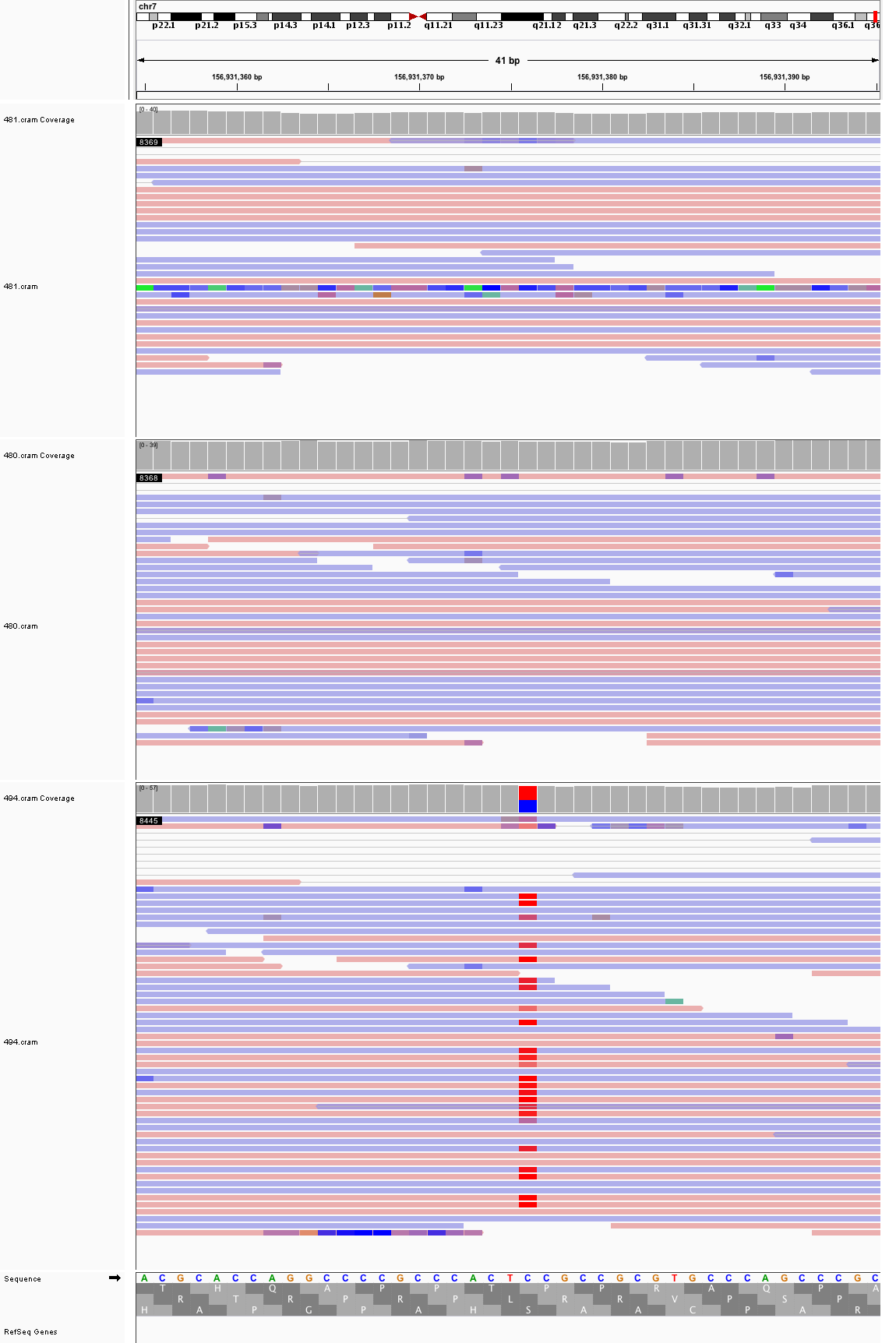

### chr8_1,630,006_1,630,046.png

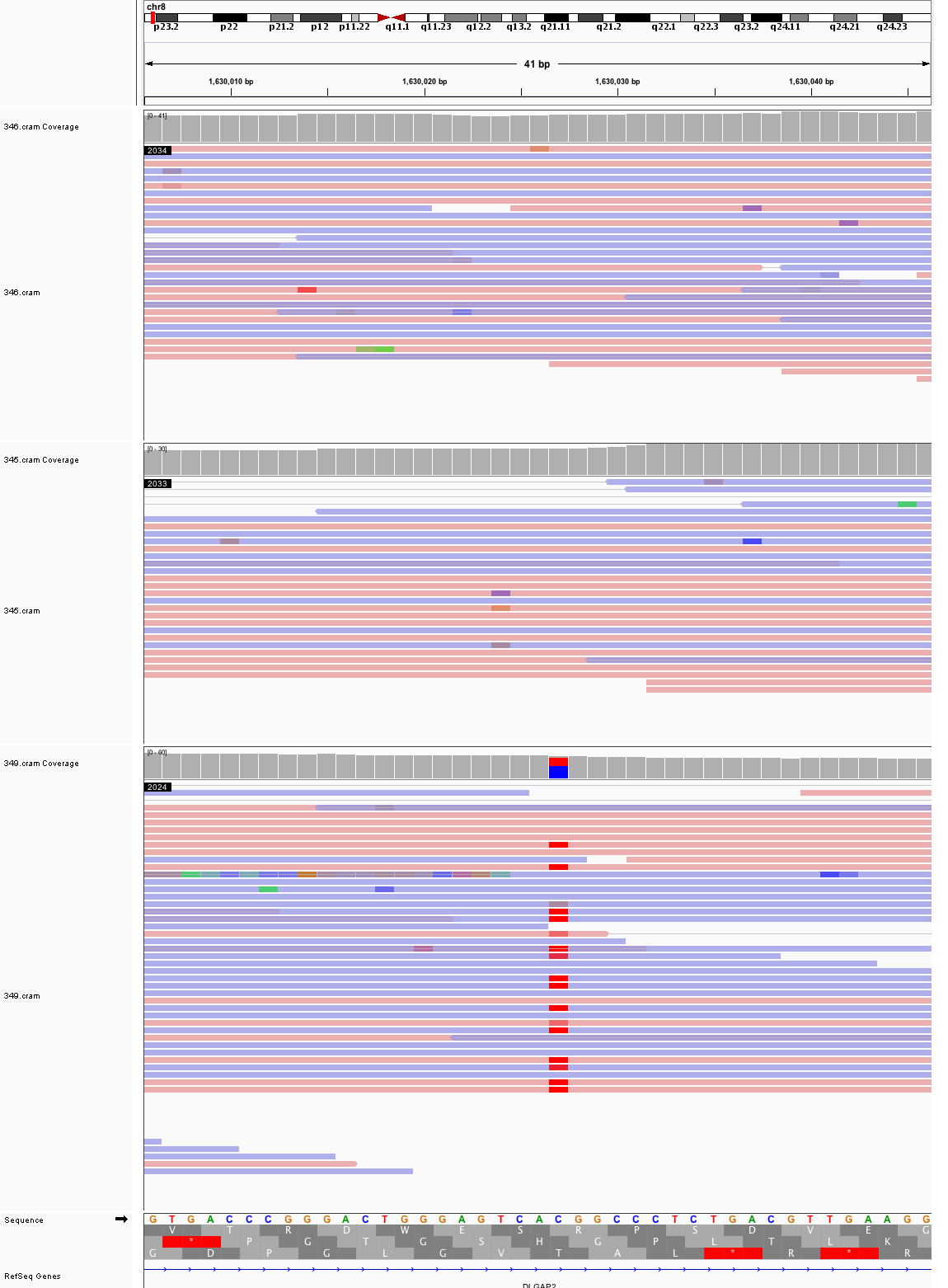

### chr8_93,396,295_93,396,335.png

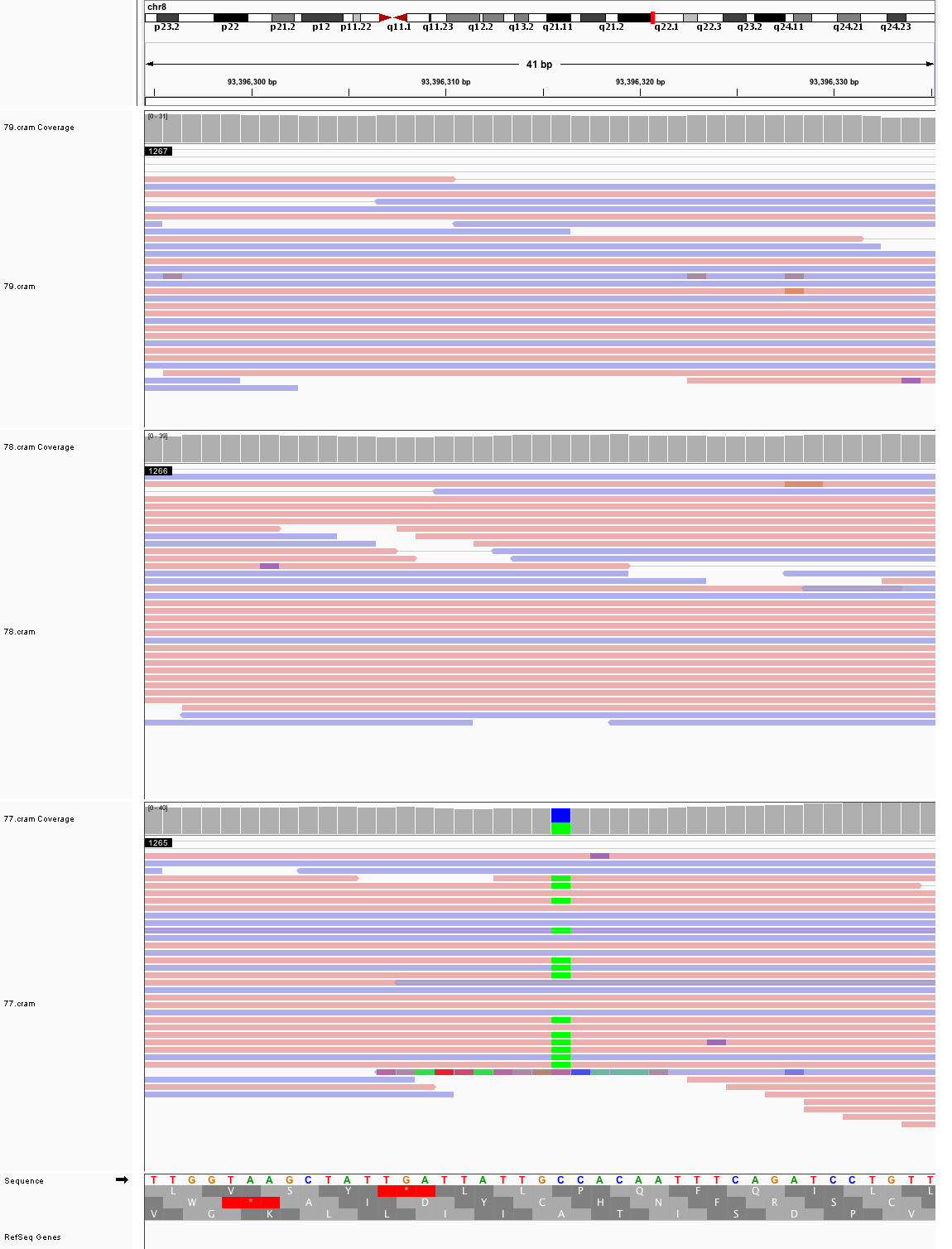

### chr9_90,374,329_90,374,369.png

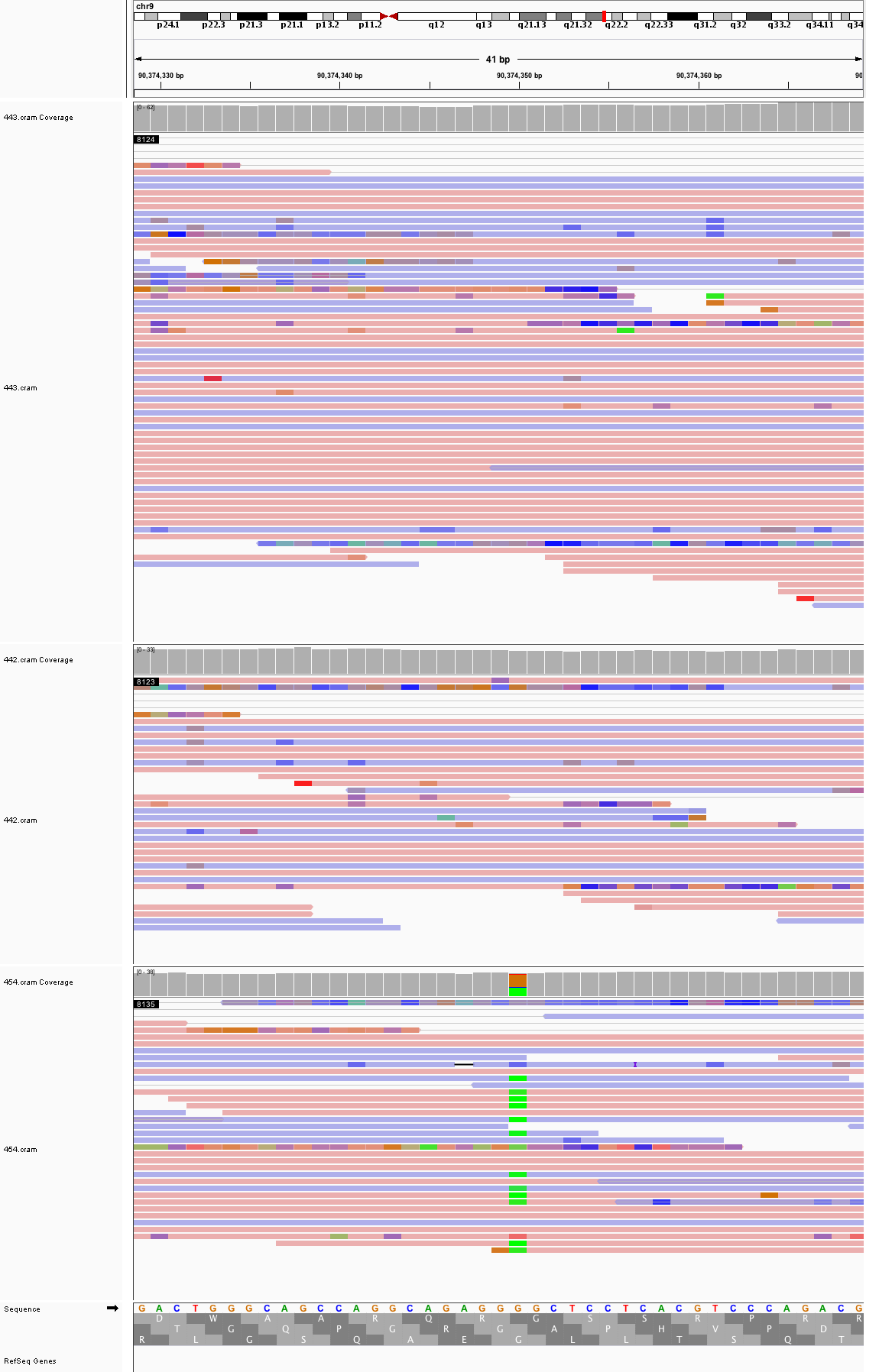

### chr9_112,696,809_112,696,849.png

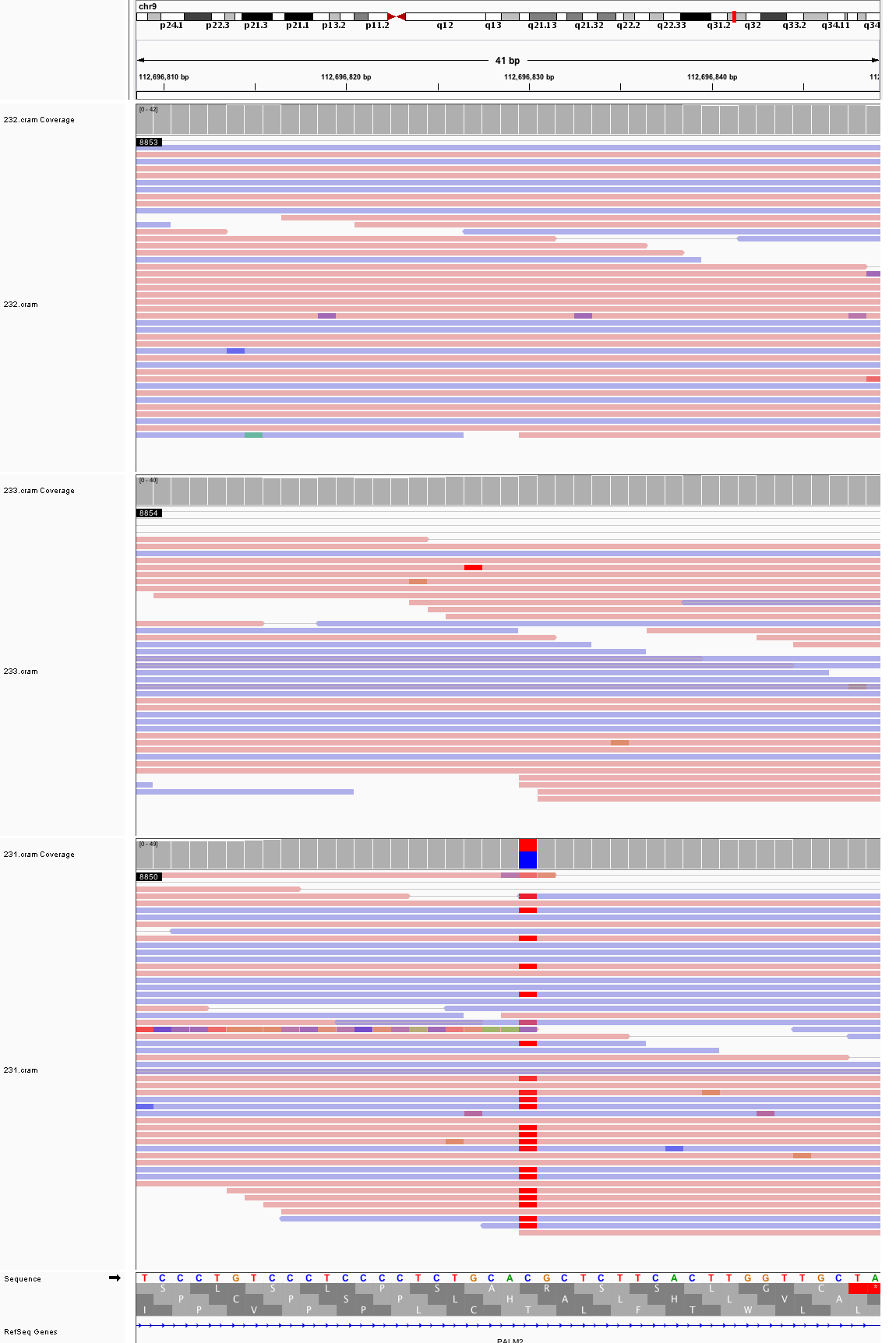

### chr10_12,787,063_12,787,103.png

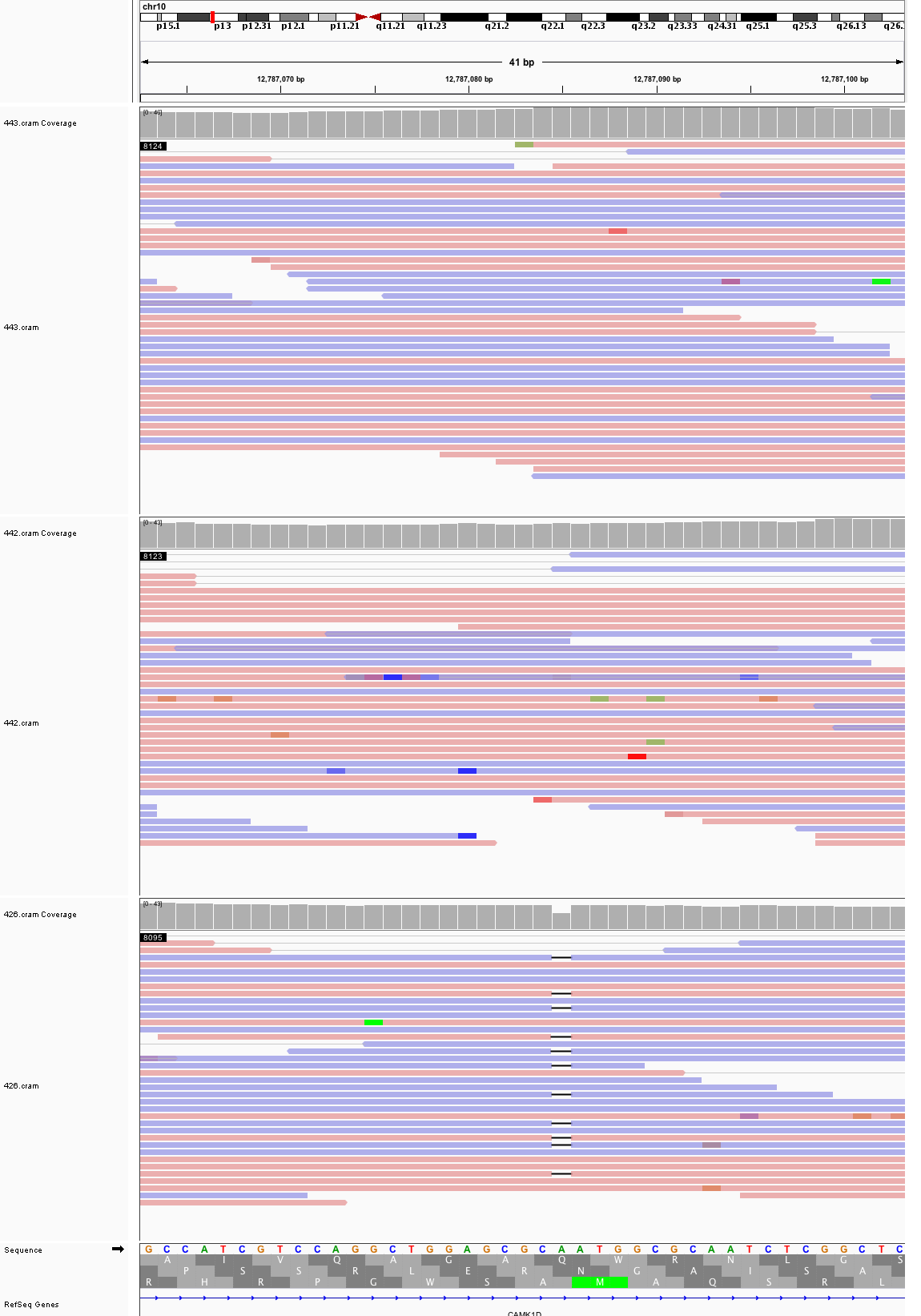

### chr10_78,700,013_78,700,053.png

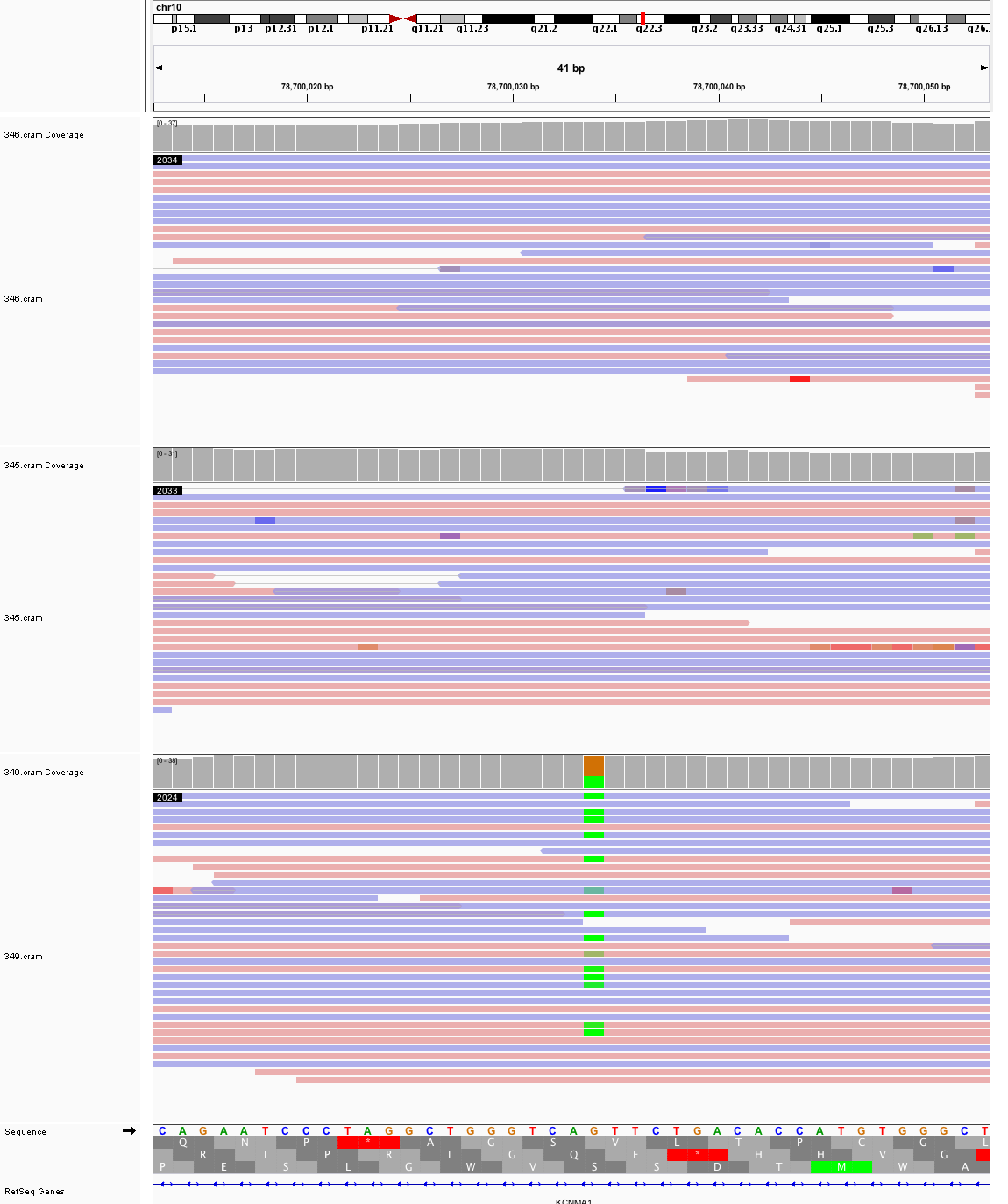

### chr10_79,908,170_79,908,210.png

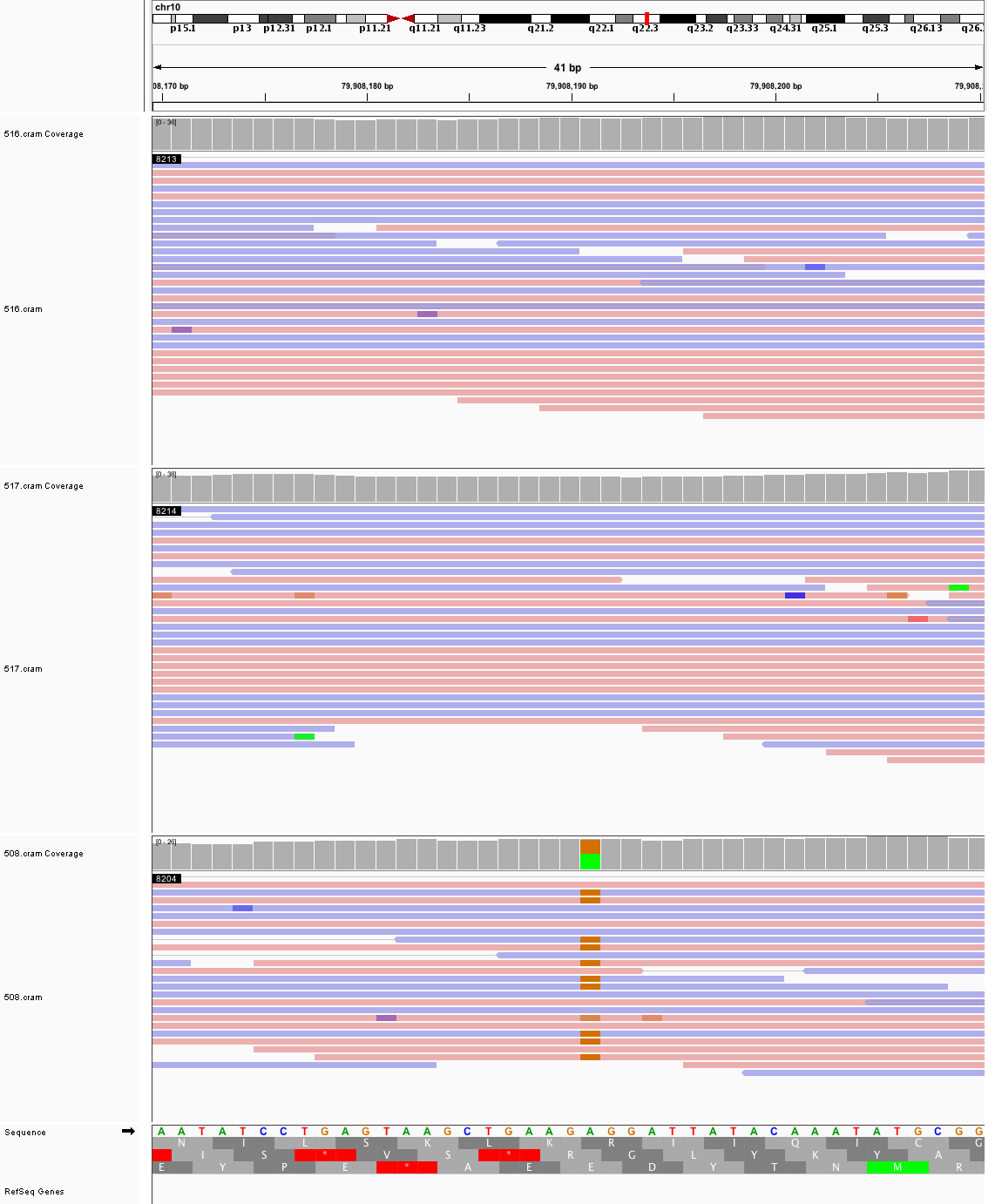

### chr11_114,820,023_114,820,063.png

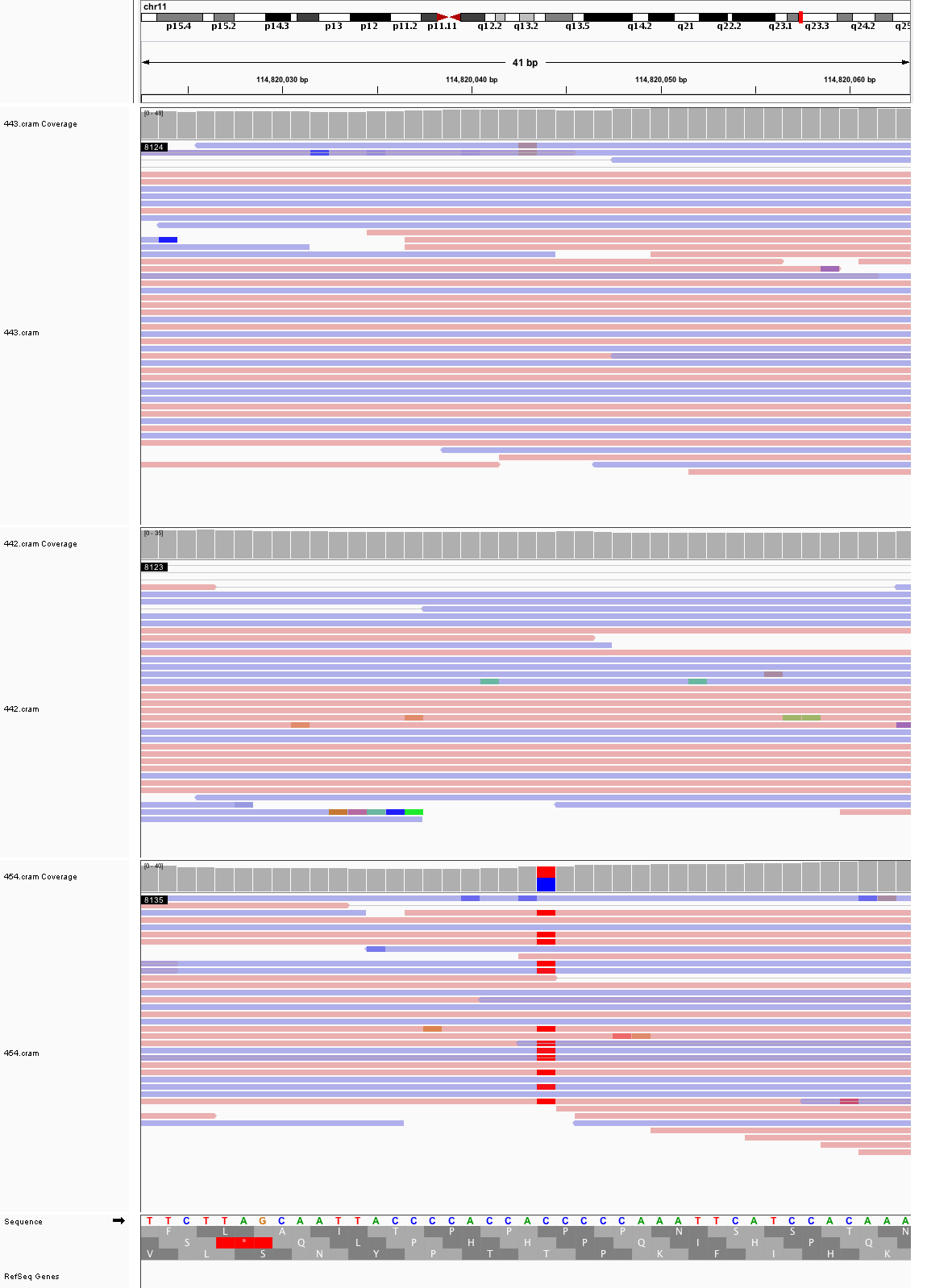

### chr11_121,670,536_121,670,576.png

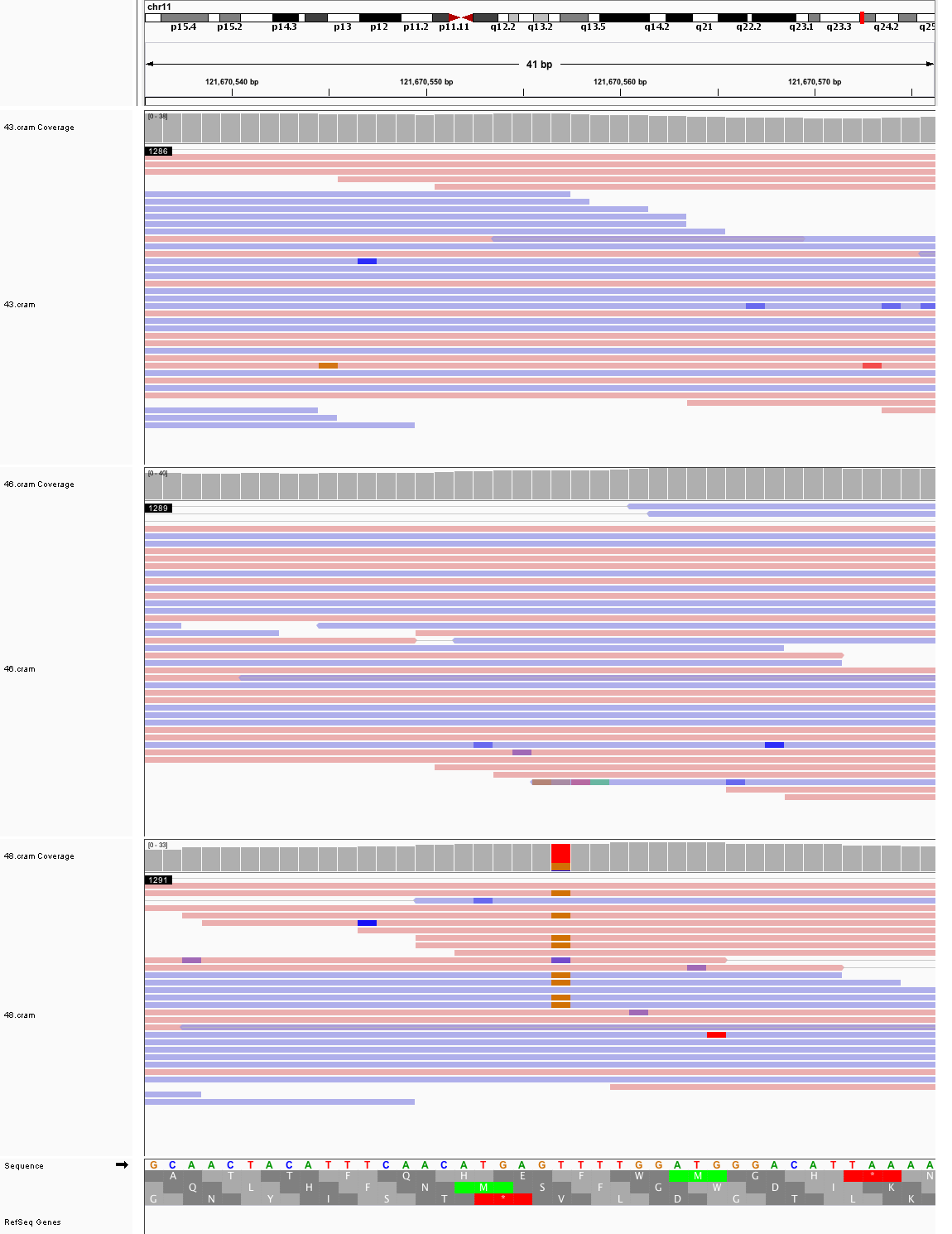

### chr12_18,560,283_18,560,323.png

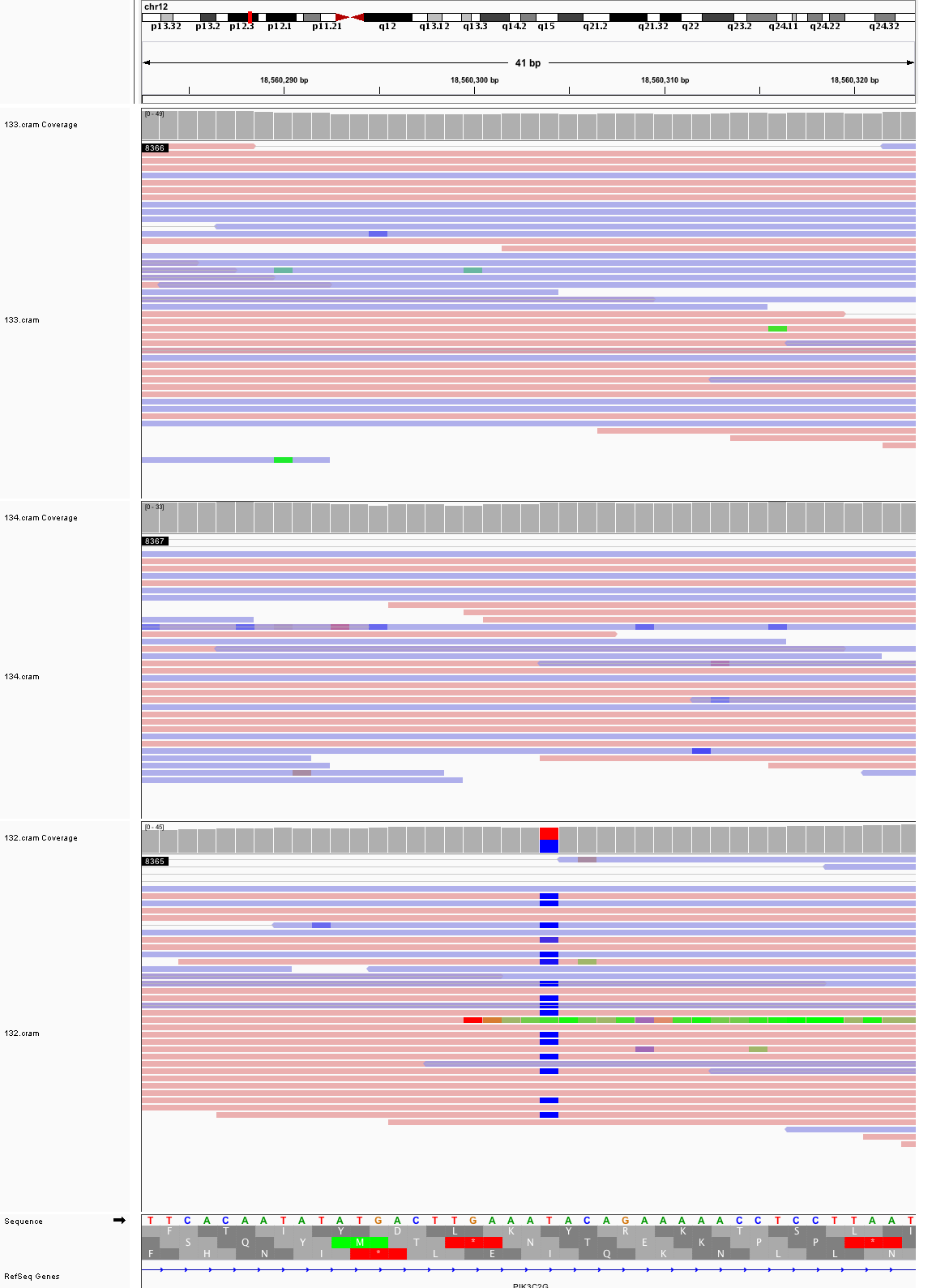

### chr12_27,255,242_27,255,282.png

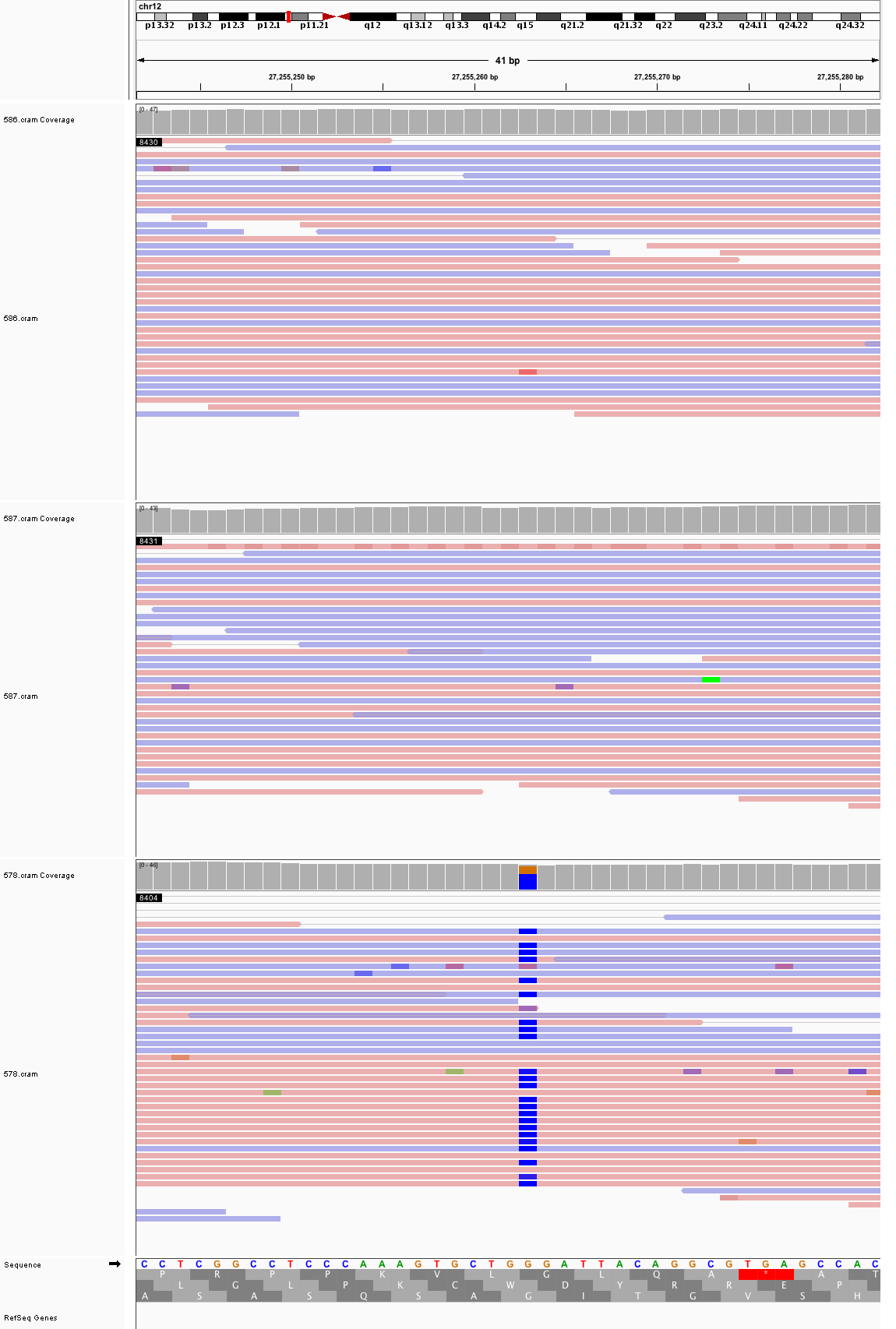

### chr13_44,236,828_44,236,868.png

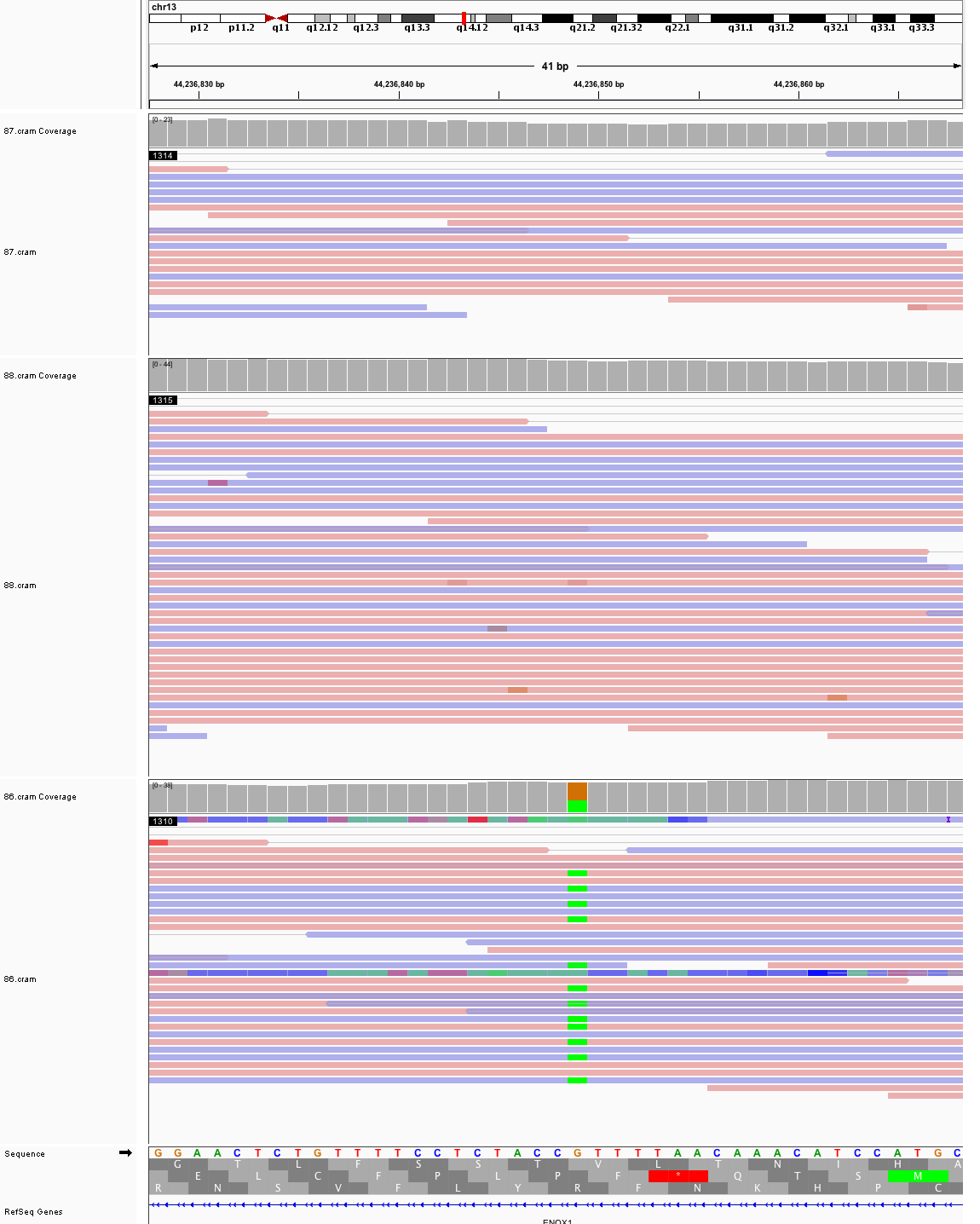

### chr14_51,447,577_51,447,617.png

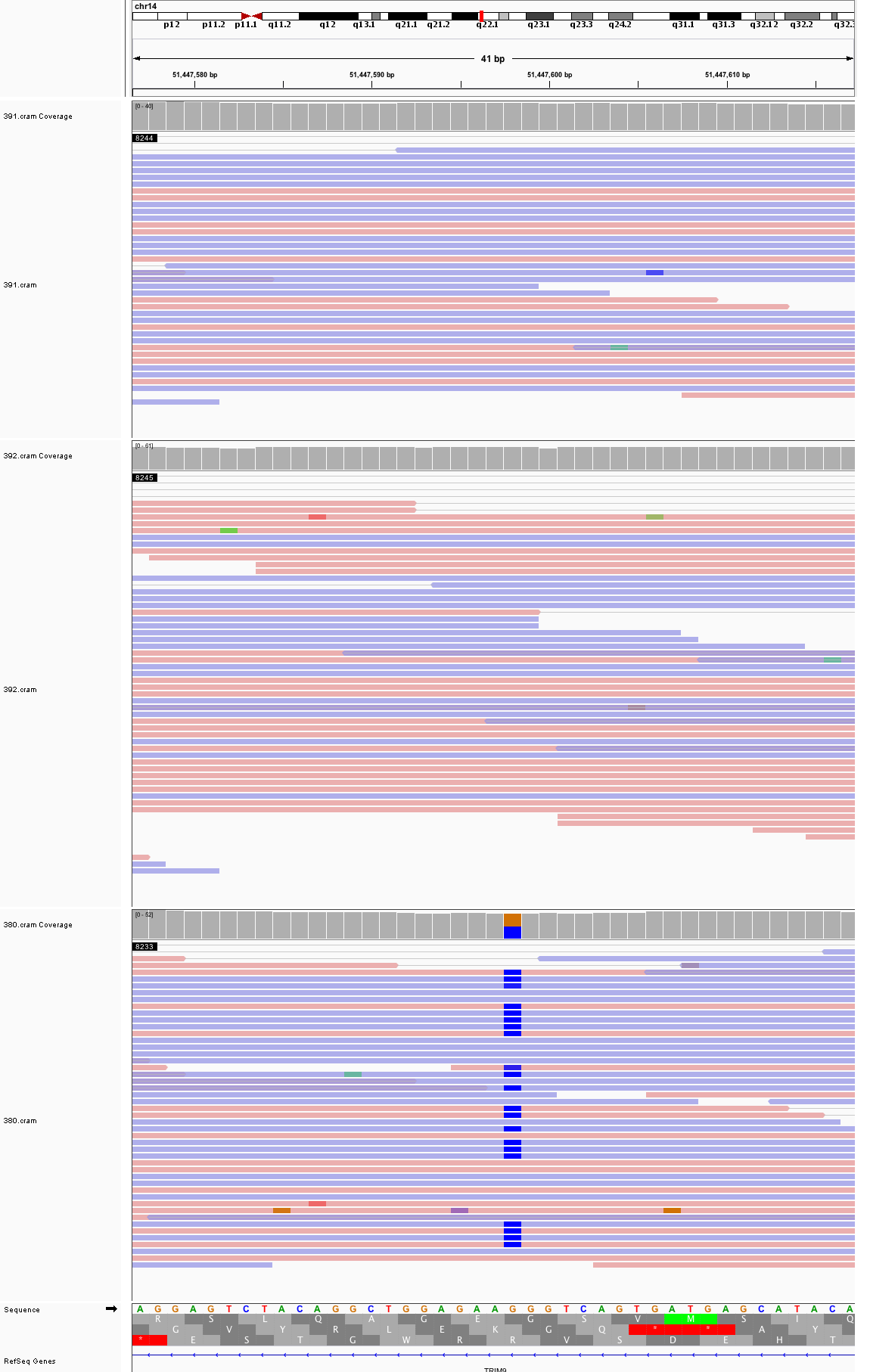

### chr14_65,491,989_65,492,029.png

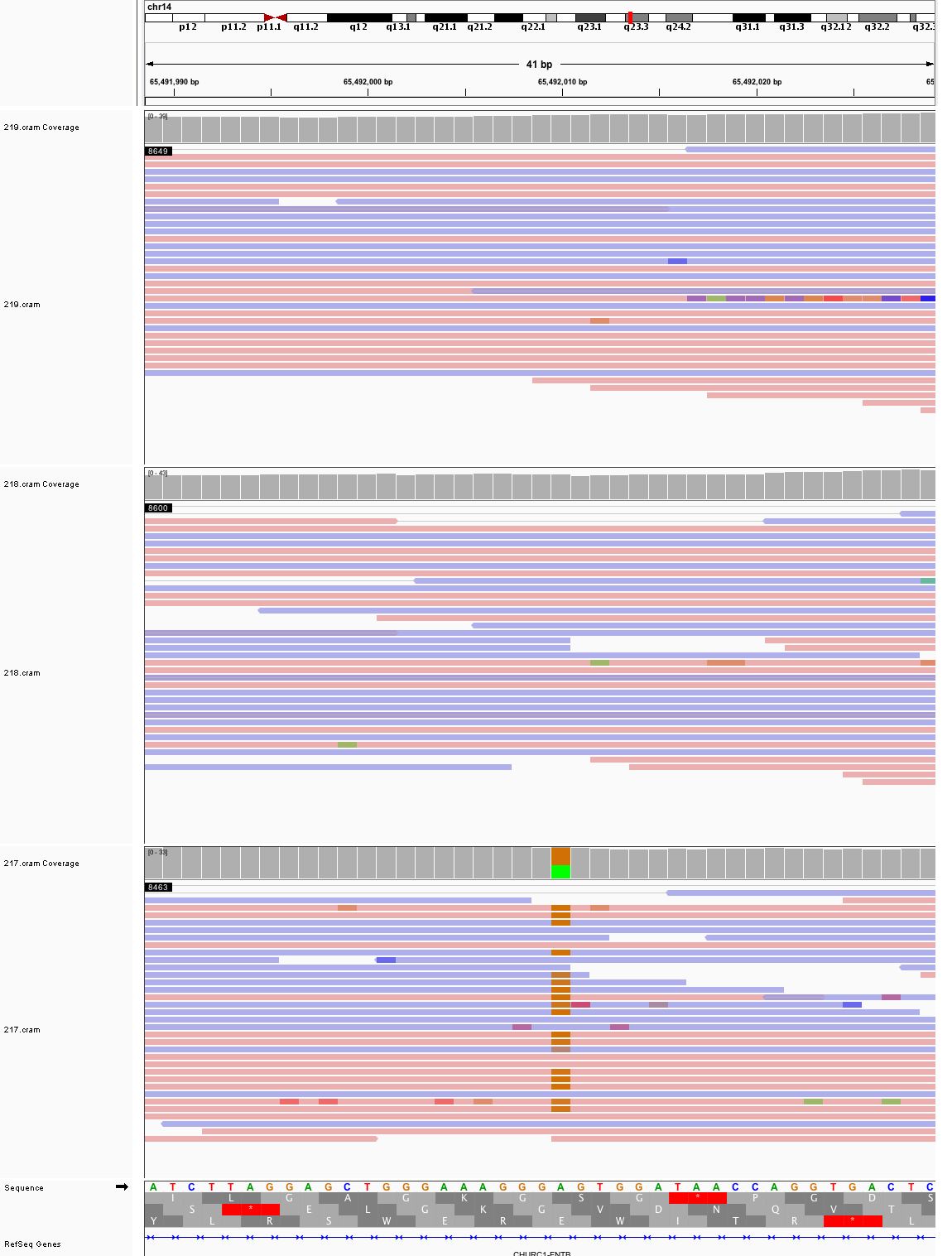

### chr15_49,572,253_49,572,293.png

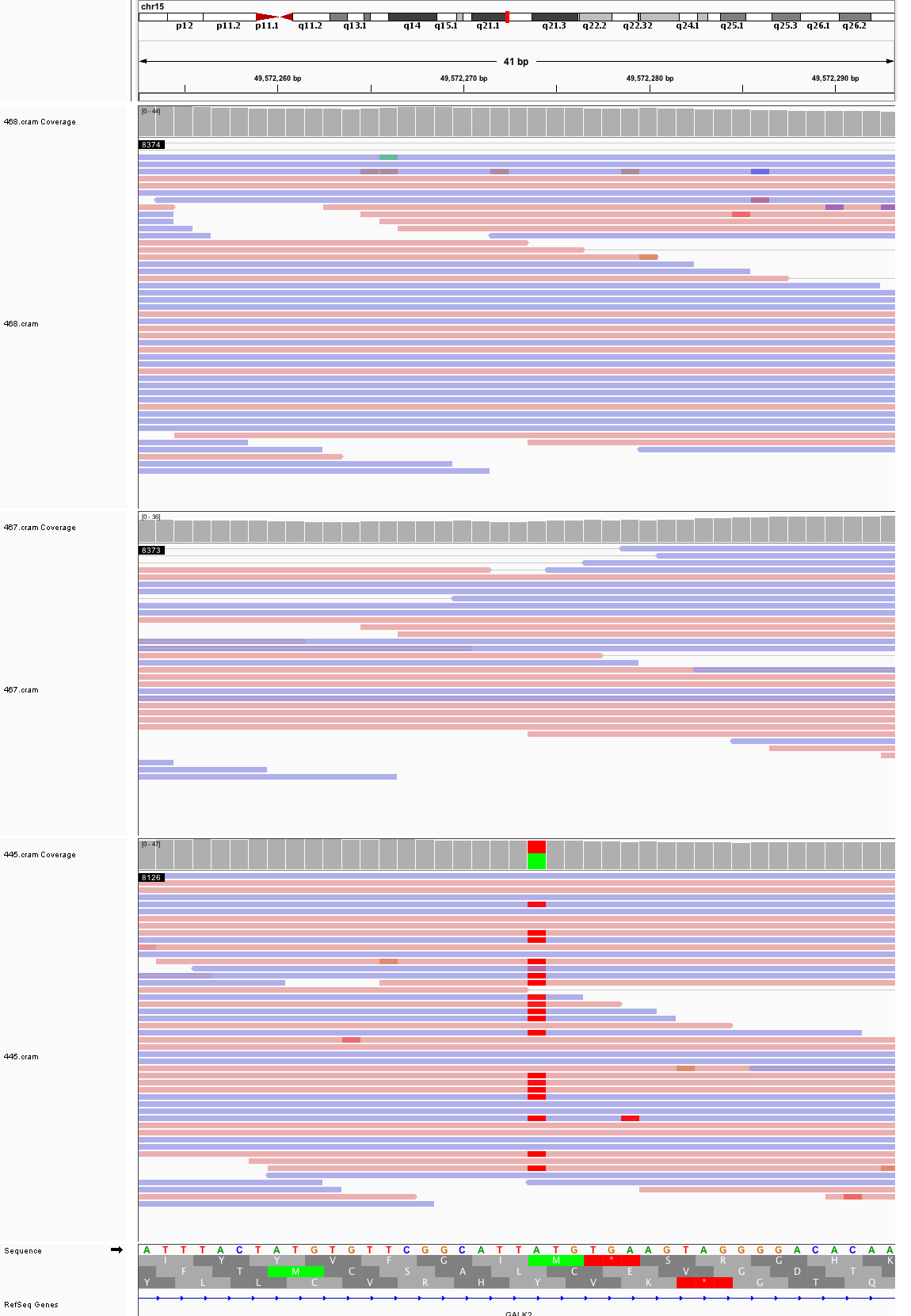

### chr15_62,915,136_62,915,176.png

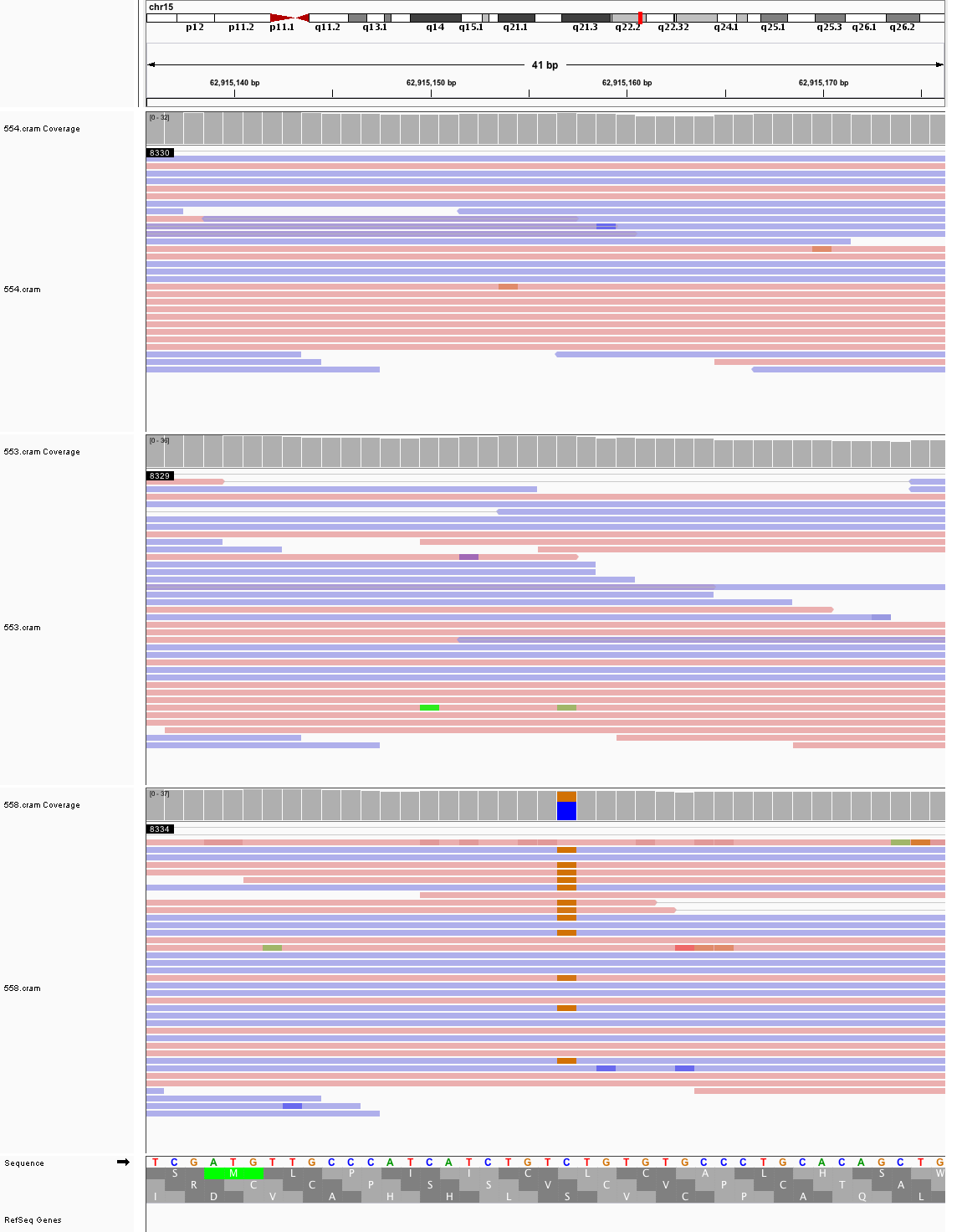

### chr15_100,847,960_100,848,000.png

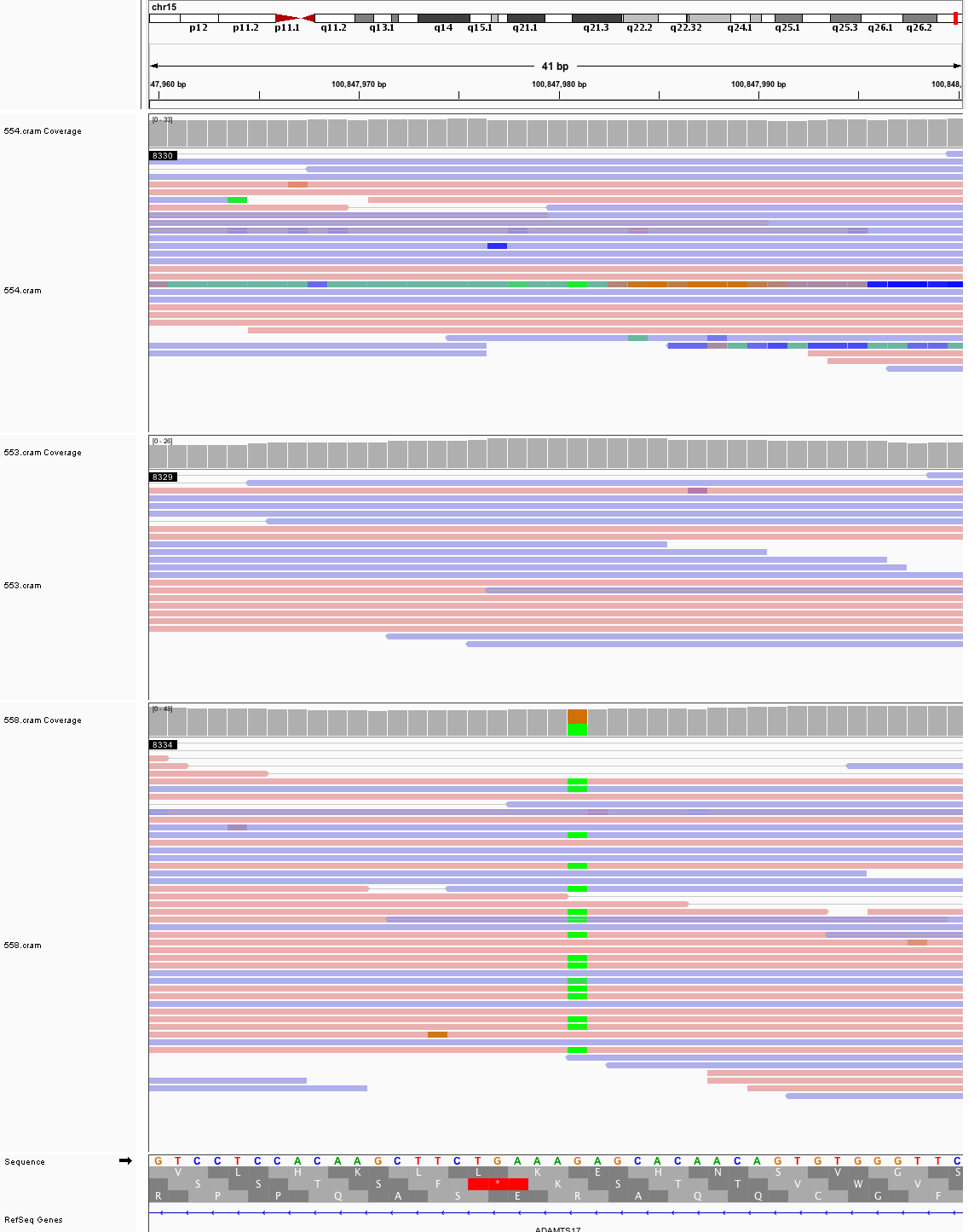

### chr16_65,110,553_65,110,593.png

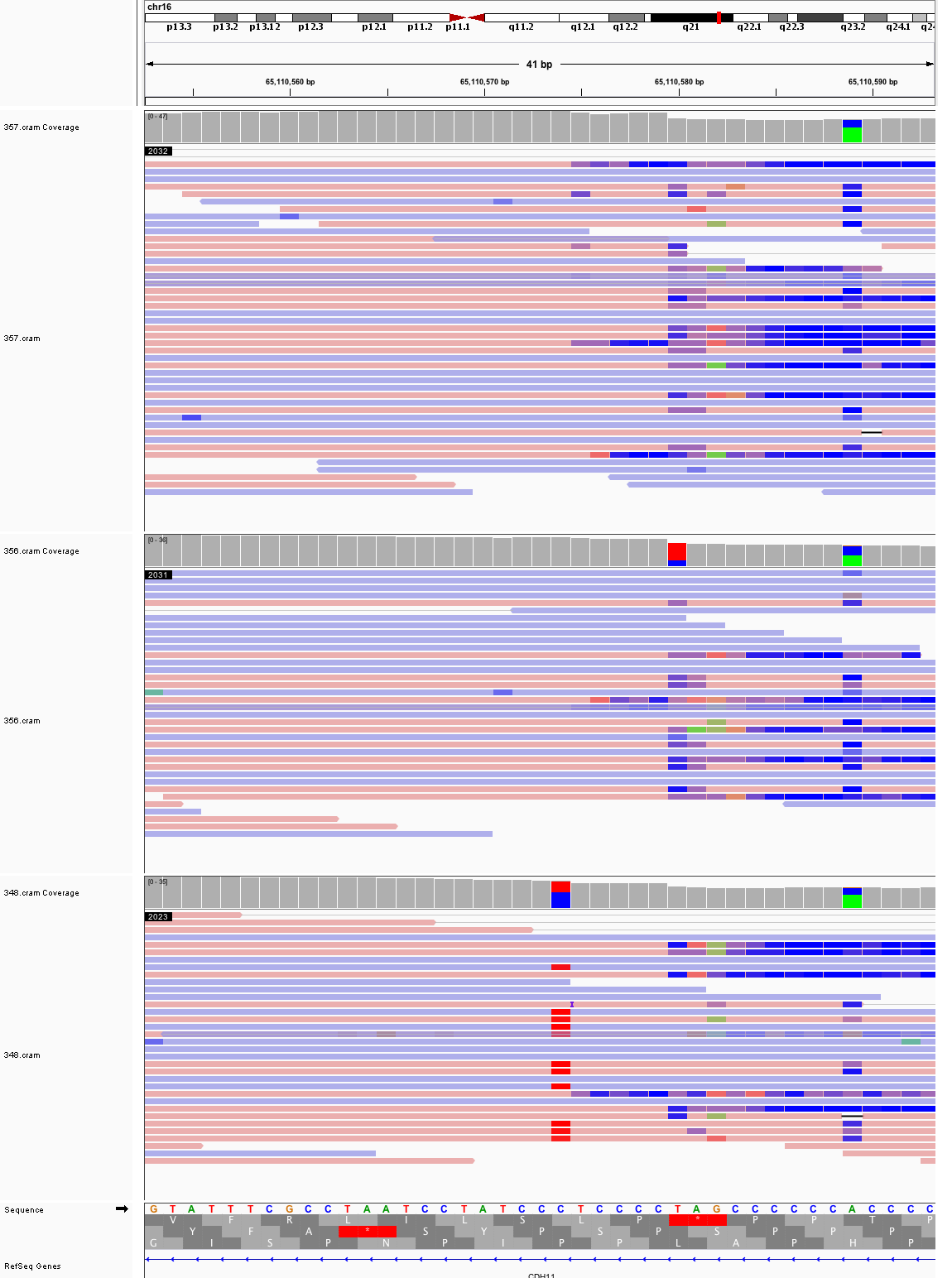

### chr17_3,681,324_3,681,364.png

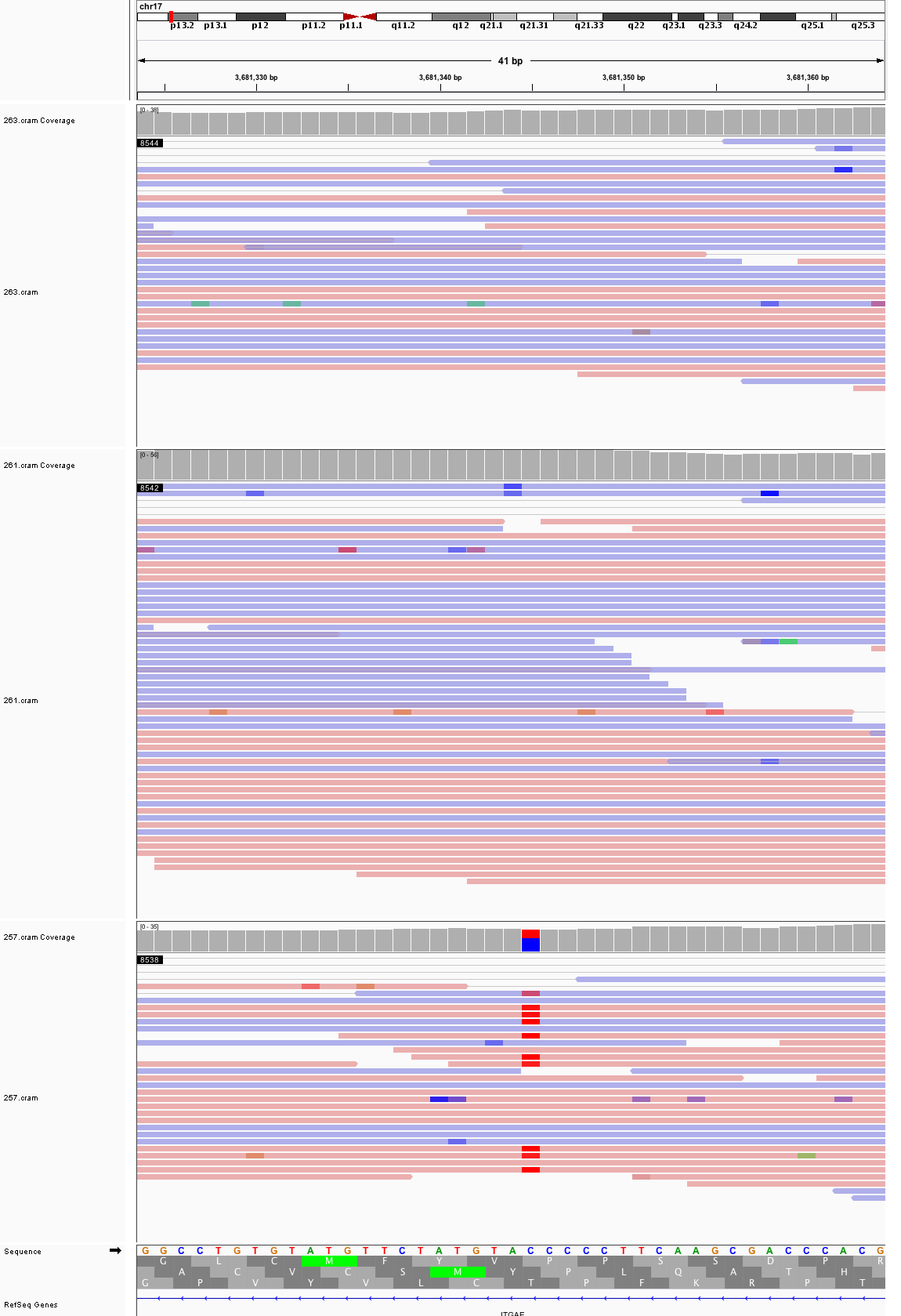

### chr17_25,506,457_25,506,497.png

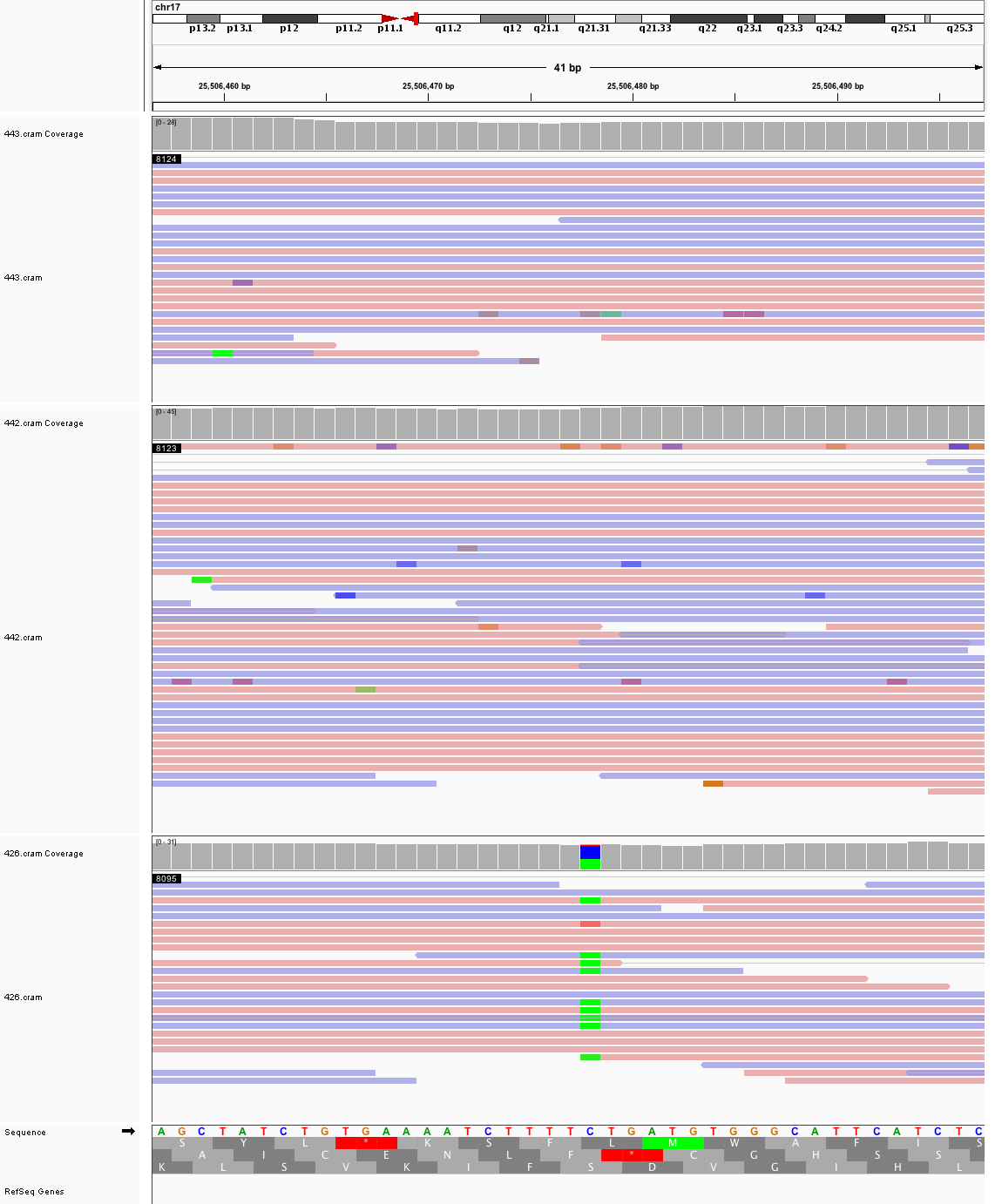

### chr18_21,655,028_21,655,068.png

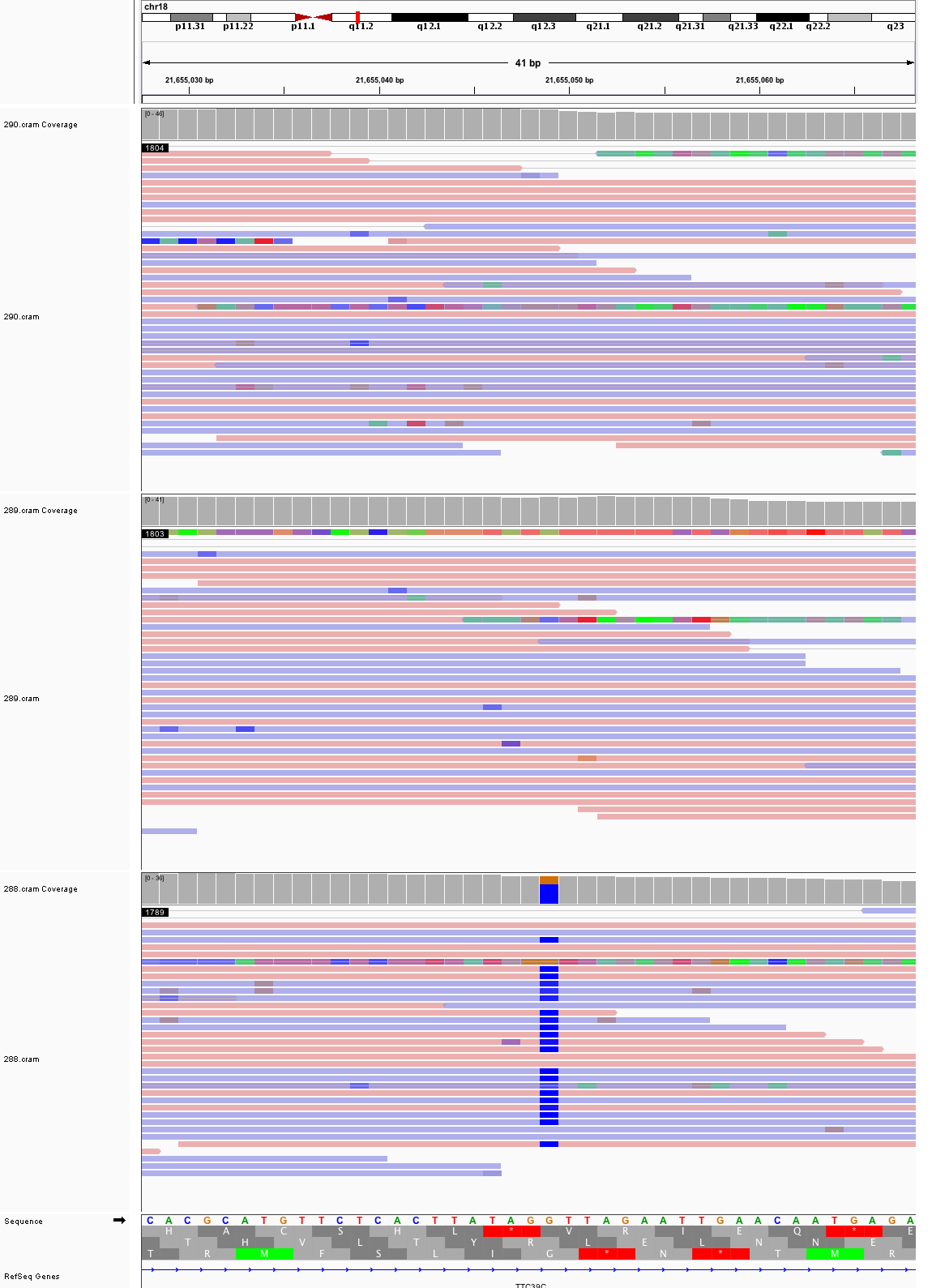

### chr18_52,677,563_52,677,603.png

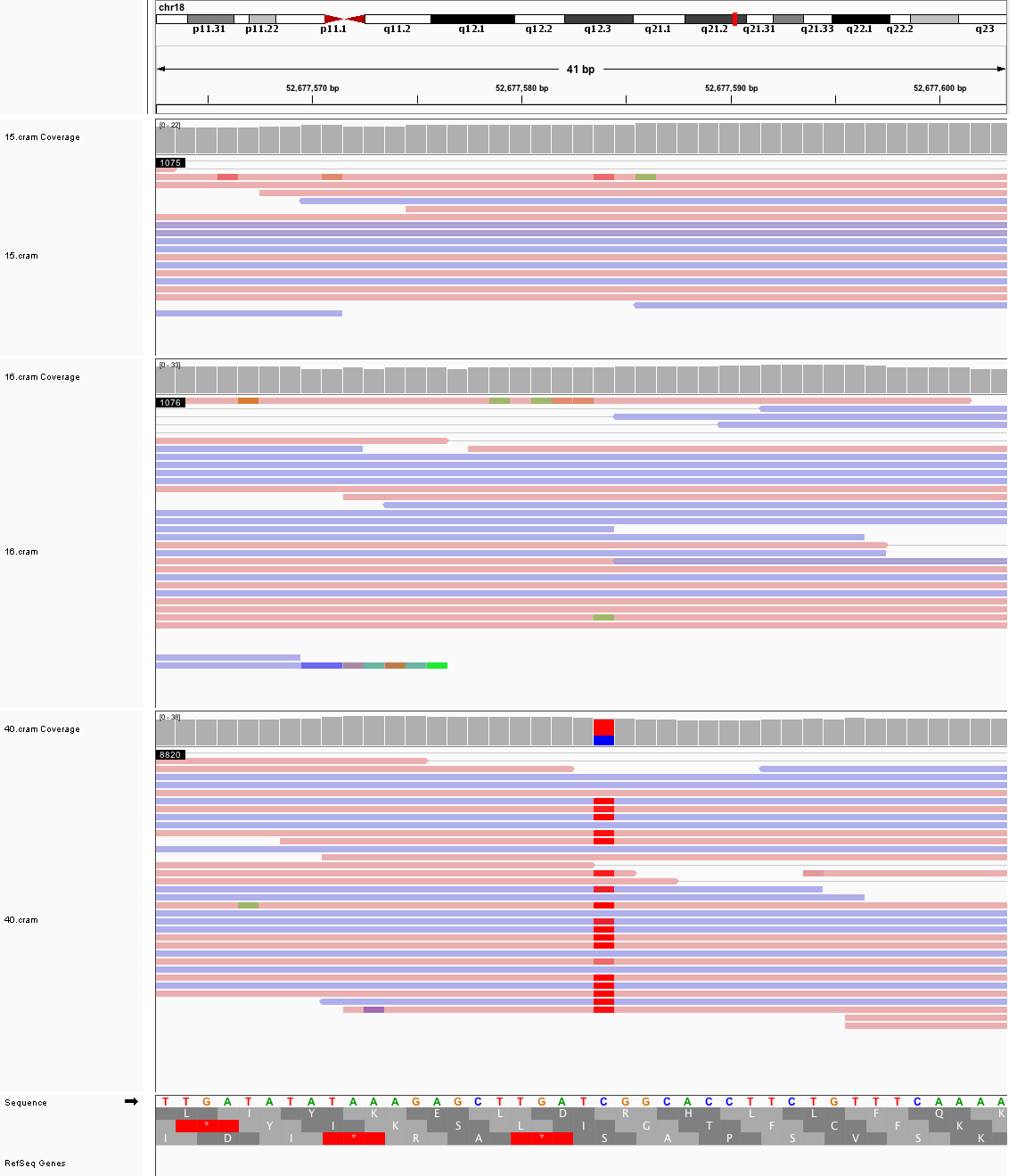

### chr19_34,522,317_34,522,357.png

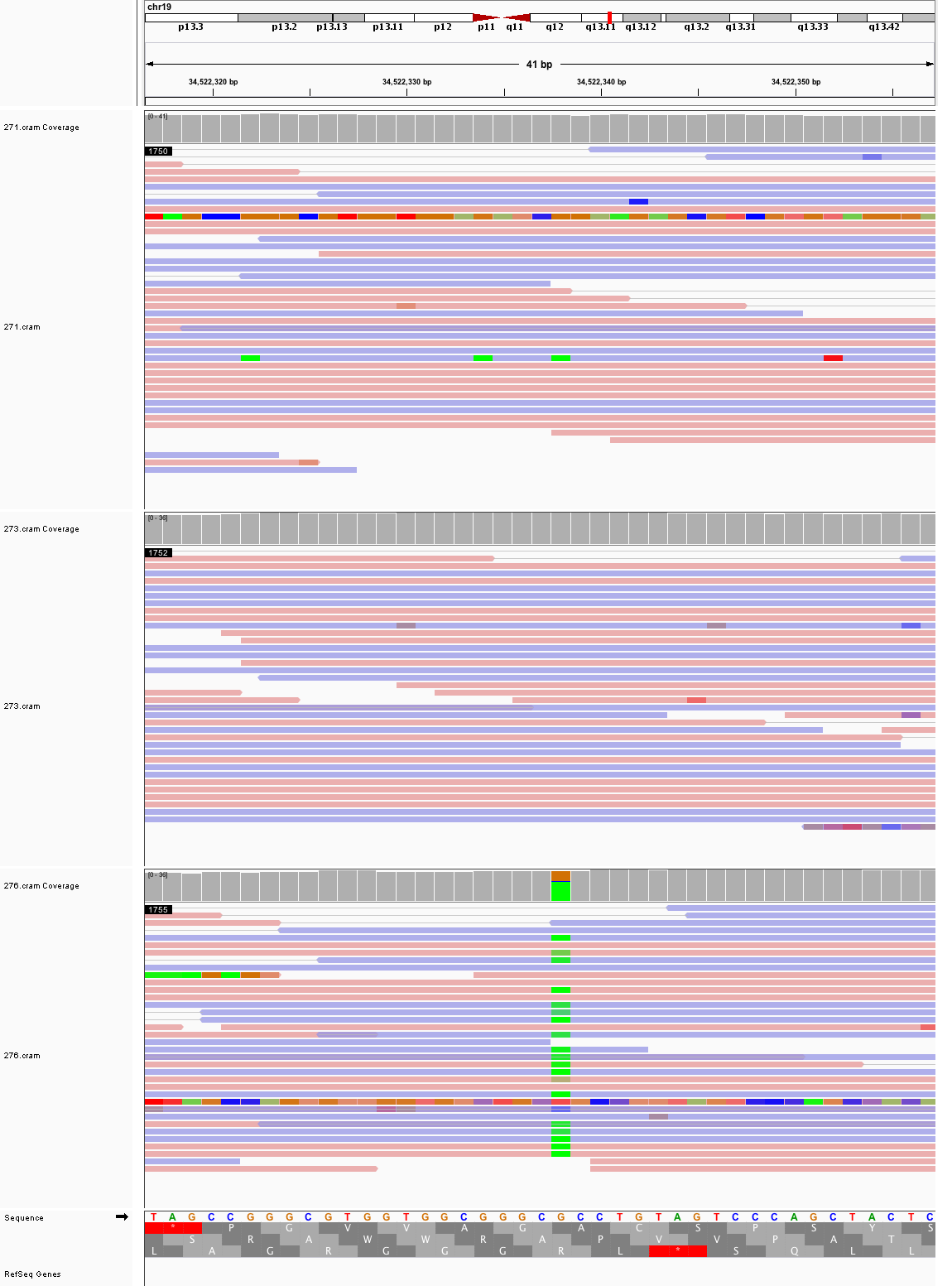

### chr19_44,639,177_44,639,217.png

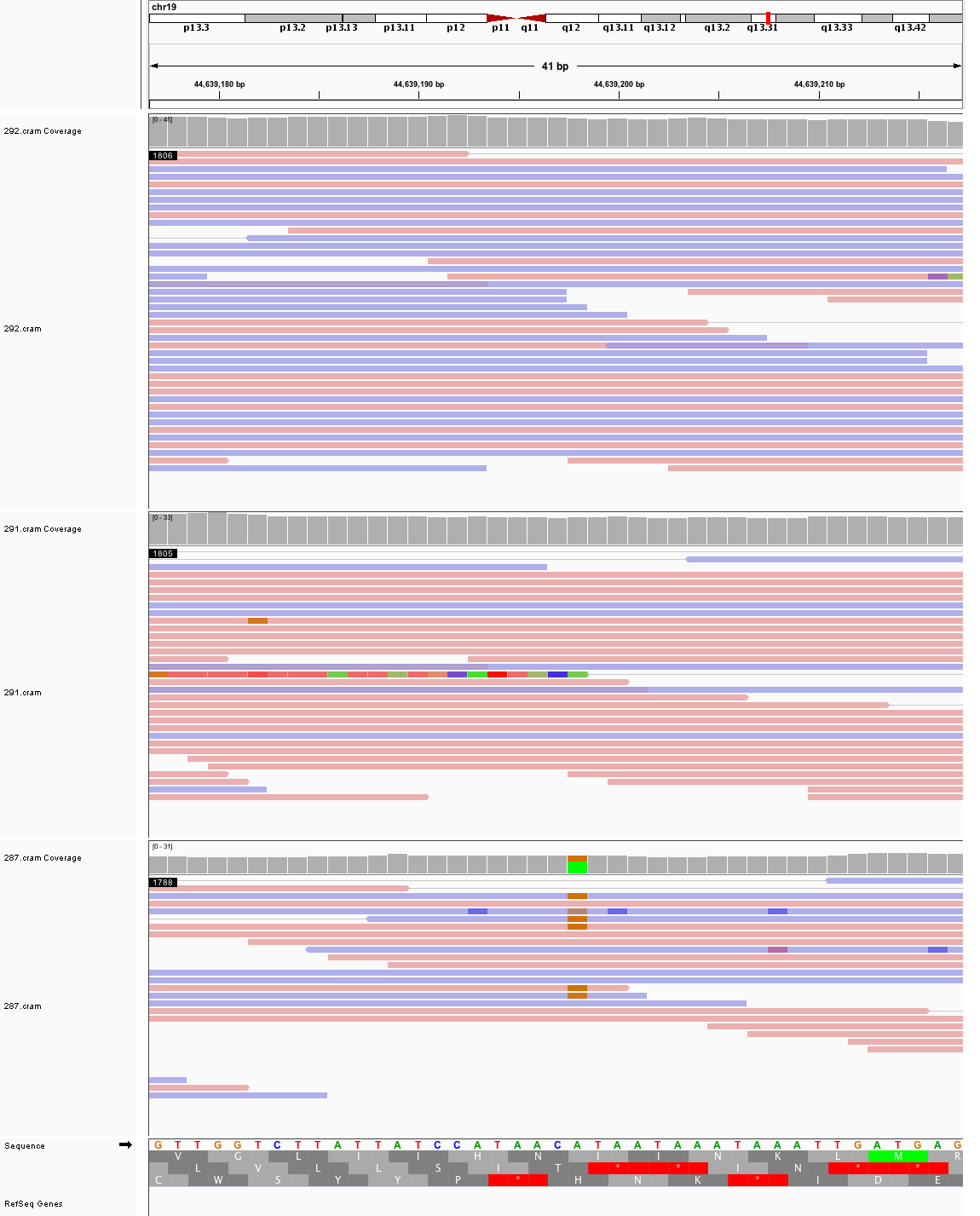

### chr19_48,014,501_48,014,541.png

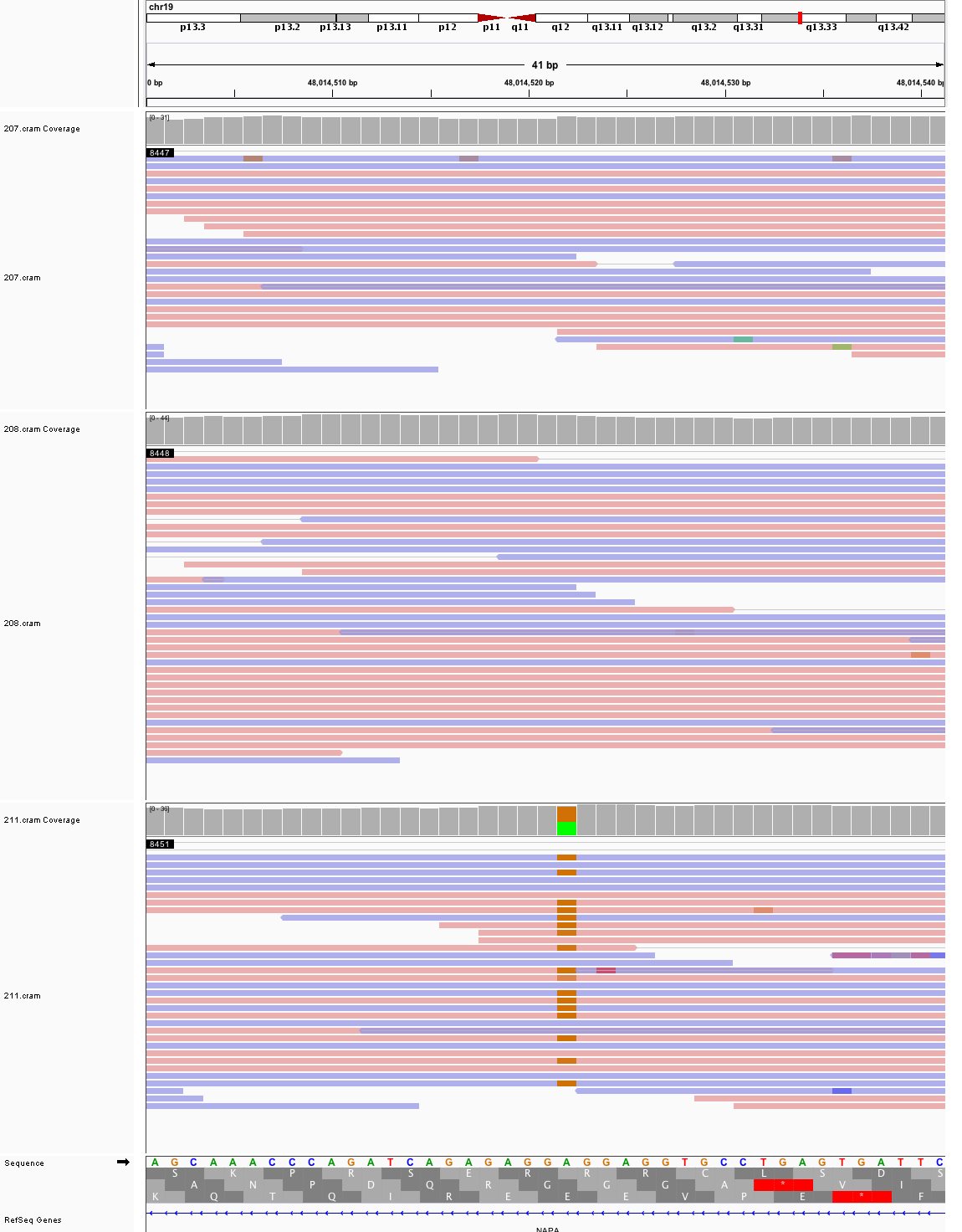

### chr22_41,346,679_41,346,719.png

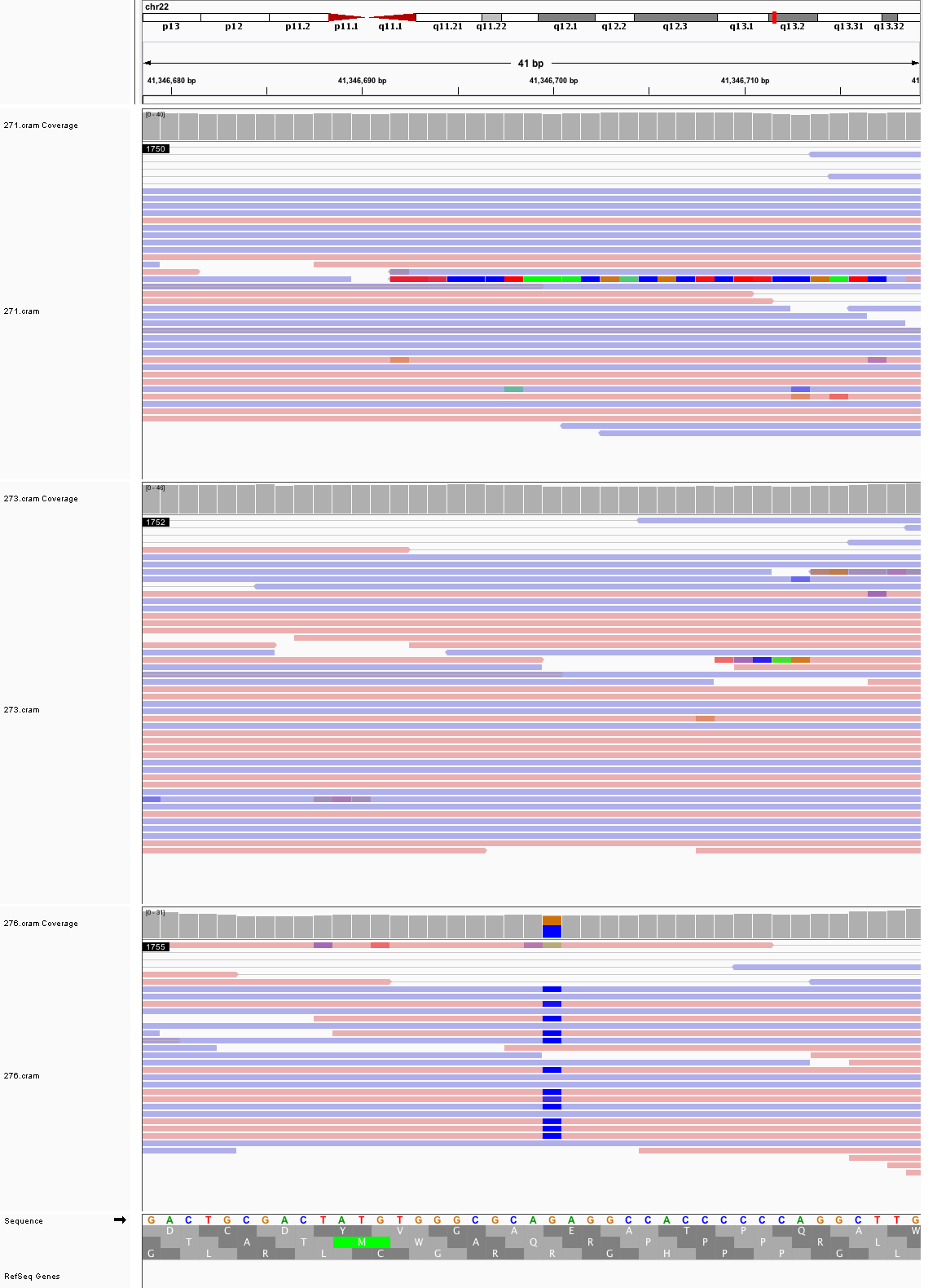

### chrX_15,727,930_15,727,970.png

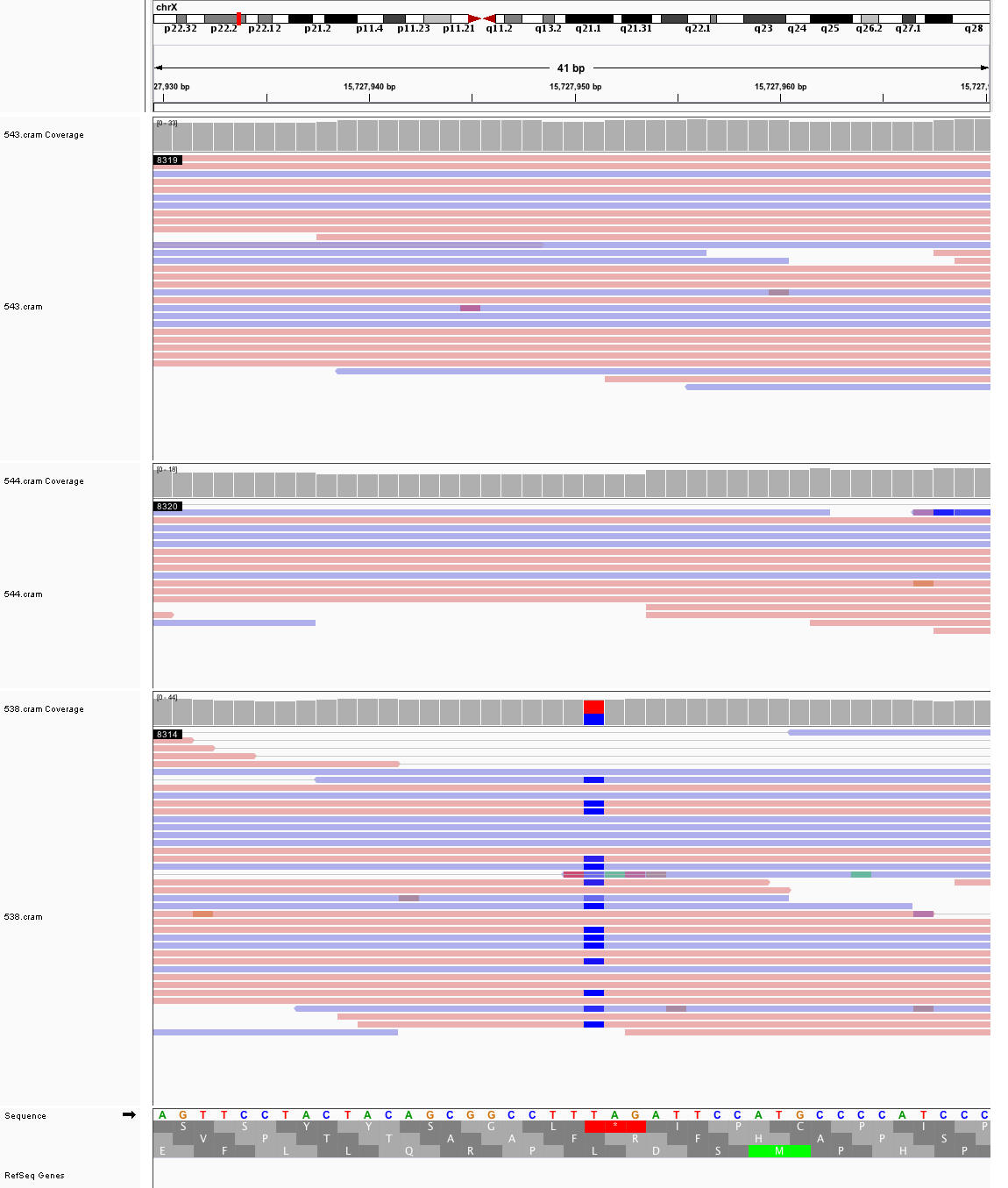

### chrX_50,391,451_50,391,491.png

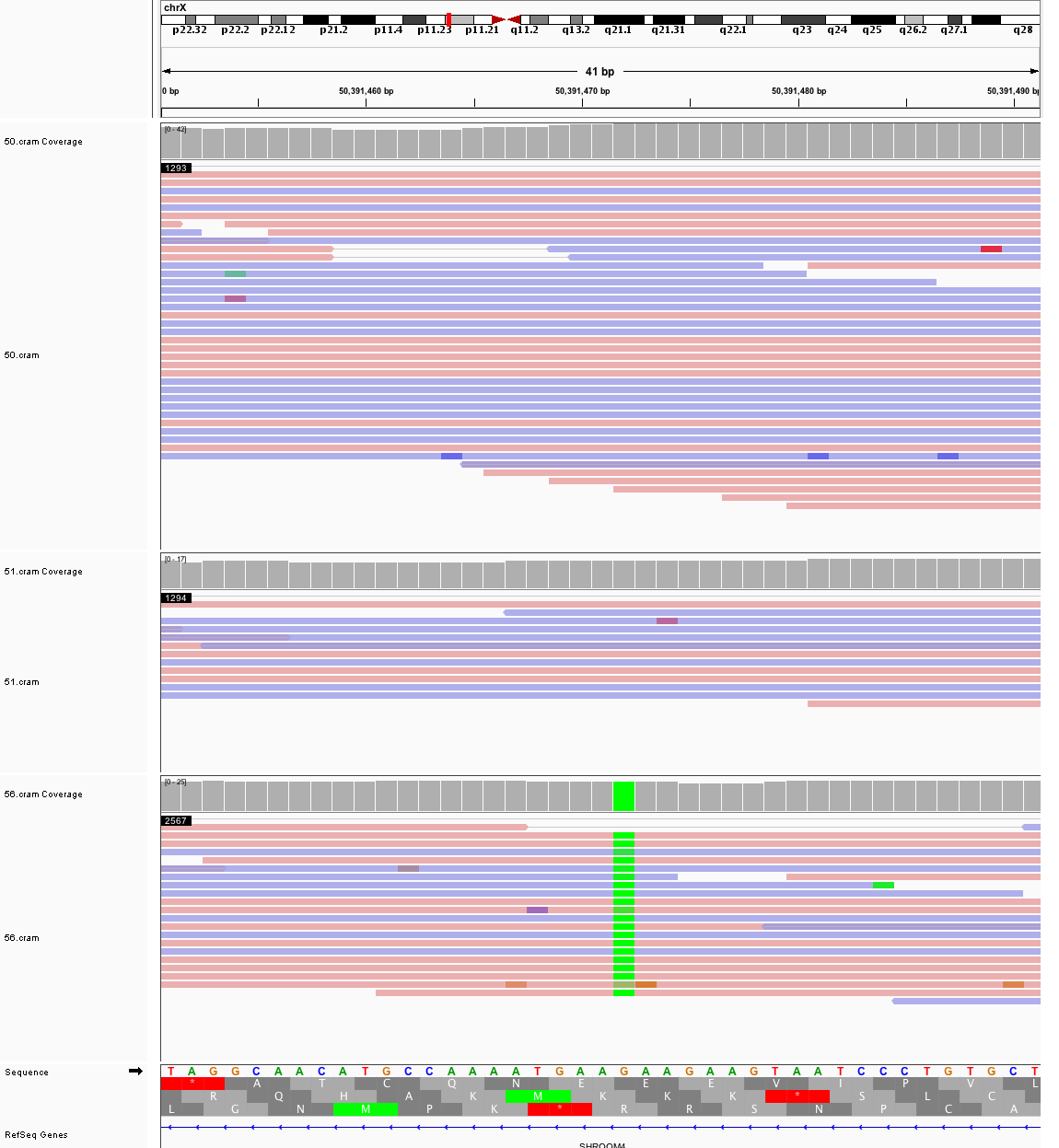
