## Supplementary figures and images for "Large, three-generation CEPH families reveal post-zygotic mosaicism and variability in germline mutation accumulation"

### chr1_178,468,189_178,468,229.png

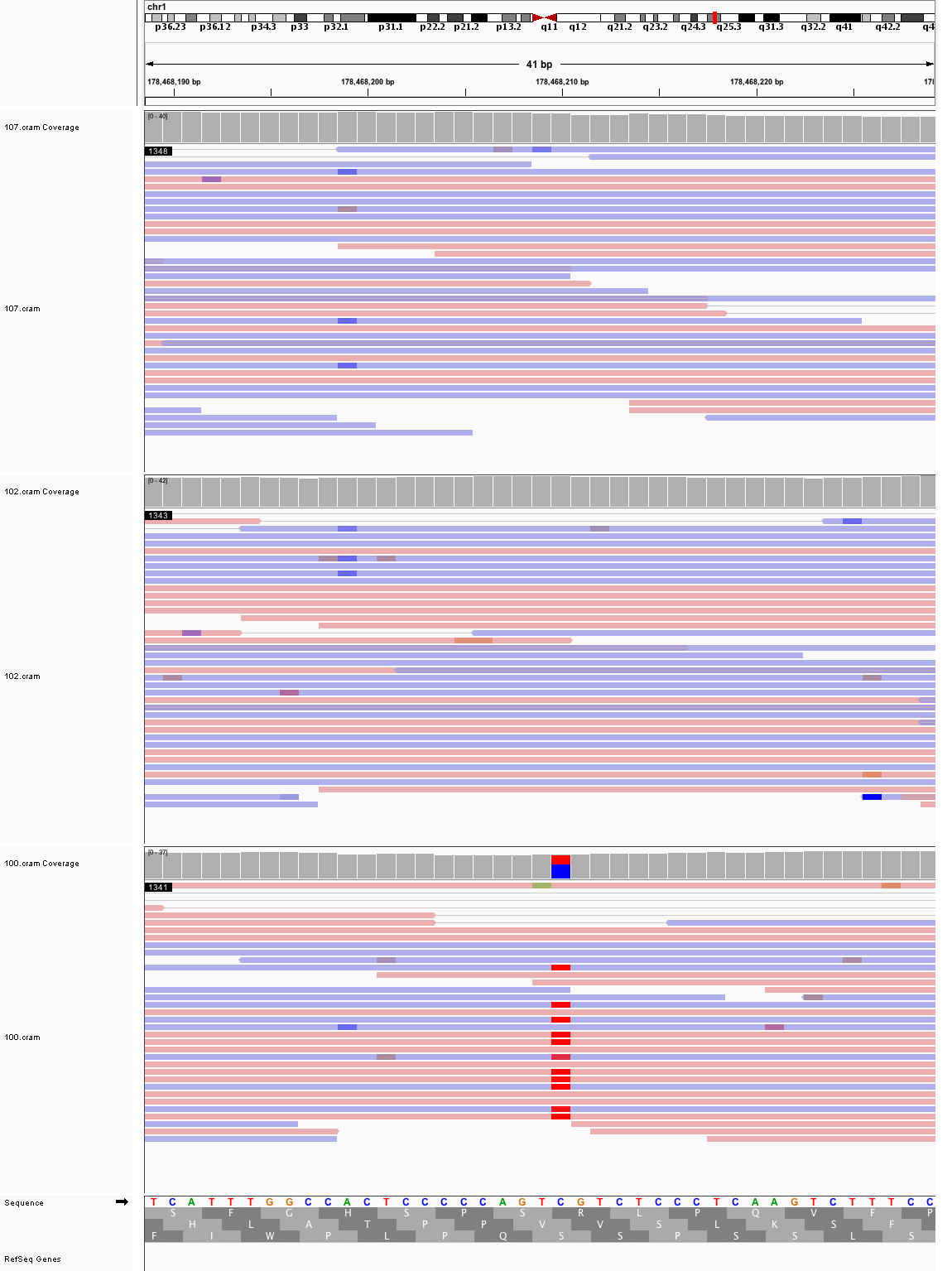

### chr1_202,099,419_202,099,459.png

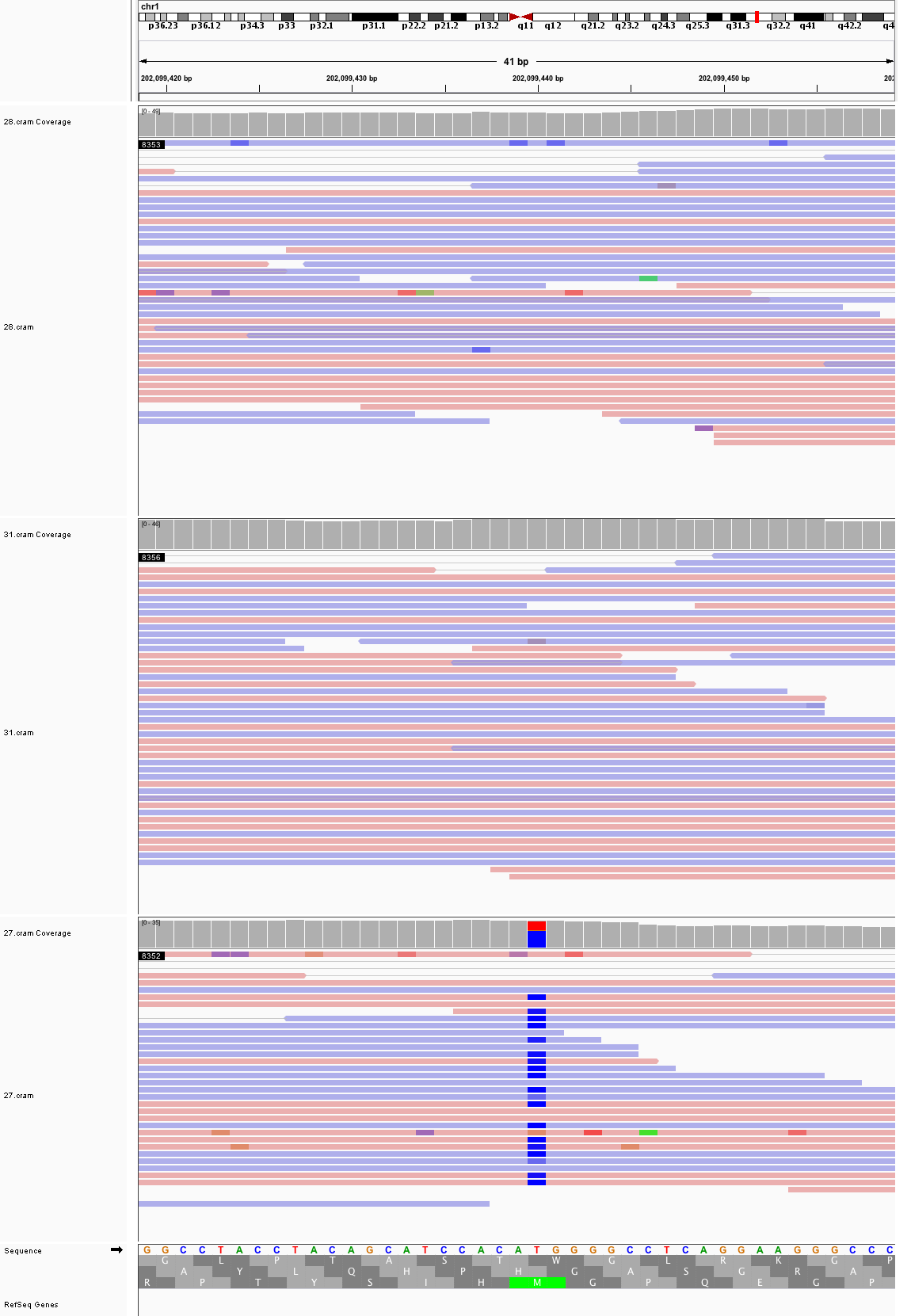

### chr1_226,654,620_226,654,660.png

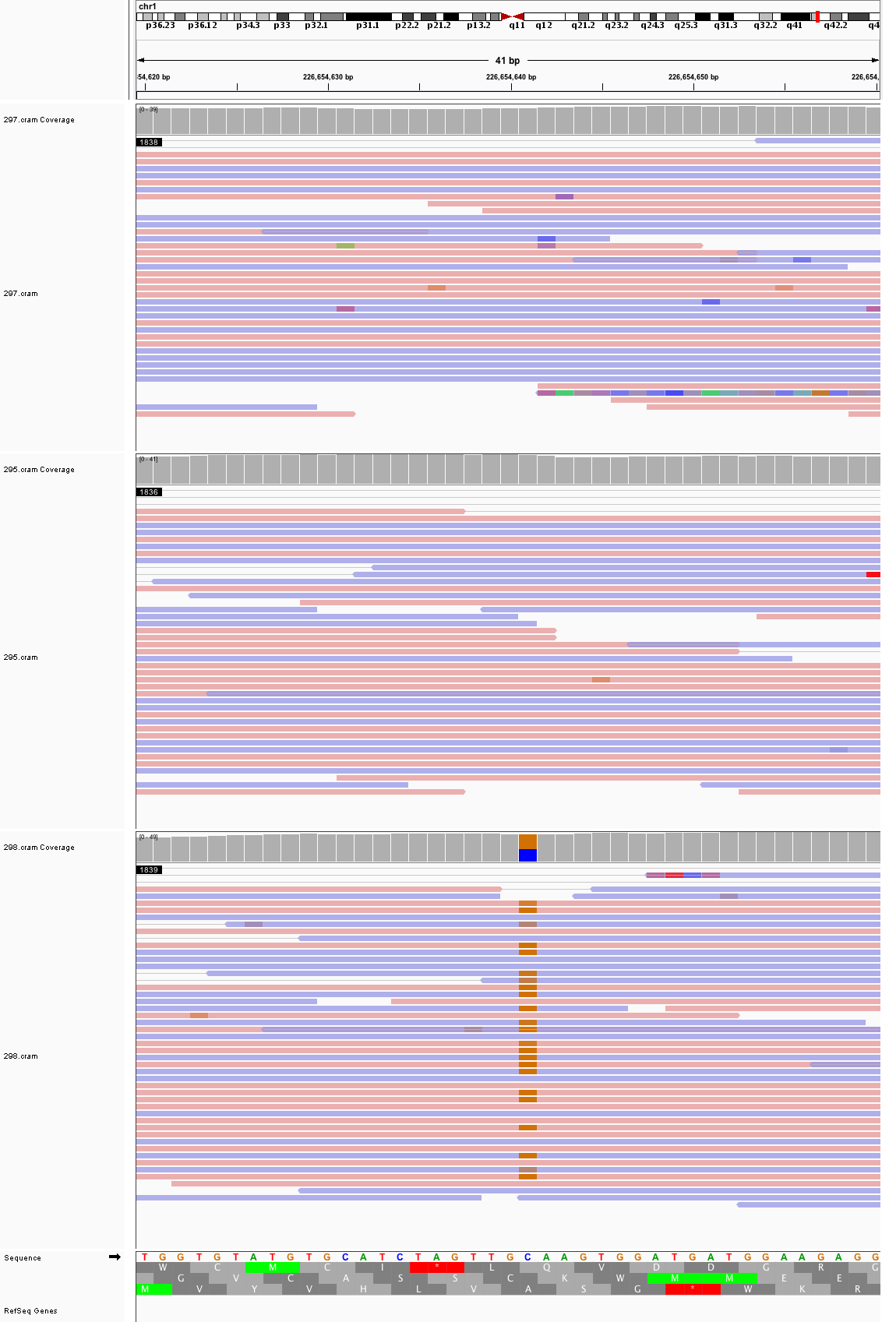

### chr1_235,909,366_235,909,406.png

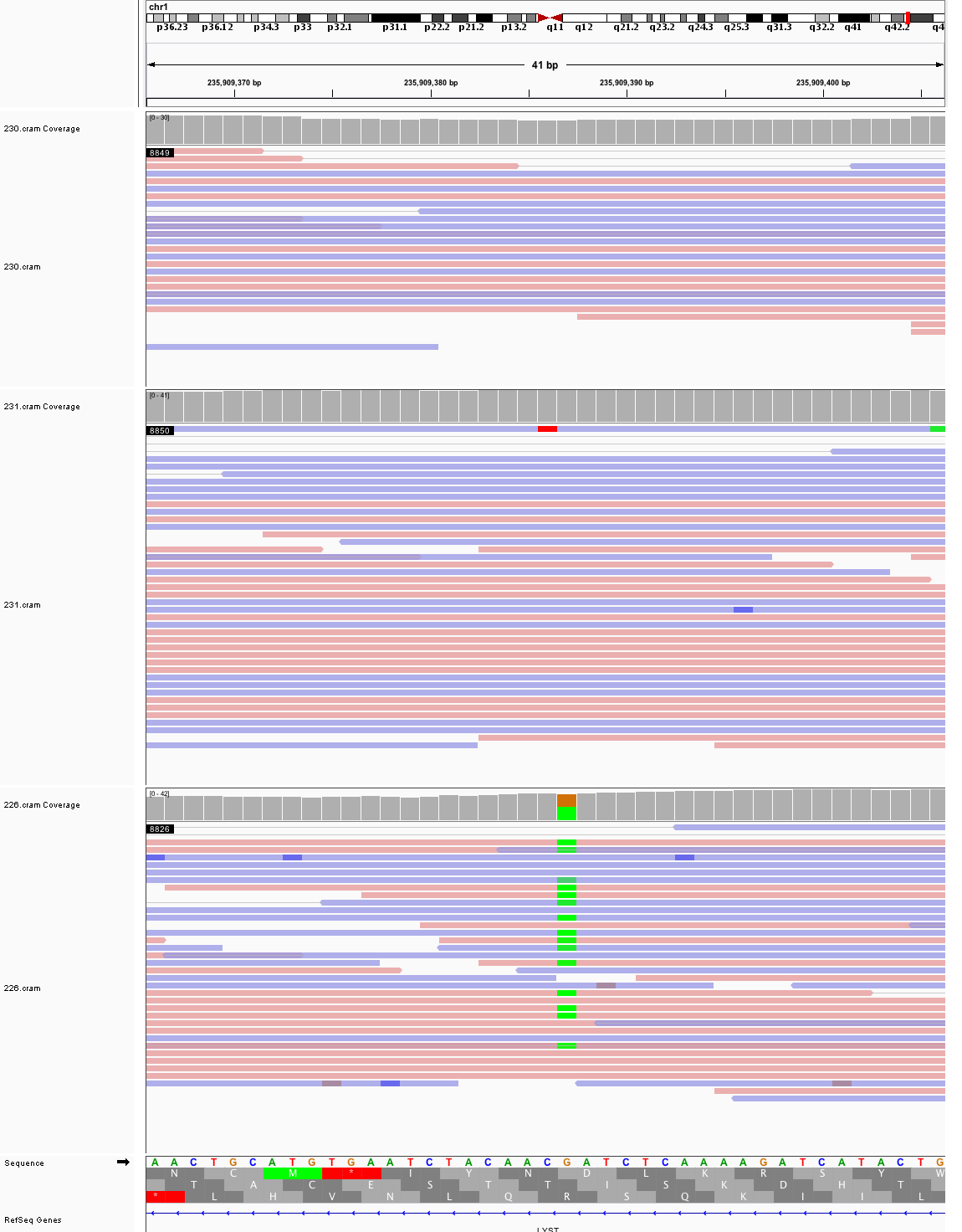

### chr1_238,651,017_238,651,057.png

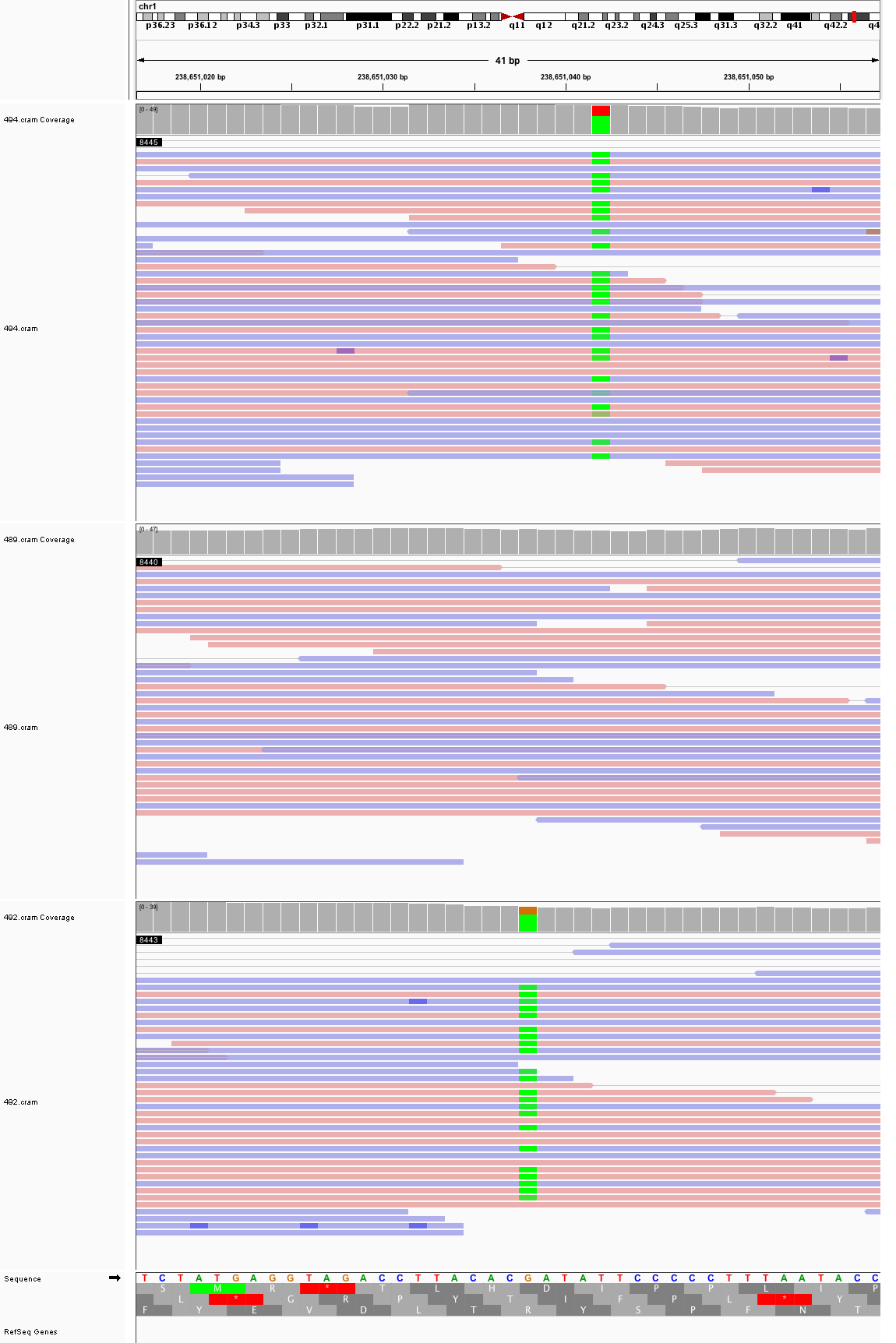

### chr2_58,508,130_58,508,170.png

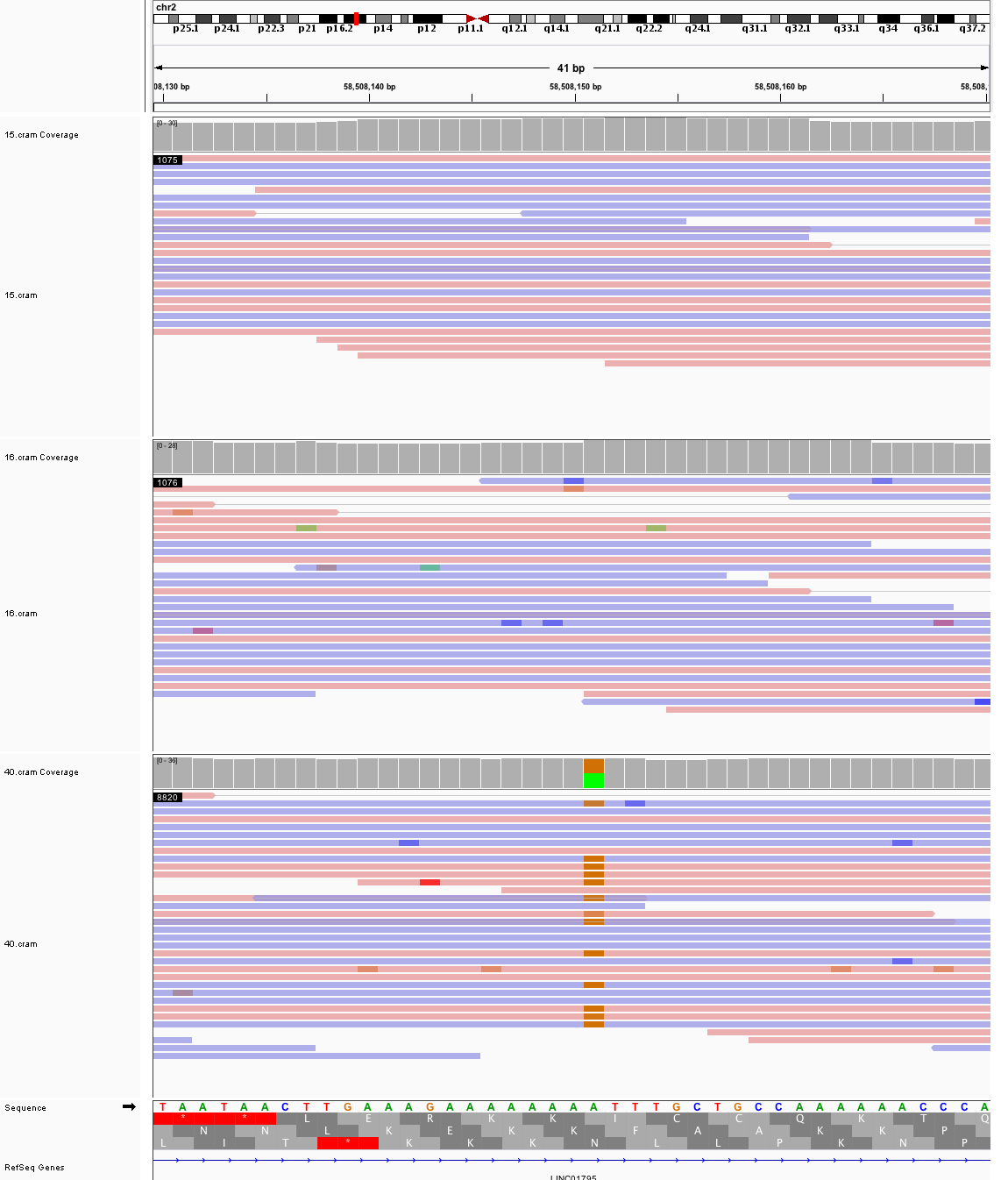

### chr2_100,714,918_100,714,958.png

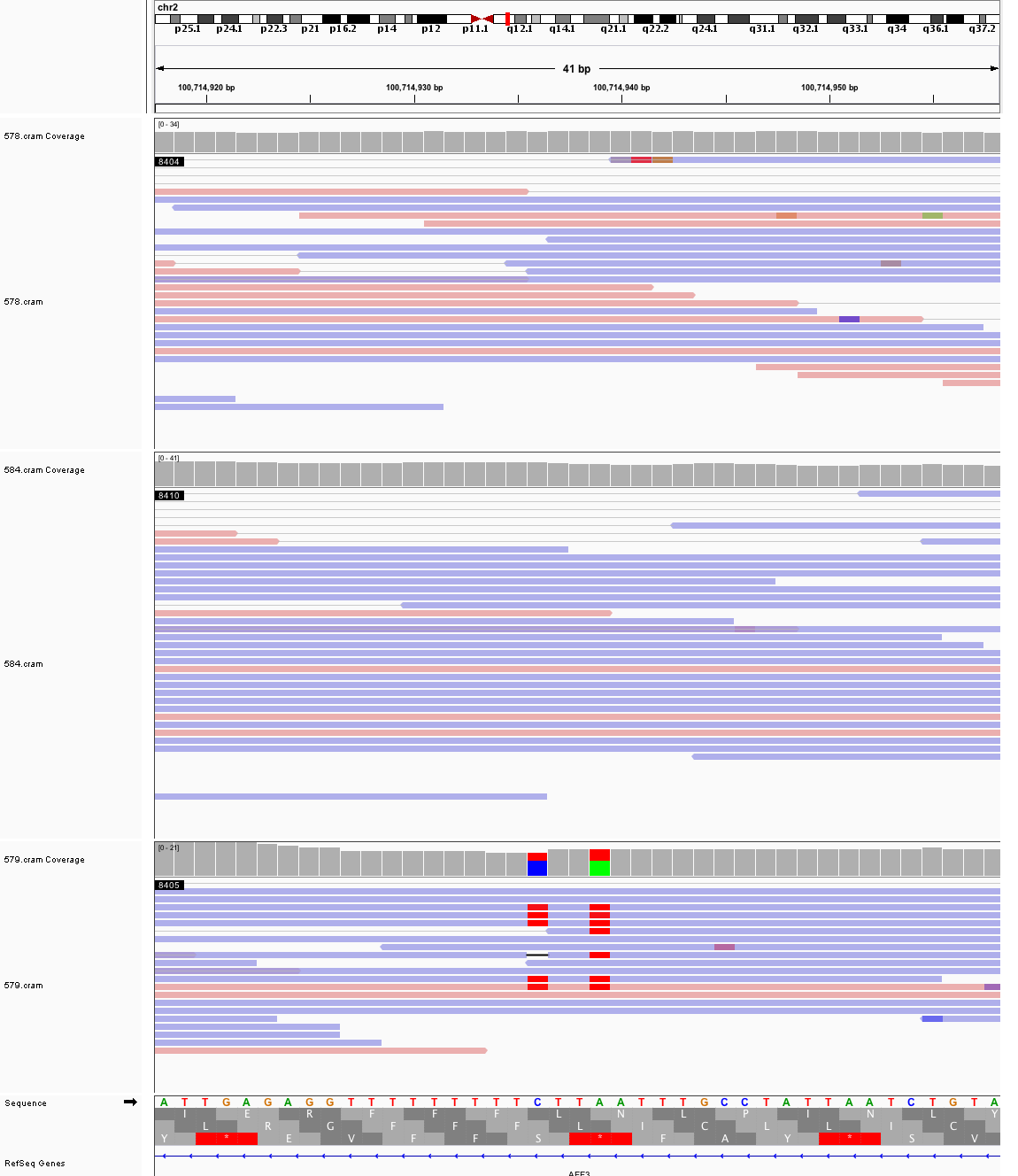

### chr2_101,541,898_101,541,938.png

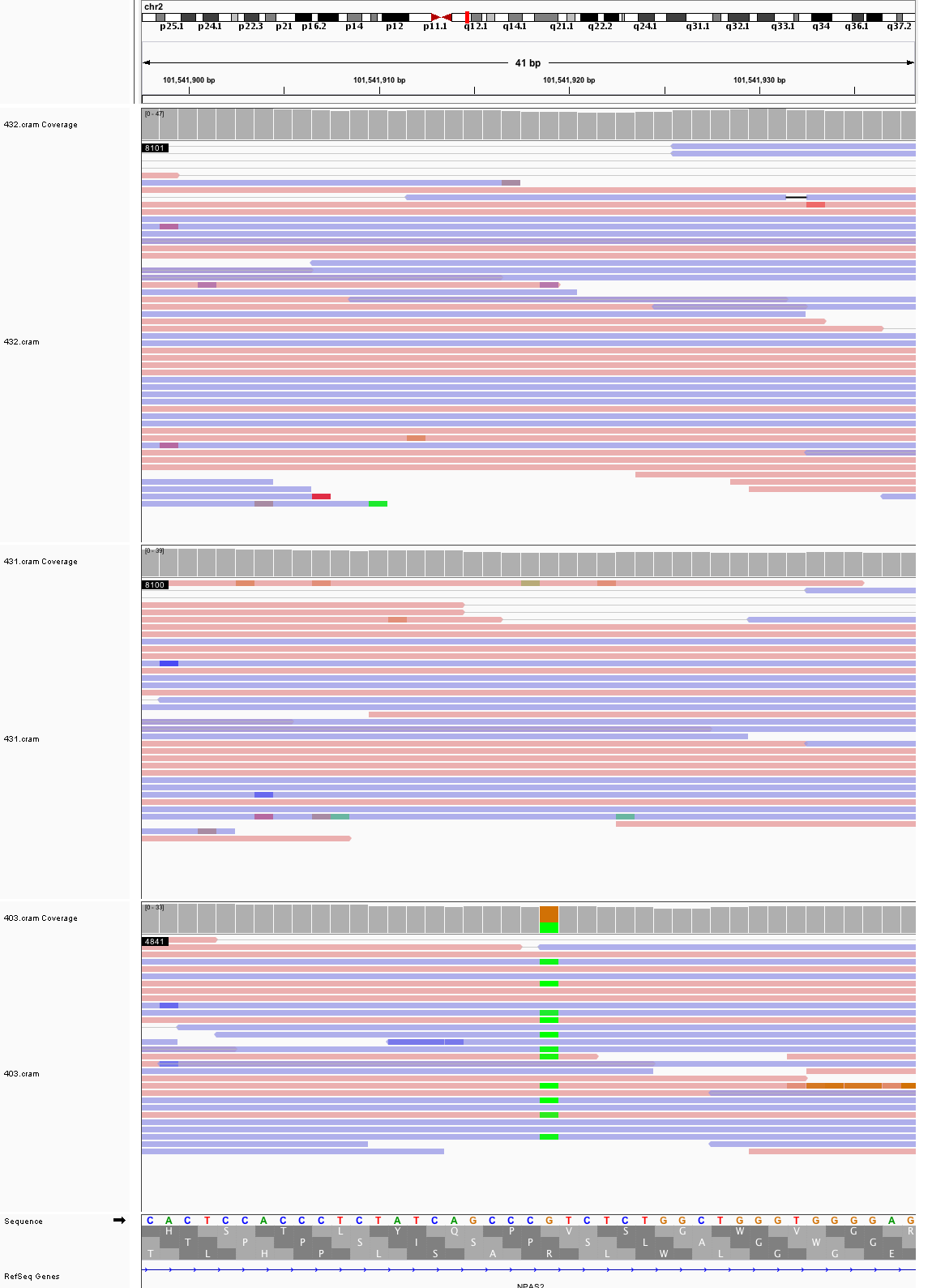

### chr2_103,500,693_103,500,733.png

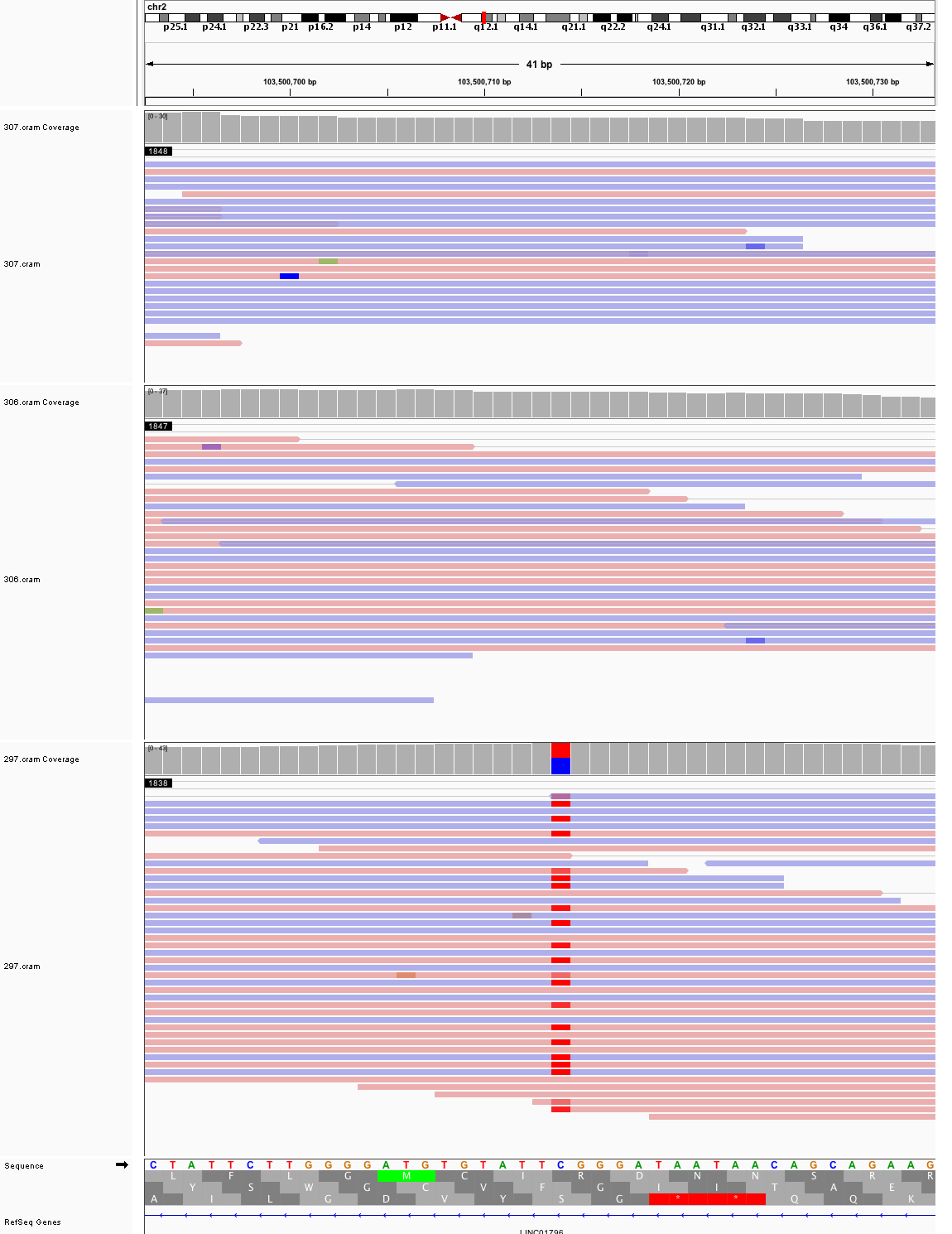

### chr2_123,491,858_123,491,898.png

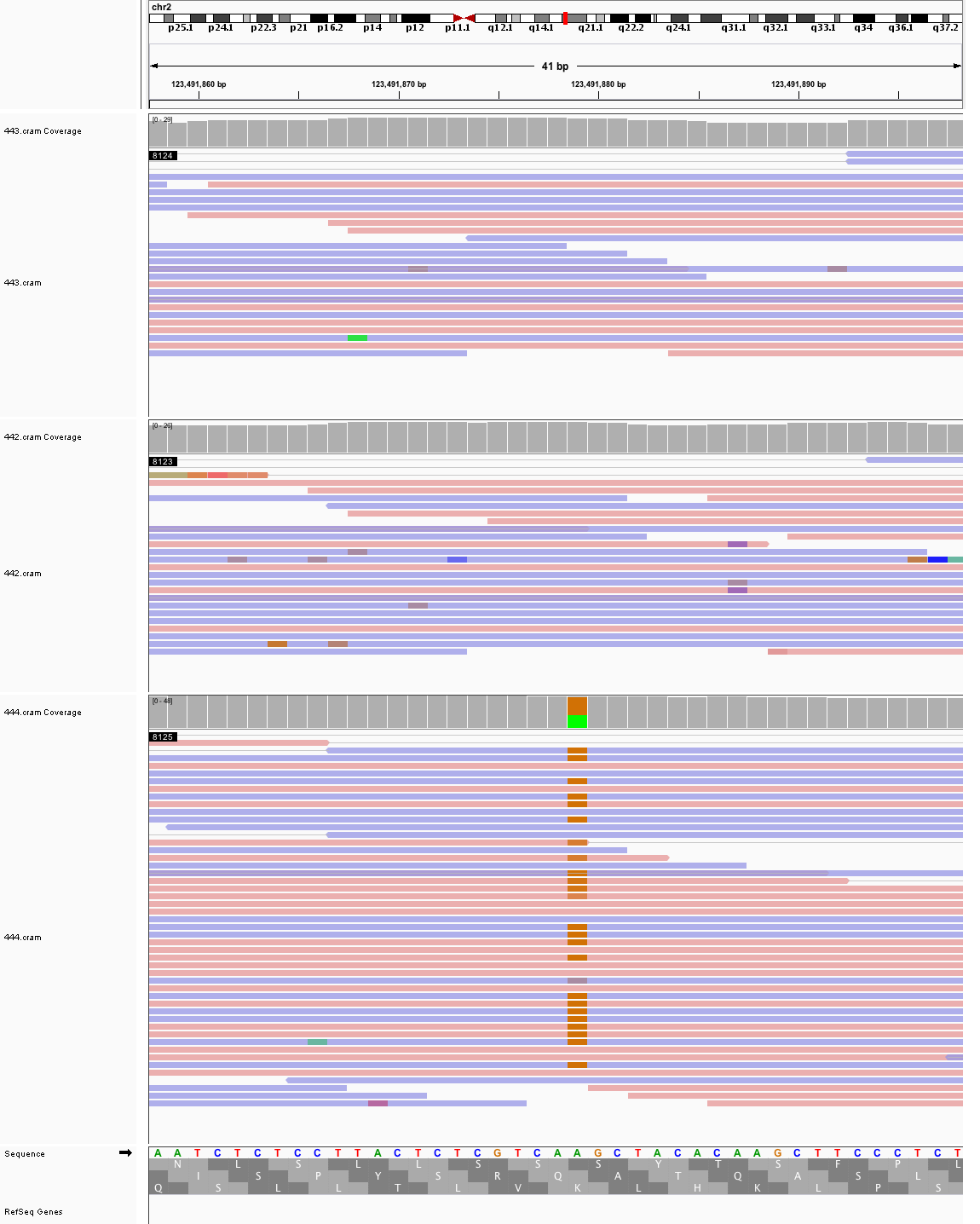

### chr2_138,139,012_138,139,052.png

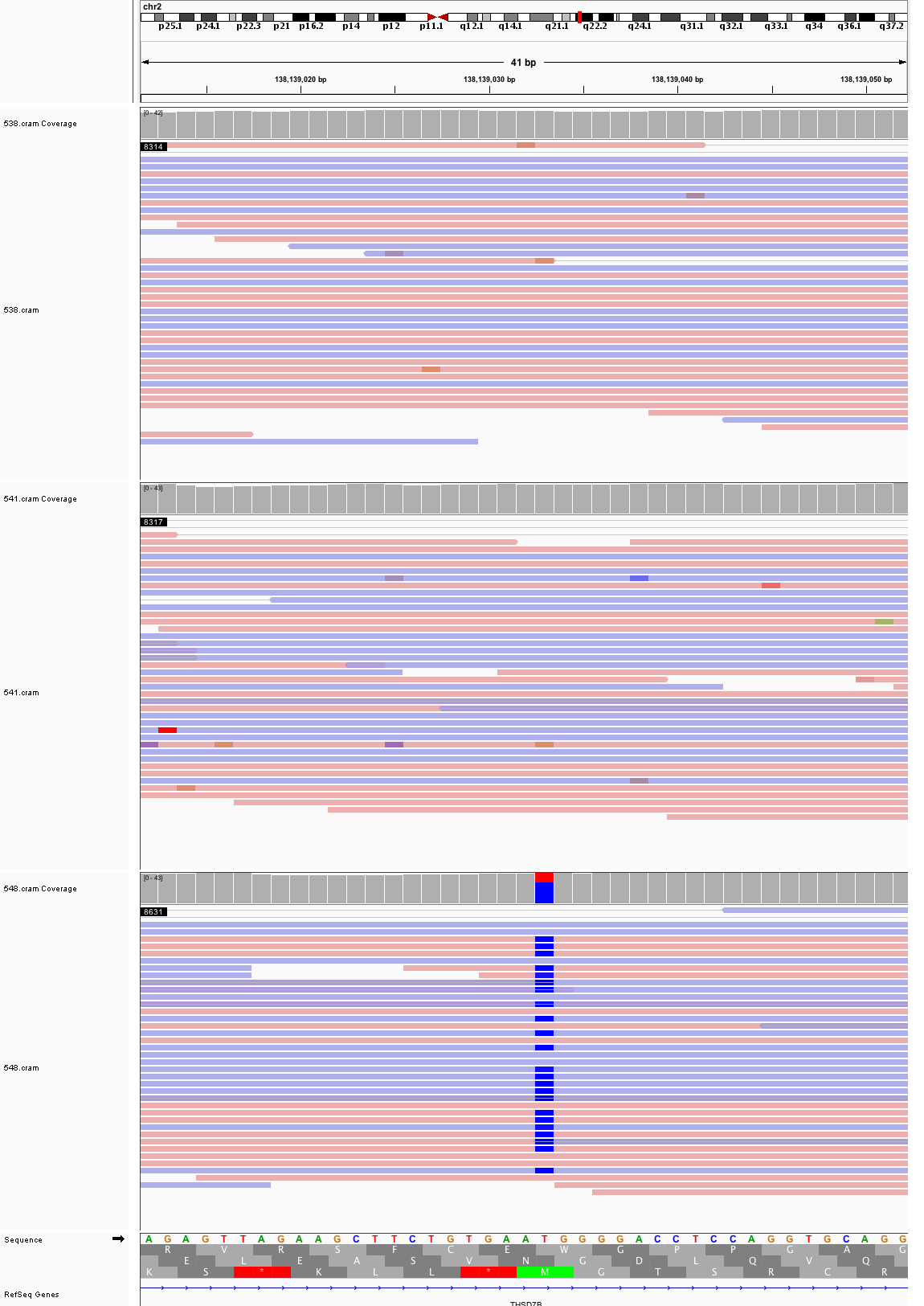

### chr2_164,024,421_164,024,461.png

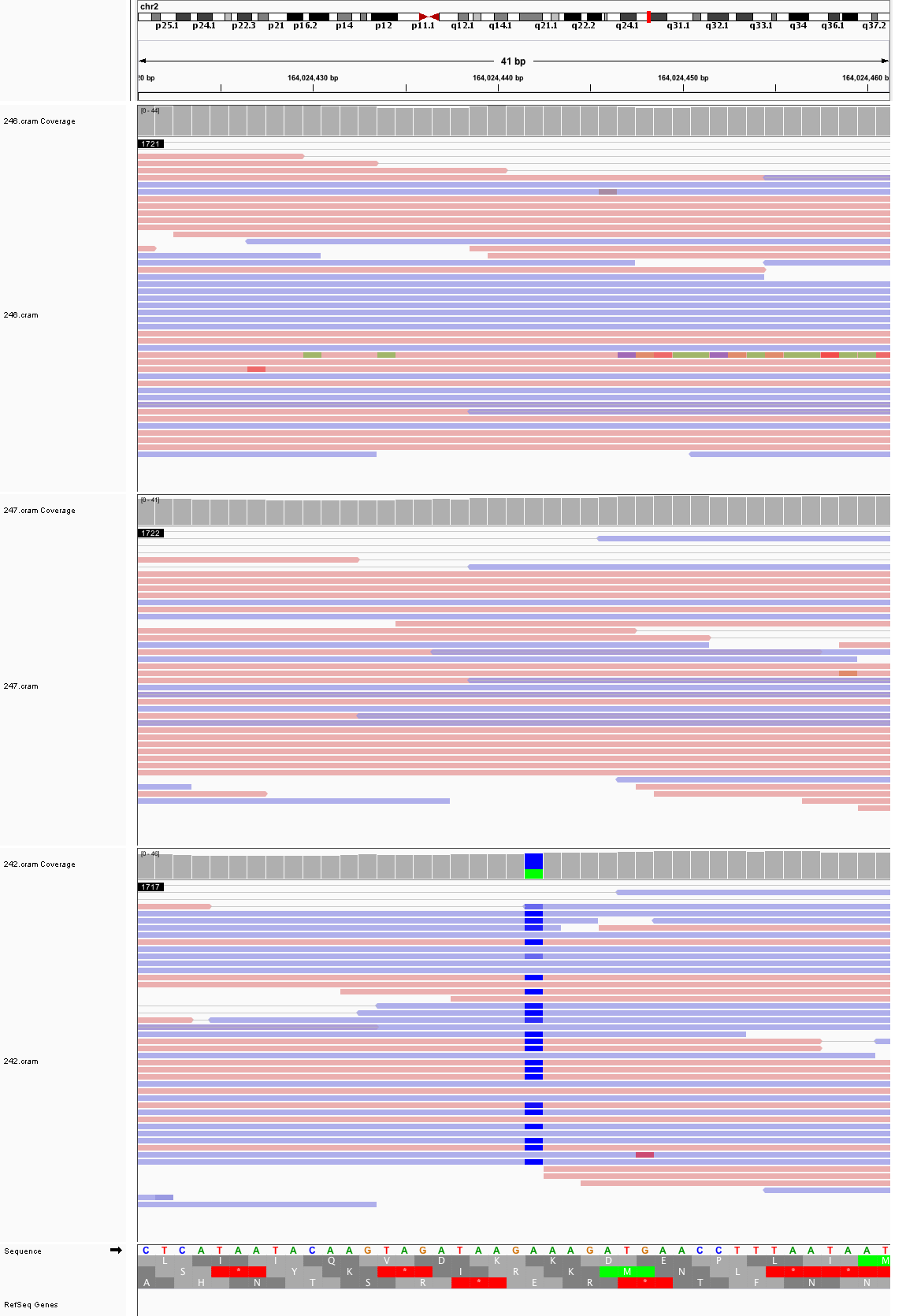

### chr3_23,148,751_23,148,791.png

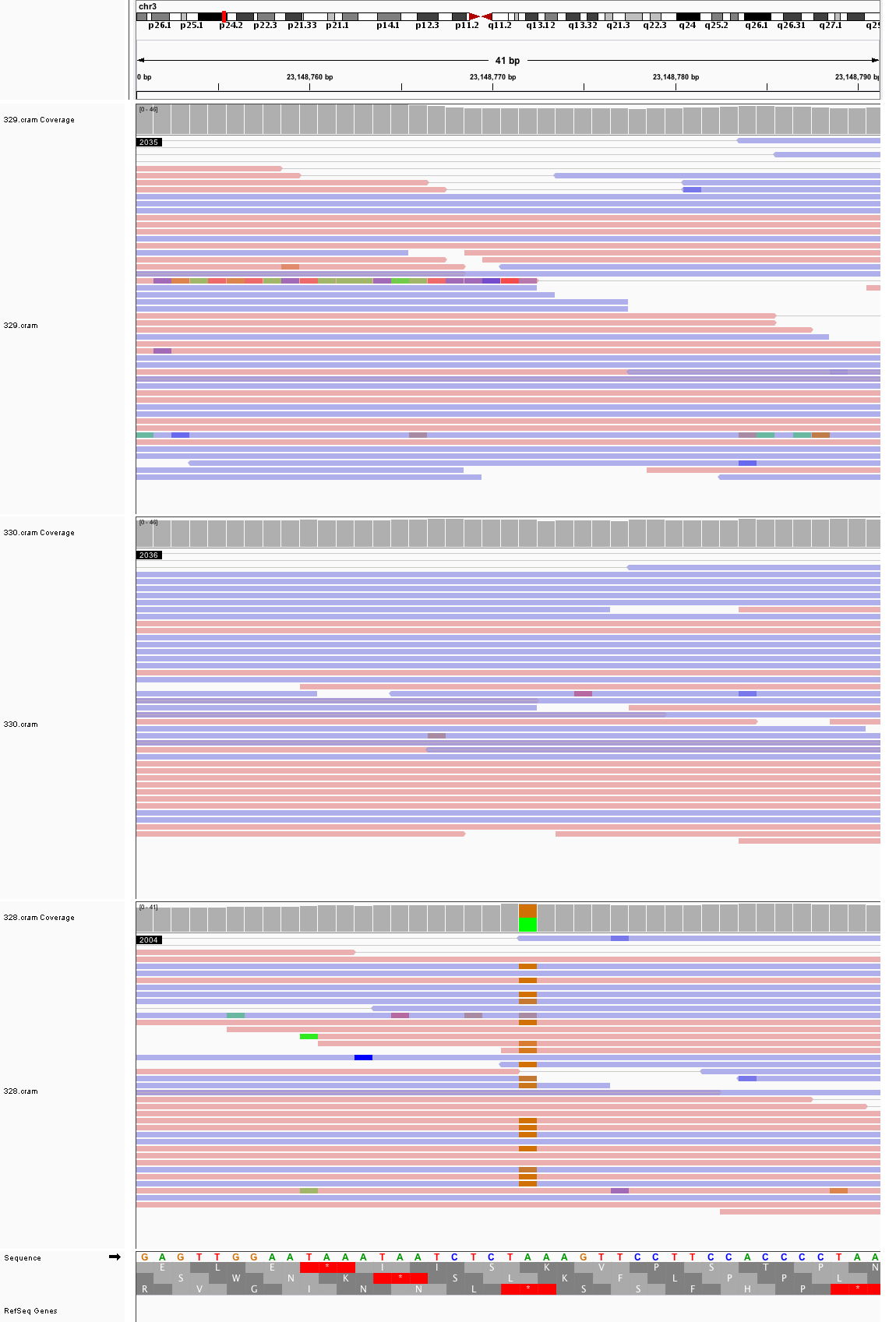

### chr3_34,561,128_34,561,168.png

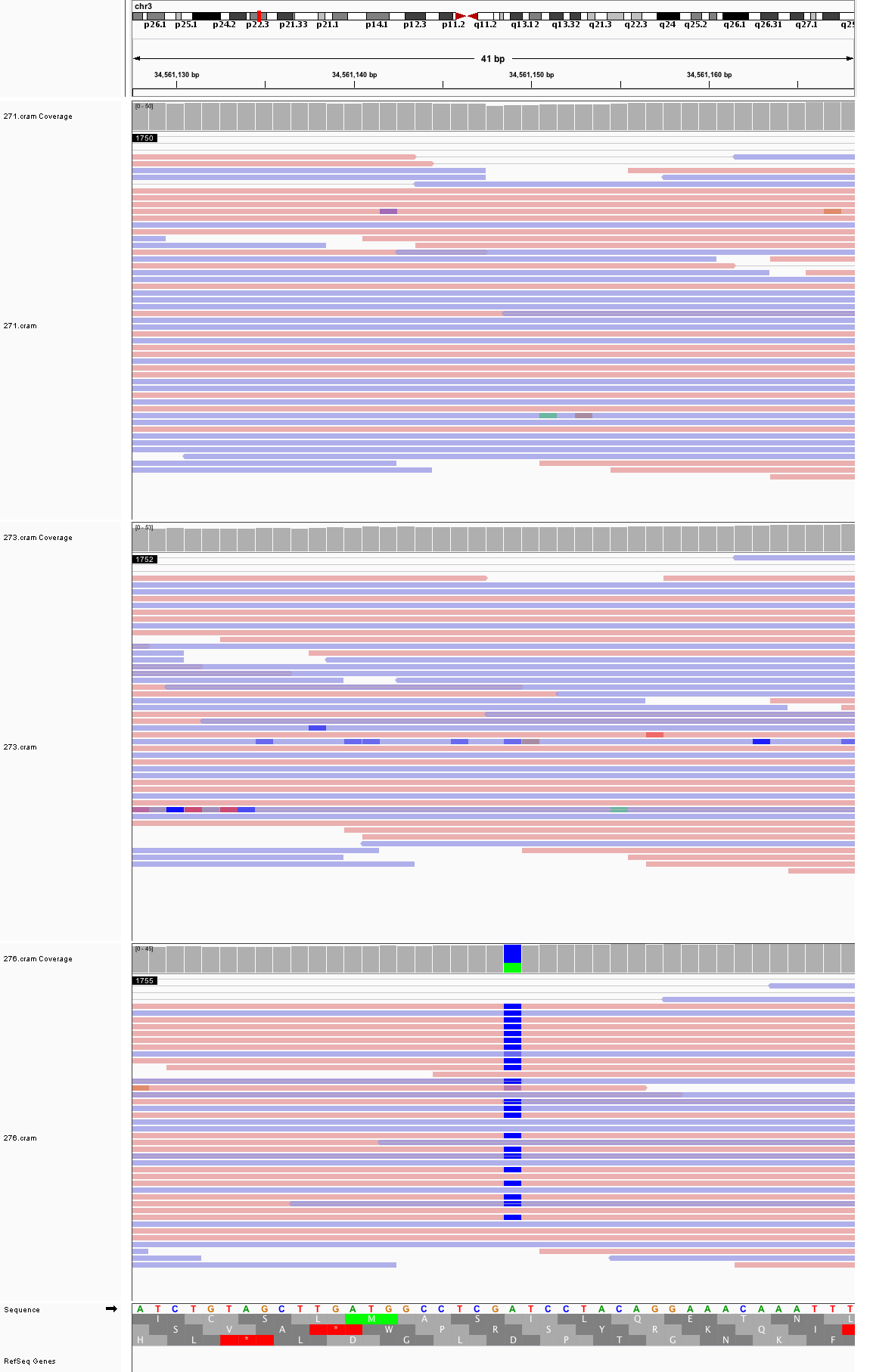

### chr3_38,799,470_38,799,510.png

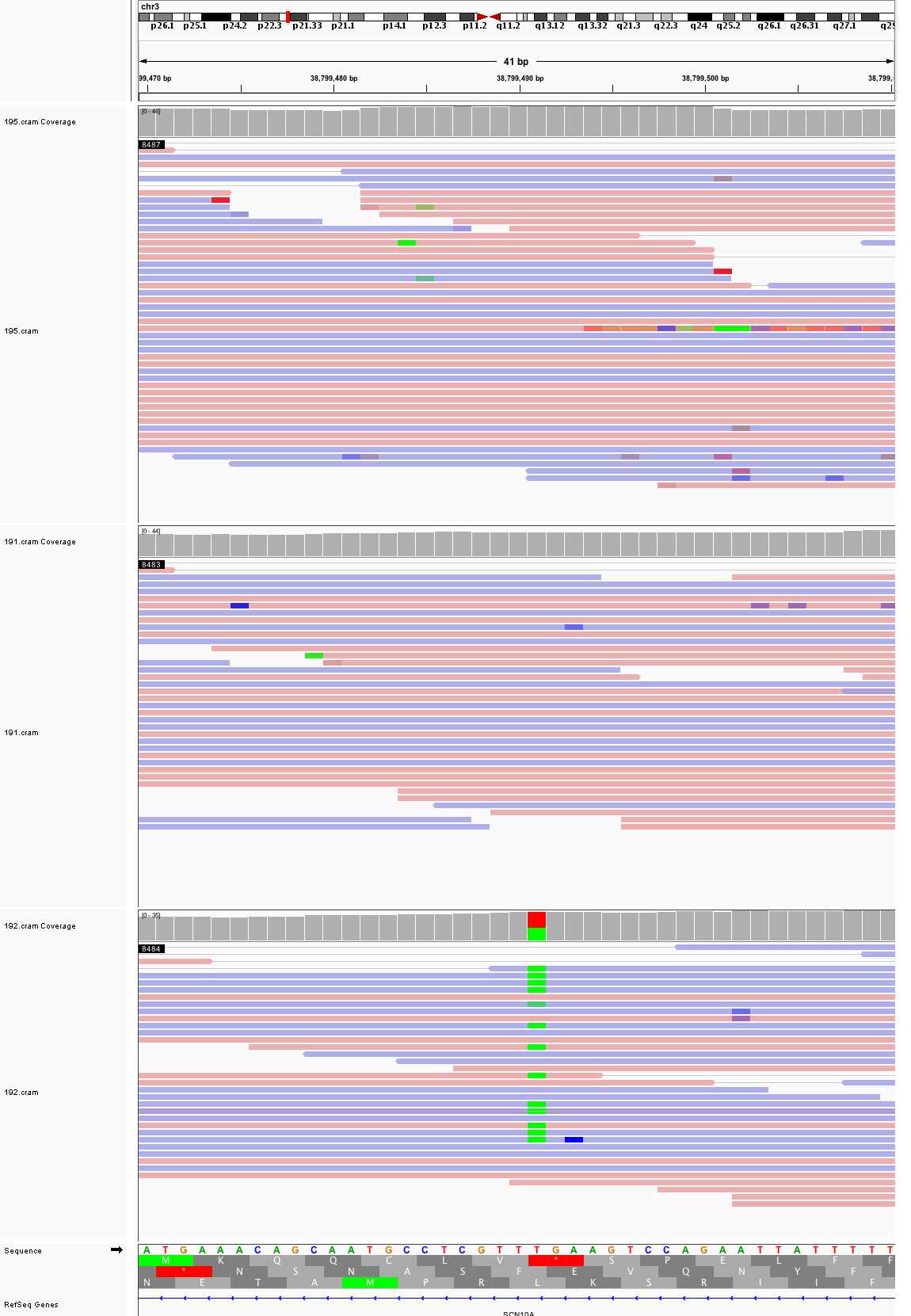

### chr3_45,036,320_45,036,360.png

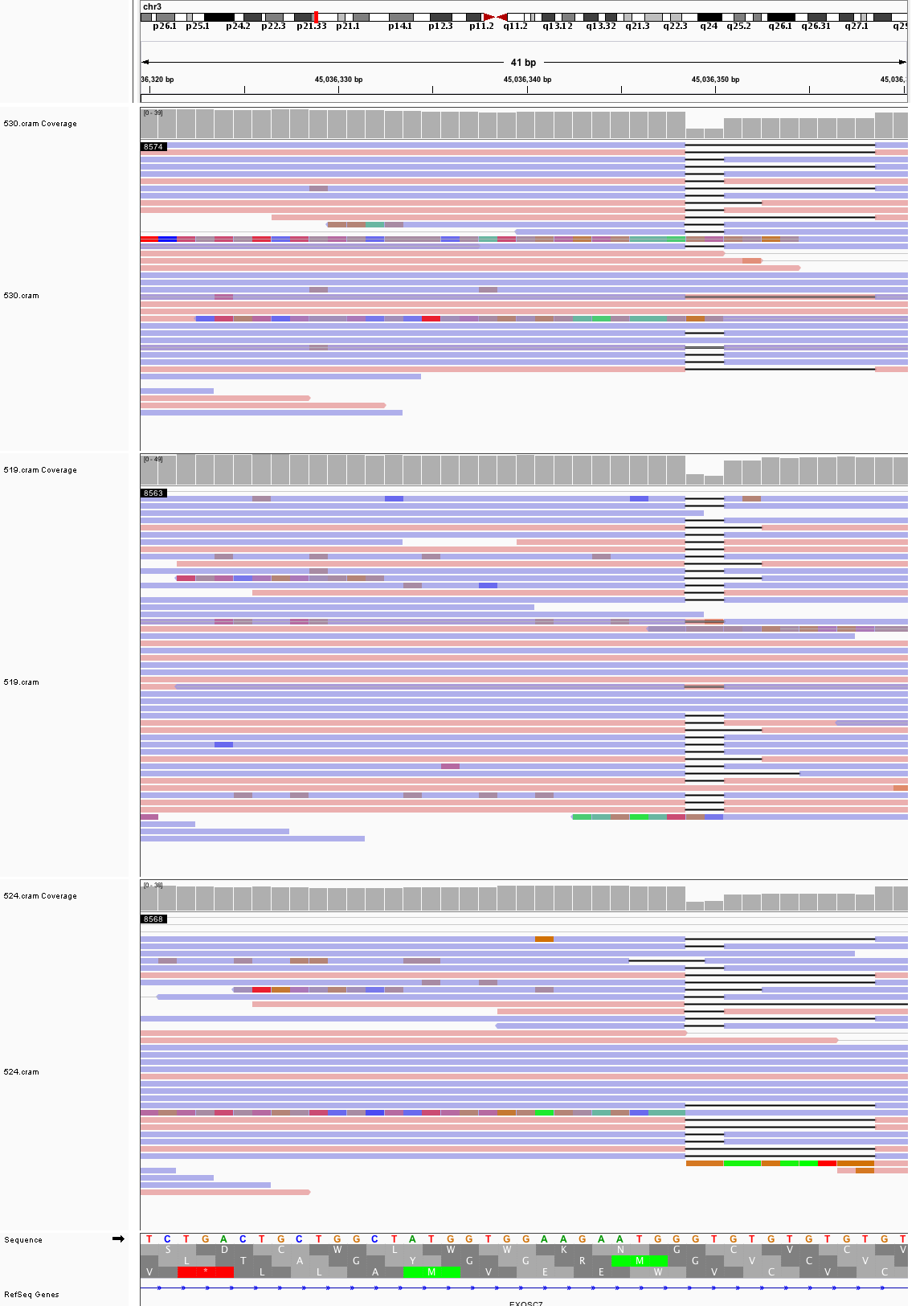

### chr3_45,586,868_45,586,908.png

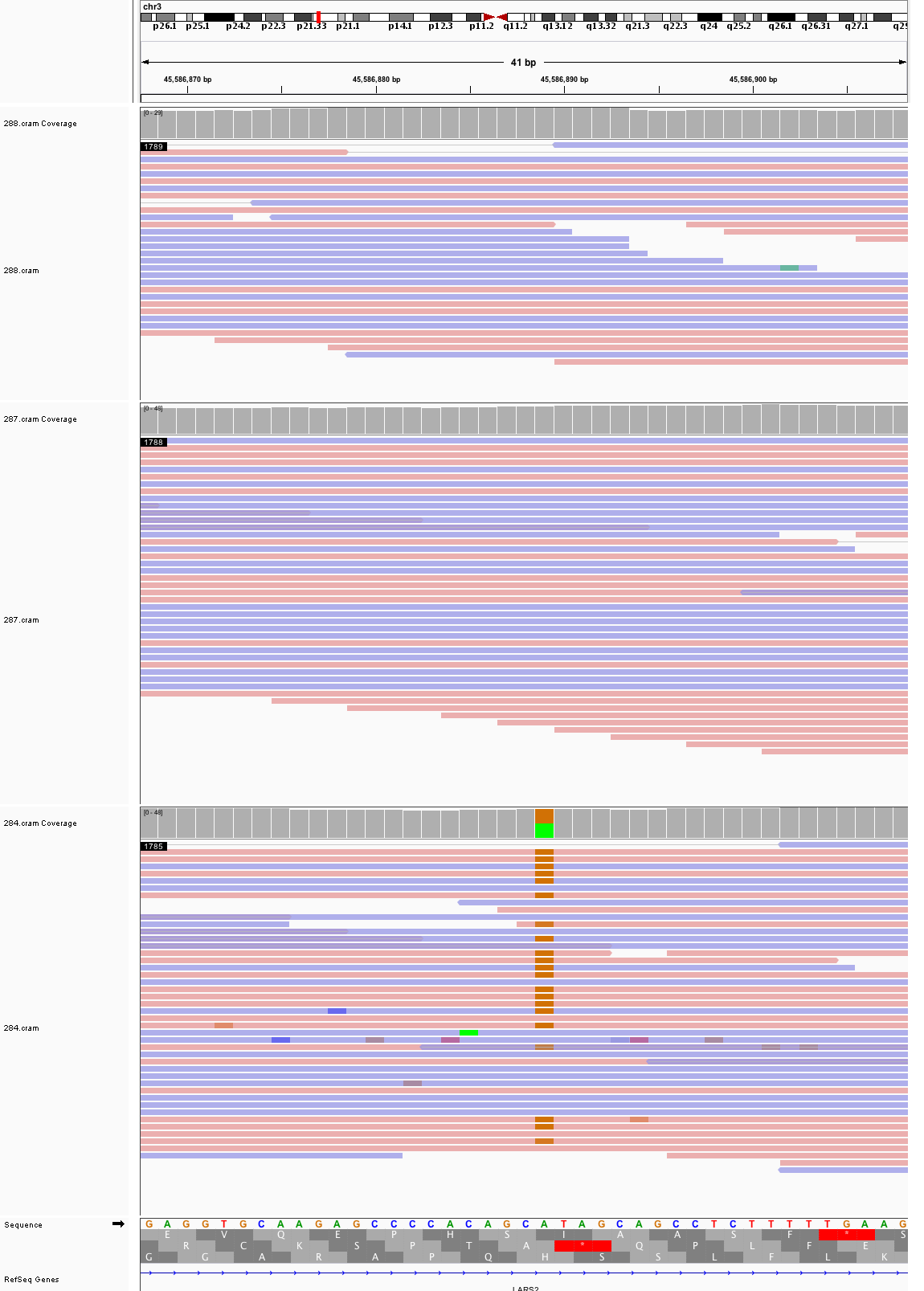

### chr3_78,634,981_78,635,021.png

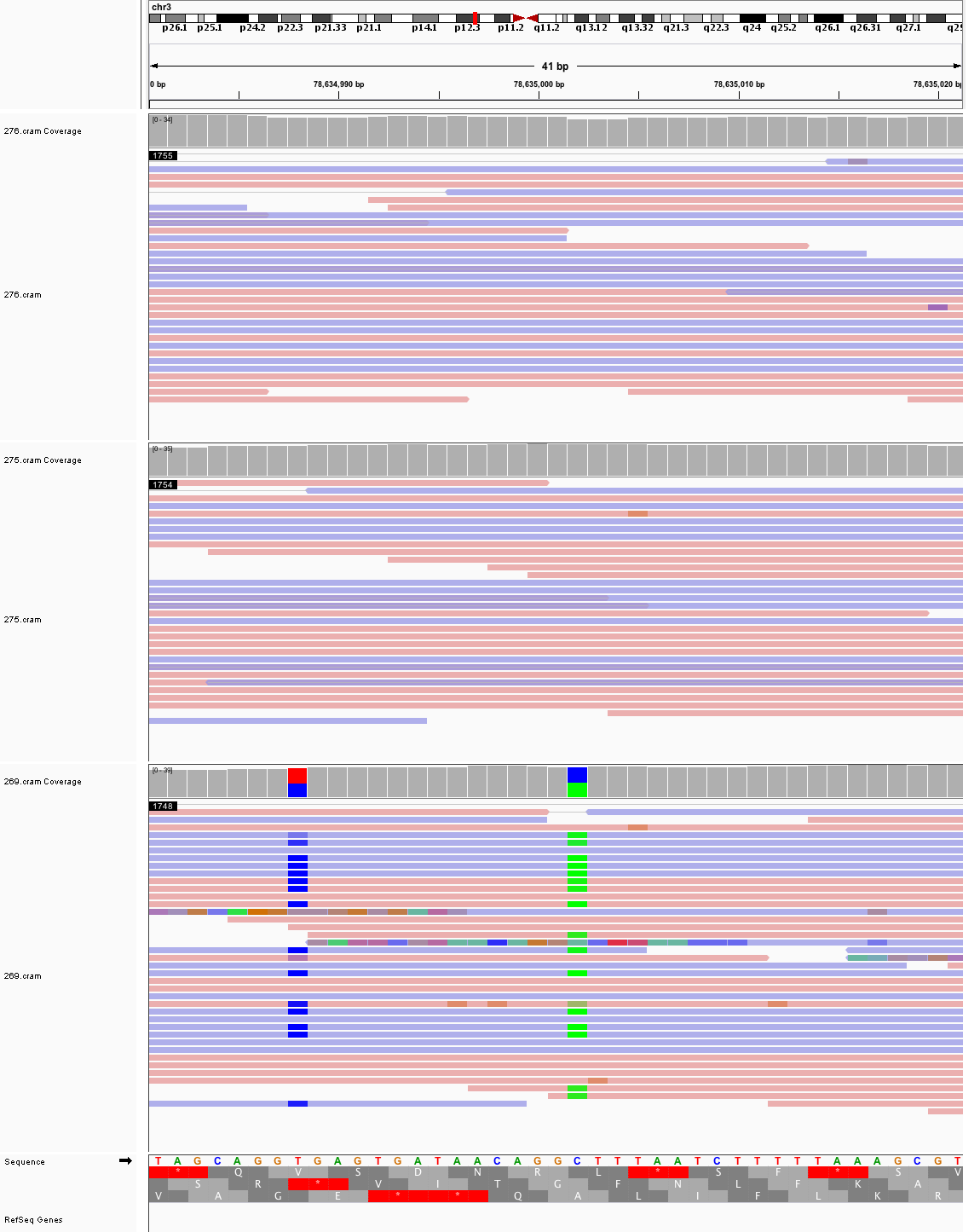

### chr3_94,599,682_94,599,722.png

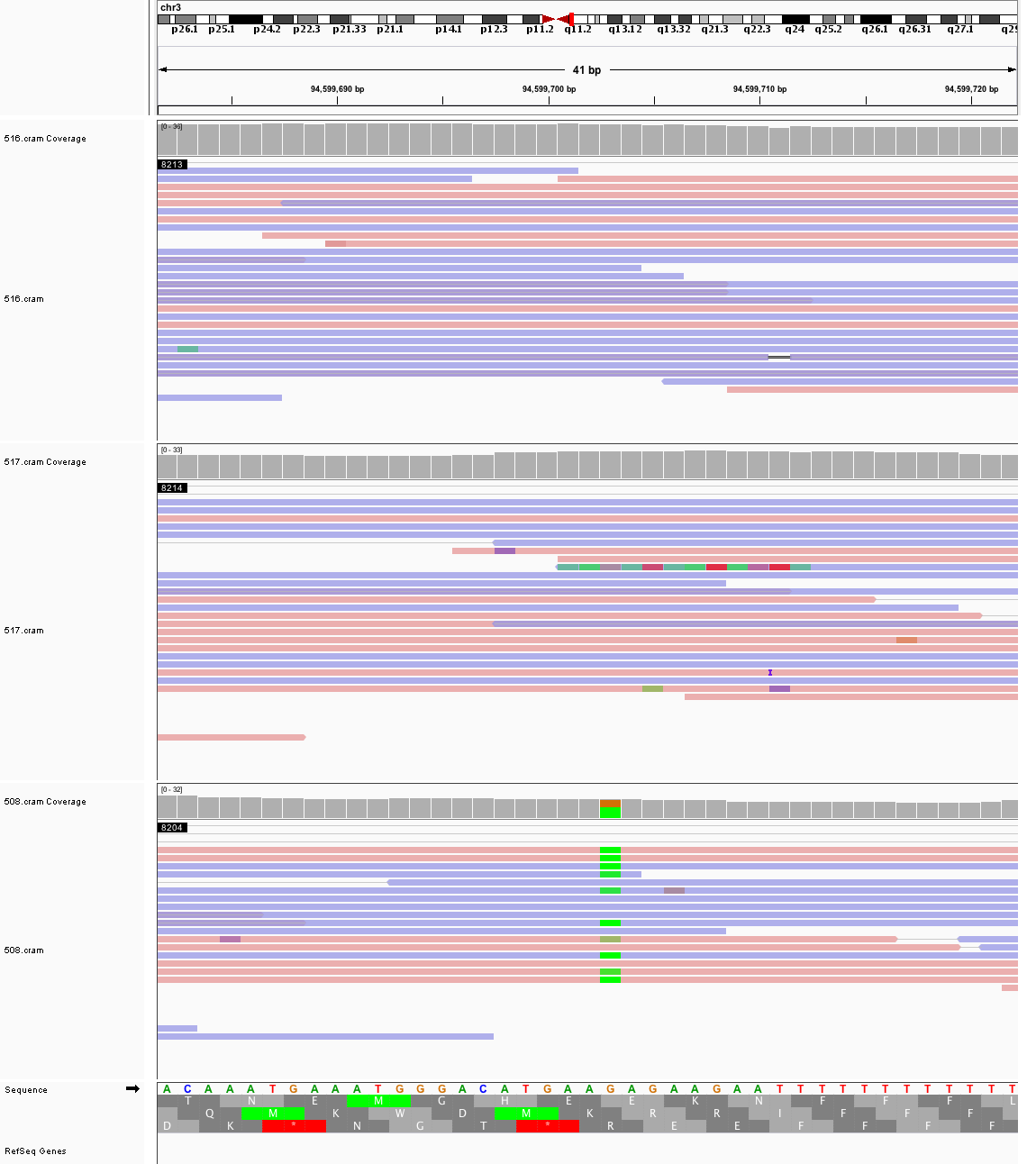

### chr3_97,165,411_97,165,451.png

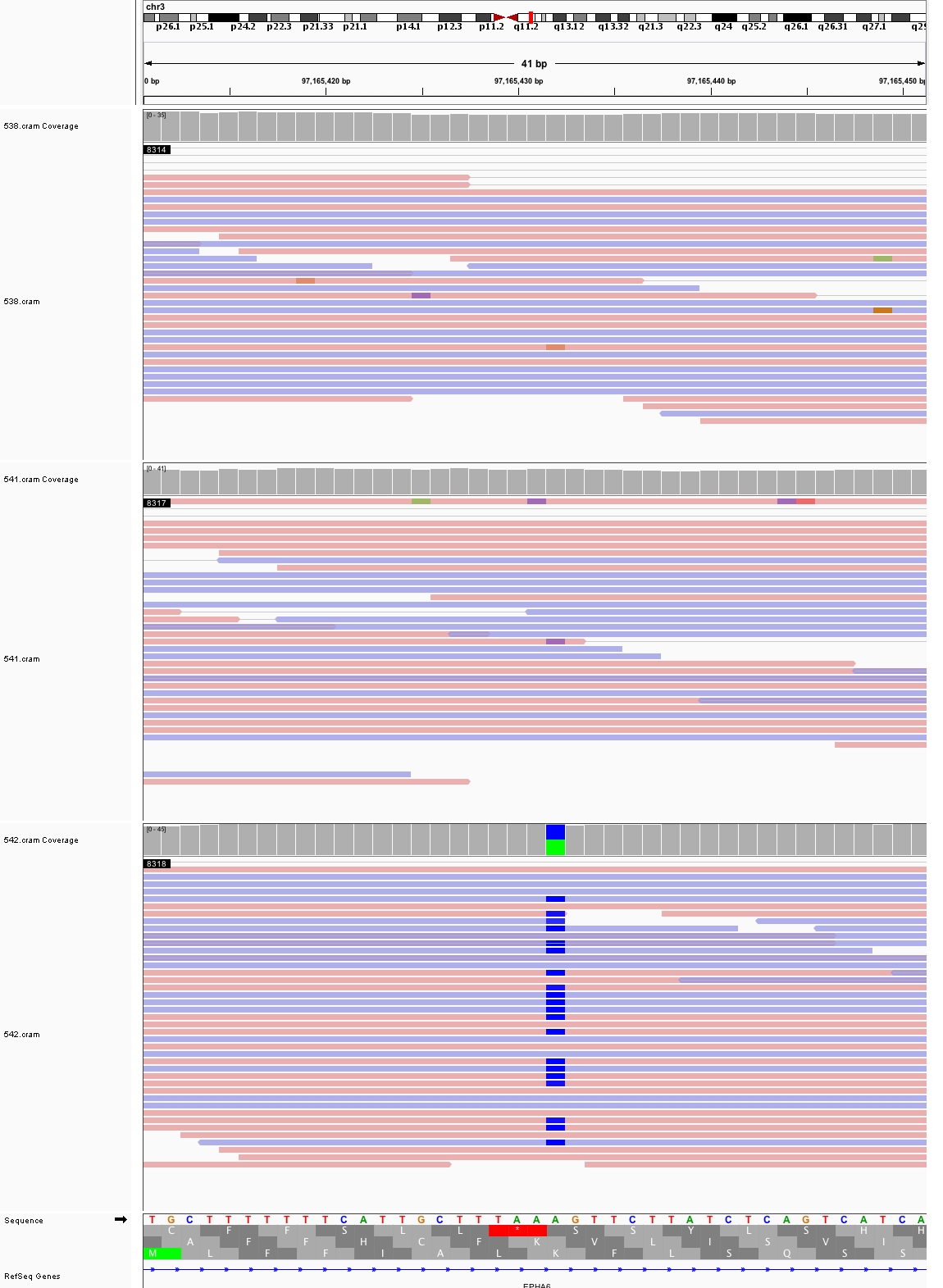

### chr3_104,980,146_104,980,186.png

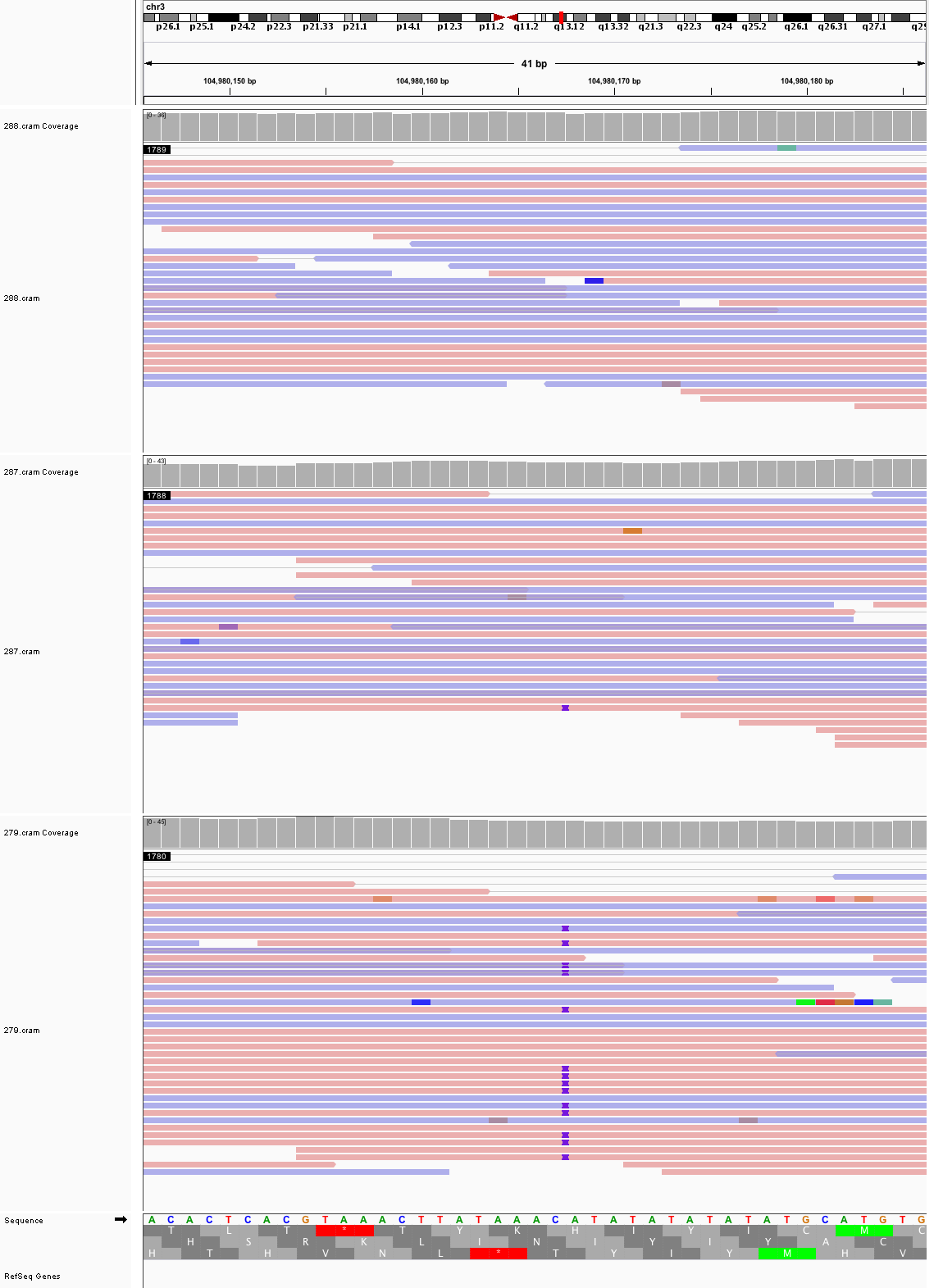

### chr3_113,327,219_113,327,259.png

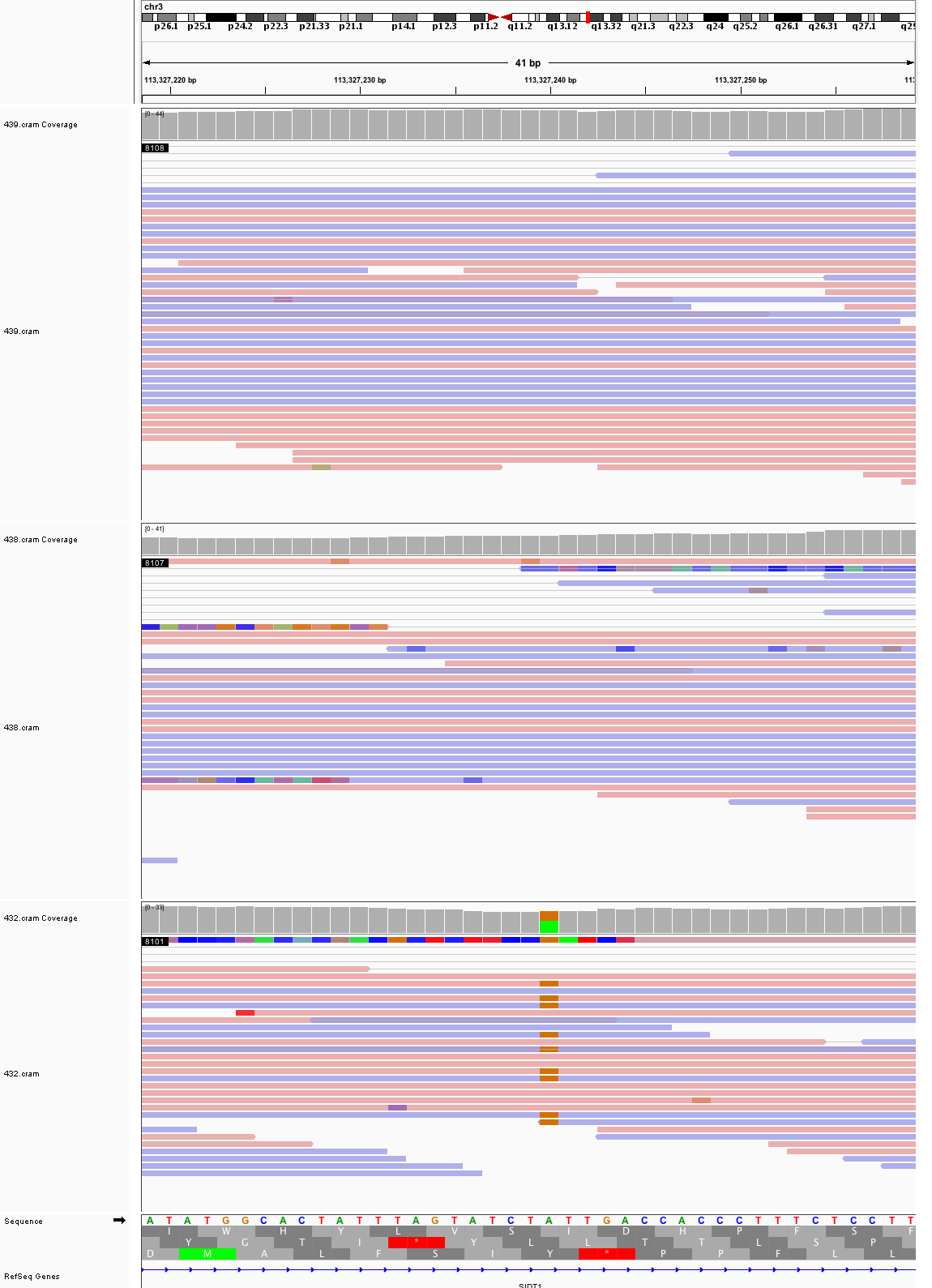

### chr3_129,662,703_129,662,743.png

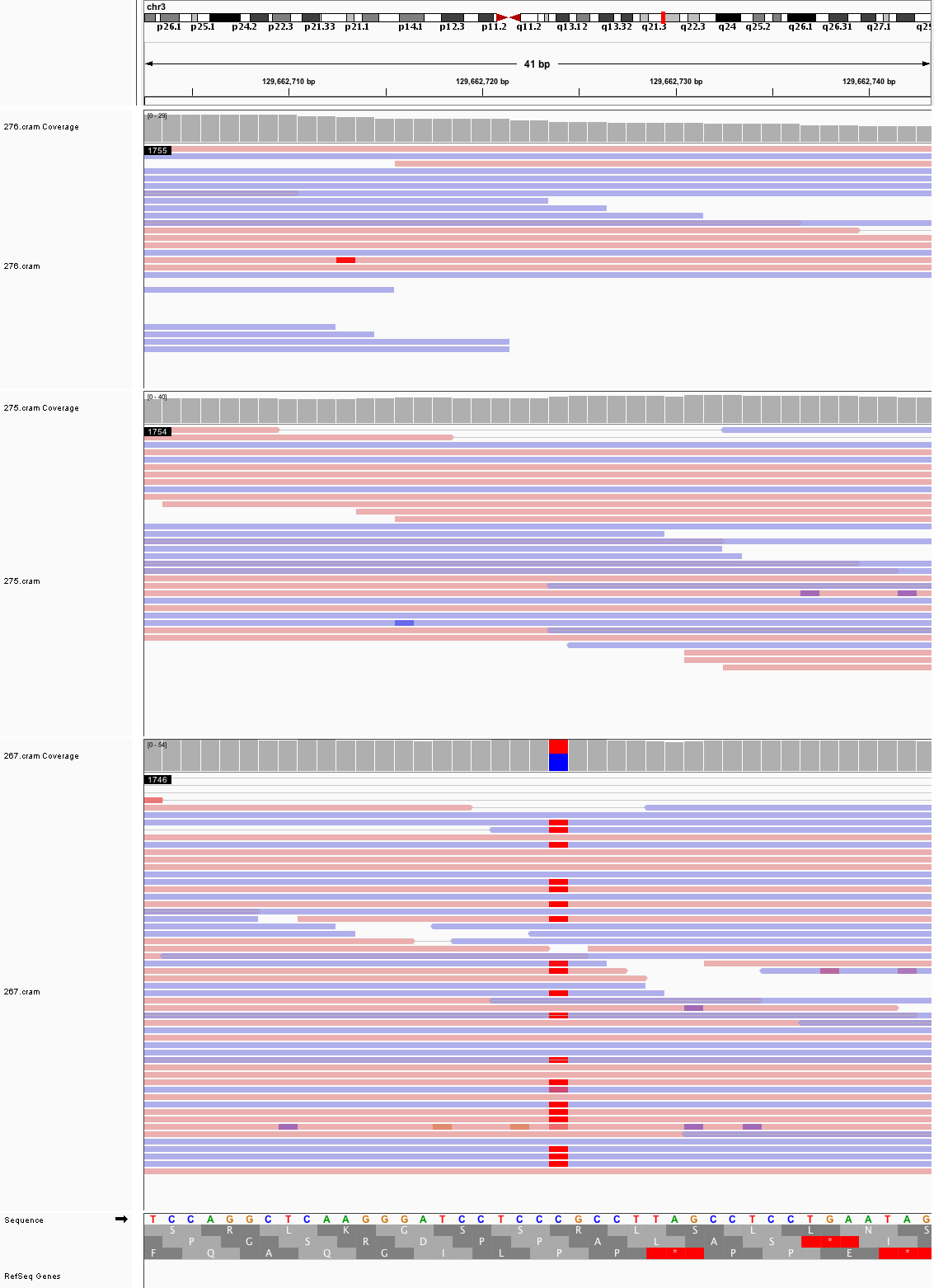

### chr3_131,024,756_131,024,796.png

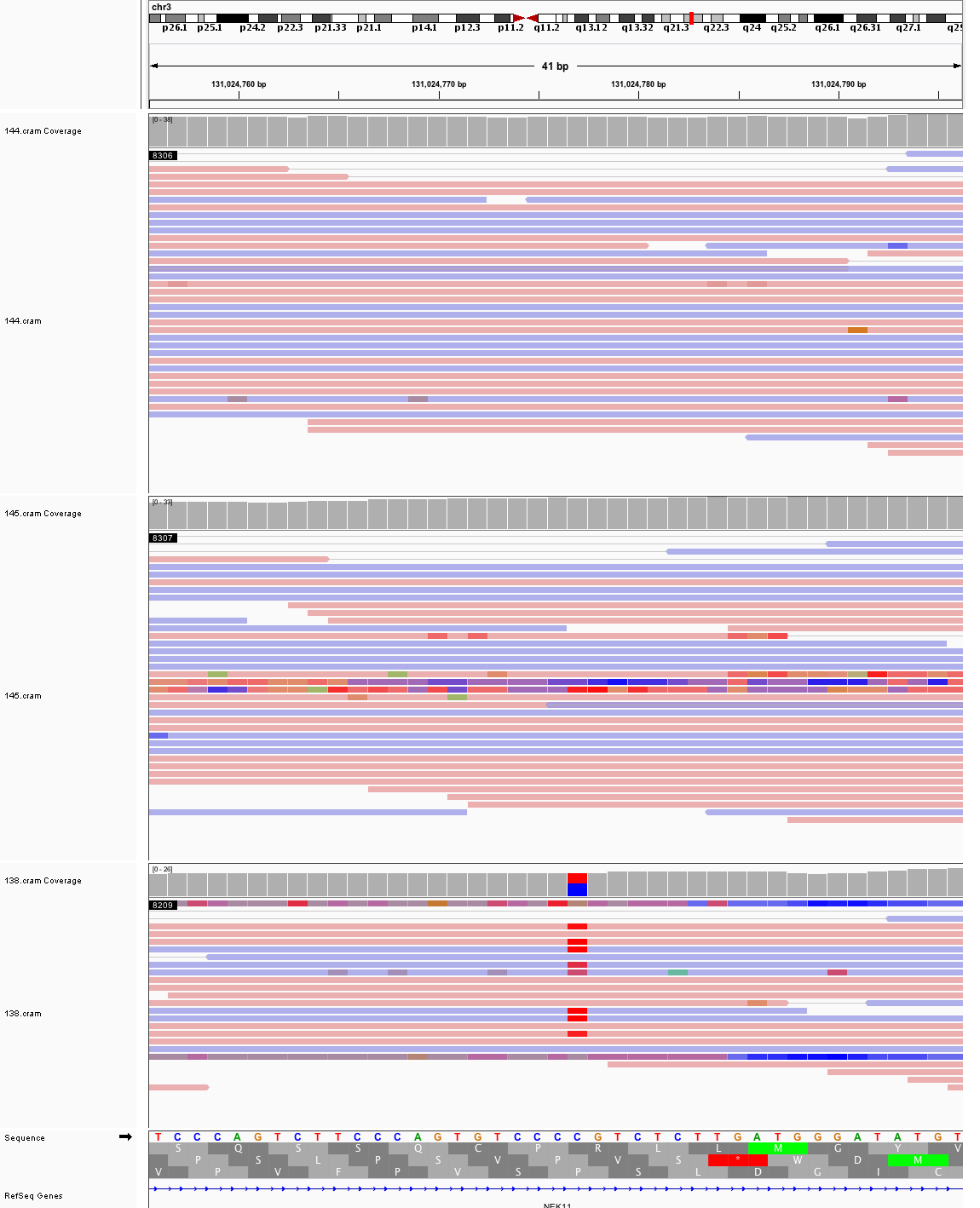

### chr3_188,794,337_188,794,377.png

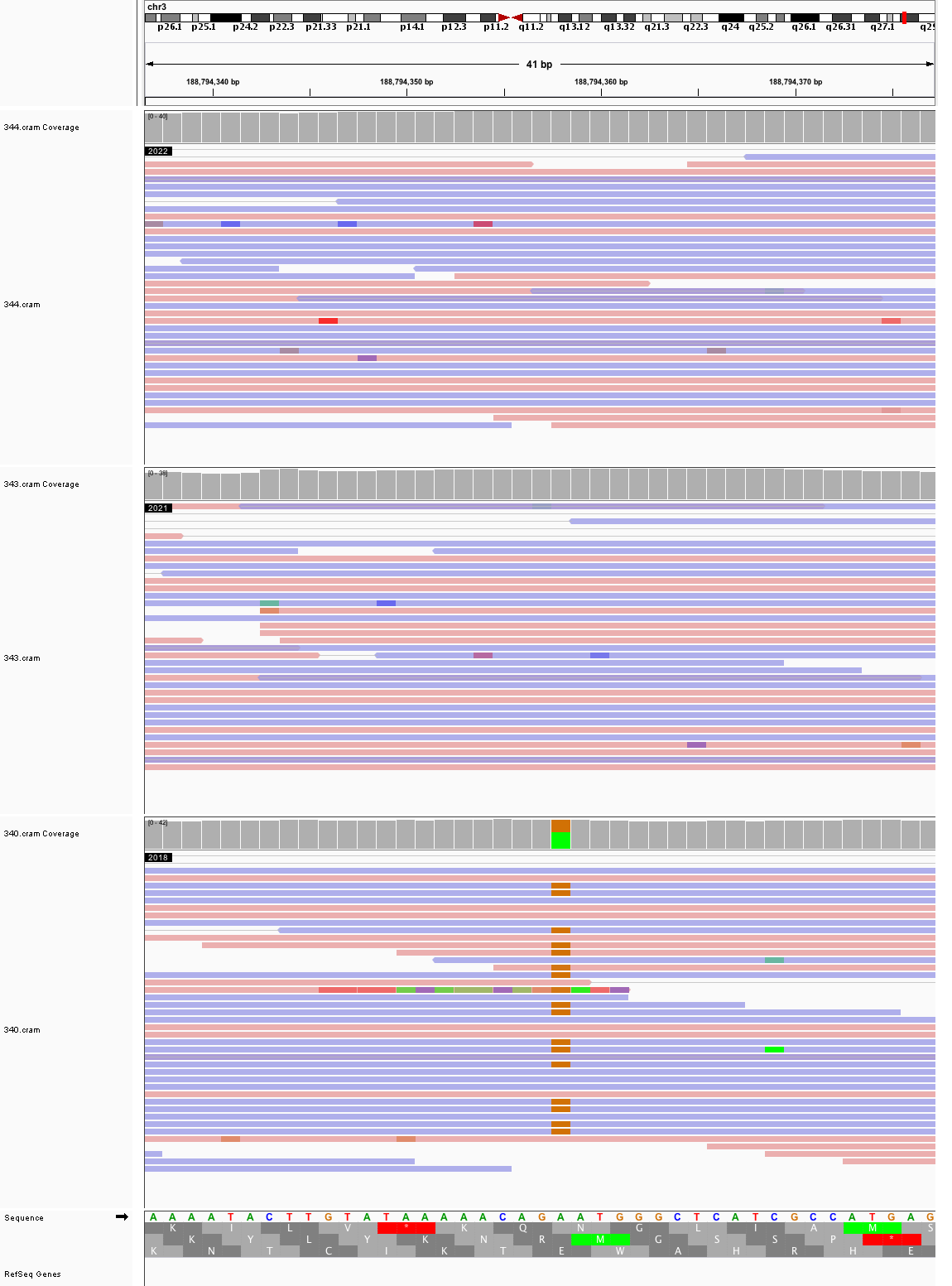

### chr4_37,886,881_37,886,921.png

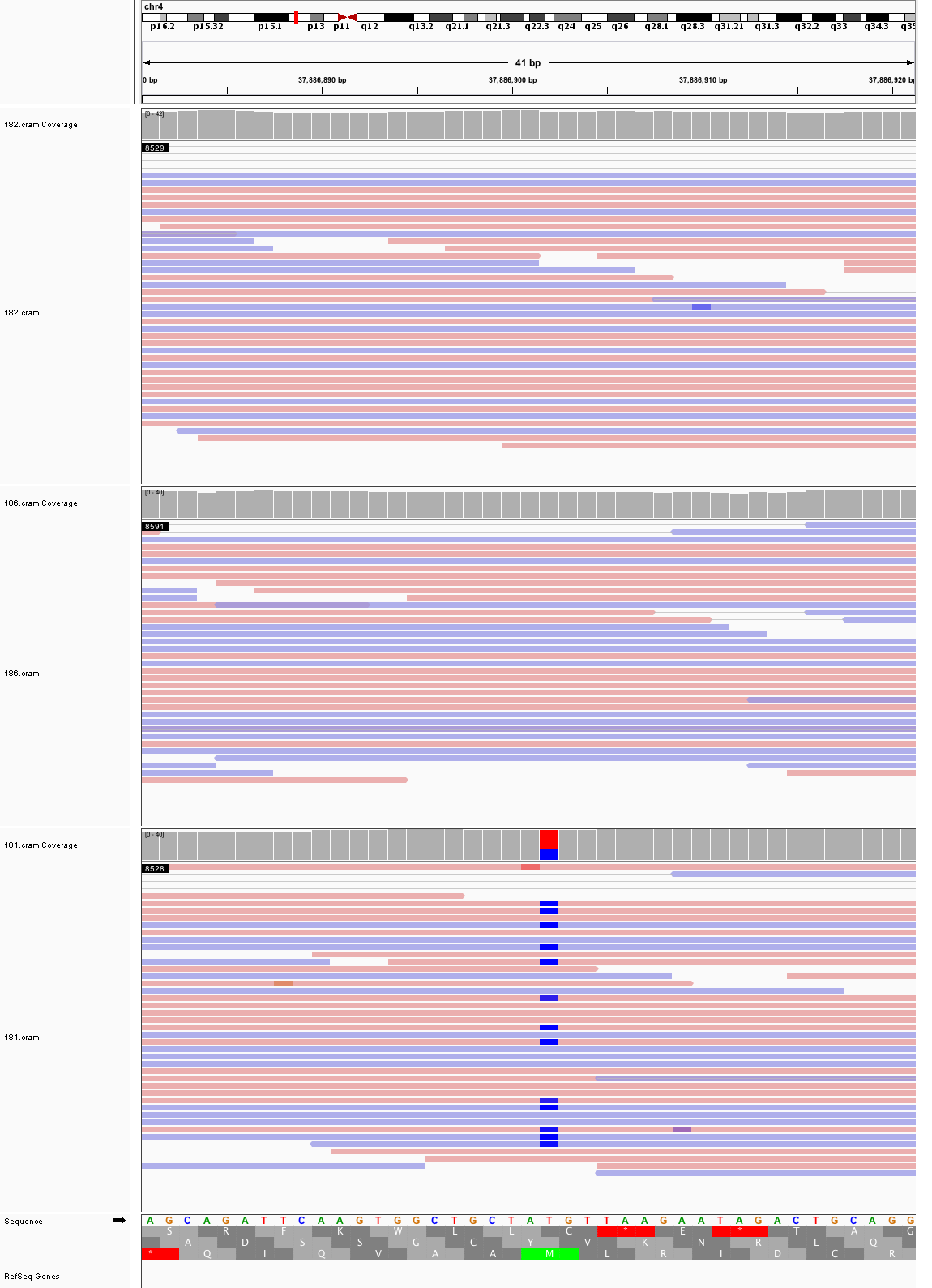

### chr4_55,150,604_55,150,644.png

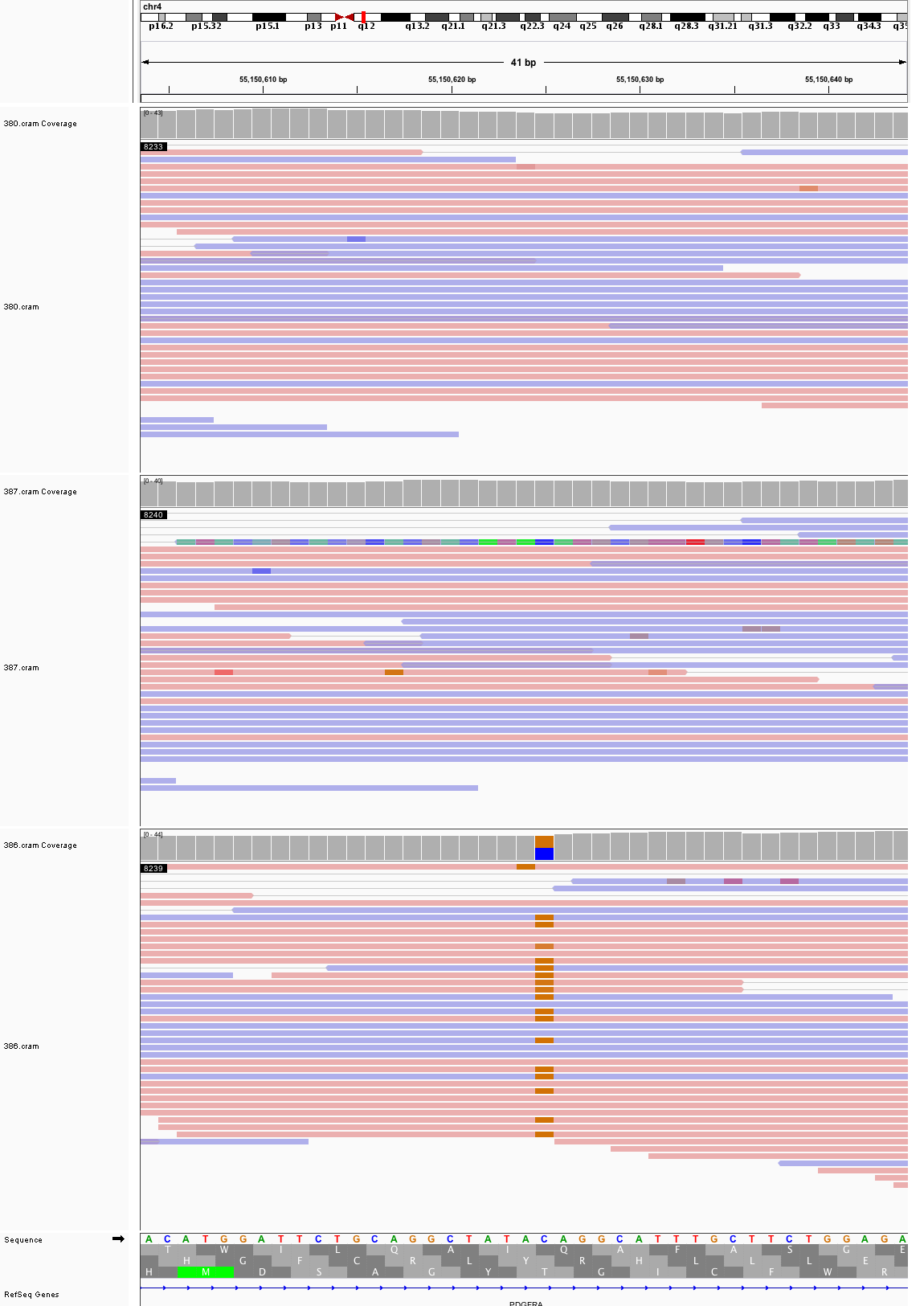

### chr4_80,848,289_80,848,329.png

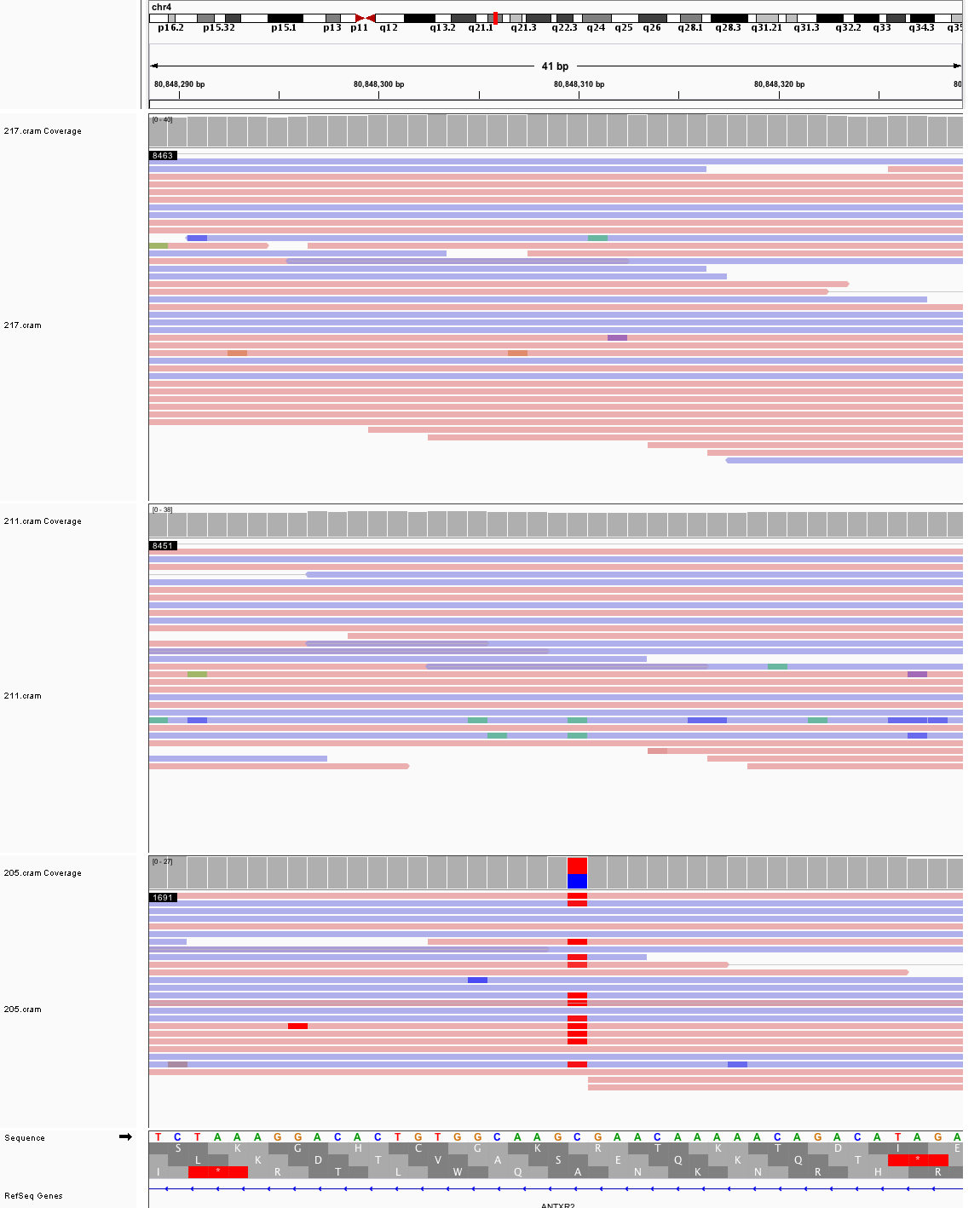

### chr4_91,648,322_91,648,362.png

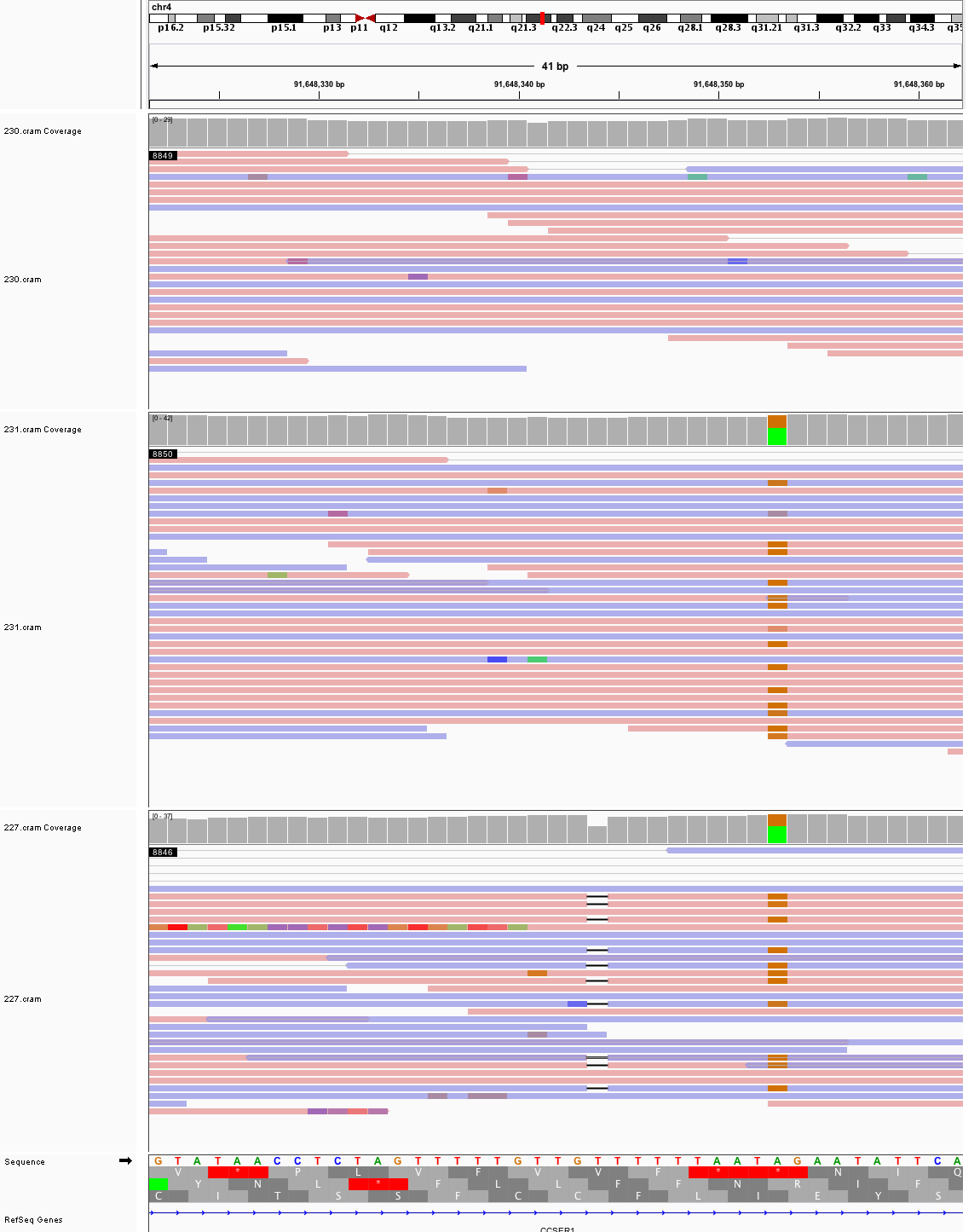

### chr4_92,300,576_92,300,616.png

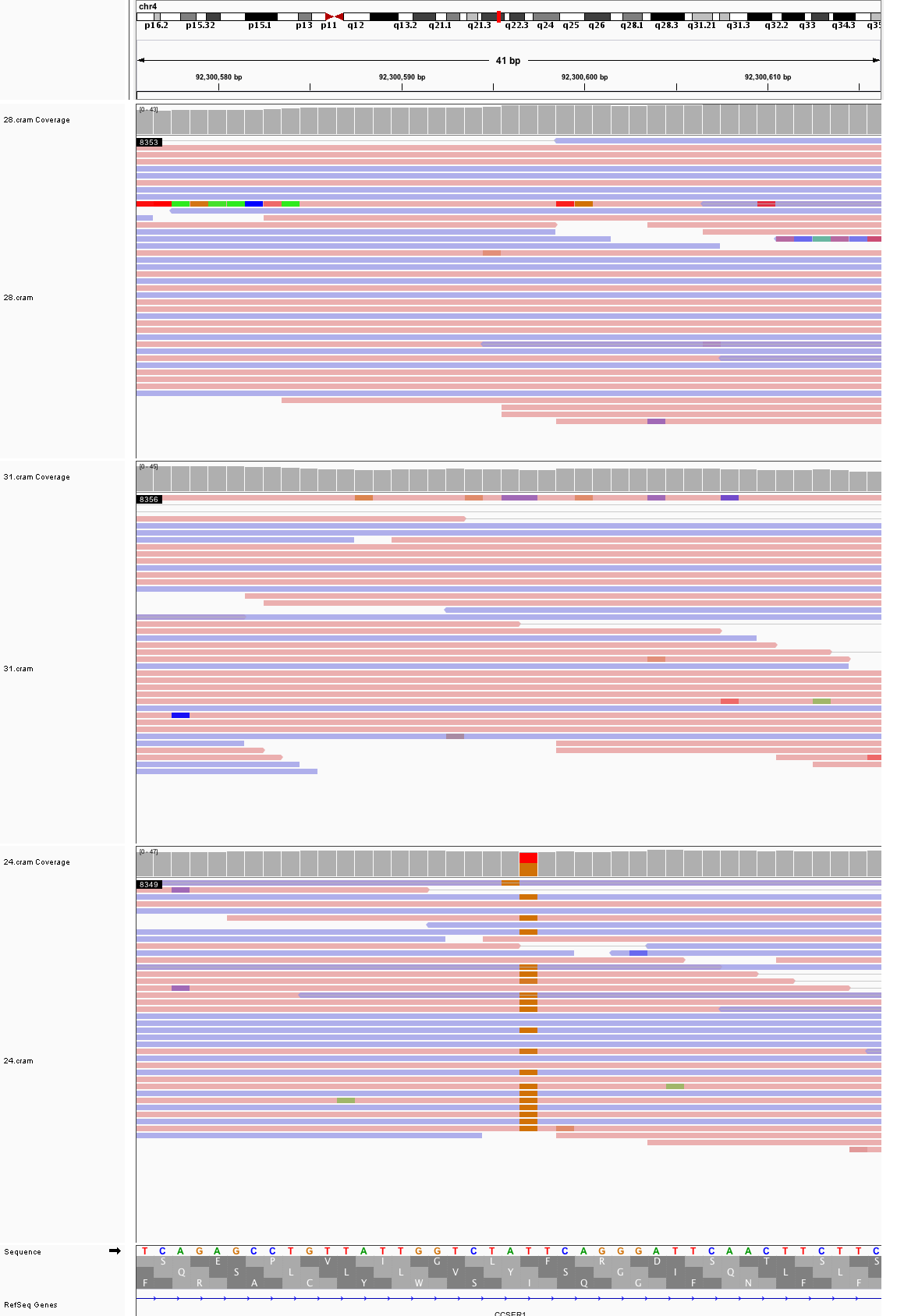

### chr4_134,944,407_134,944,447.png

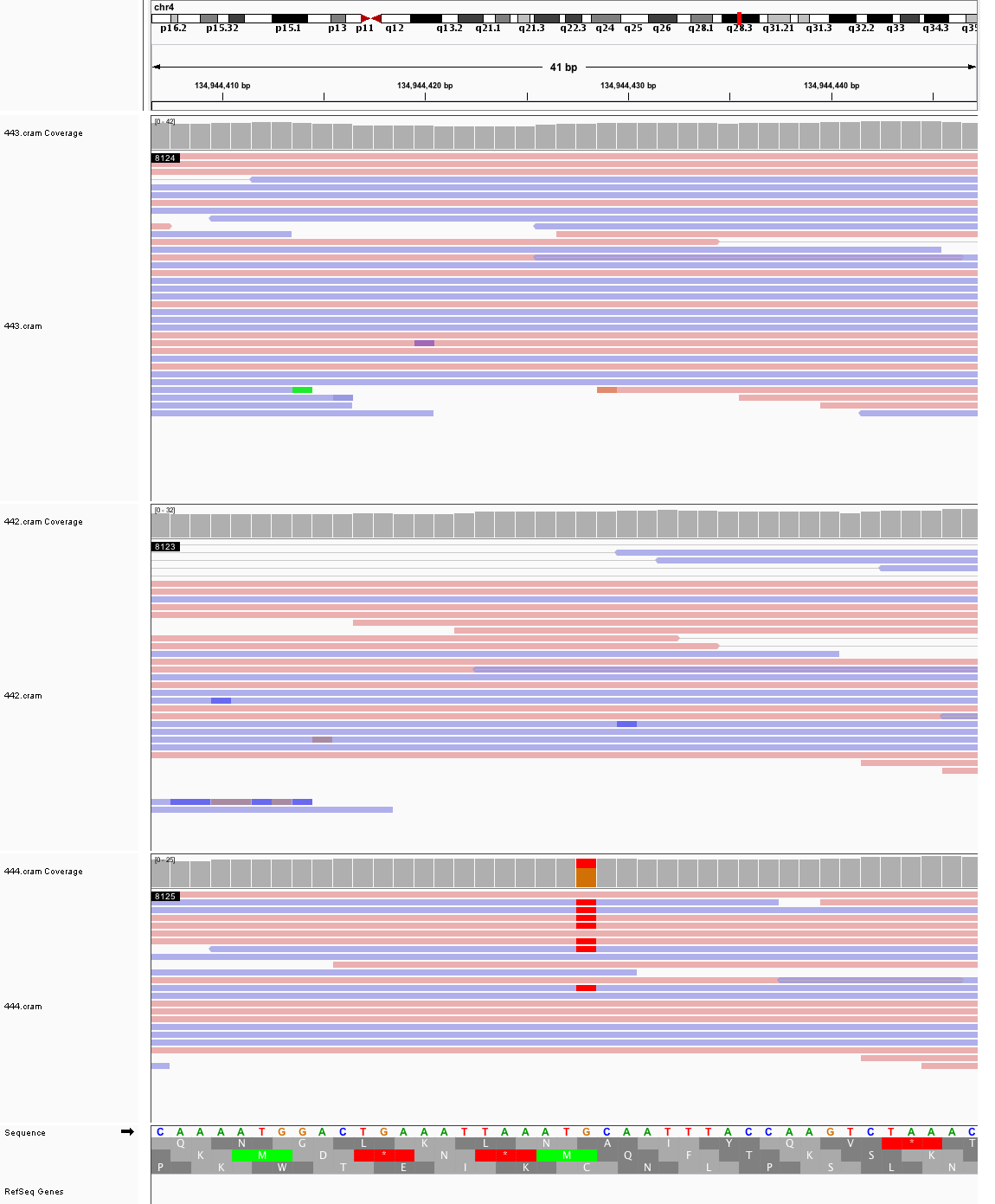

### chr5_7,919,866_7,919,906.png

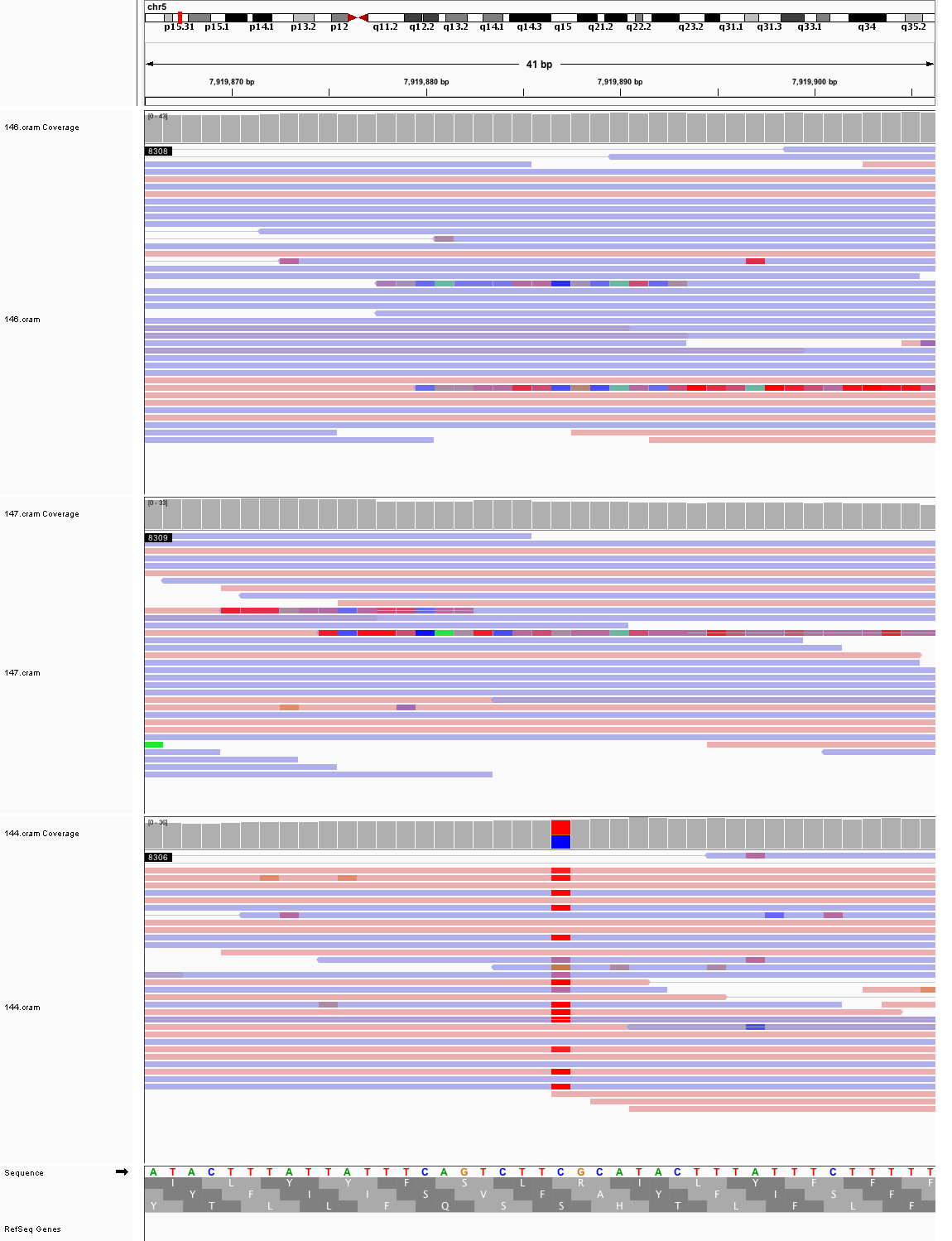

### chr5_110,524,004_110,524,044.png

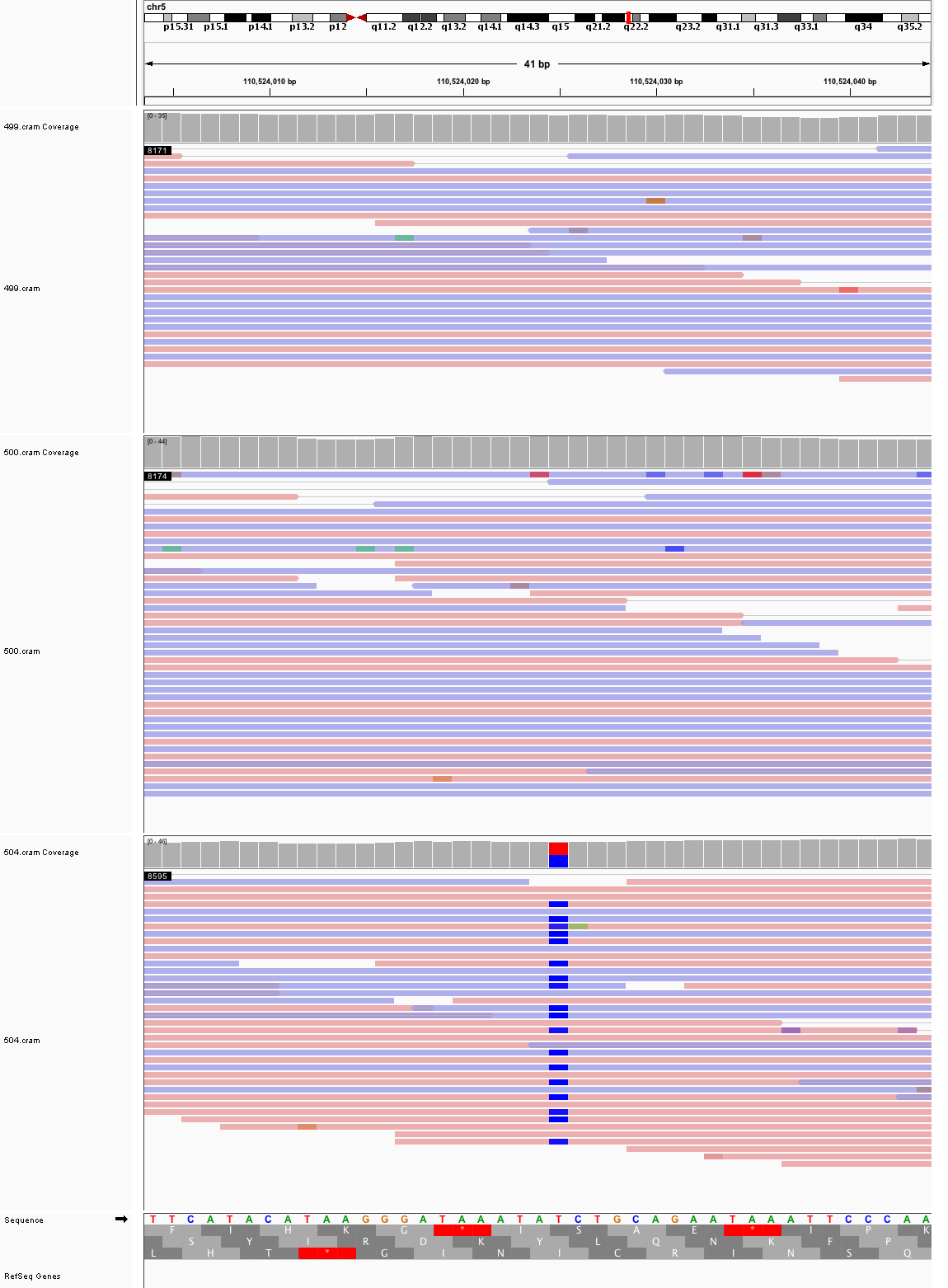

### chr5_117,856,699_117,856,739.png

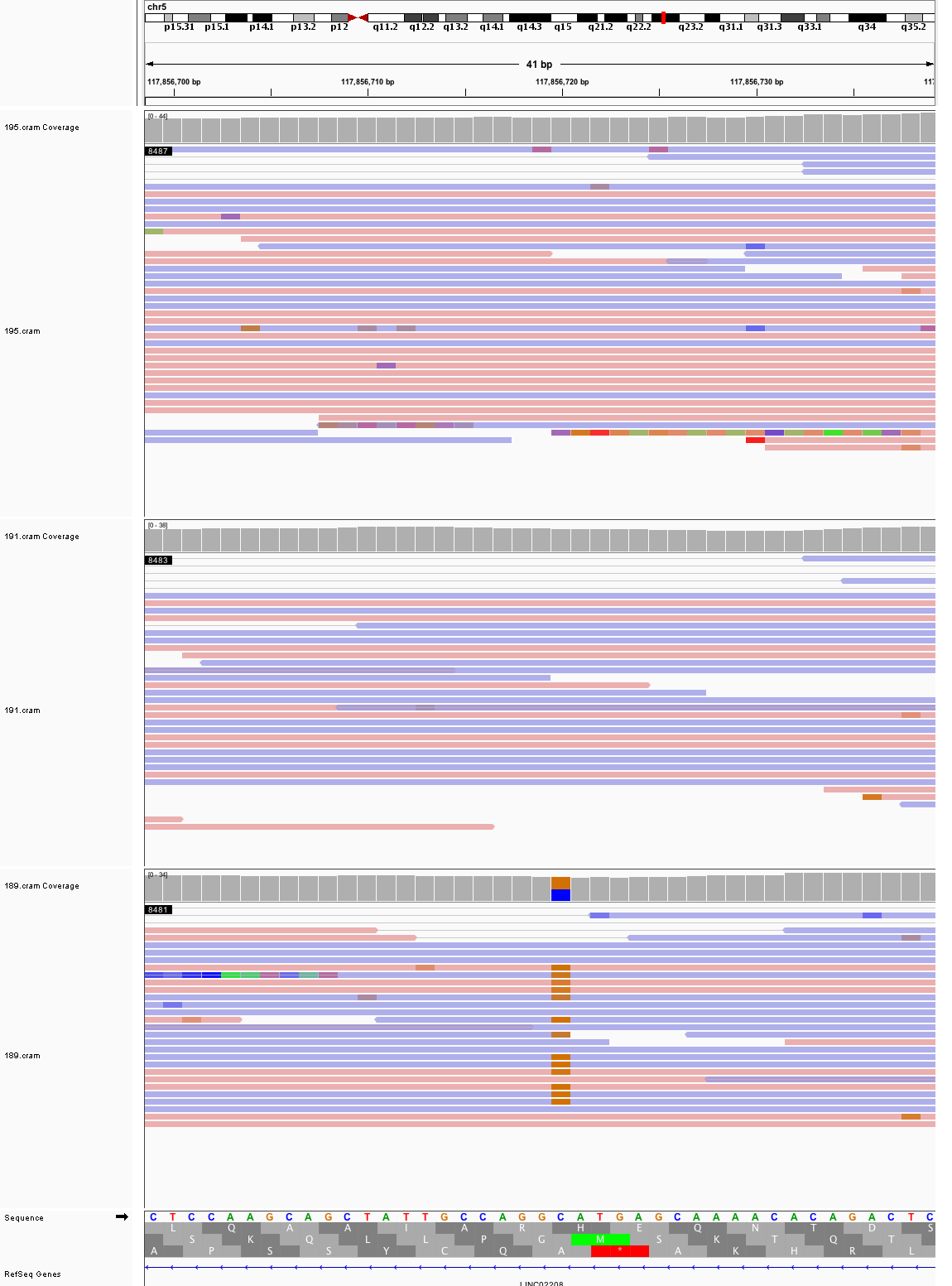

### chr5_123,158,305_123,158,345.png

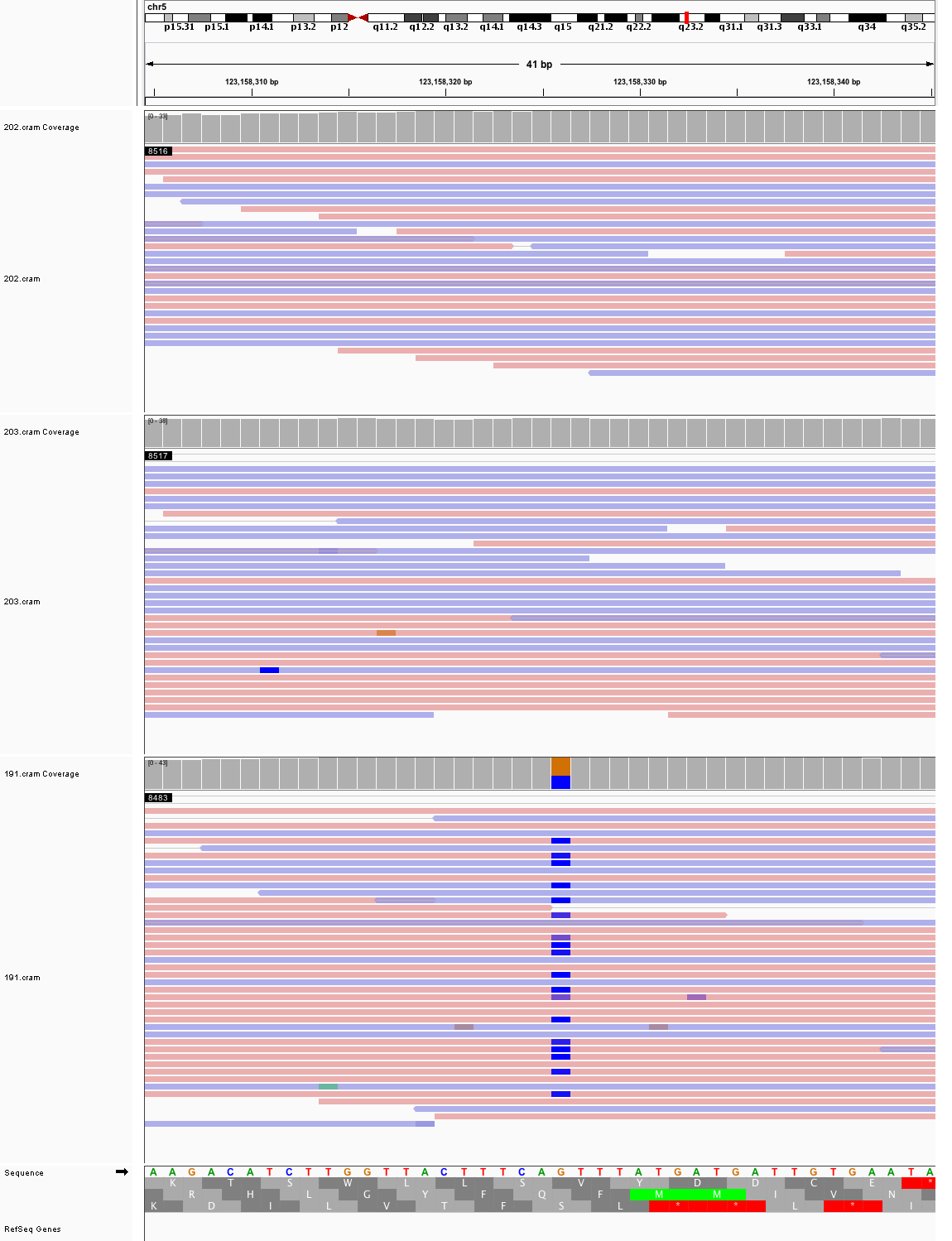

### chr6_1,621,142_1,621,182.png

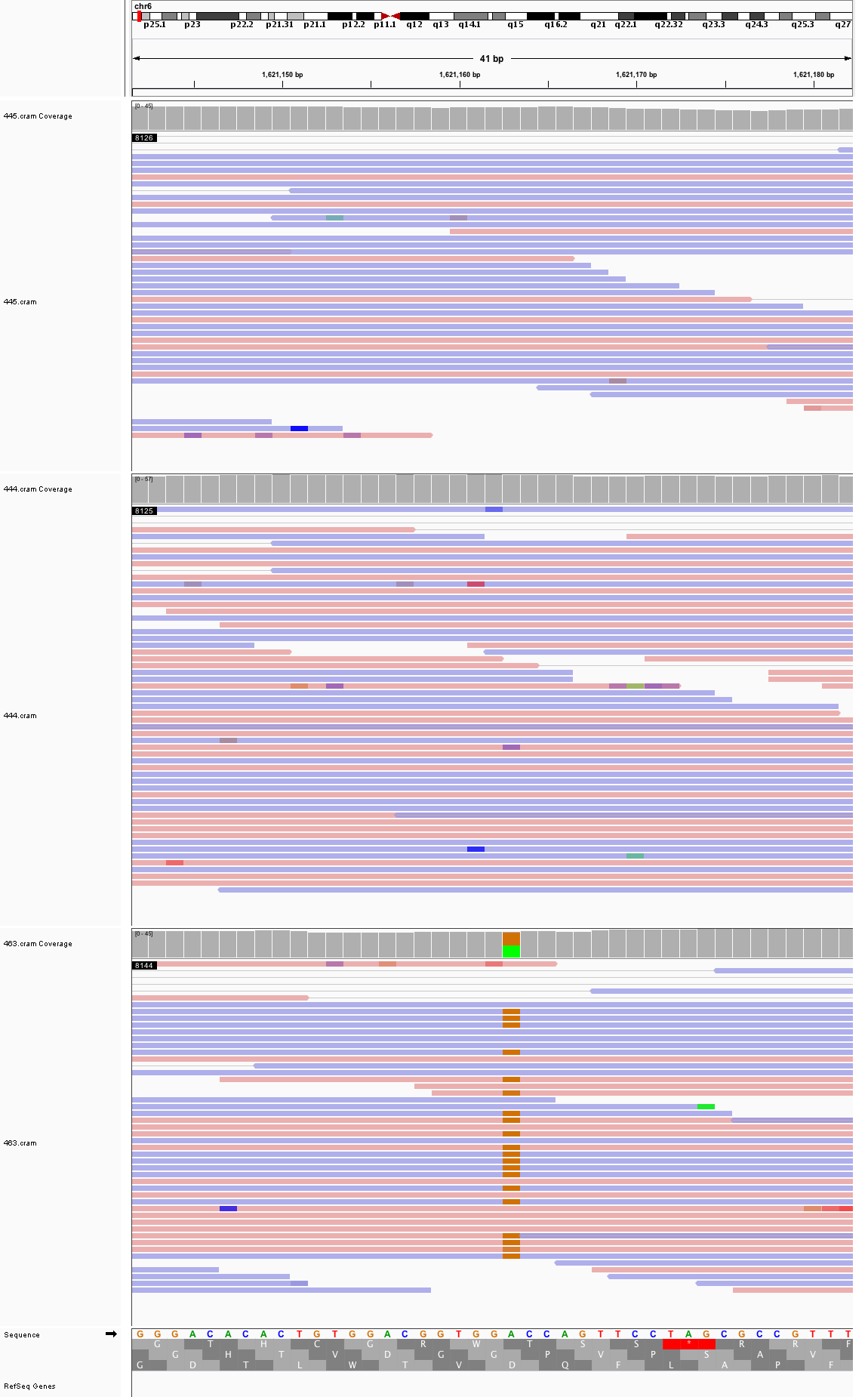

### chr6_2,236,059_2,236,099.png

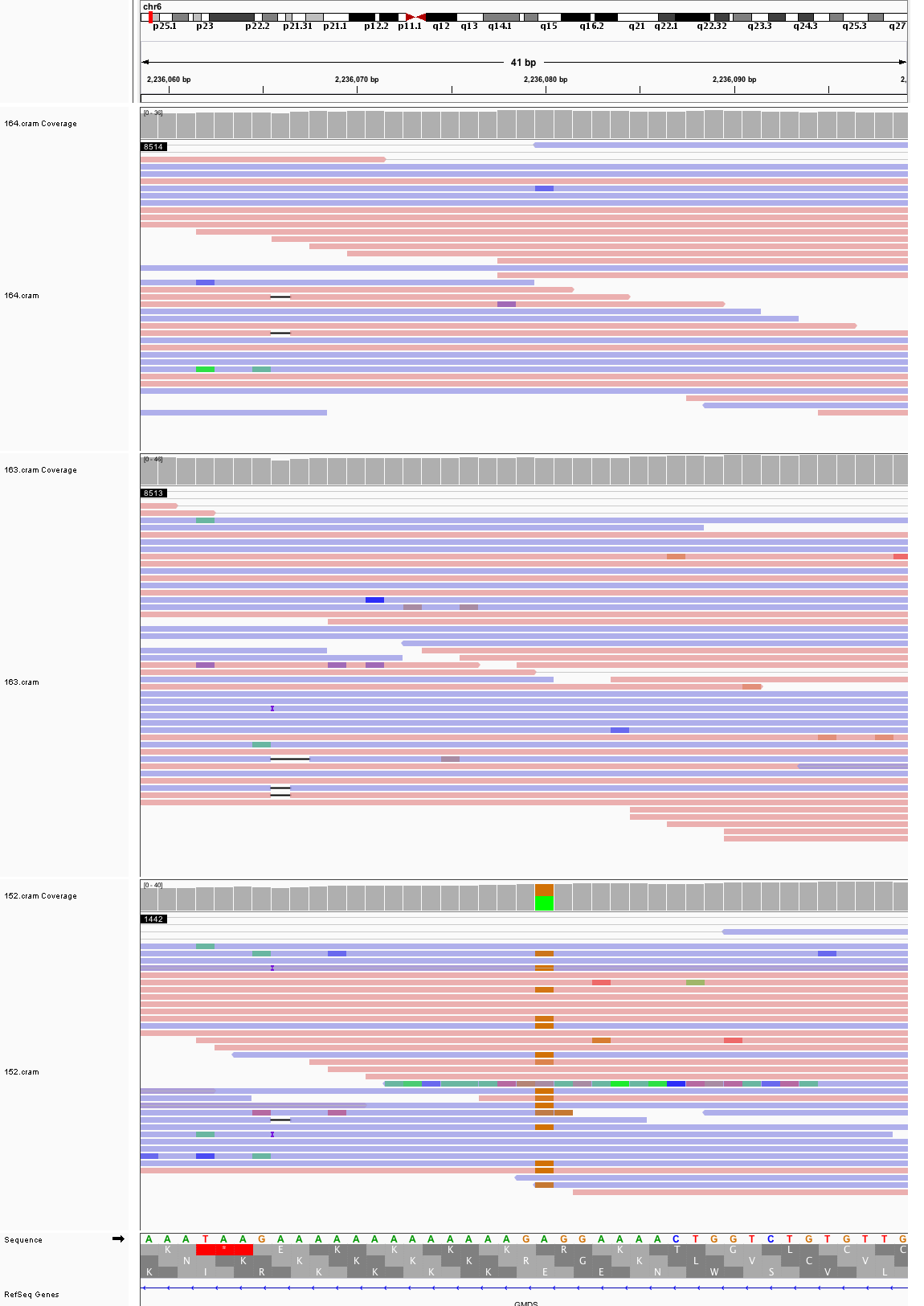

### chr6_66,092,913_66,092,953.png

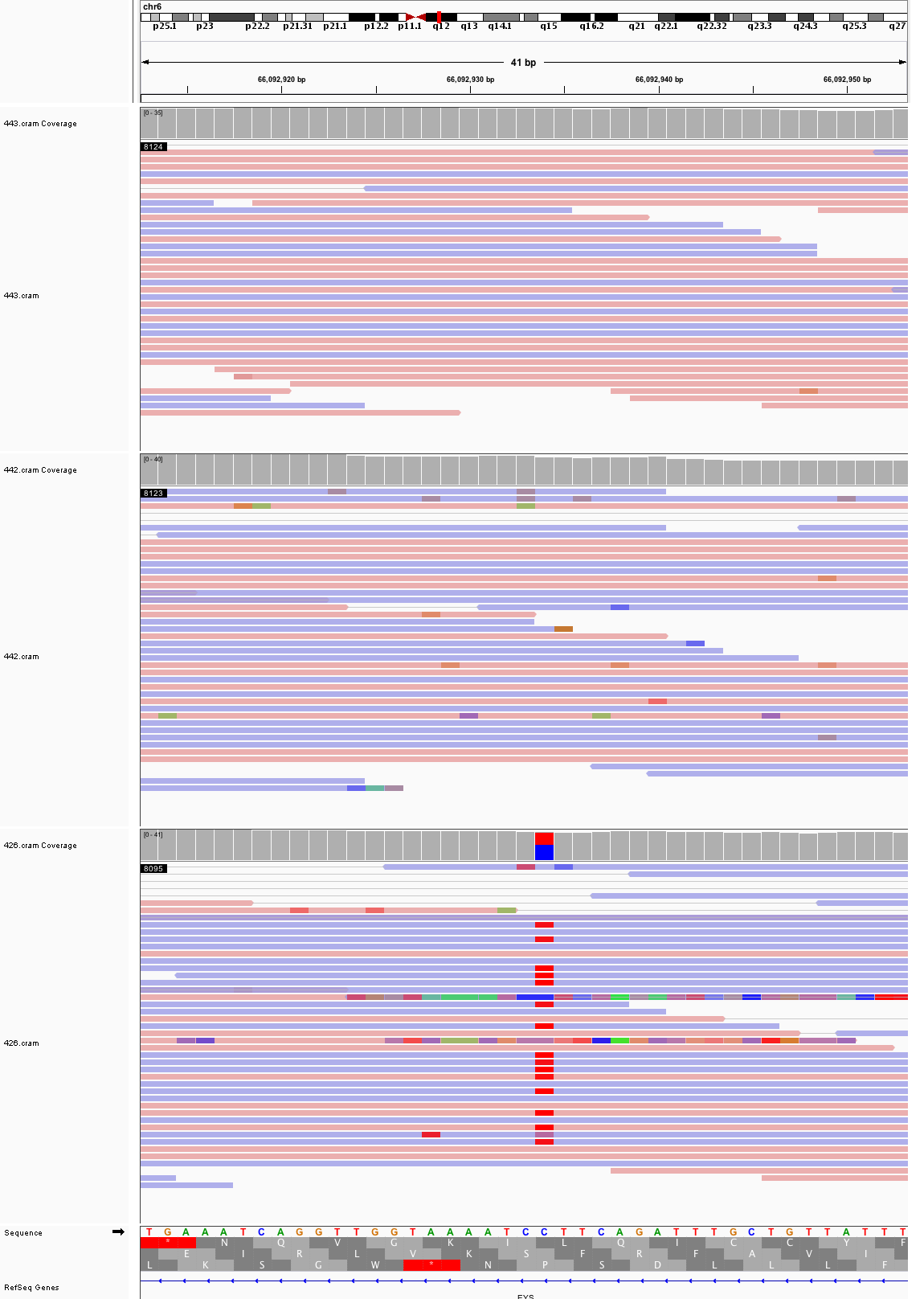

### chr6_102,772,745_102,772,785.png

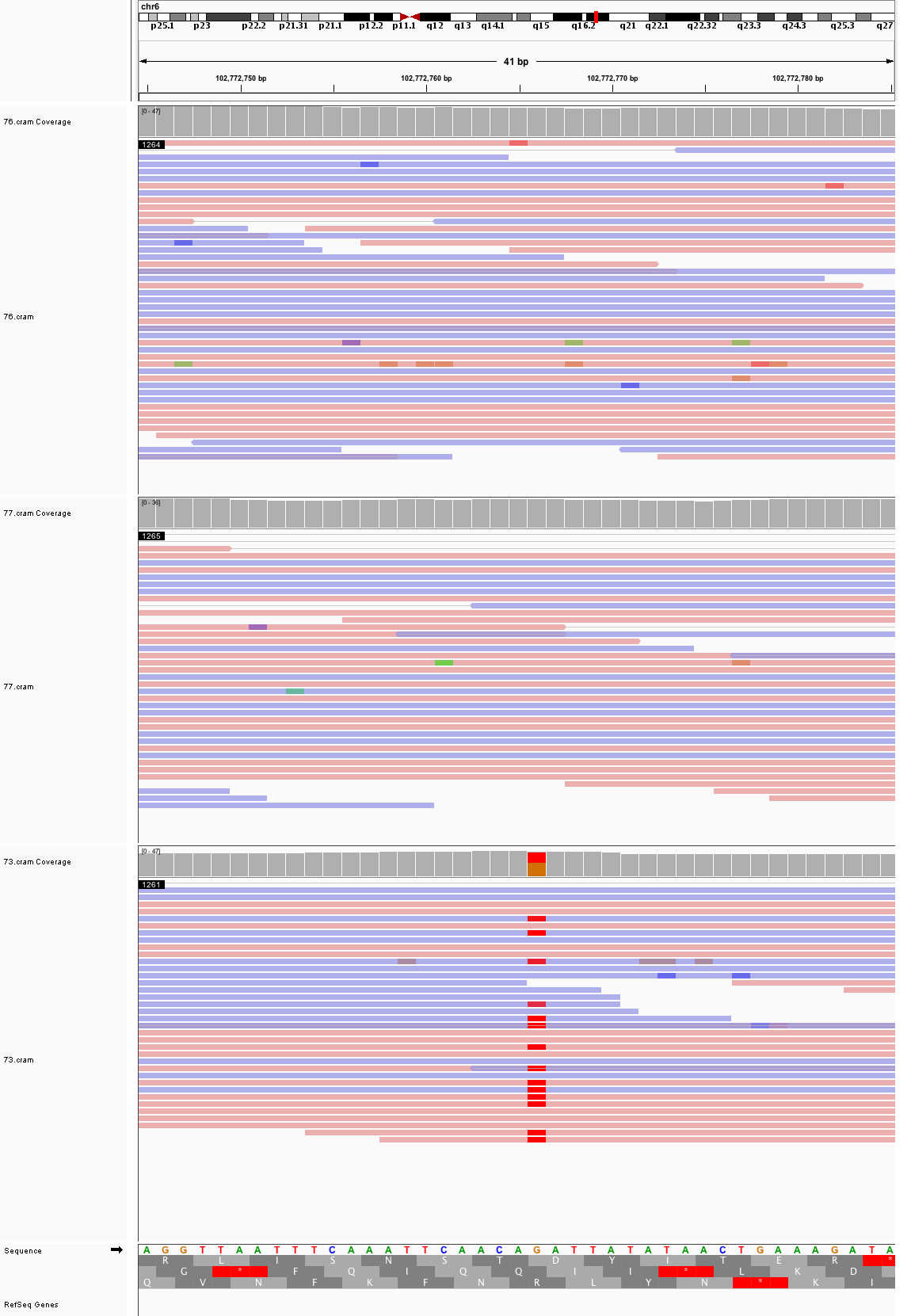

### chr6_105,275,899_105,275,939.png

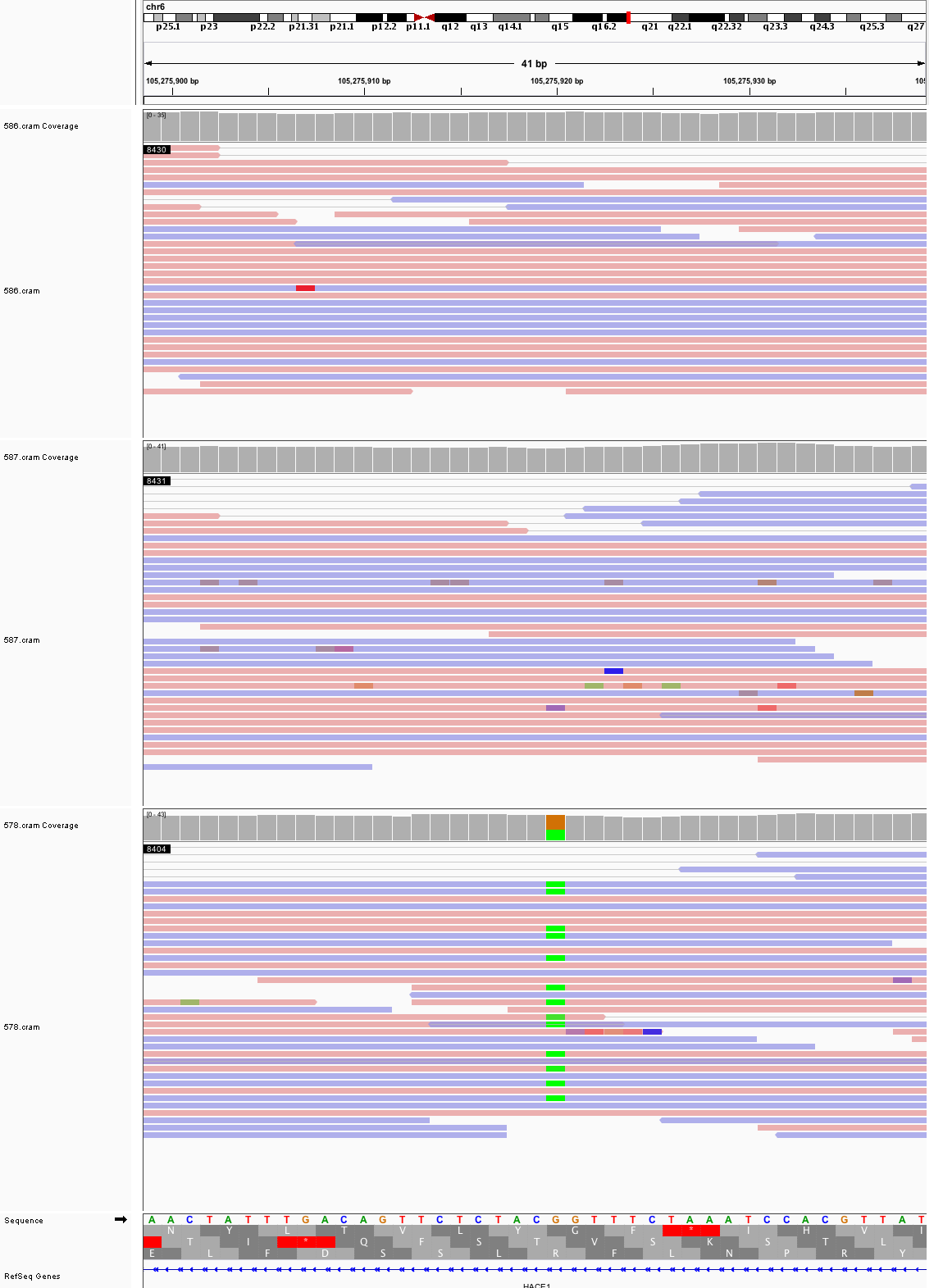

### chr6_143,929,881_143,929,921.png

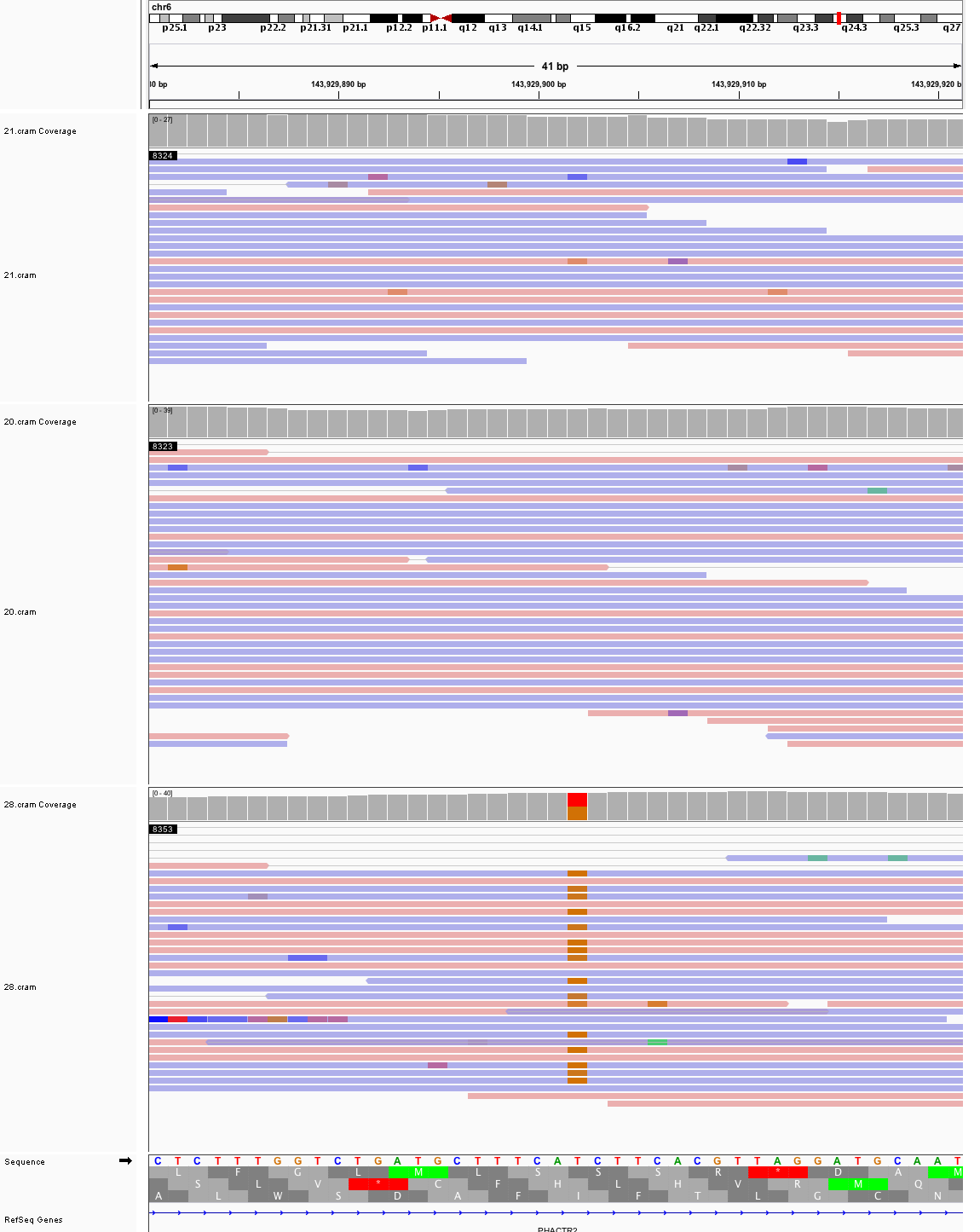

### chr6_146,286,194_146,286,234.png

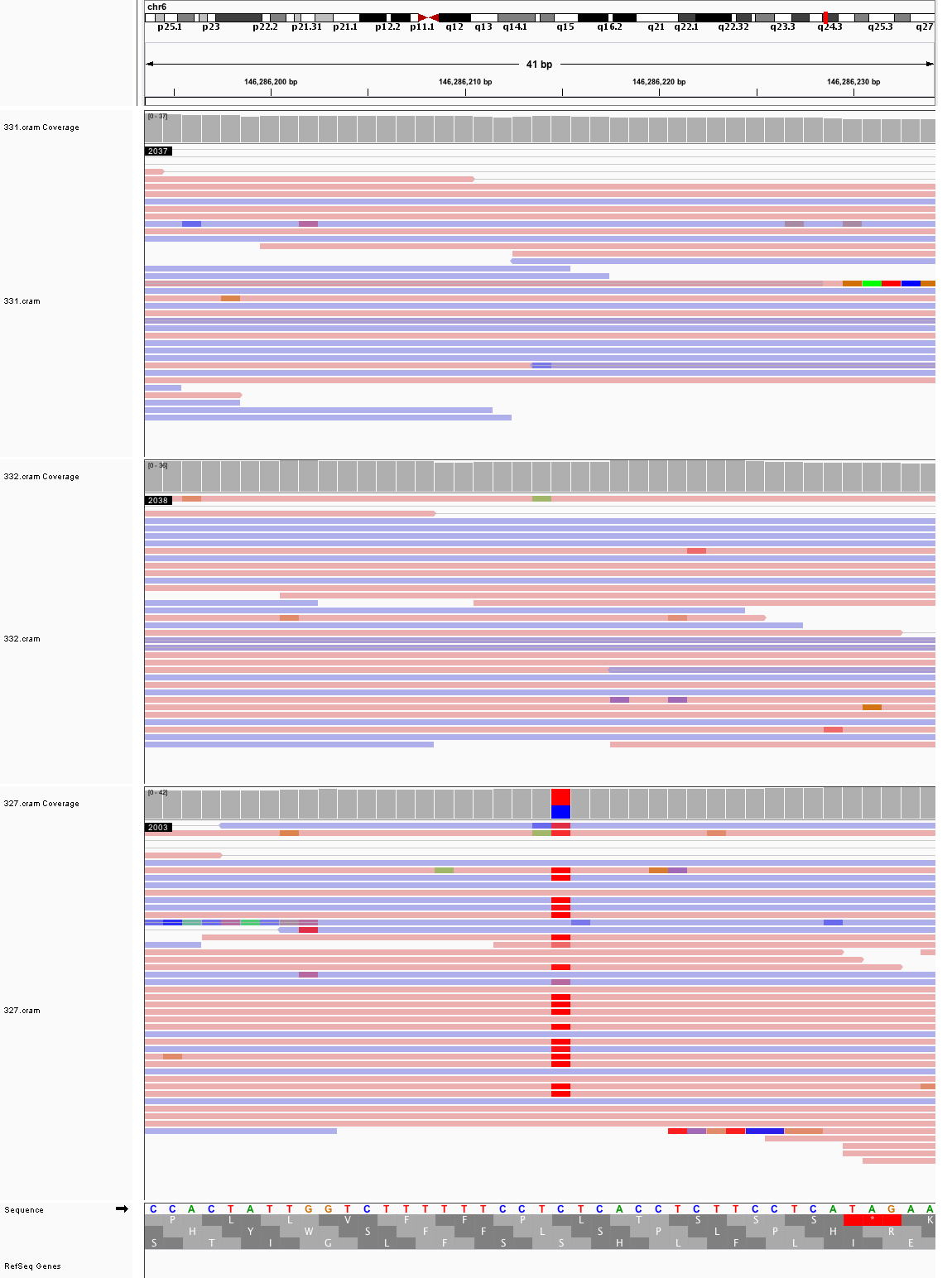

### chr6_148,594,824_148,594,864.png

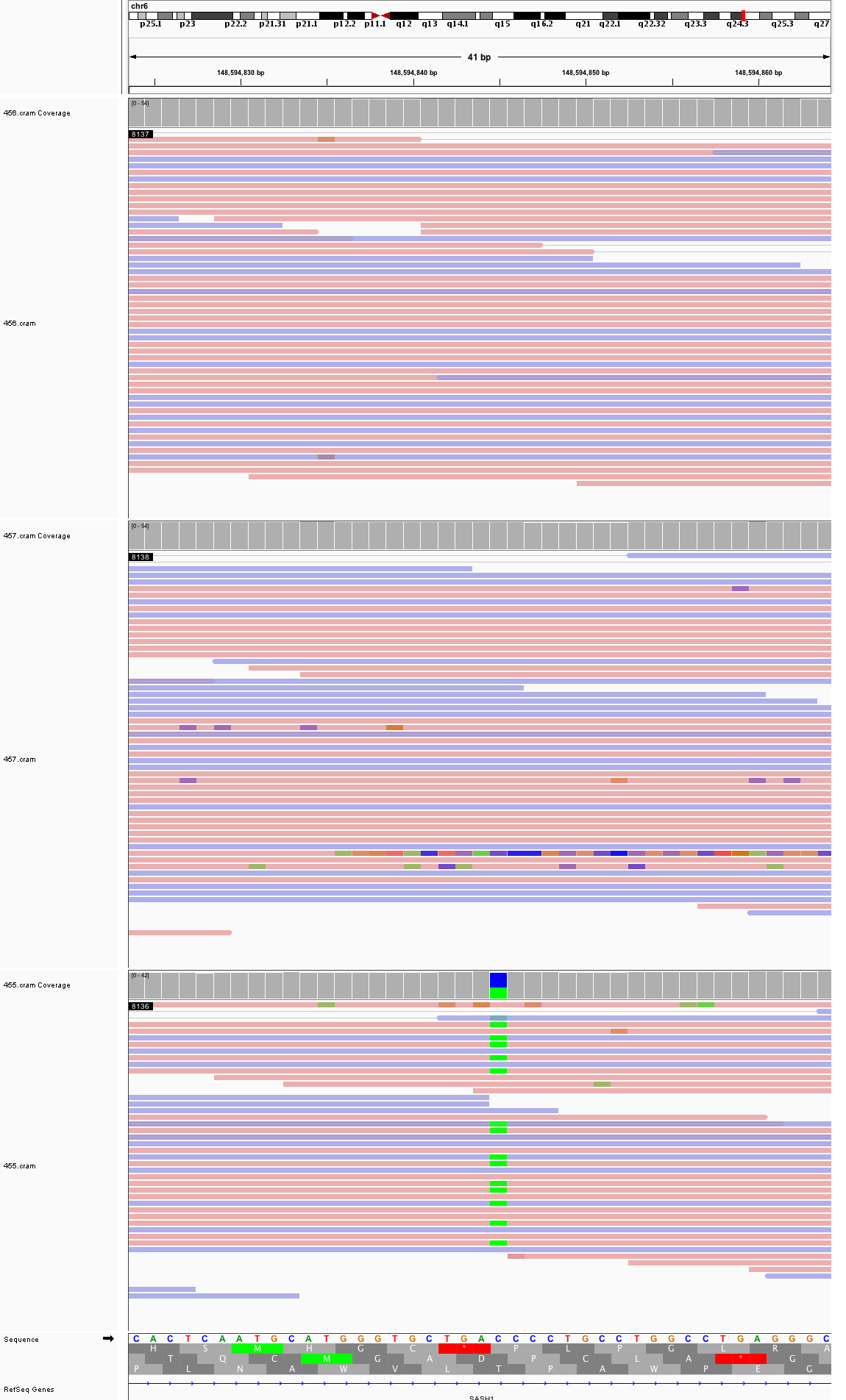

### chr6_167,115,463_167,115,503.png

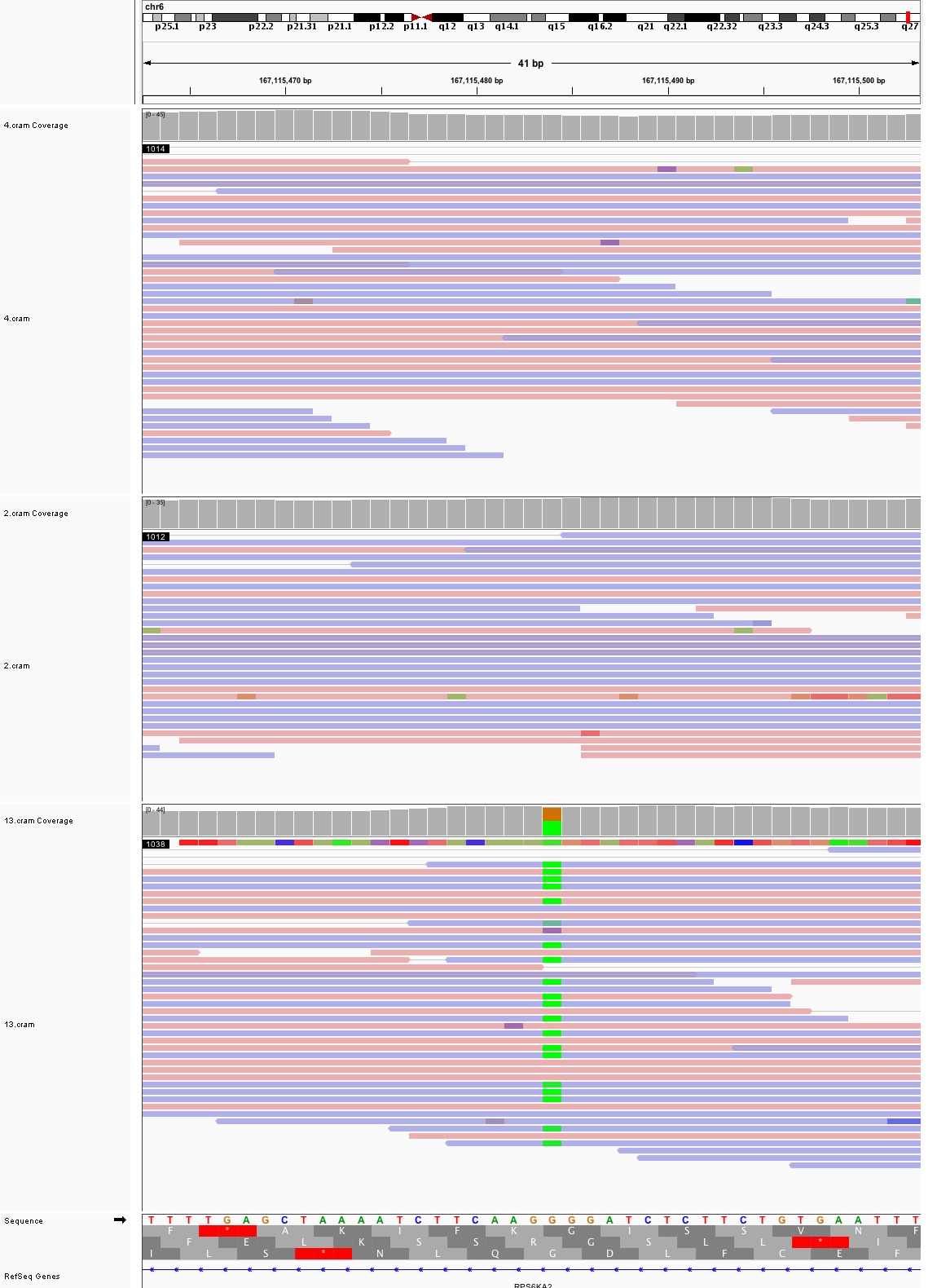

### chr7_65,944,023_65,944,063.png

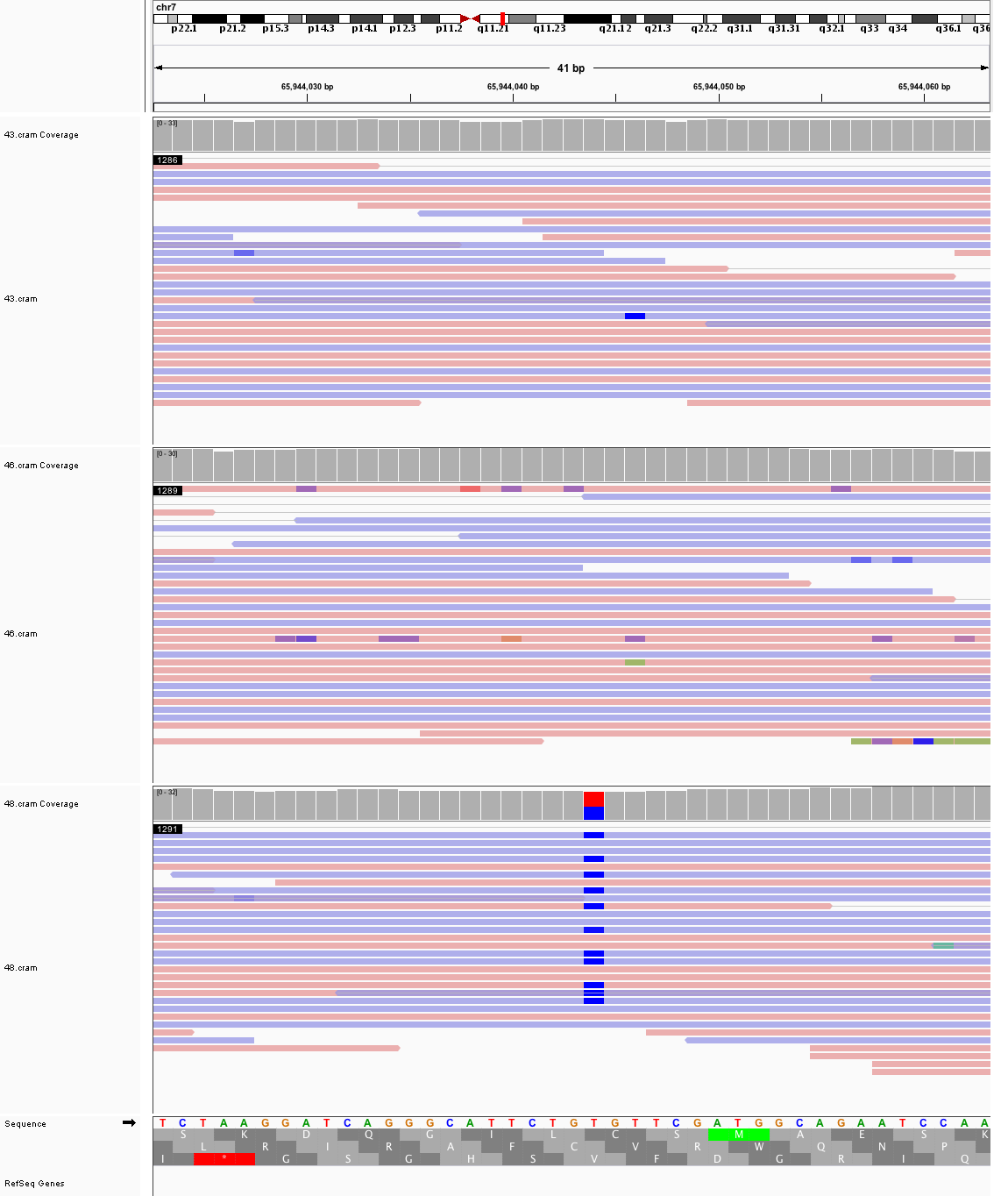

### chr7_123,290,740_123,290,780.png

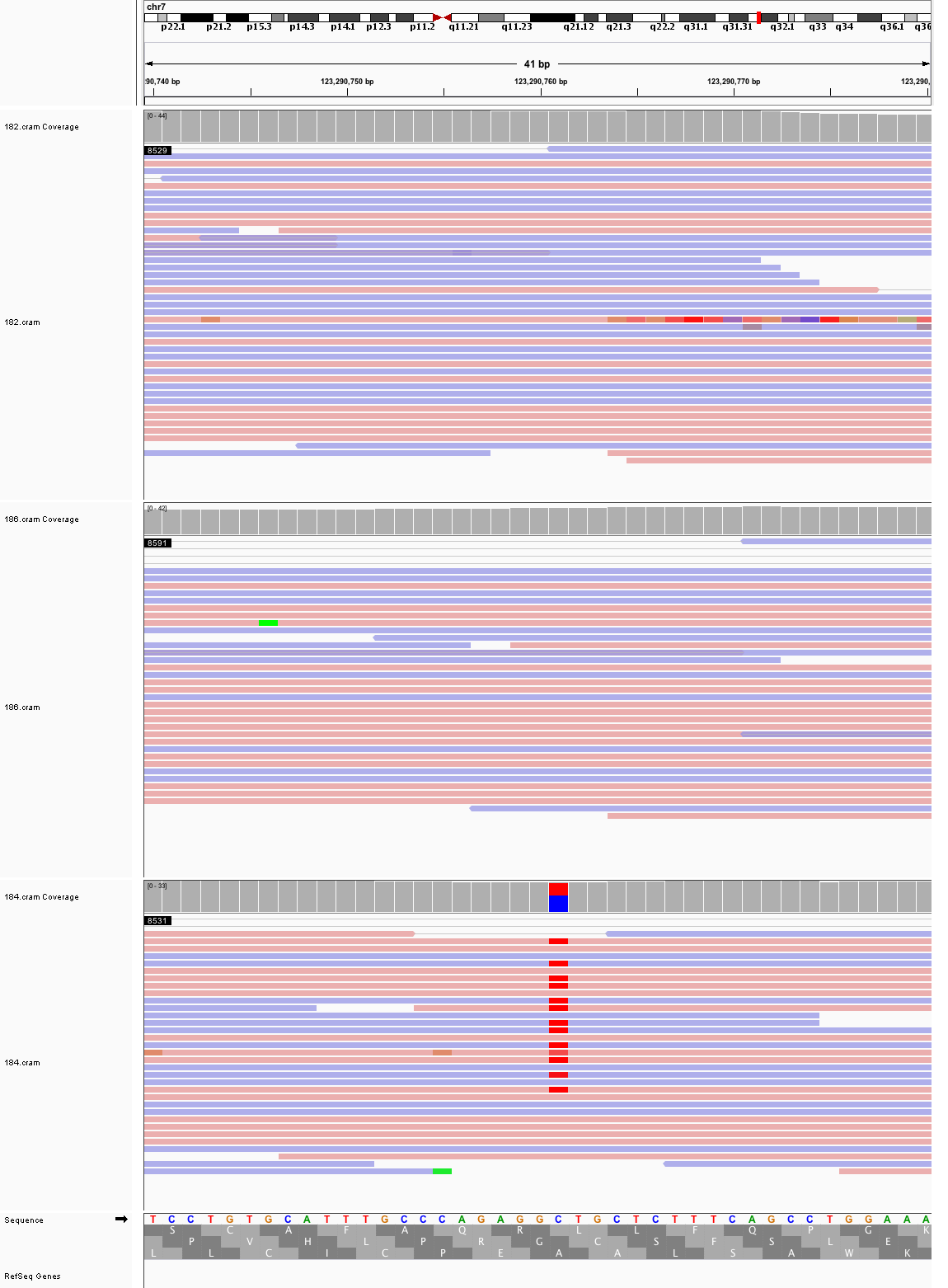

### chr7_140,630,032_140,630,072.png

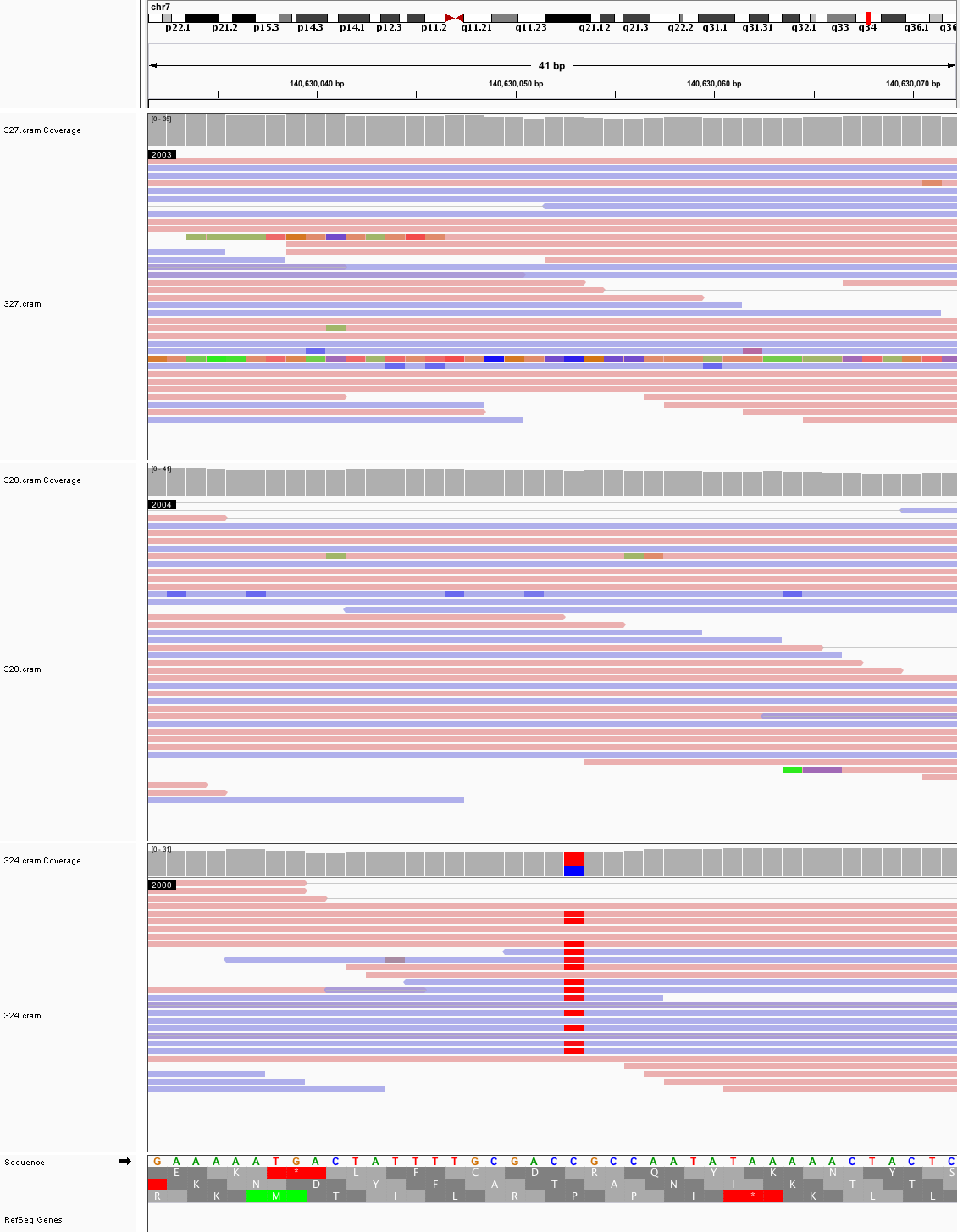

### chr8_20,915,131_20,915,171.png

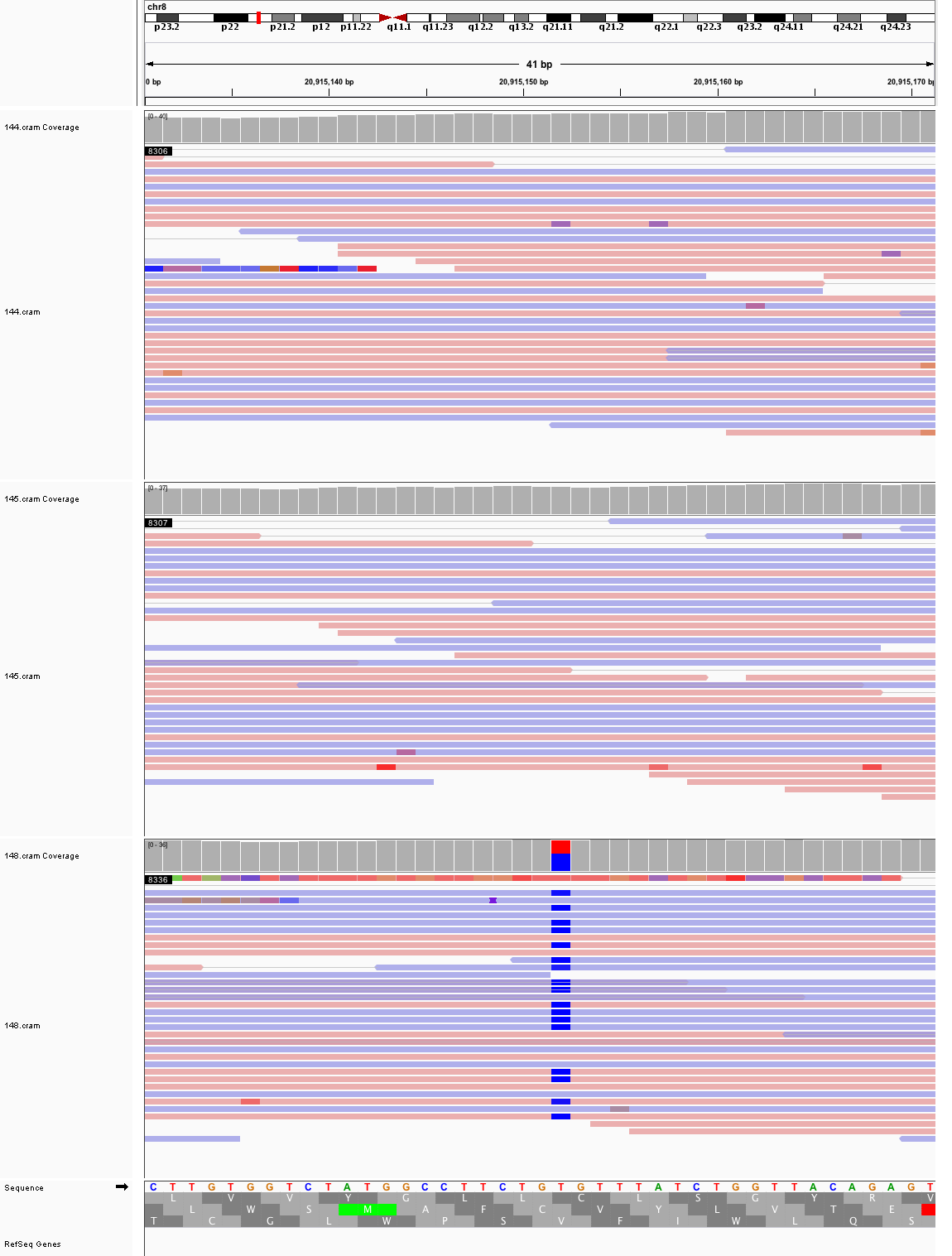

### chr8_23,566,778_23,566,818.png

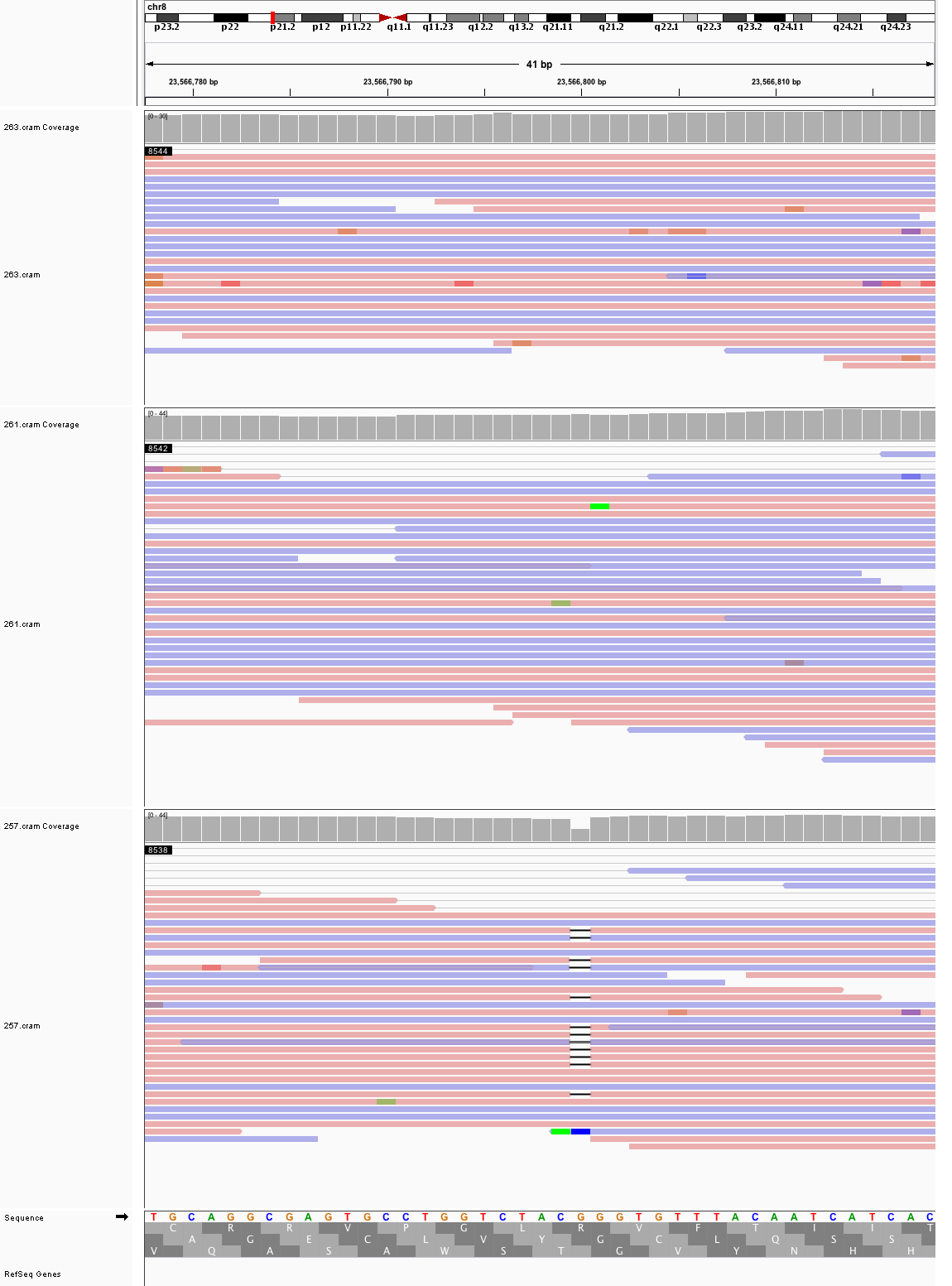

### chr8_56,319,840_56,319,880.png

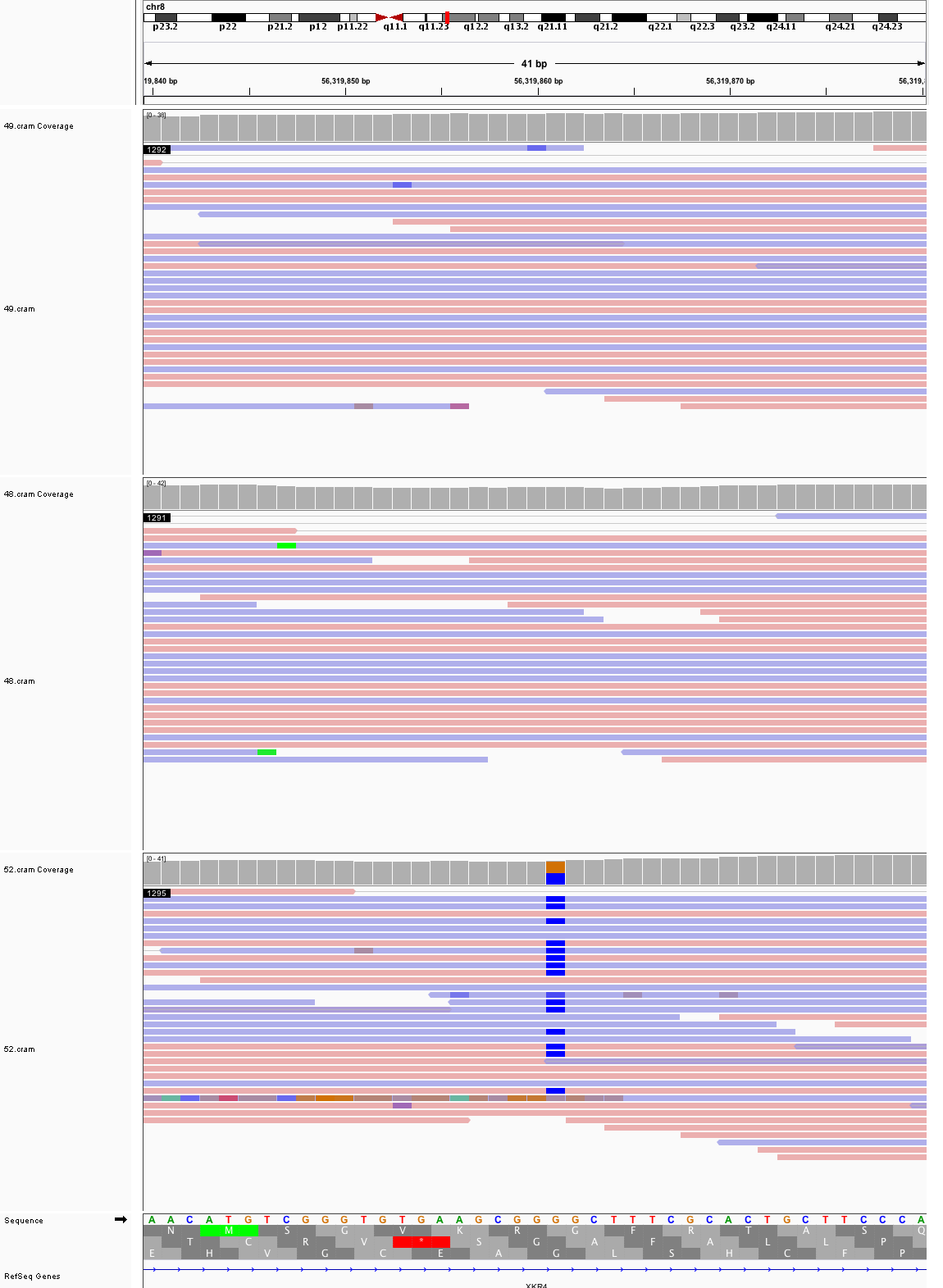

### chr8_94,267,505_94,267,545.png

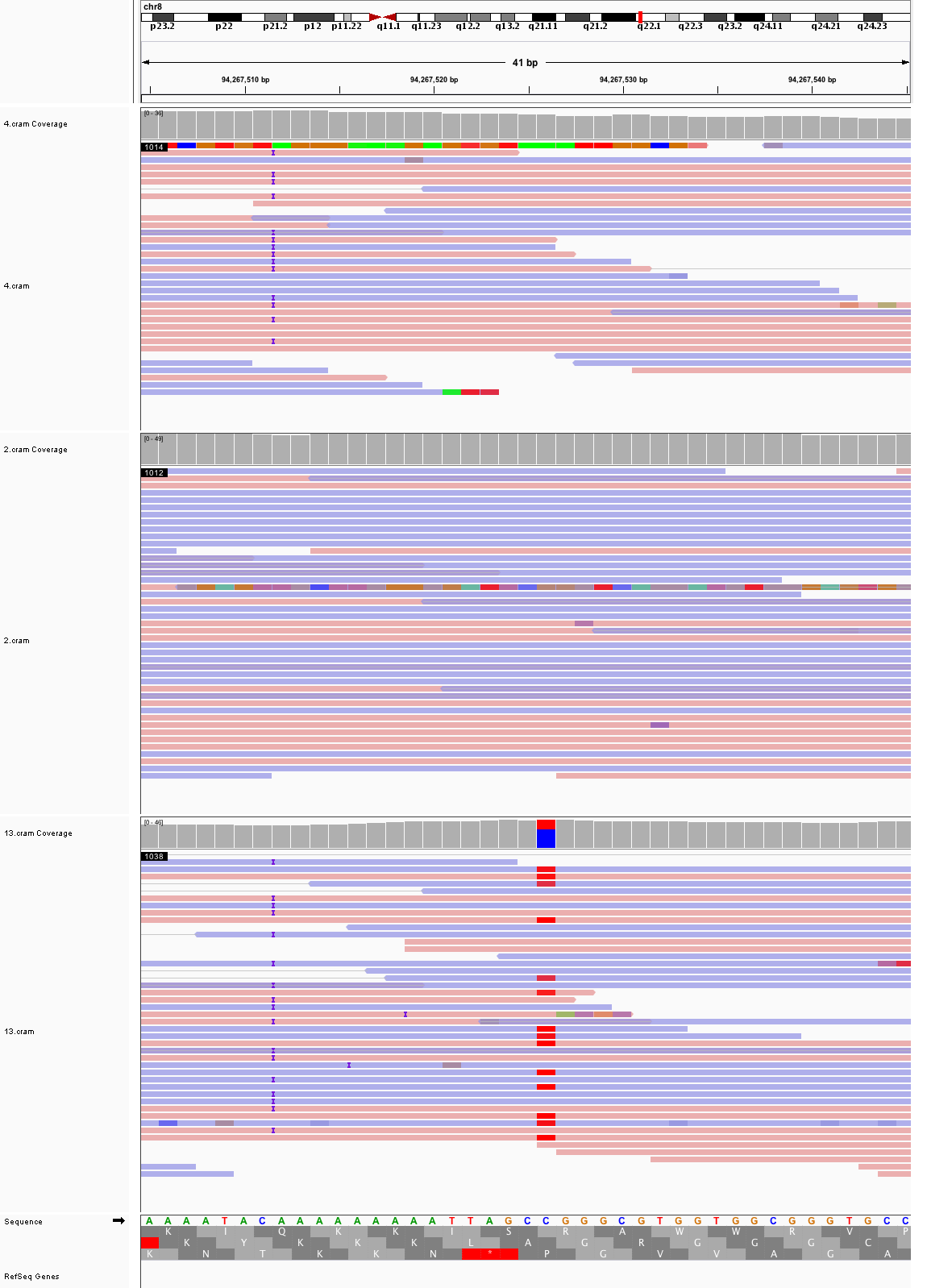

### chr8_121,574,492_121,574,532.png

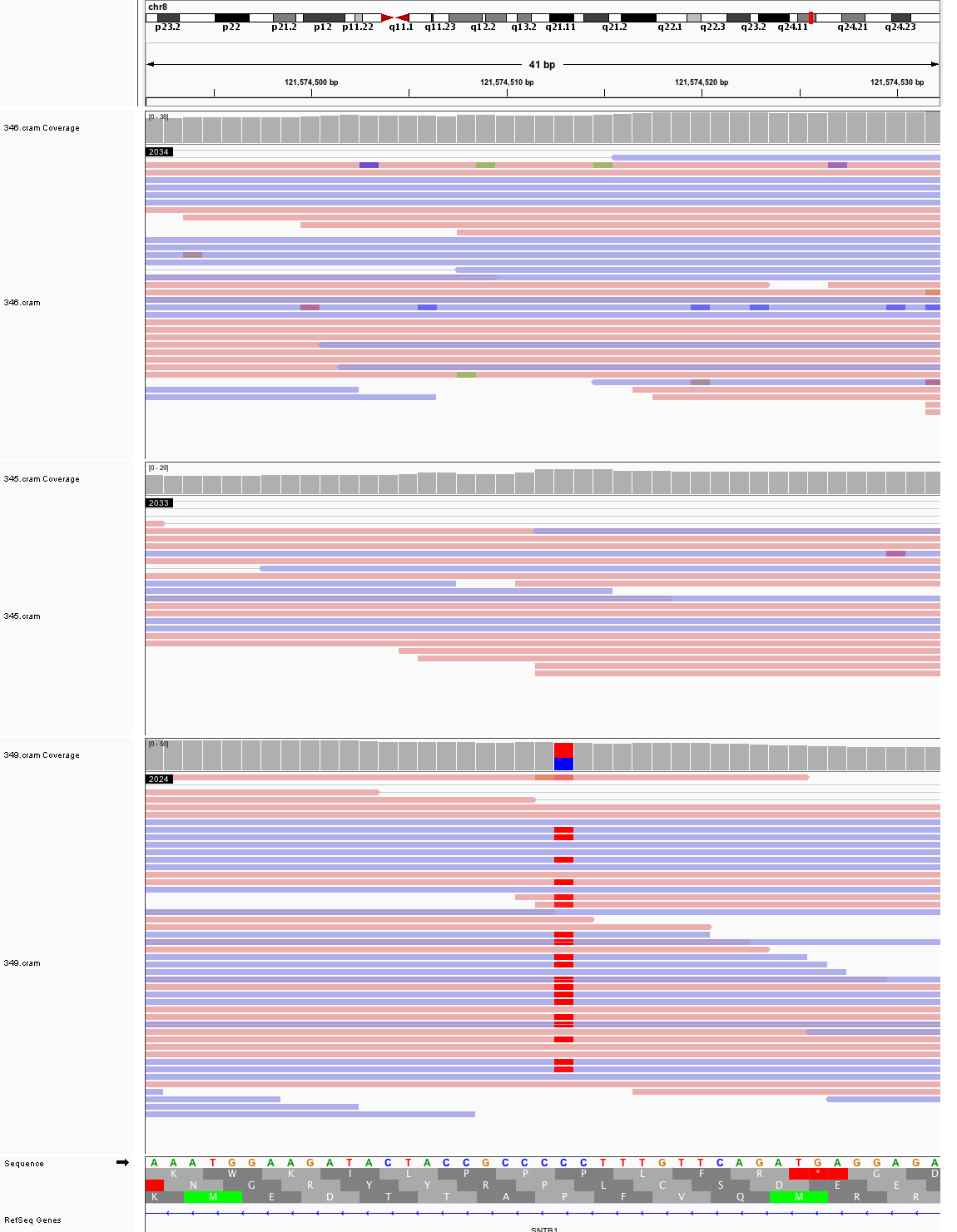

### chr9_23,353,467_23,353,507.png

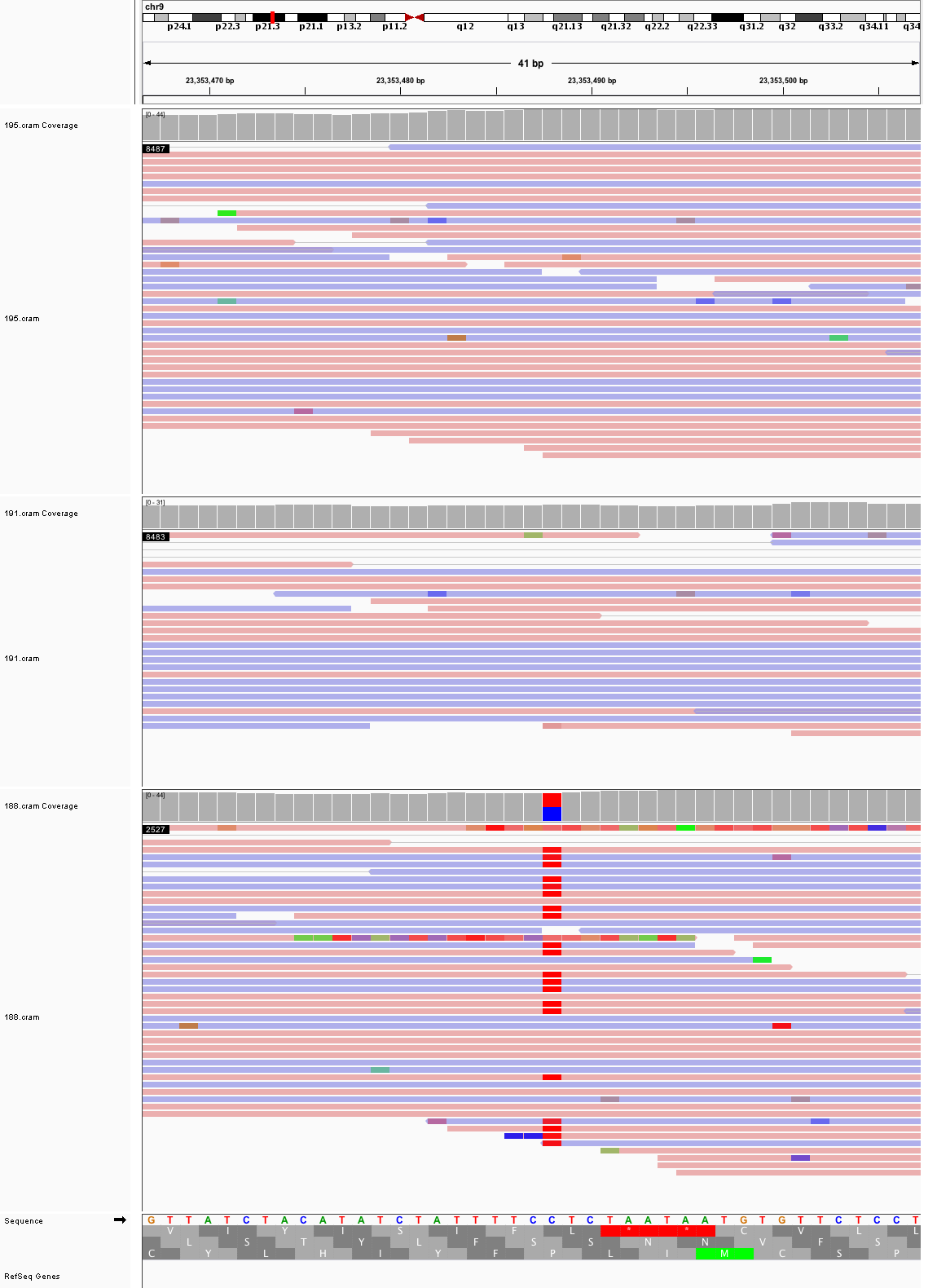

### chr9_23,942,987_23,943,027.png

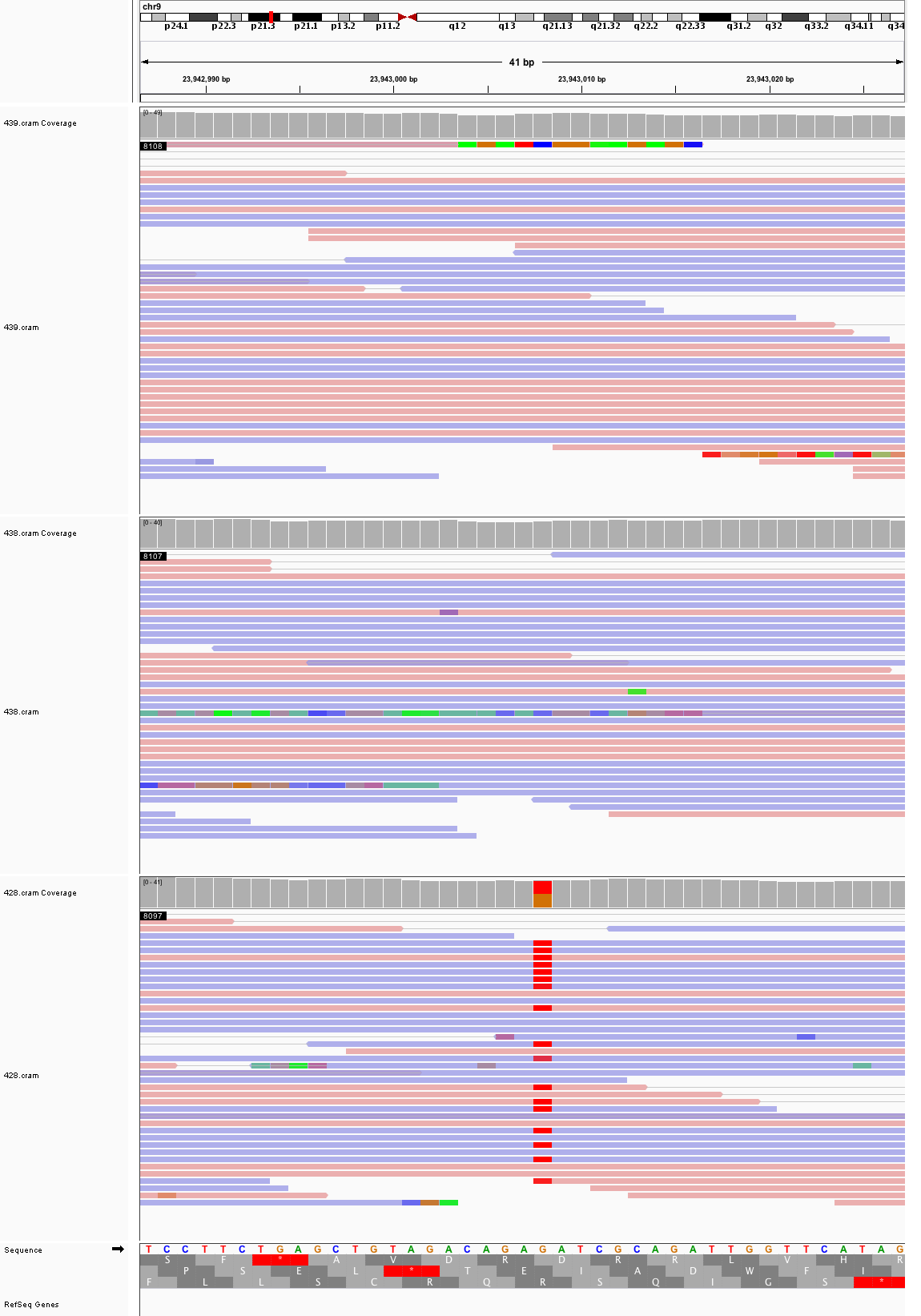

### chr9_26,014,107_26,014,147.png

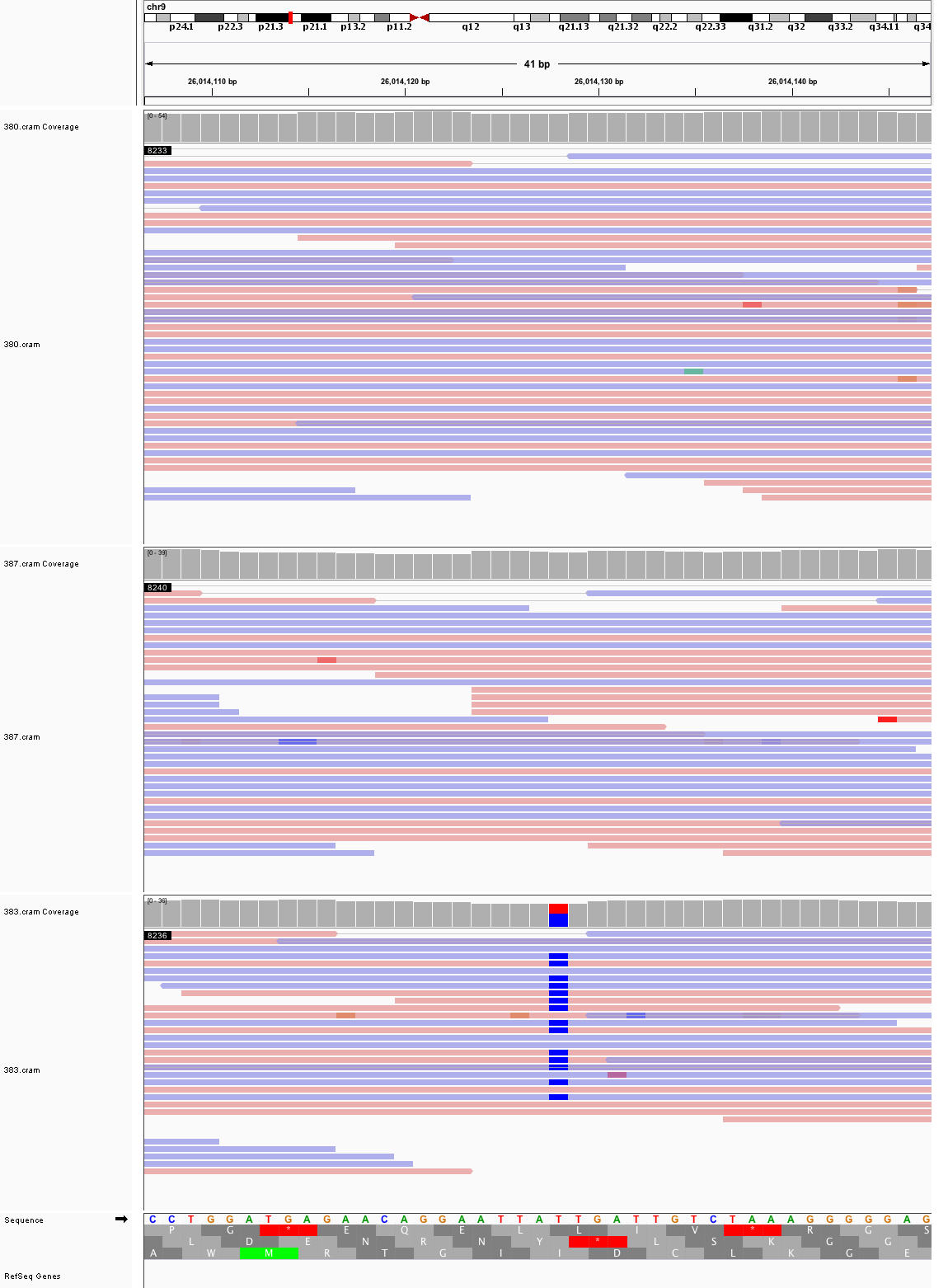

### chr9_68,232,749_68,232,789.png

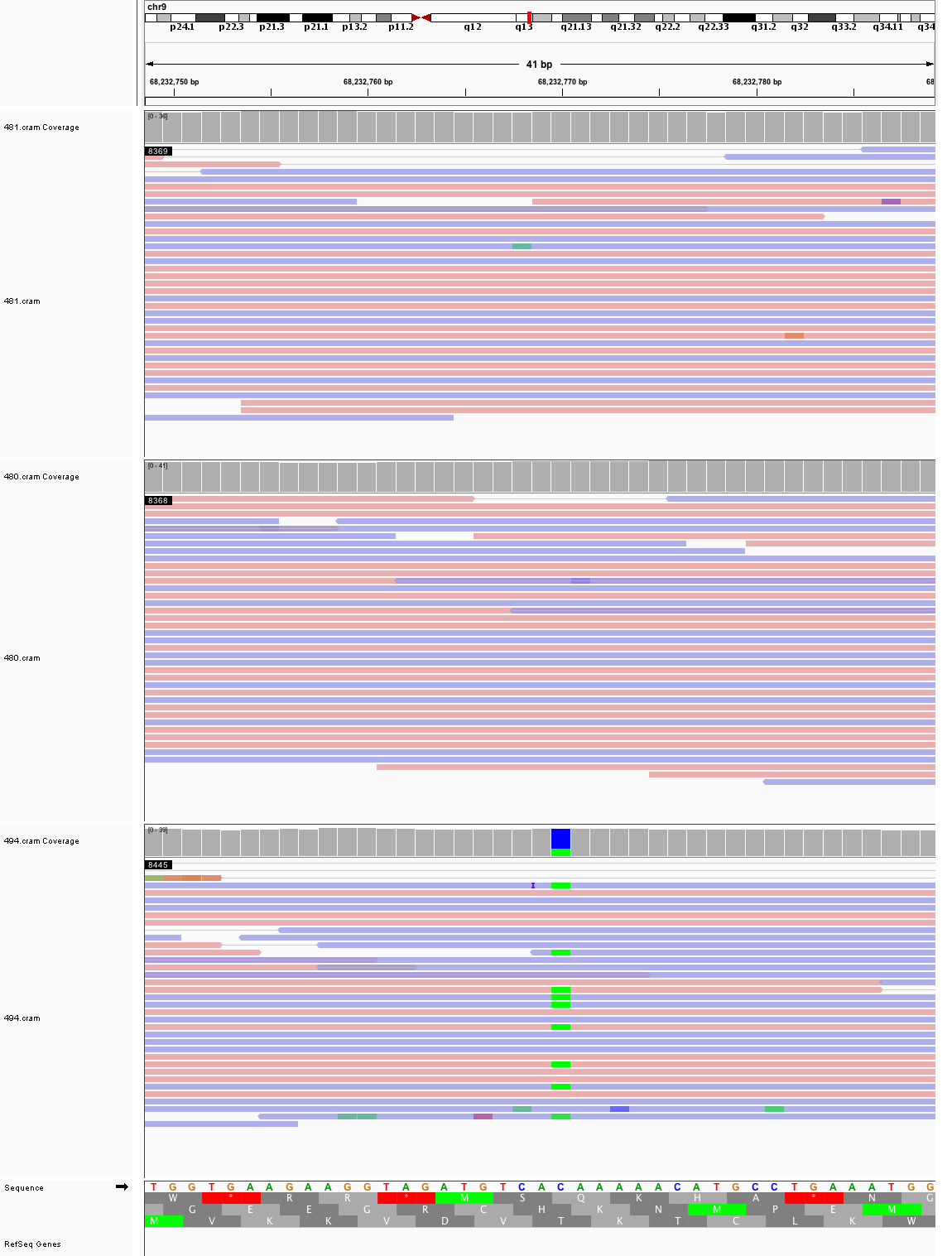

### chr9_84,505,465_84,505,505.png

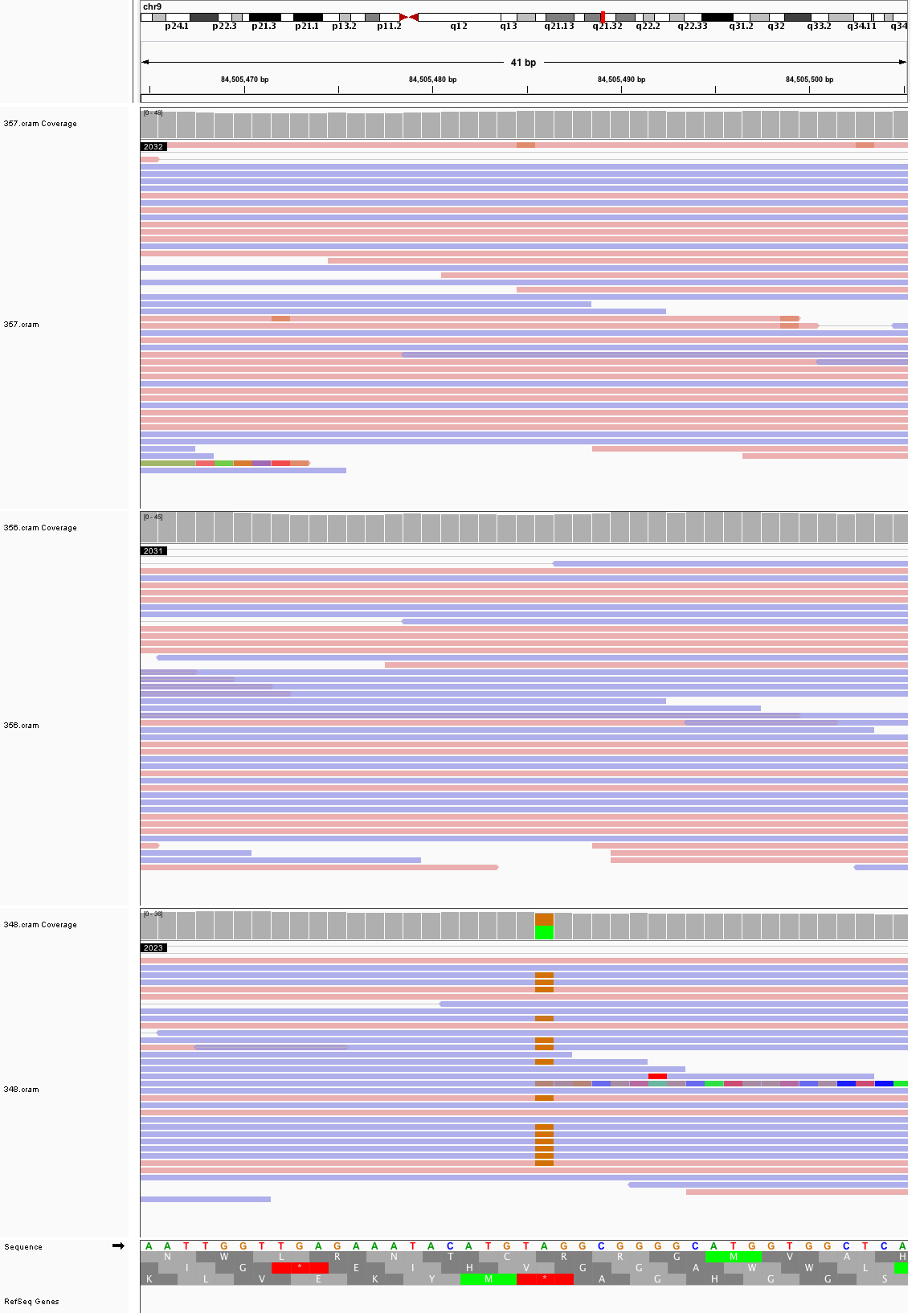

### chr9_85,494,122_85,494,162.png

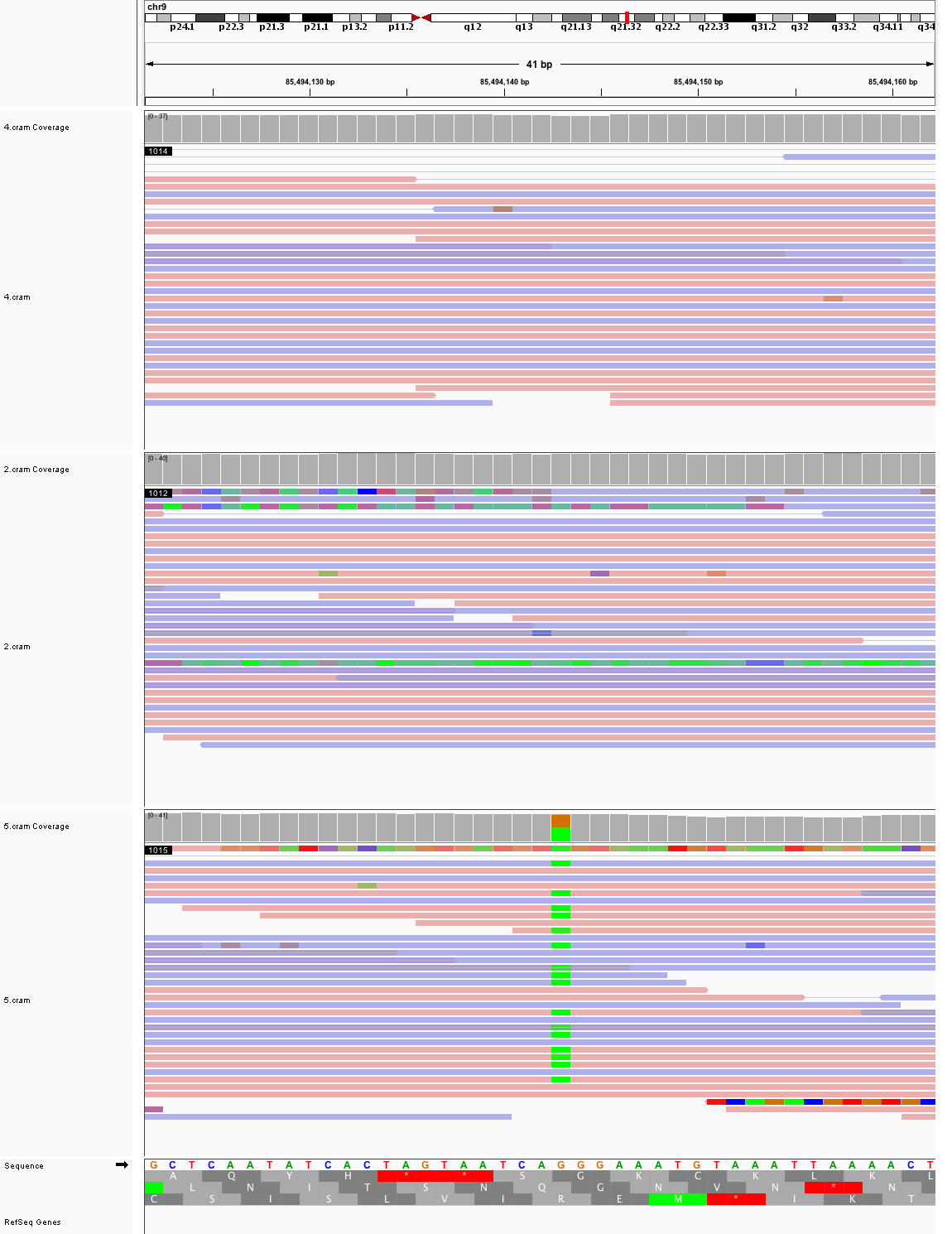

### chr9_89,035,942_89,035,982.png

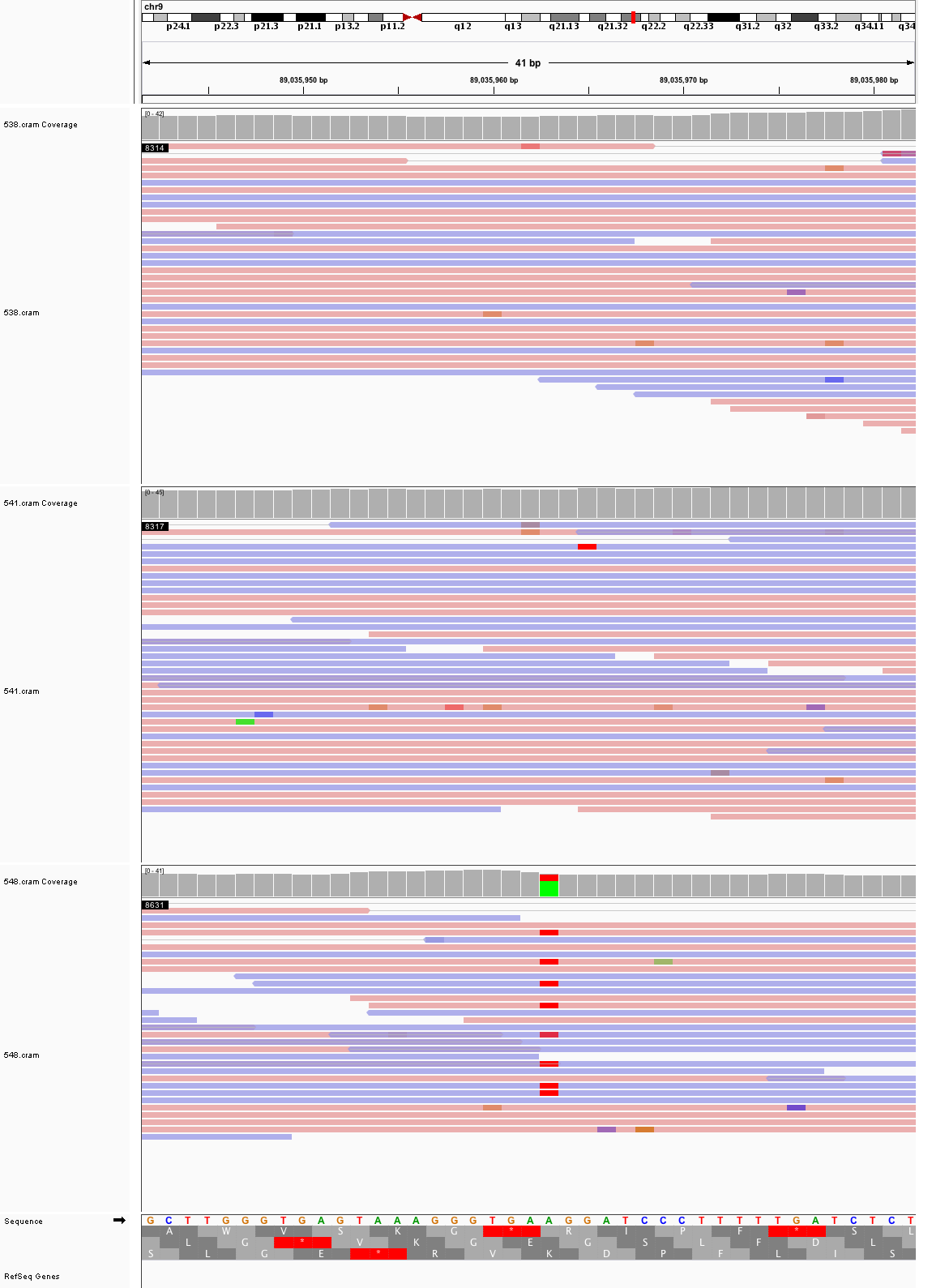

### chr9_114,782,560_114,782,600.png

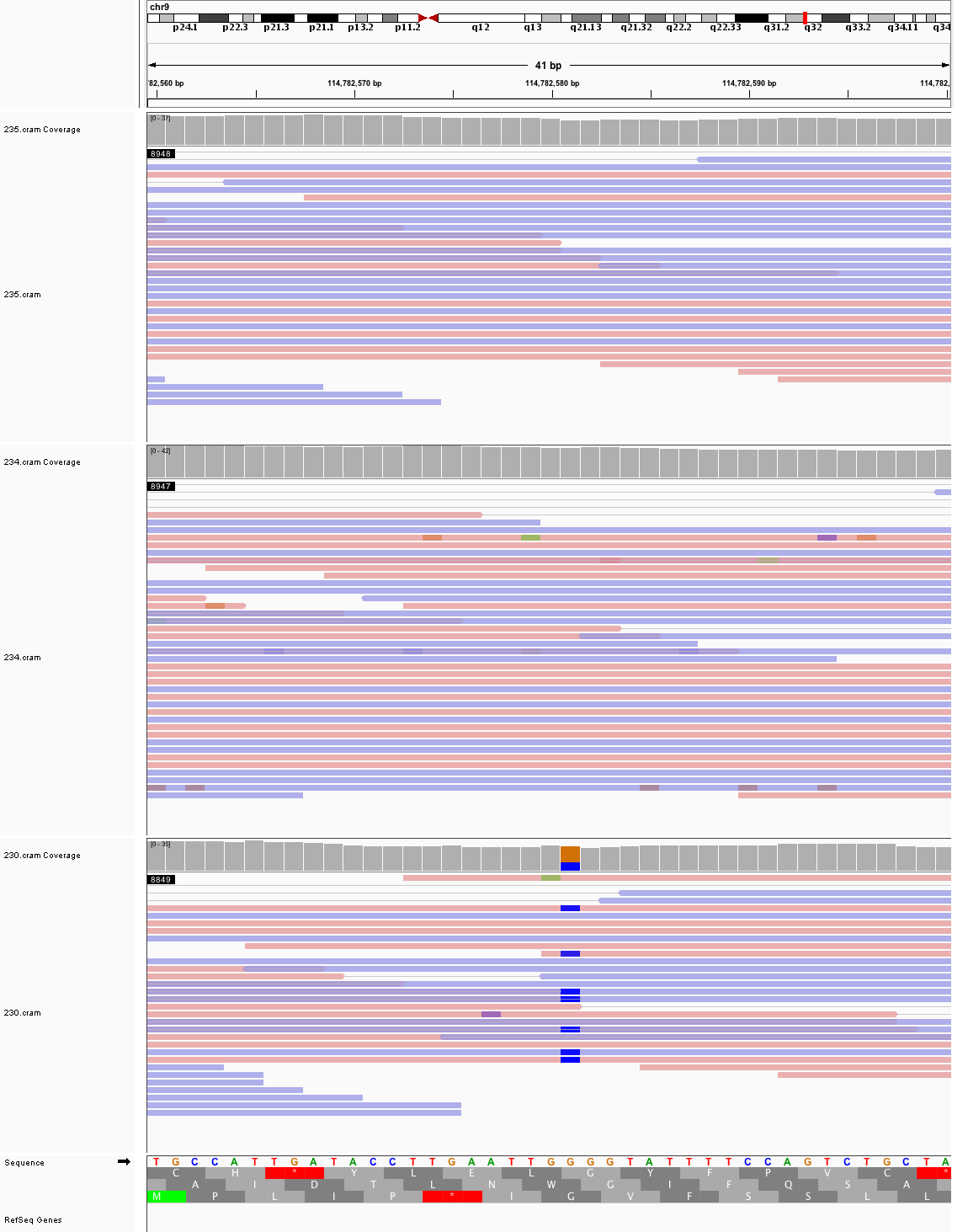

### chr9_139,914,900_139,914,940.png

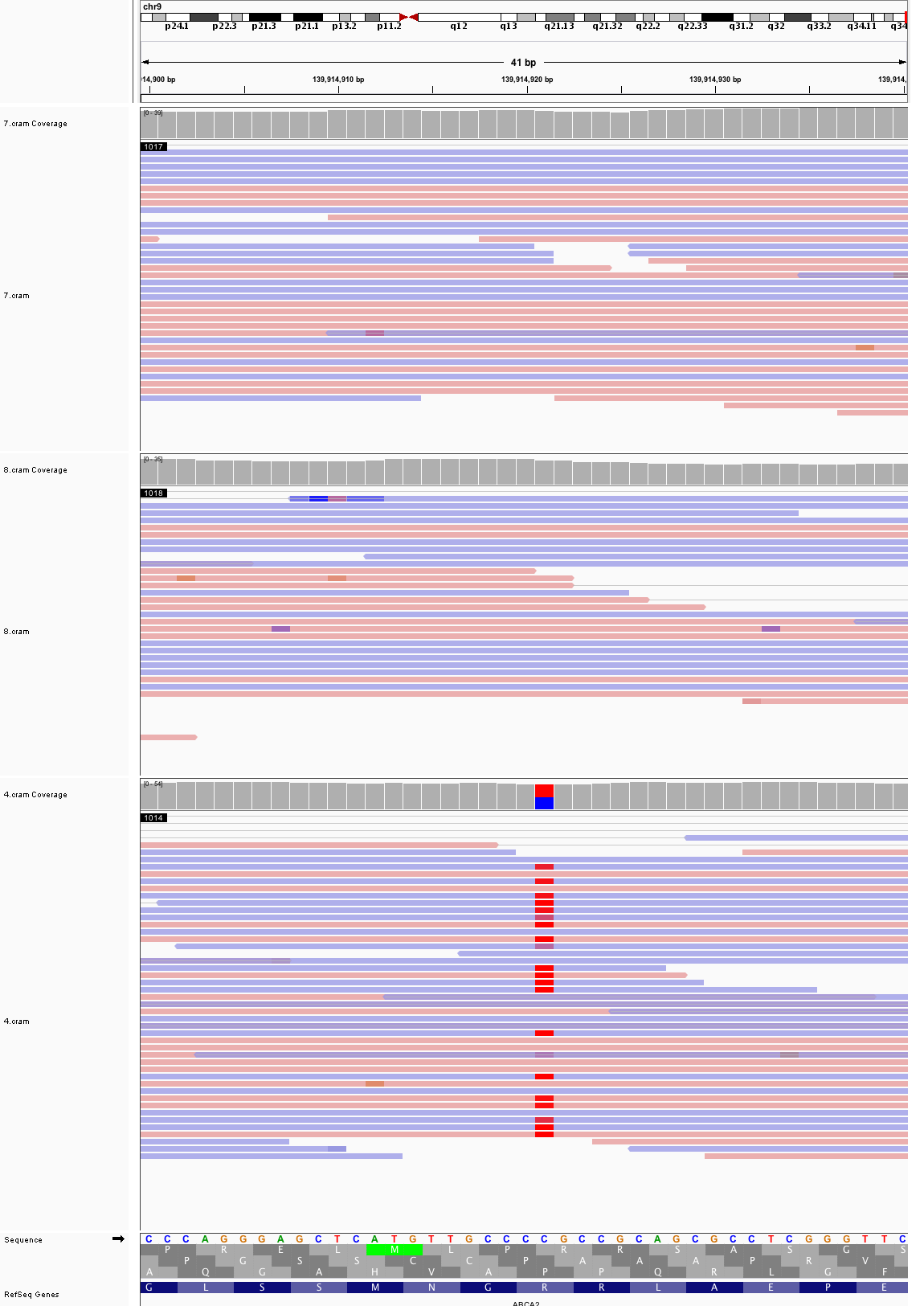

### chr10_36,620,311_36,620,351.png

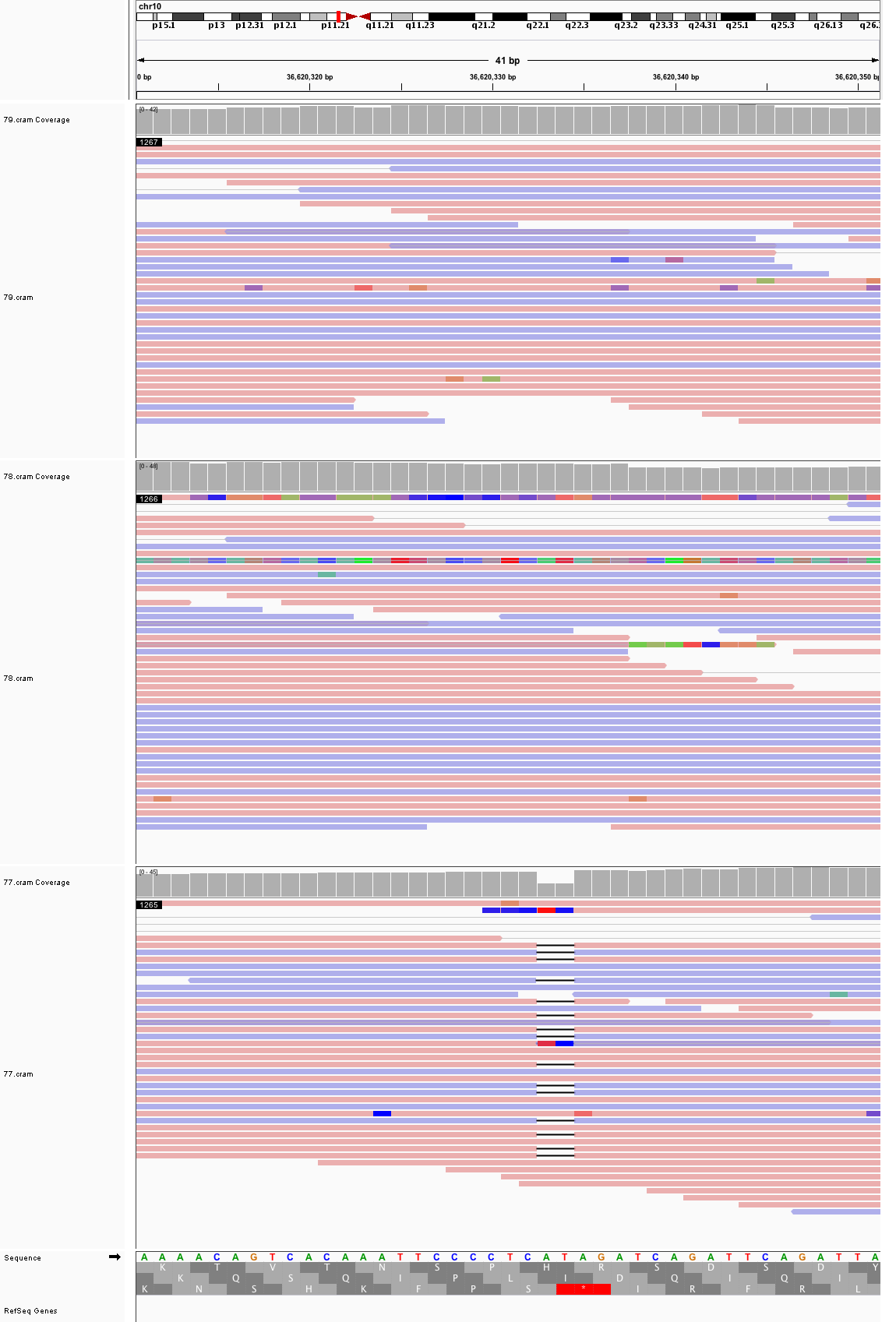

### chr10_53,663,171_53,663,211.png

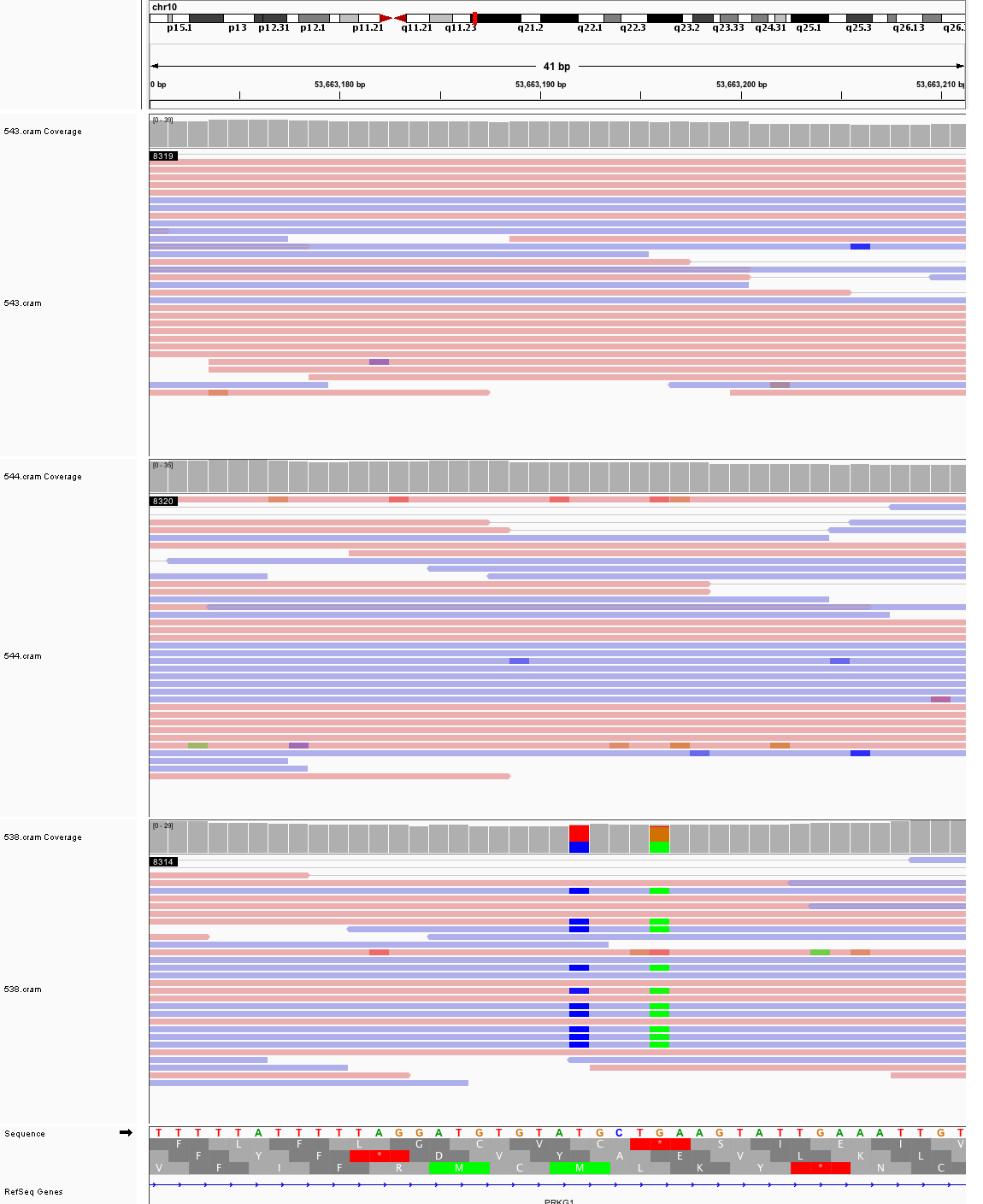

### chr10_108,567,404_108,567,444.png

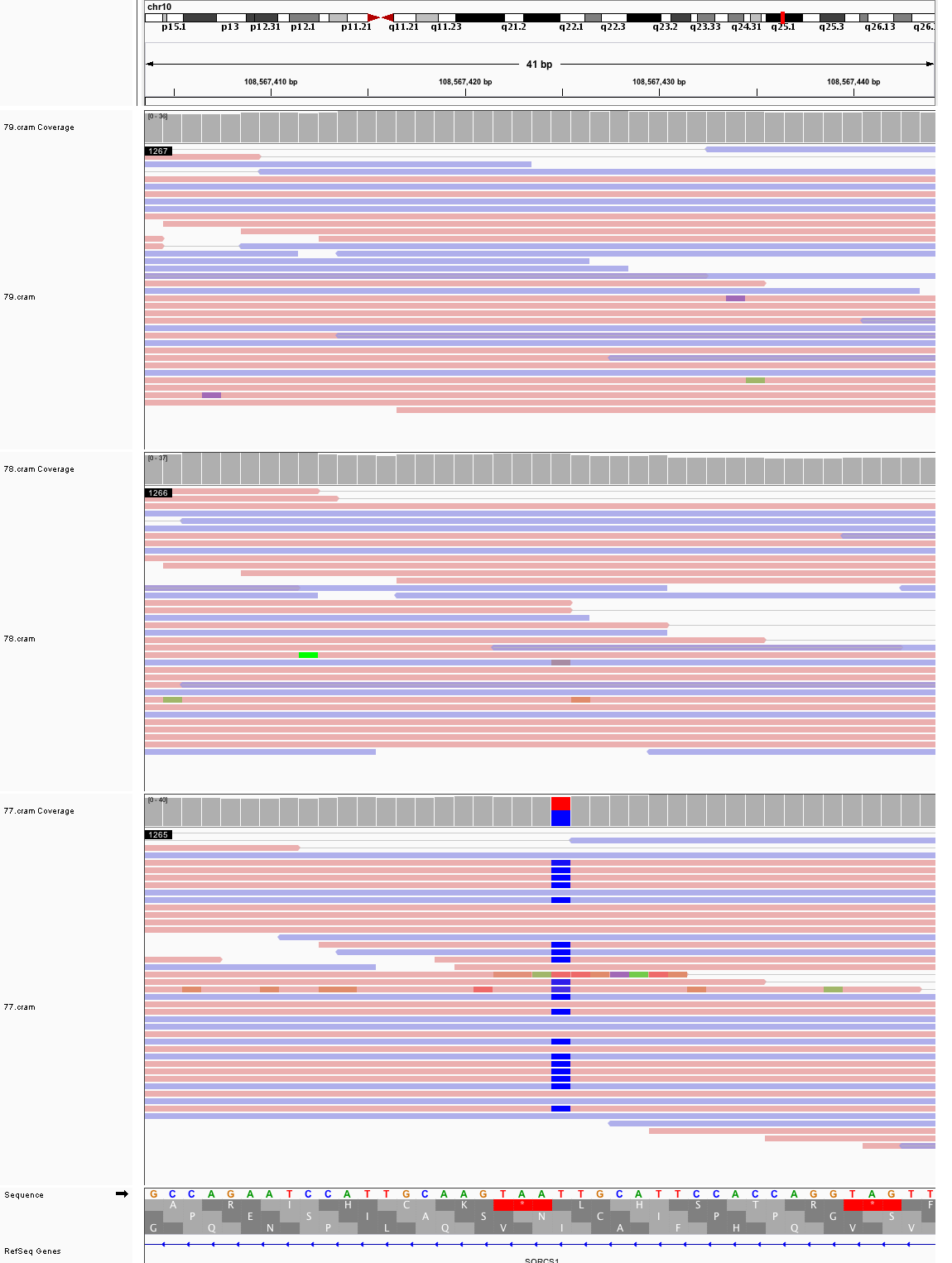

### chr11_16,799,136_16,799,176.png

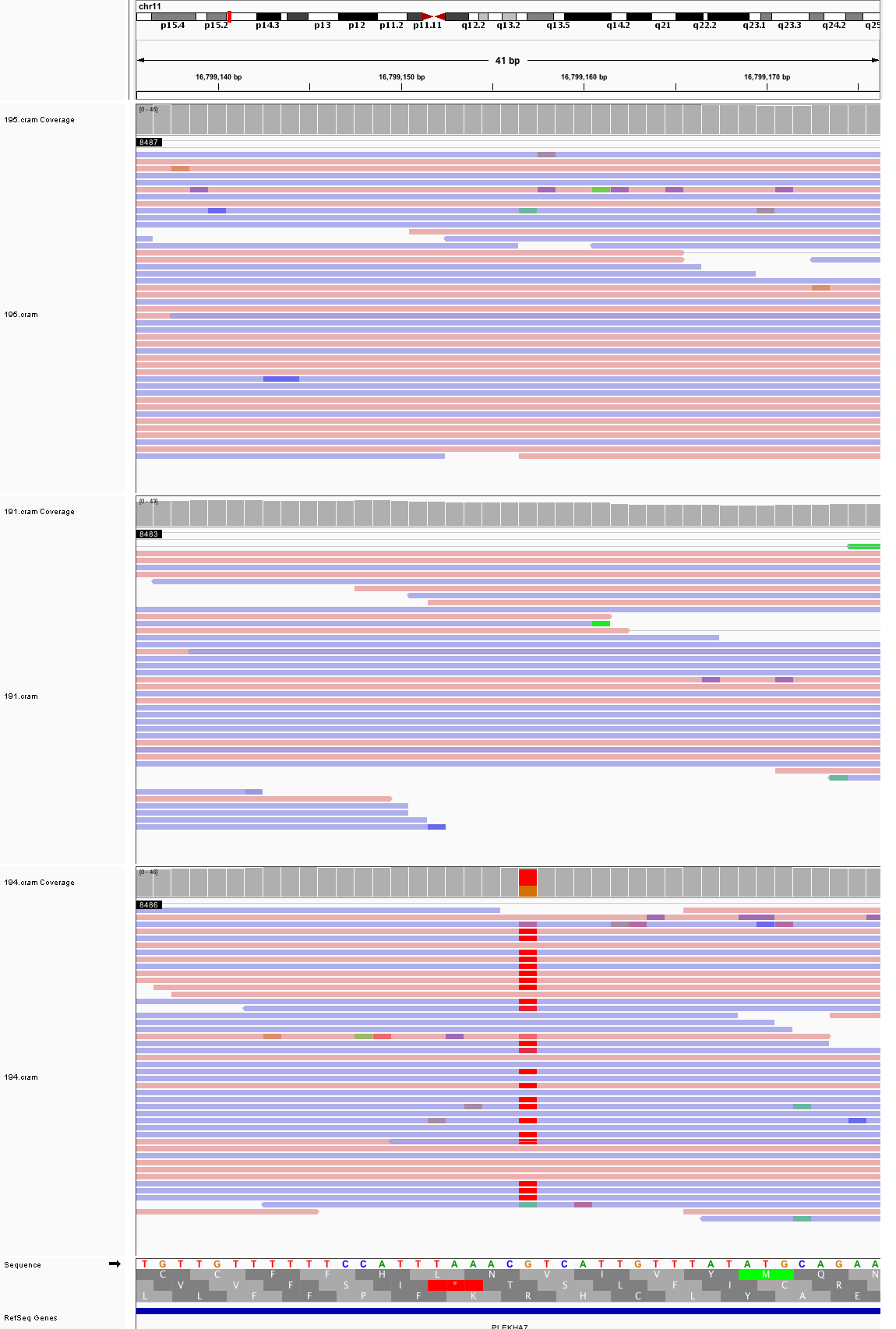

### chr11_76,993,067_76,993,107.png

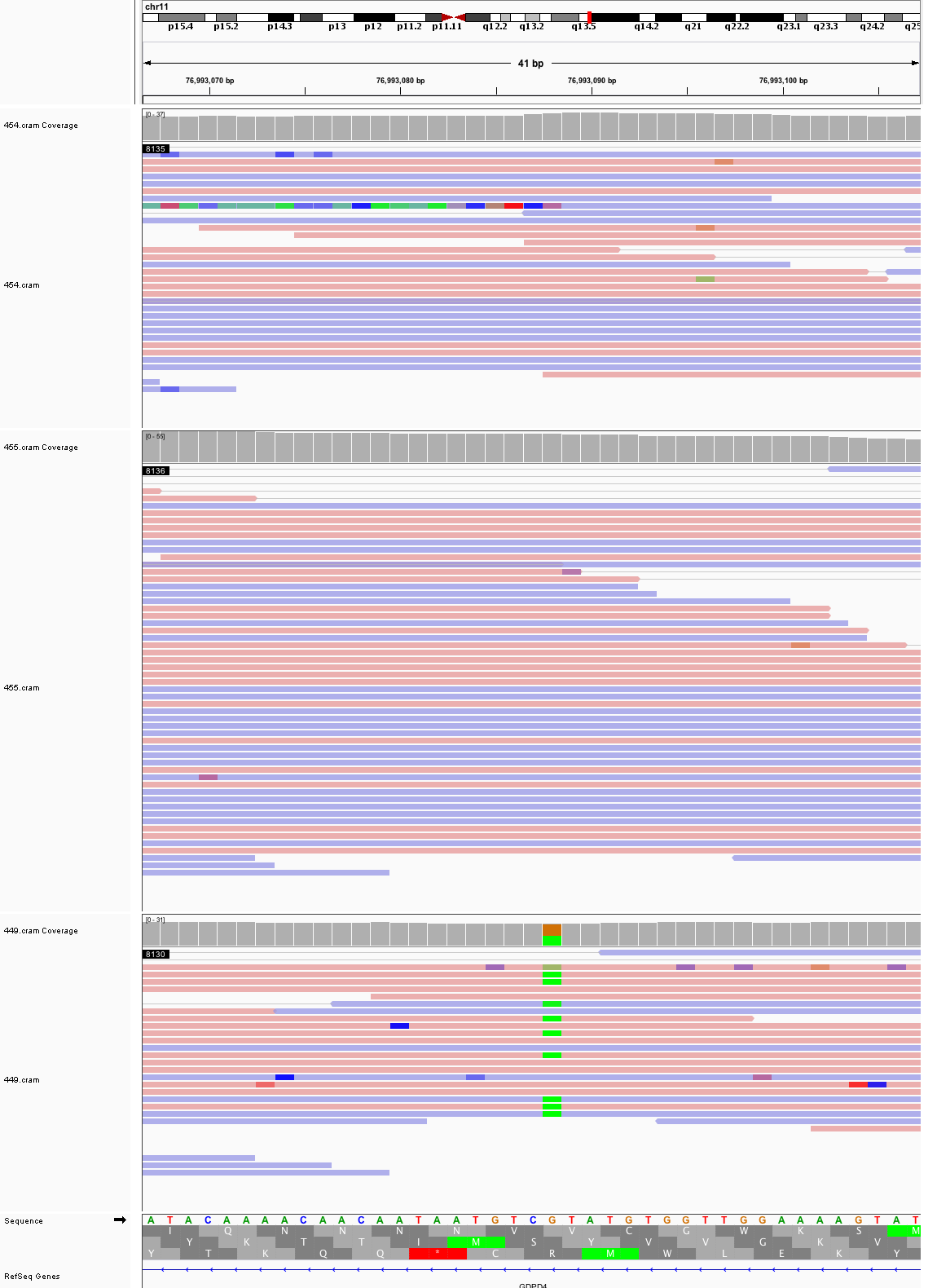

### chr11_87,928,865_87,928,905.png

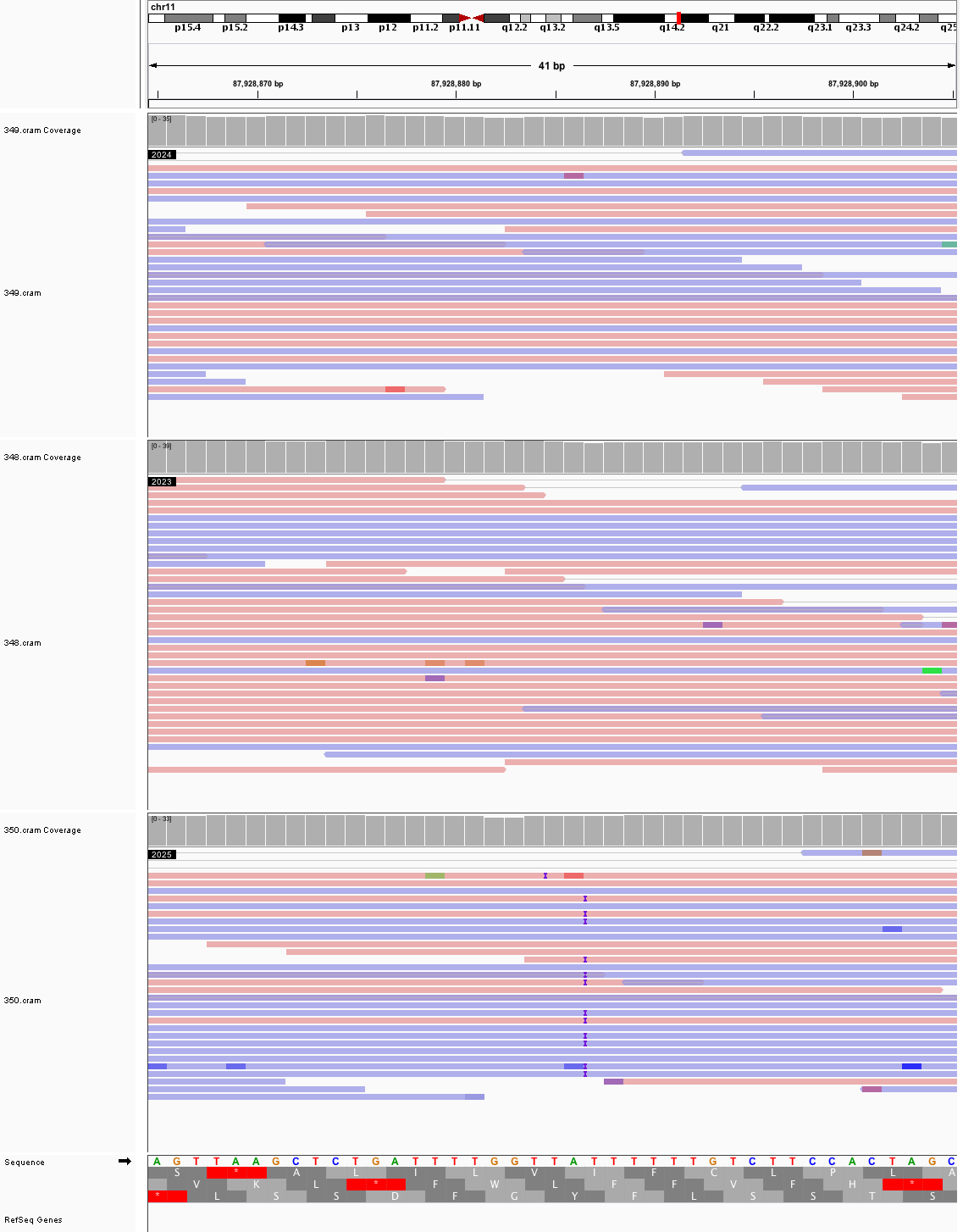

### chr11_102,659,946_102,659,986.png

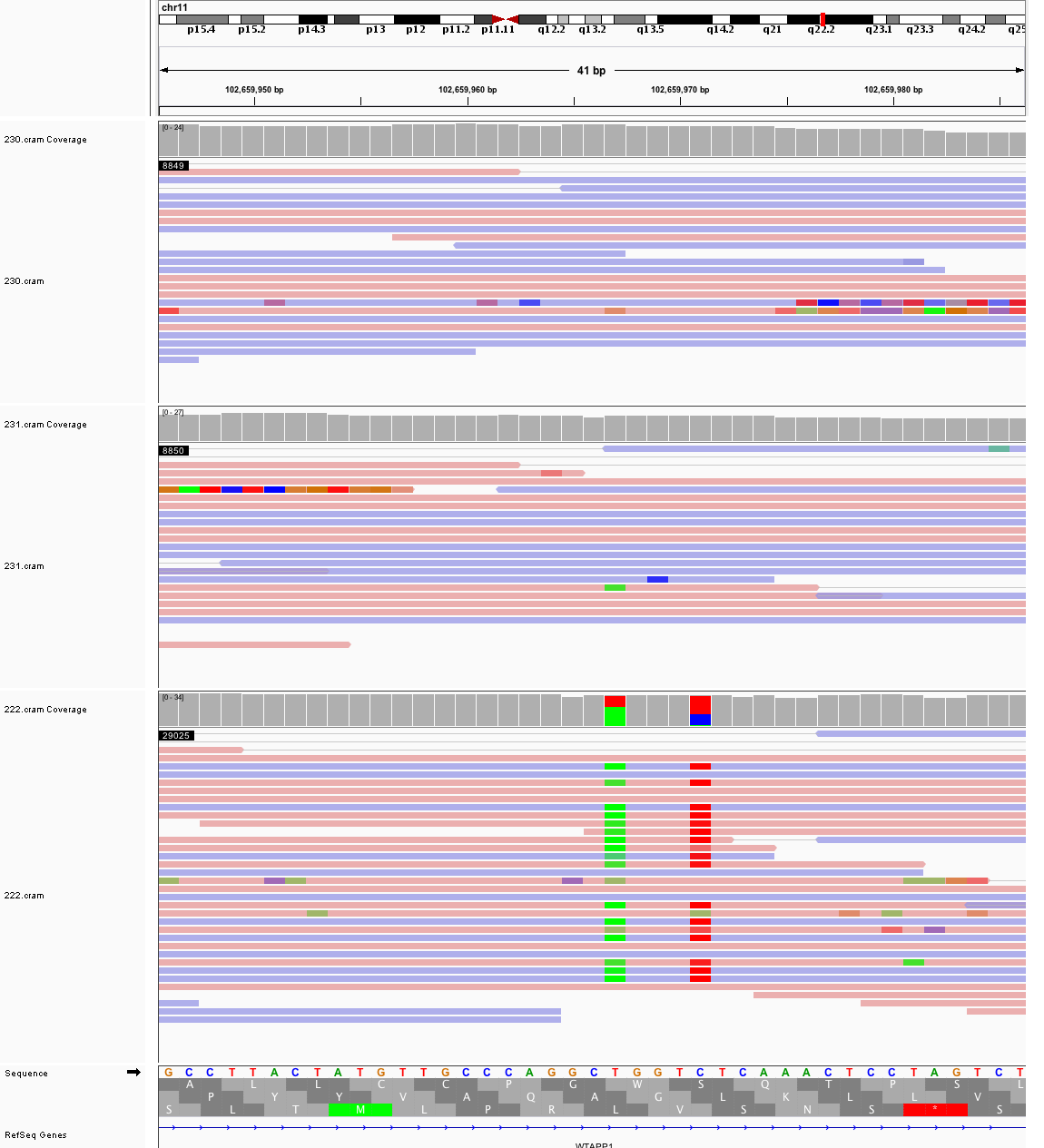

### chr11_104,948,755_104,948,795.png

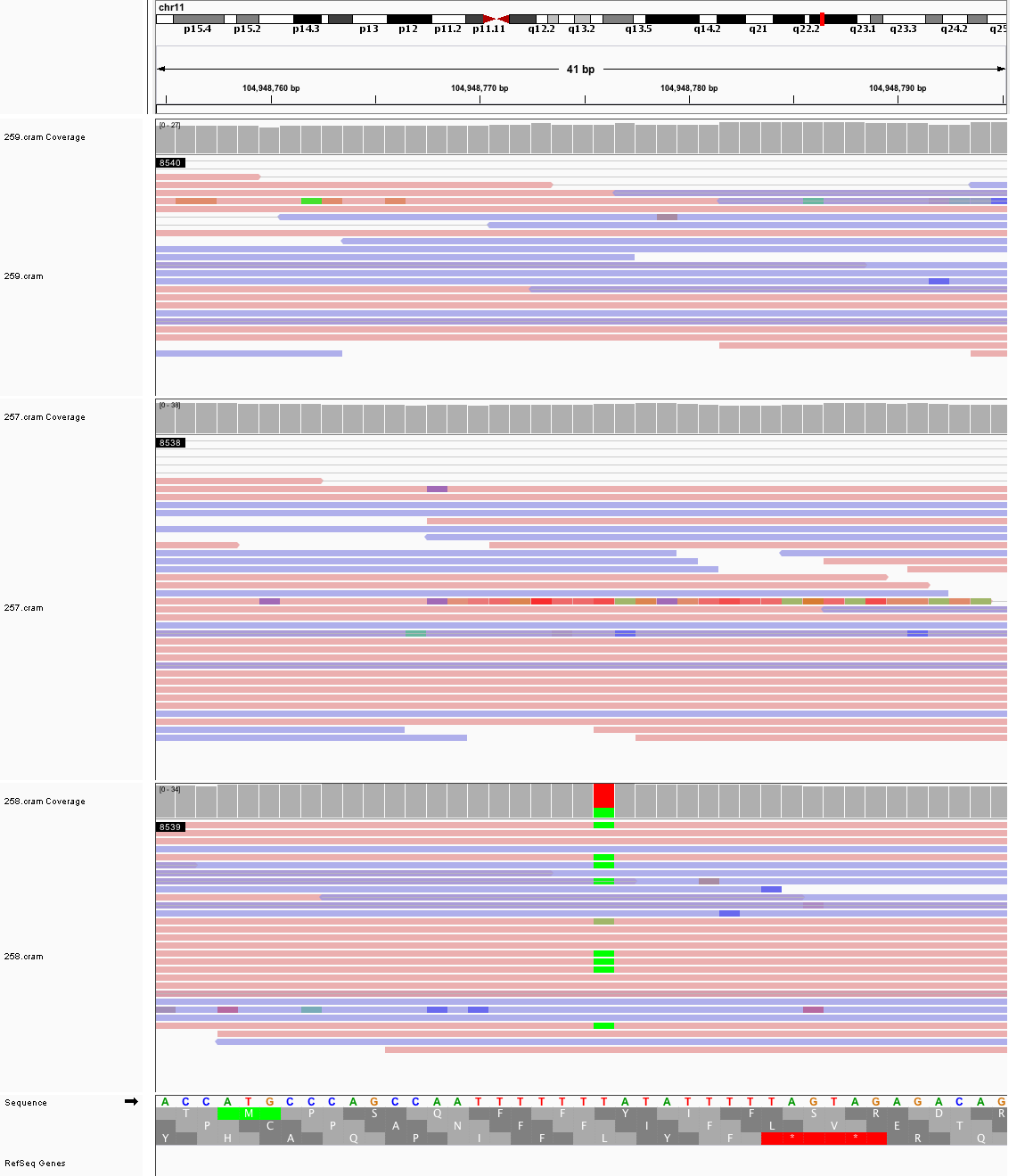

### chr11_108,223,311_108,223,351.png

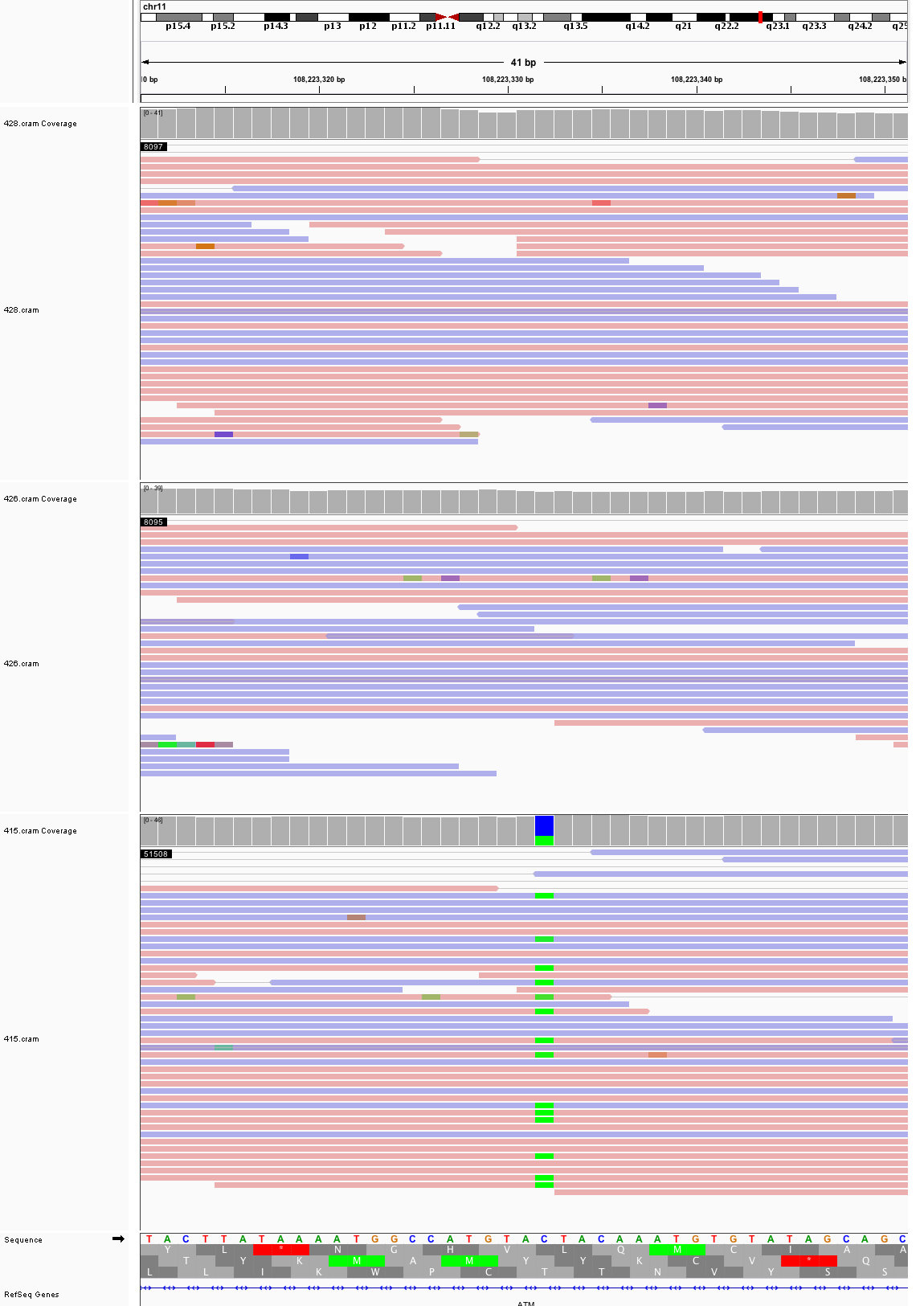

### chr11_113,123,787_113,123,827.png

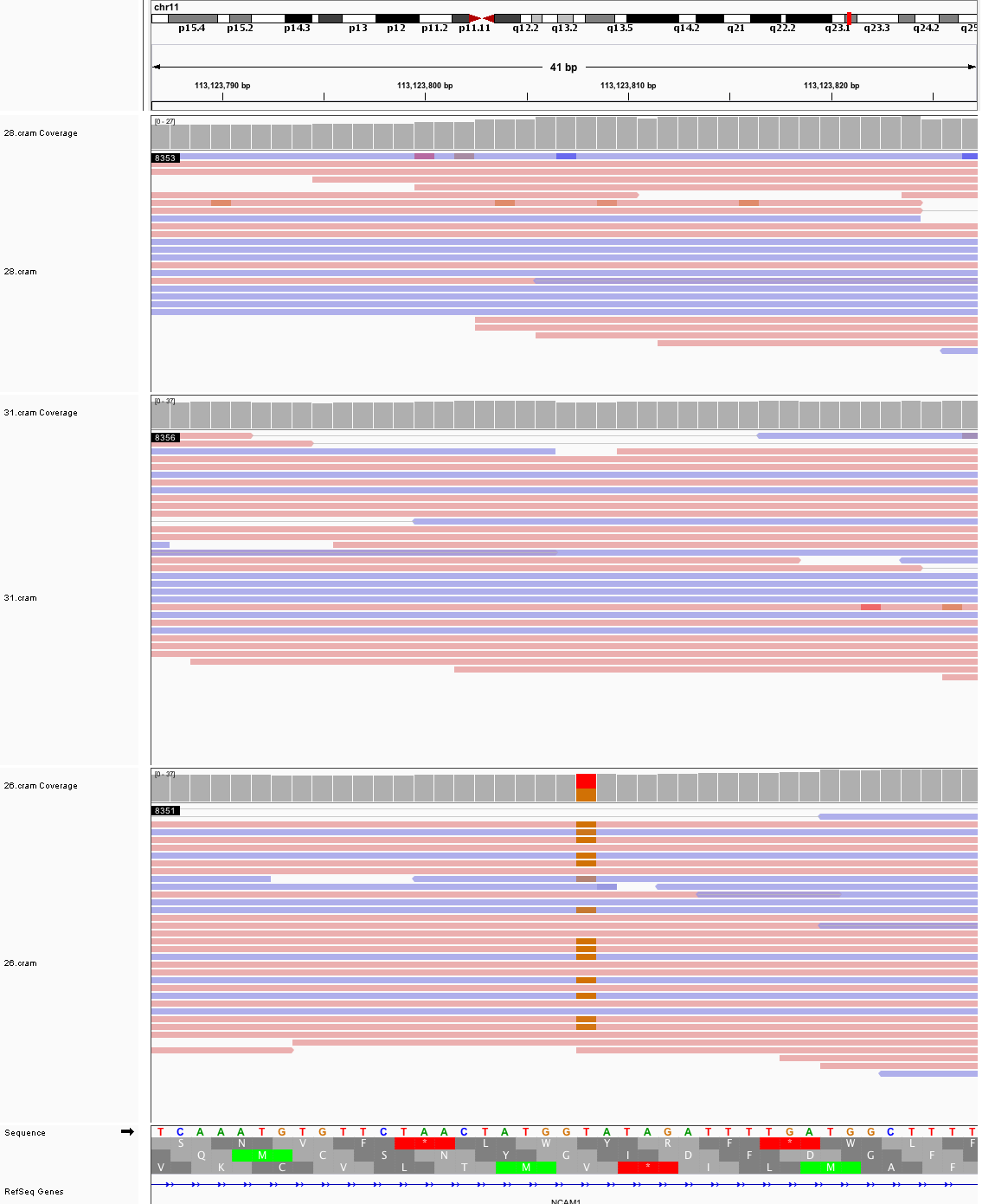

### chr11_128,618,461_128,618,501.png

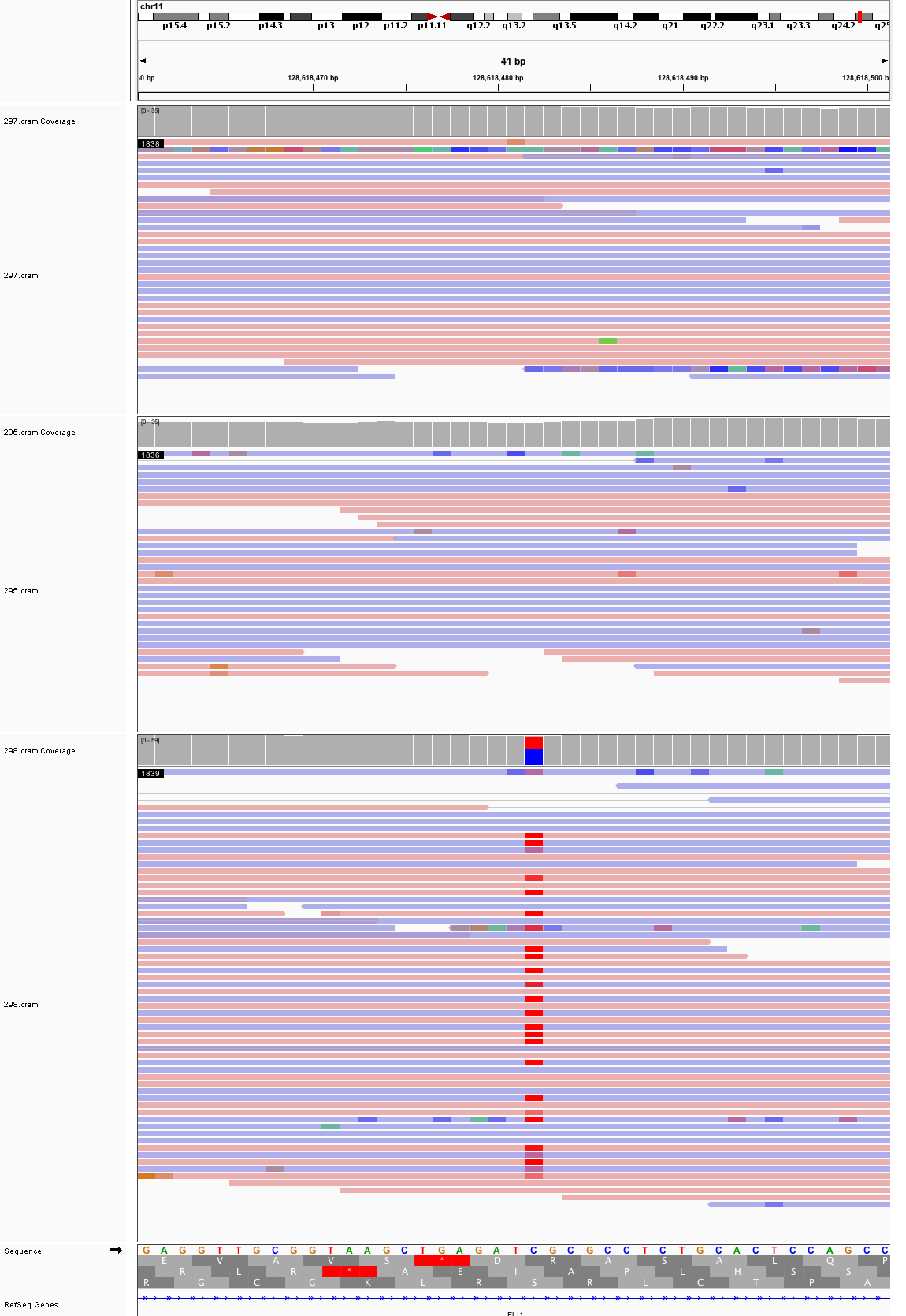

### chr11_130,606,145_130,606,185.png

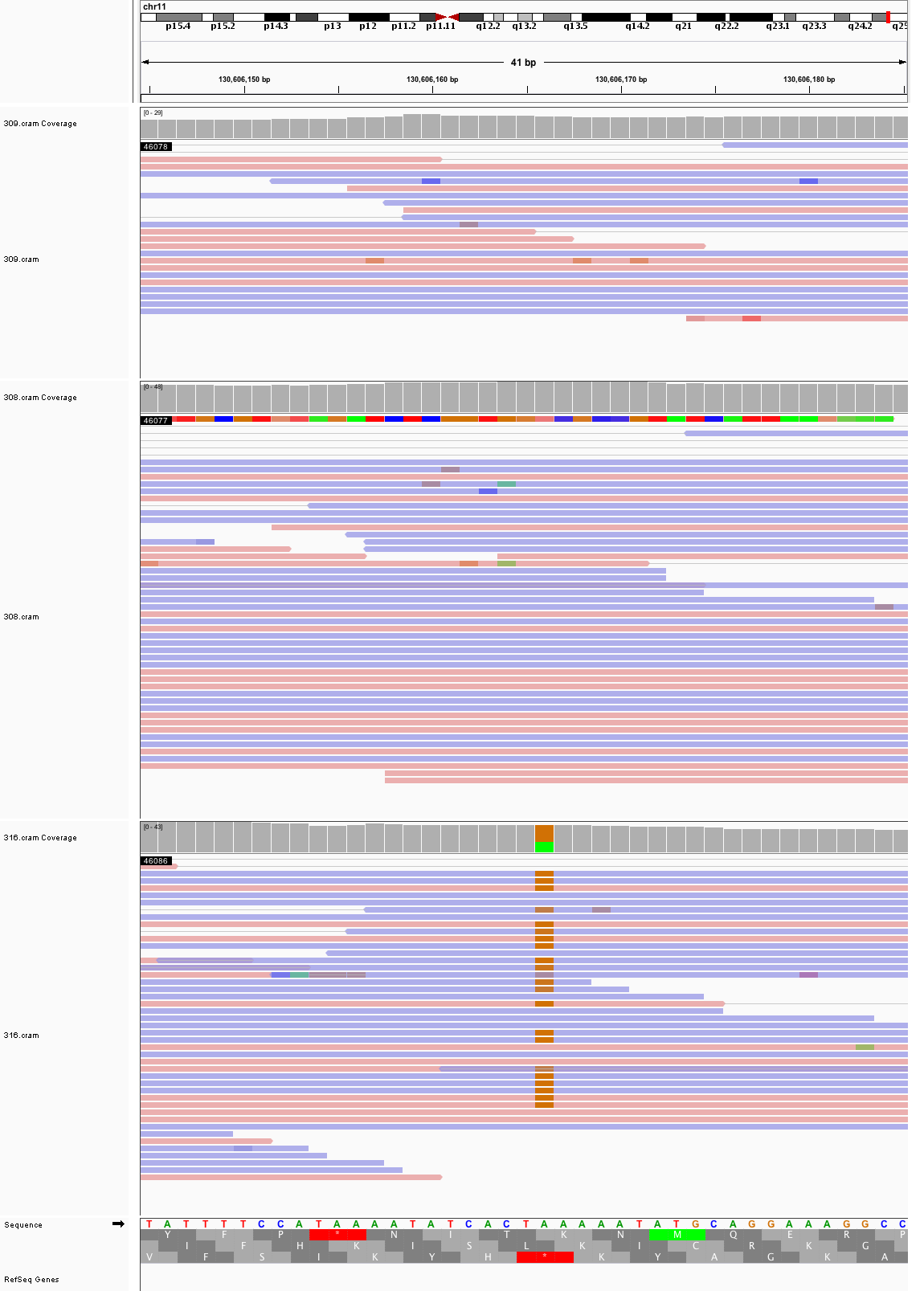

### chr12_23,752,121_23,752,161.png

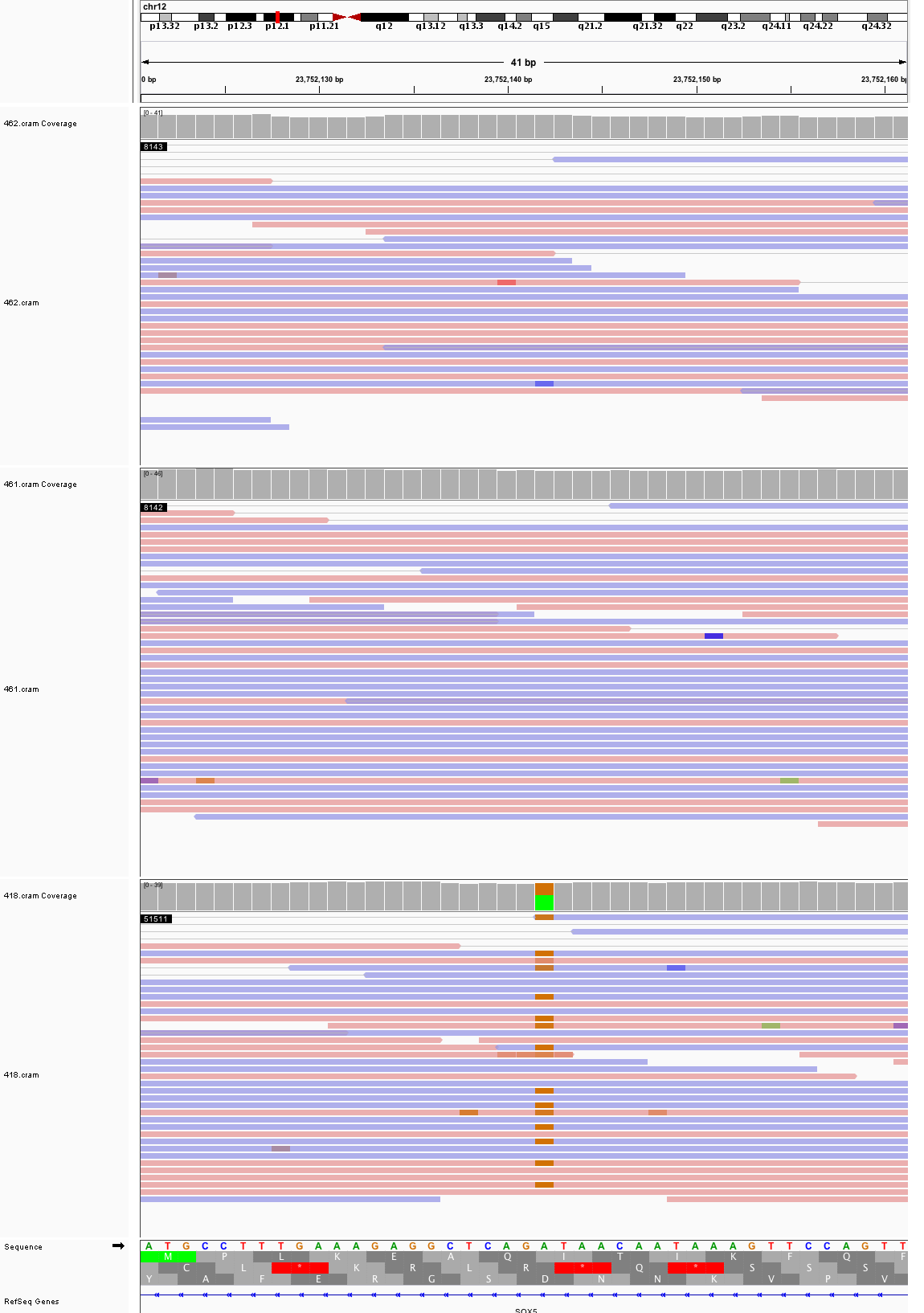

### chr12_80,282,270_80,282,310.png

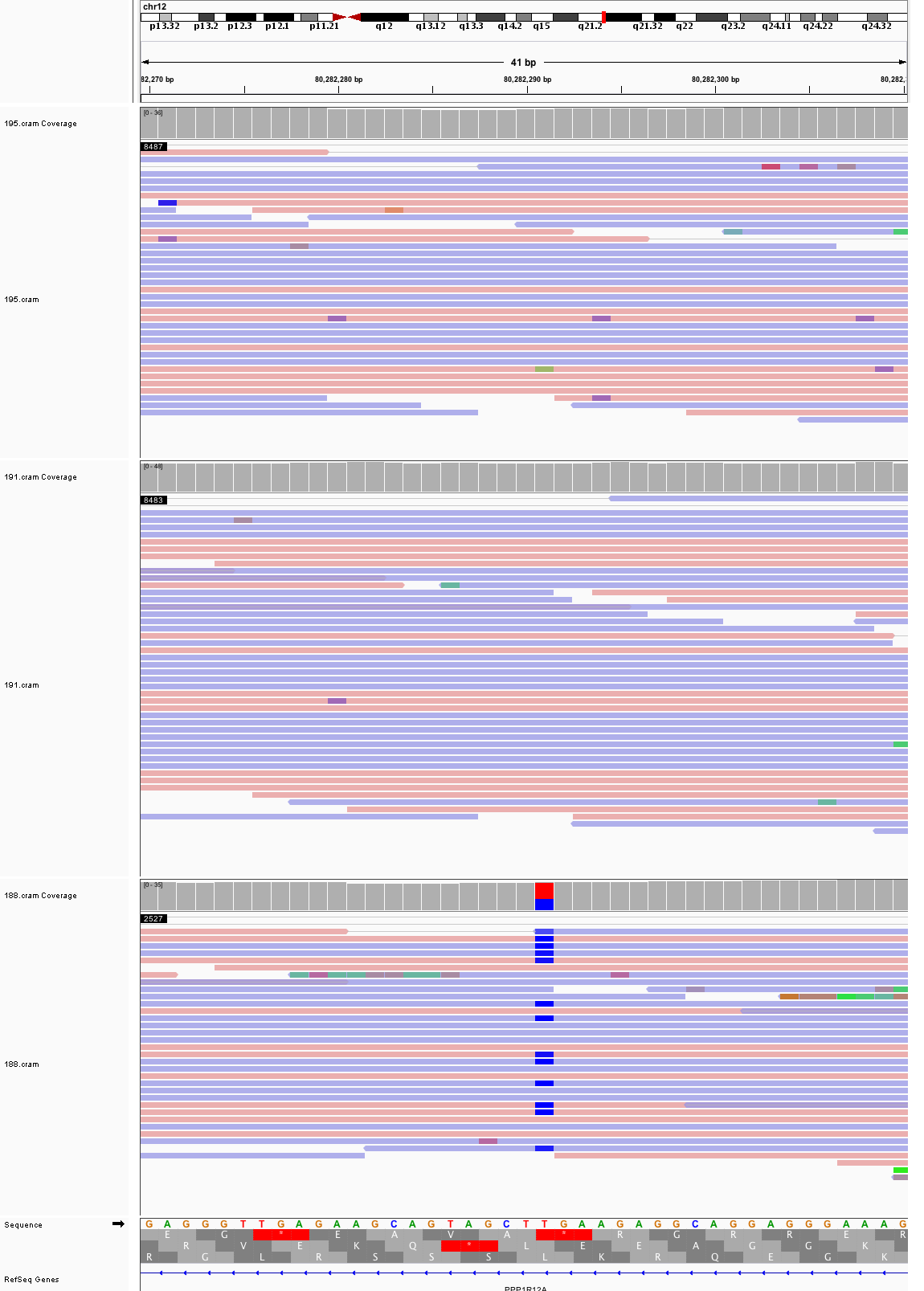

### chr12_106,483,276_106,483,316.png

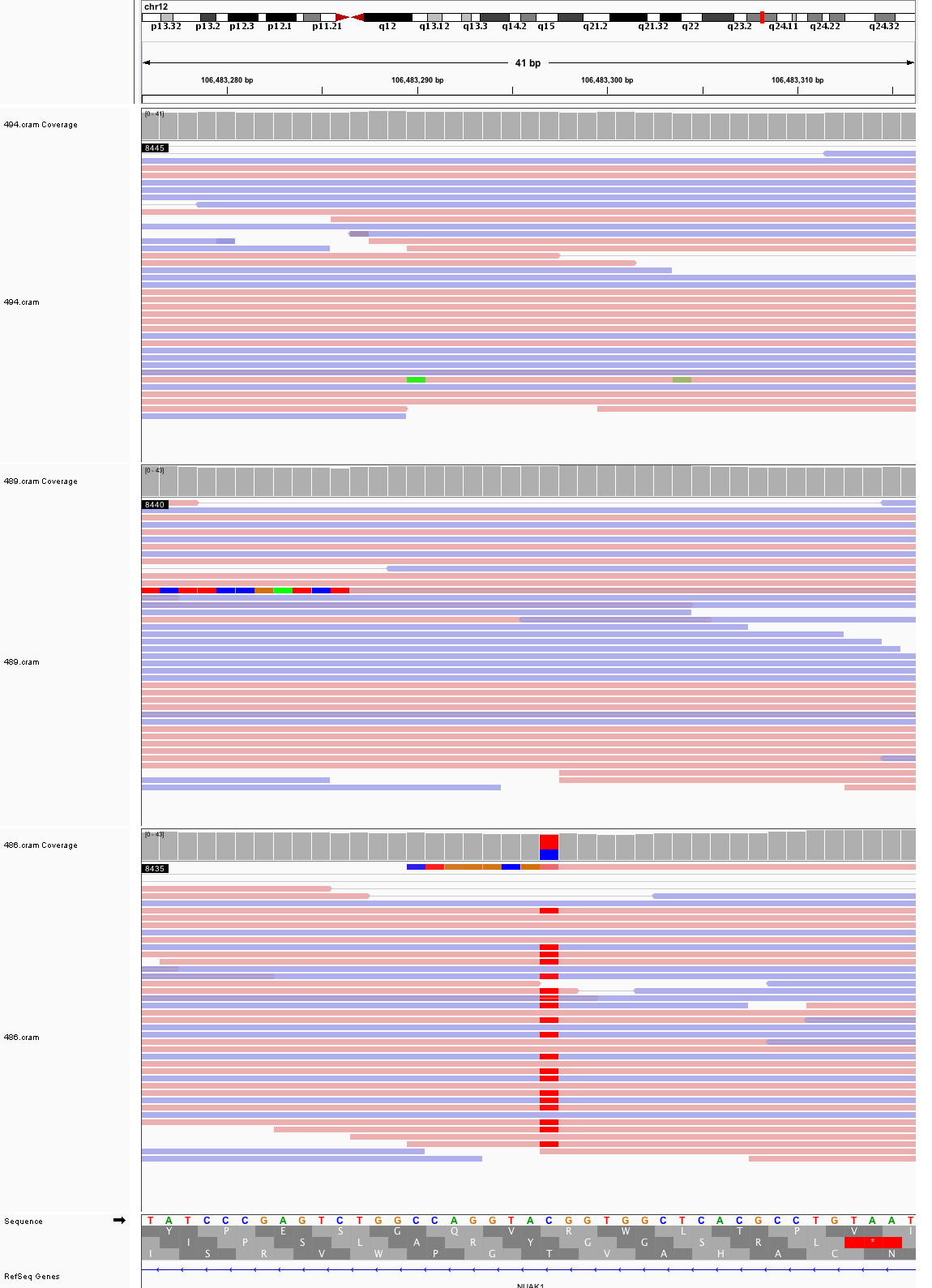

### chr13_86,324,400_86,324,440.png

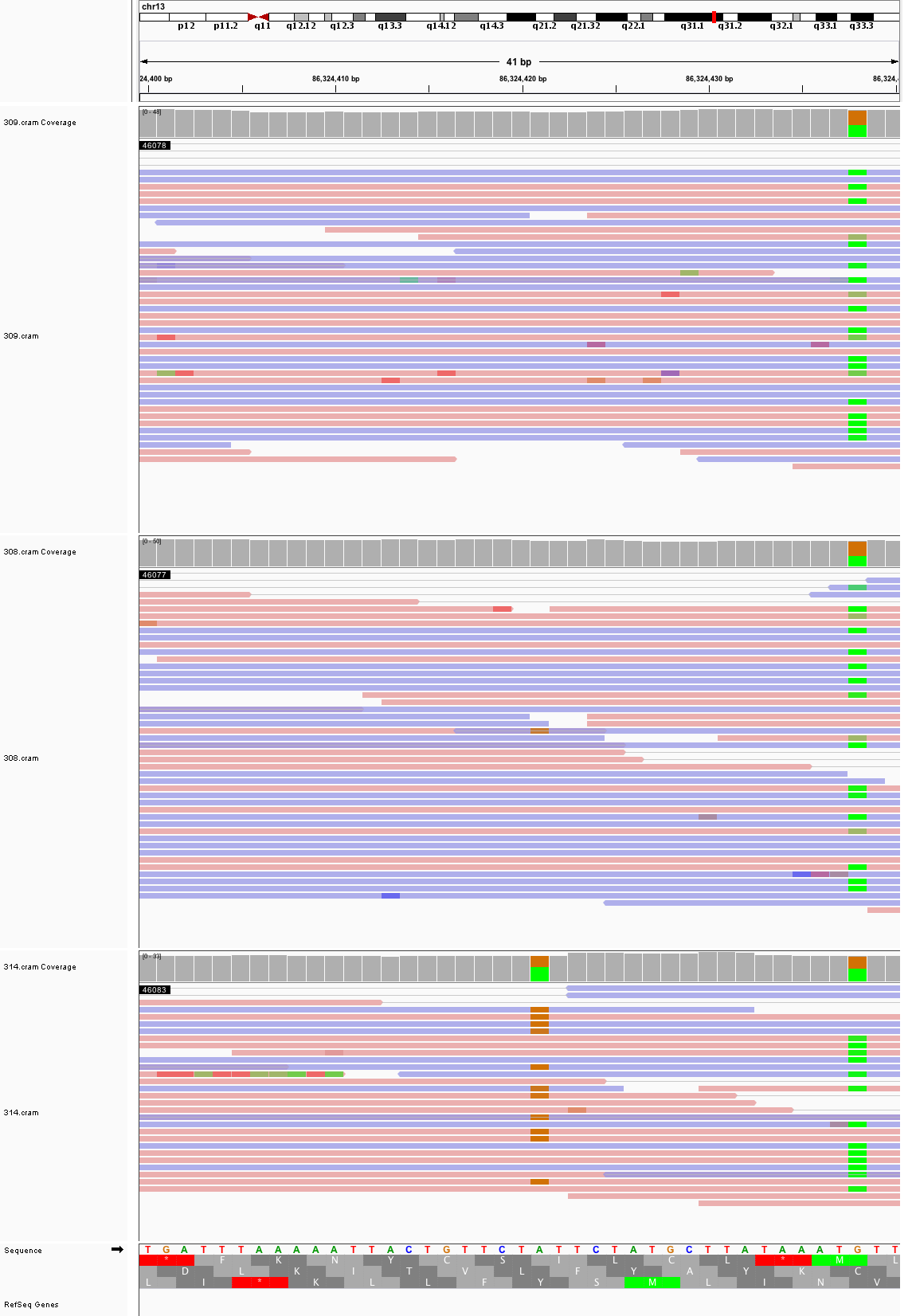

### chr14_89,044,514_89,044,554.png

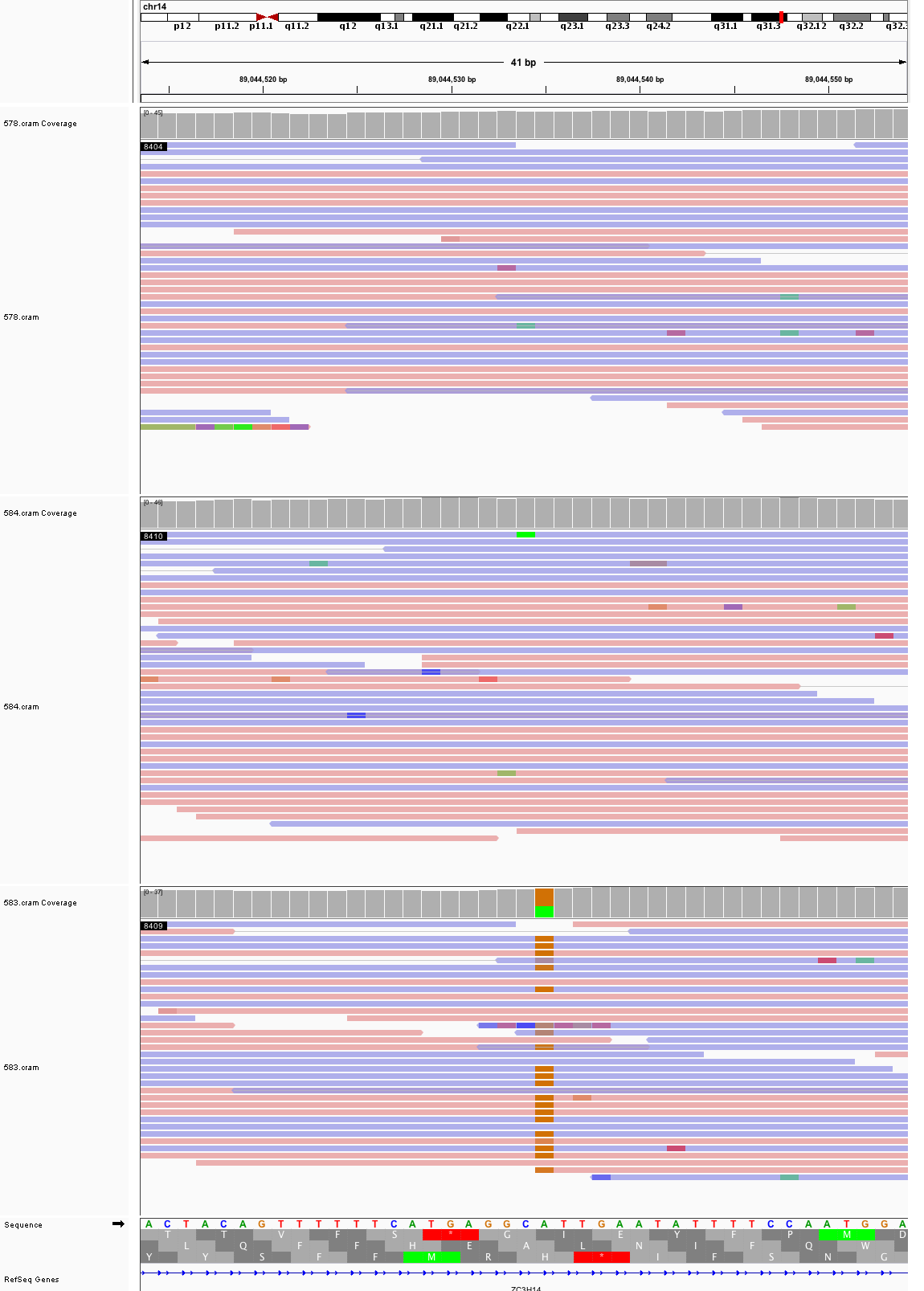

### chr14_97,101,302_97,101,342.png

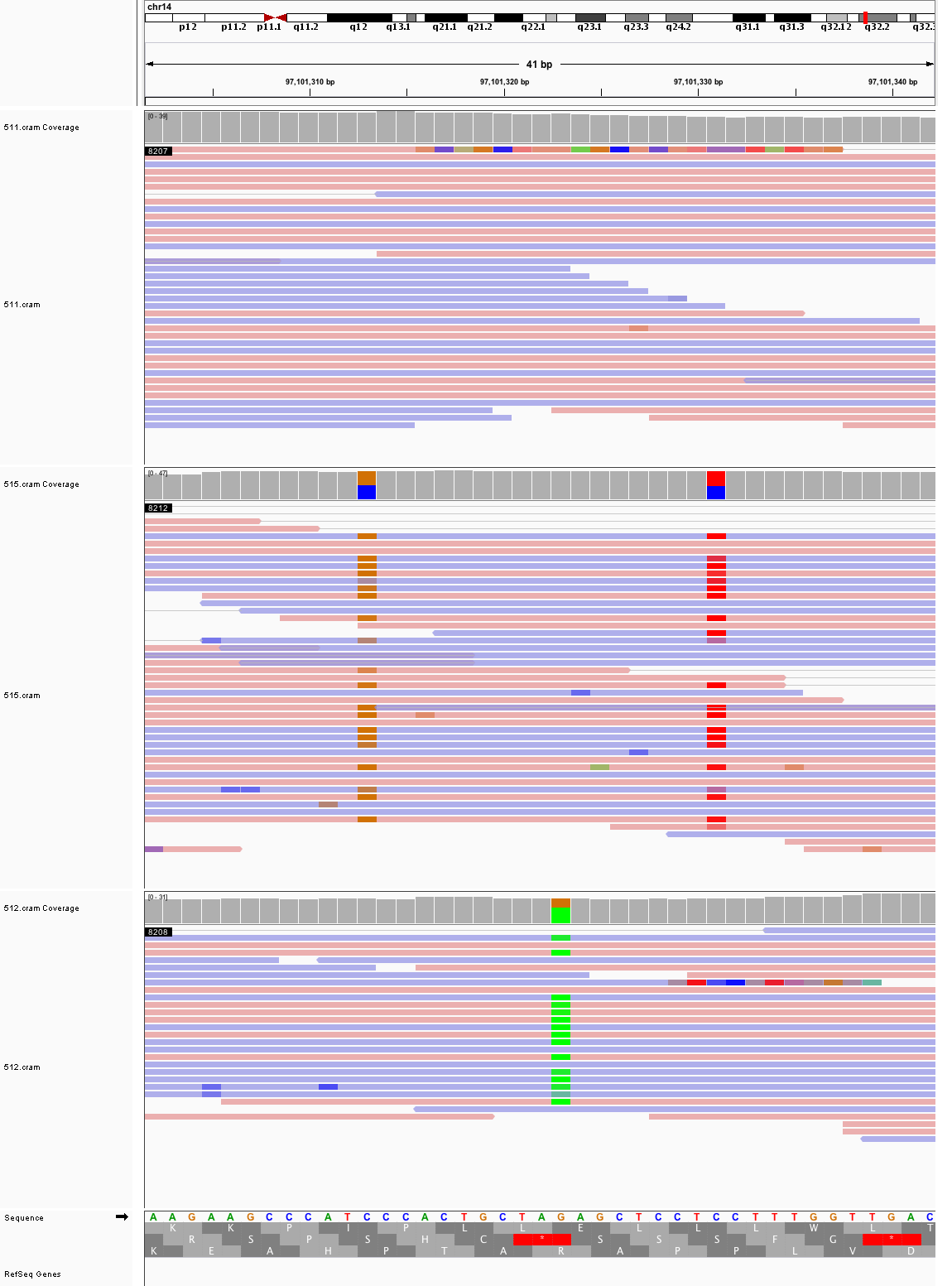

### chr15_40,280,276_40,280,316.png

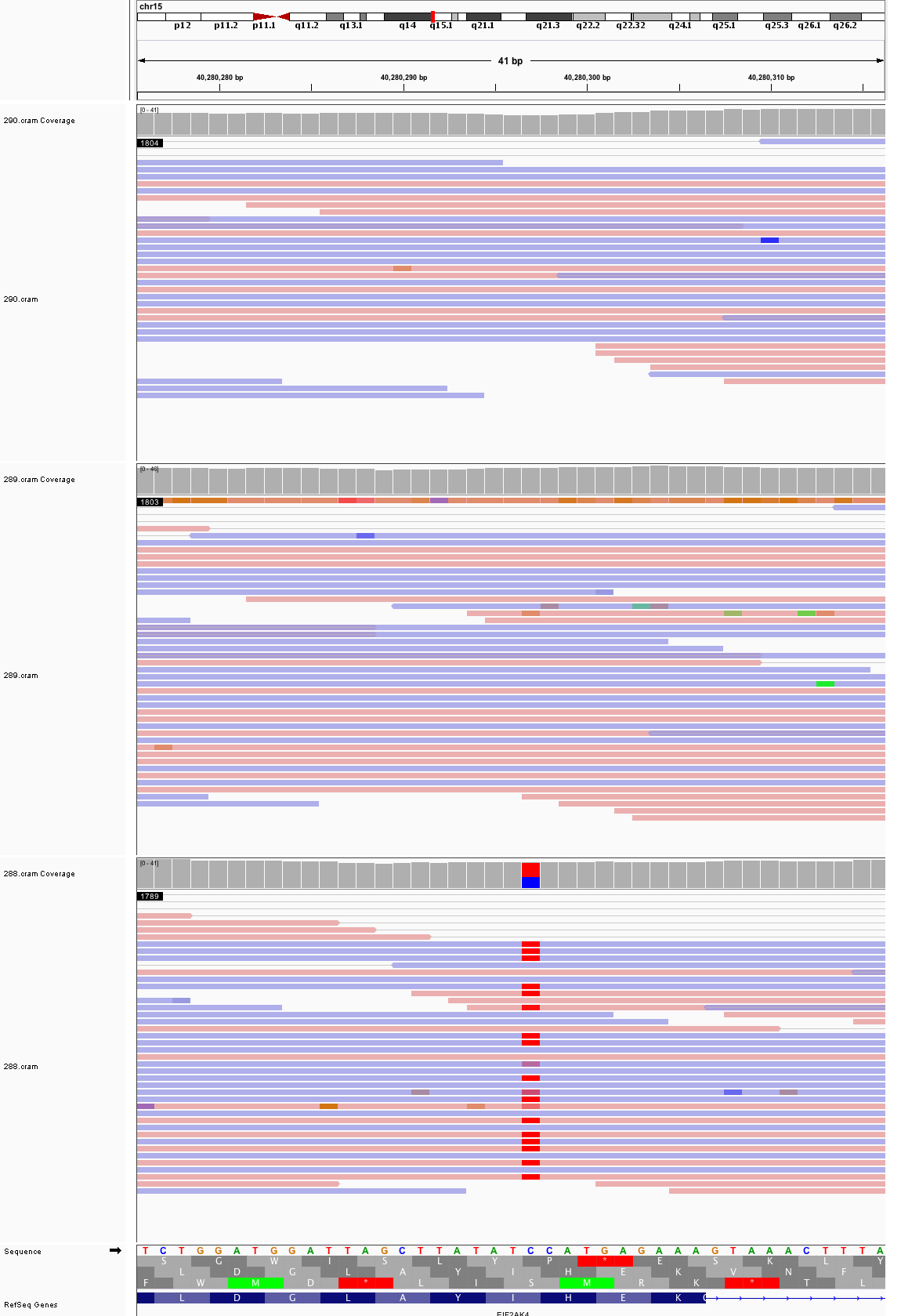

### chr15_49,214,483_49,214,523.png

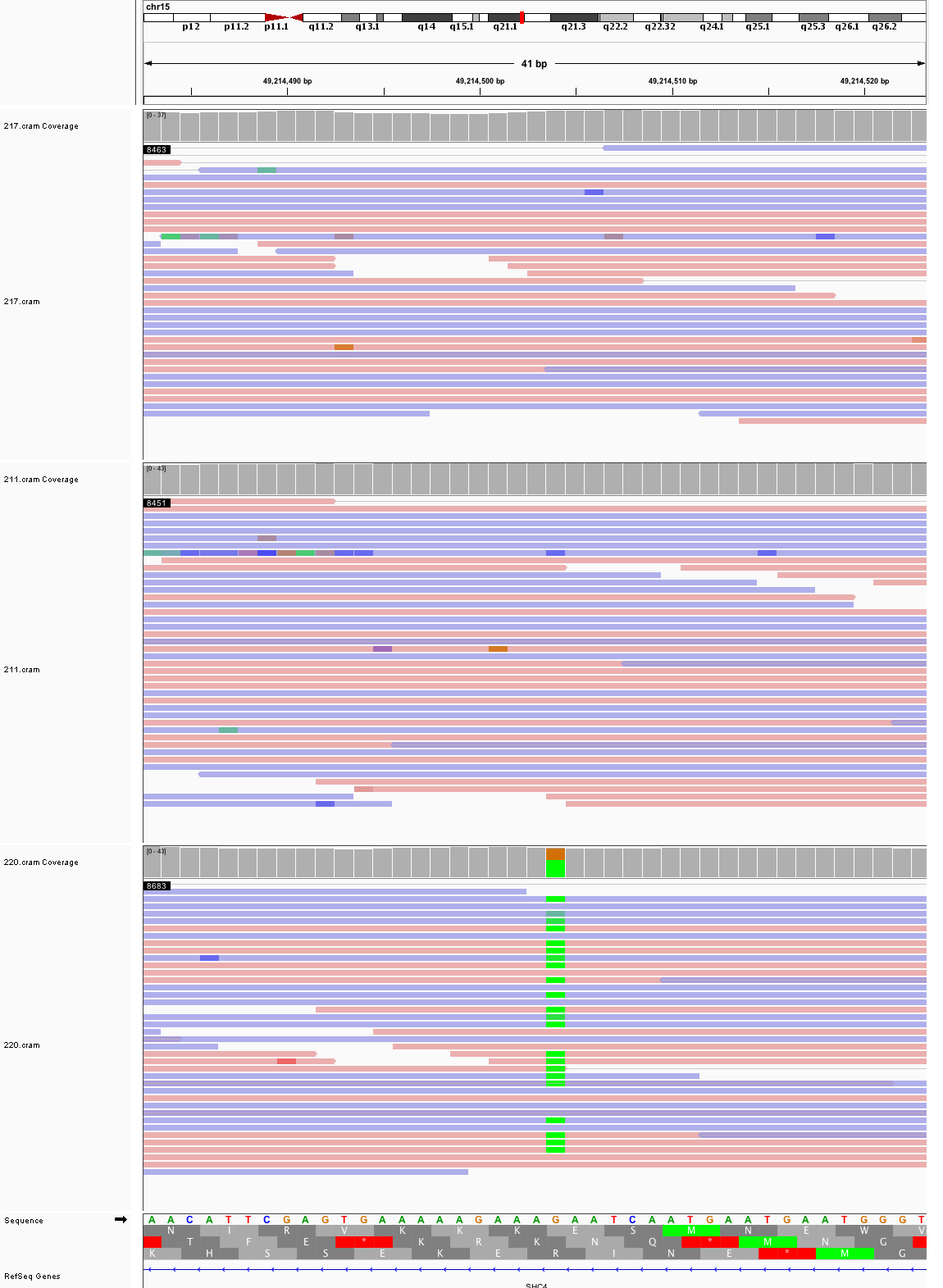

### chr15_51,656,451_51,656,491.png

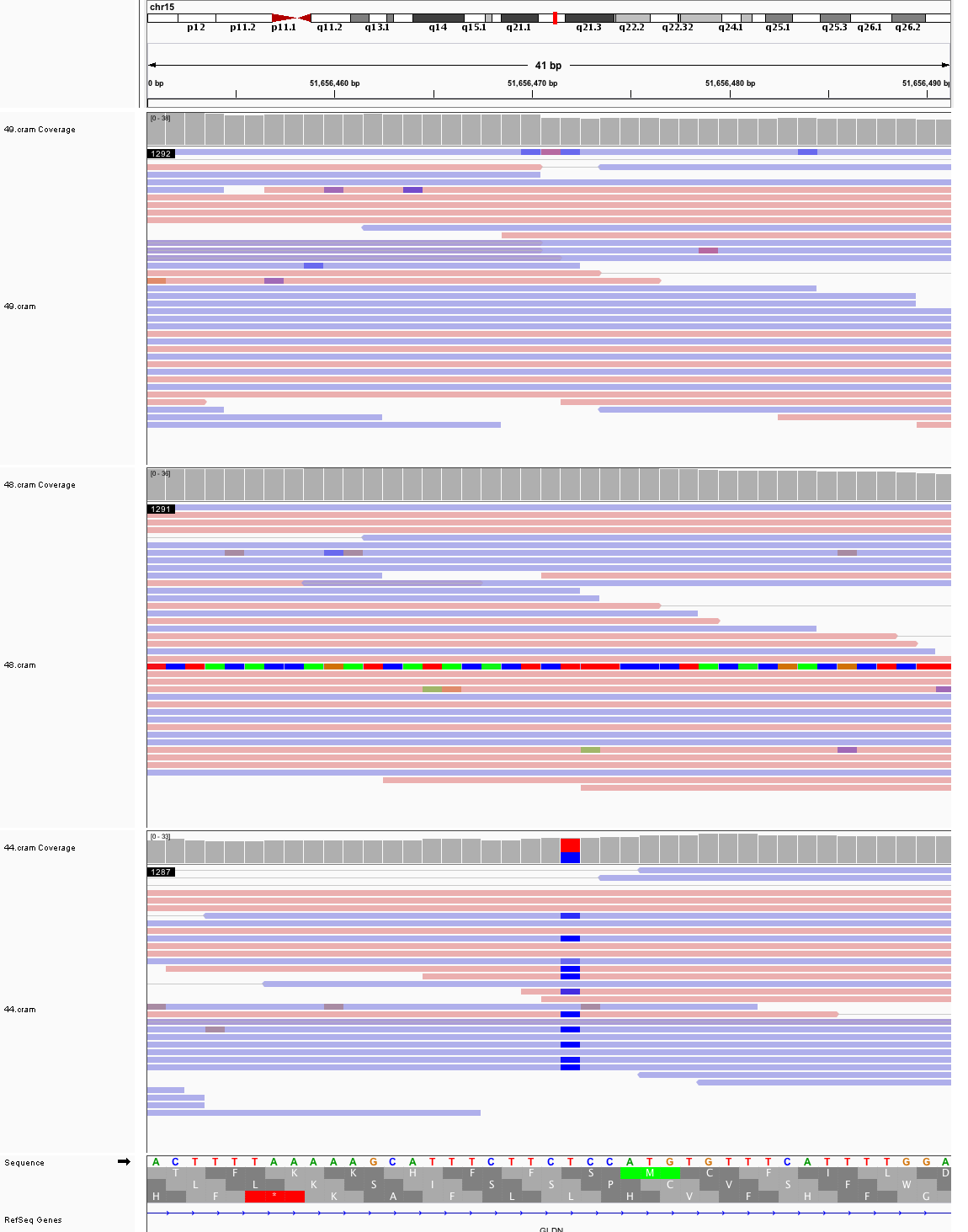

### chr15_79,603,123_79,603,163.png

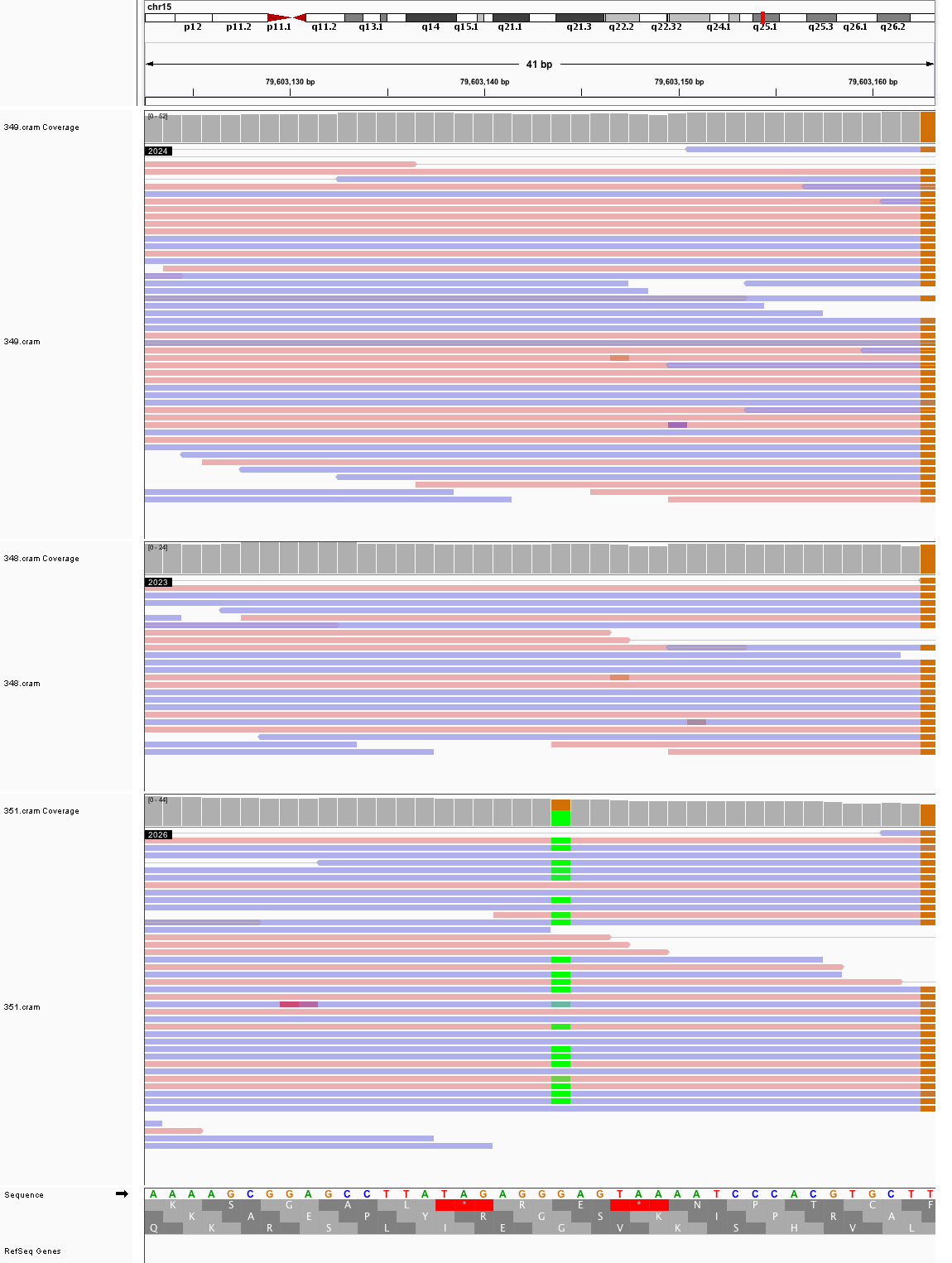

### chr15_94,291,170_94,291,210.png

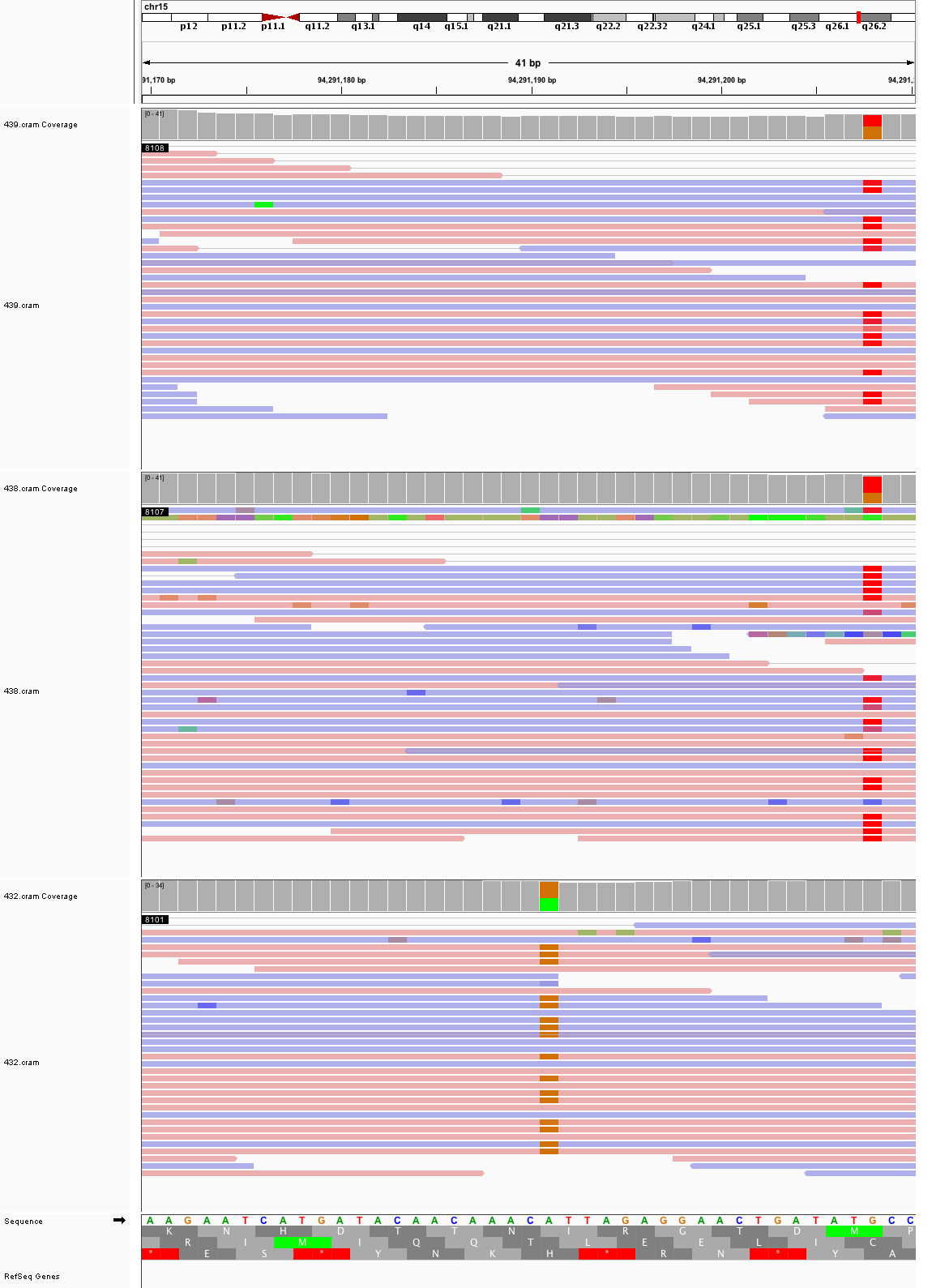

### chr16_55,899,030_55,899,070.png

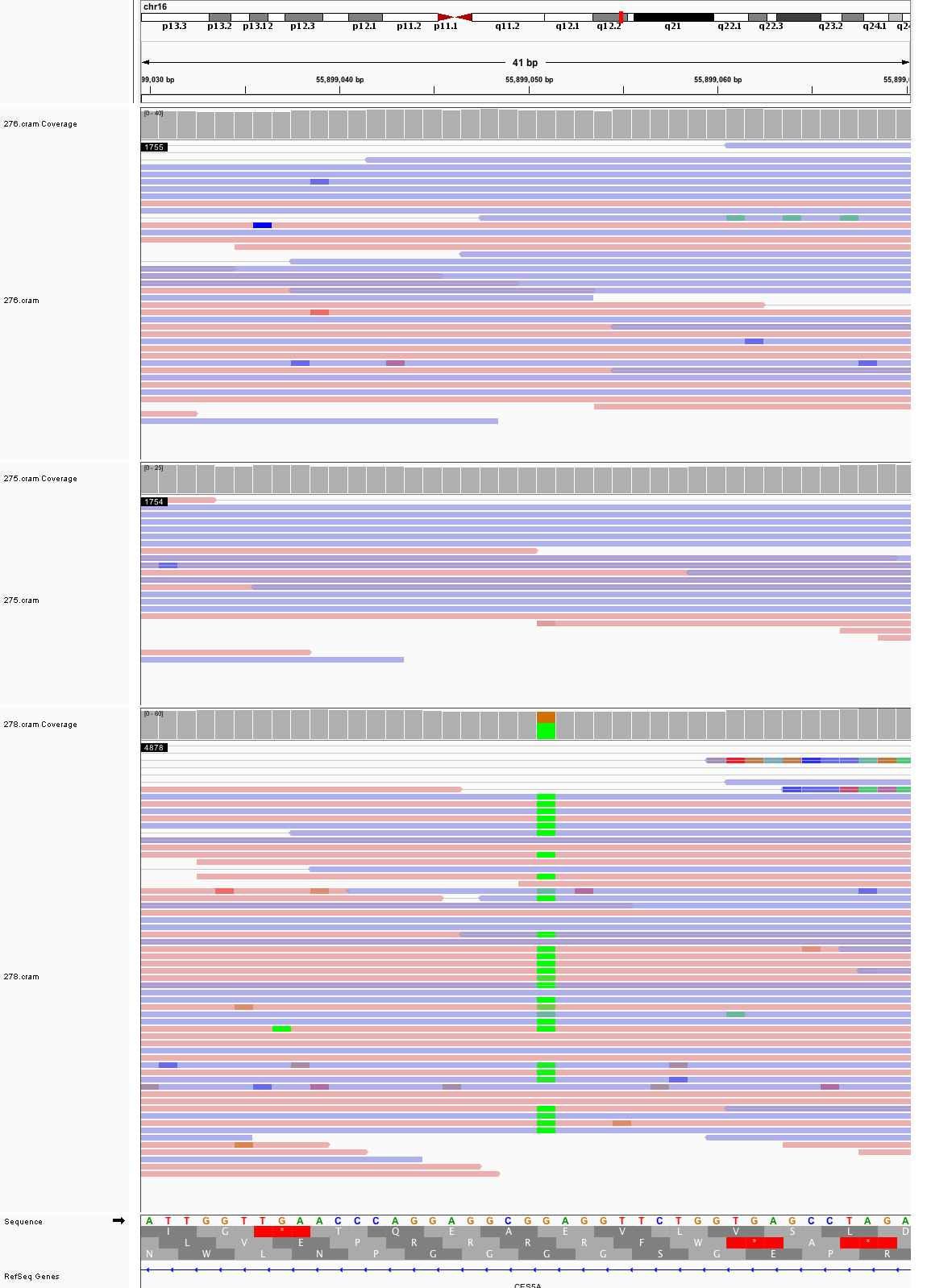

### chr16_88,947,085_88,947,125.png

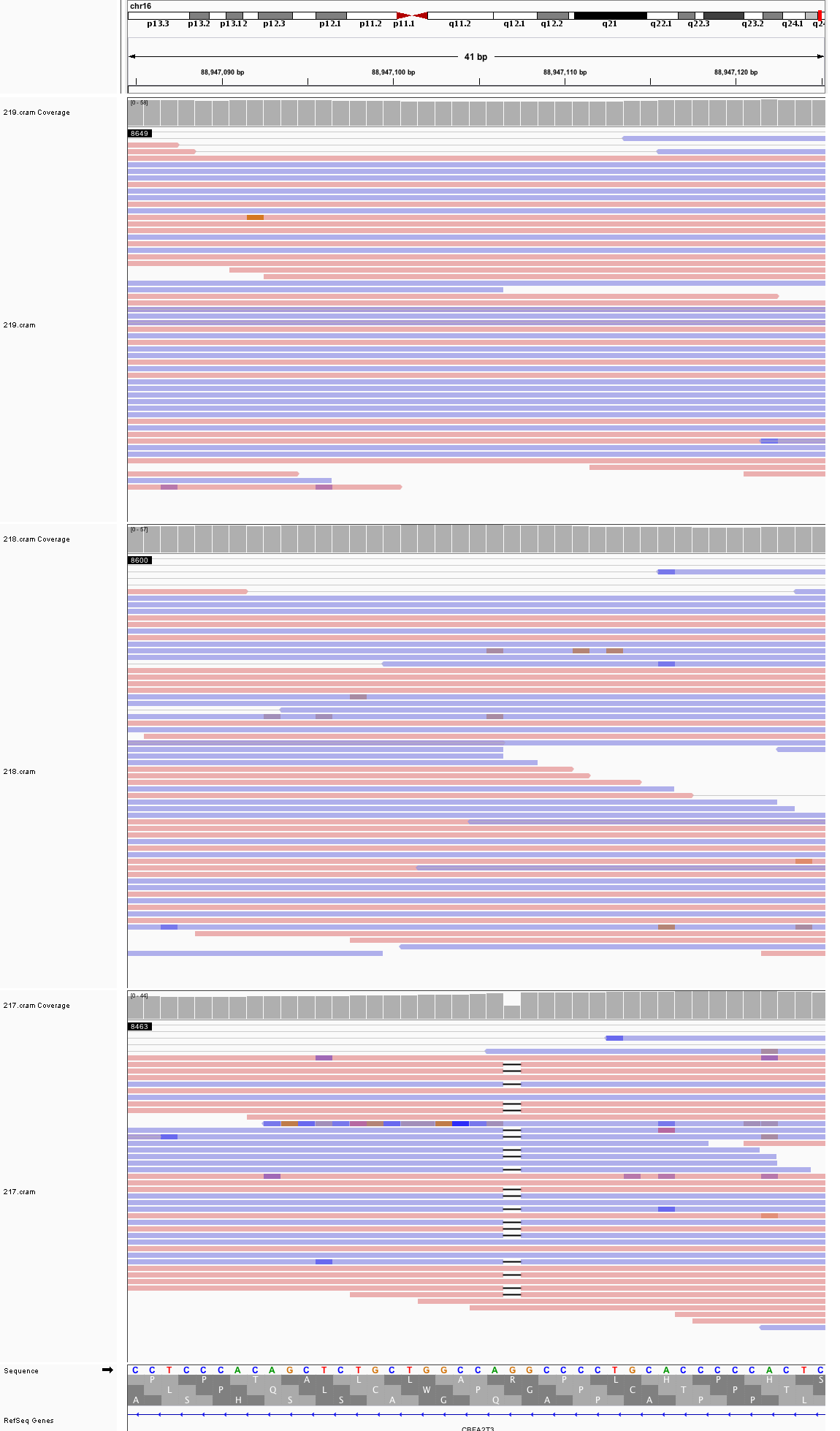

### chr17_7,373,449_7,373,489.png

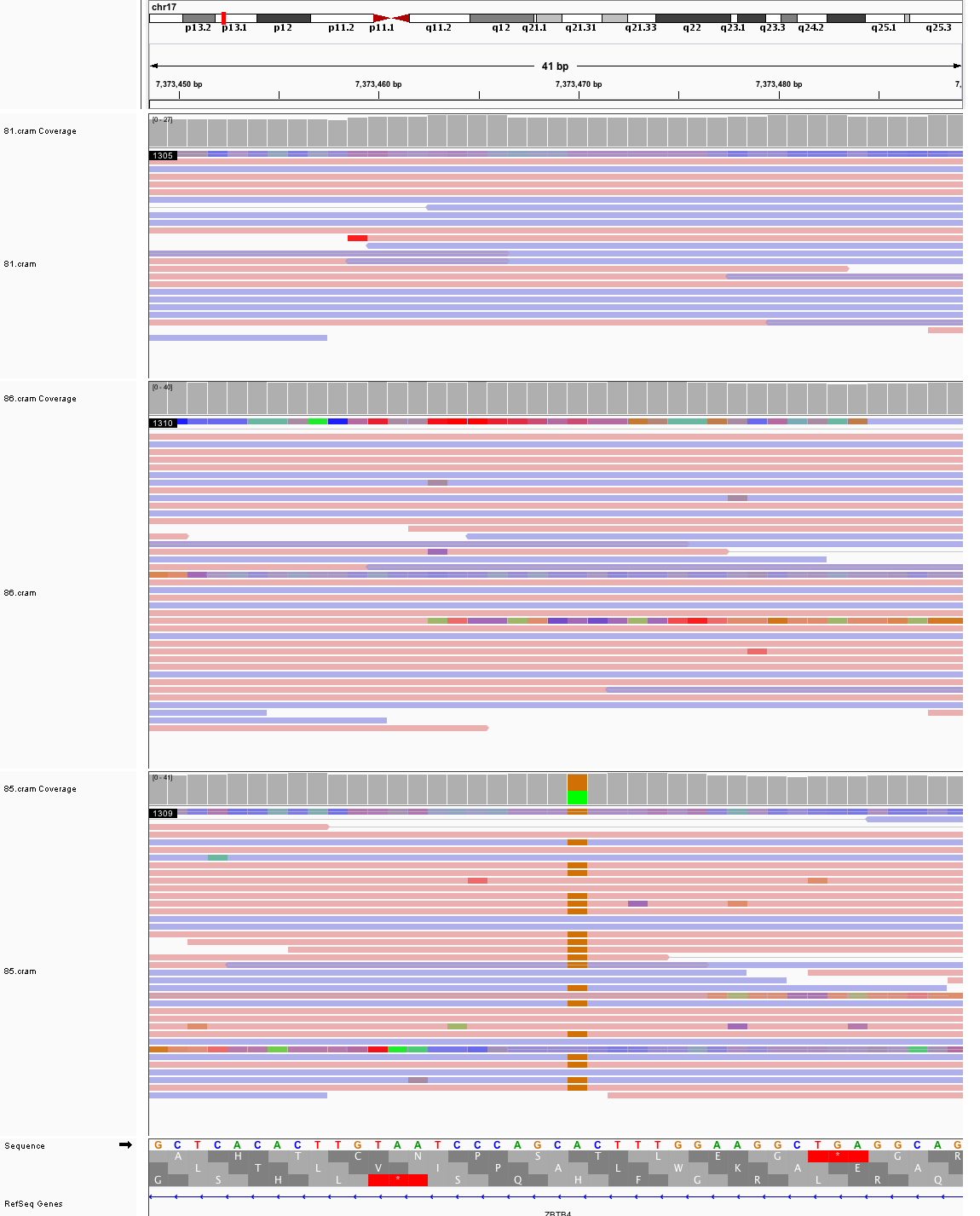

### chr17_14,703,883_14,703,923.png

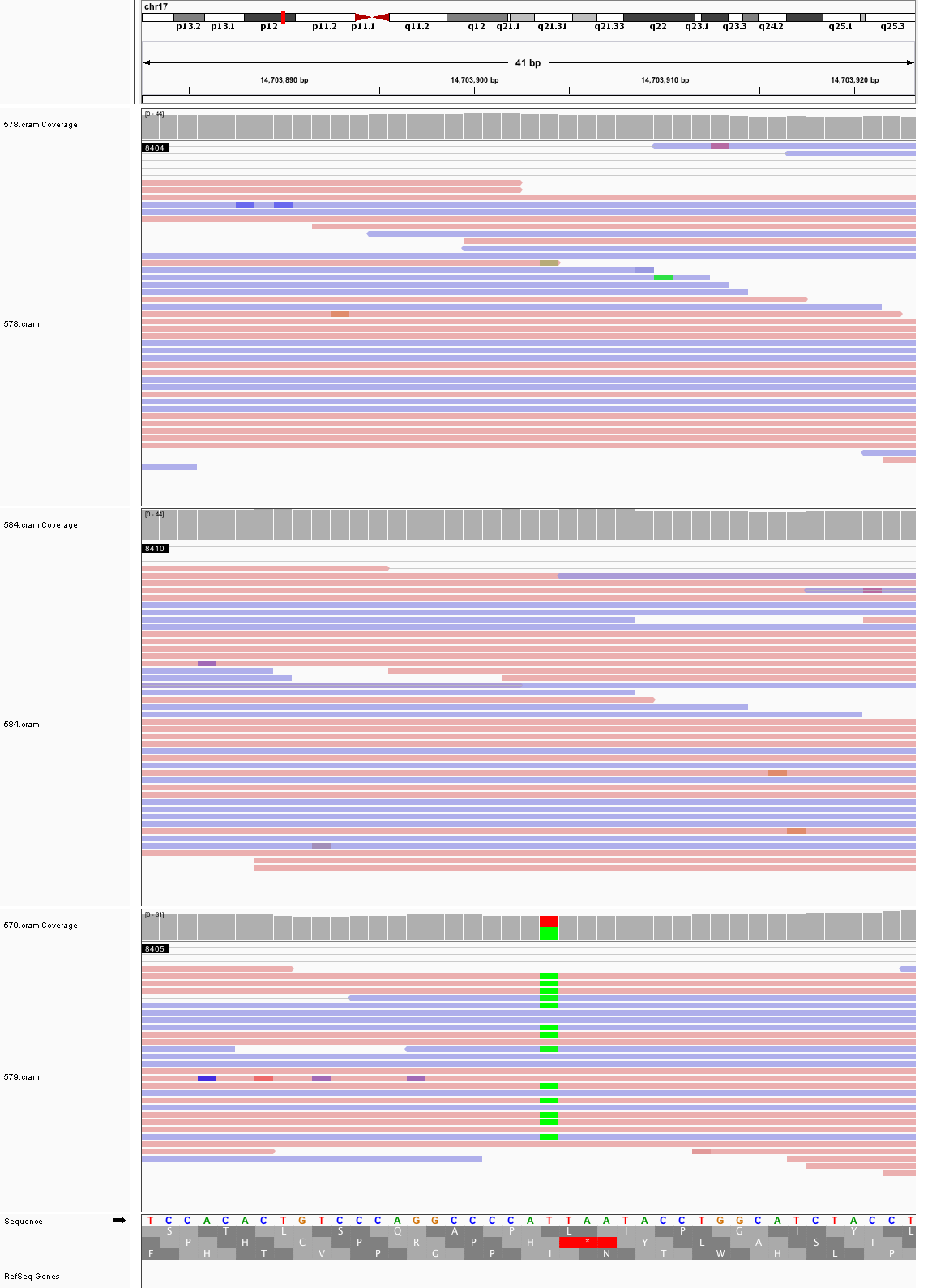

### chr17_18,881,666_18,881,706.png

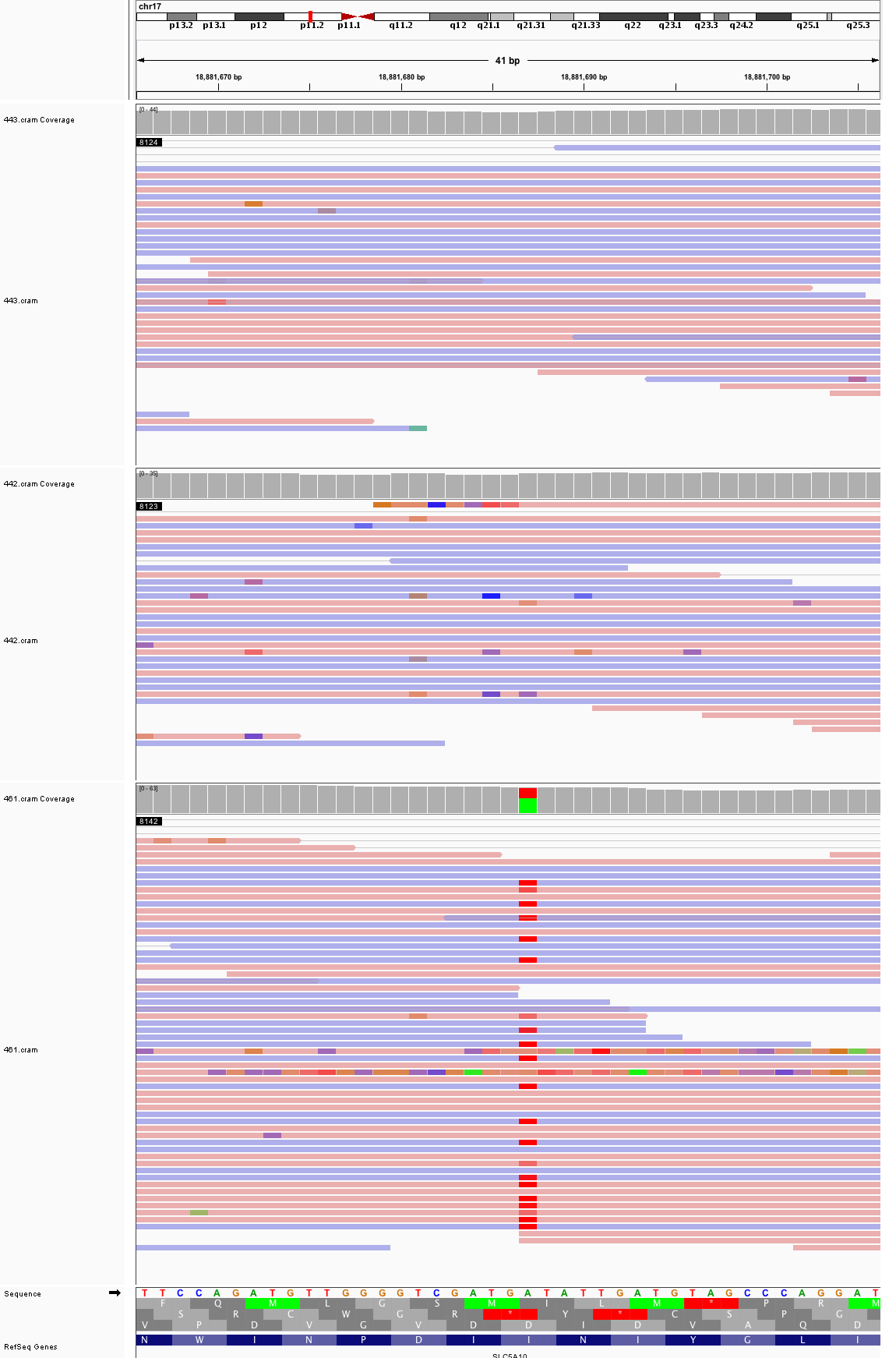

### chr17_79,232,876_79,232,916.png

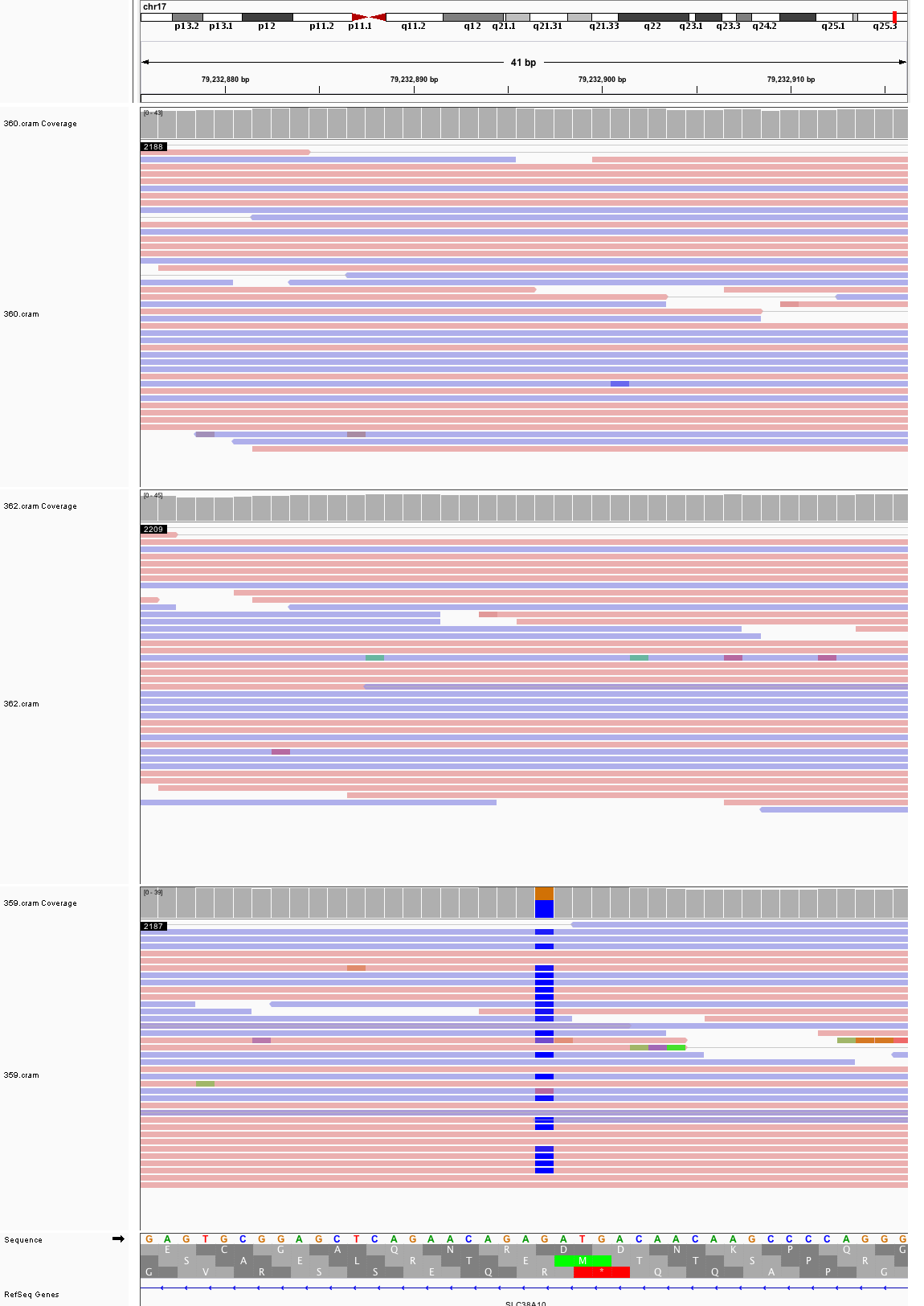

### chr18_22,276,199_22,276,239.png

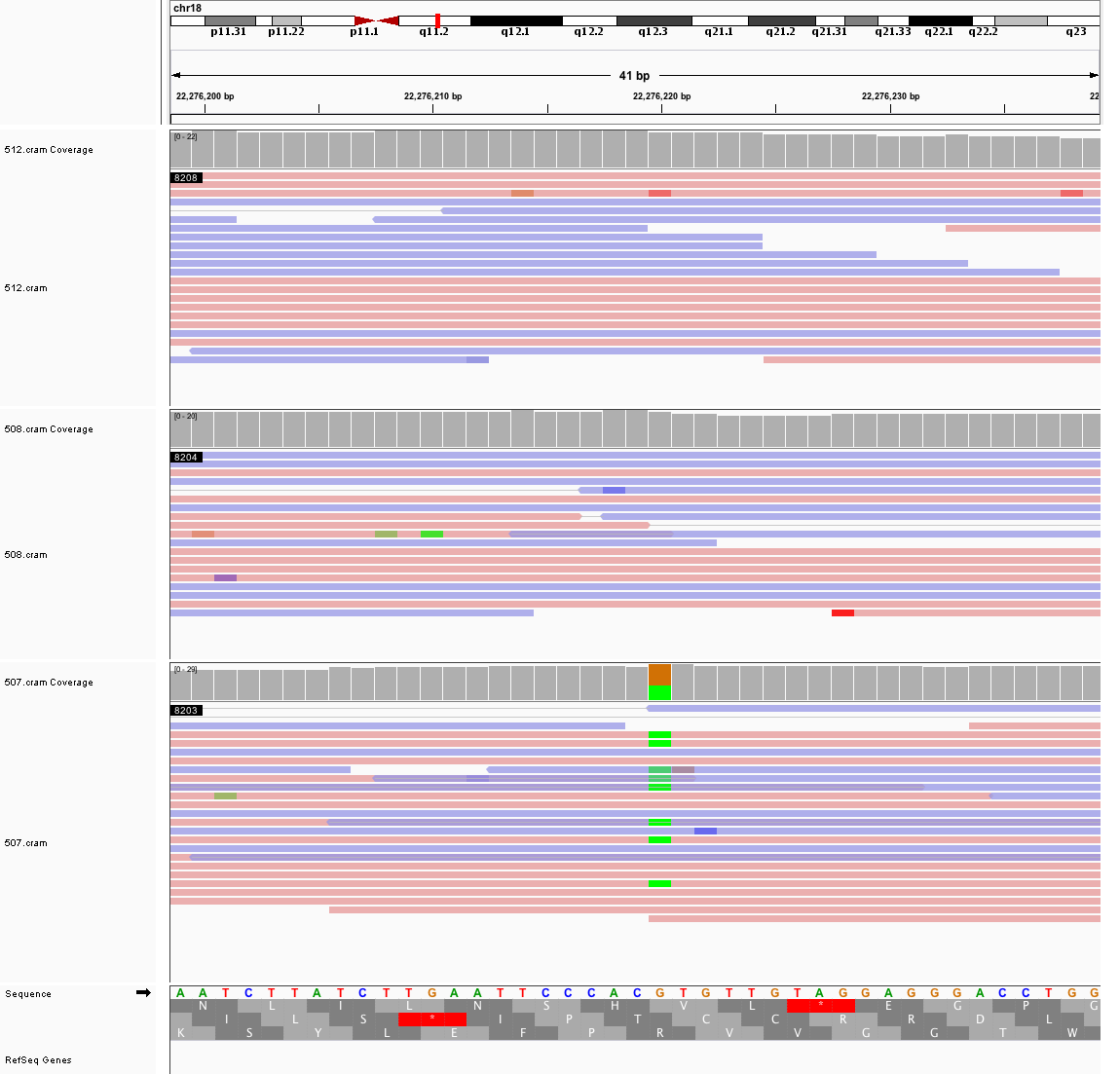

### chr18_38,901,672_38,901,712.png

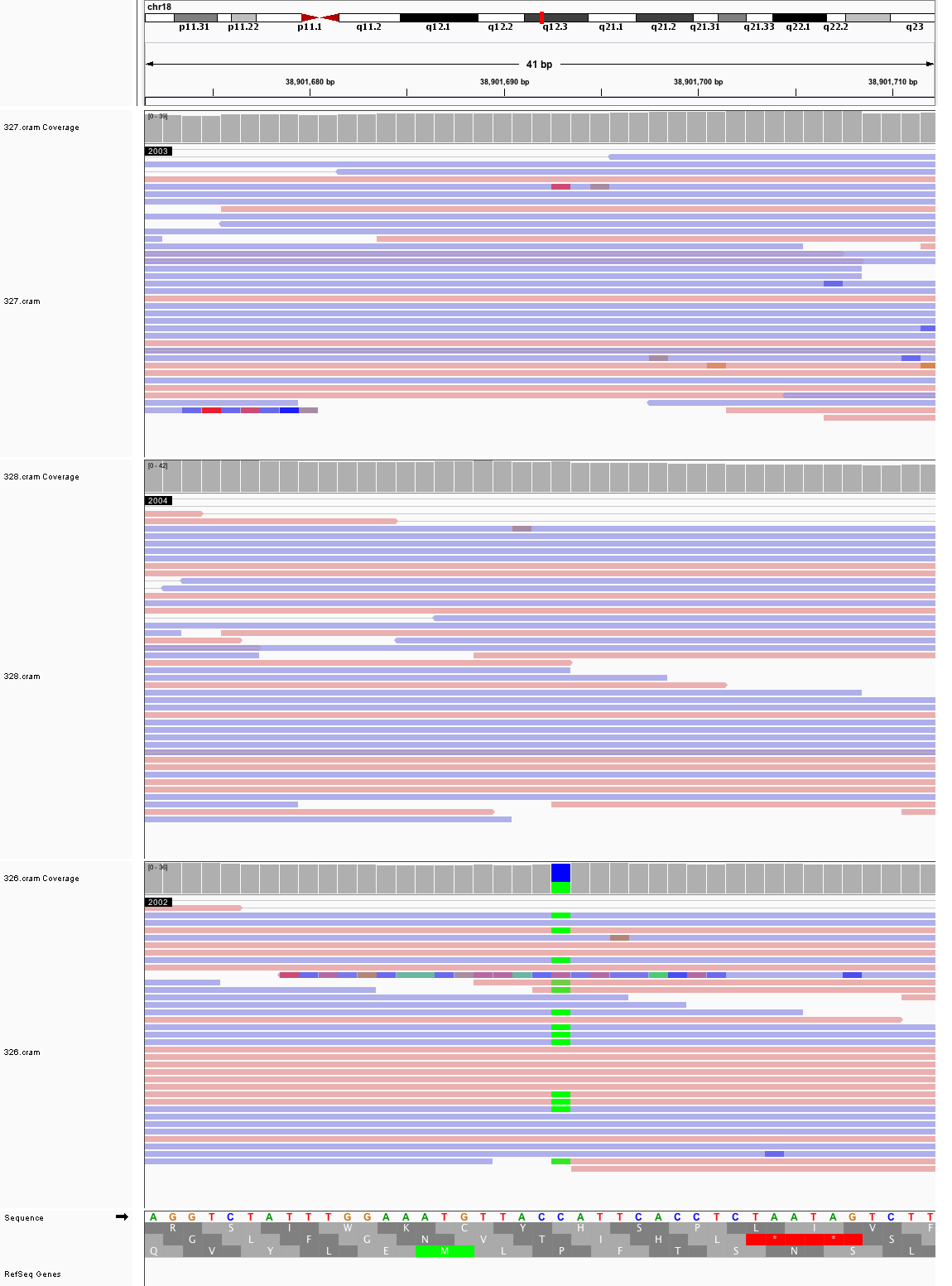

### chr18_46,776,822_46,776,862.png

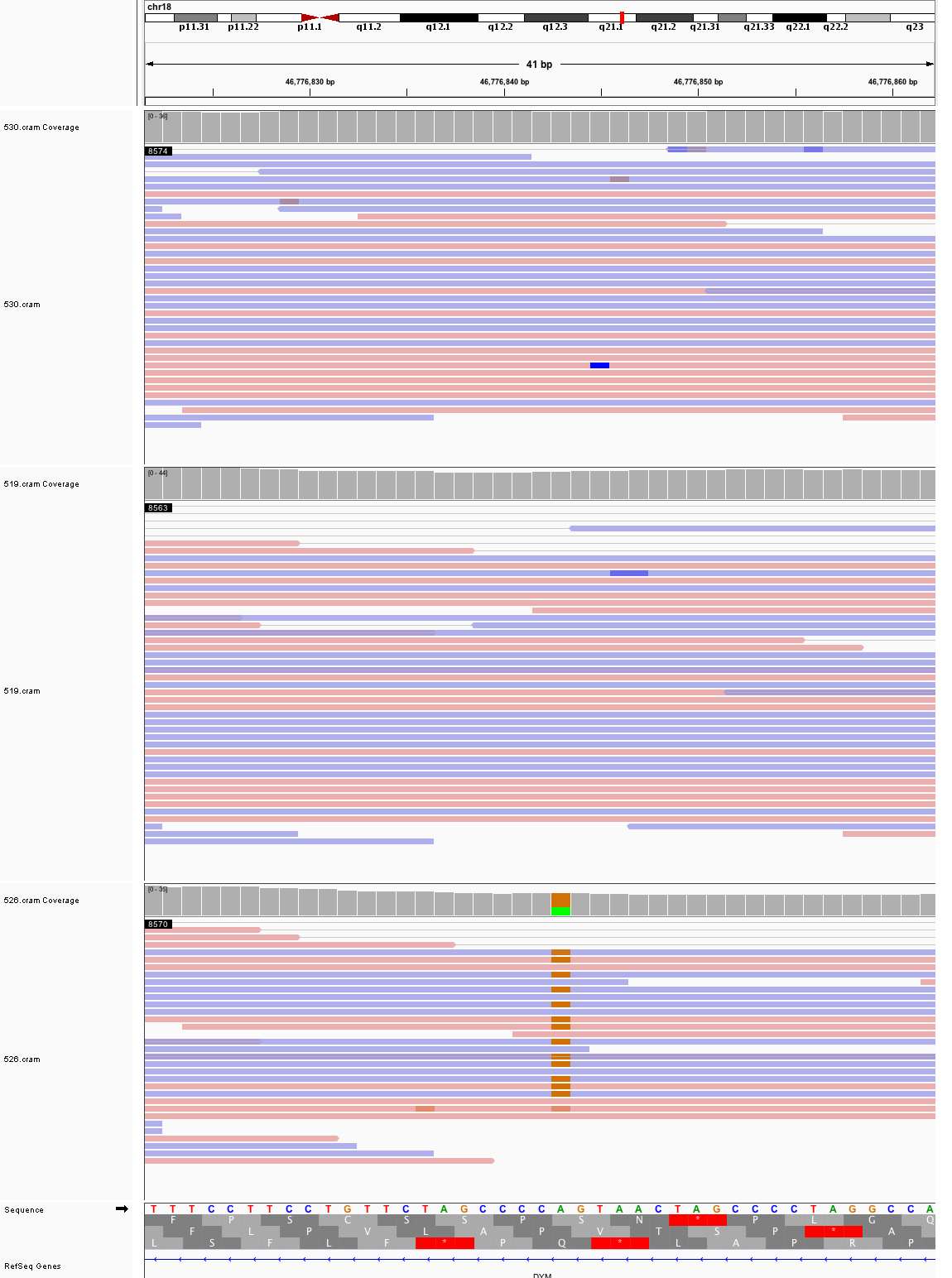

### chr20_7,289,206_7,289,246.png

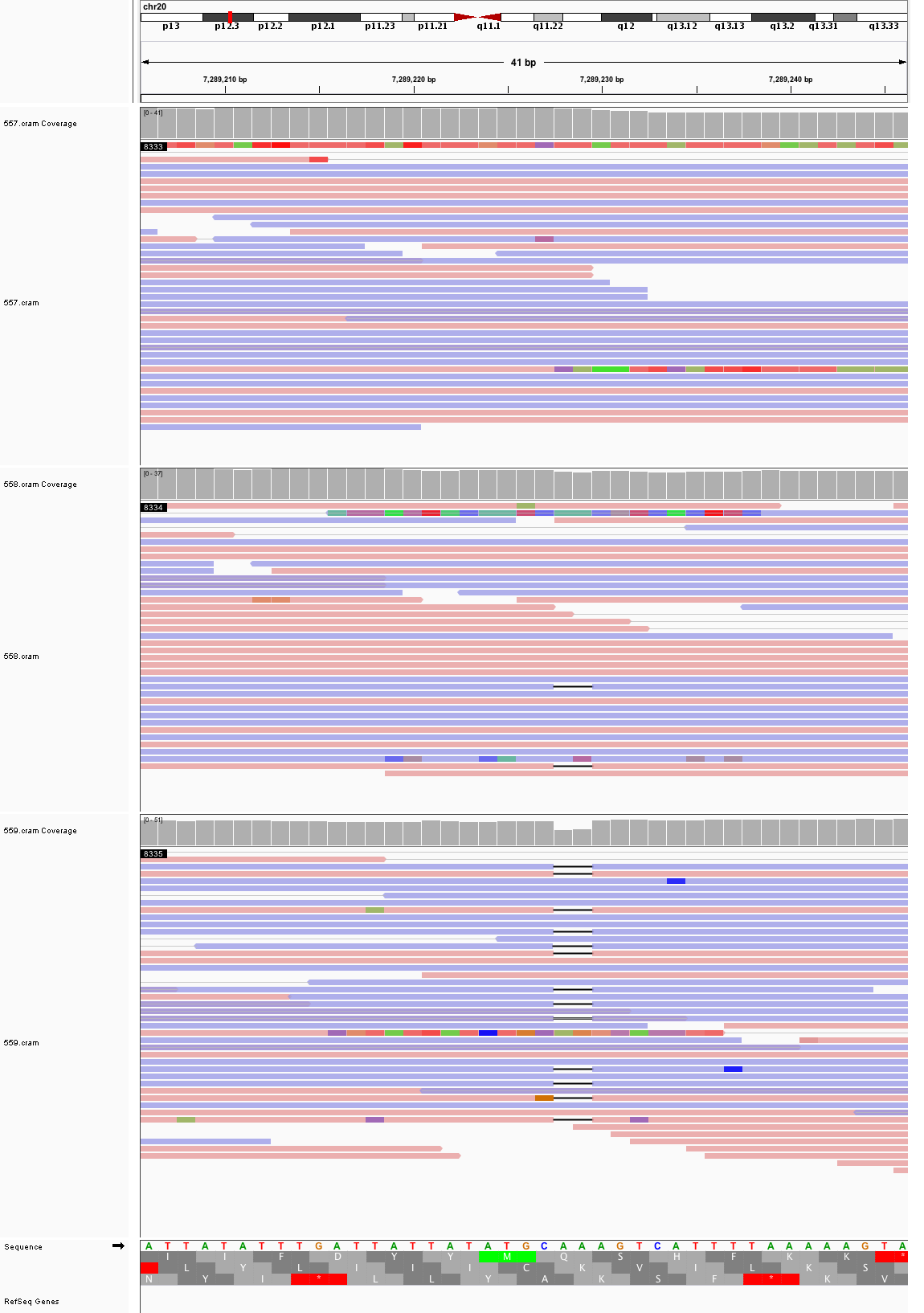

### chr20_17,110,435_17,110,475.png

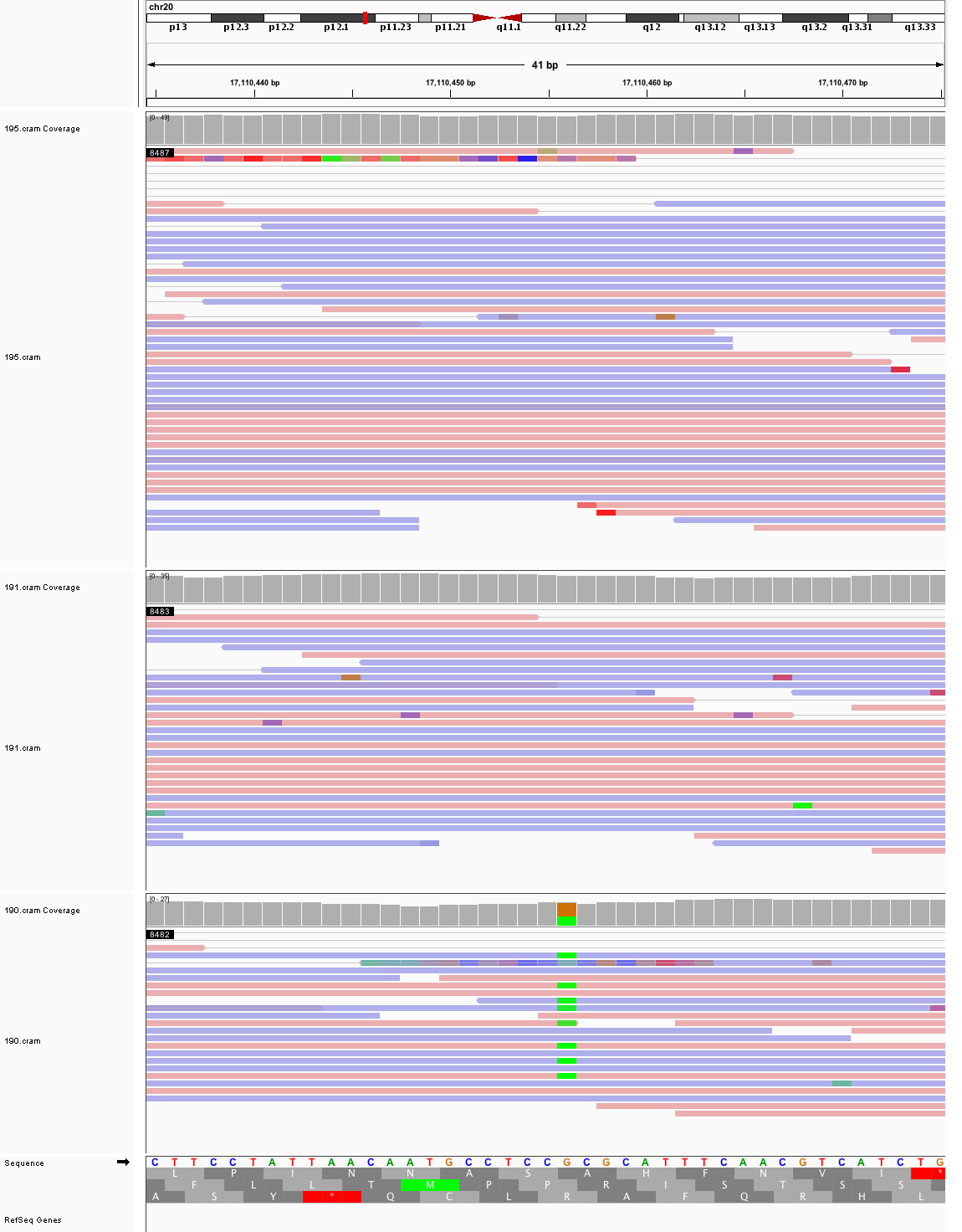

### chr20_31,646,936_31,646,976.png

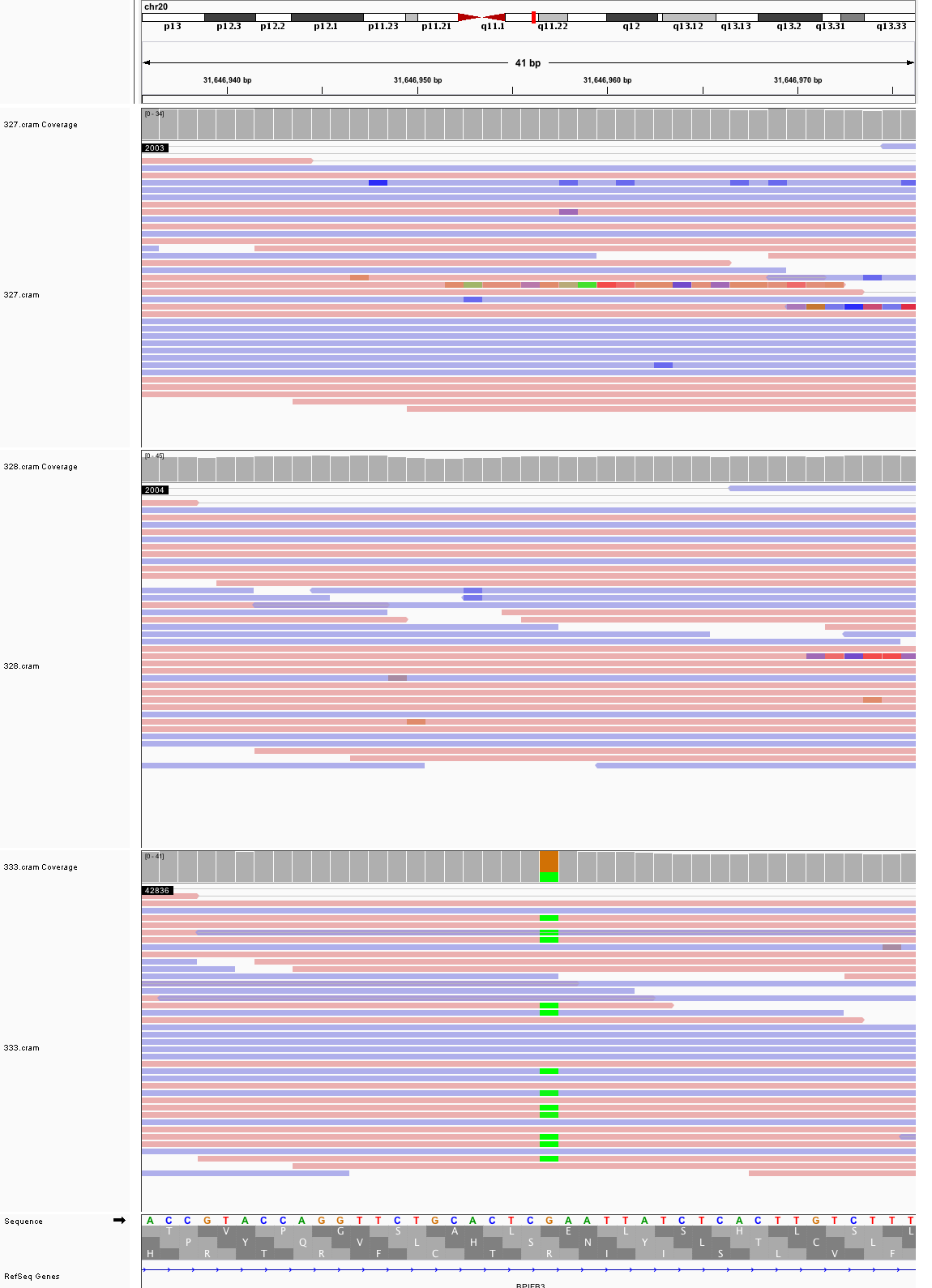

### chr21_20,165,333_20,165,373.png

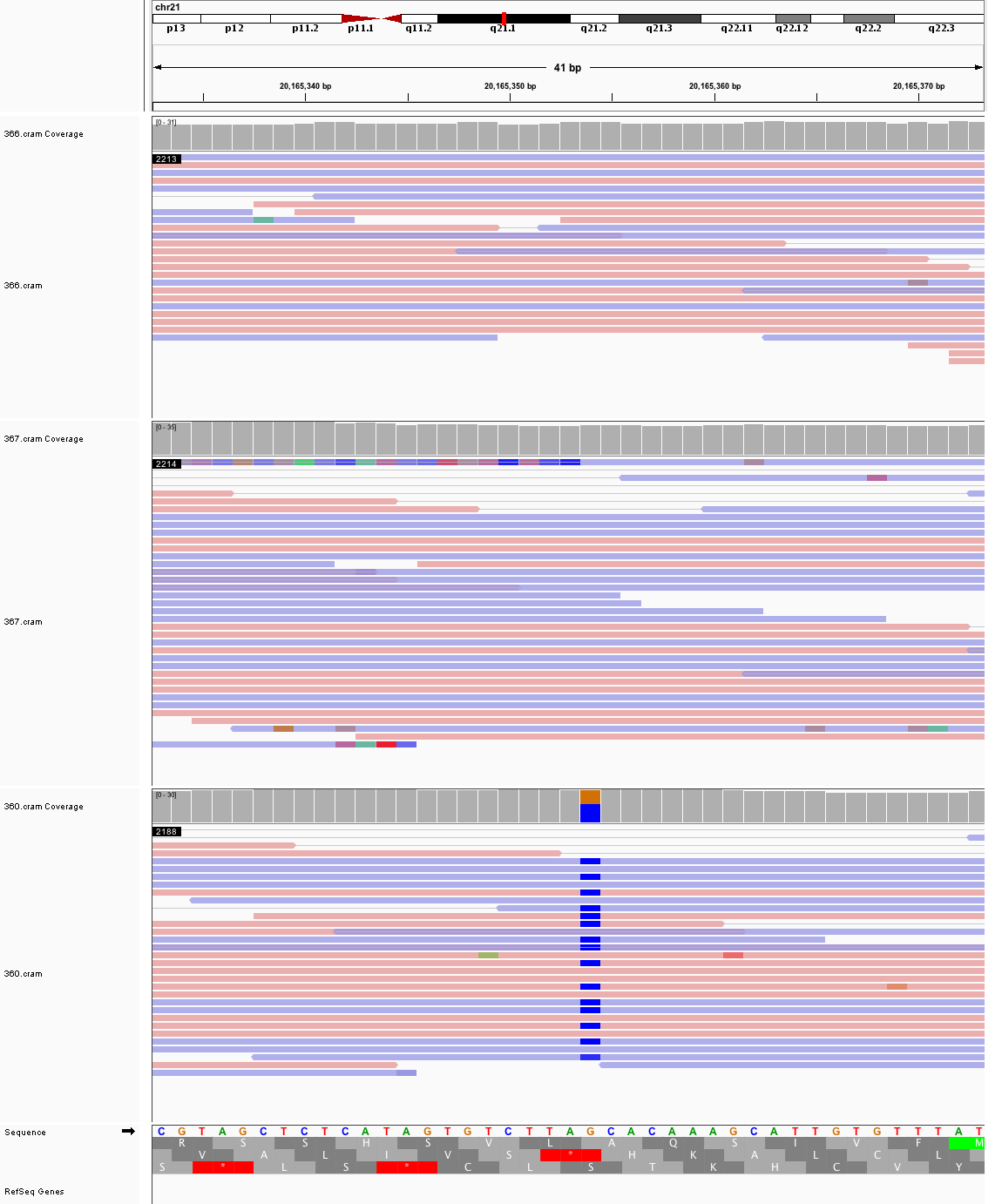

### chr21_20,848,492_20,848,532.png

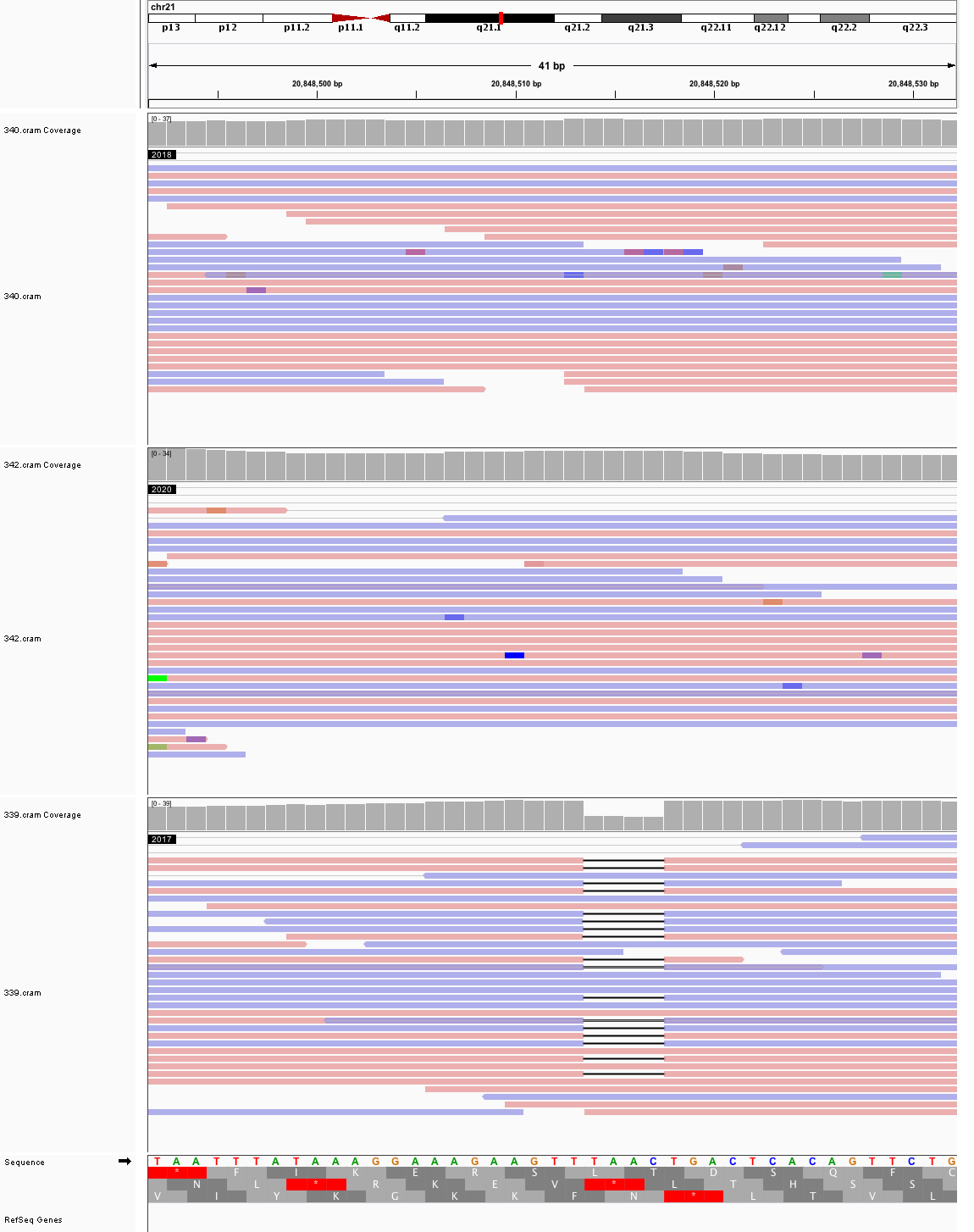

### chr21_26,065,548_26,065,588.png

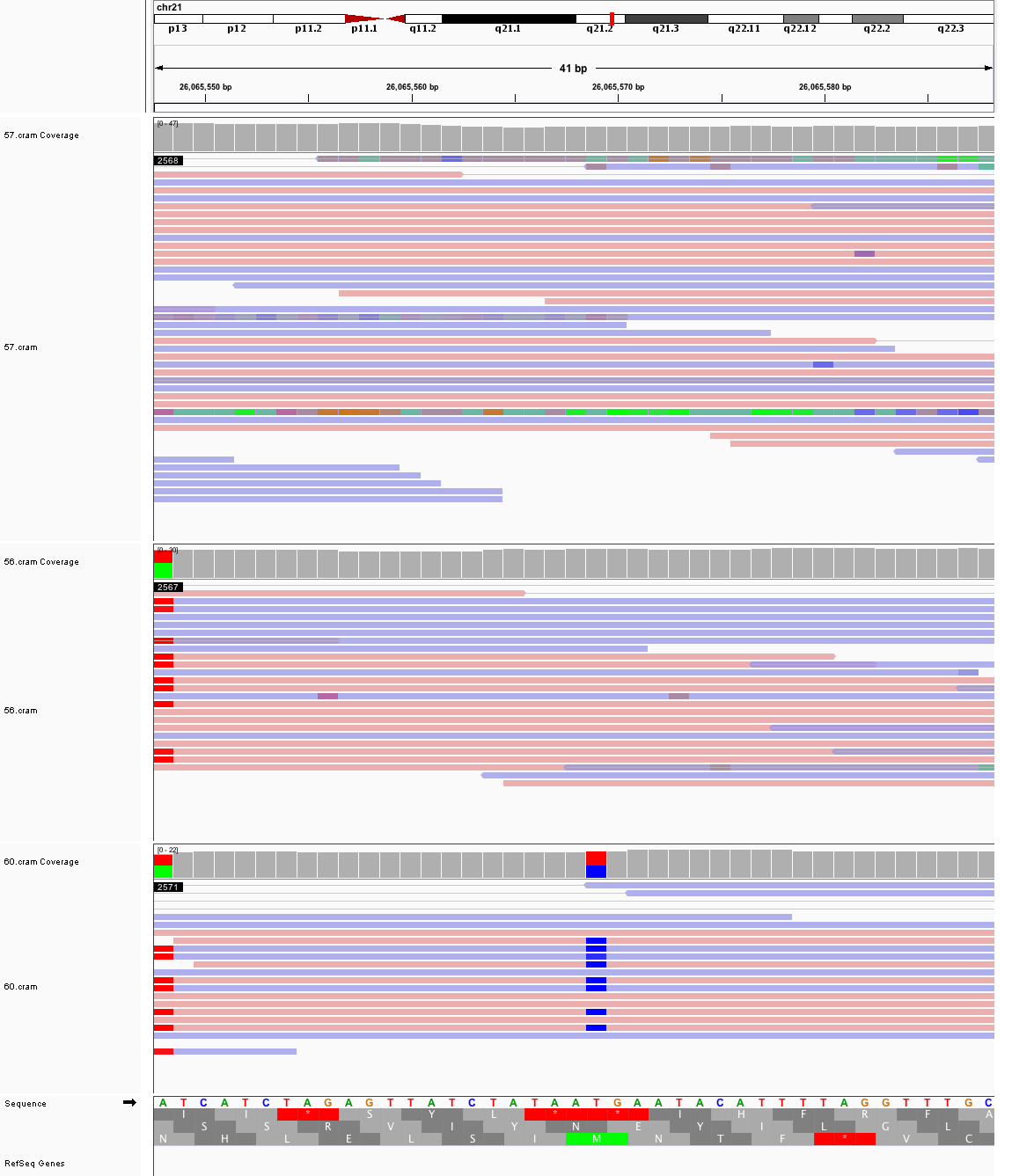

### chrX_117,489,209_117,489,249.png

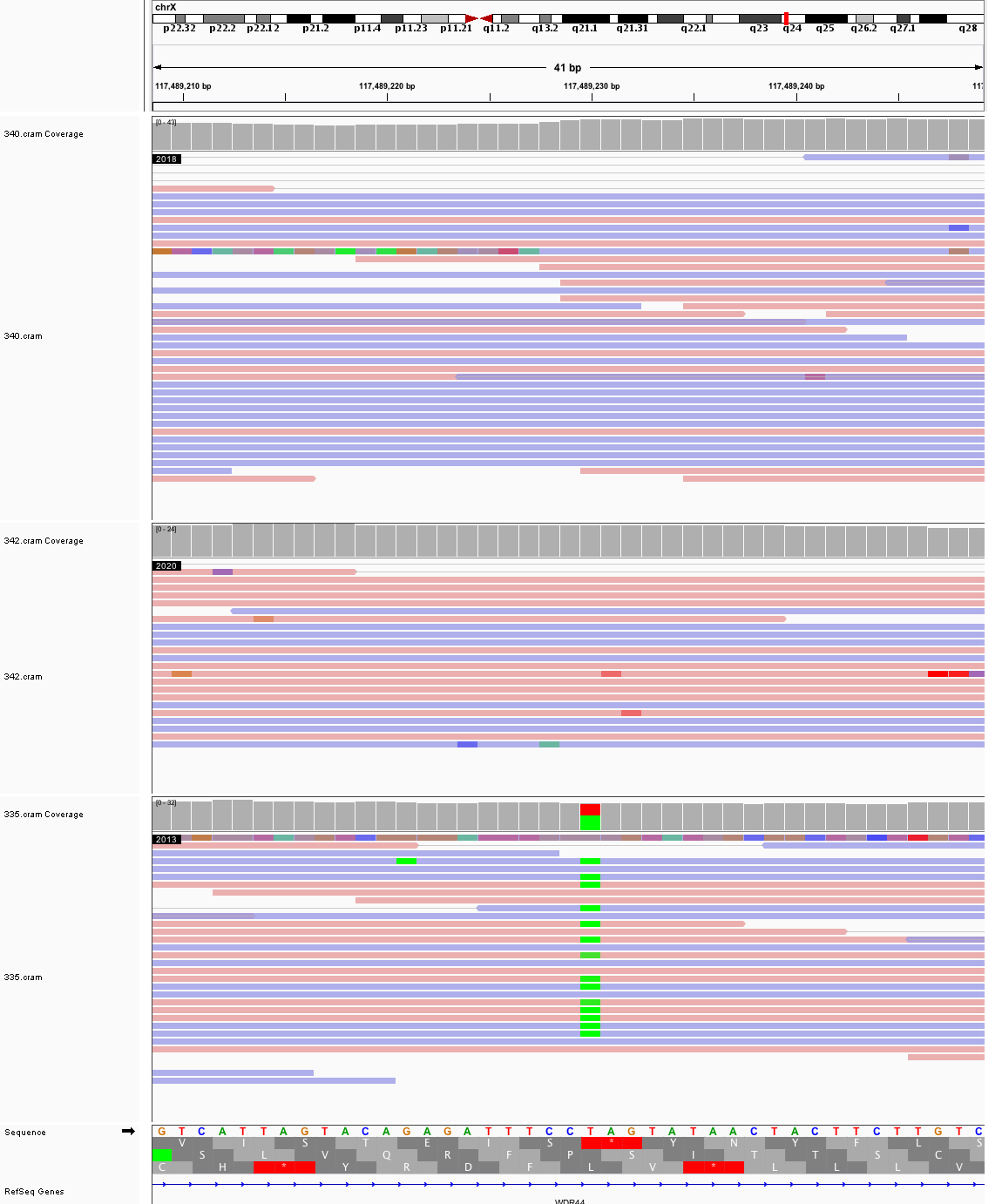
