## Supplementary figures and images for "Large, three-generation CEPH families reveal post-zygotic mosaicism and variability in germline mutation accumulation"

### chr1_102,272,741_102,272,781.png

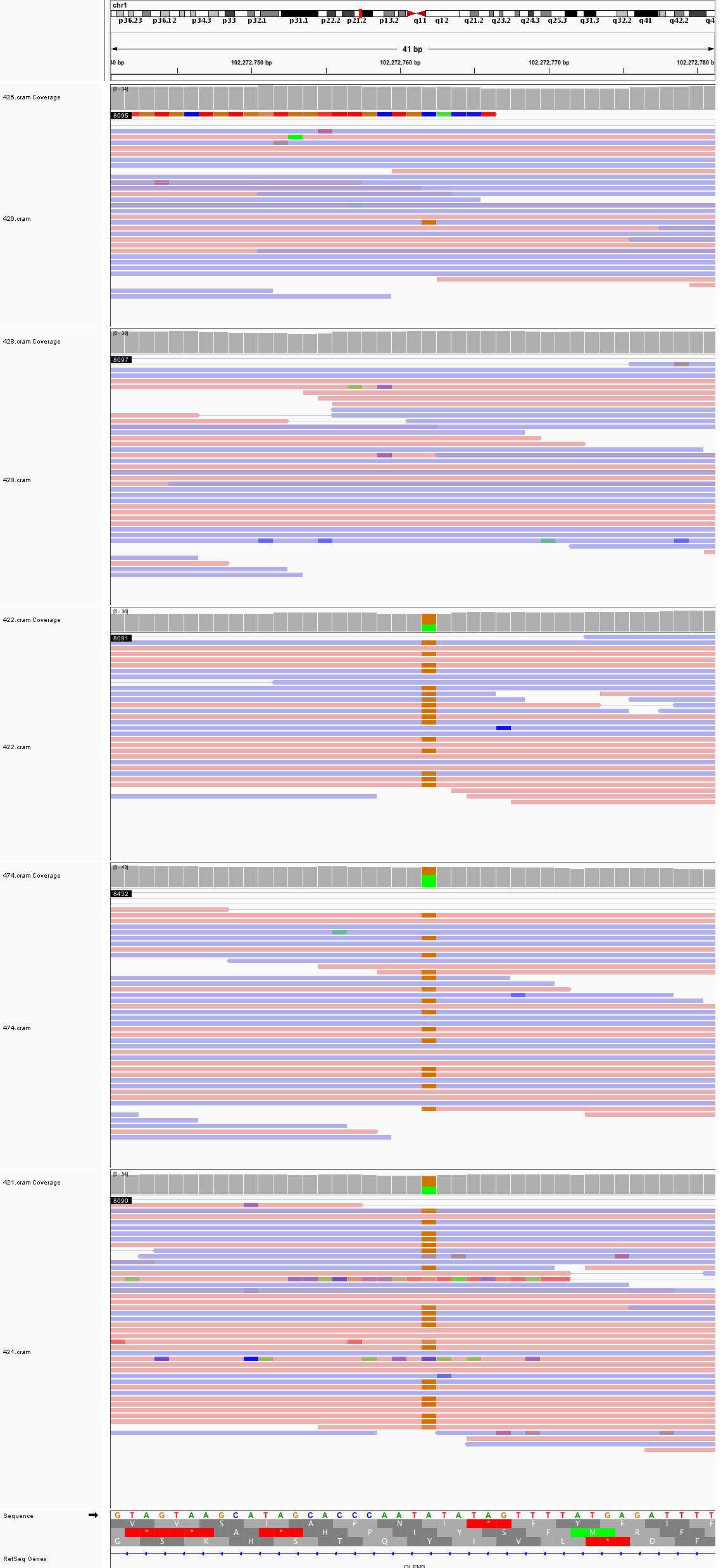

### chr1_116,075,439_116,075,479.png

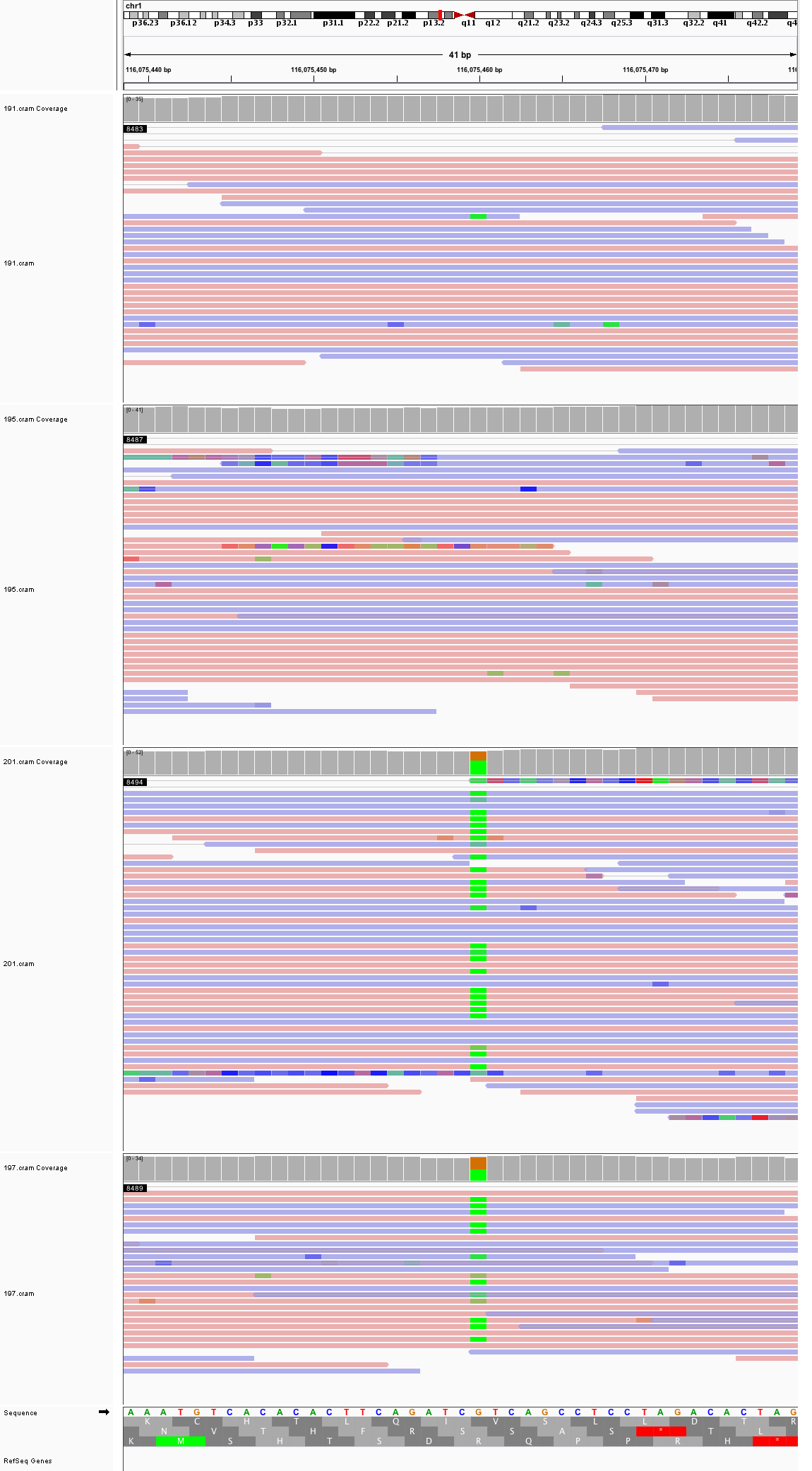

### chr1_184,186,590_184,186,630.png

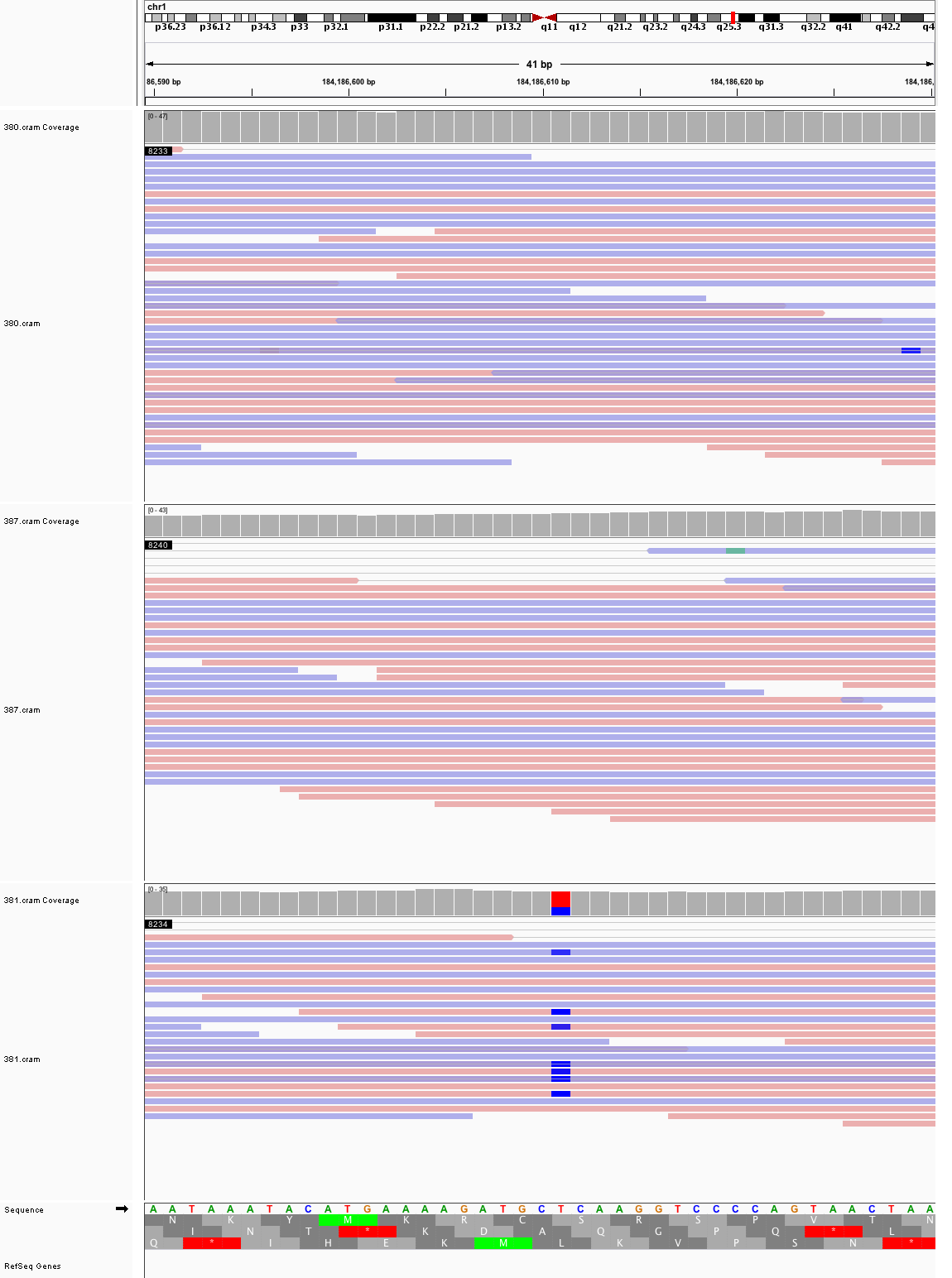

### chr1_239,790,929_239,790,969.png

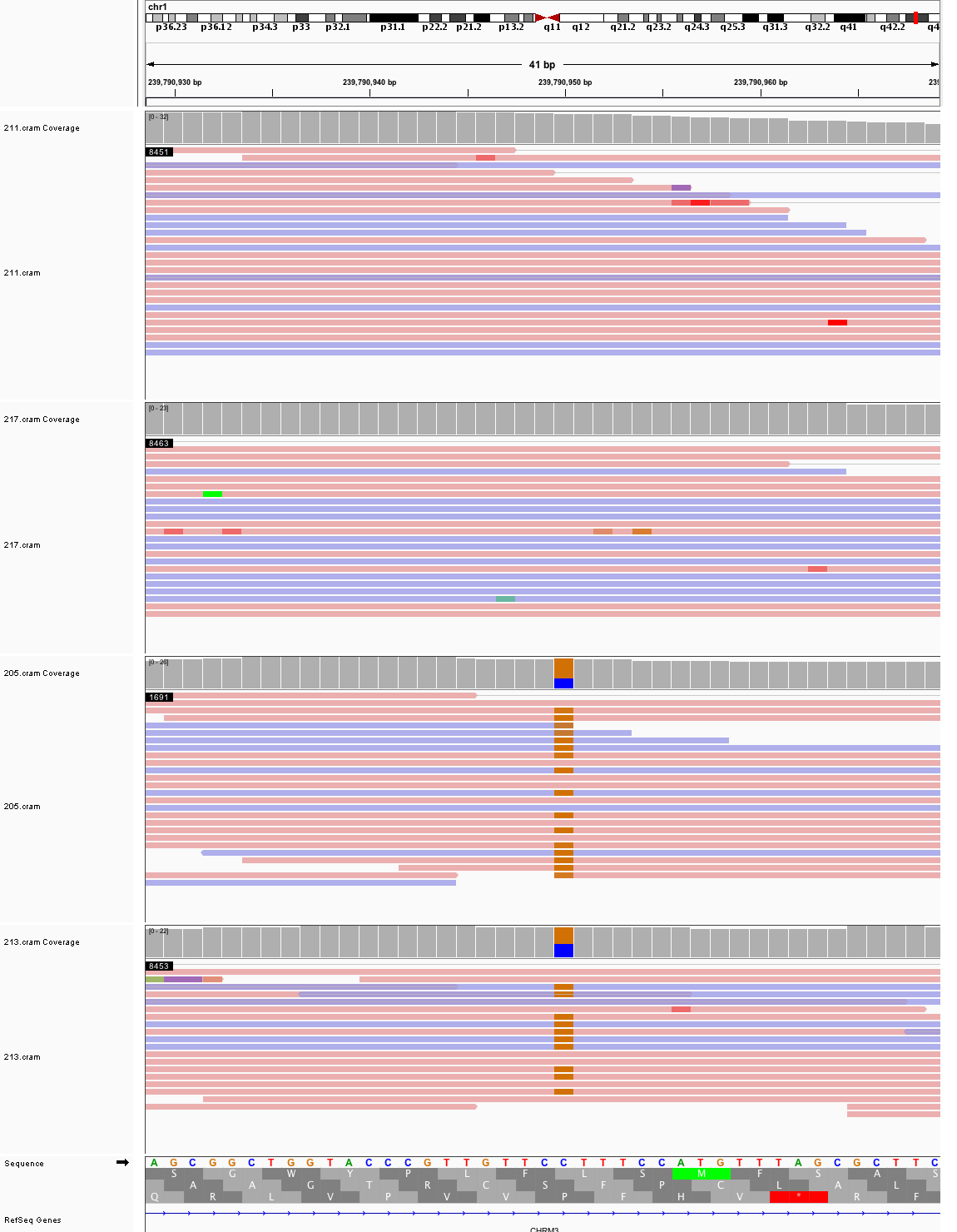

### chr1_241,079,587_241,079,627.png

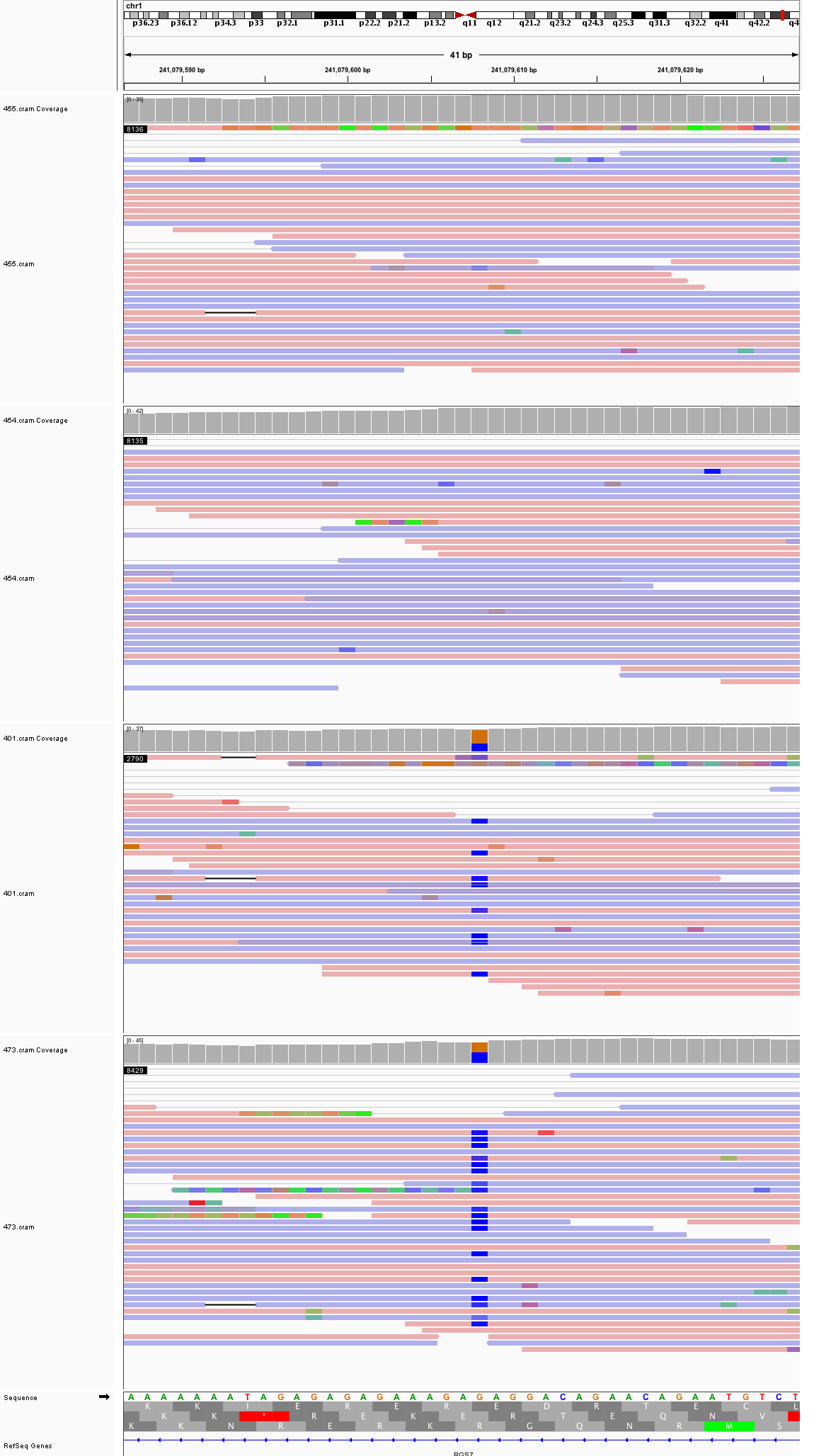

### chr2_3,699,969_3,700,009.png

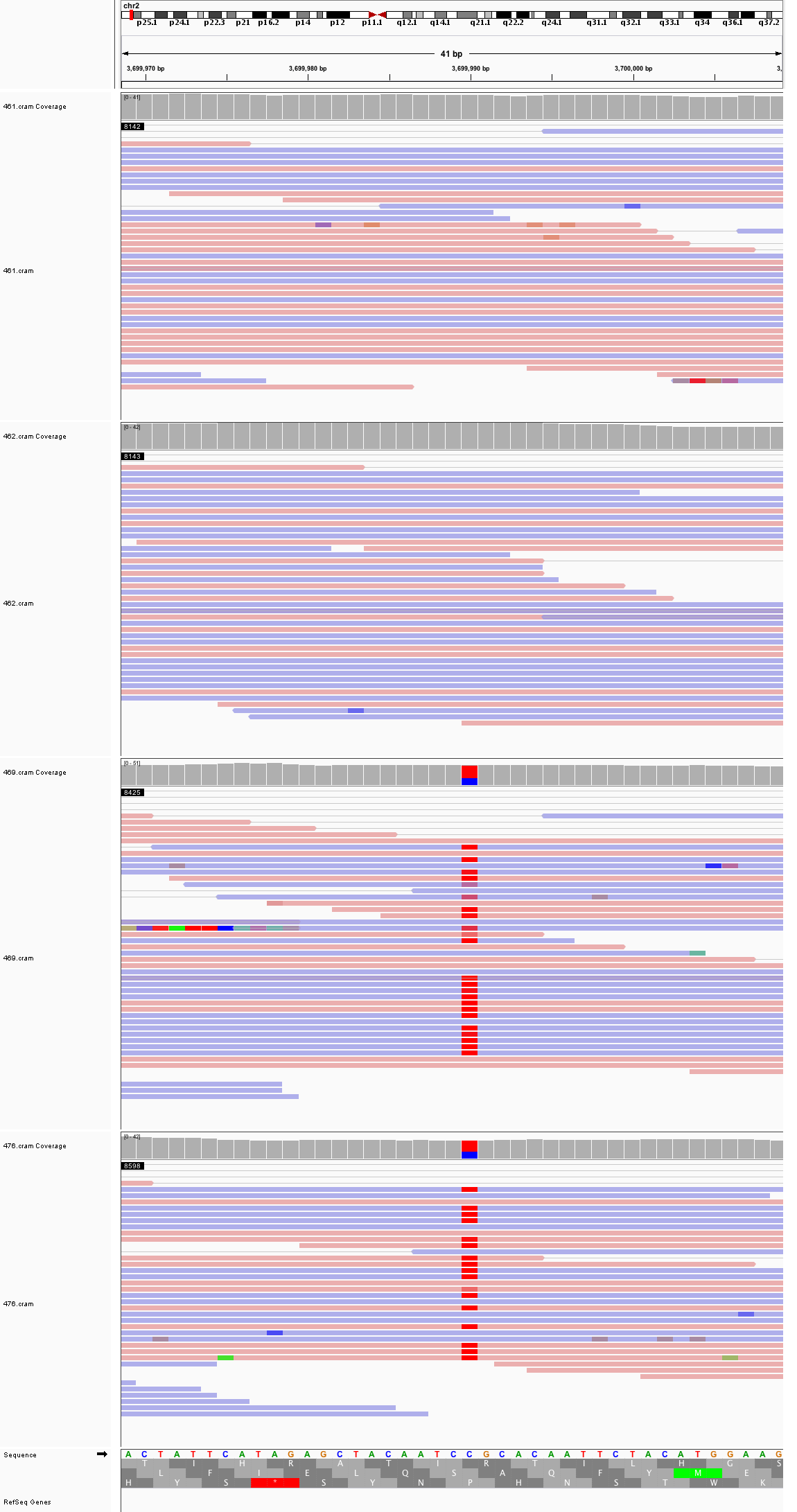

### chr2_16,835,745_16,835,785.png

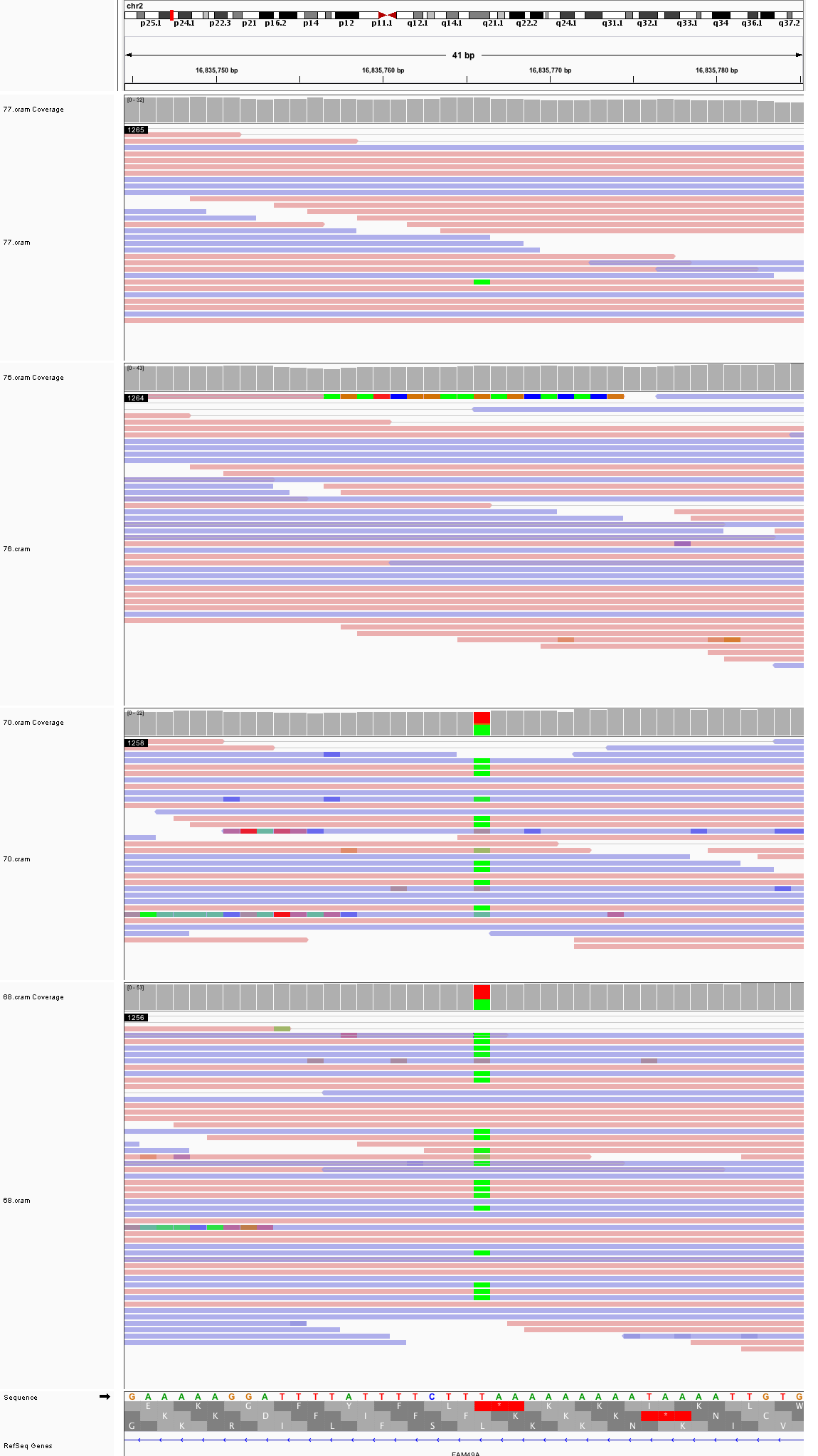

### chr2_31,543,729_31,543,769.png

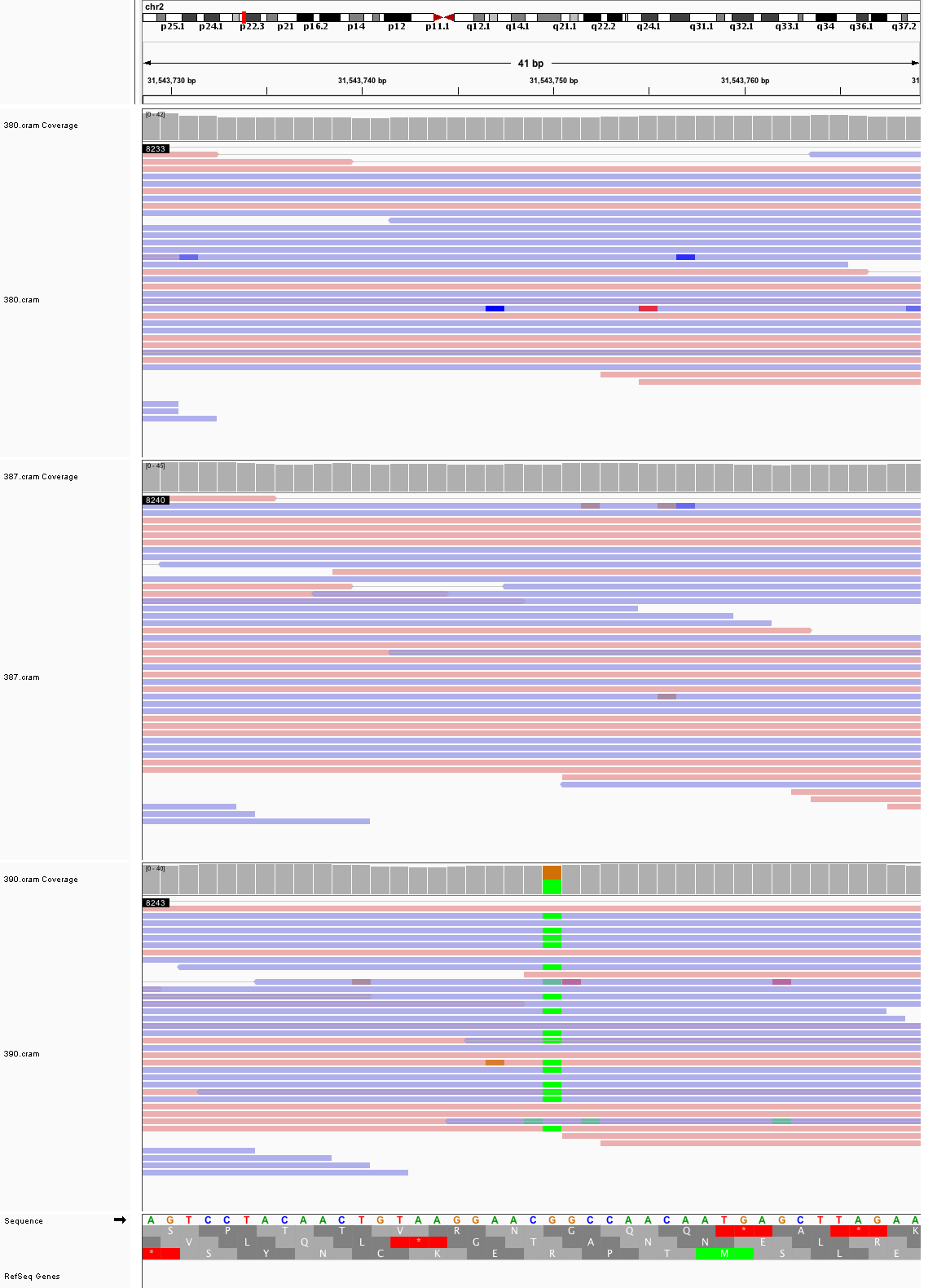

### chr2_44,544,090_44,544,130.png

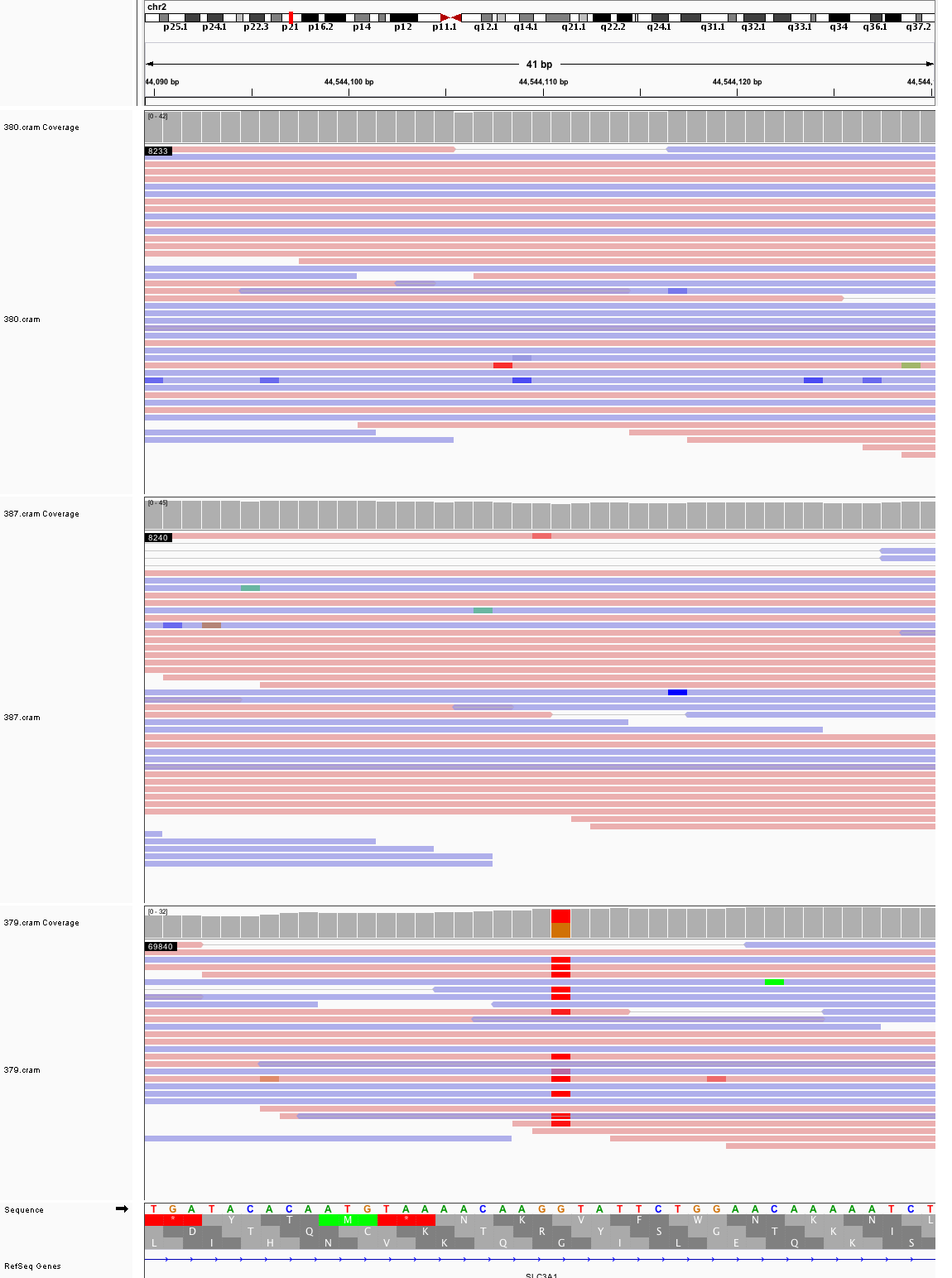

### chr2_121,134,796_121,134,836.png

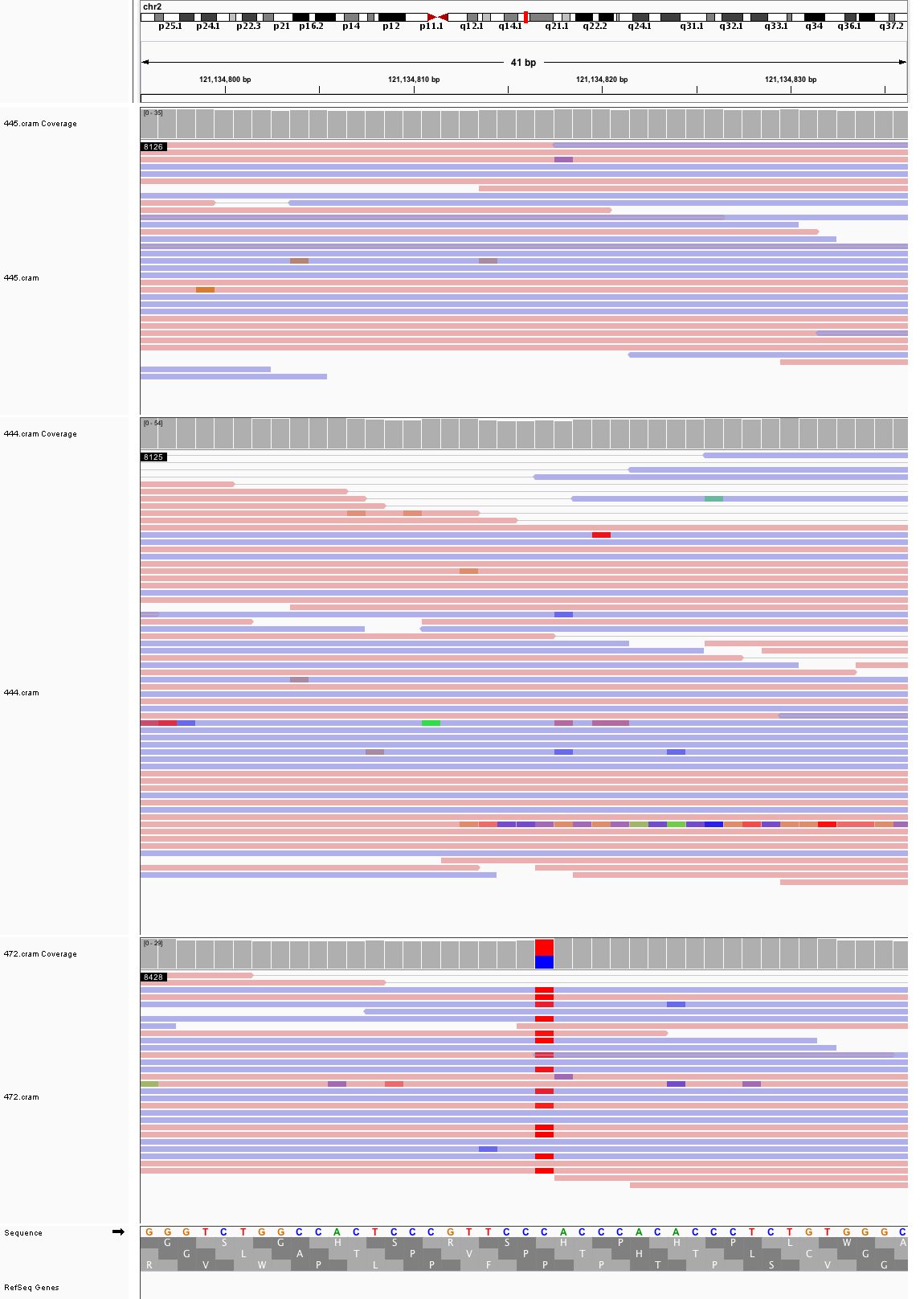

### chr2_148,430,009_148,430,049.png

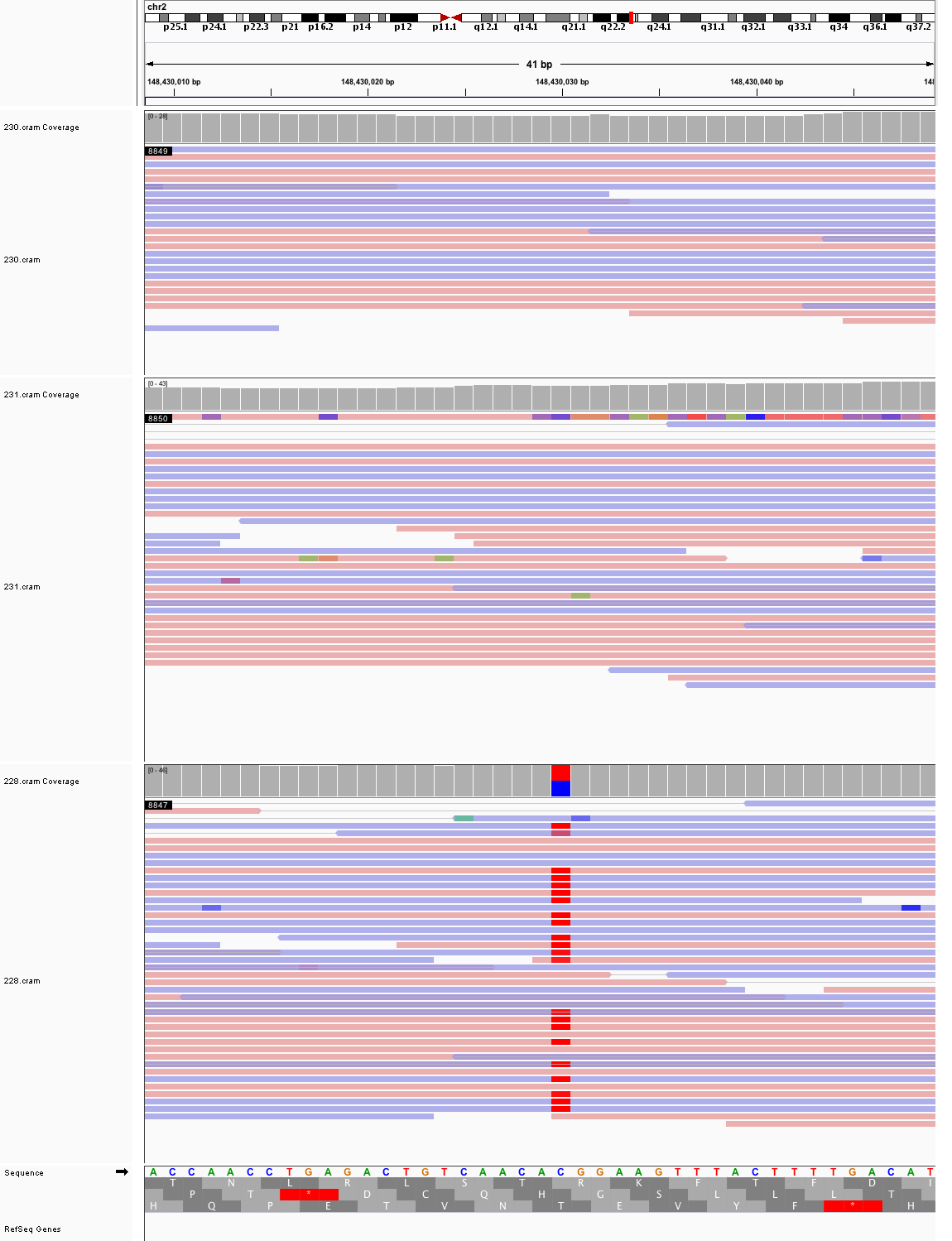

### chr2_163,786,495_163,786,535.png

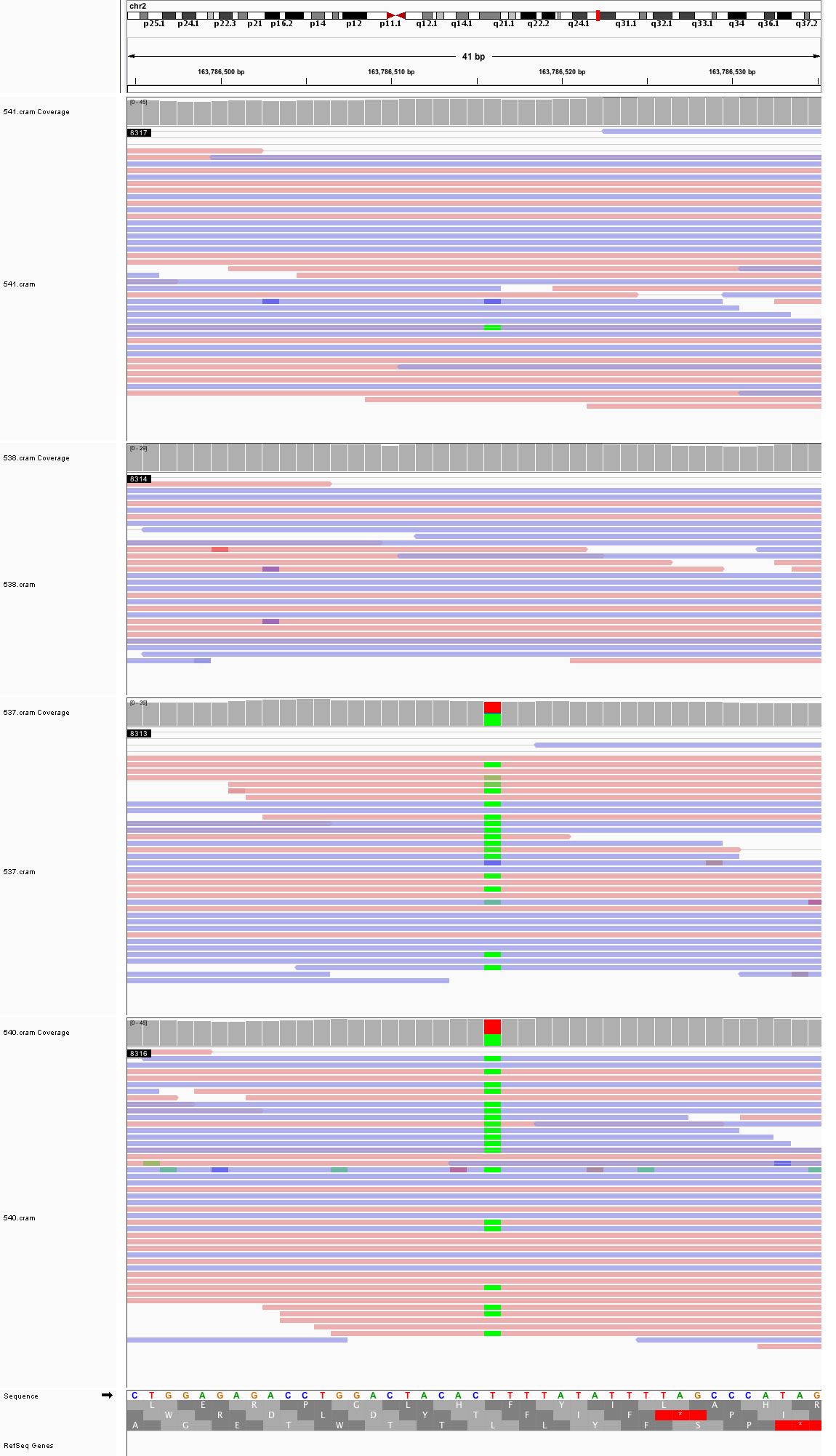

### chr2_183,662,015_183,662,055.png

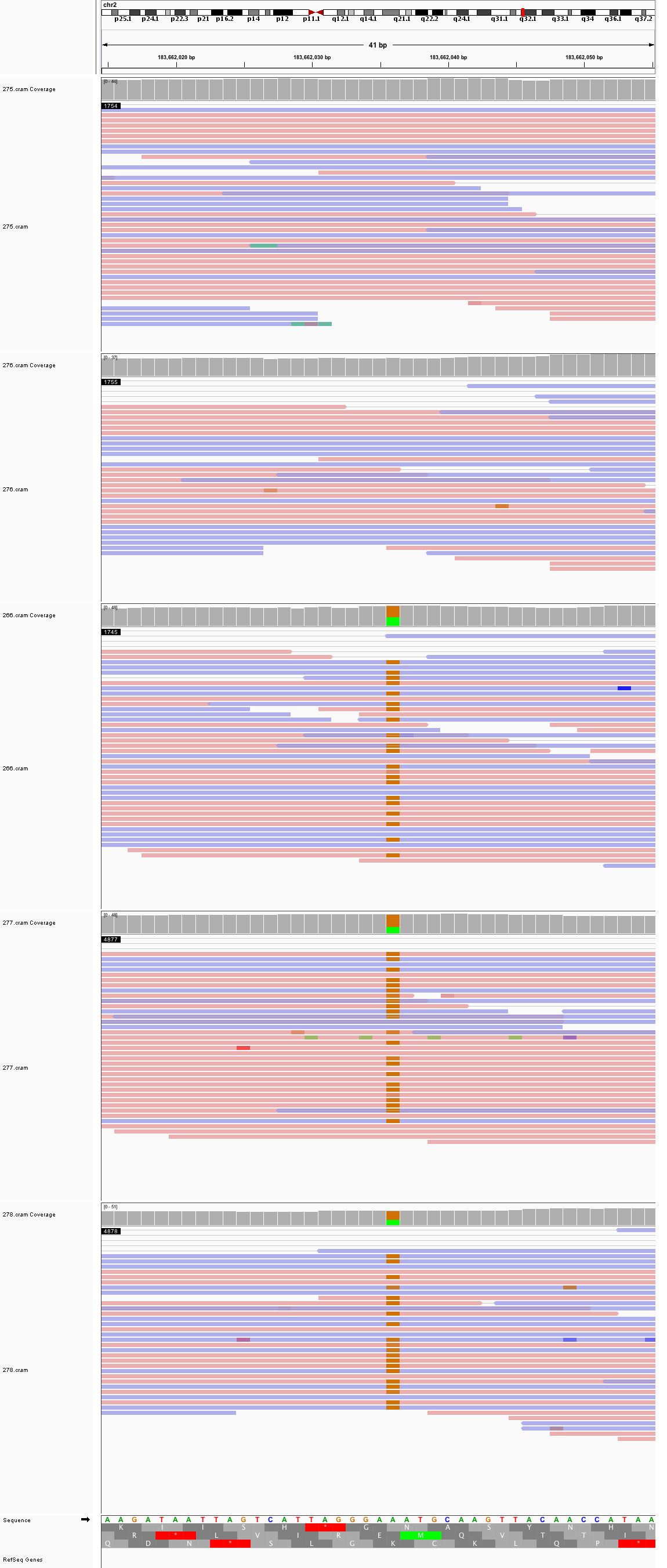

### chr2_185,409,856_185,409,896.png

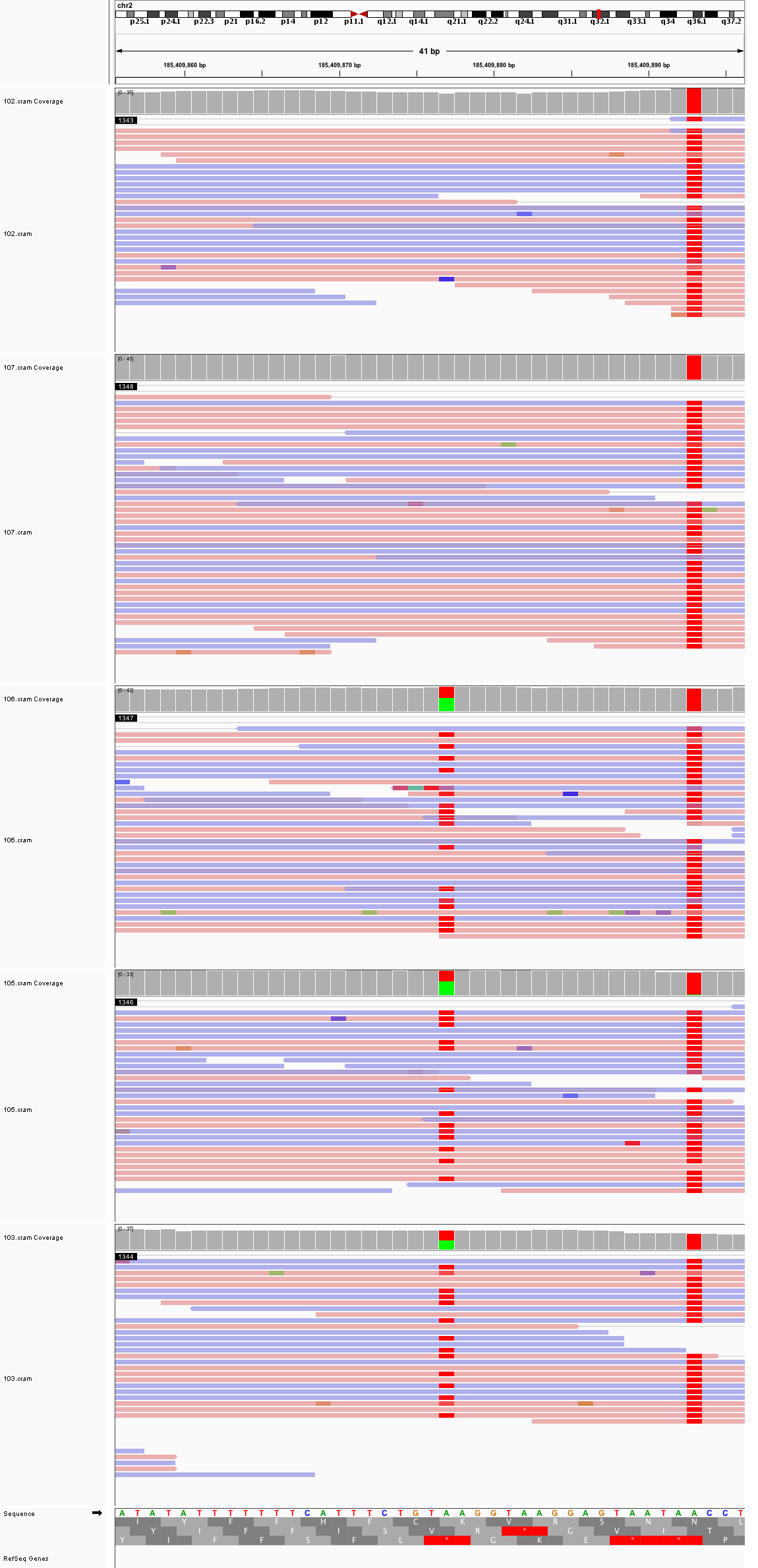

### chr2_209,084,430_209,084,470.png

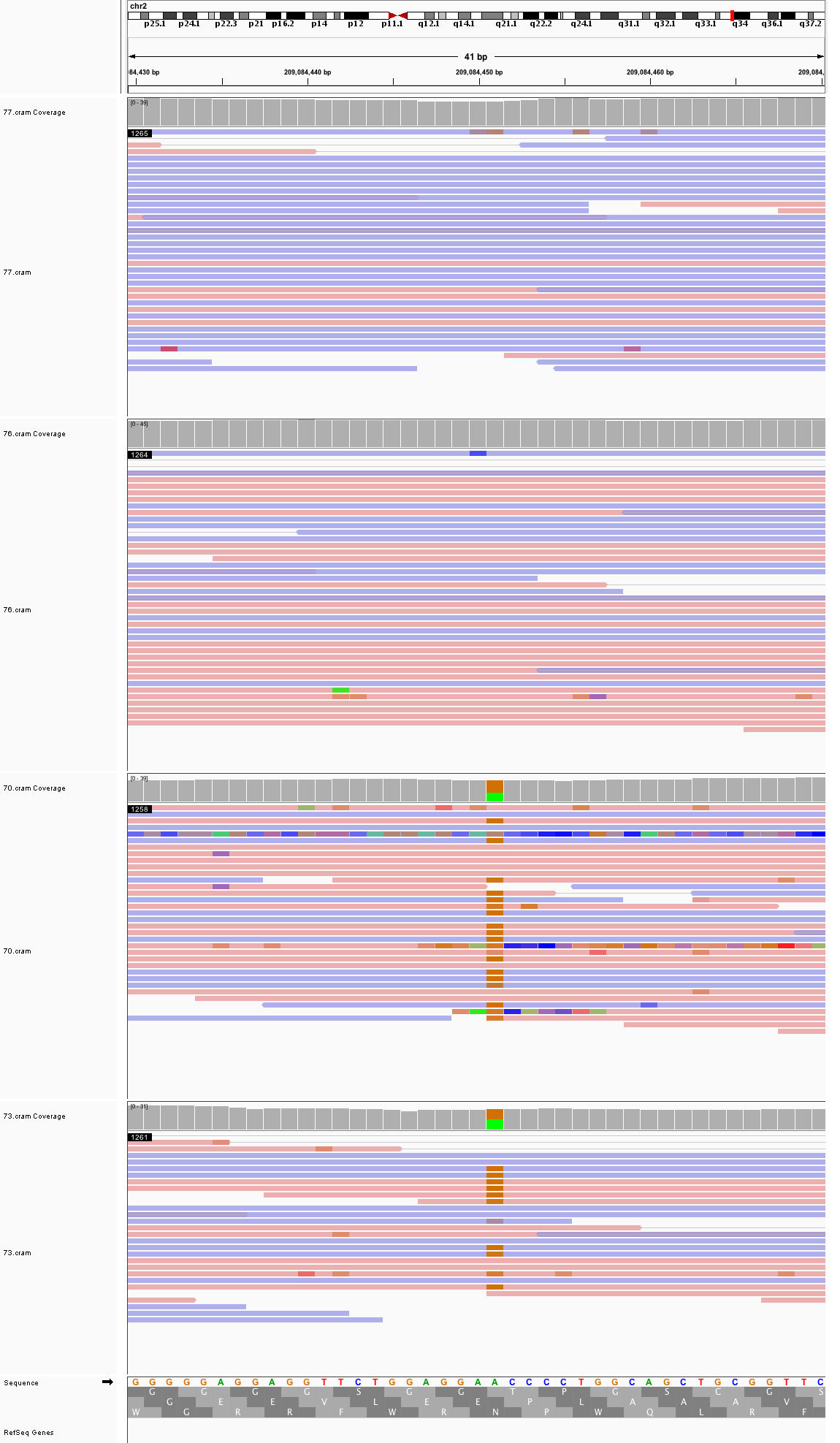

### chr2_220,426,637_220,426,677.png

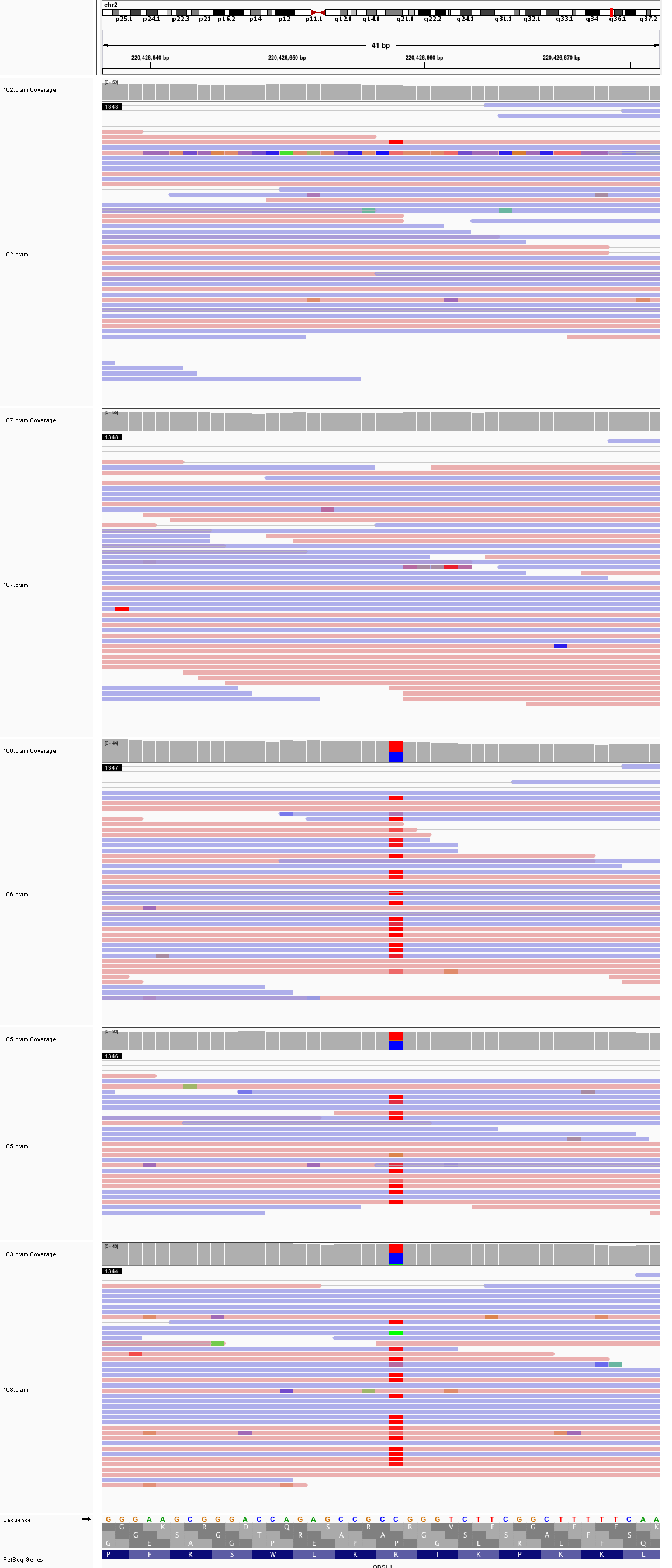

### chr2_222,592,763_222,592,803.png

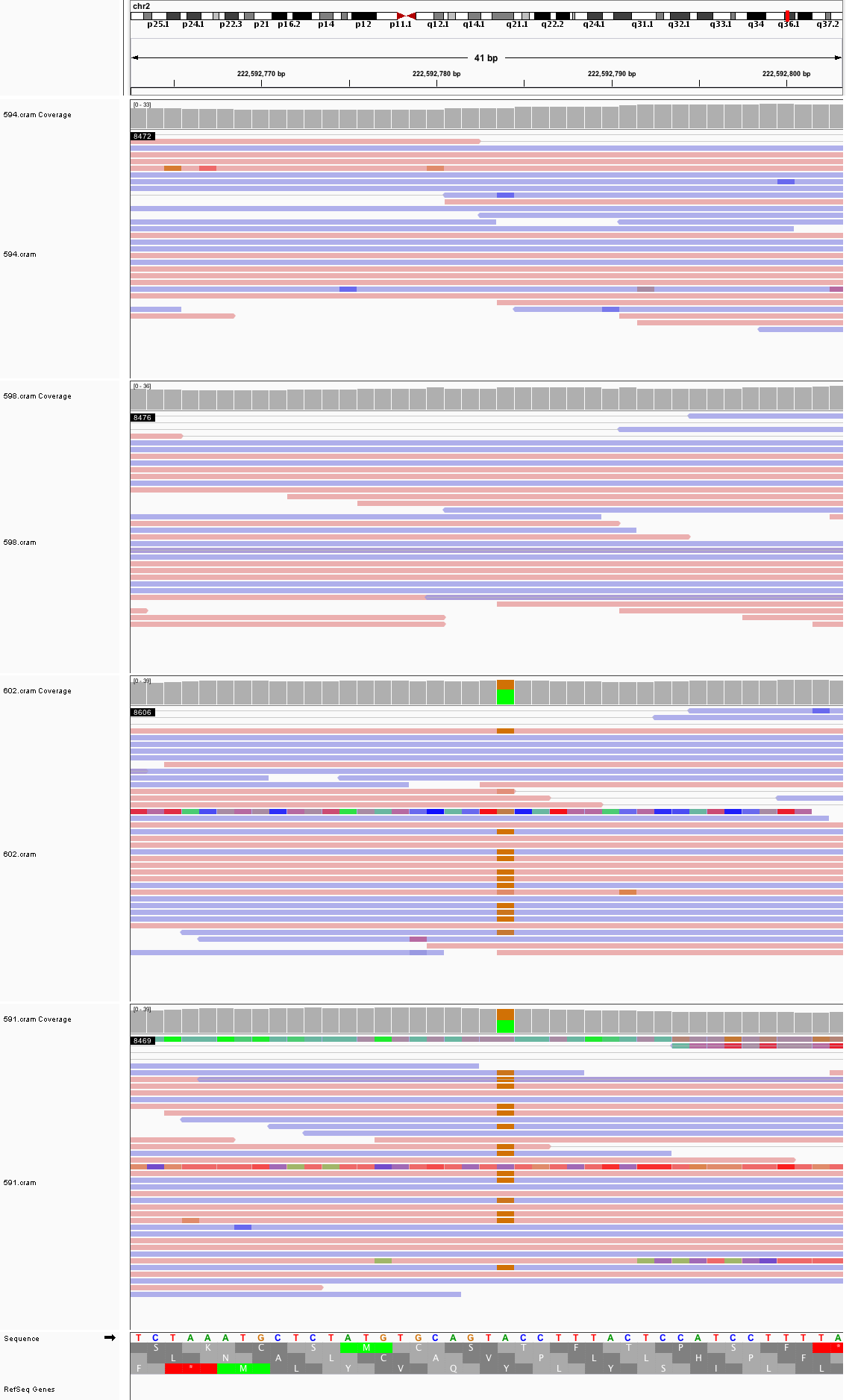

### chr2_230,789,490_230,789,530.png

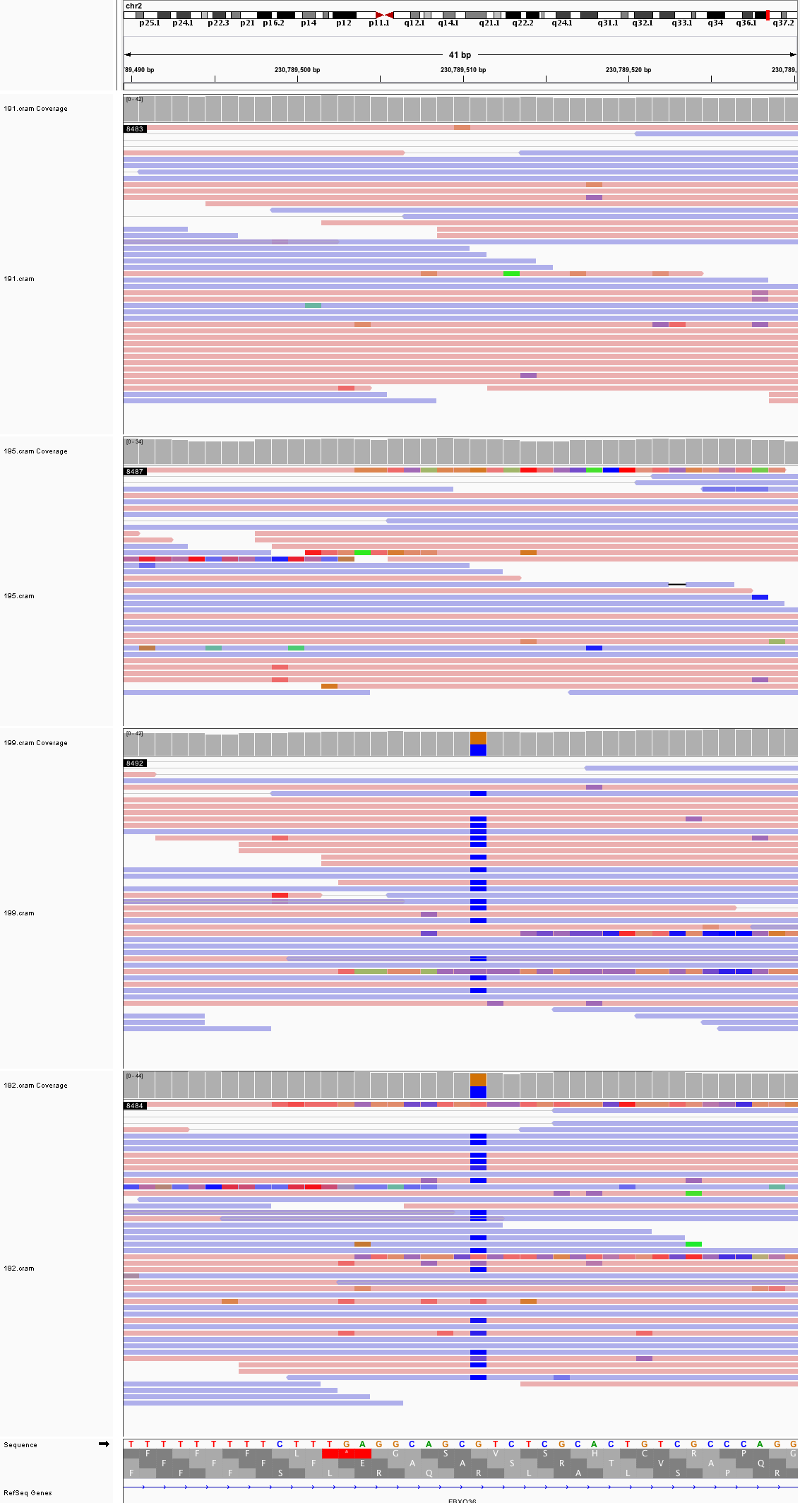

### chr2_231,591,503_231,591,543.png

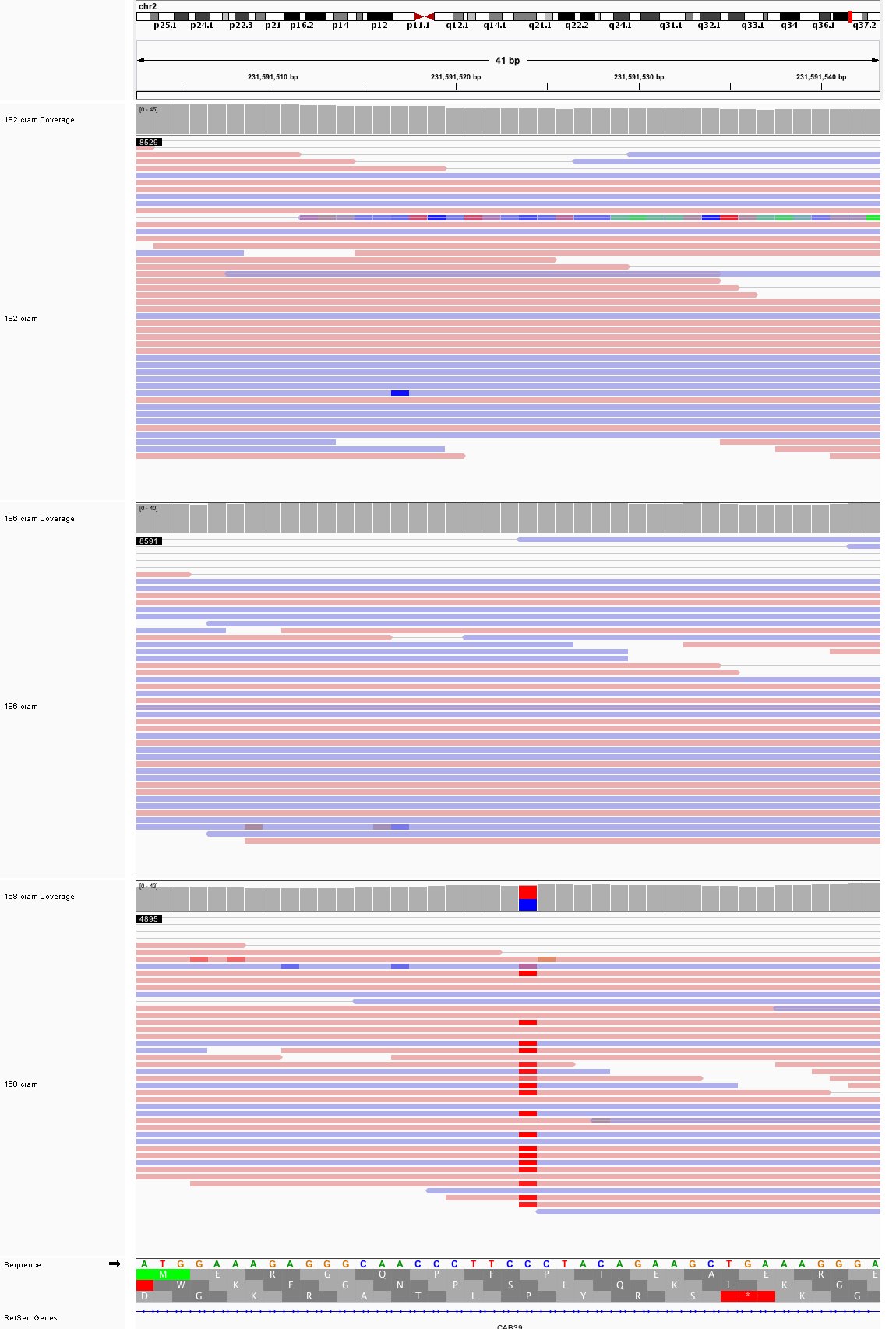

### chr3_20,254,383_20,254,423.png

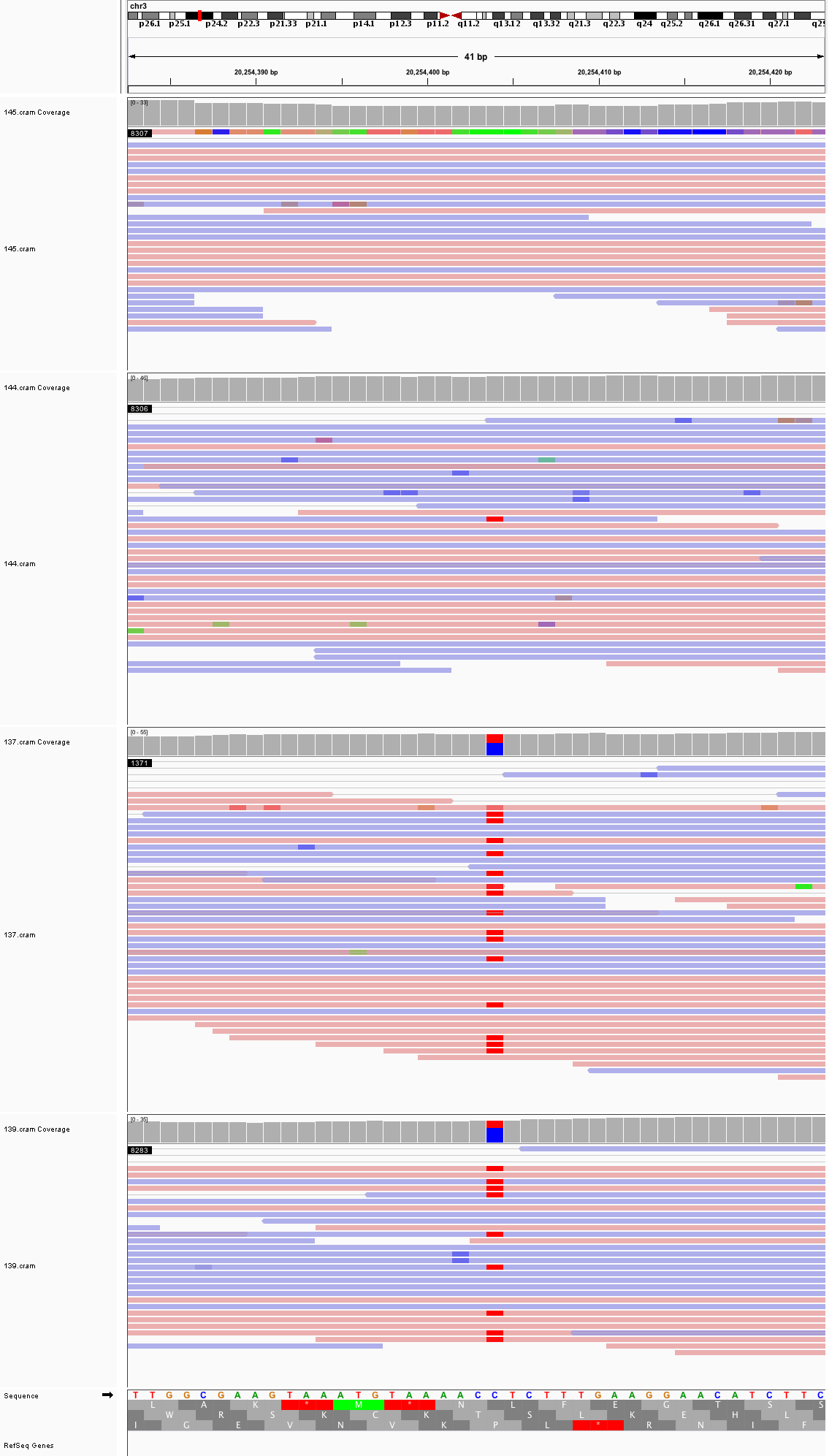

### chr3_20,764,822_20,764,862.png

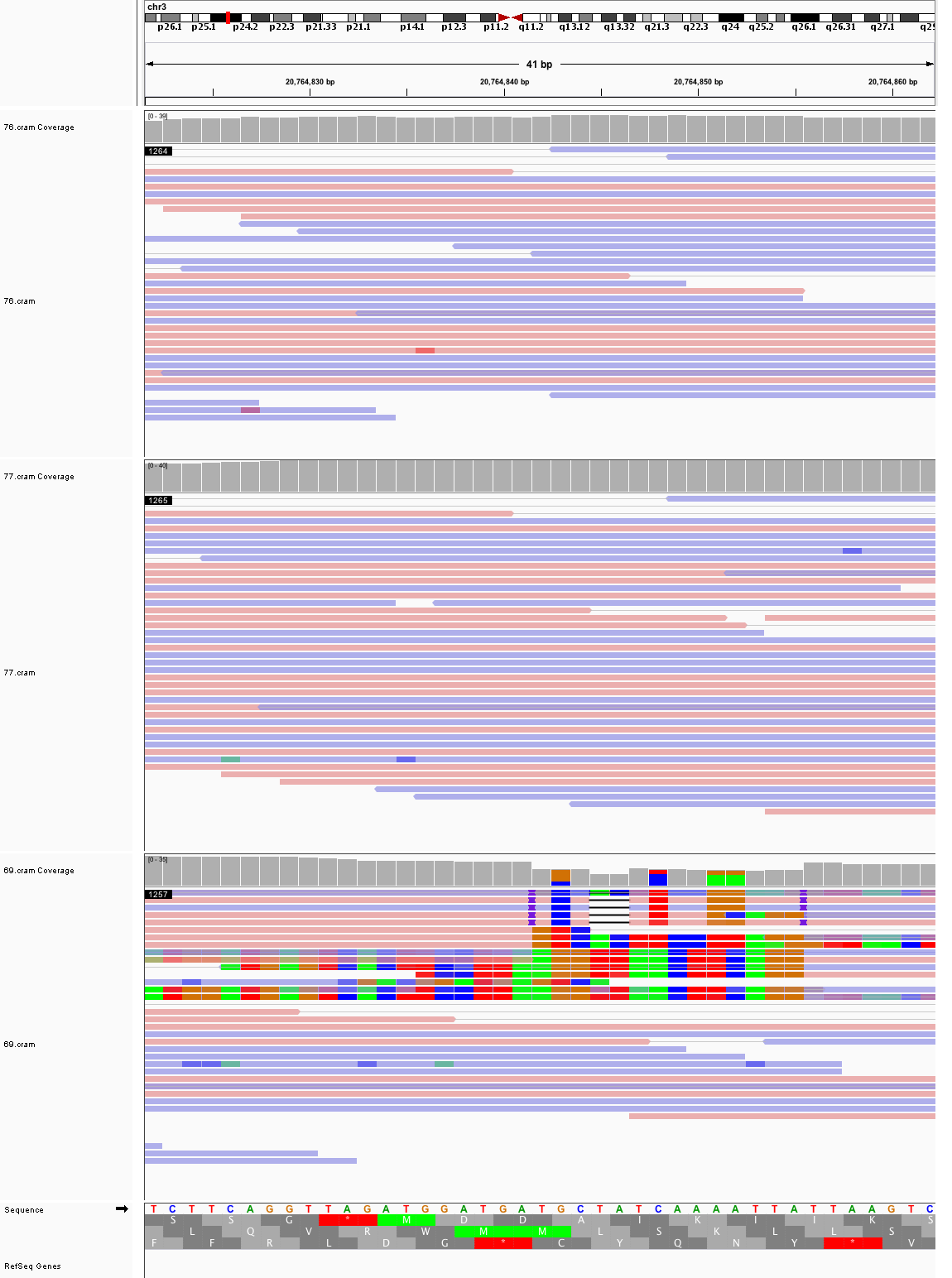

### chr3_37,841,468_37,841,508.png

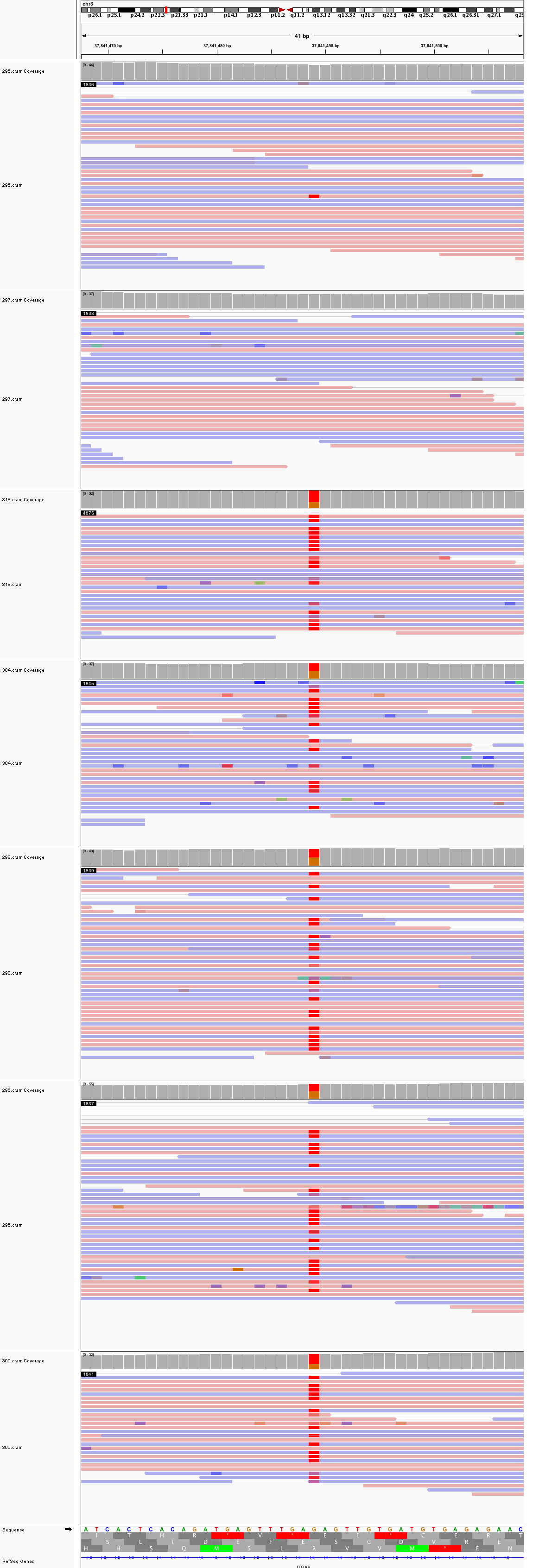

### chr3_123,849,555_123,849,595.png

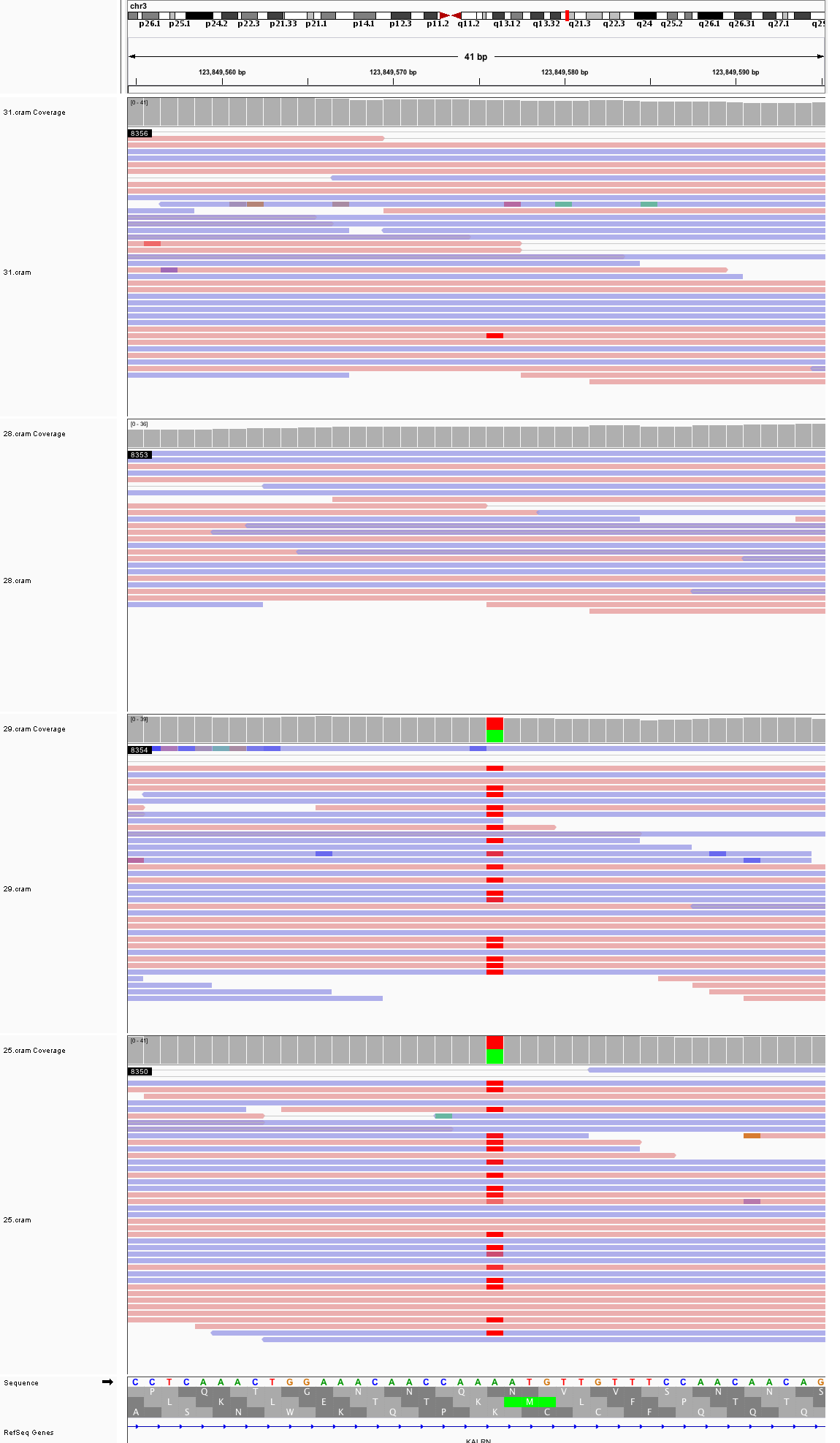

### chr3_128,747,710_128,747,750.png

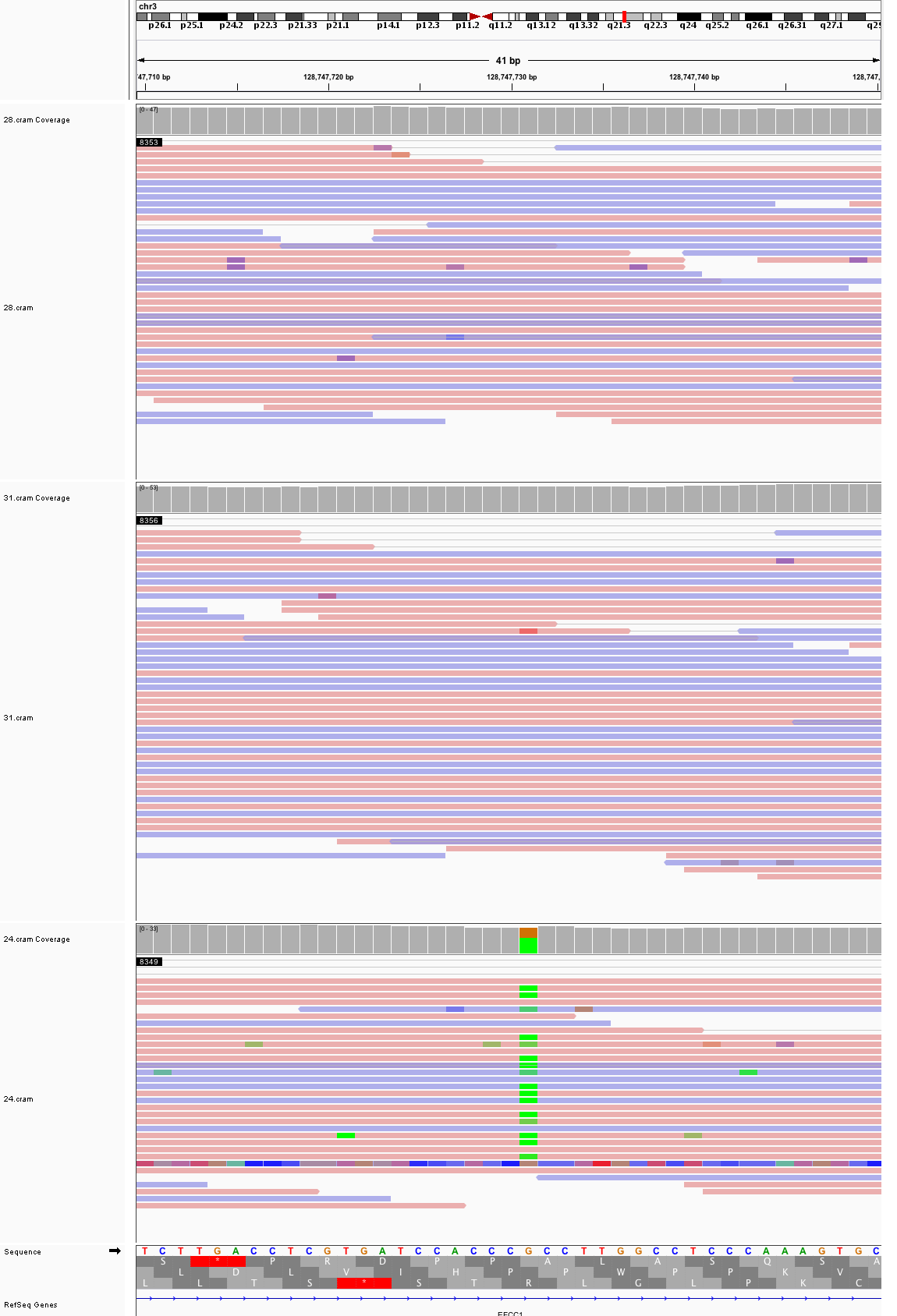

### chr3_136,130,012_136,130,052.png

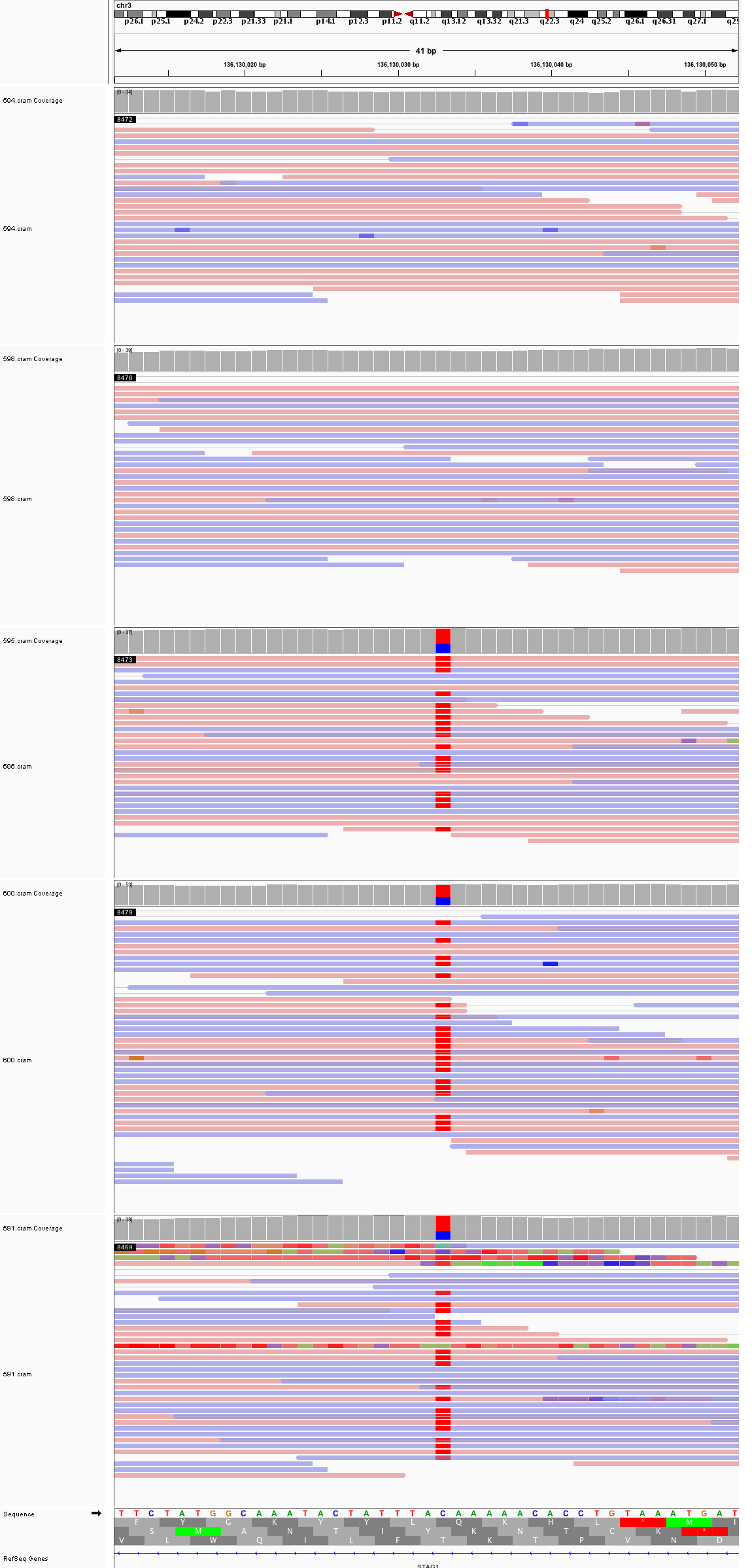

### chr3_187,220,154_187,220,194.png

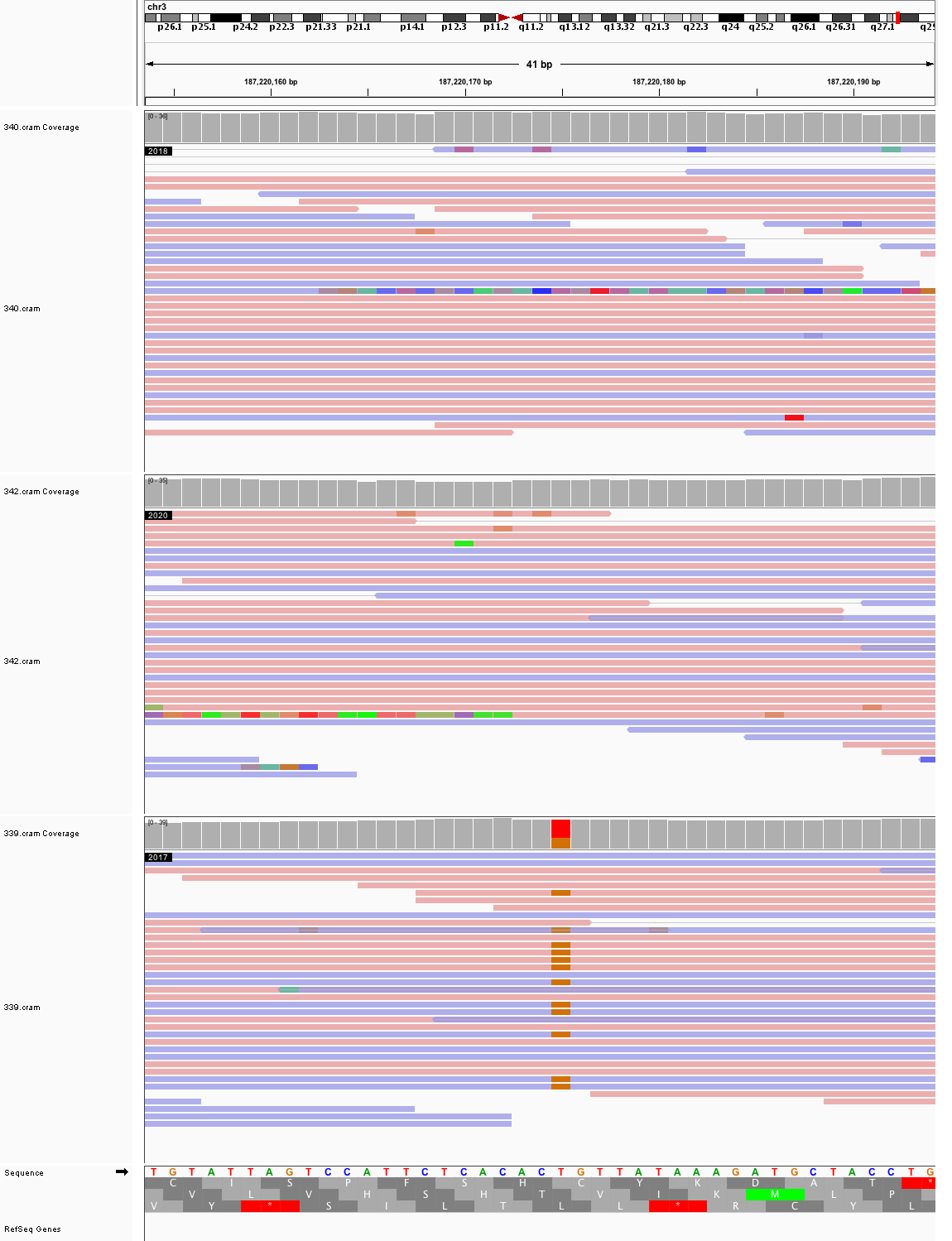

### chr4_13,461,087_13,461,127.png

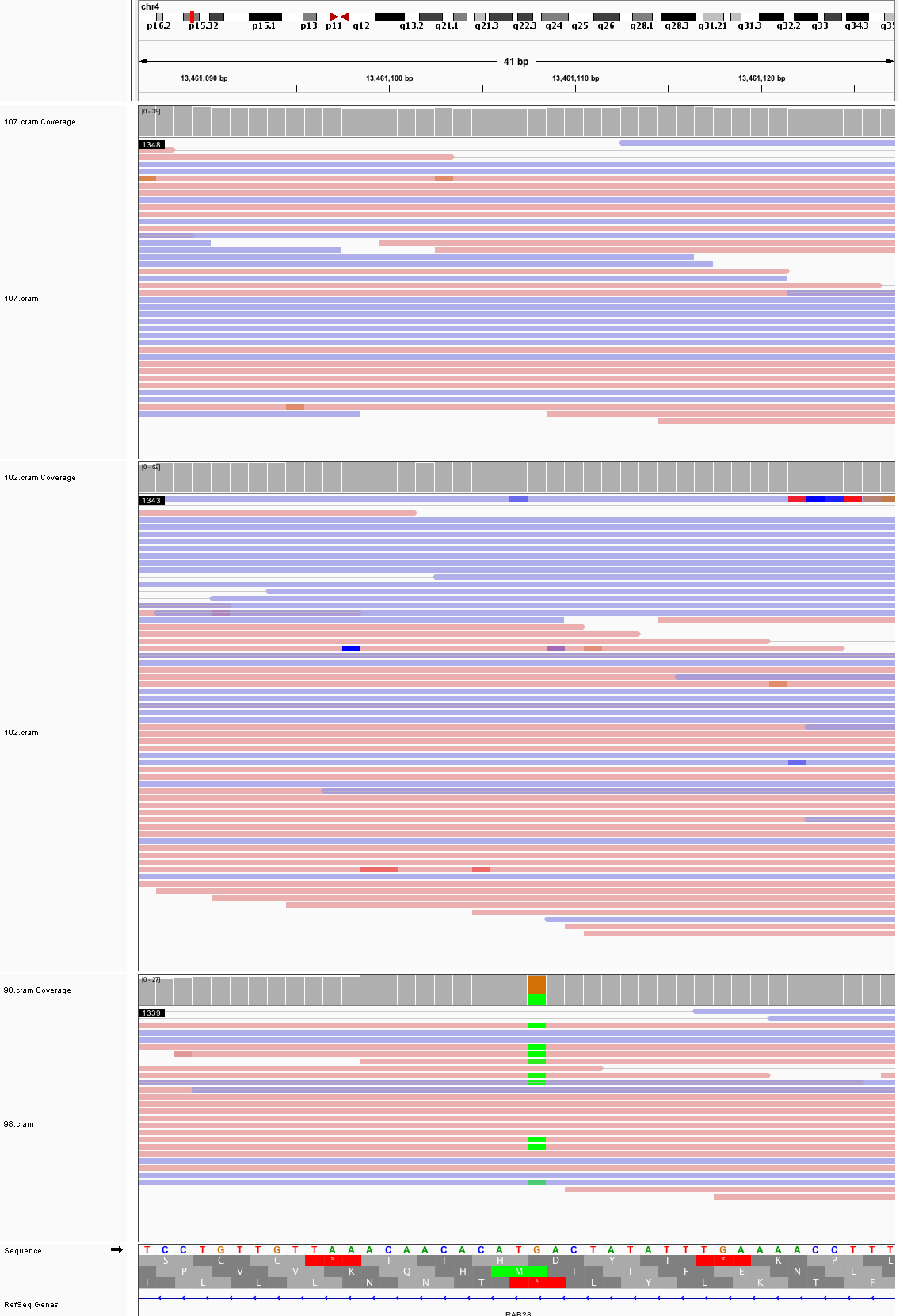

### chr4_33,166,413_33,166,453.png

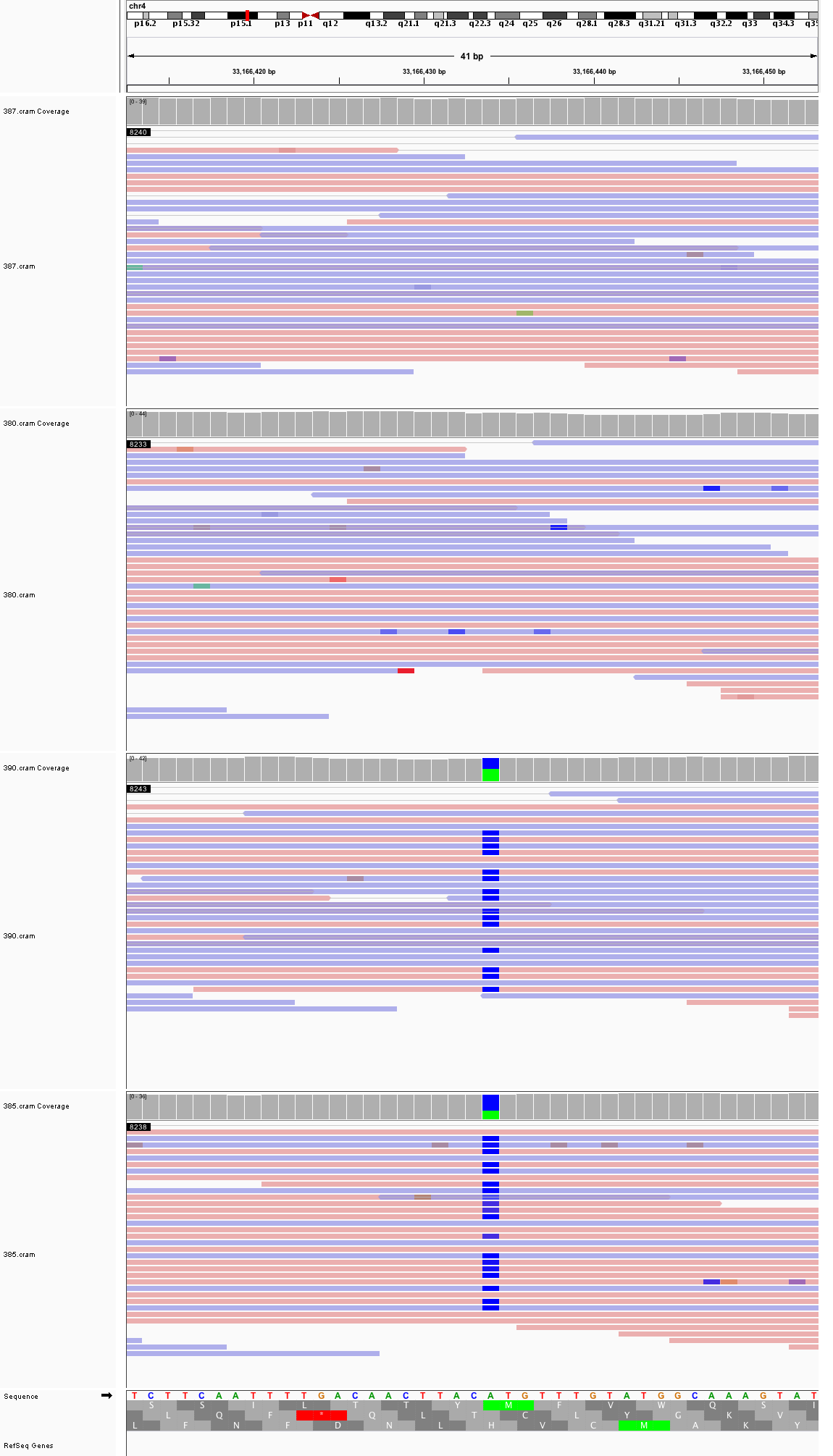

### chr4_66,904,505_66,904,545.png

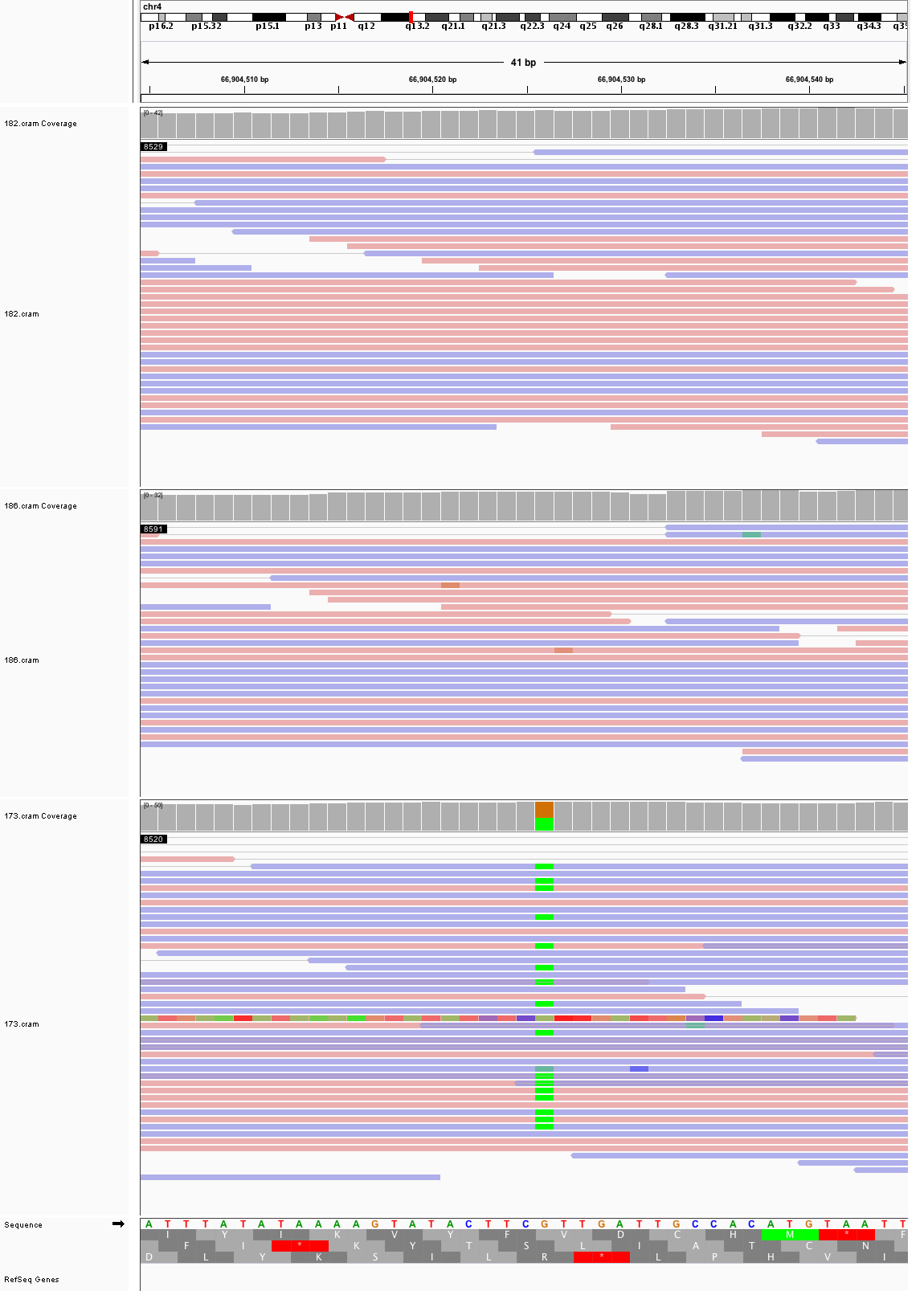

### chr4_106,654,923_106,654,963.png

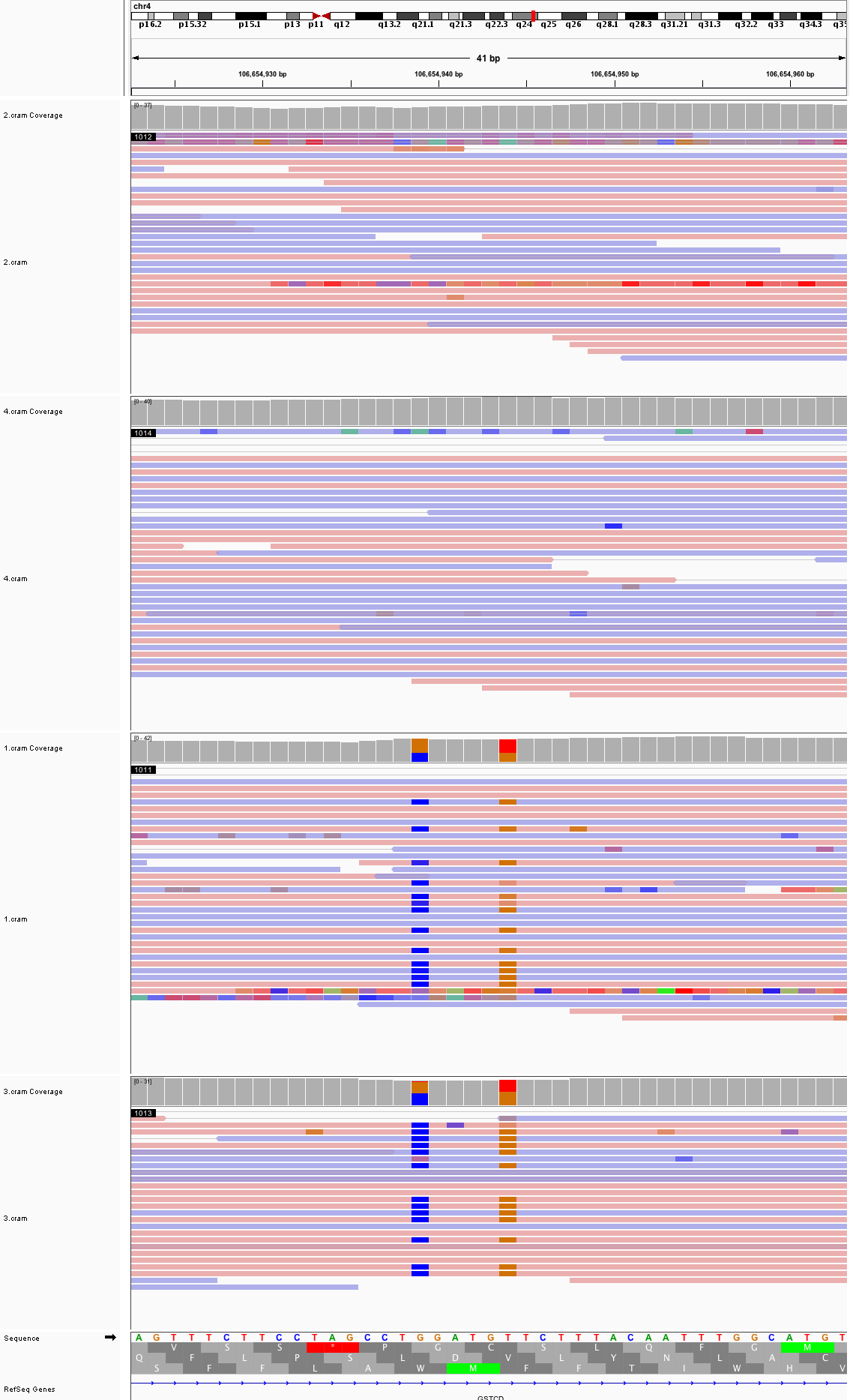

### chr4_143,683,468_143,683,508.png

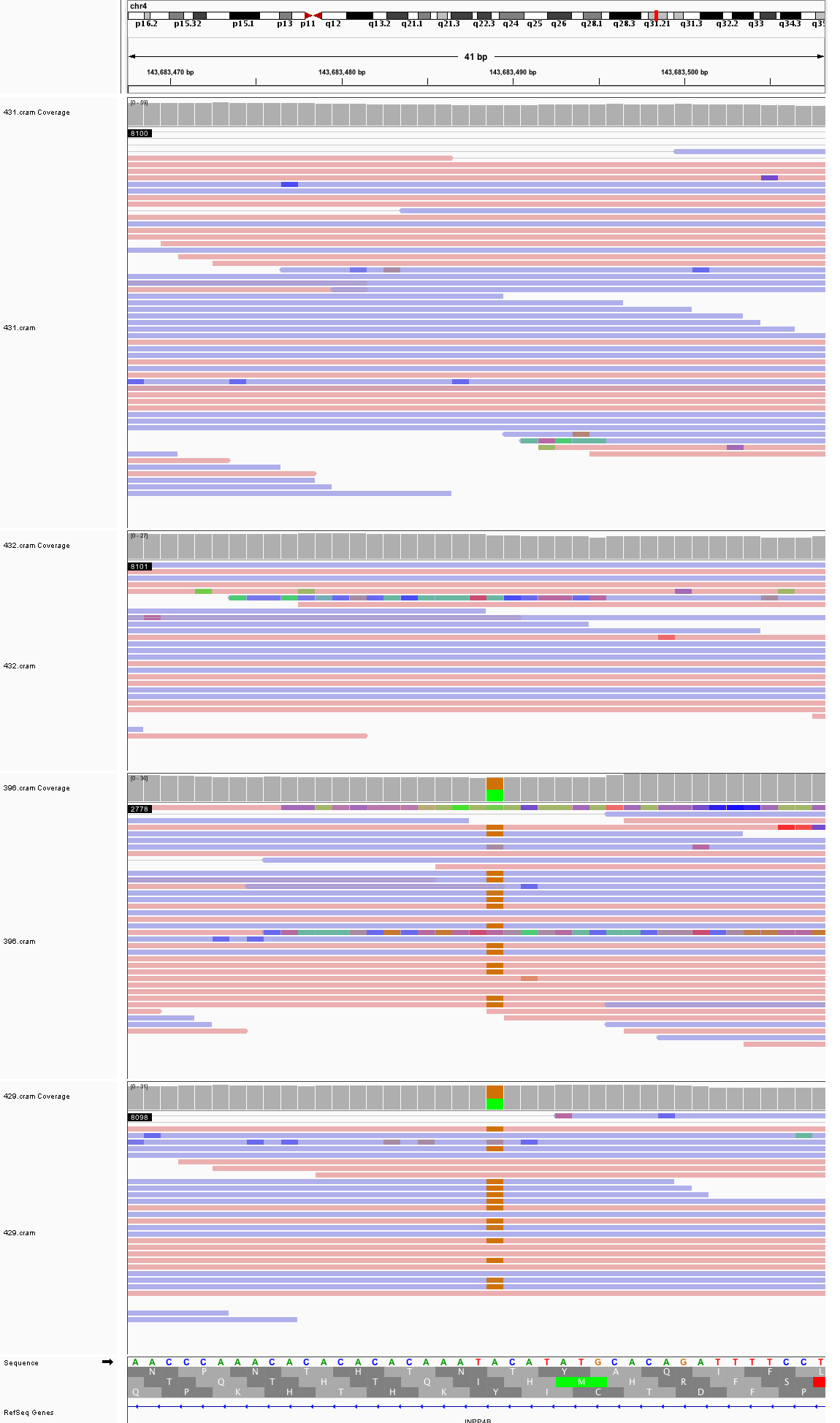

### chr4_176,443,194_176,443,234.png

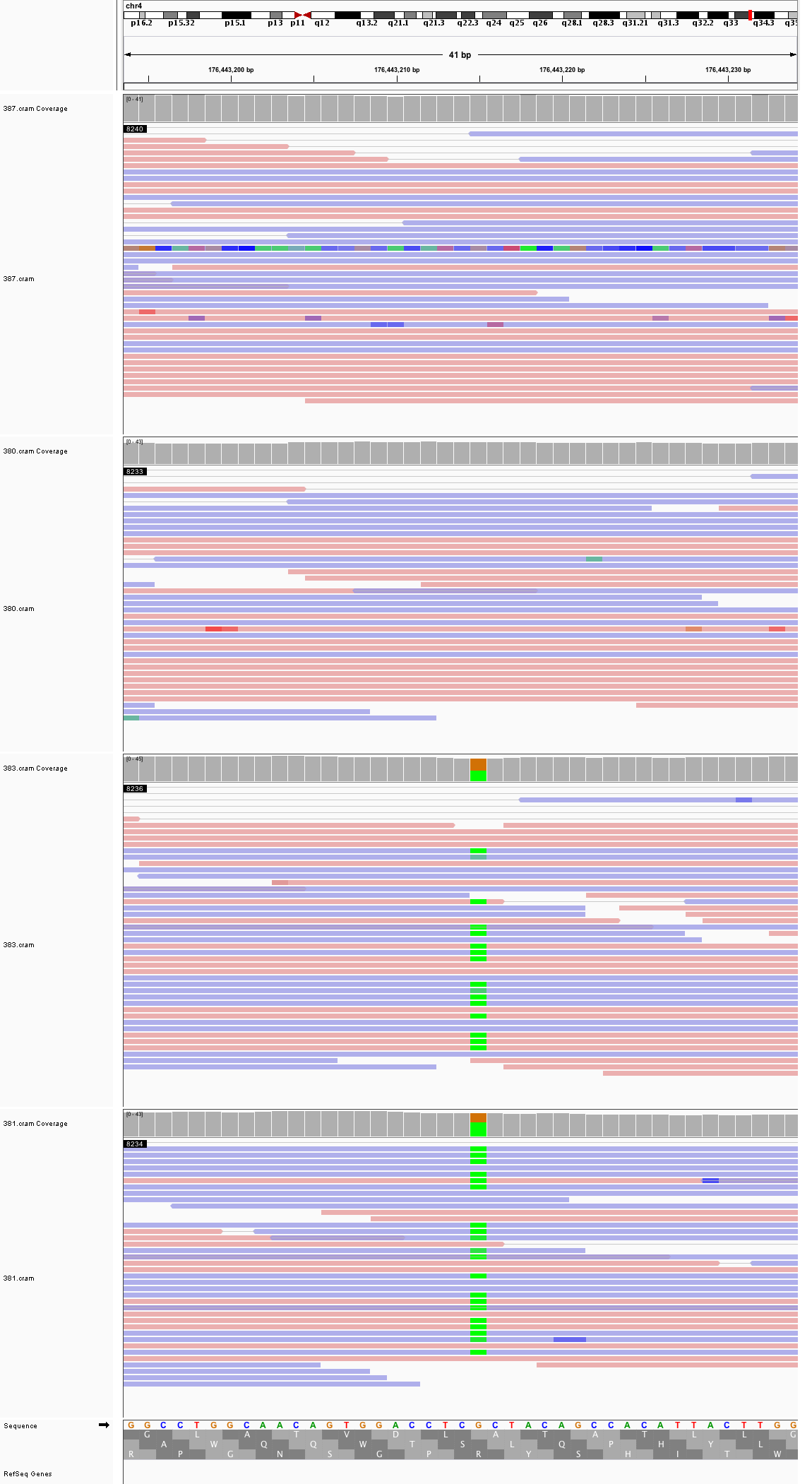

### chr5_9,747,950_9,747,990.png

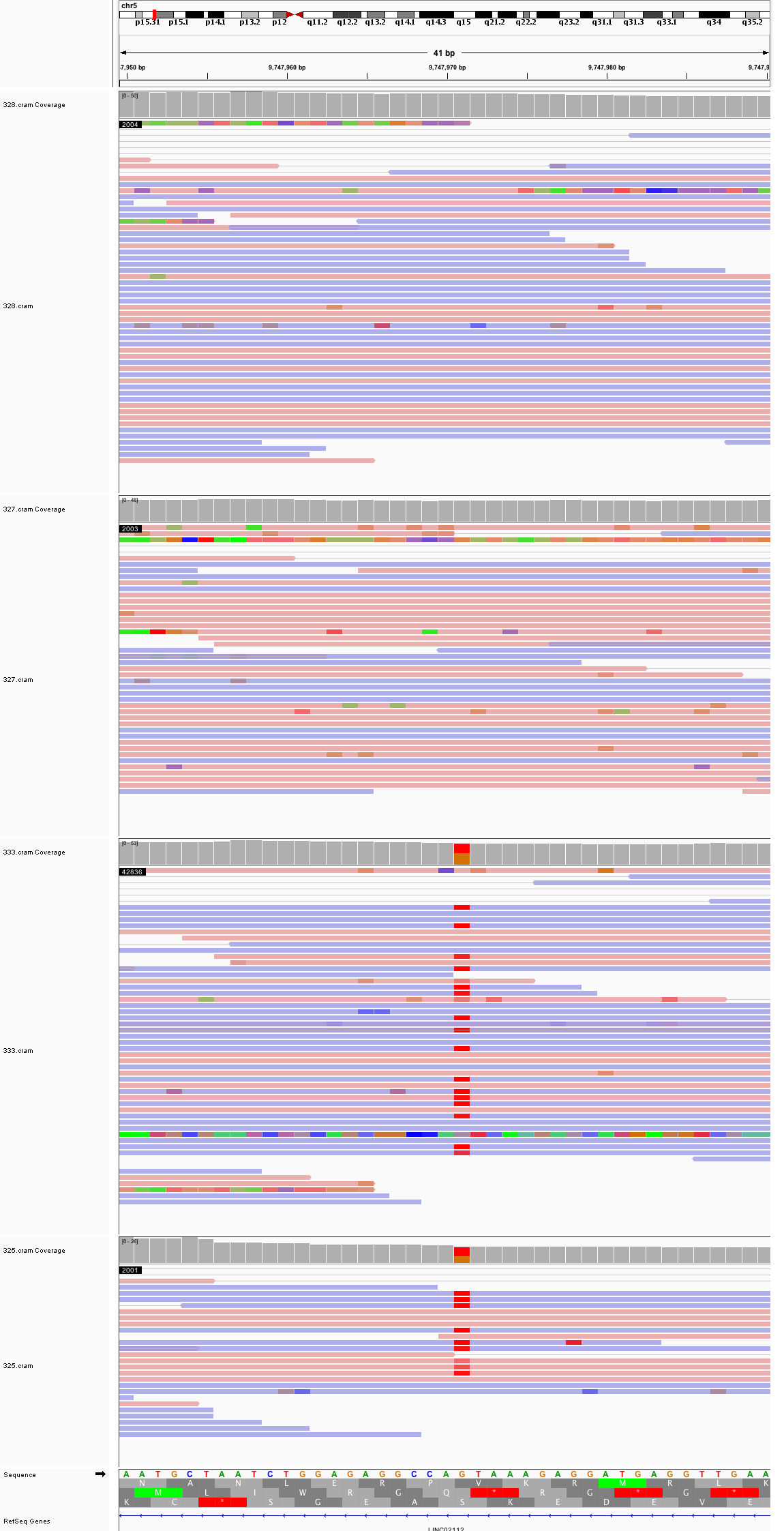

### chr5_62,260,770_62,260,810.png

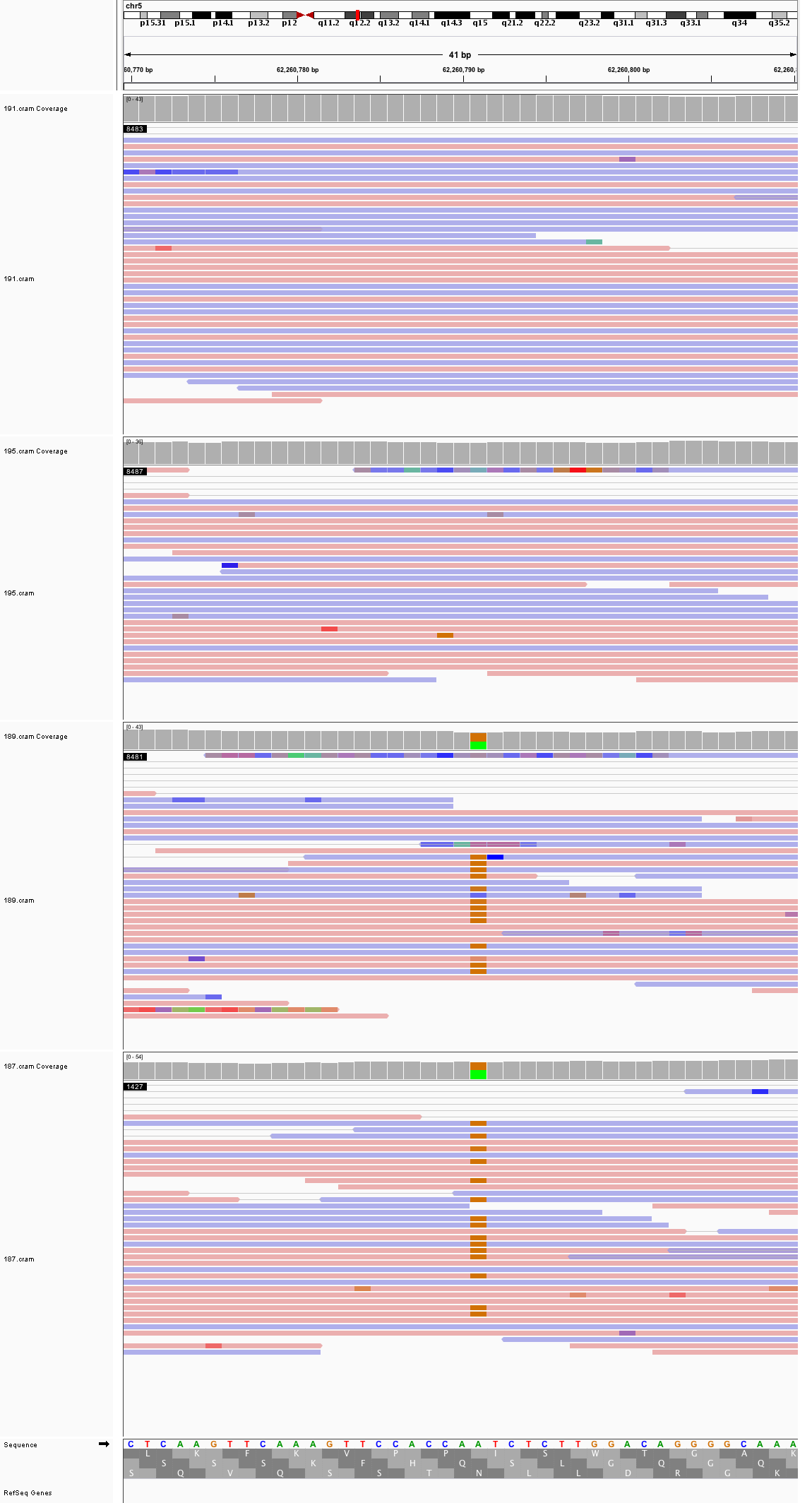

### chr5_78,664,059_78,664,099.png

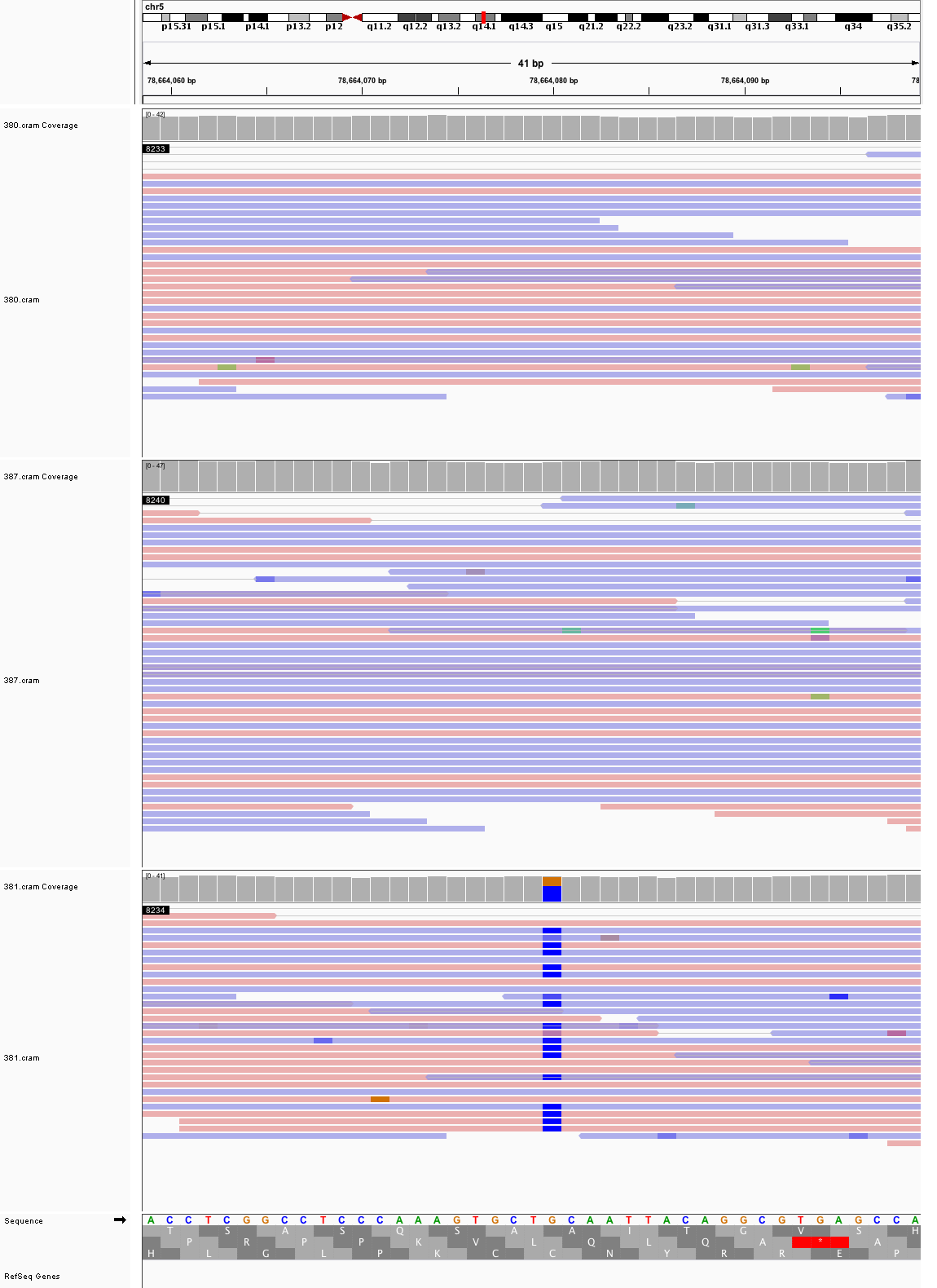

### chr5_113,268,636_113,268,676.png

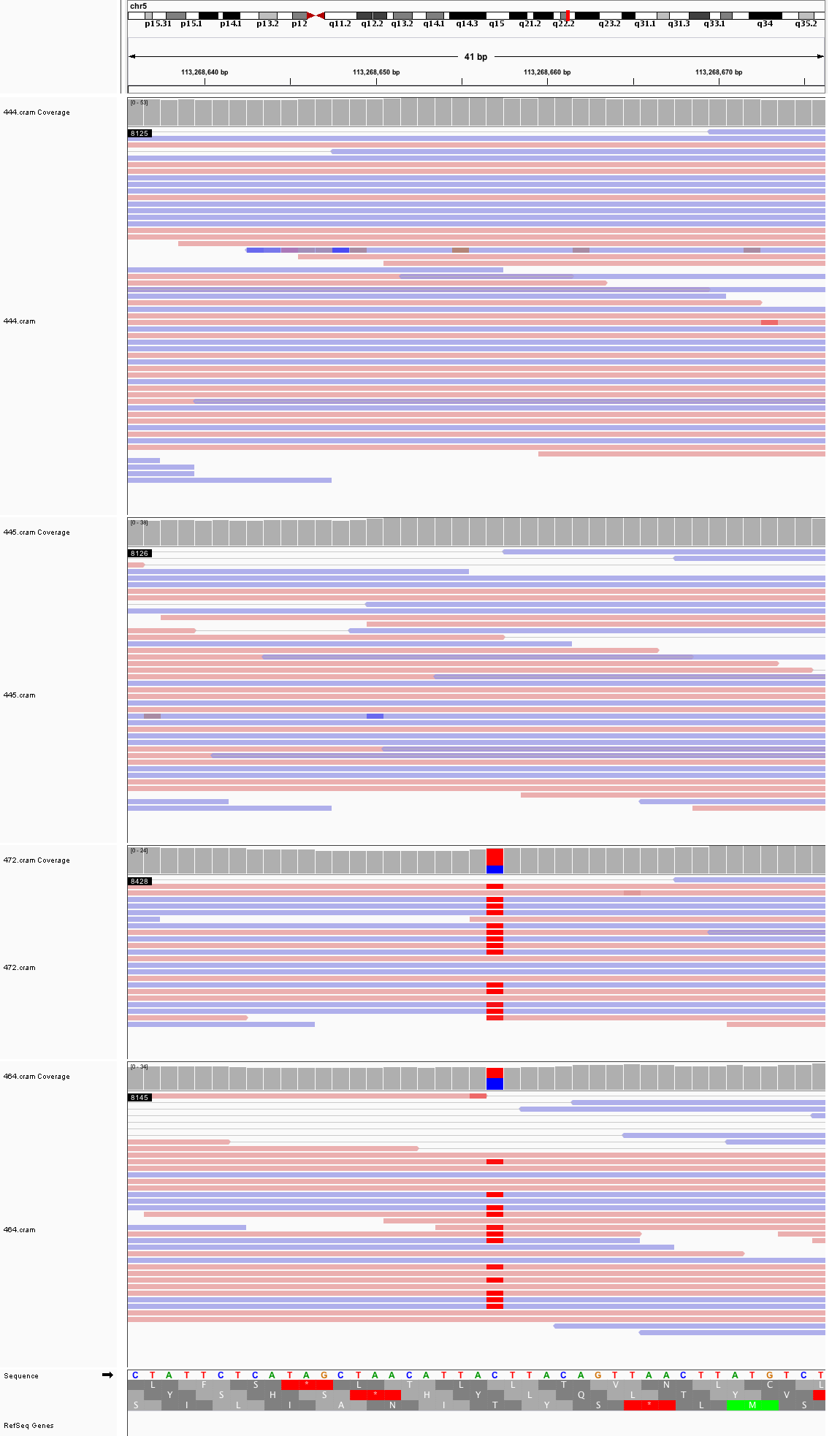

### chr5_144,293,597_144,293,637.png

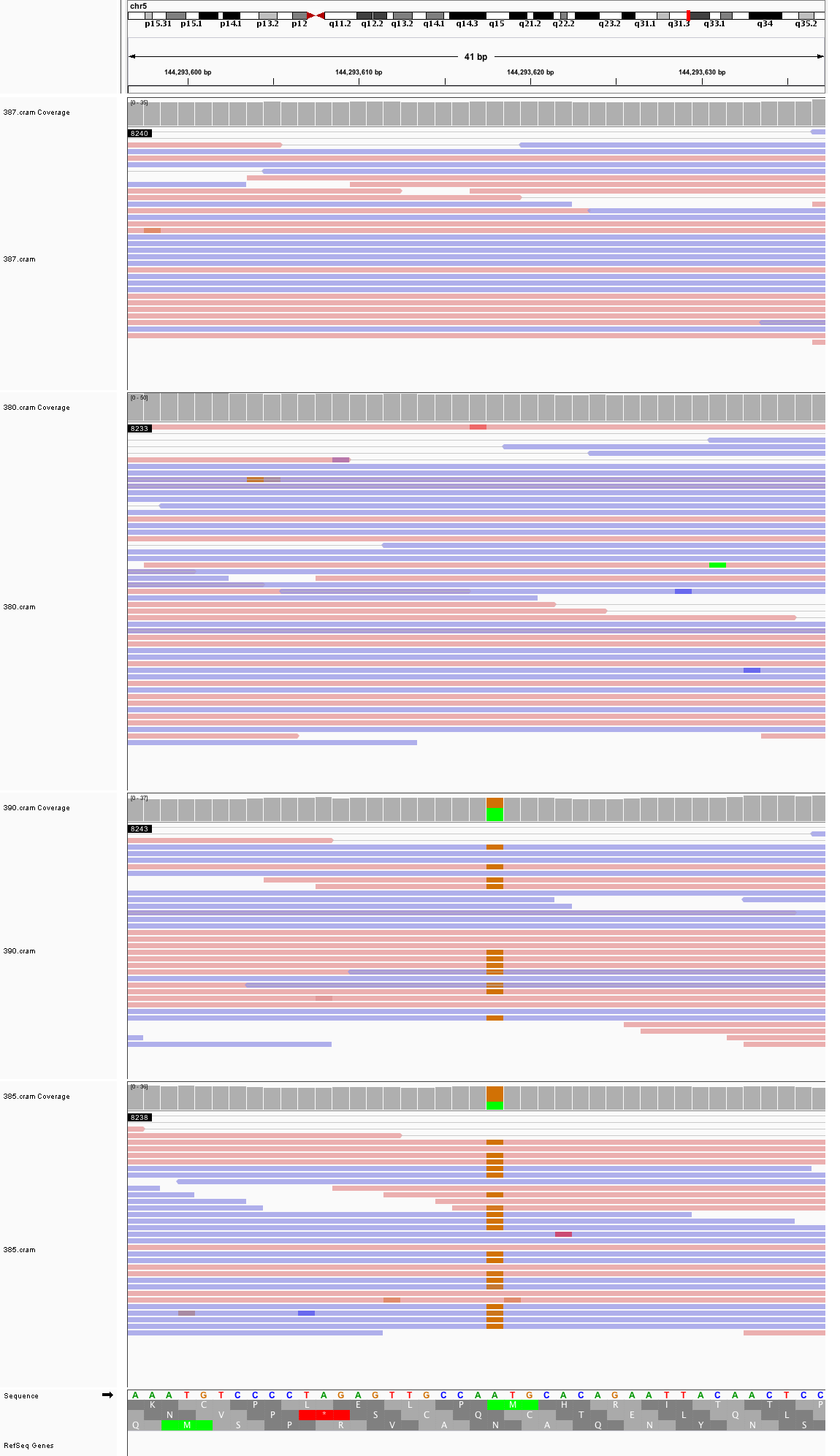

### chr6_8,107,118_8,107,158.png

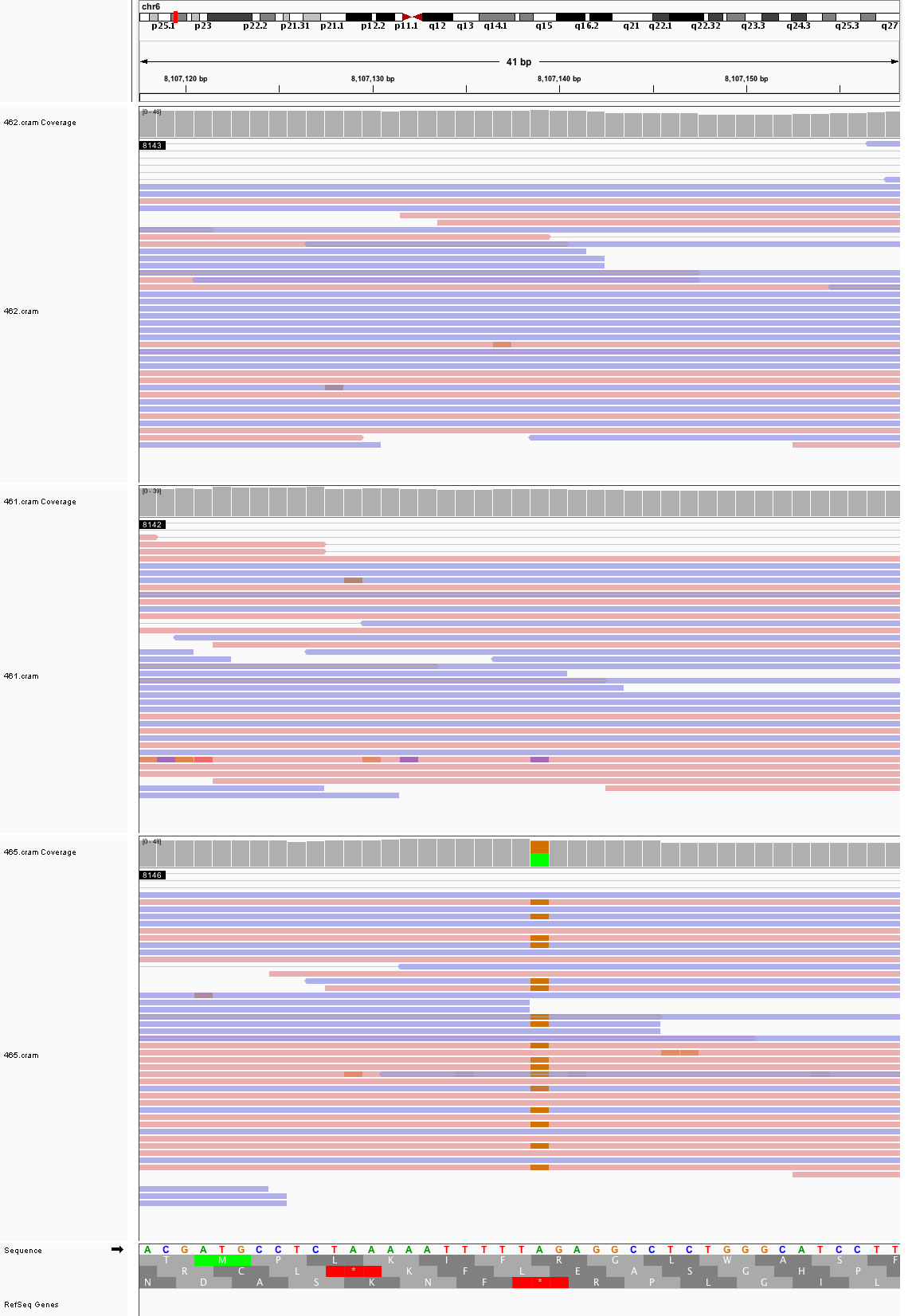

### chr6_10,538,448_10,538,488.png

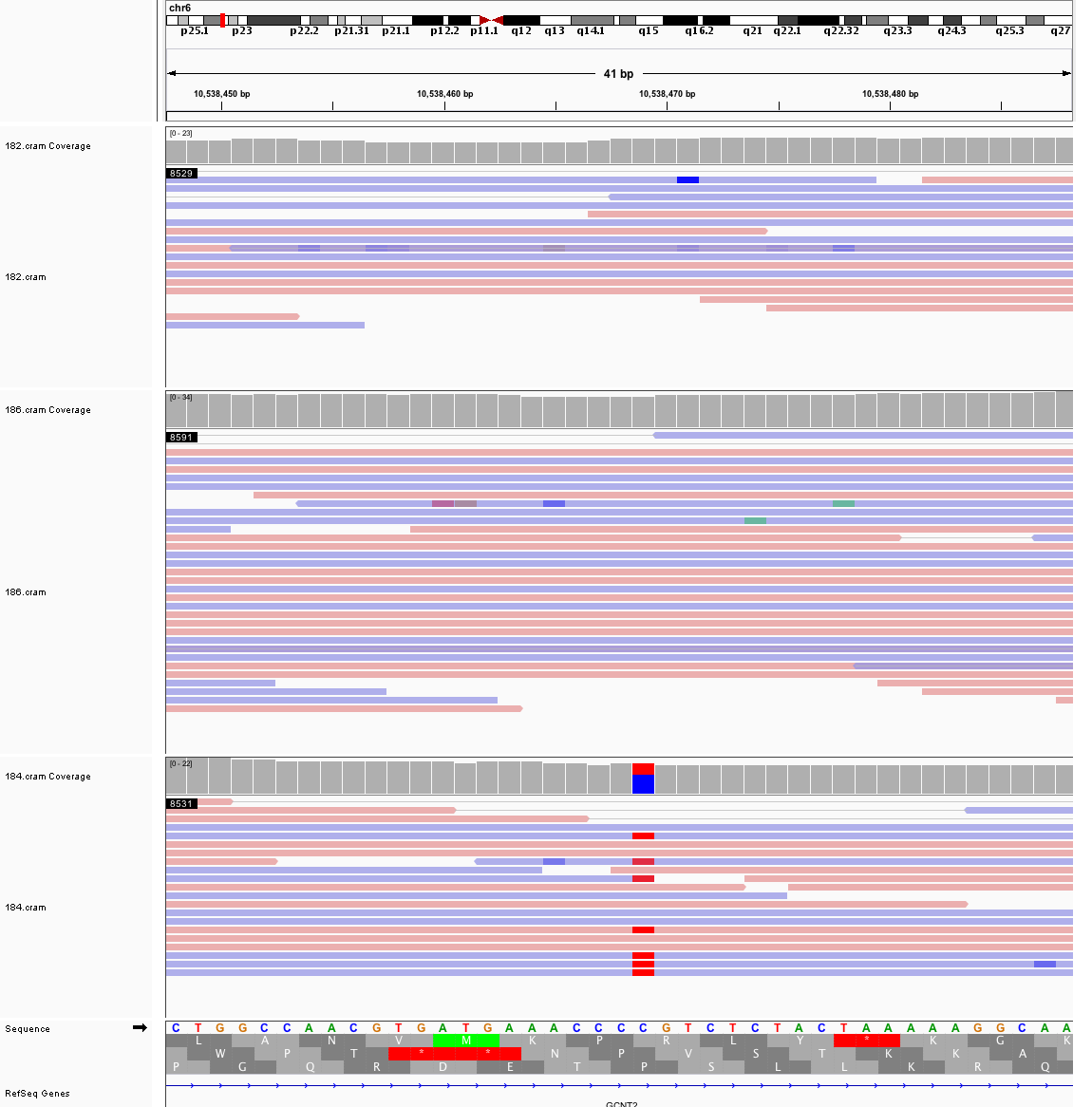

### chr6_16,418,007_16,418,047.png

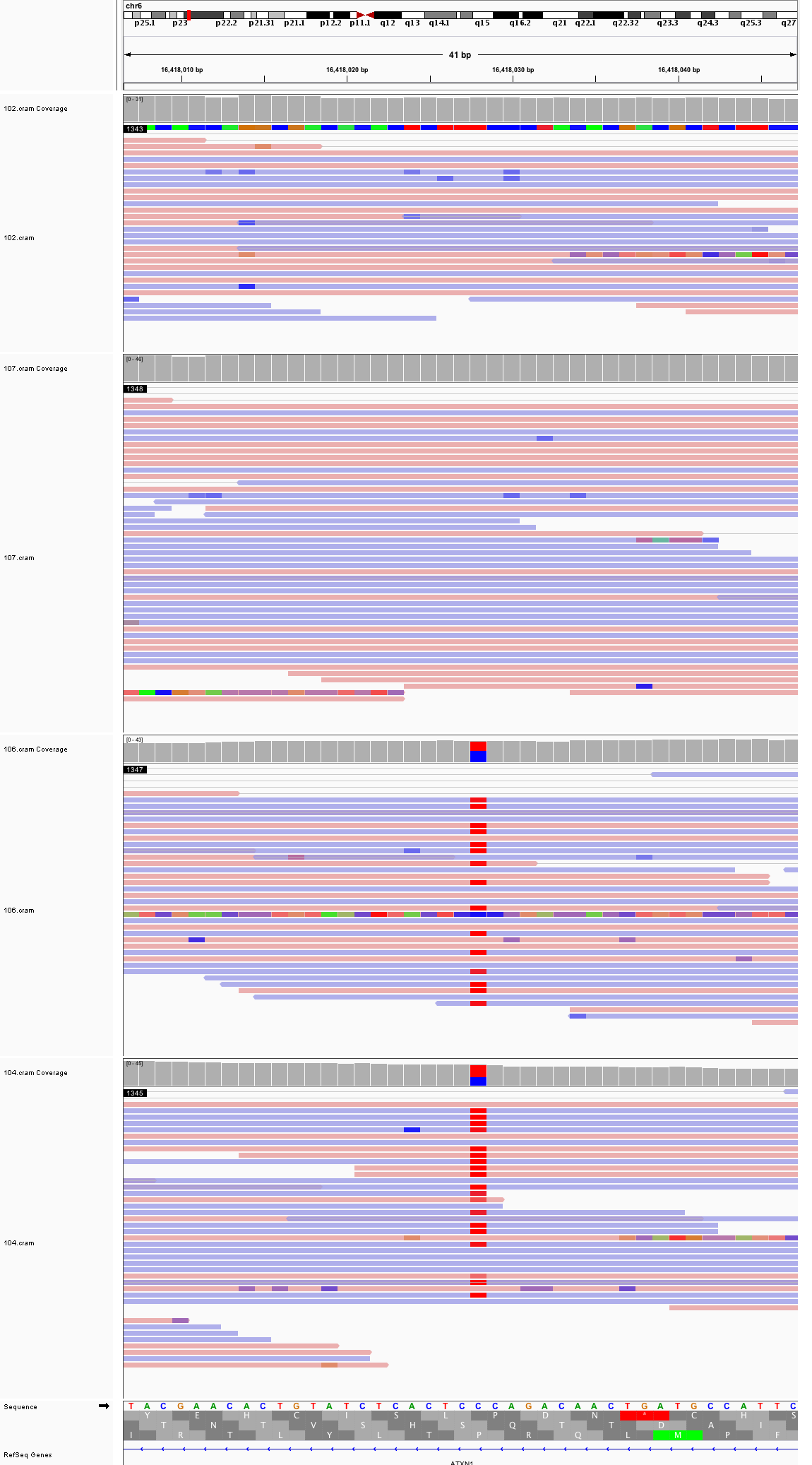

### chr6_62,711,131_62,711,171.png

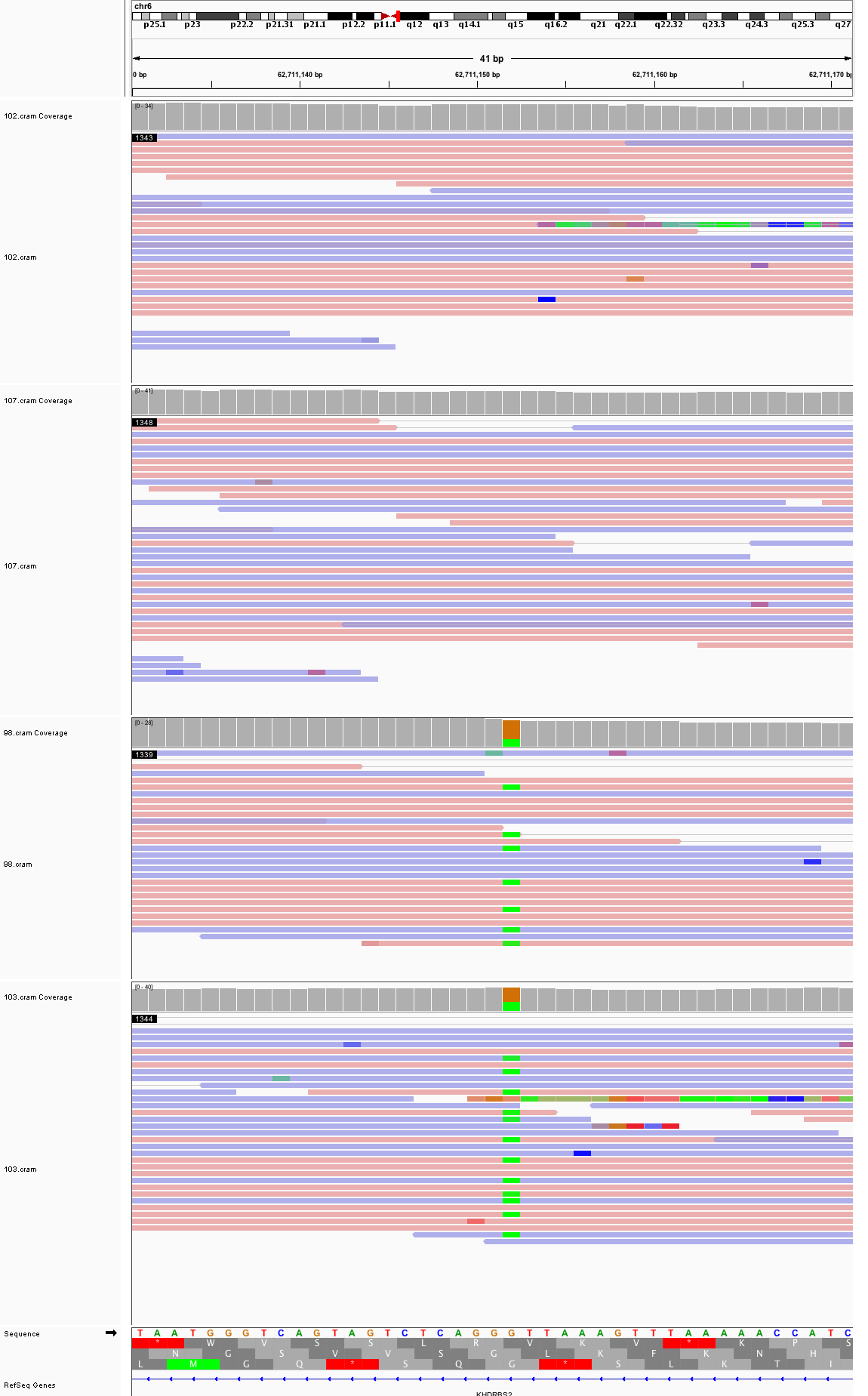

### chr6_69,804,468_69,804,508.png

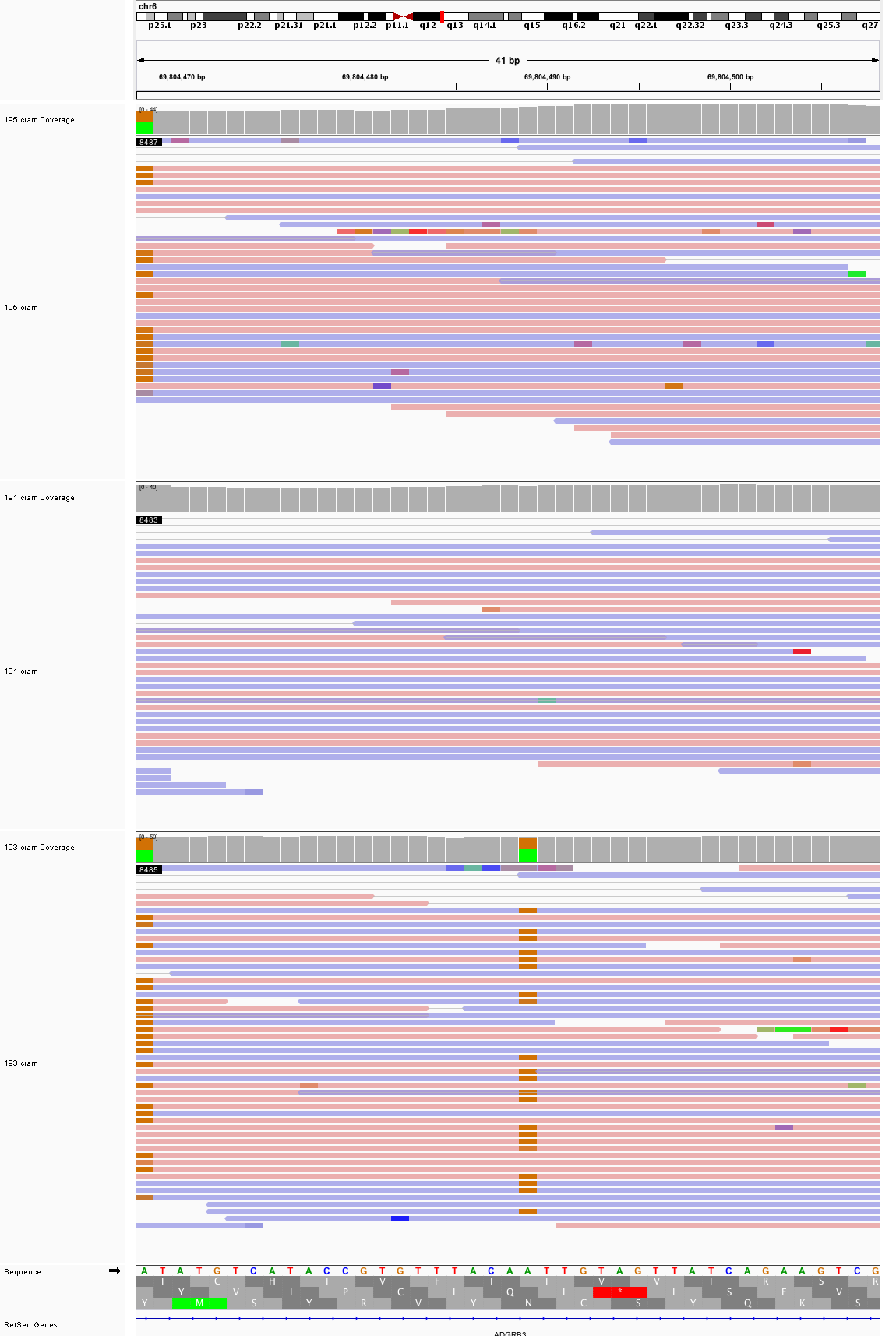

### chr7_61,790,870_61,790,910.png

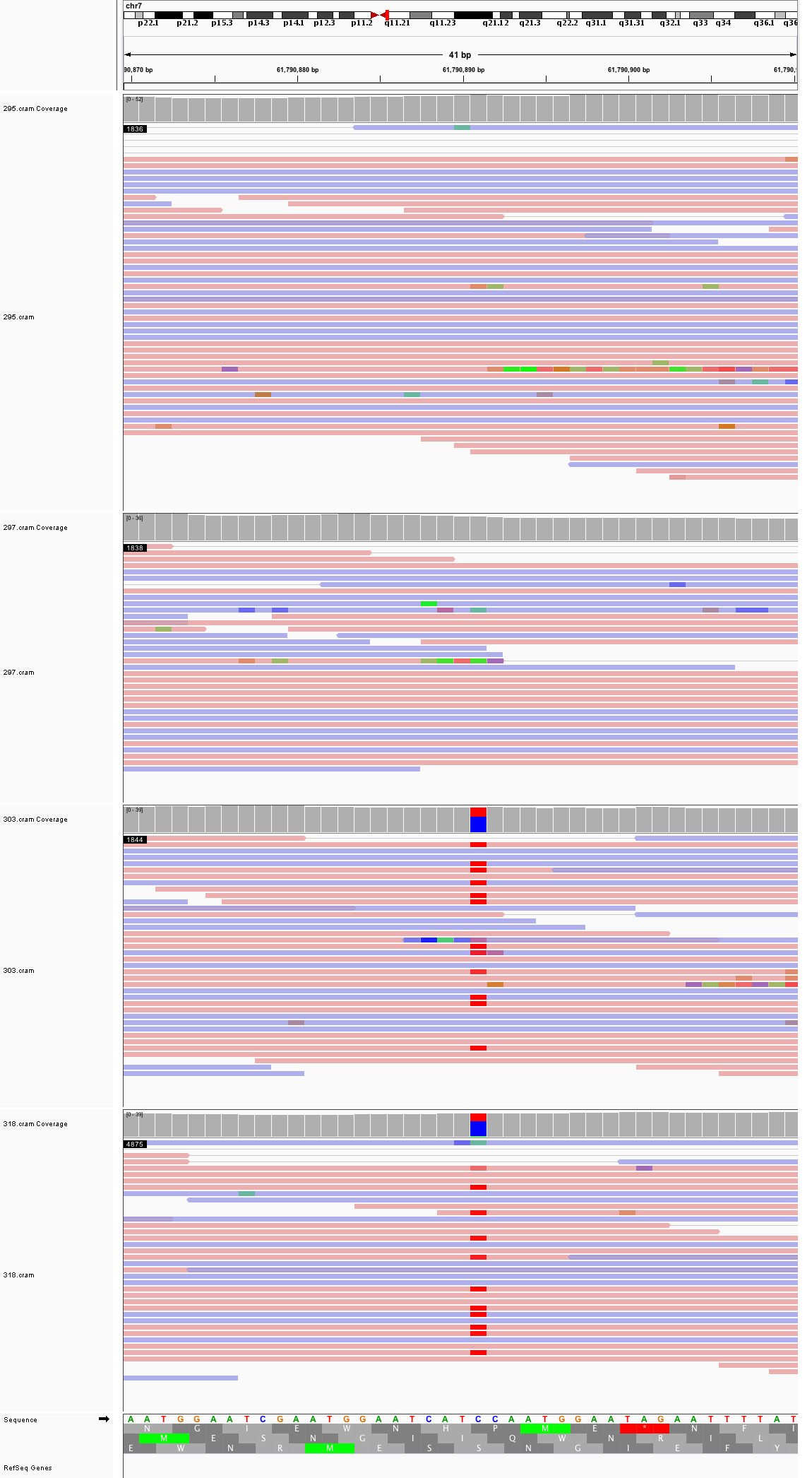

### chr7_63,781,278_63,781,318.png

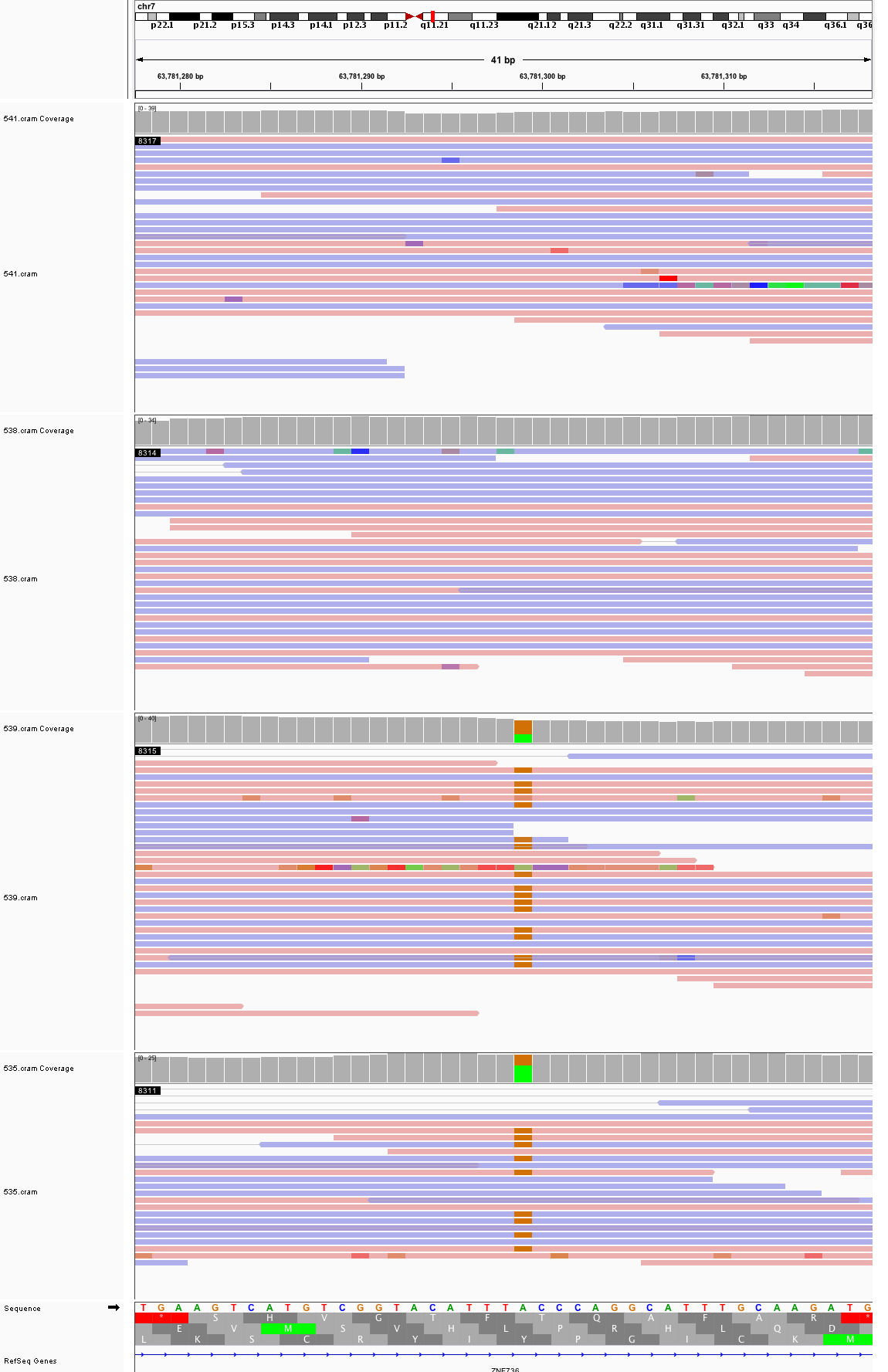

### chr7_87,718,209_87,718,249.png

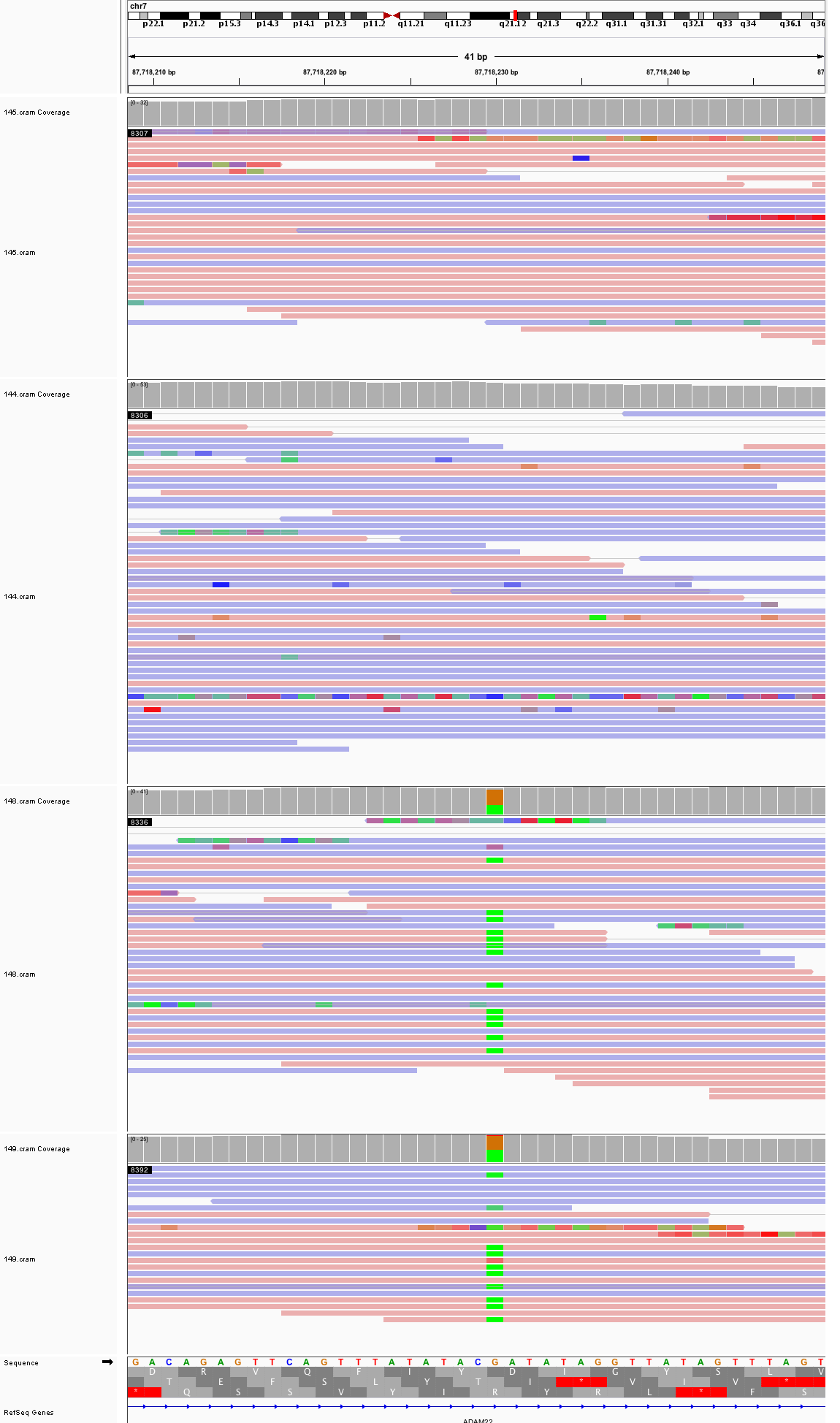

### chr7_102,115,414_102,115,454.png

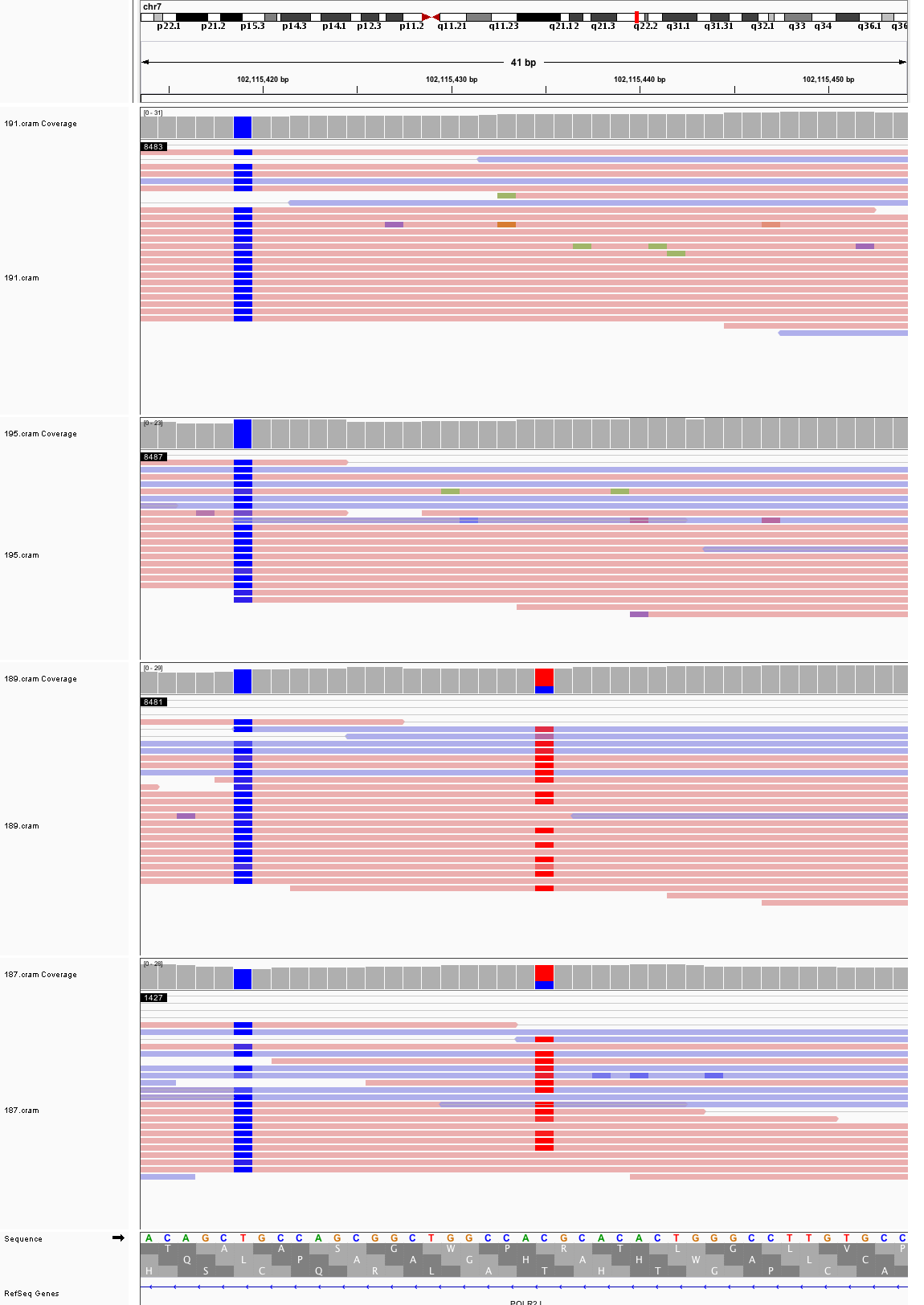

### chr7_154,513,219_154,513,259.png

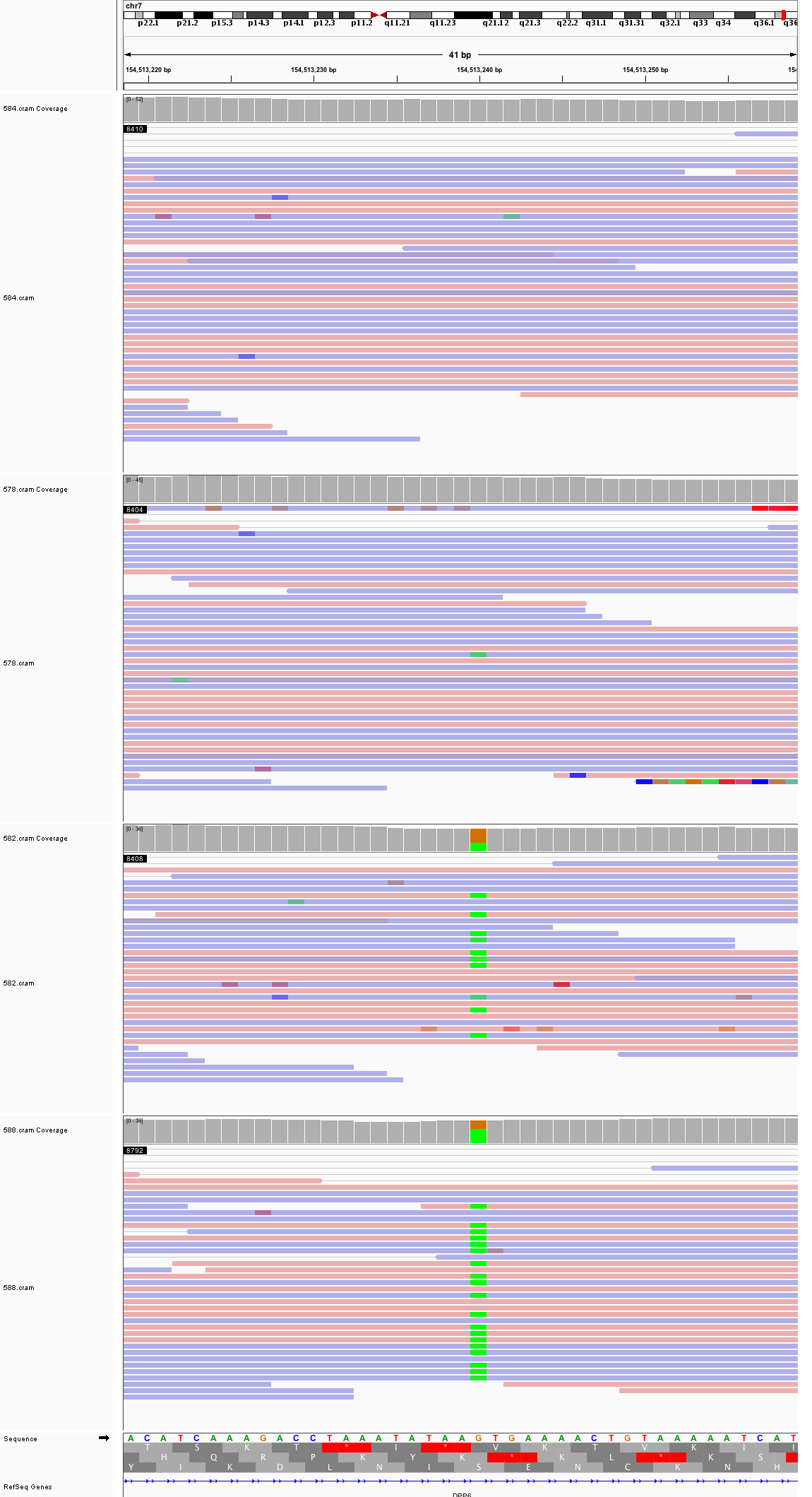

### chr8_4,406,373_4,406,413.png

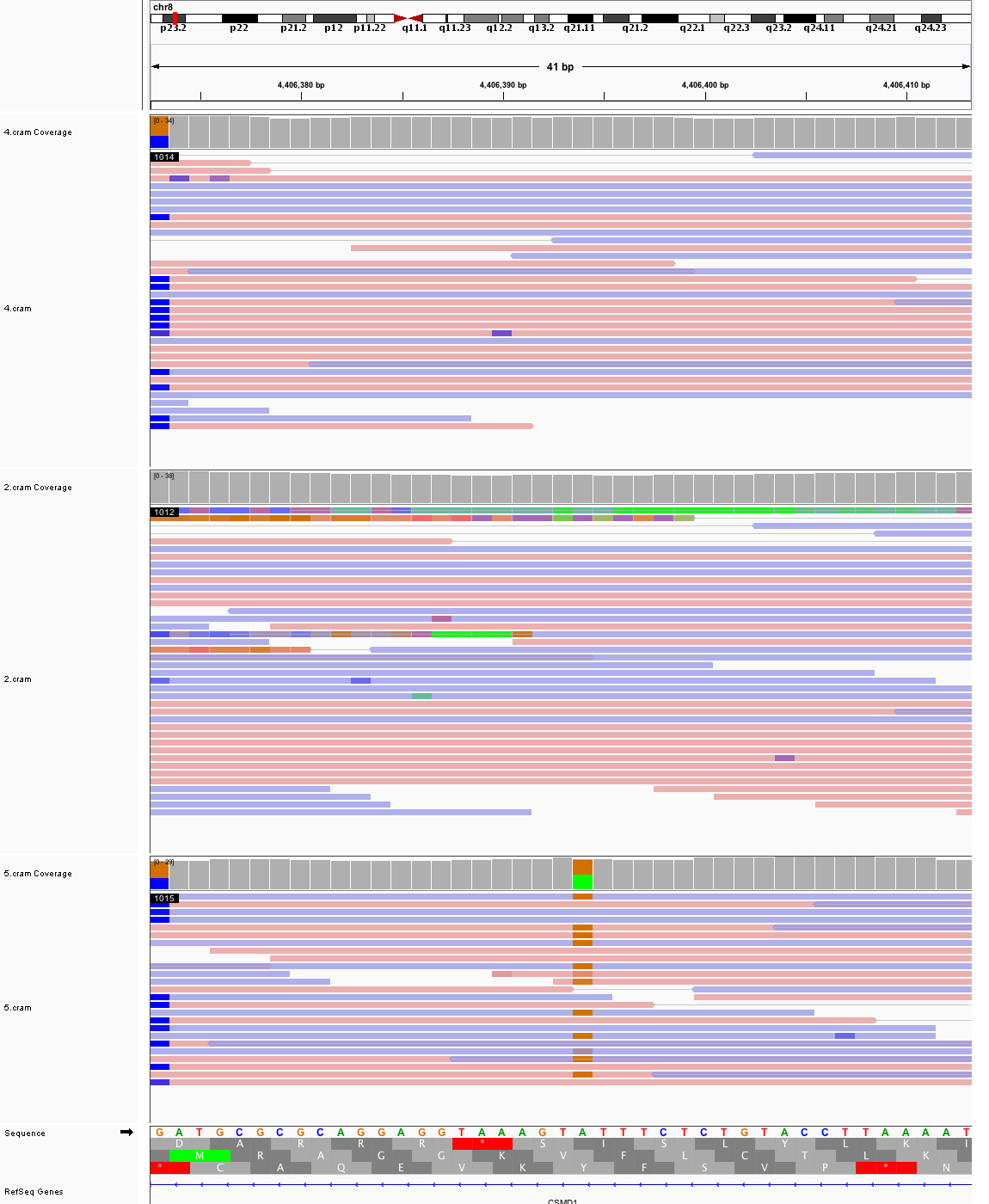

### chr8_19,257,448_19,257,488.png

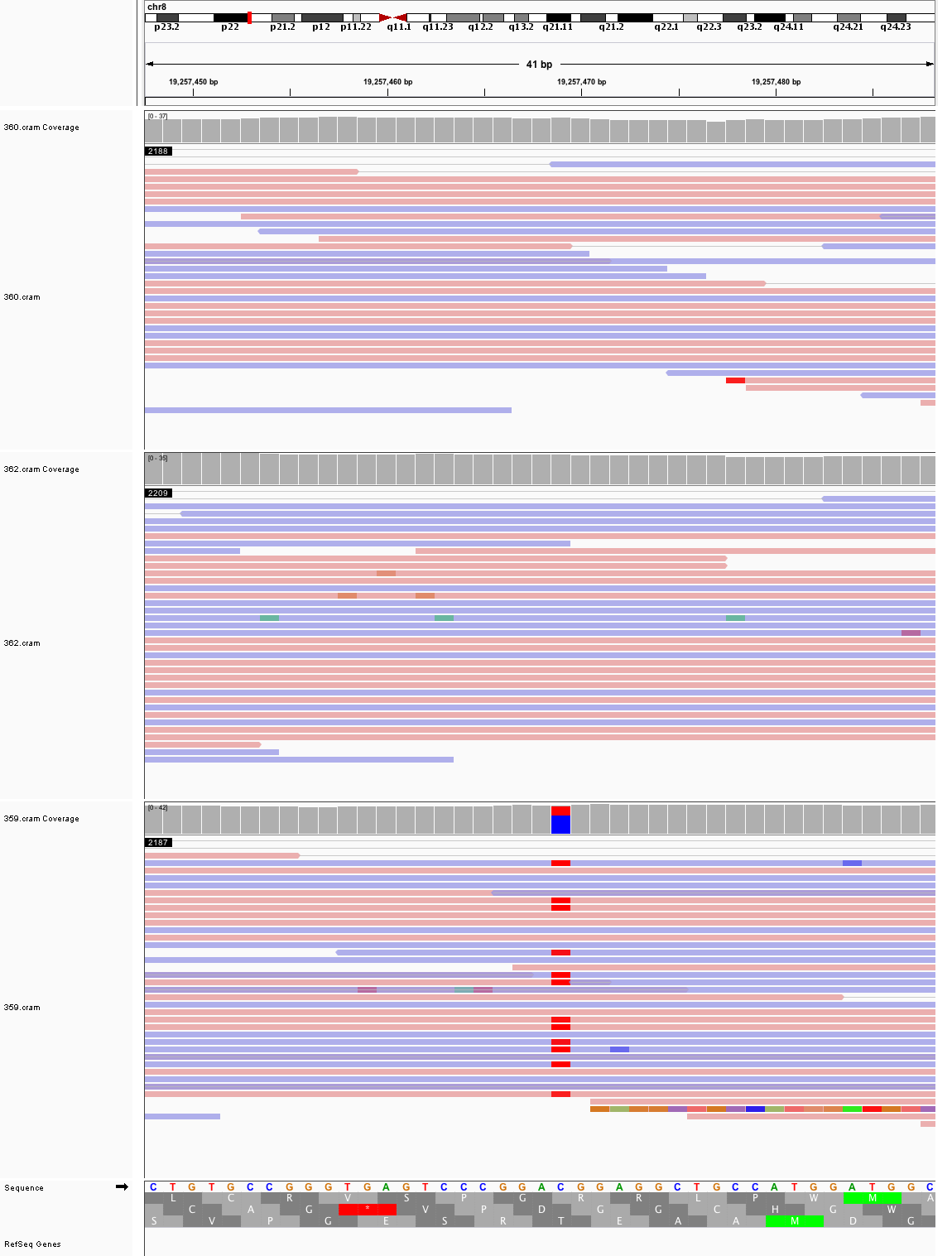

### chr8_31,597,535_31,597,575.png

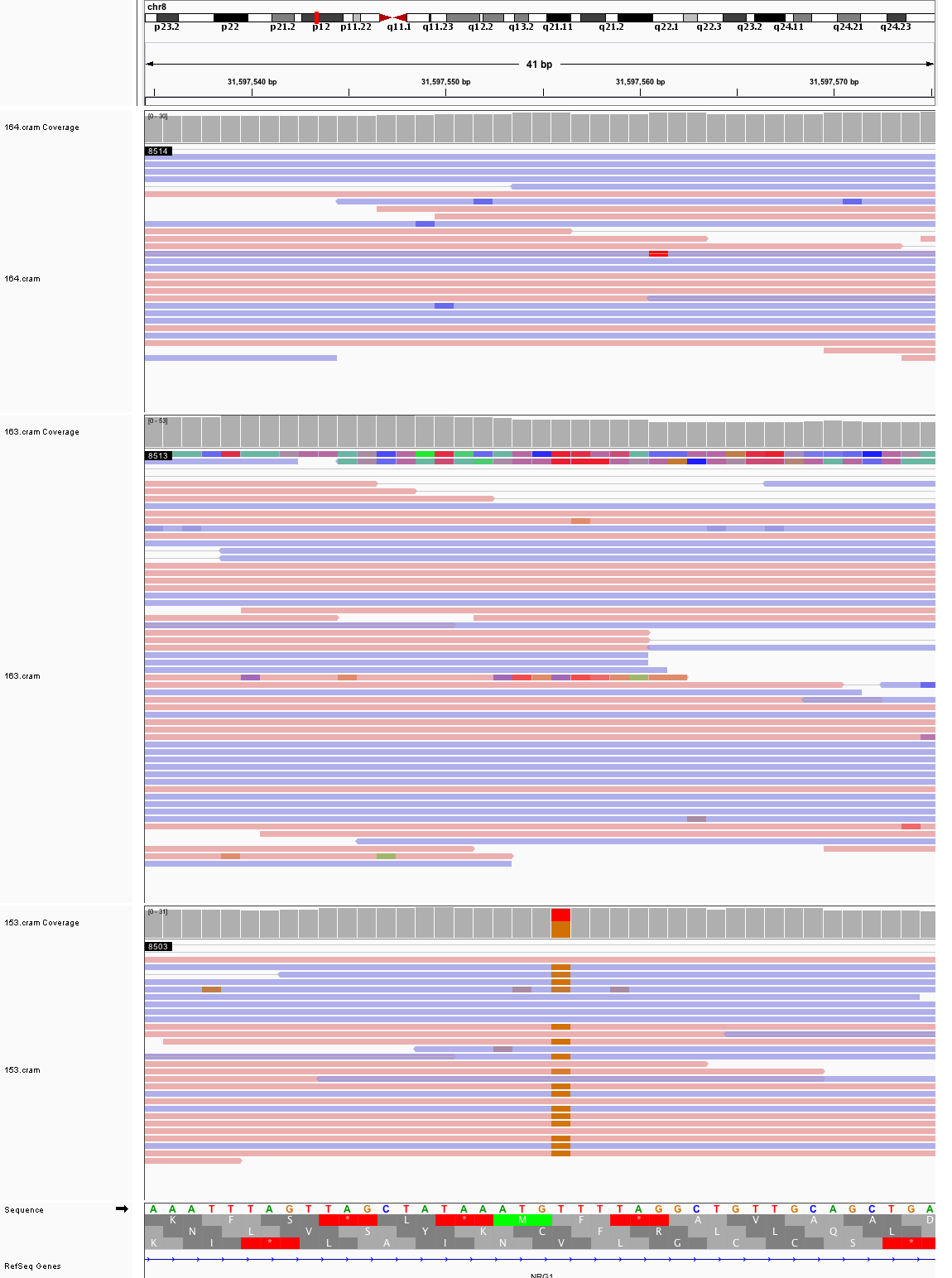

### chr8_43,704,348_43,704,388.png

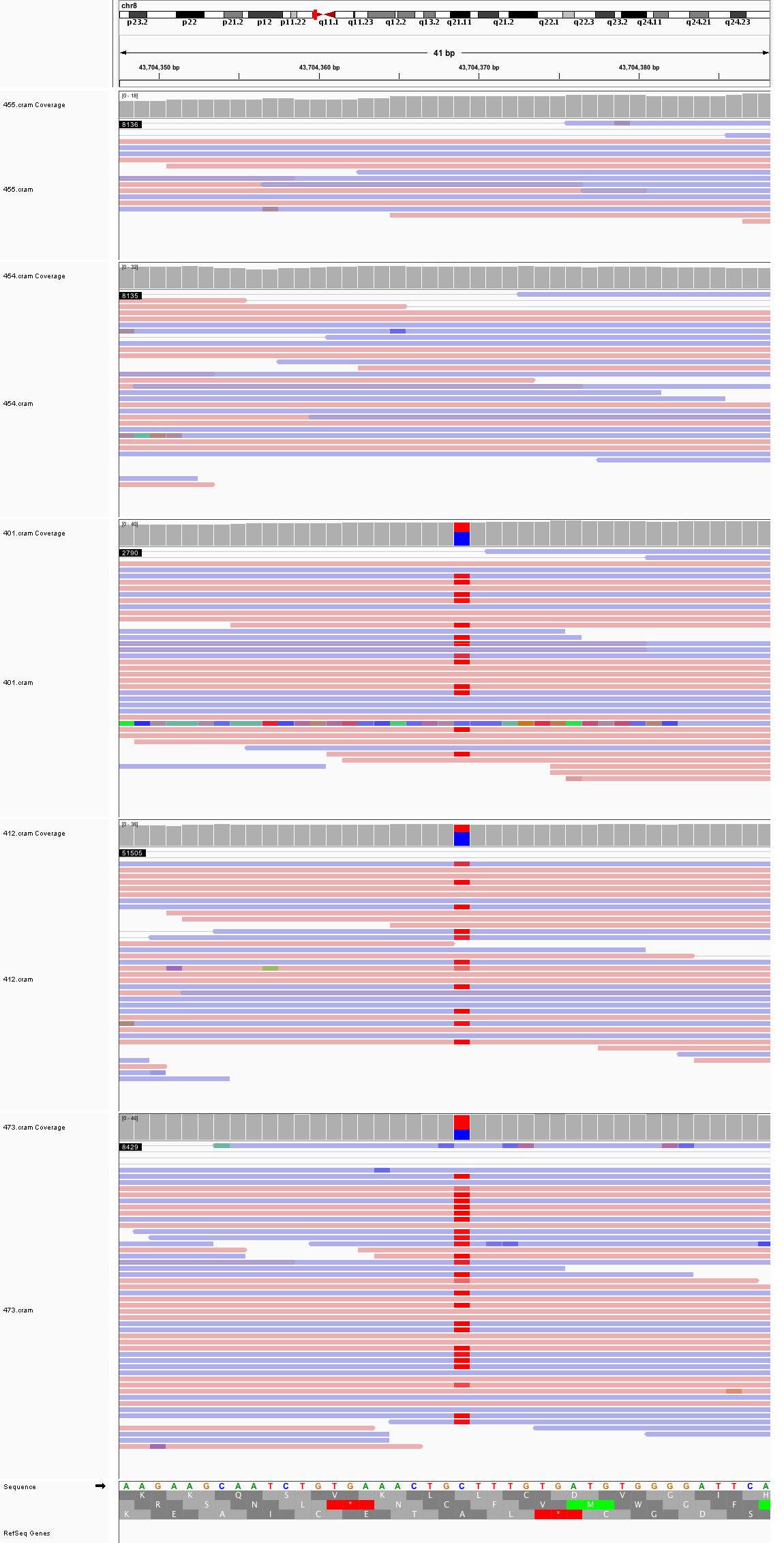

### chr8_50,812,575_50,812,615.png

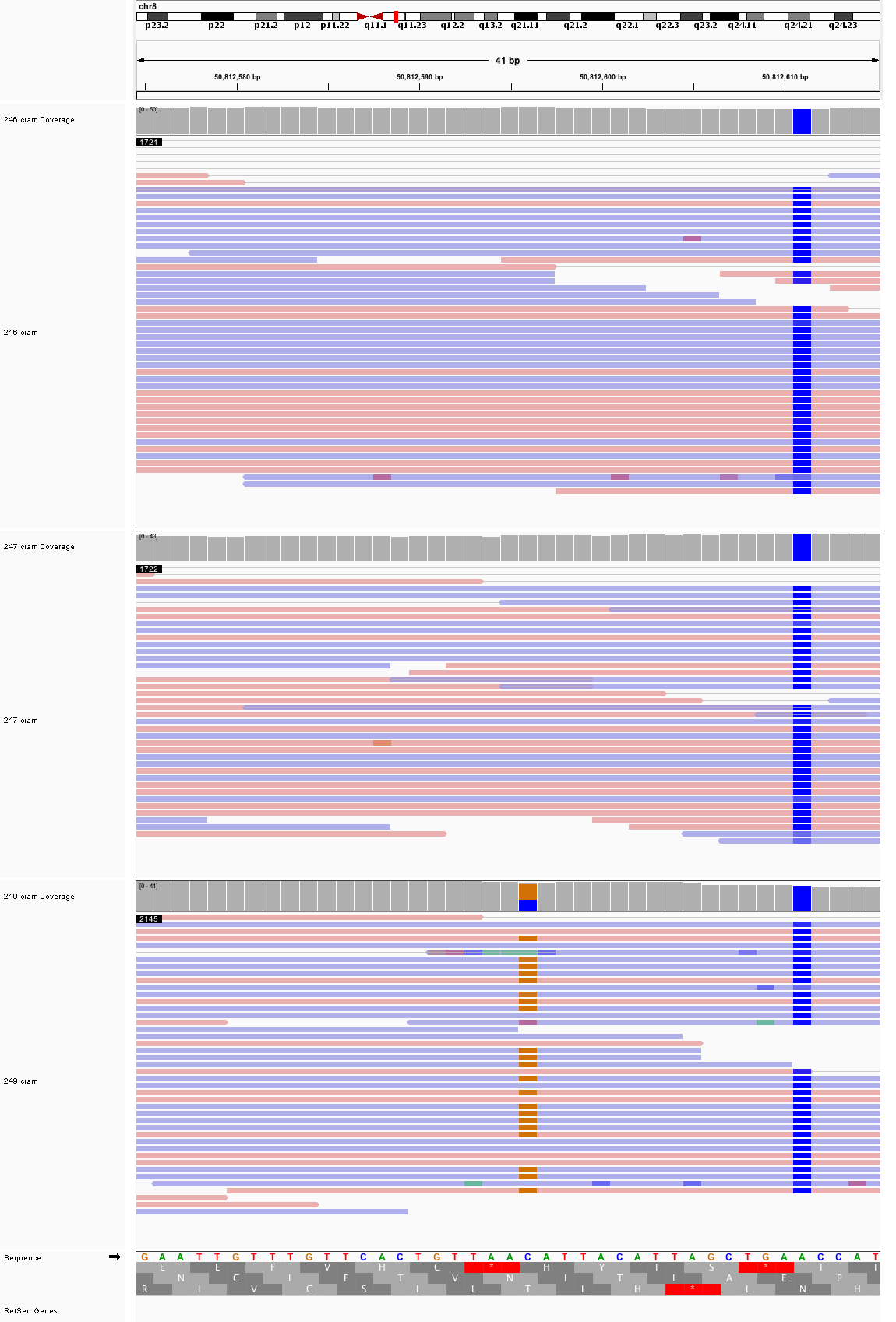

### chr8_60,766,017_60,766,057.png

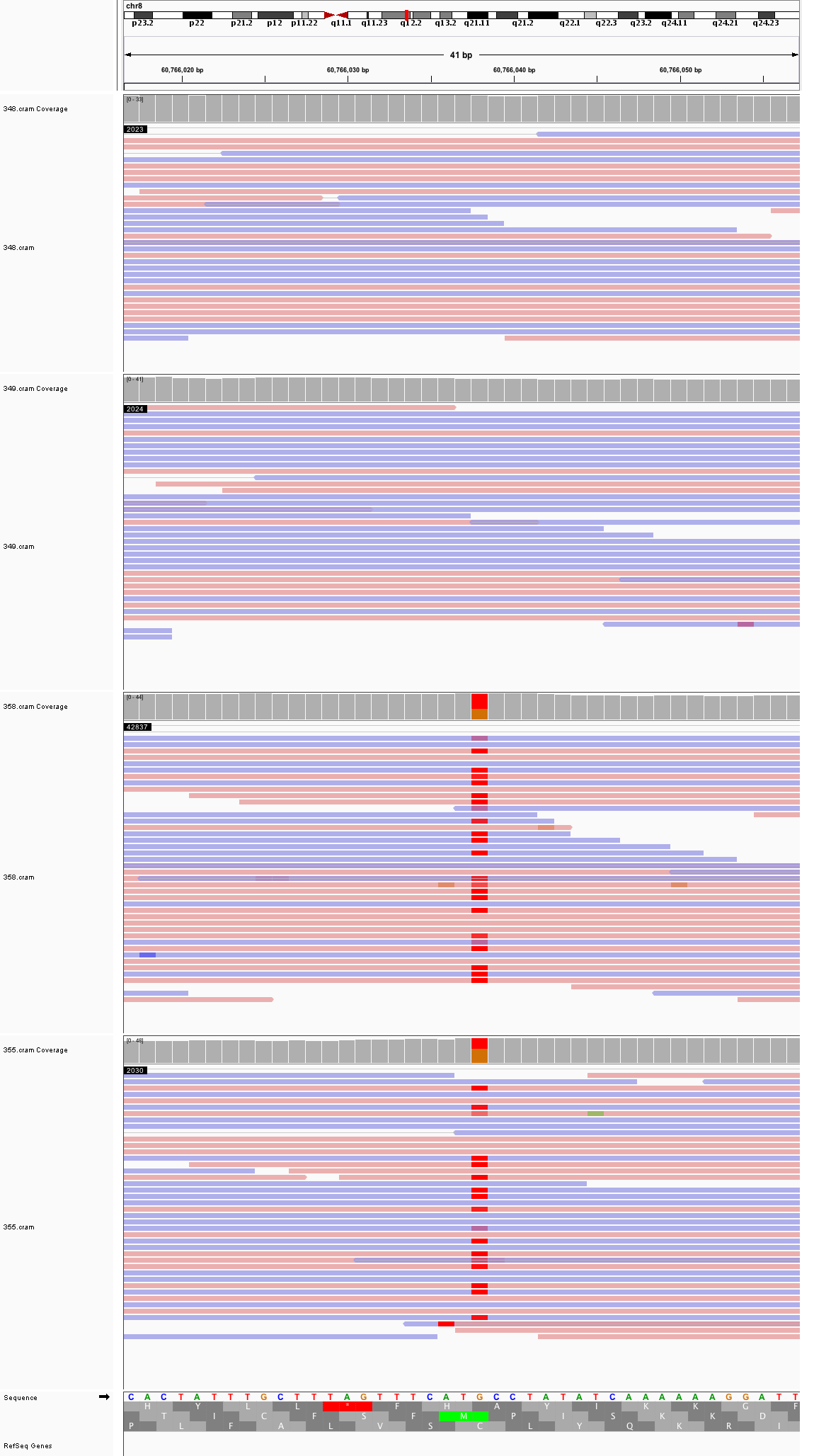

### chr8_116,140,002_116,140,042.png

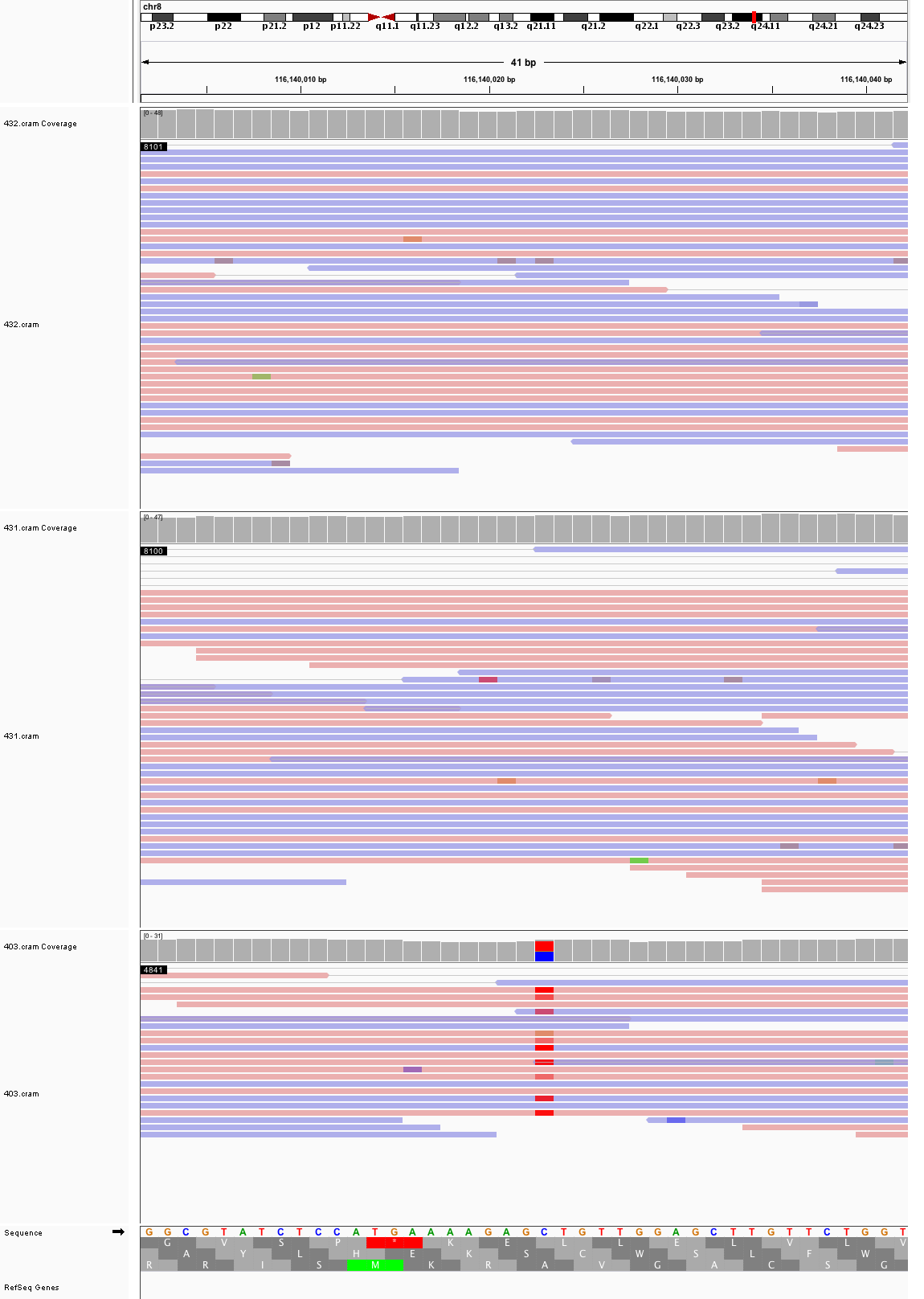

### chr8_118,642,818_118,642,858.png

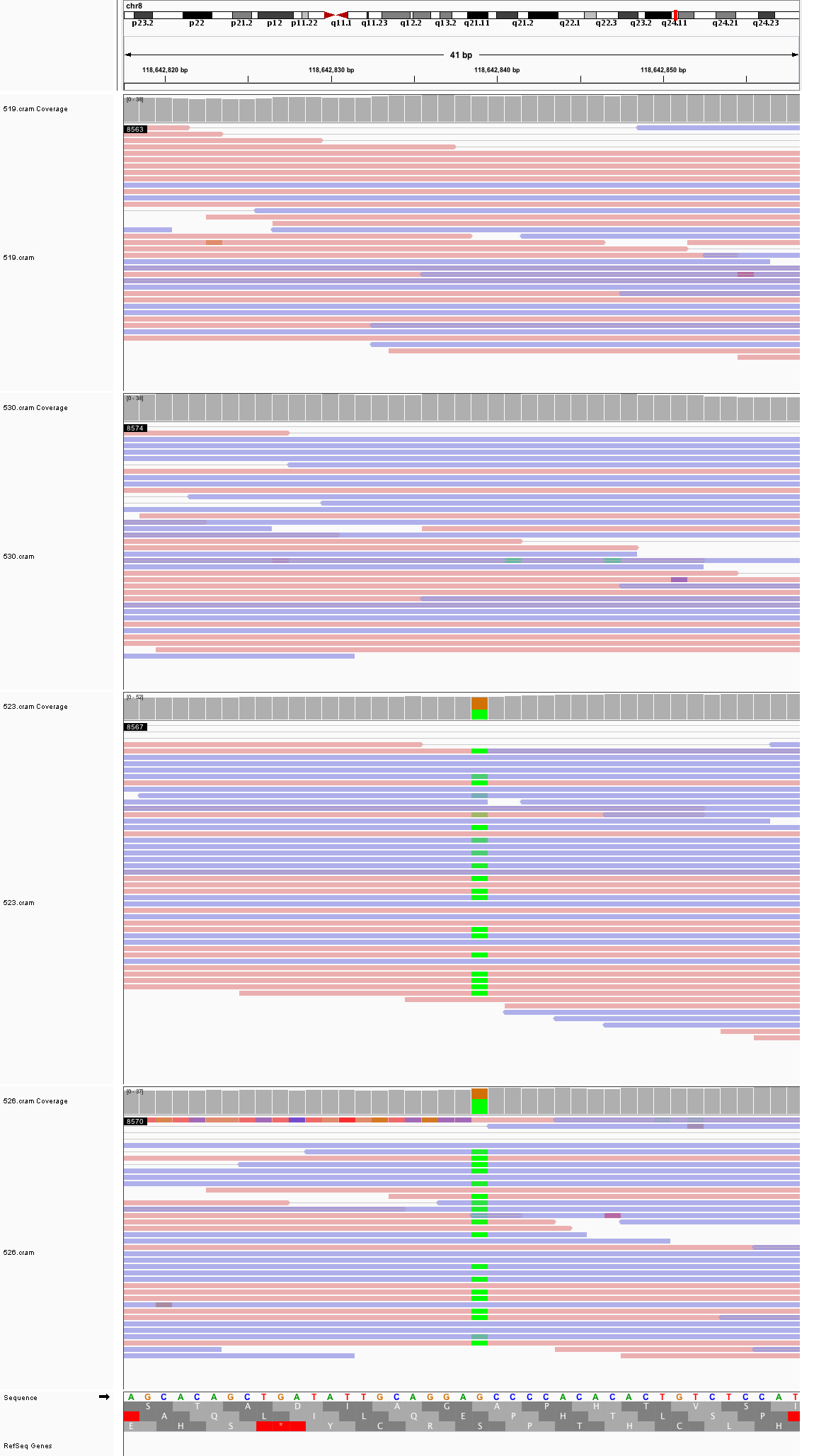

### chr8_143,322,450_143,322,490.png

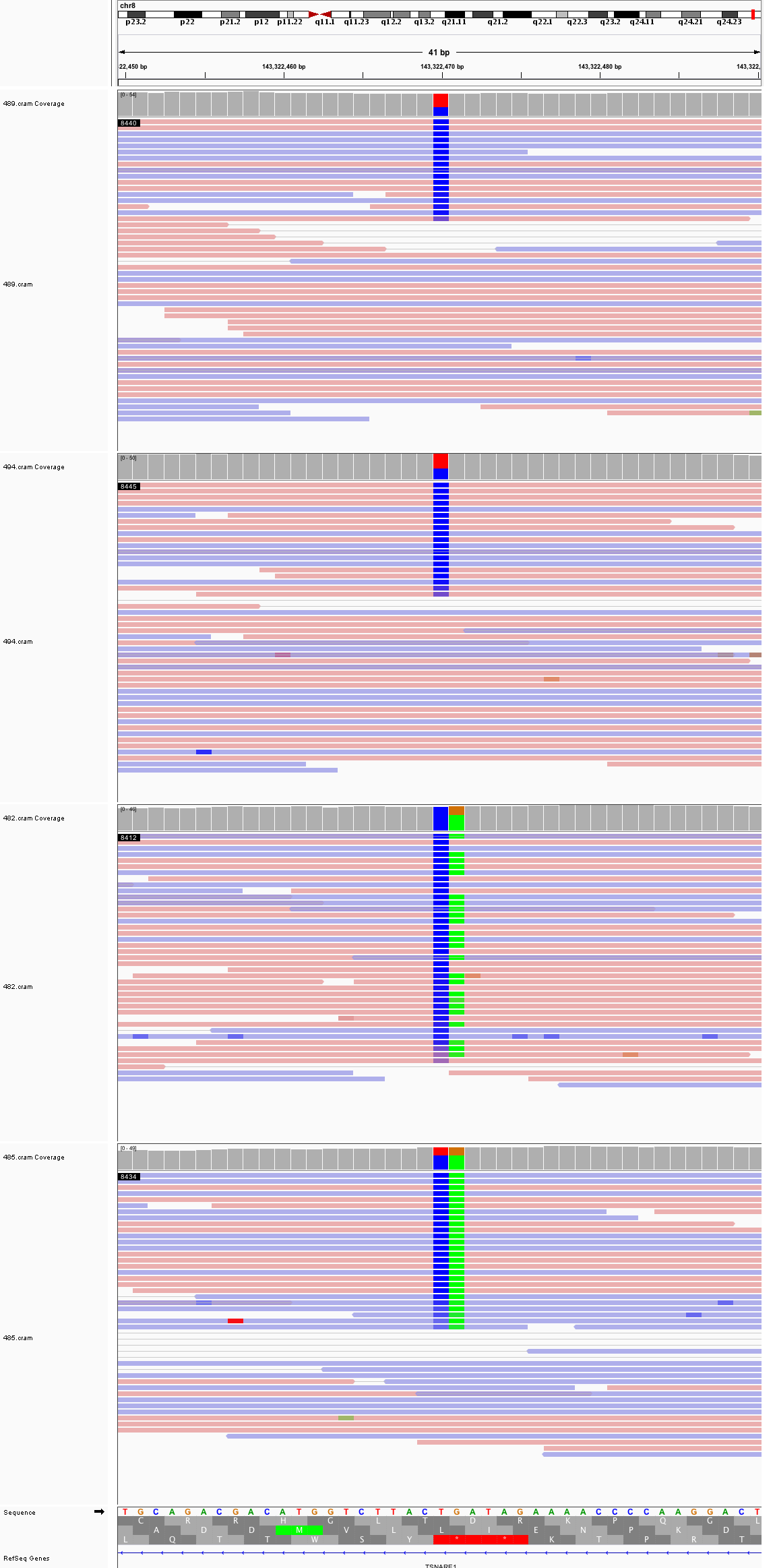

### chr9_3,813,283_3,813,323.png

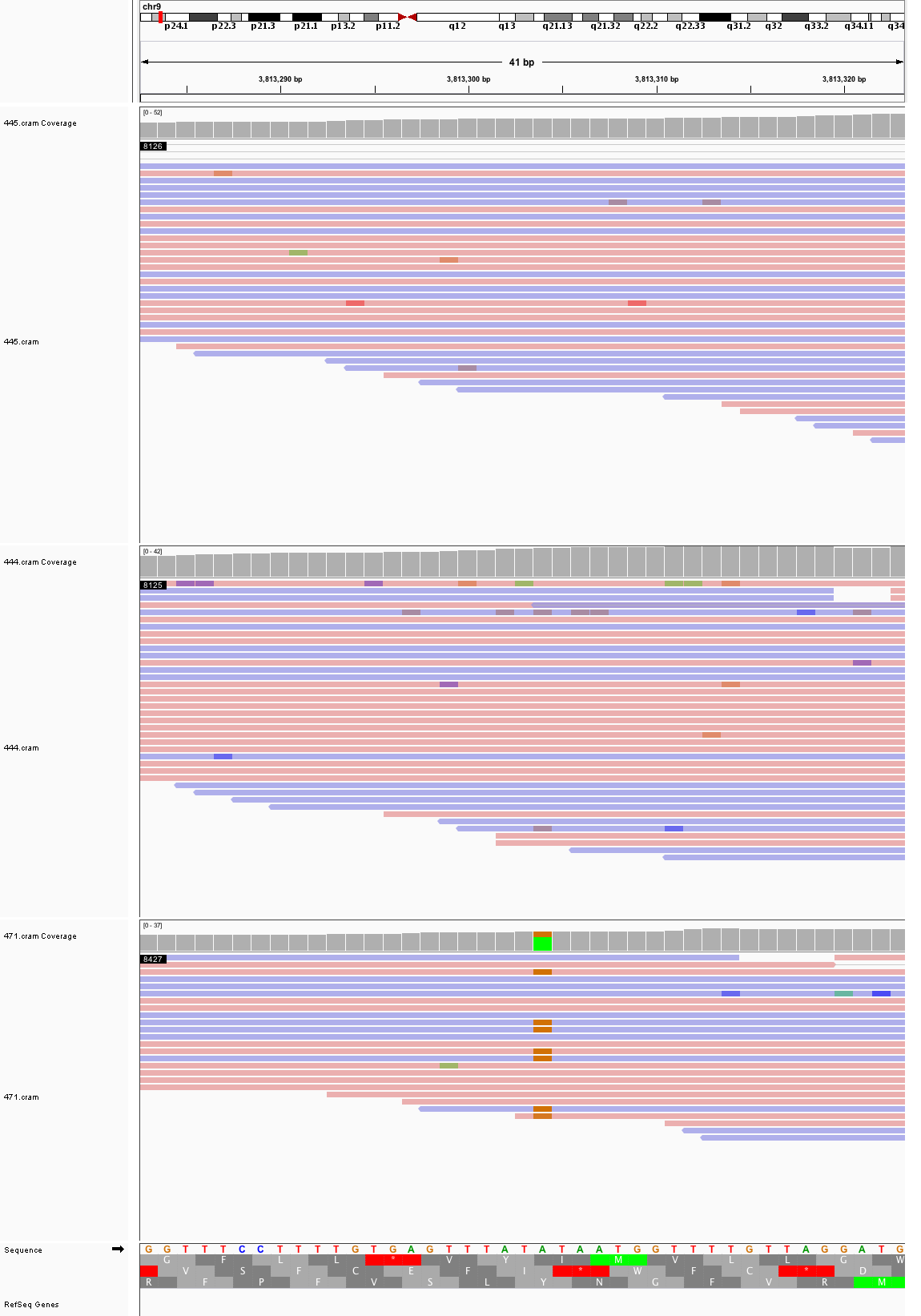

### chr9_44,363,780_44,363,820.png

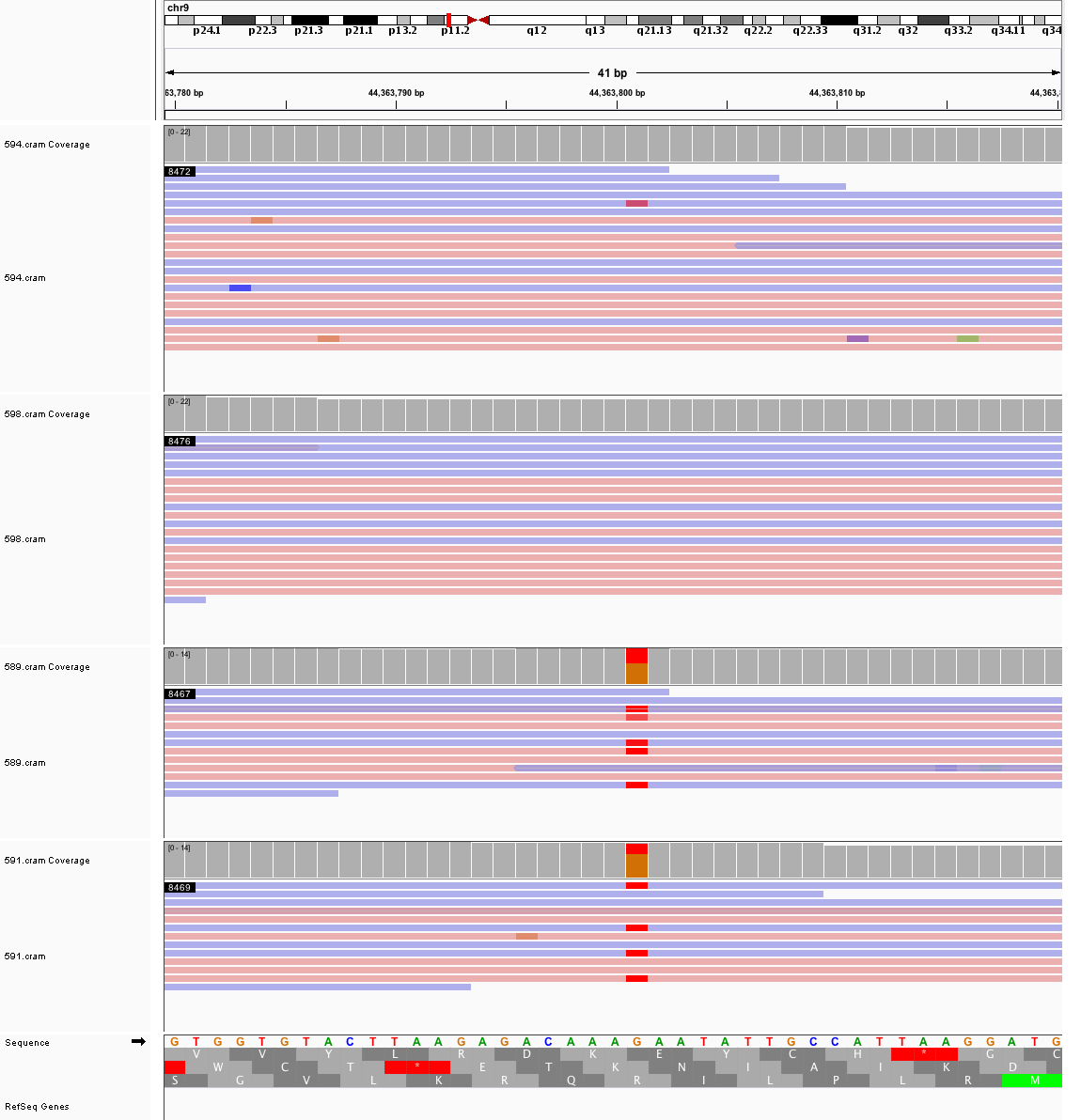

### chr9_80,274,029_80,274,069.png

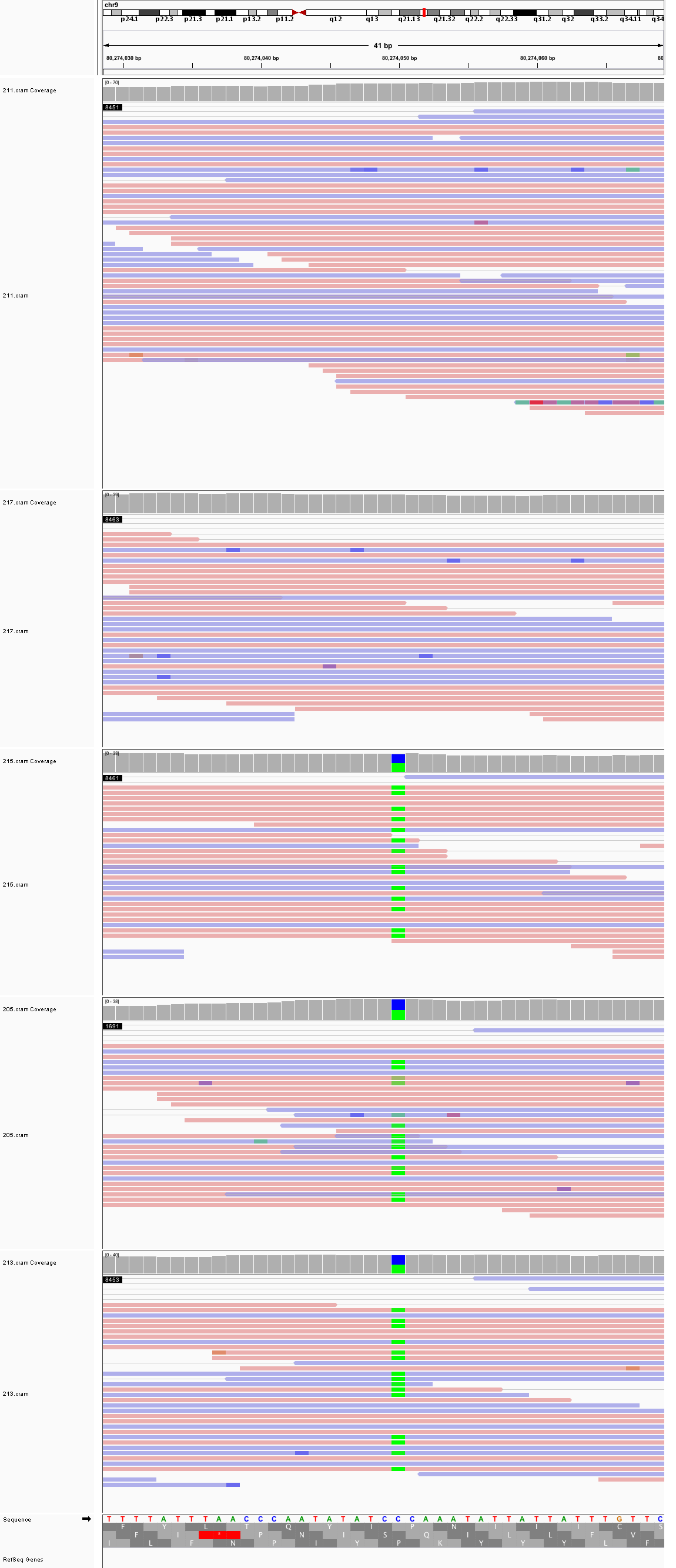

### chr9_106,254,255_106,254,295.png

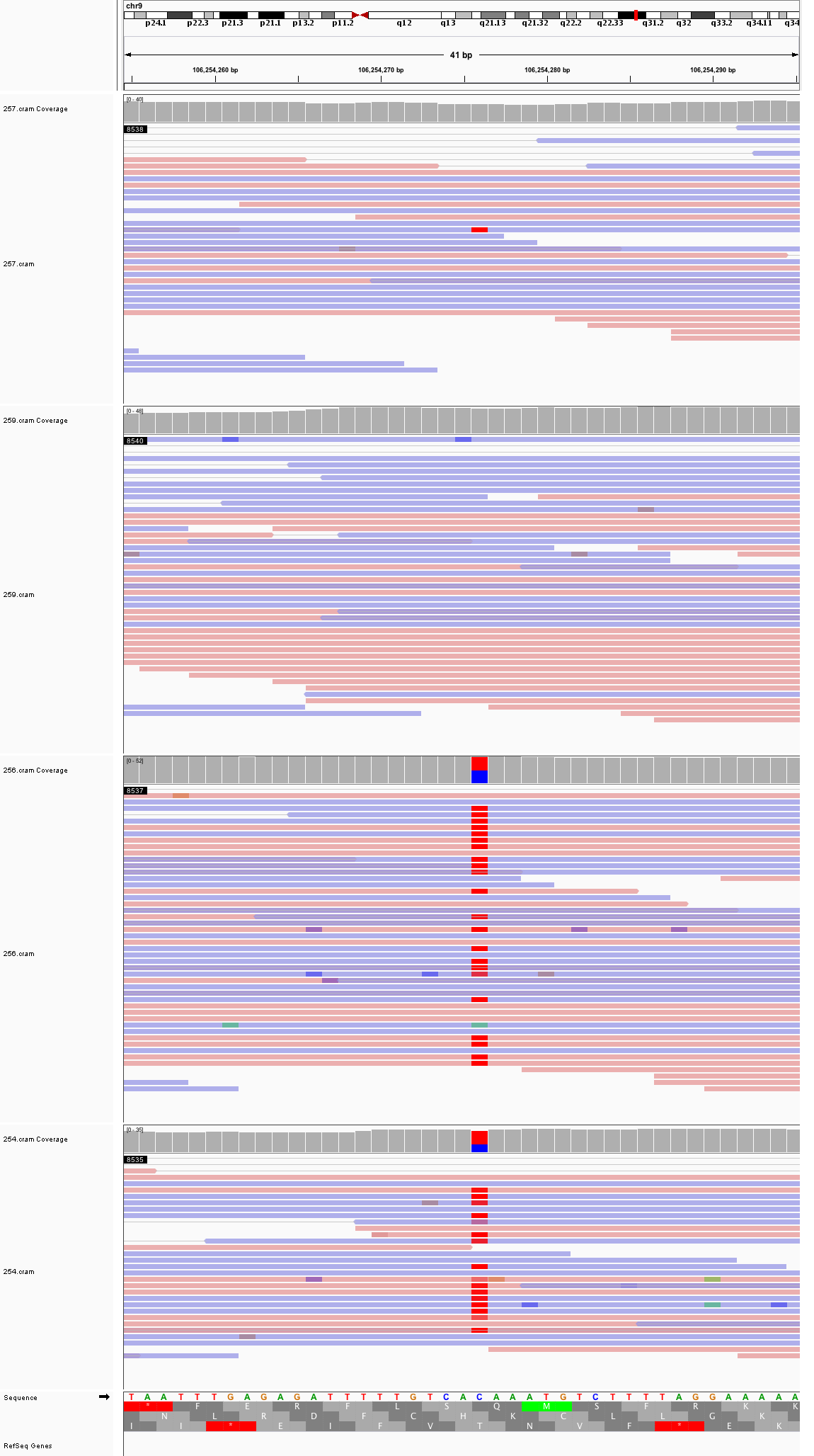

### chr9_116,204,746_116,204,786.png

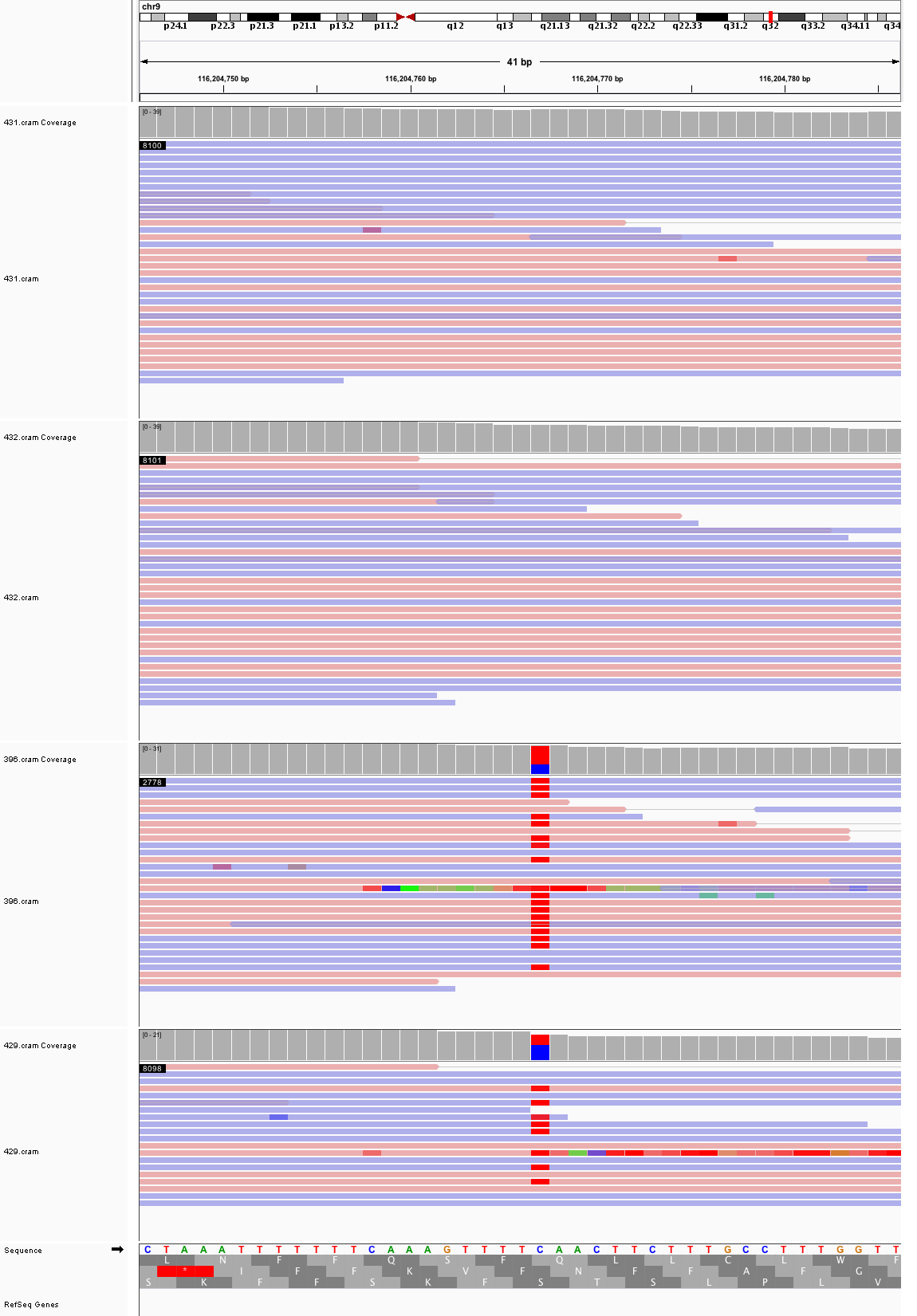

### chr9_127,680,788_127,680,828.png

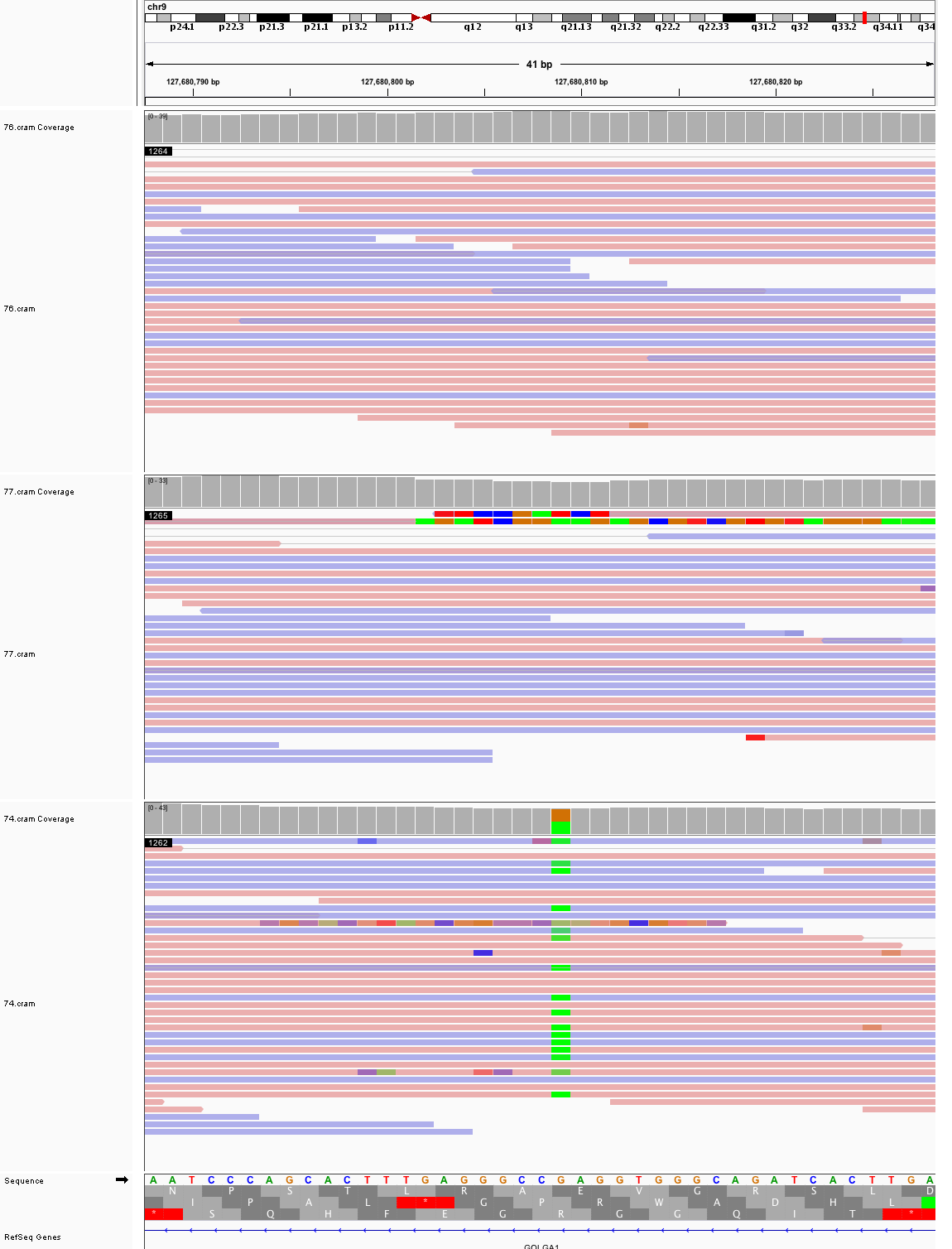

### chr10_33,554,850_33,554,890.png

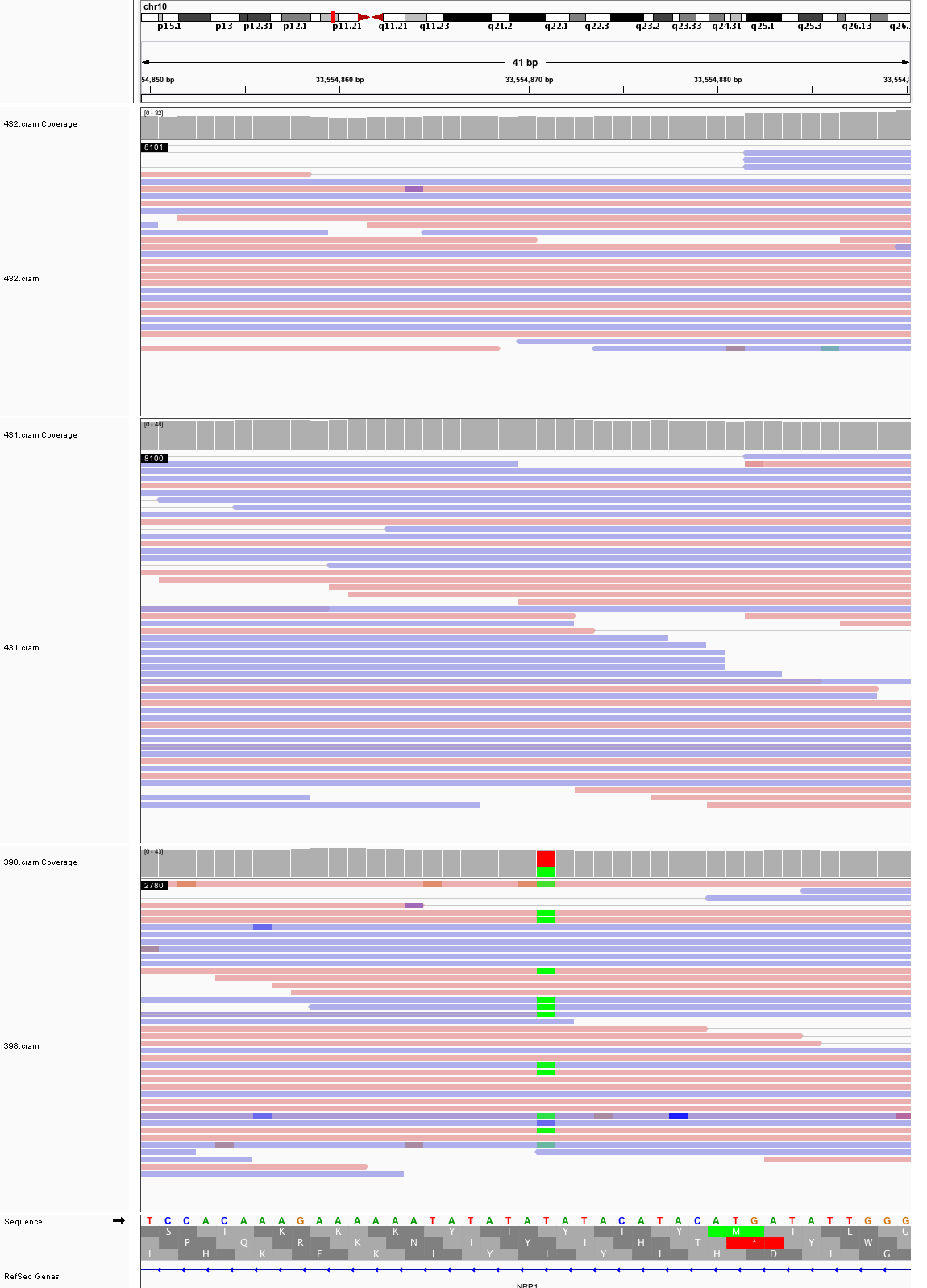

### chr10_80,520,922_80,520,962.png

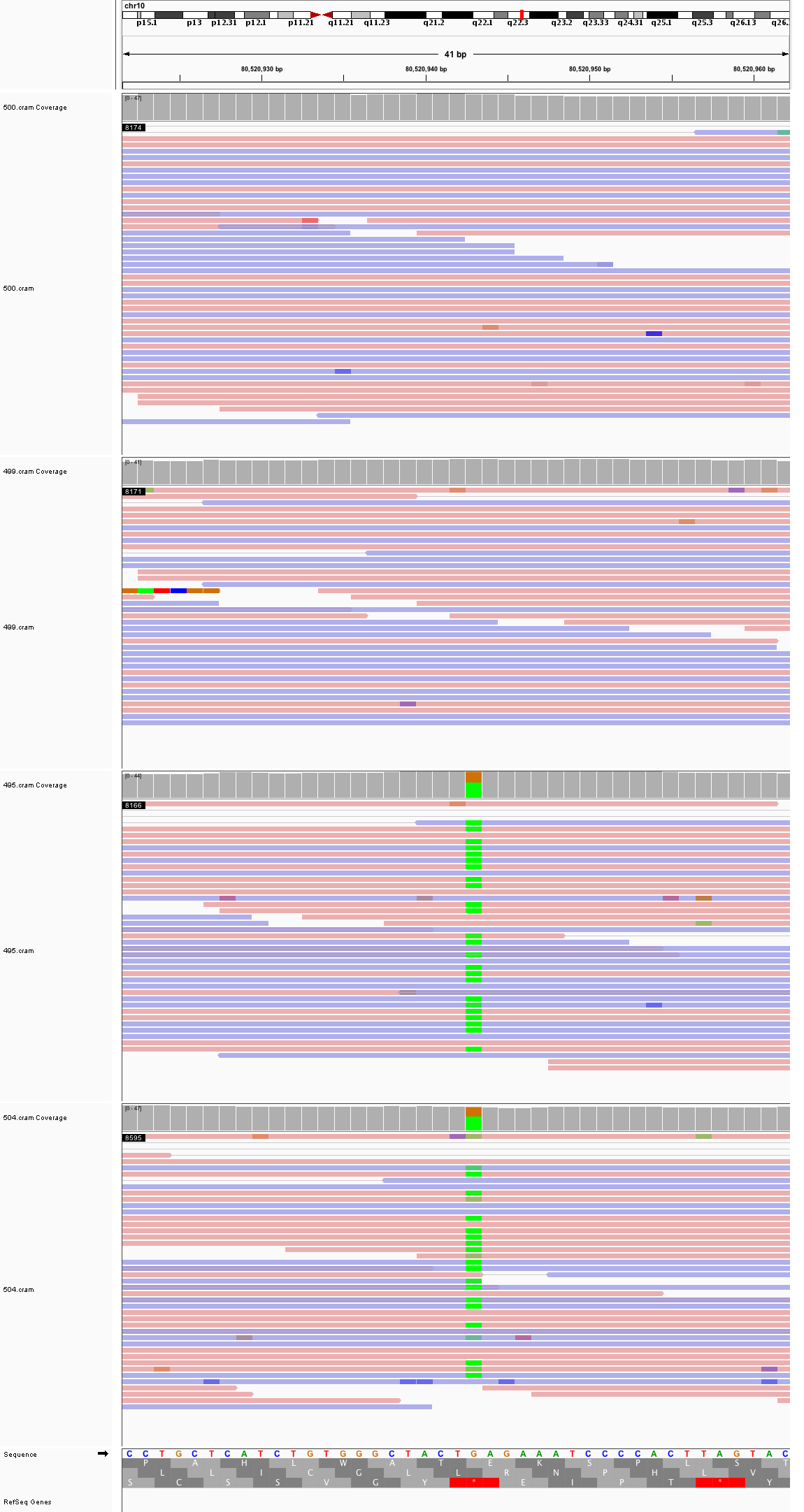

### chr10_107,499,596_107,499,636.png

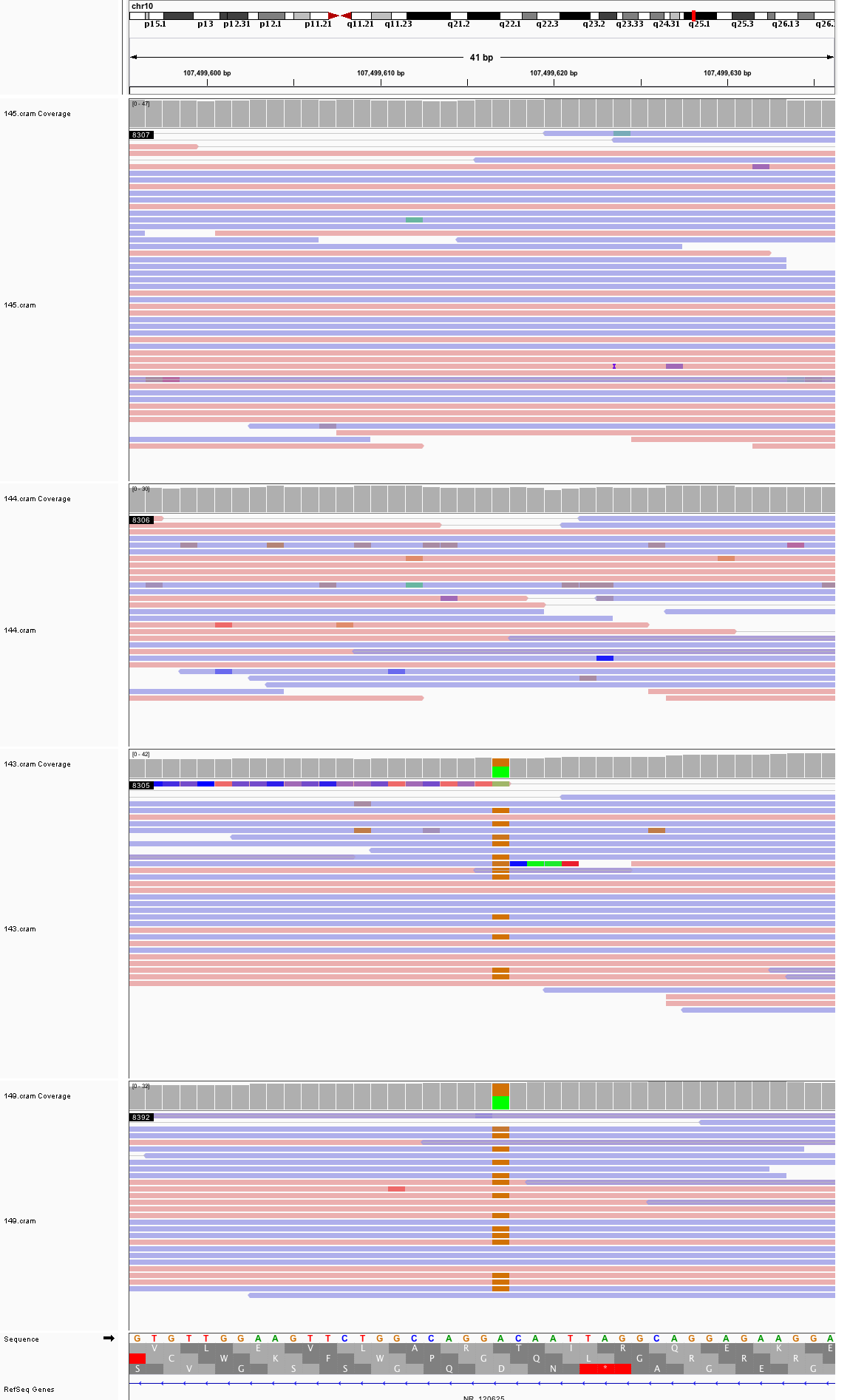

### chr10_117,108,232_117,108,272.png

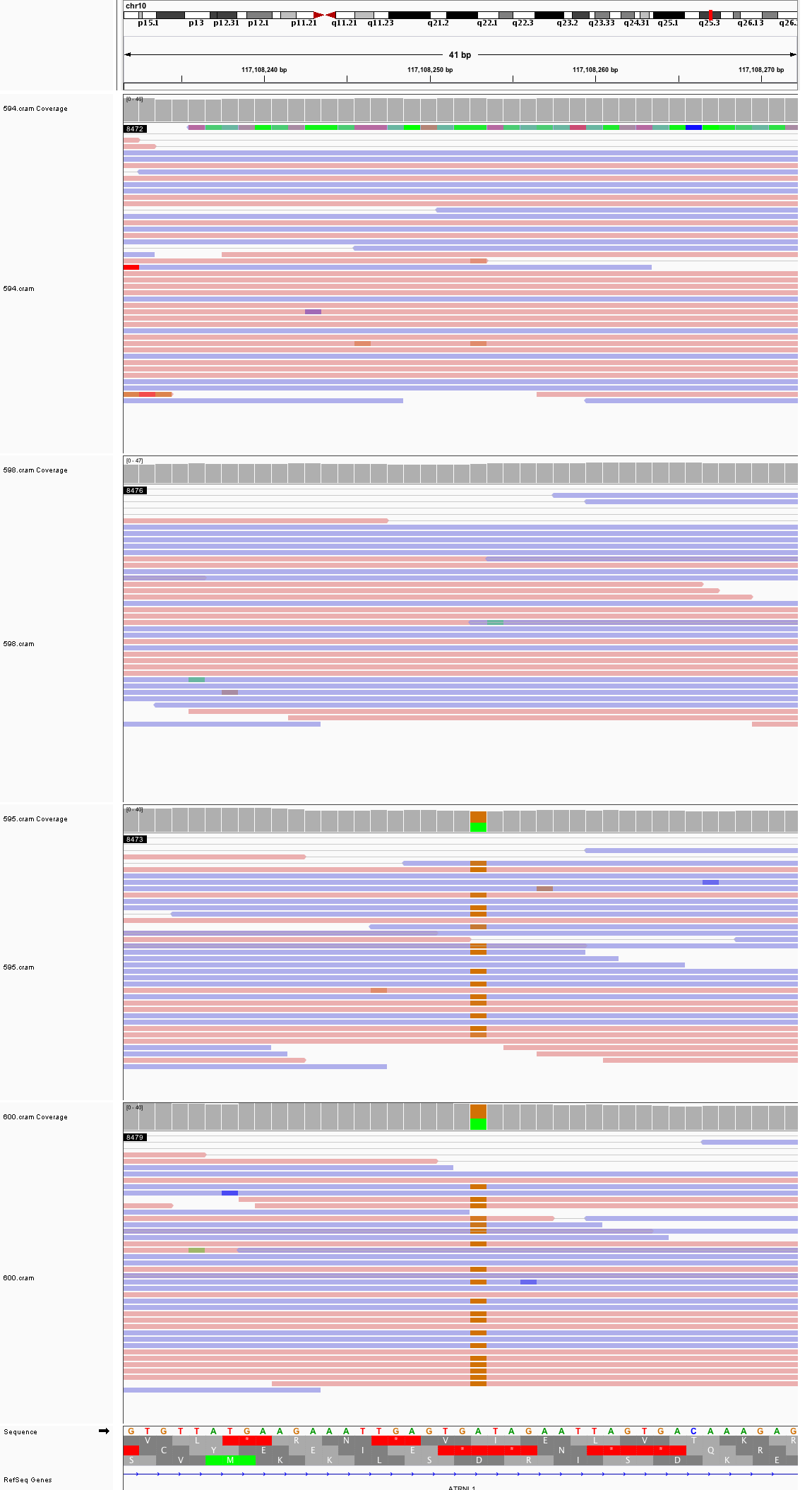

### chr10_128,315,054_128,315,094.png

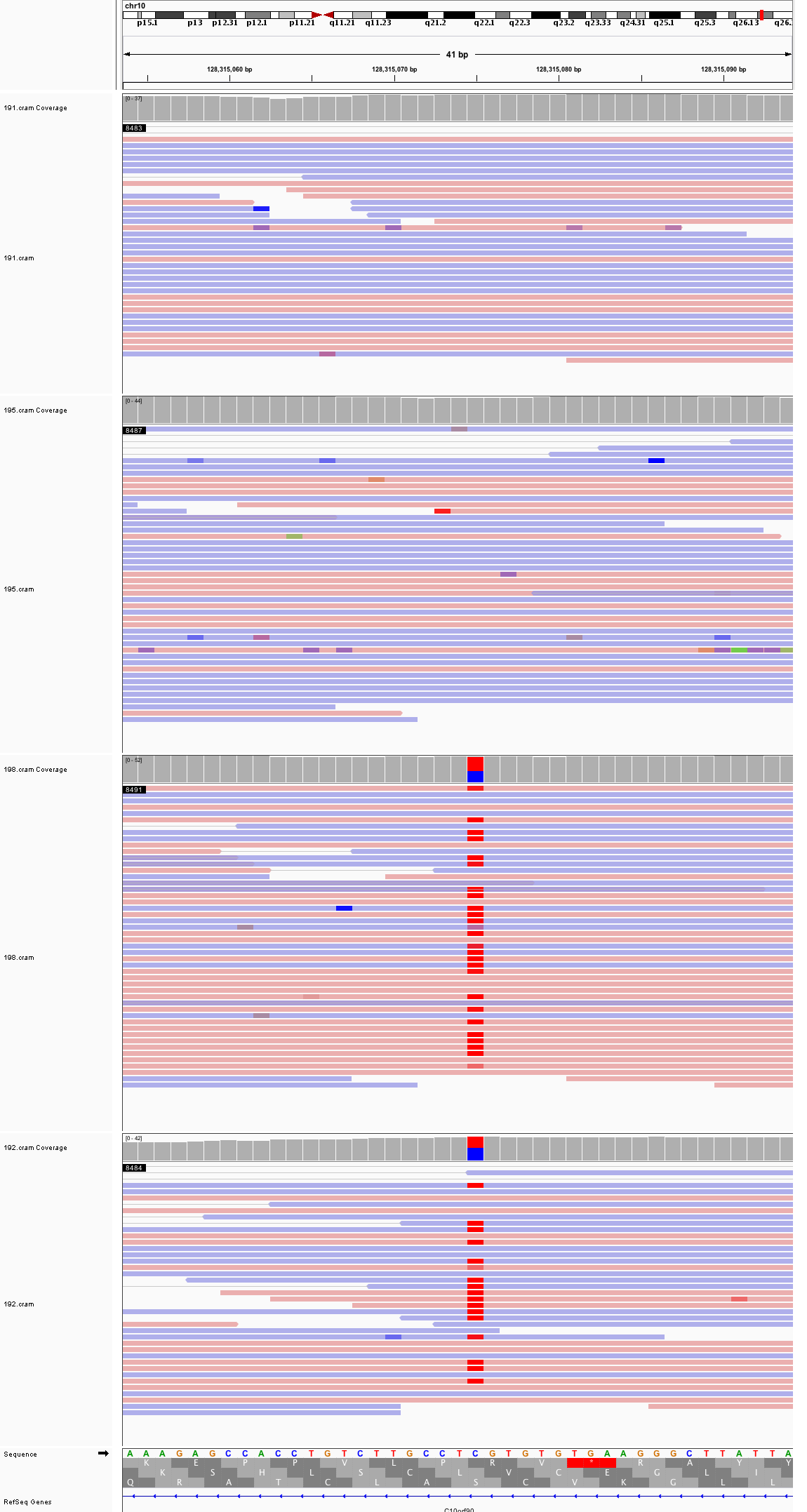

### chr10_133,923,850_133,923,890.png

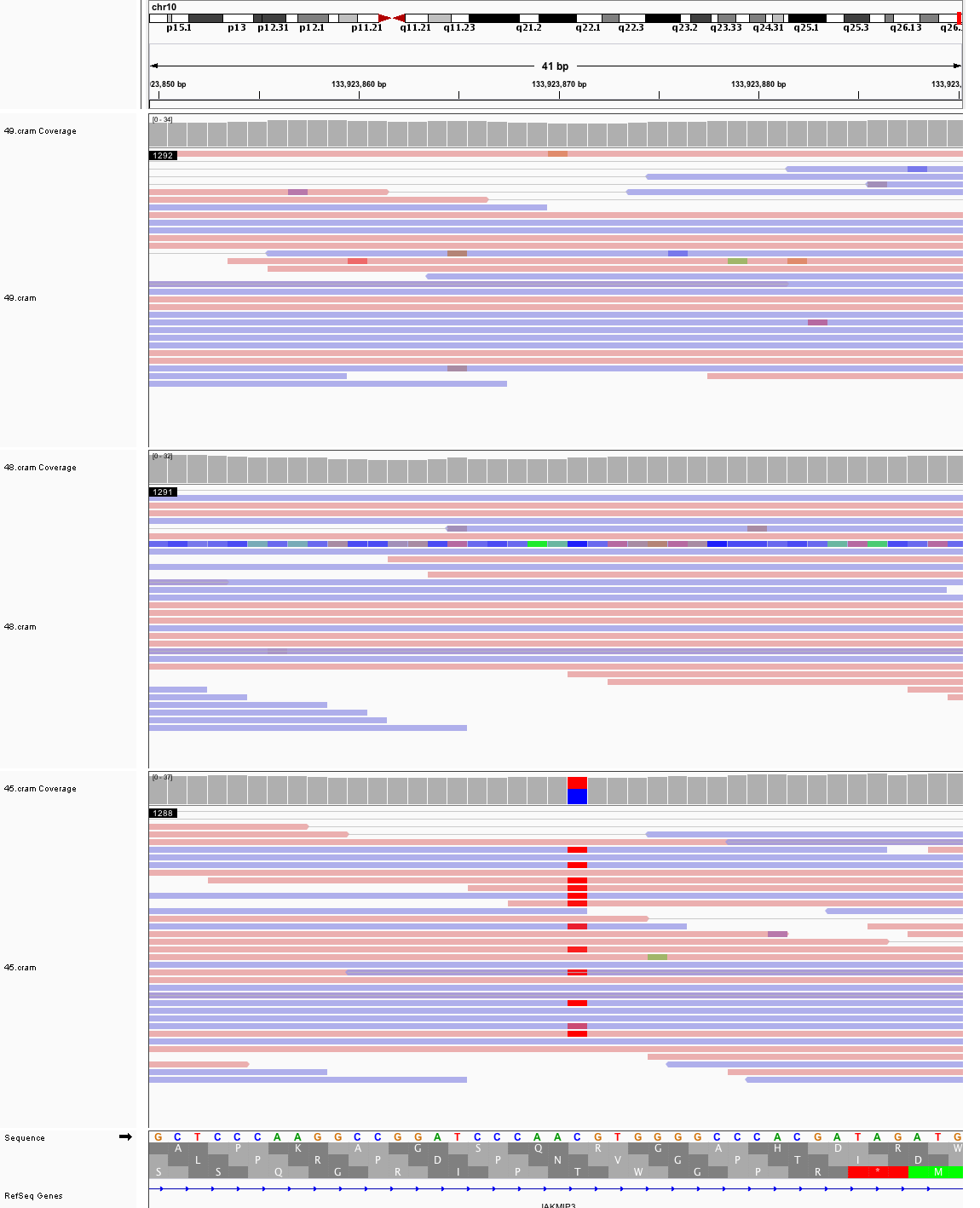

### chr11_28,393,621_28,393,661.png

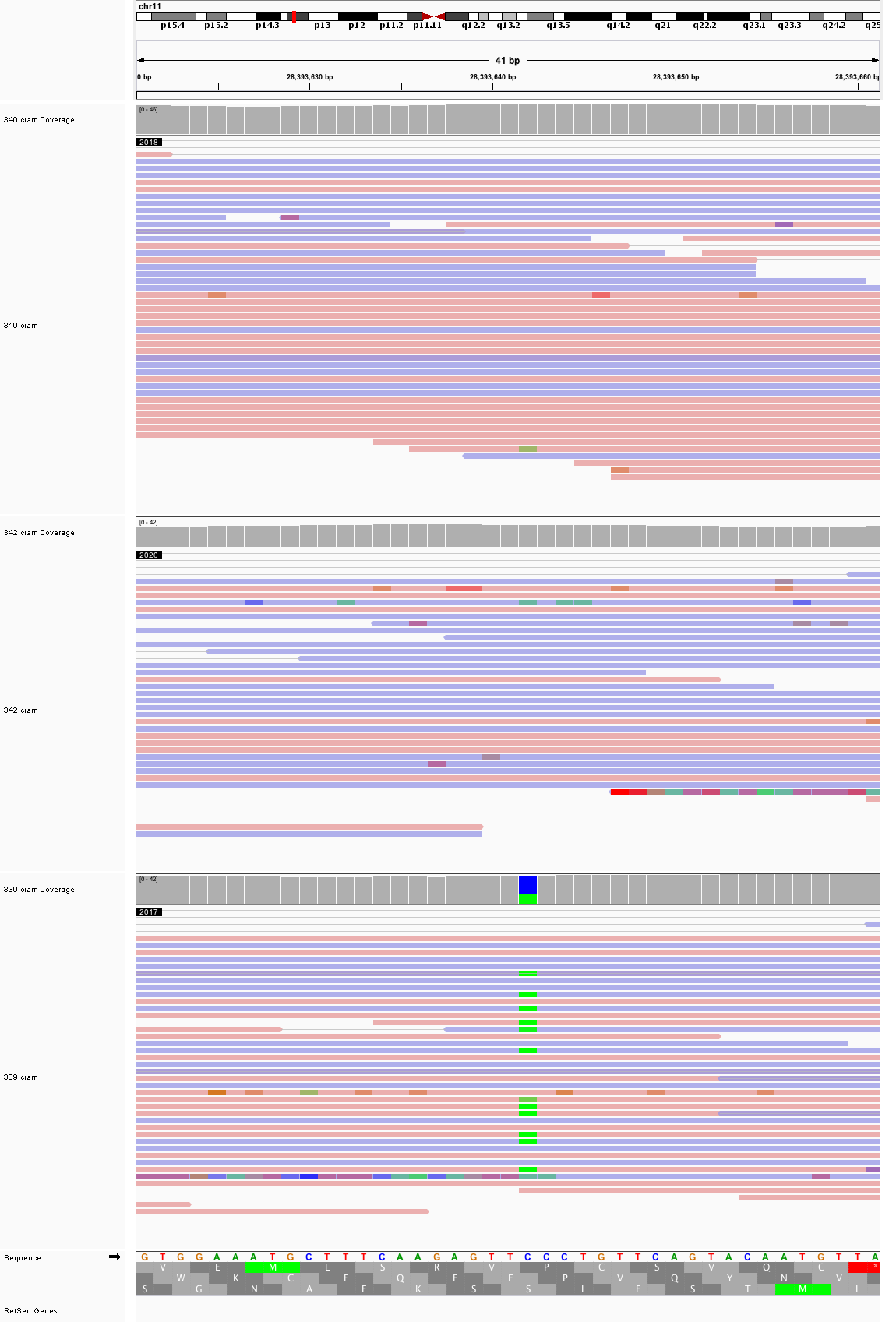

### chr11_76,765,650_76,765,690.png

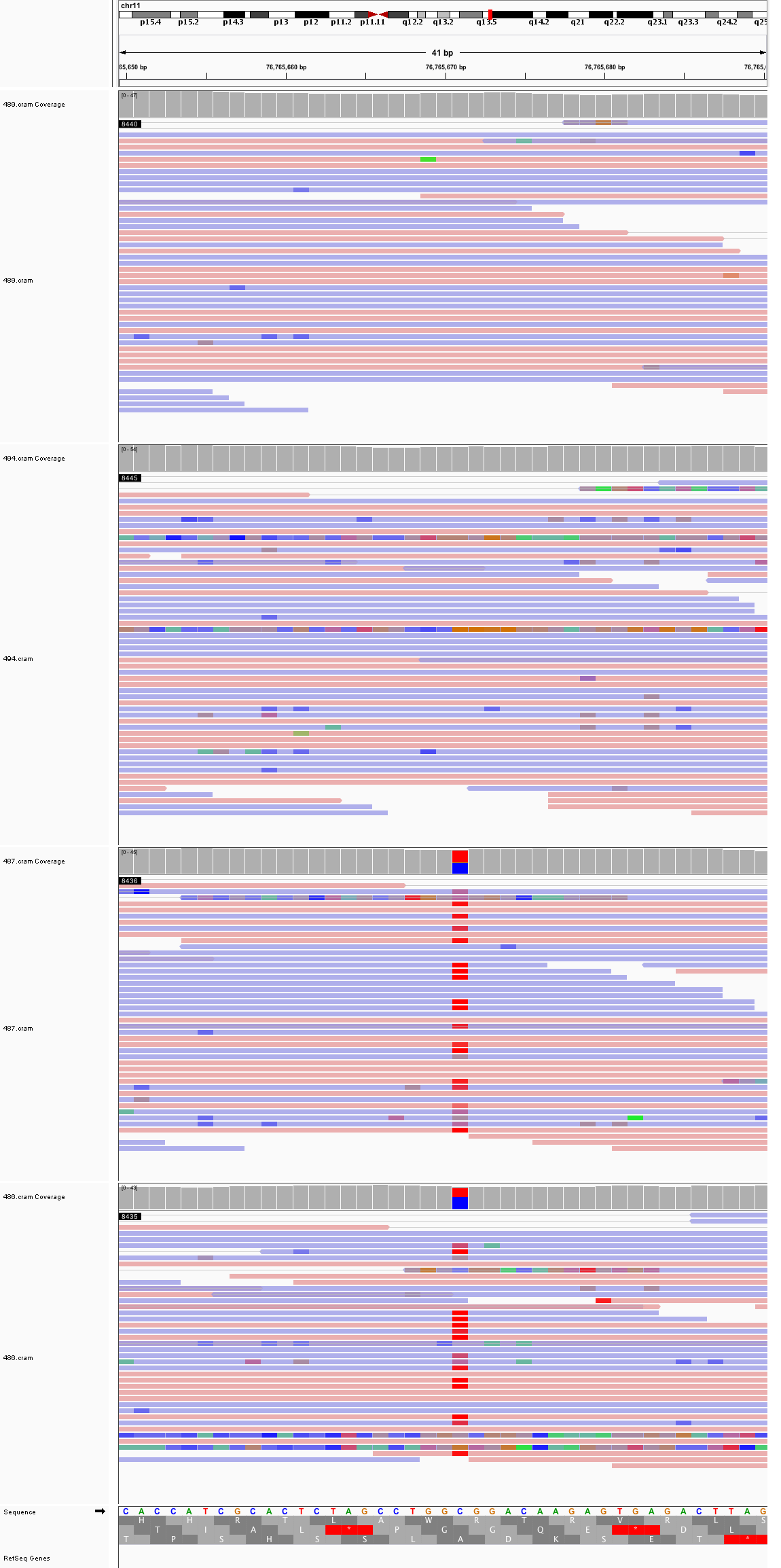

### chr11_112,679,691_112,679,731.png

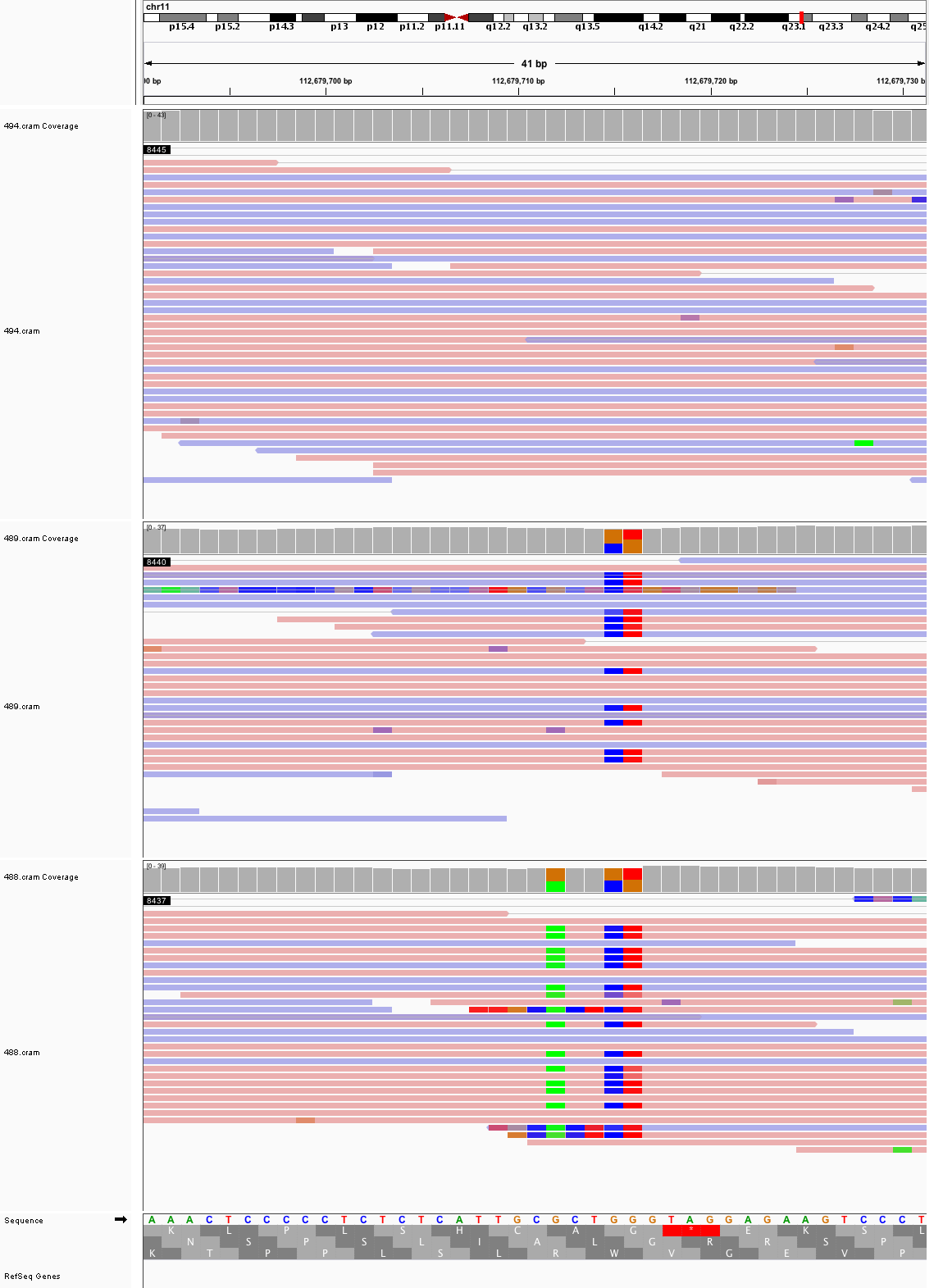

### chr12_16,764,451_16,764,491.png

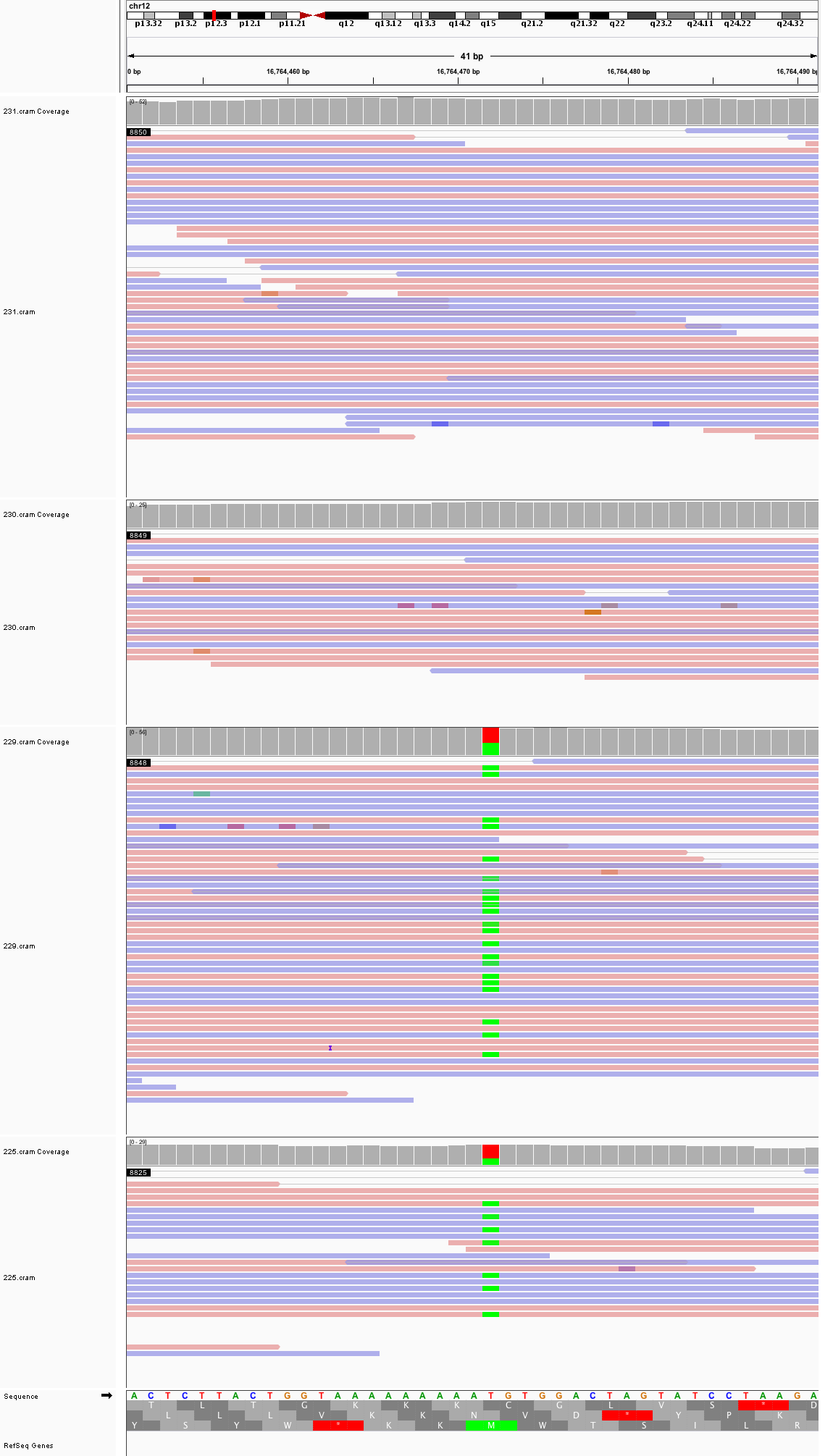

### chr12_41,080,642_41,080,682.png

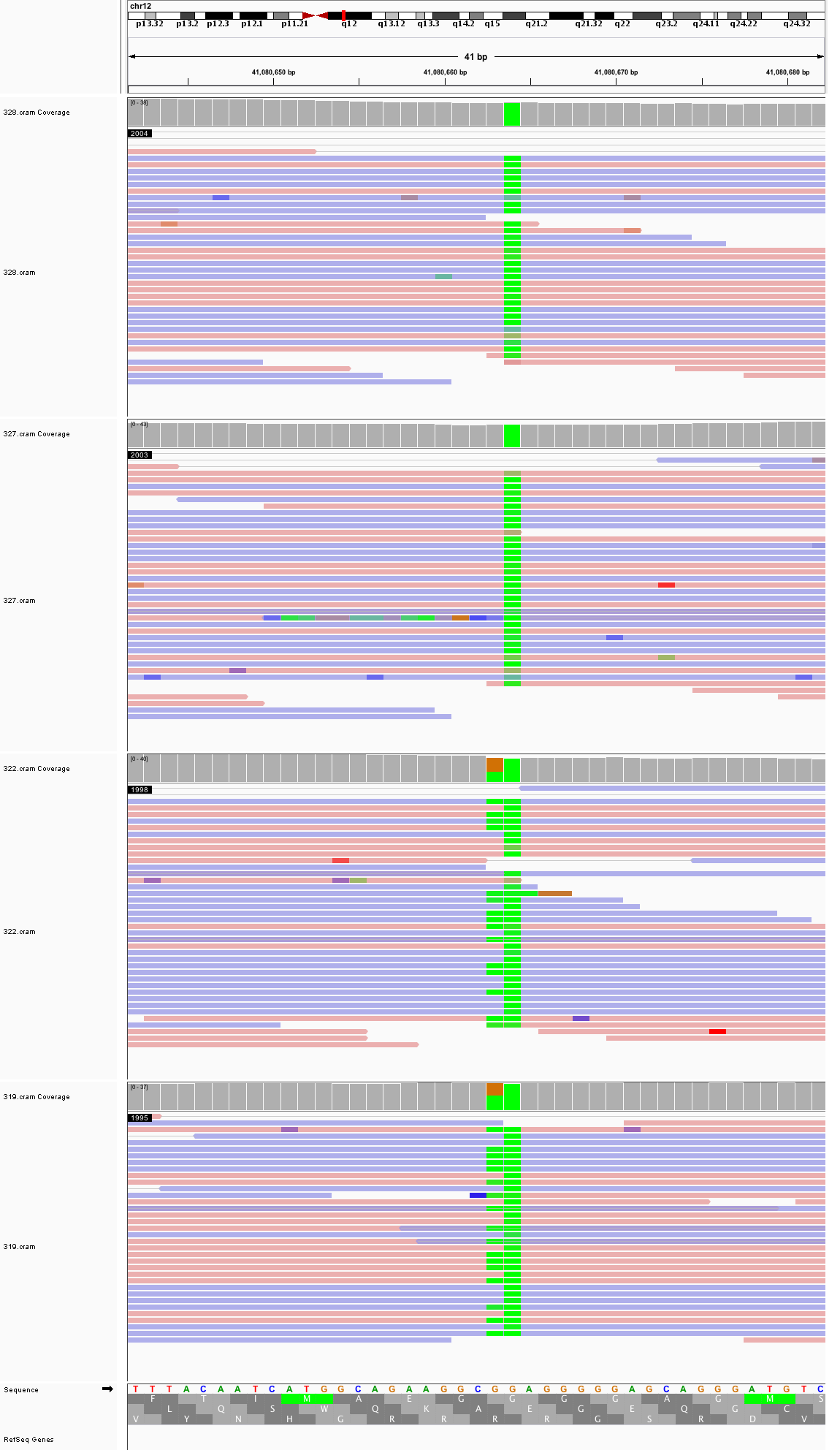

### chr12_44,622,367_44,622,407.png

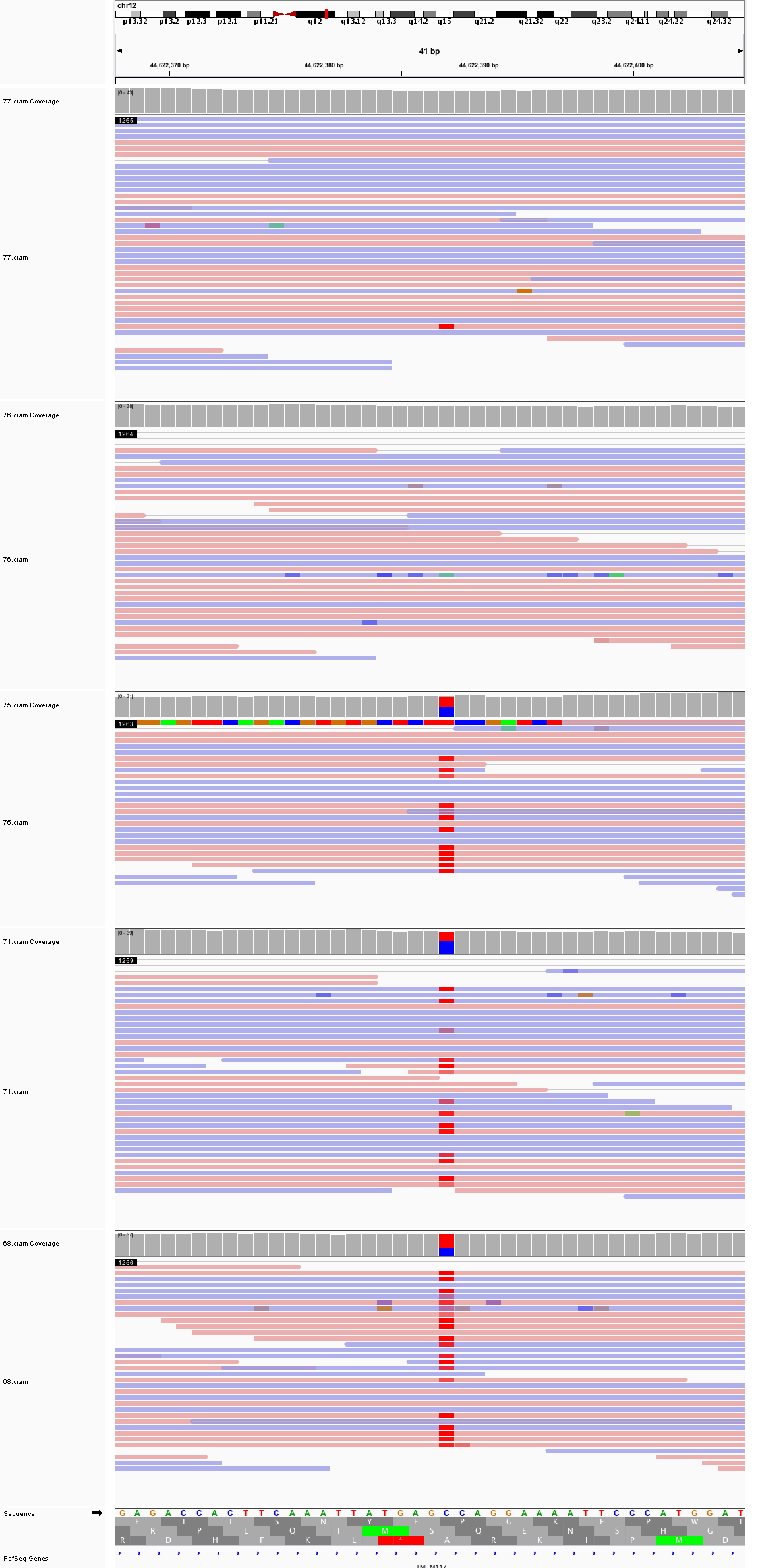

### chr12_55,732,054_55,732,094.png

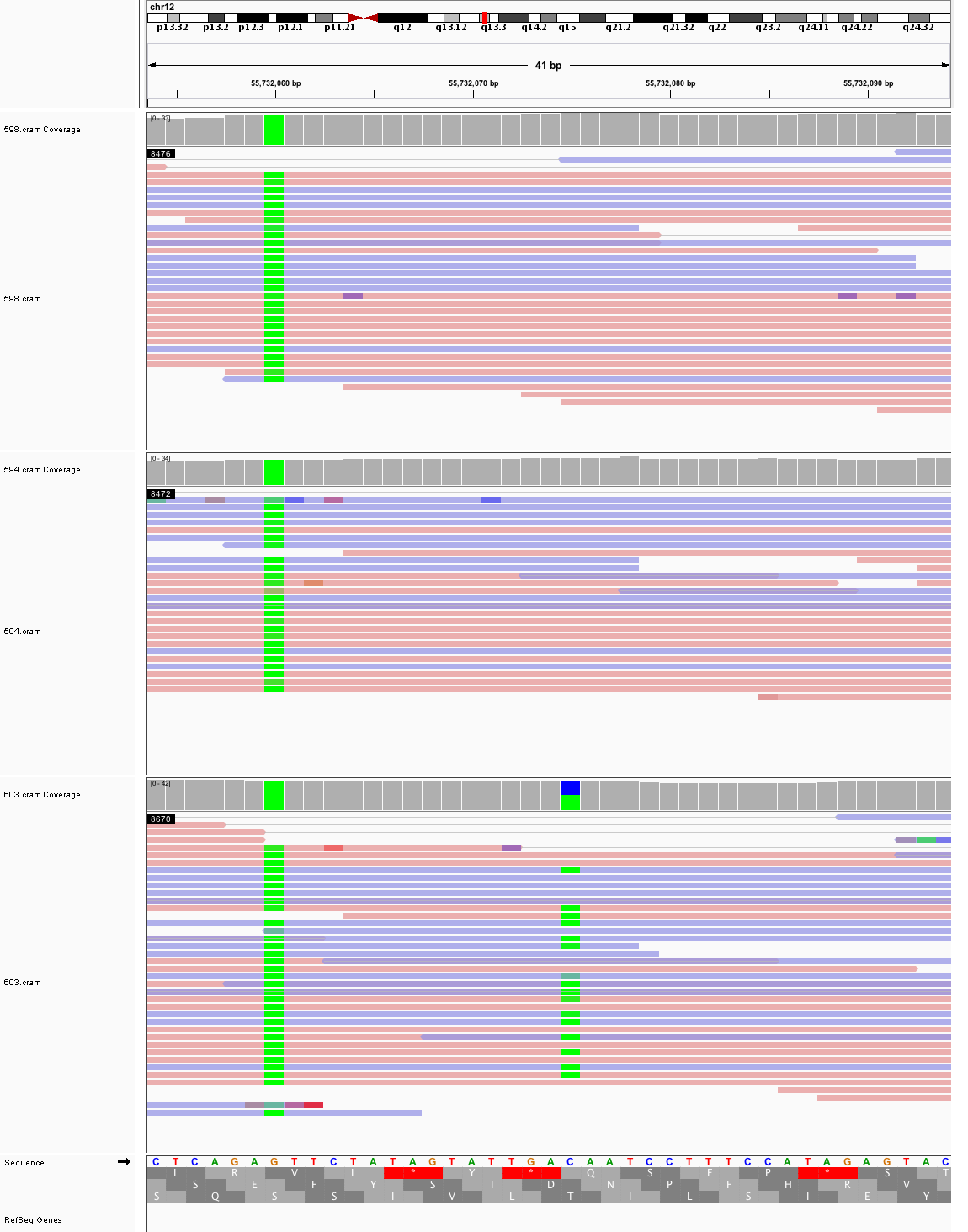

### chr12_55,958,831_55,958,871.png

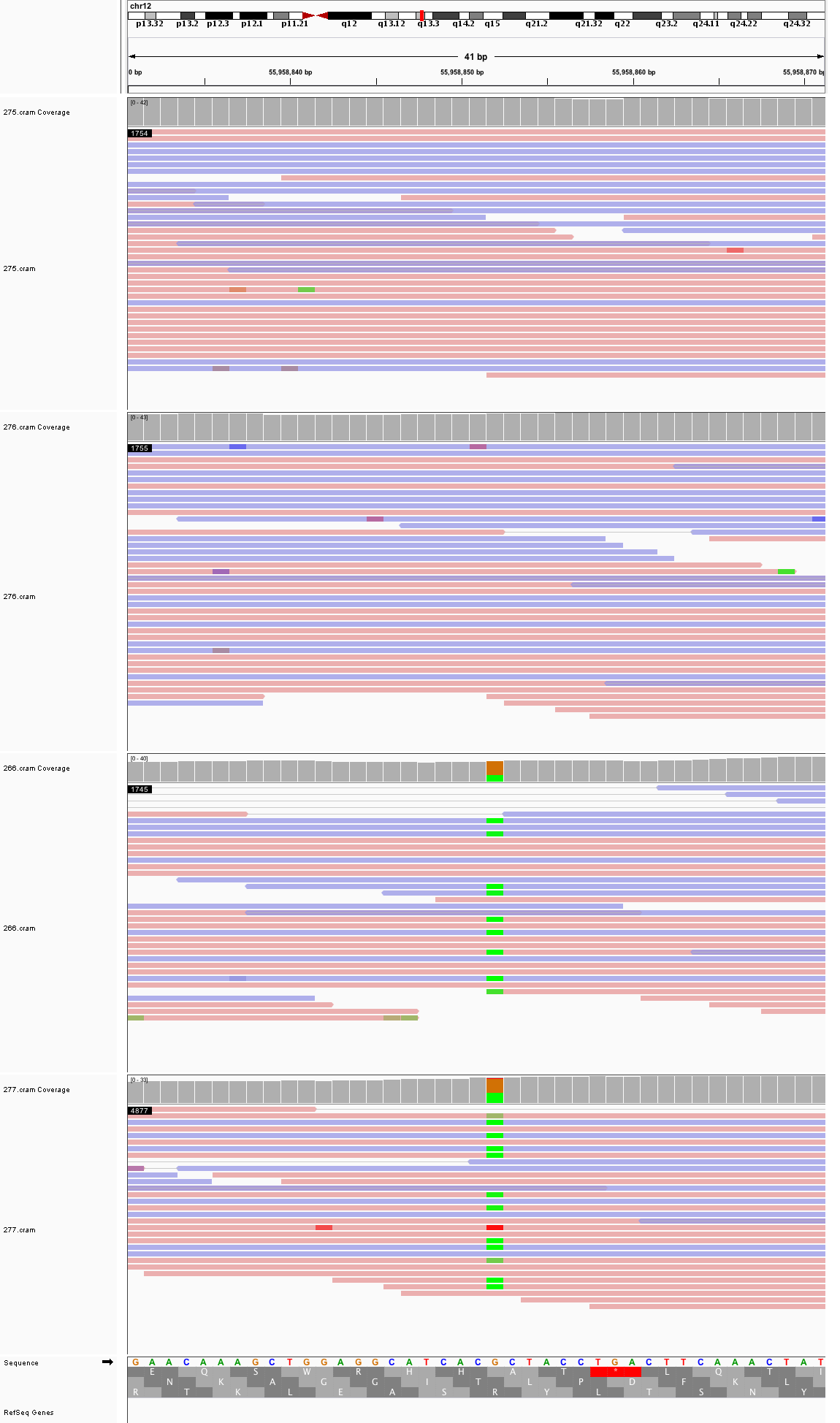

### chr13_53,395,378_53,395,418.png

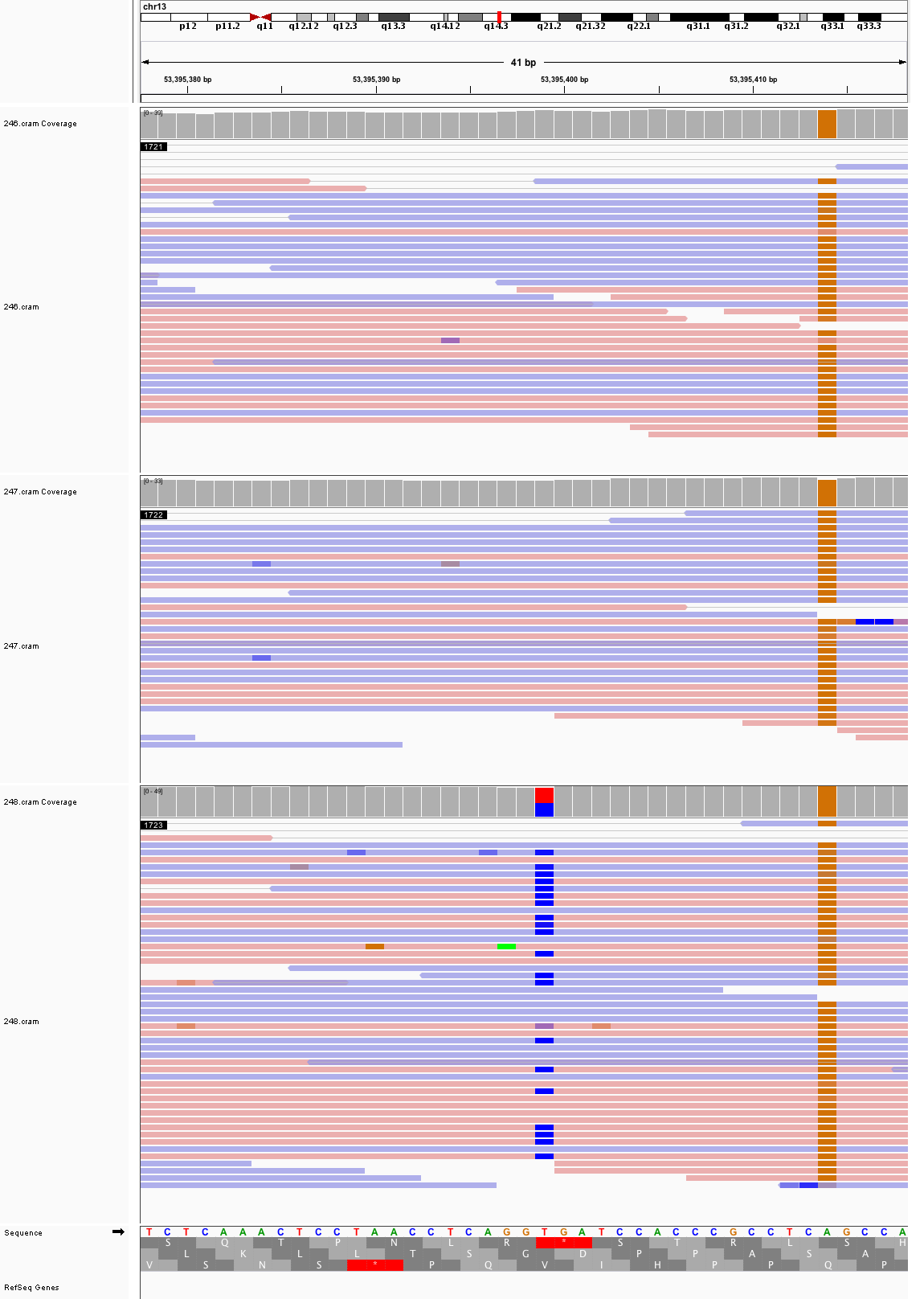

### chr13_87,301,573_87,301,613.png

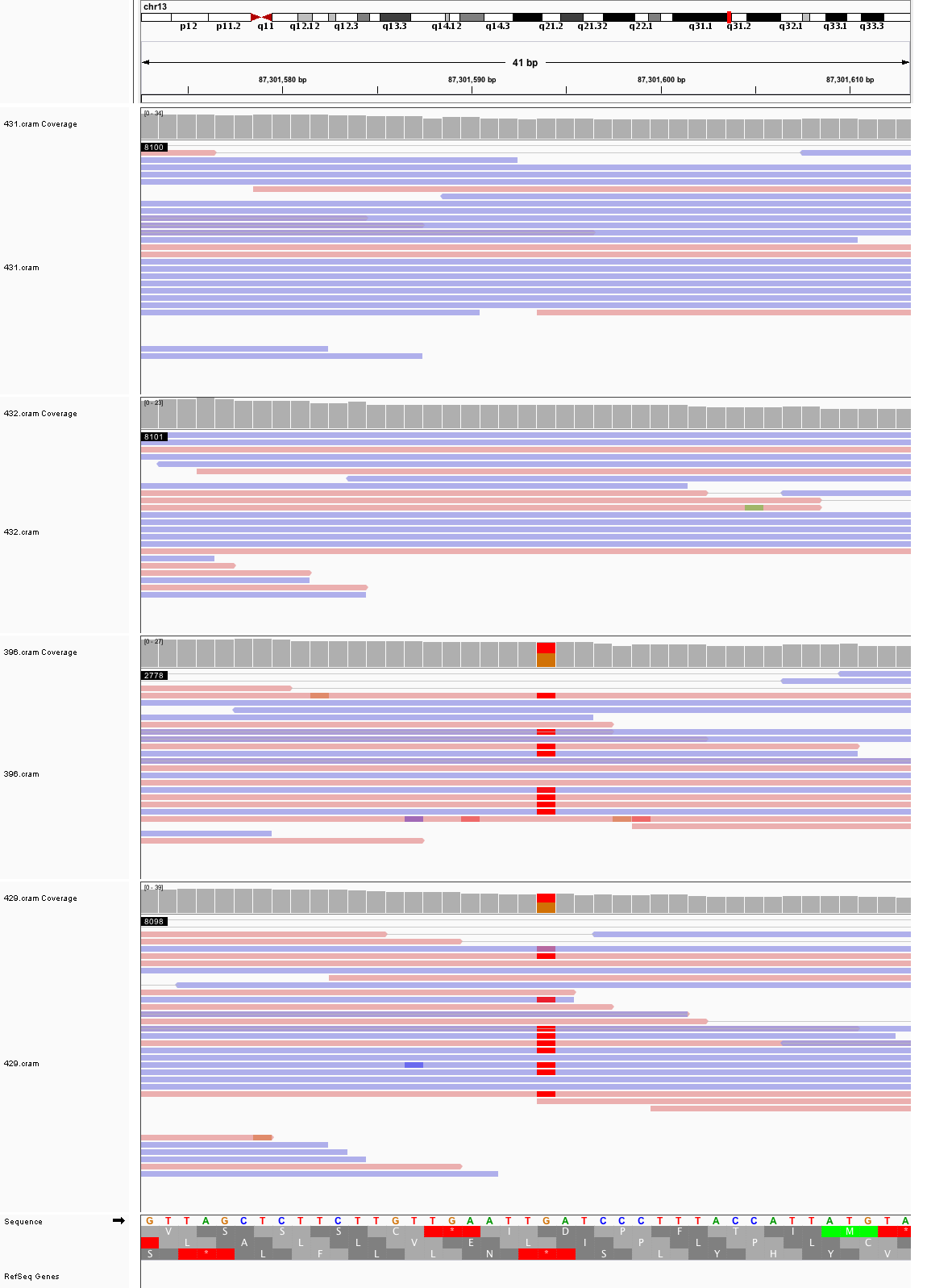

### chr13_109,017,557_109,017,597.png

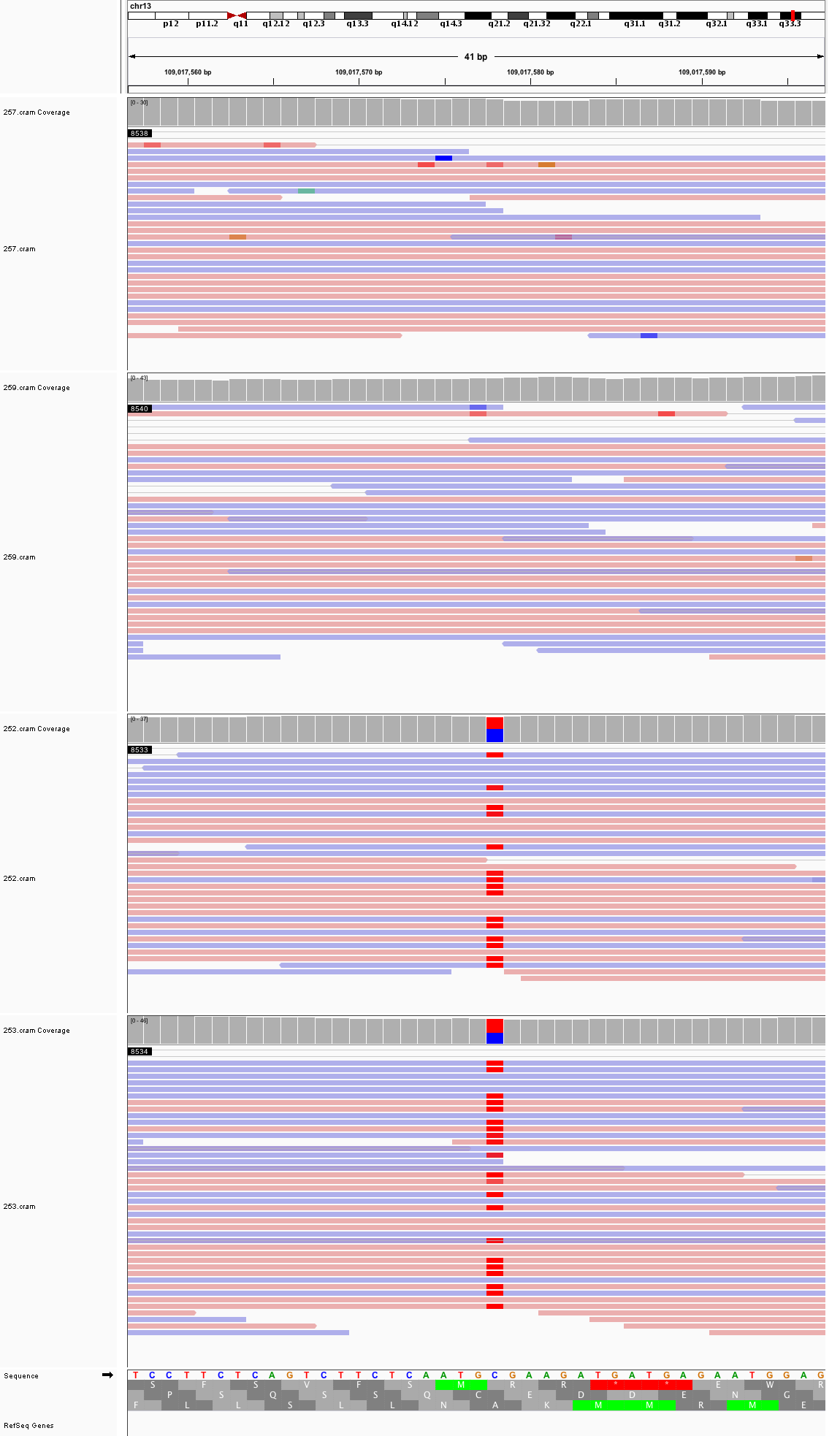

### chr14_26,034,854_26,034,894.png

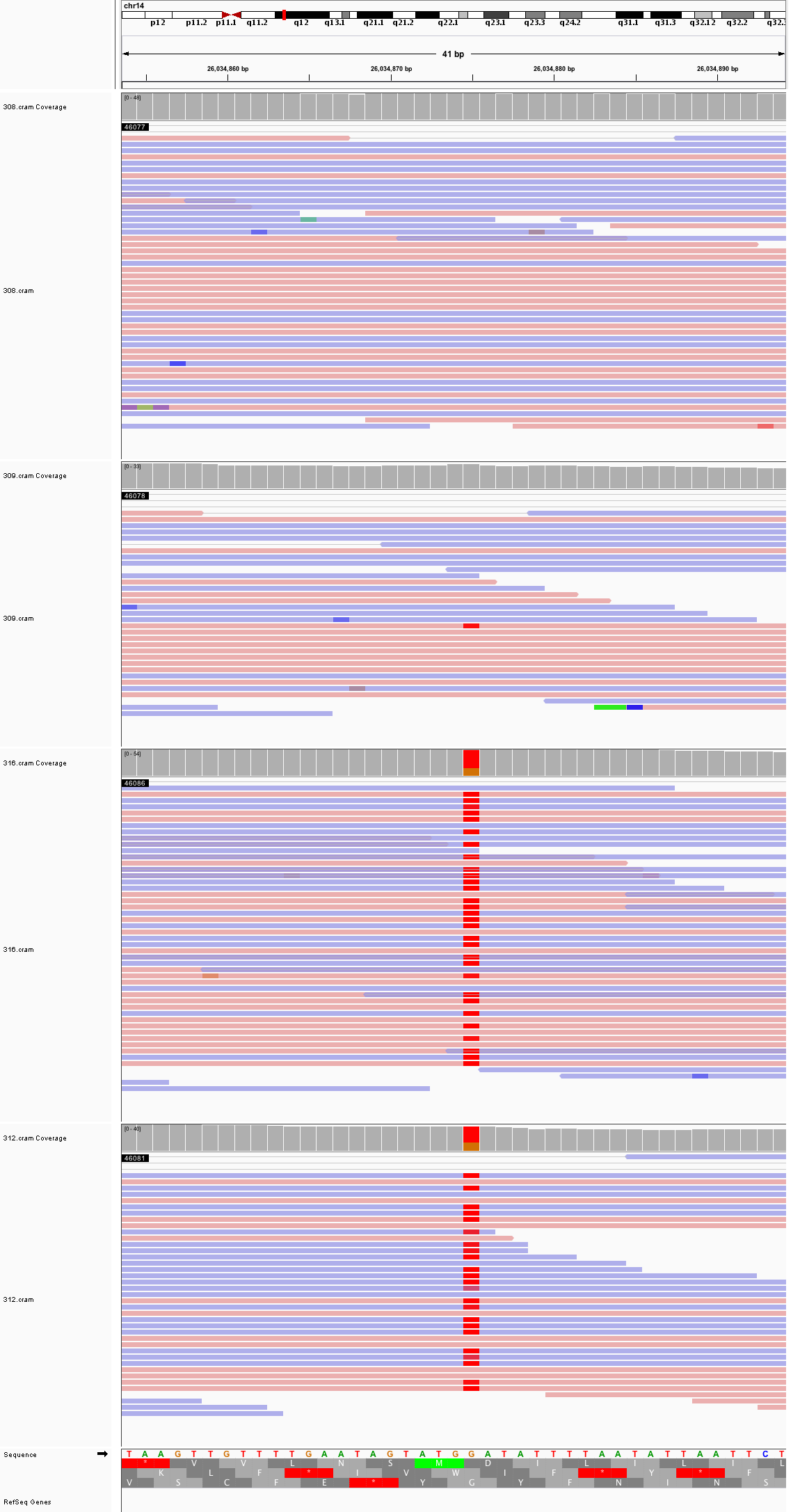

### chr14_106,782,048_106,782,088.png

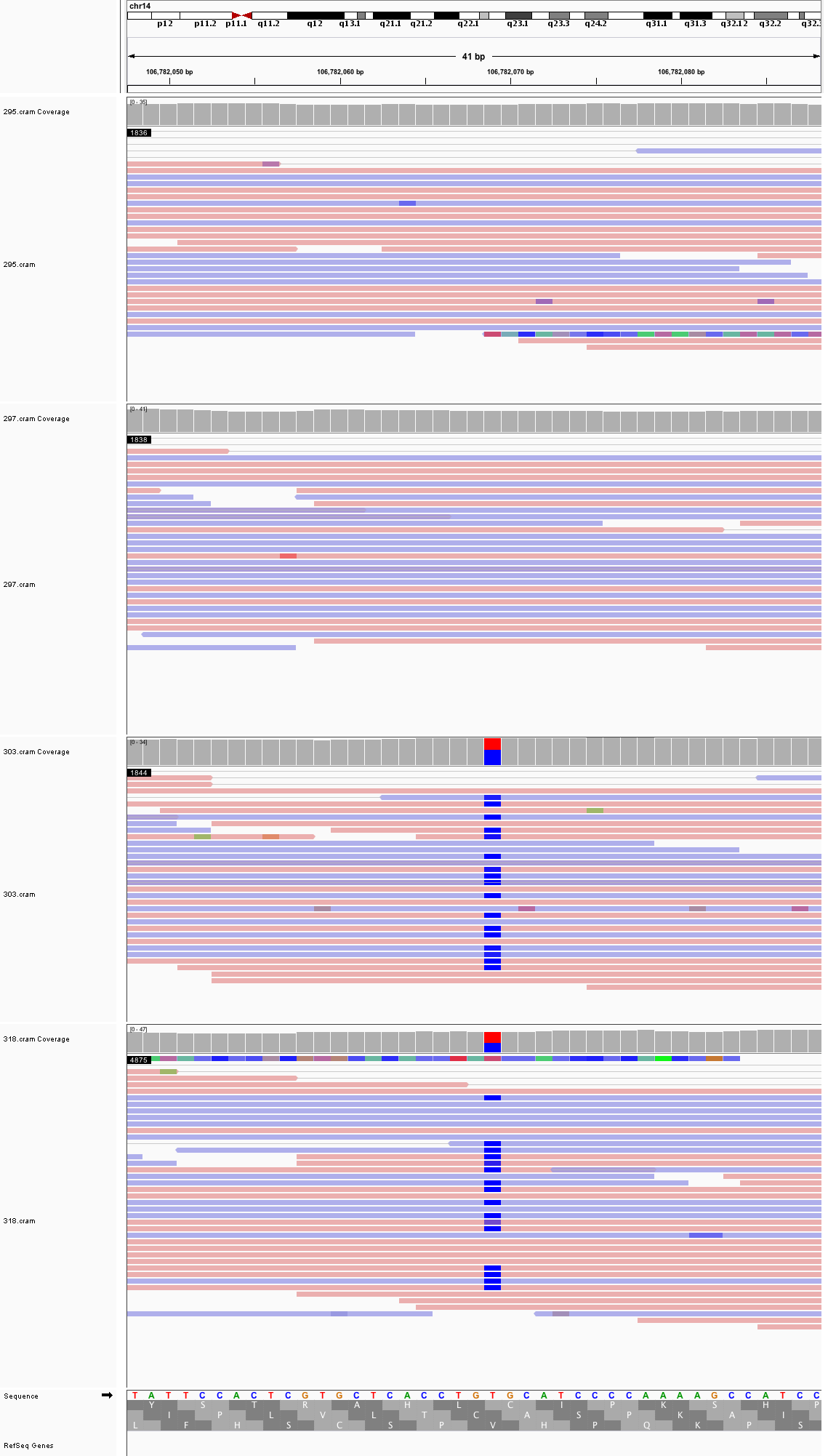

### chr15_25,467,310_25,467,350.png

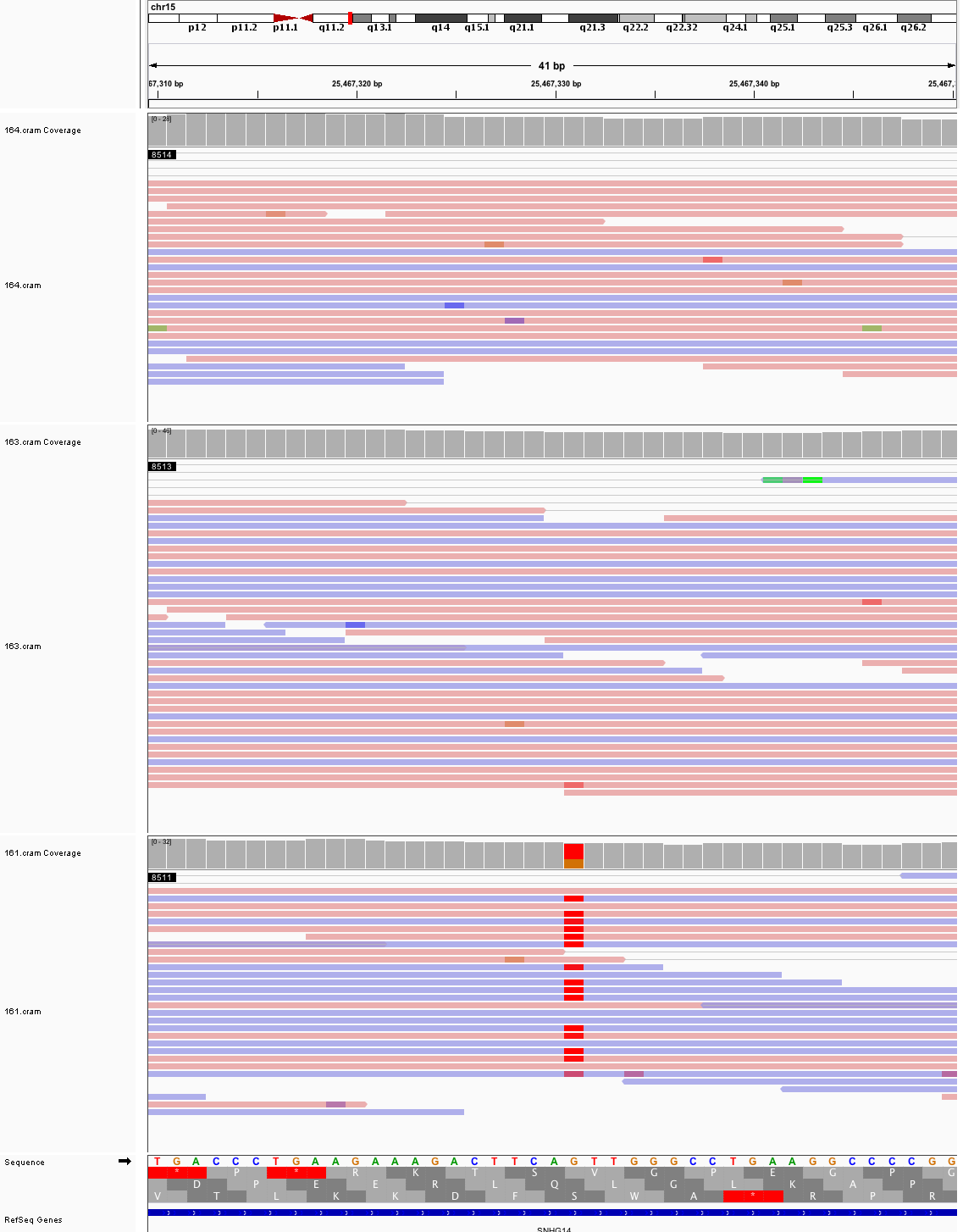

### chr15_48,244,132_48,244,172.png

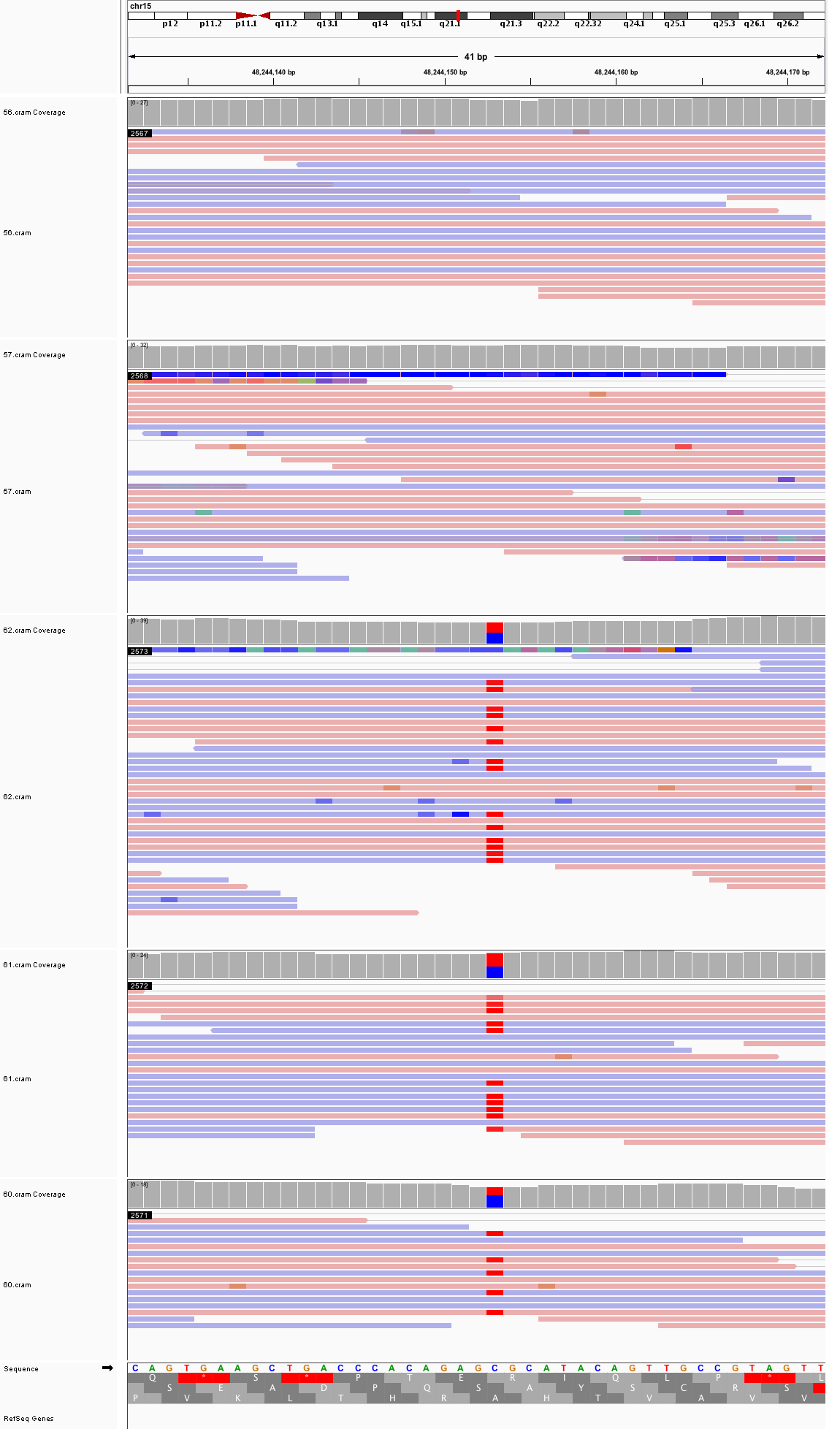

### chr15_89,575,369_89,575,409.png

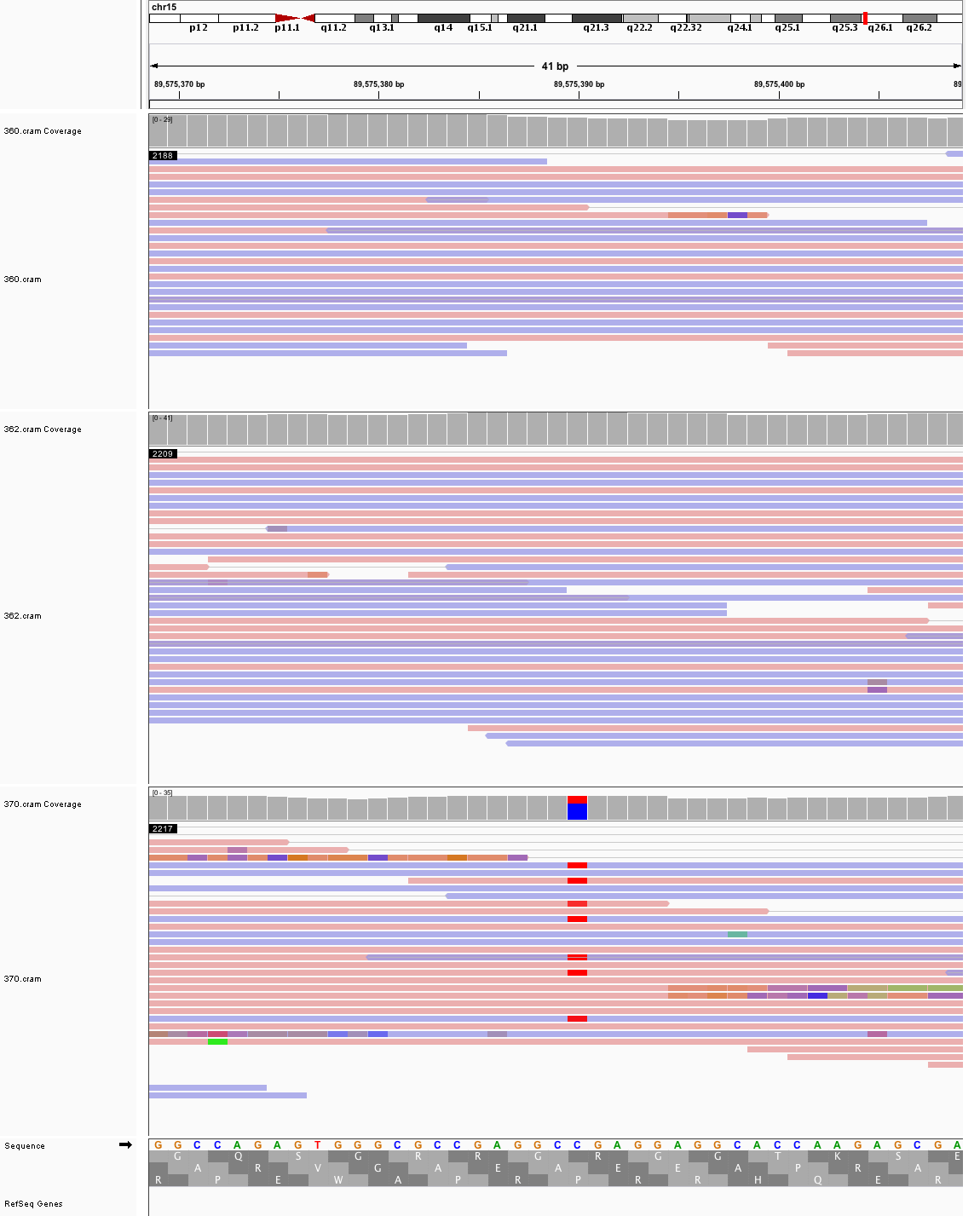

### chr17_2,733,437_2,733,477.png

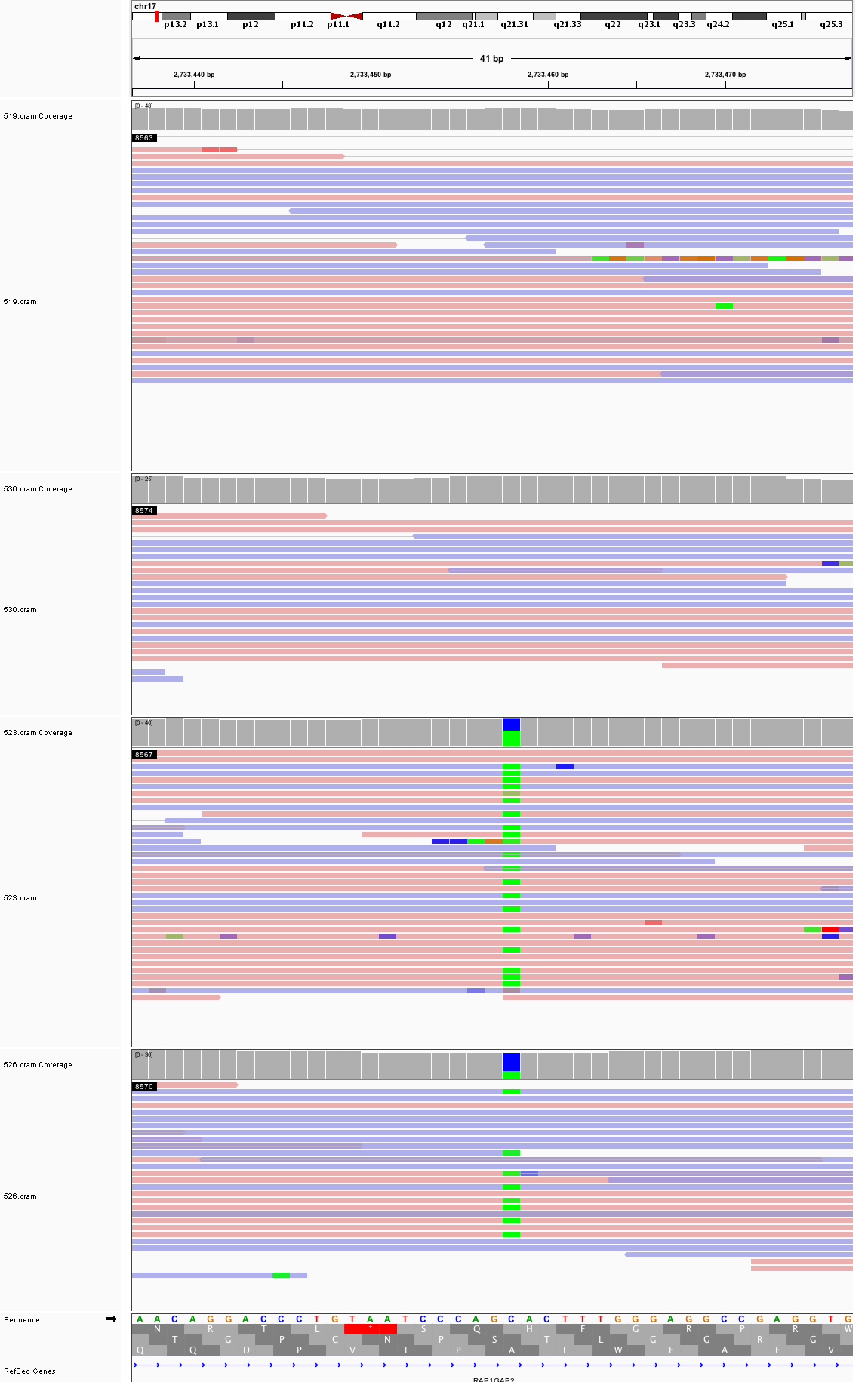

### chr17_12,982,605_12,982,645.png

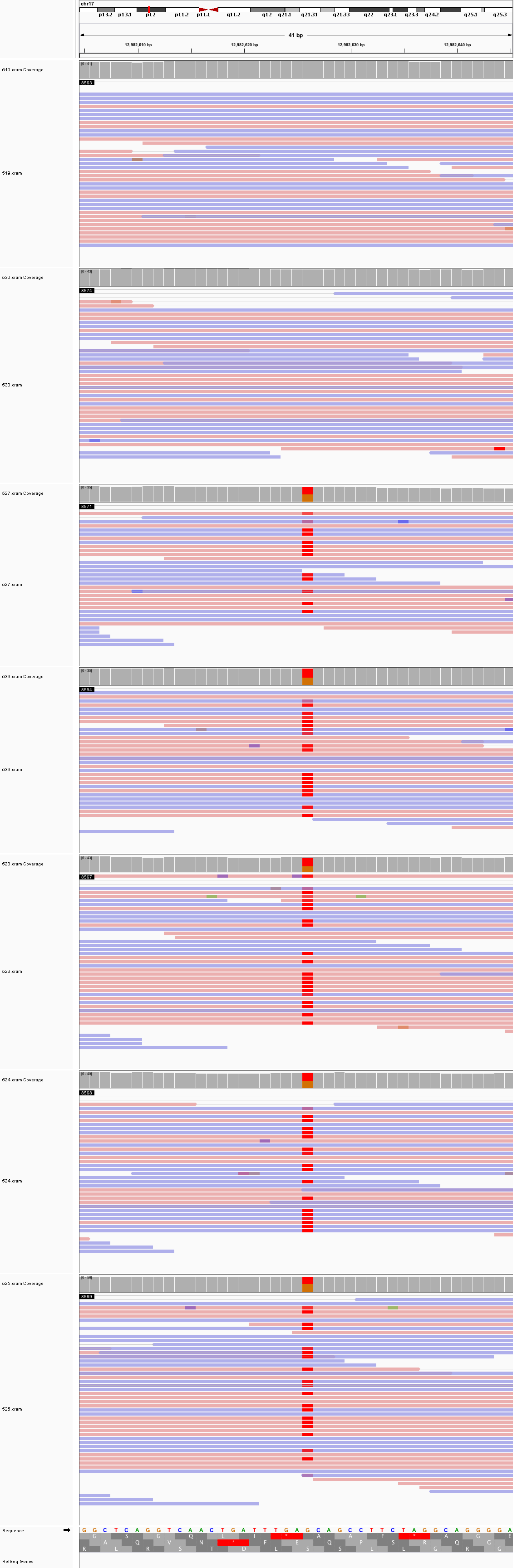

### chr17_48,679,864_48,679,904.png

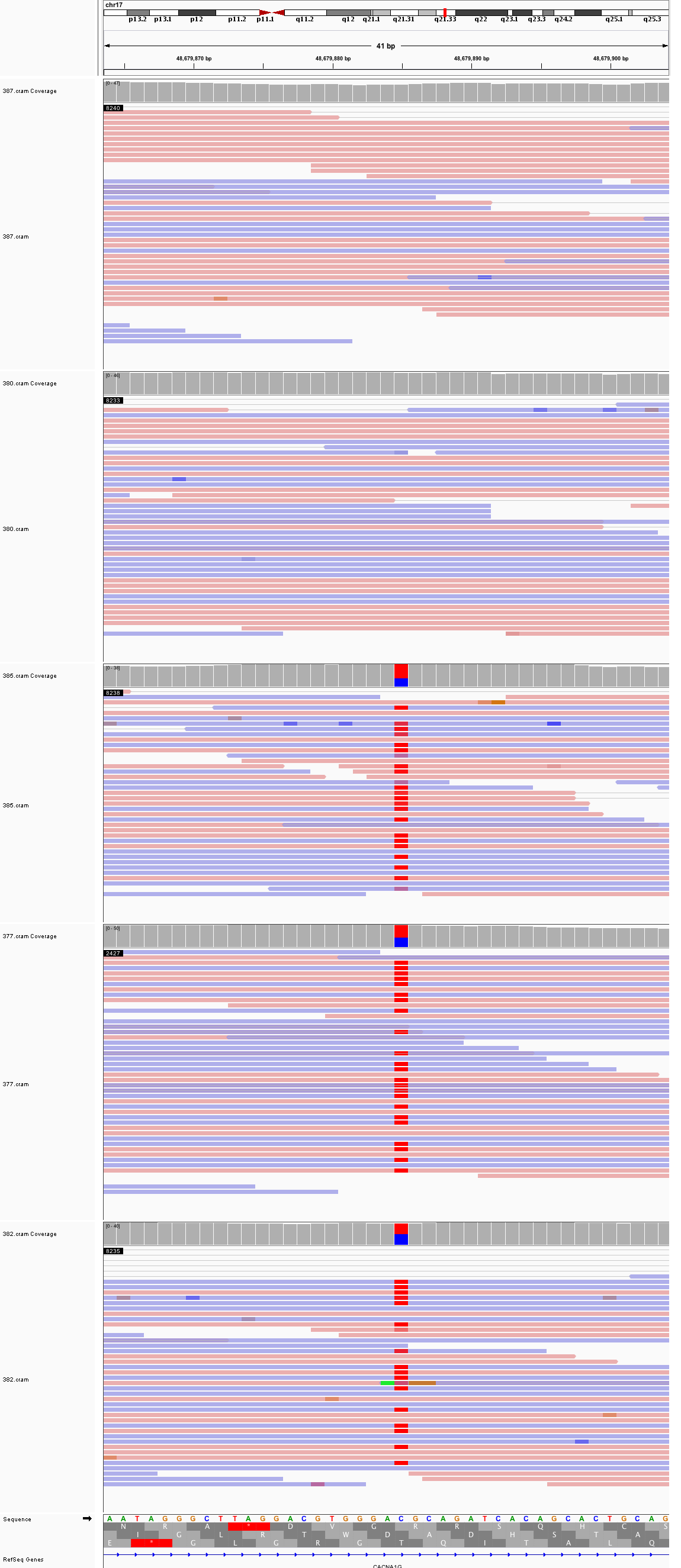

### chr17_79,350,891_79,350,931.png

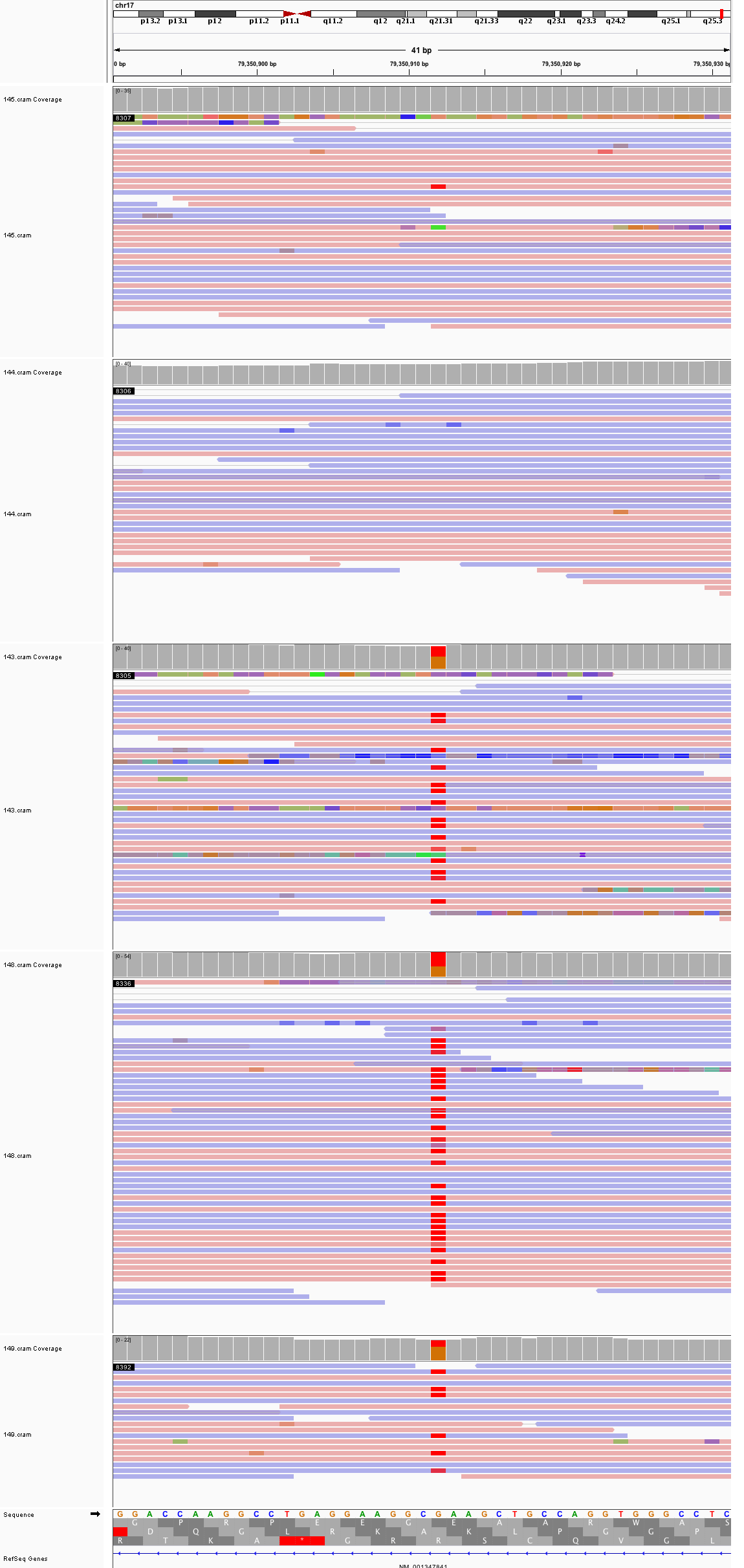

### chr18_22,867,951_22,867,991.png

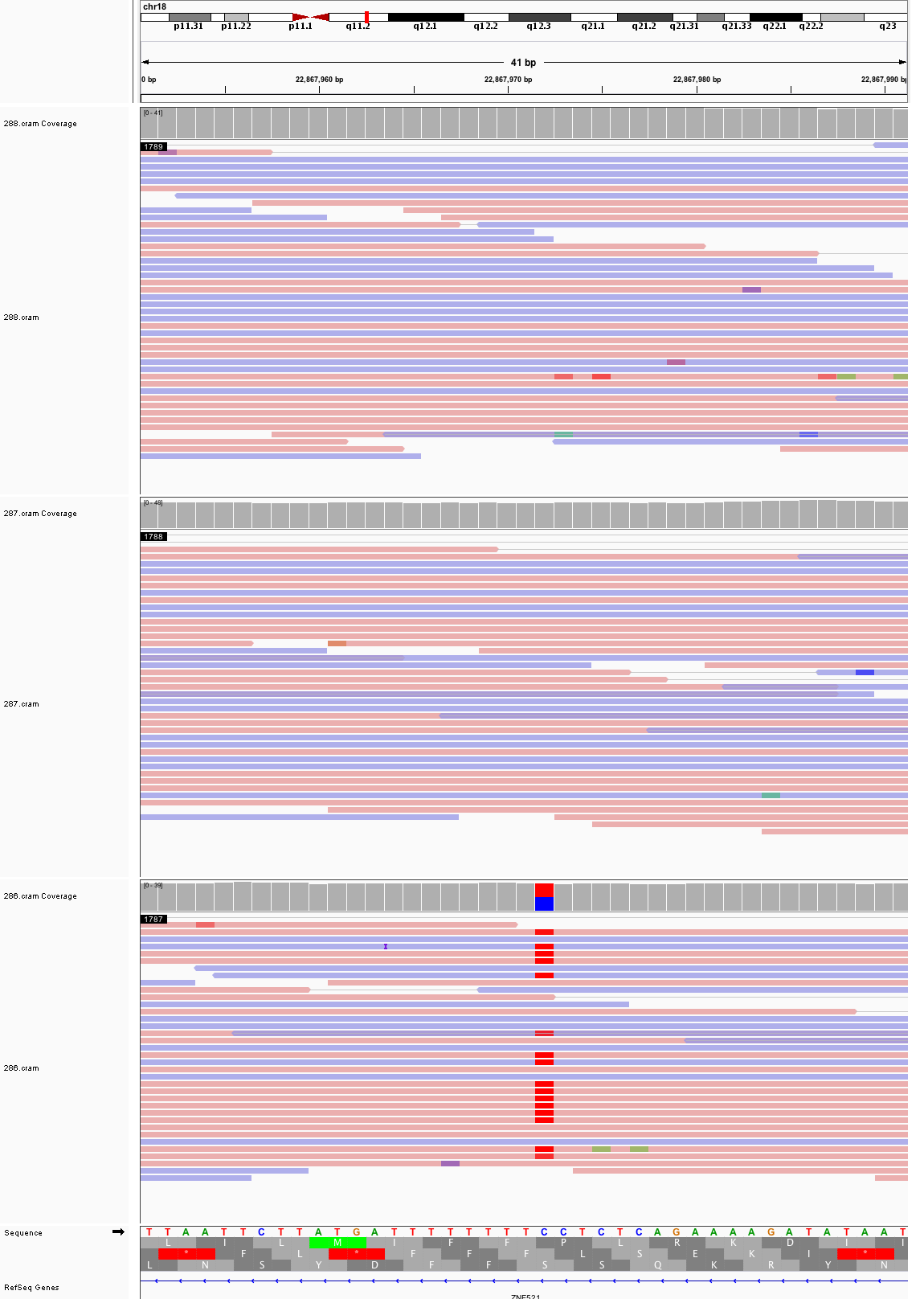

### chr19_33,717,892_33,717,932.png

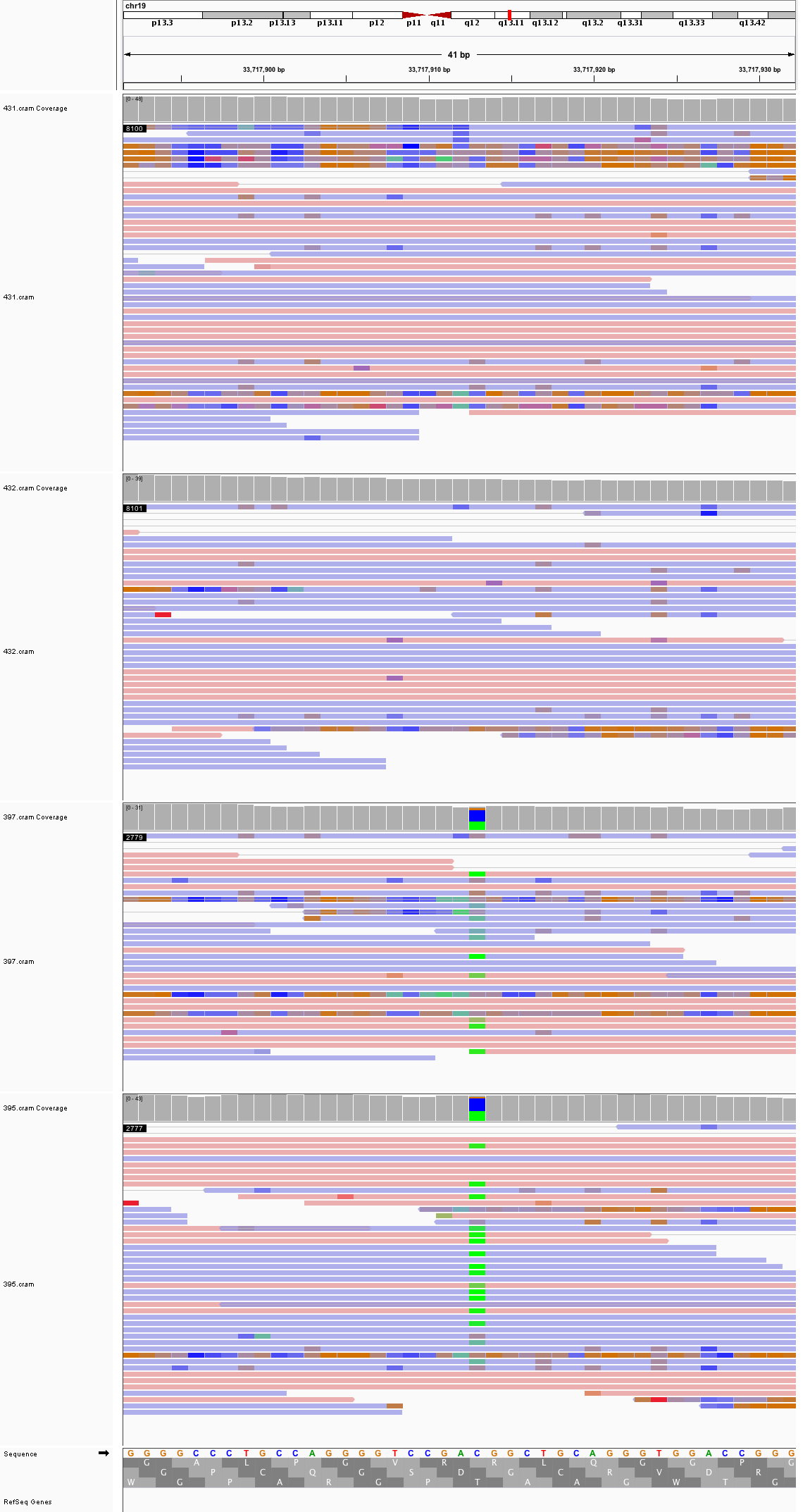

### chr19_54,504,968_54,505,008.png

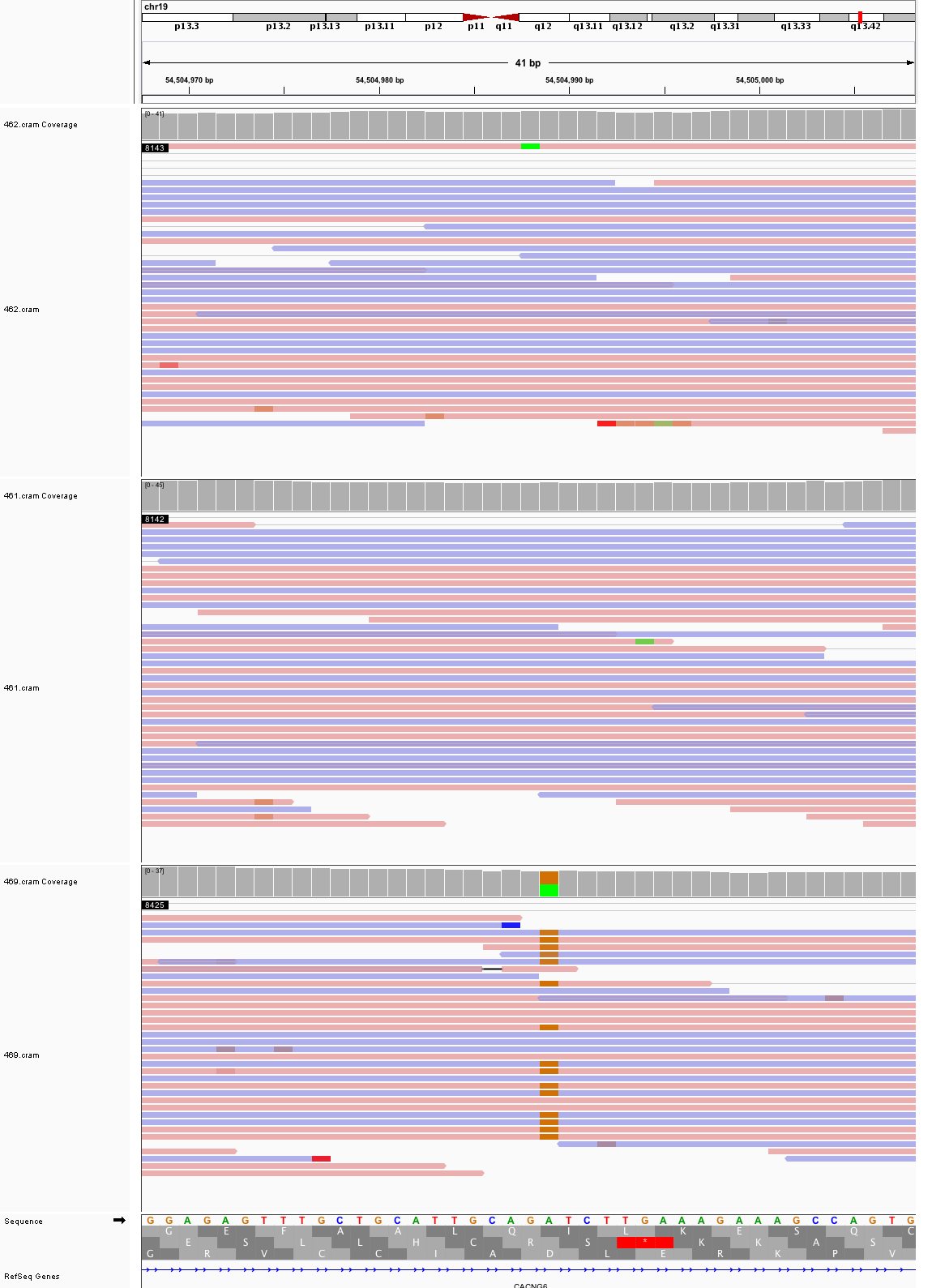

### chr20_33,756,190_33,756,230.png

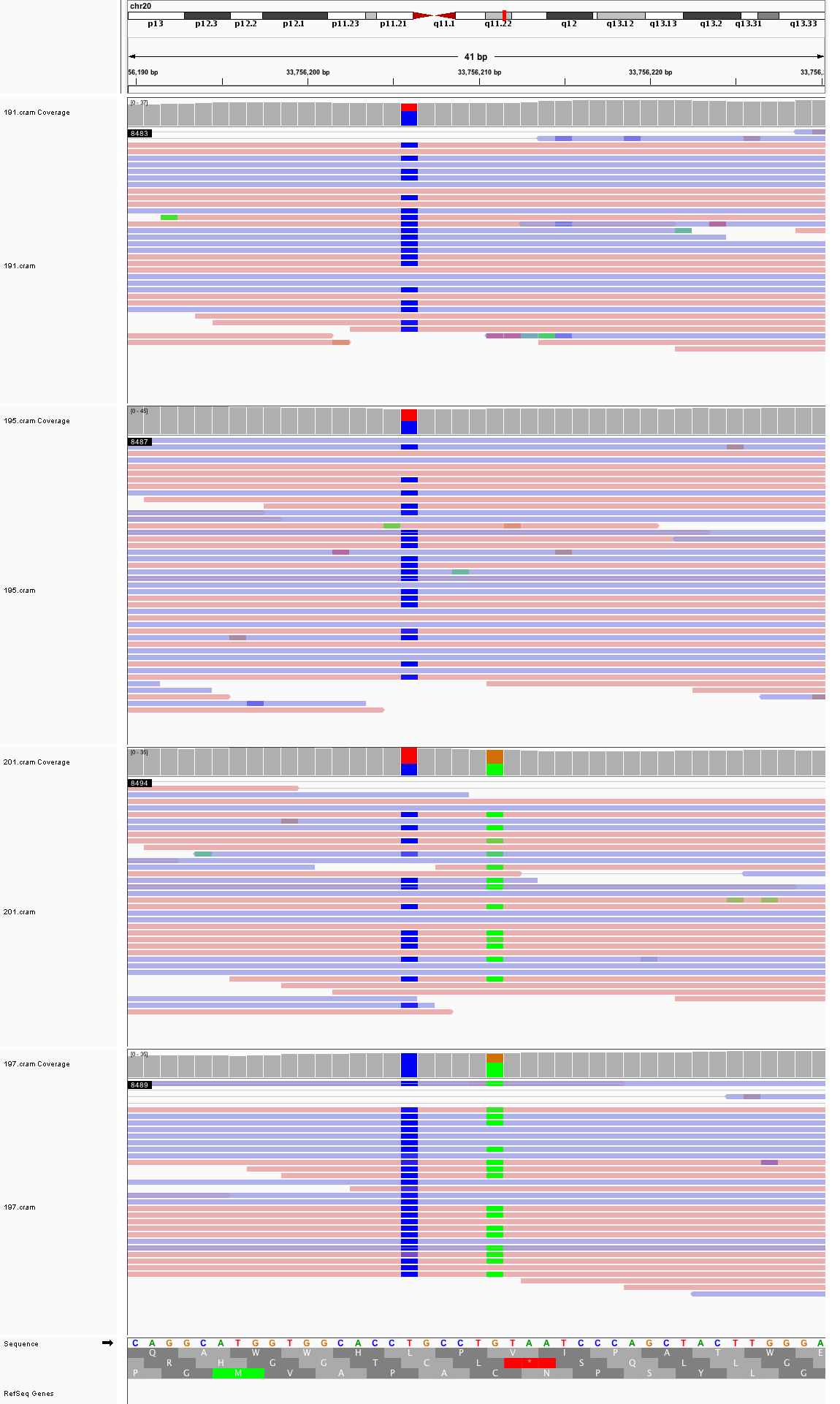

### chr20_46,790,166_46,790,206.png

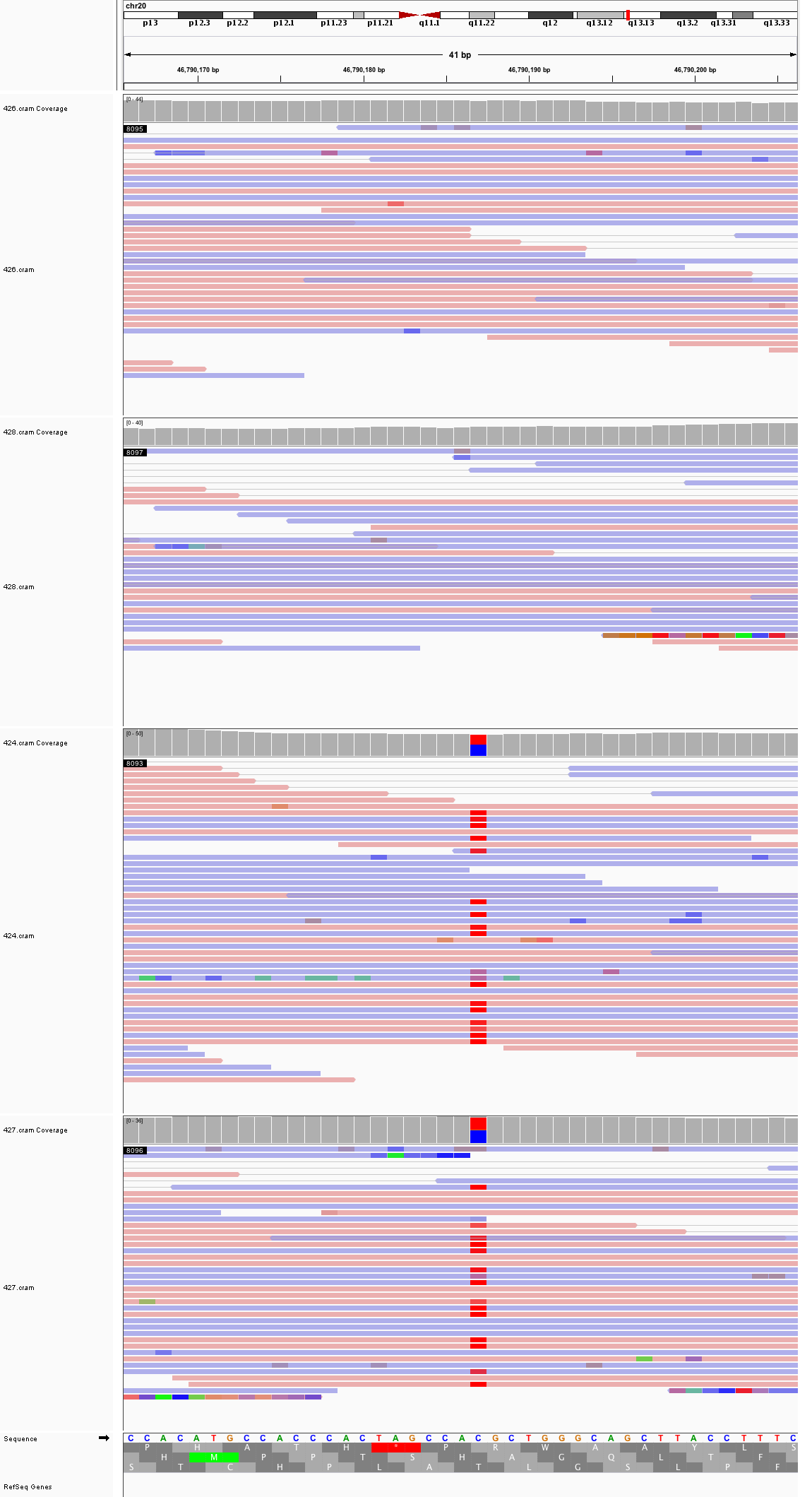

### chr21_28,658,406_28,658,446.png

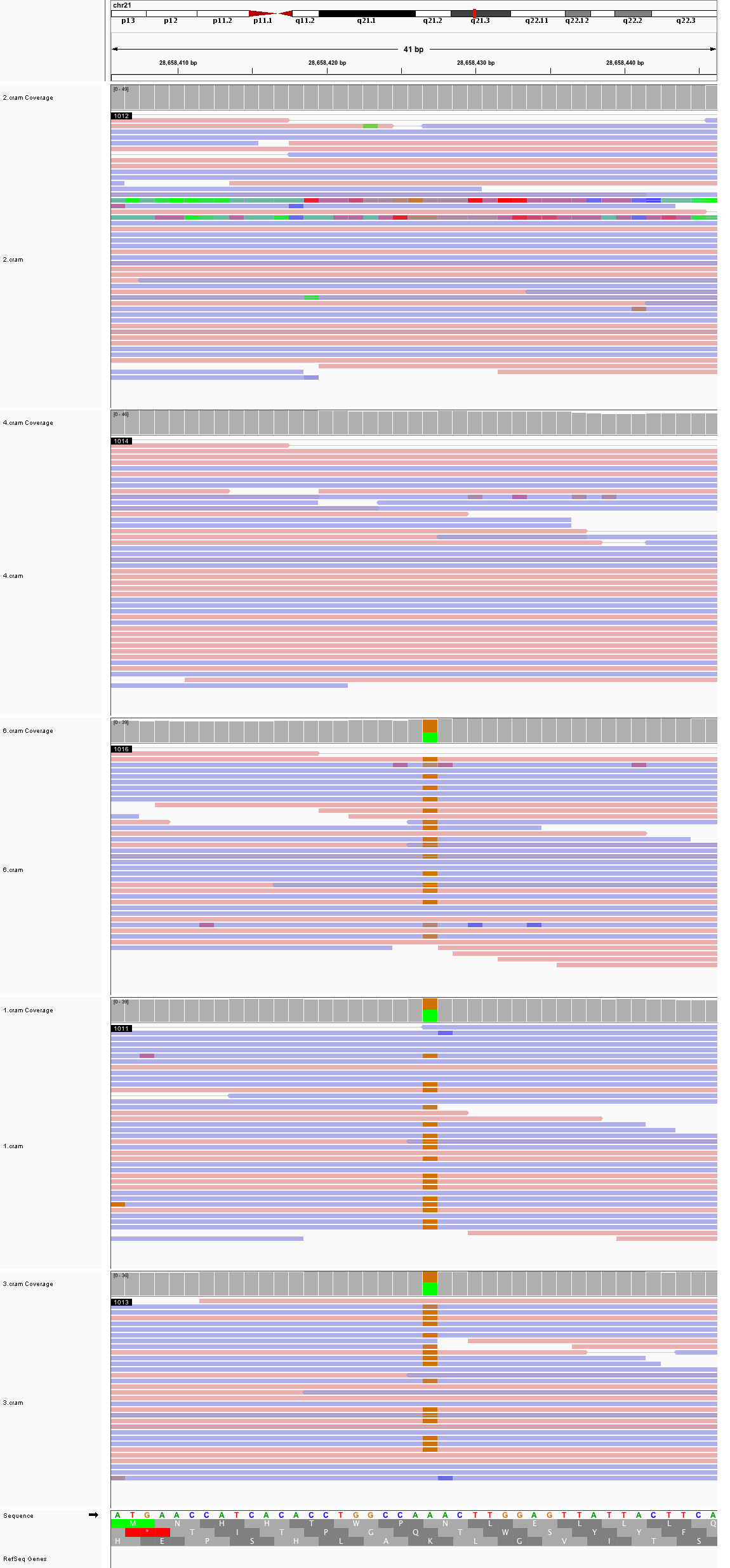

### chr22_19,776,310_19,776,350.png

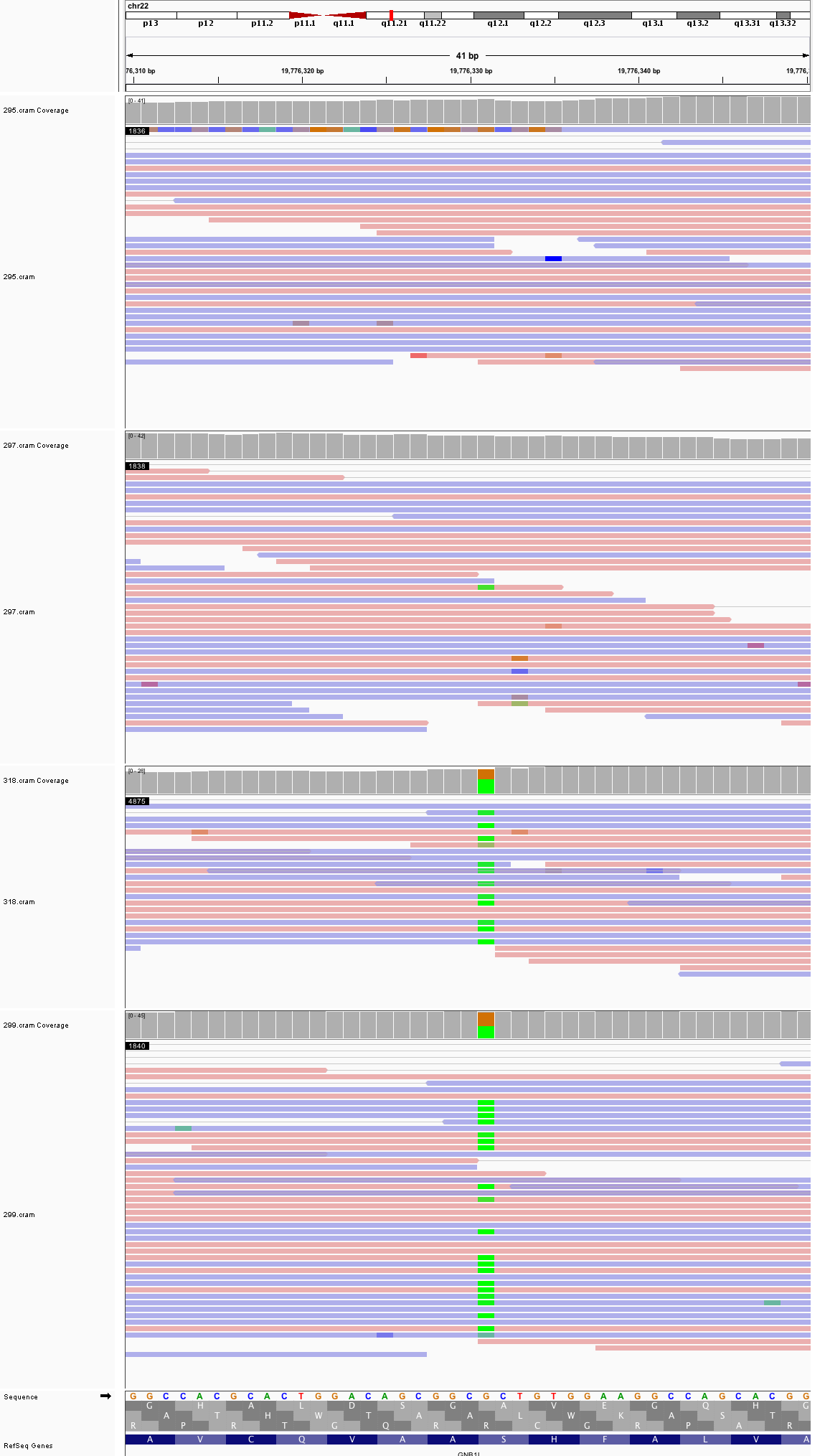

### chr22_46,889,660_46,889,700.png

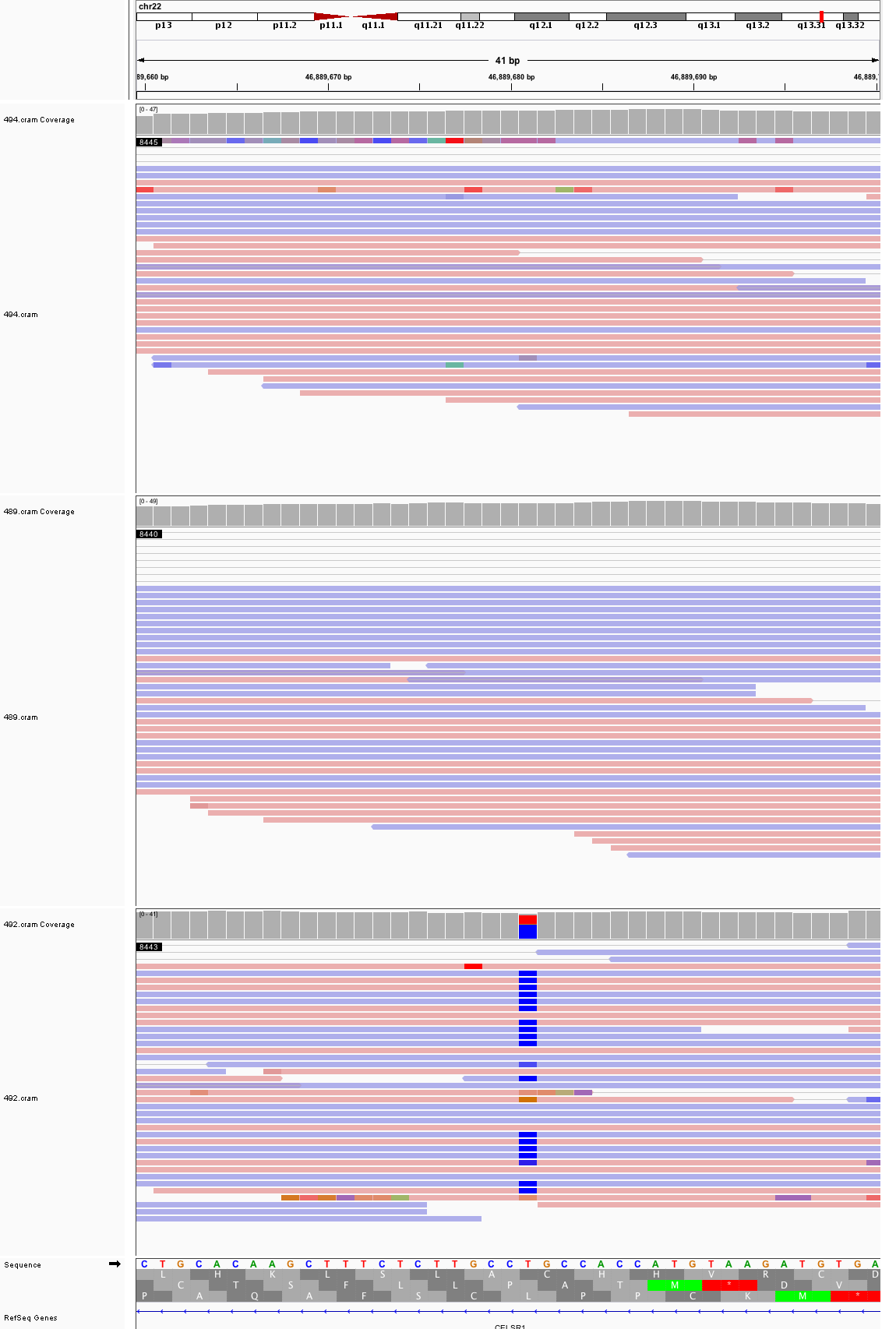

### chrX_31,992,333_31,992,373.png

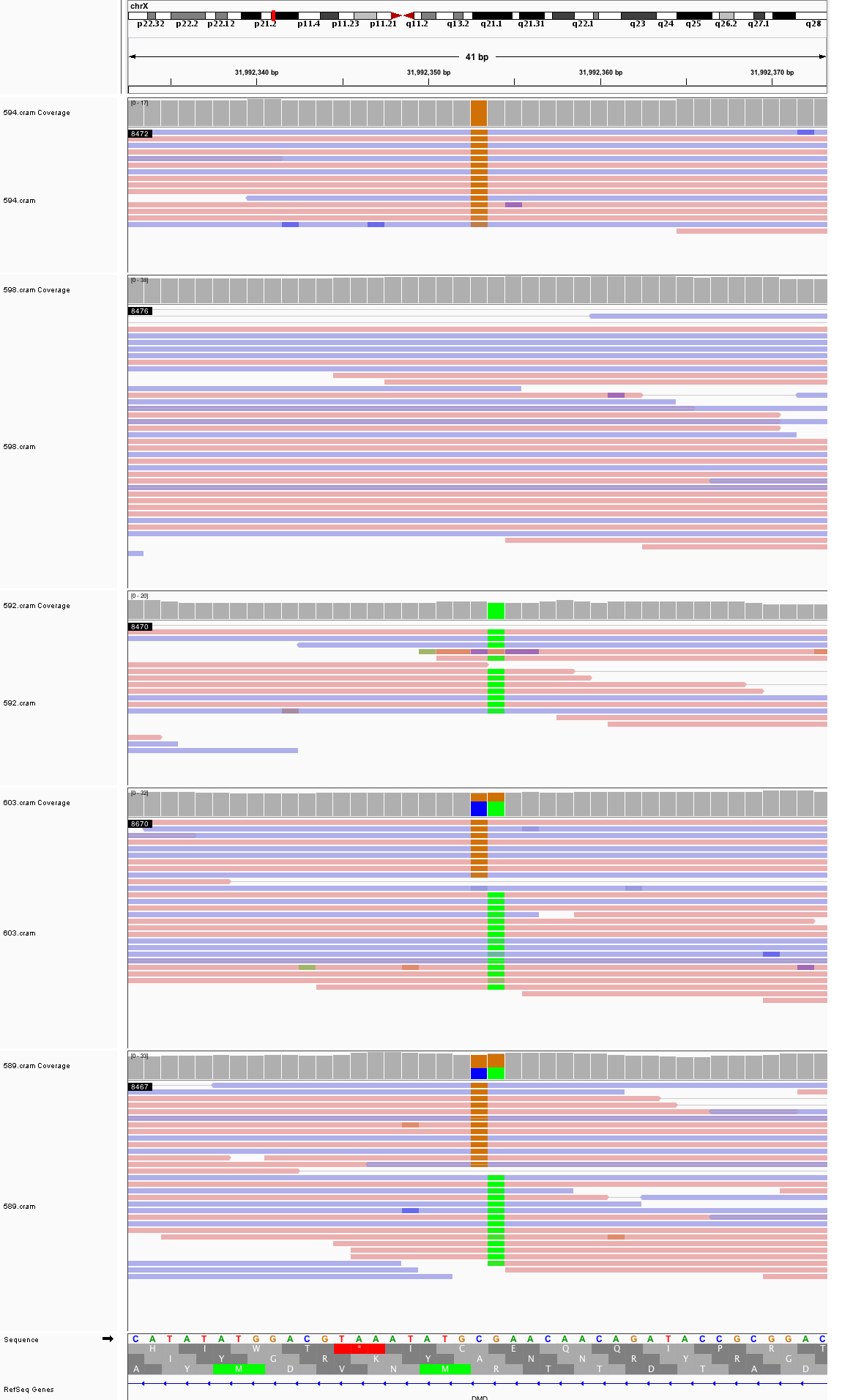

### chrX_40,397,691_40,397,731.png

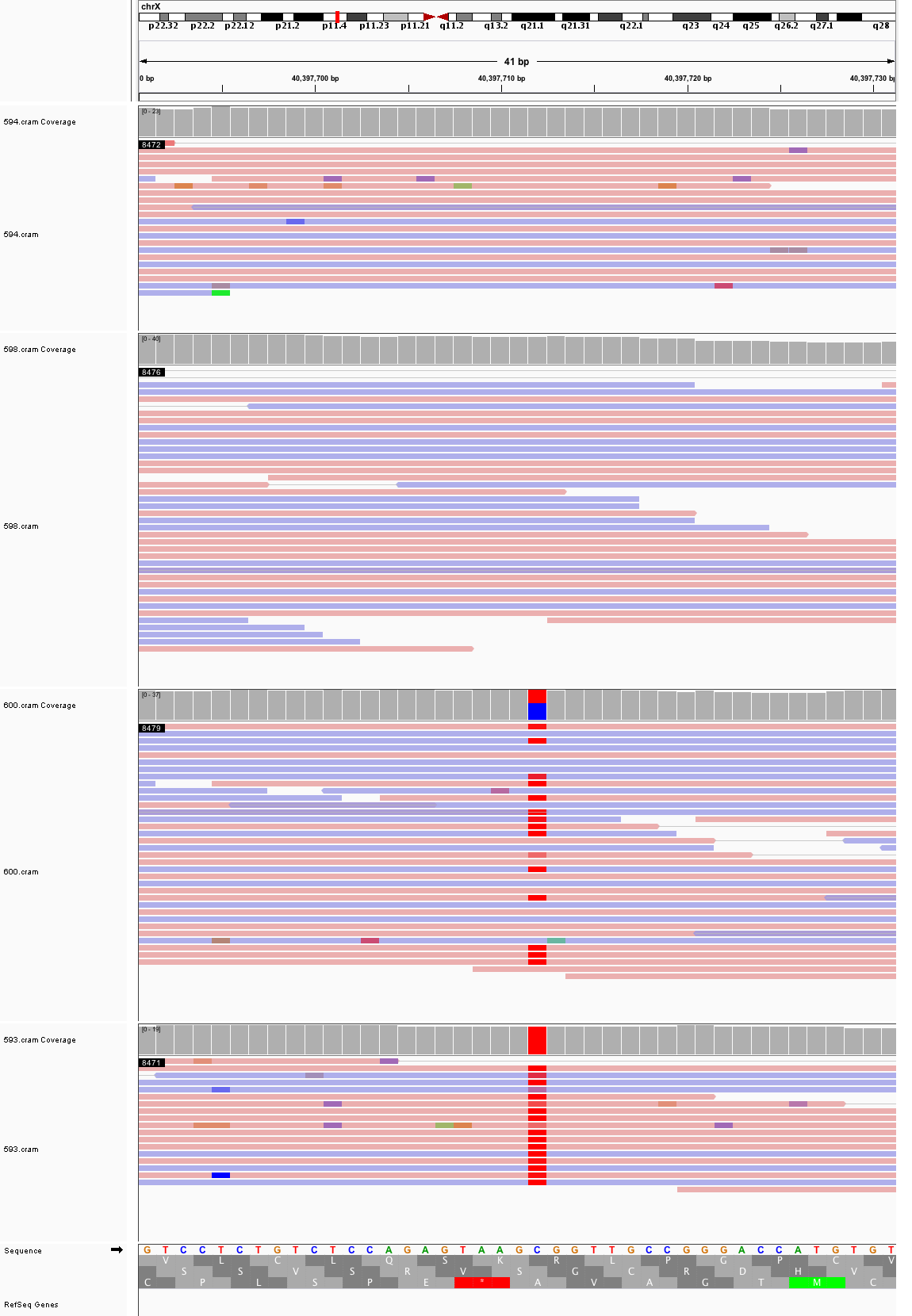

### chrX_79,157,197_79,157,237.png

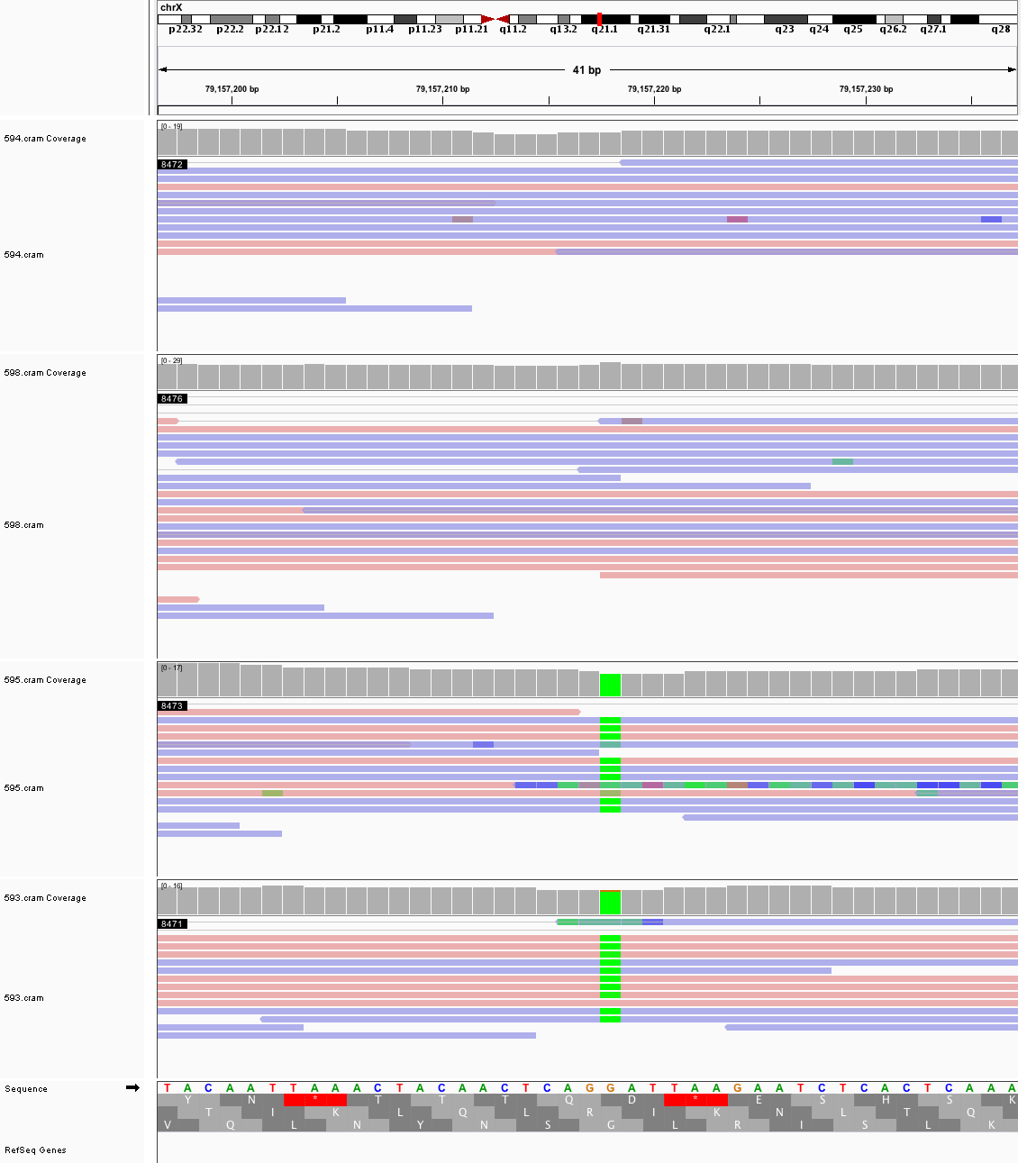

### chrX_126,409,263_126,409,303.png

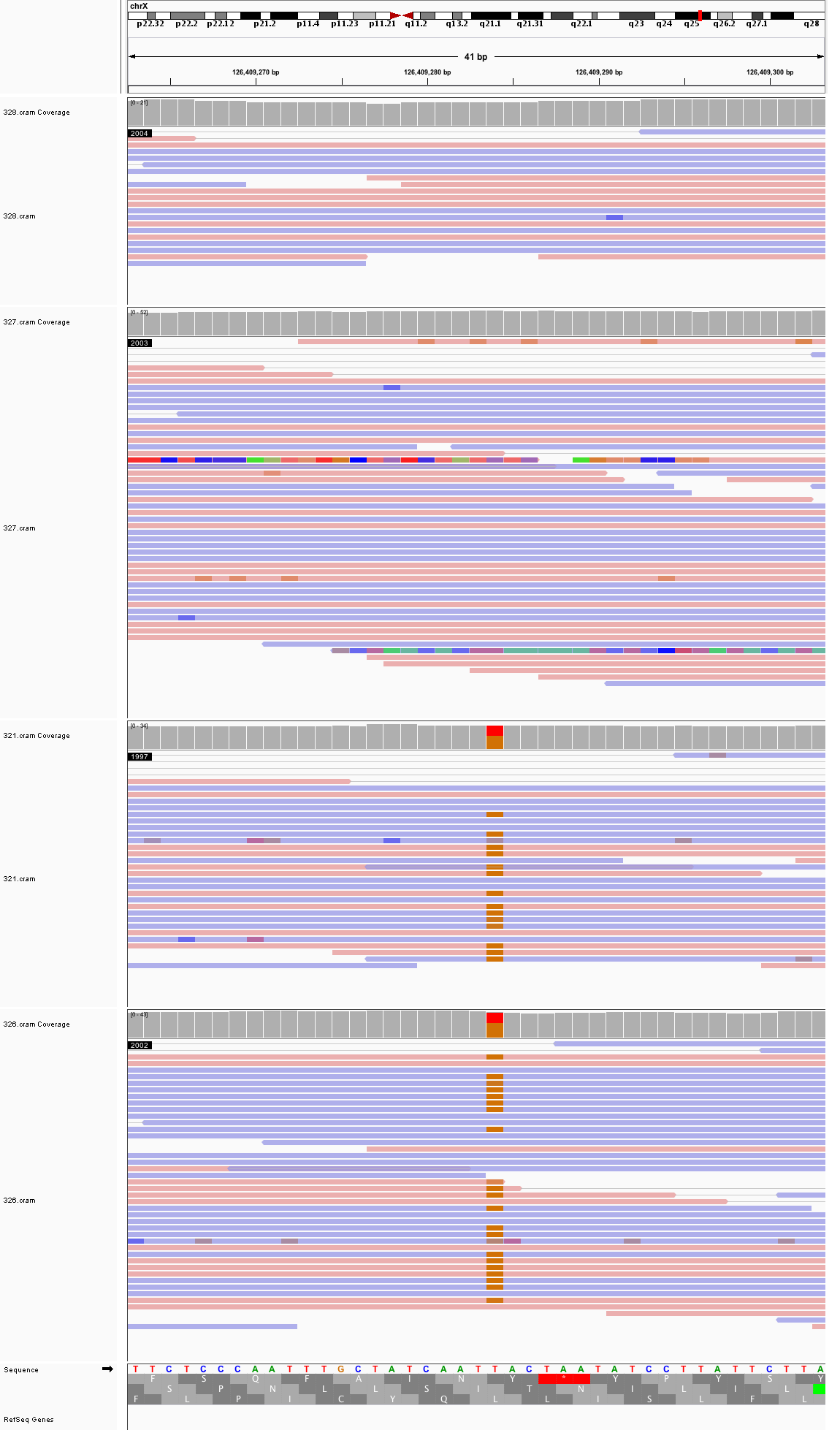
