## Supplementary figures and images for "Large, three-generation CEPH families reveal post-zygotic mosaicism and variability in germline mutation accumulation"

### chr1_2,860,815_2,860,855.png

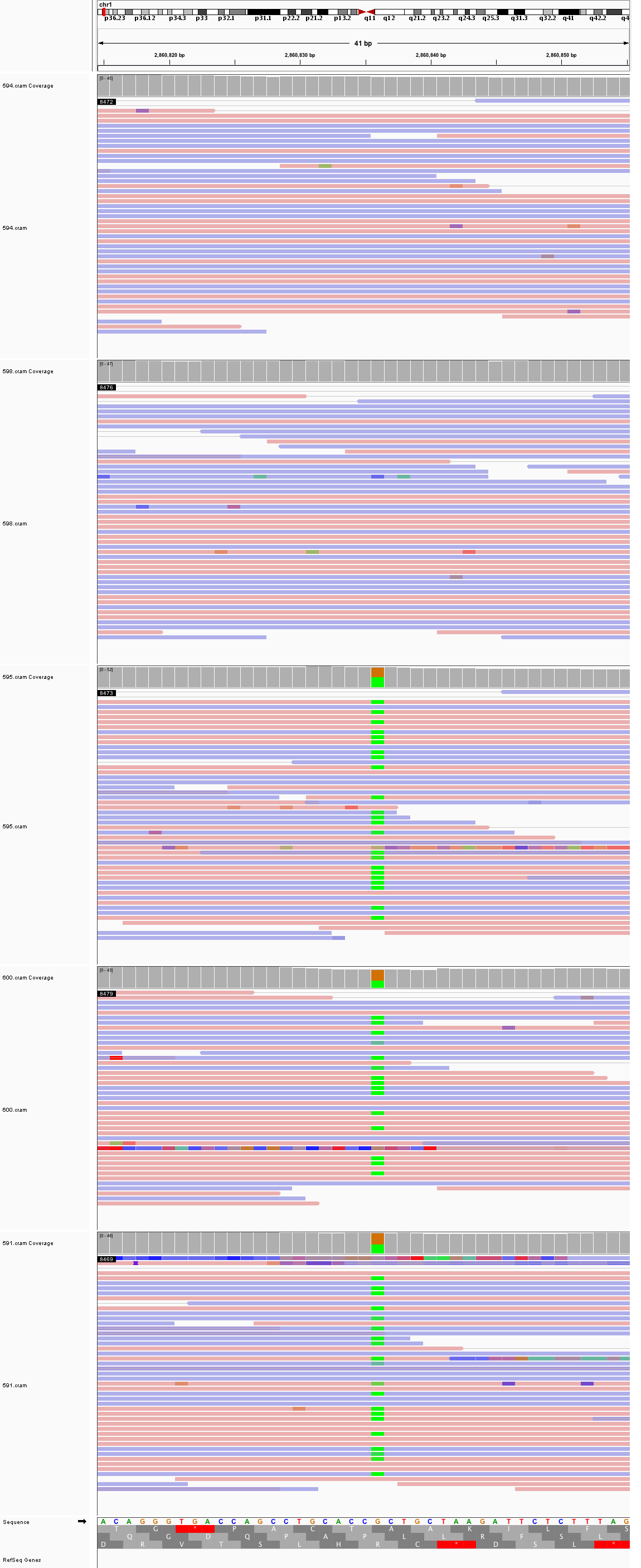

### chr1_69,846,519_69,846,559.png

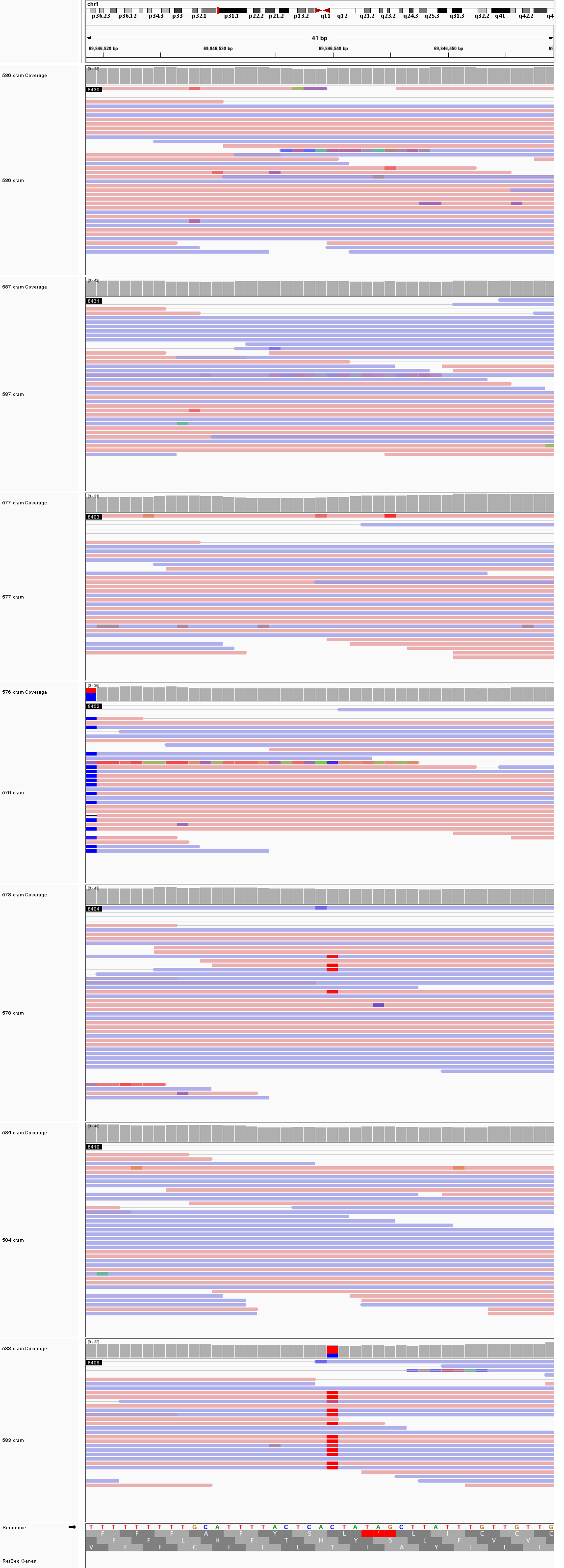

### chr1_78,000,486_78,000,526.png

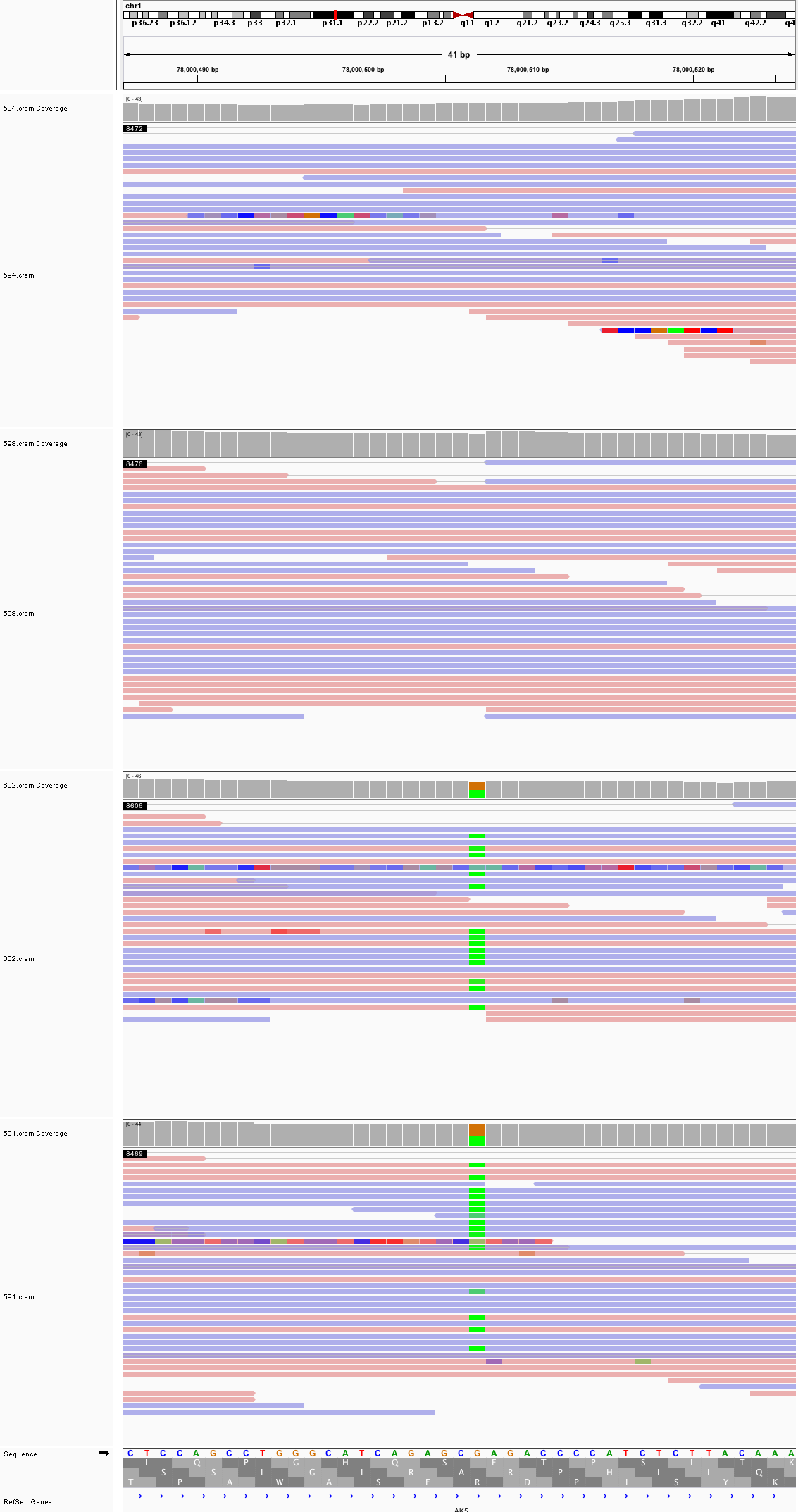

### chr1_86,573,074_86,573,114.png

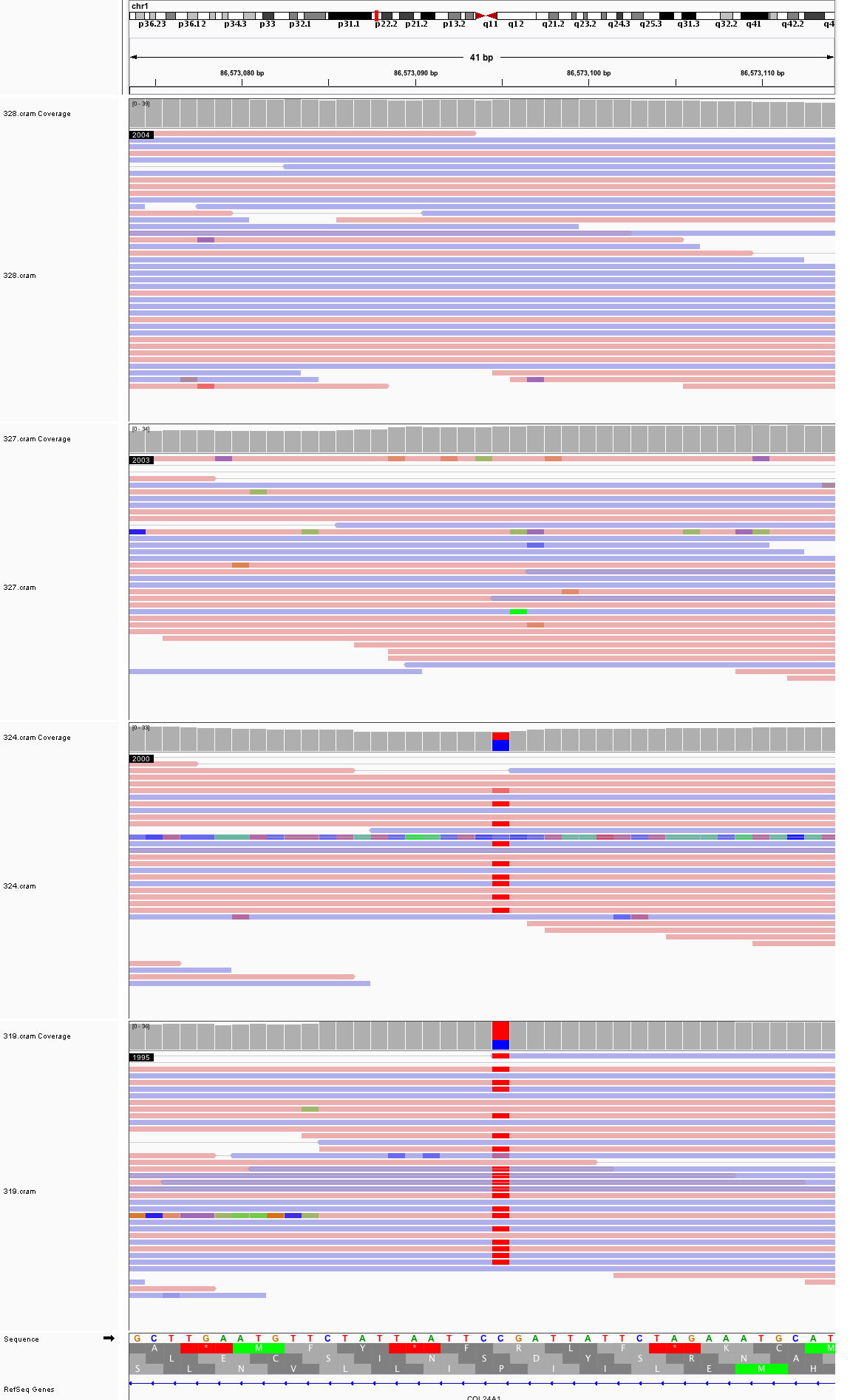

### chr1_103,732,753_103,732,793.png

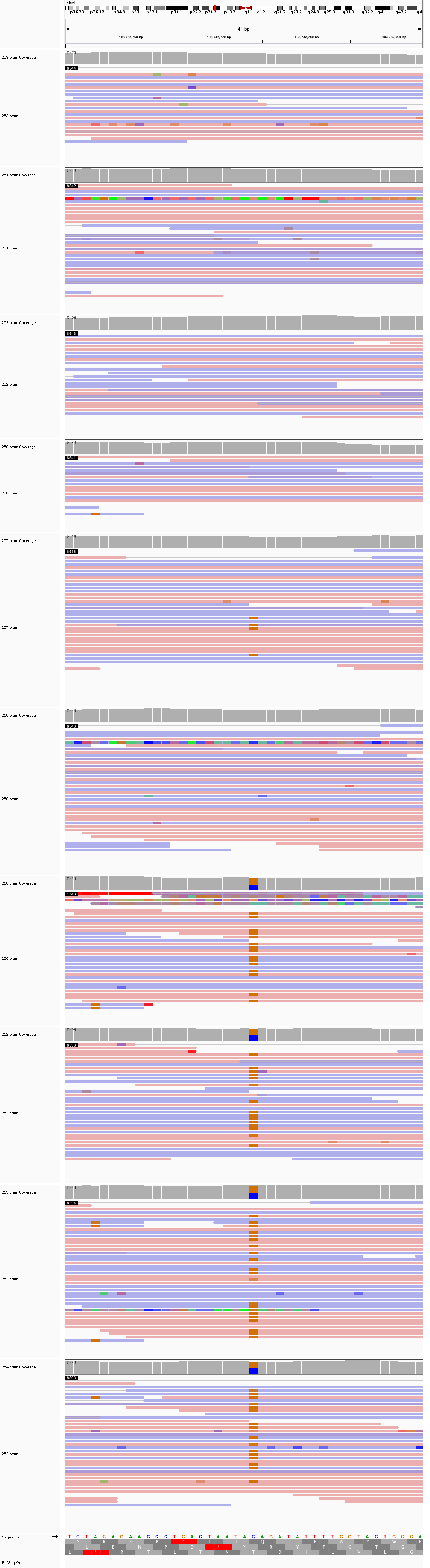

### chr1_155,733,697_155,733,737.png

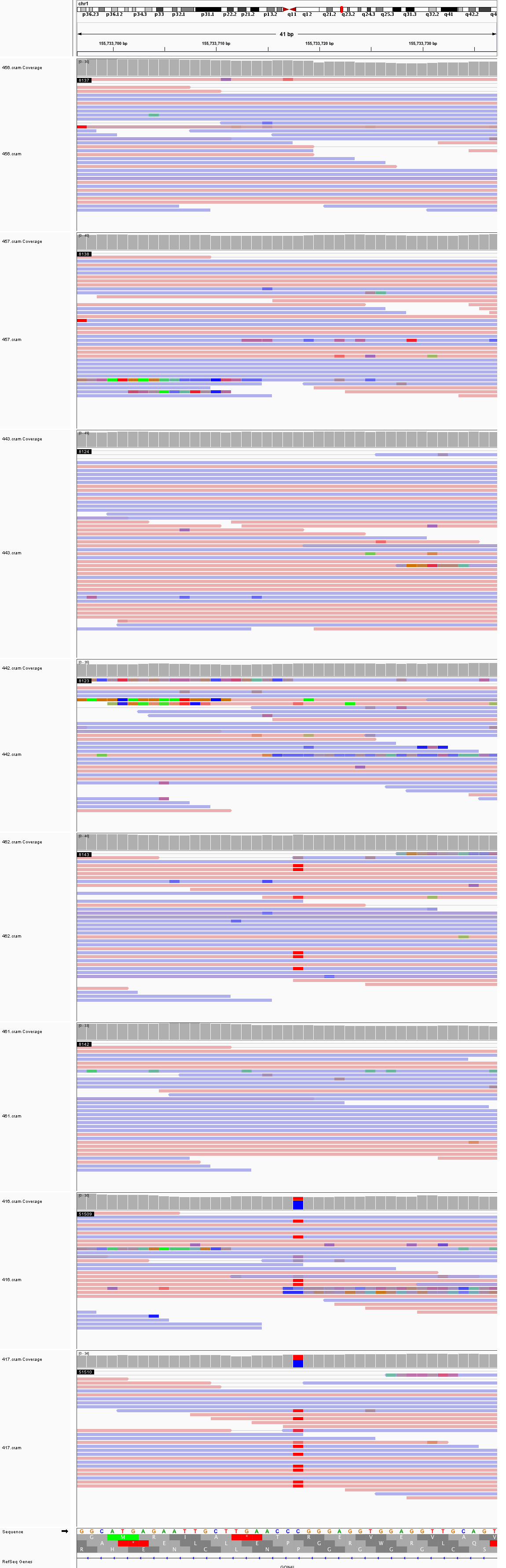

### chr1_173,846,775_173,846,815.png

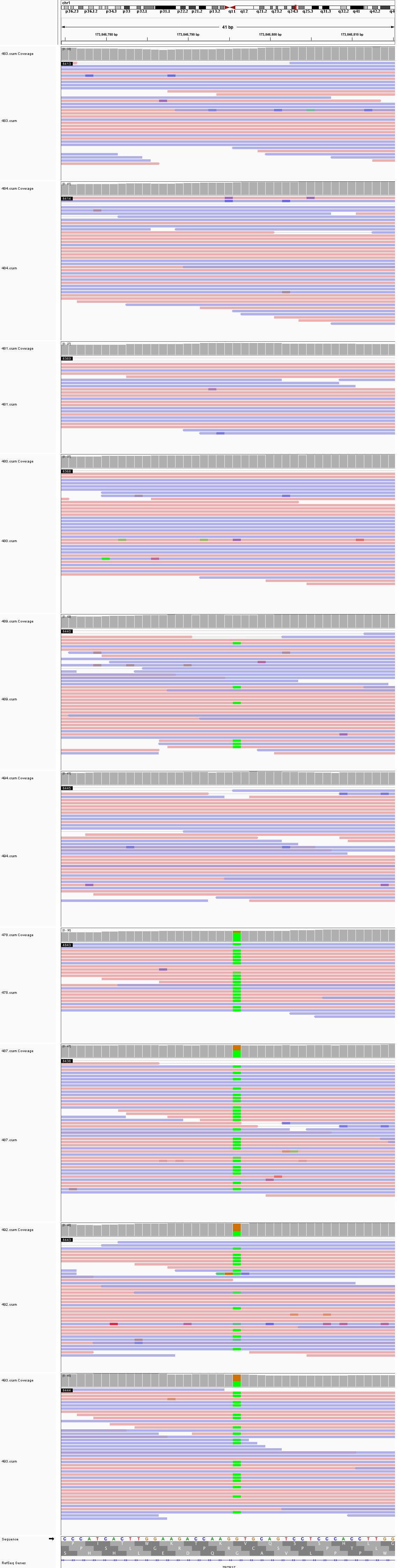

### chr1_176,263,747_176,263,787.png

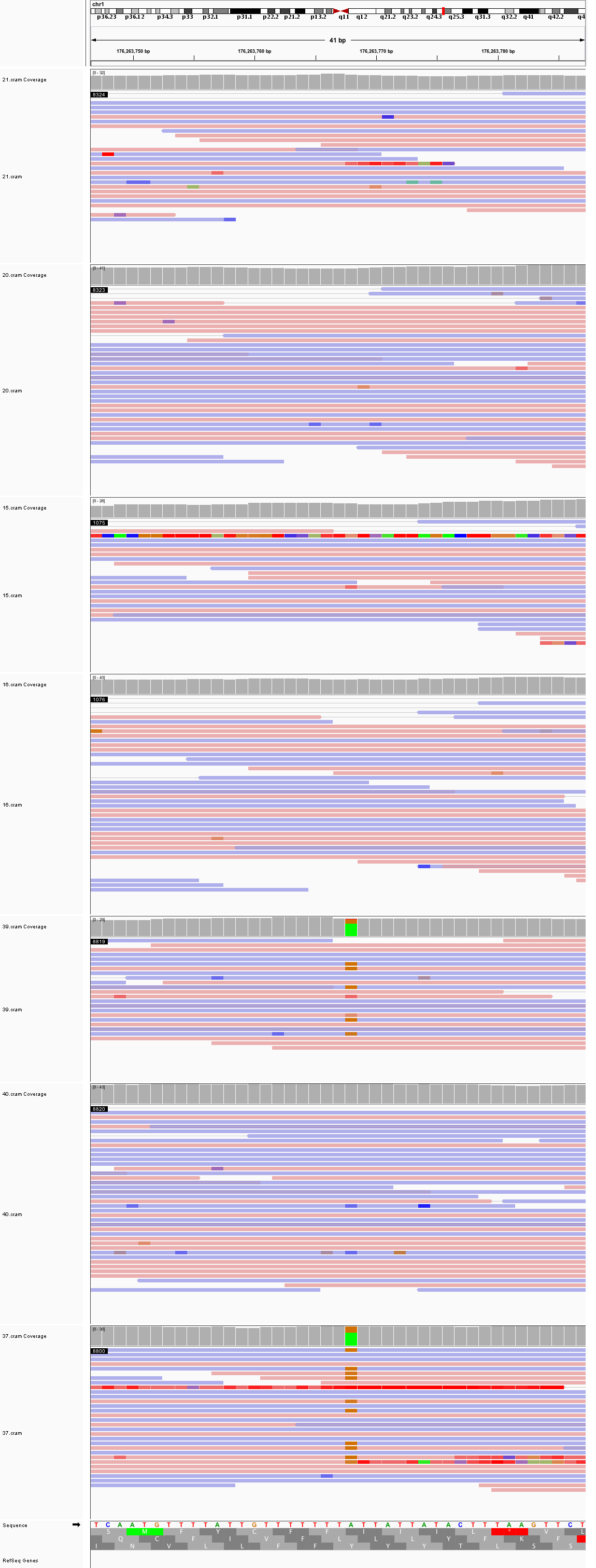

### chr1_183,870,398_183,870,438.png

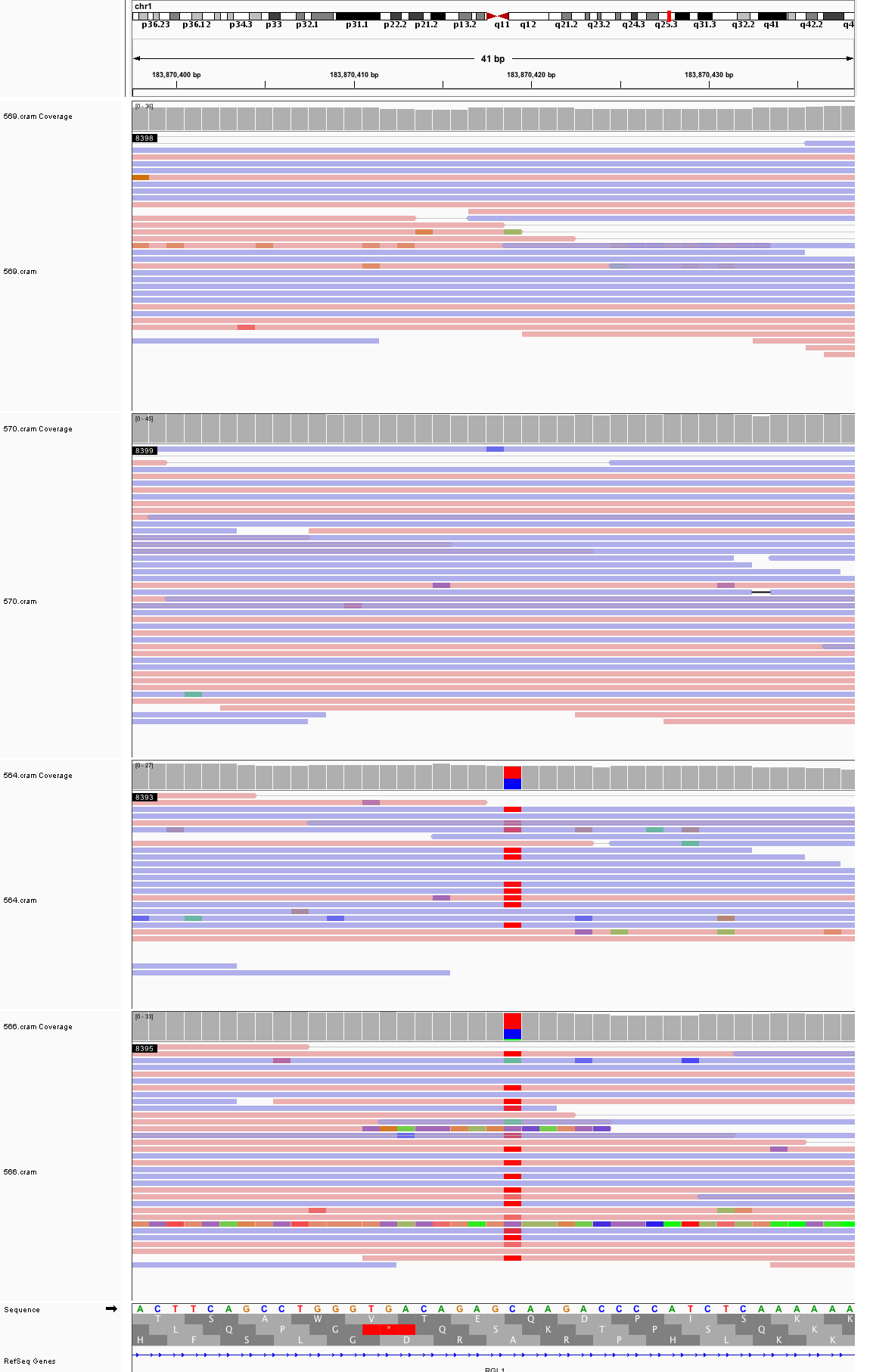

### chr1_223,339,016_223,339,056.png

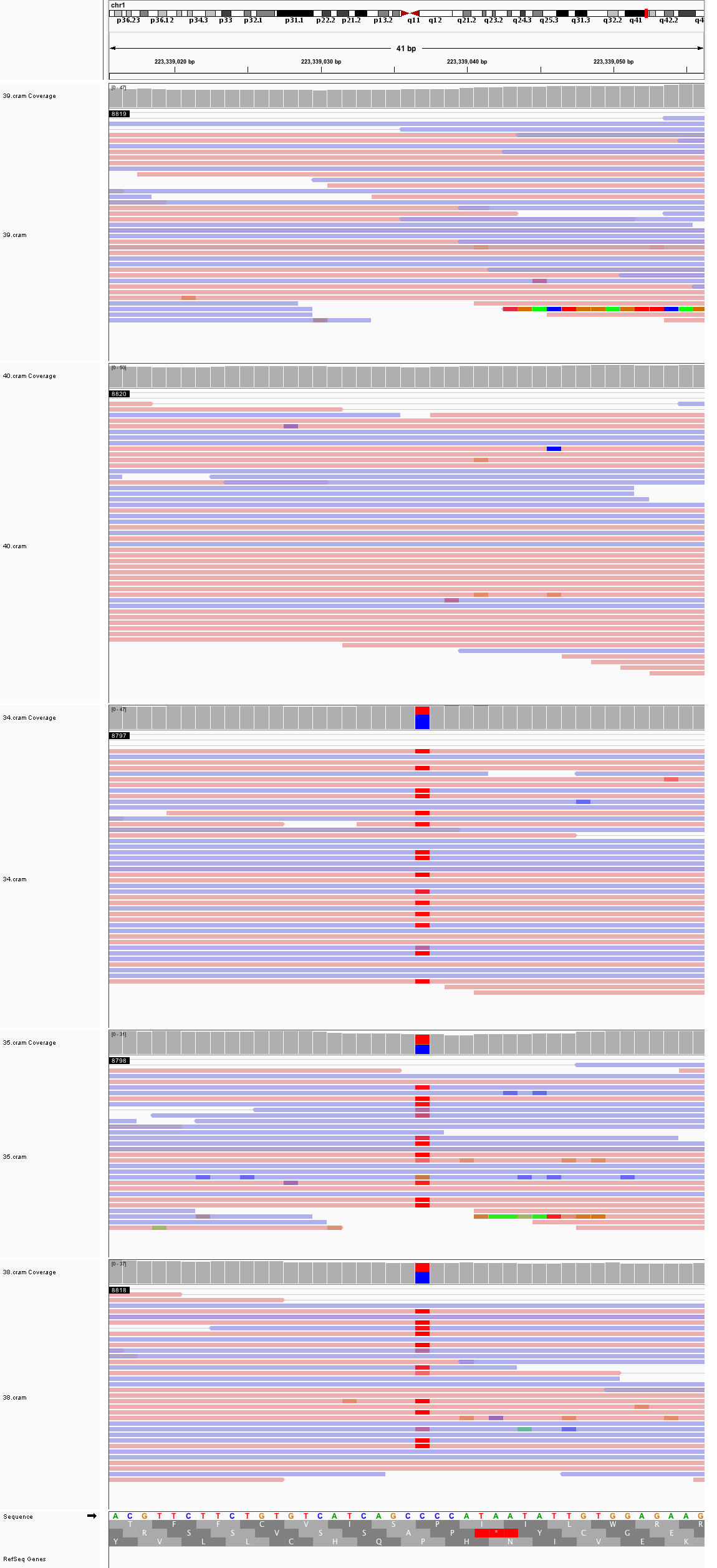

### chr1_224,535,552_224,535,592.png

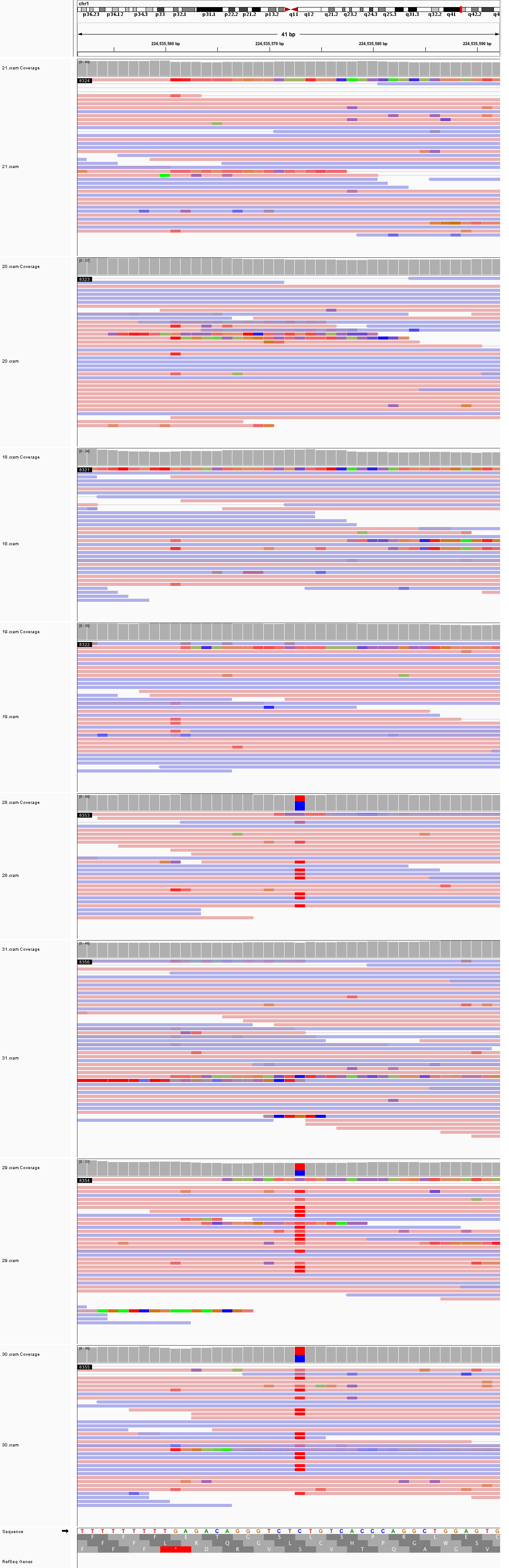

### chr2_6,464,742_6,464,782.png

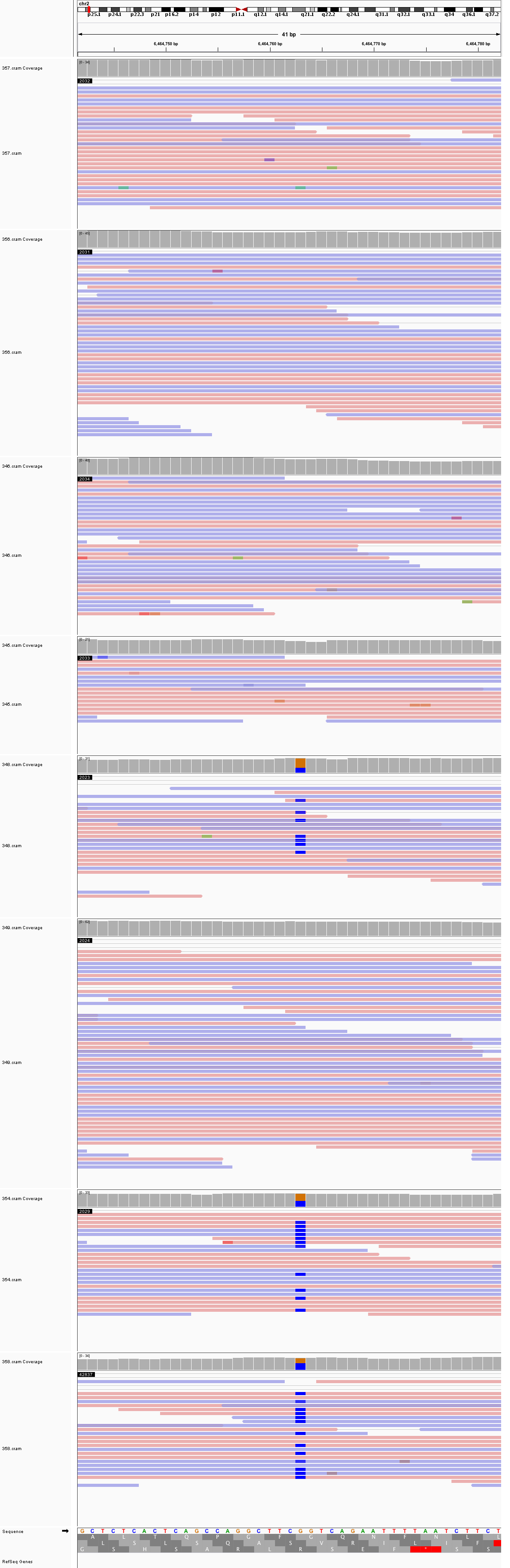

### chr2_8,629,119_8,629,159.png

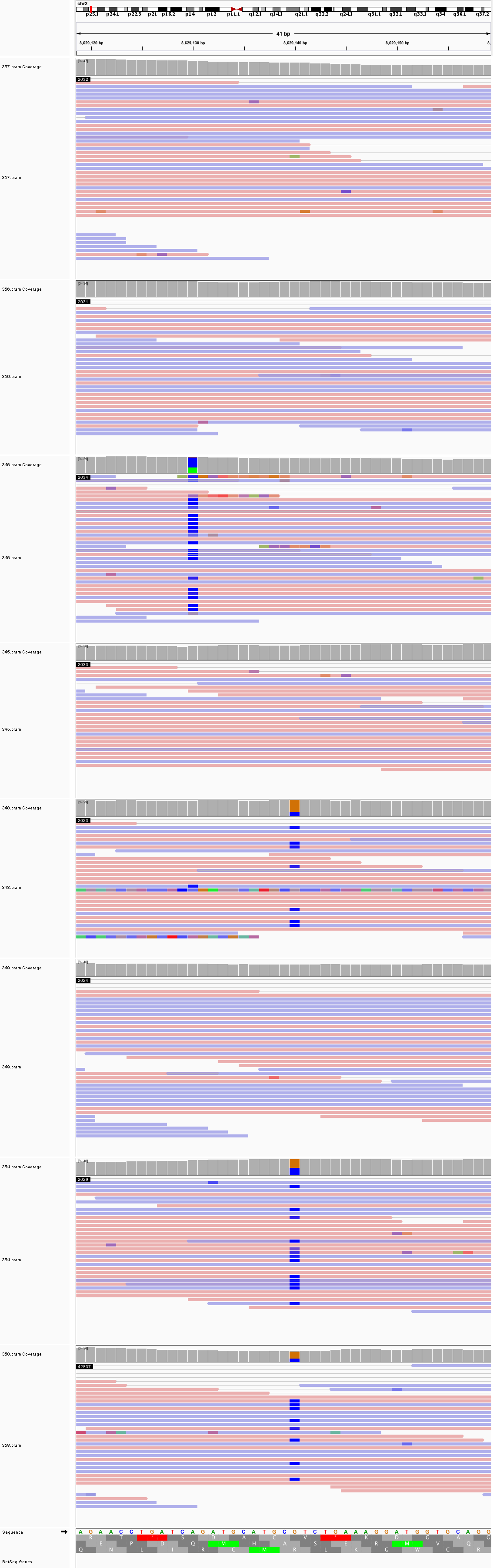

### chr2_32,999,615_32,999,655.png

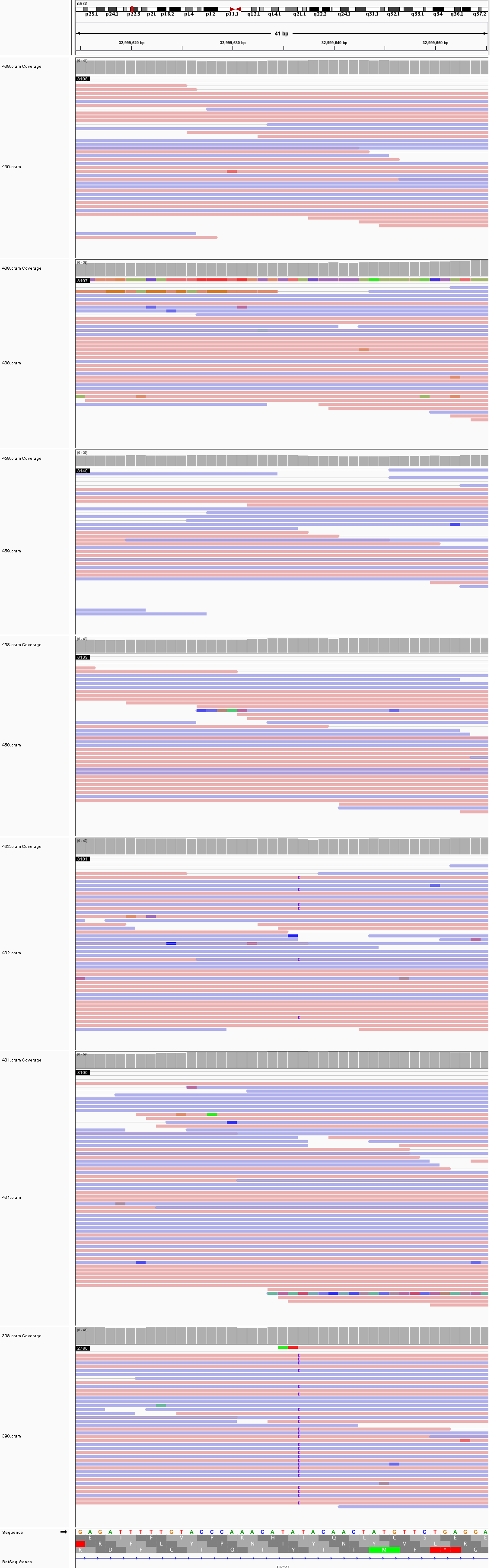

### chr2_77,732,491_77,732,531.png

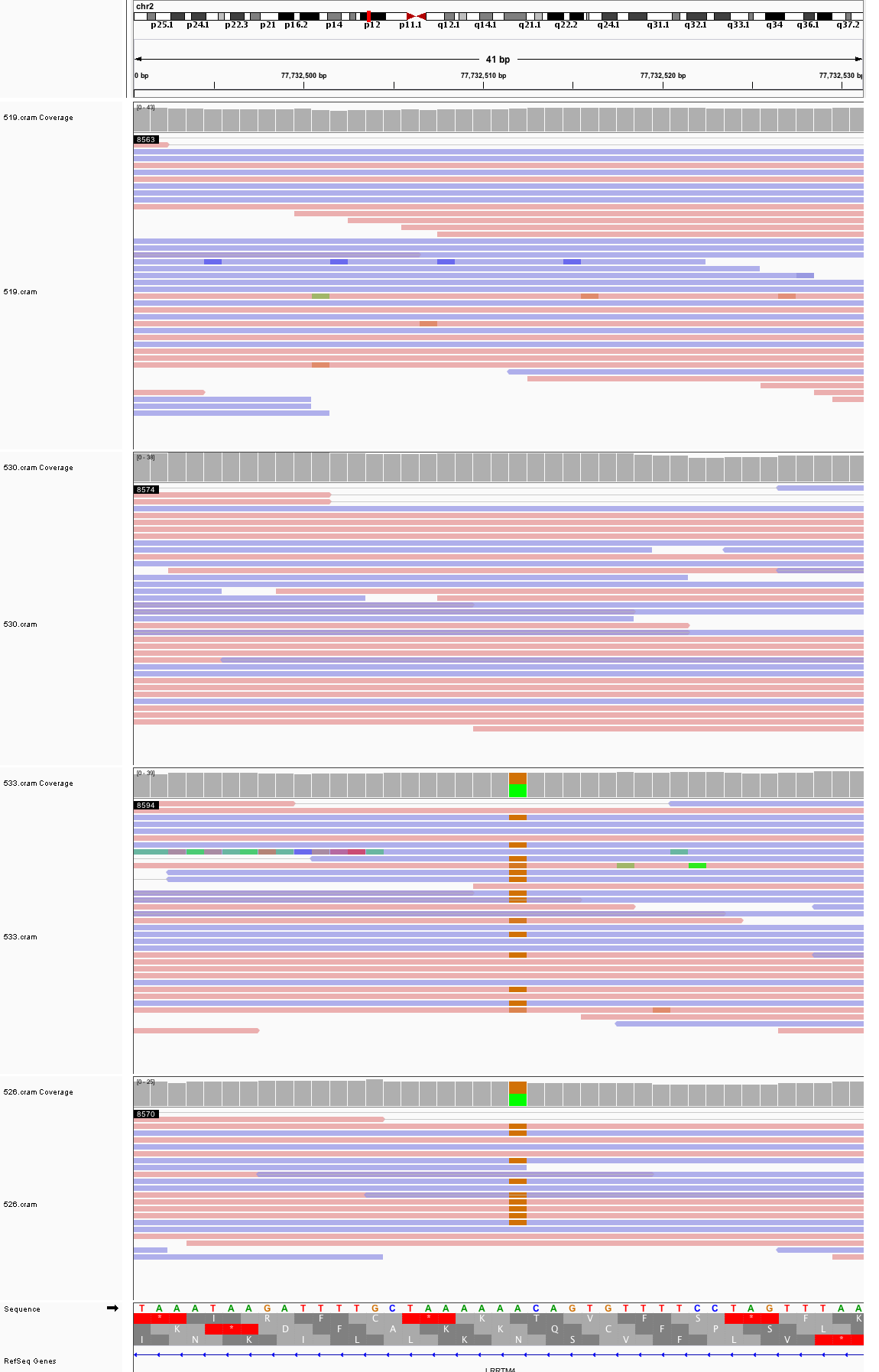

### chr2_112,886,798_112,886,838.png

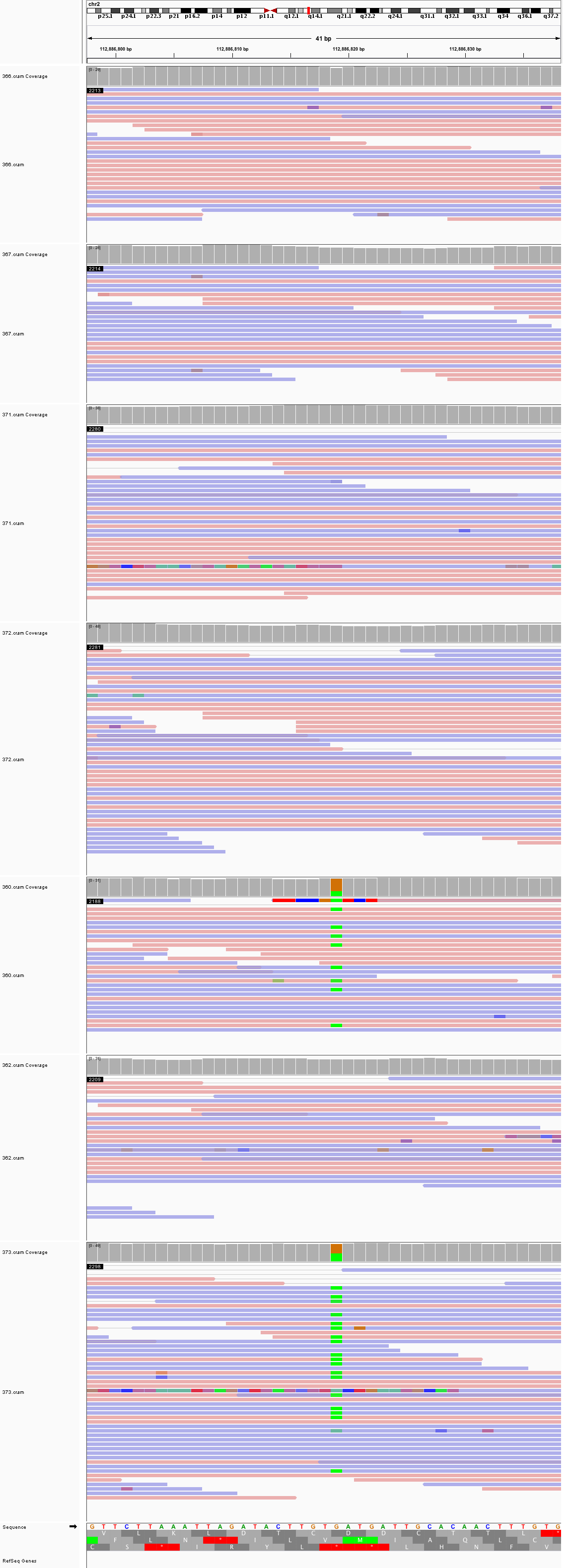

### chr2_123,683,033_123,683,073.png

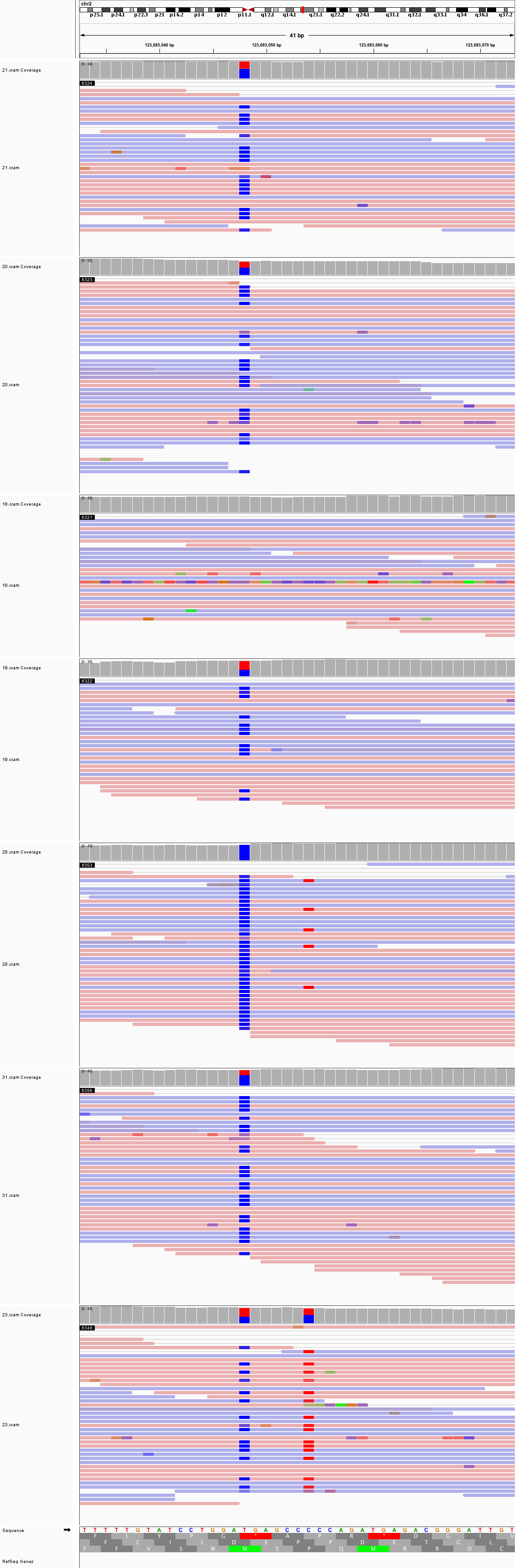

### chr3_38,755,474_38,755,514.png

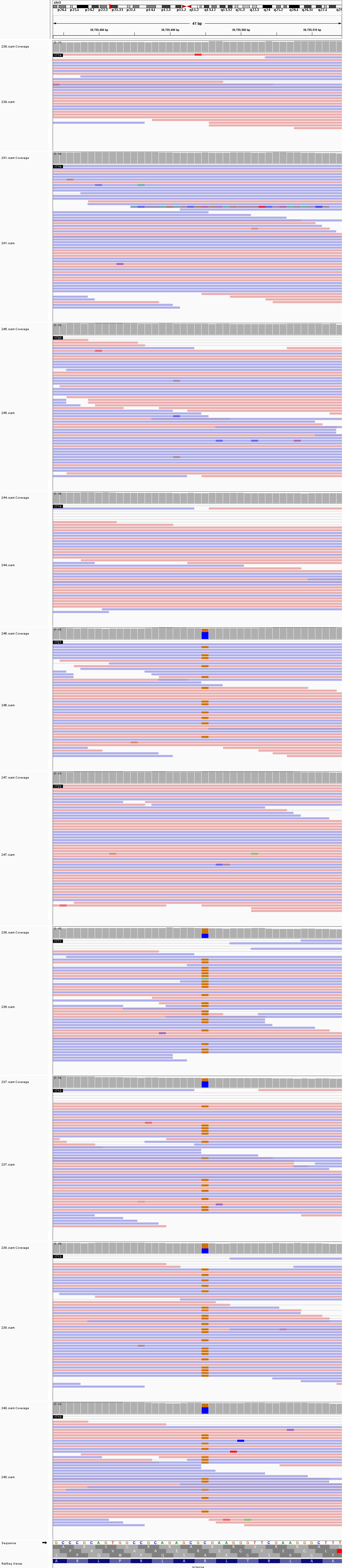

### chr3_52,255,509_52,255,549.png

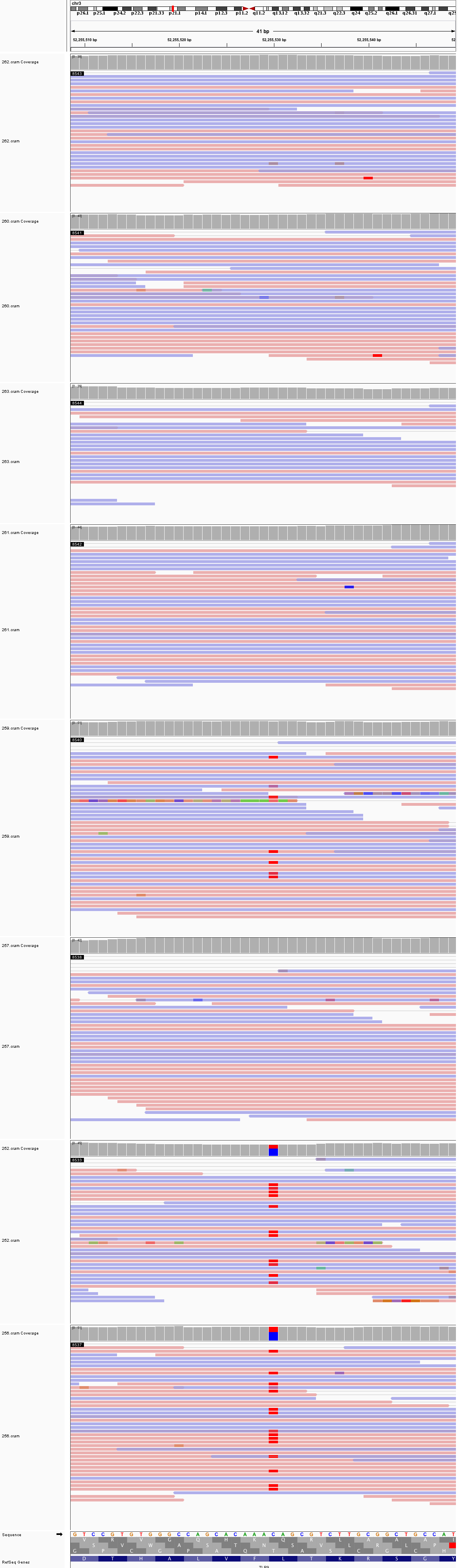

### chr3_103,494,592_103,494,632.png

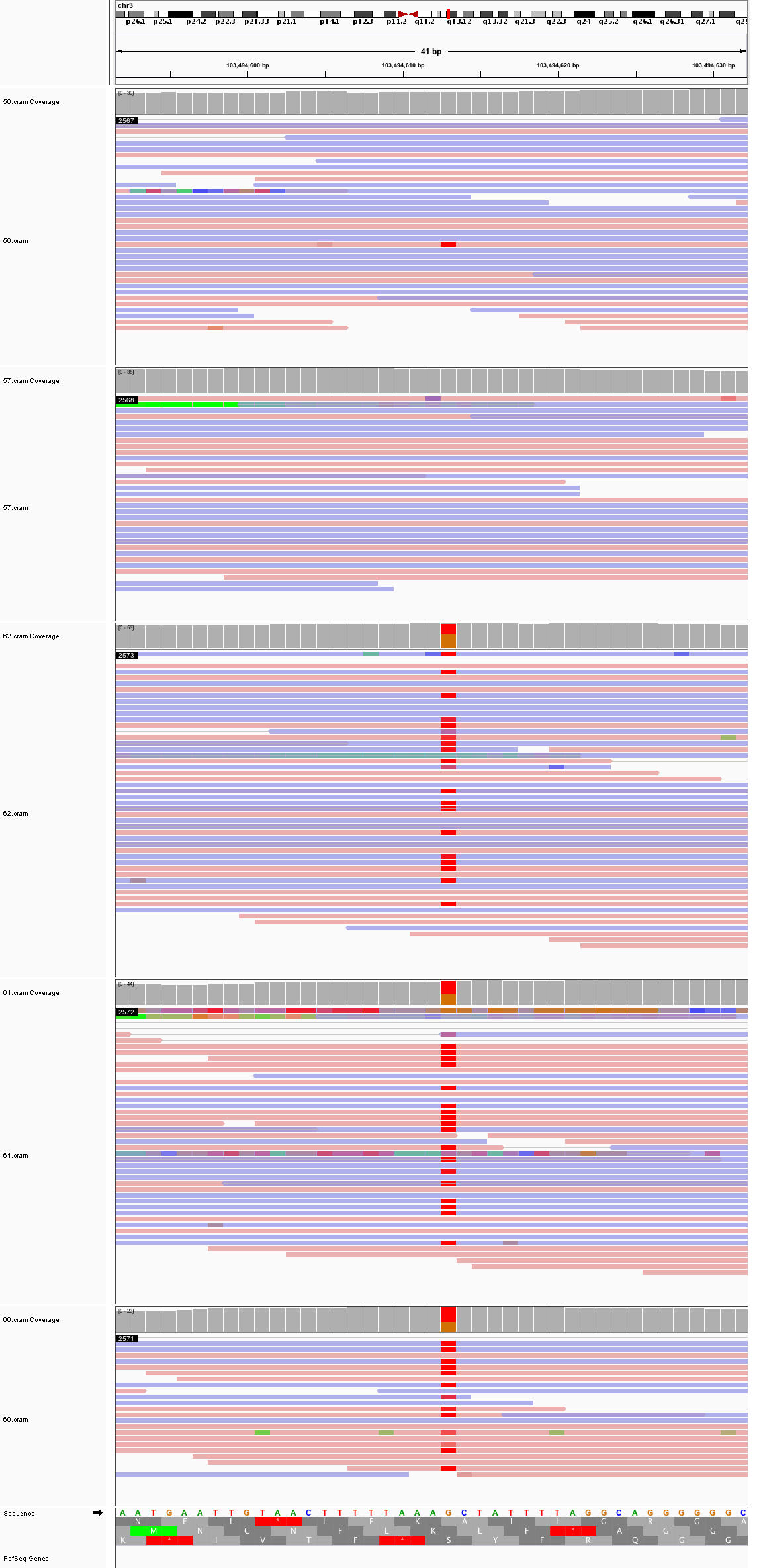

### chr3_121,250,881_121,250,921.png

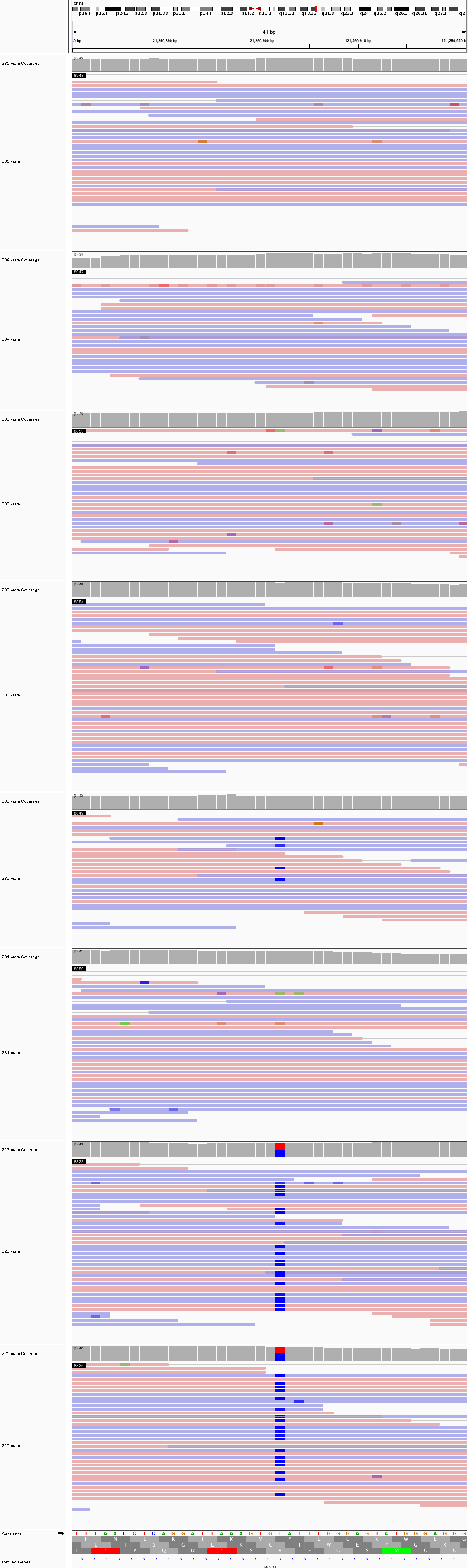

### chr3_136,173,285_136,173,325.png

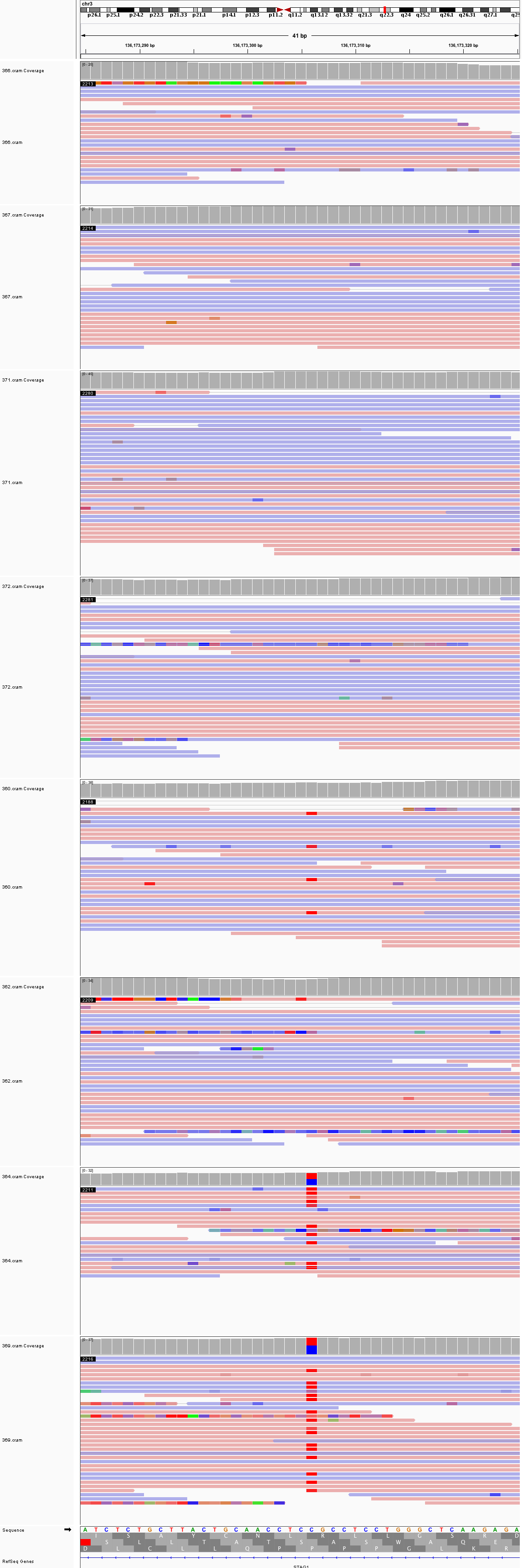

### chr3_148,974,042_148,974,082.png

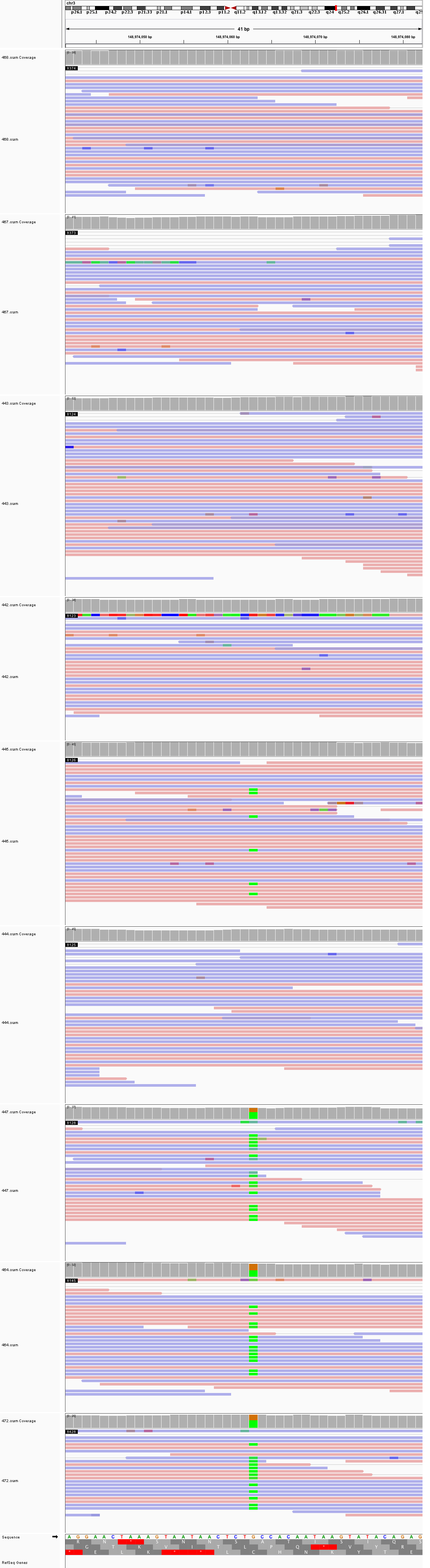

### chr3_180,351,909_180,351,949.png

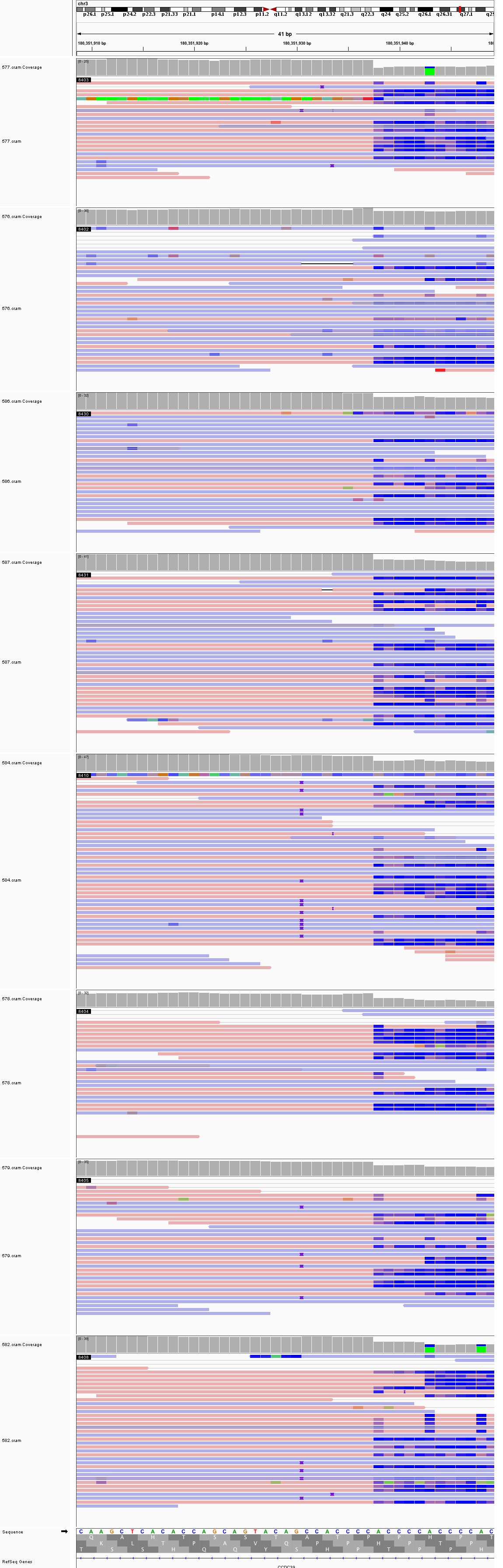

### chr4_60,477,654_60,477,694.png

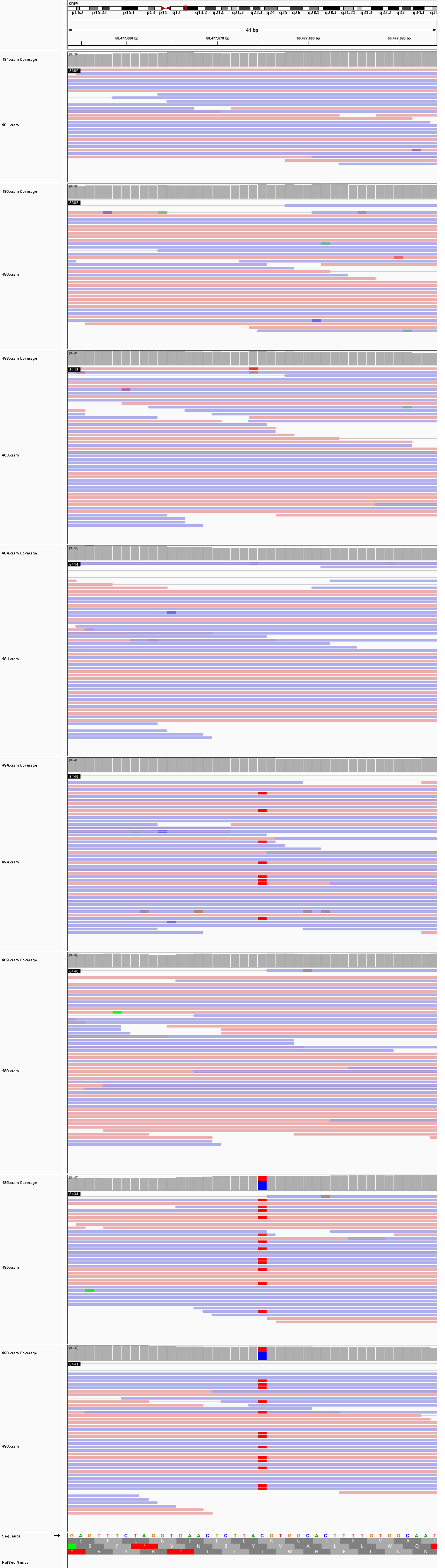

### chr4_61,807,494_61,807,534.png

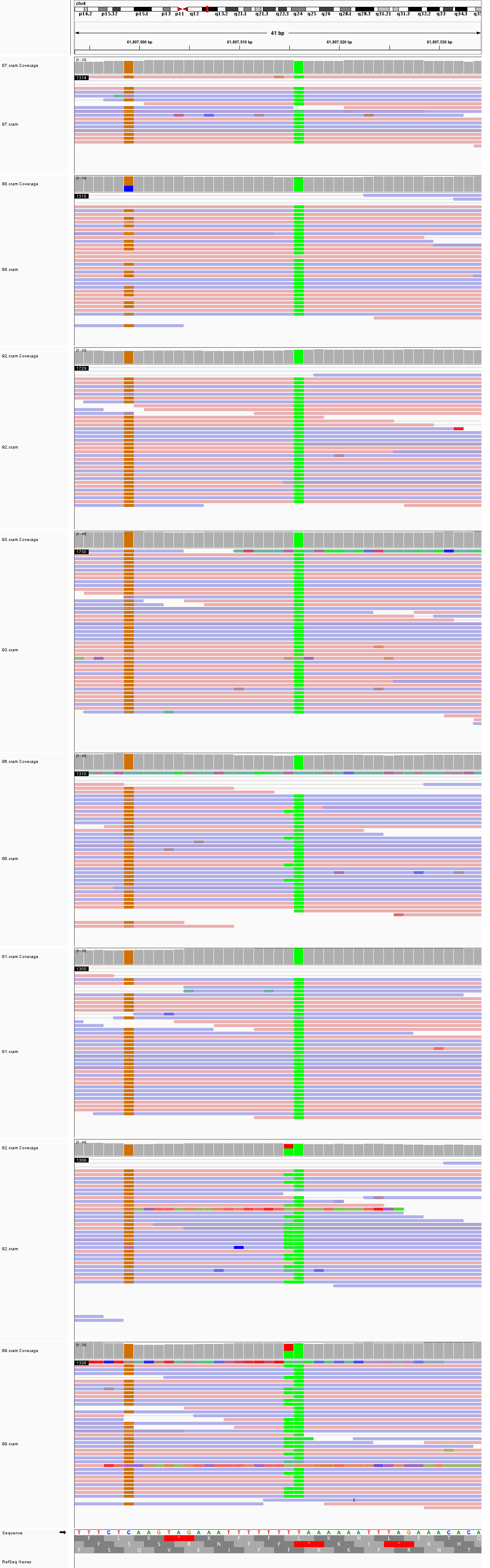

### chr4_65,725,295_65,725,335.png

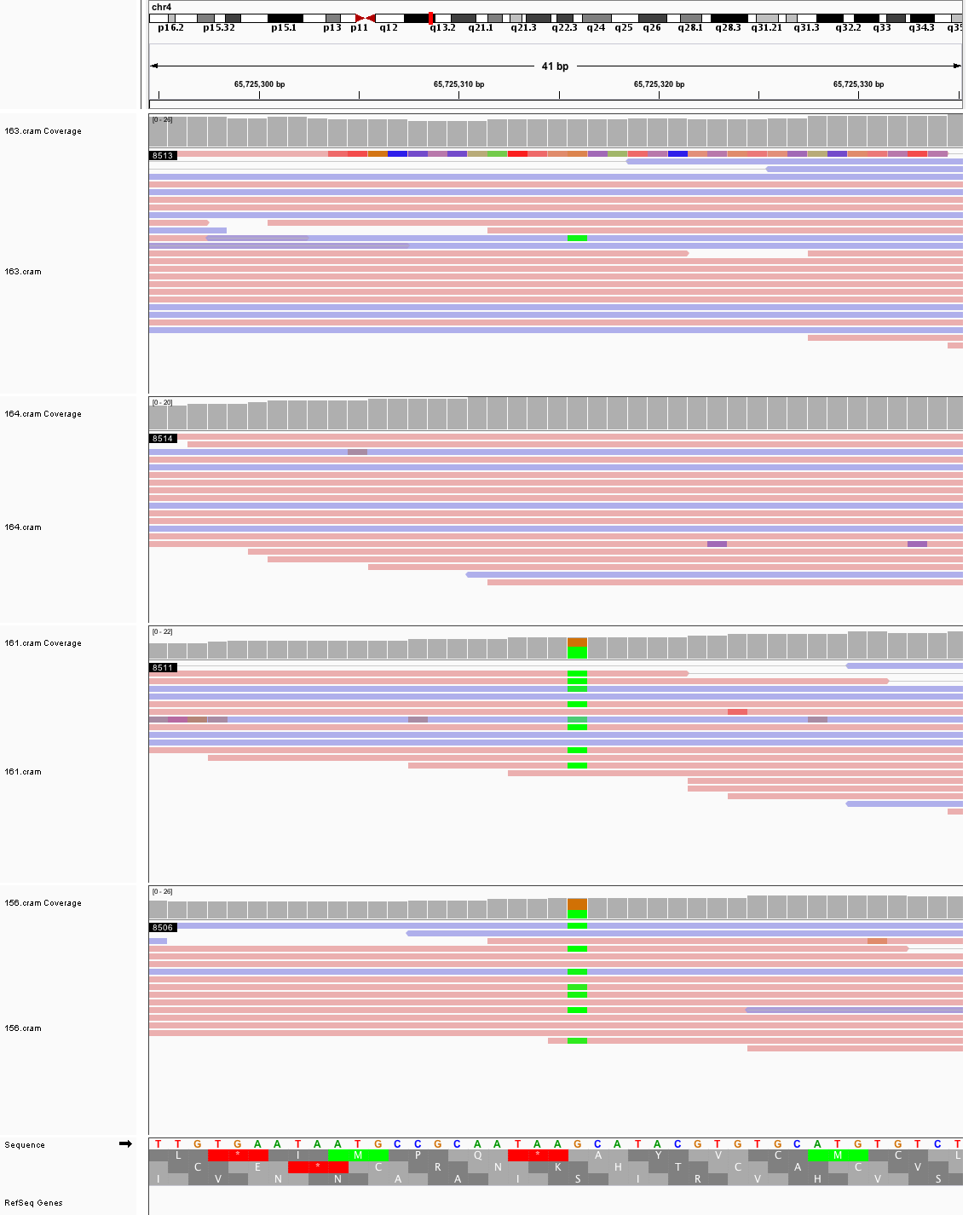

### chr4_148,409,376_148,409,416.png

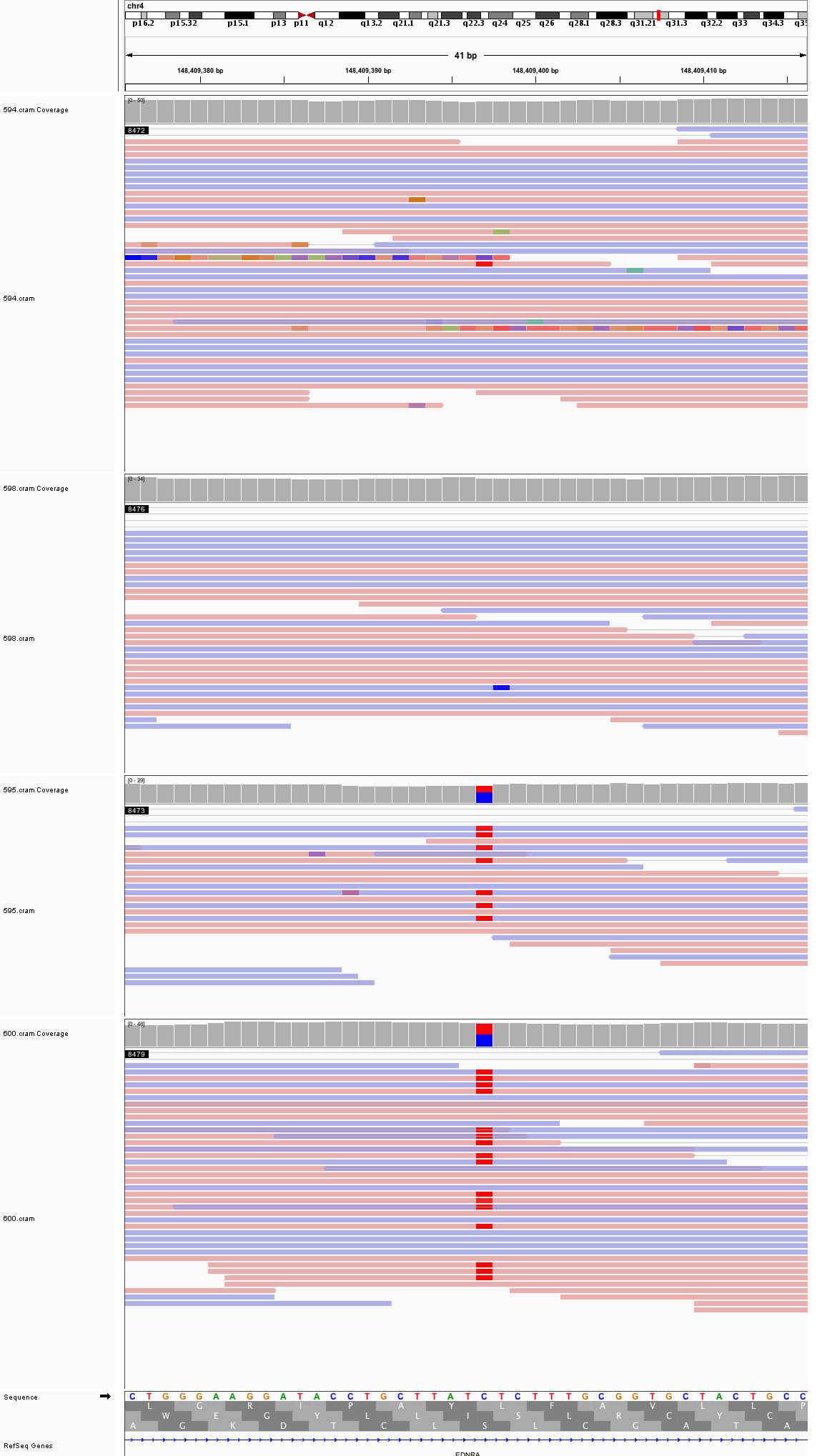

### chr4_163,417,899_163,417,939.png

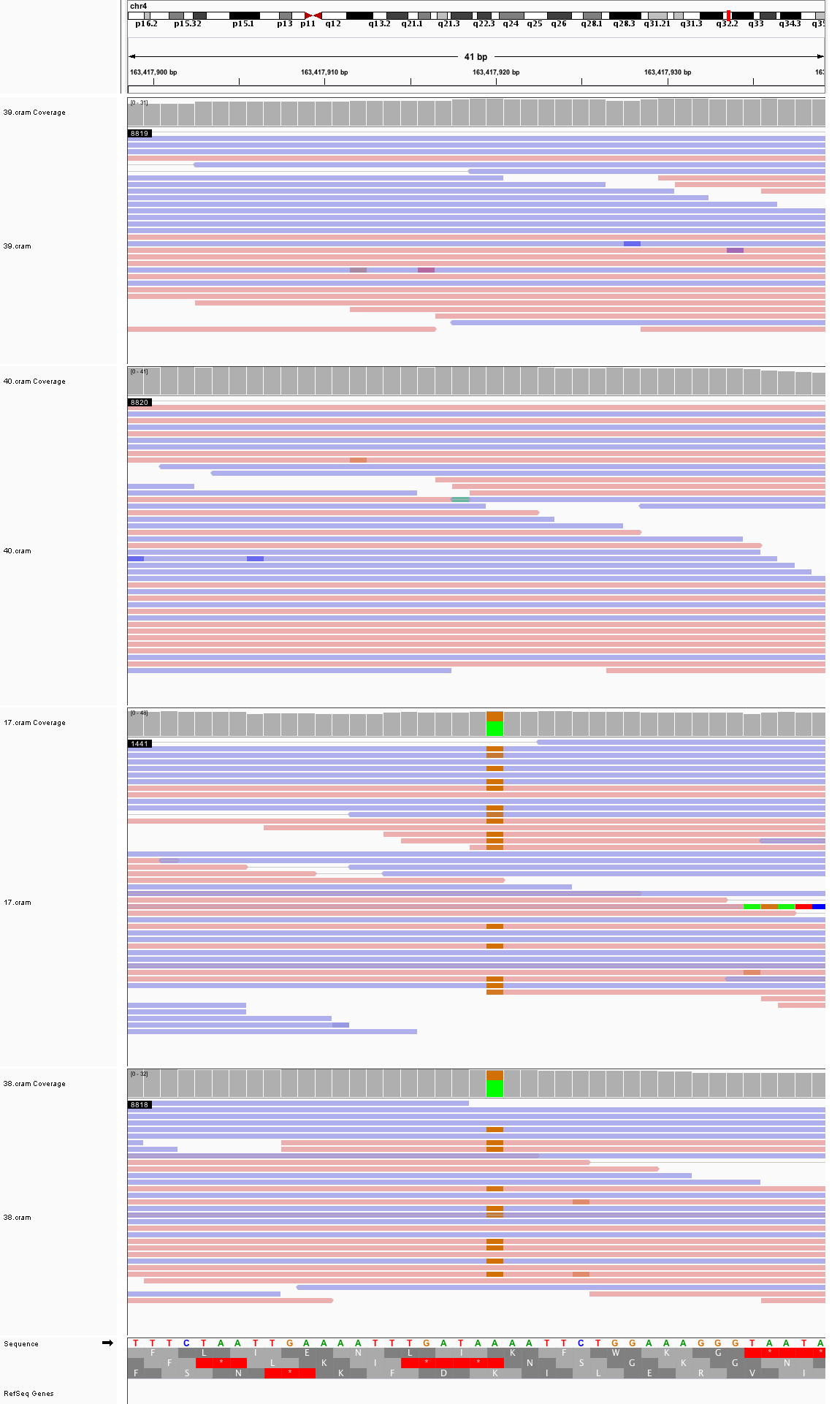

### chr4_177,098,768_177,098,808.png

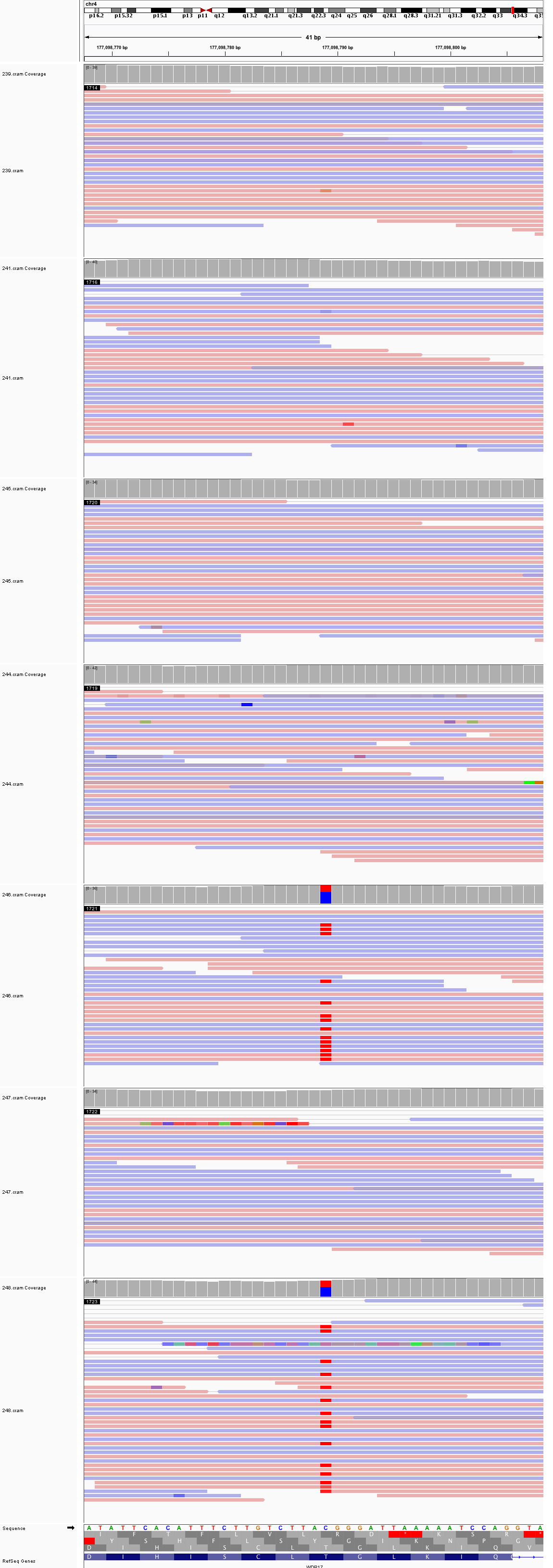

### chr5_27,444,081_27,444,121.png

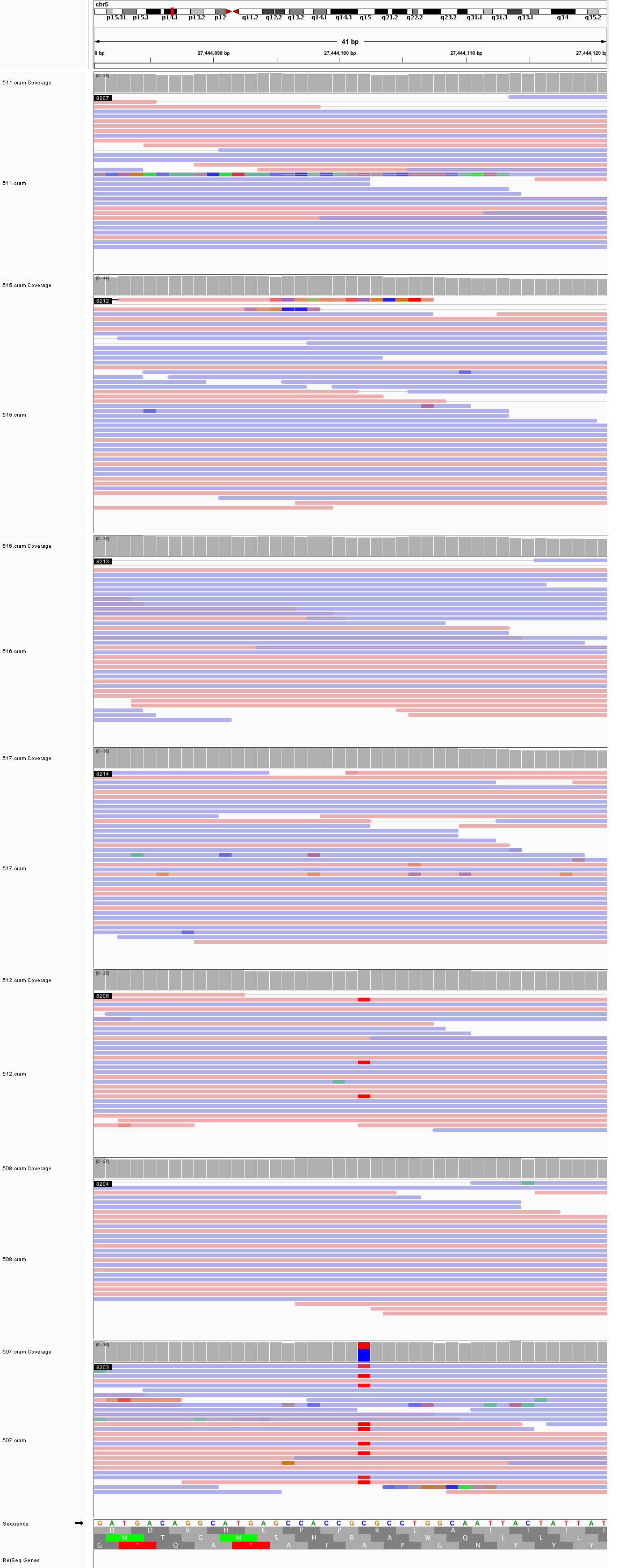

### chr5_150,536,177_150,536,217.png

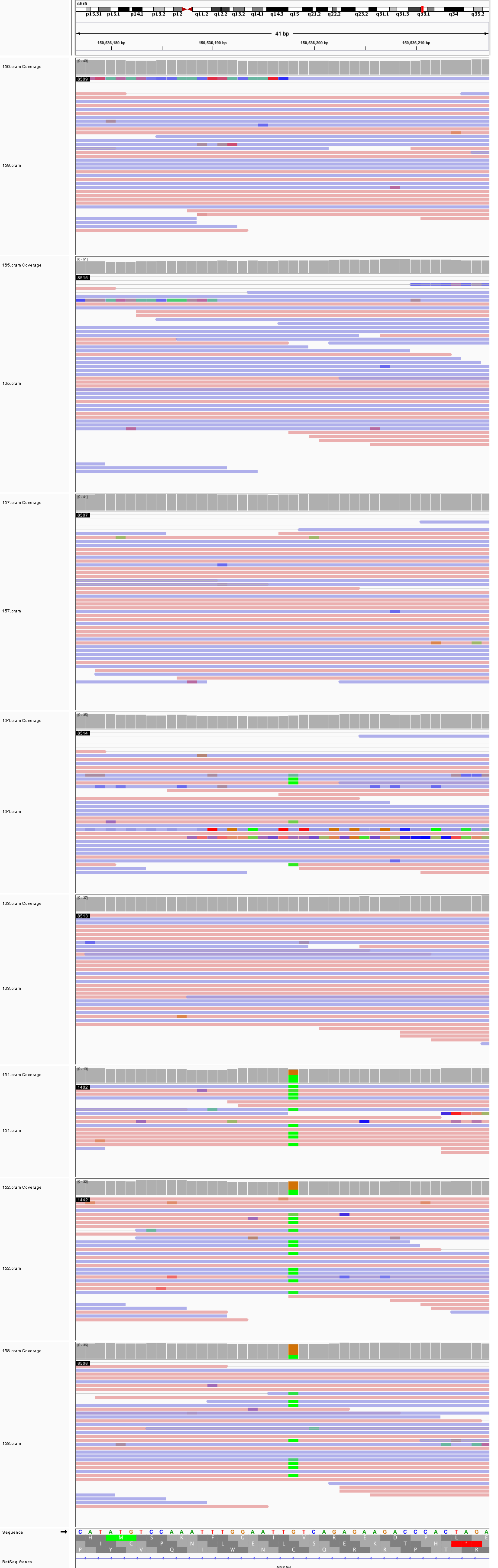

### chr5_155,129,893_155,129,933.png

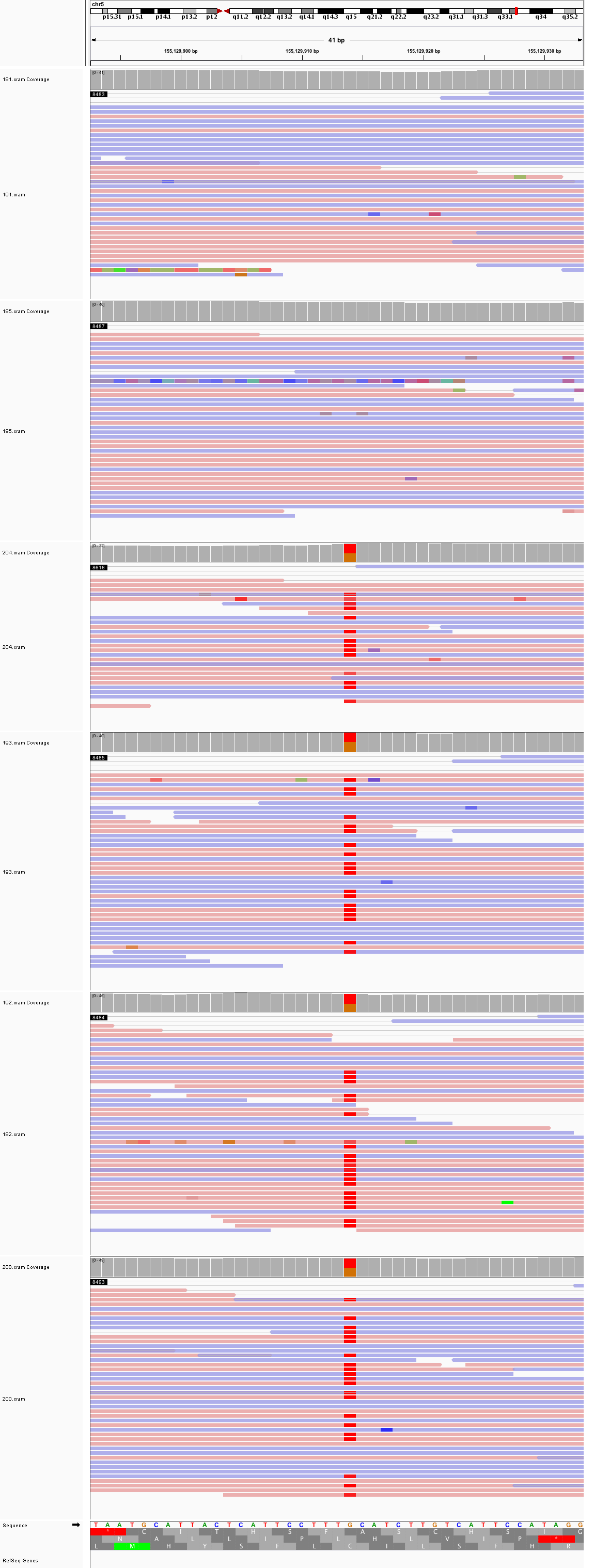

### chr6_31,654,580_31,654,620.png

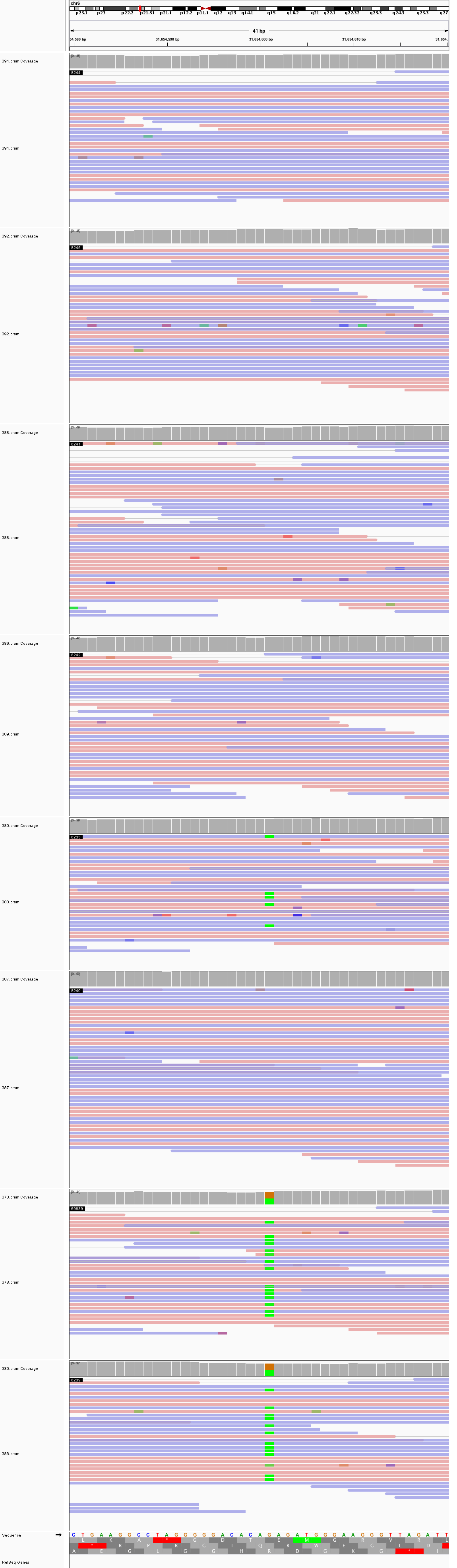

### chr6_145,921,825_145,921,865.png

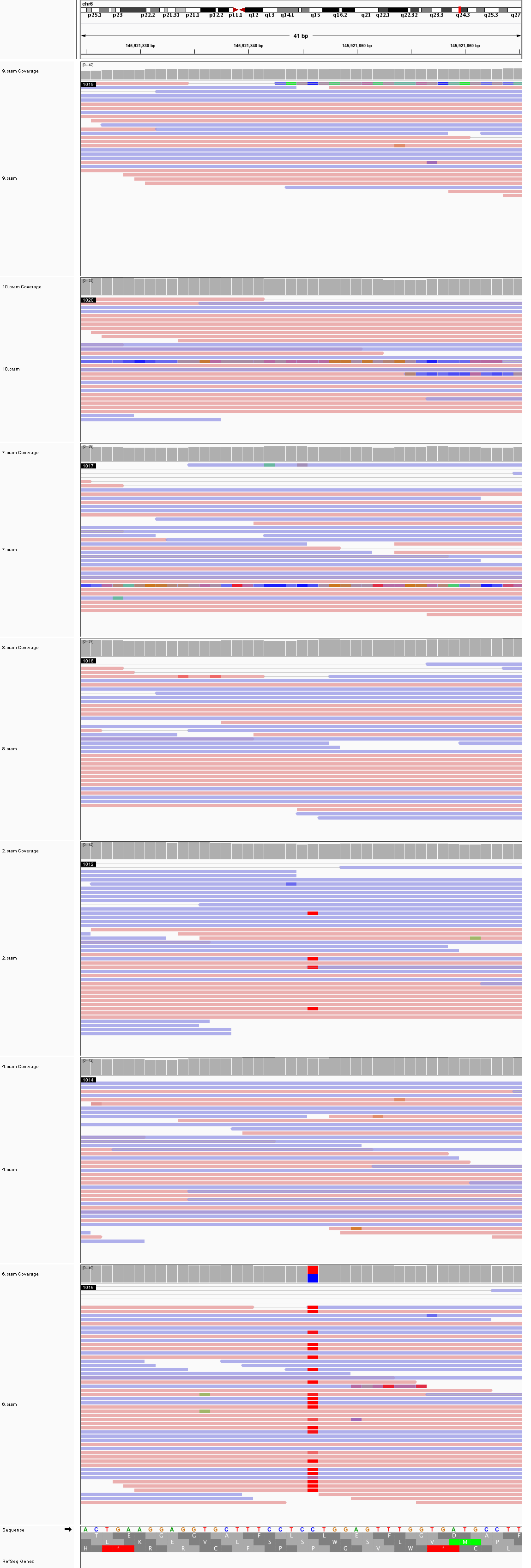

### chr6_165,543,860_165,543,900.png

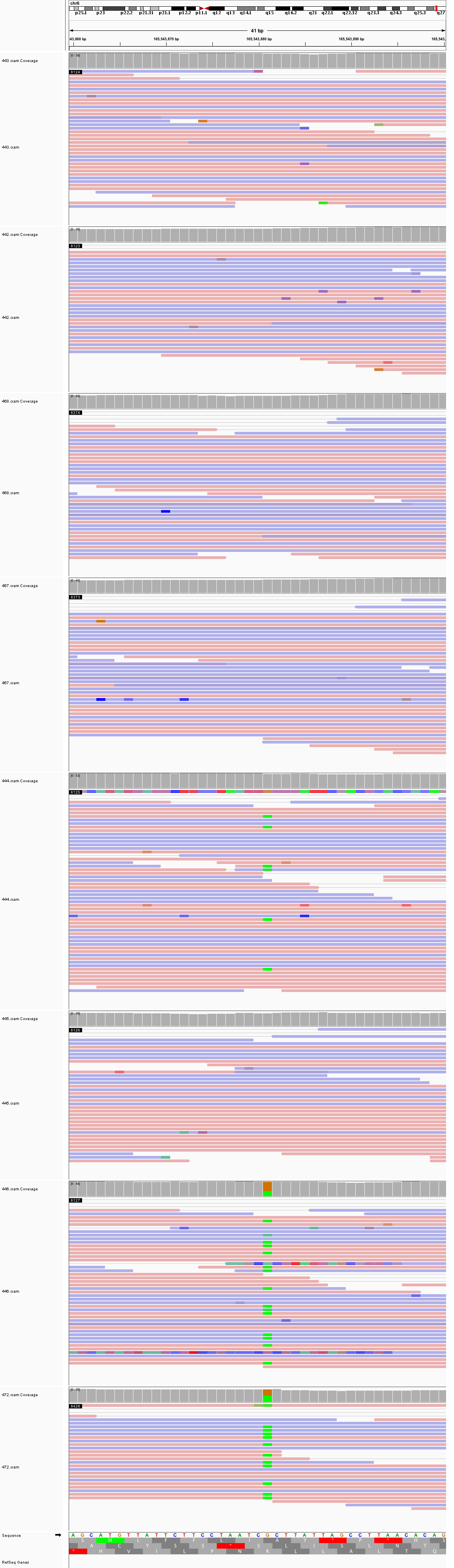

### chr7_25,489,755_25,489,795.png

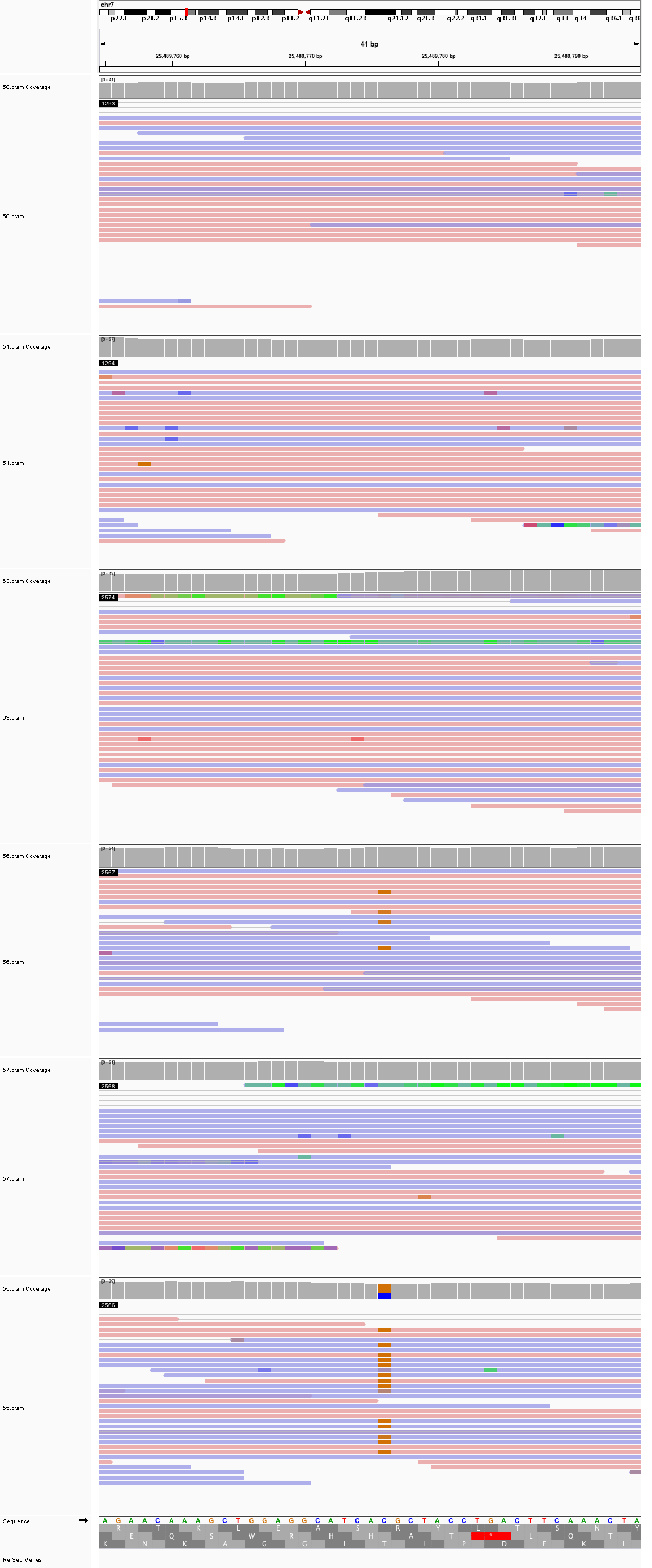

### chr7_72,004,505_72,004,545.png

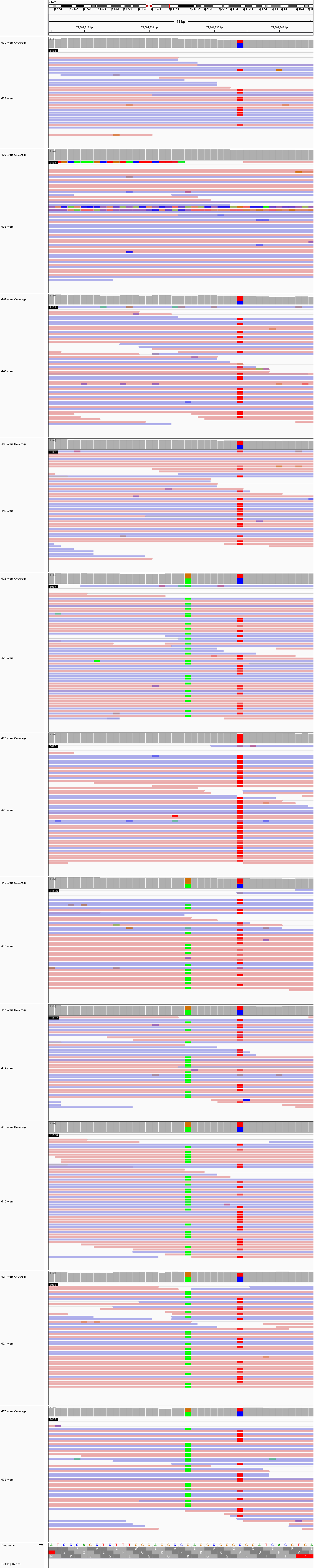

### chr7_100,525,480_100,525,520.png

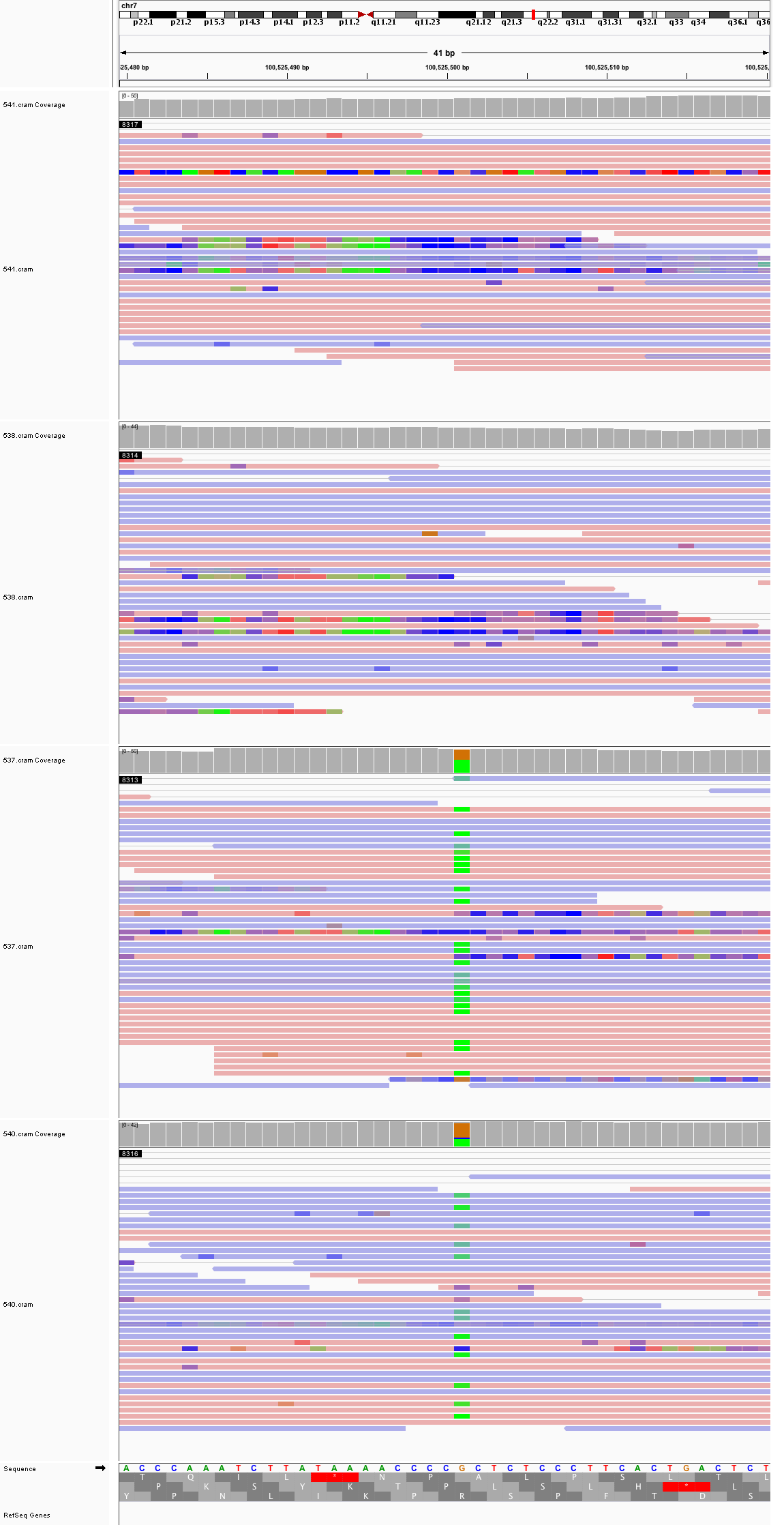

### chr7_142,108,182_142,108,222.png

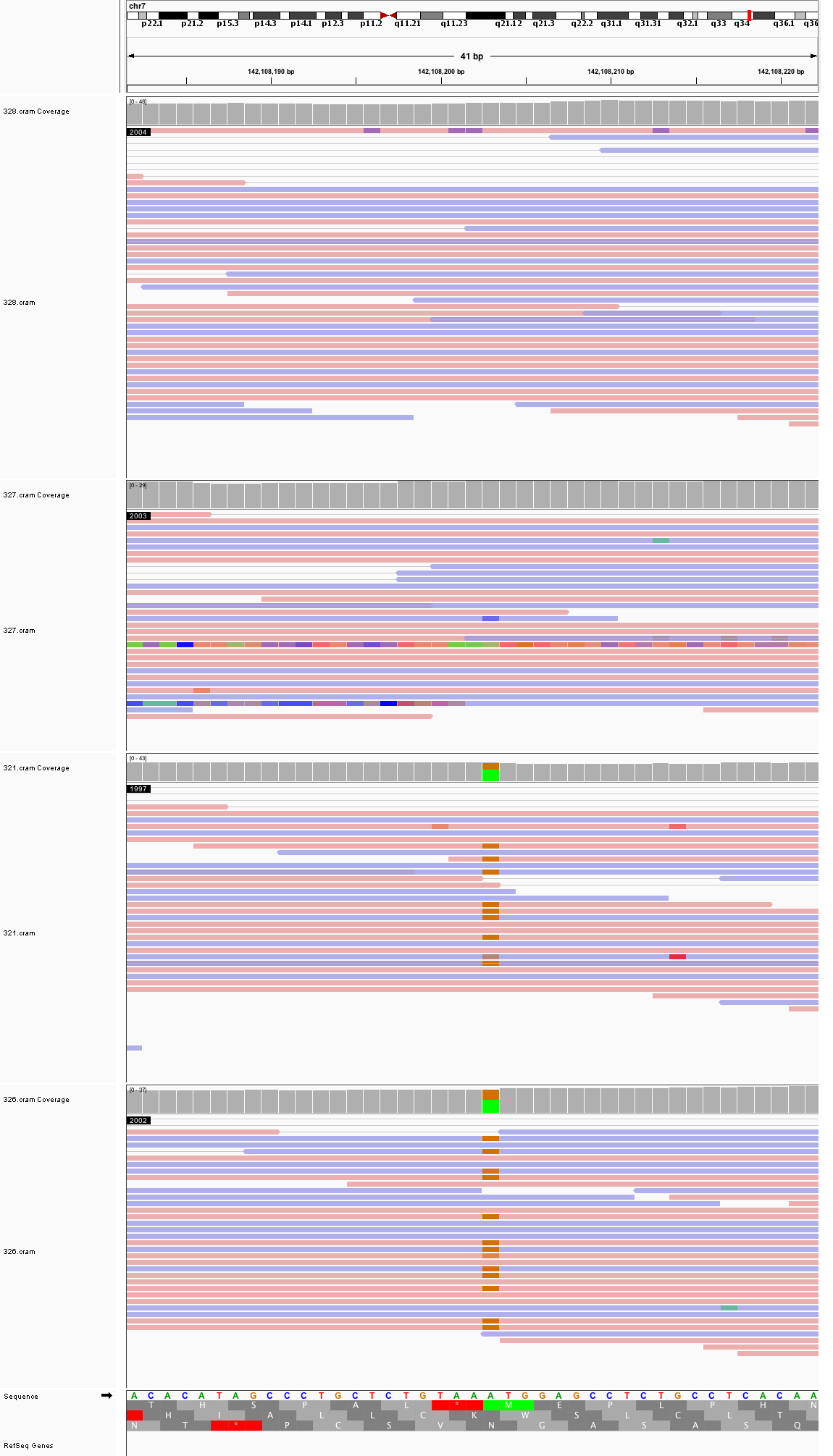

### chr7_146,245,437_146,245,477.png

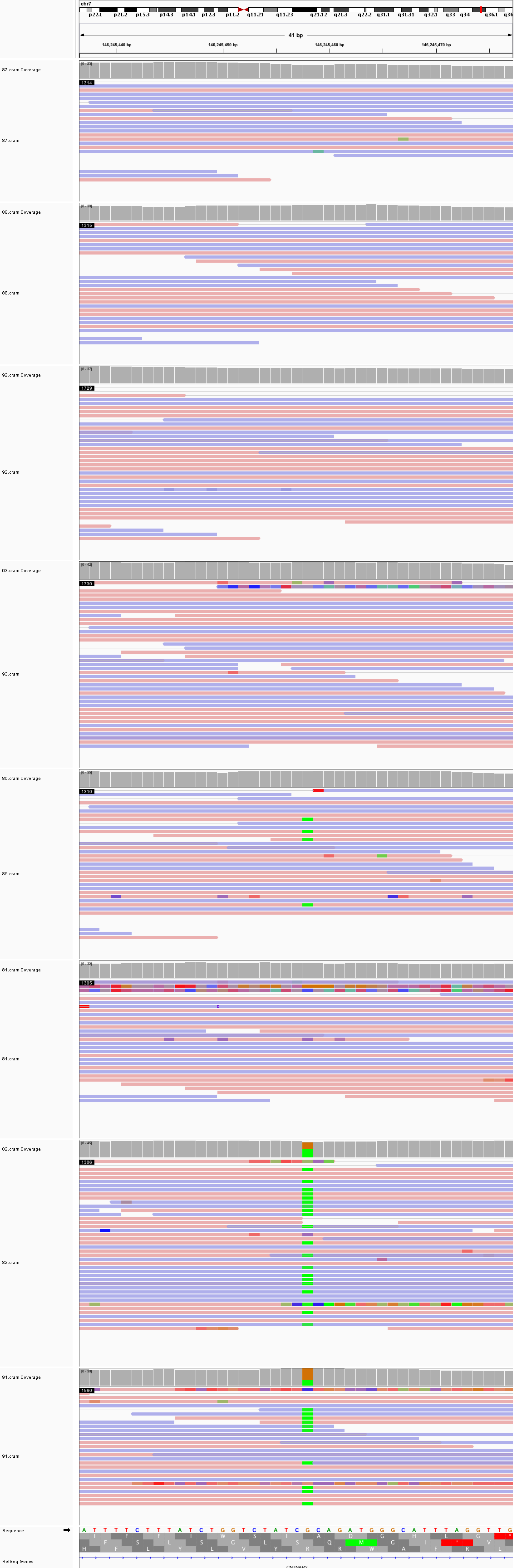

### chr7_147,903,022_147,903,062.png

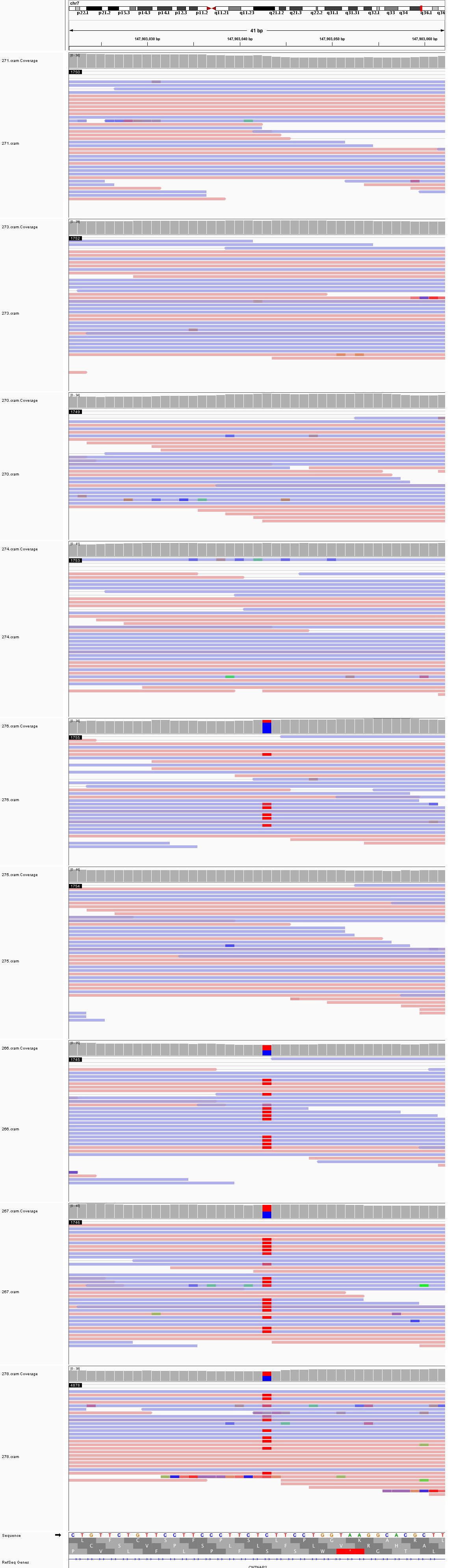

### chr7_155,342,382_155,342,422.png

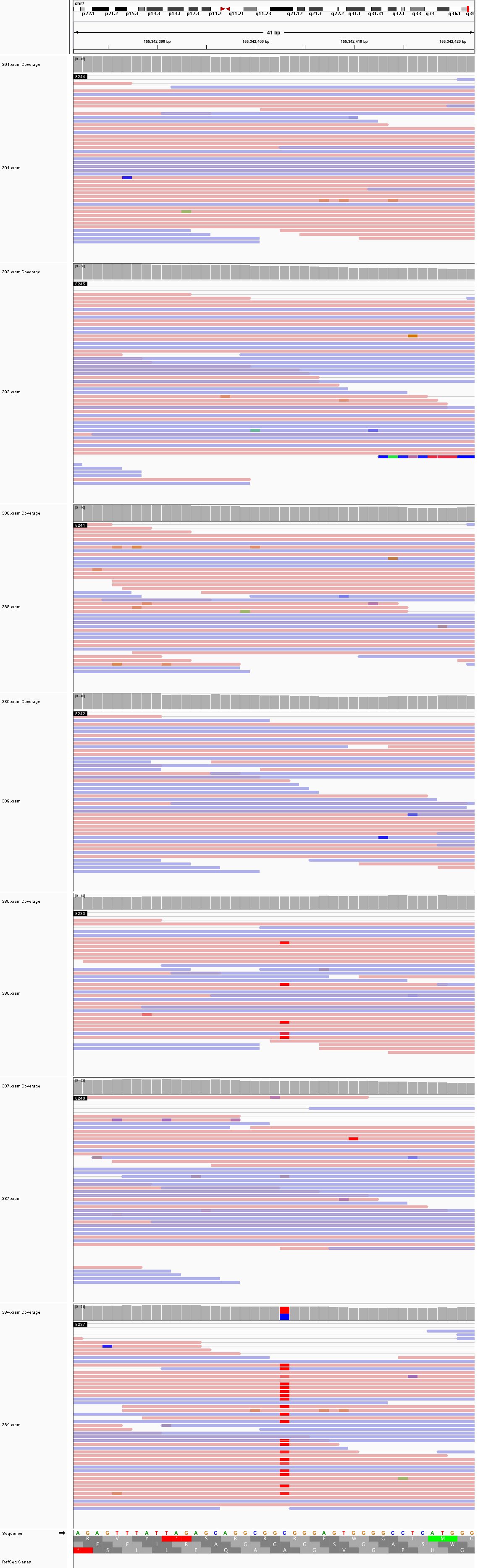

### chr7_158,104,384_158,104,424.png

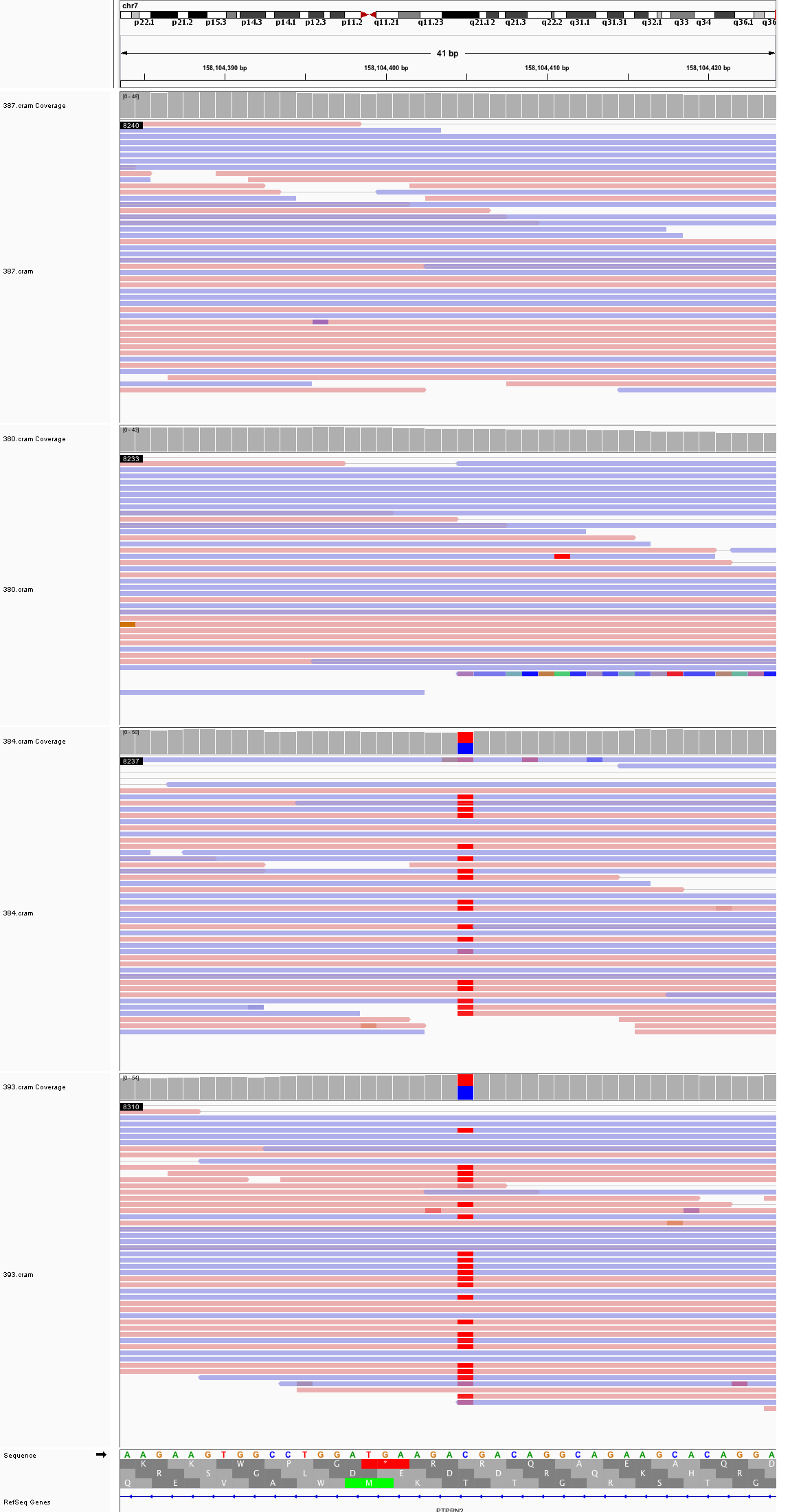

### chr8_5,877,598_5,877,638.png

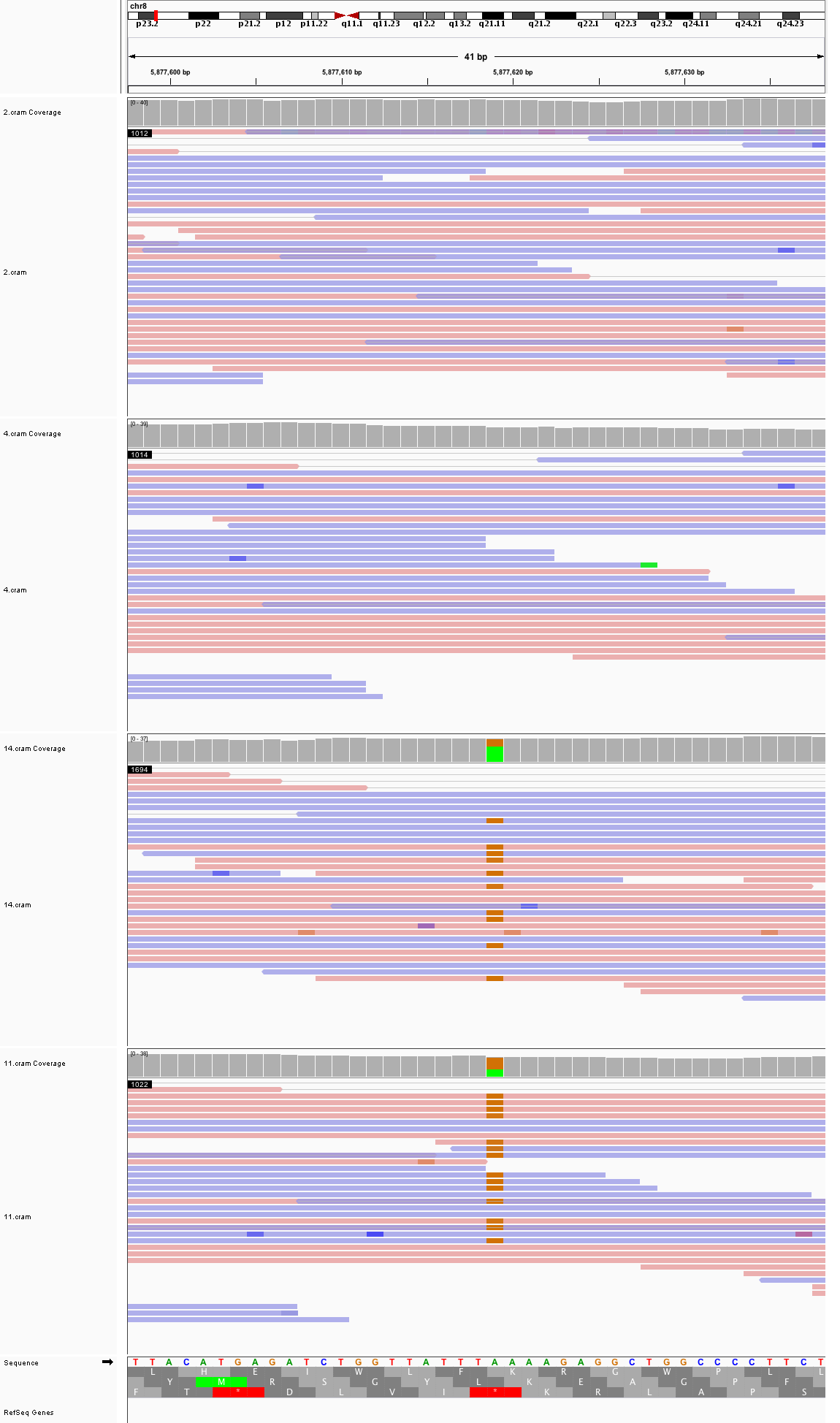

### chr8_12,939,118_12,939,158.png

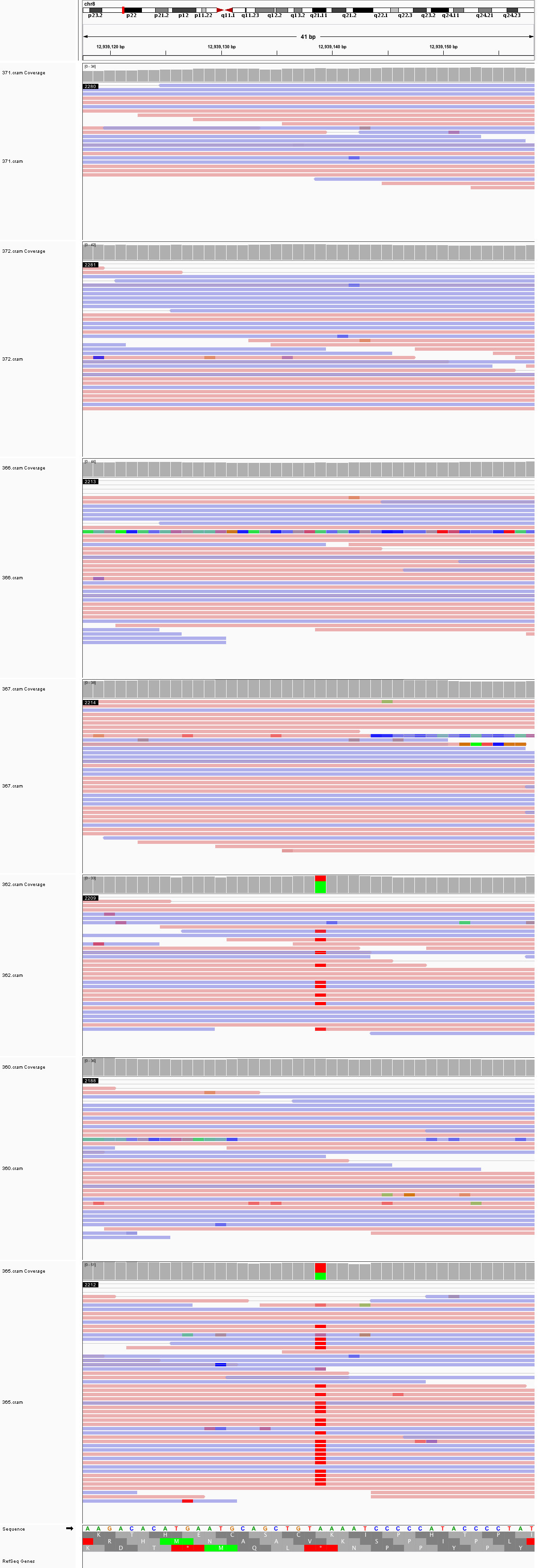

### chr8_49,711,339_49,711,379.png

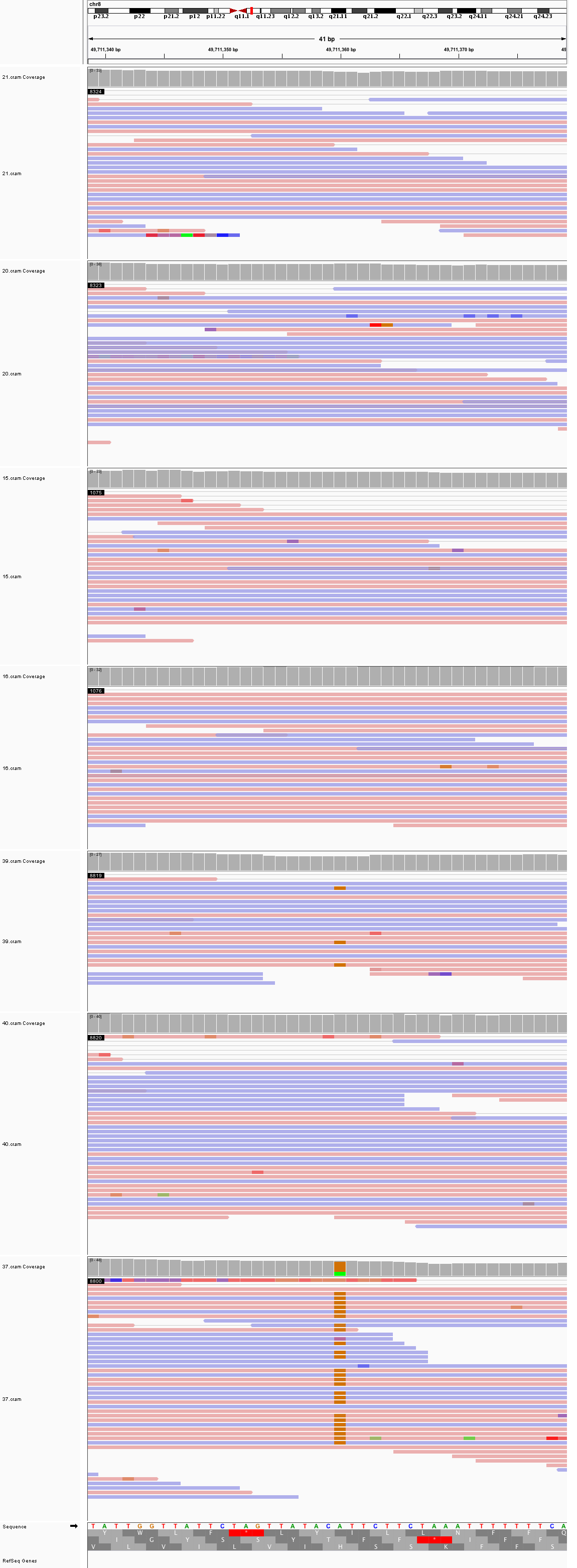

### chr8_83,980,202_83,980,242.png

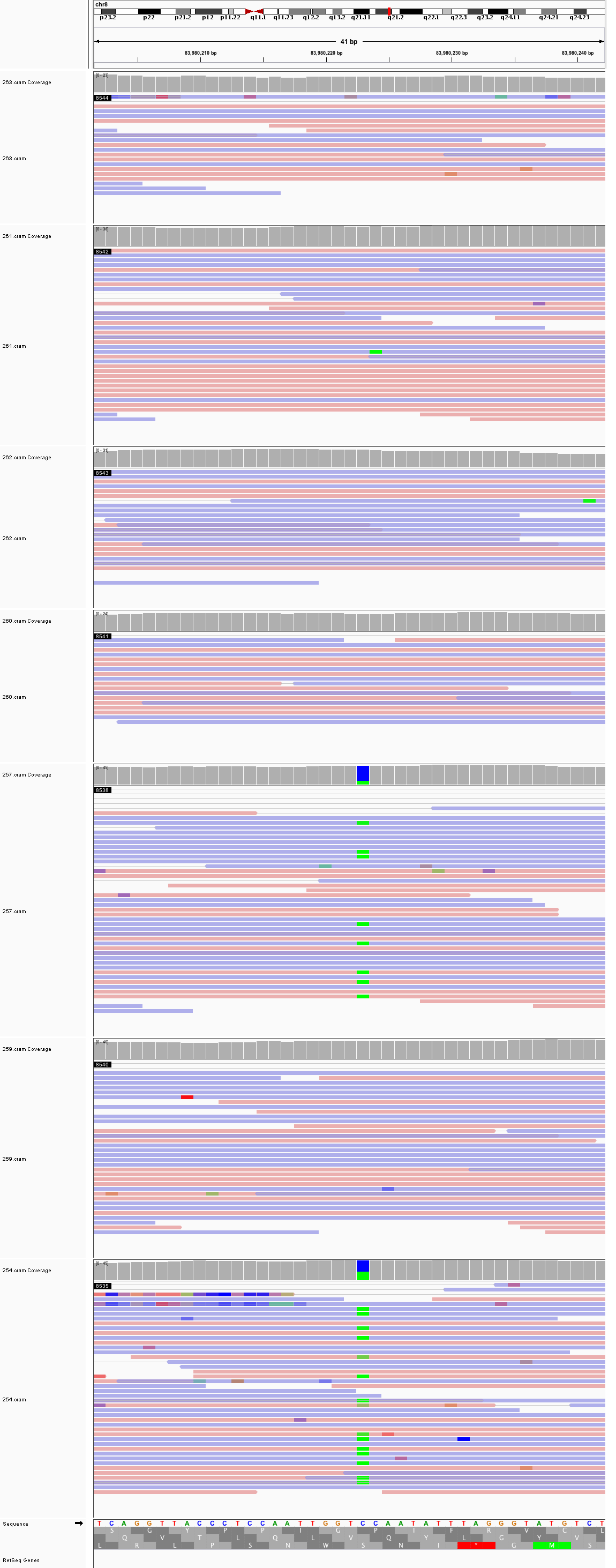

### chr8_124,866,590_124,866,630.png

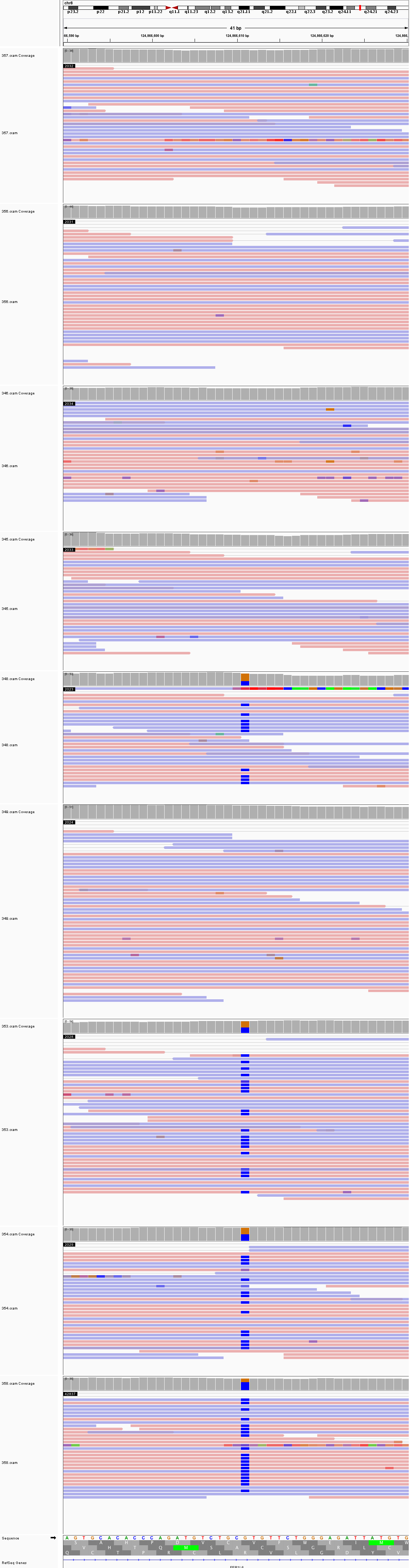

### chr9_12,141,212_12,141,252.png

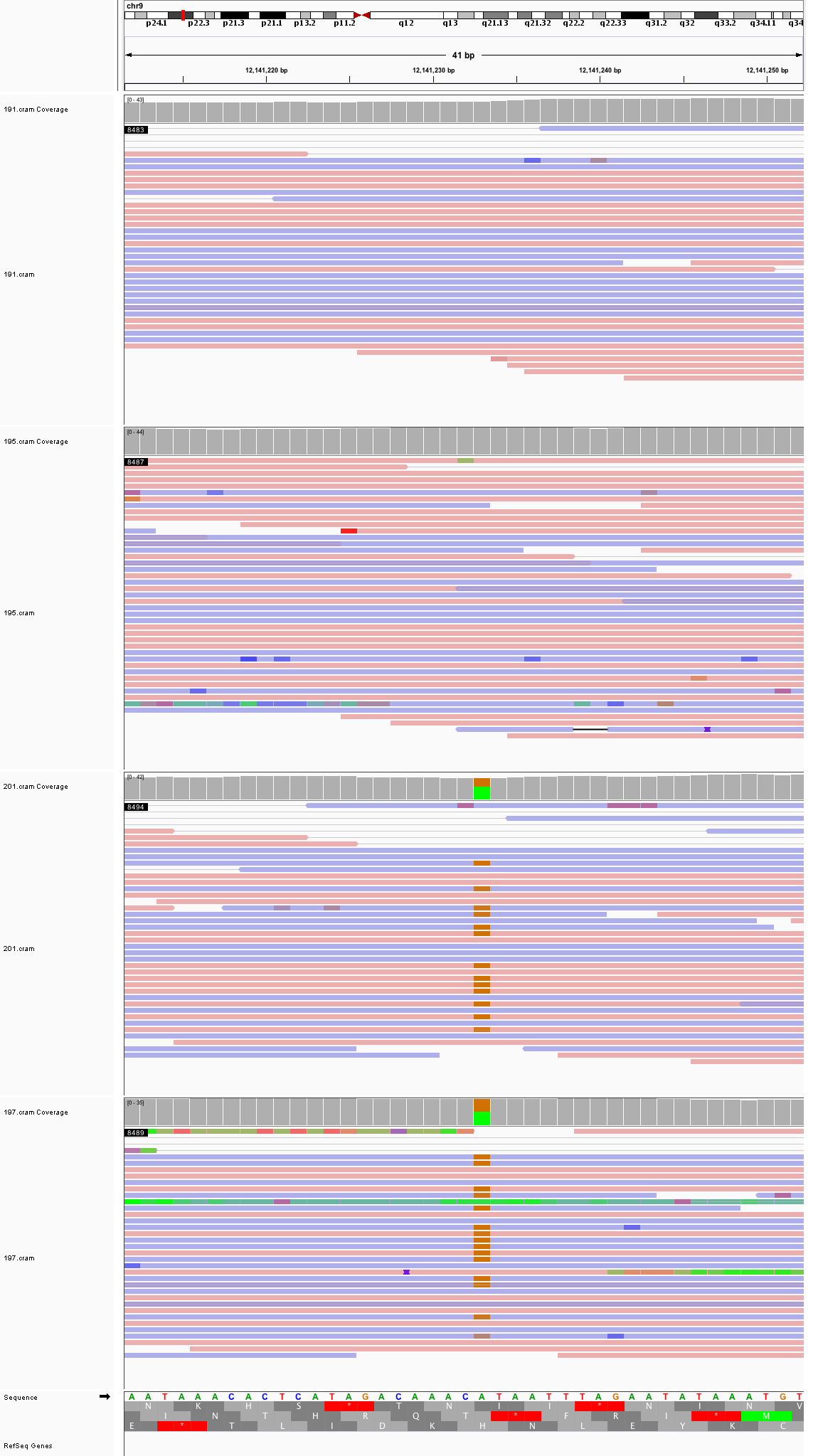

### chr9_14,994,863_14,994,903.png

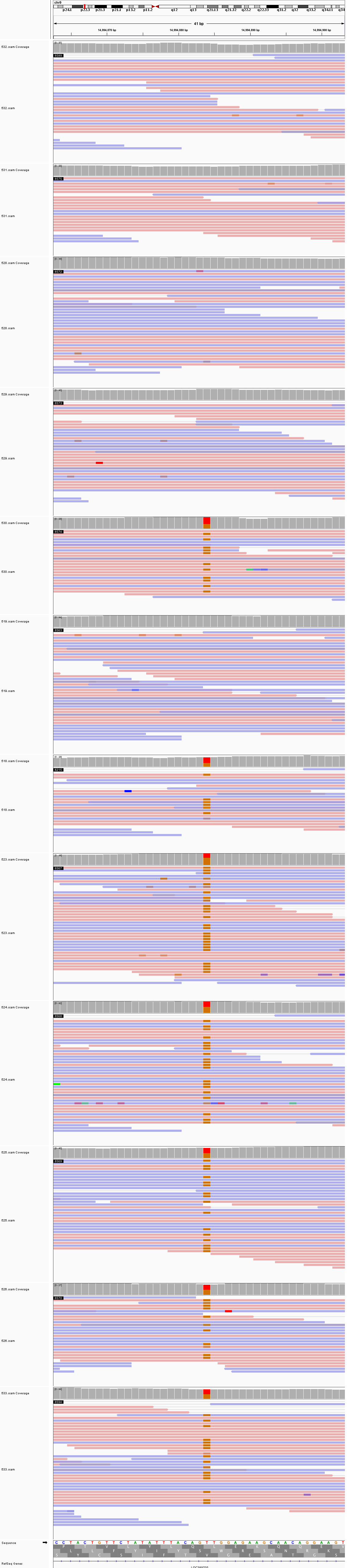

### chr9_102,711,196_102,711,236.png

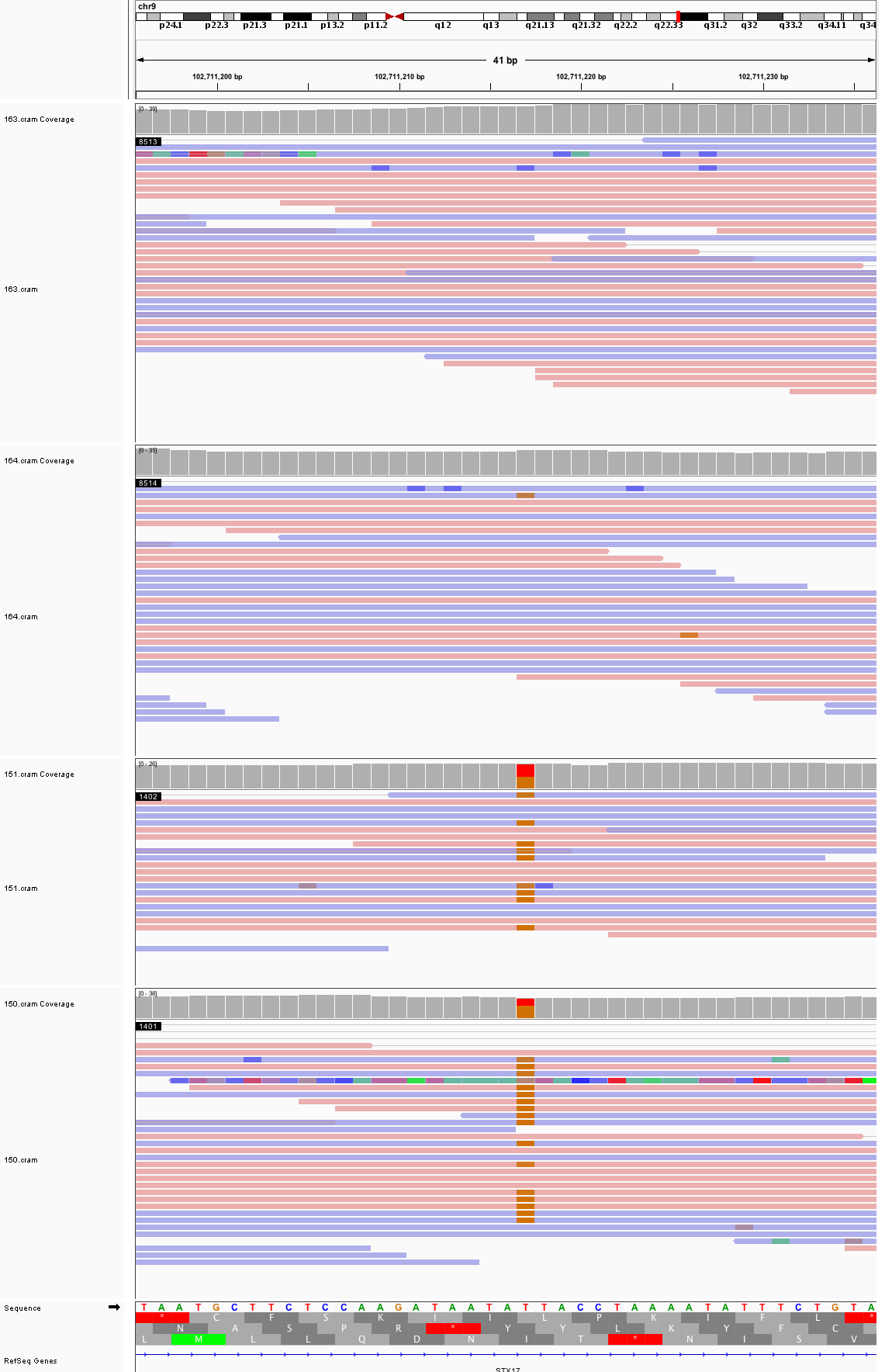

### chr9_124,126,304_124,126,344.png

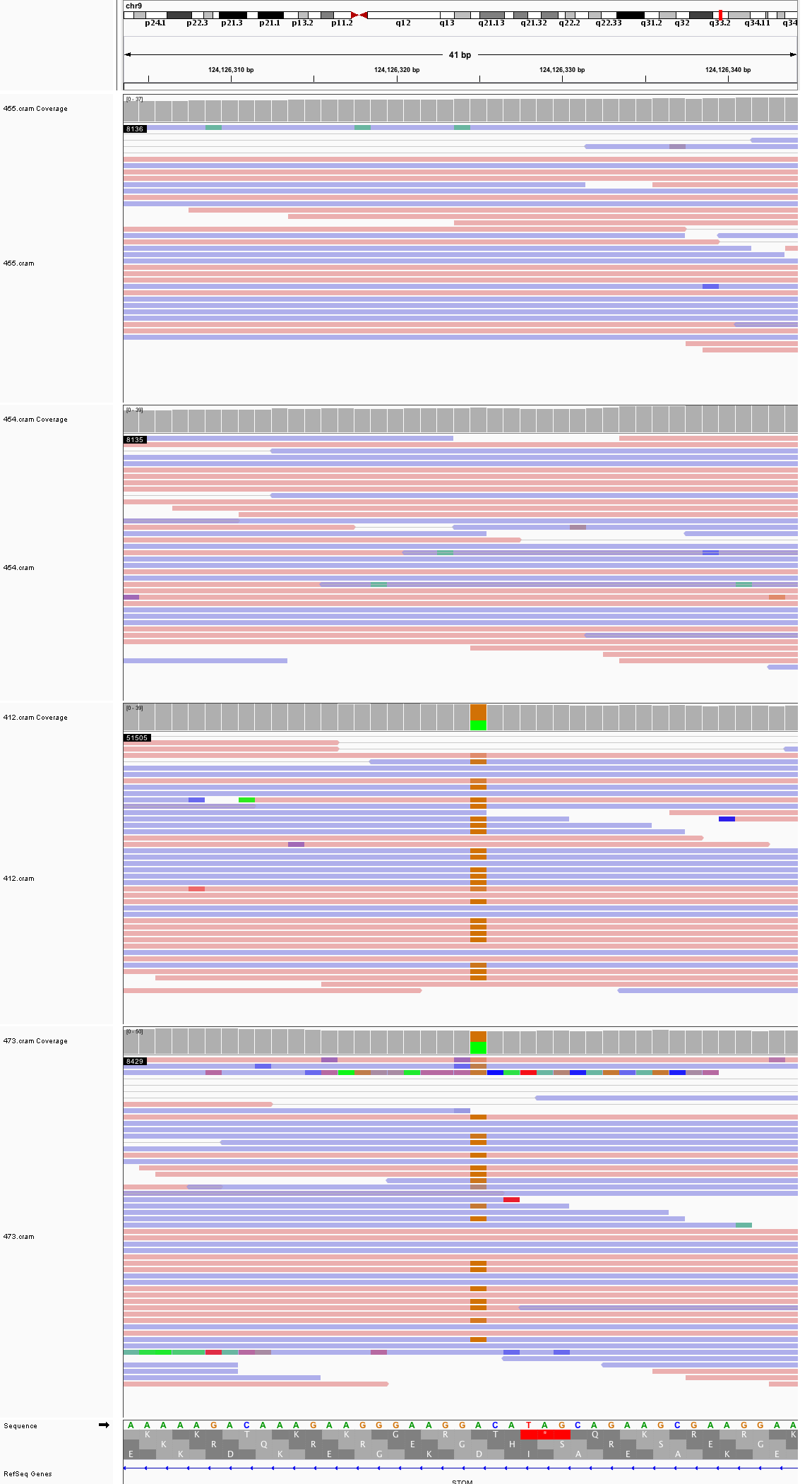

### chr9_130,038,715_130,038,755.png

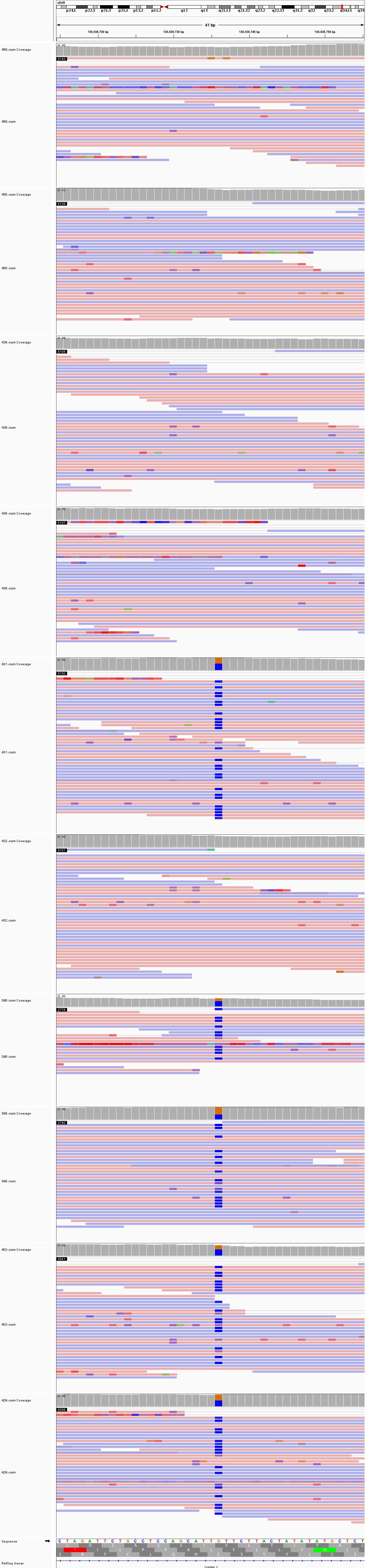

### chr9_140,524,310_140,524,350.png

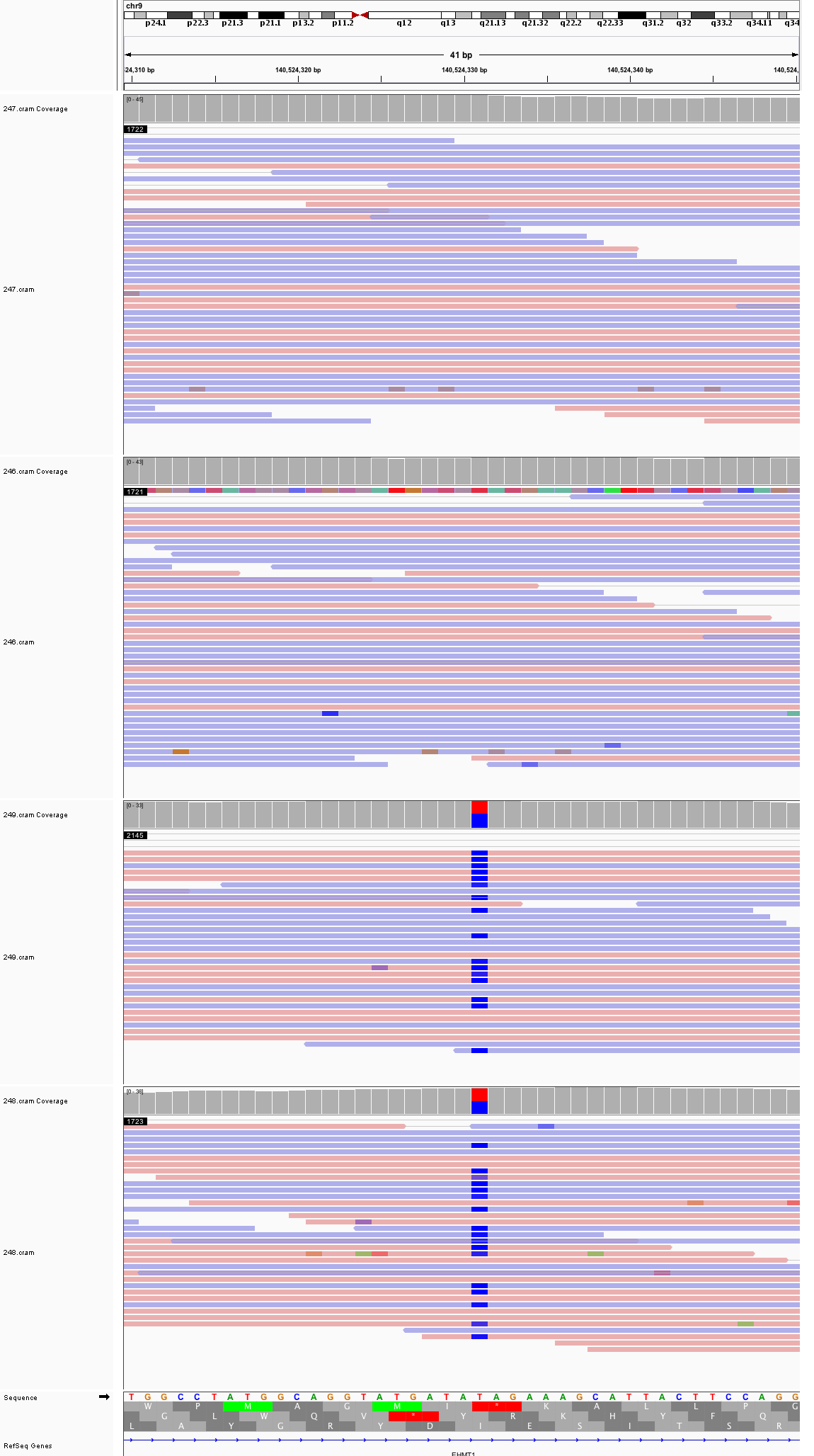

### chr10_20,827,901_20,827,941.png

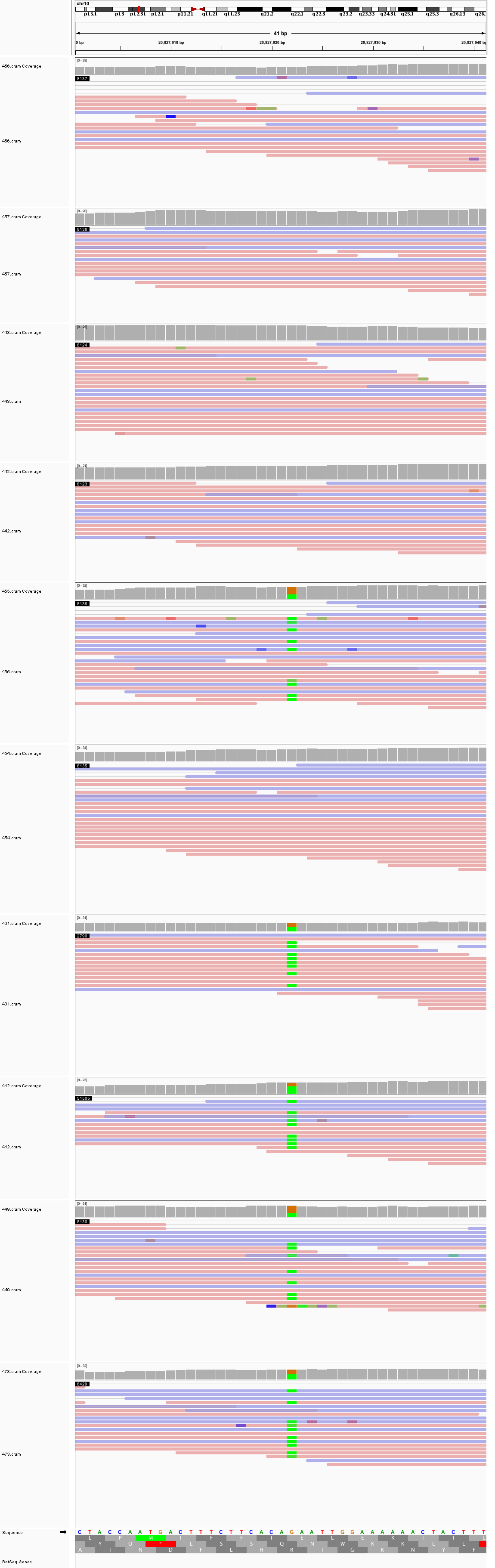

### chr10_23,925,108_23,925,148.png

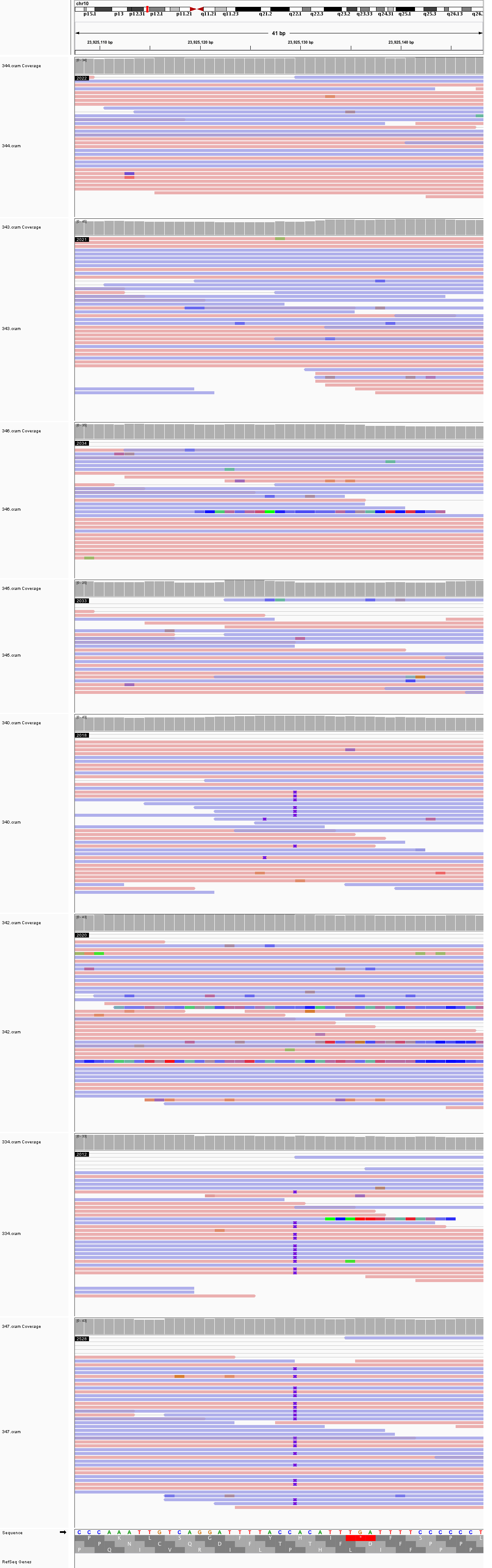

### chr10_36,602,107_36,602,147.png

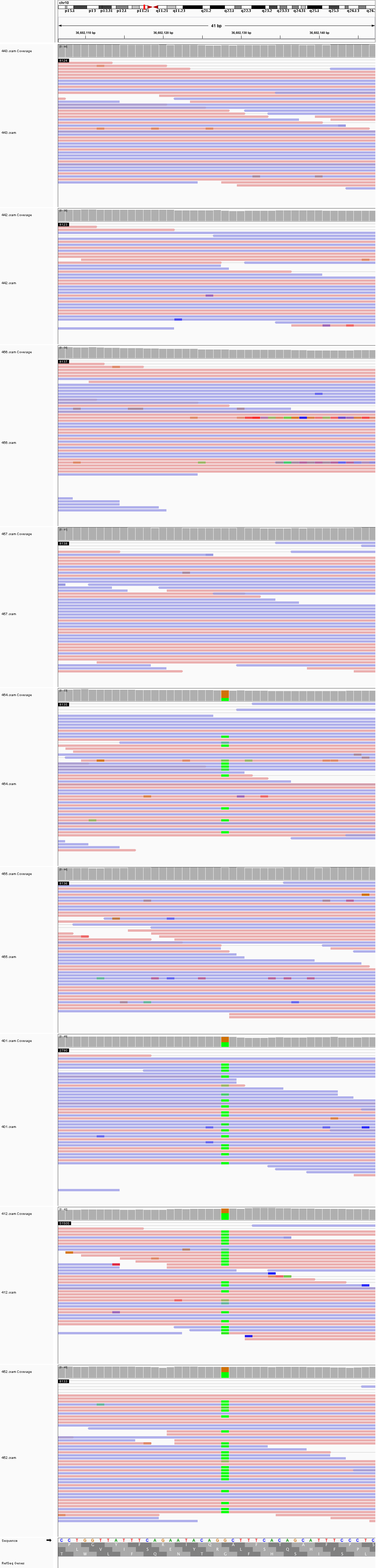

### chr10_47,538,904_47,538,944.png

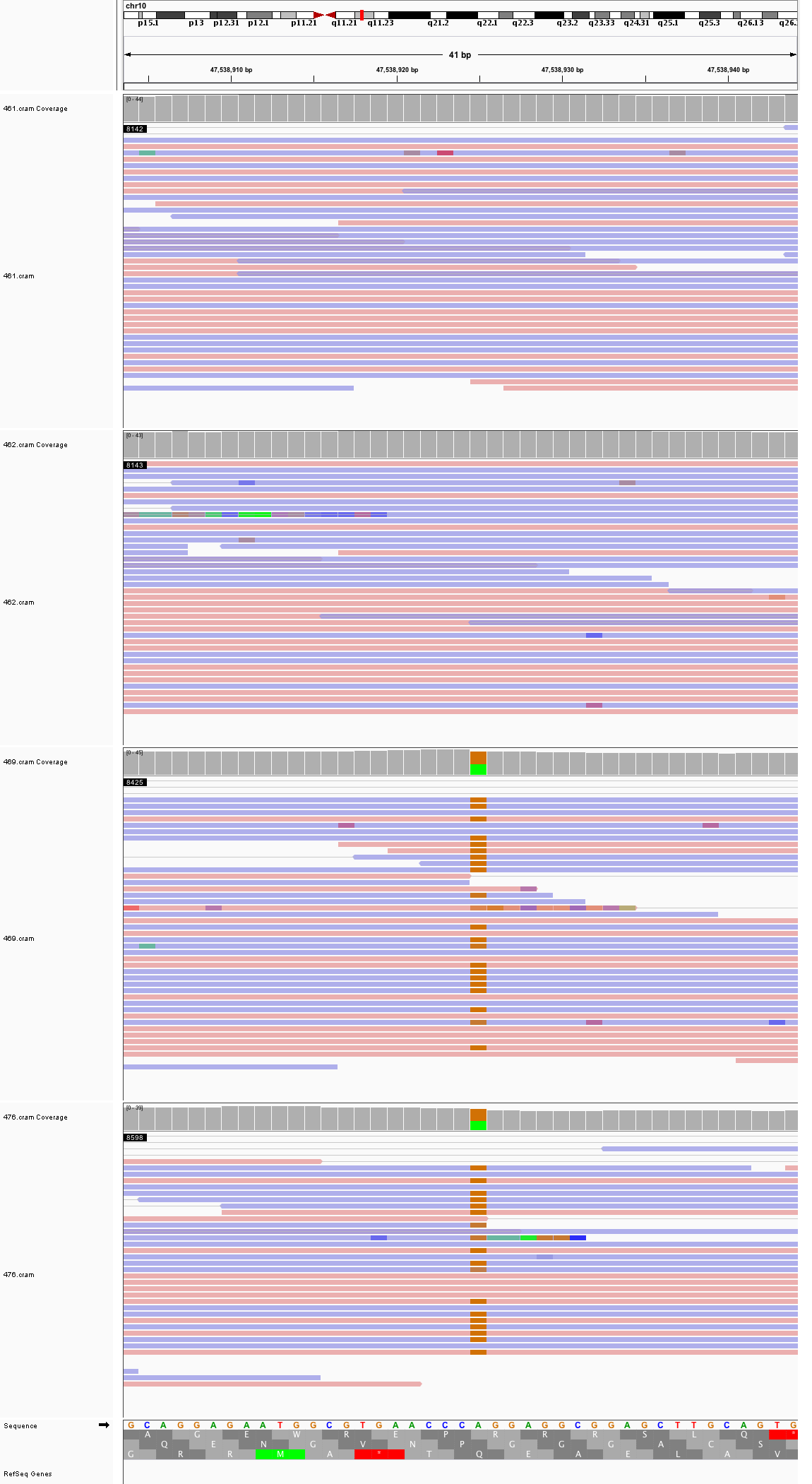

### chr10_56,276,478_56,276,518.png

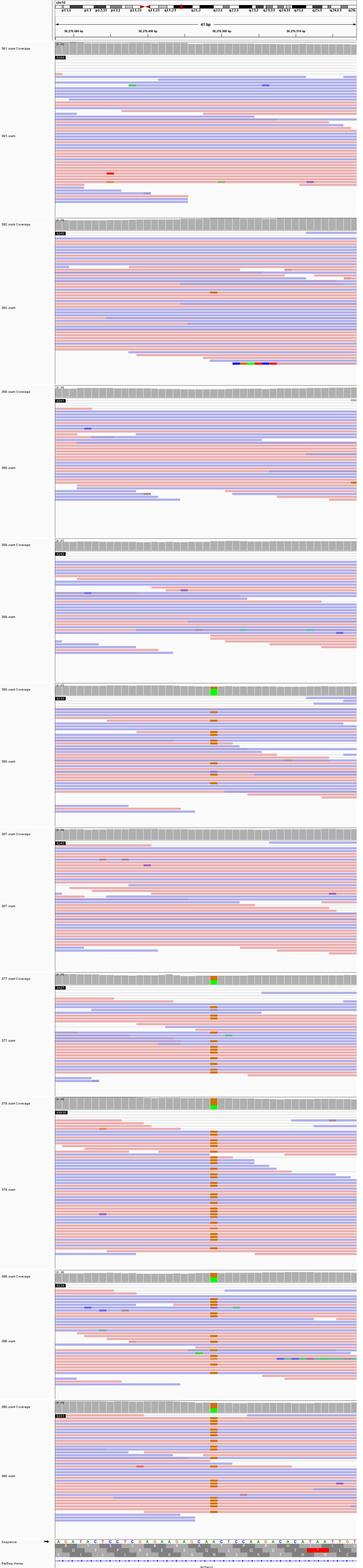

### chr10_64,122,770_64,122,810.png

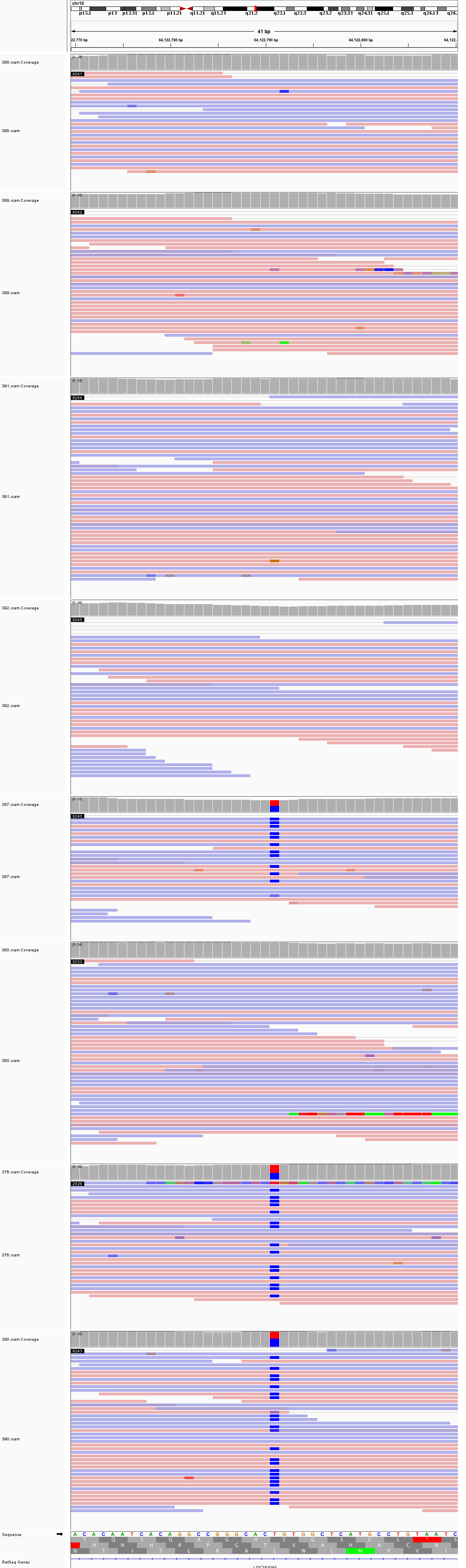

### chr11_6,750,434_6,750,474.png

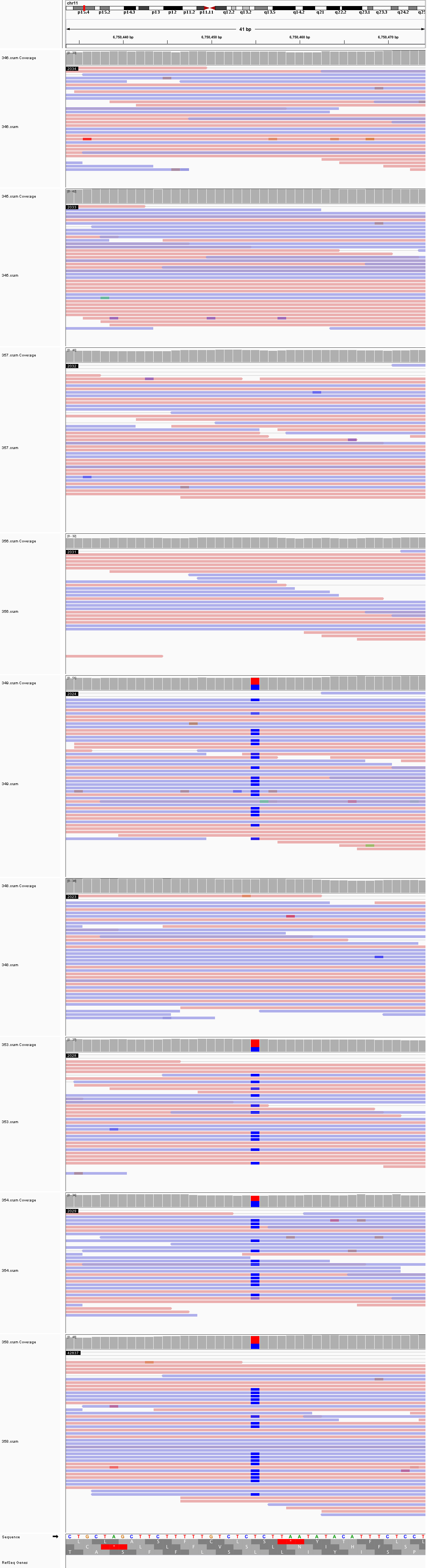

### chr11_19,139,146_19,139,186.png

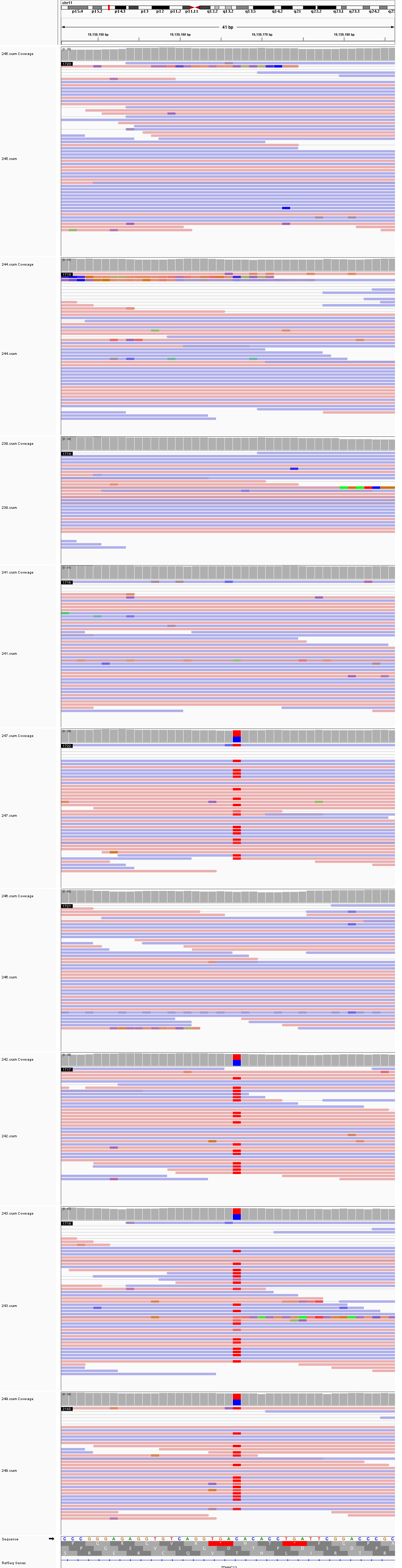

### chr11_83,607,485_83,607,525.png

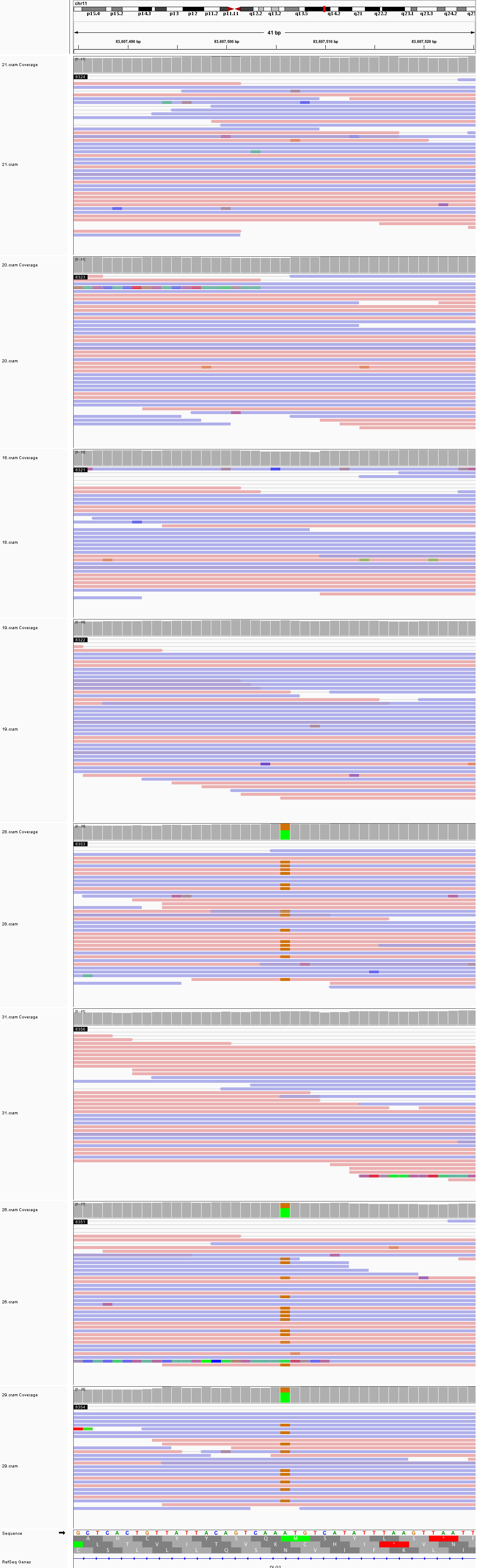

### chr11_120,739,231_120,739,271.png

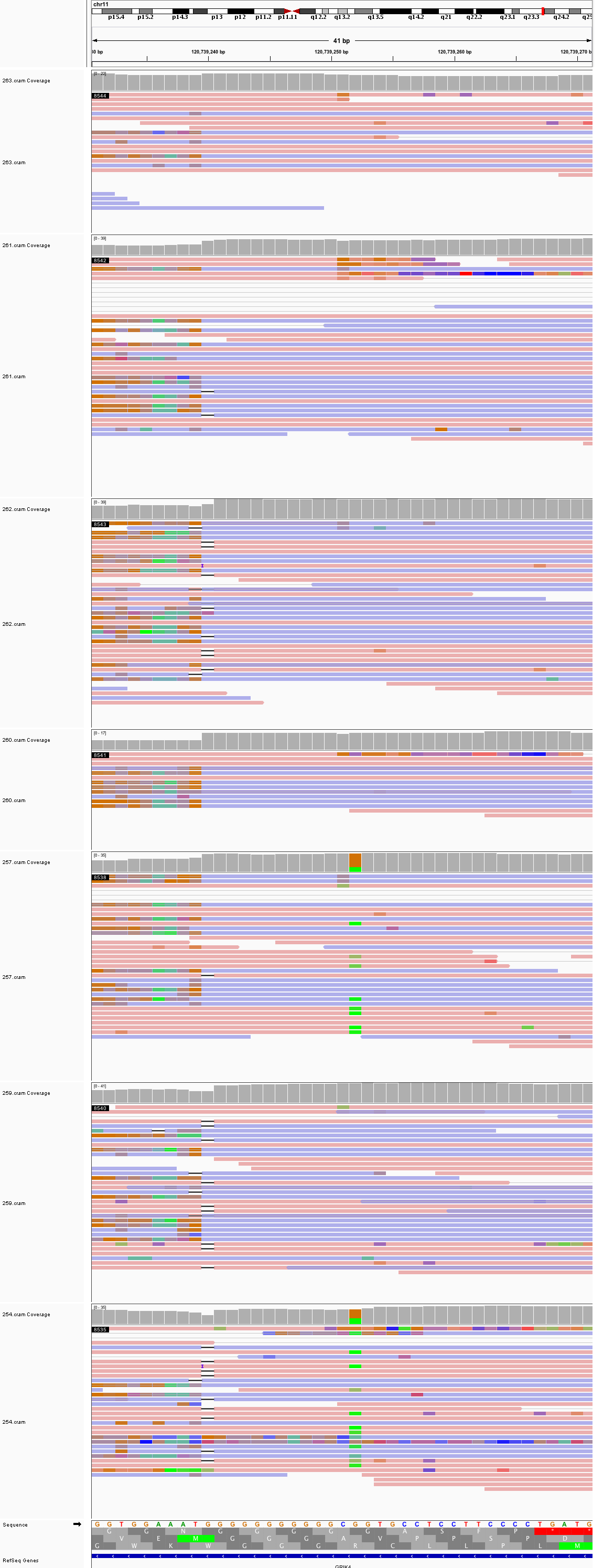

### chr12_22,931,838_22,931,878.png

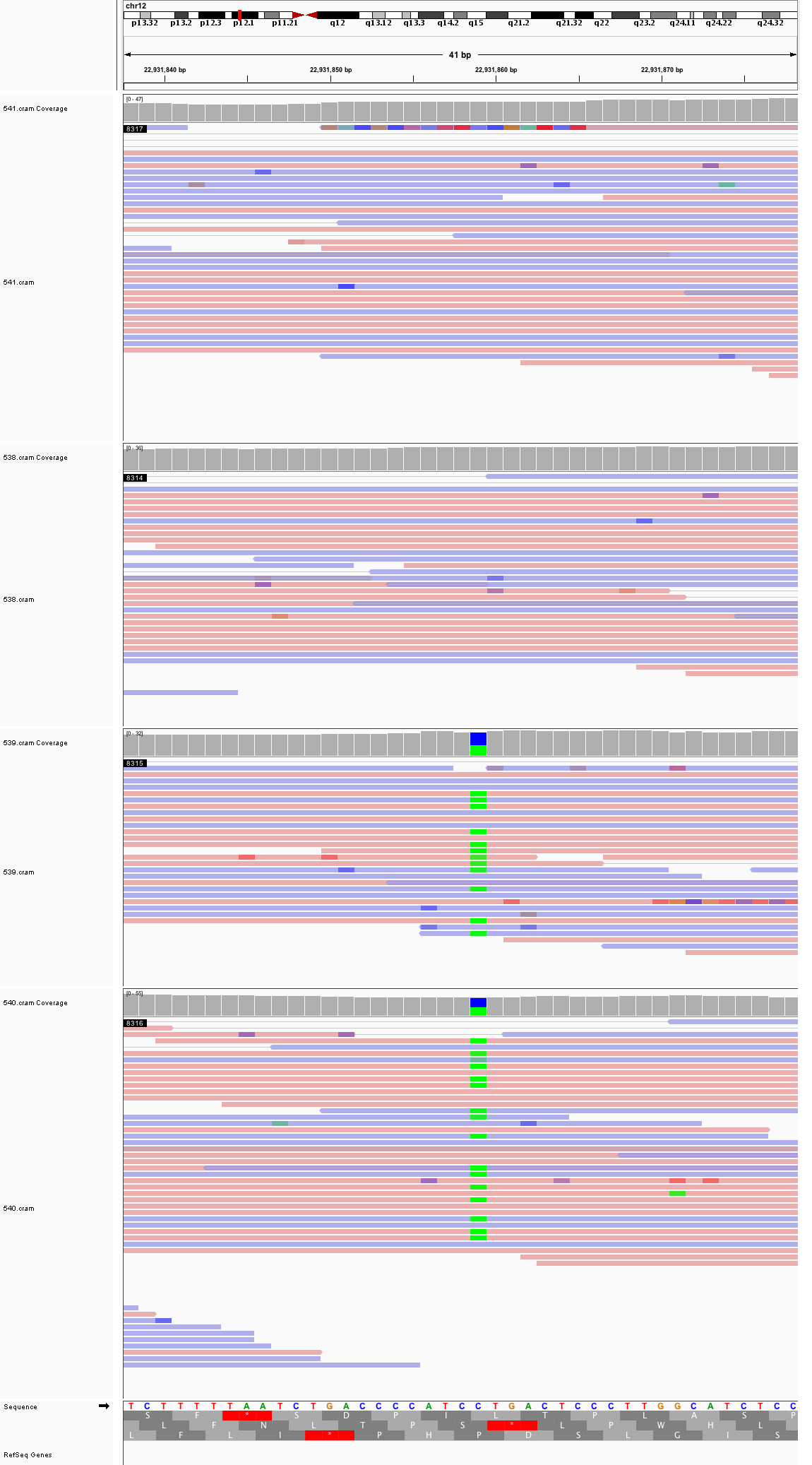

### chr12_30,284,139_30,284,179.png

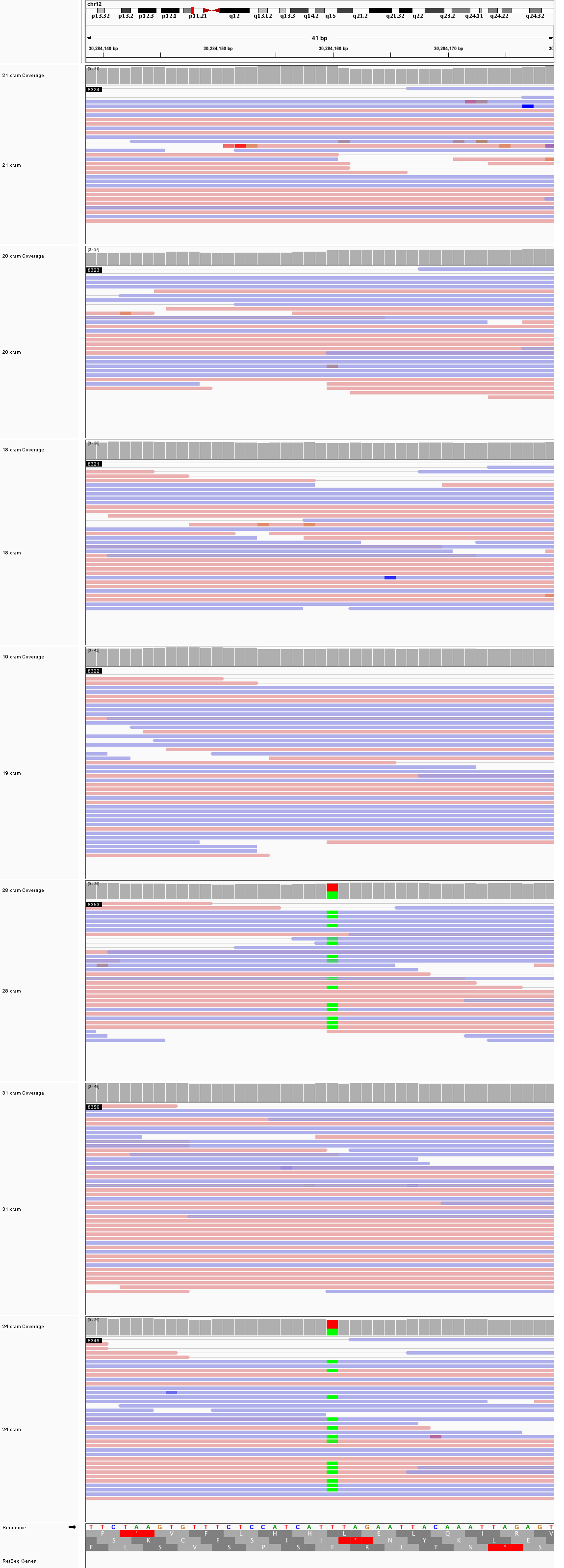

### chr12_66,312,973_66,313,013.png

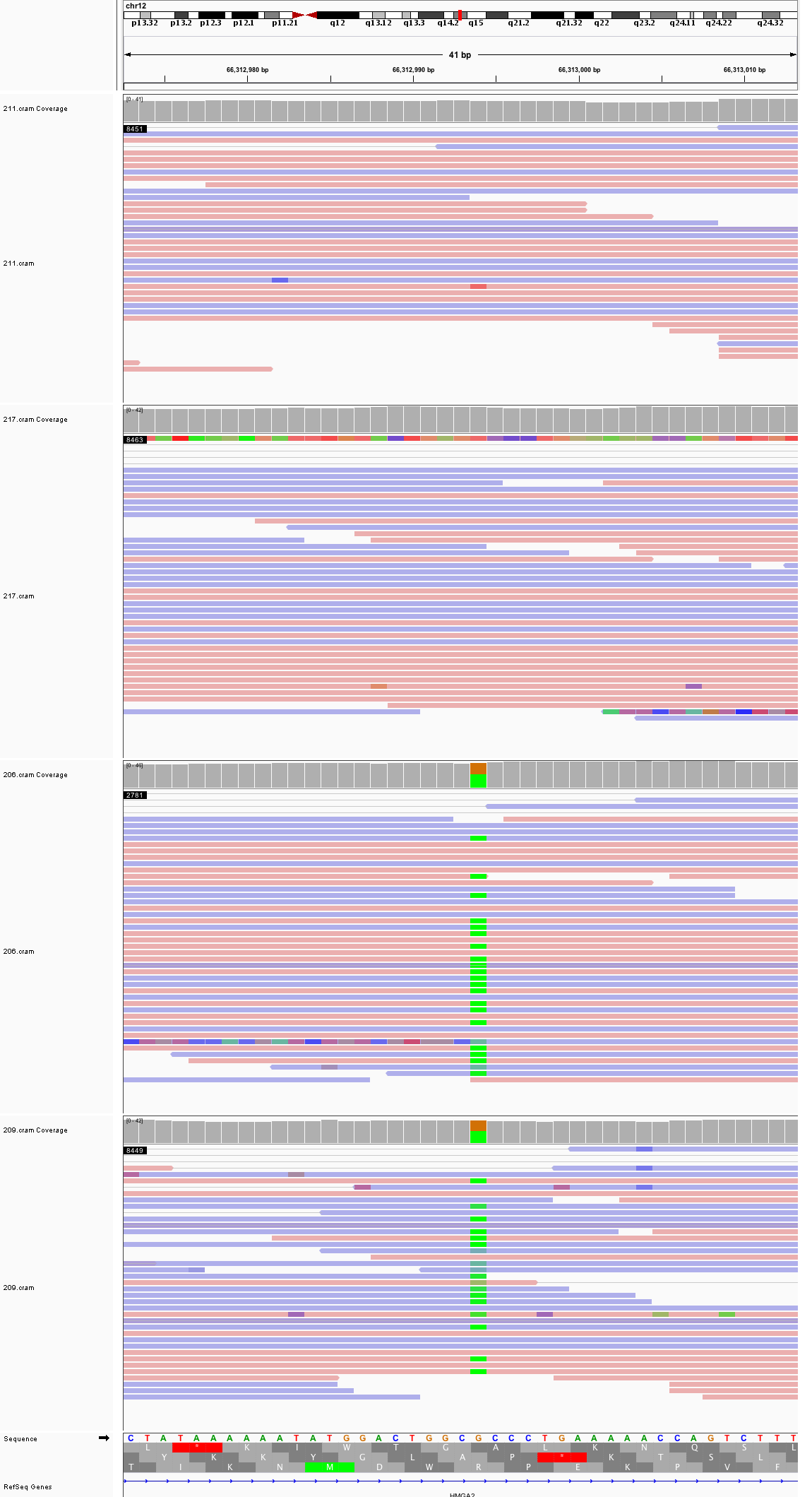

### chr12_88,030,686_88,030,726.png

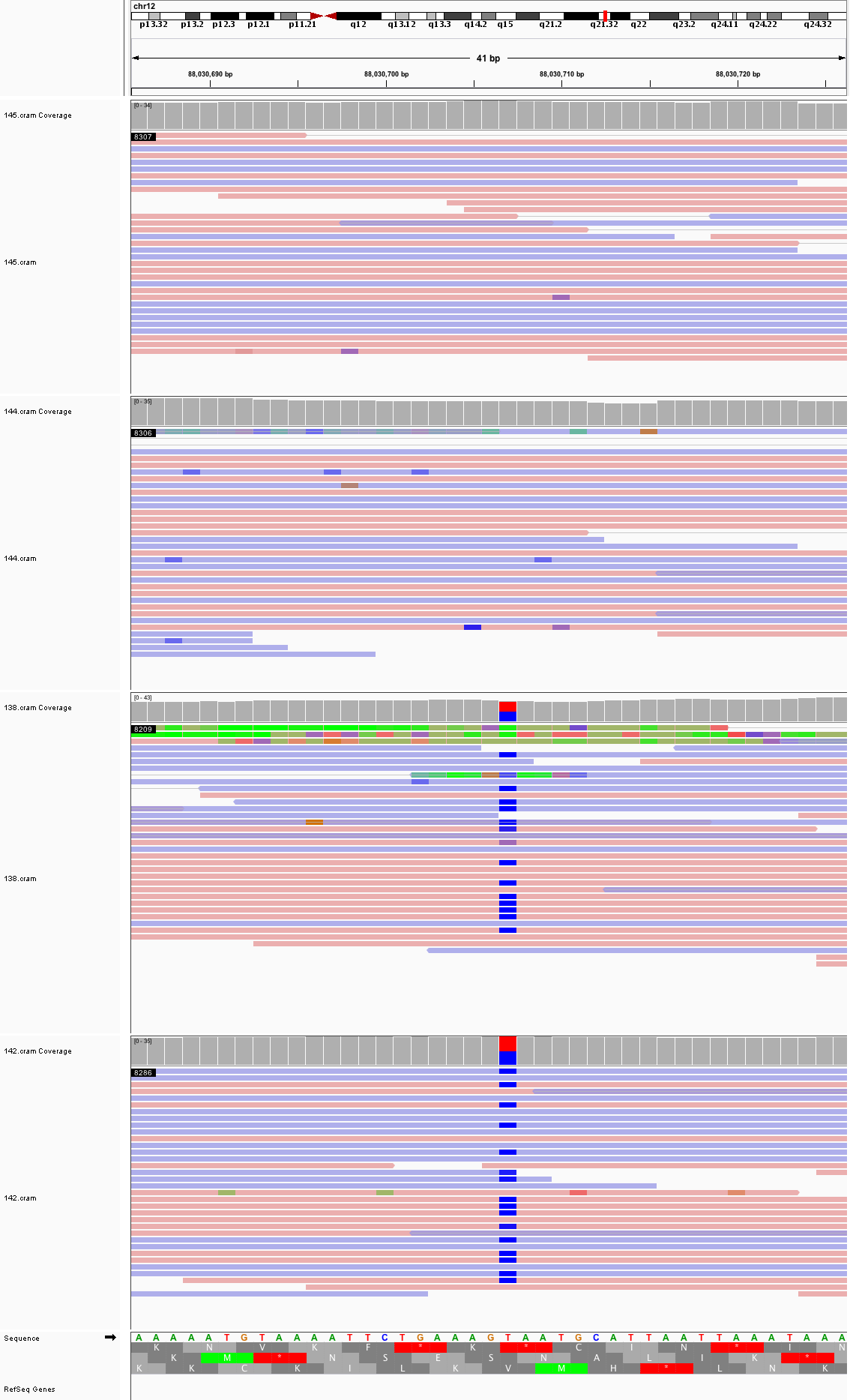

### chr12_130,288,442_130,288,482.png

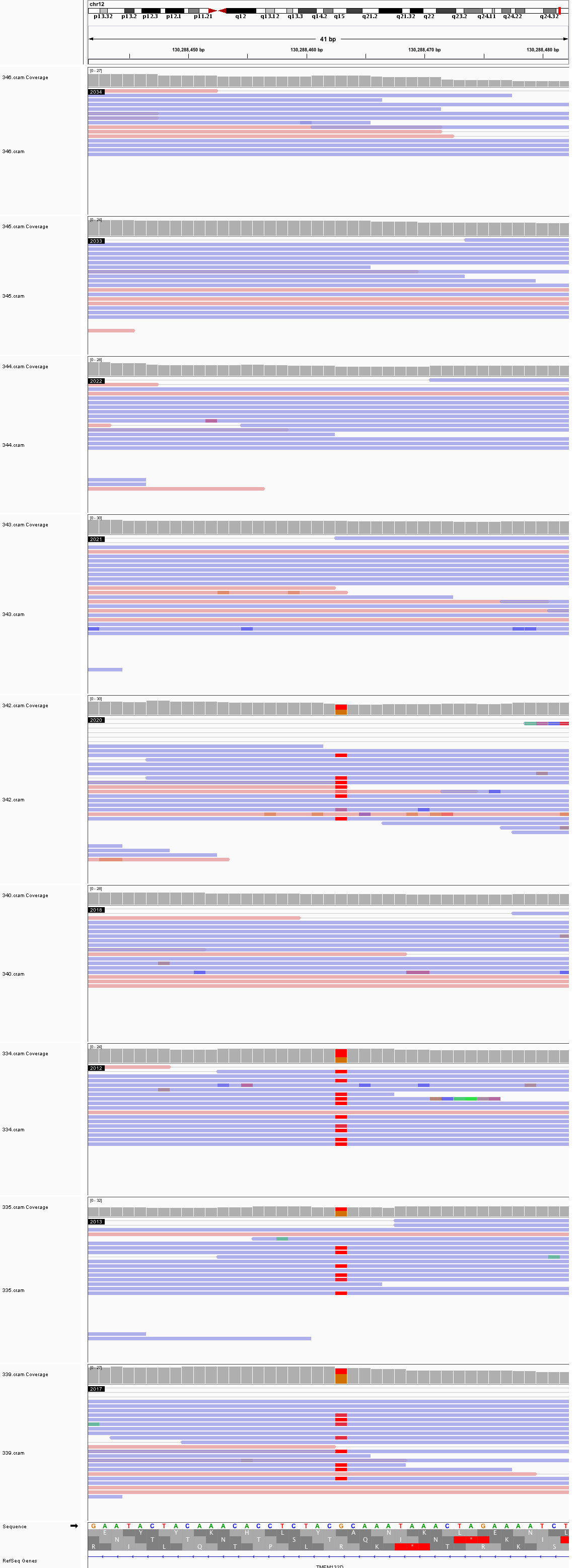

### chr13_23,719,182_23,719,222.png

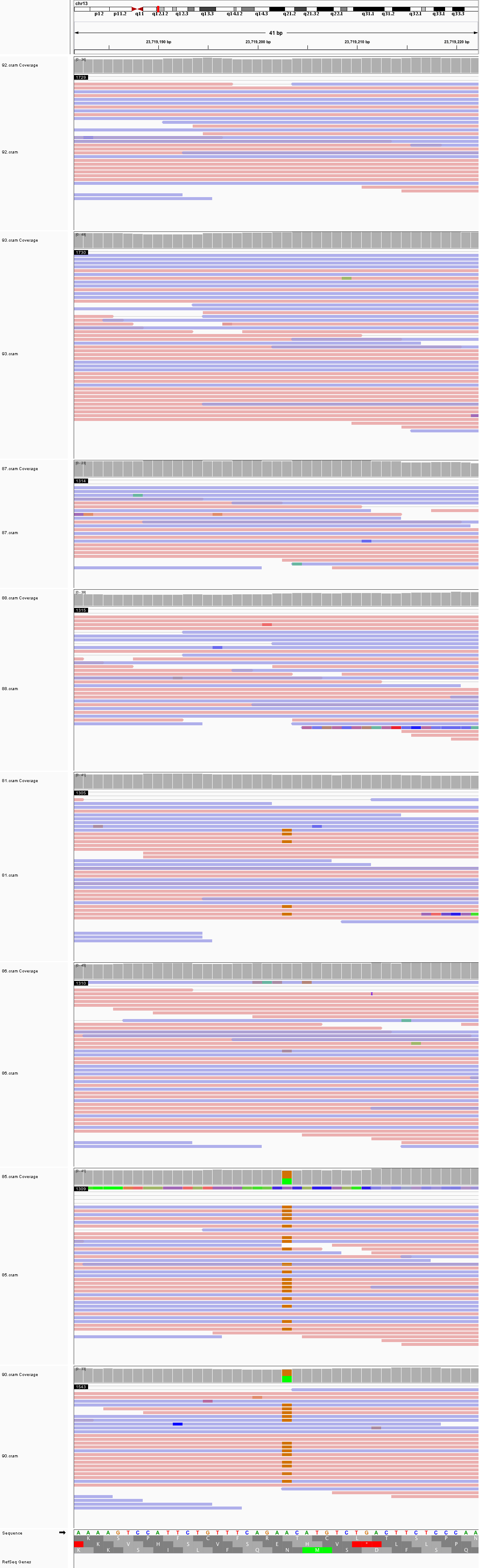

### chr13_41,823,505_41,823,545.png

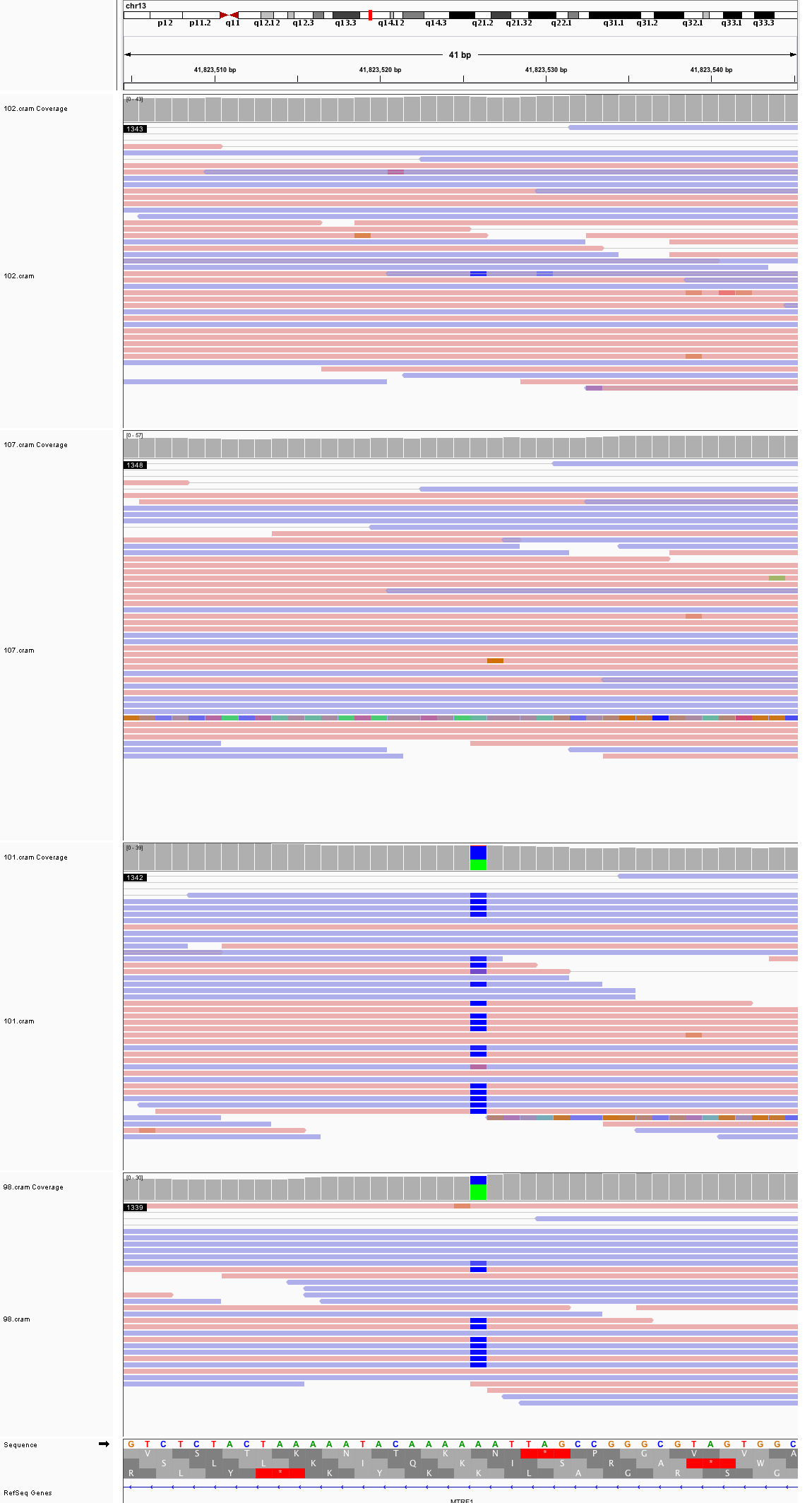

### chr13_85,653,178_85,653,218.png

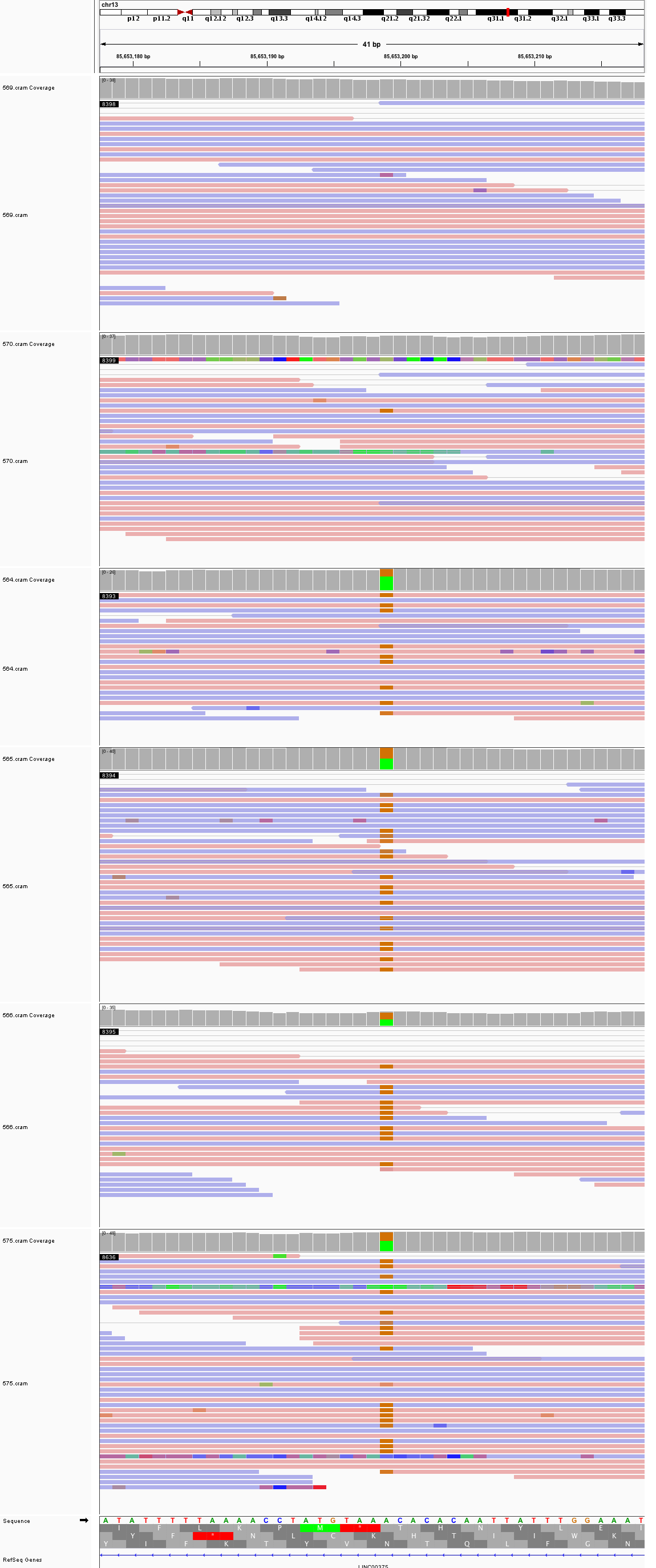

### chr13_89,380,385_89,380,425.png

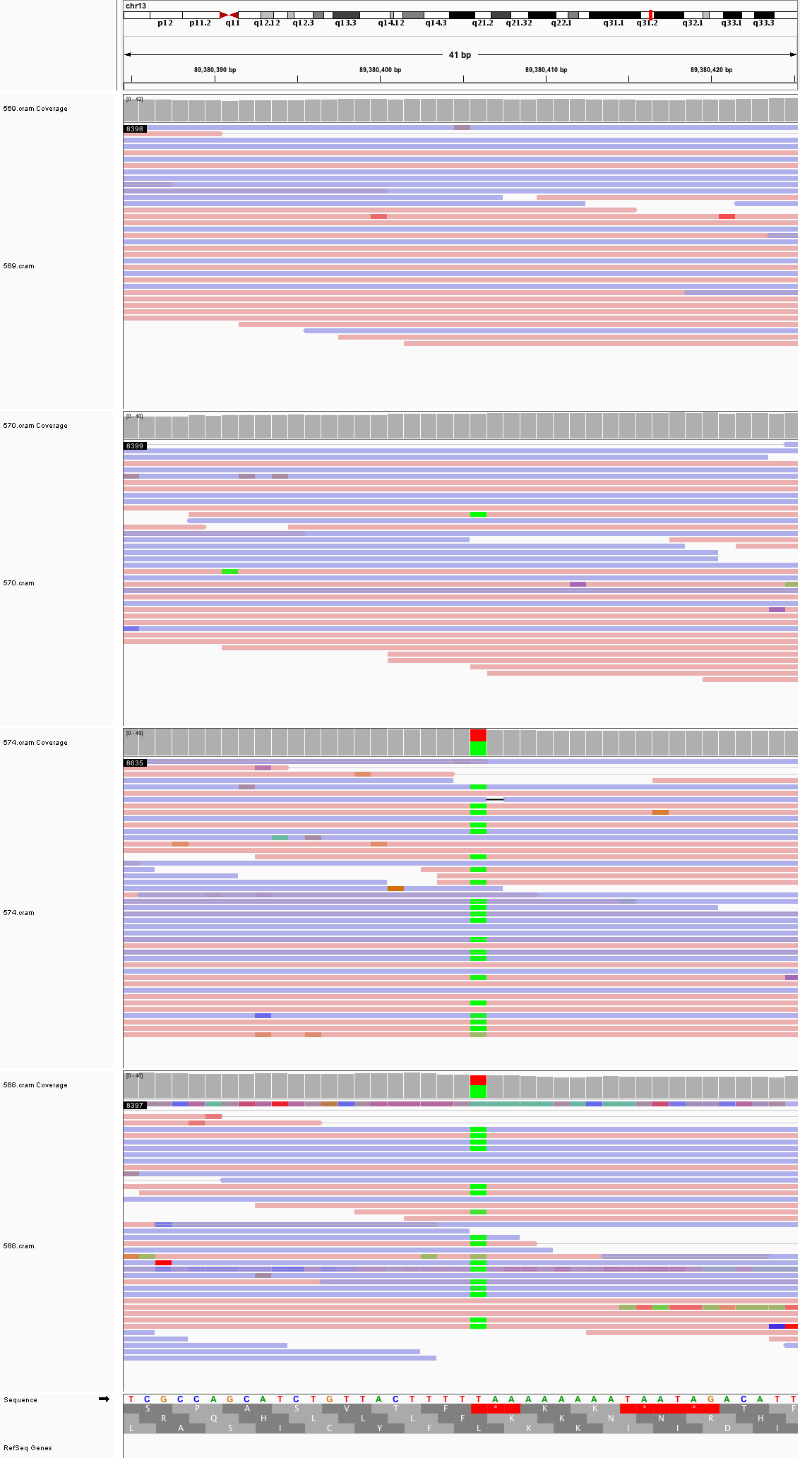

### chr14_26,268,149_26,268,189.png

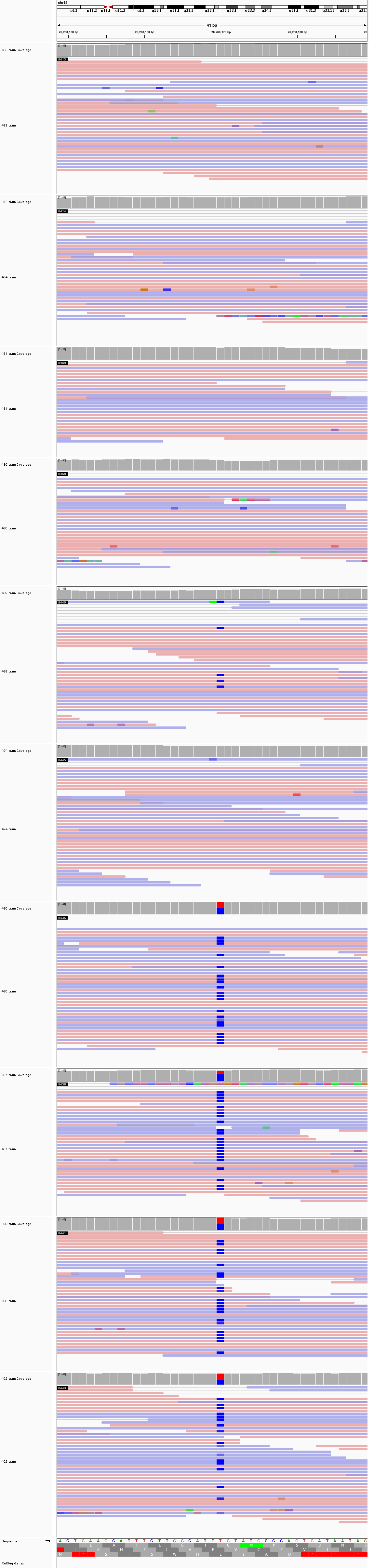

### chr14_26,383,184_26,383,224.png

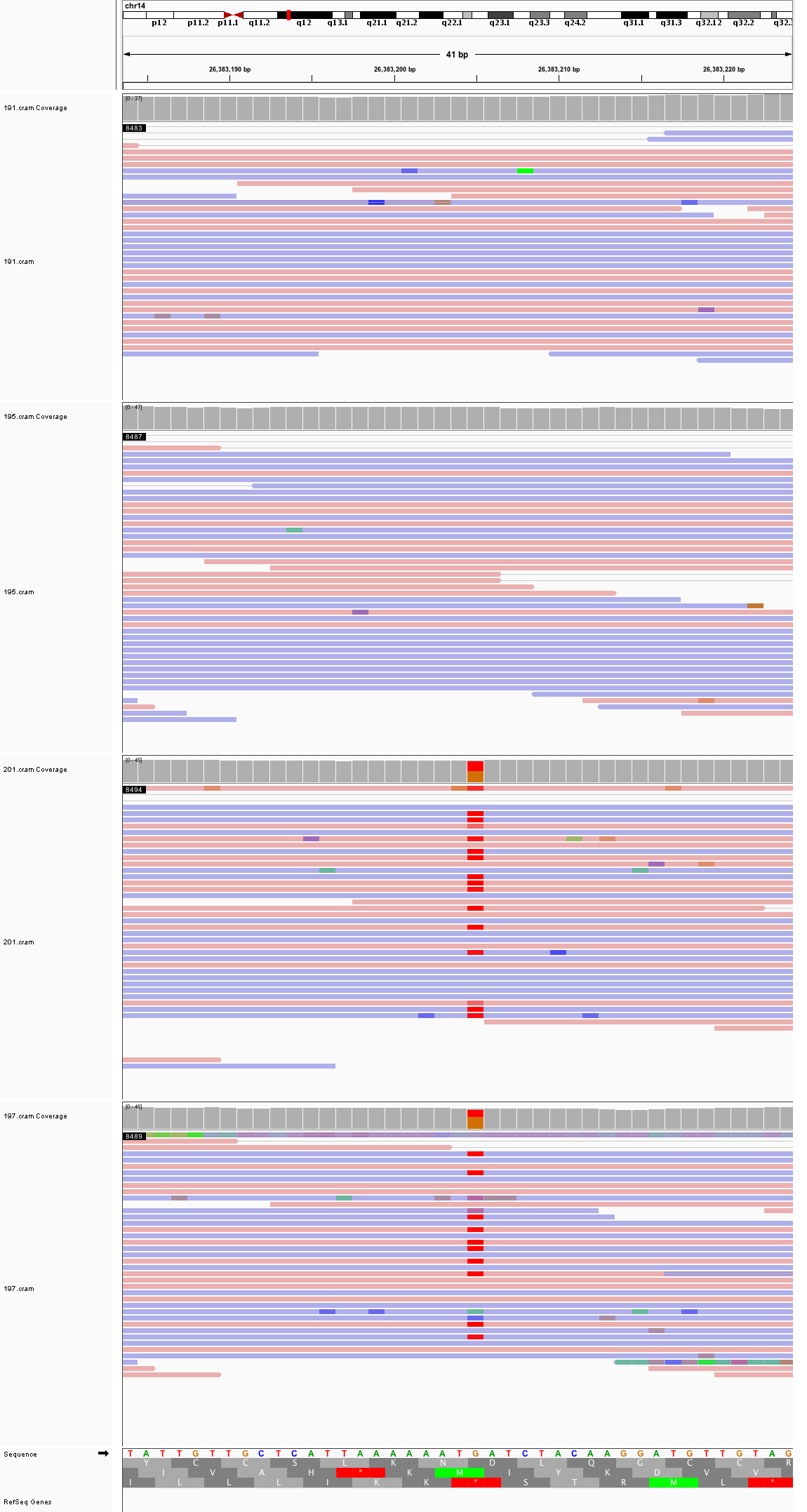

### chr14_78,923,756_78,923,796.png

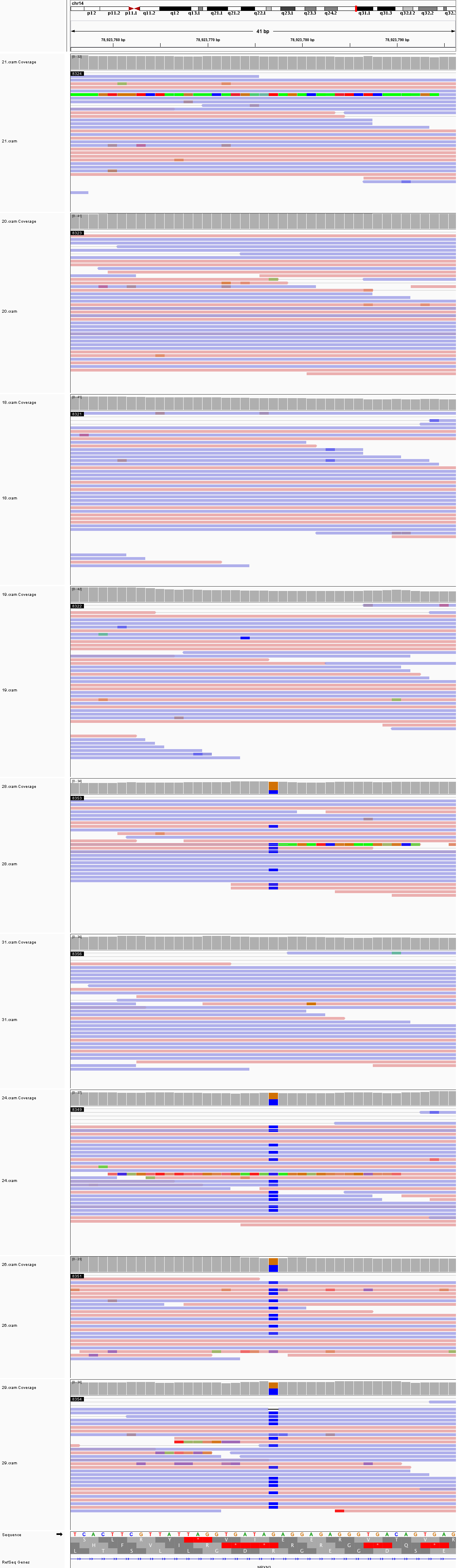

### chr14_84,507,704_84,507,744.png

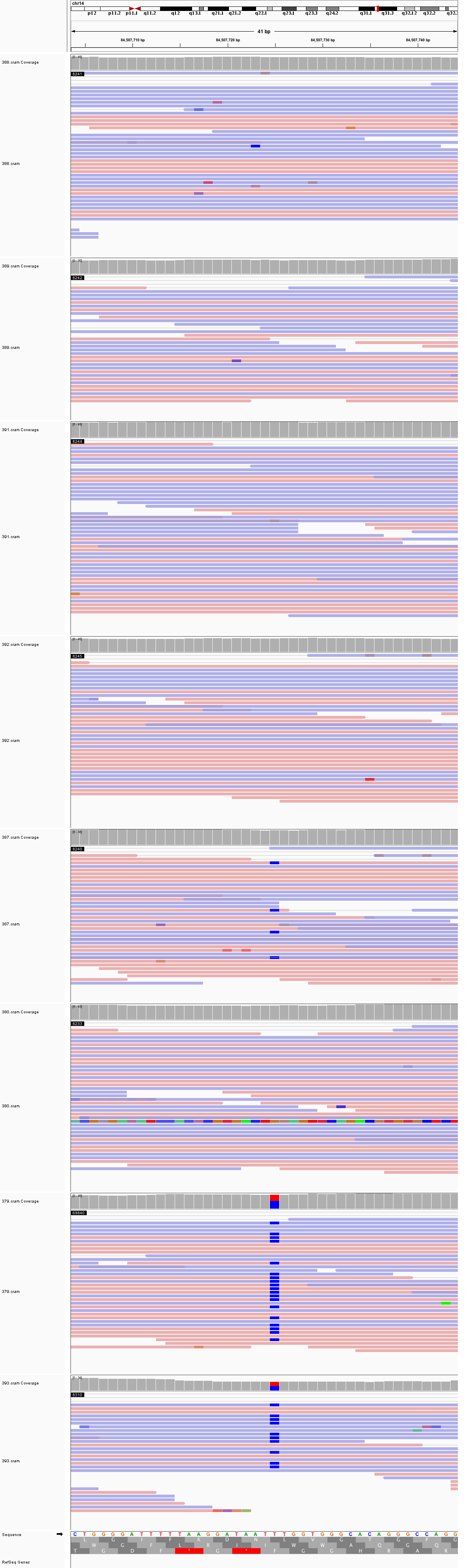

### chr15_20,257,537_20,257,577.png

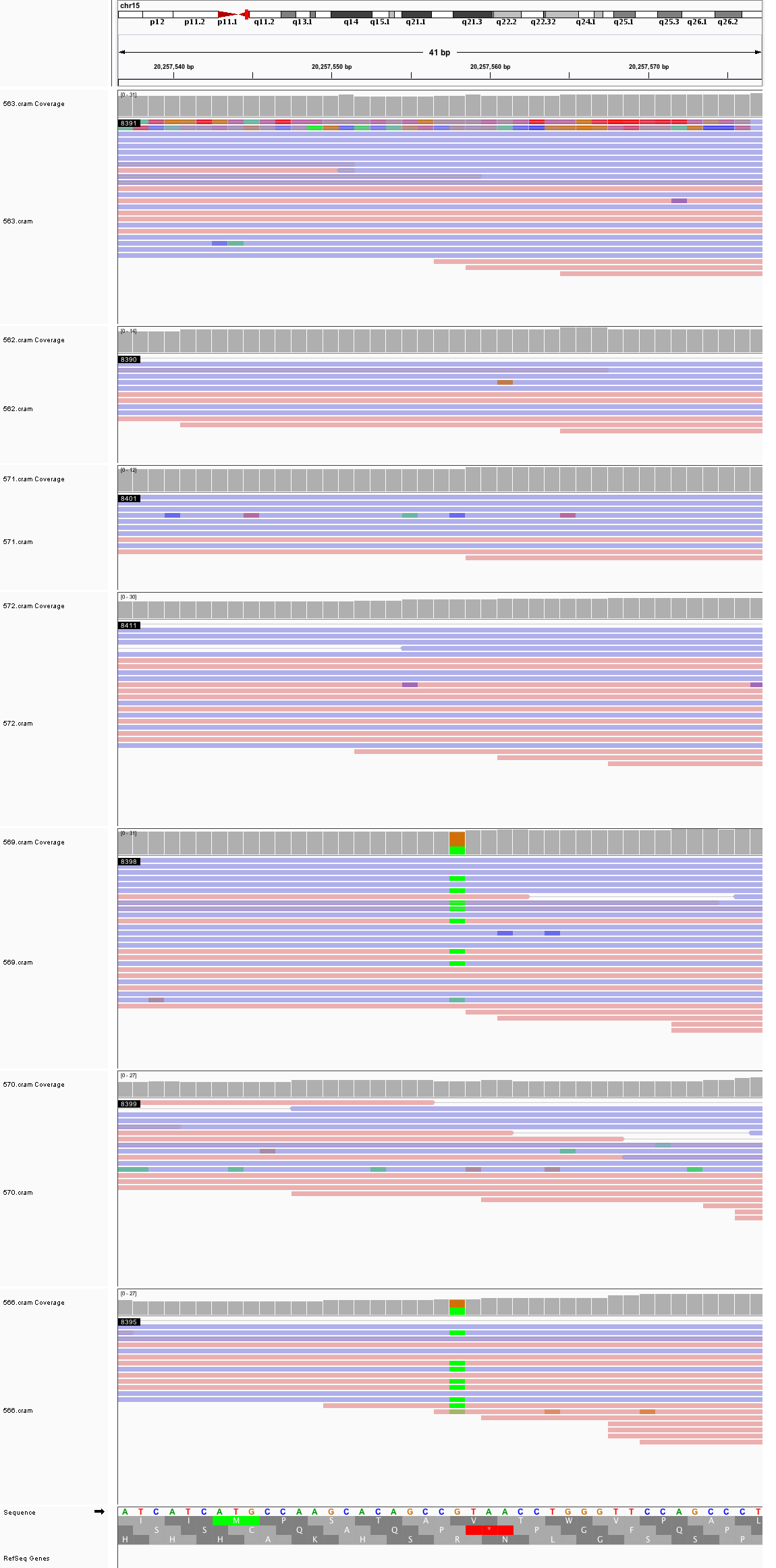

### chr15_32,092,842_32,092,882.png

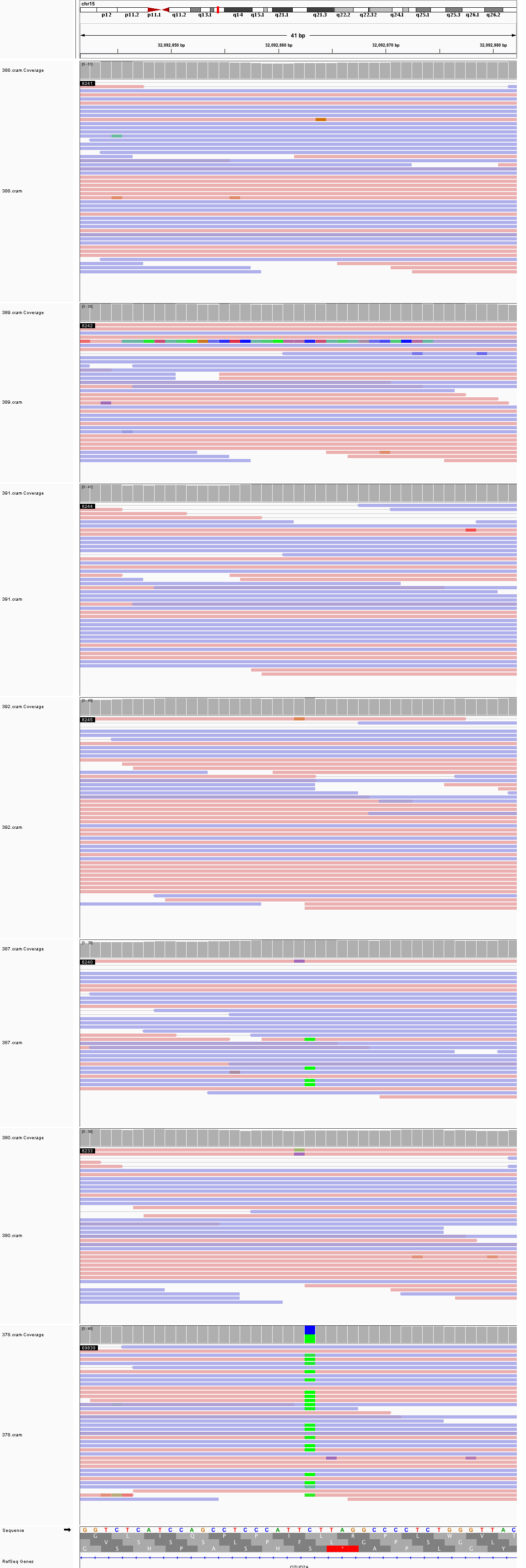

### chr15_41,643,773_41,643,813.png

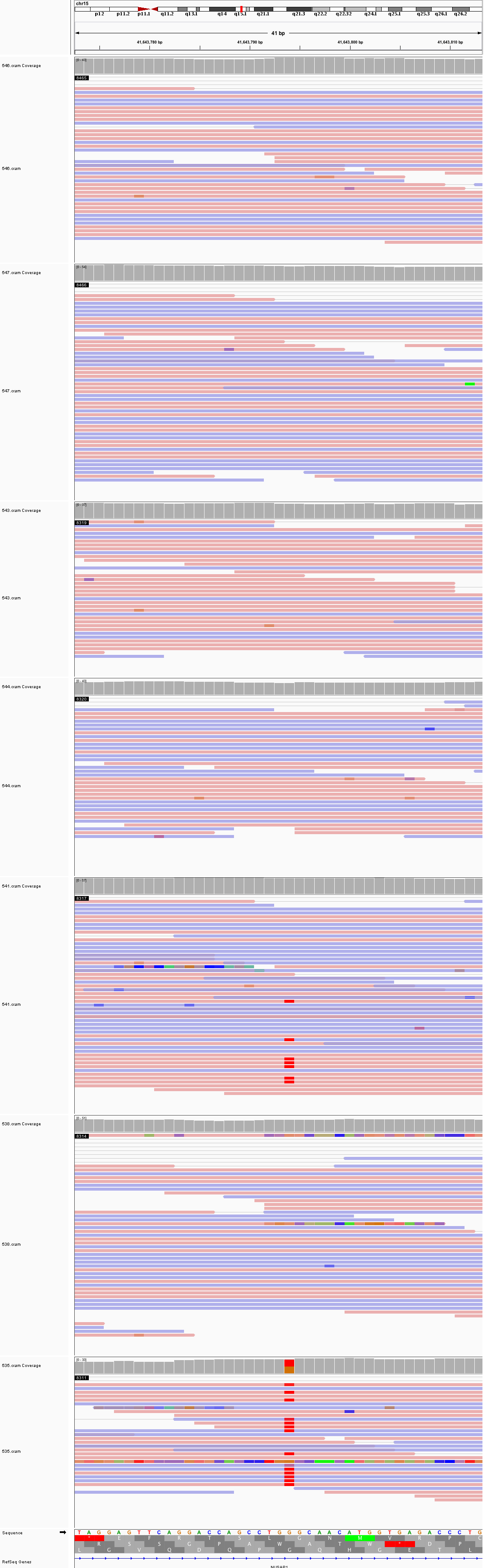

### chr15_52,318,893_52,318,933.png

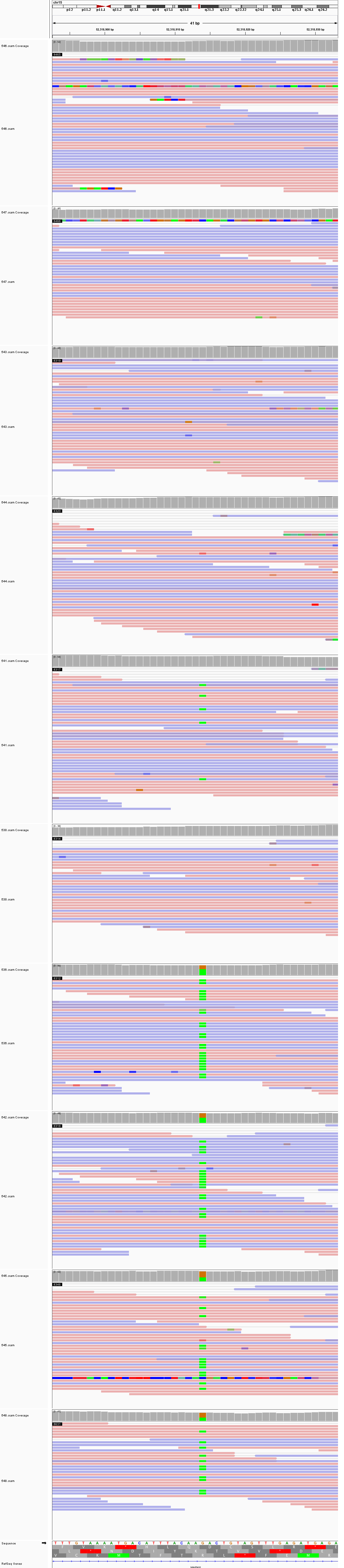

### chr15_64,831,843_64,831,883.png

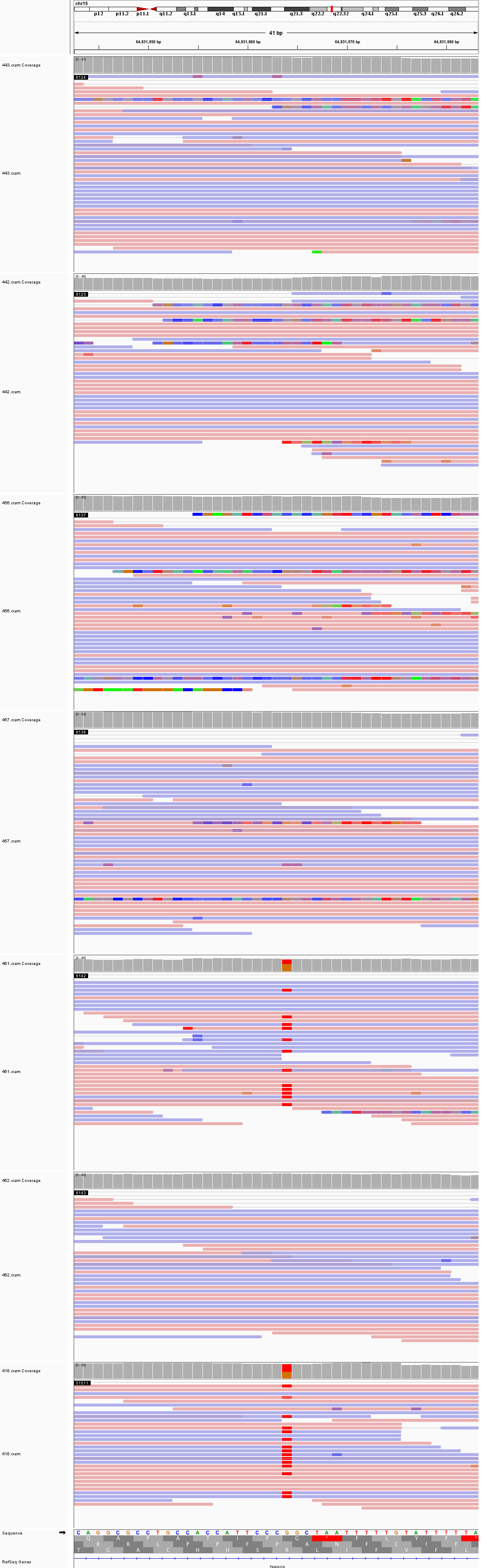

### chr16_15,040,474_15,040,514.png

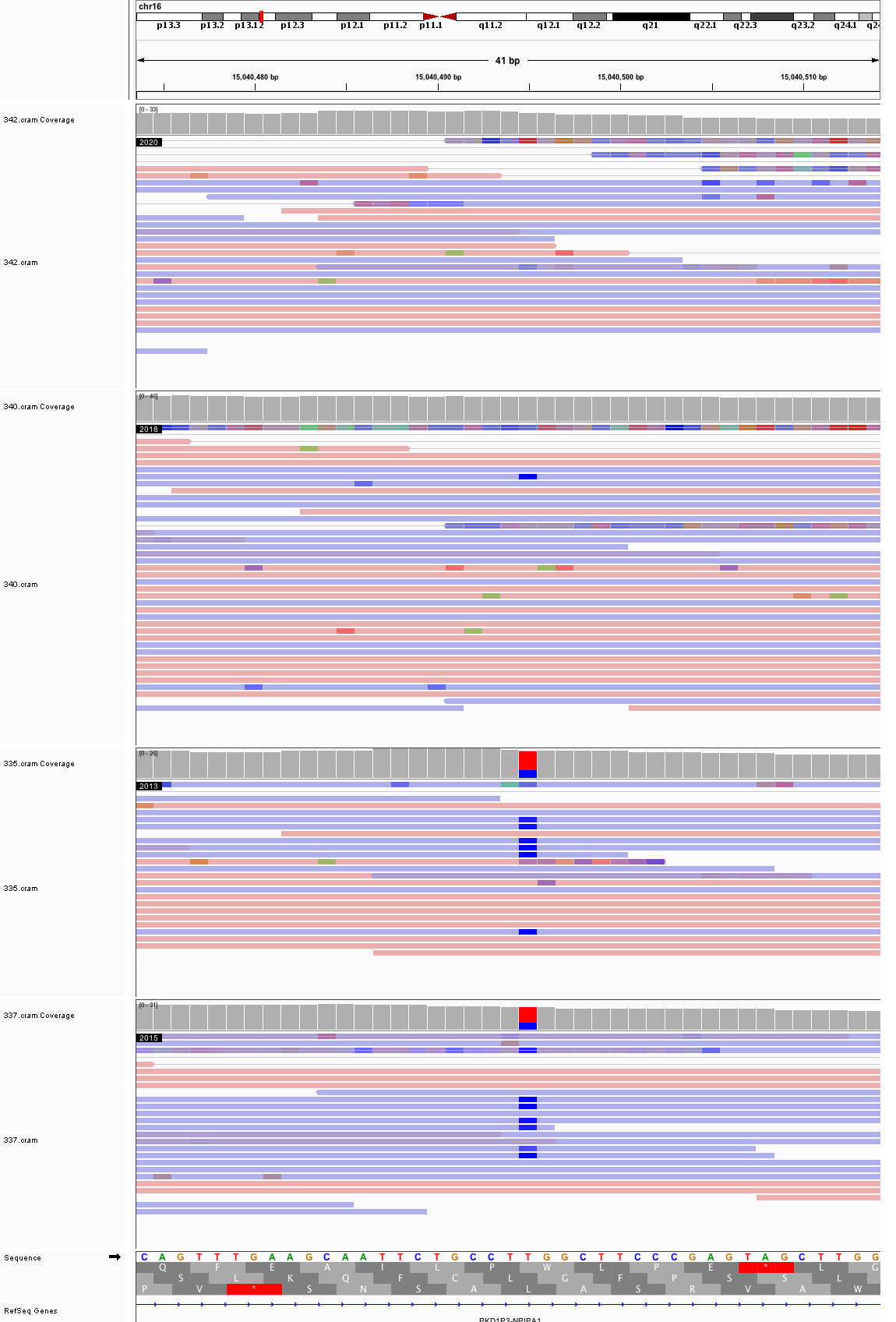

### chr16_18,274,645_18,274,685.png

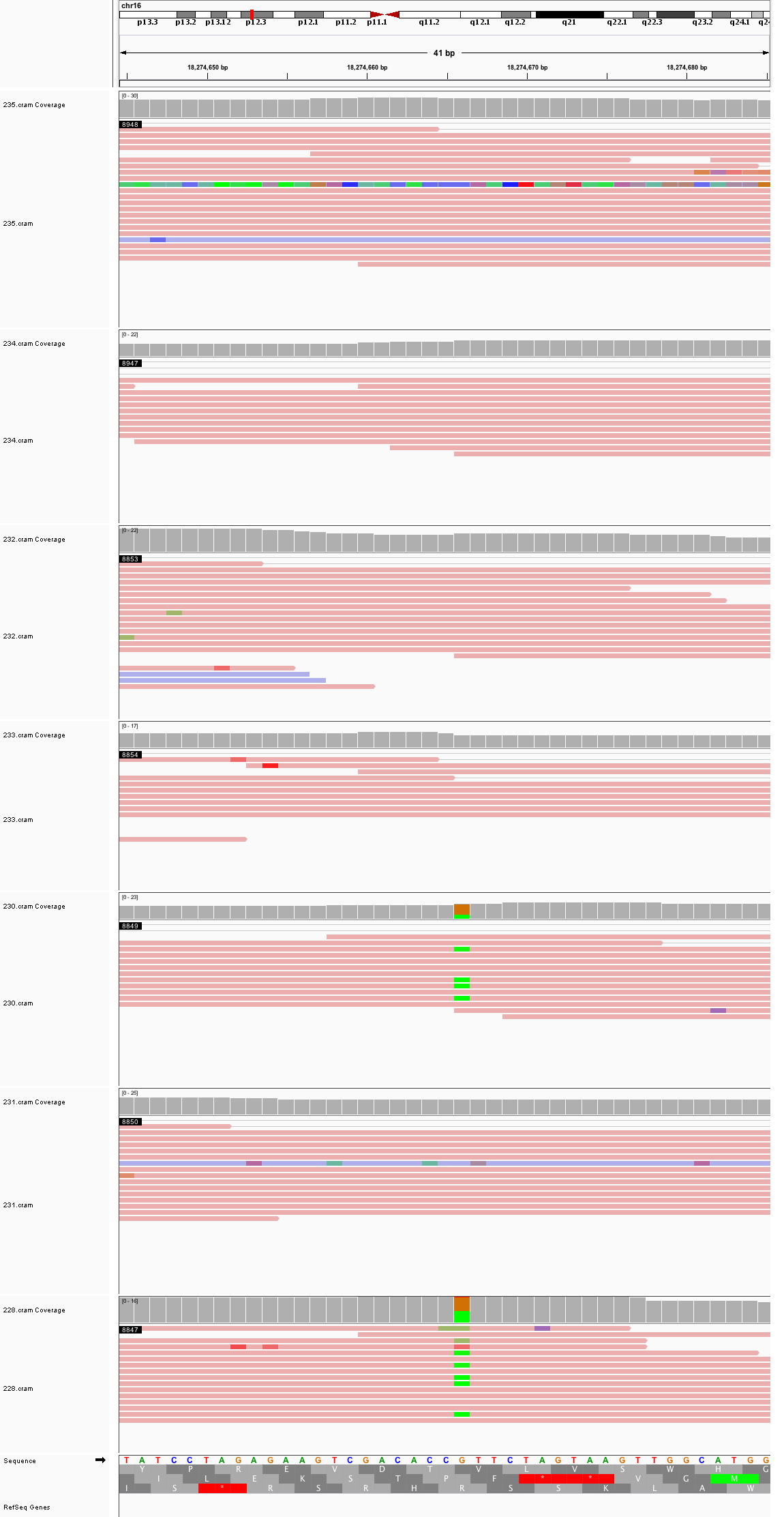

### chr16_18,341,905_18,341,945.png

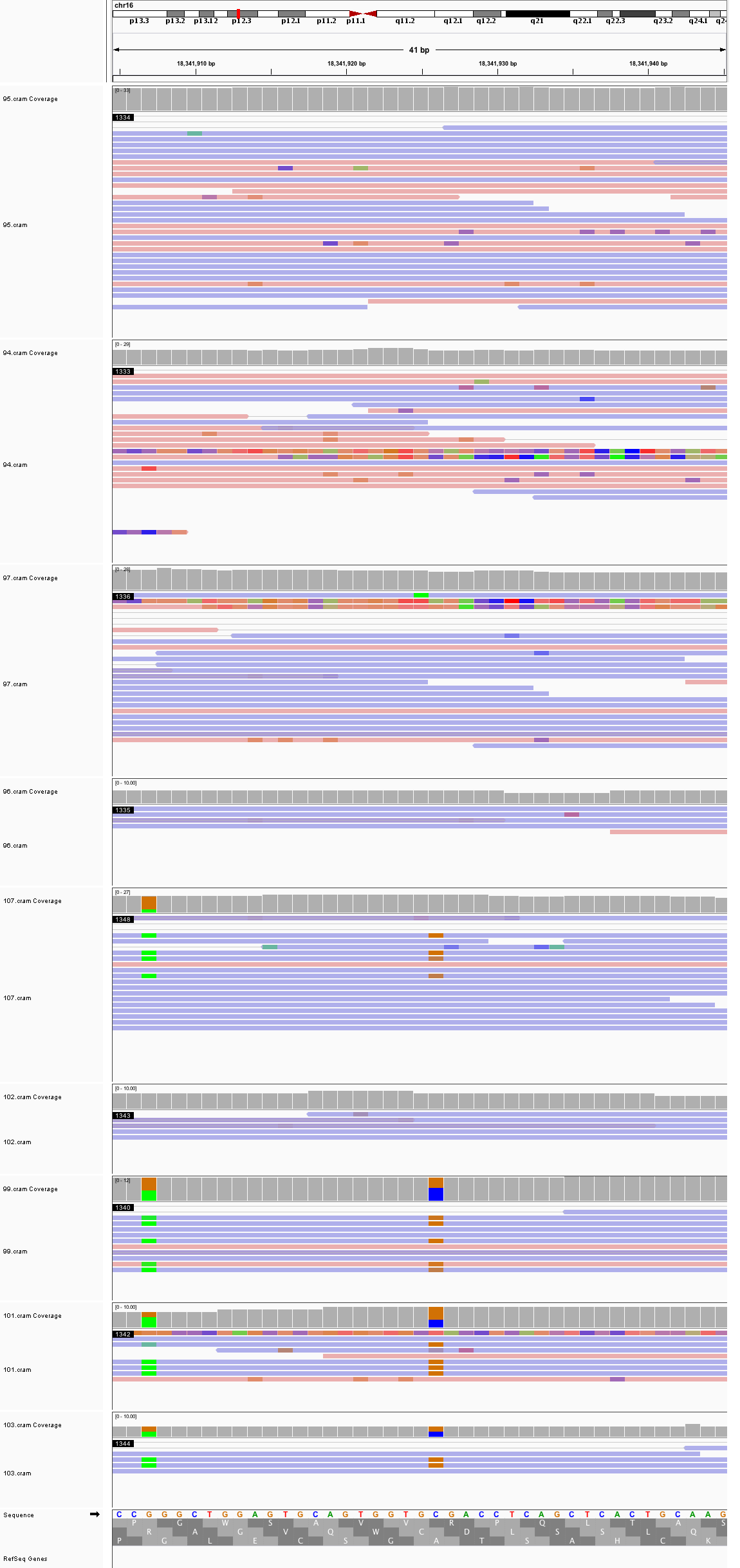

### chr16_62,148,786_62,148,826.png

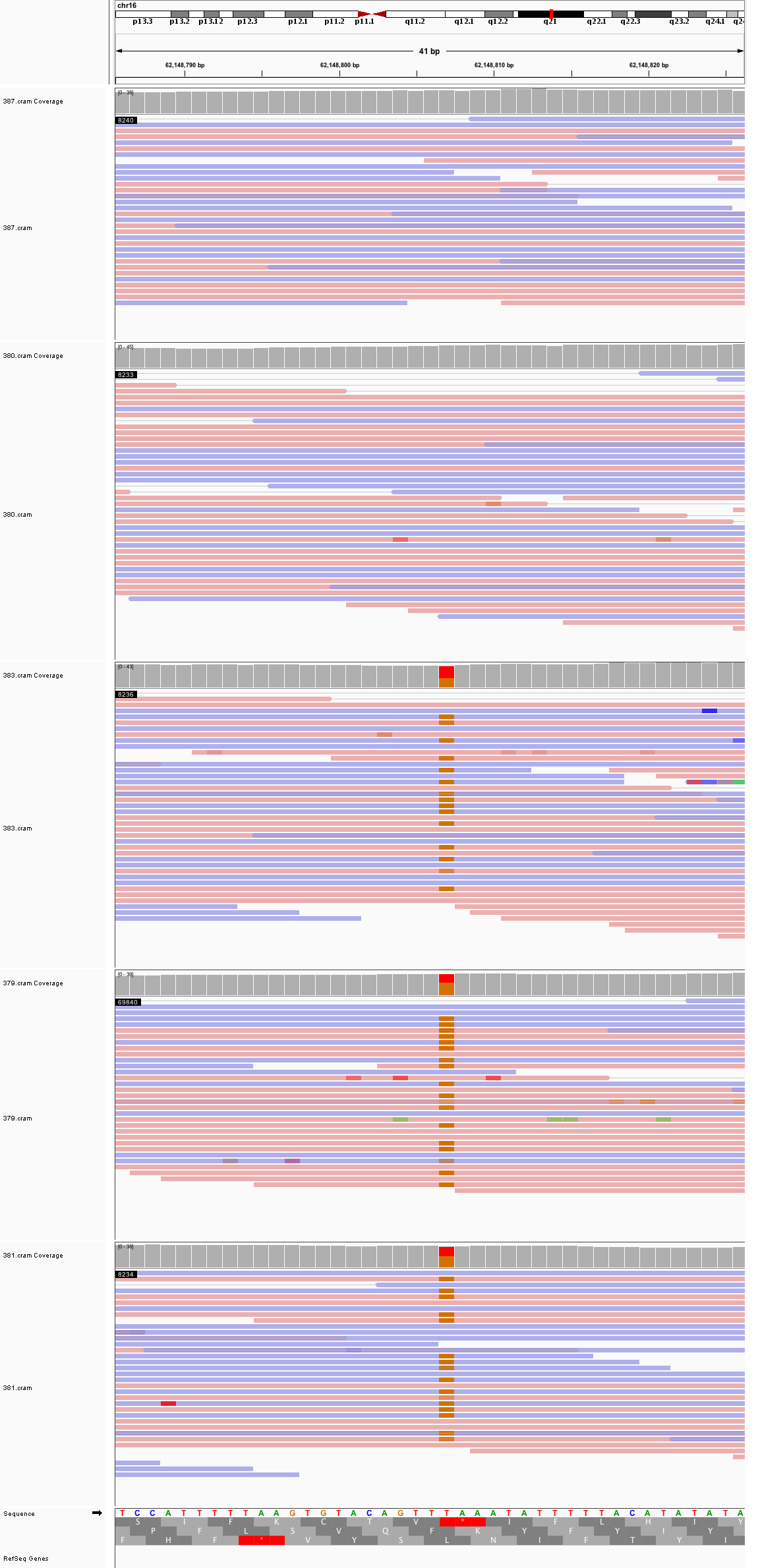

### chr16_74,677,729_74,677,769.png

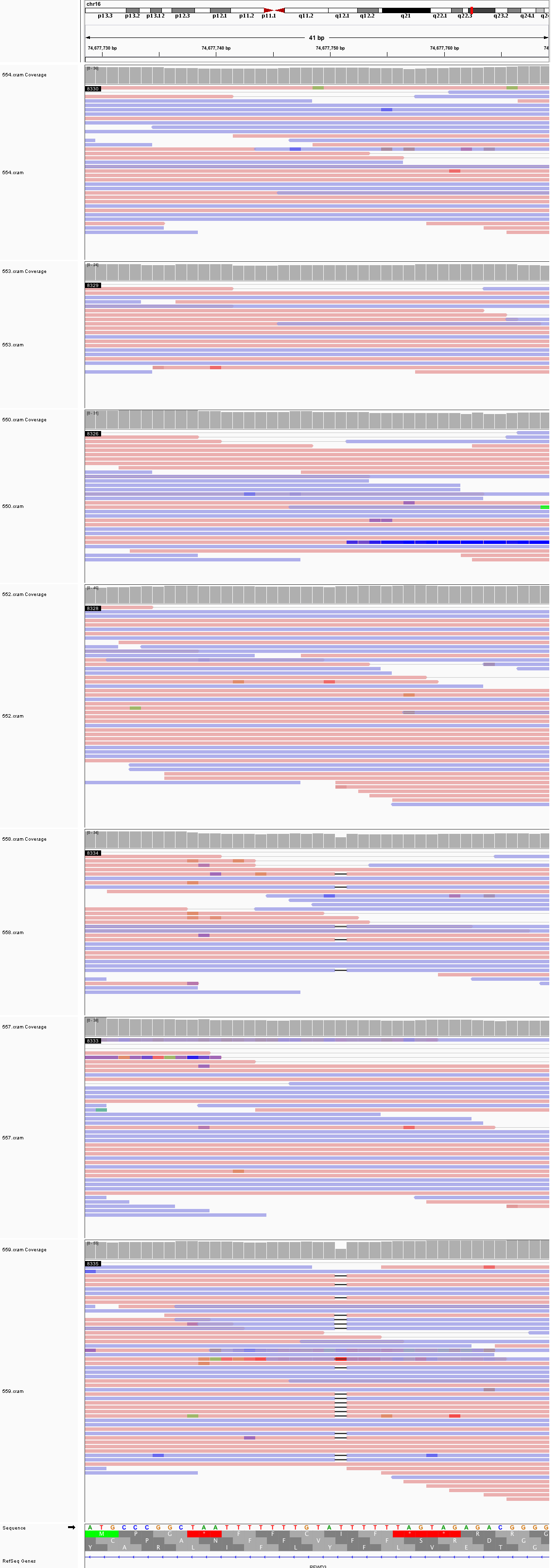

### chr17_17,760,580_17,760,620.png

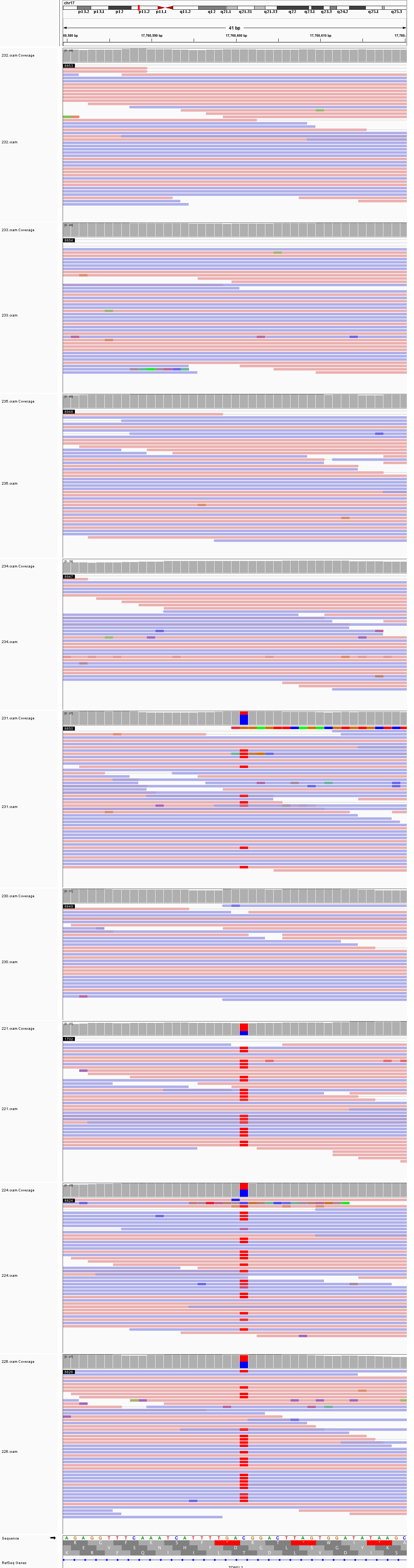

### chr17_21,337,441_21,337,481.png

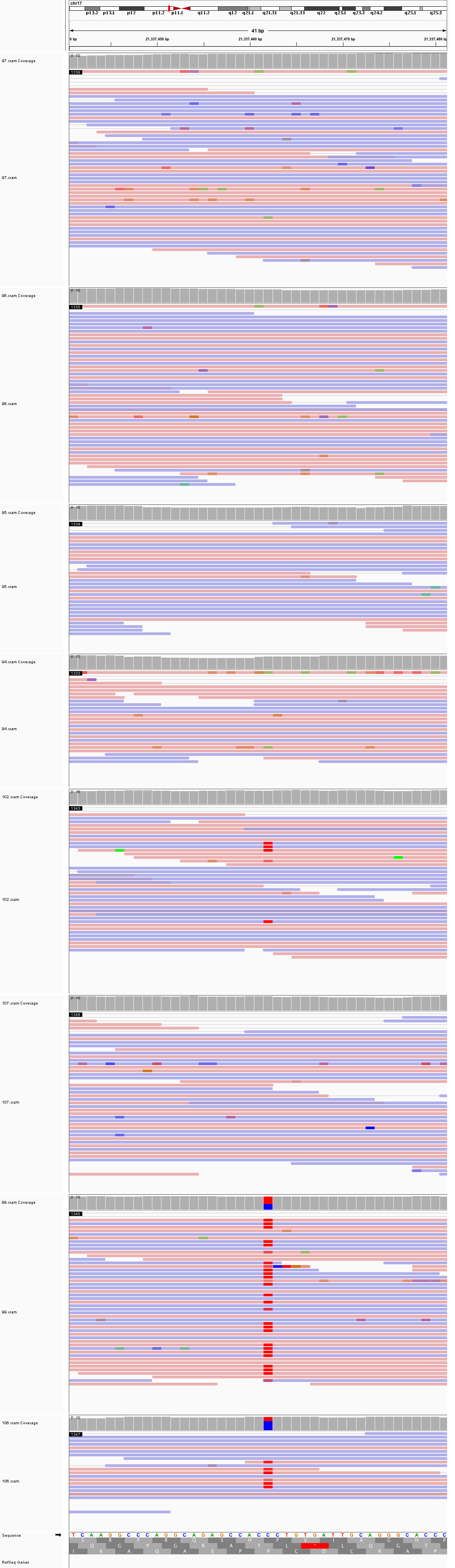

### chr17_62,001,144_62,001,184.png

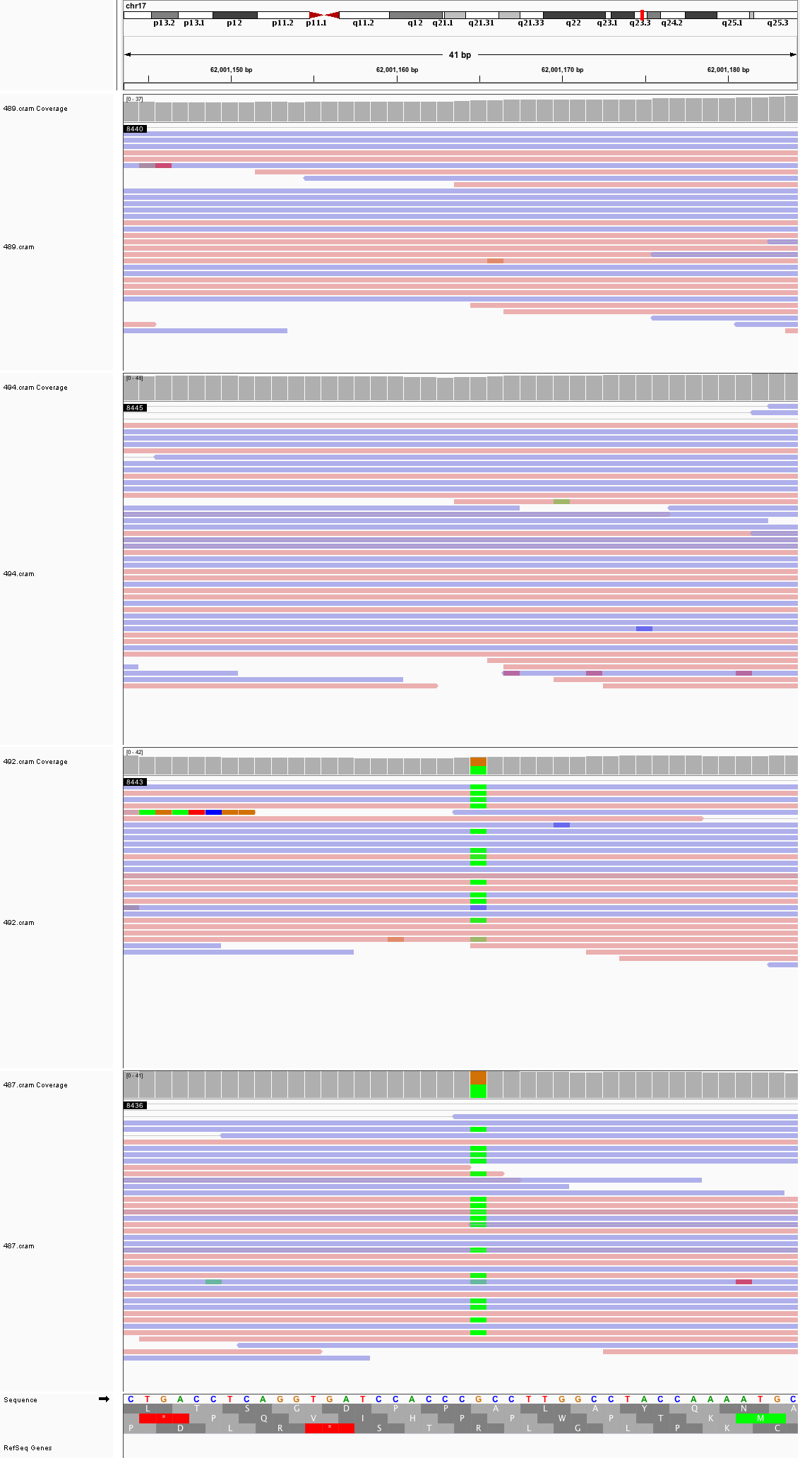

### chr17_79,069,065_79,069,105.png

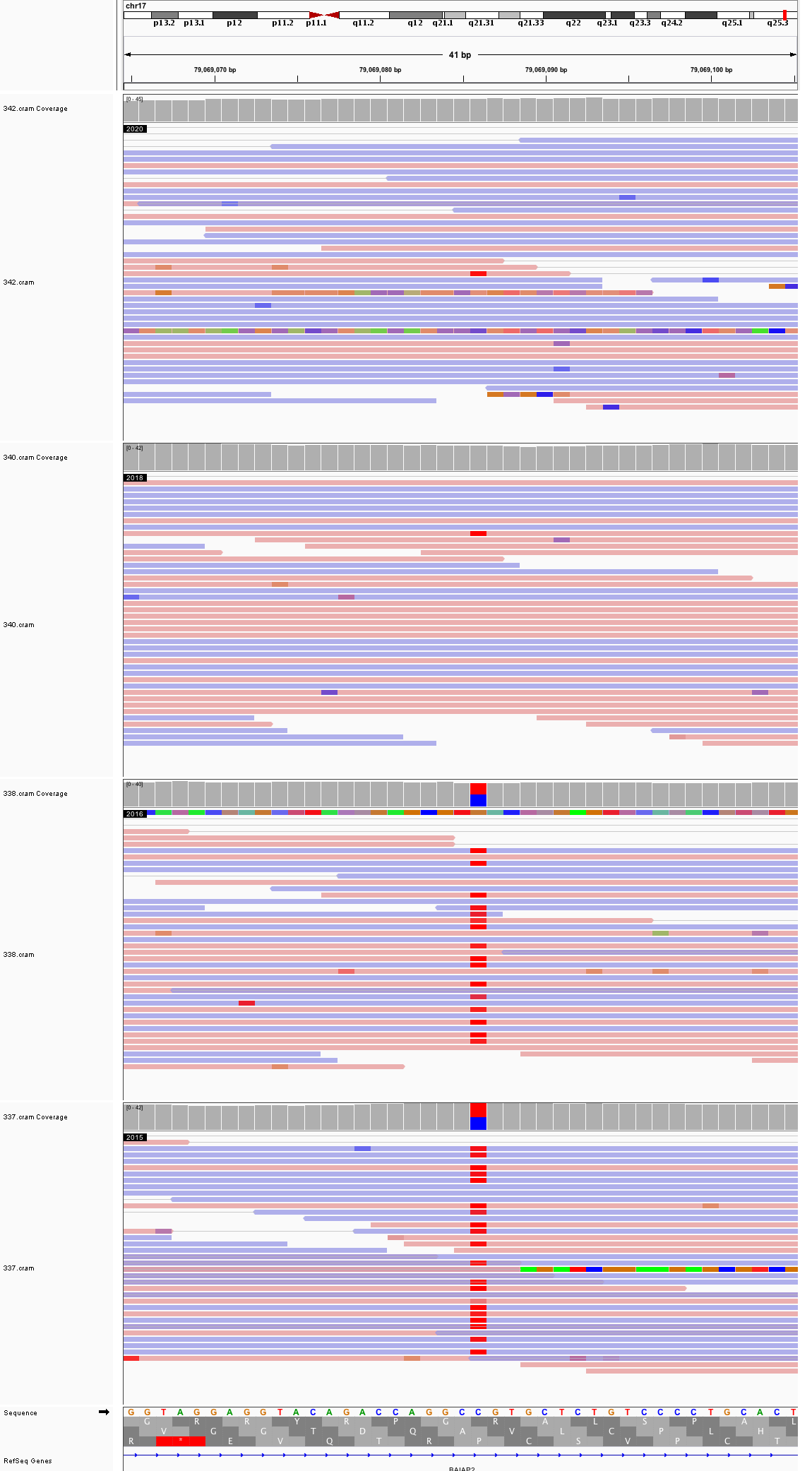

### chr18_6,070,553_6,070,593.png

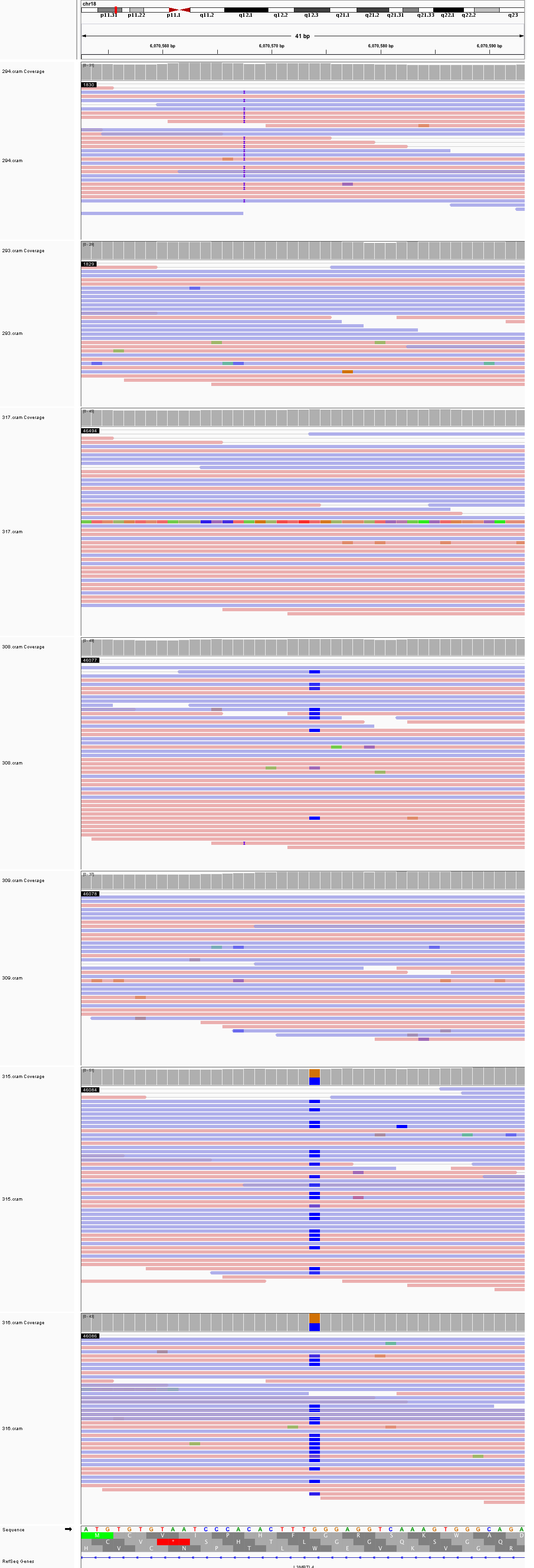

### chr18_63,353,177_63,353,217.png

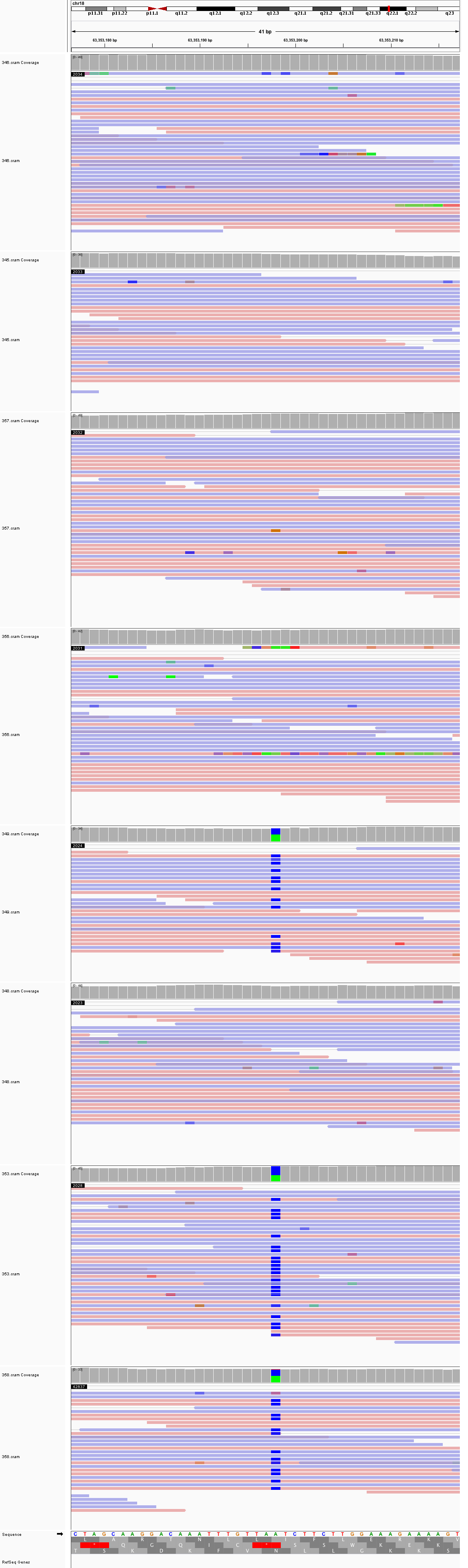

### chr18_67,177,300_67,177,340.png

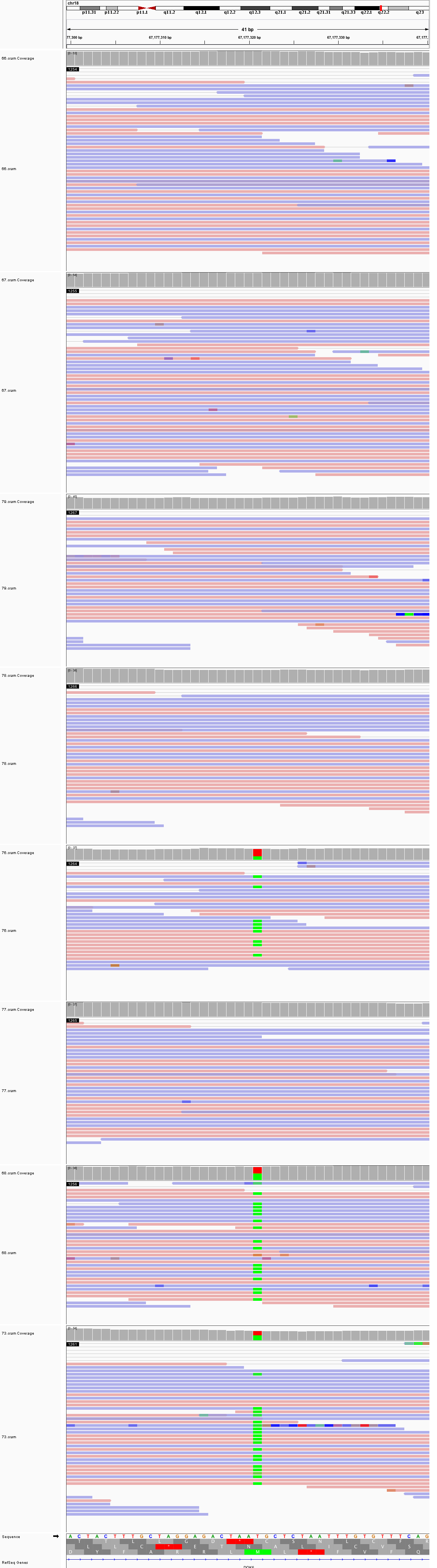

### chr19_42,668,635_42,668,675.png

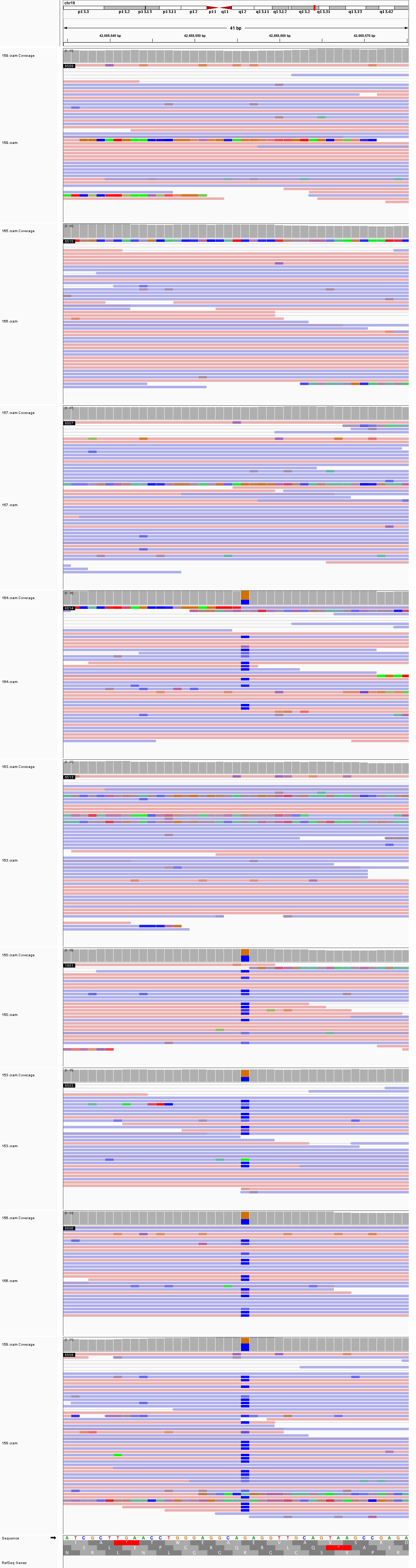

### chr19_840,697_840,737.png

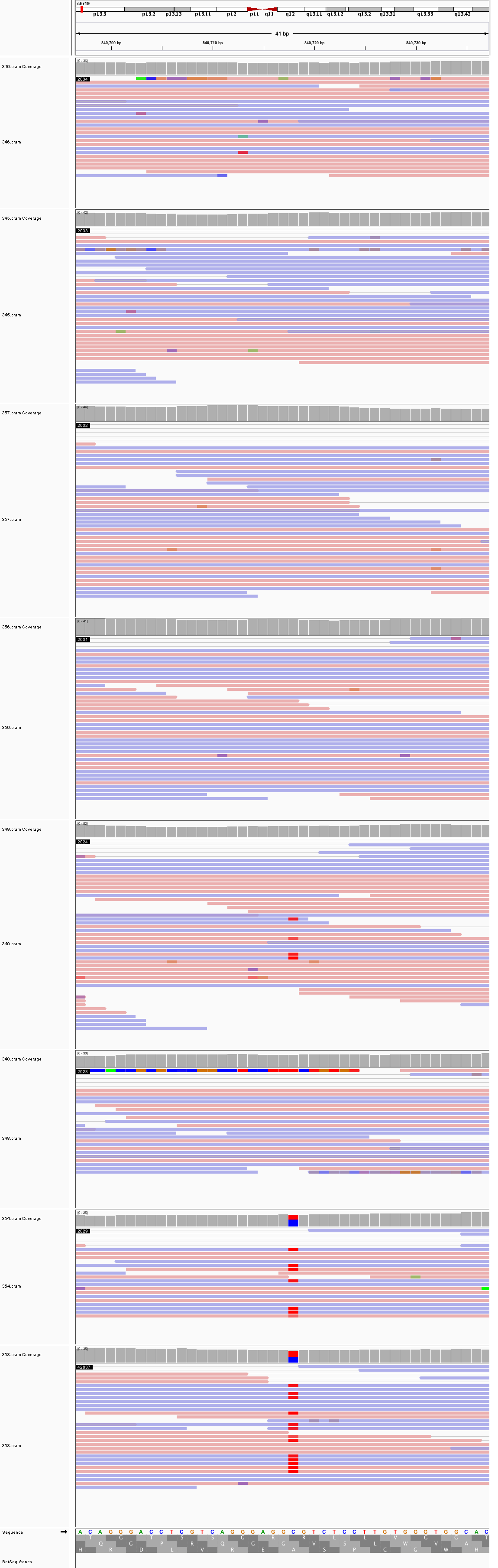

### chr21_29,349,992_29,350,032.png

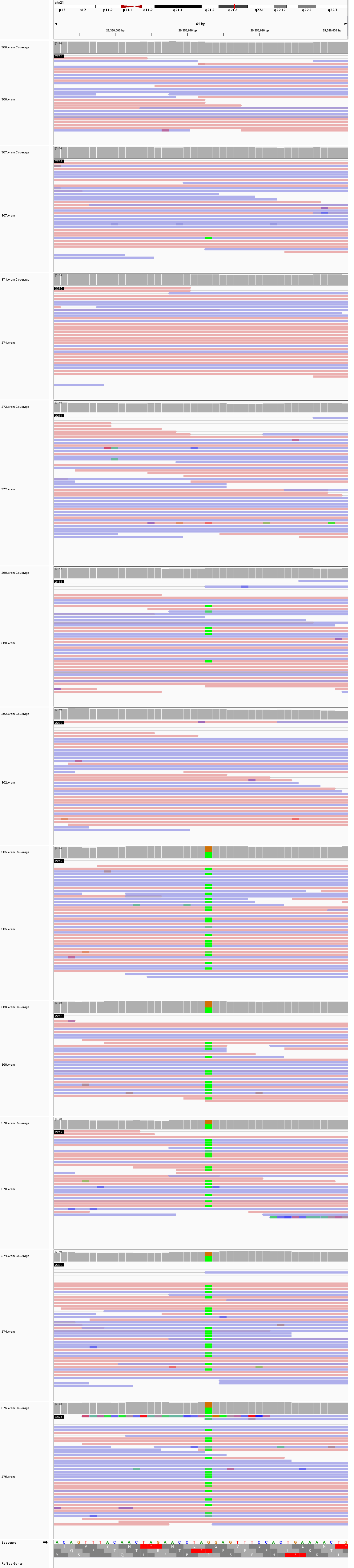

### chrX_24,645,645_24,645,685.png

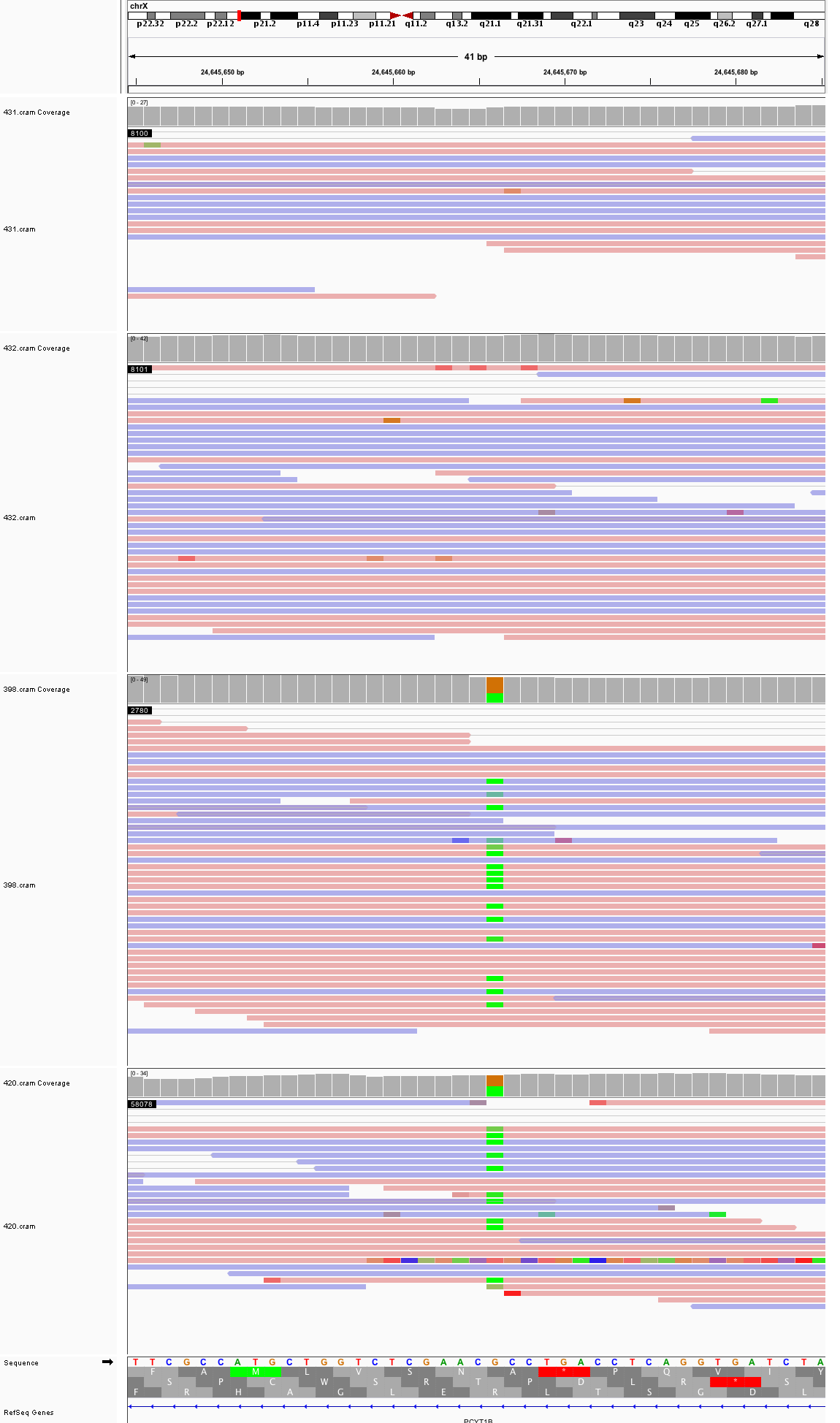

### chrX_127,388,545_127,388,585.png

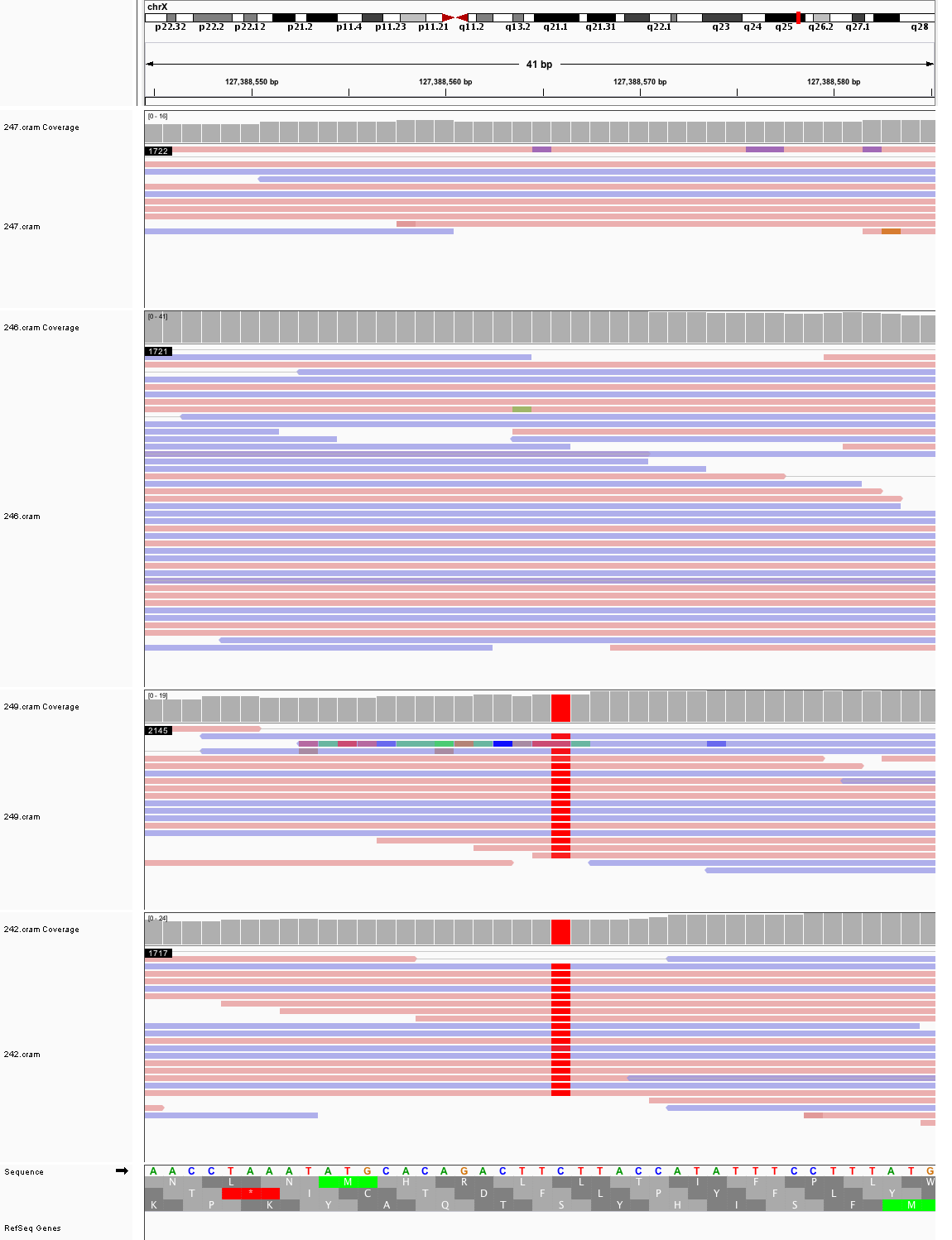
