## Supplementary figures and images for "Large, three-generation CEPH families reveal post-zygotic mosaicism and variability in germline mutation accumulation"

### chr1_27,473,916_27,473,956.png

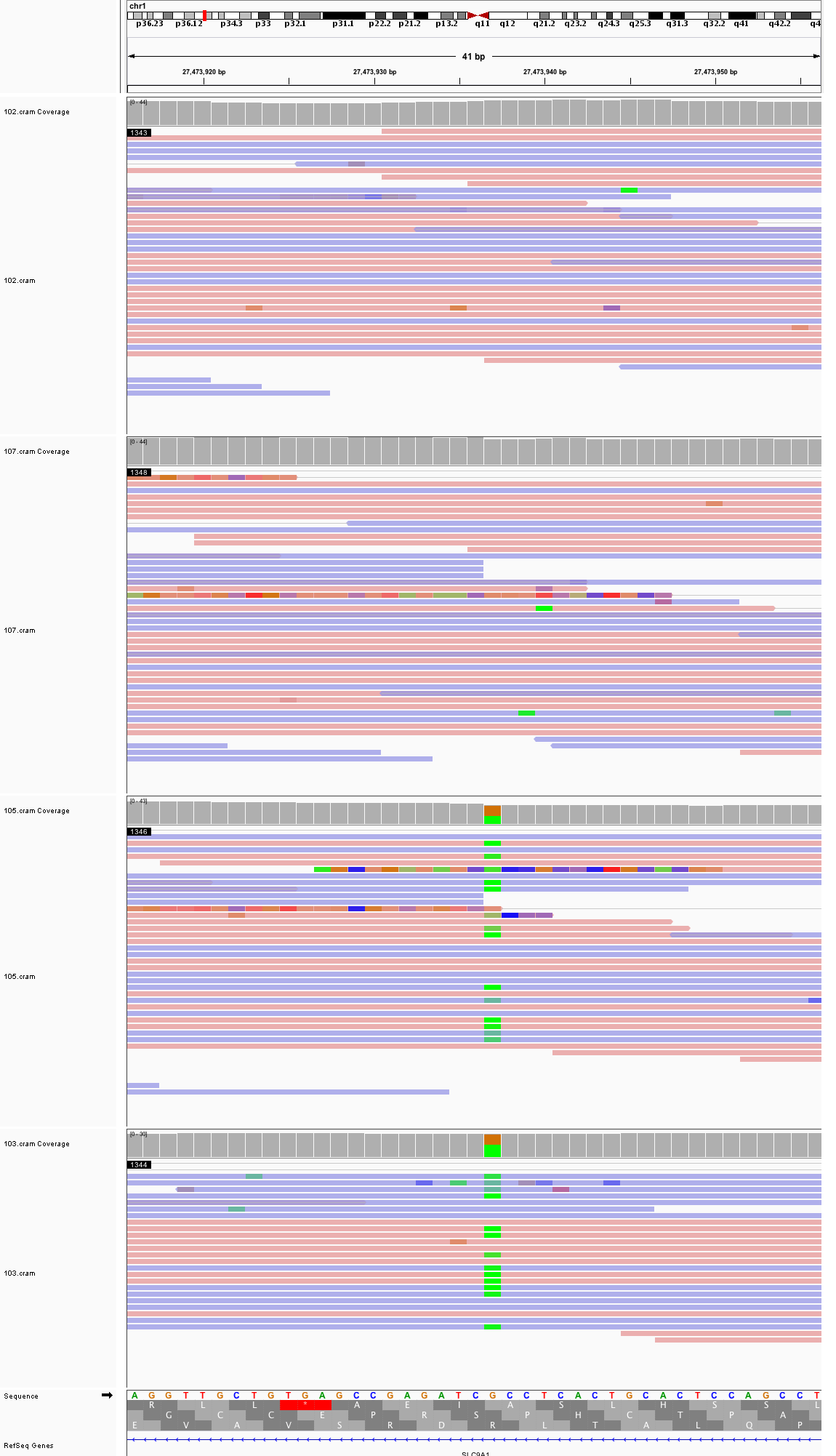

### chr1_33,270,195_33,270,235.png

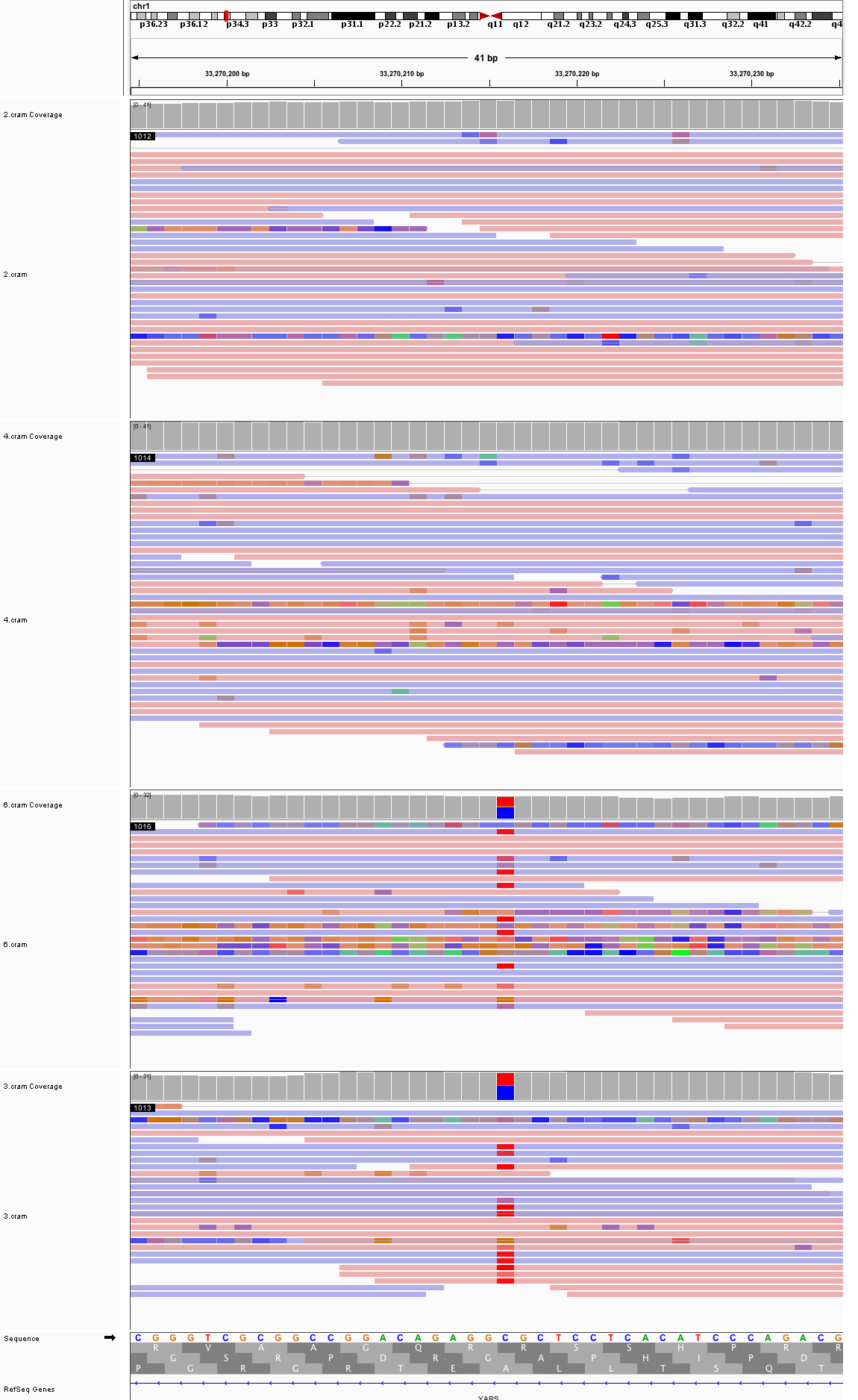

### chr1_224,608,759_224,608,799.png

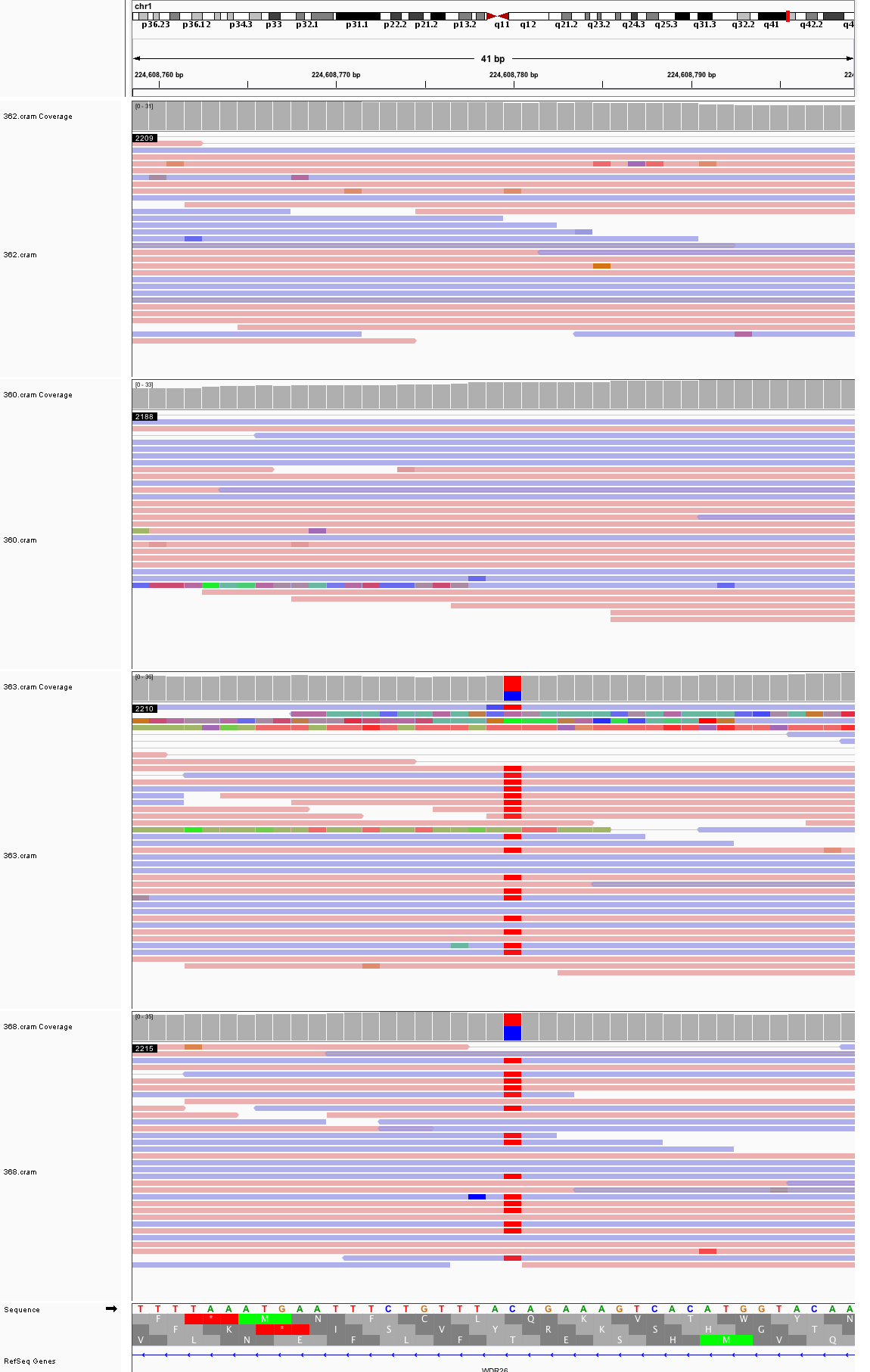

### chr1_235,501,960_235,502,000.png

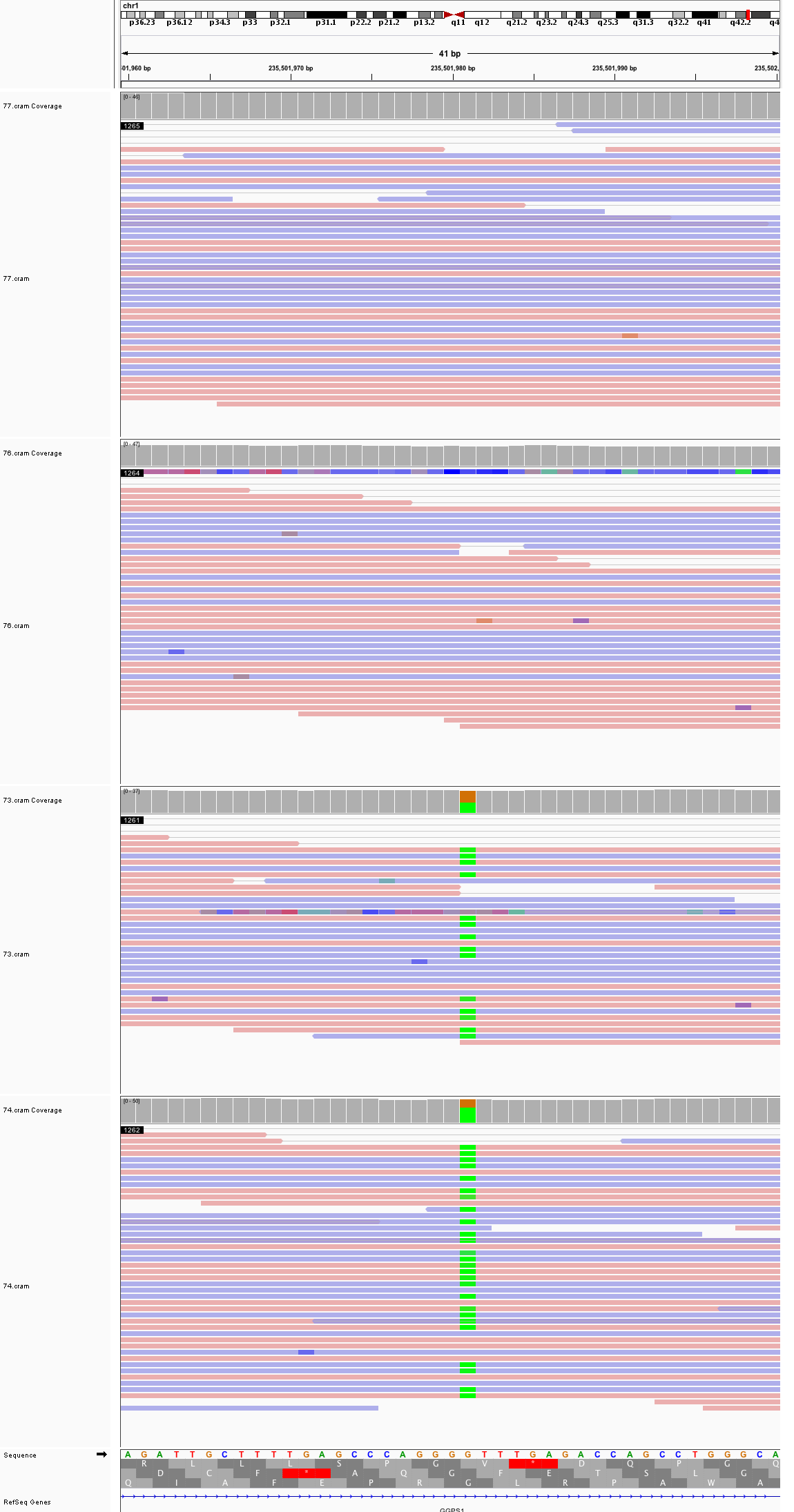

### chr2_12,712,380_12,712,420.png

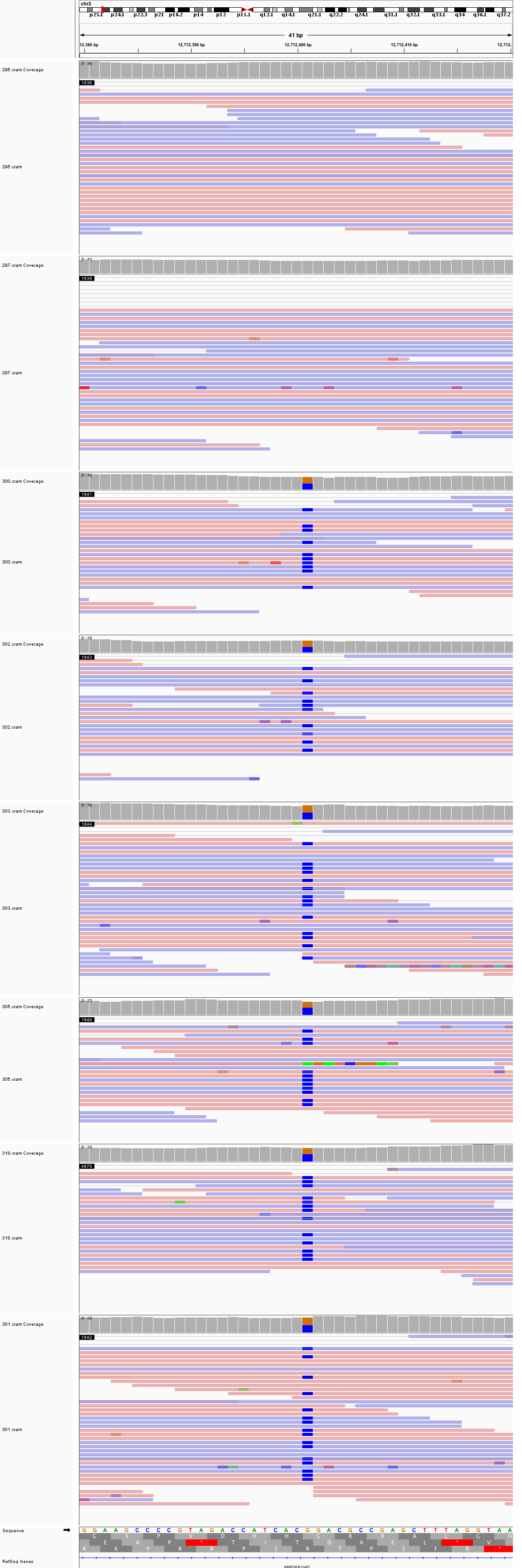

### chr2_54,569,426_54,569,466.png

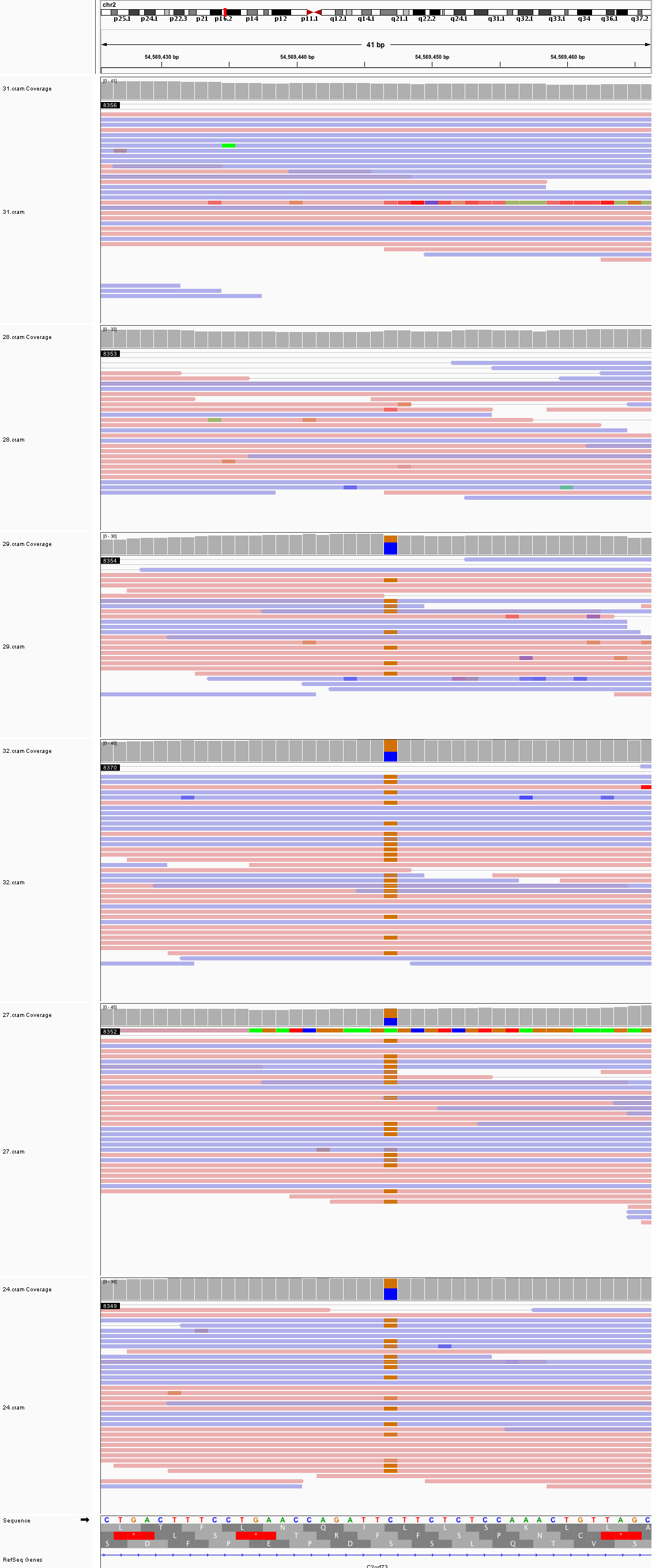

### chr2_55,103,085_55,103,125.png

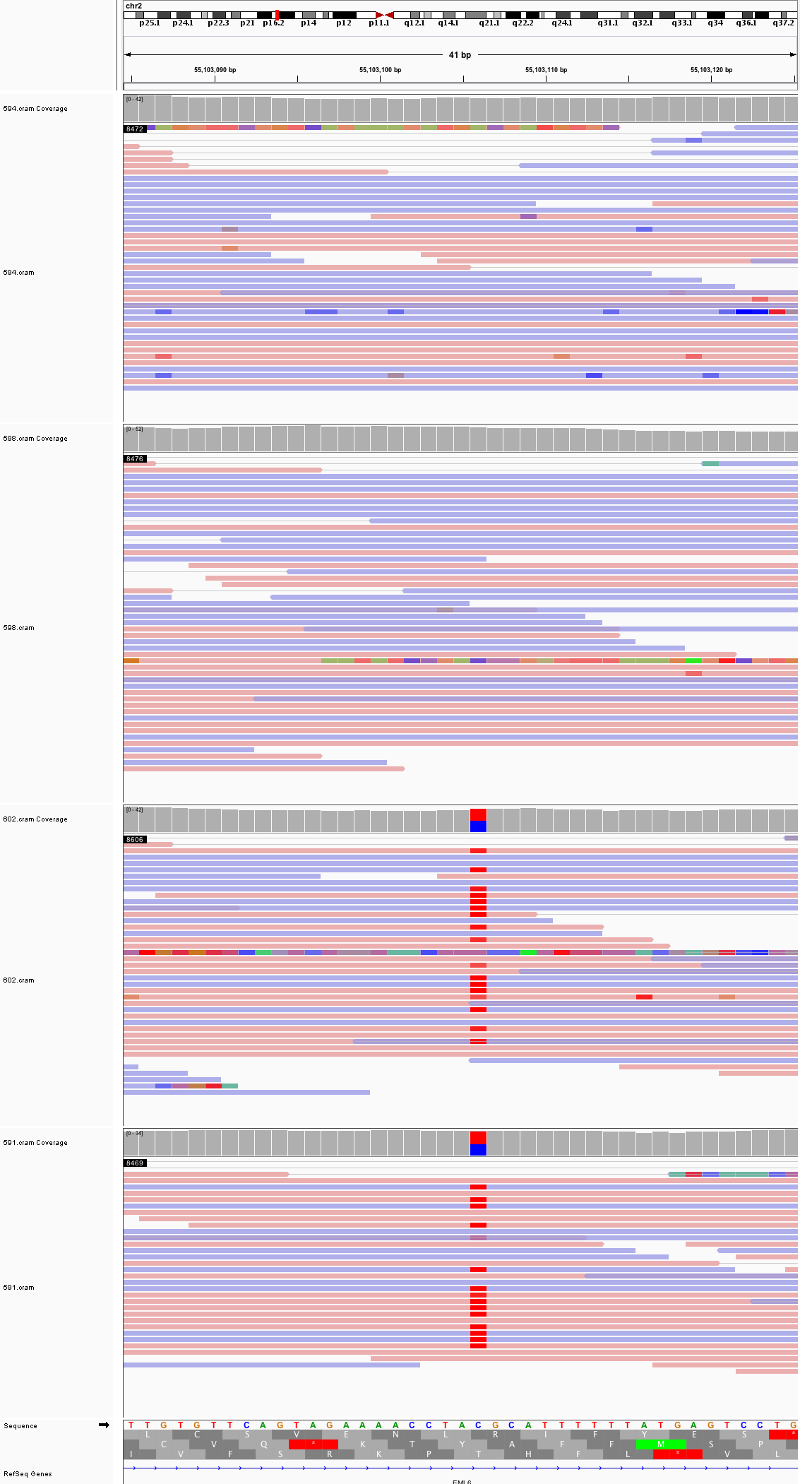

### chr2_63,679,582_63,679,622.png

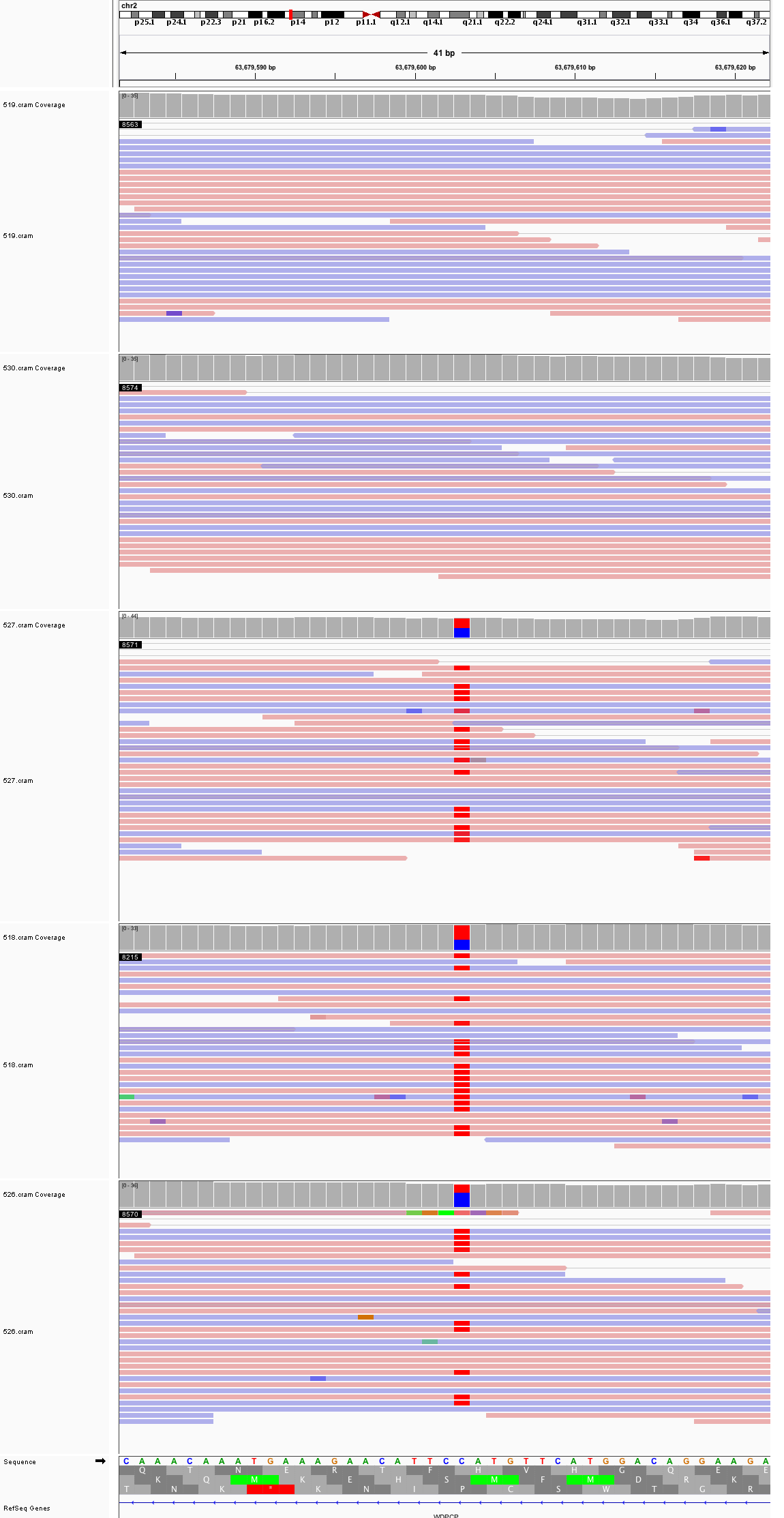

### chr2_75,845,385_75,845,425.png

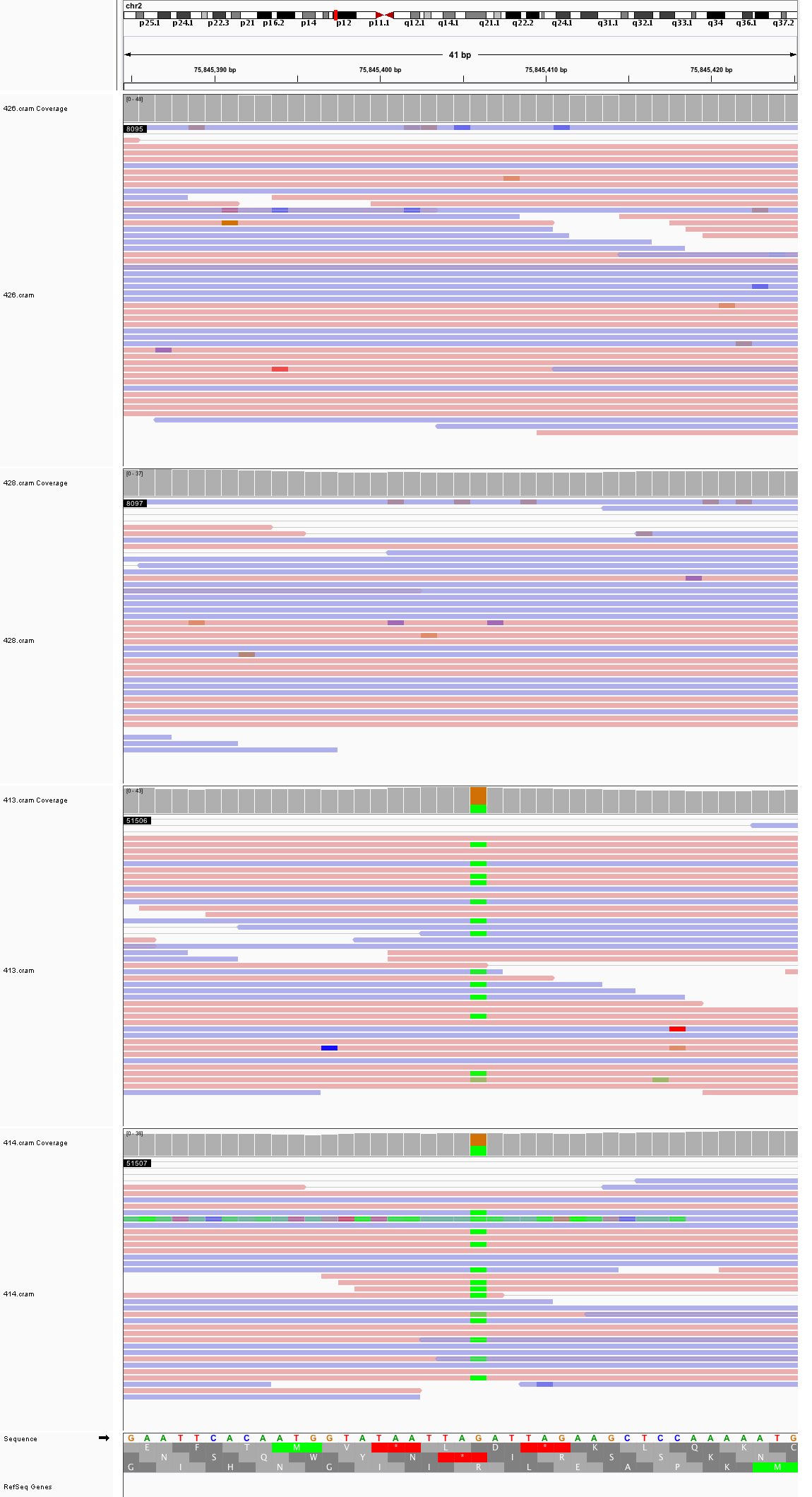

### chr2_118,523,975_118,524,015.png

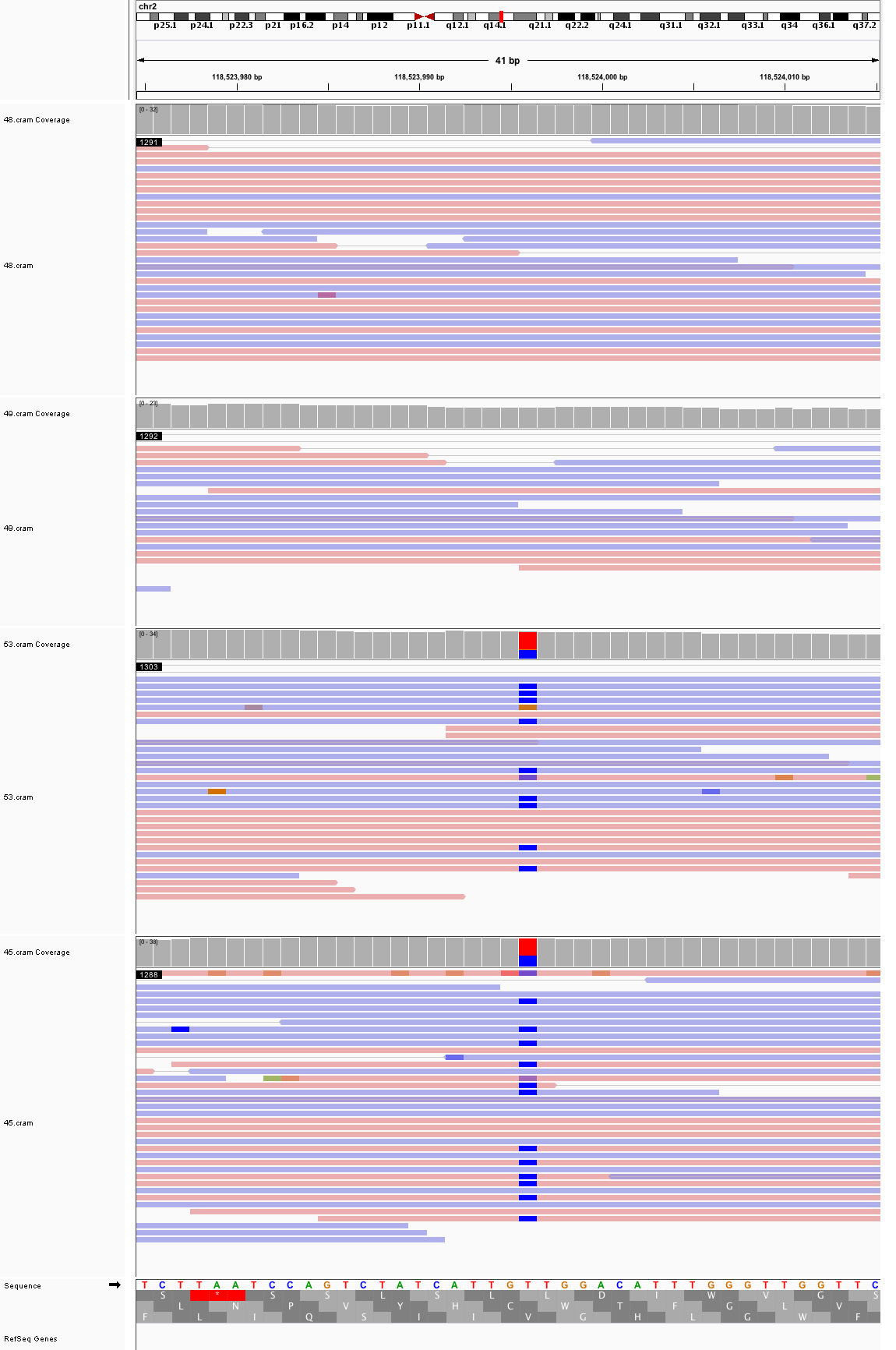

### chr2_141,781,343_141,781,383.png

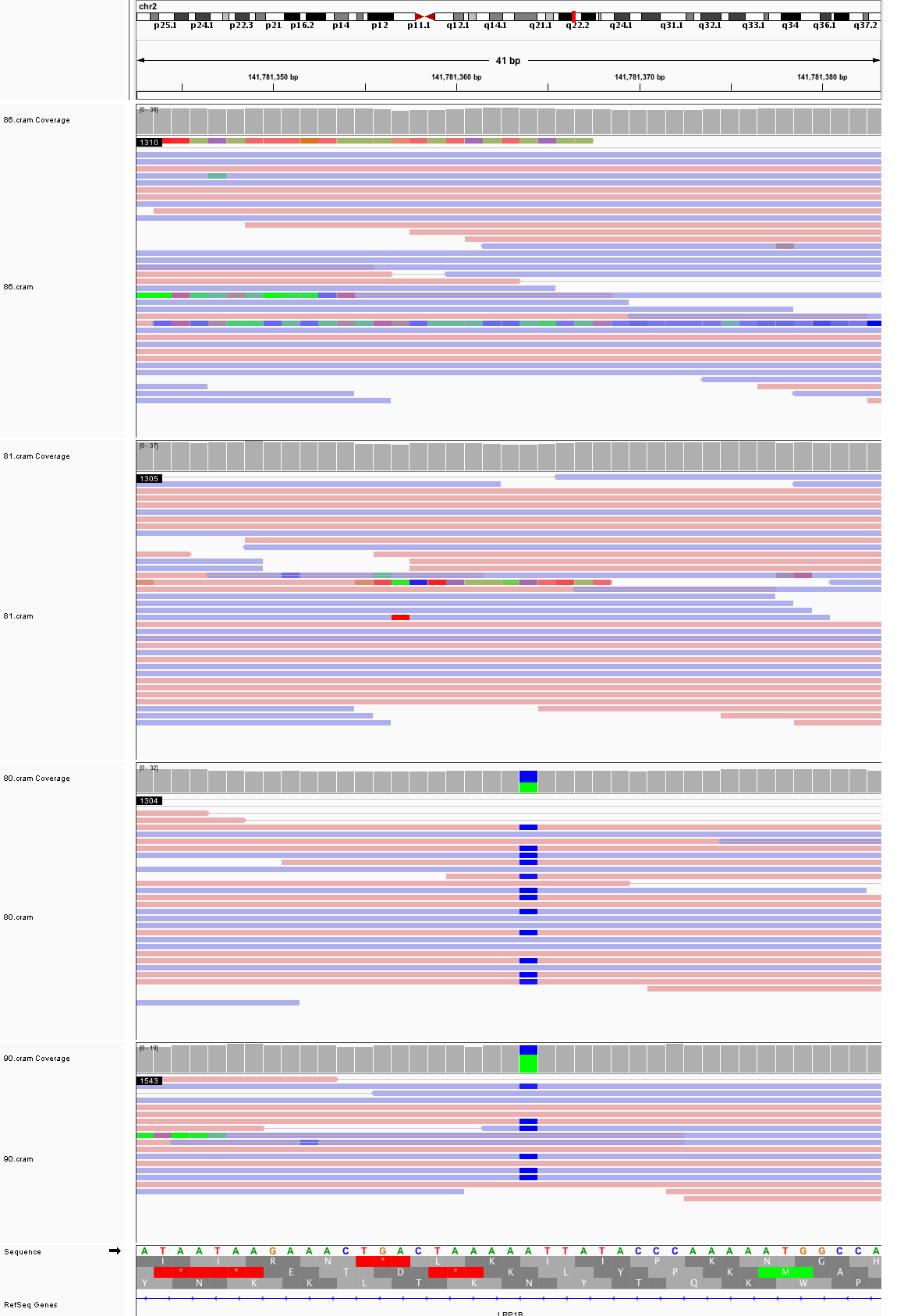

### chr2_150,625,598_150,625,638.png

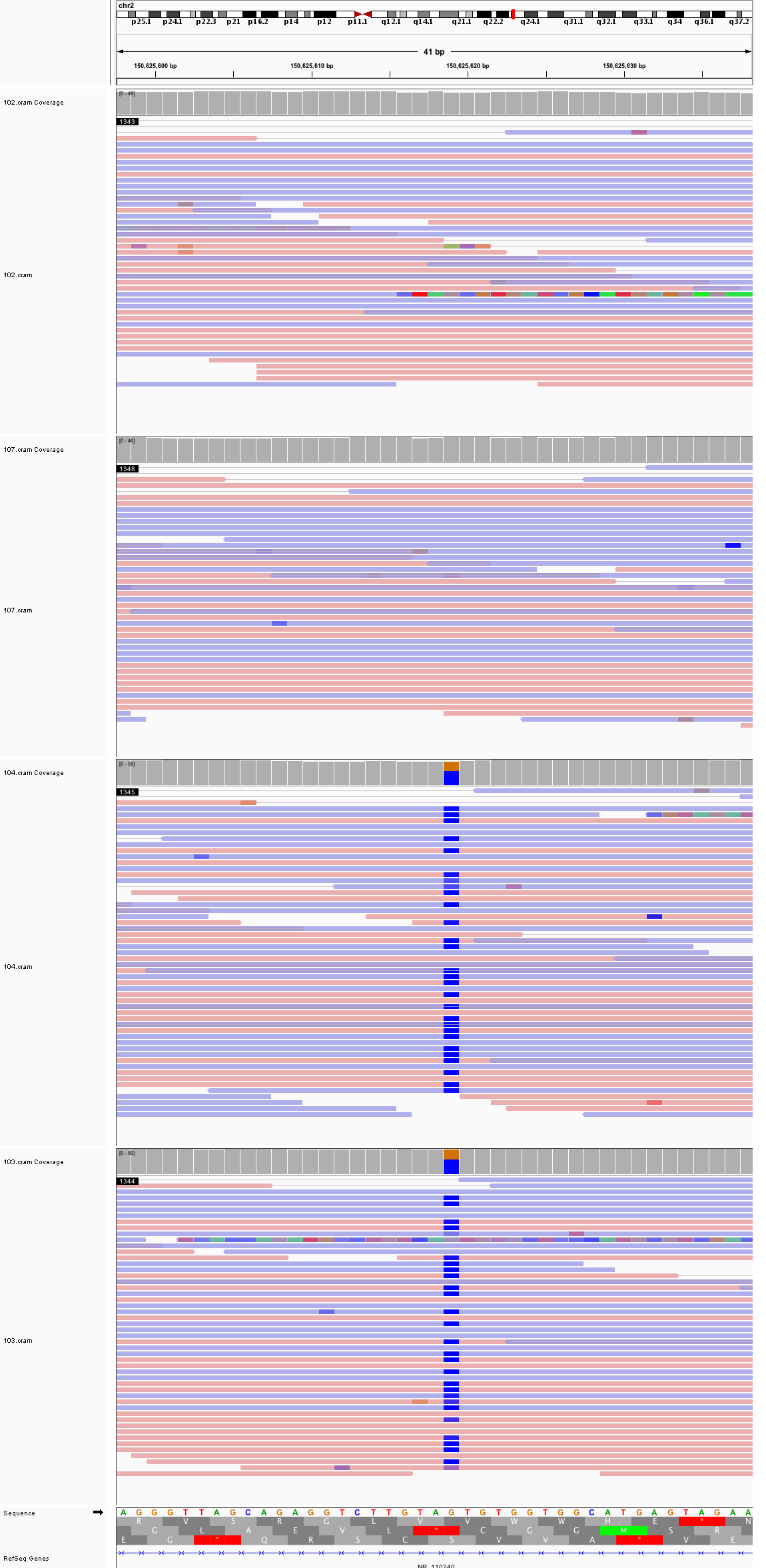

### chr2_156,822,516_156,822,556.png

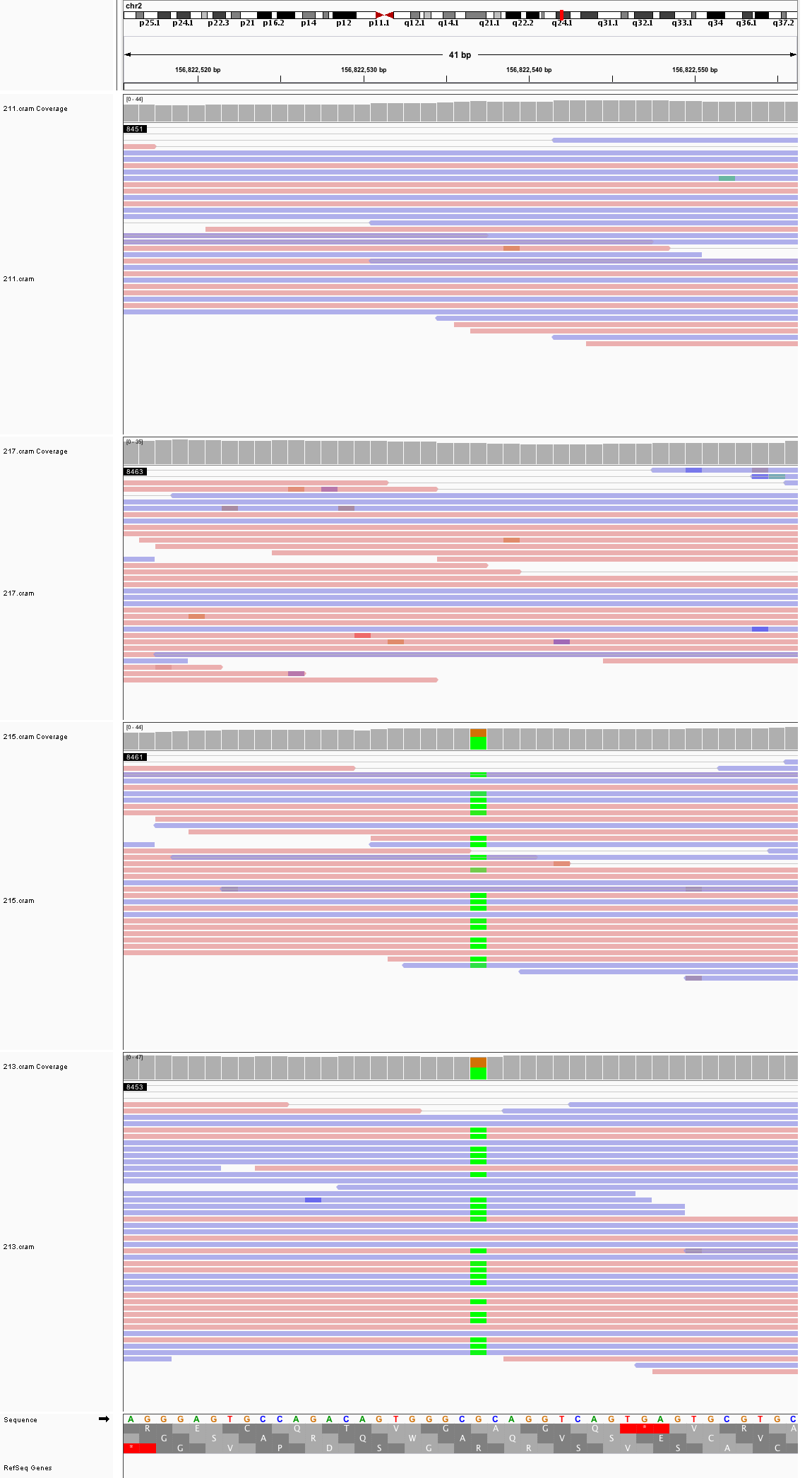

### chr2_159,742,525_159,742,565.png

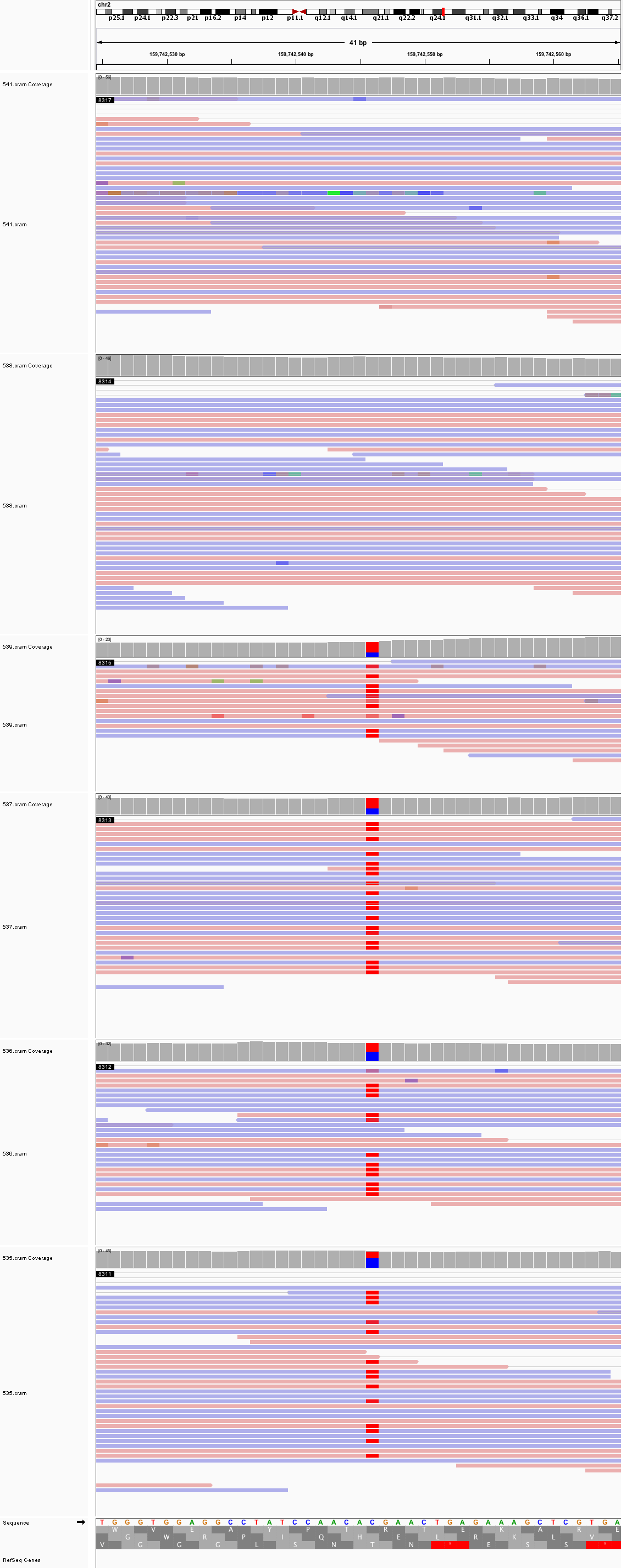

### chr2_163,874,558_163,874,598.png

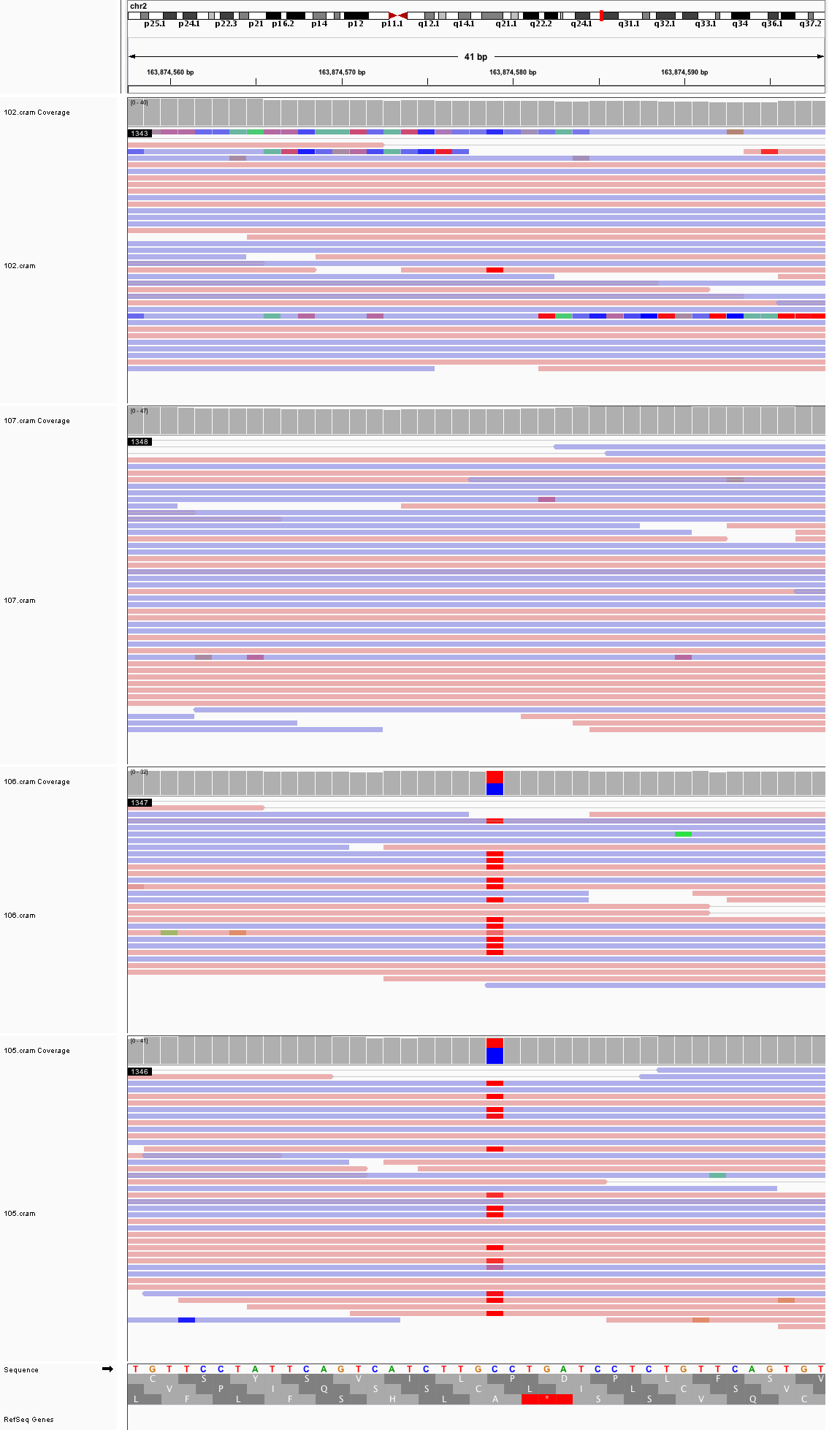

### chr2_165,977,707_165,977,747.png

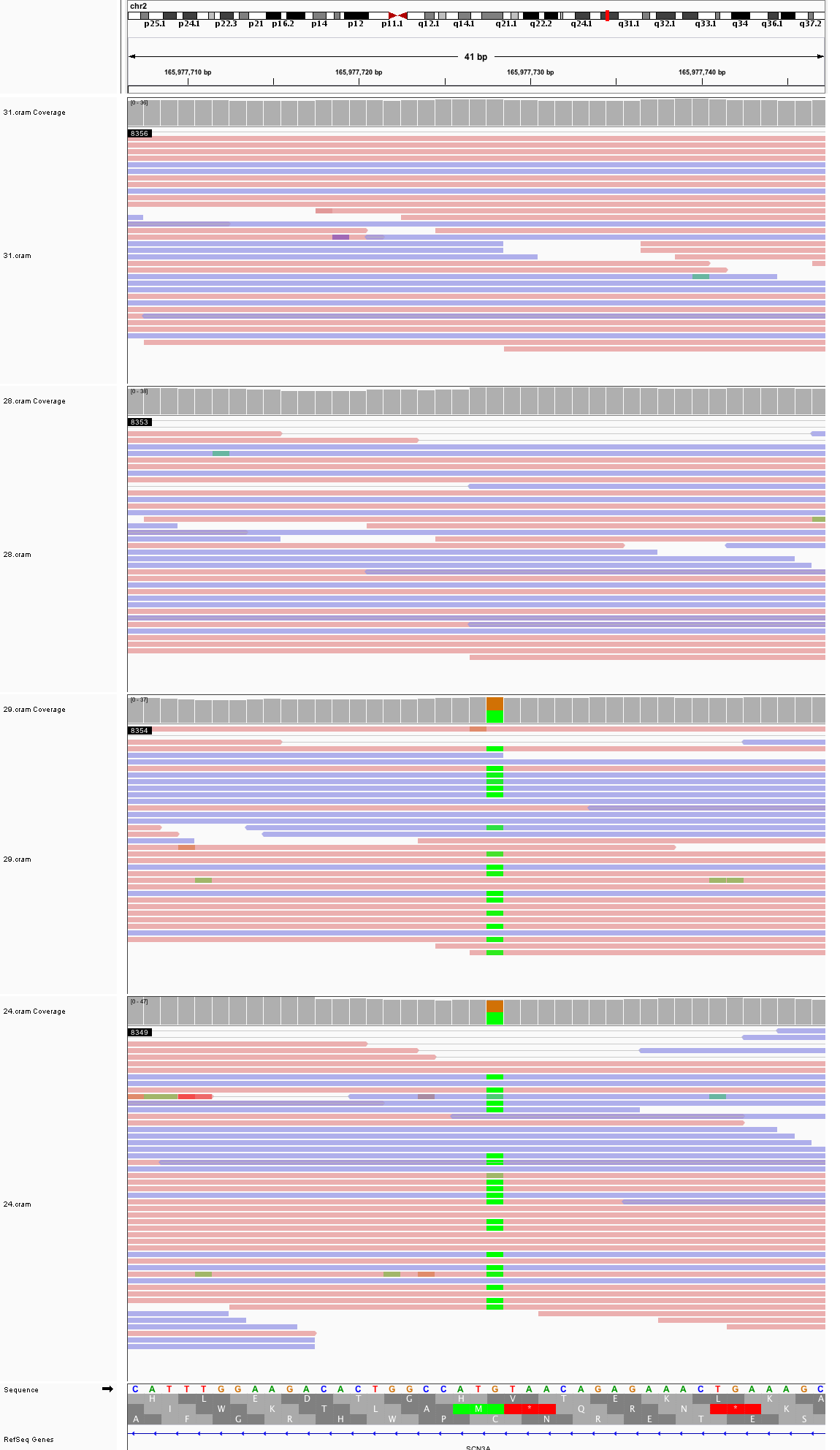

### chr2_167,500,553_167,500,593.png

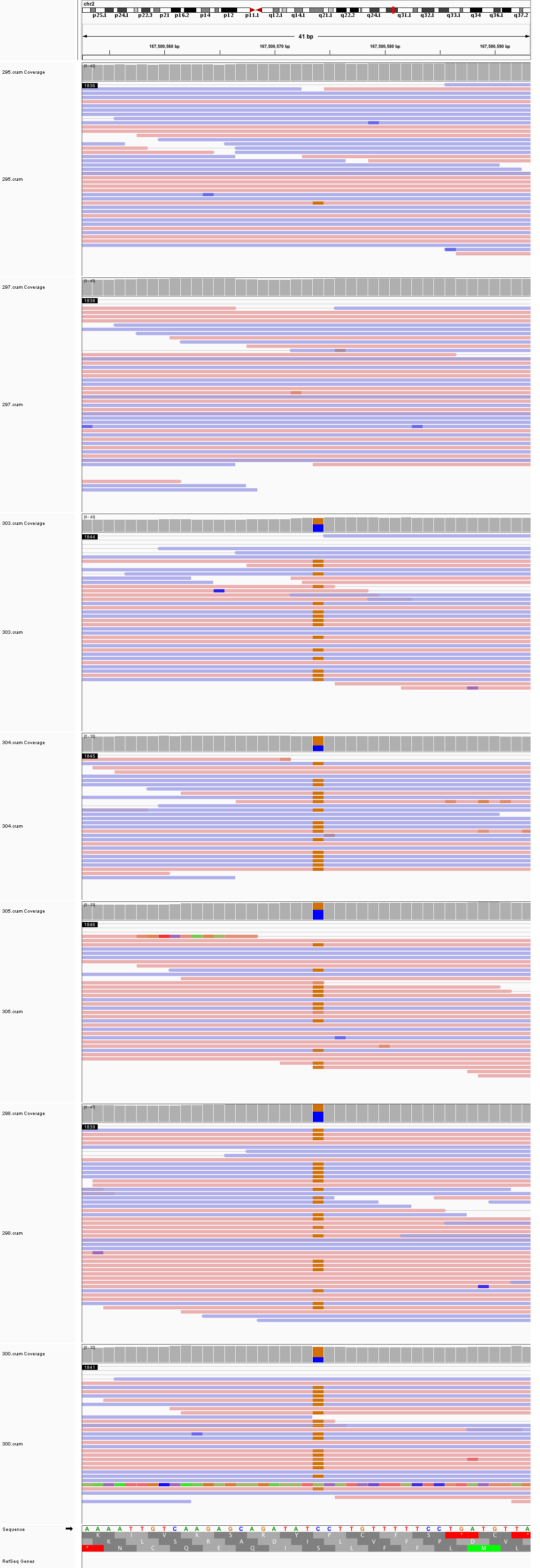

### chr2_199,388,393_199,388,433.png

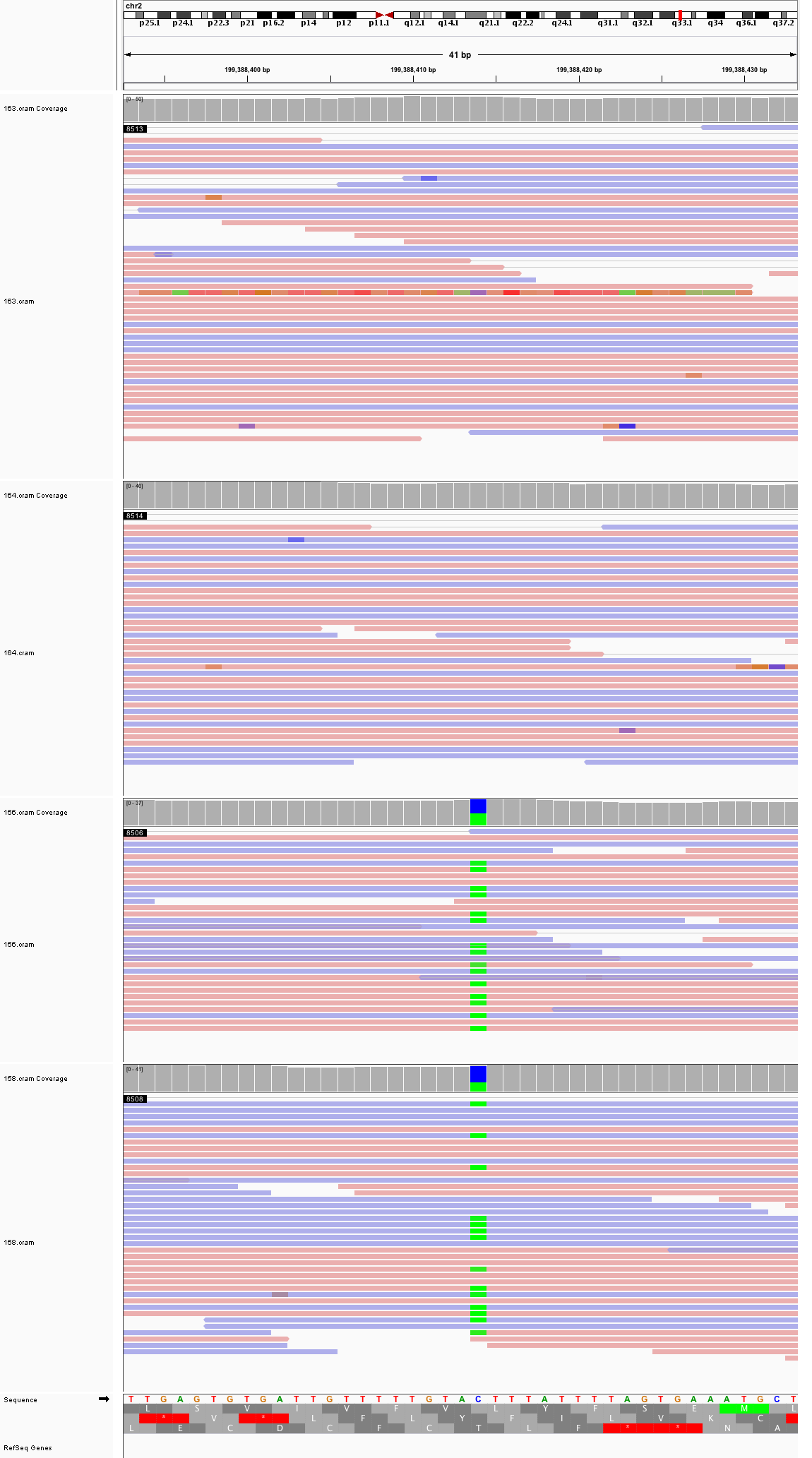

### chr2_225,962,047_225,962,087.png

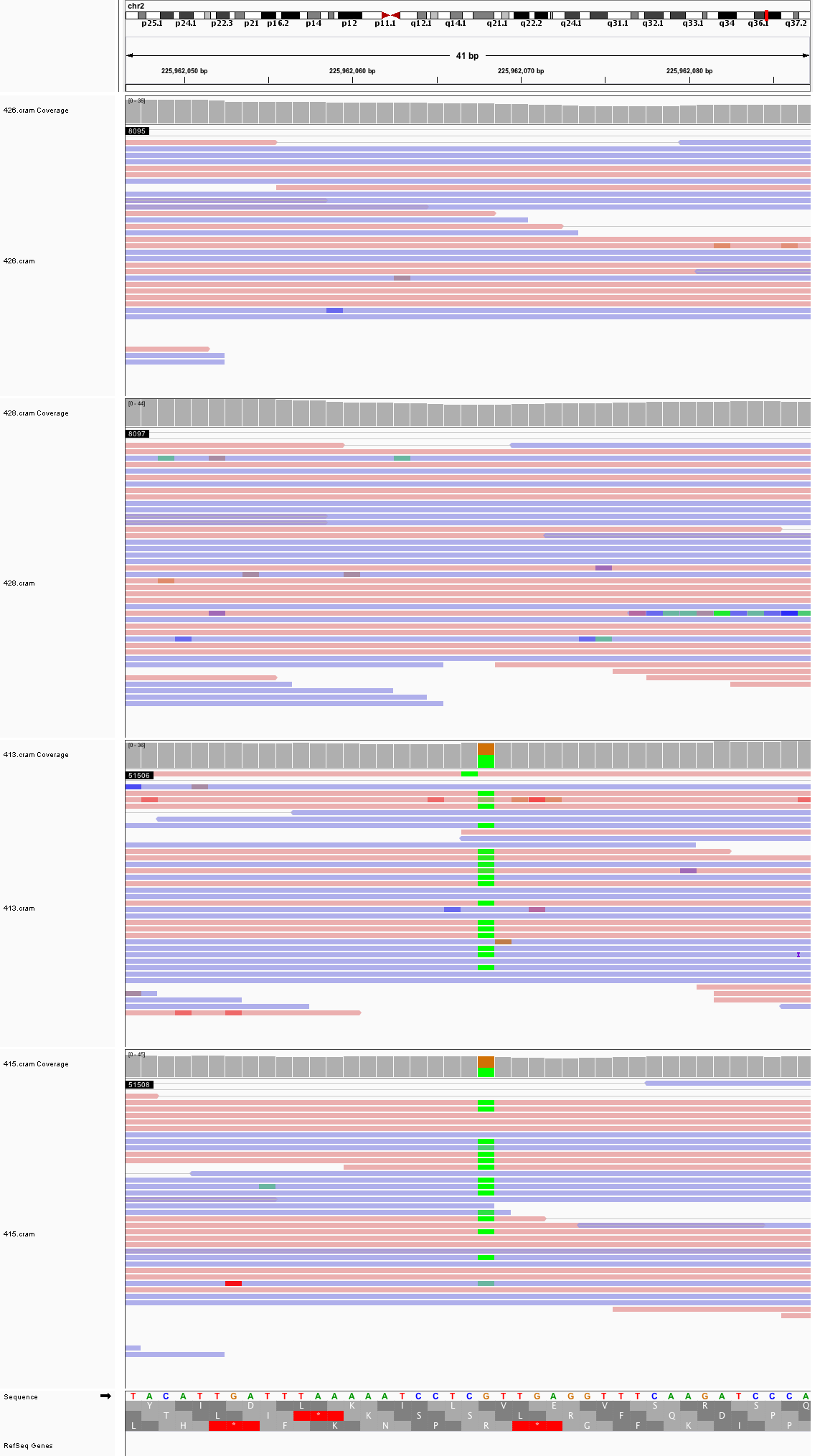

### chr2_239,476,521_239,476,561.png

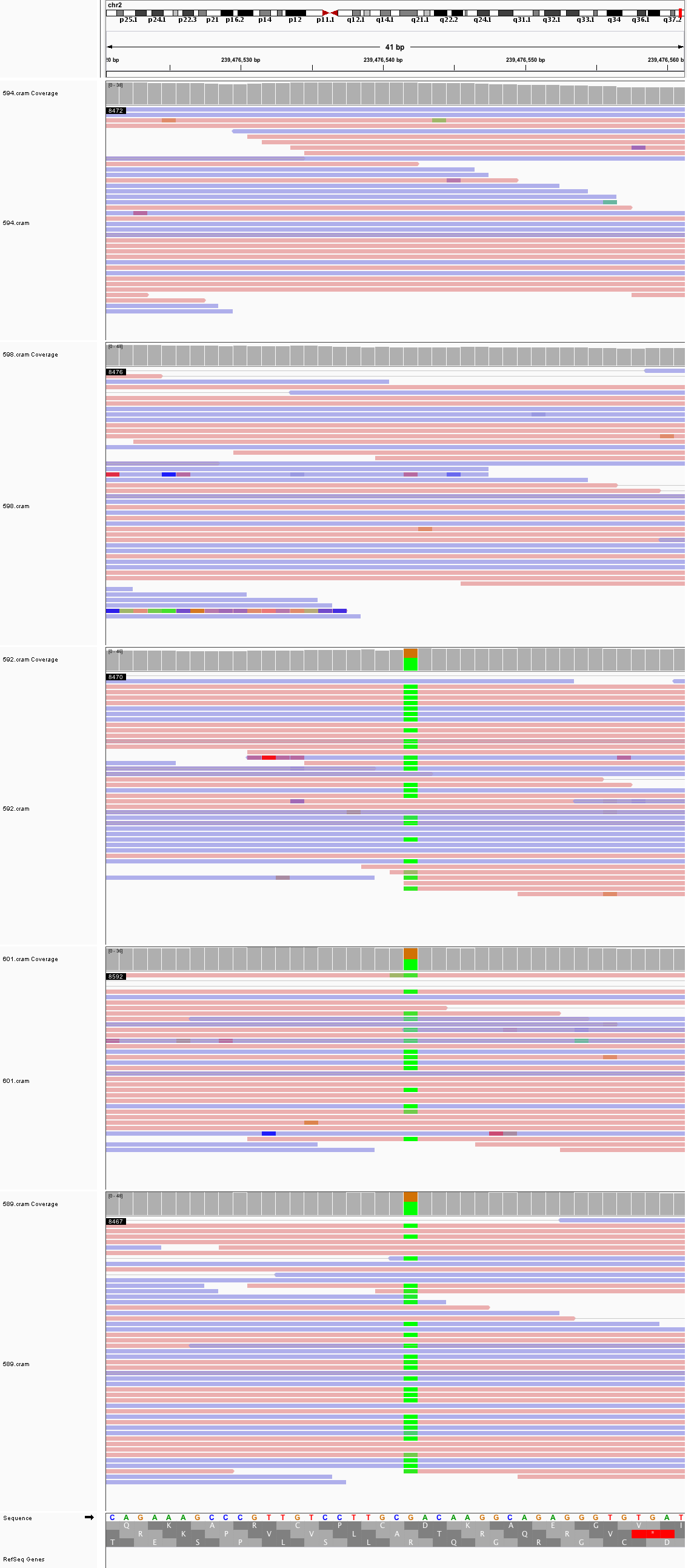

### chr2_242,164,047_242,164,087.png

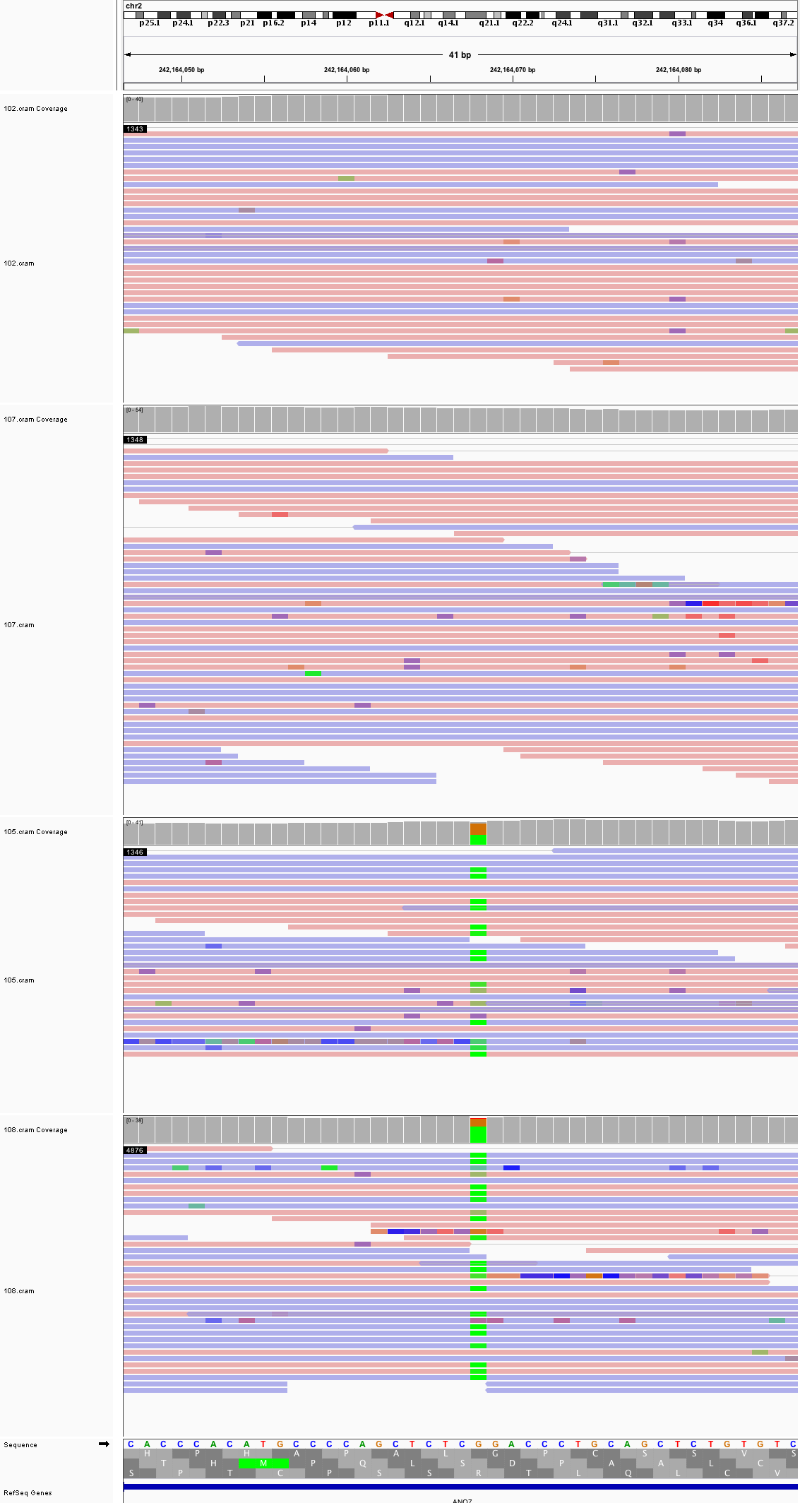

### chr3_2,268,991_2,269,031.png

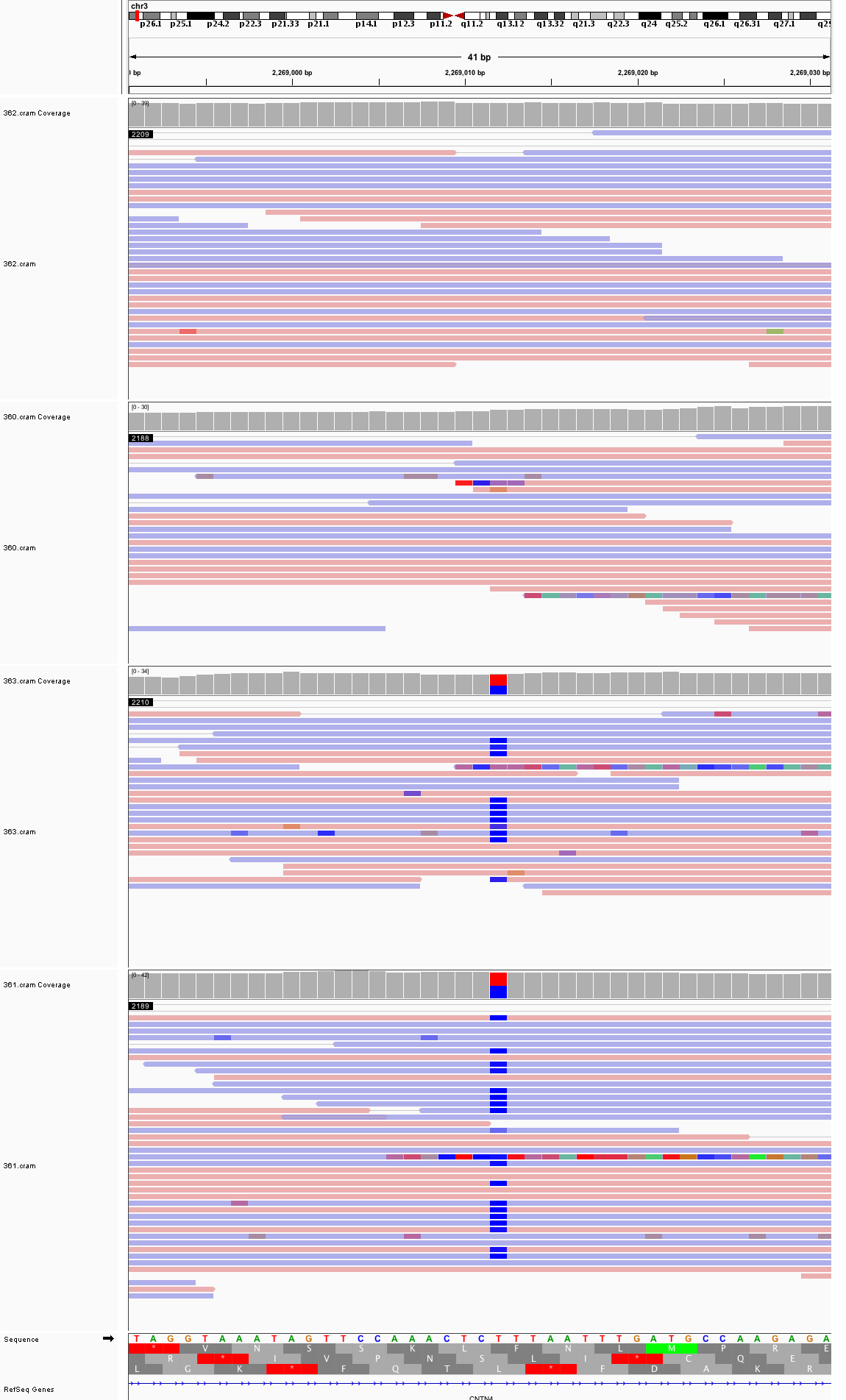

### chr3_119,566,351_119,566,391.png

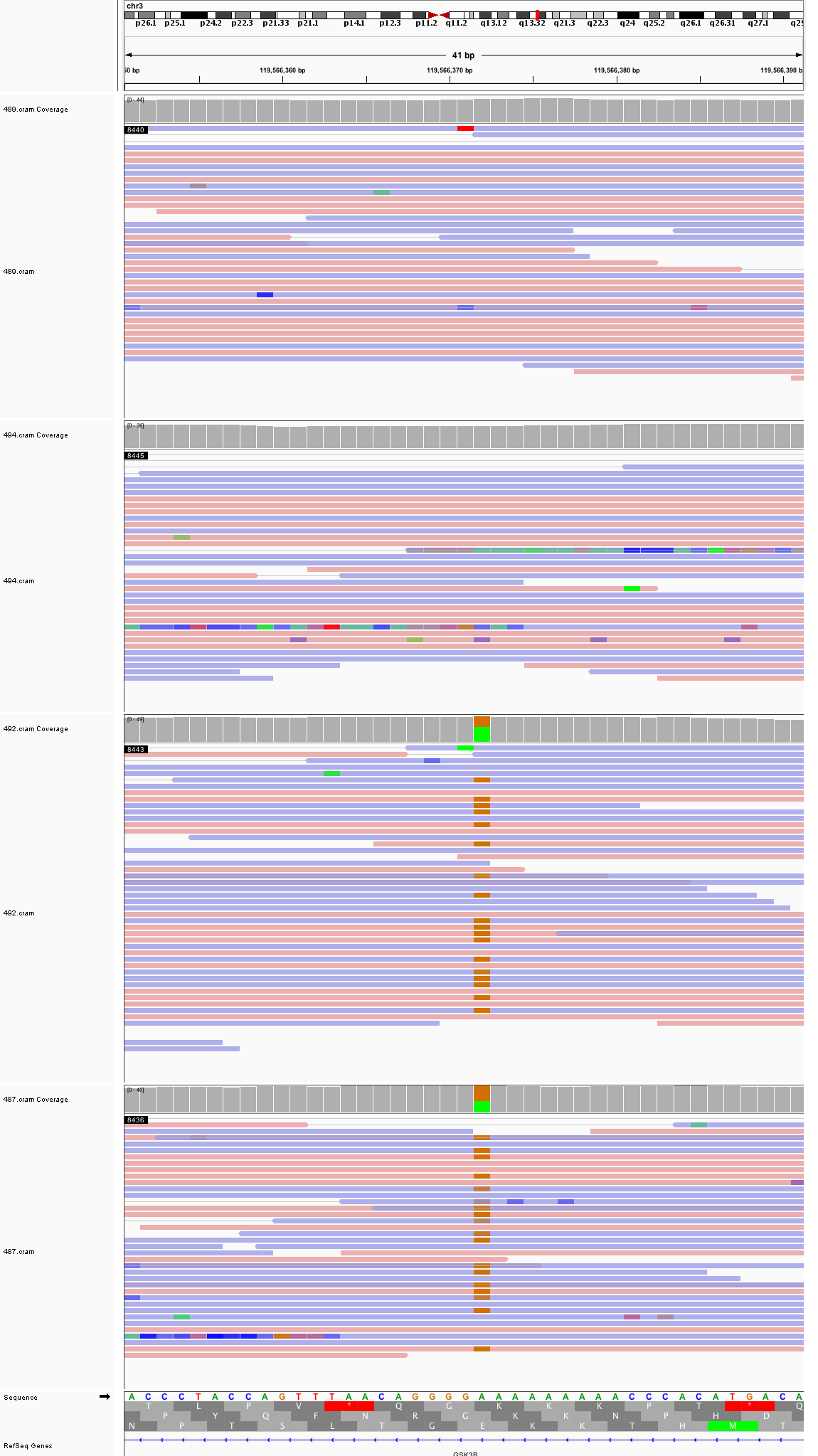

### chr3_126,374,277_126,374,317.png

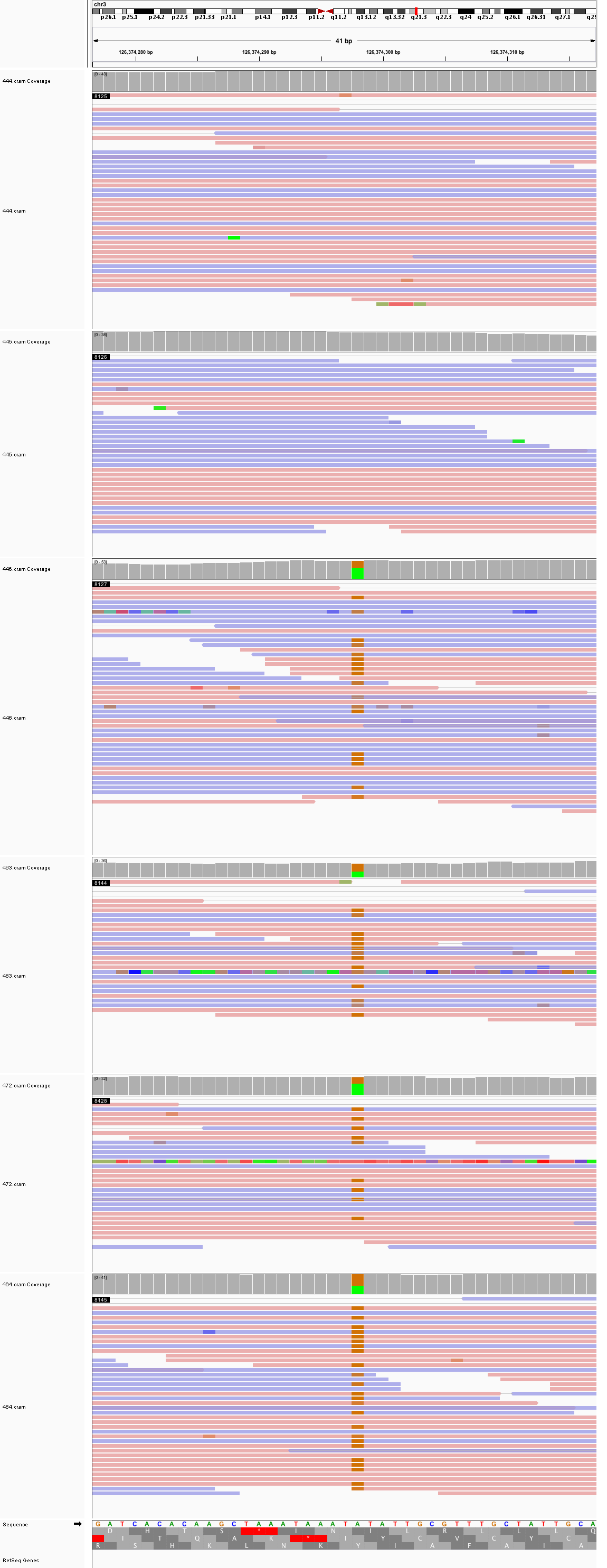

### chr3_173,123,973_173,124,013.png

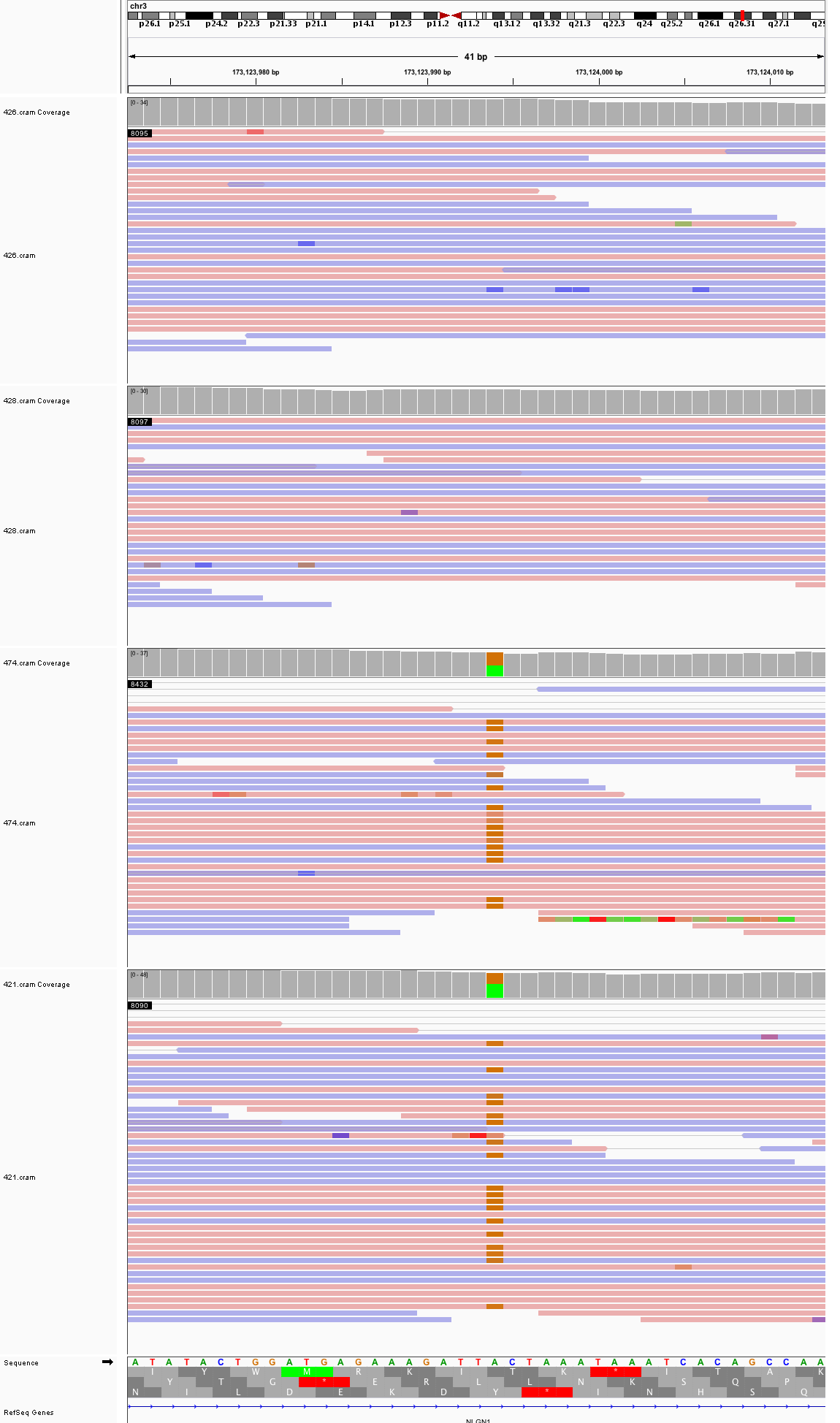

### chr3_185,133,111_185,133,151.png

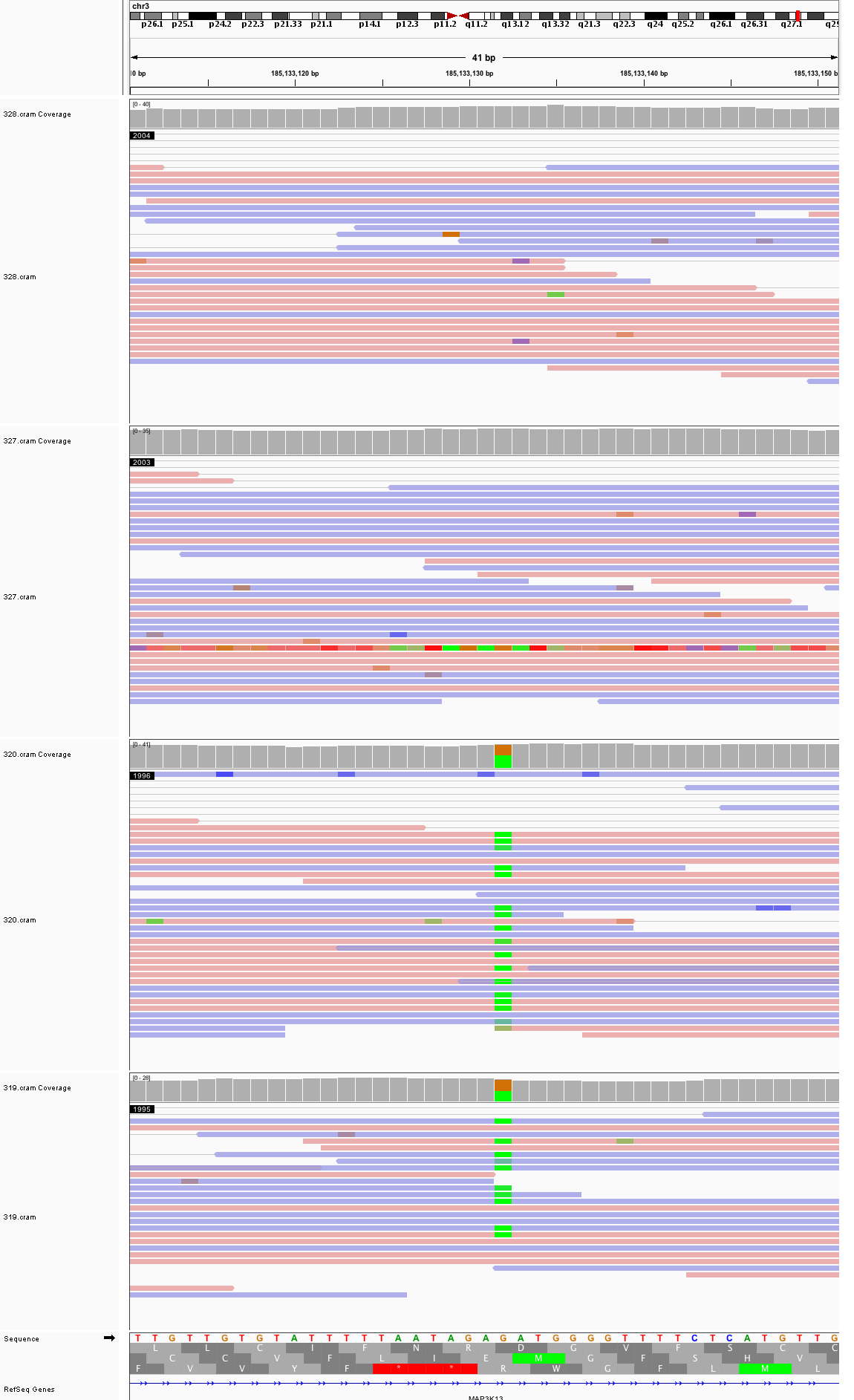

### chr4_14,636,632_14,636,672.png

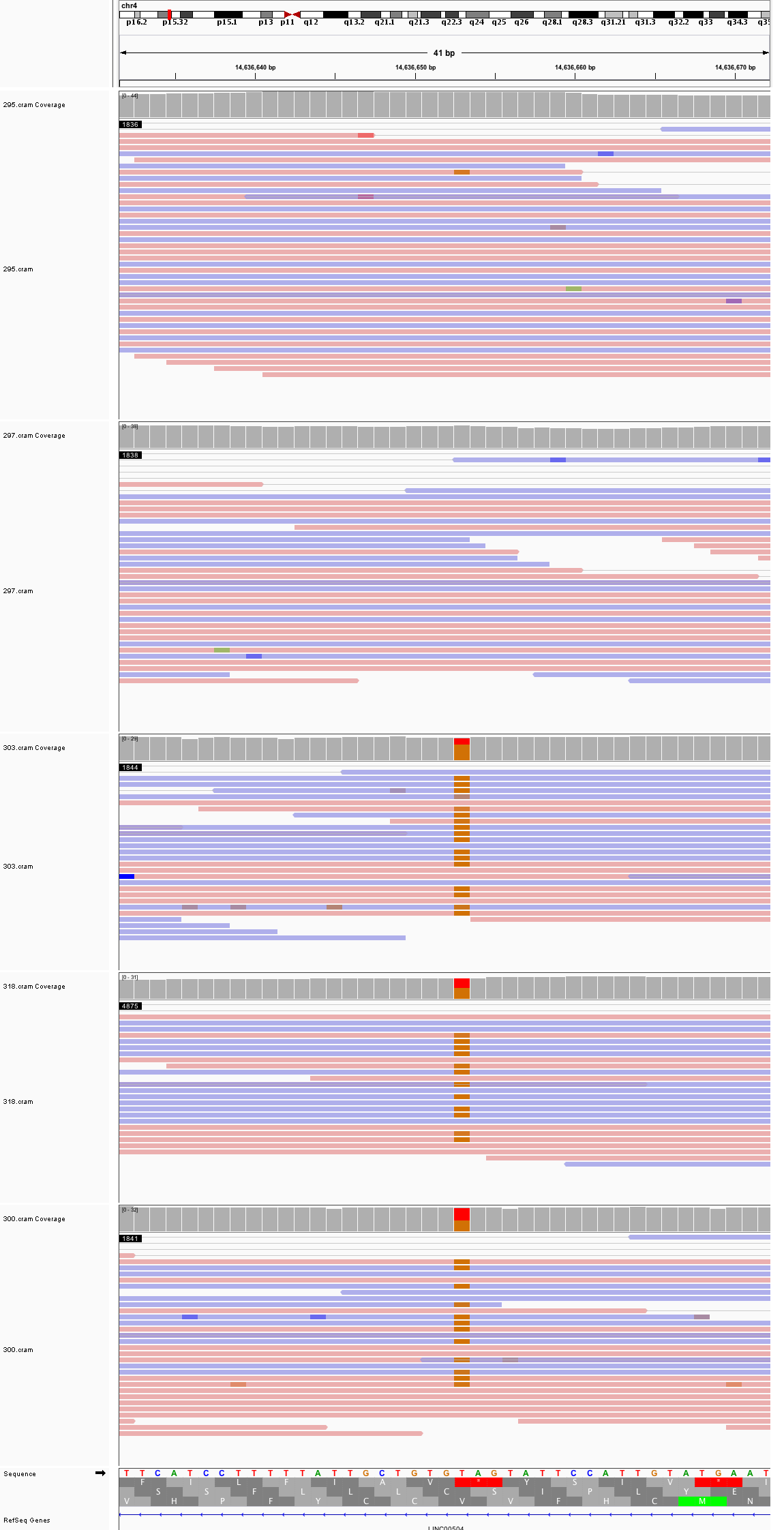

### chr4_39,208,044_39,208,084.png

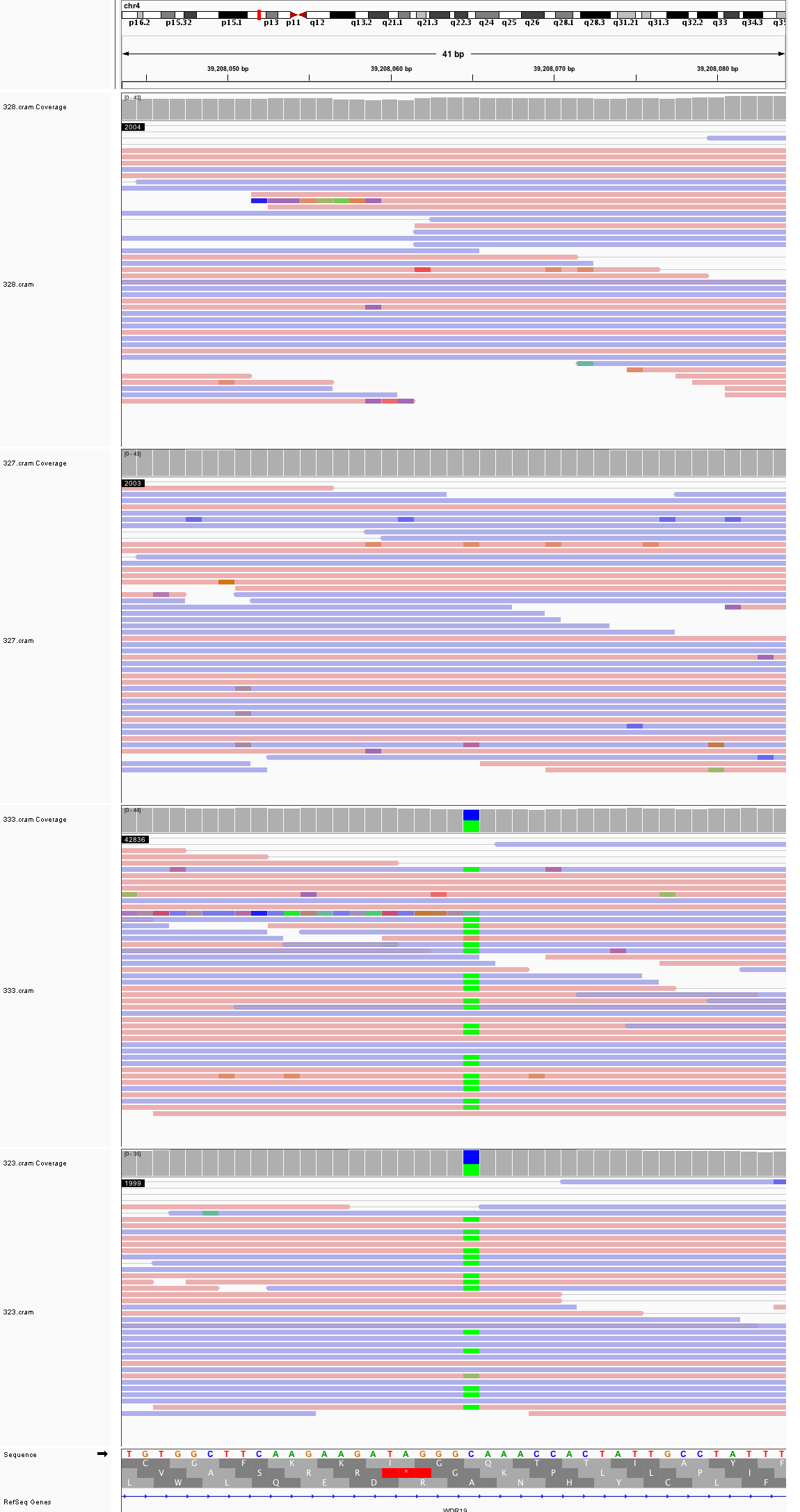

### chr4_63,480,961_63,481,001.png

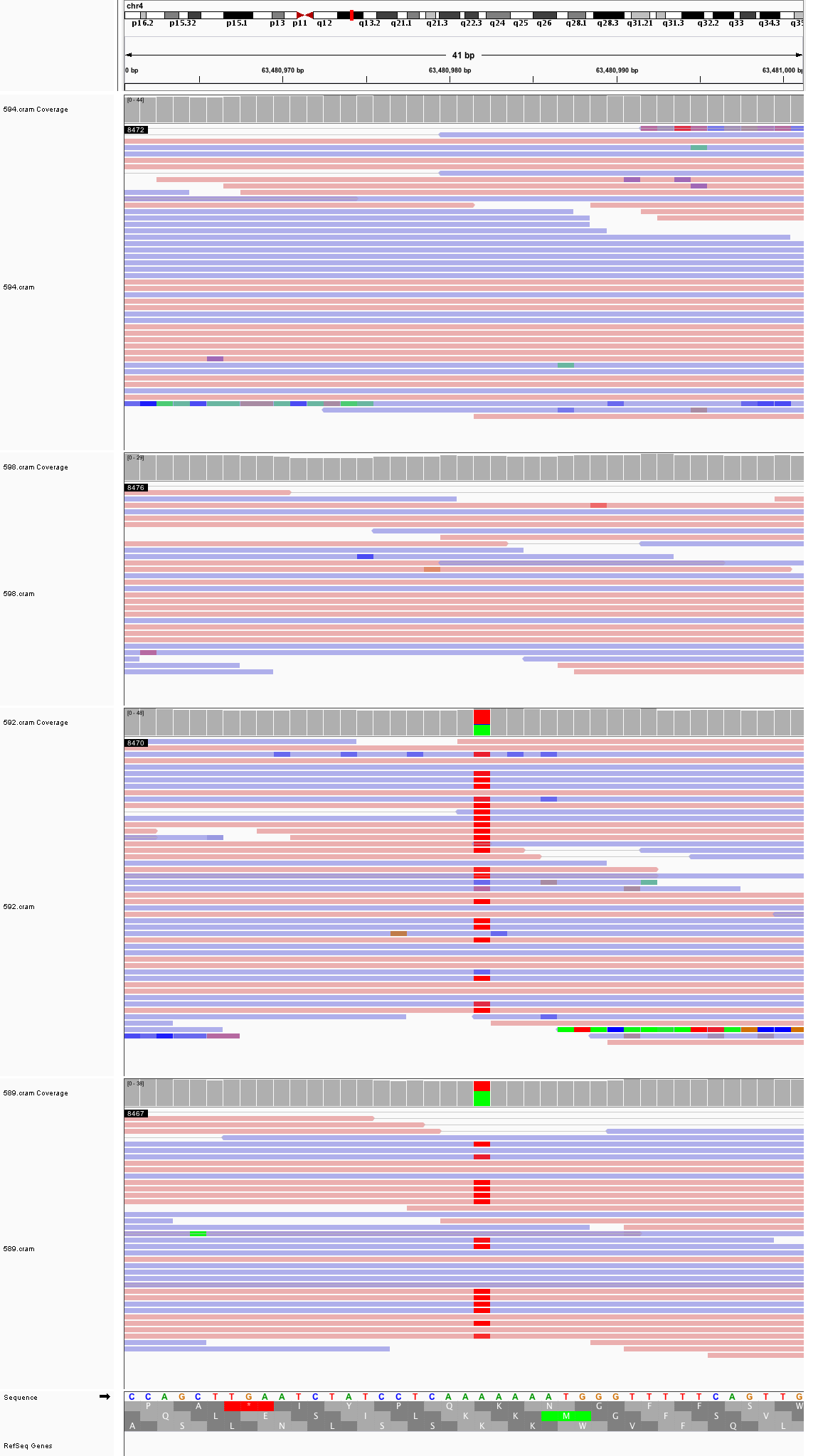

### chr4_71,378,123_71,378,163.png

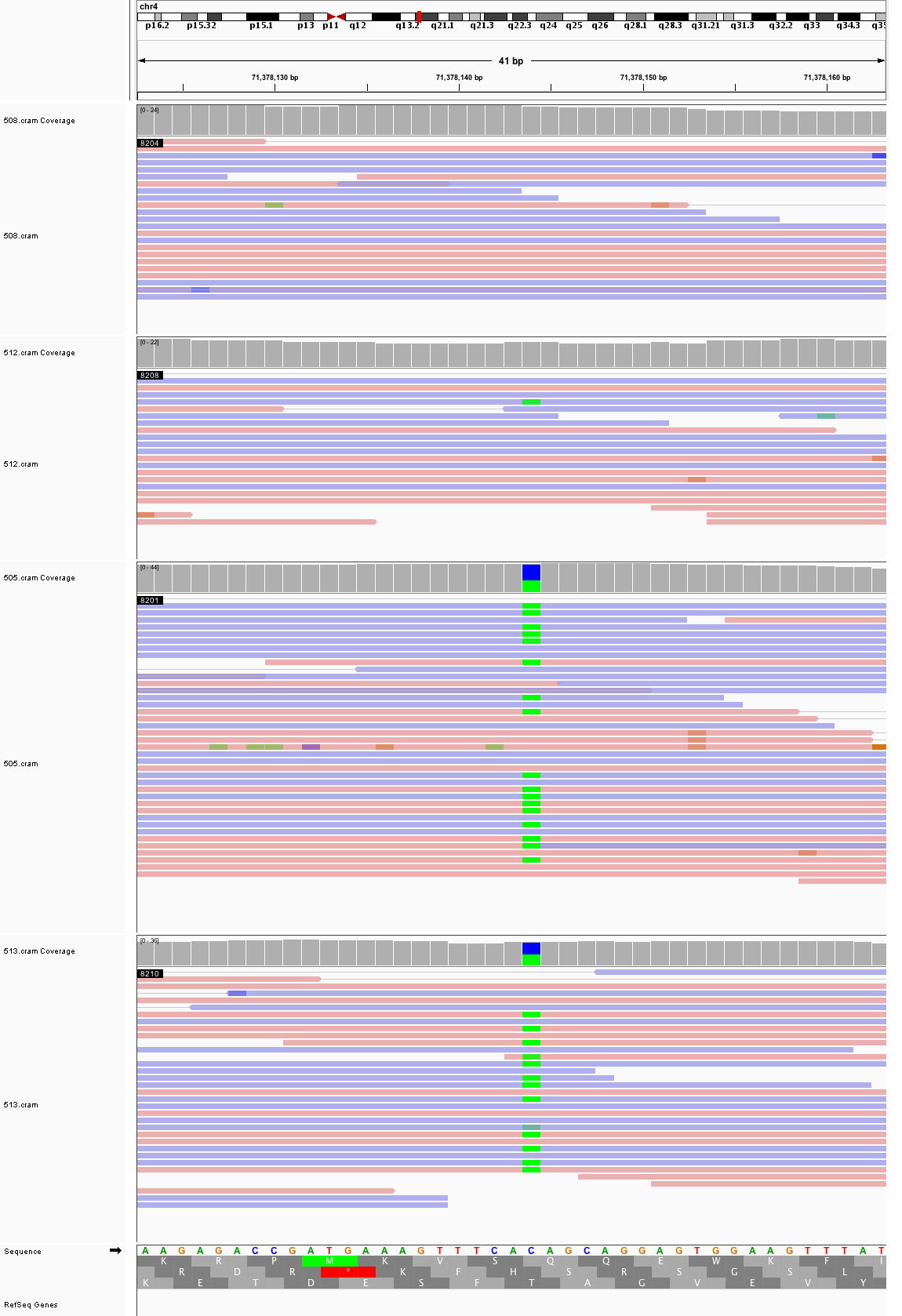

### chr4_106,654,918_106,654,958.png

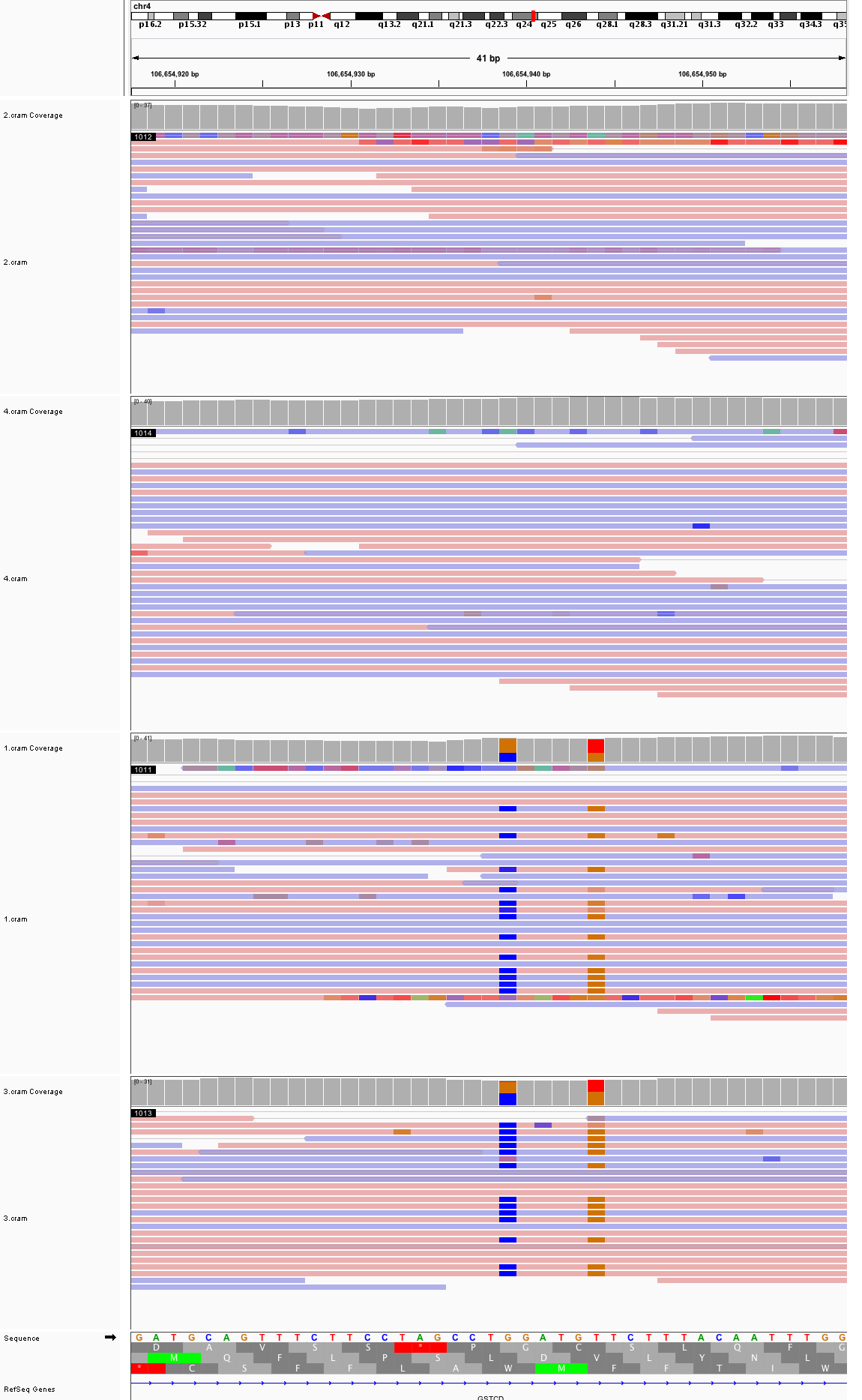

### chr4_139,430,385_139,430,425.png

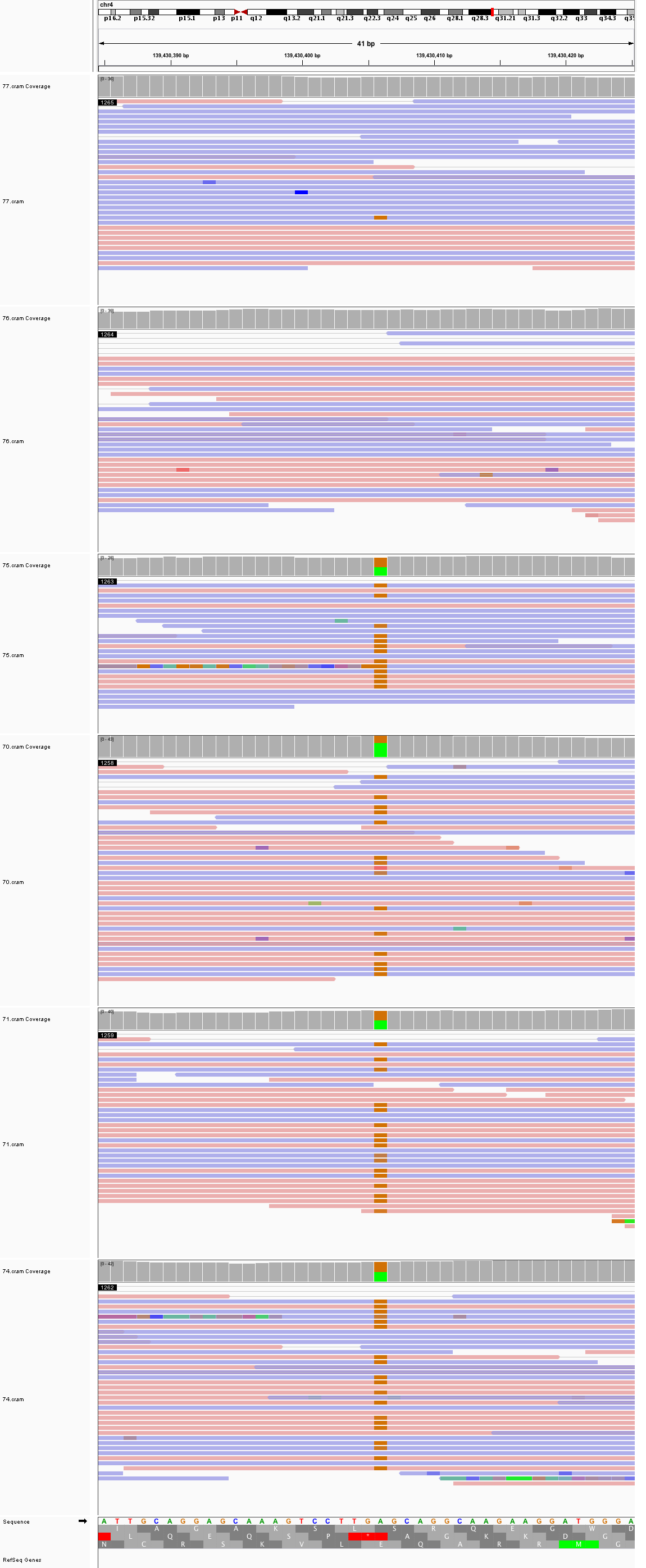

### chr4_150,465,787_150,465,827.png

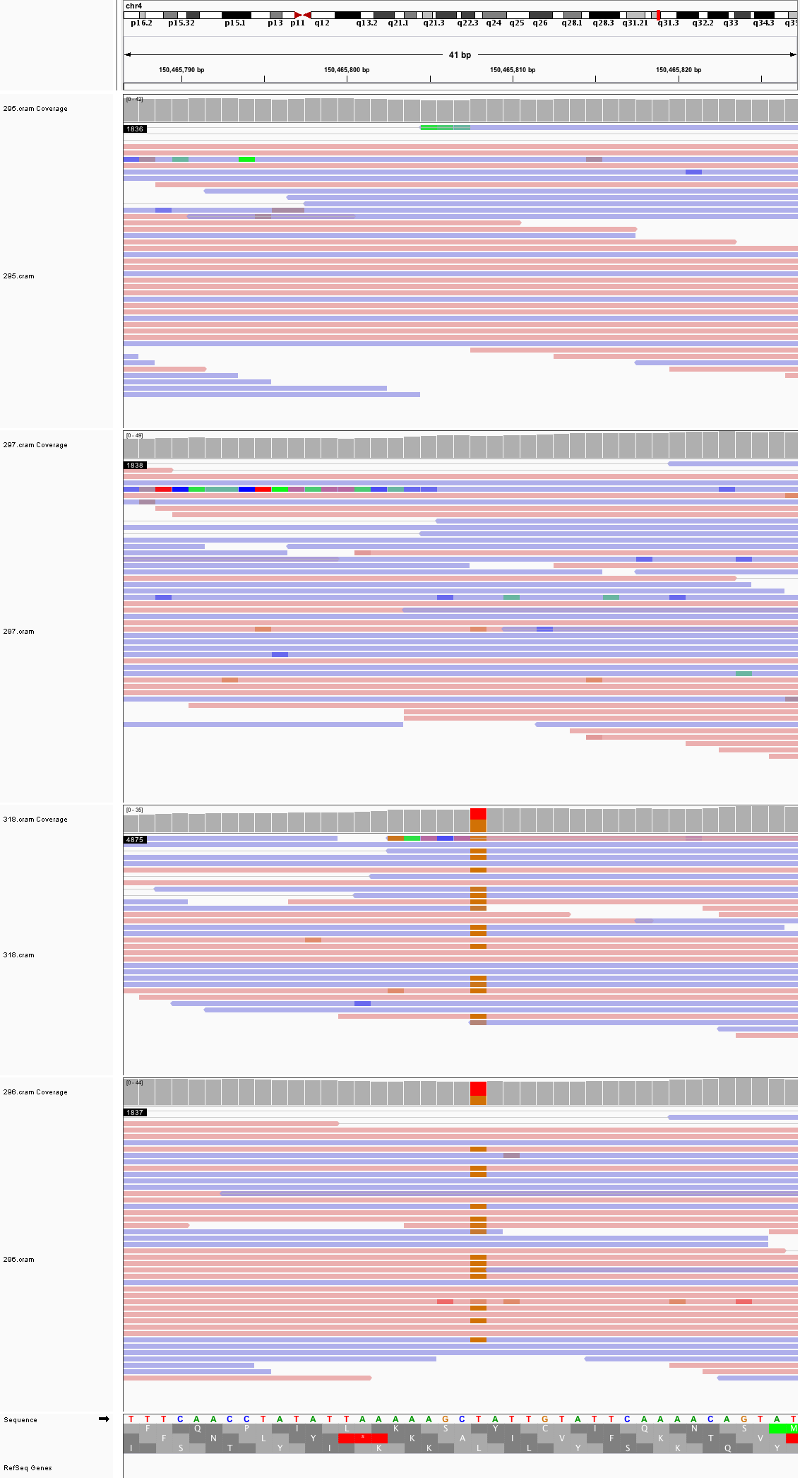

### chr4_167,426,284_167,426,324.png

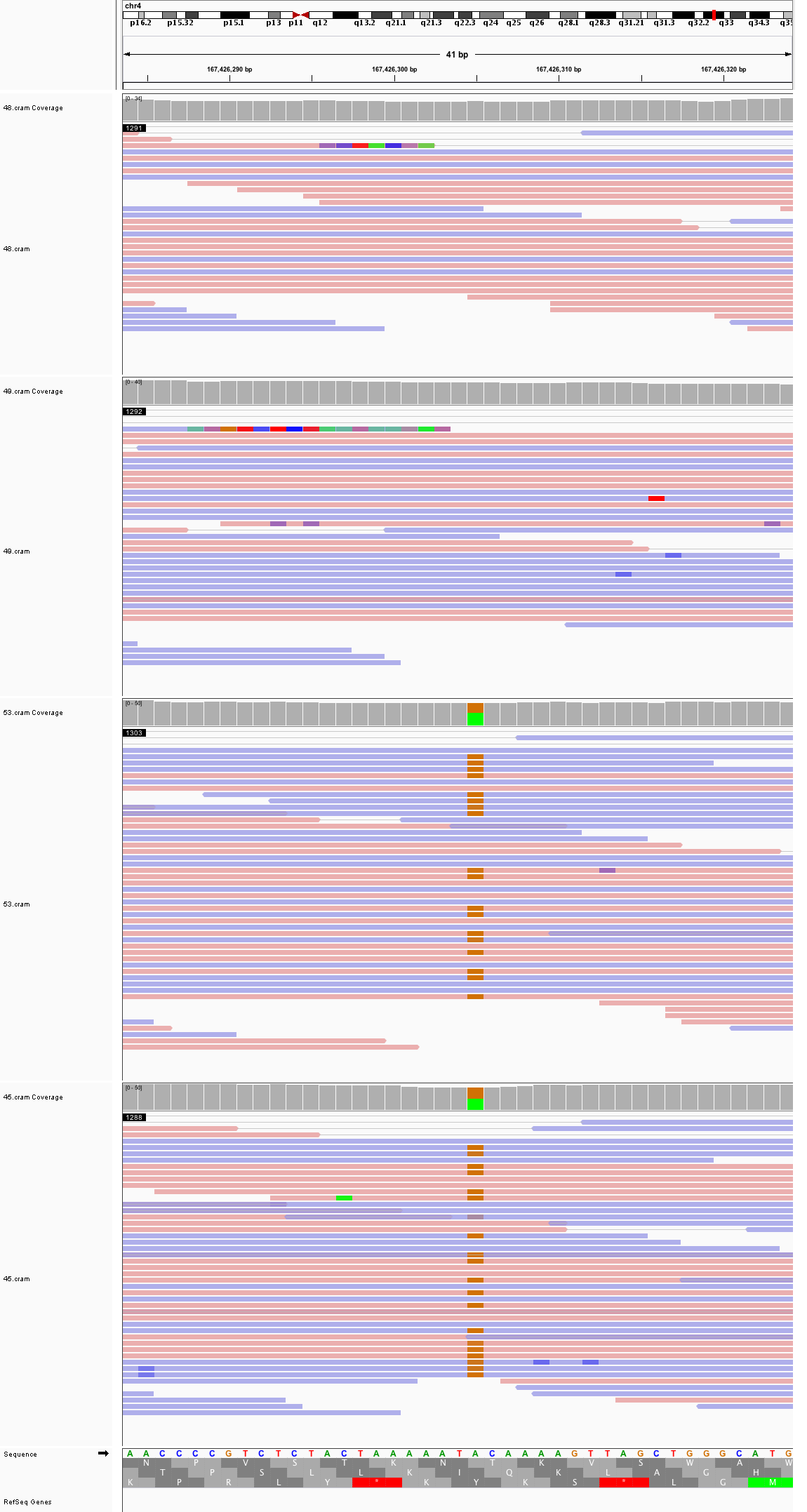

### chr5_2,359,958_2,359,998.png

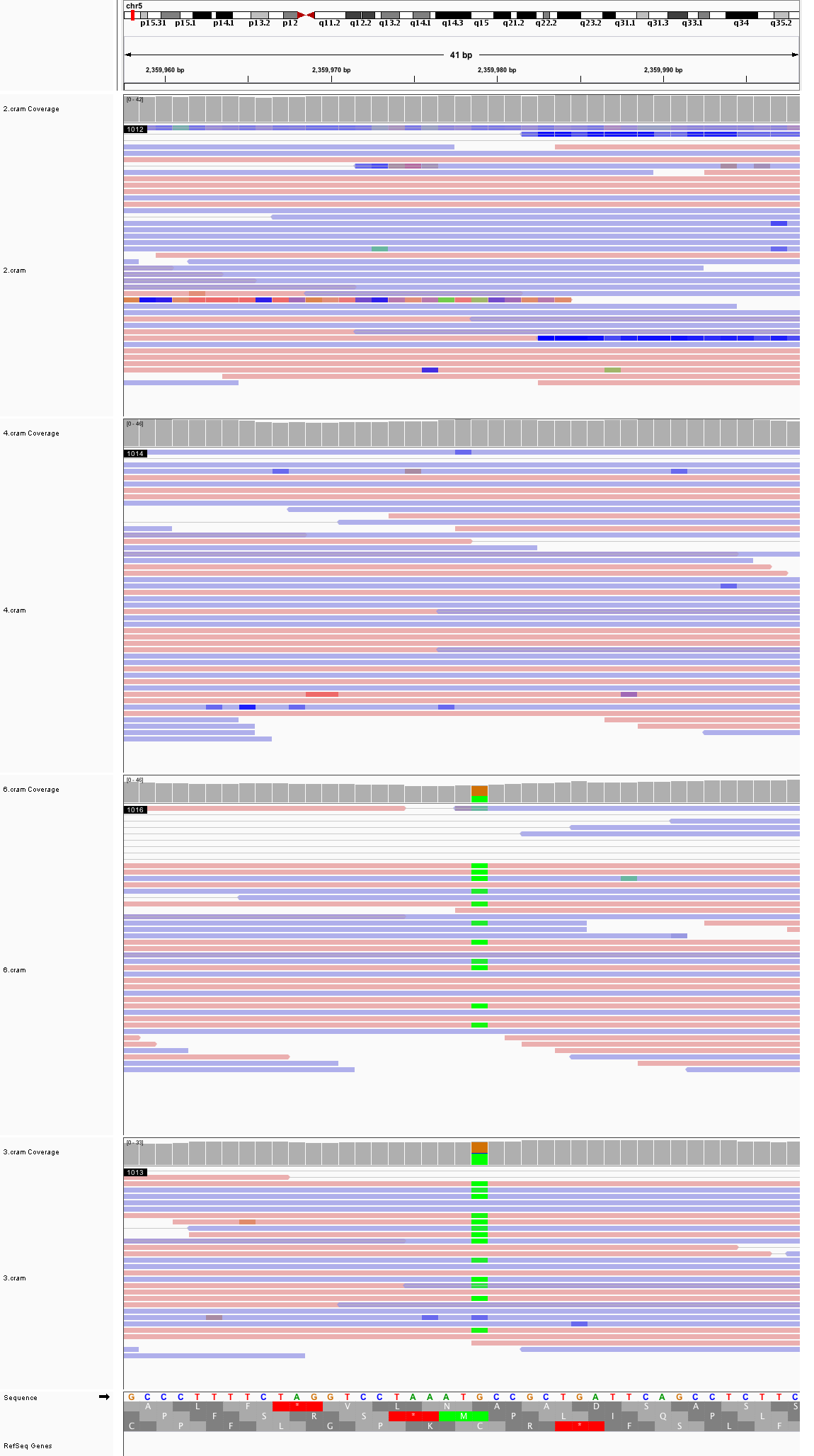

### chr5_76,707,962_76,708,002.png

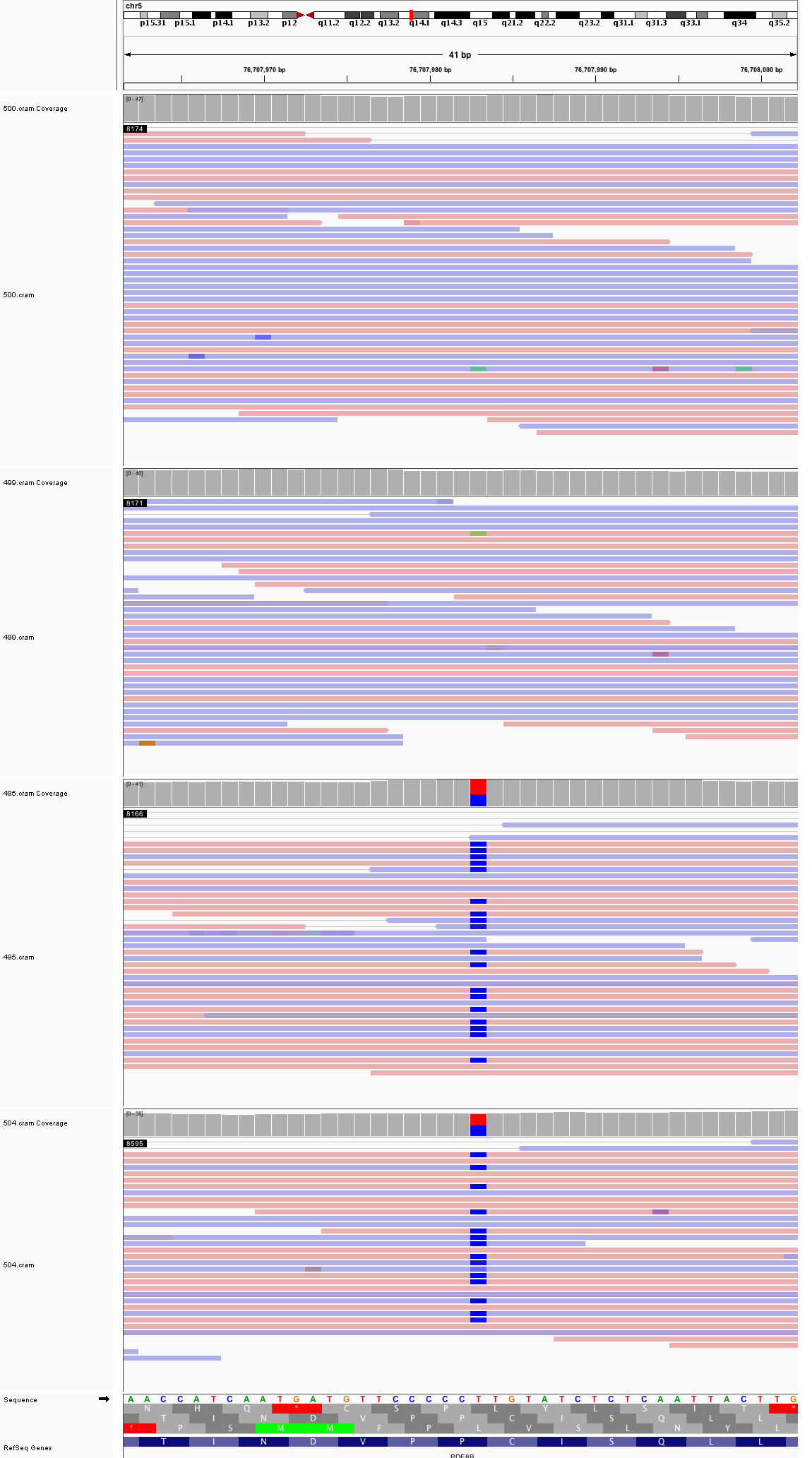

### chr5_83,112,543_83,112,583.png

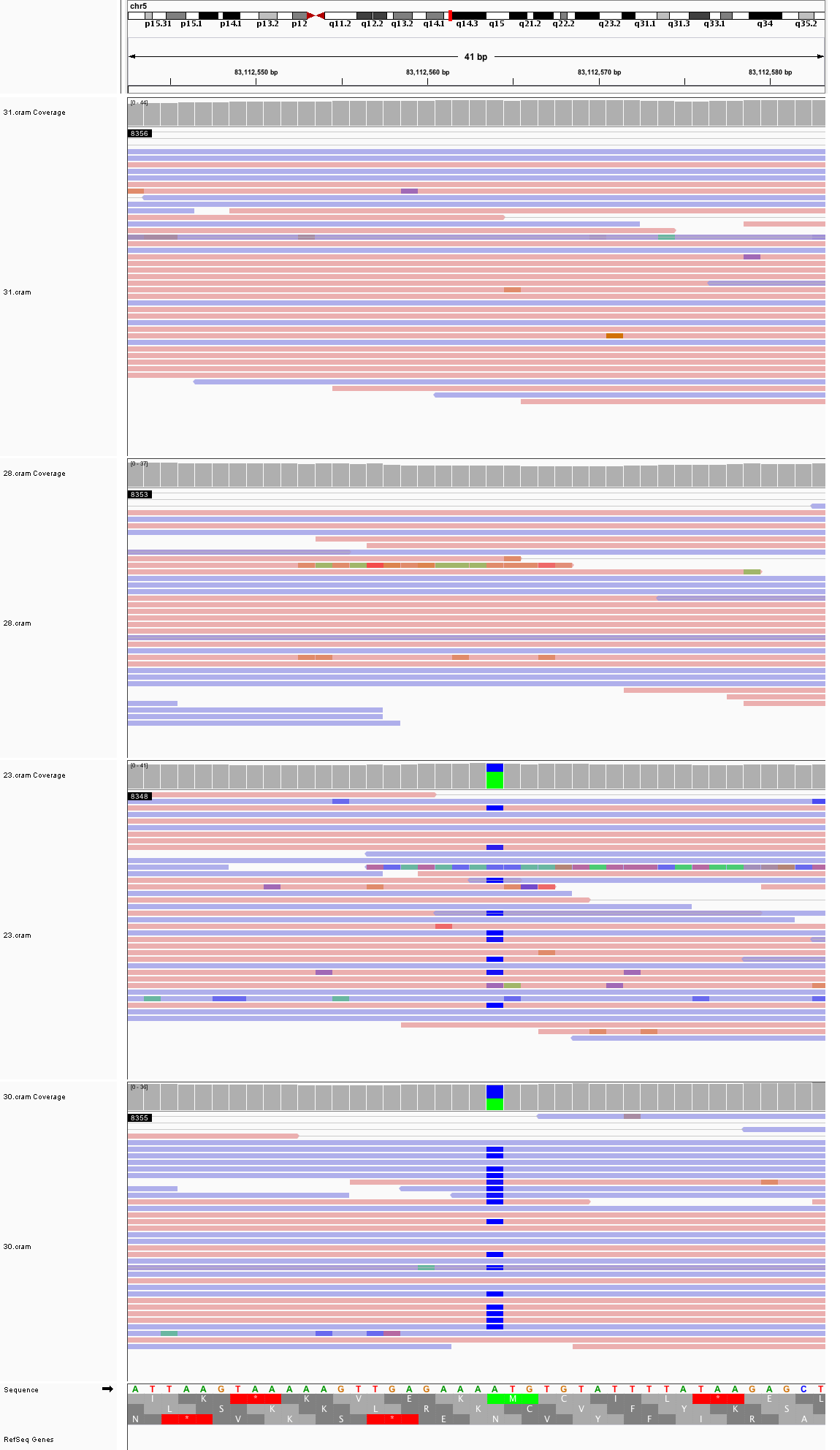

### chr5_99,533,226_99,533,266.png

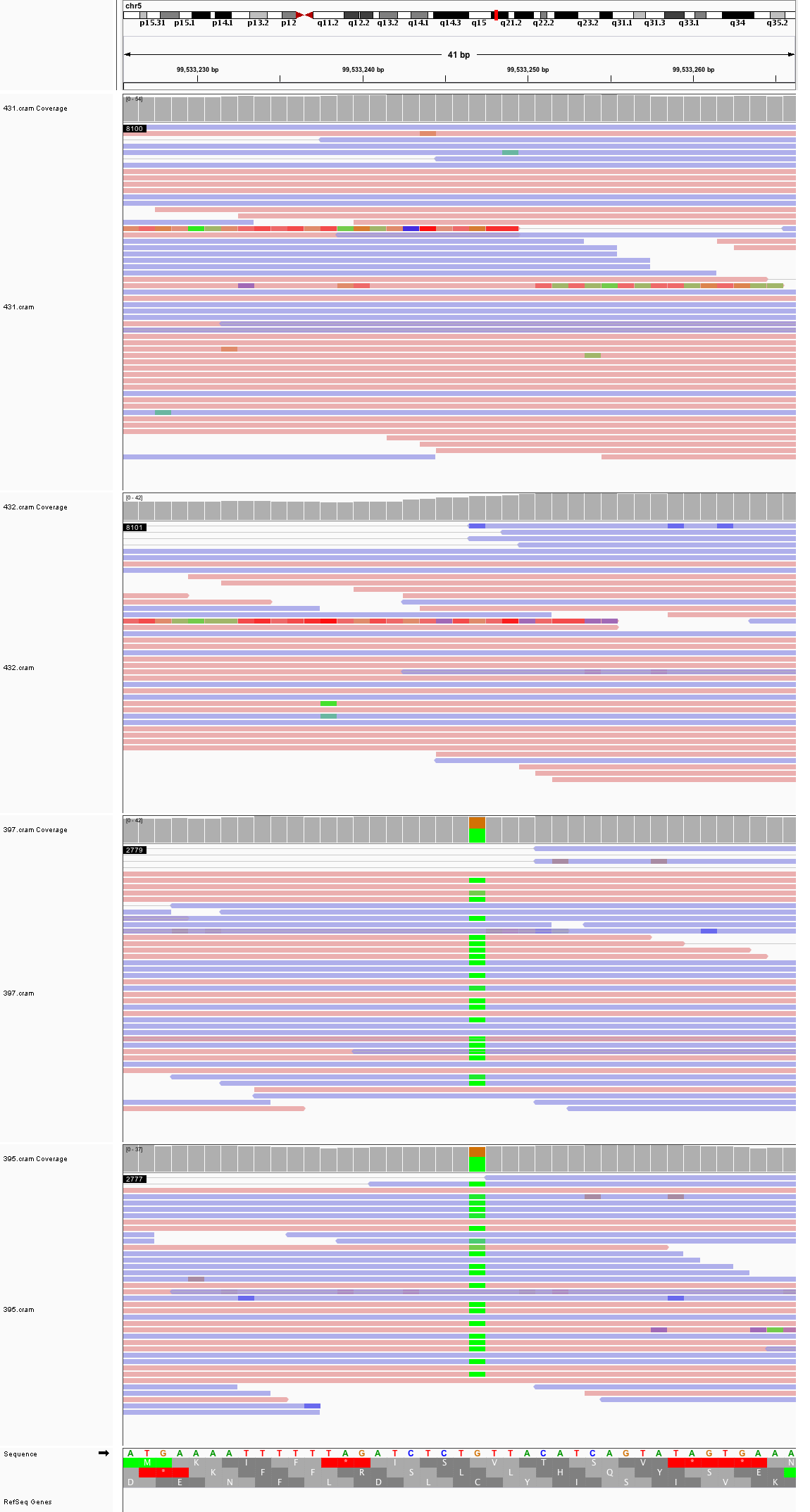

### chr5_175,948,678_175,948,718.png

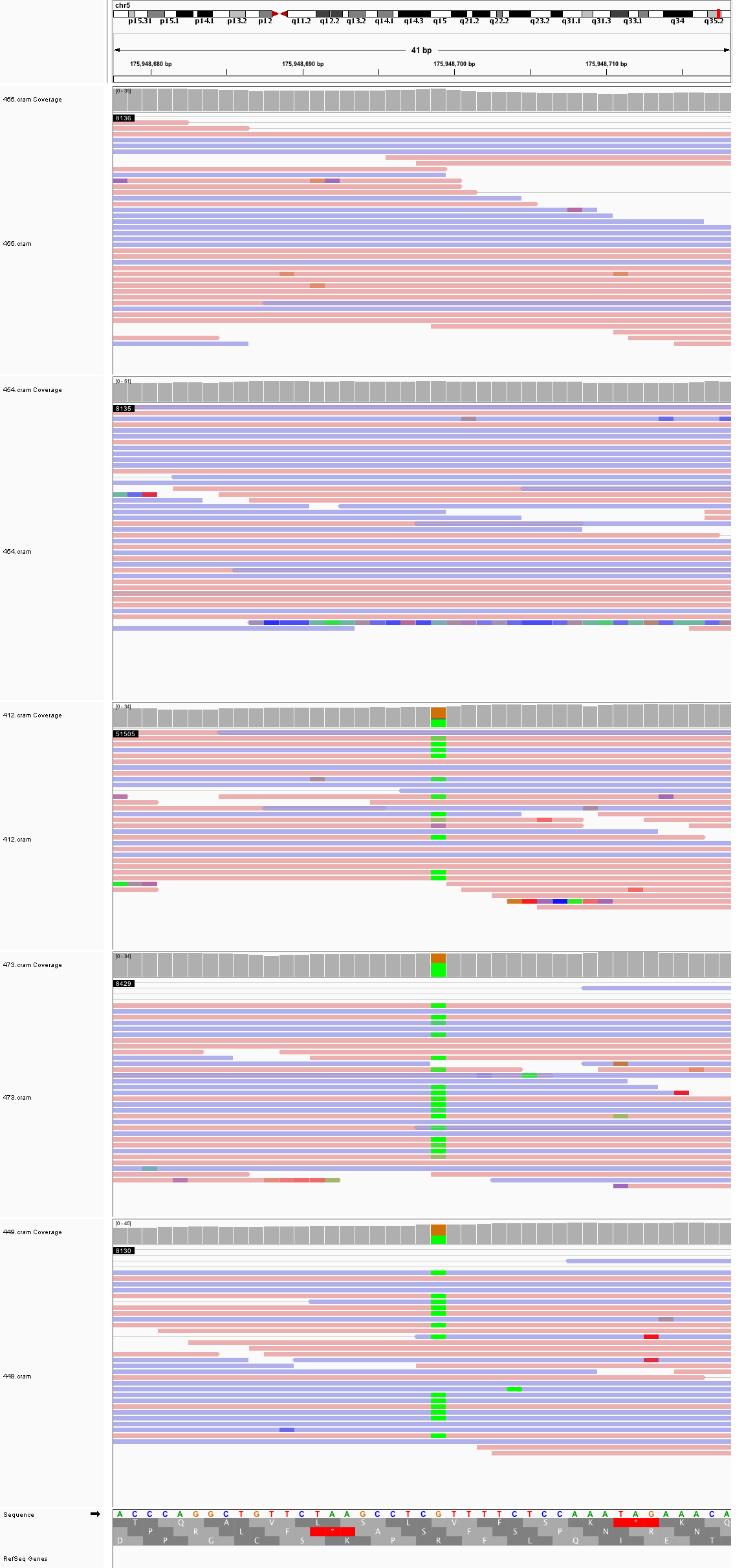

### chr6_9,830,850_9,830,890.png

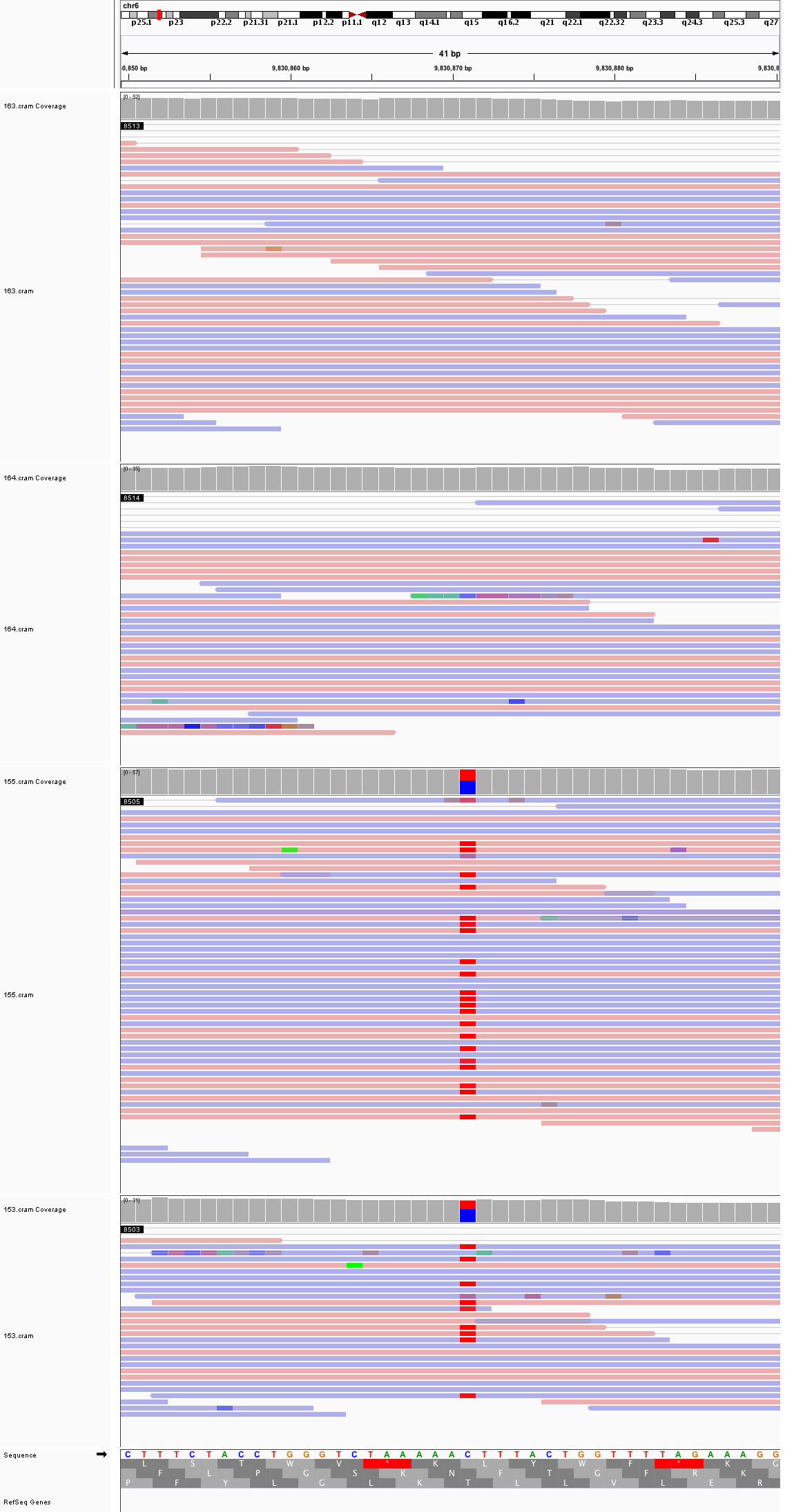

### chr6_25,583,759_25,583,799.png

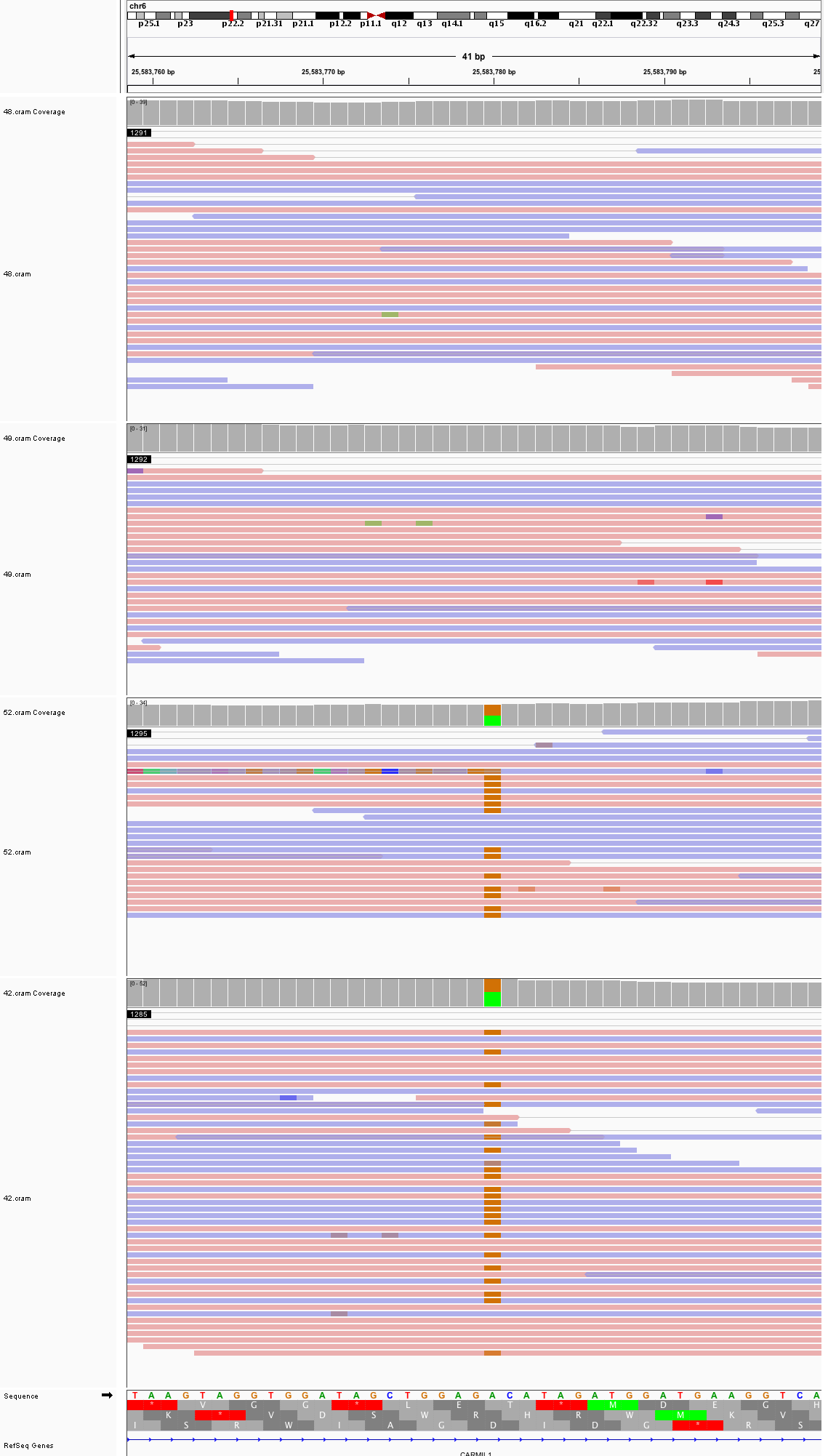

### chr6_121,608,044_121,608,084.png

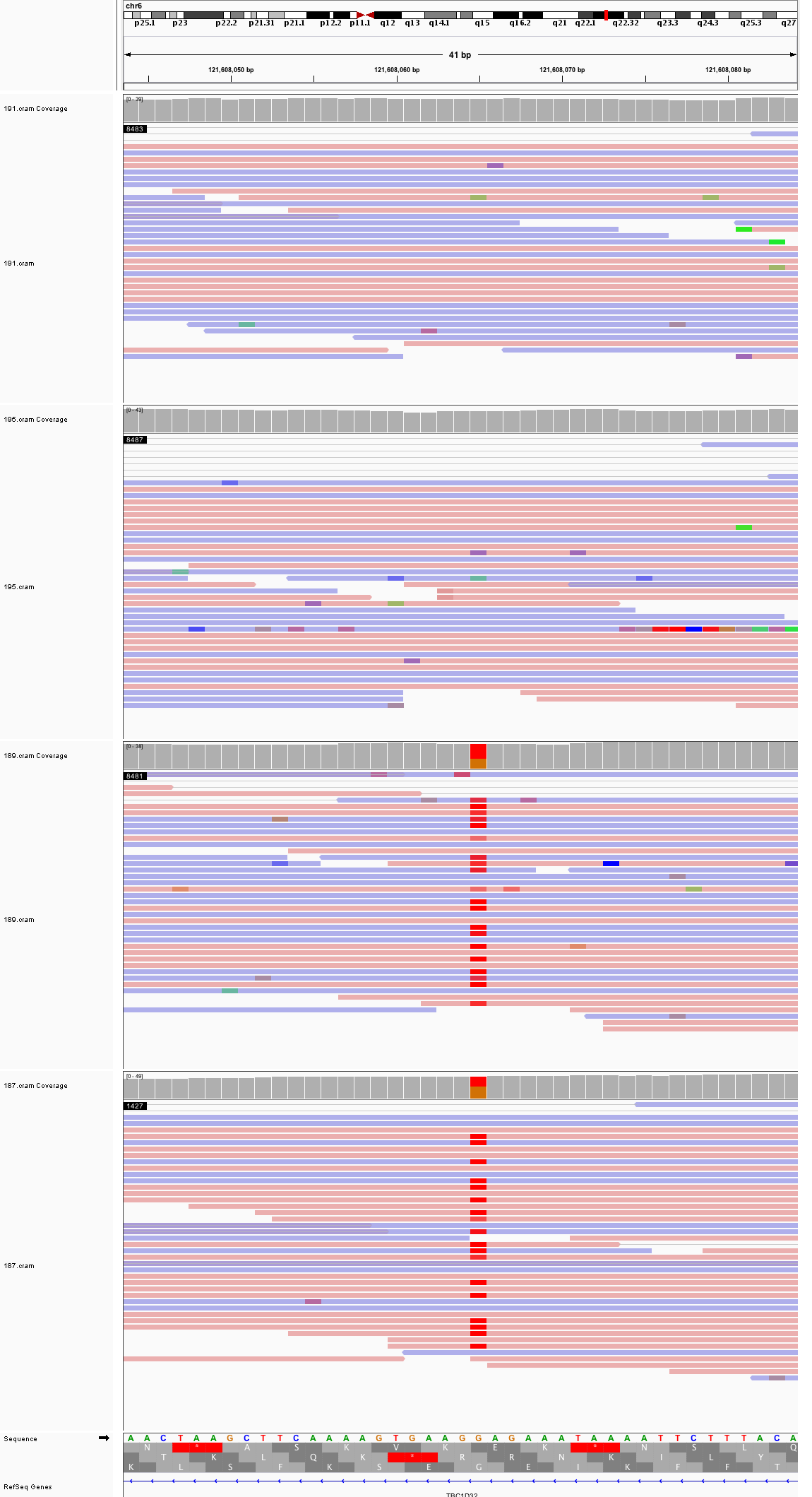

### chr6_130,877,895_130,877,935.png

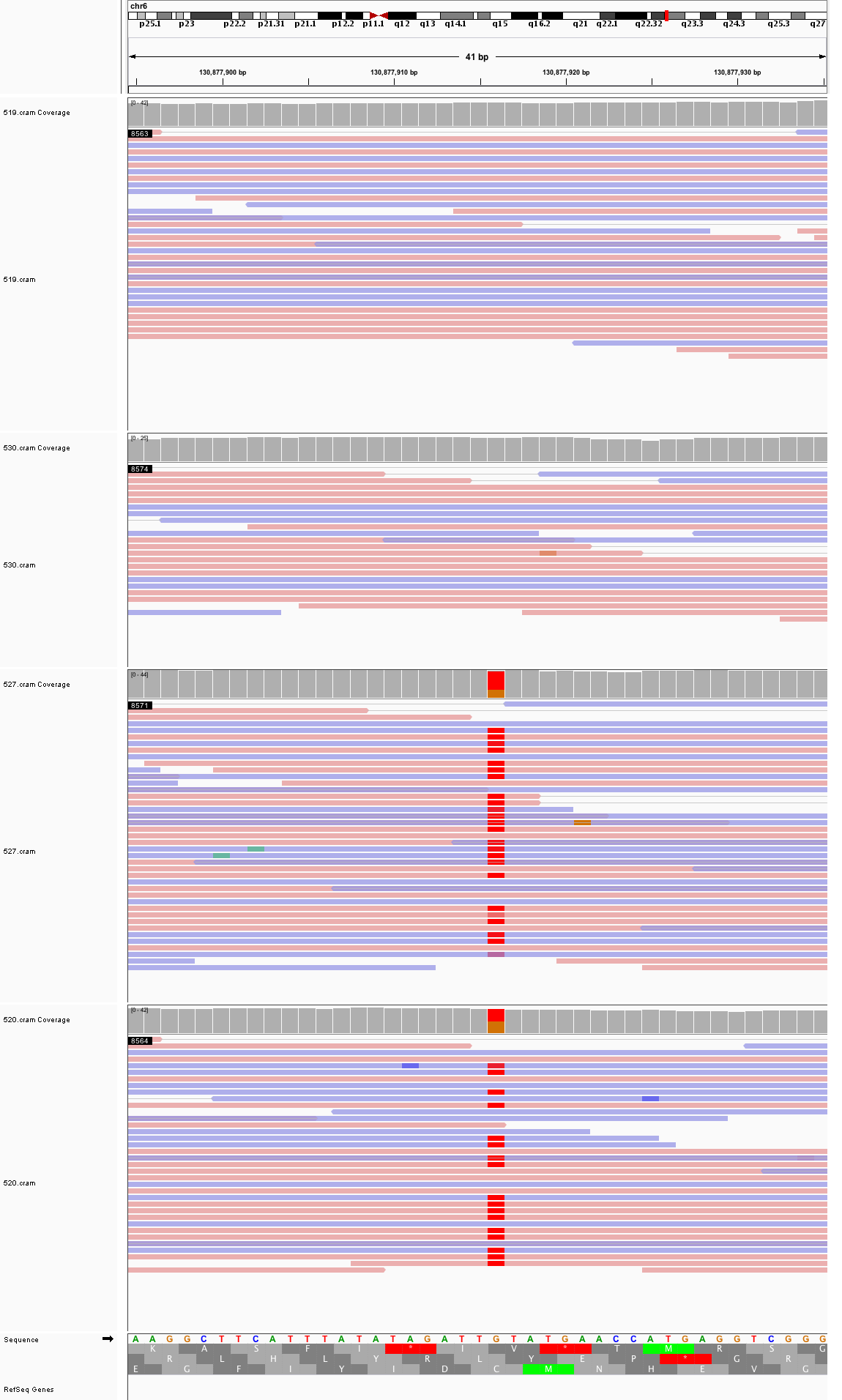

### chr6_145,115,403_145,115,443.png

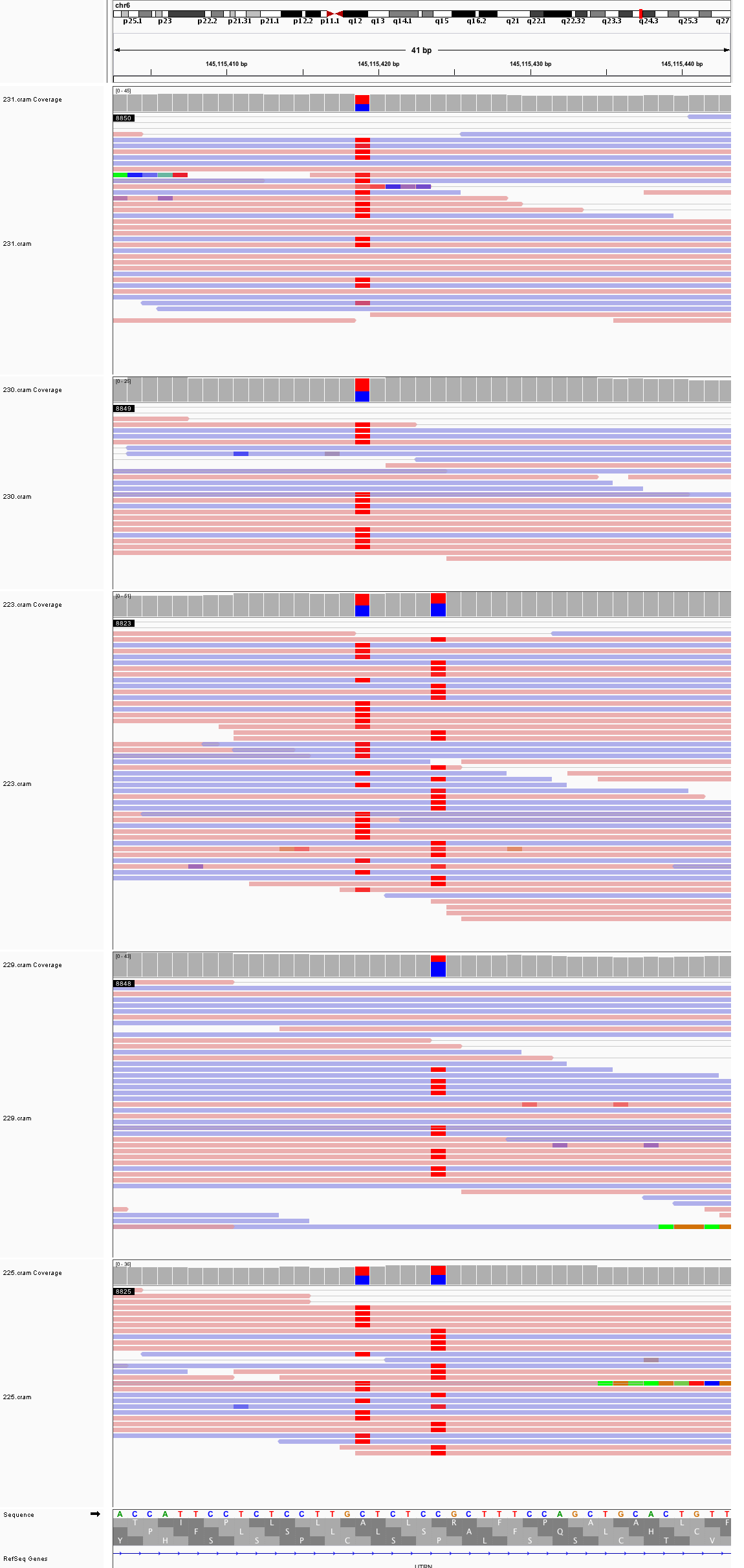

### chr7_18,946,233_18,946,273.png

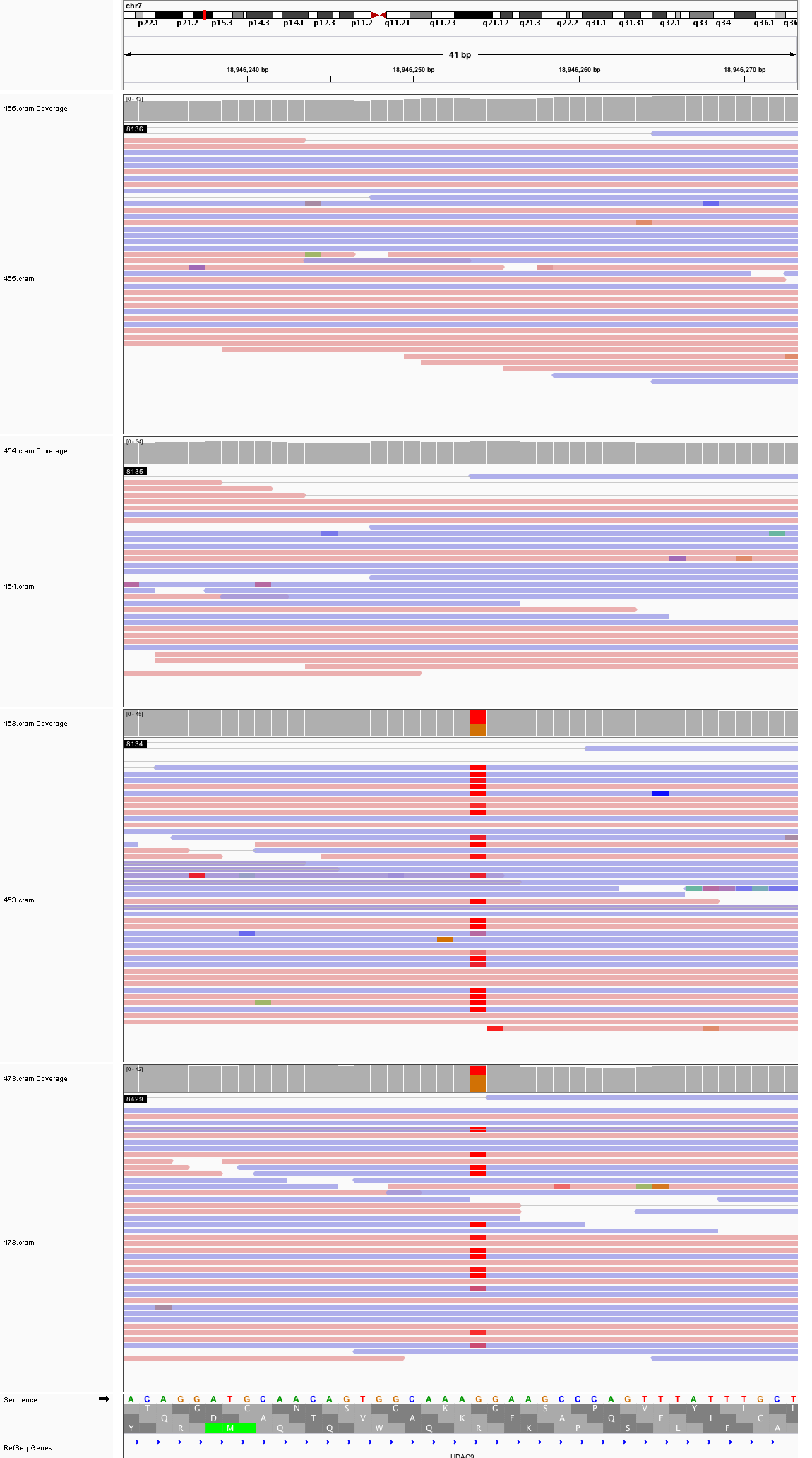

### chr7_75,160,773_75,160,813.png

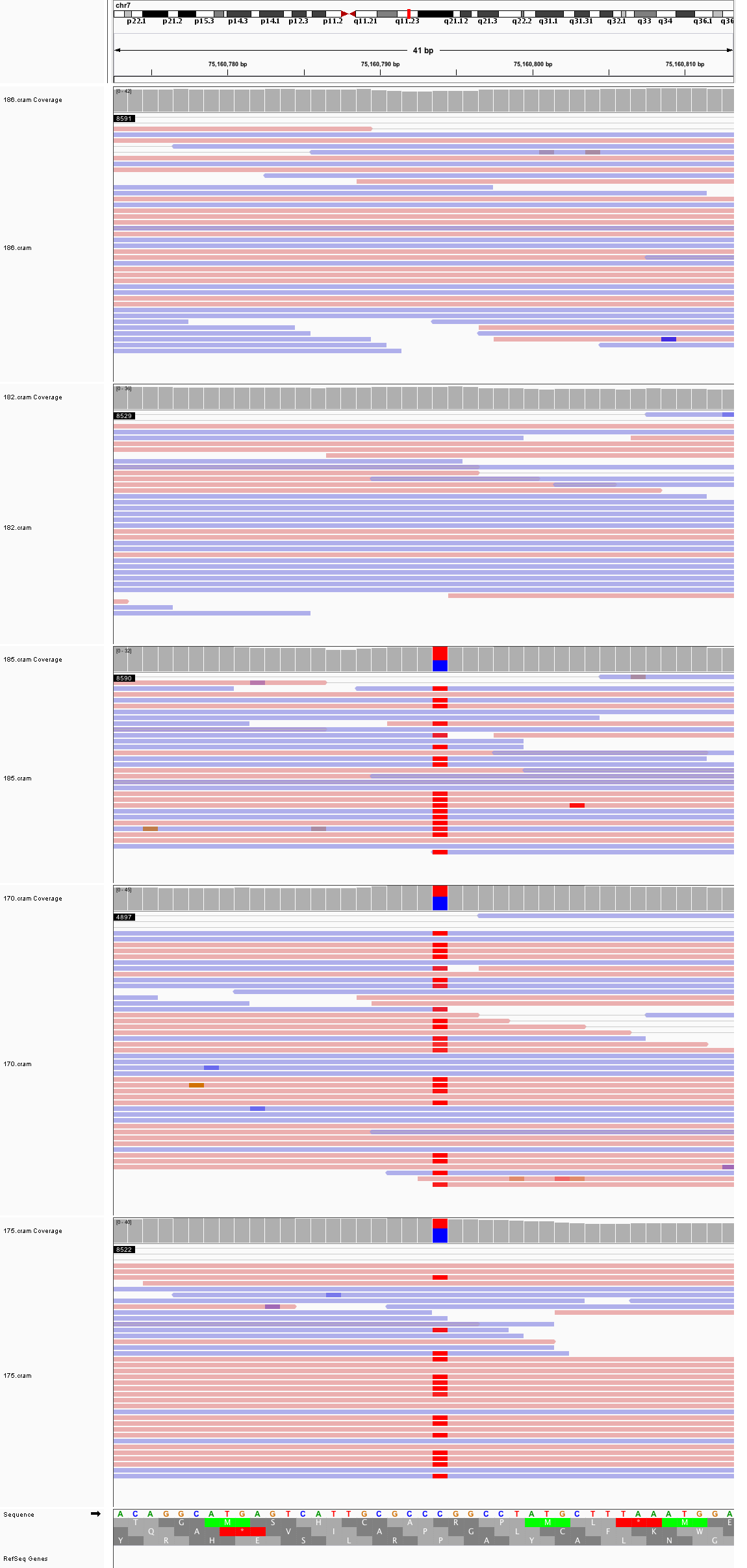

### chr7_151,283,178_151,283,218.png

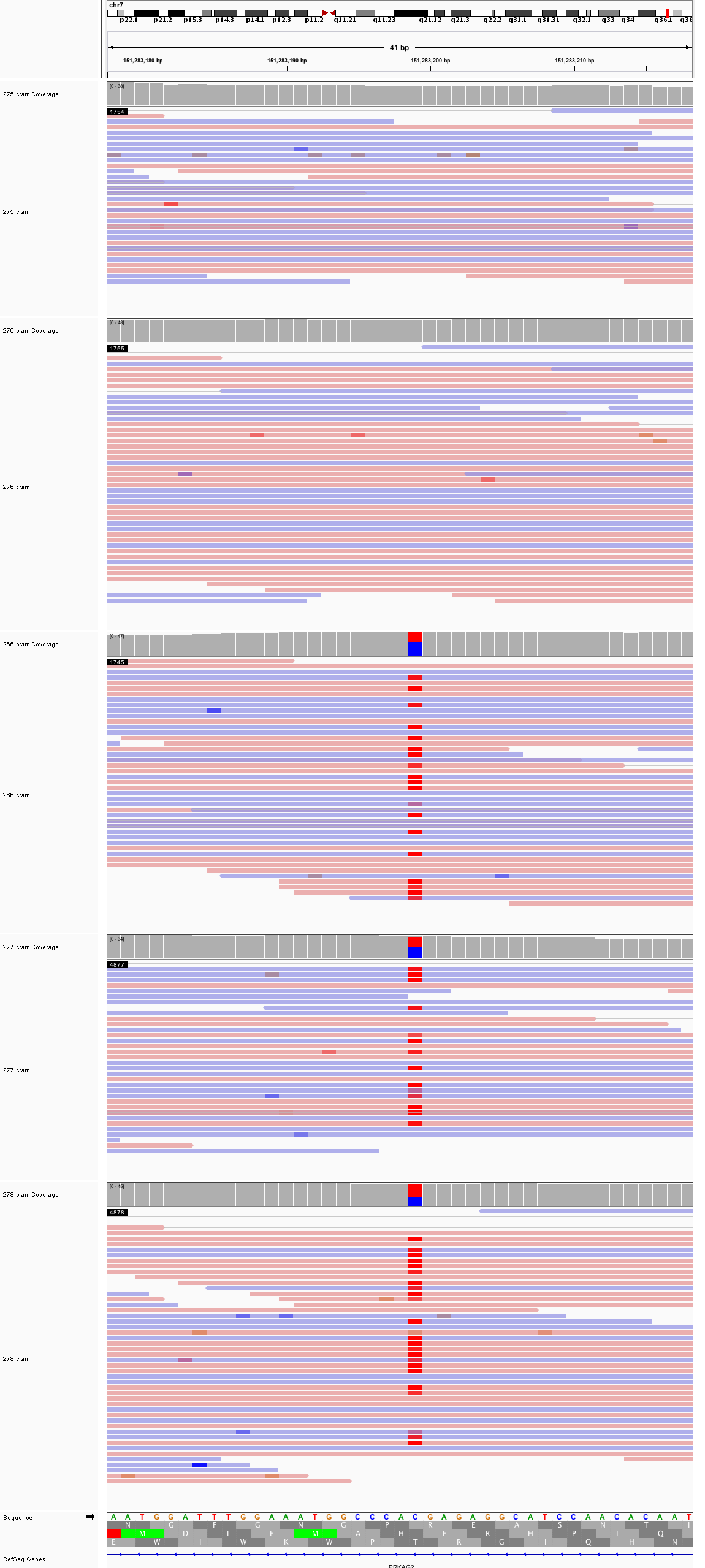

### chr8_129,726,894_129,726,934.png

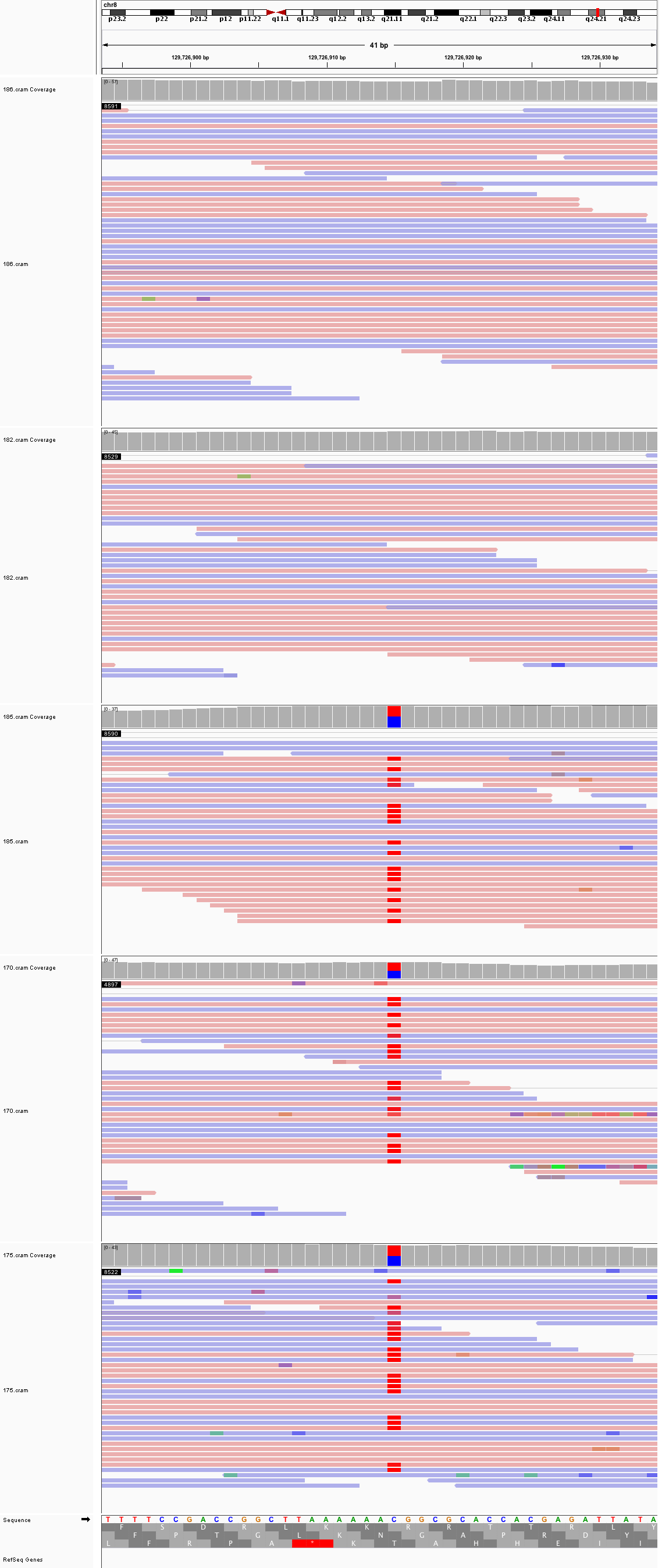

### chr8_139,346,085_139,346,125.png

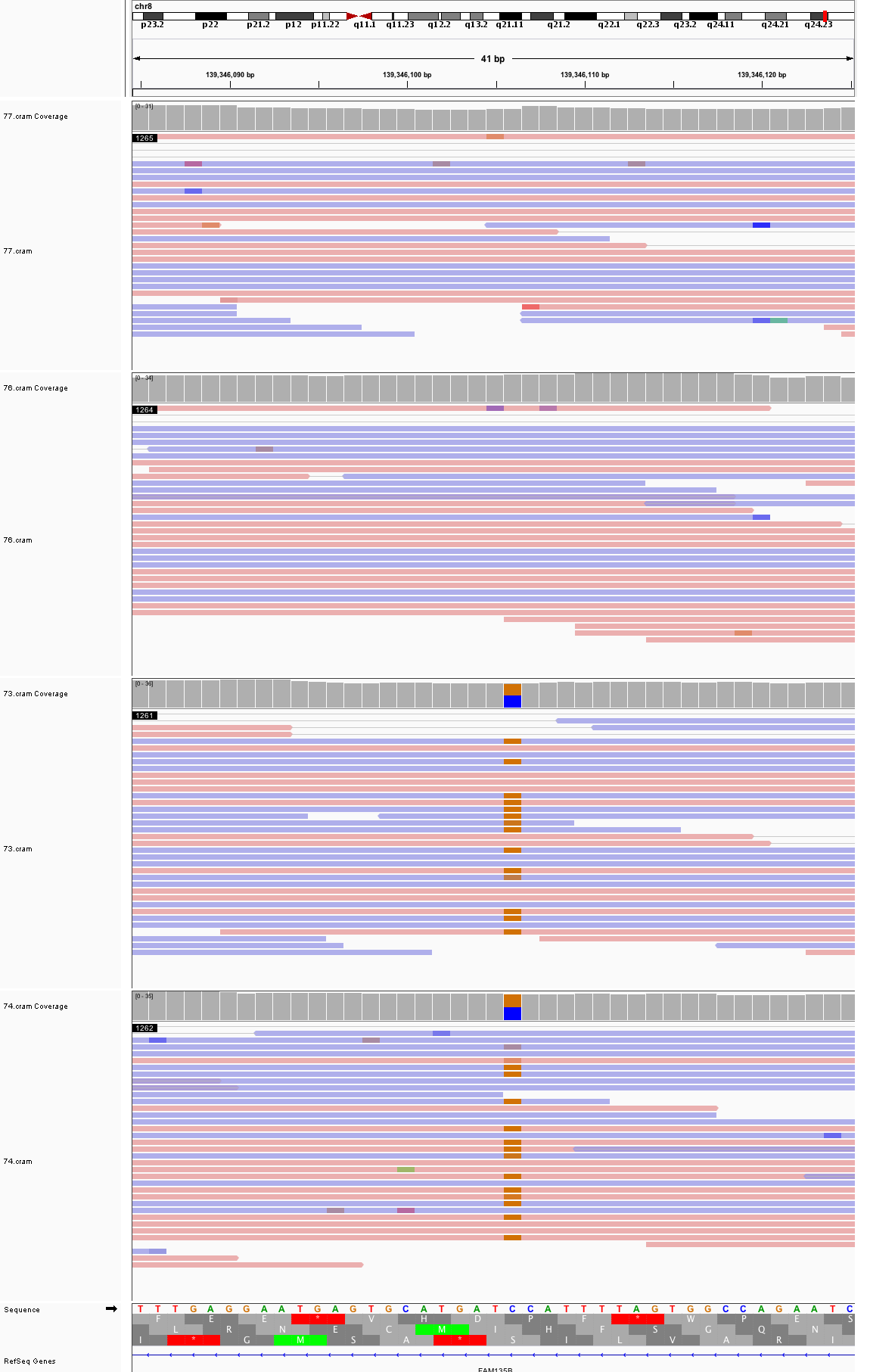

### chr8_141,881,491_141,881,531.png

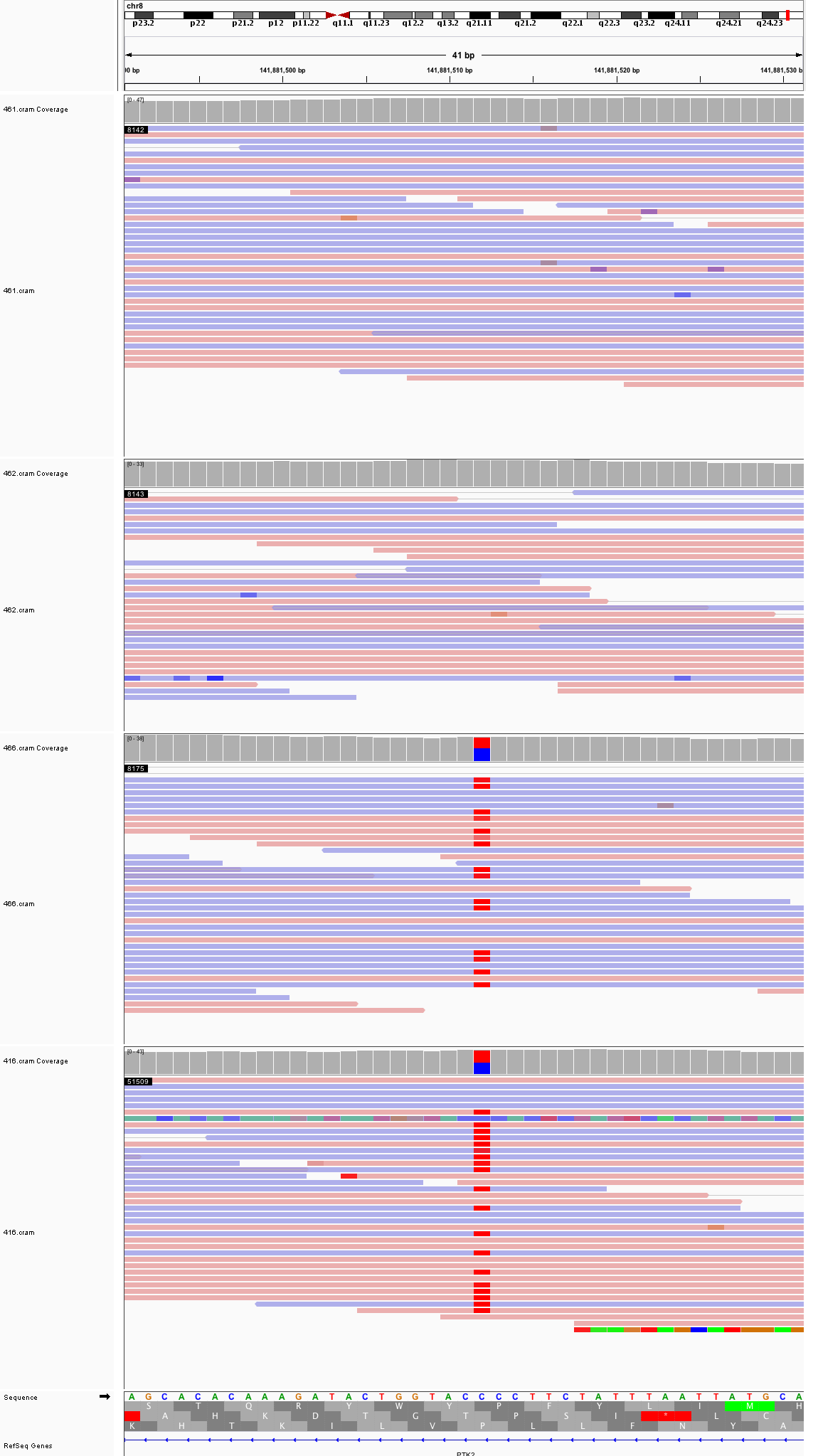

### chr9_1,675,968_1,676,008.png

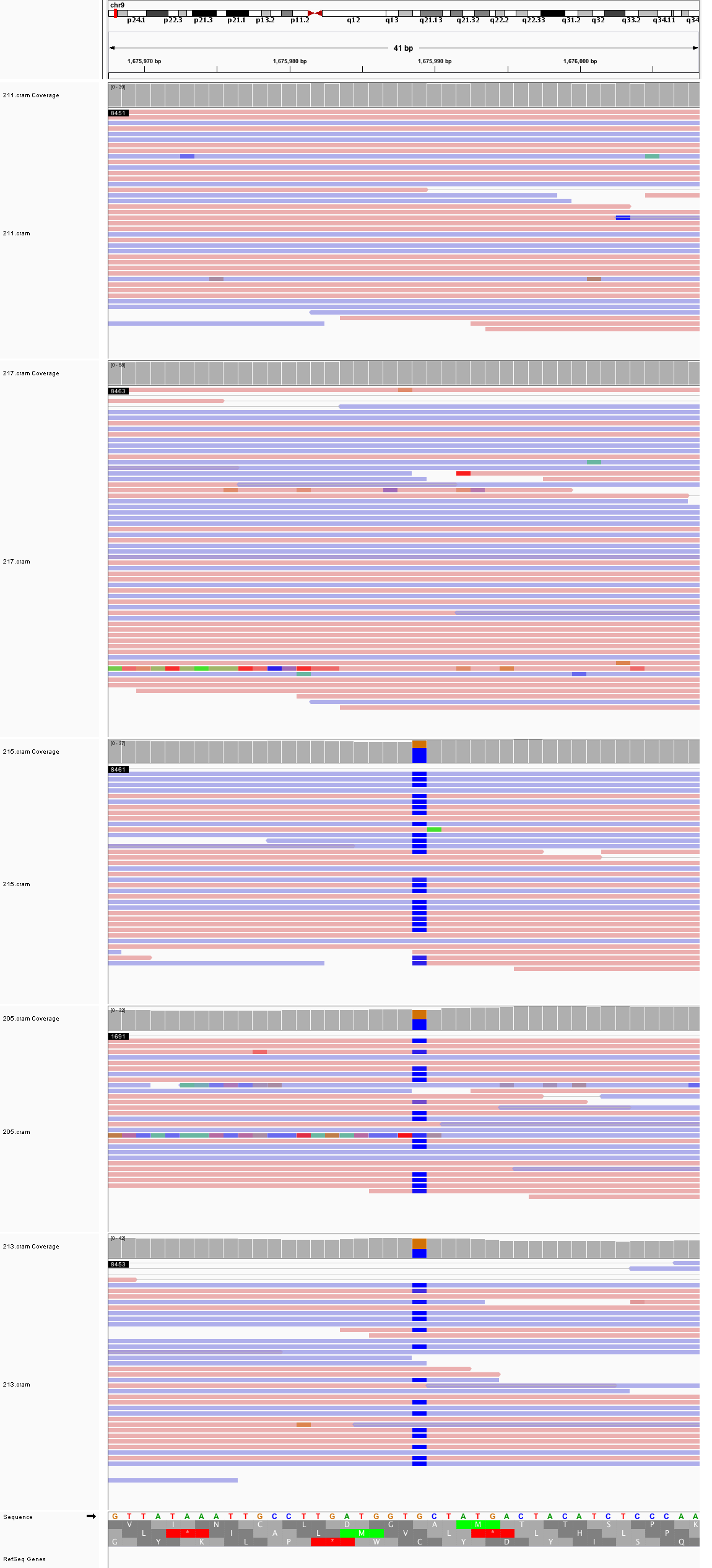

### chr9_35,972,413_35,972,453.png

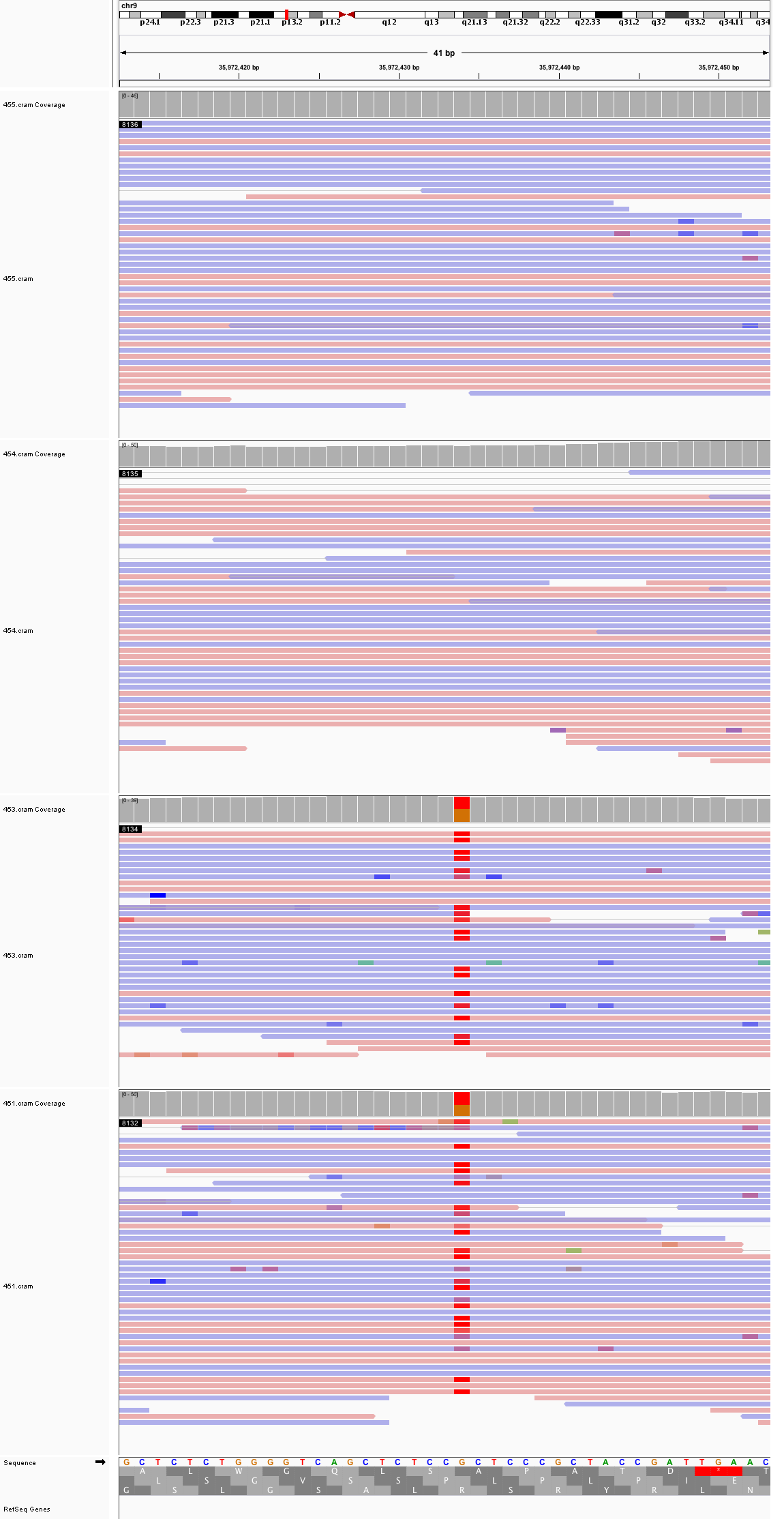

### chr9_37,969,874_37,969,914.png

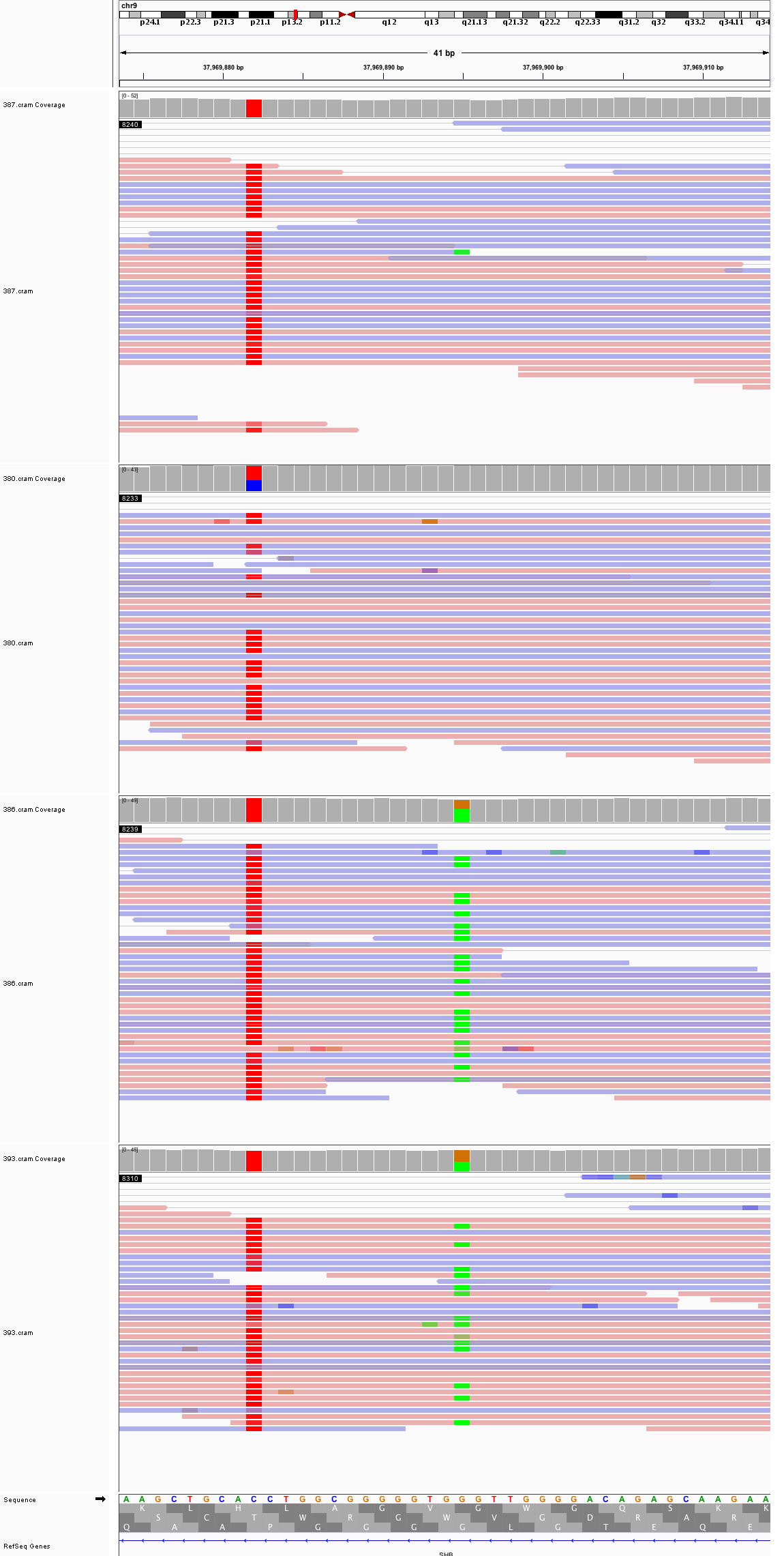

### chr9_82,055_82,095.png

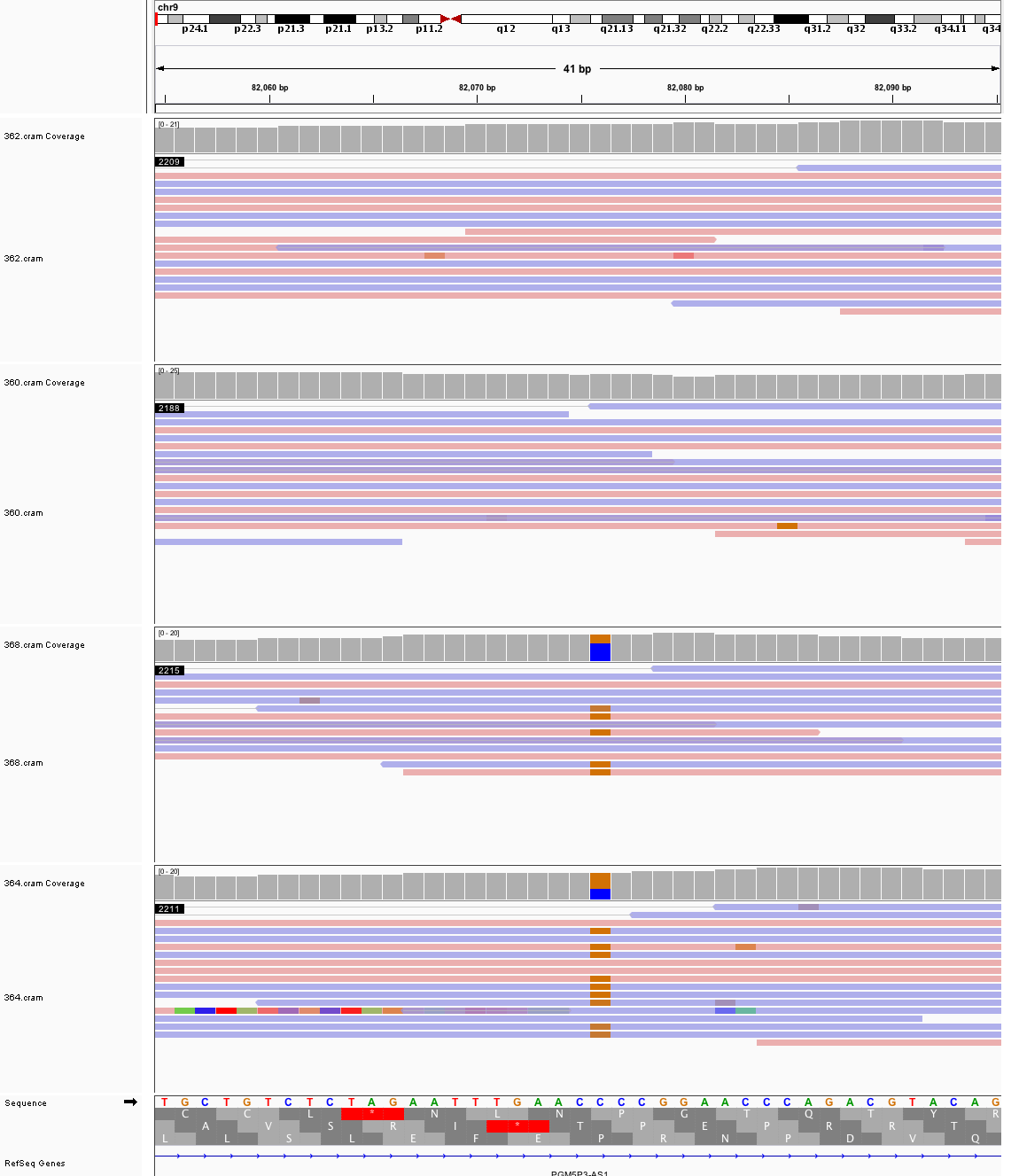

### chr9_104,822,465_104,822,505.png

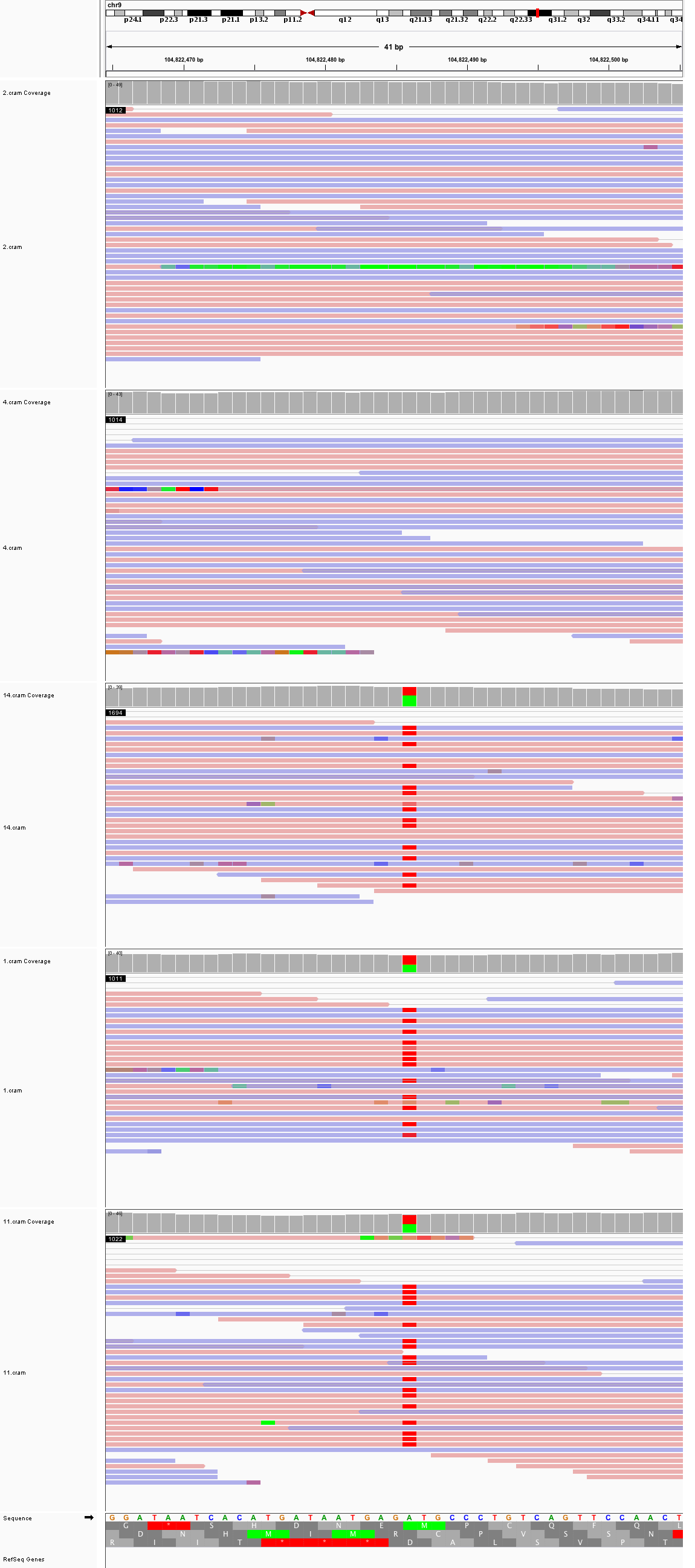

### chr9_115,447,299_115,447,339.png

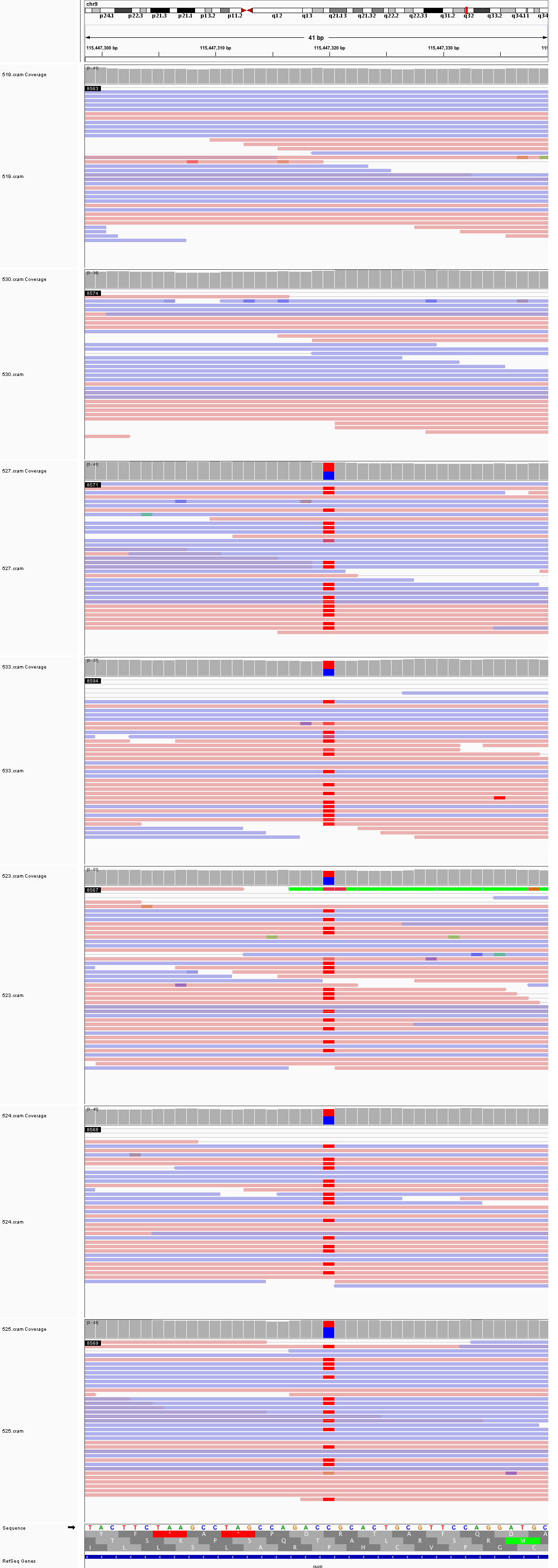

### chr9_131,638,913_131,638,953.png

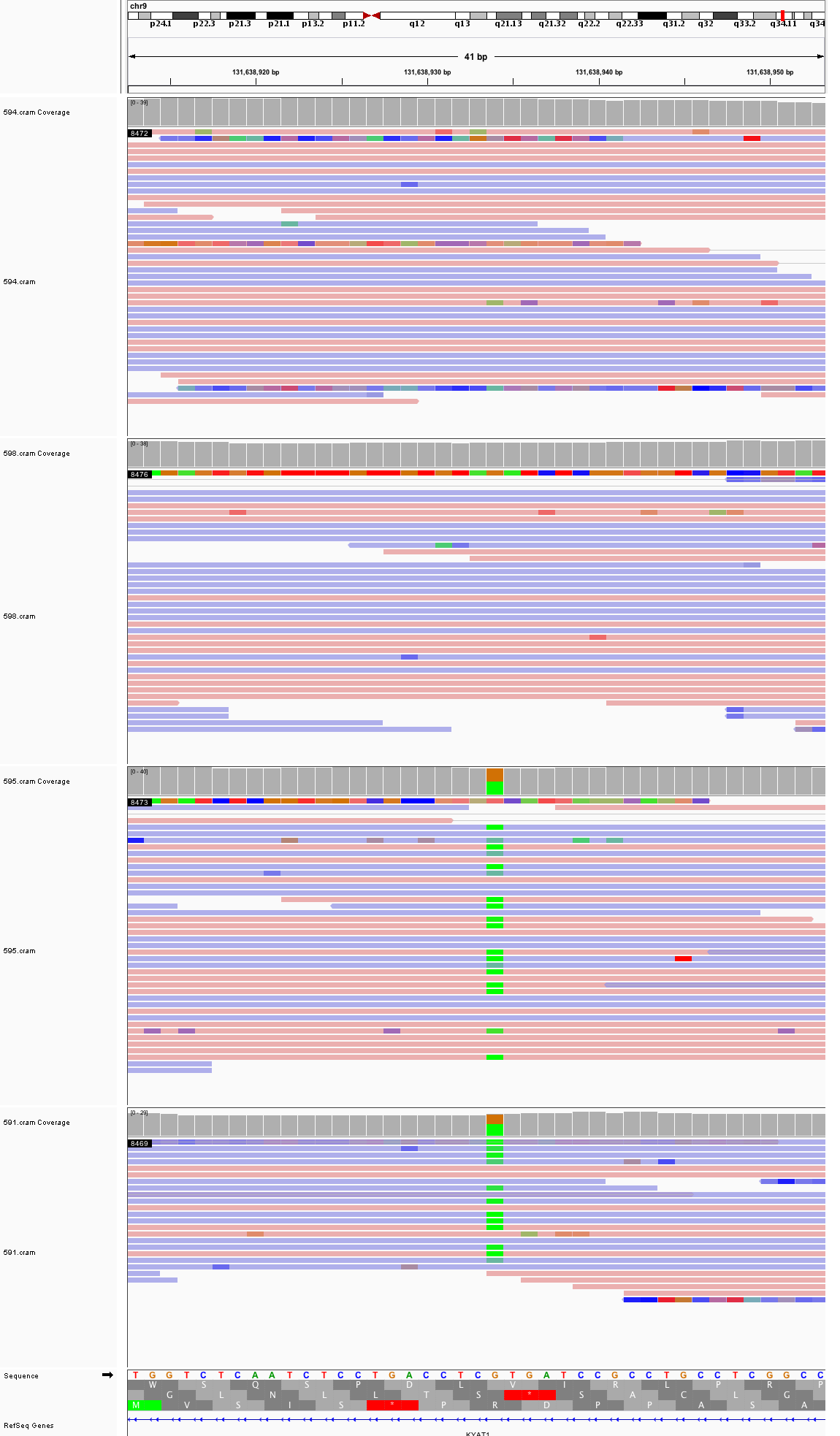

### chr10_26,224,242_26,224,282.png

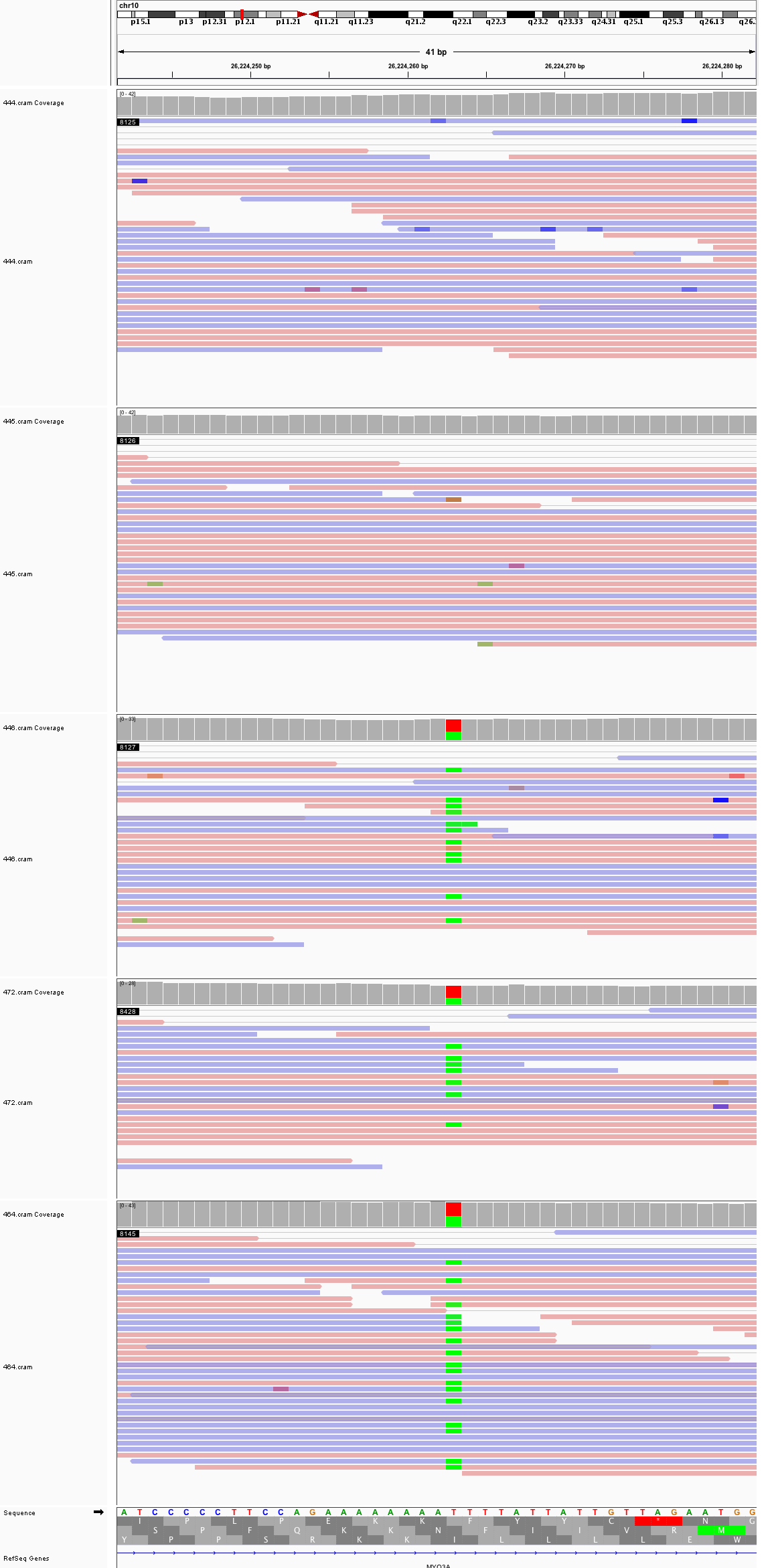

### chr10_91,194,038_91,194,078.png

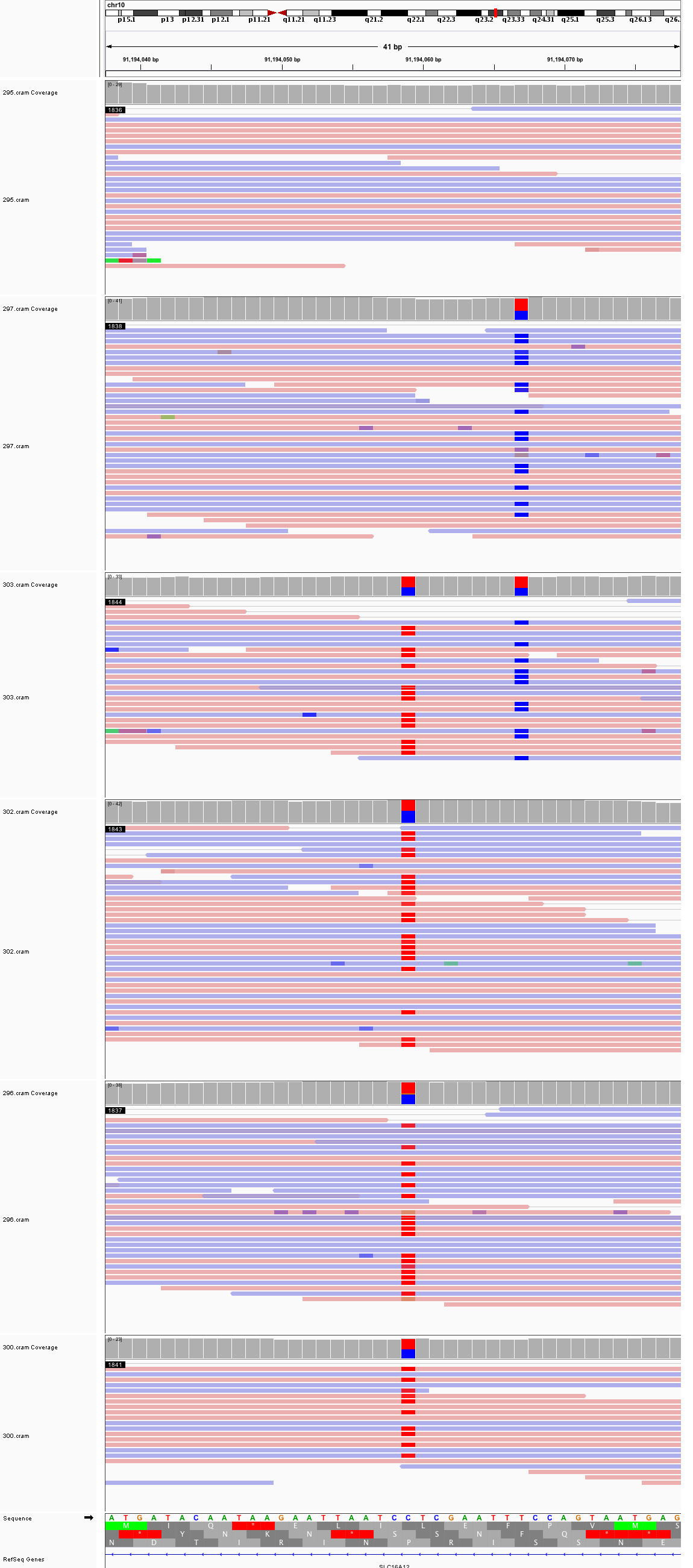

### chr10_111,616,526_111,616,566.png

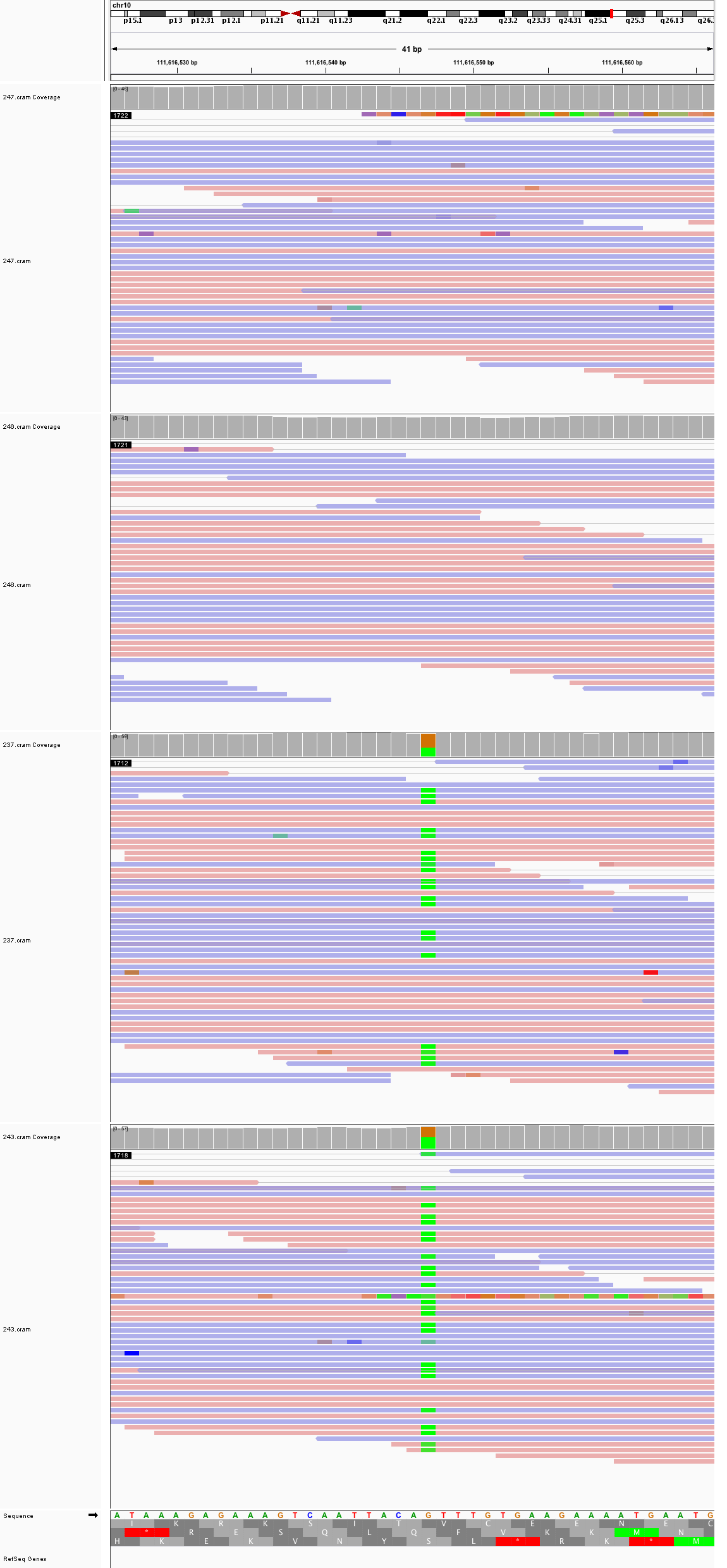

### chr10_131,146,539_131,146,579.png

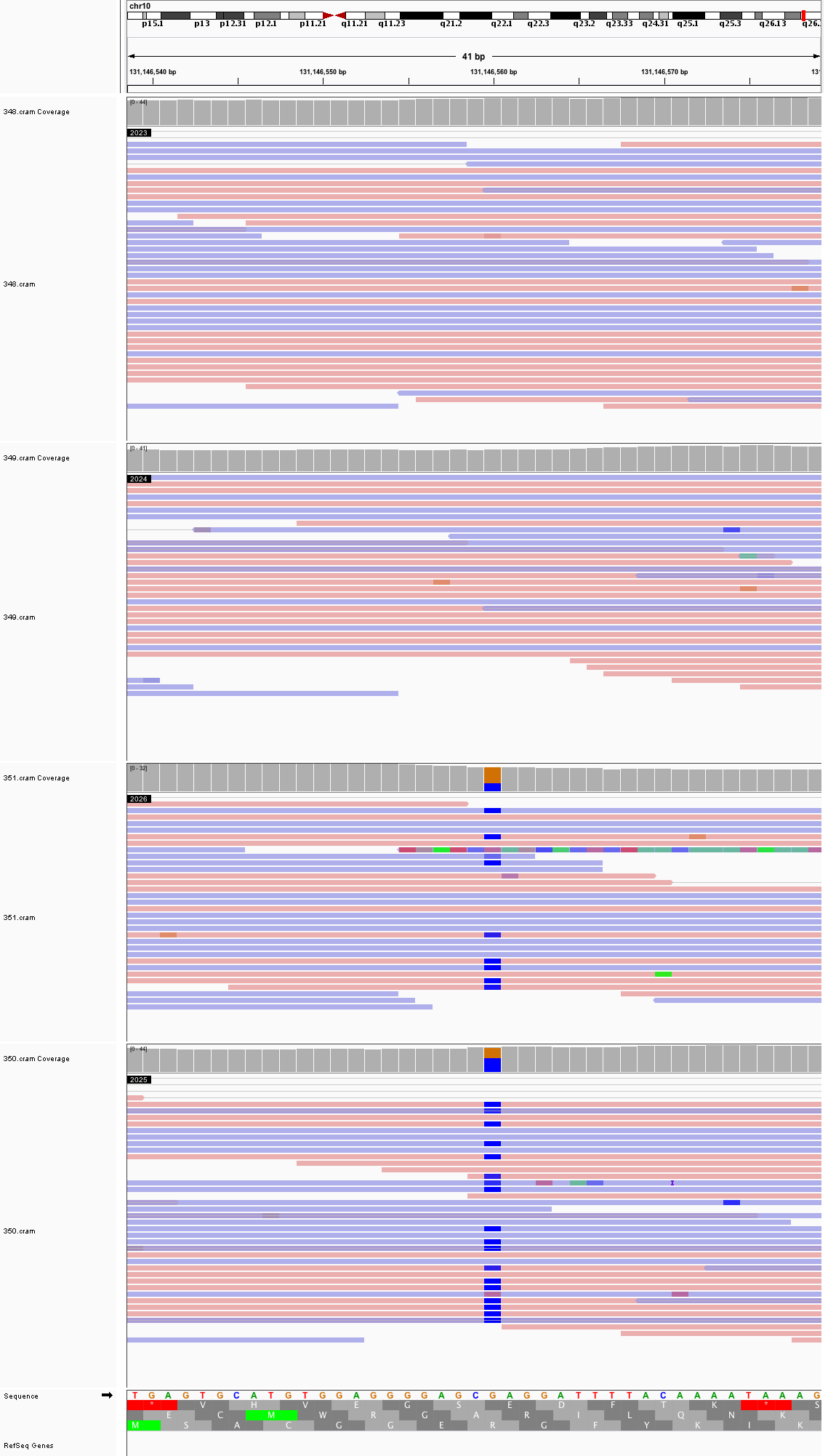

### chr10_134,480,428_134,480,468.png

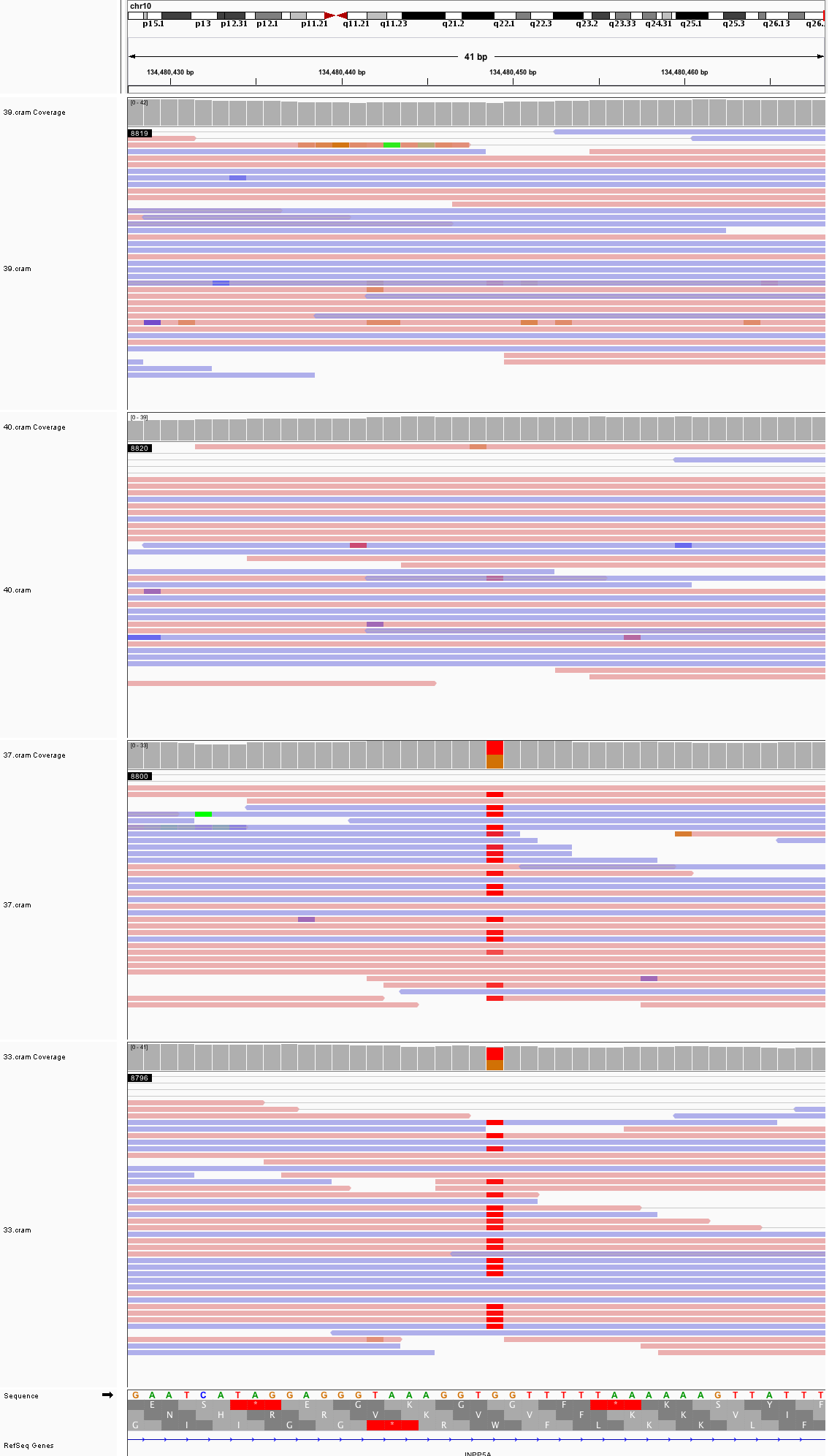

### chr11_50,032,059_50,032,099.png

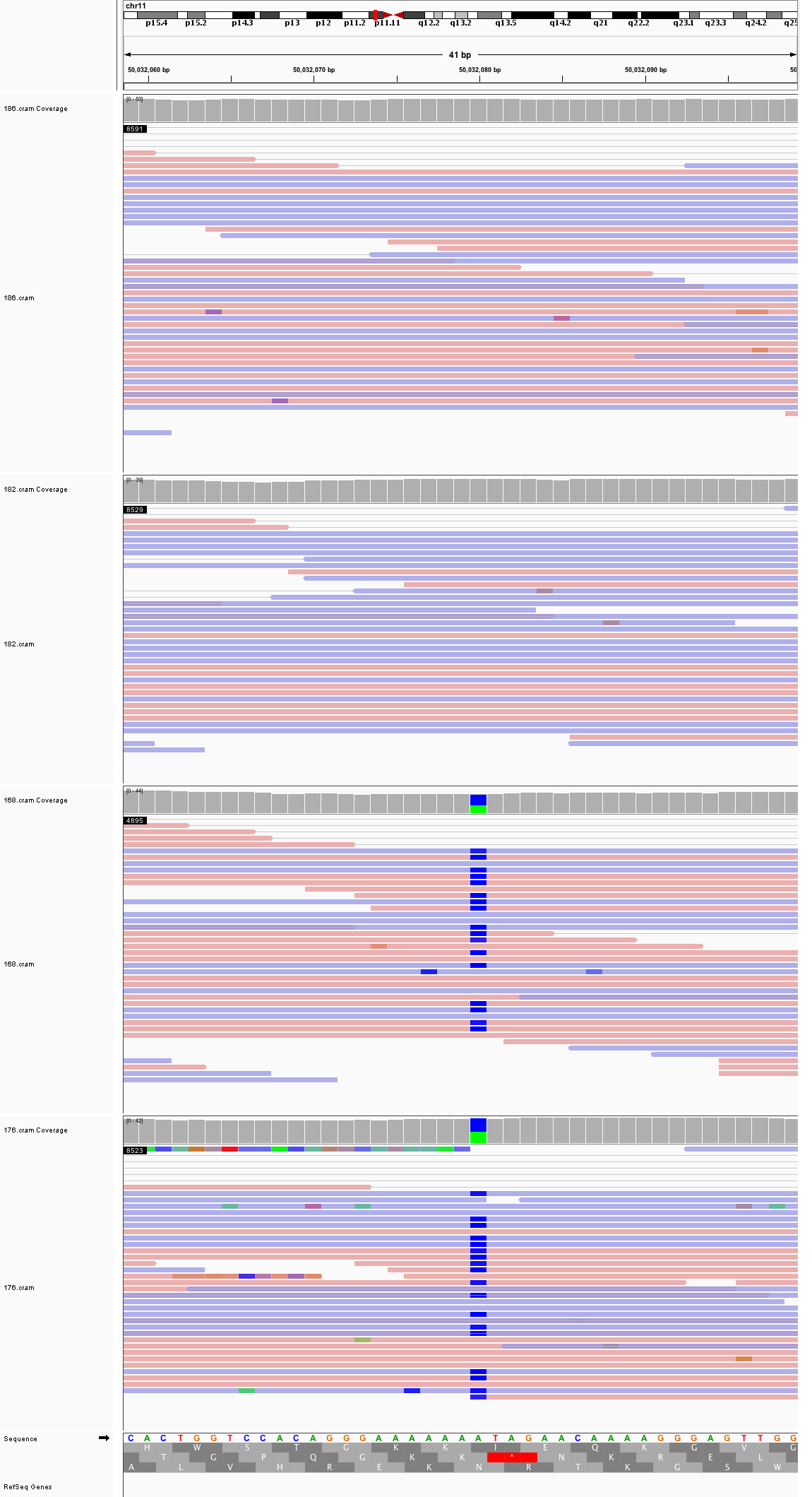

### chr11_64,684,139_64,684,179.png

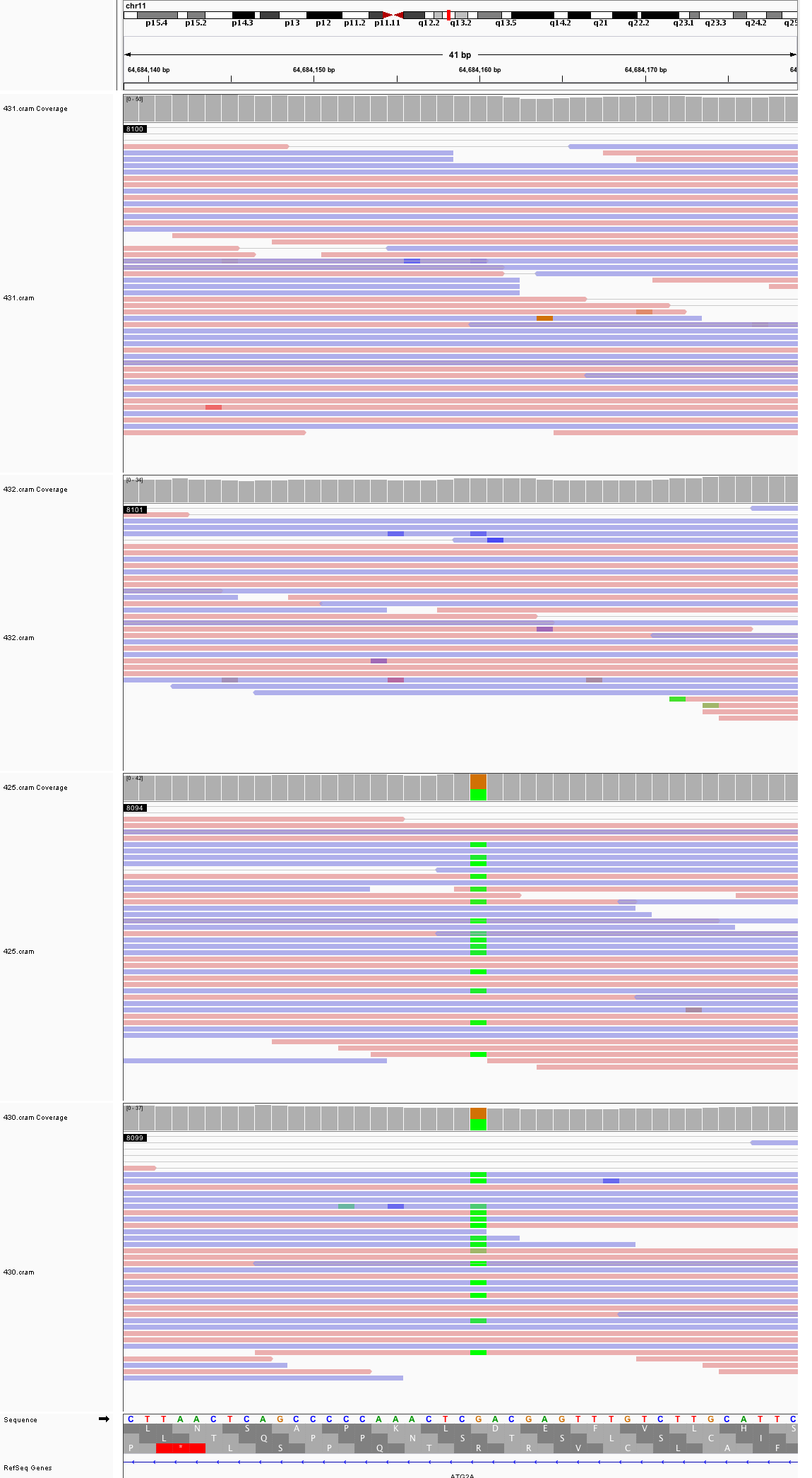

### chr11_64,867,319_64,867,359.png

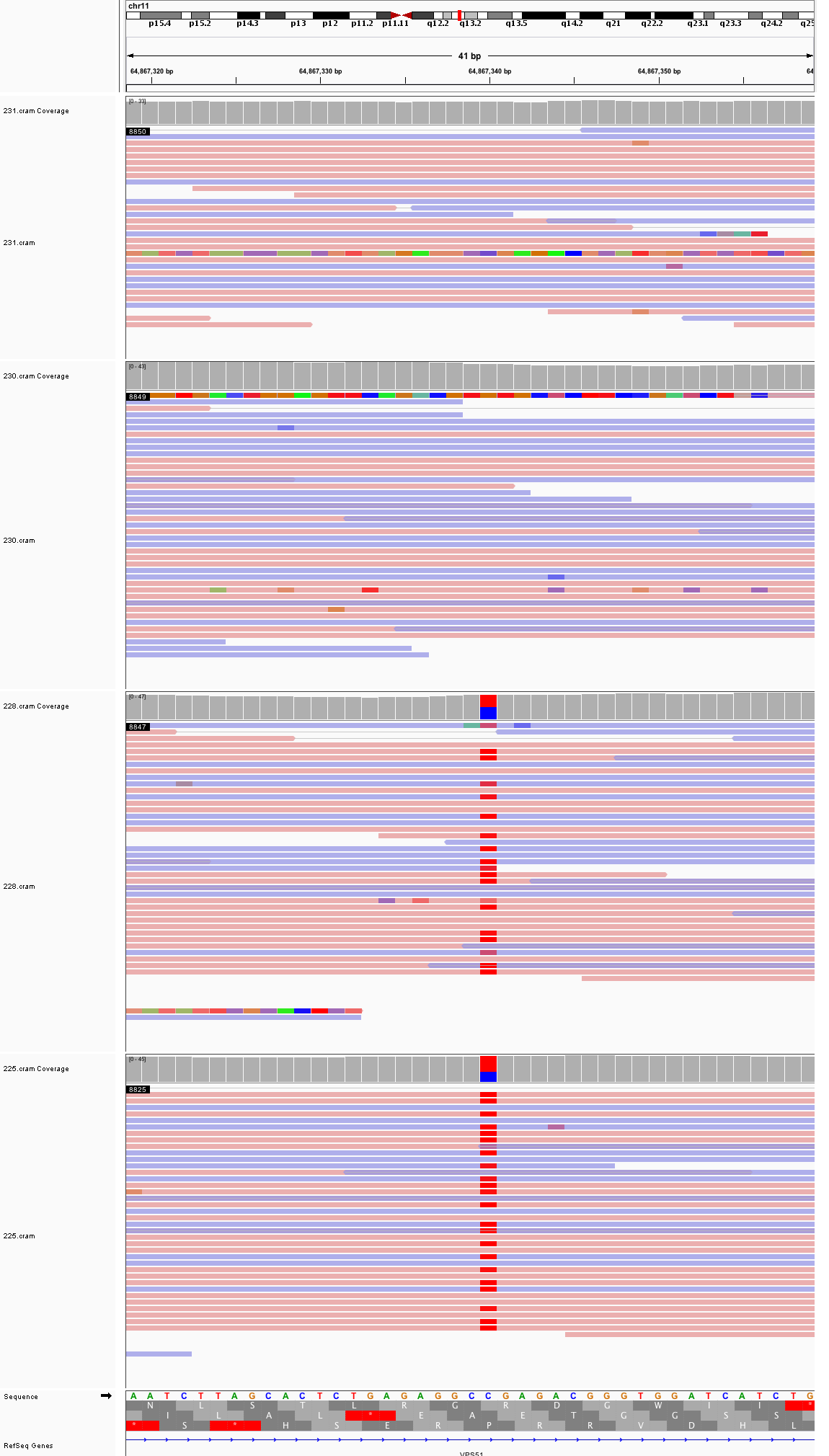

### chr12_70,267,211_70,267,251.png

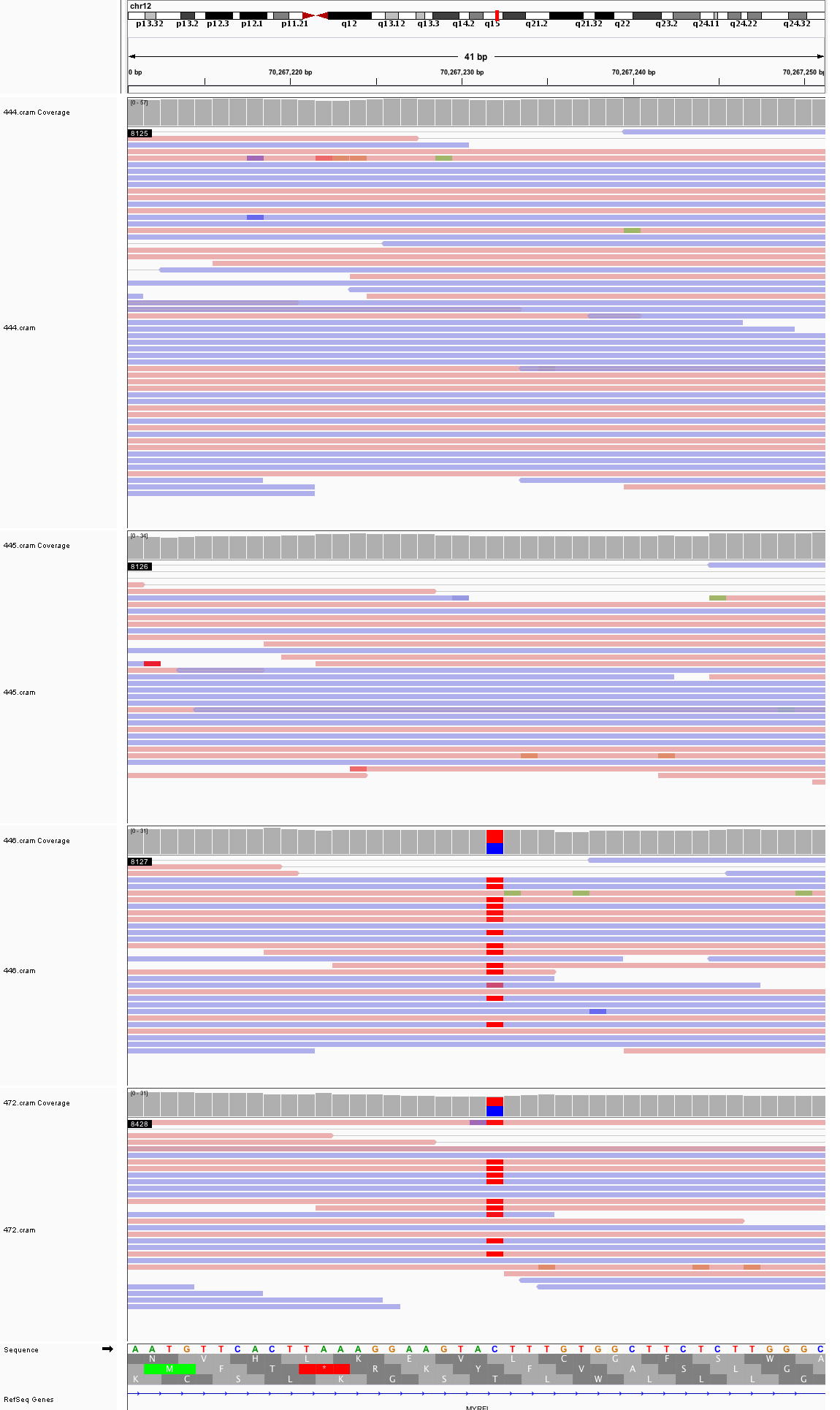

### chr12_85,265,105_85,265,145.png

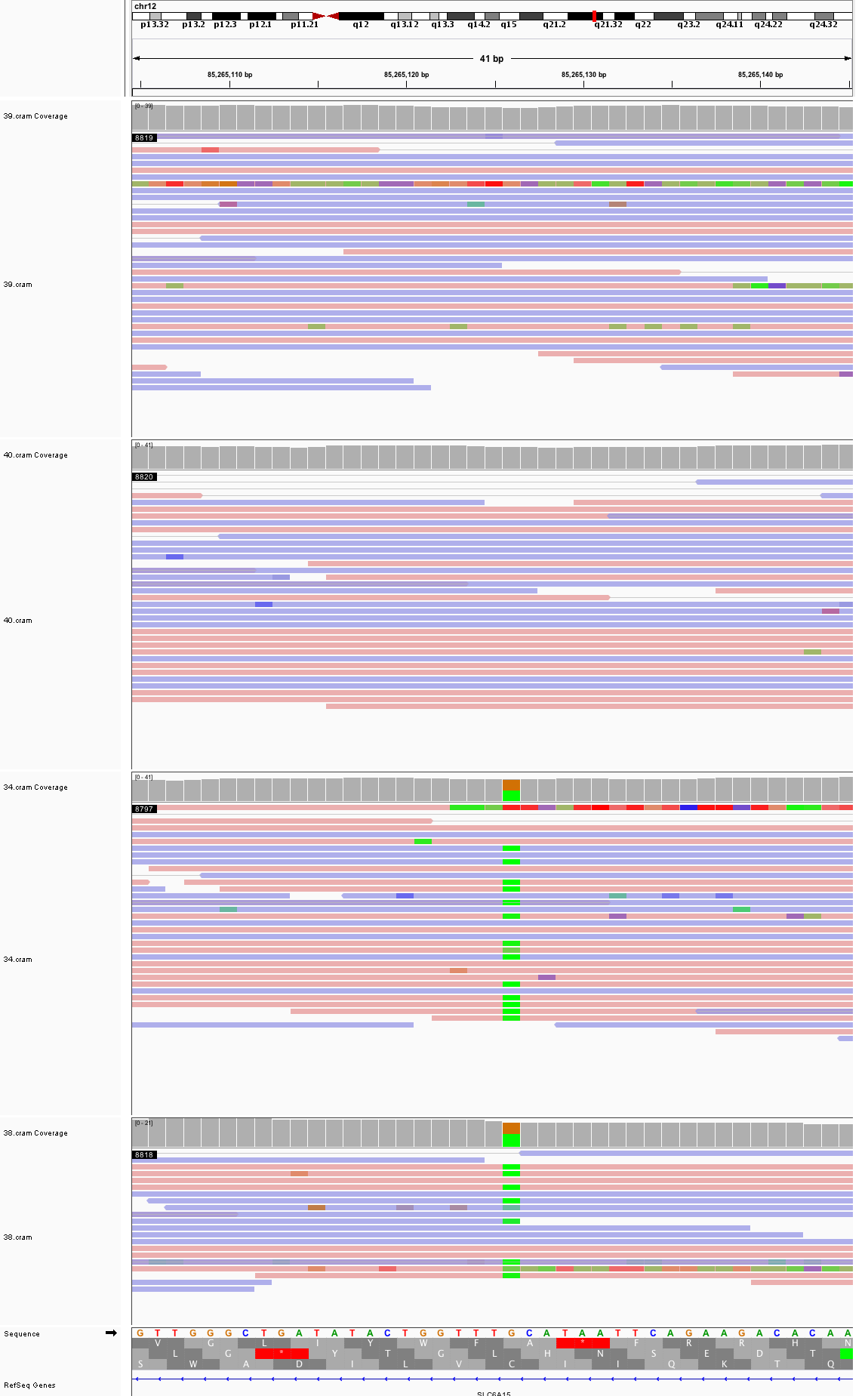

### chr12_98,392,427_98,392,467.png

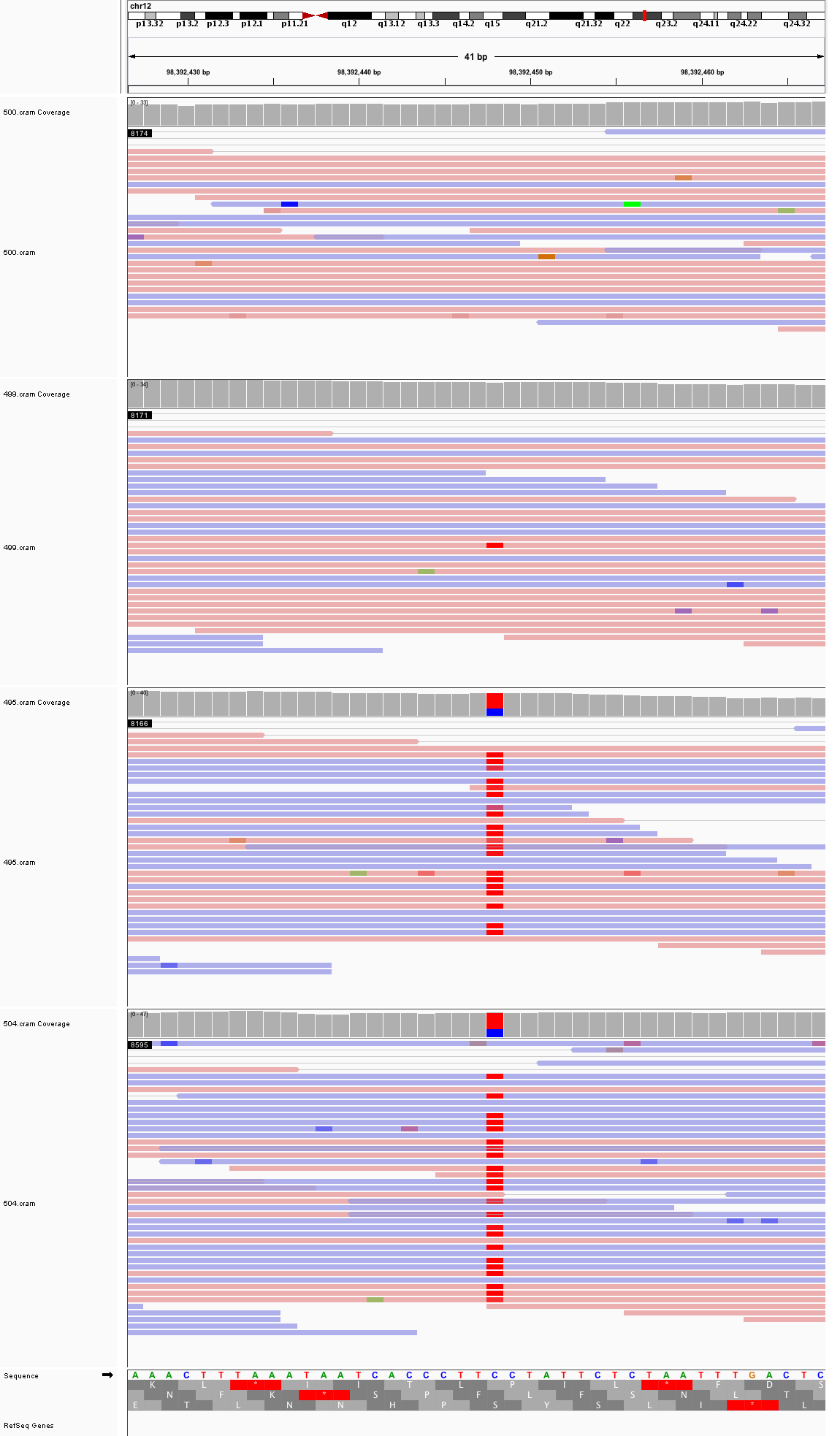

### chr12_105,853,378_105,853,418.png

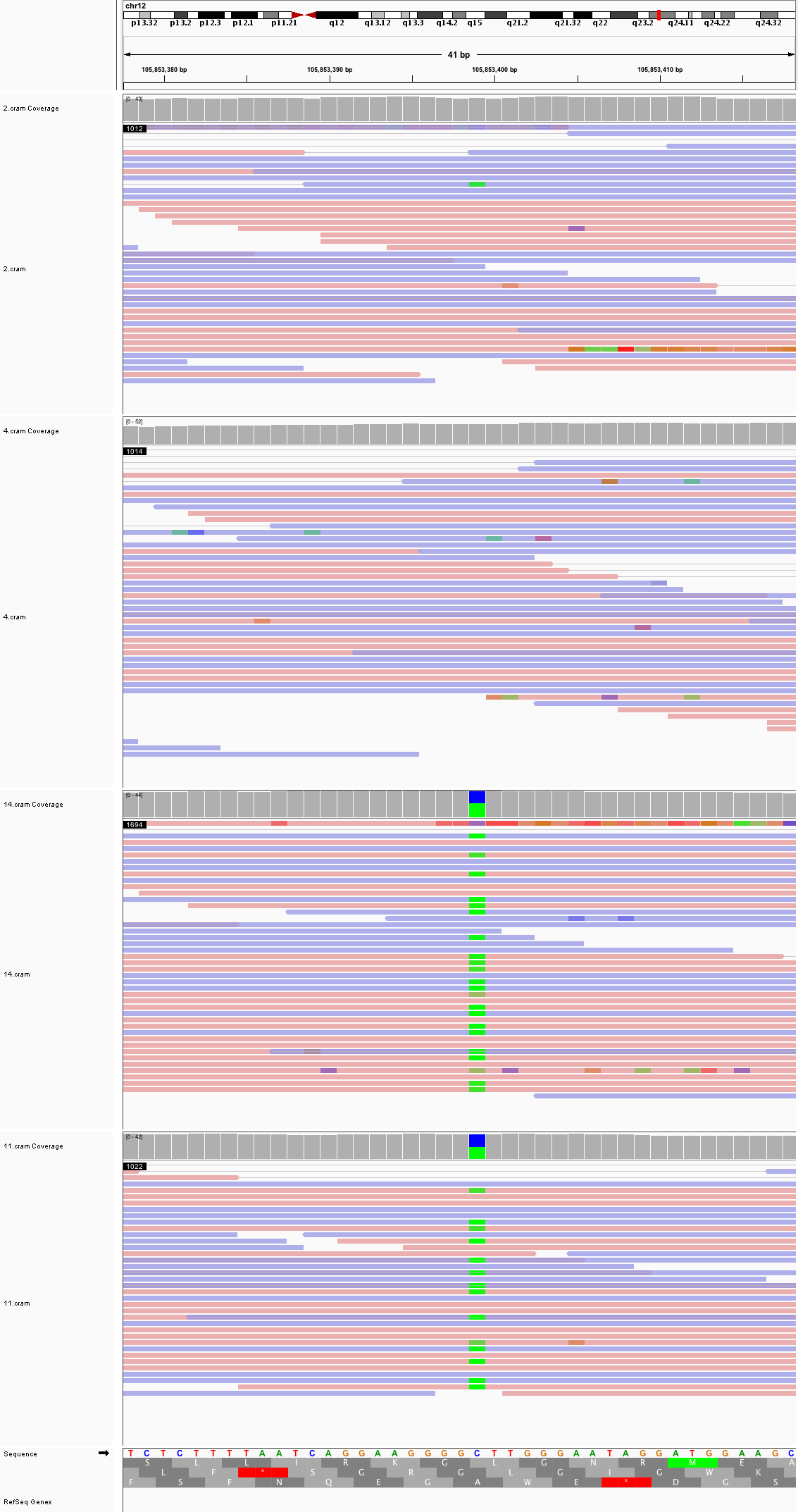

### chr12_109,920,273_109,920,313.png

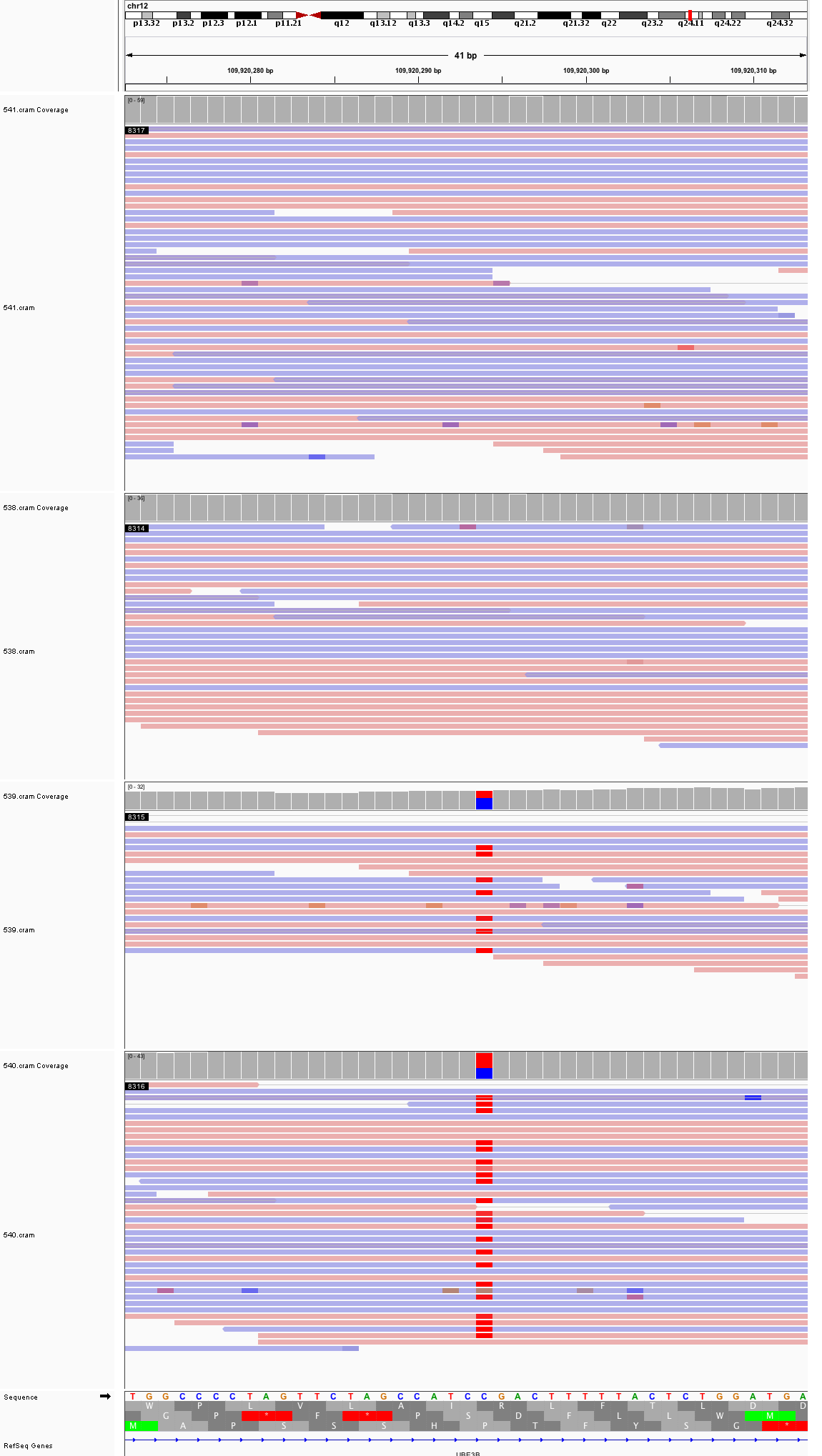

### chr13_23,450,122_23,450,162.png

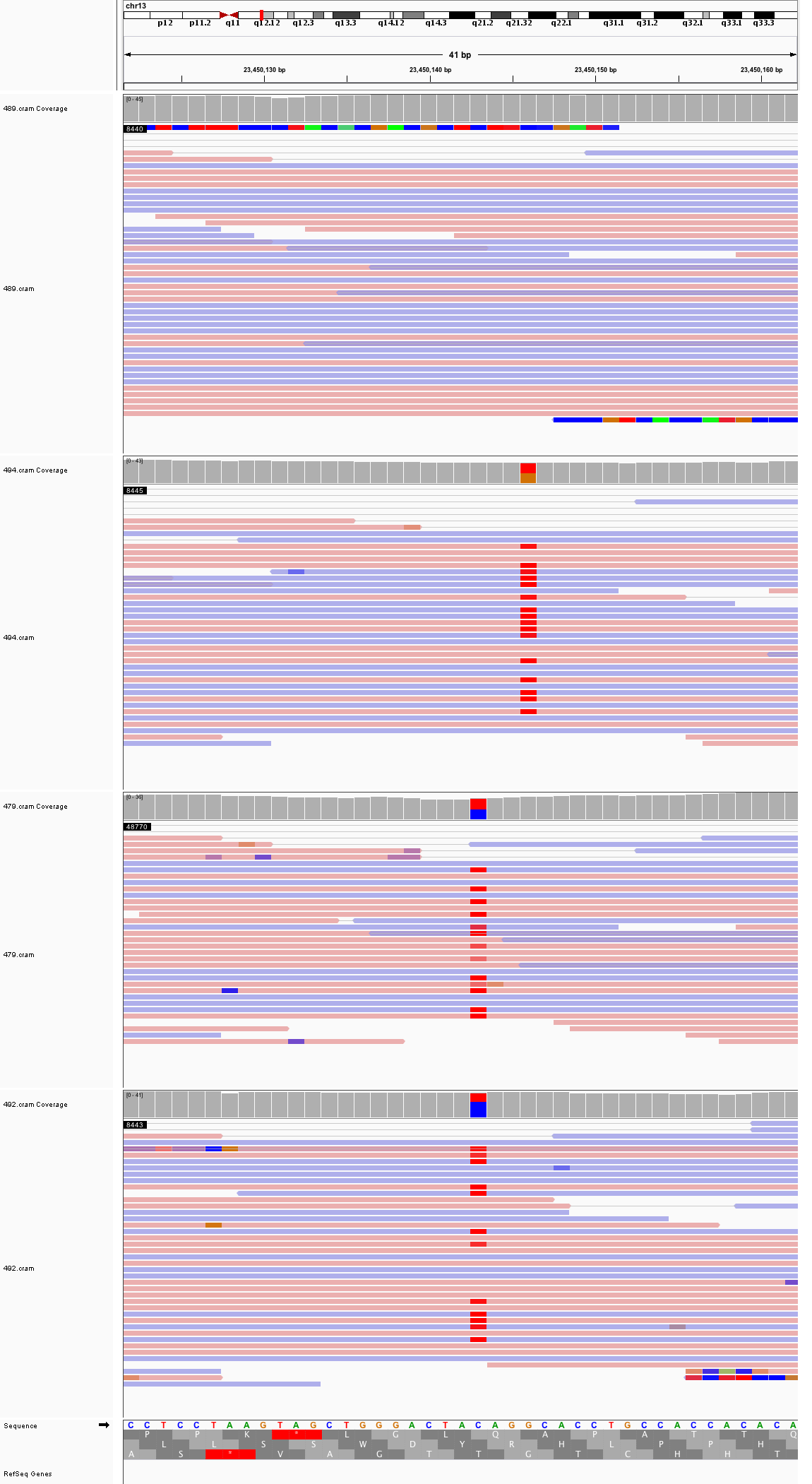

### chr13_84,539,010_84,539,050.png

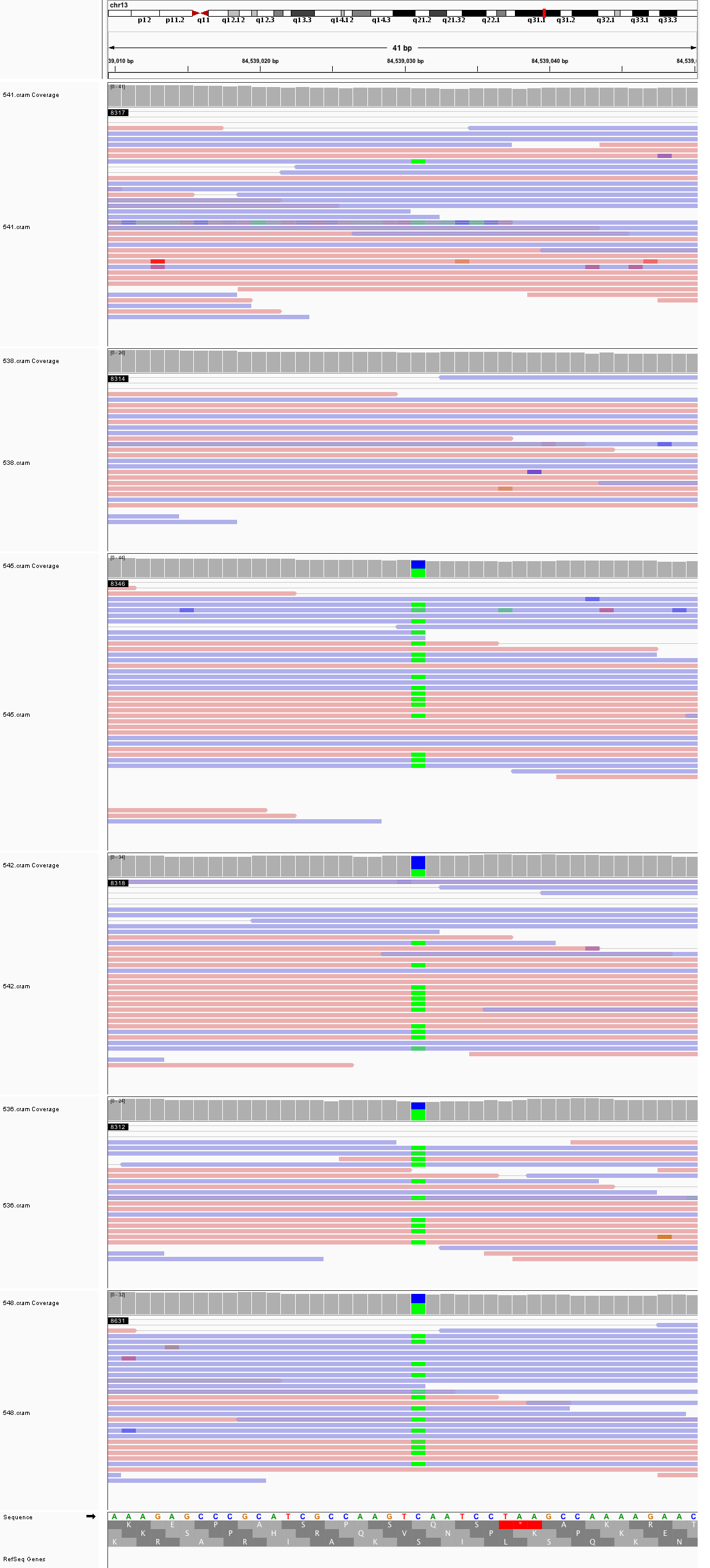

### chr14_26,835,787_26,835,827.png

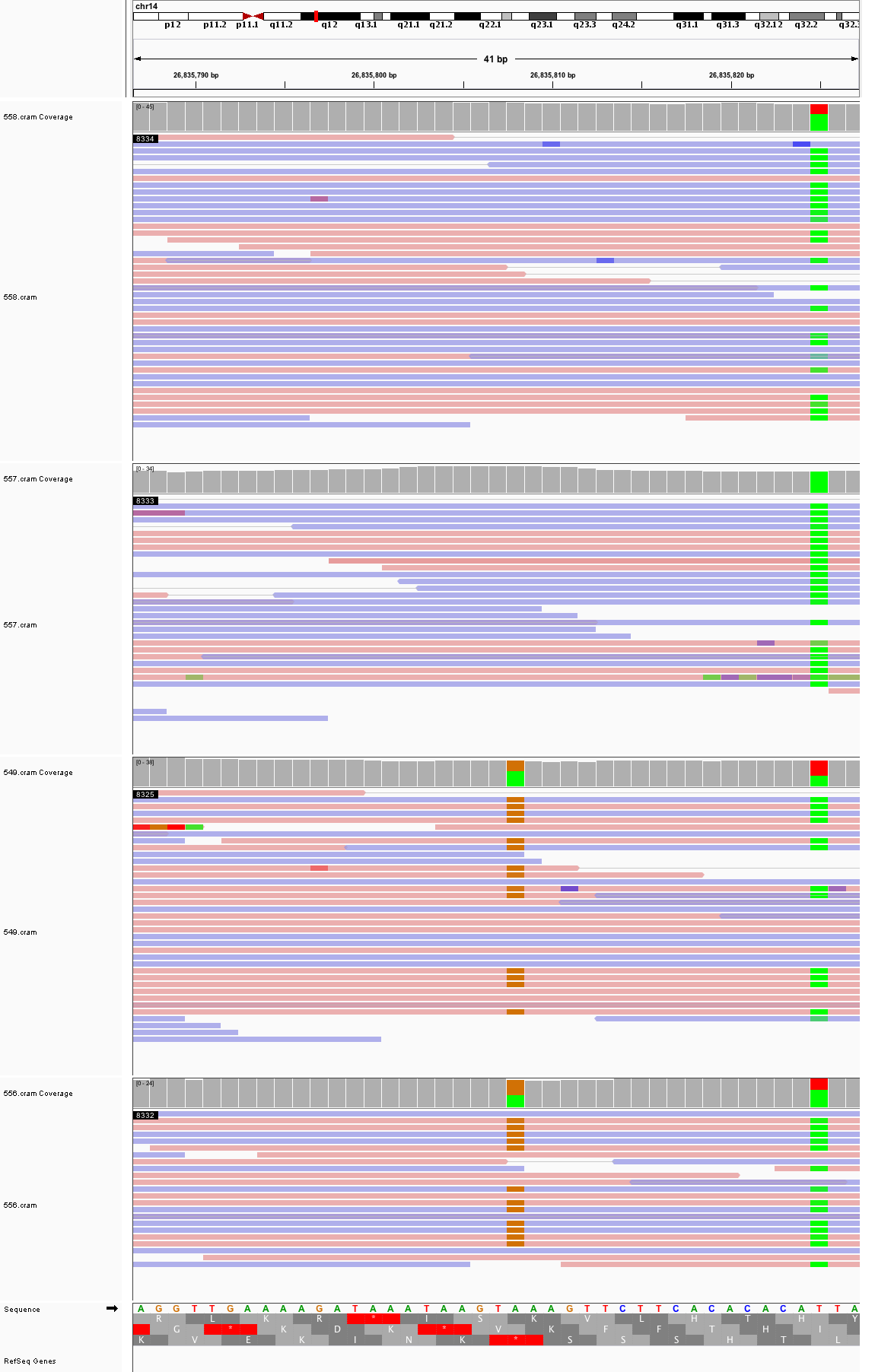

### chr14_33,223,740_33,223,780.png

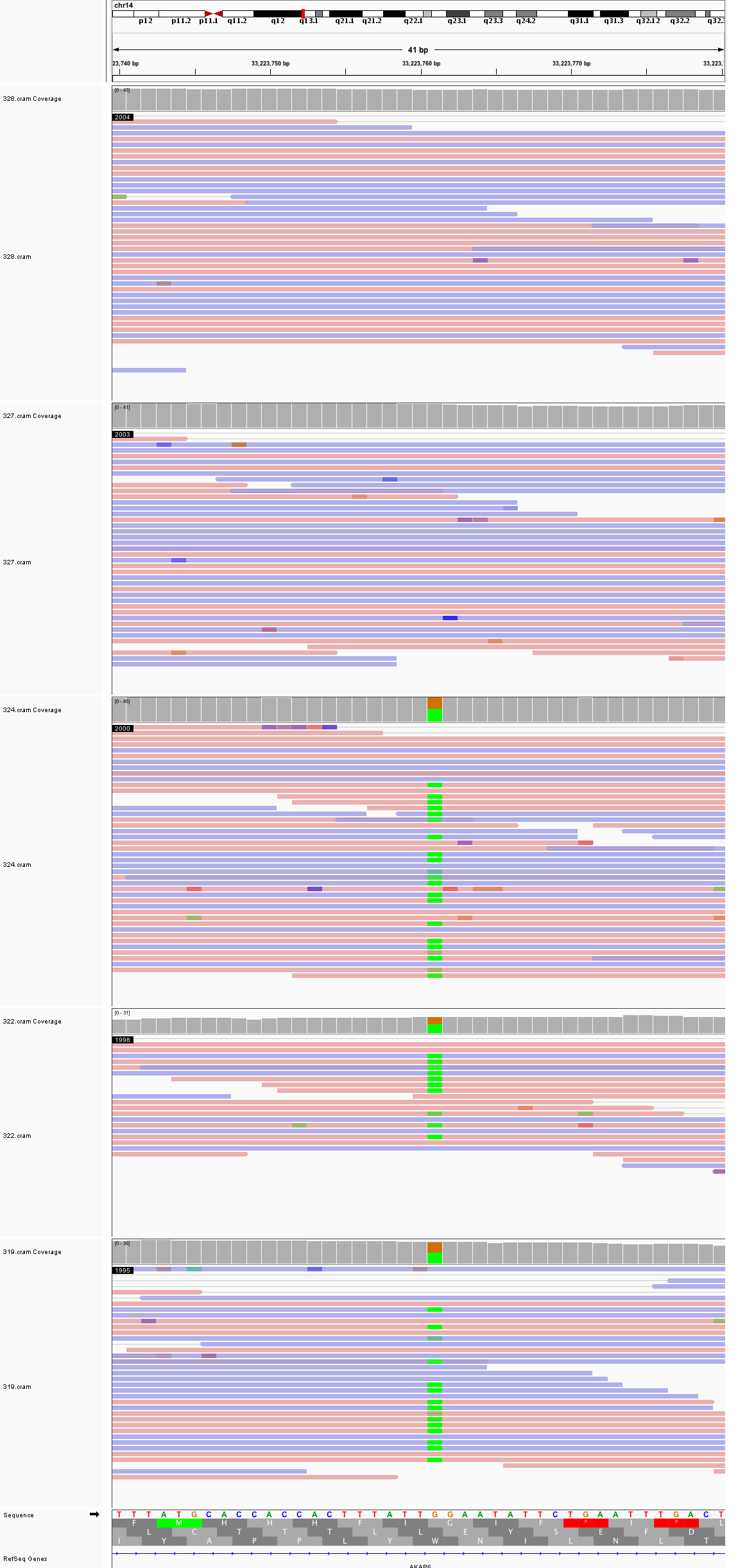

### chr16_32,977,446_32,977,486.png

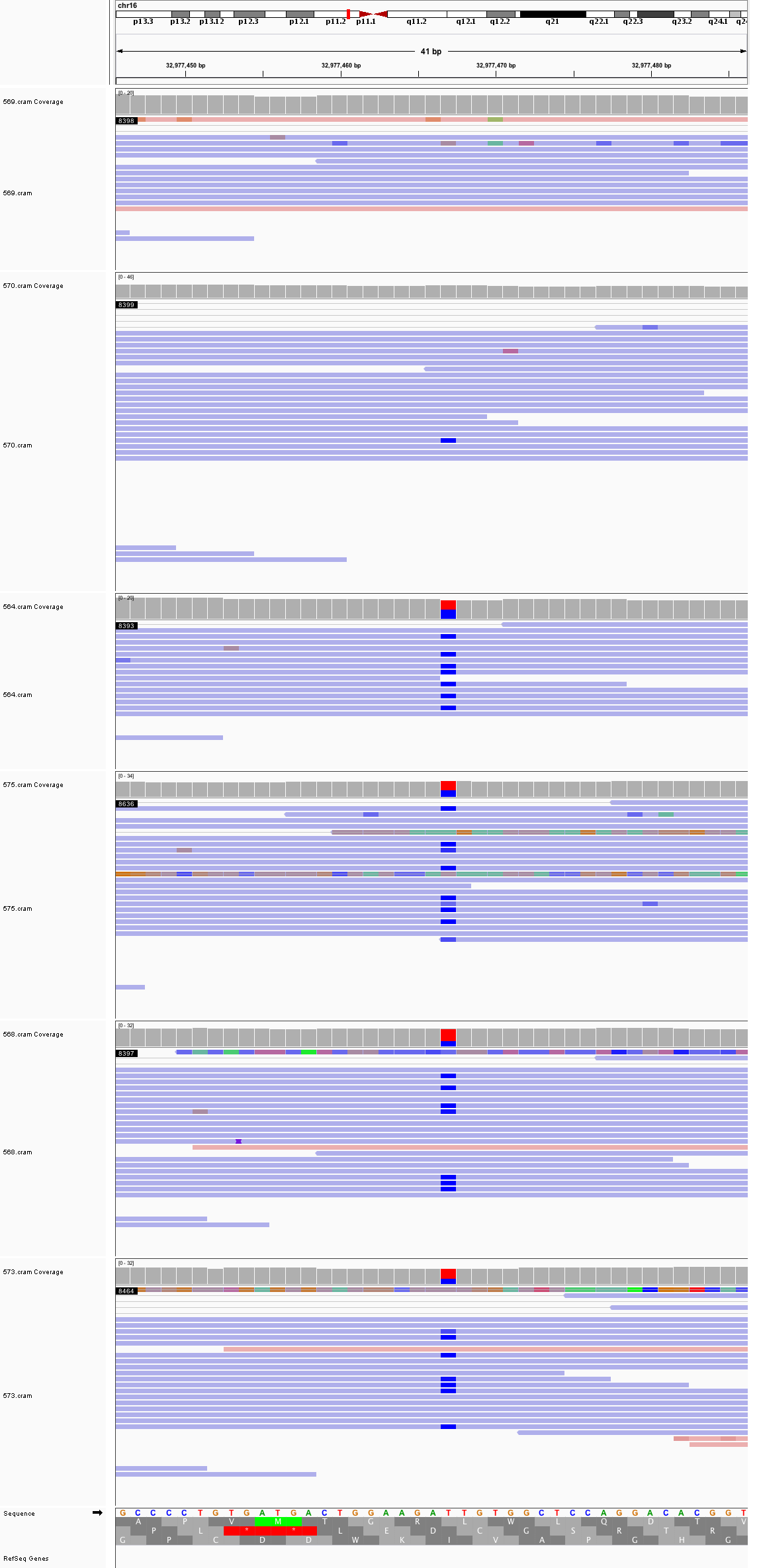

### chr16_54,751,663_54,751,703.png

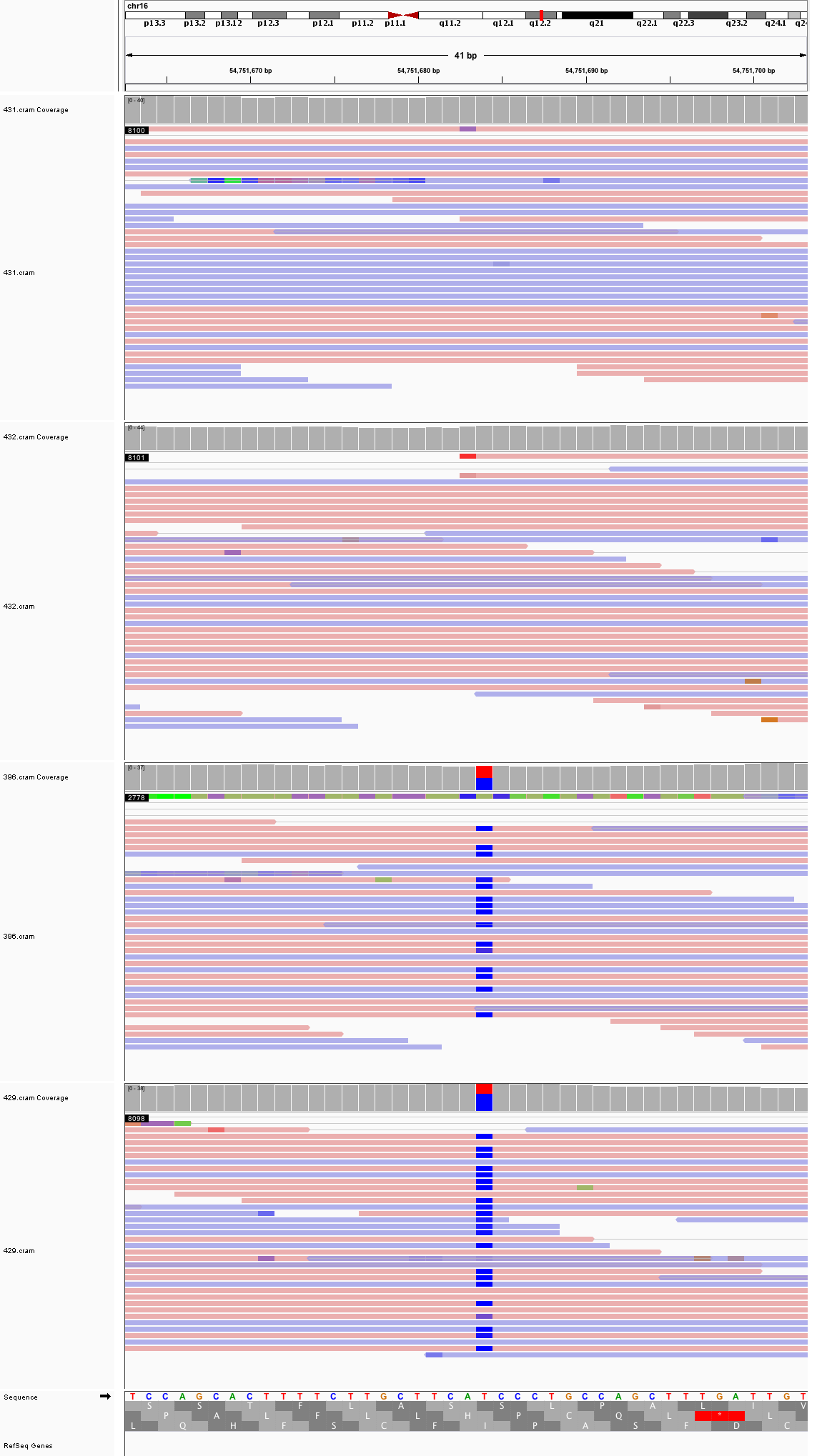

### chr16_55,444,225_55,444,265.png

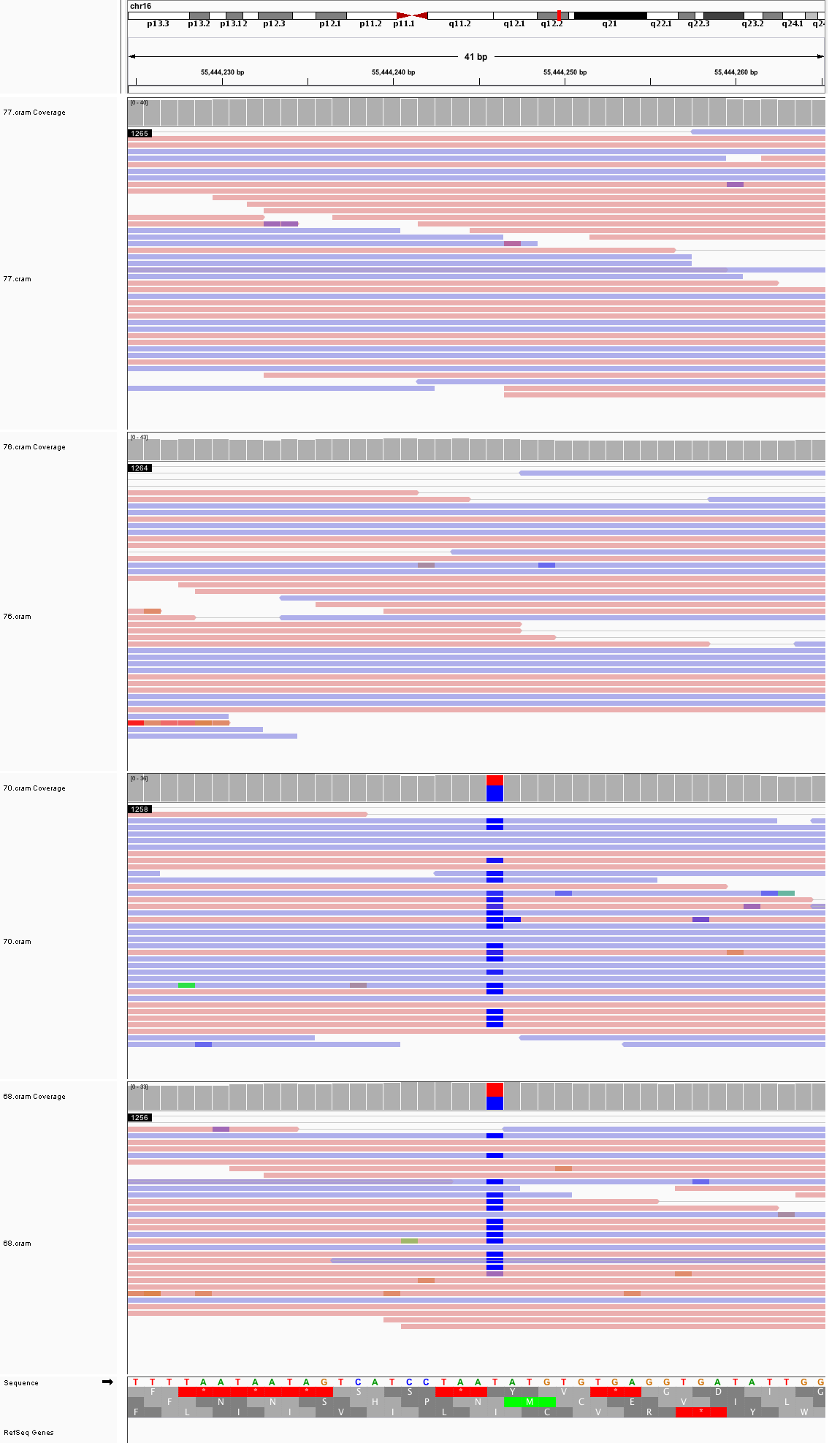

### chr16_66,834_66,874.png

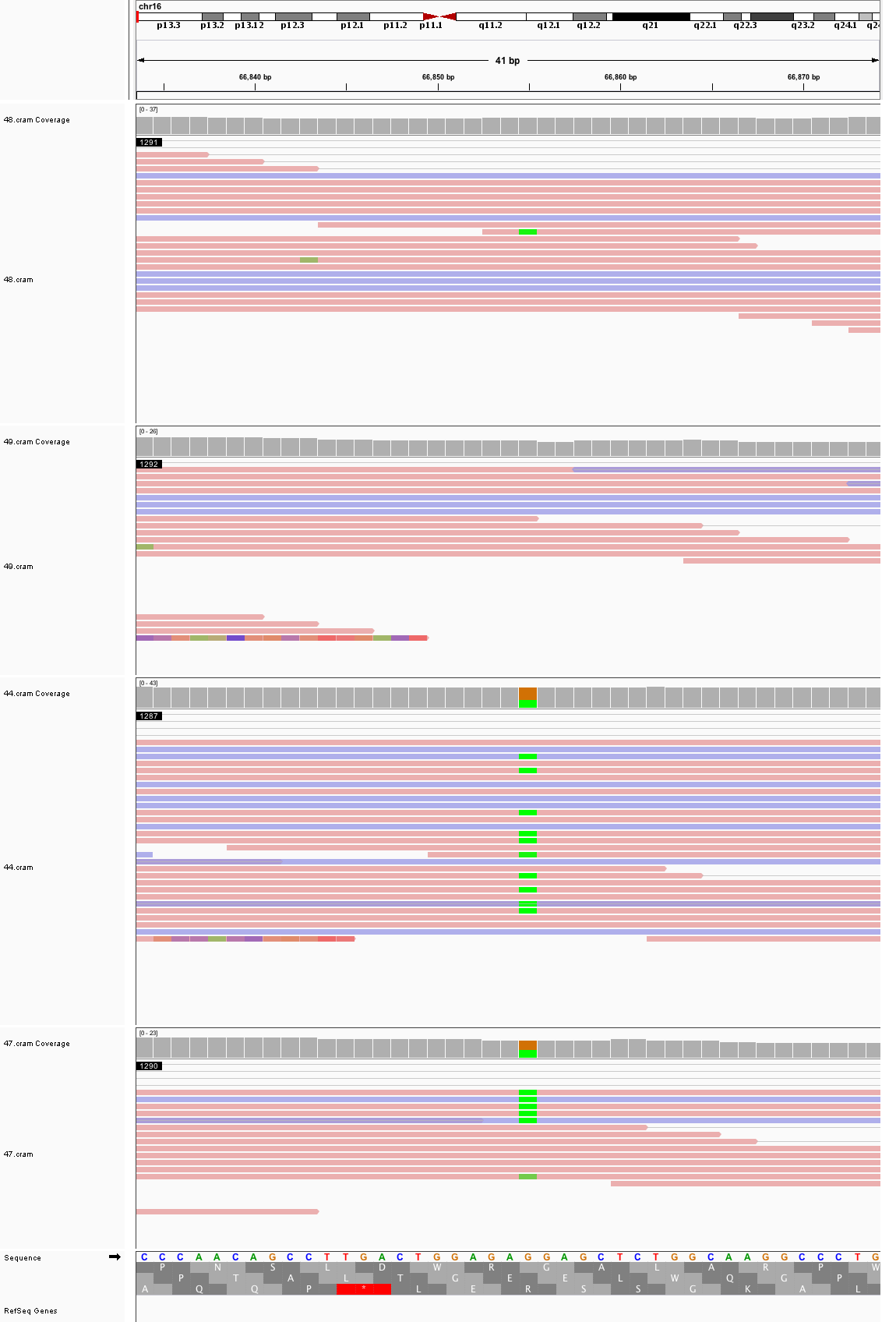

### chr16_76,760,763_76,760,803.png

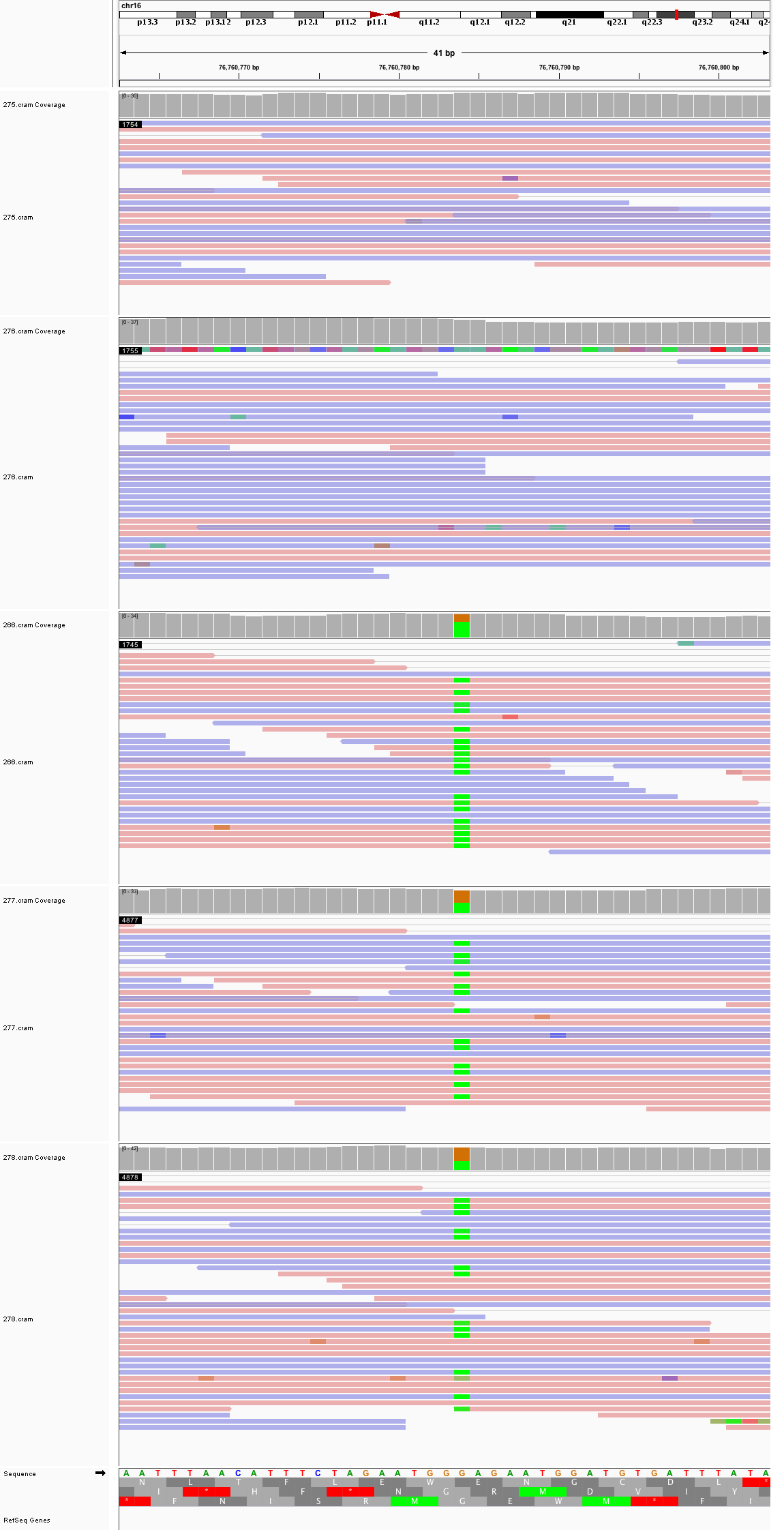

### chr17_4,644,349_4,644,389.png

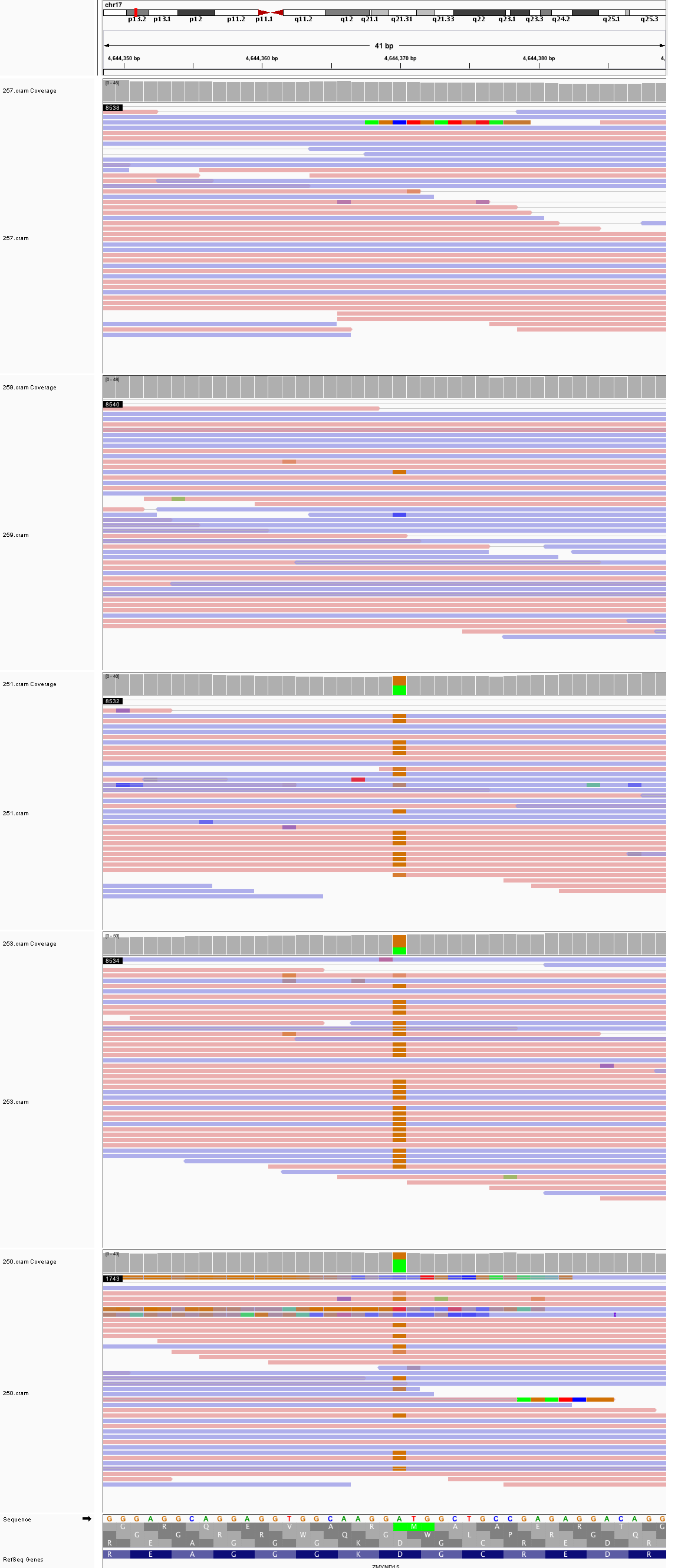

### chr17_45,709,182_45,709,222.png

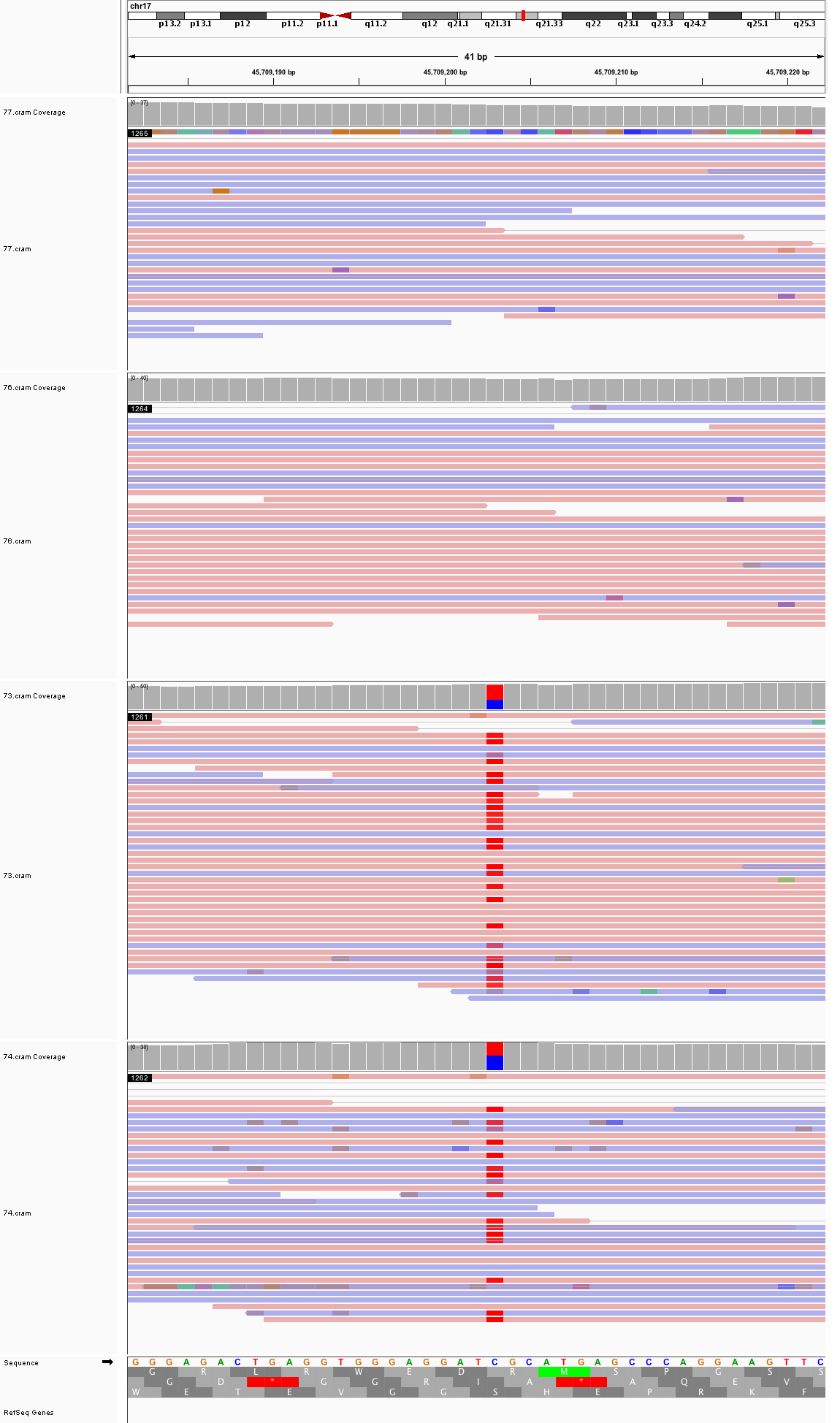

### chr17_77,353,918_77,353,958.png

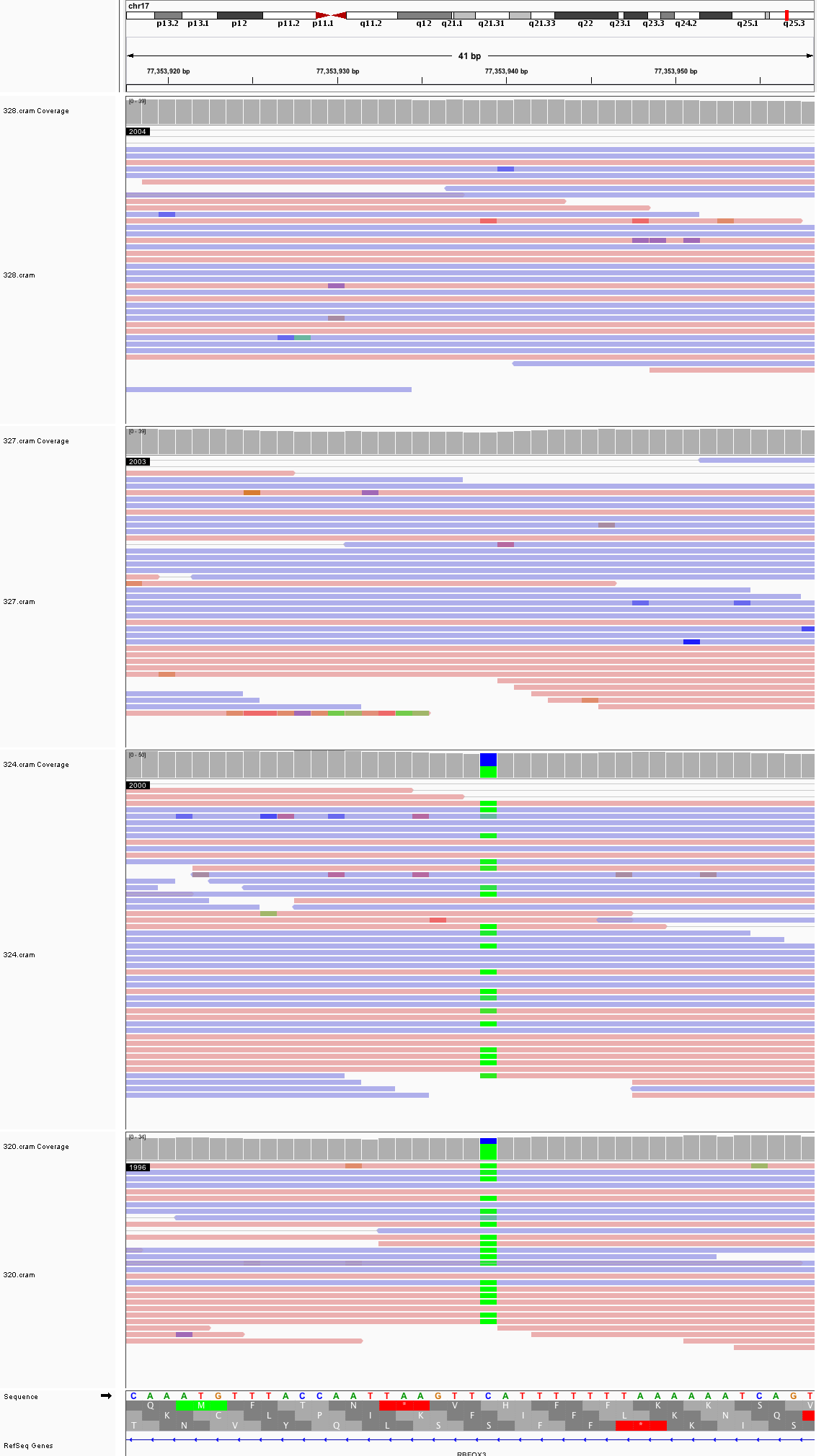

### chr17_78,167,742_78,167,782.png

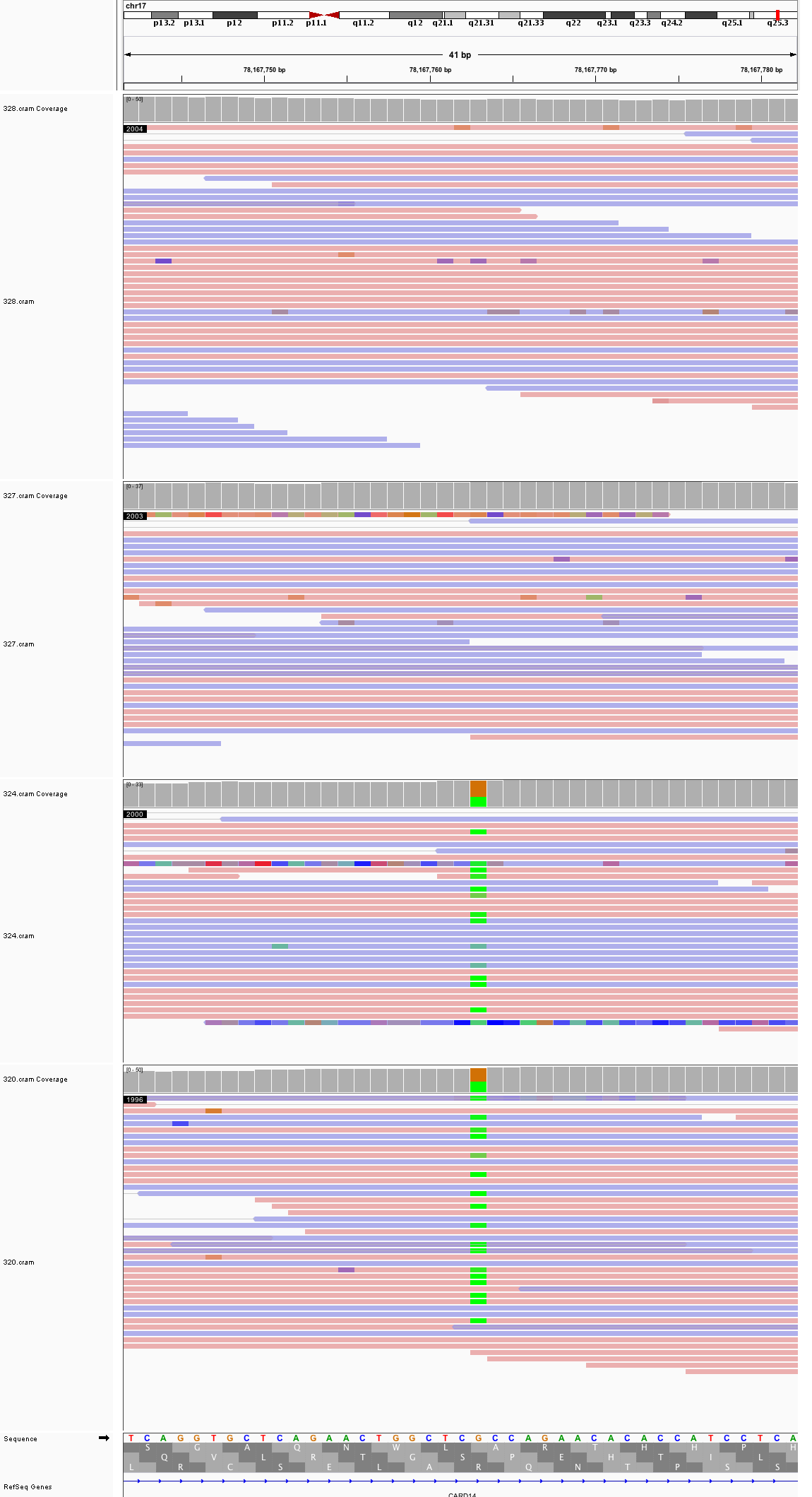

### chr17_78,798,510_78,798,550.png

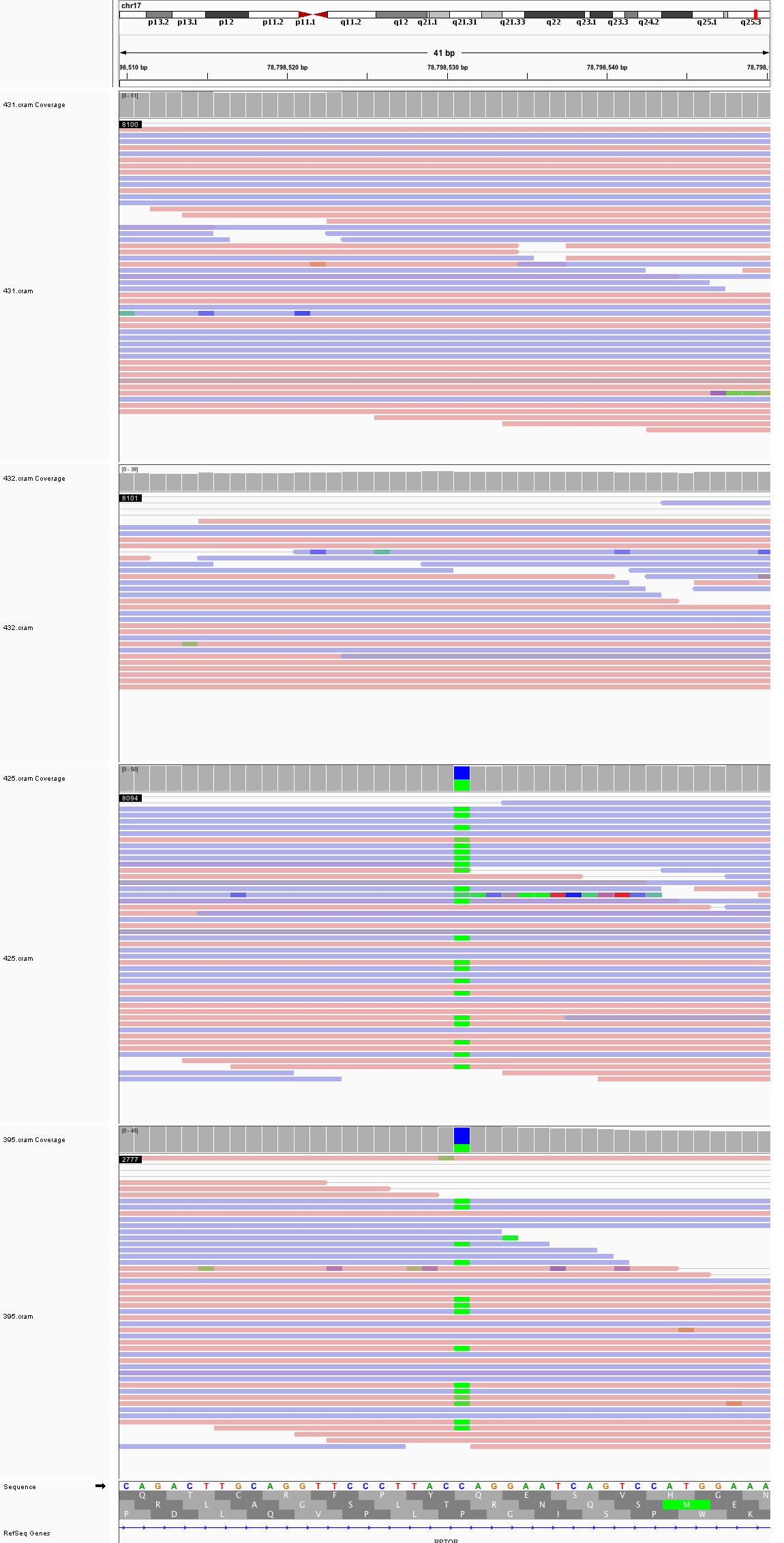

### chr18_4,682,771_4,682,811.png

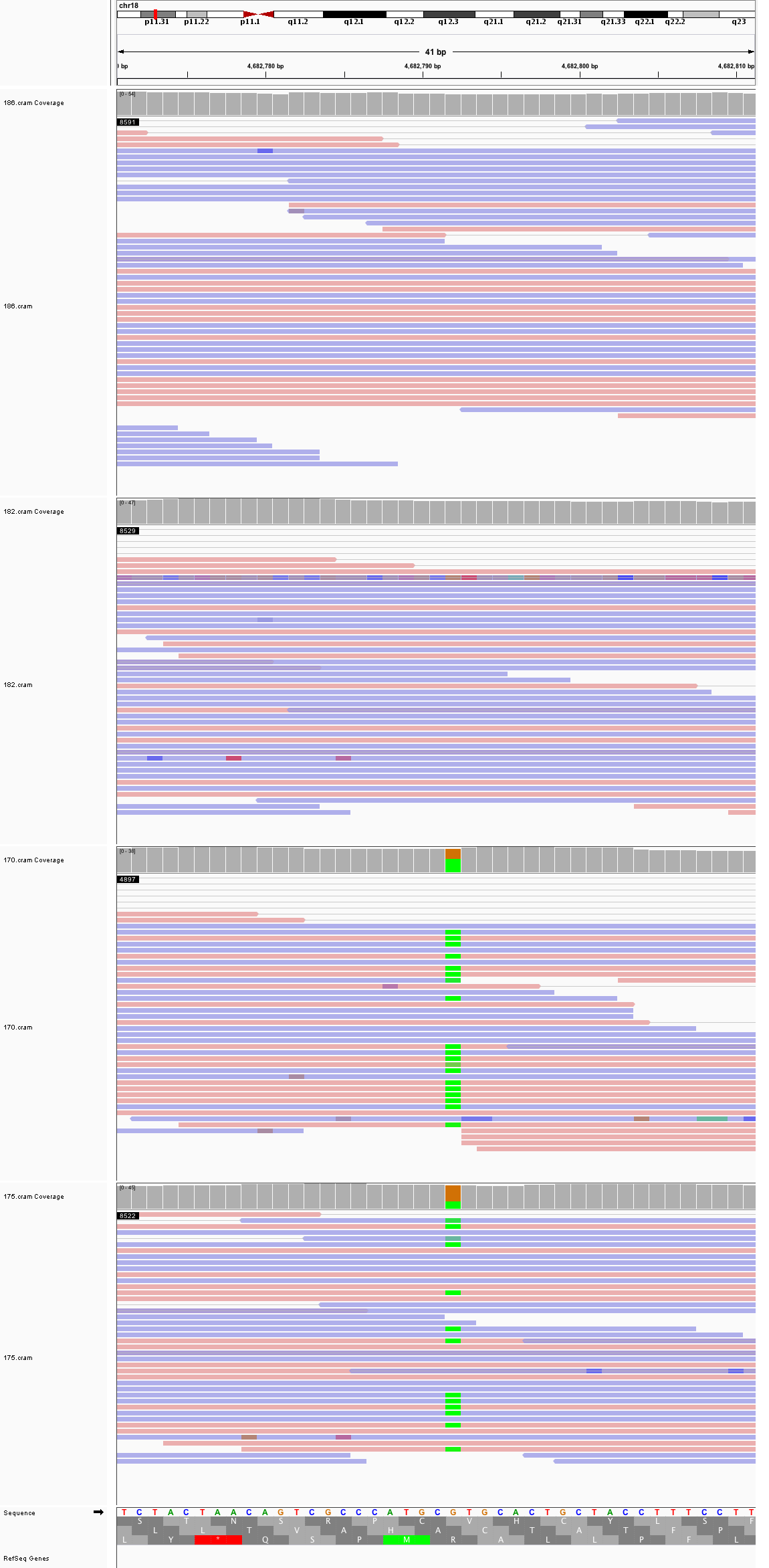

### chr18_7,893,239_7,893,279.png

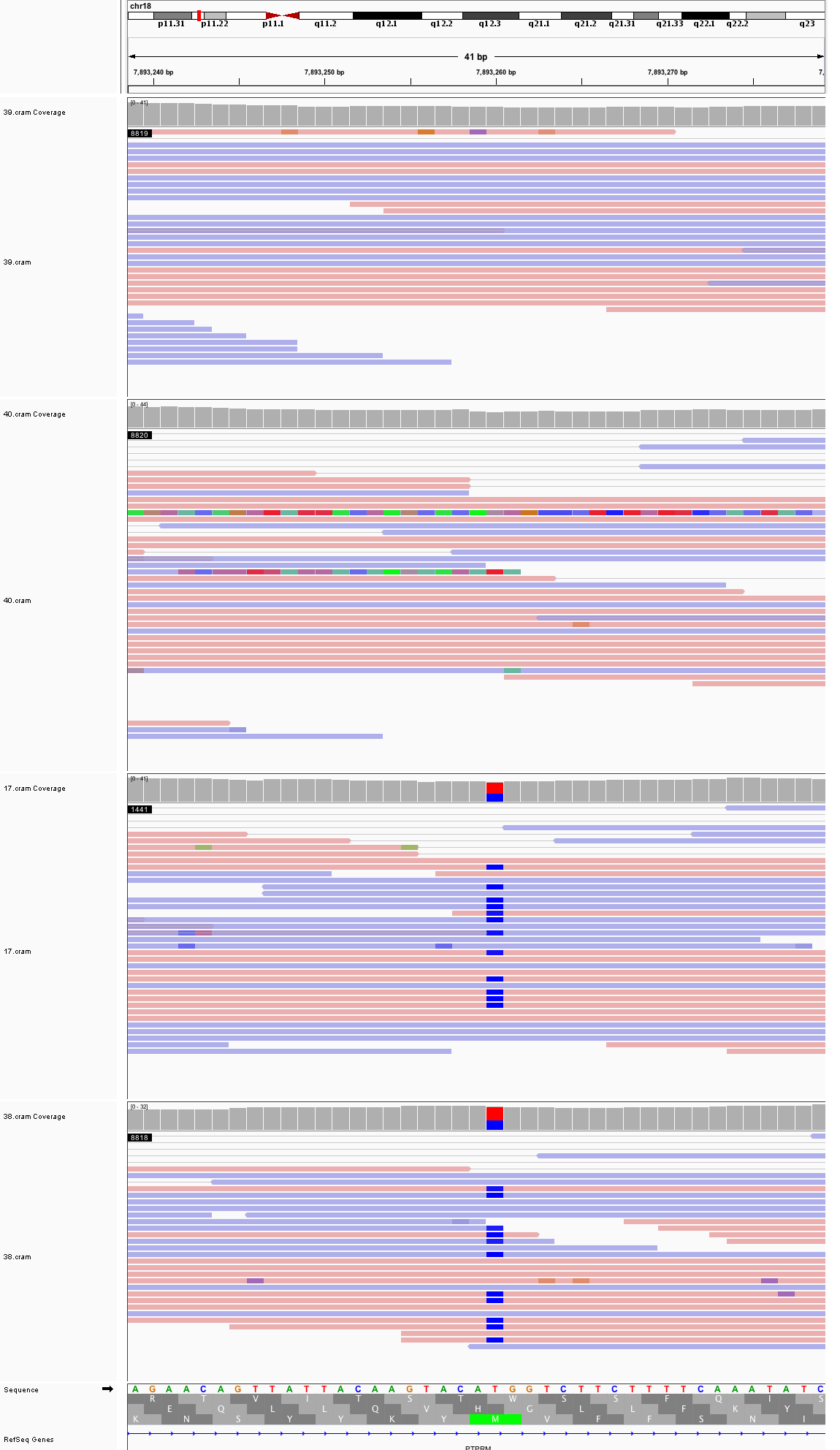

### chr18_27,425,993_27,426,033.png

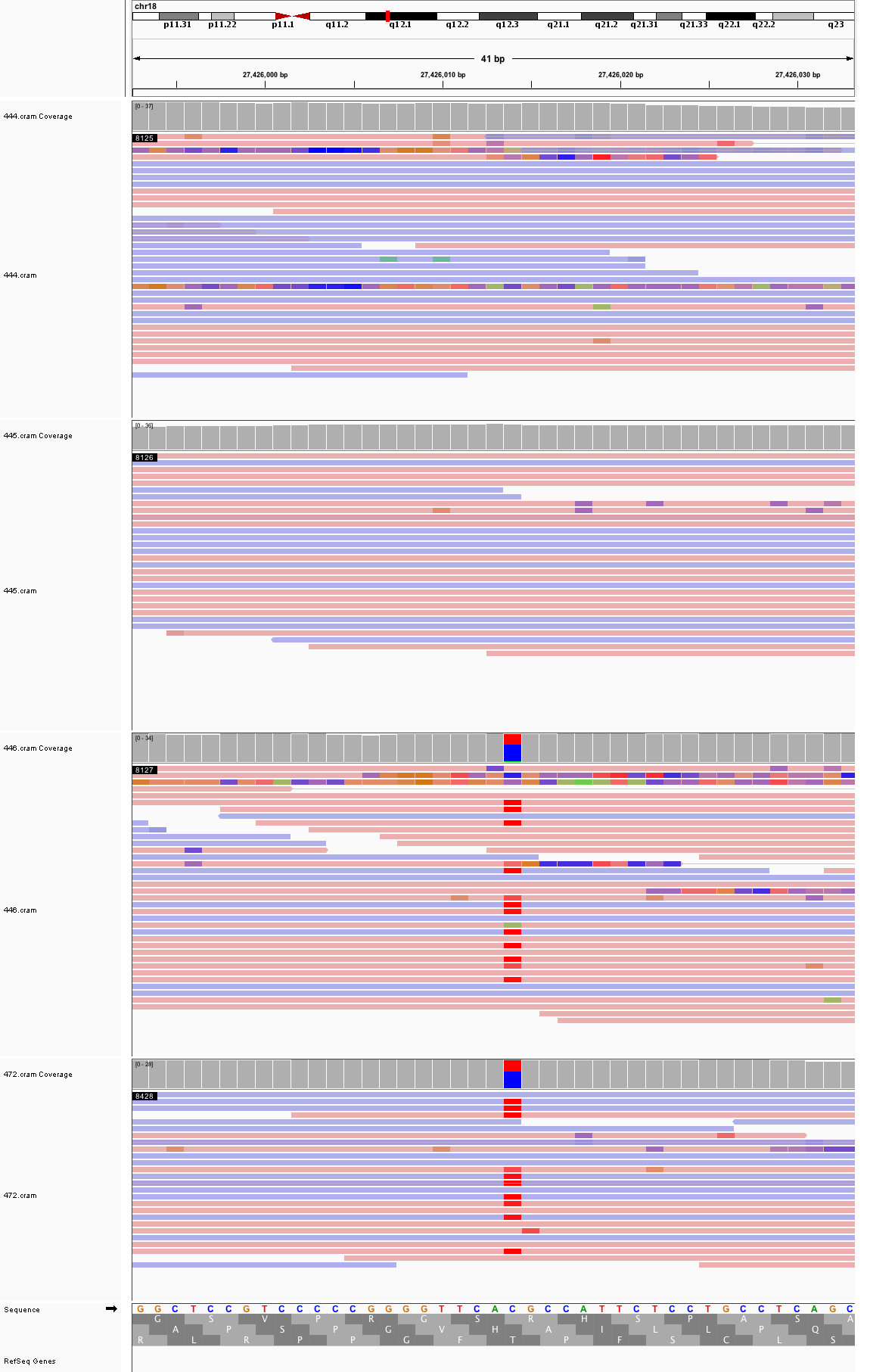

### chr18_43,703,777_43,703,817.png

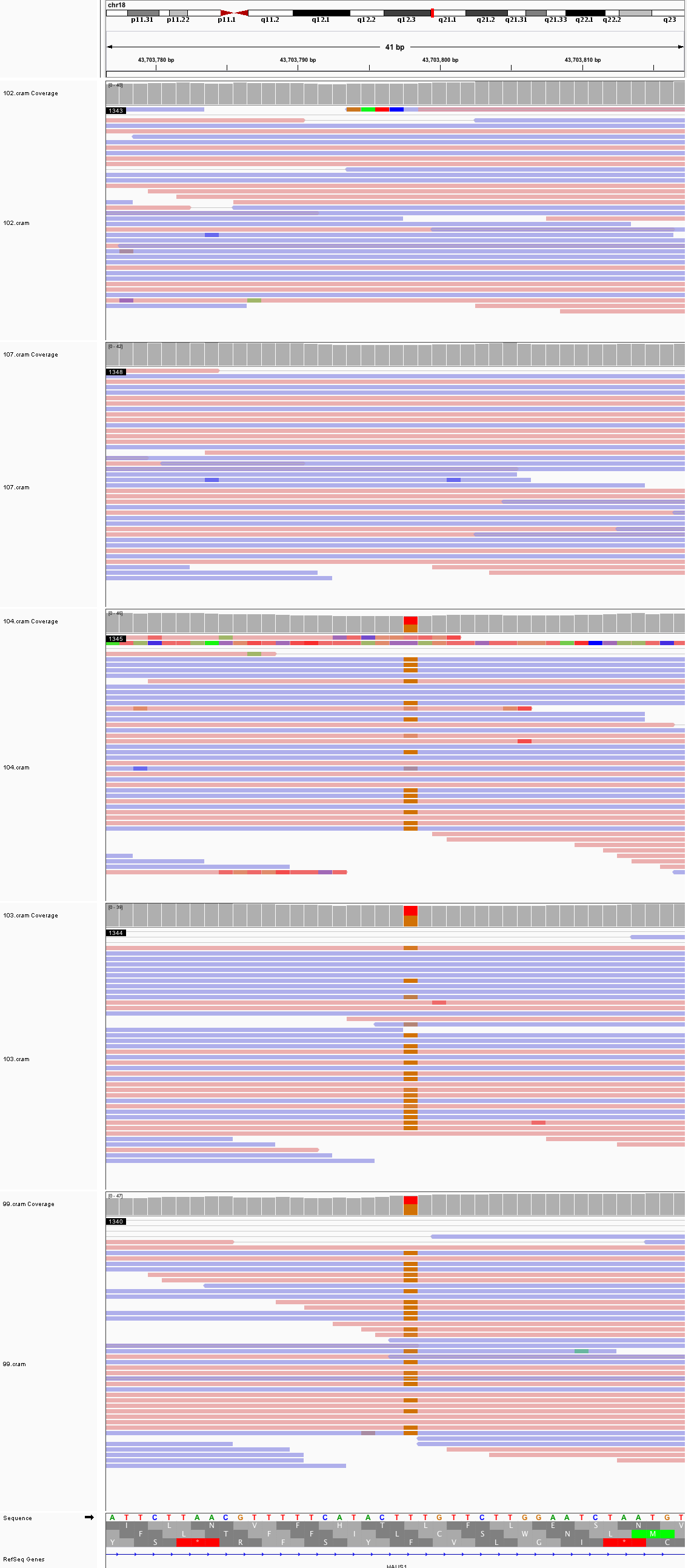

### chr19_7,945,606_7,945,646.png

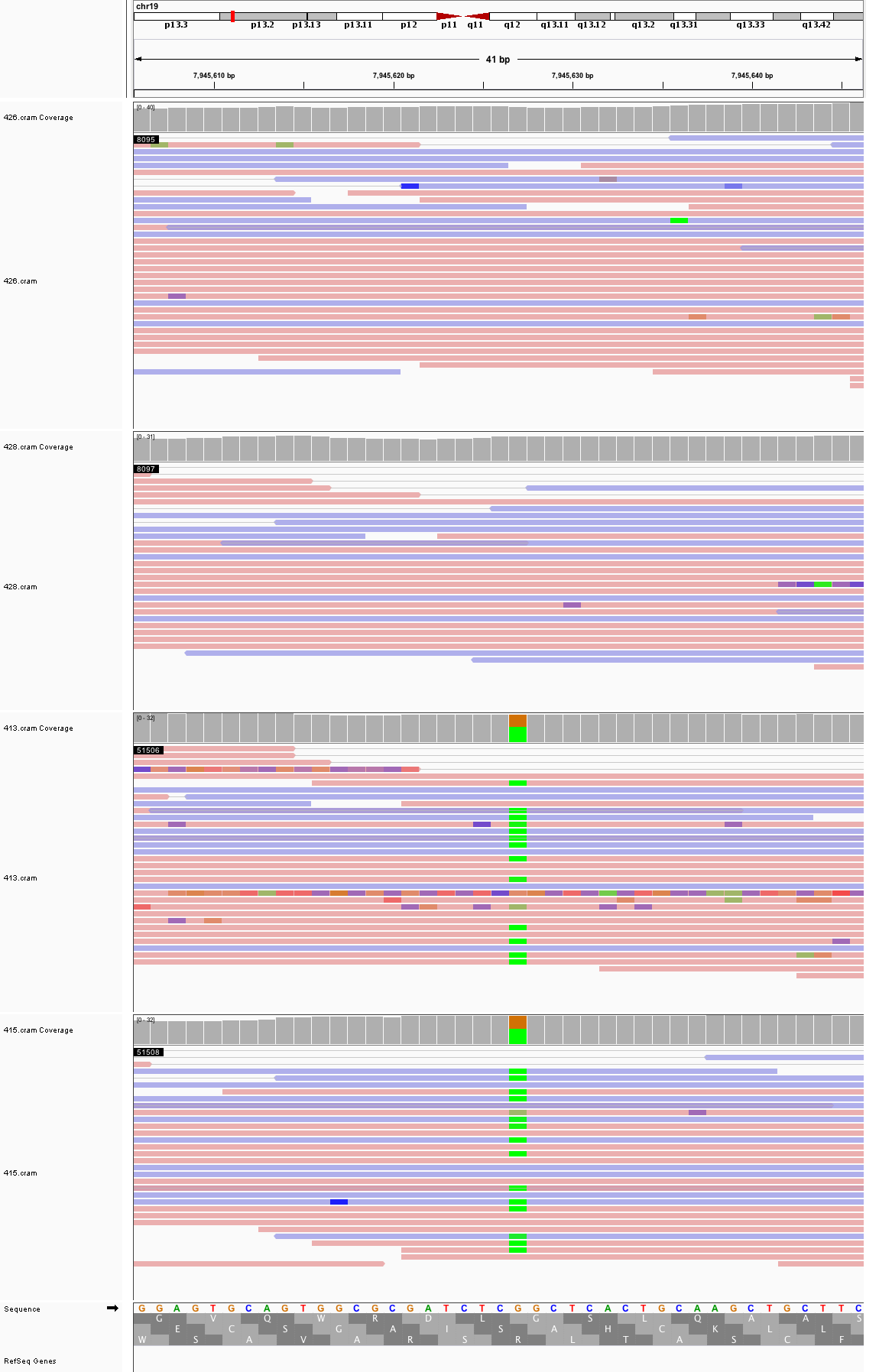

### chr19_52,865,352_52,865,392.png

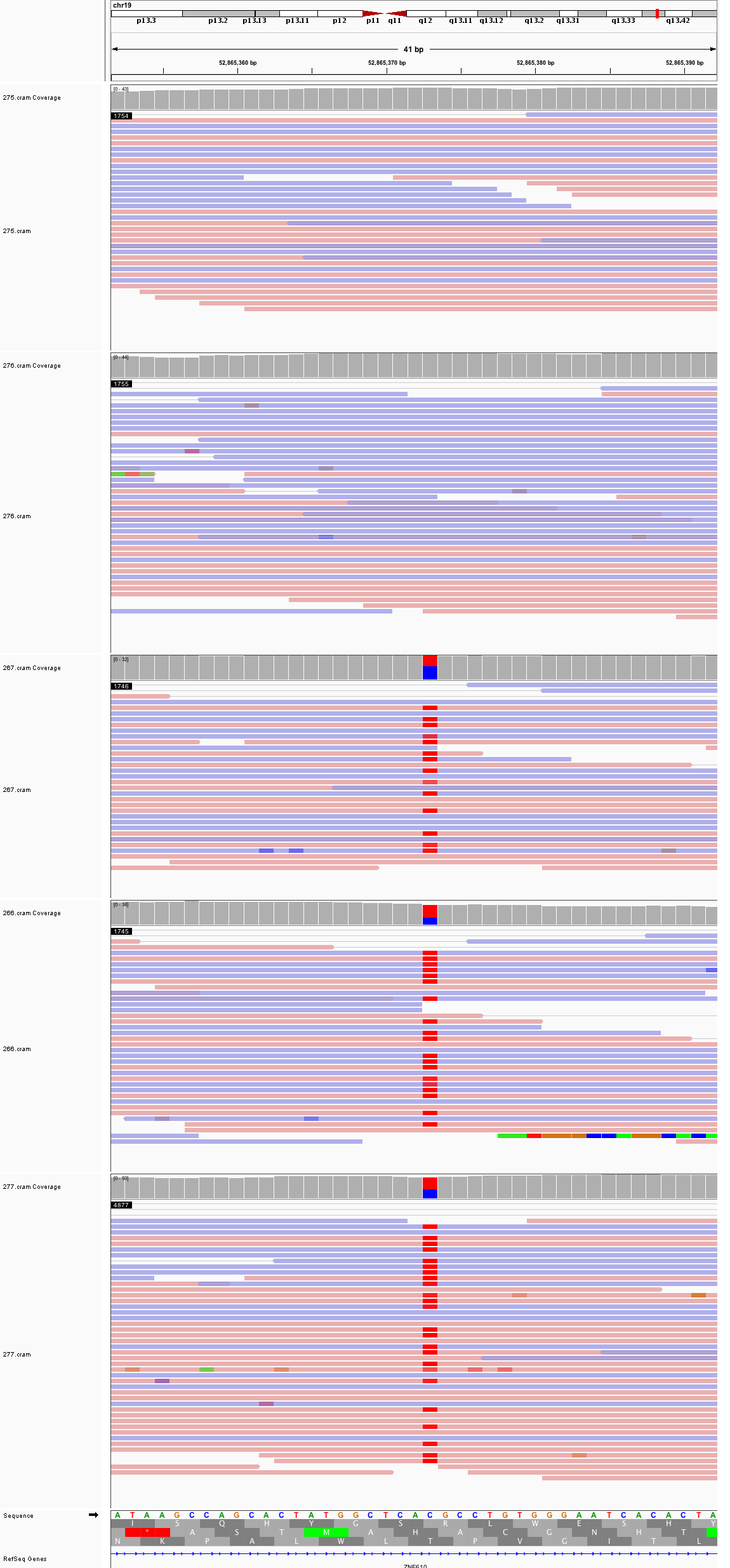

### chr20_7,067,683_7,067,723.png

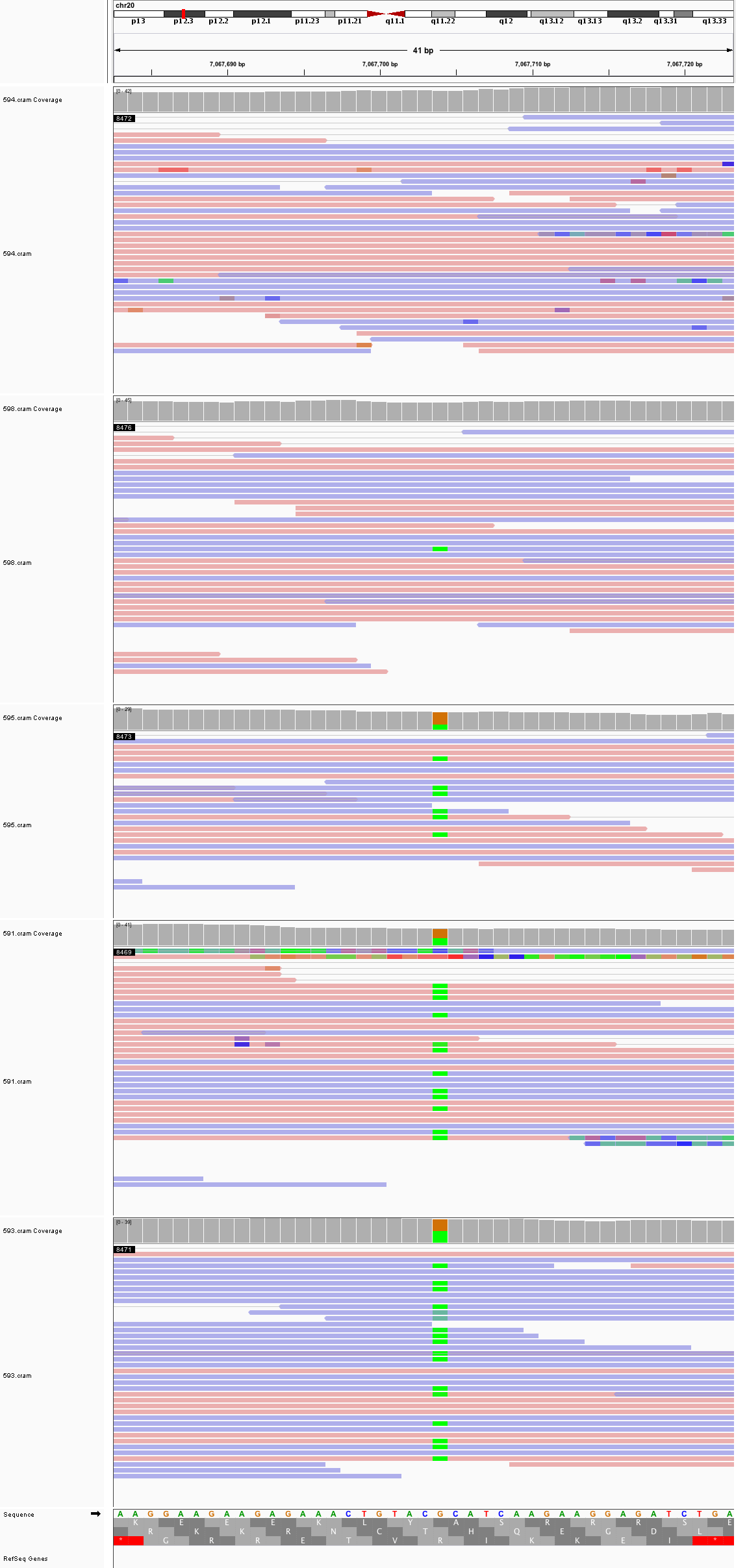

### chr20_62,477,685_62,477,725.png

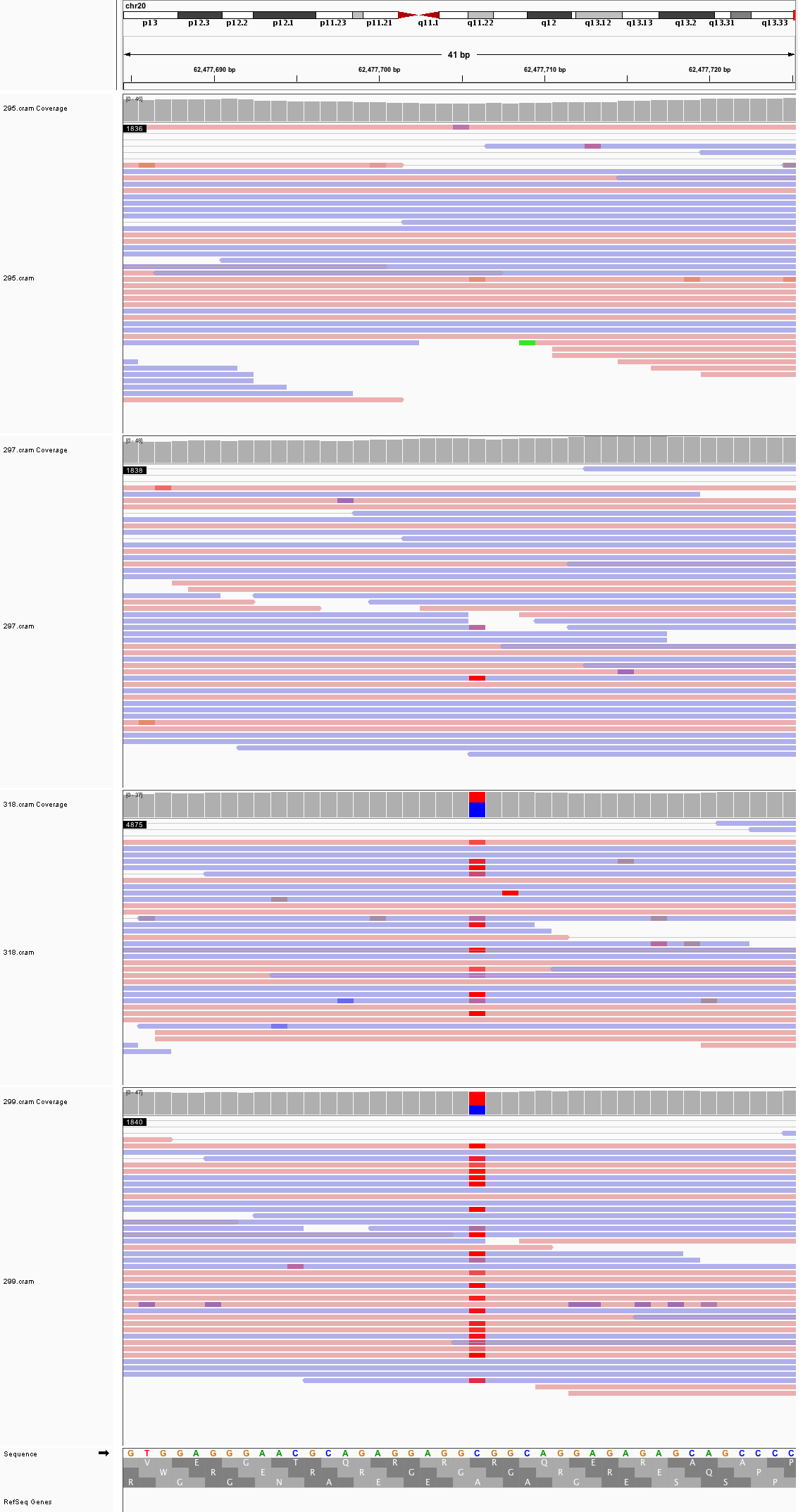

### chr21_24,143,545_24,143,585.png

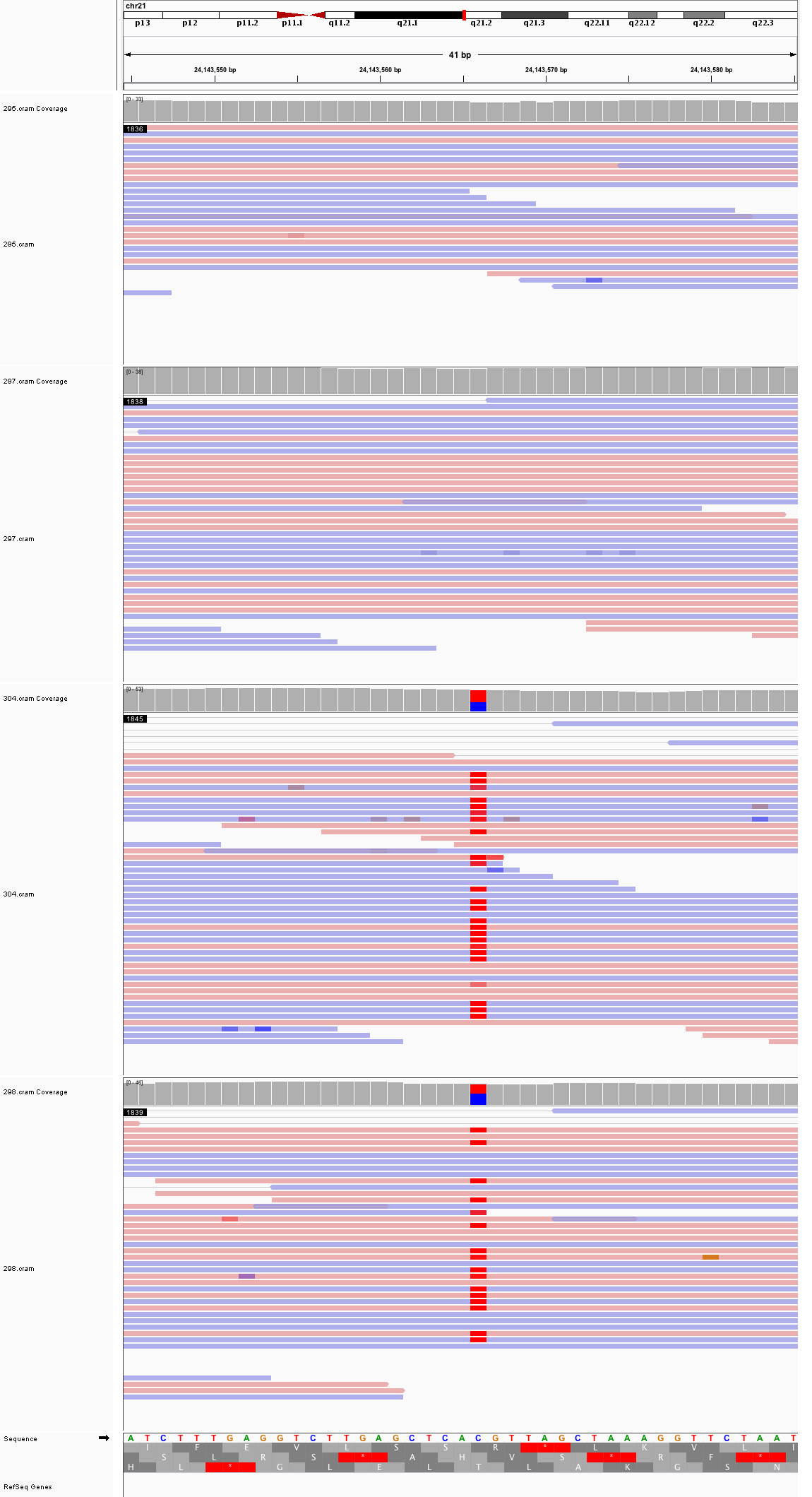

### chr22_28,571,649_28,571,689.png

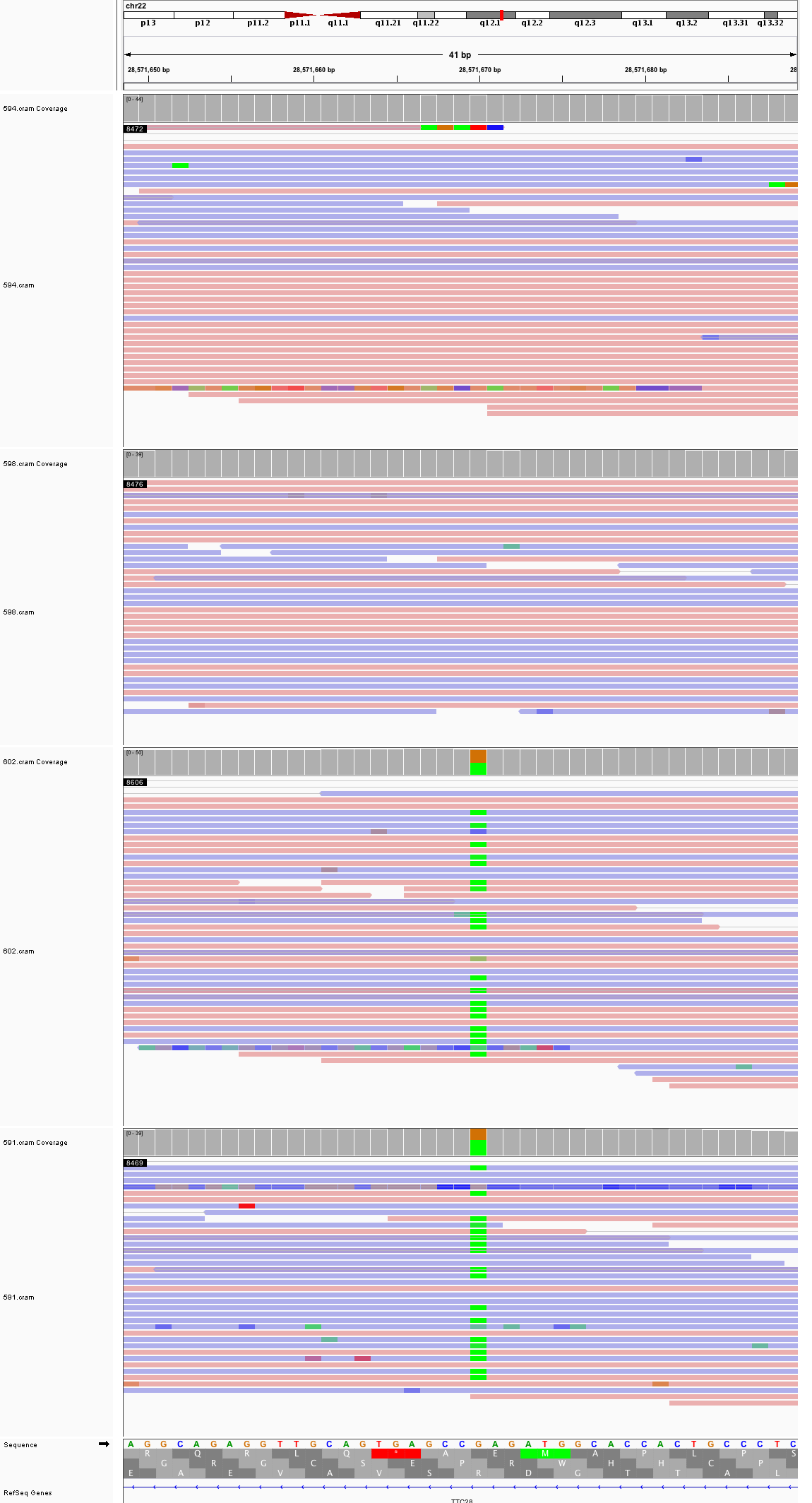

### chrX_11,432,195_11,432,235.png

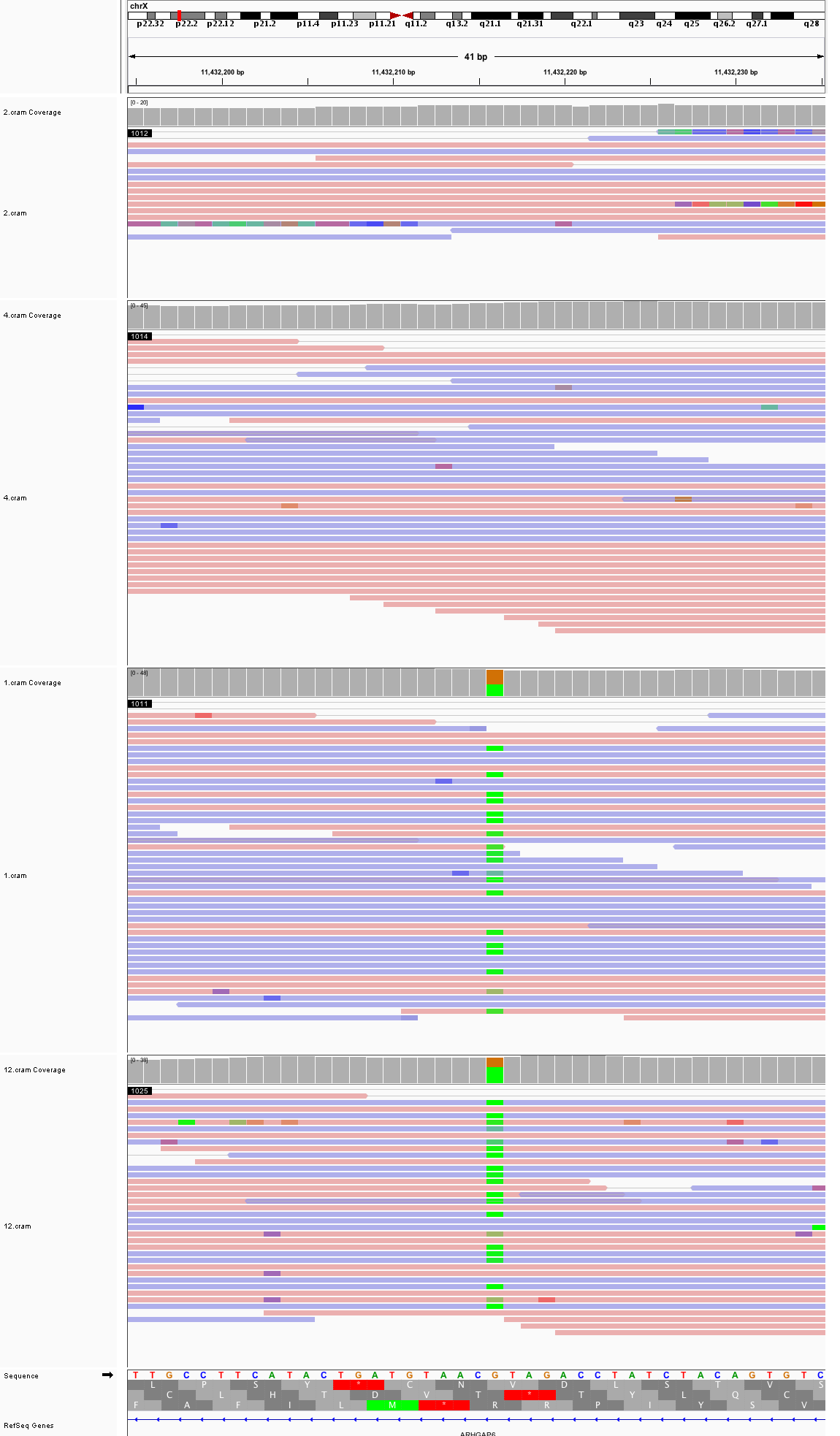

### chrX_24,562,081_24,562,121.png

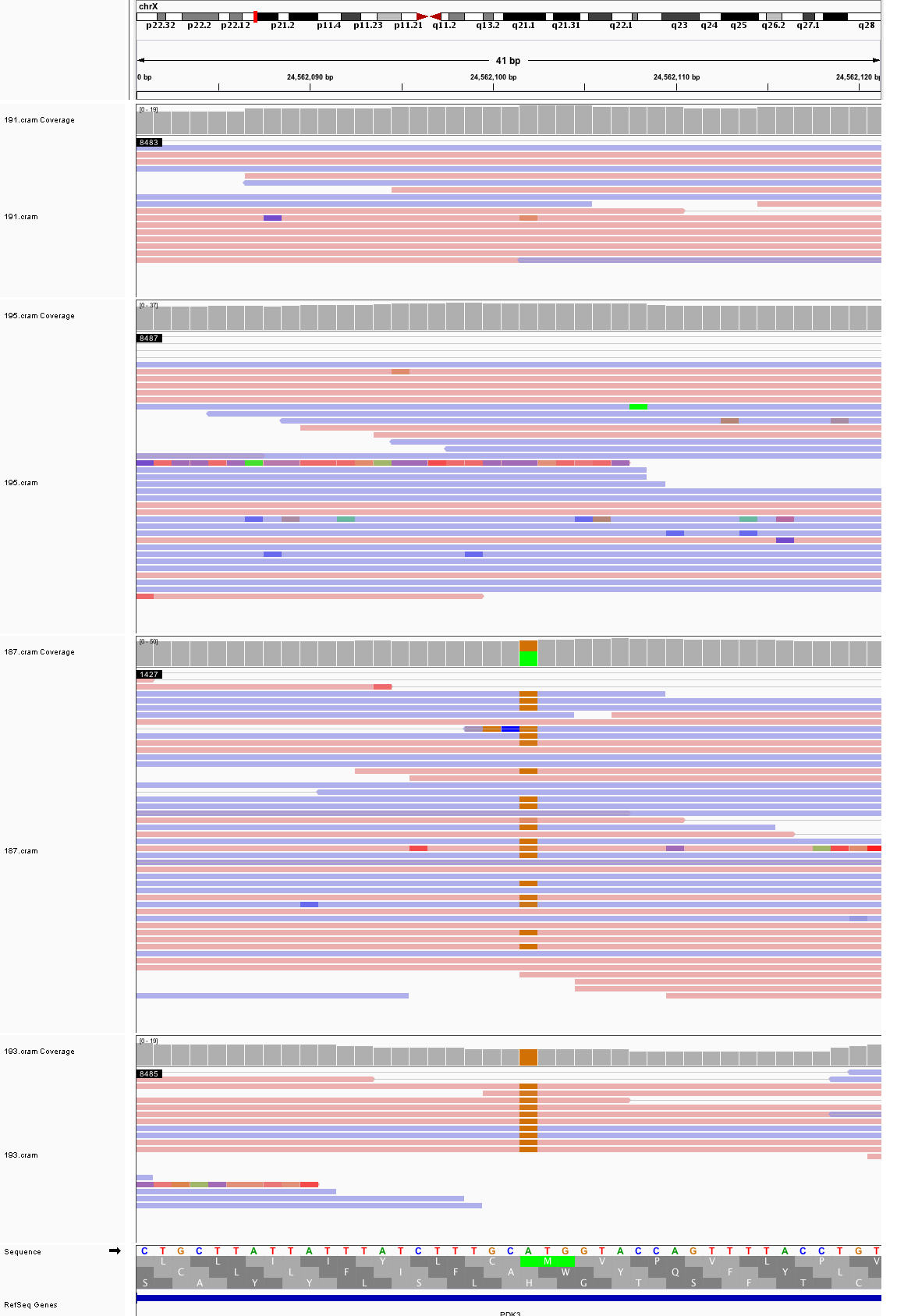

### chrX_52,741,615_52,741,655.png

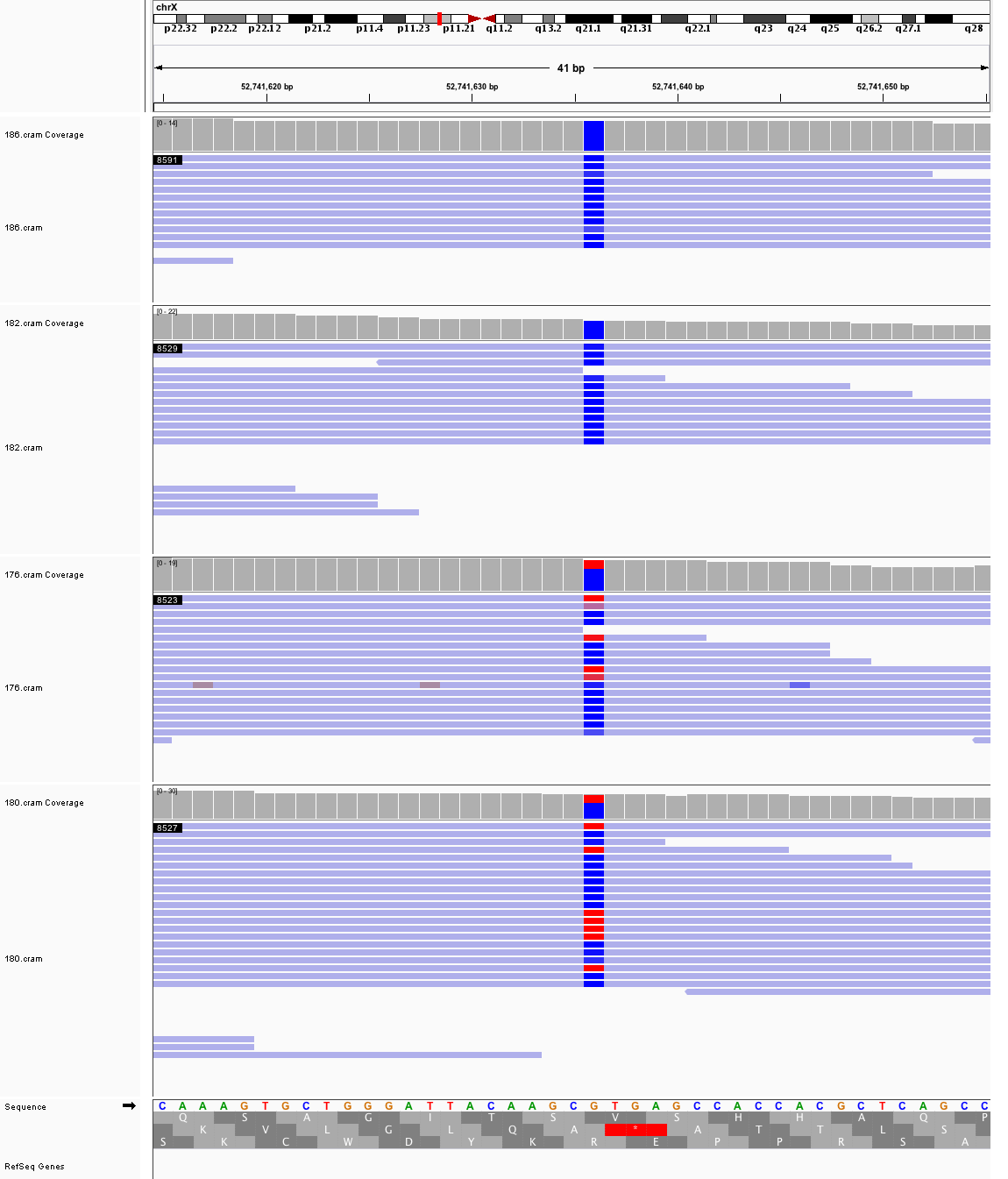

### chrX_84,907,715_84,907,755.png

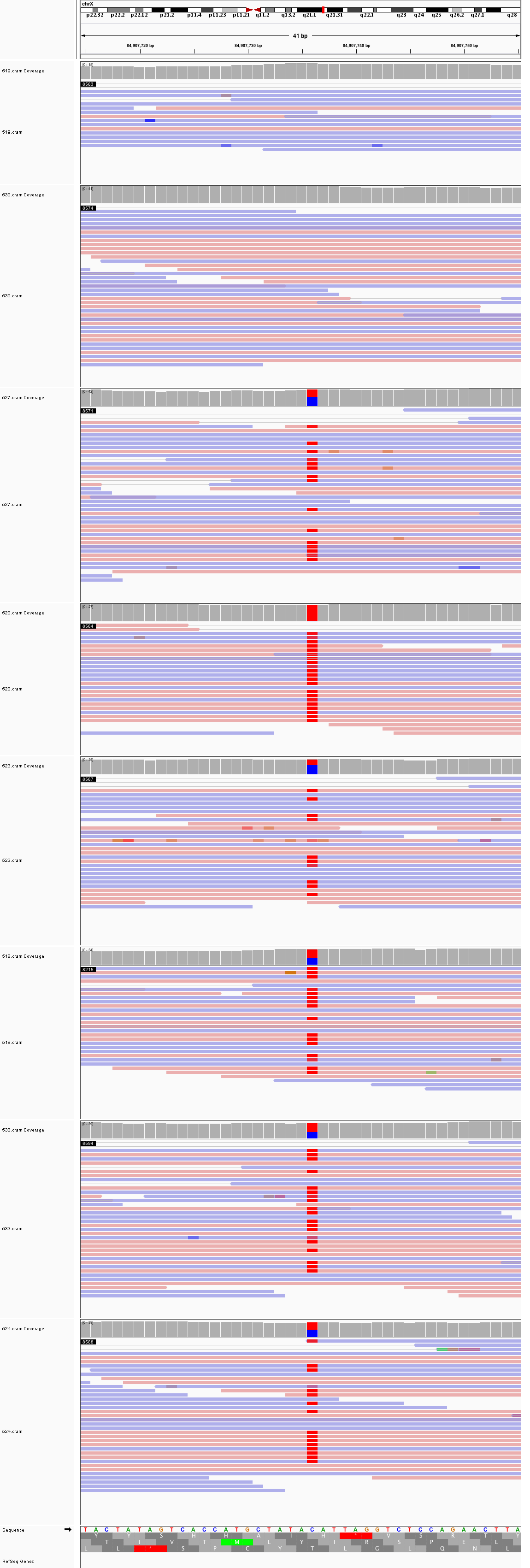

### chrX_114,715,985_114,716,025.png

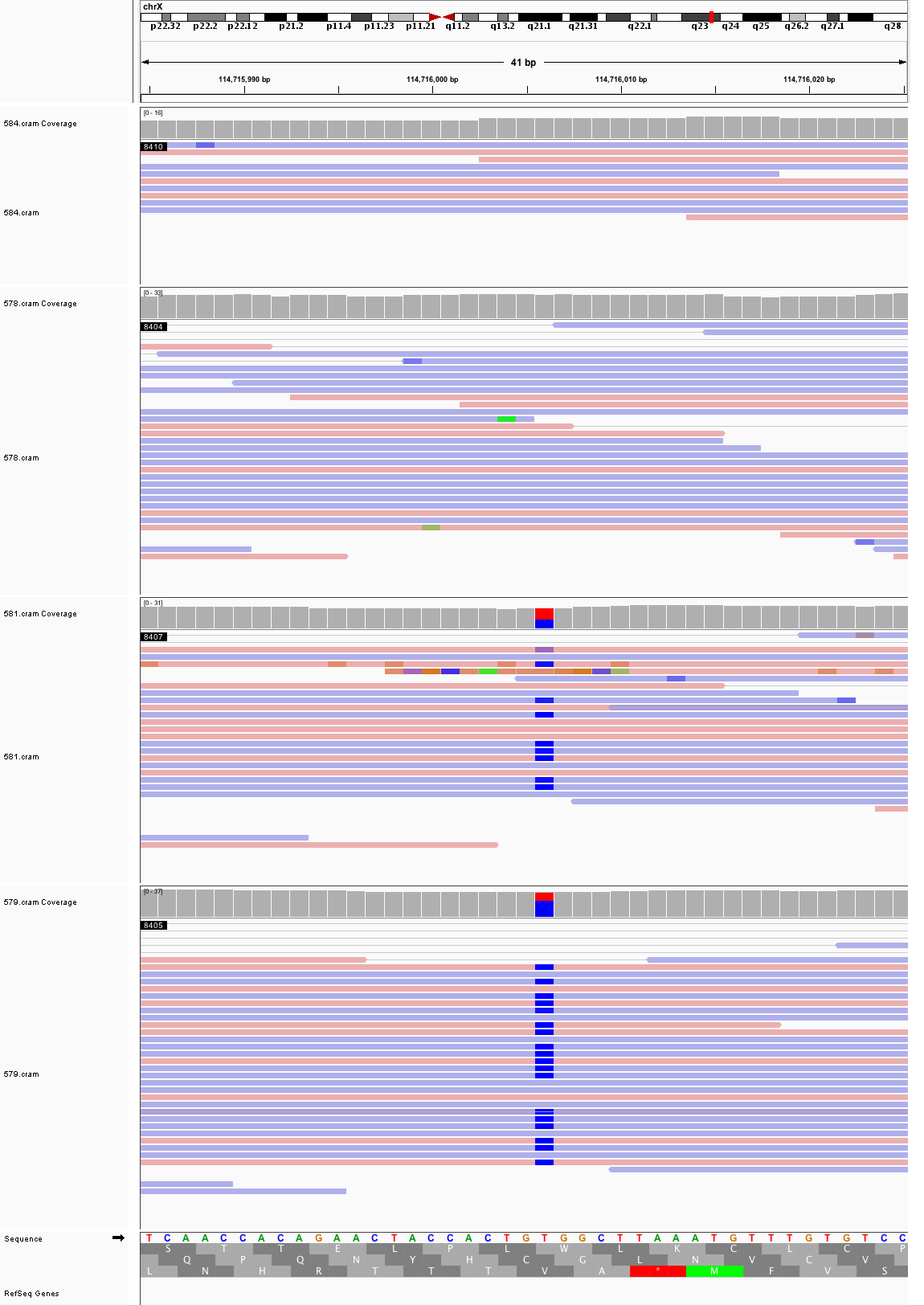

### chrX_133,864,603_133,864,643.png

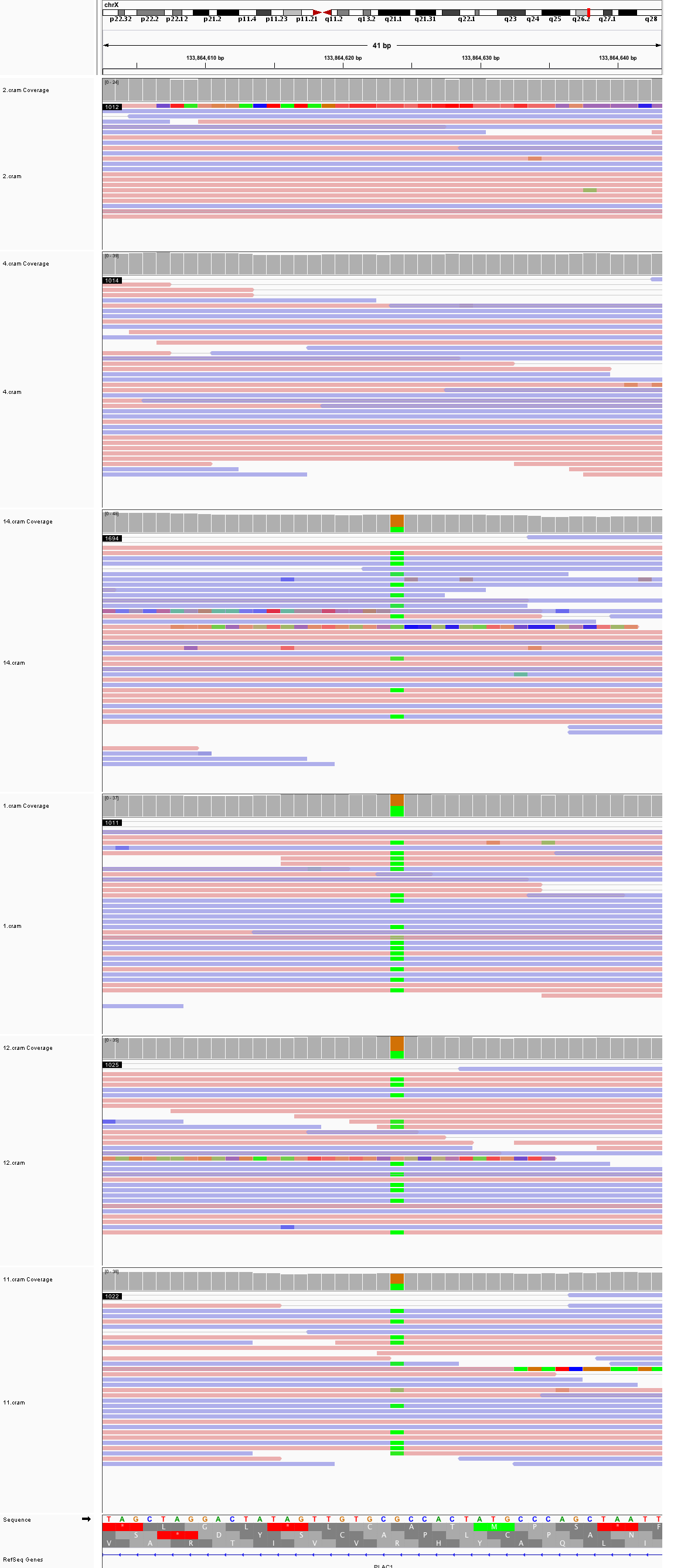
