## Supplementary figures and images for "Large, three-generation CEPH families reveal post-zygotic mosaicism and variability in germline mutation accumulation"

### chr1_21,998,569_21,998,609.png

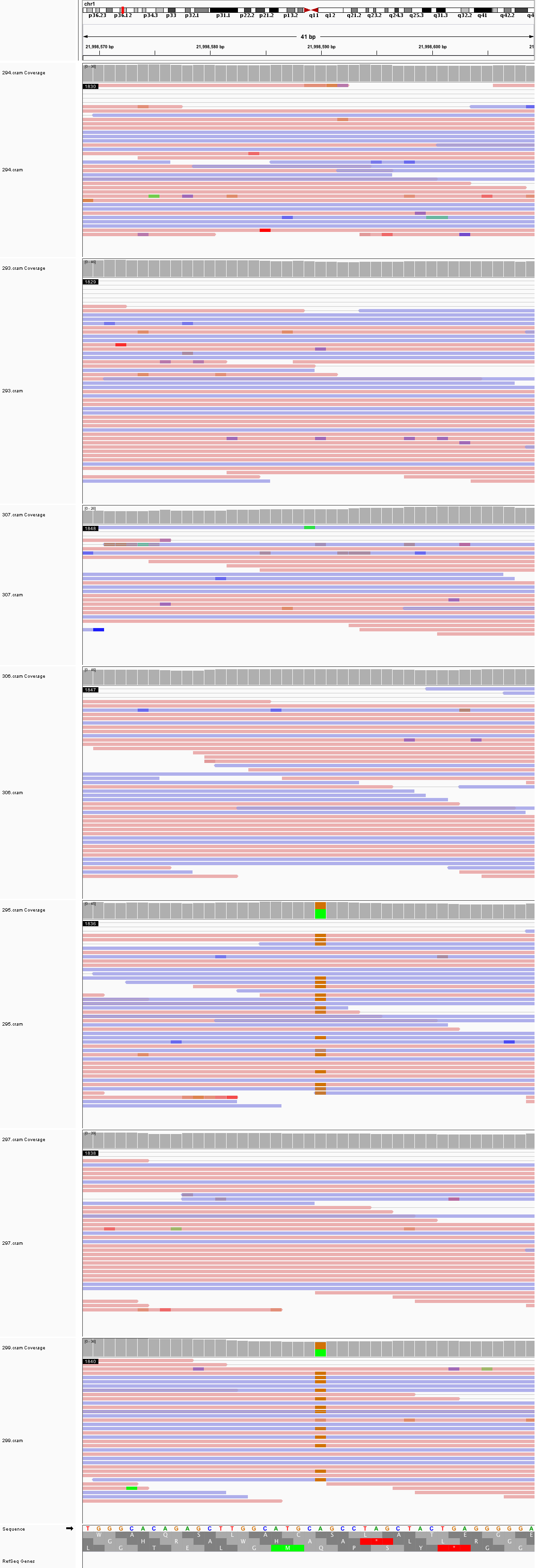

### chr1_68,859,937_68,859,977.png

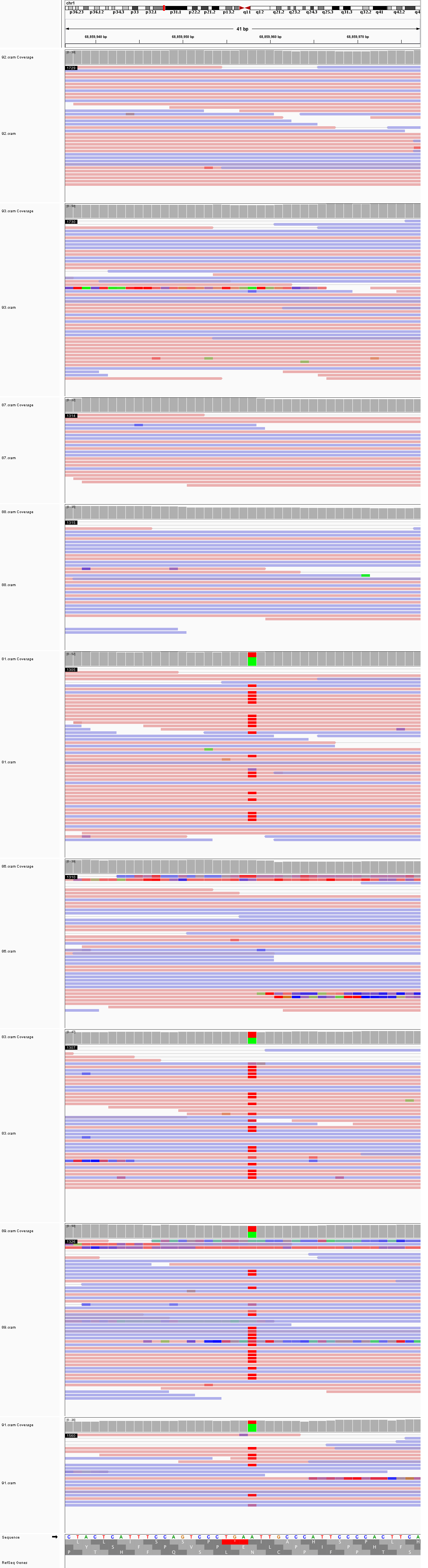

### chr1_219,001,437_219,001,477.png

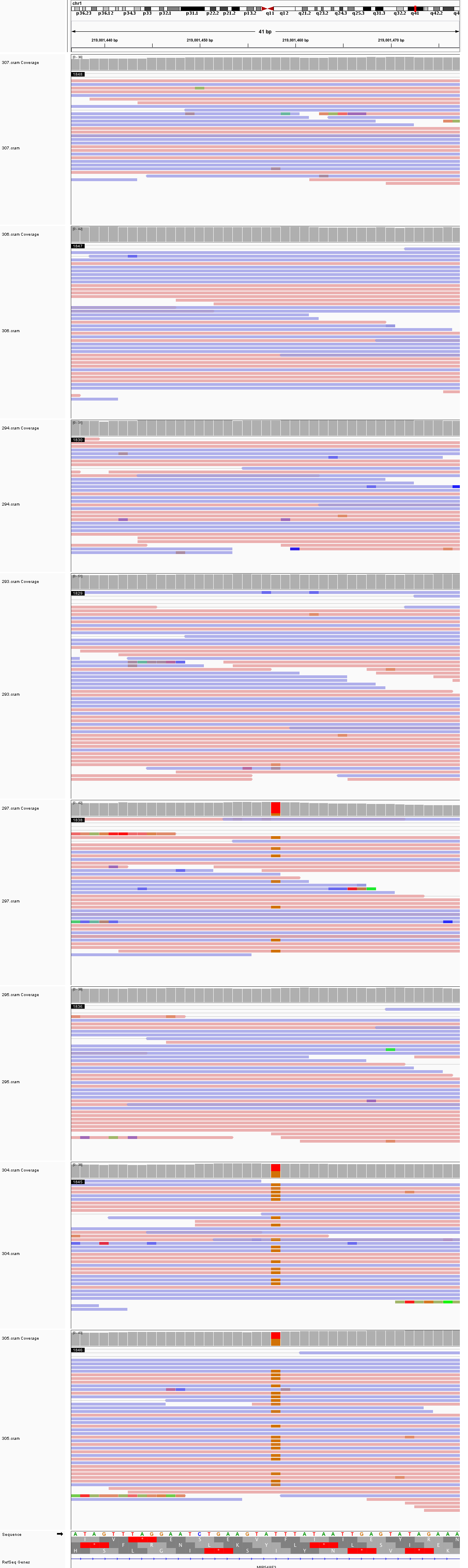

### chr2_6,751,084_6,751,124.png

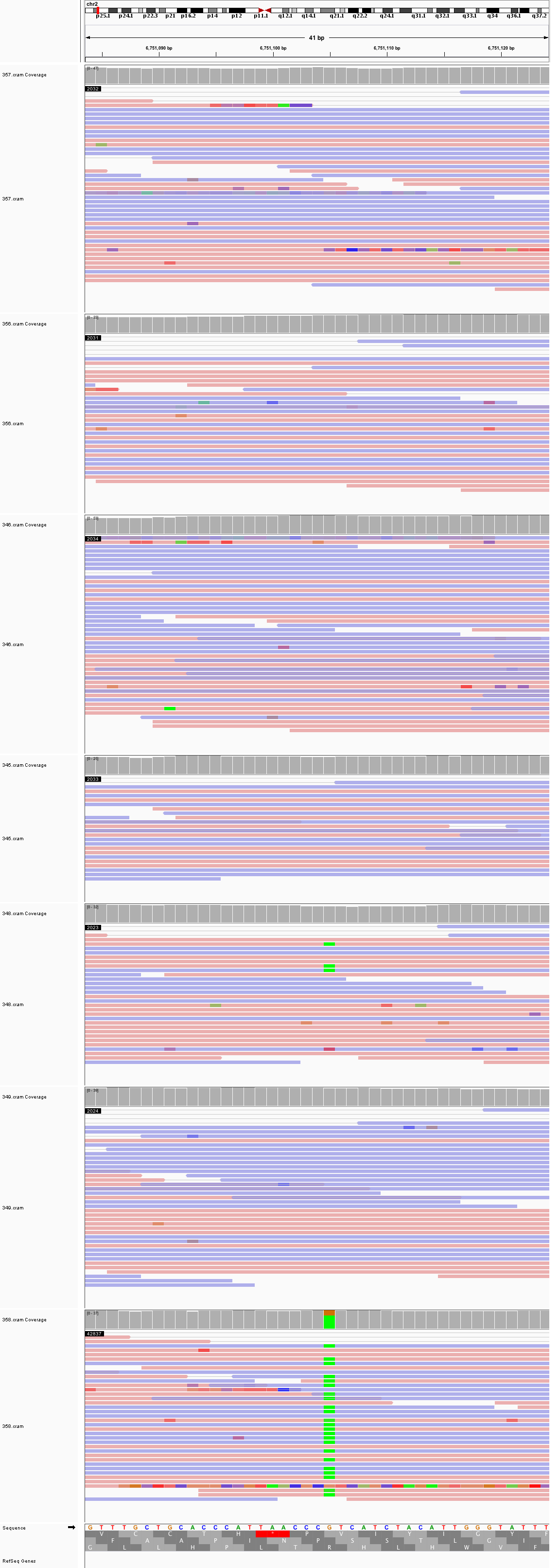

### chr2_109,070,215_109,070,255.png

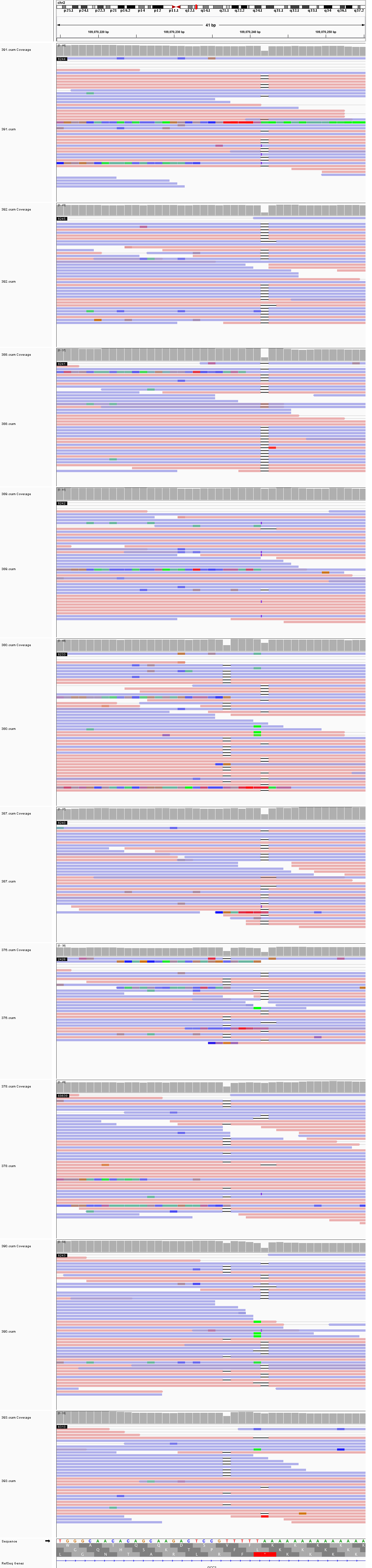

### chr3_65,833,782_65,833,822.png

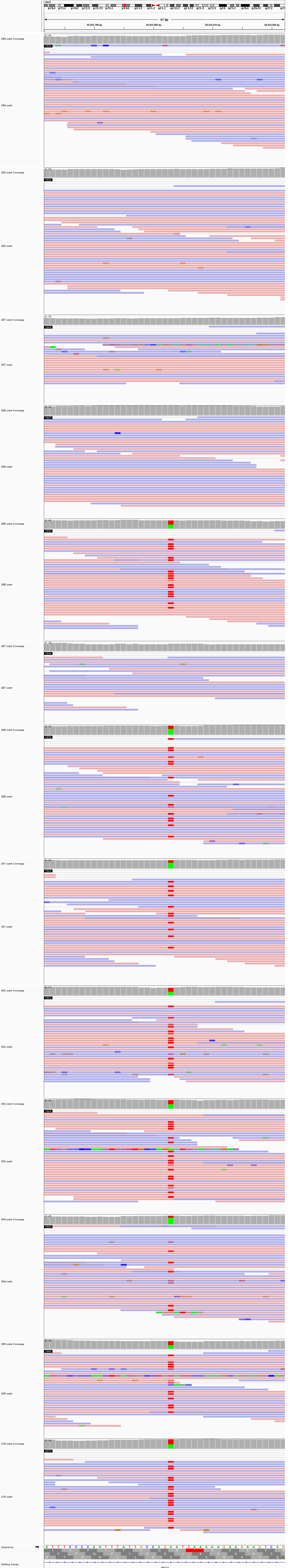

### chr3_79,030,576_79,030,616.png

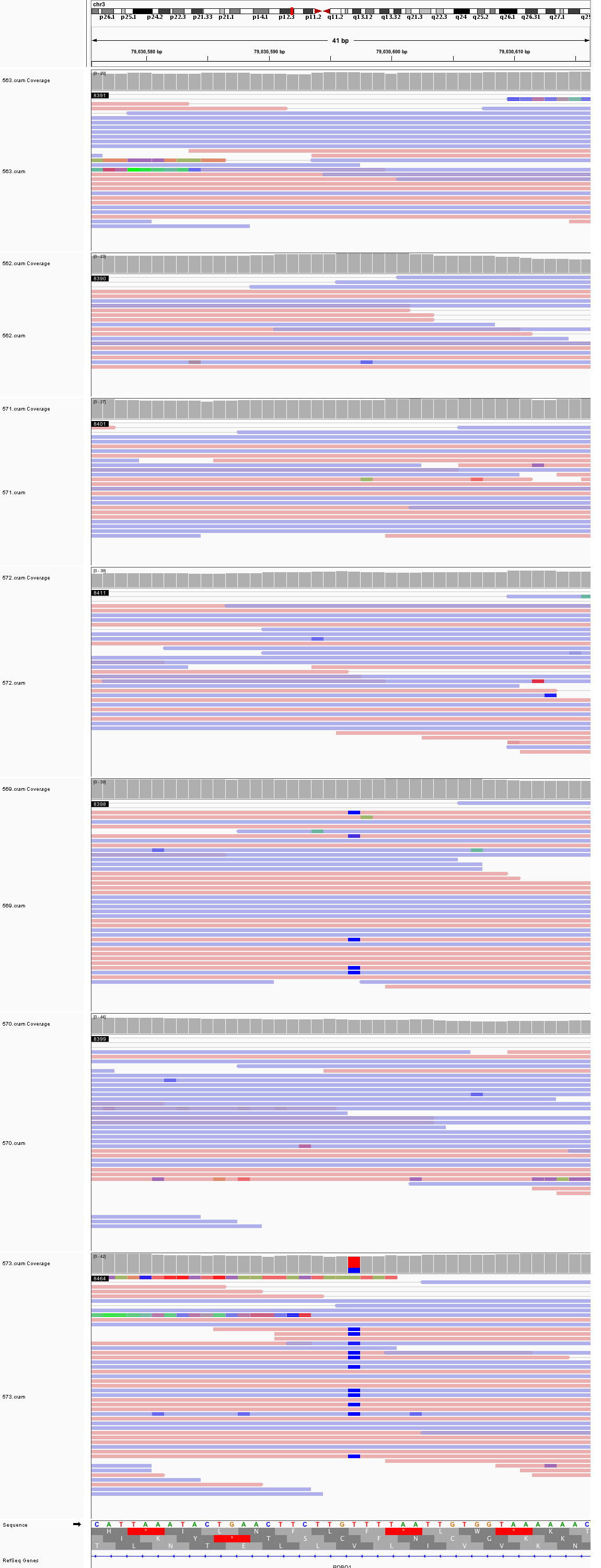

### chr3_108,536,418_108,536,458.png

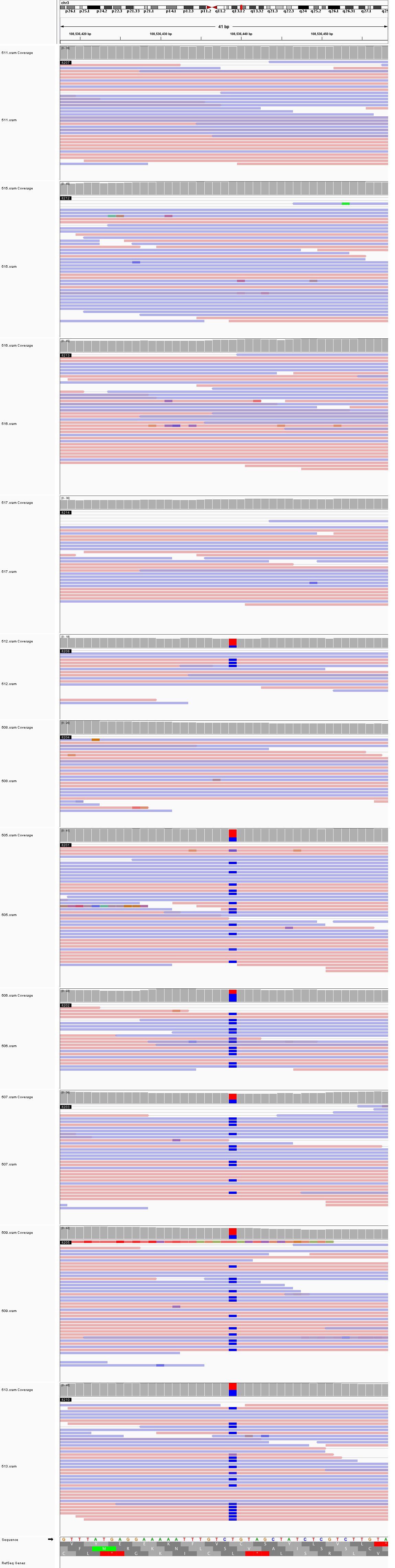

### chr4_9,634,106_9,634,146.png

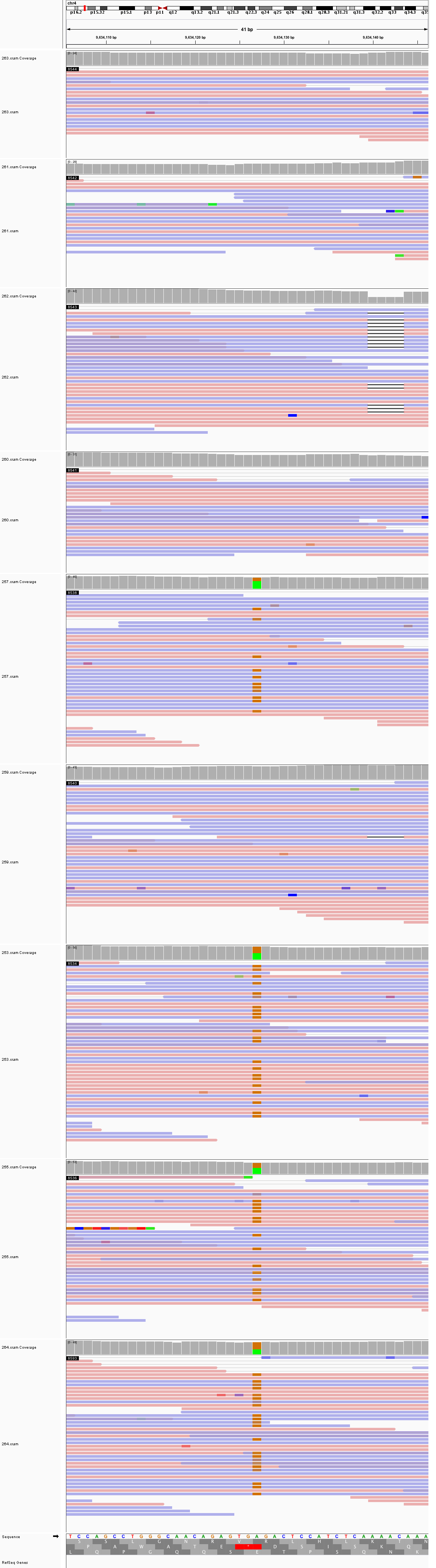

### chr4_33,890,069_33,890,109.png

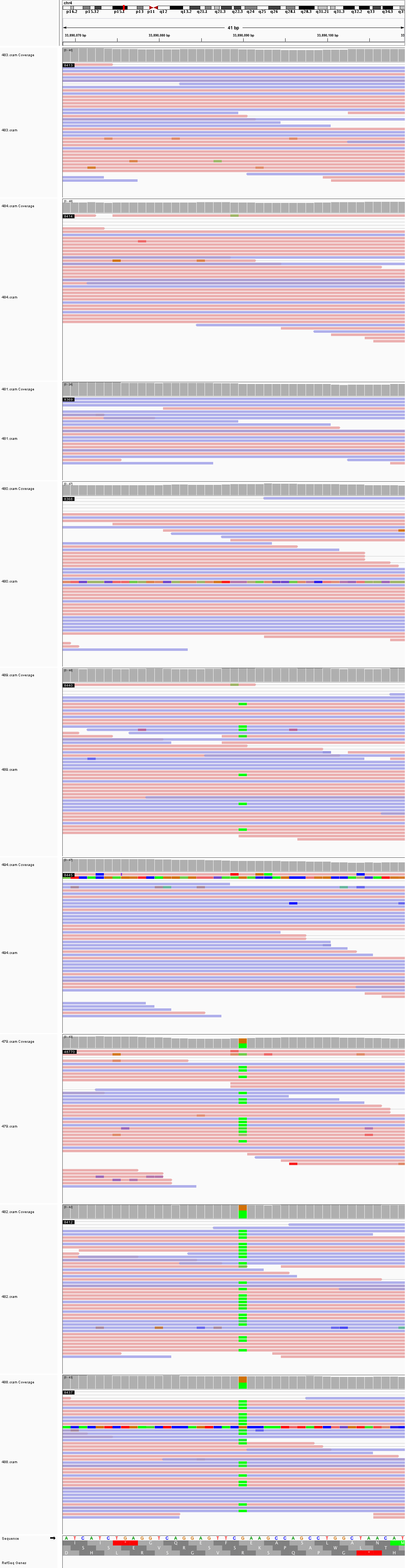

### chr4_54,568,593_54,568,633.png

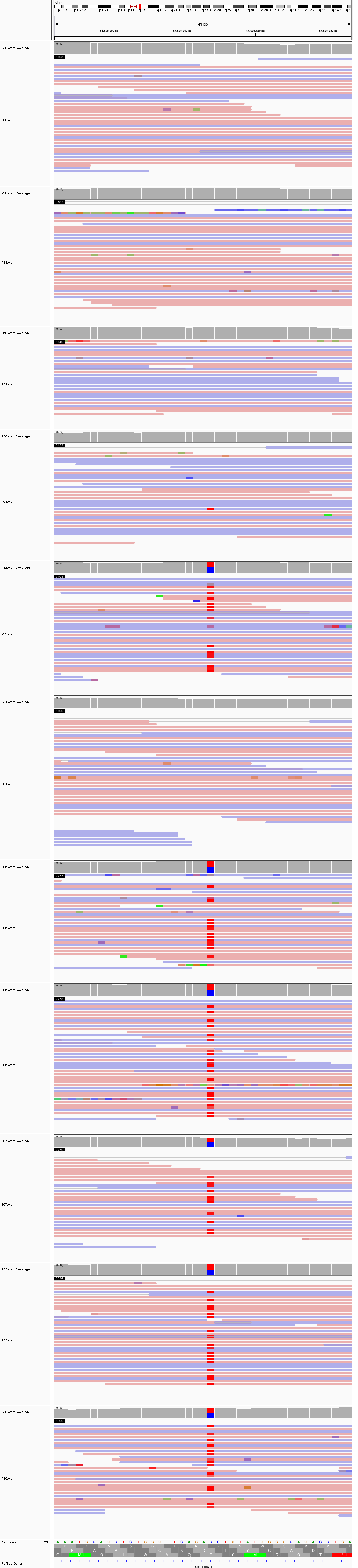

### chr4_118,531,597_118,531,637.png

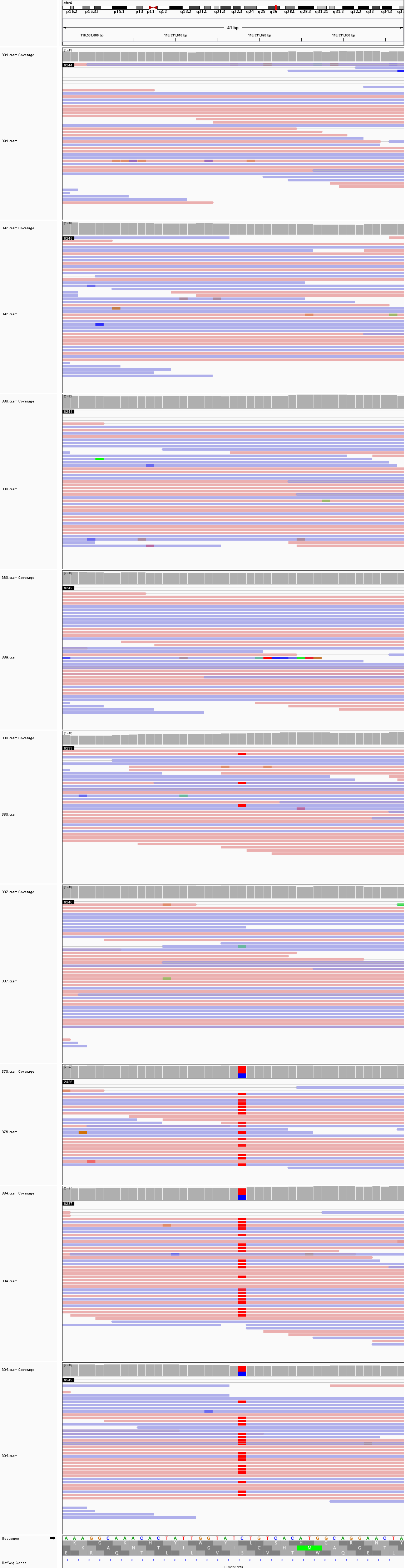

### chr5_25,949,220_25,949,260.png

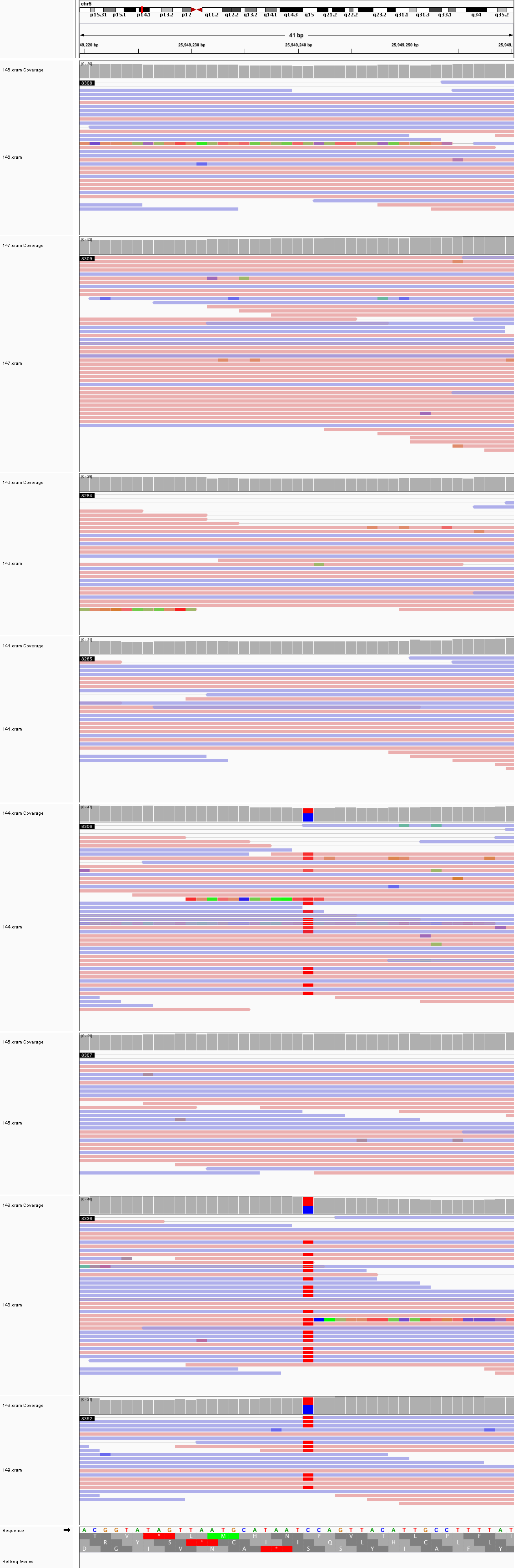

### chr5_52,665,158_52,665,198.png

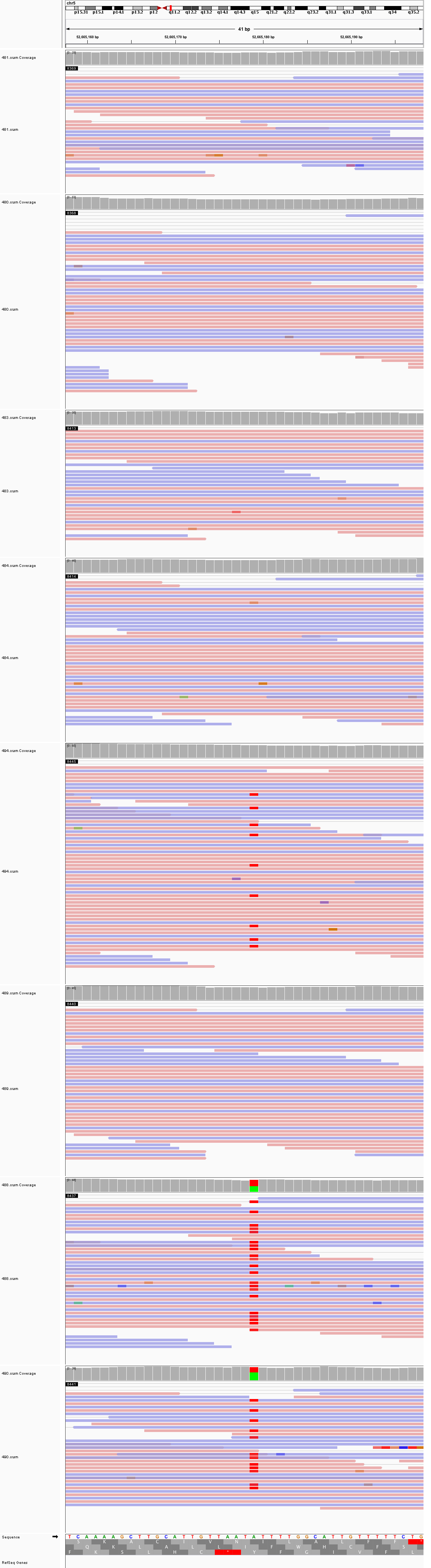

### chr6_44,857,300_44,857,340.png

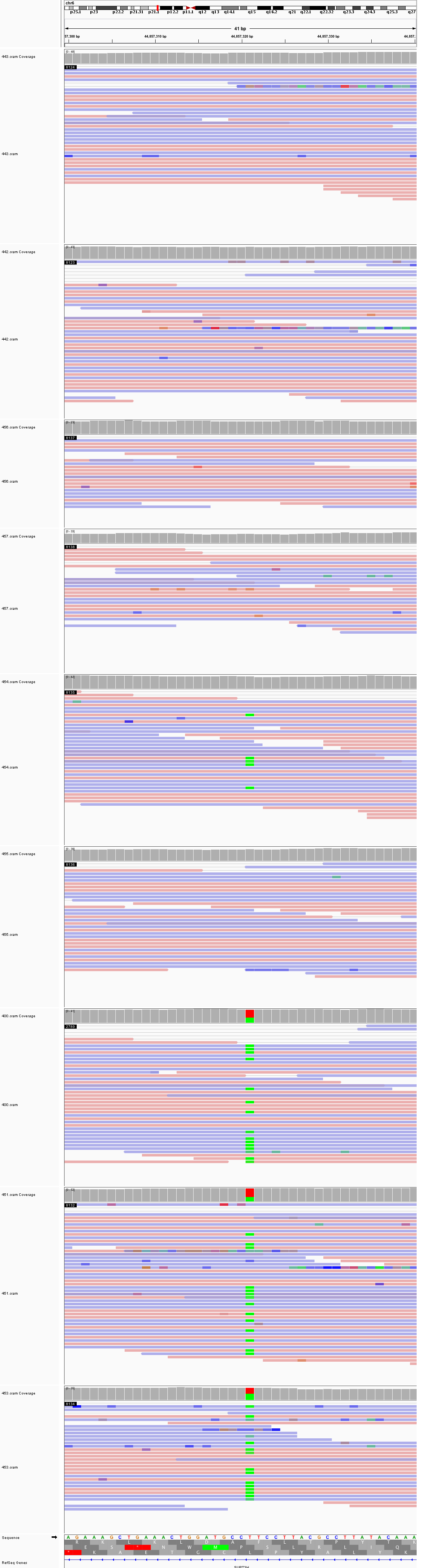

### chr8_53,067,876_53,067,916.png

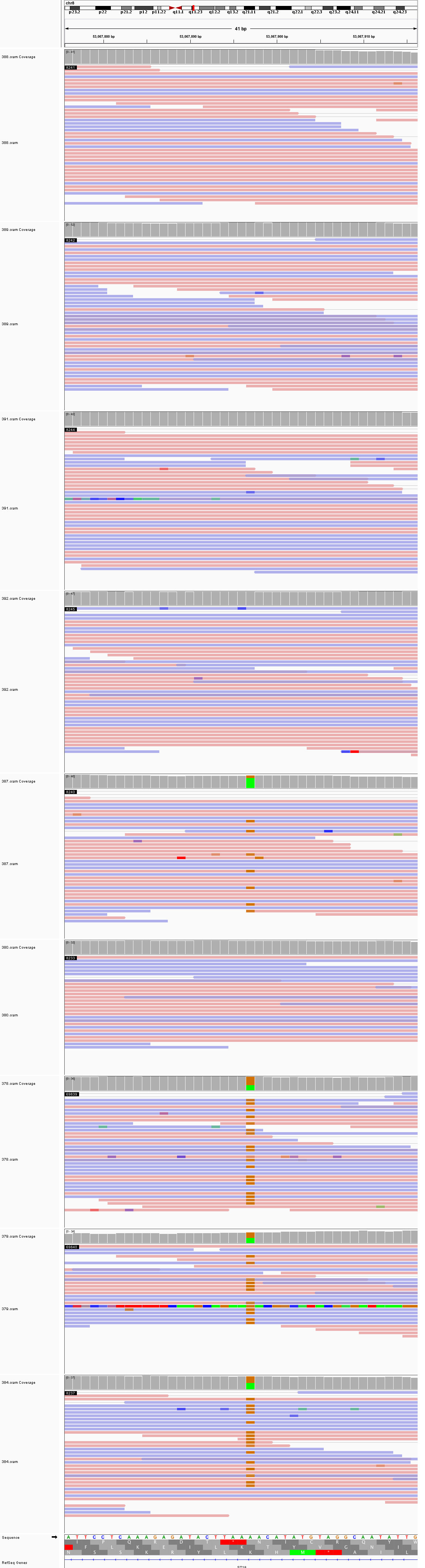

### chr8_99,257,806_99,257,846.png

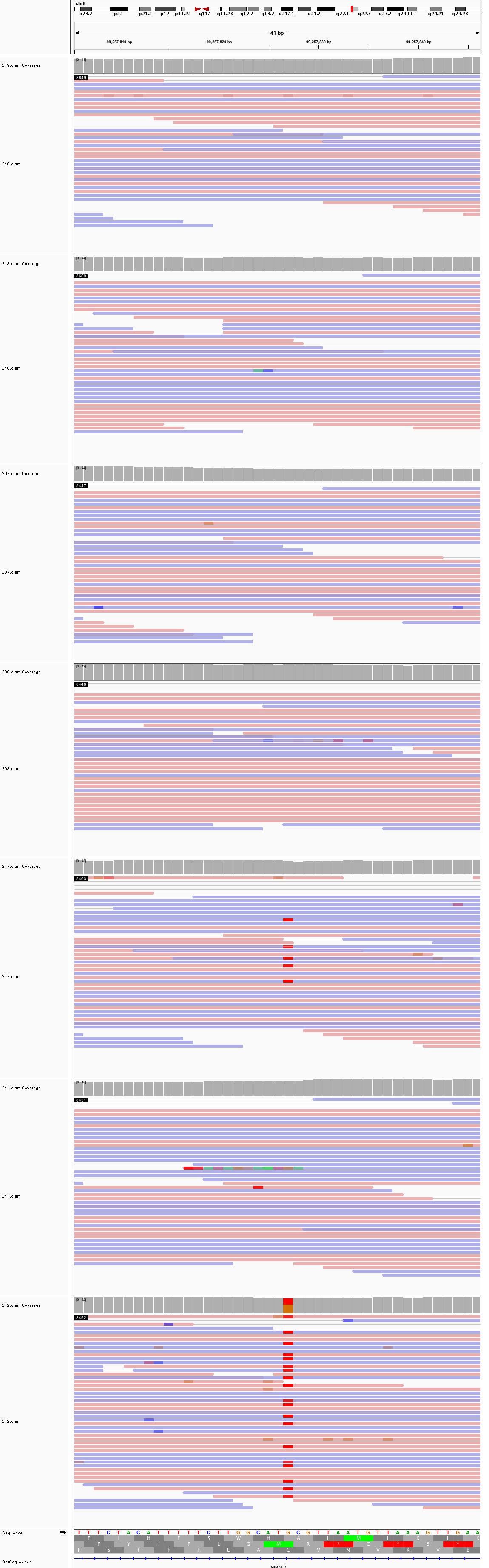

### chr10_36,949,458_36,949,498.png

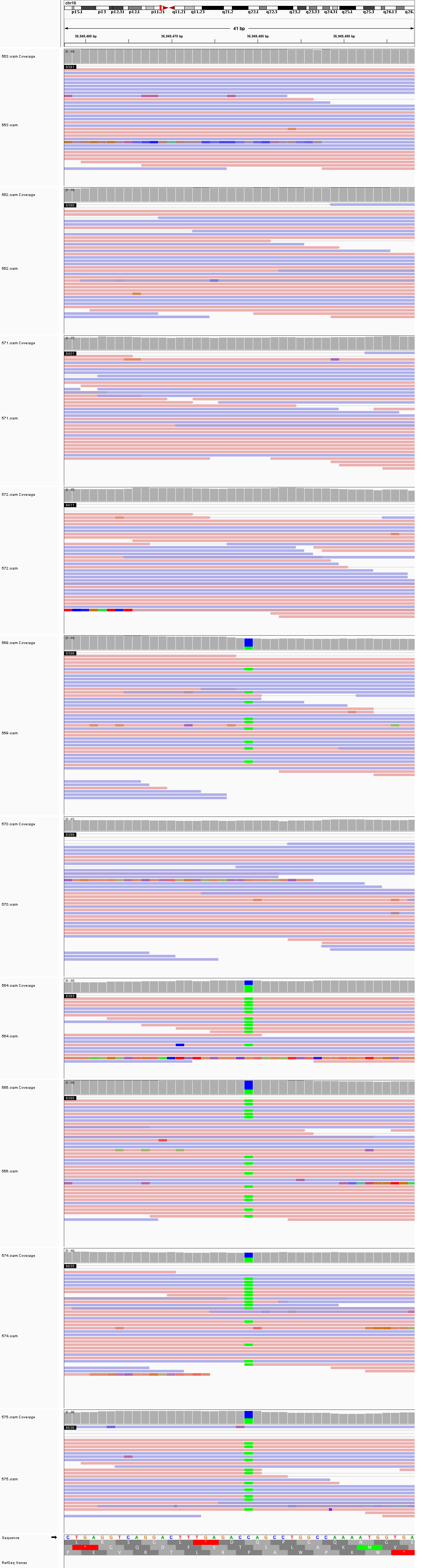

### chr10_130,402,010_130,402,050.png

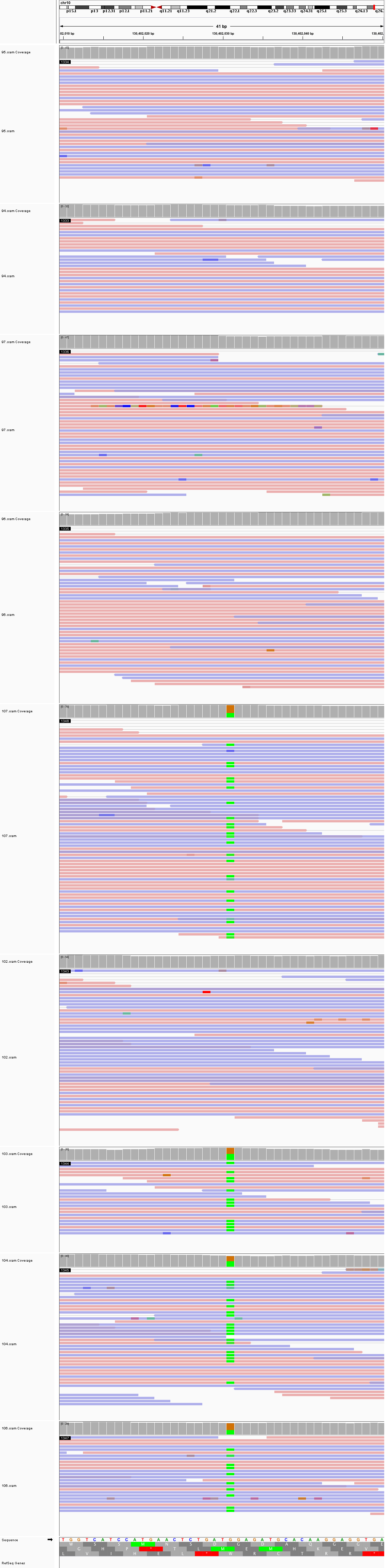

### chr11_17,616,087_17,616,127.png

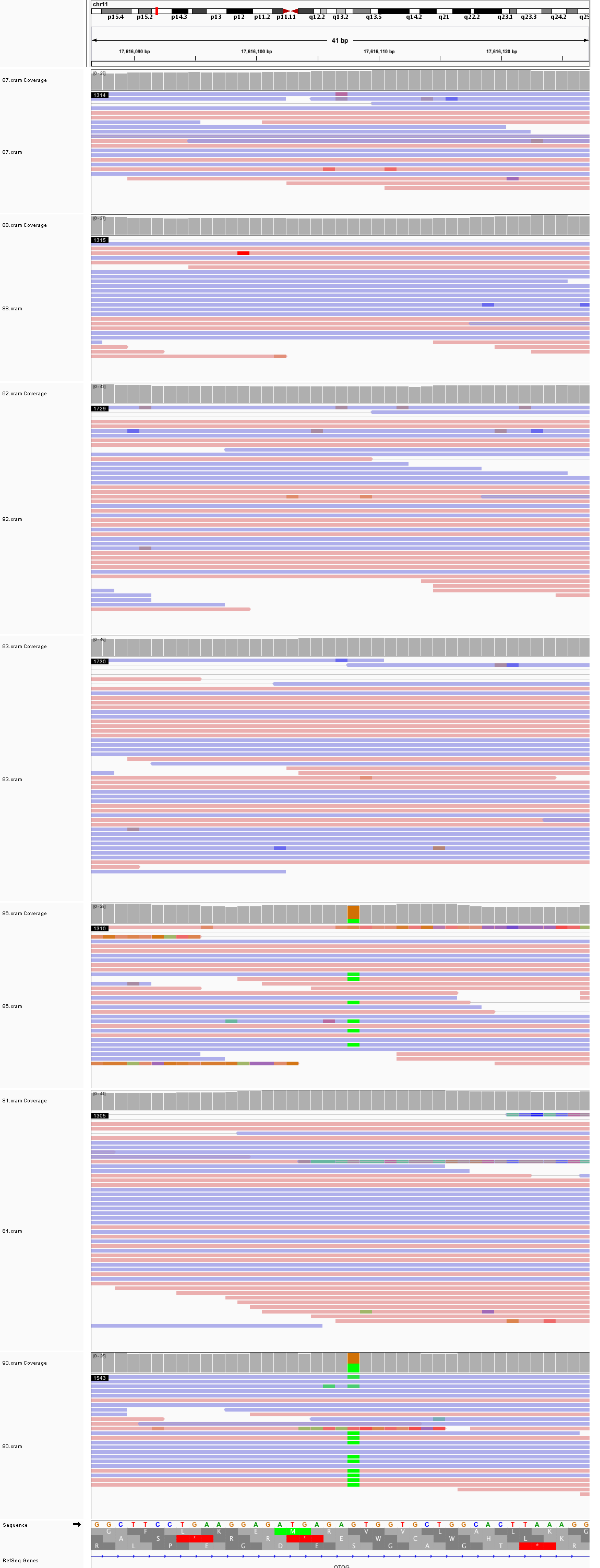

### chr11_86,967,764_86,967,804.png

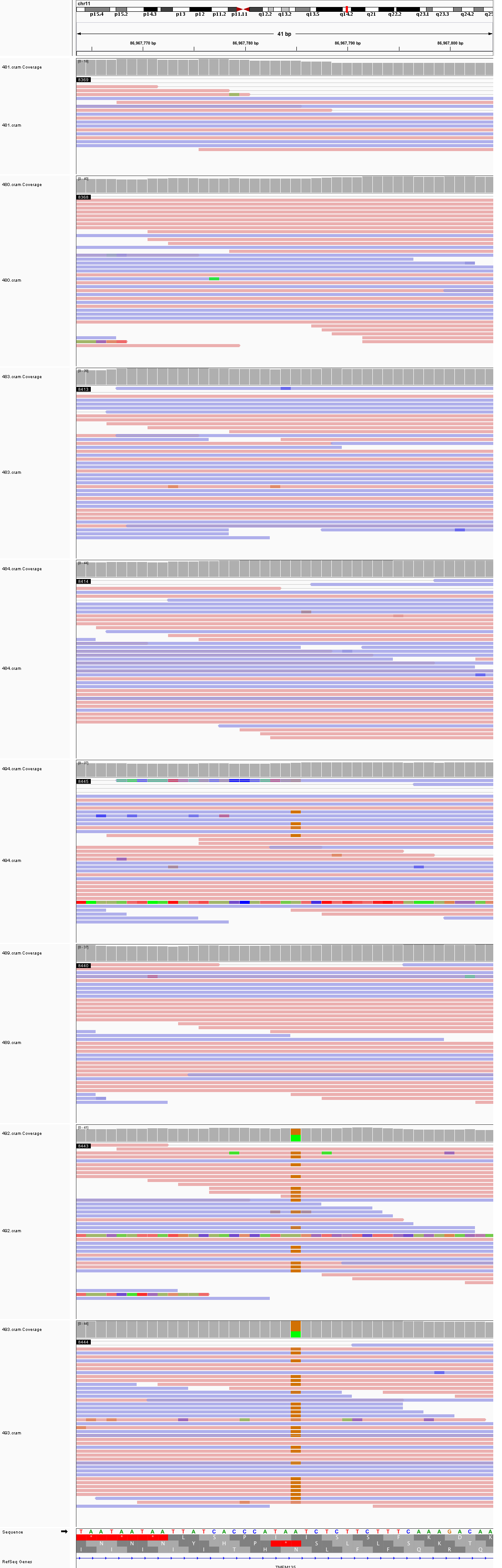

### chr11_134,209,239_134,209,279.png

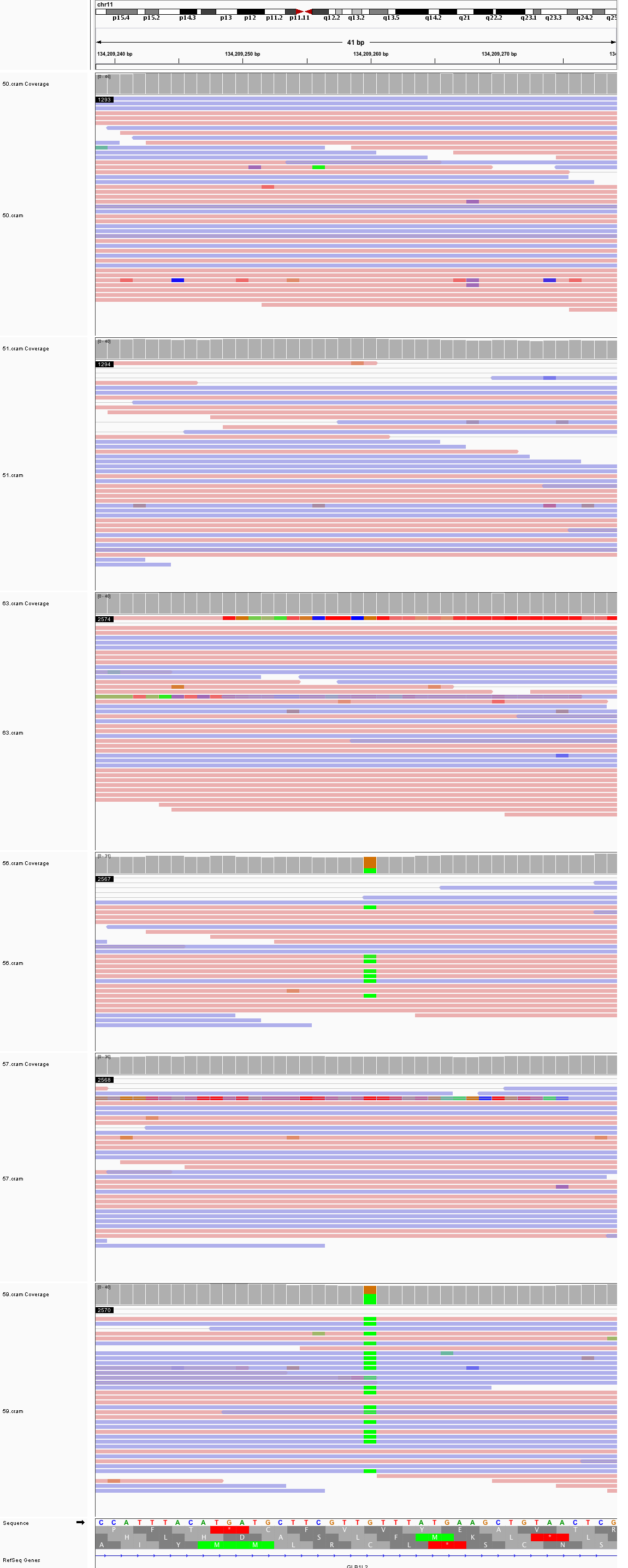

### chr12_46,995,253_46,995,293.png

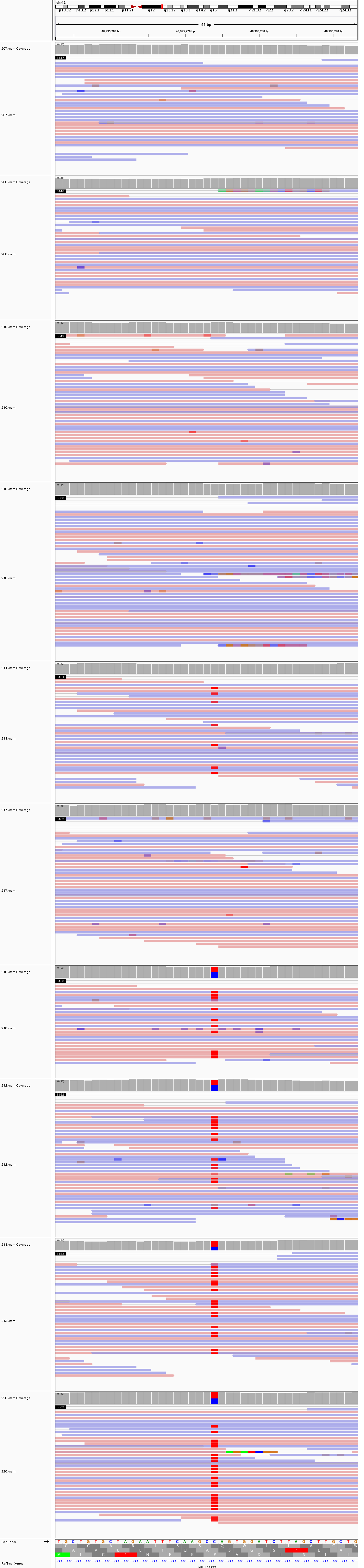

### chr12_117,570,598_117,570,638.png

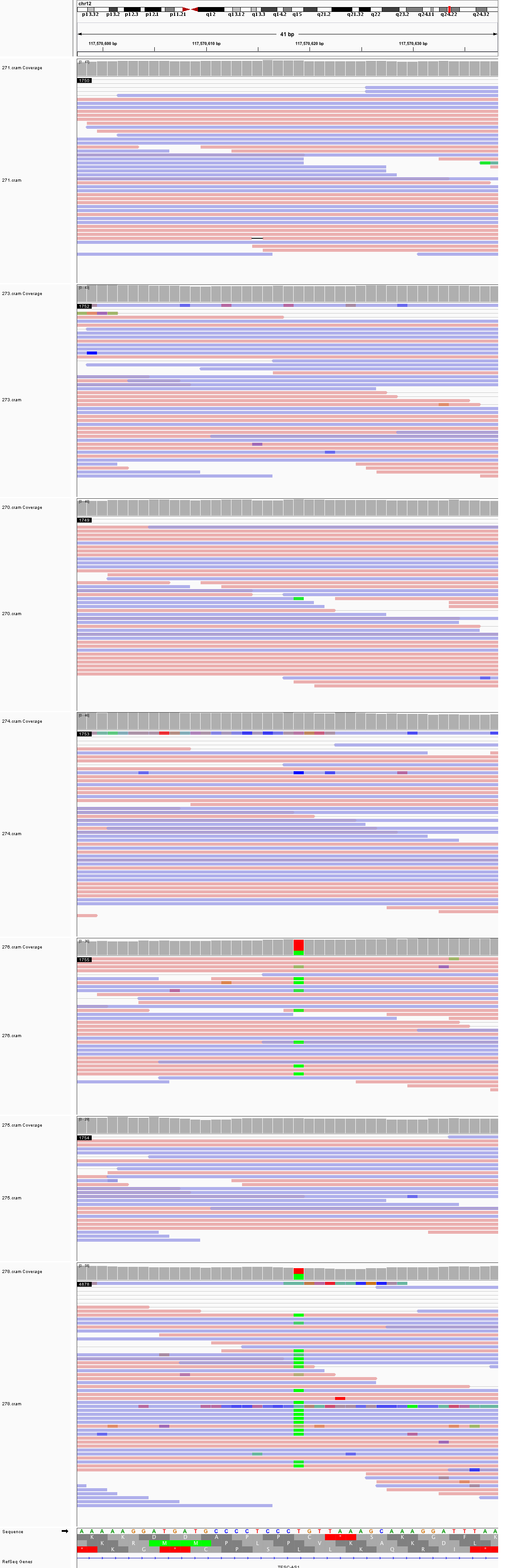

### chr12_132,979,729_132,979,769.png

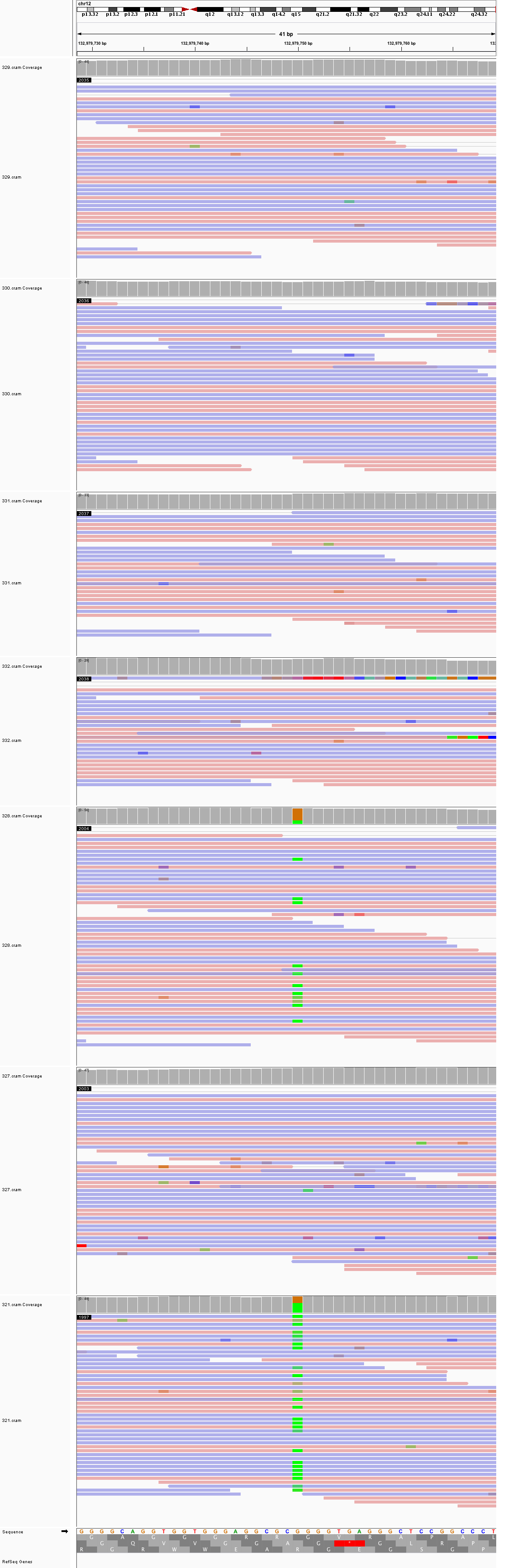

### chr13_57,193,407_57,193,447.png

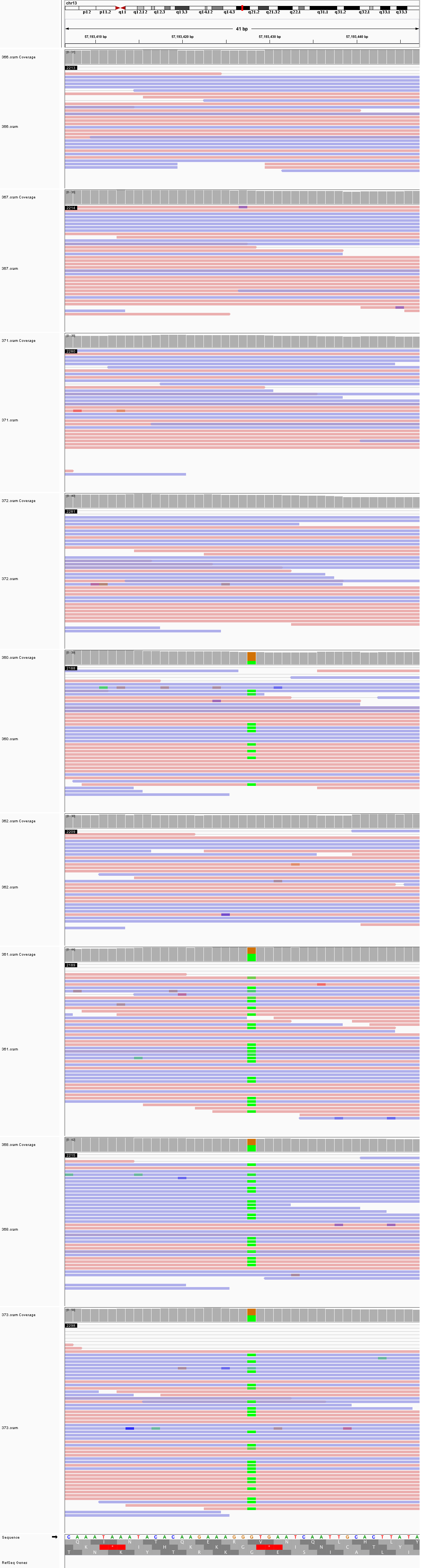

### chr16_28,662,312_28,662,352.png

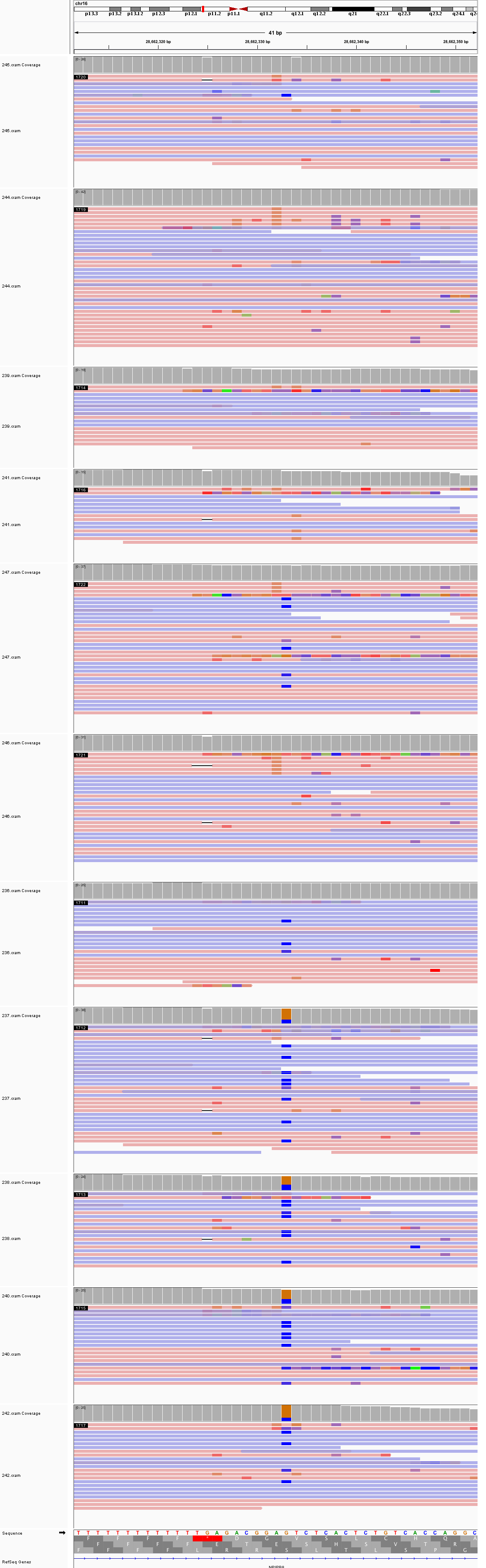

### chr16_59,030,712_59,030,752.png

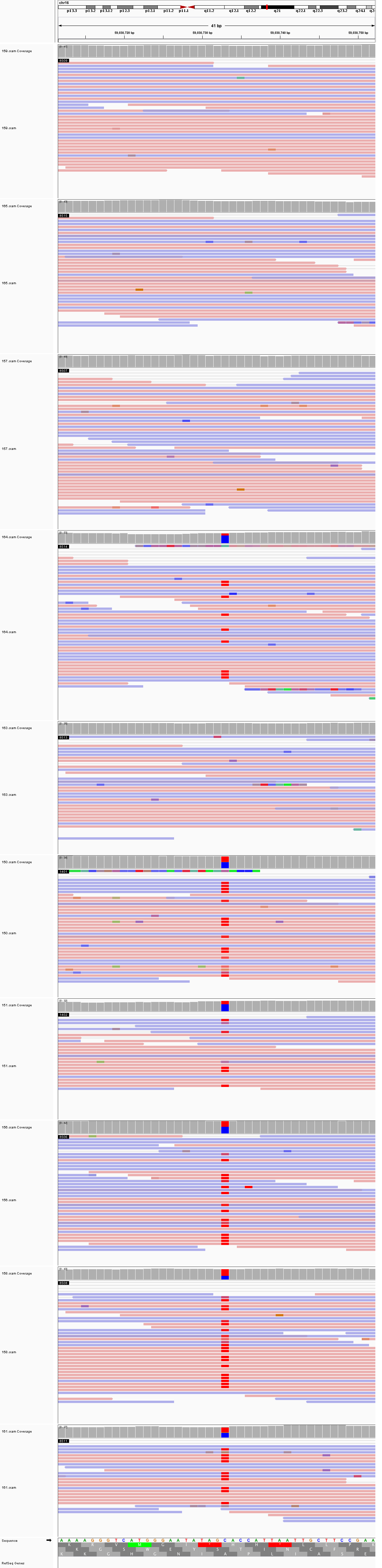

### chr16_73,997,224_73,997,264.png

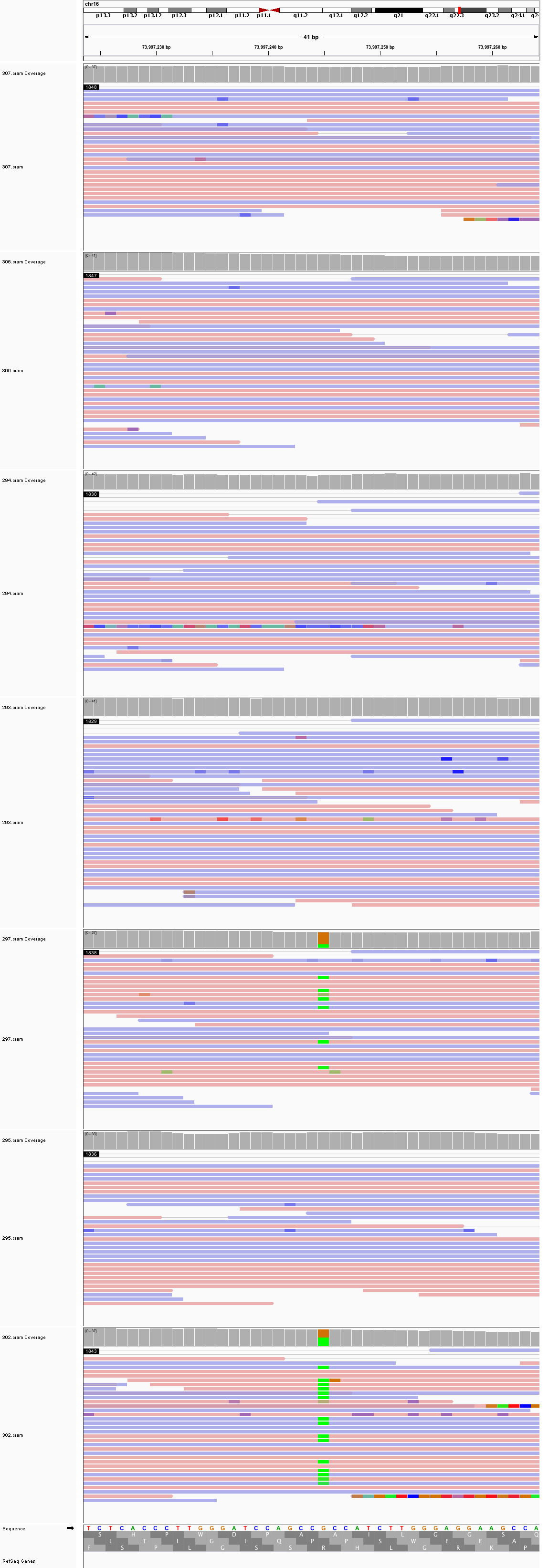

### chr19_22,985,684_22,985,724.png

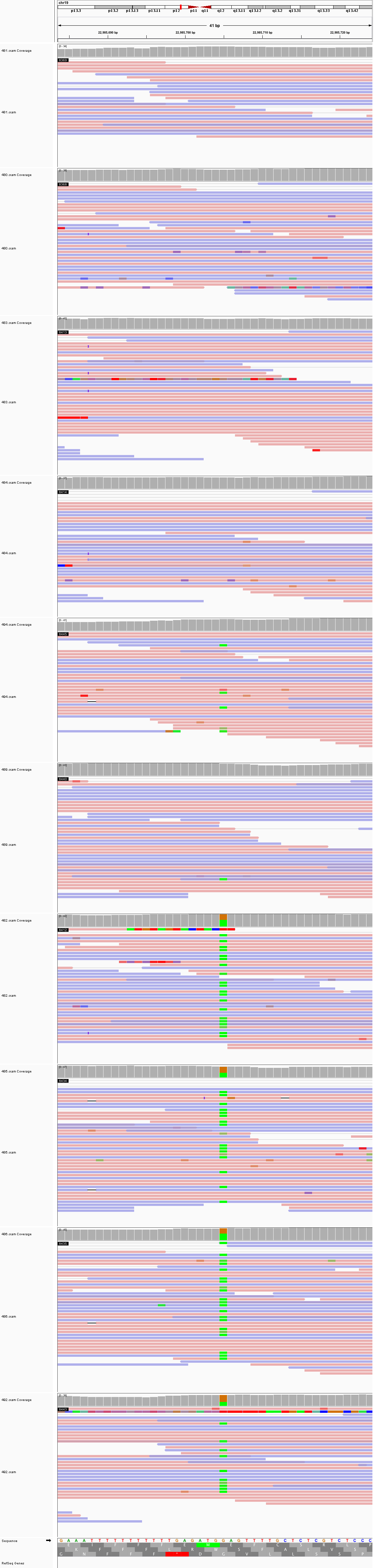

### chr19_46,001,911_46,001,951.png

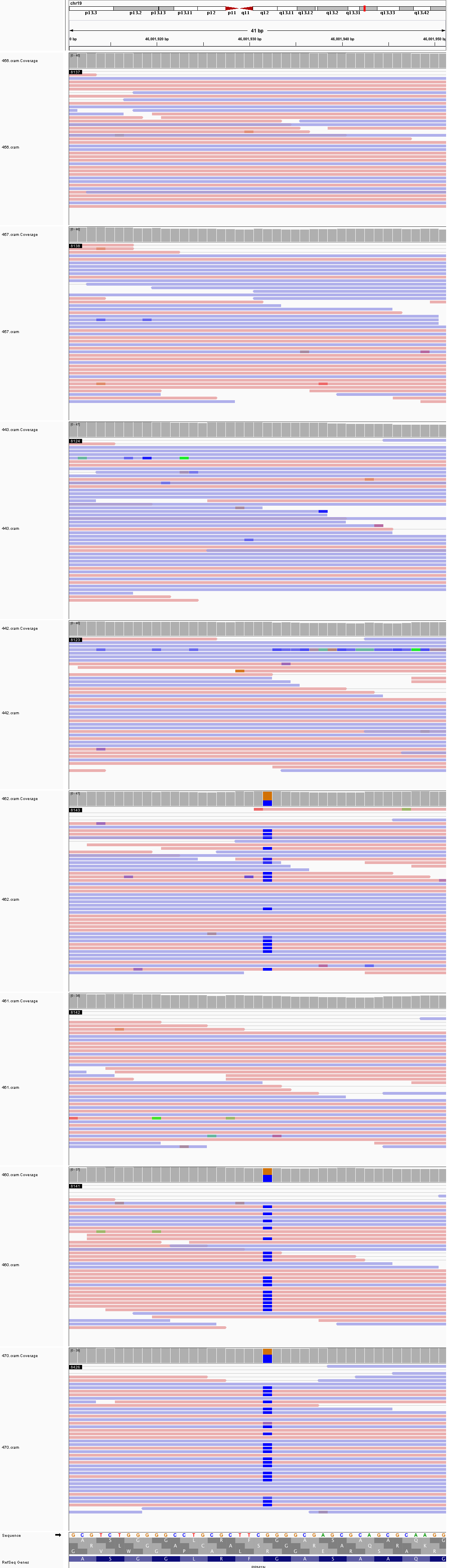

### chr19_56,078,958_56,078,998.png

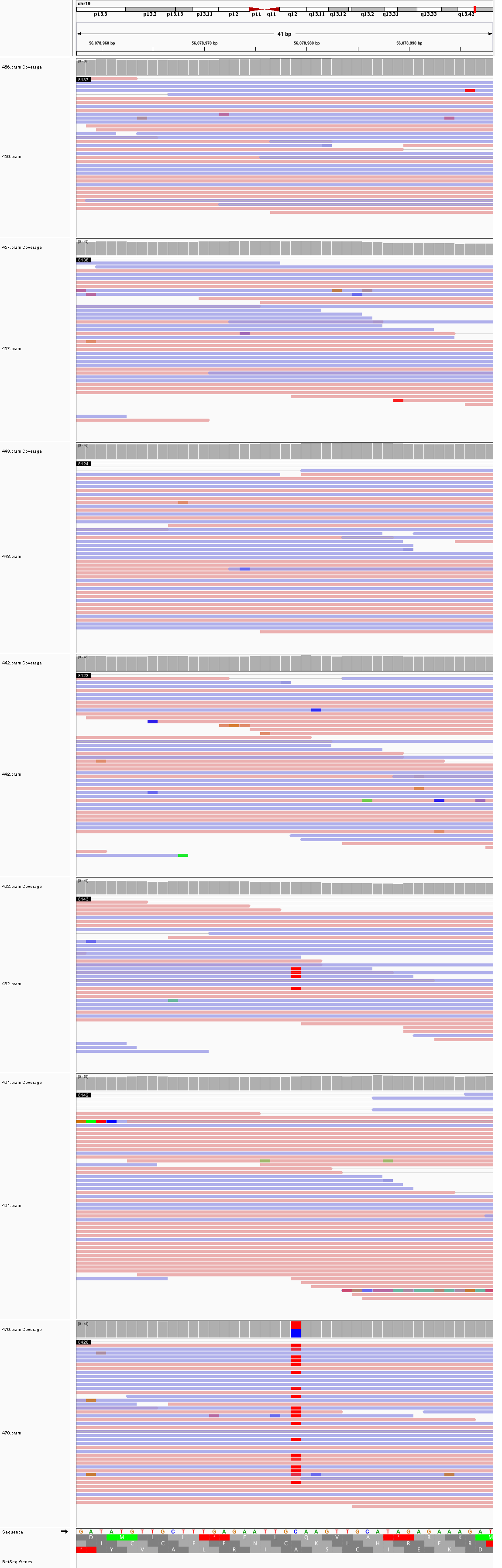

### chr20_30,698,163_30,698,203.png

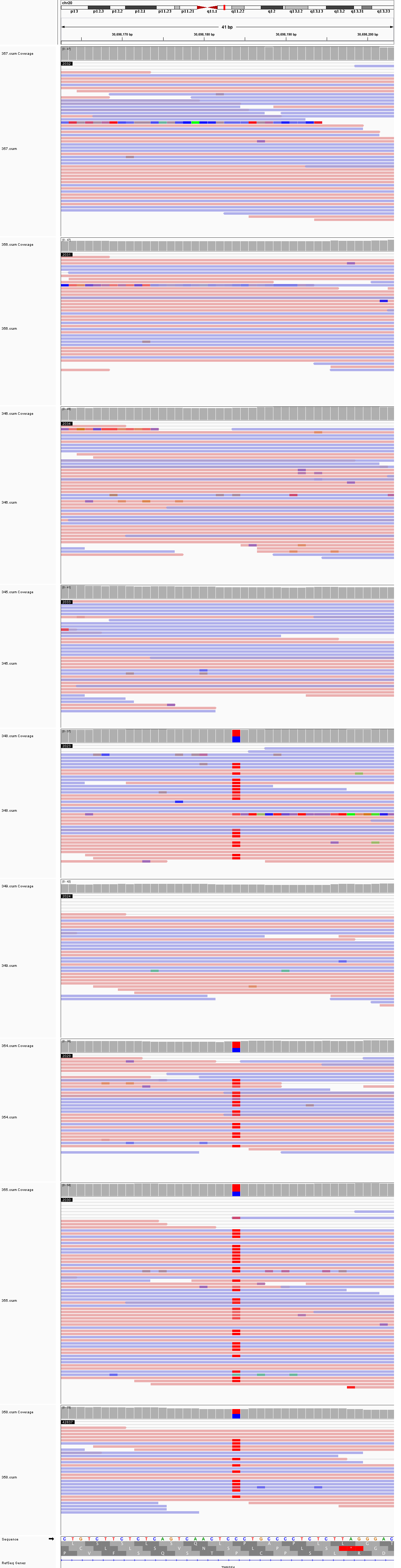
