## Supplementary figures and images for "Large, three-generation CEPH families reveal post-zygotic mosaicism and variability in germline mutation accumulation"

### chr1_10,290,743_10,290,783.png

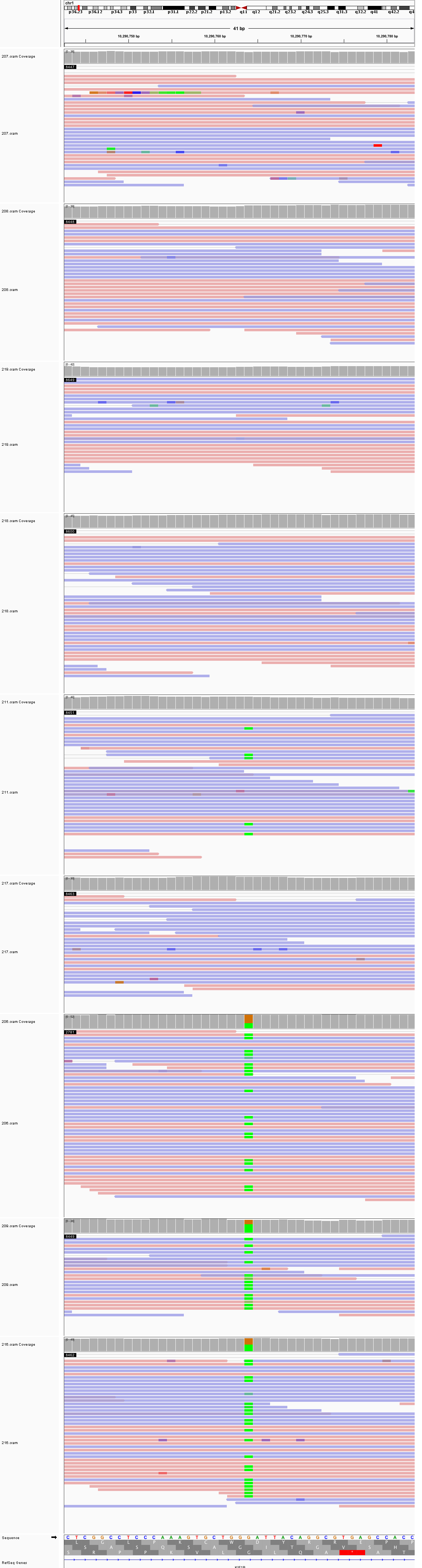

### chr1_11,185,218_11,185,258.png

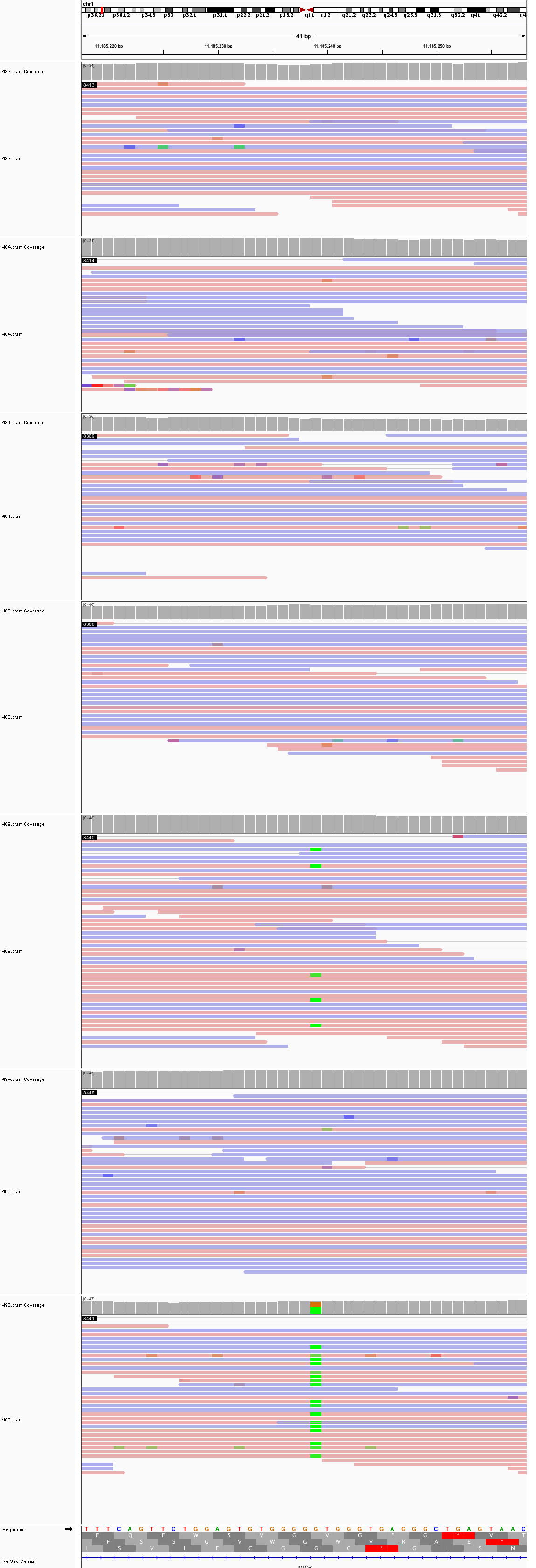

### chr1_24,536,590_24,536,630.png

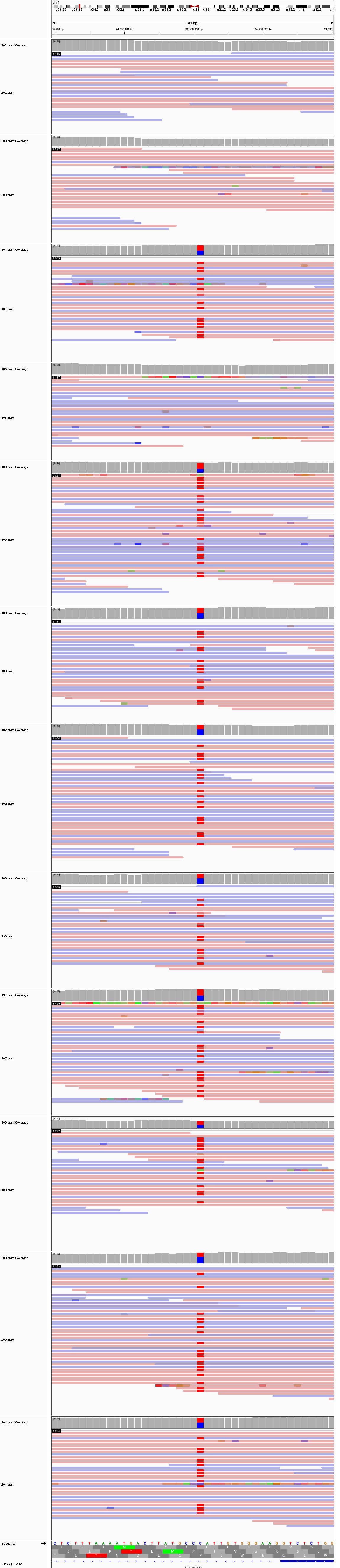

### chr1_29,251,413_29,251,453.png

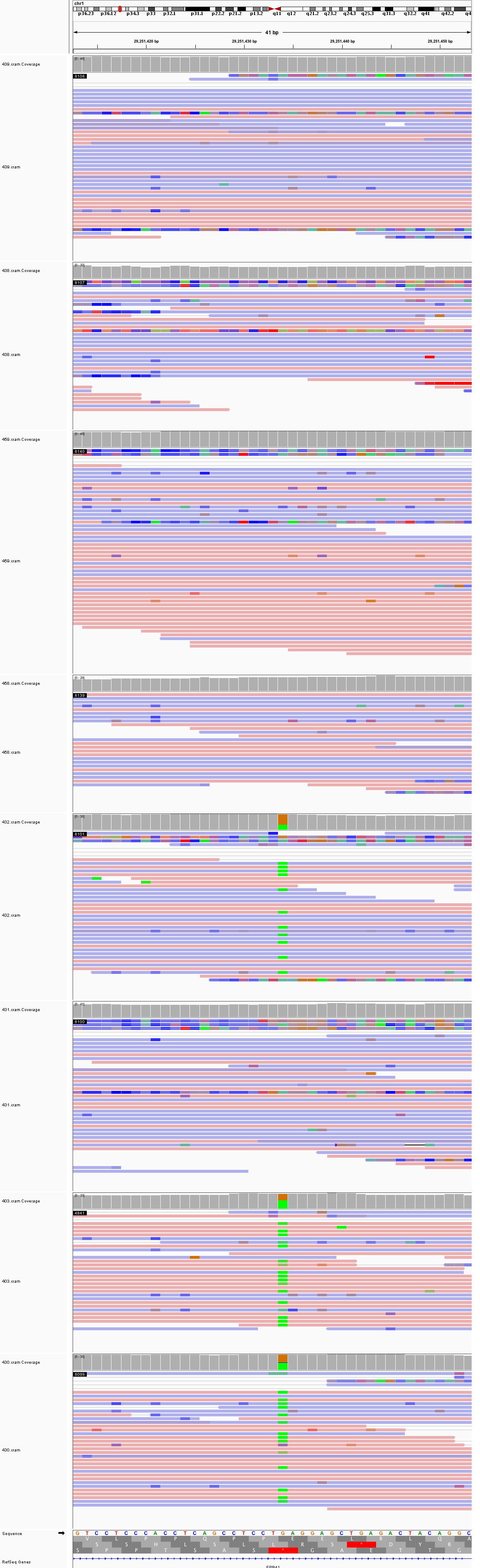

### chr1_36,518,585_36,518,625.png

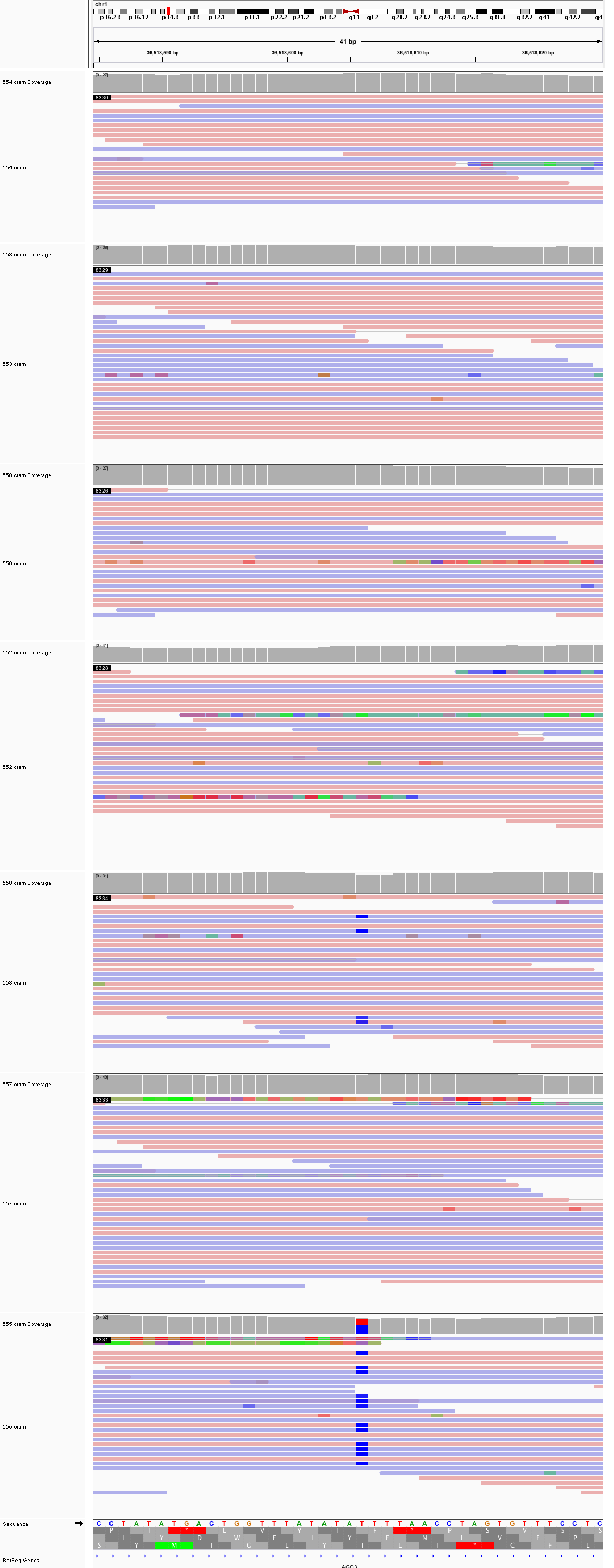

### chr1_113,317,224_113,317,264.png

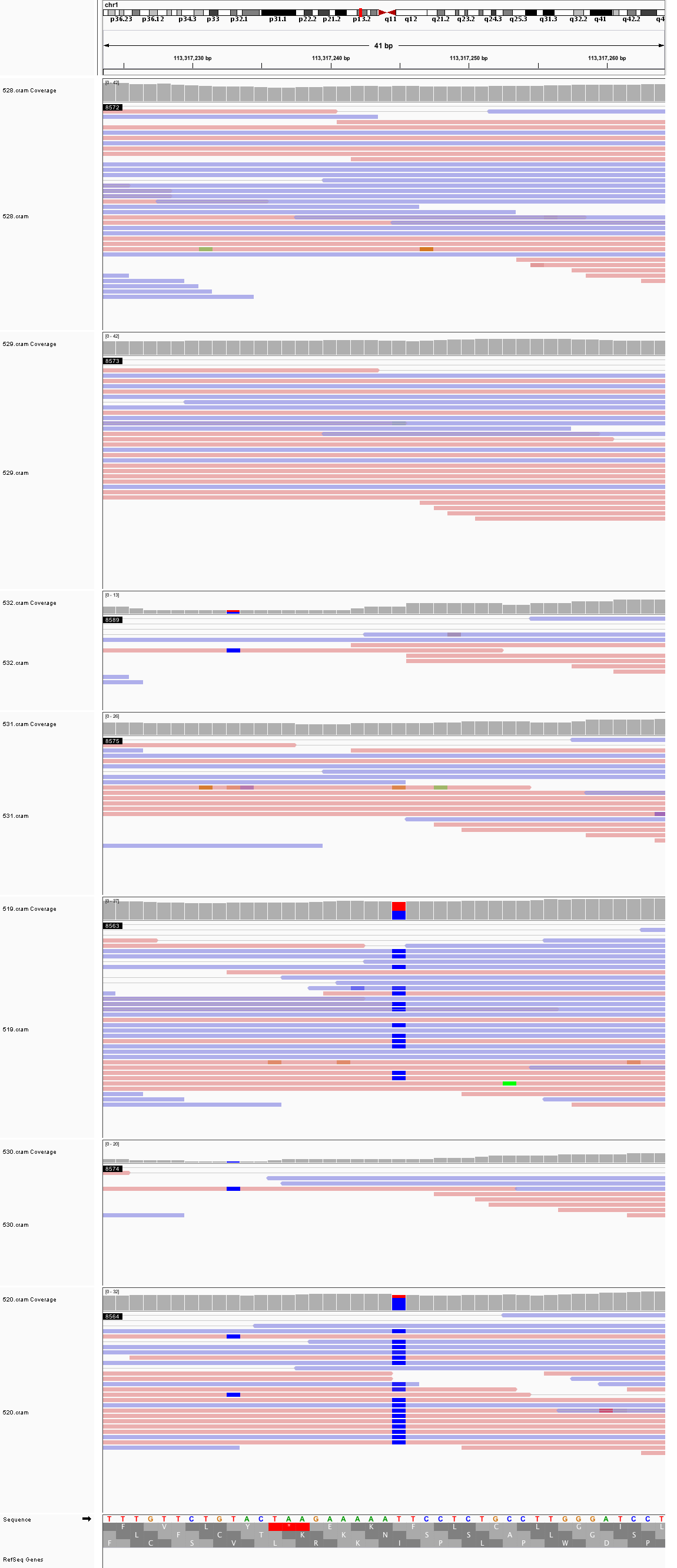

### chr1_116,795,419_116,795,459.png

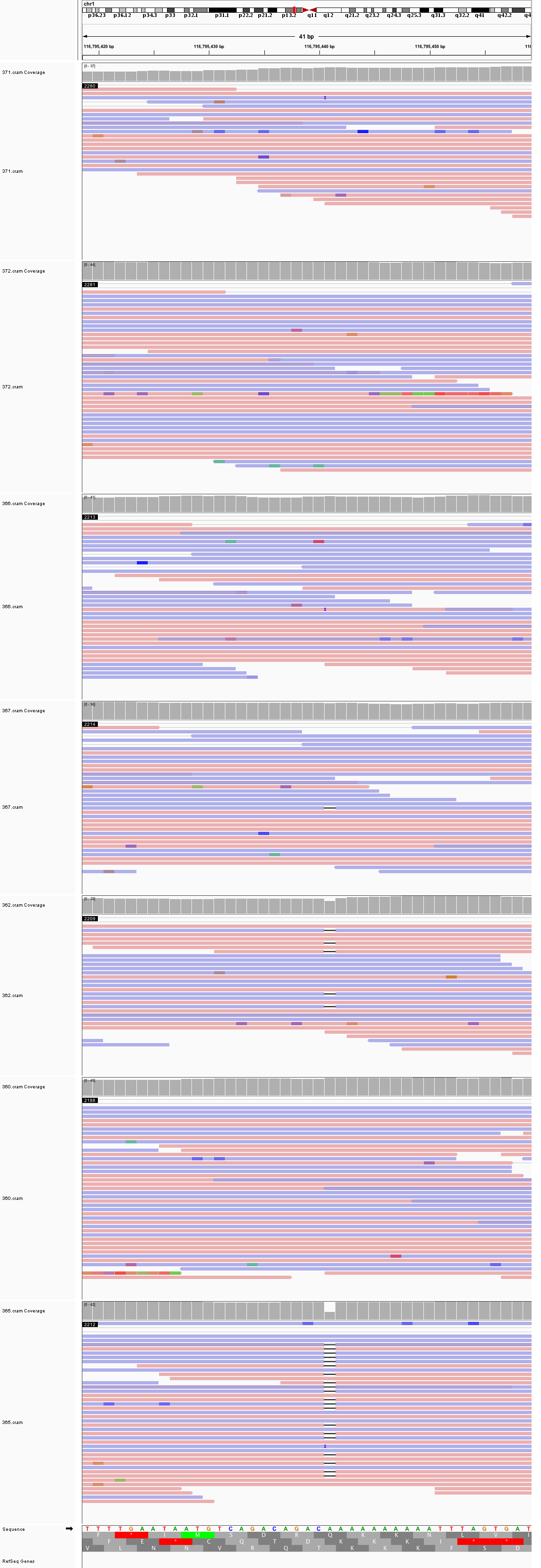

### chr1_174,370,058_174,370,098.png

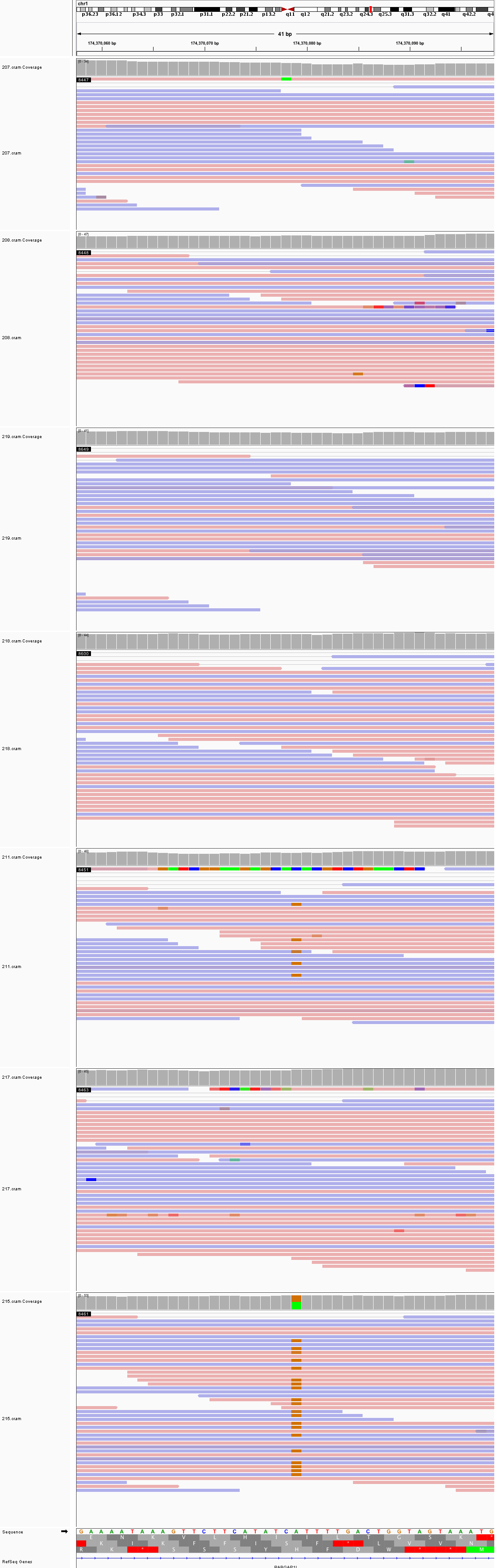

### chr1_190,673,786_190,673,826.png

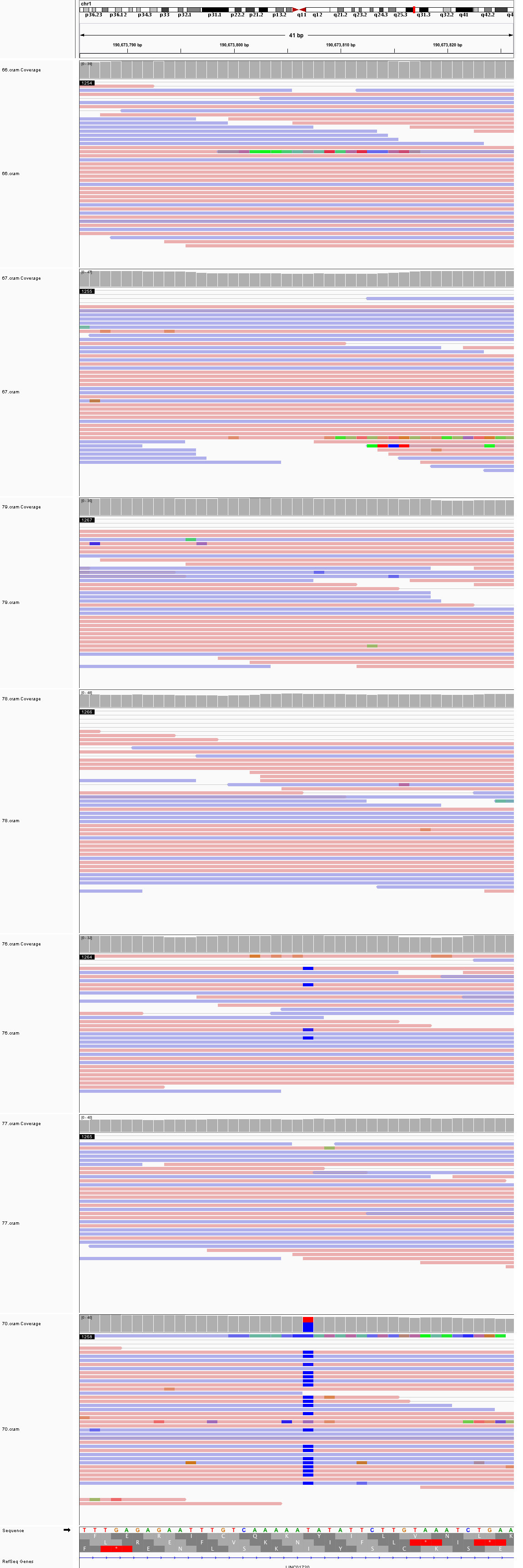

### chr2_13,392,169_13,392,209.png

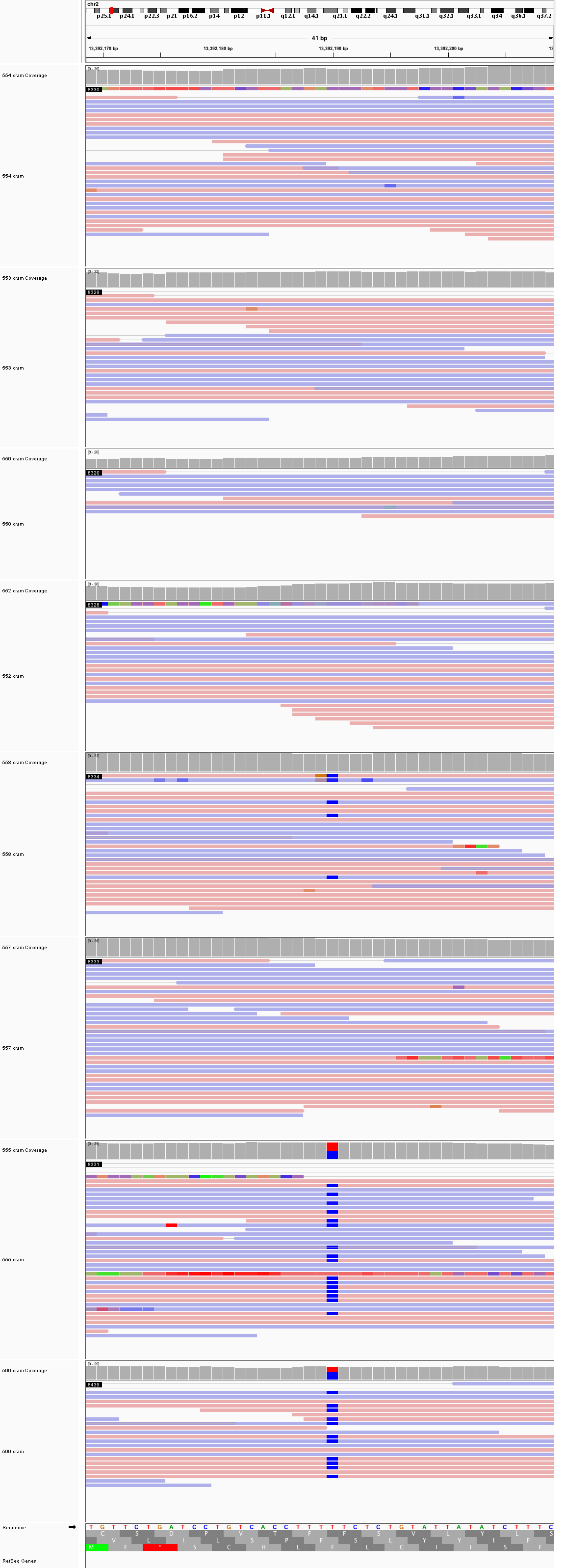

### chr2_34,560,497_34,560,537.png

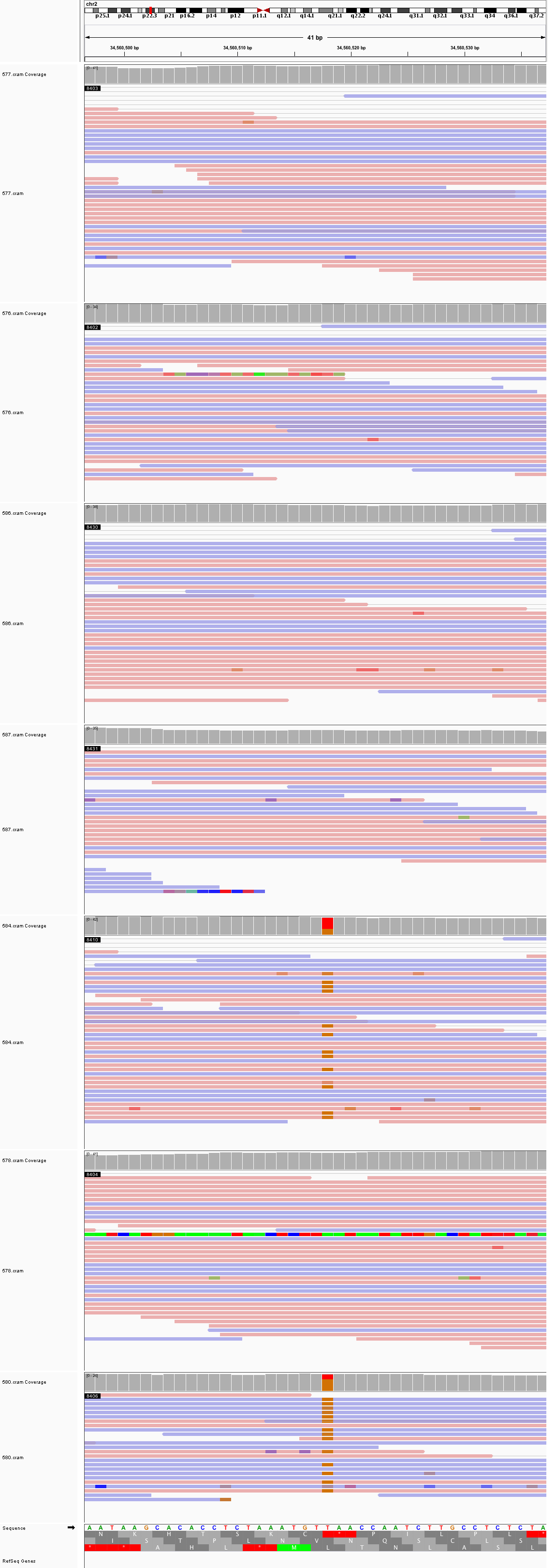

### chr2_39,220,392_39,220,432.png

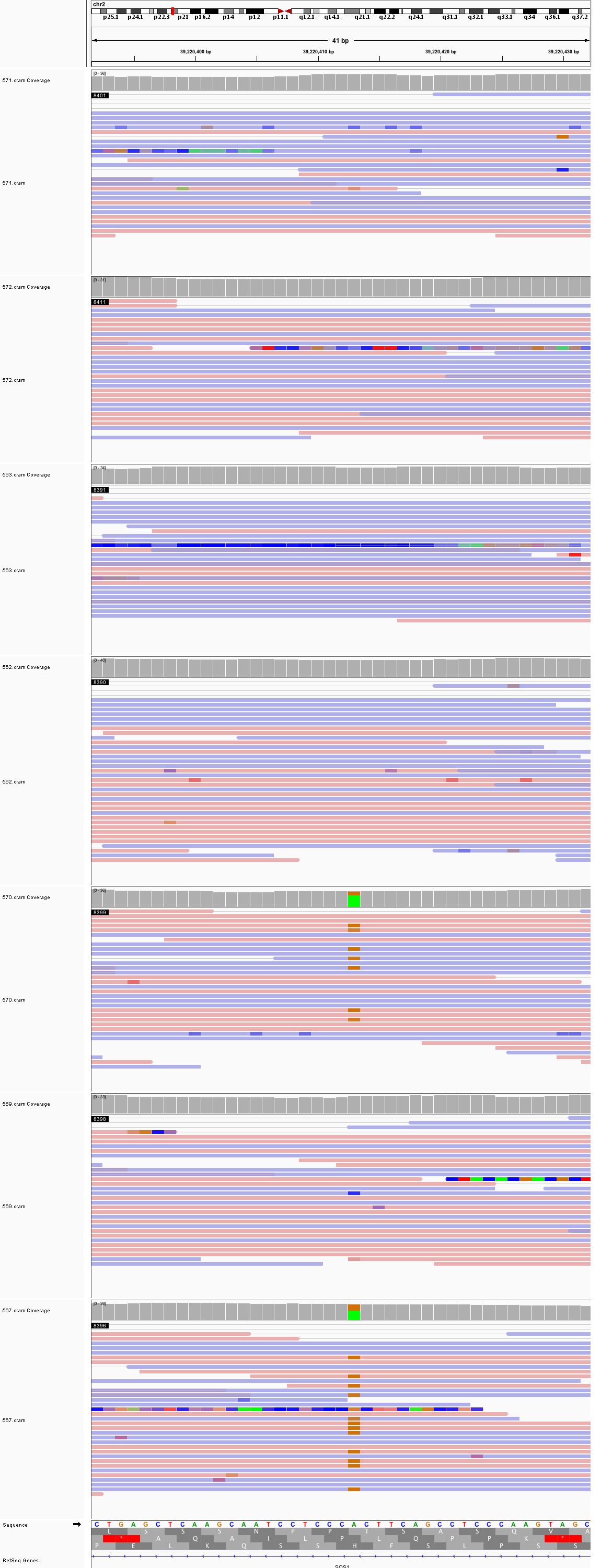

### chr2_58,177,158_58,177,198.png

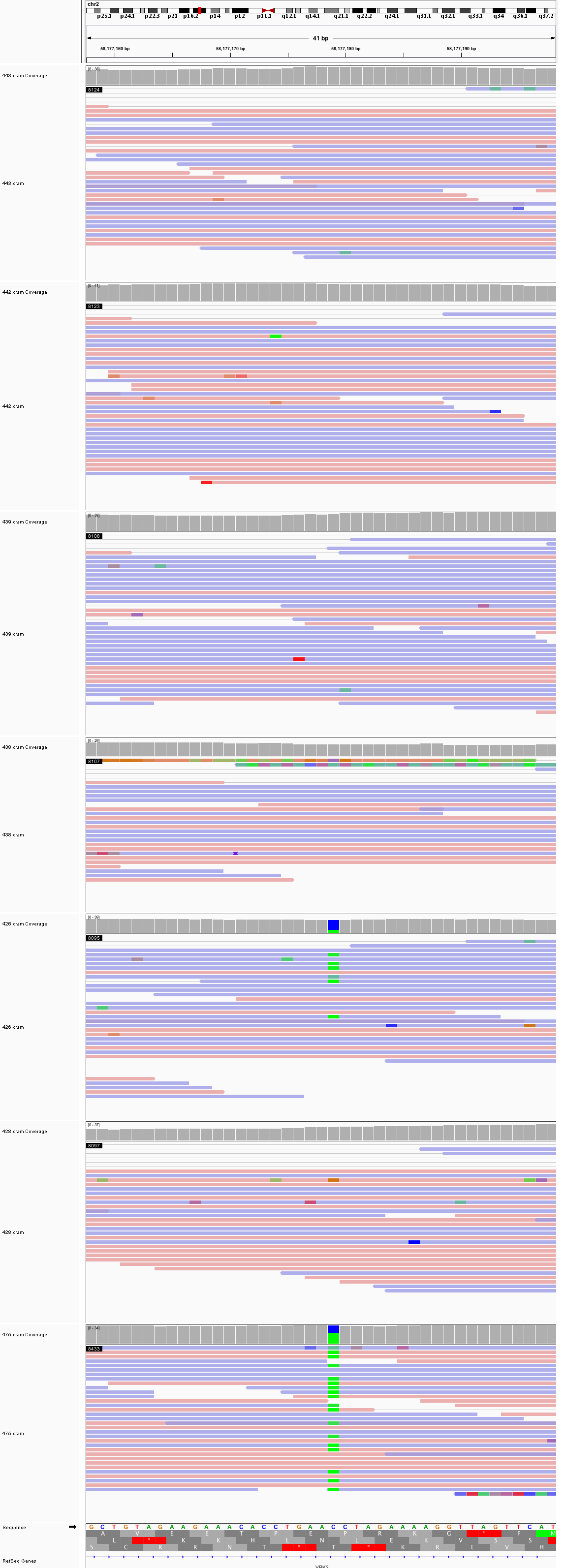

### chr2_127,193,708_127,193,748.png

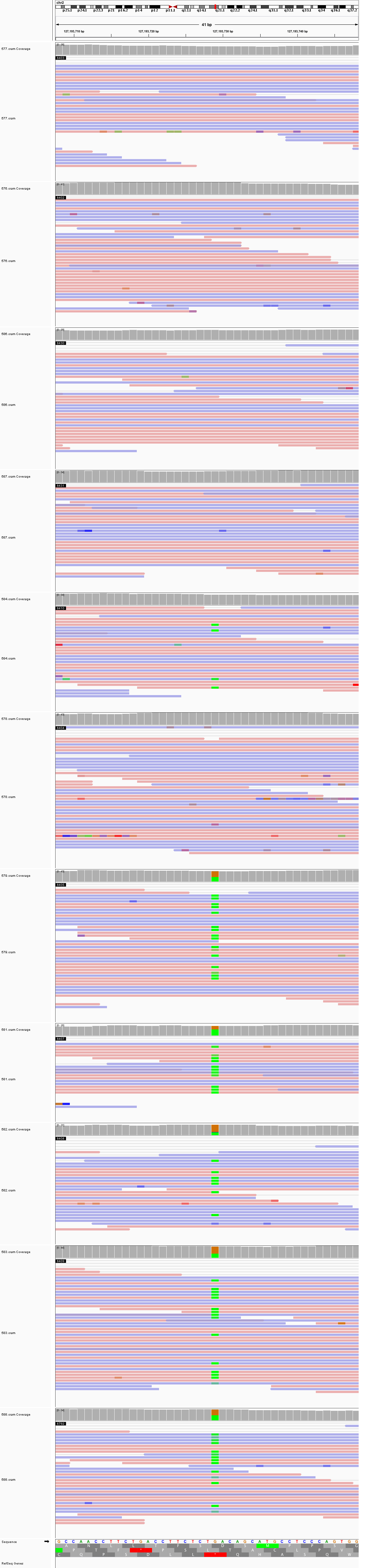

### chr2_134,141,556_134,141,596.png

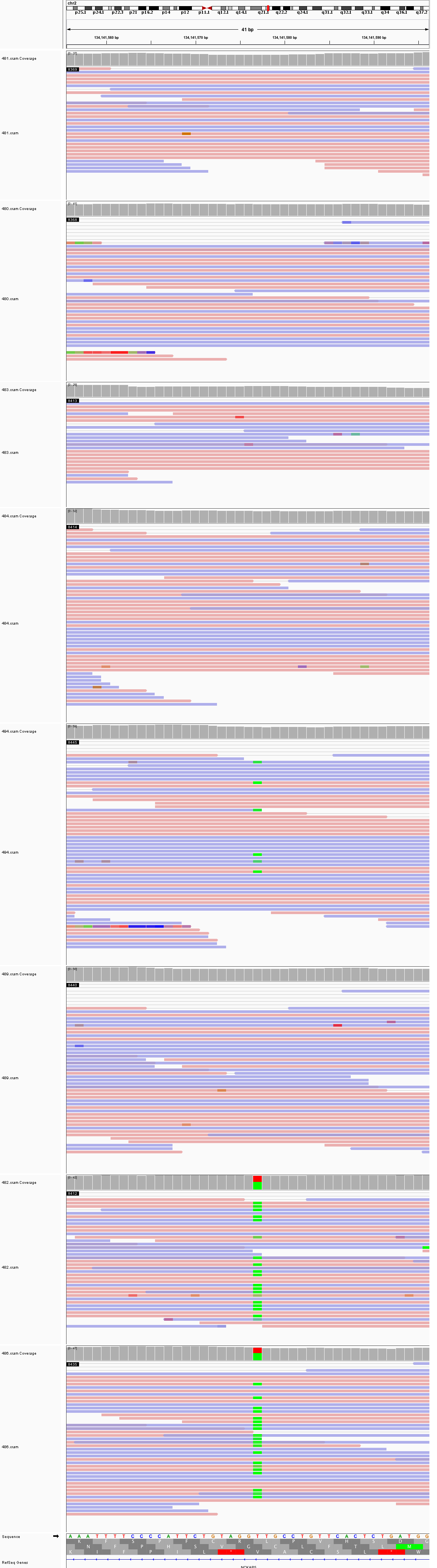

### chr2_138,289,943_138,289,983.png

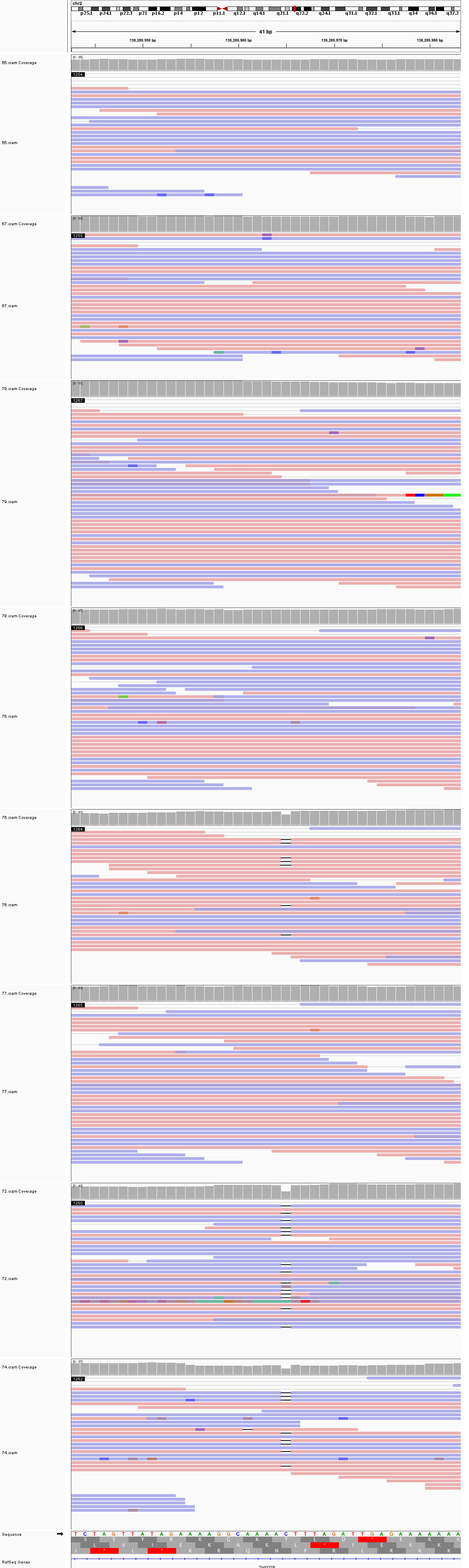

### chr2_191,347,892_191,347,932.png

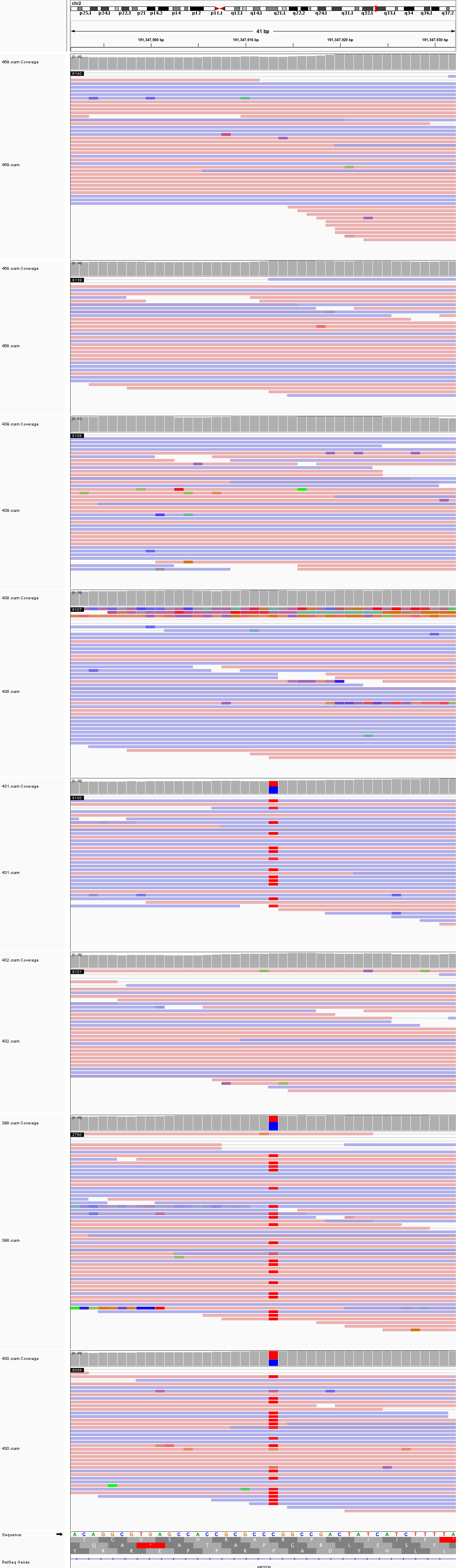

### chr2_194,980,719_194,980,759.png

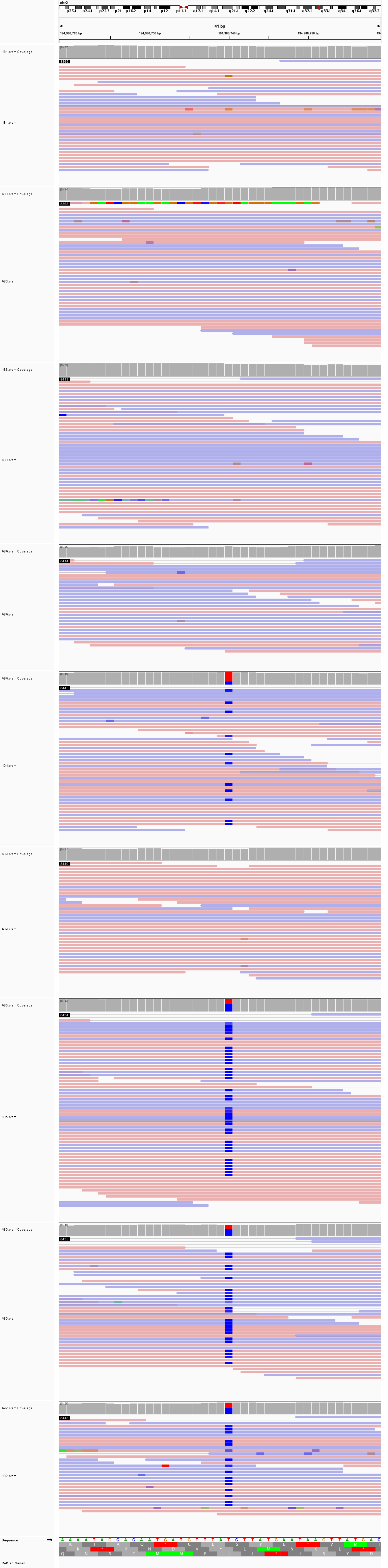

### chr2_222,009,741_222,009,781.png

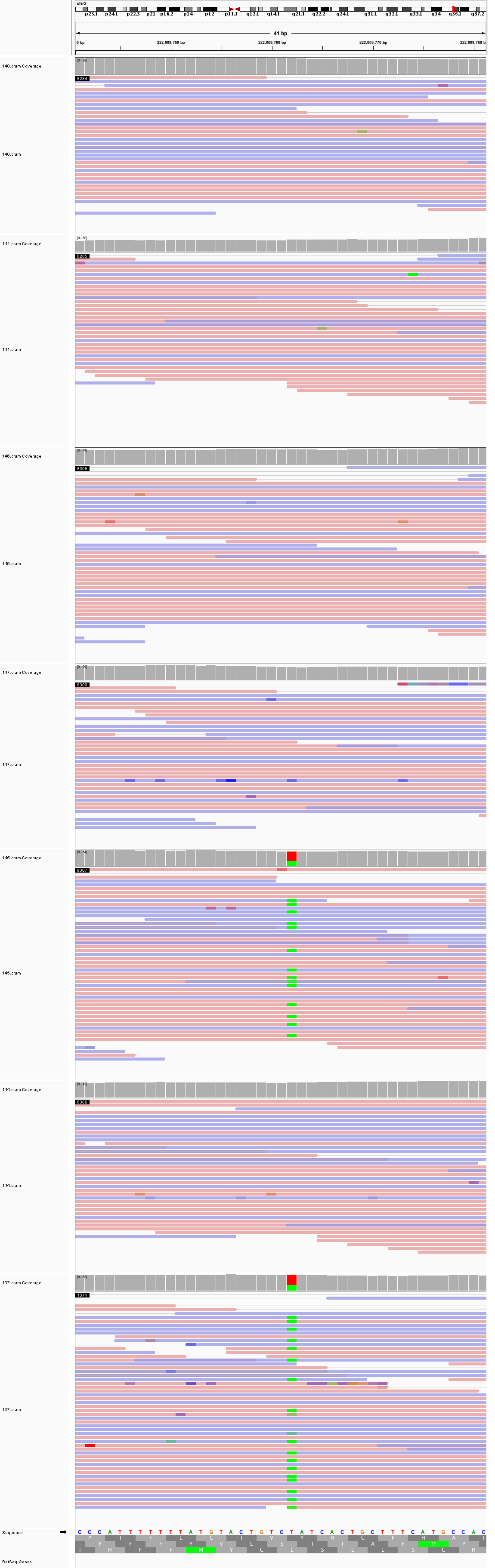

### chr2_232,345,972_232,346,012.png

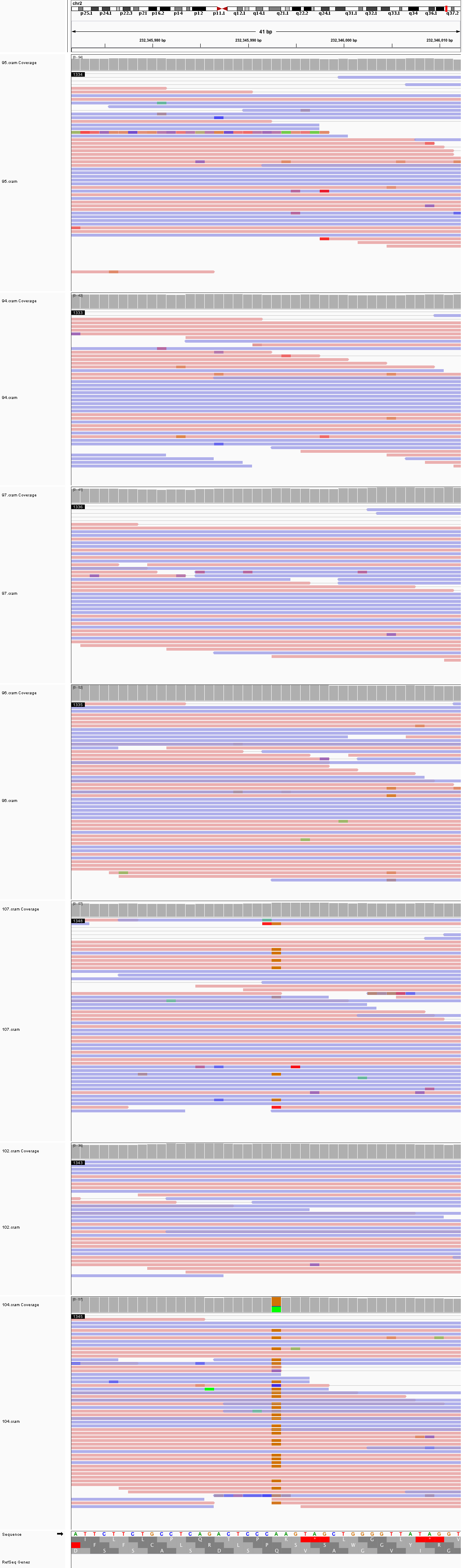

### chr2_242,728,136_242,728,176.png

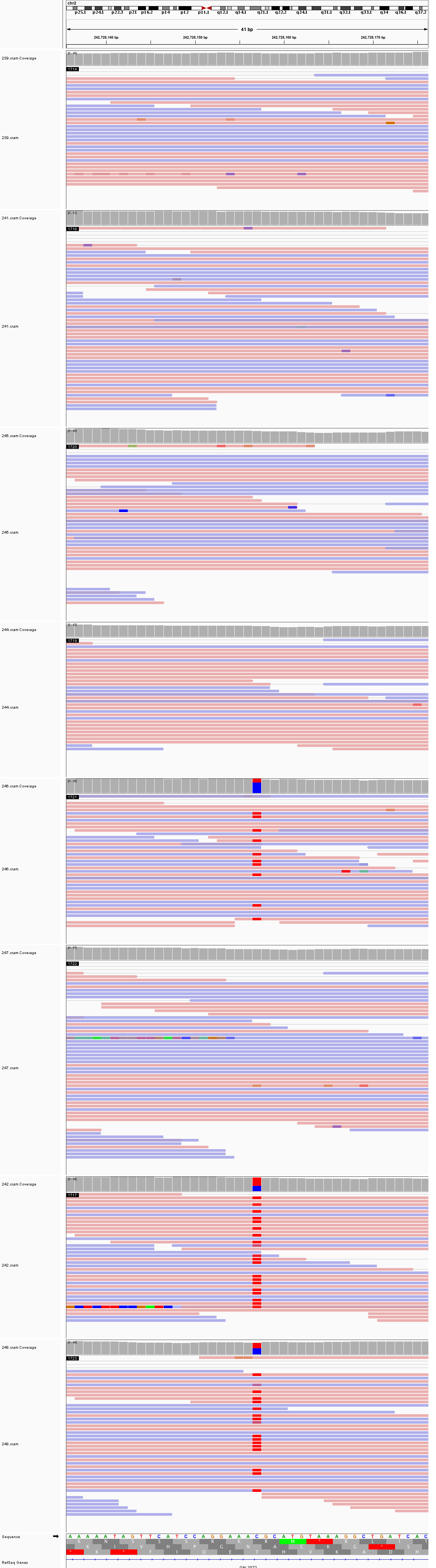

### chr3_33,970,919_33,970,959.png

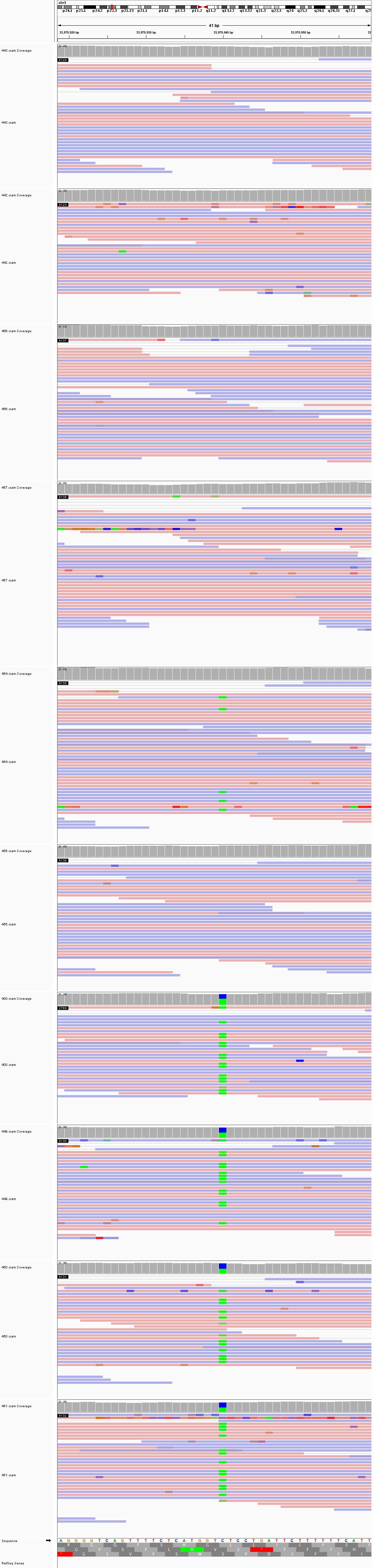

### chr3_34,525,067_34,525,107.png

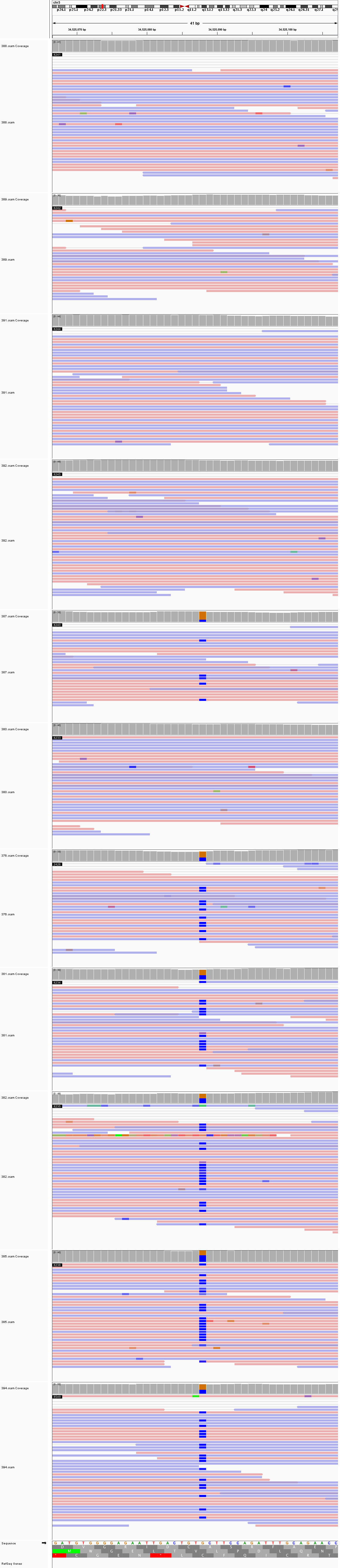

### chr3_36,975,262_36,975,302.png

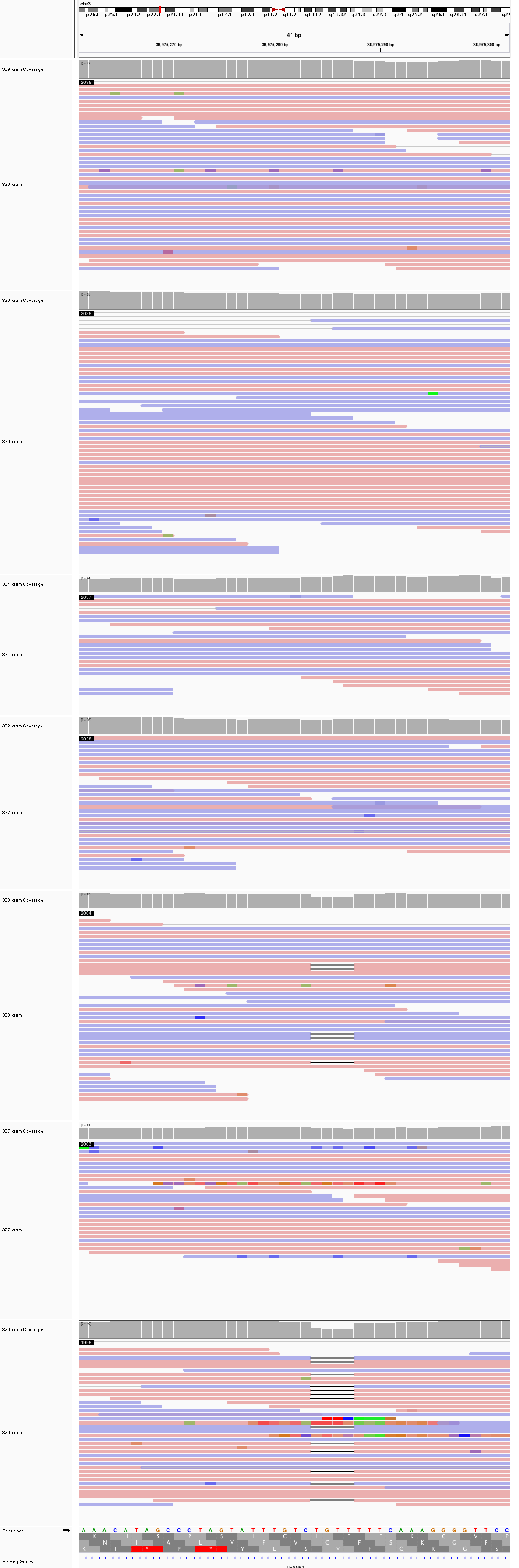

### chr3_40,754,331_40,754,371.png

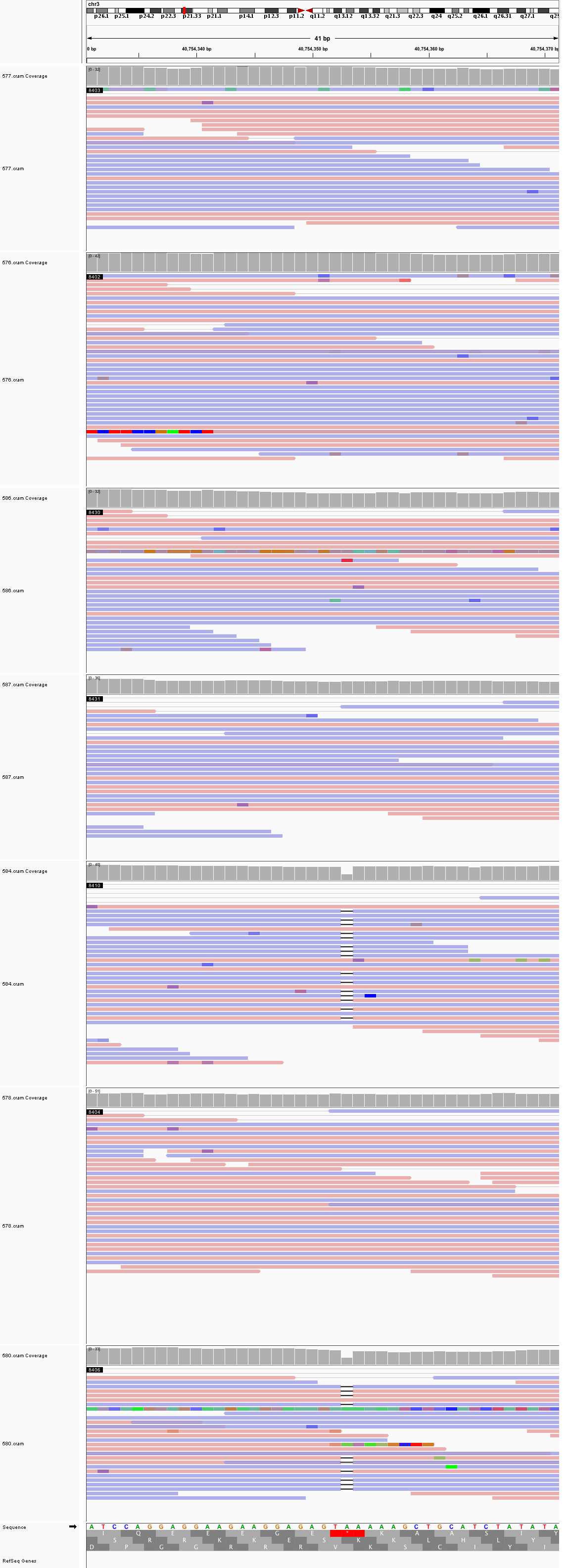

### chr3_56,565,904_56,565,944.png

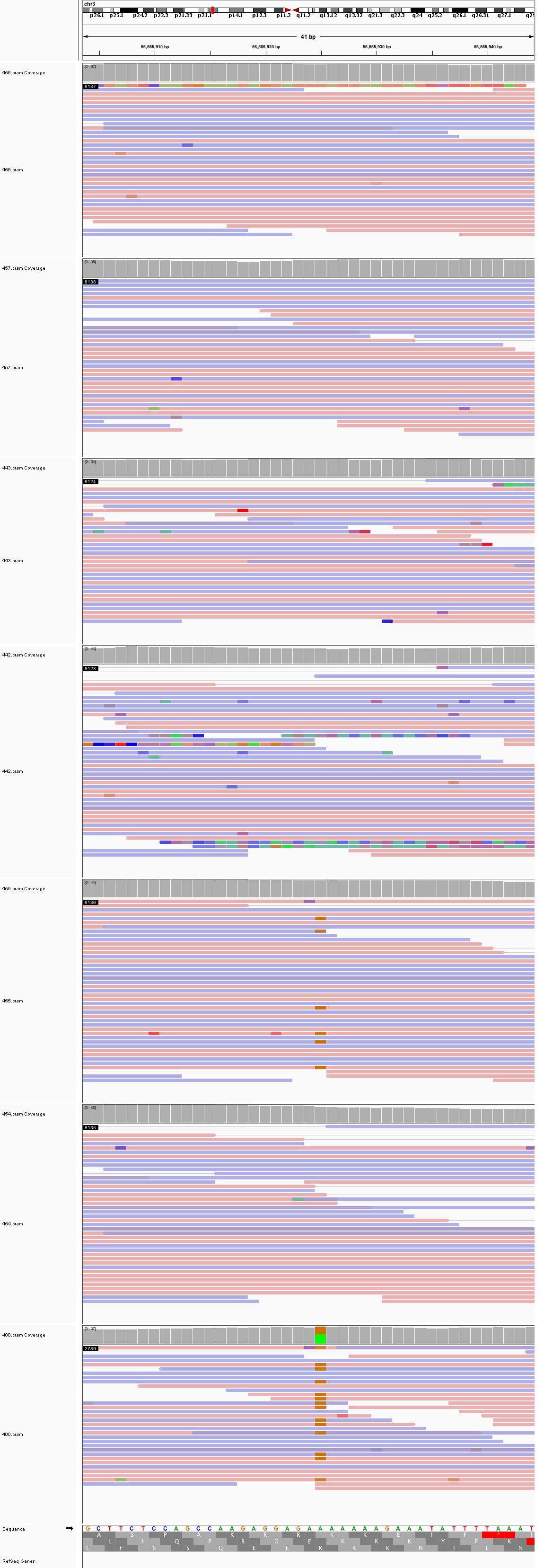

### chr3_56,998,730_56,998,770.png

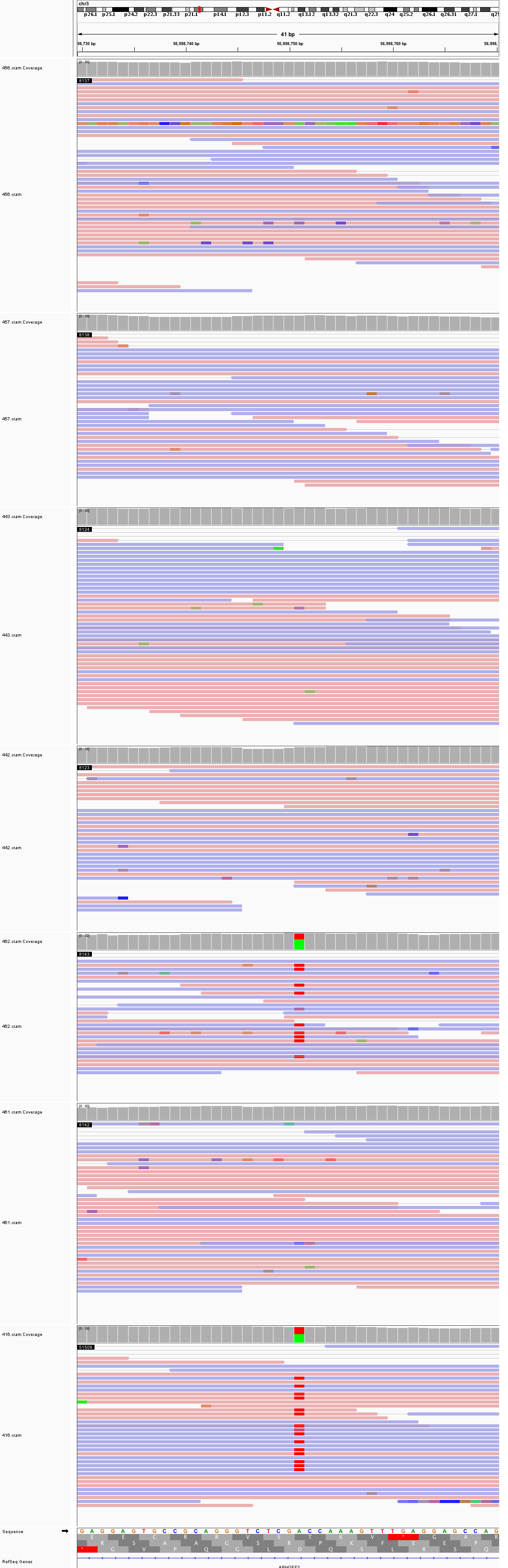

### chr3_83,453,550_83,453,590.png

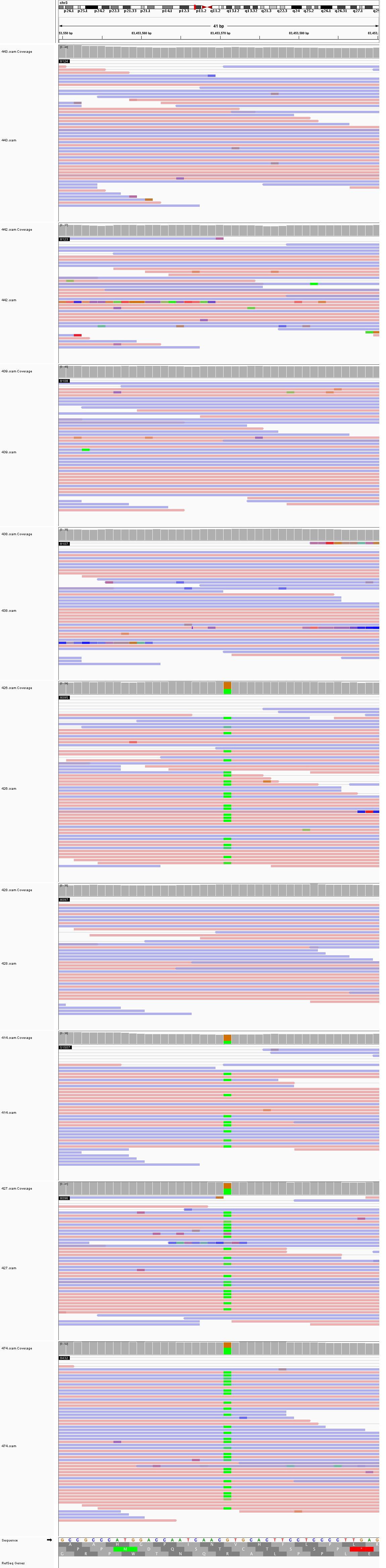

### chr3_128,755,614_128,755,654.png

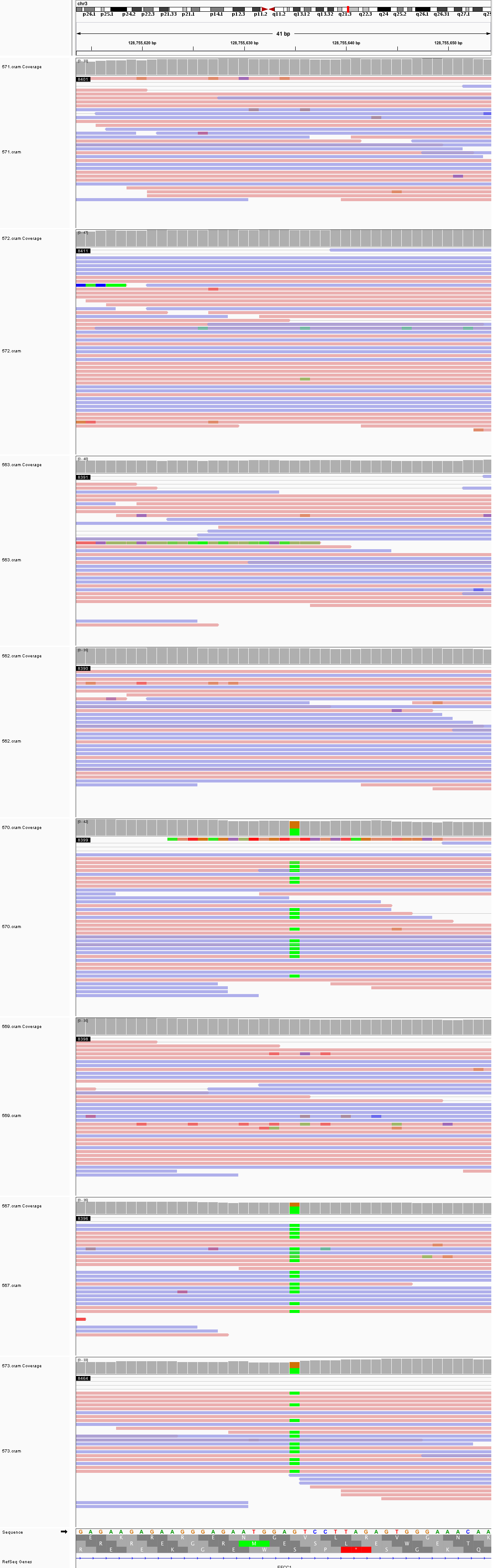

### chr3_185,177,827_185,177,867.png

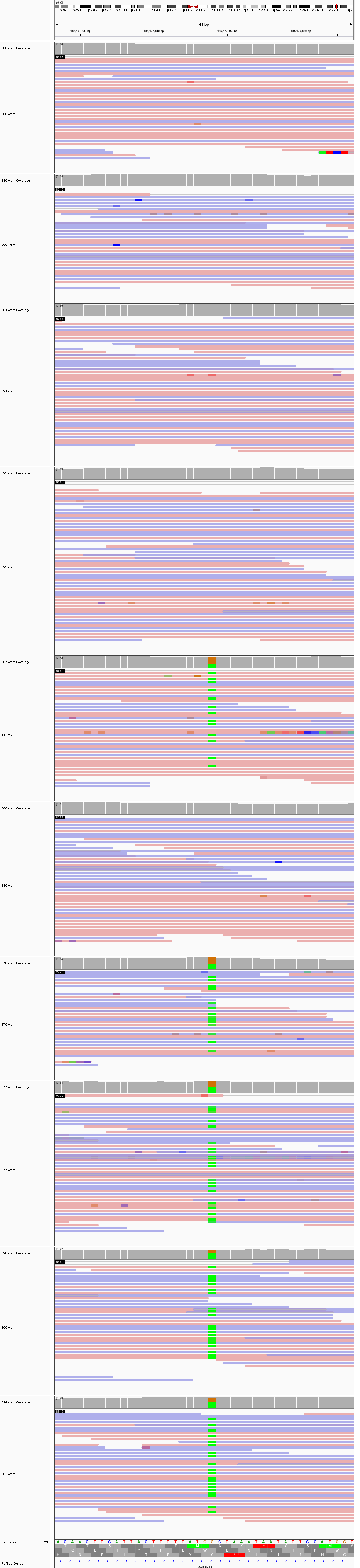

### chr4_29,859,158_29,859,198.png

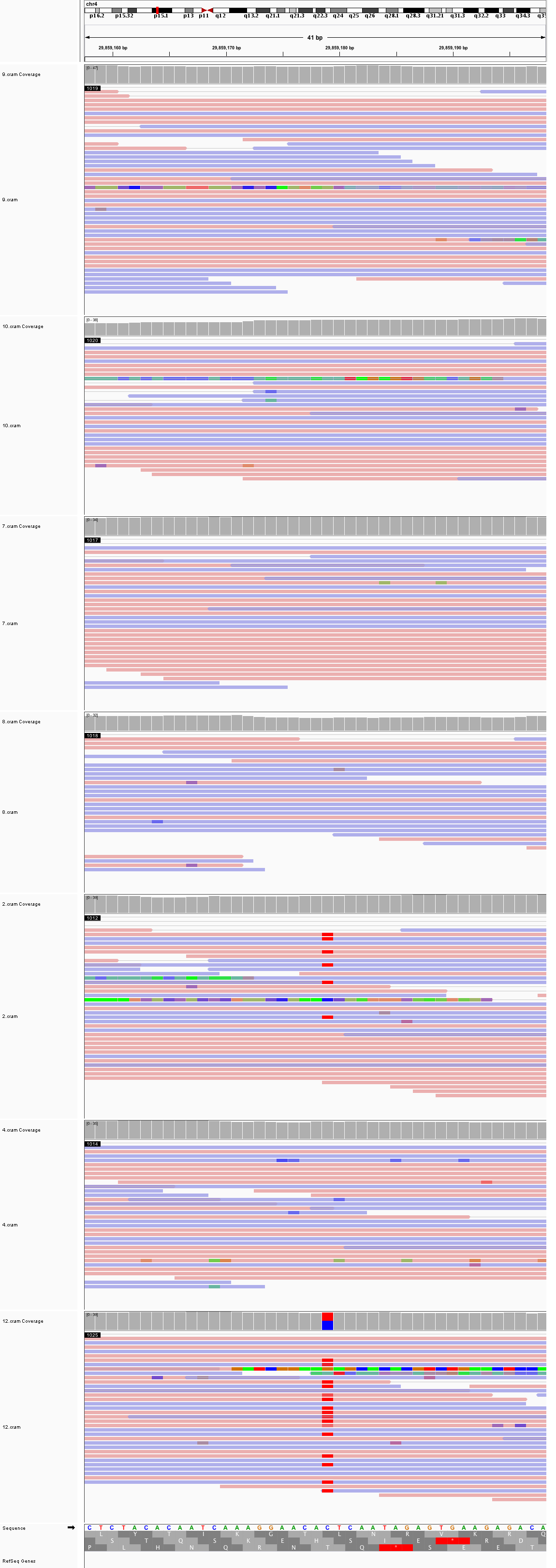

### chr4_57,912,958_57,912,998.png

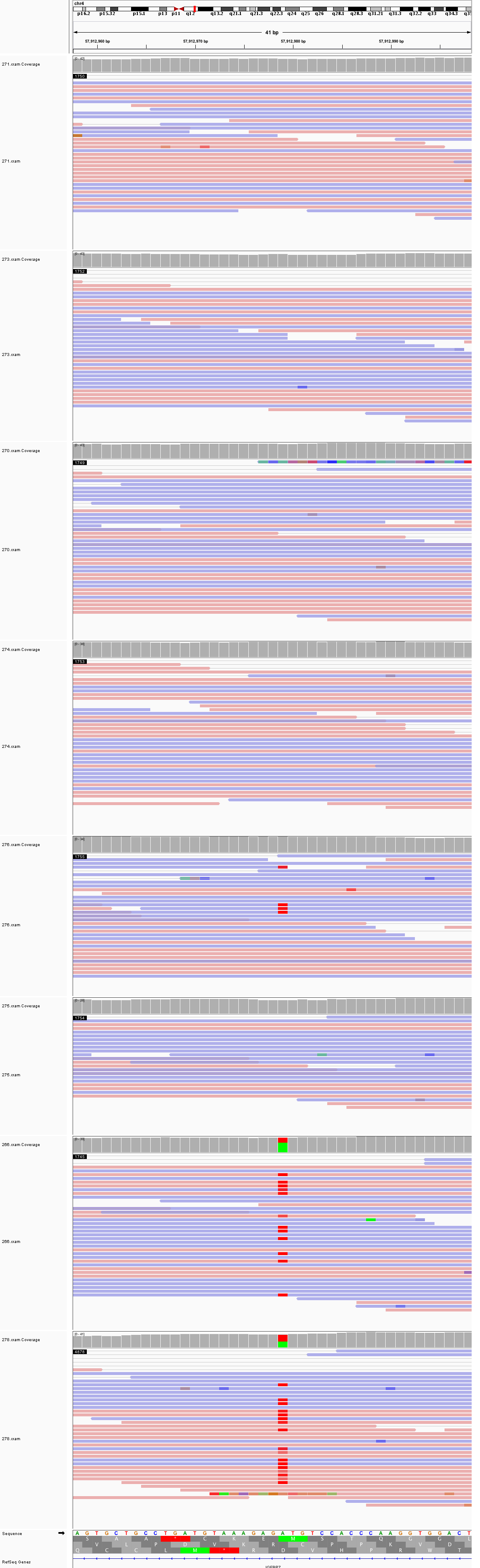

### chr4_66,176,885_66,176,925.png

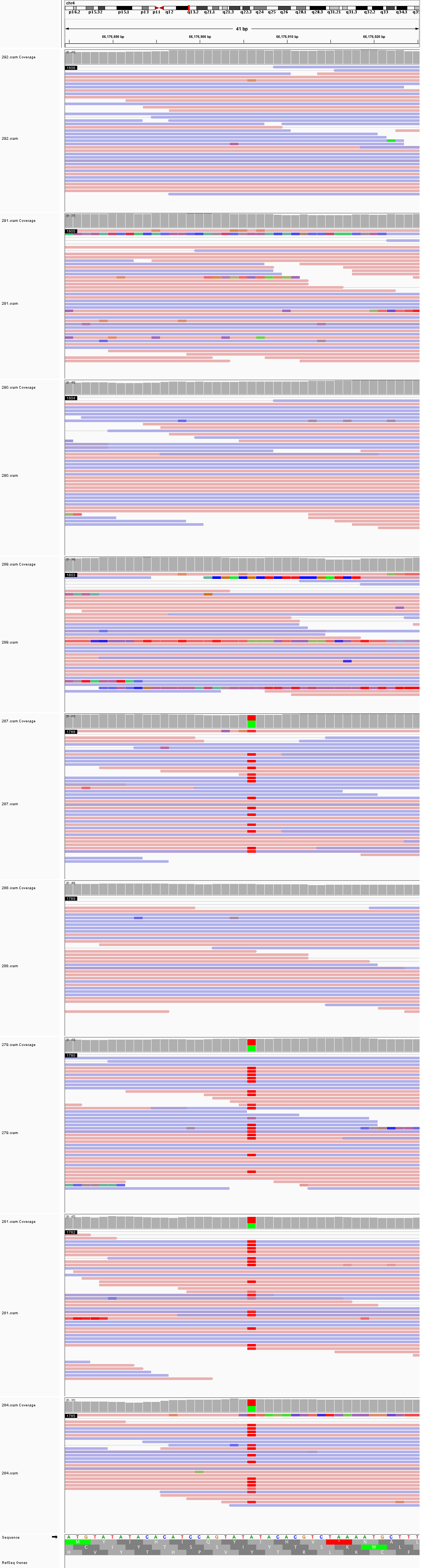

### chr4_137,046,587_137,046,627.png

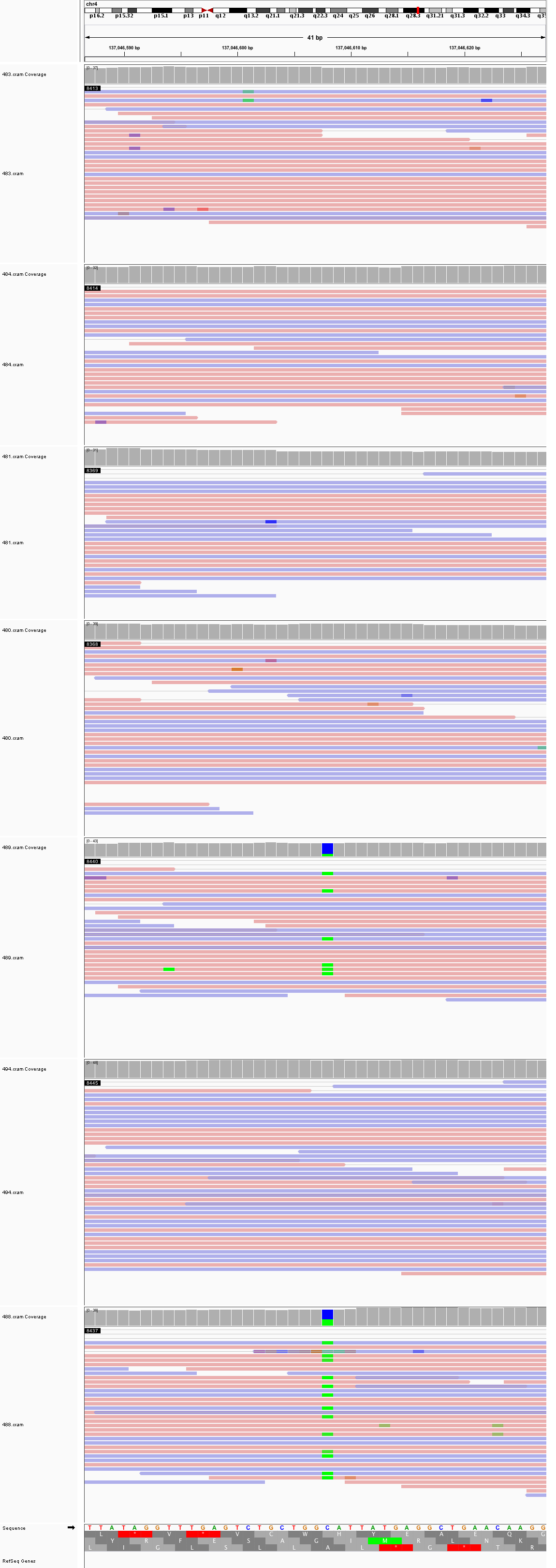

### chr4_155,983,436_155,983,476.png

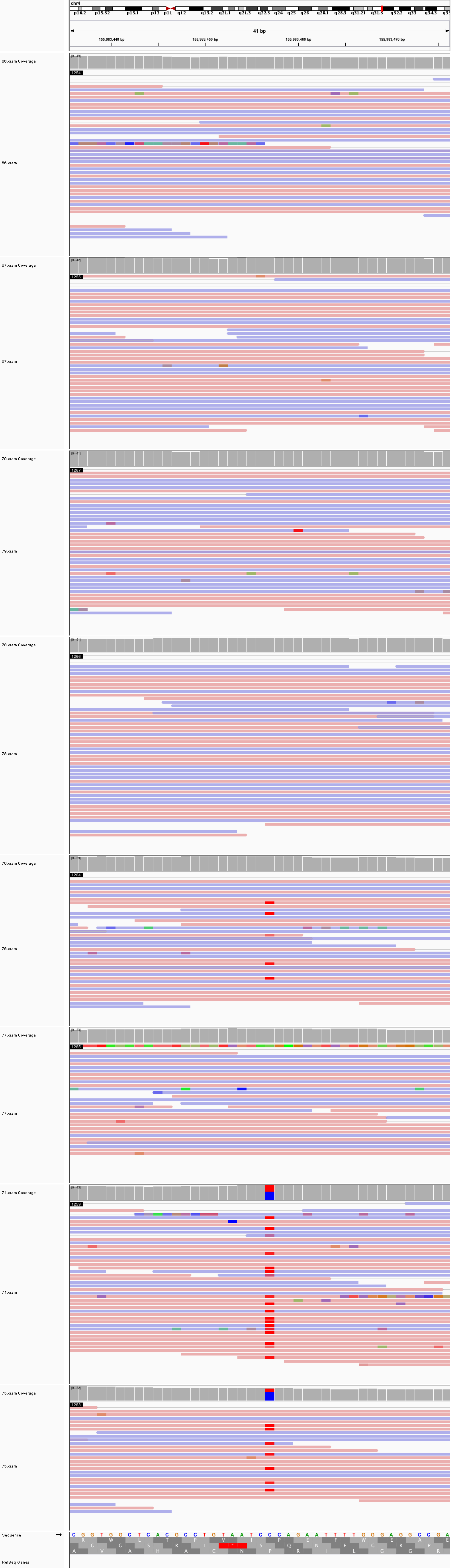

### chr5_12,788,216_12,788,256.png

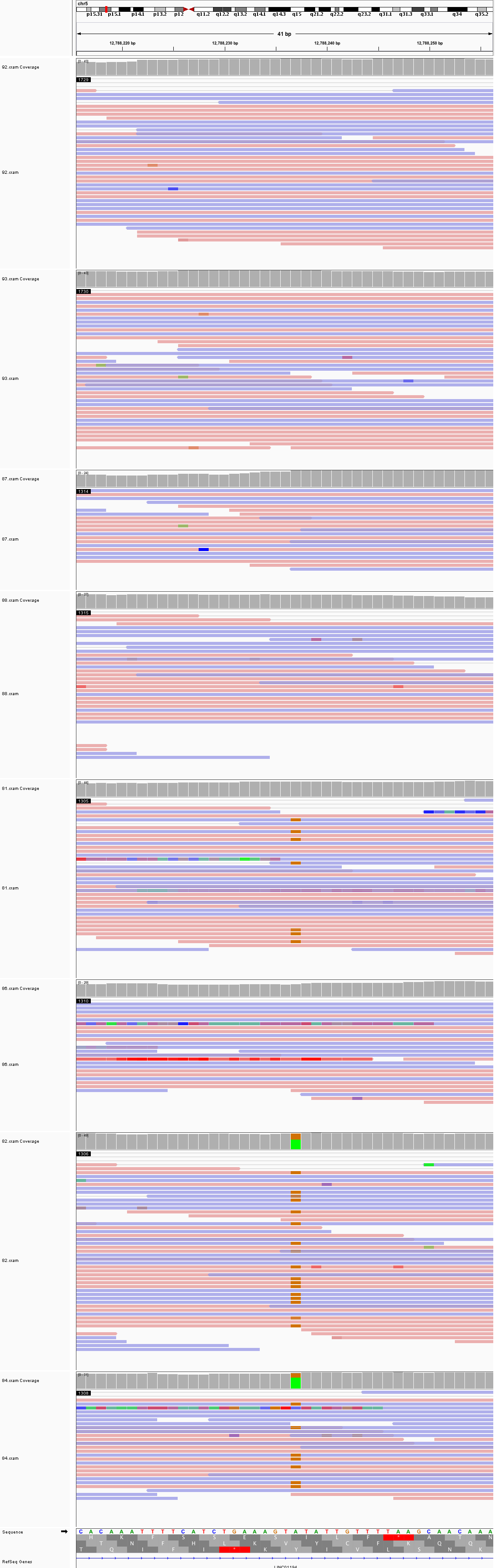

### chr5_16,336,785_16,336,825.png

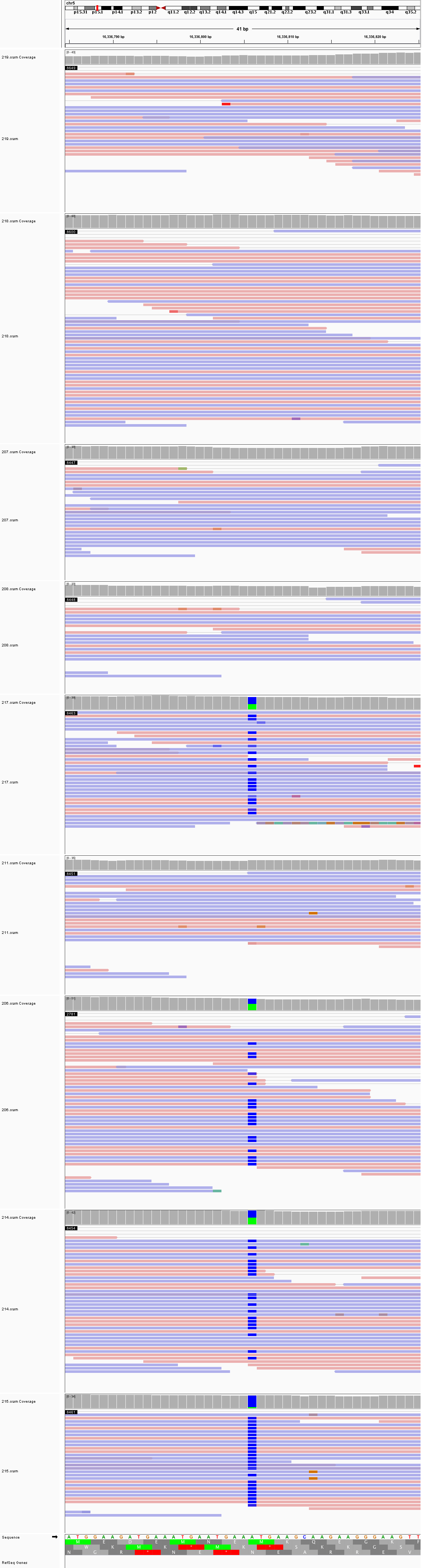

### chr5_54,558,810_54,558,850.png

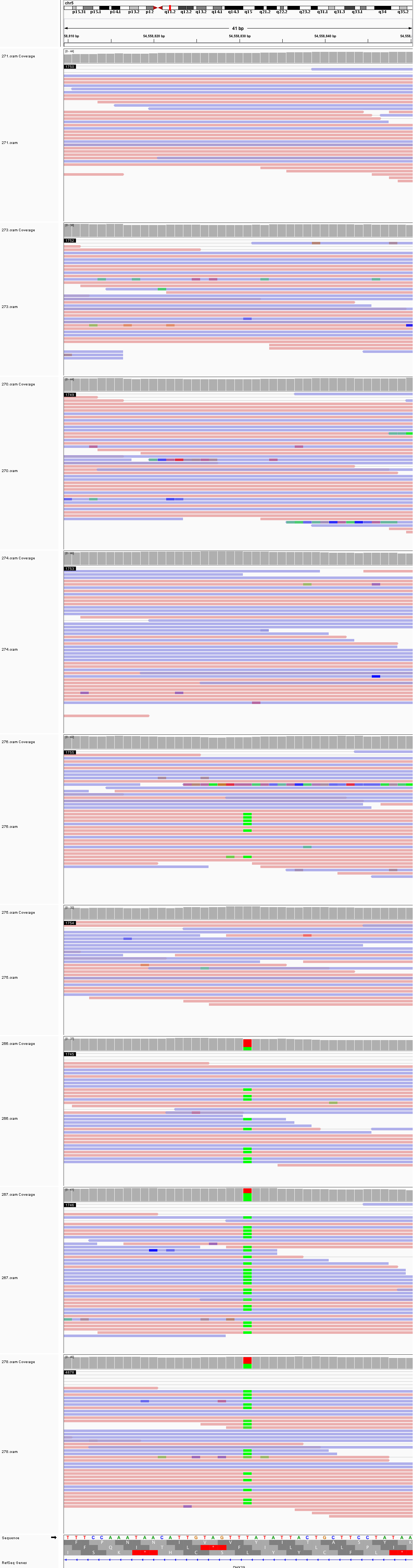

### chr5_66,236,934_66,236,974.png

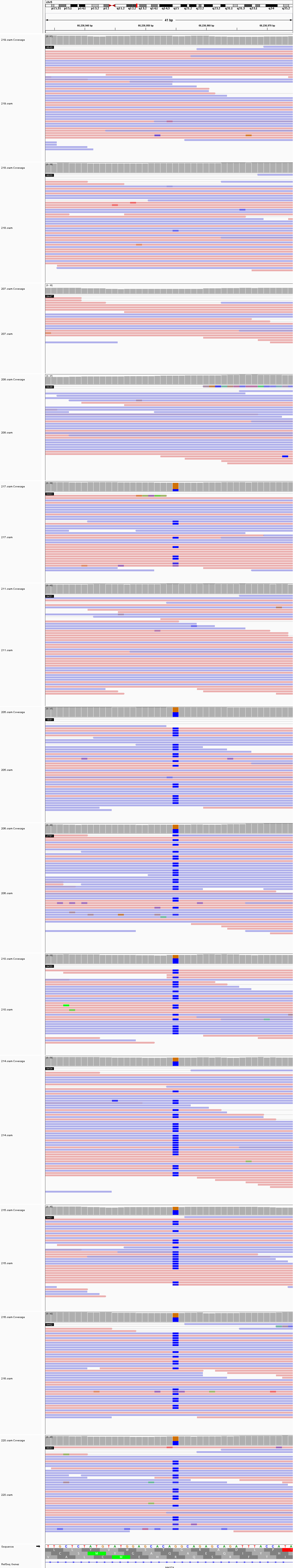

### chr5_101,455,325_101,455,365.png

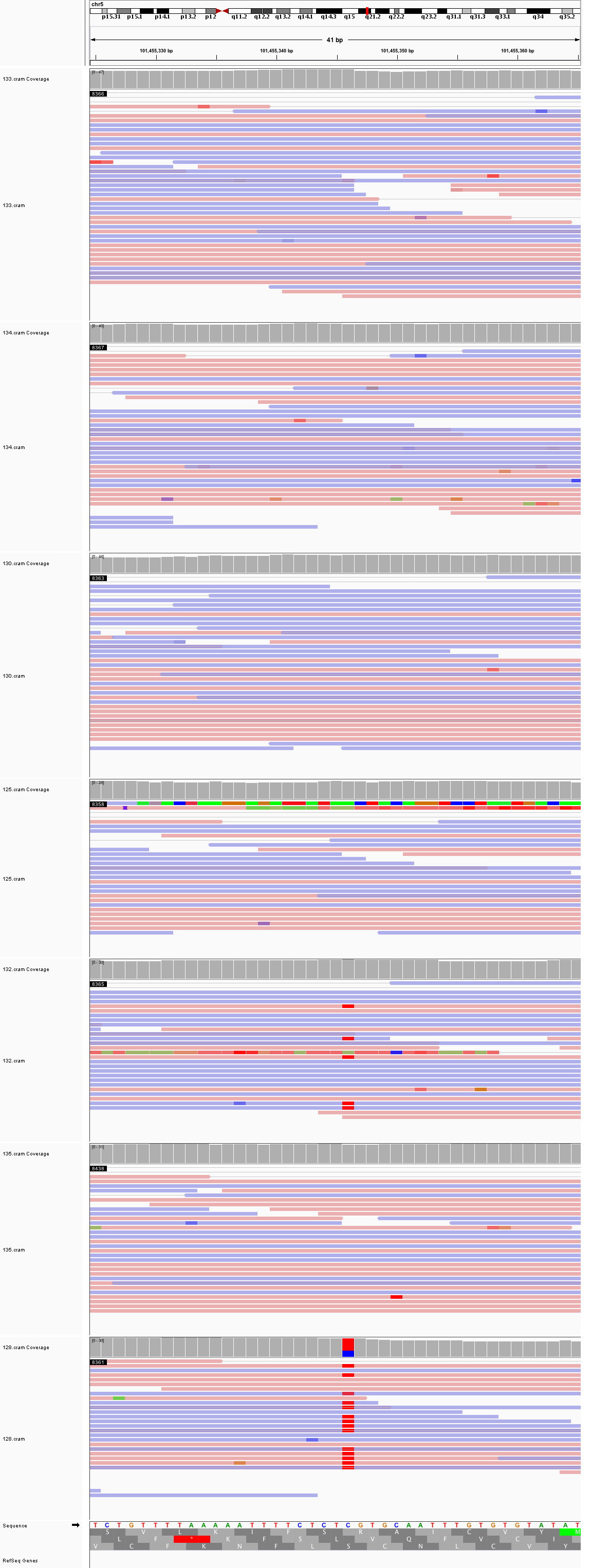

### chr5_107,090,902_107,090,942.png

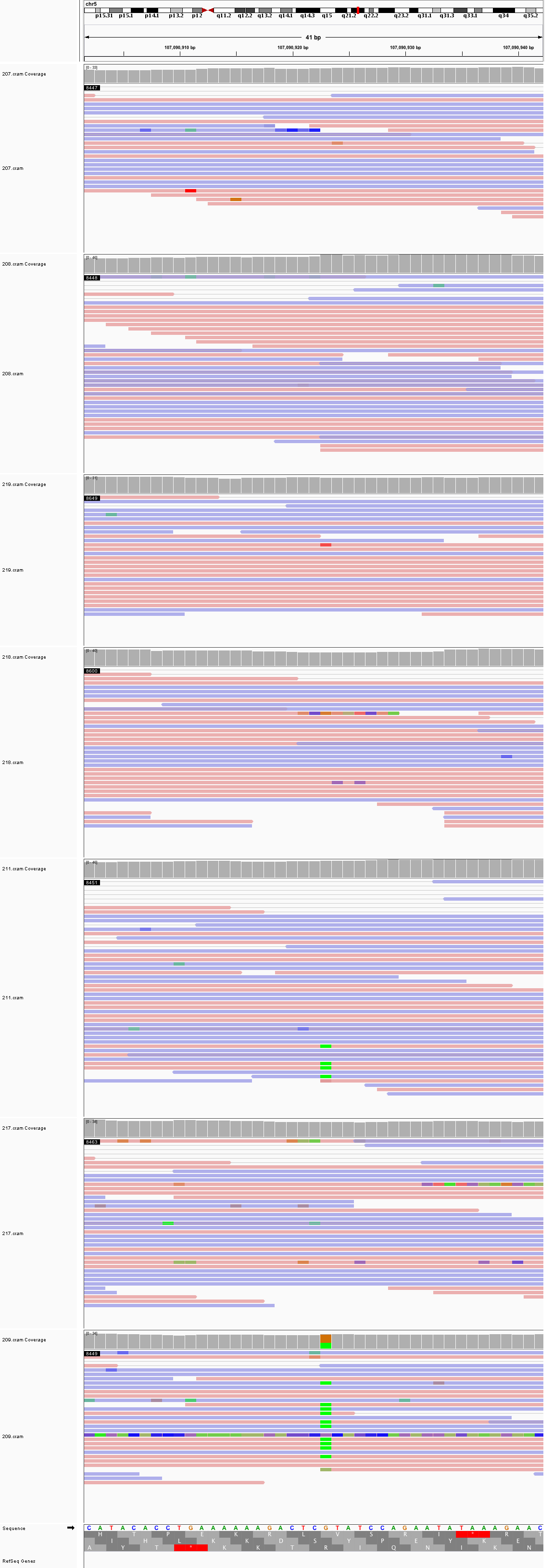

### chr5_145,826,986_145,827,026.png

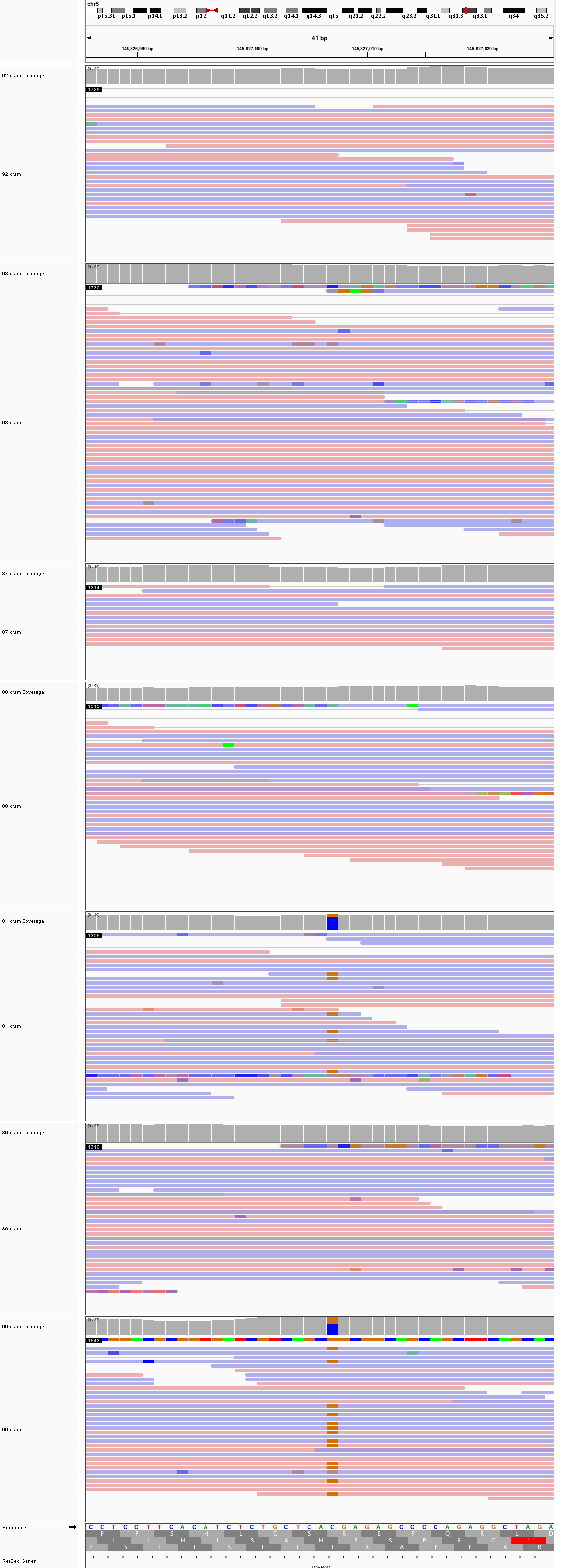

### chr5_147,256,386_147,256,426.png

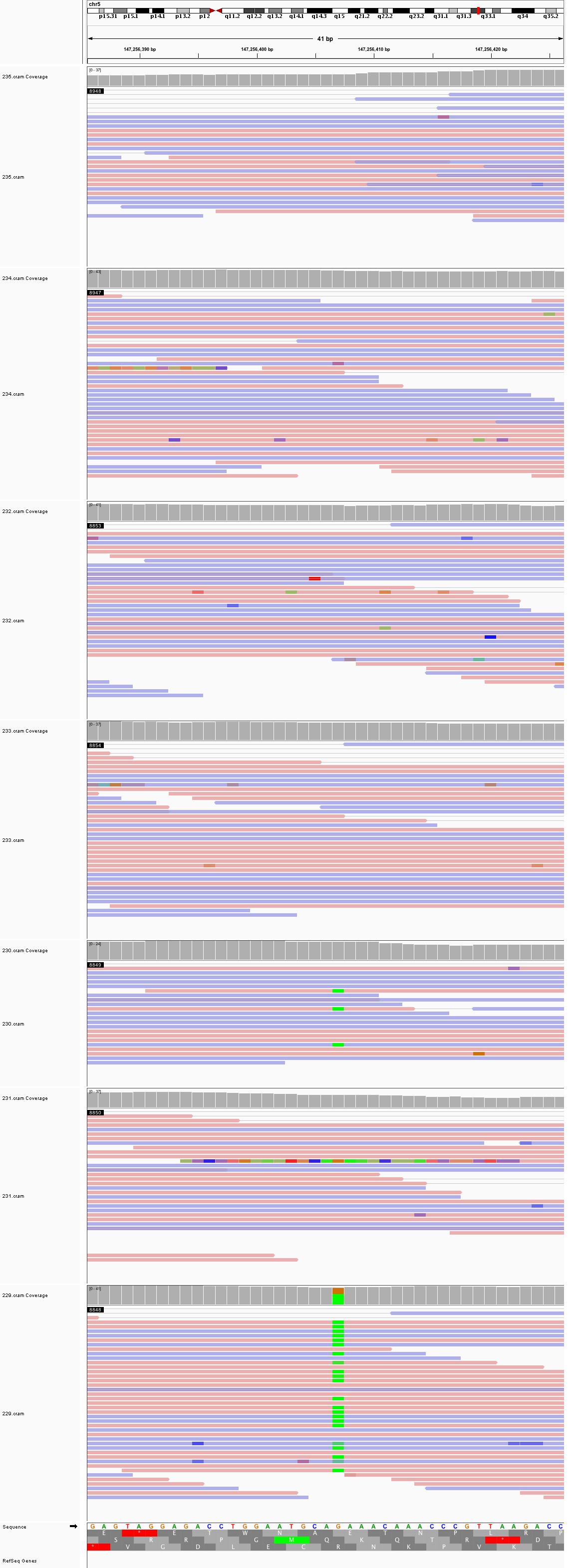

### chr5_156,167,712_156,167,752.png

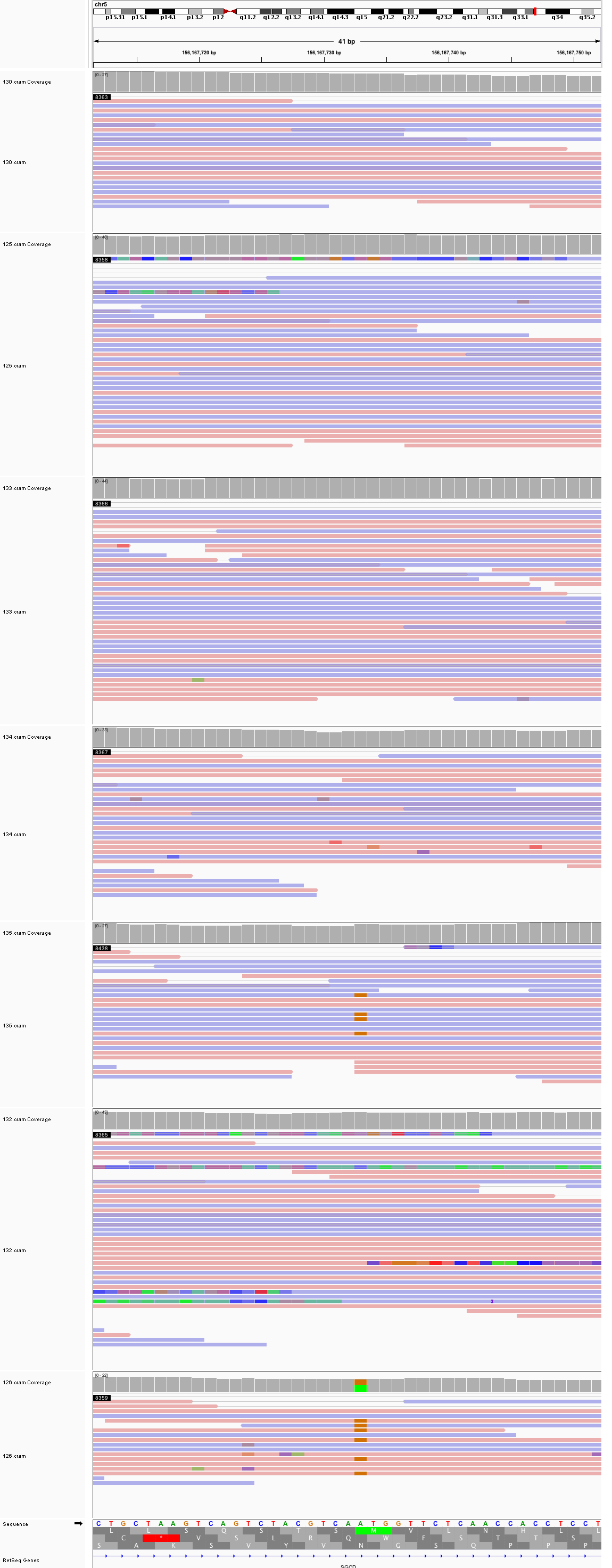

### chr5_161,959,751_161,959,791.png

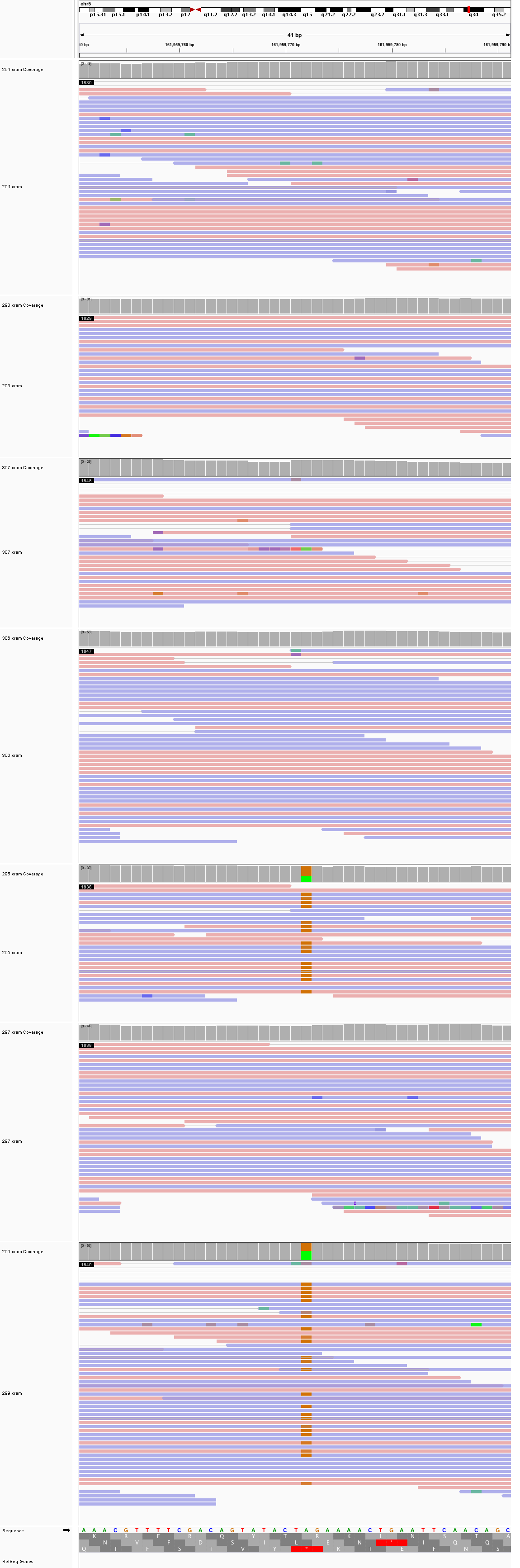

### chr6_33,872,122_33,872,162.png

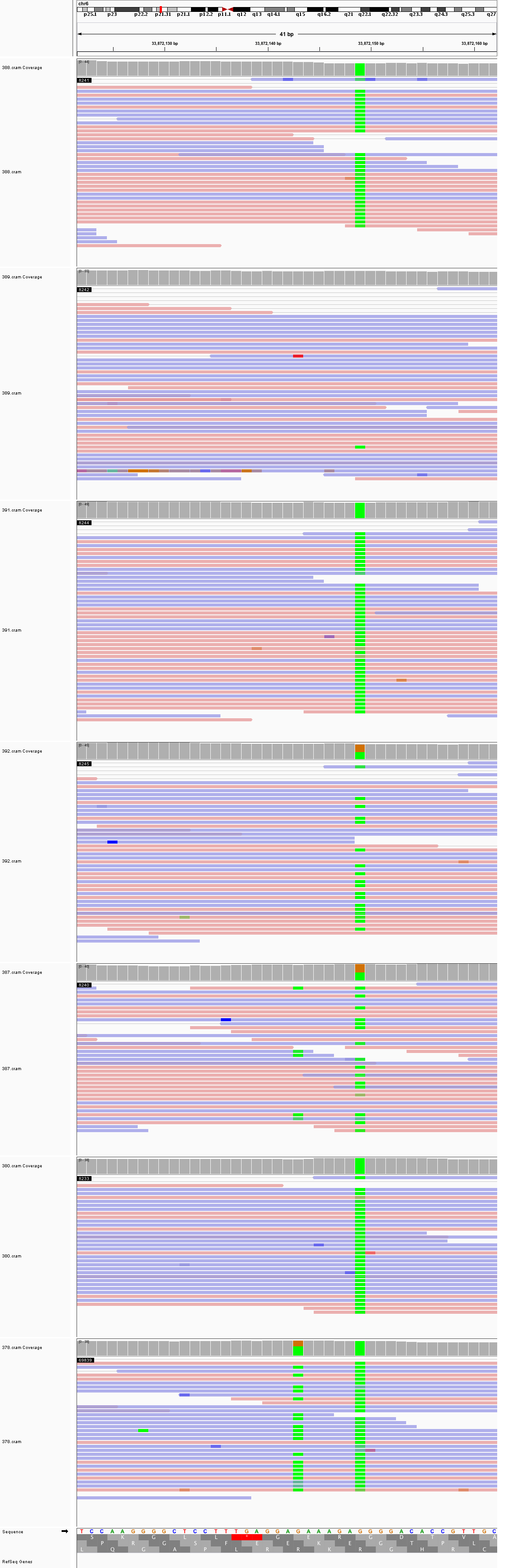

### chr6_55,283,462_55,283,502.png

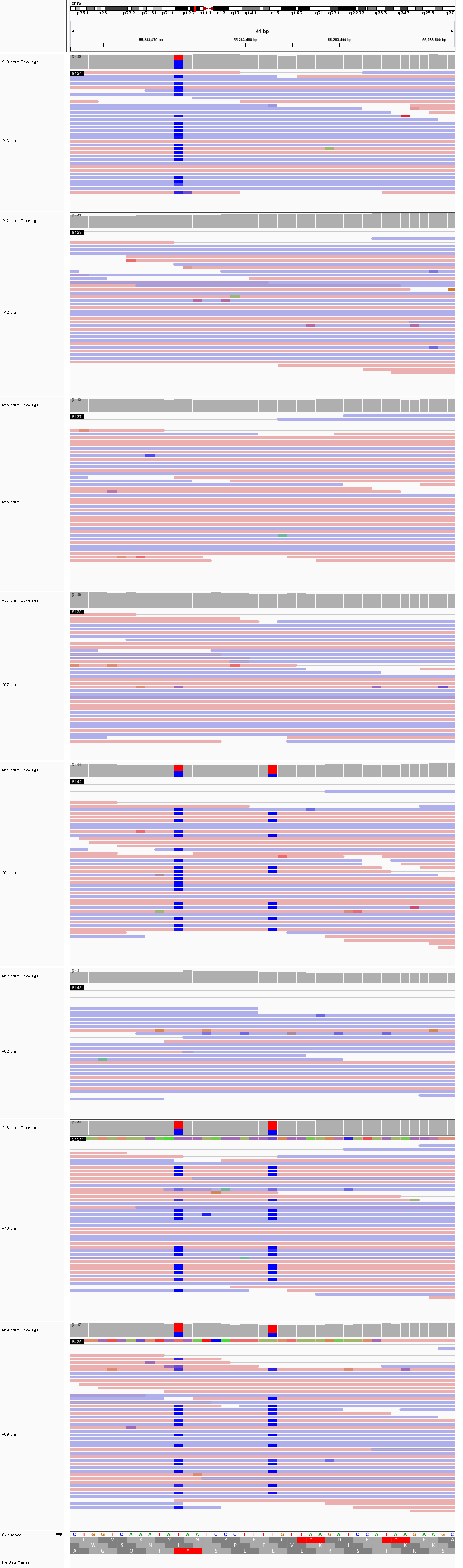

### chr6_95,232,105_95,232,145.png

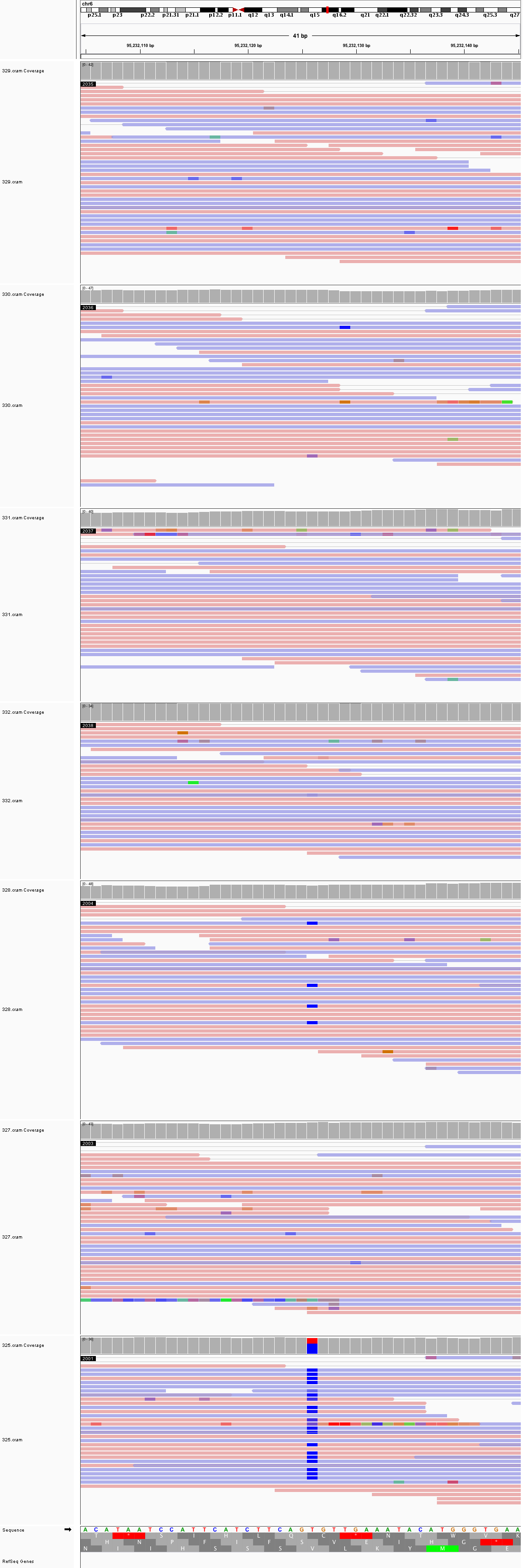

### chr6_107,038,447_107,038,487.png

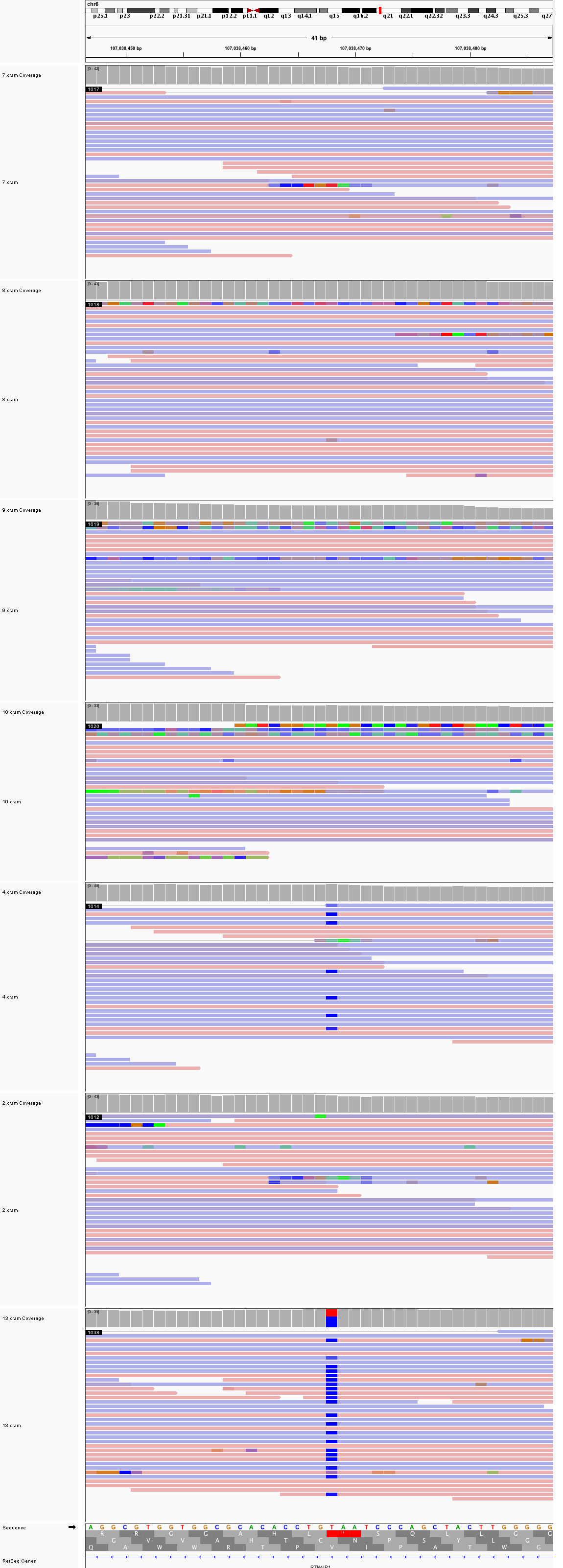

### chr6_118,123,453_118,123,493.png

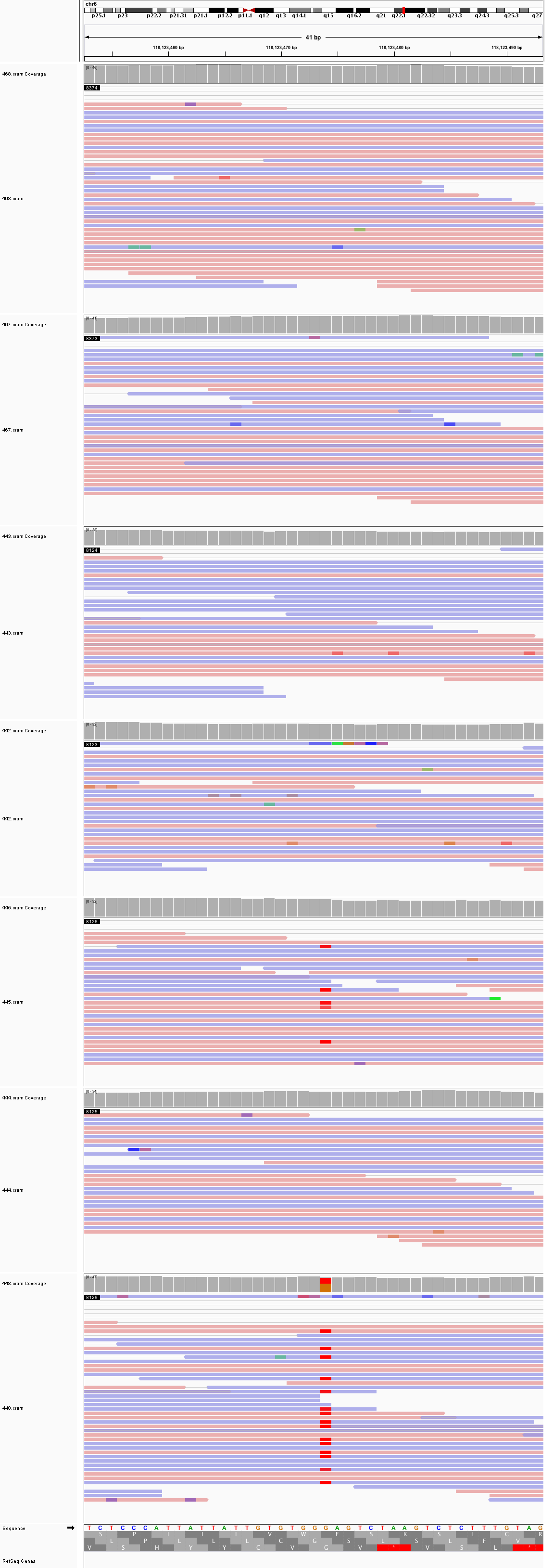

### chr7_11,681,791_11,681,831.png

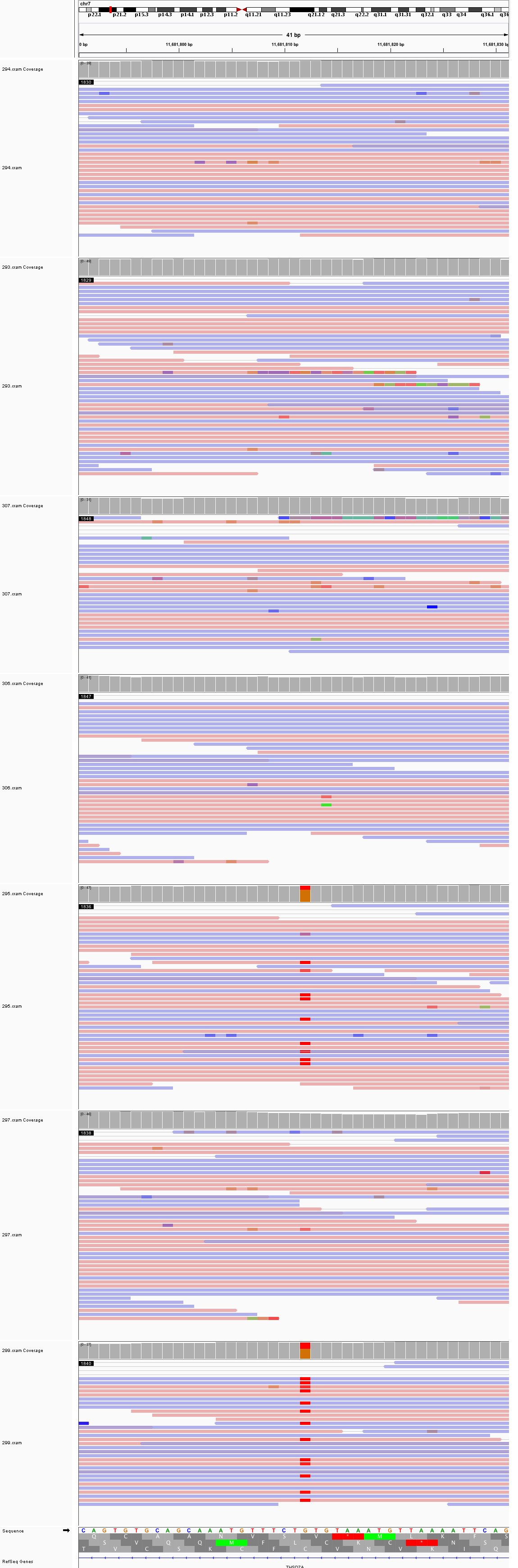

### chr7_16,051,999_16,052,039.png

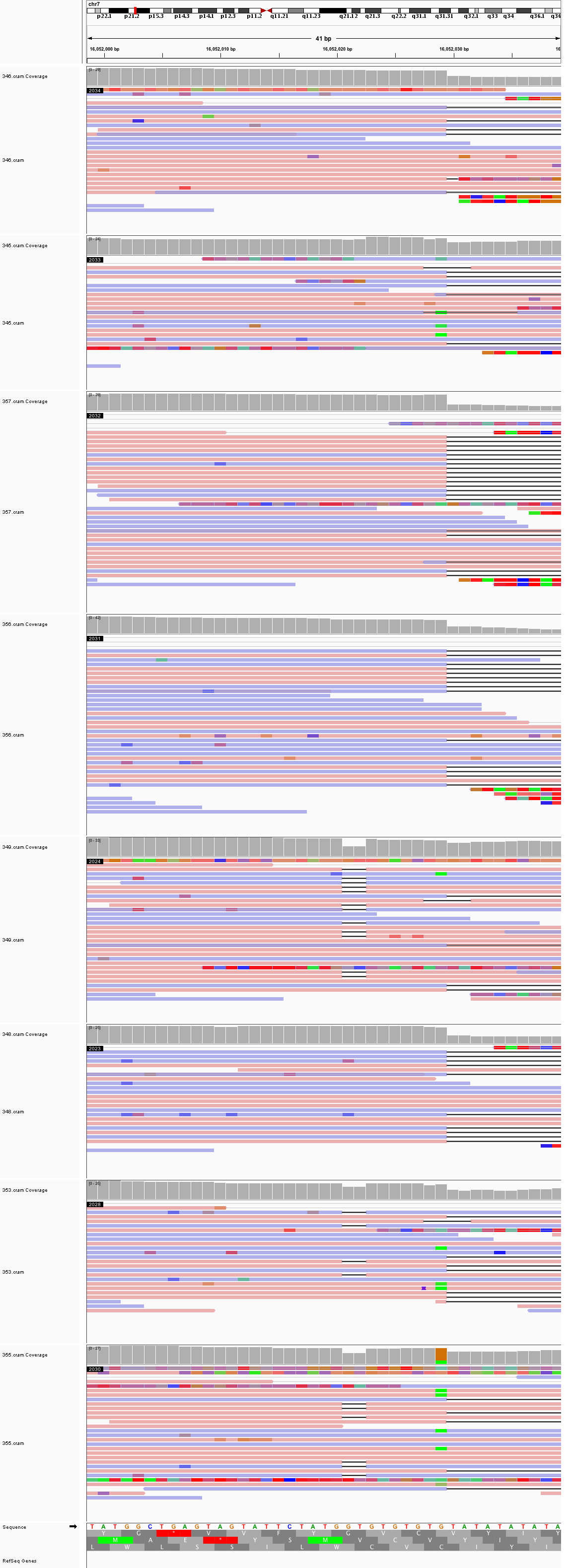

### chr7_20,072,887_20,072,927.png

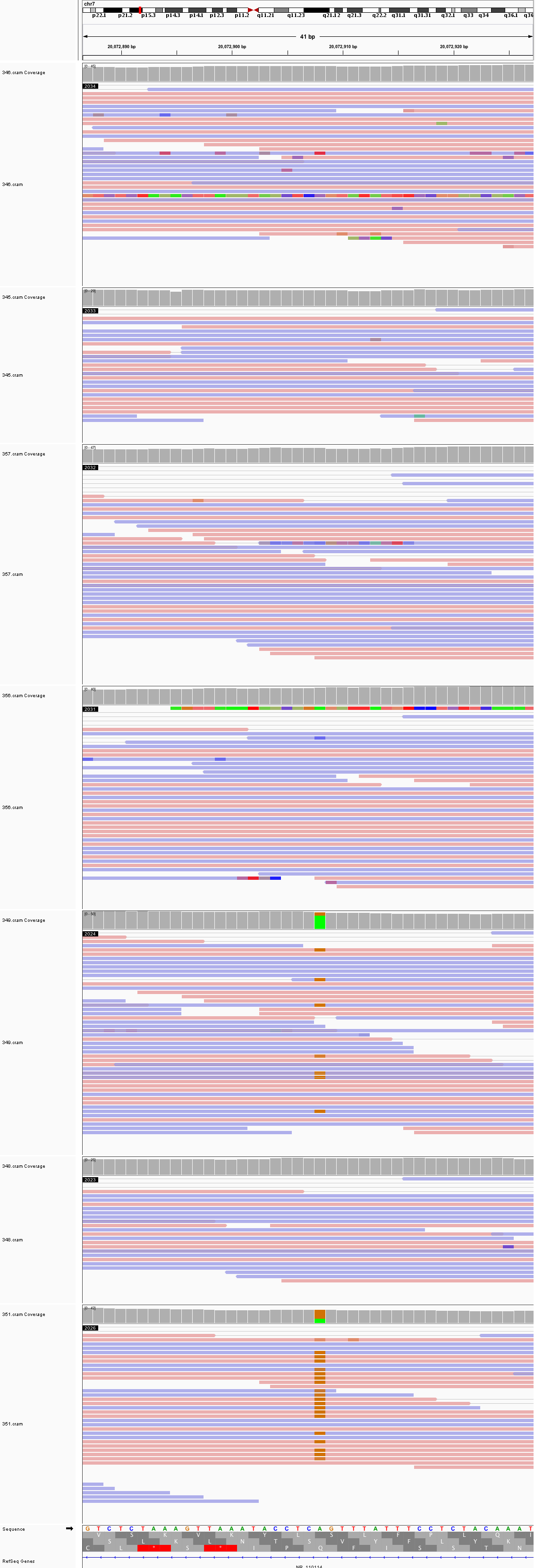

### chr7_35,686,393_35,686,433.png

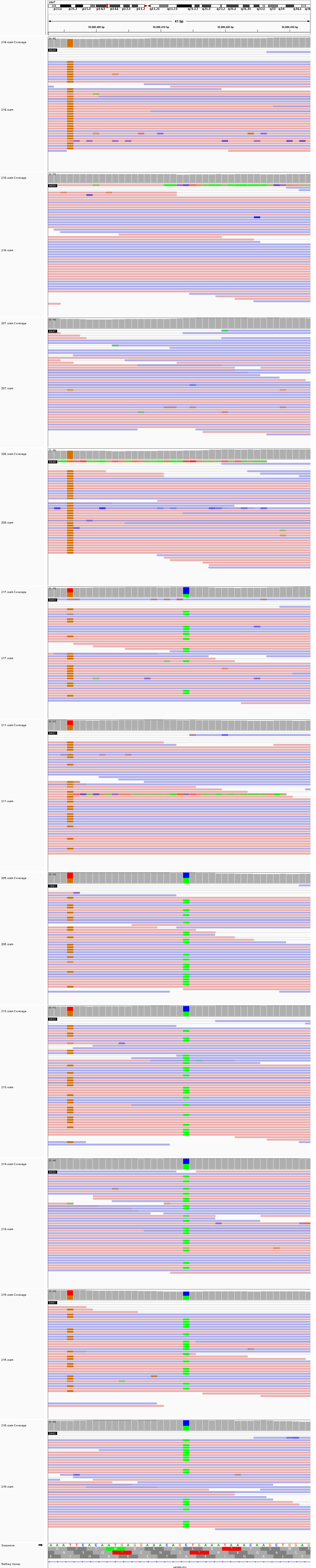

### chr7_67,286,055_67,286,095.png

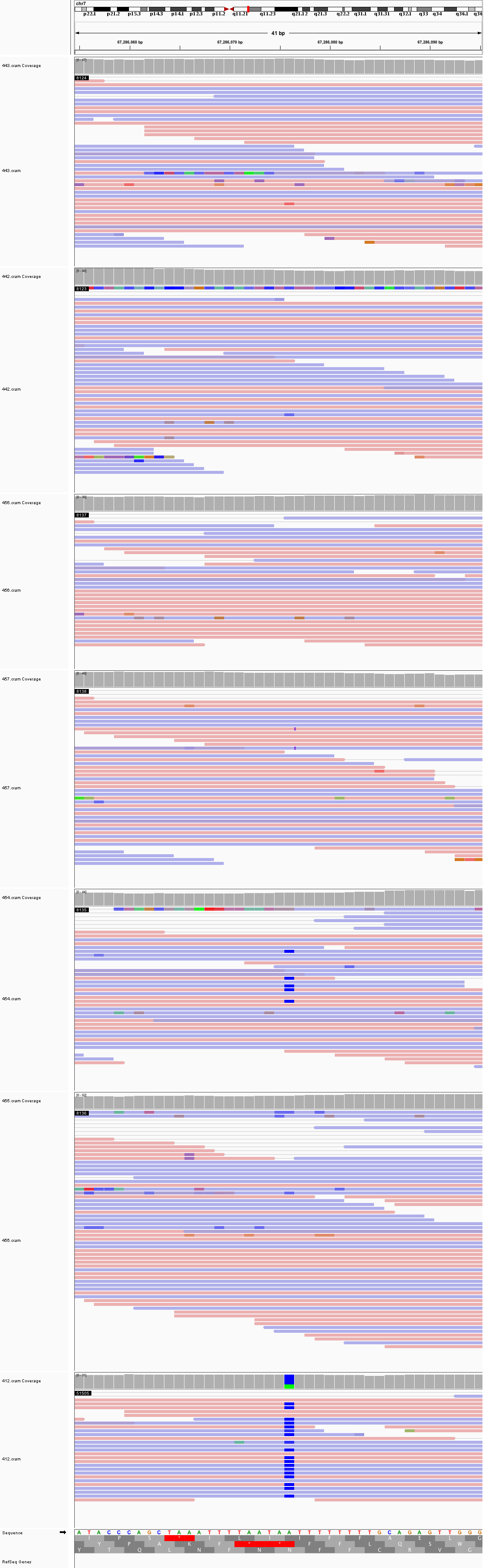

### chr7_76,655,514_76,655,554.png

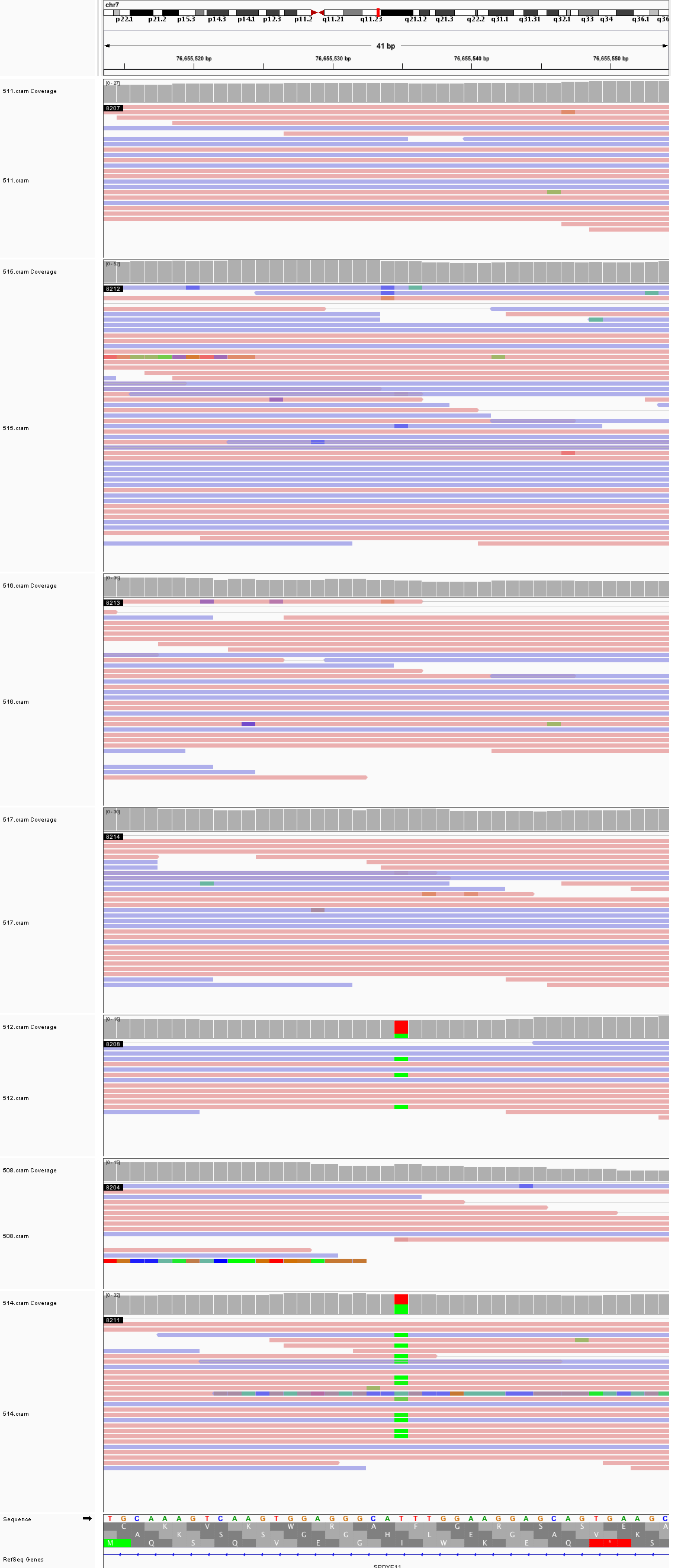

### chr7_125,936,054_125,936,094.png

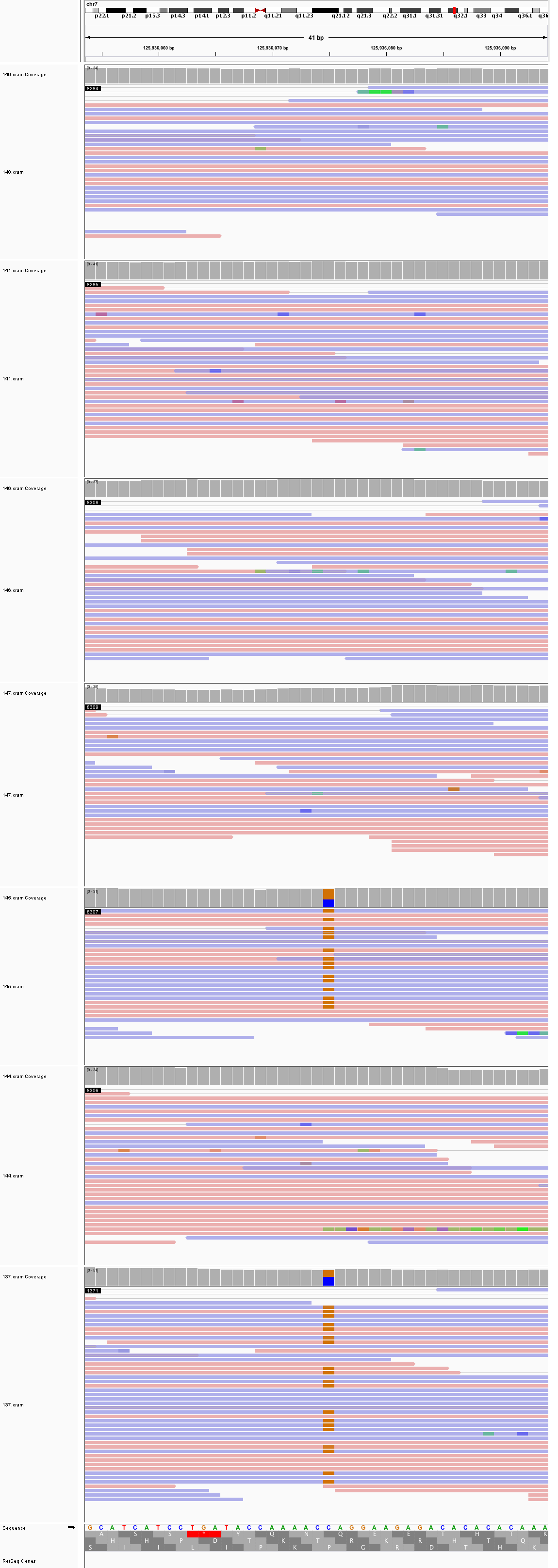

### chr7_128,095,633_128,095,673.png

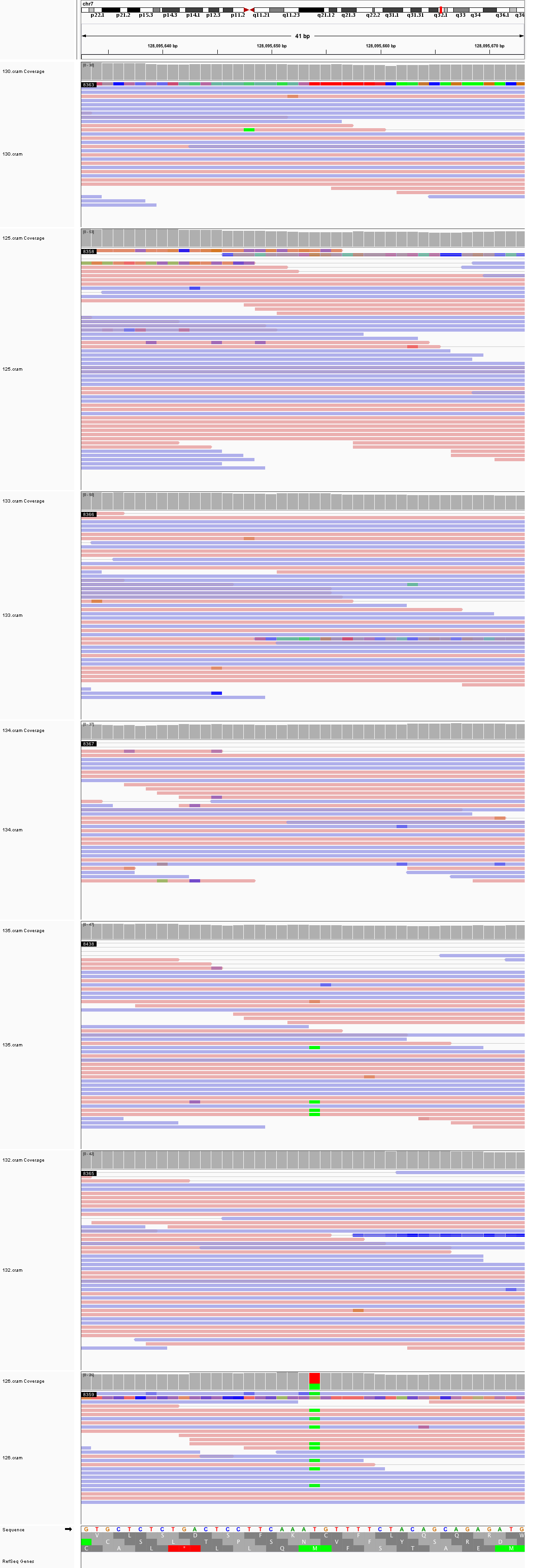

### chr7_135,593,407_135,593,447.png

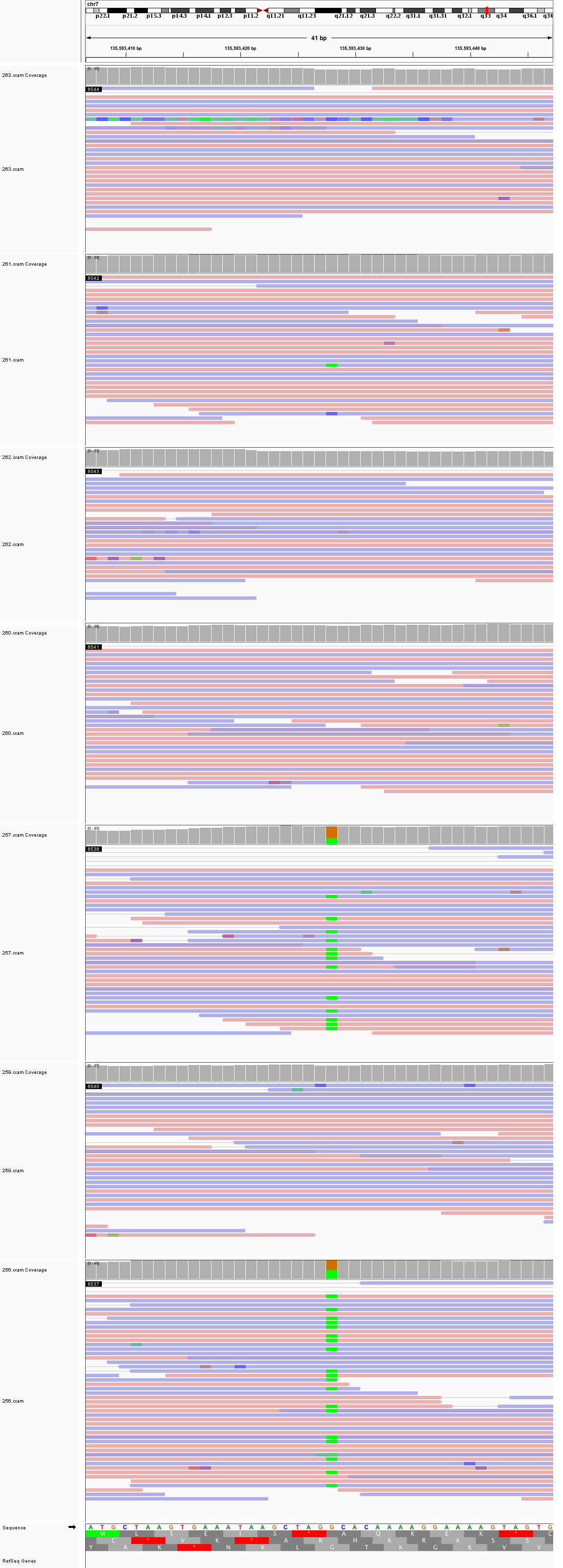

### chr8_23,098,187_23,098,227.png

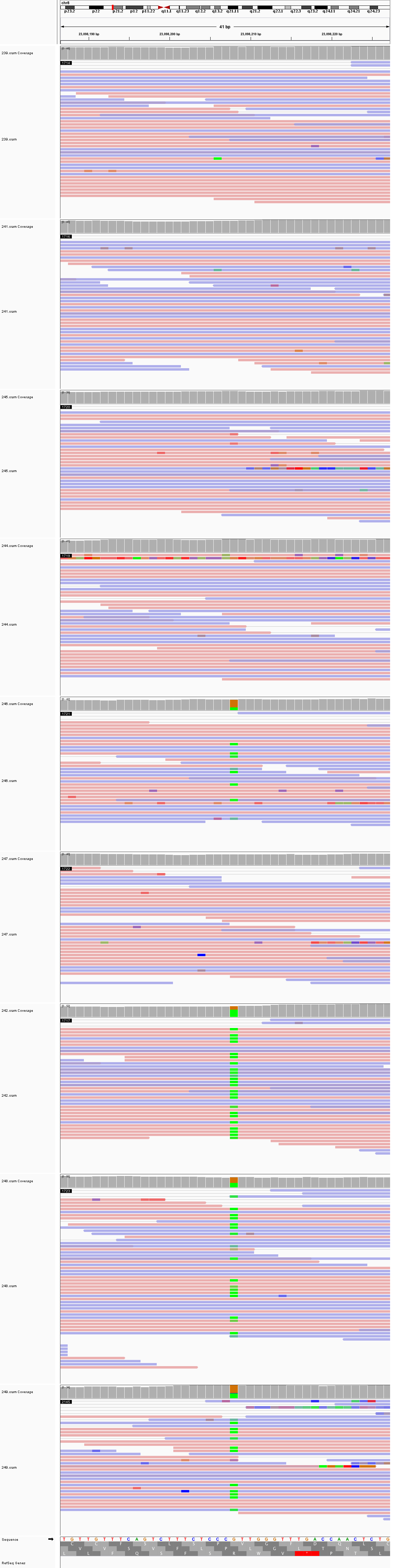

### chr8_130,821,071_130,821,111.png

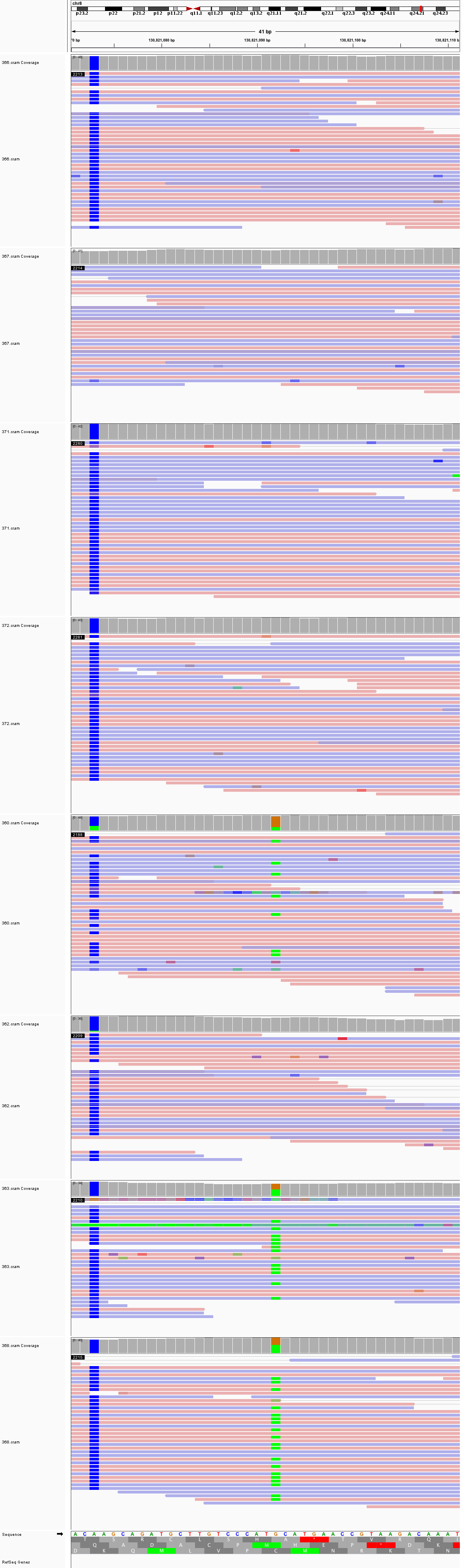

### chr8_135,382,807_135,382,847.png

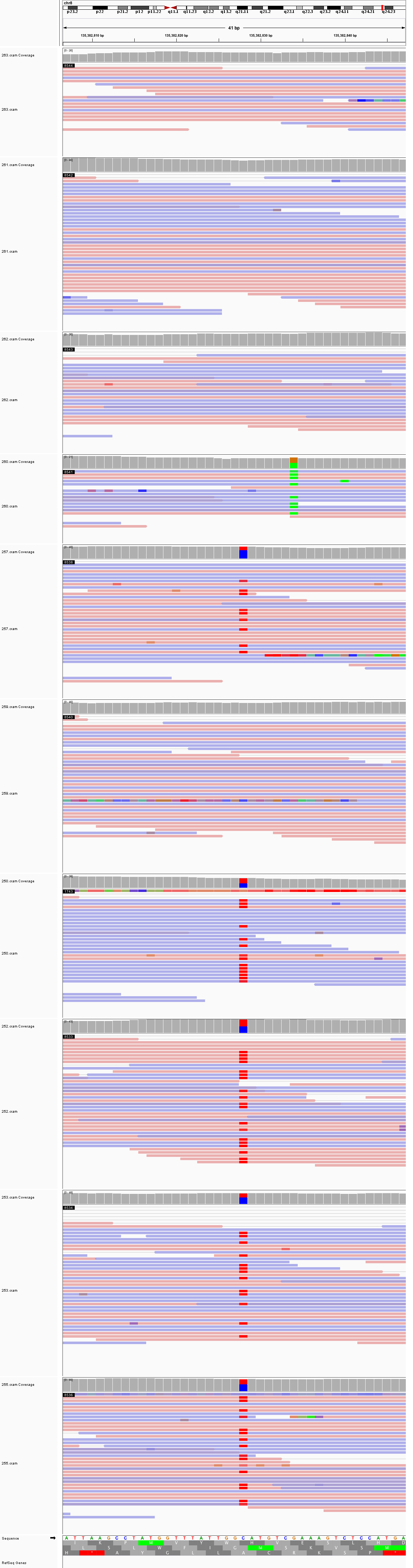

### chr8_143,745,863_143,745,903.png

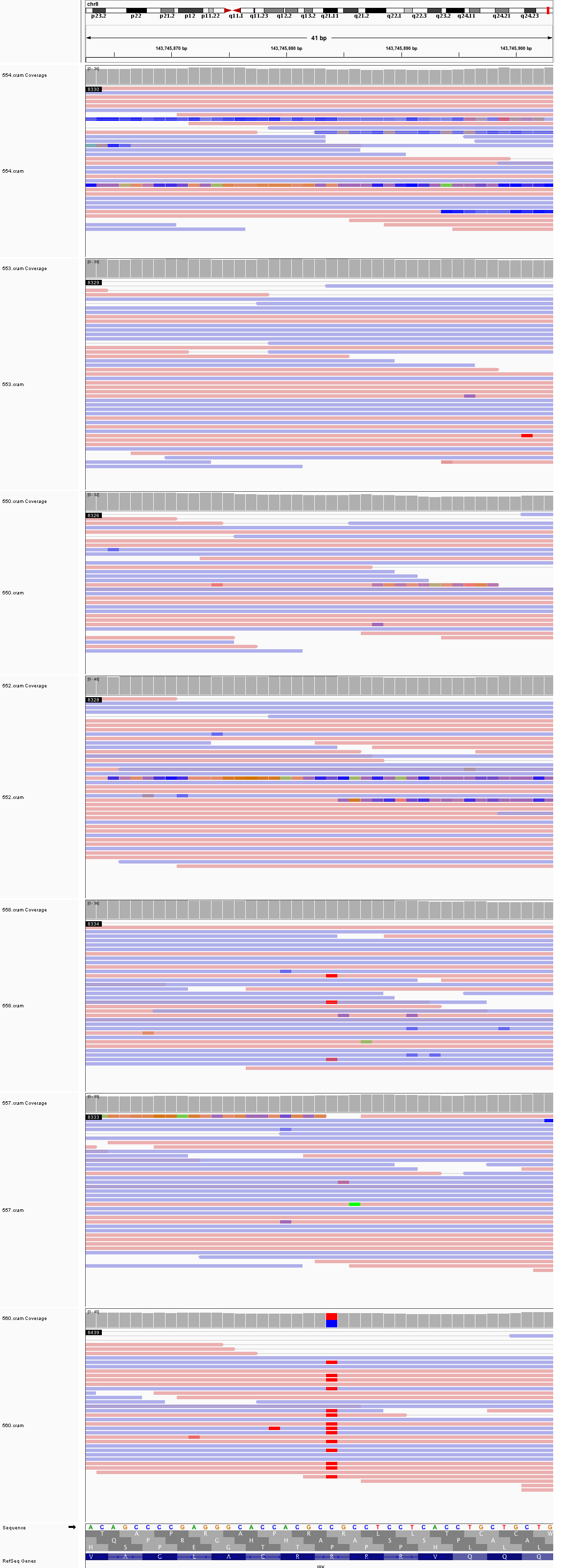

### chr9_14,992,797_14,992,837.png

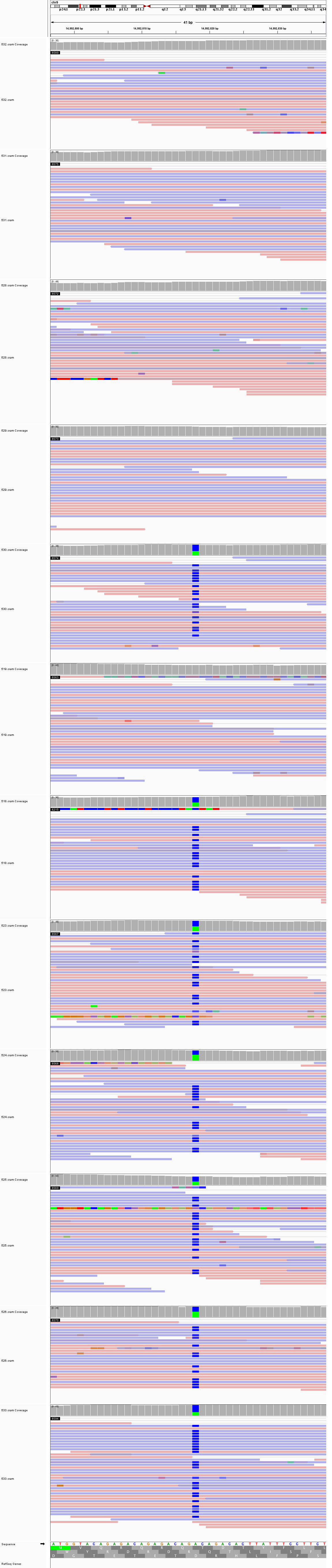

### chr9_16,782,608_16,782,648.png

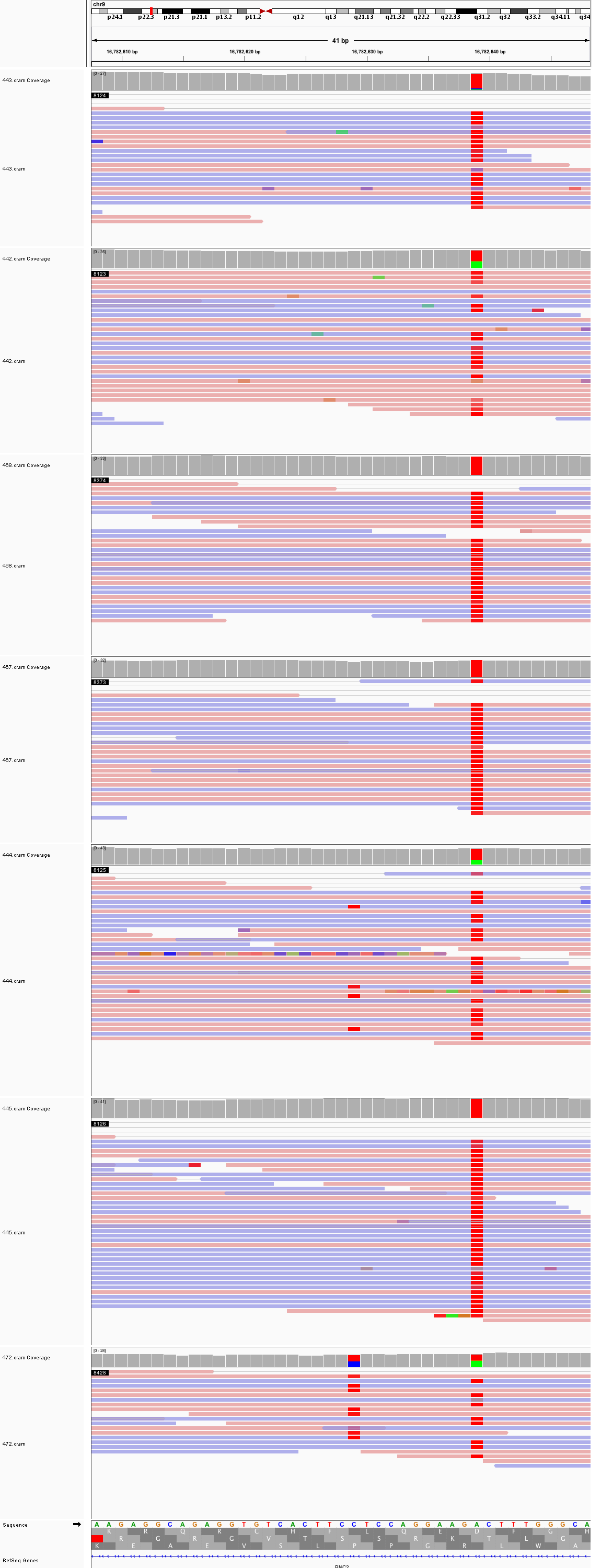

### chr9_30,298,590_30,298,630.png

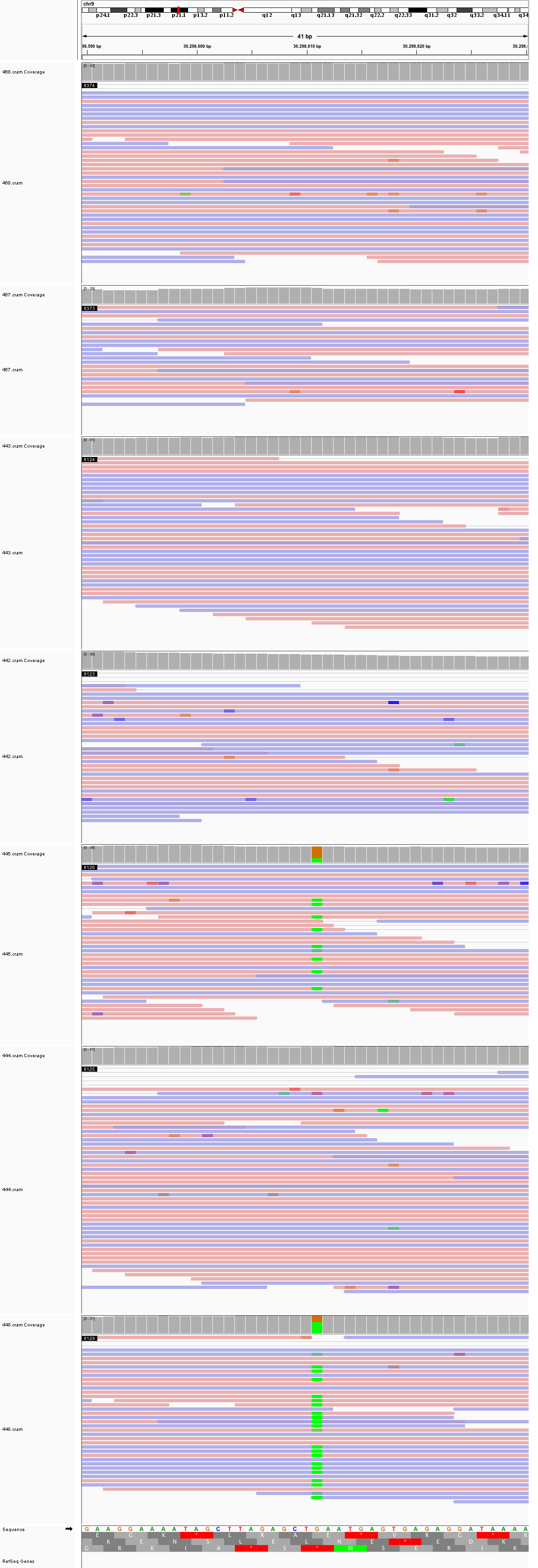

### chr9_66,789,896_66,789,936.png

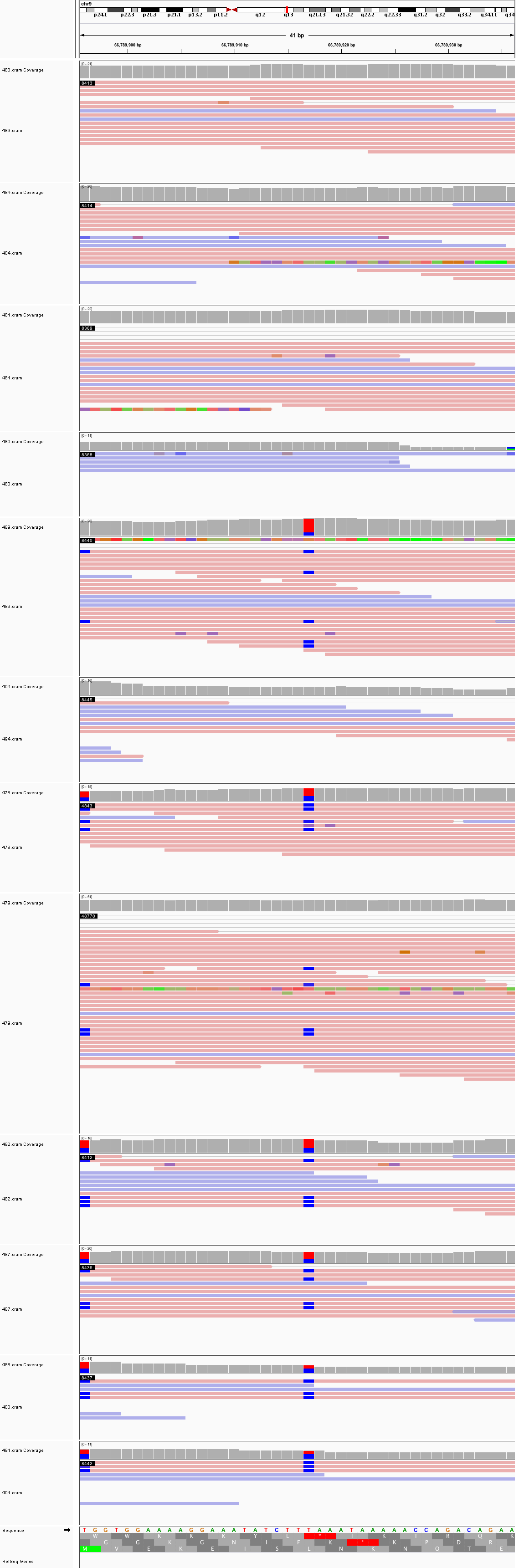

### chr9_82,691,979_82,692,019.png

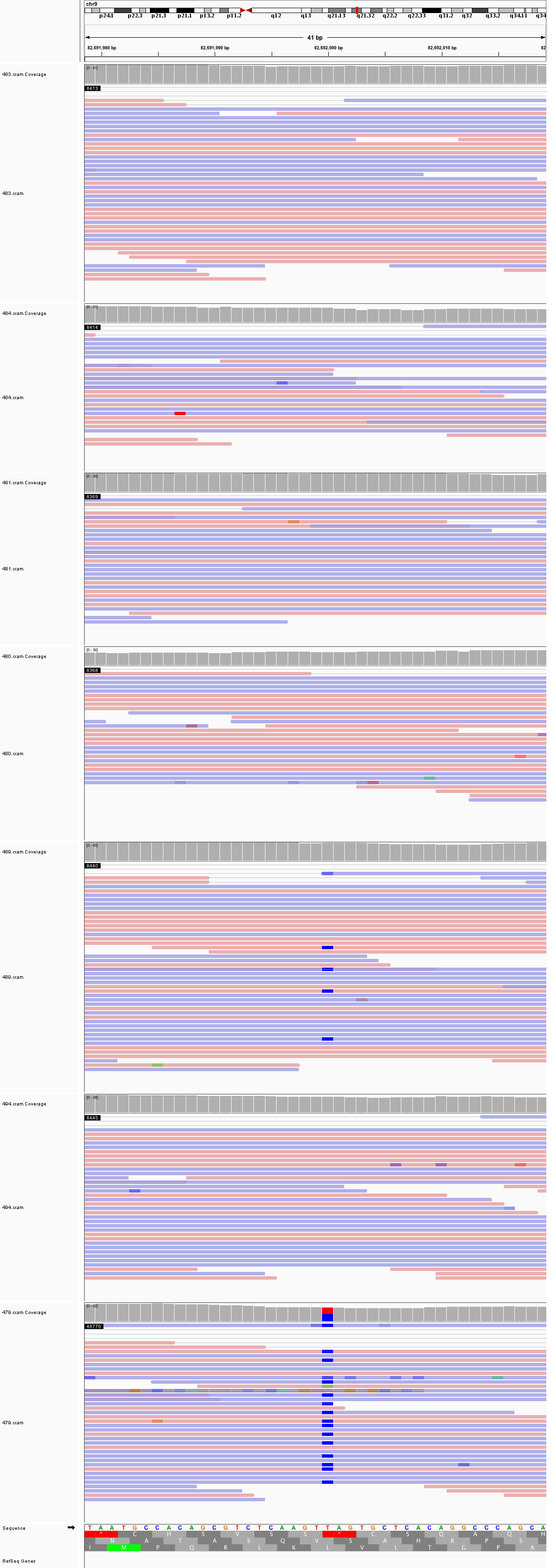

### chr9_122,523,896_122,523,936.png

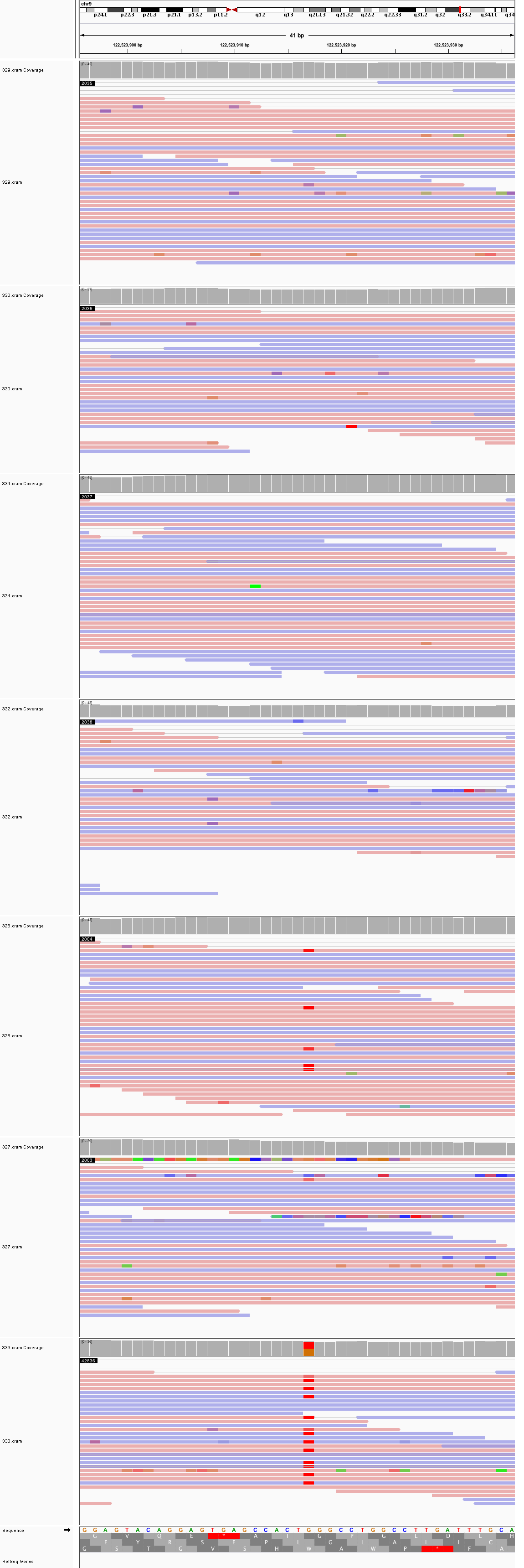

### chr10_9,590,437_9,590,477.png

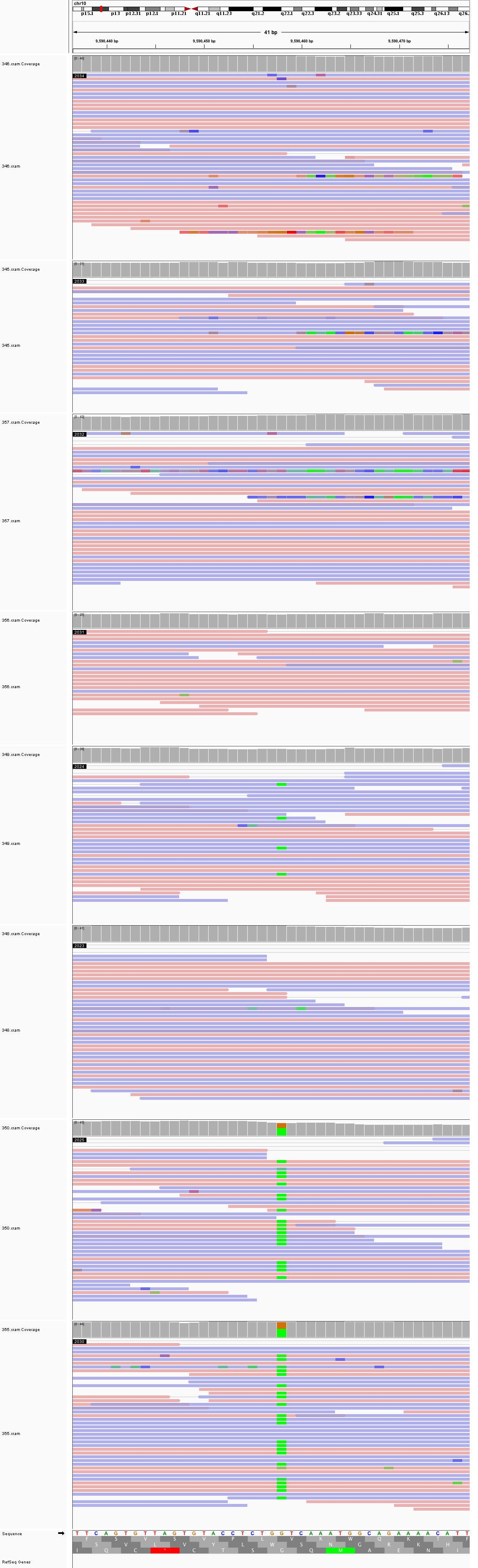

### chr10_36,937,610_36,937,650.png

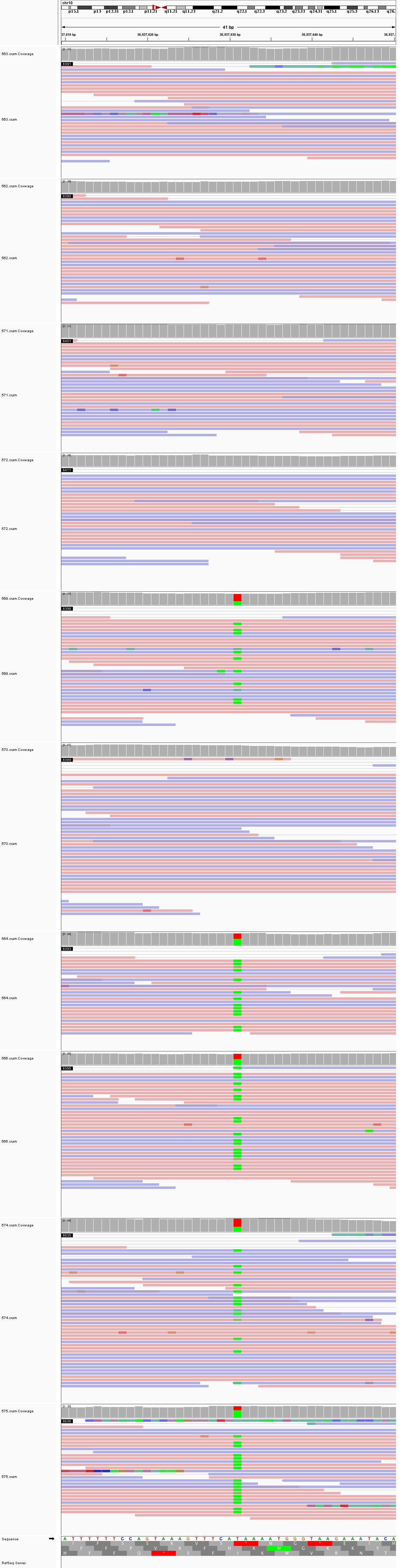

### chr12_21,433,005_21,433,045.png

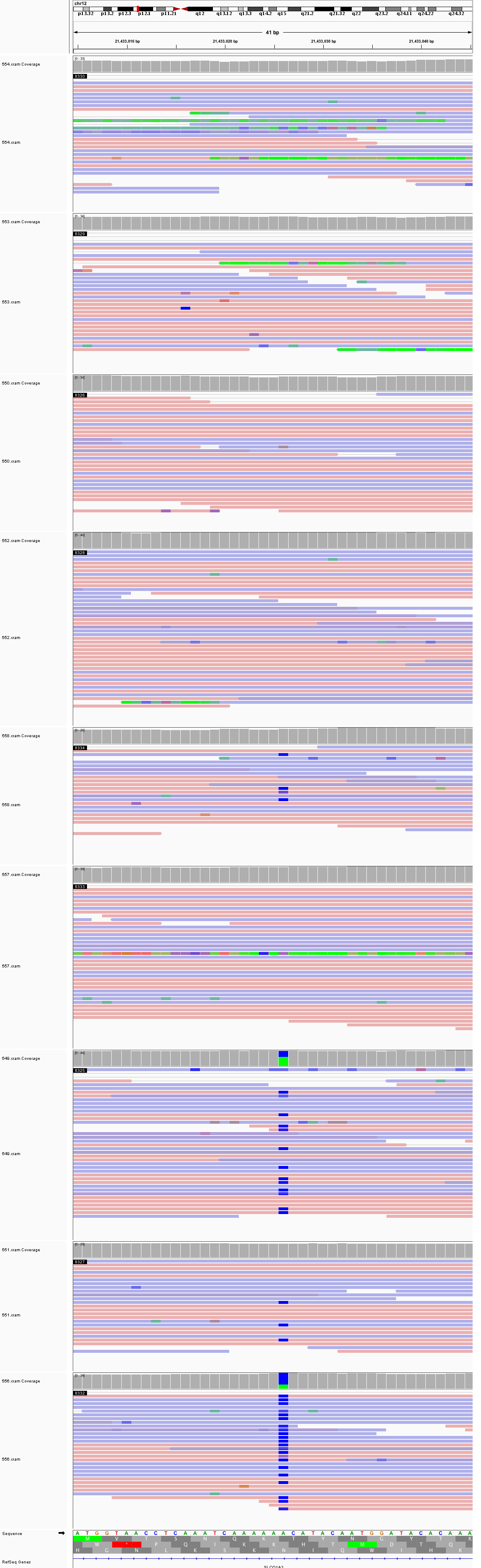

### chr12_37,939,044_37,939,084.png

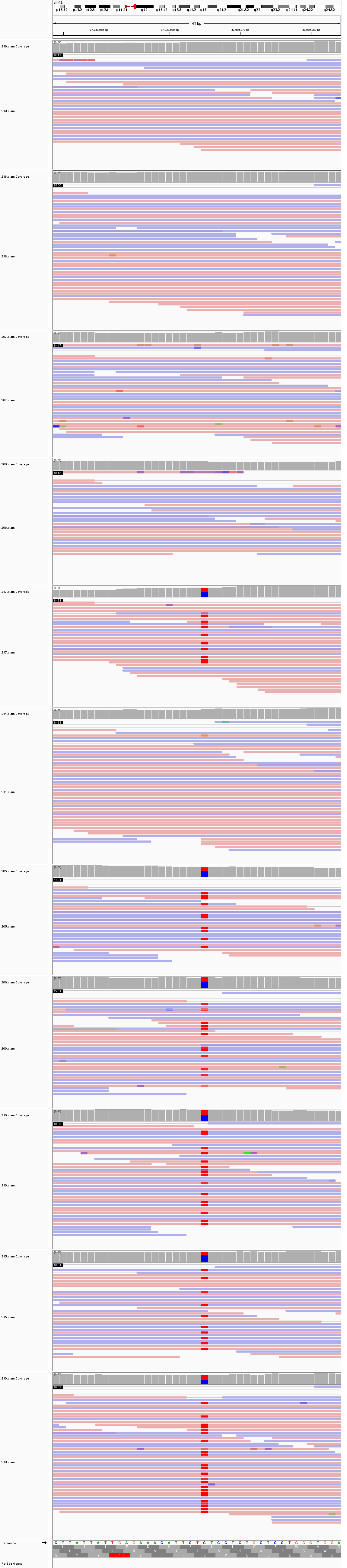

### chr12_60,906,439_60,906,479.png

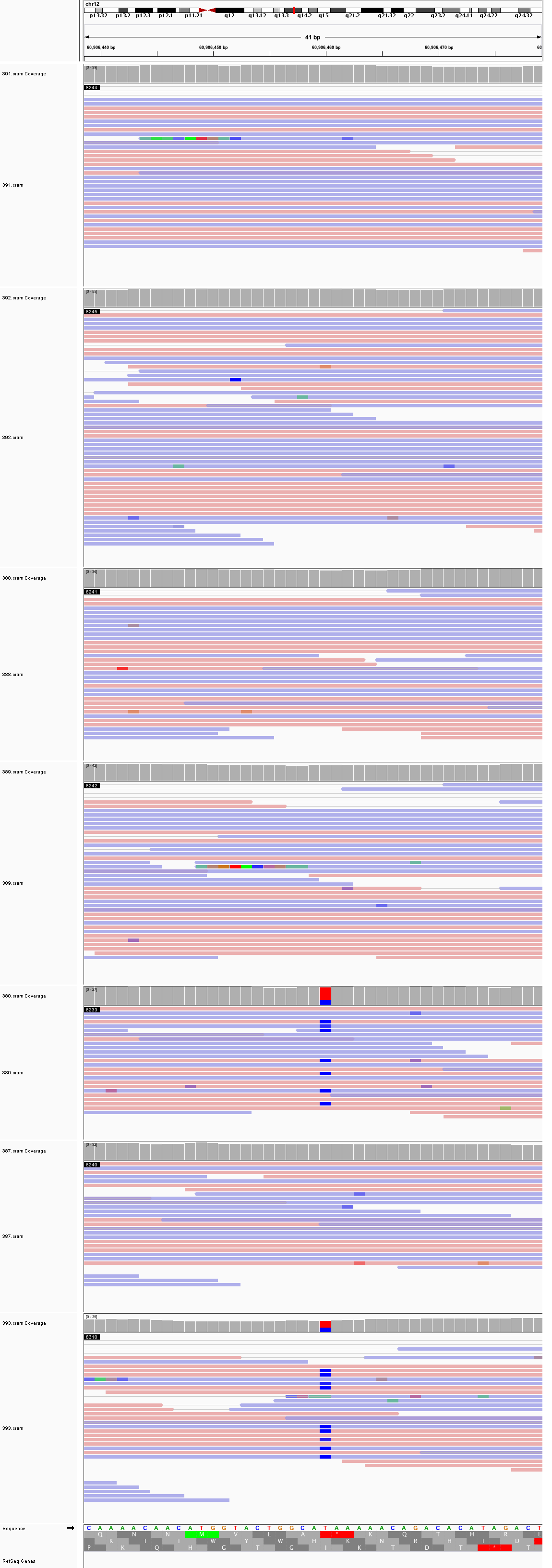

### chr12_72,526,950_72,526,990.png

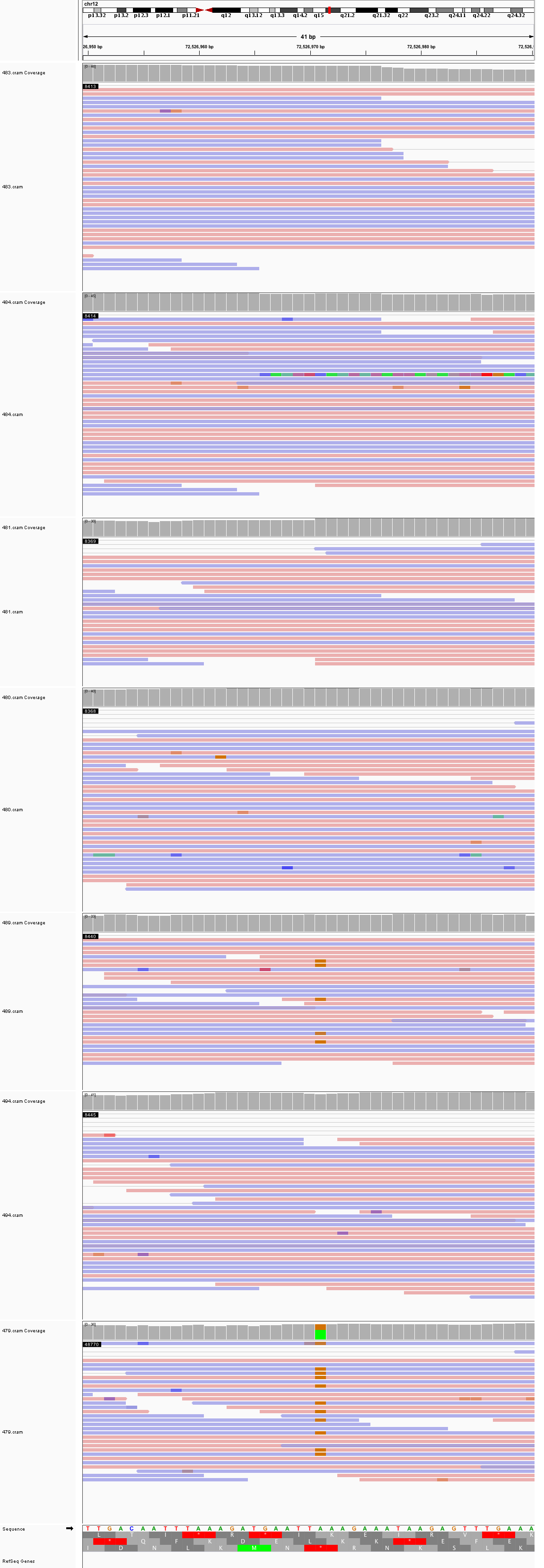

### chr12_84,549,341_84,549,381.png

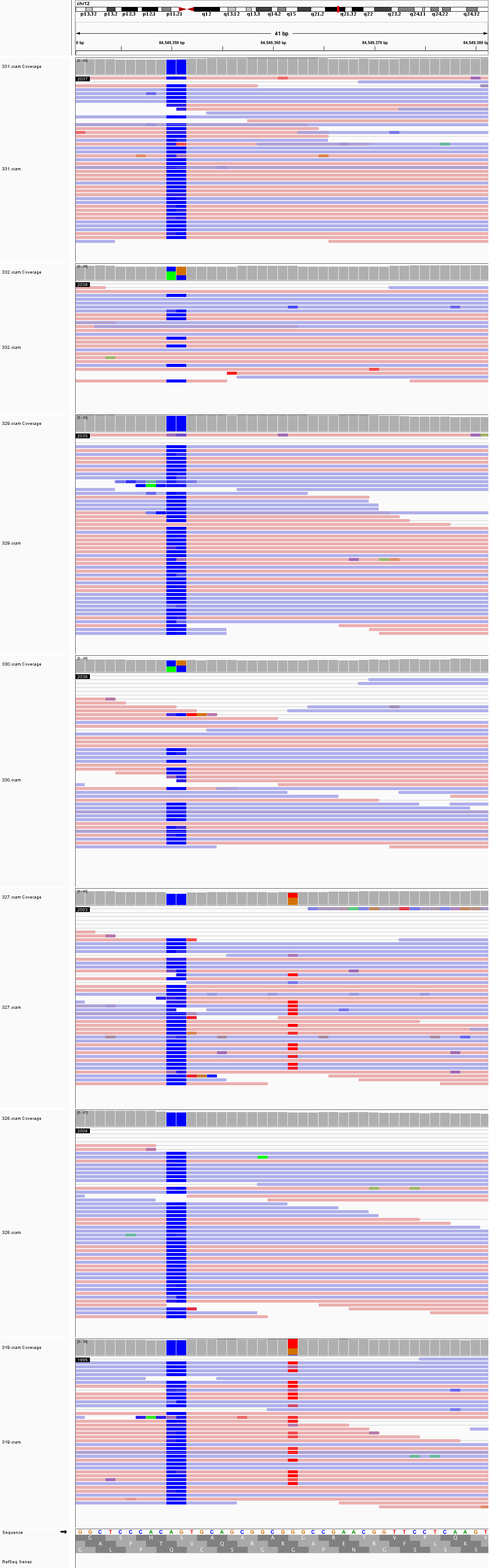

### chr12_86,145,812_86,145,852.png

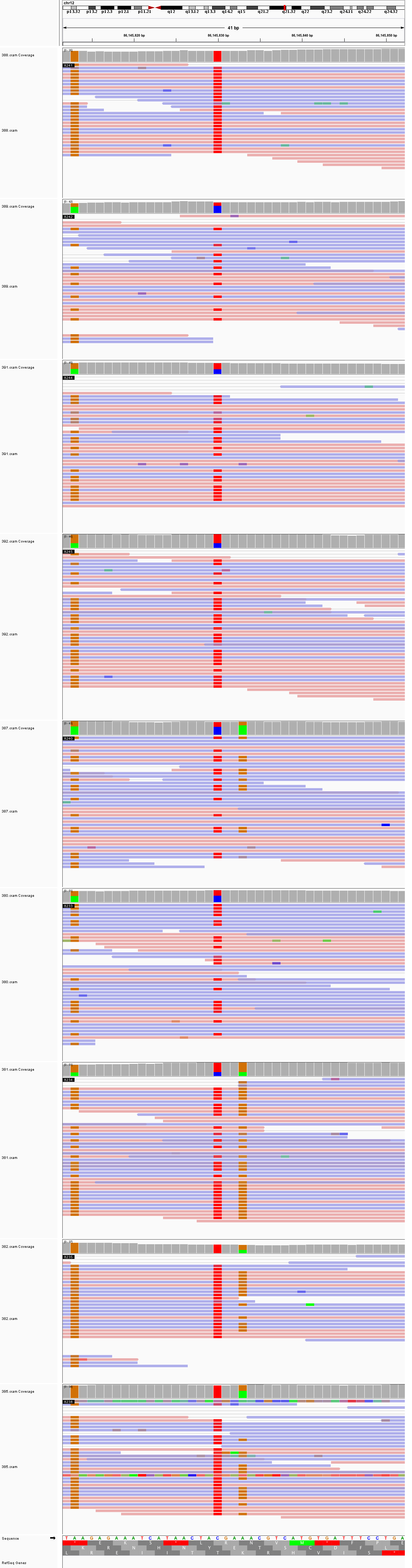

### chr13_34,442,472_34,442,512.png

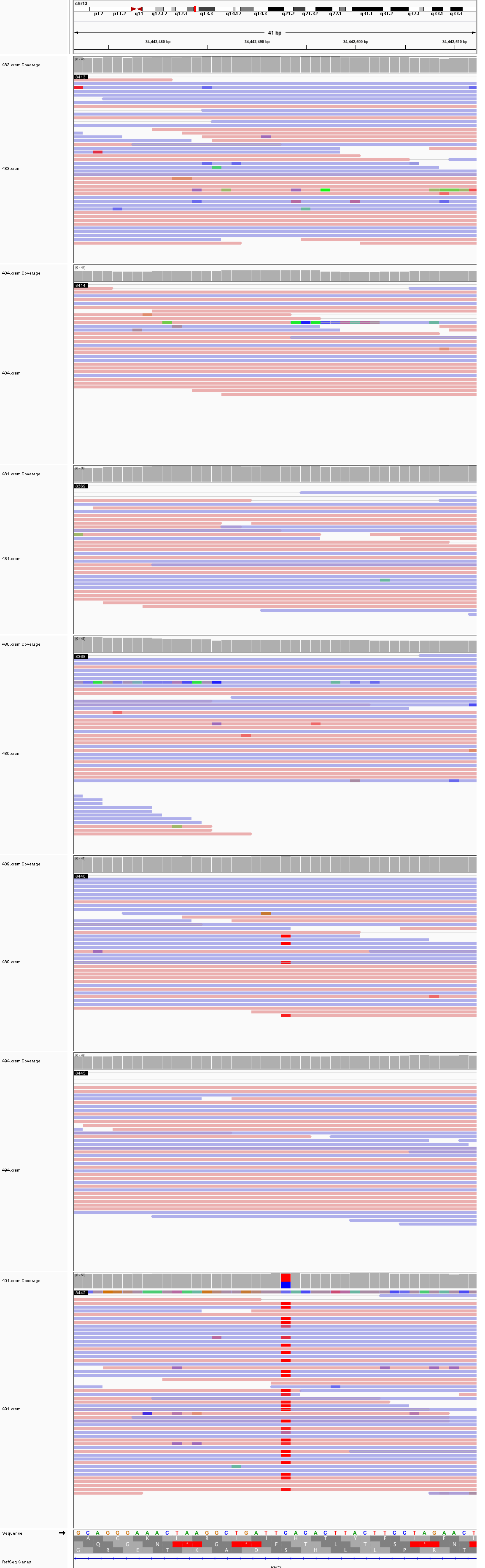

### chr13_38,843,040_38,843,080.png

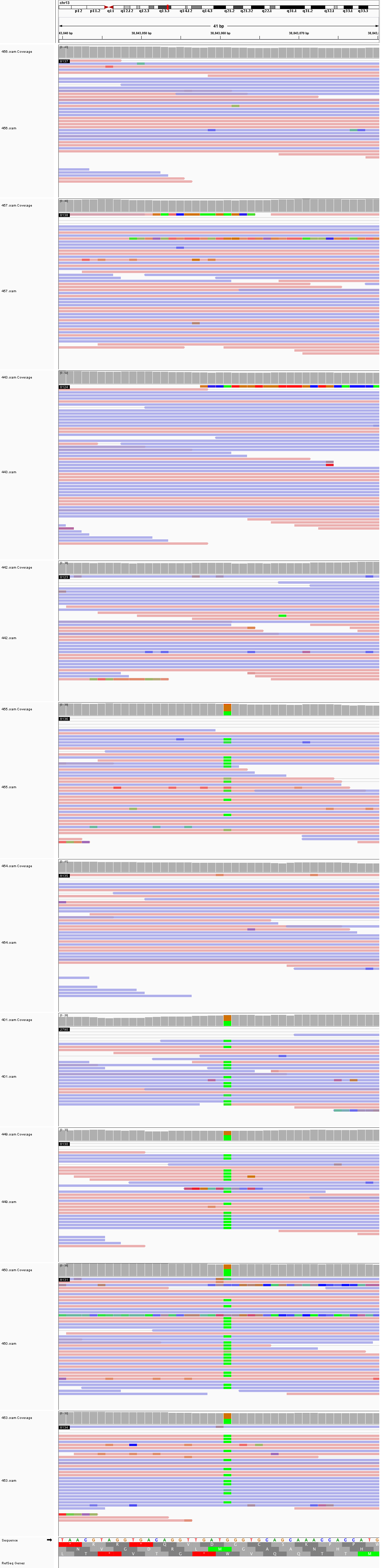

### chr13_39,737,809_39,737,849.png

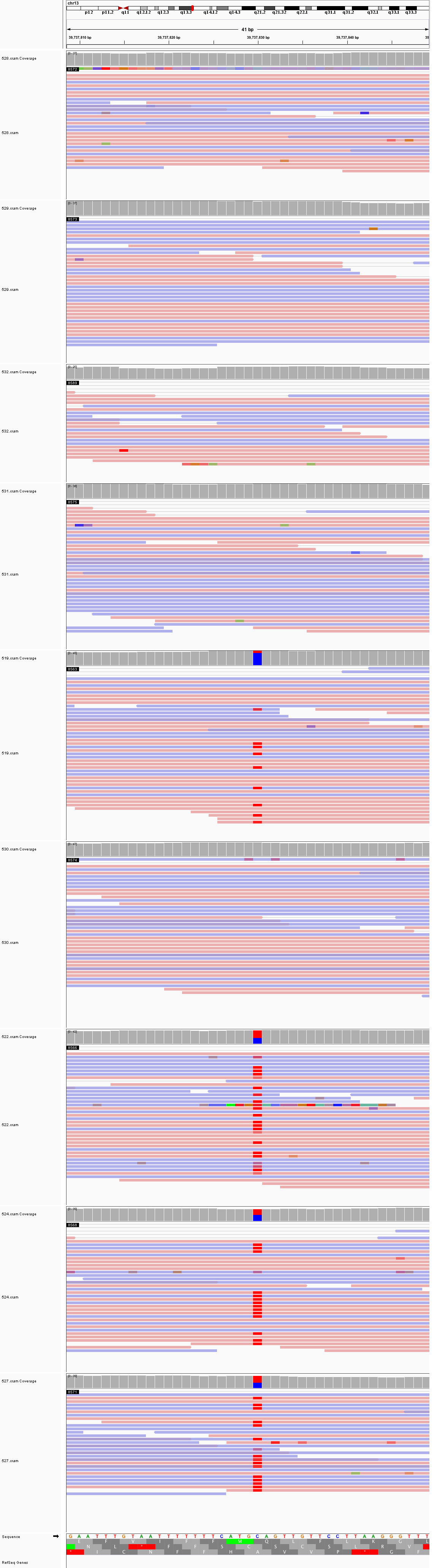

### chr13_110,168,088_110,168,128.png

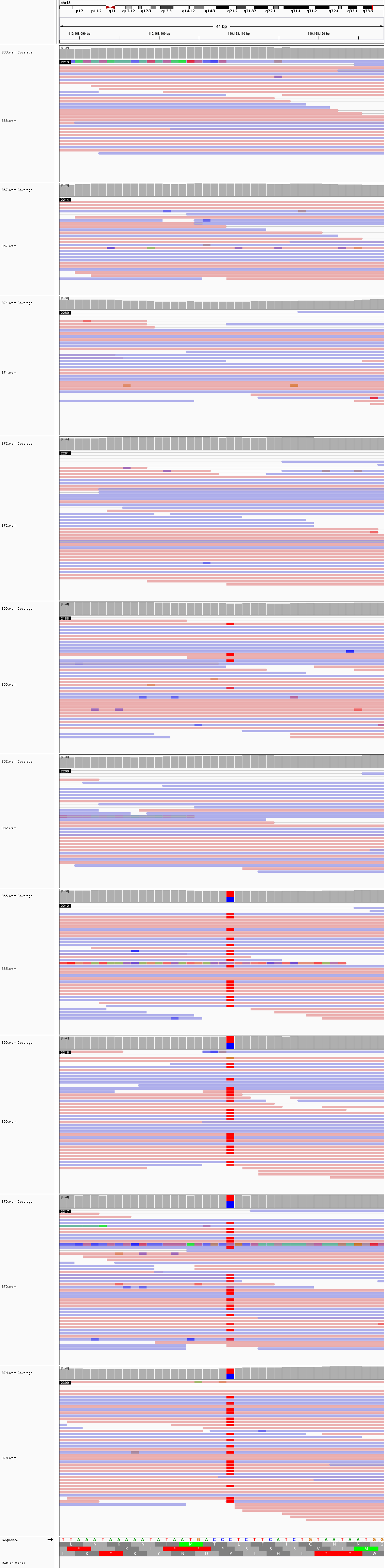

### chr13_111,384,818_111,384,858.png

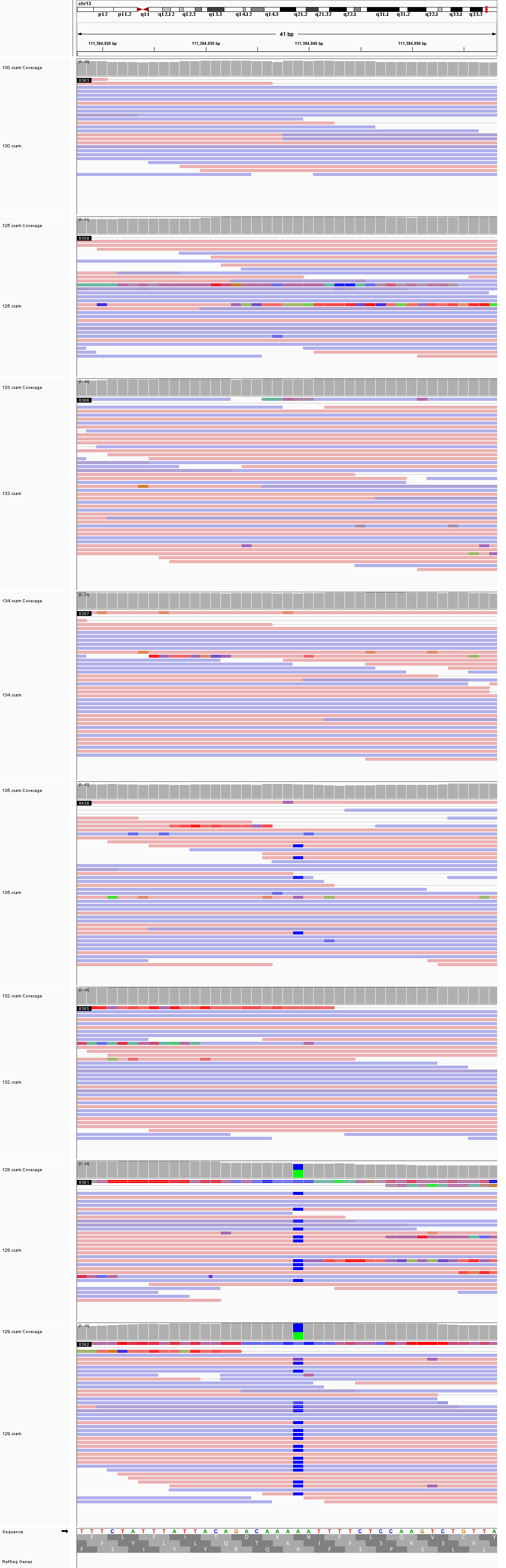

### chr14_63,964,019_63,964,059.png

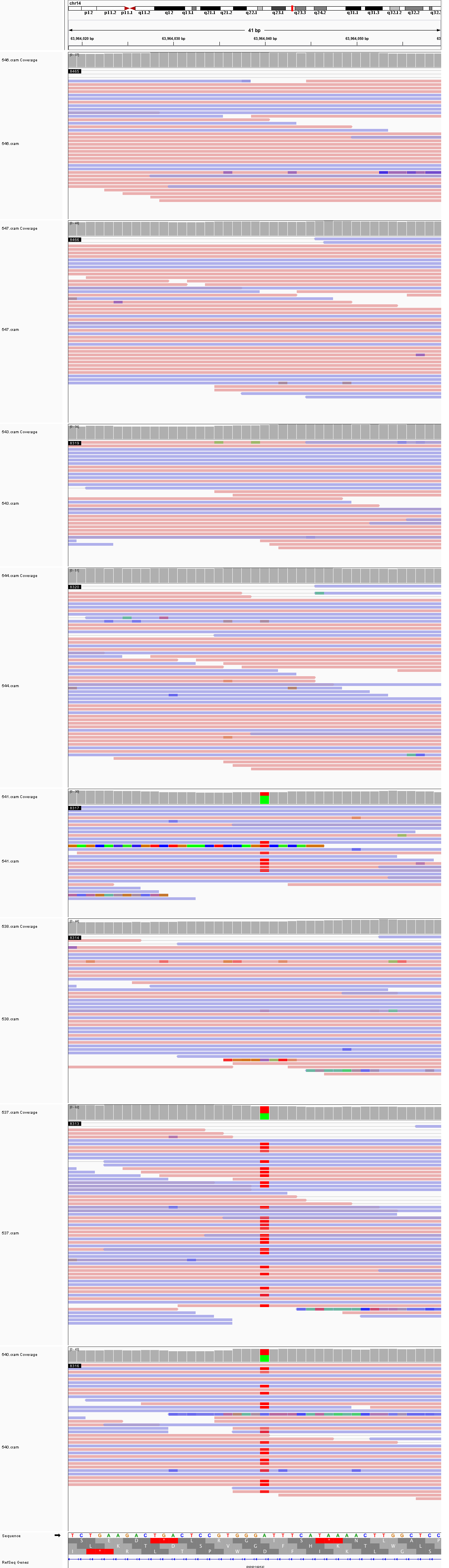

### chr14_100,168,669_100,168,709.png

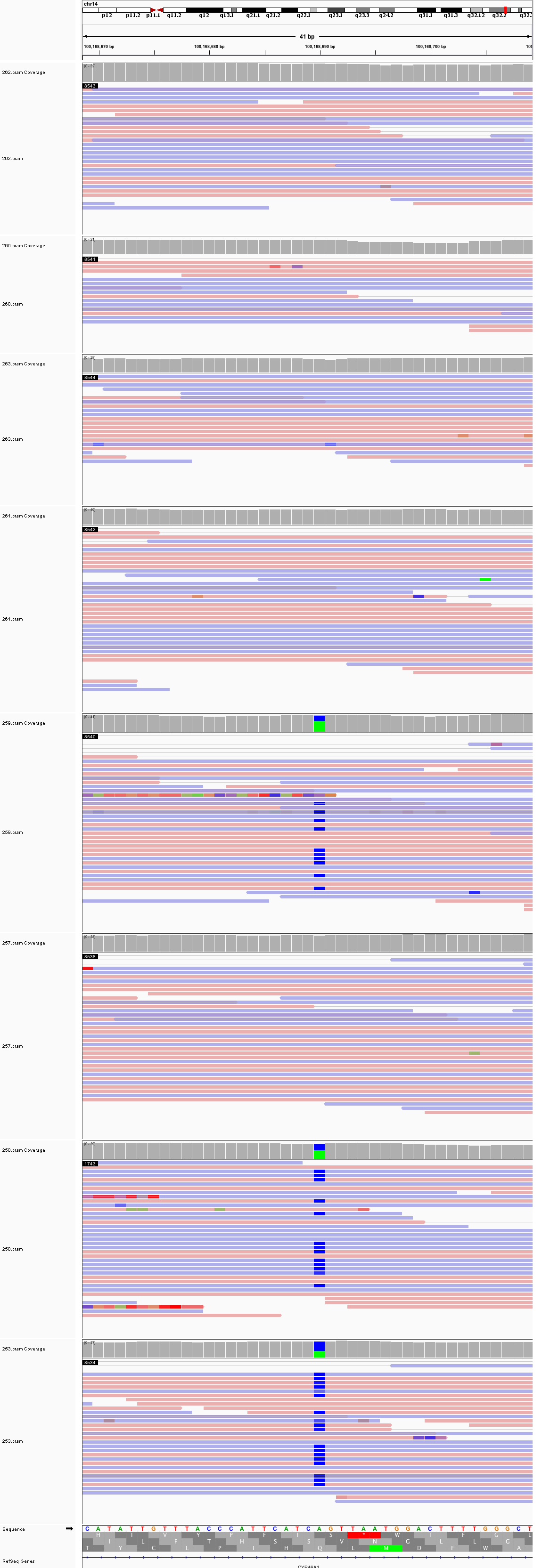

### chr15_50,735,251_50,735,291.png

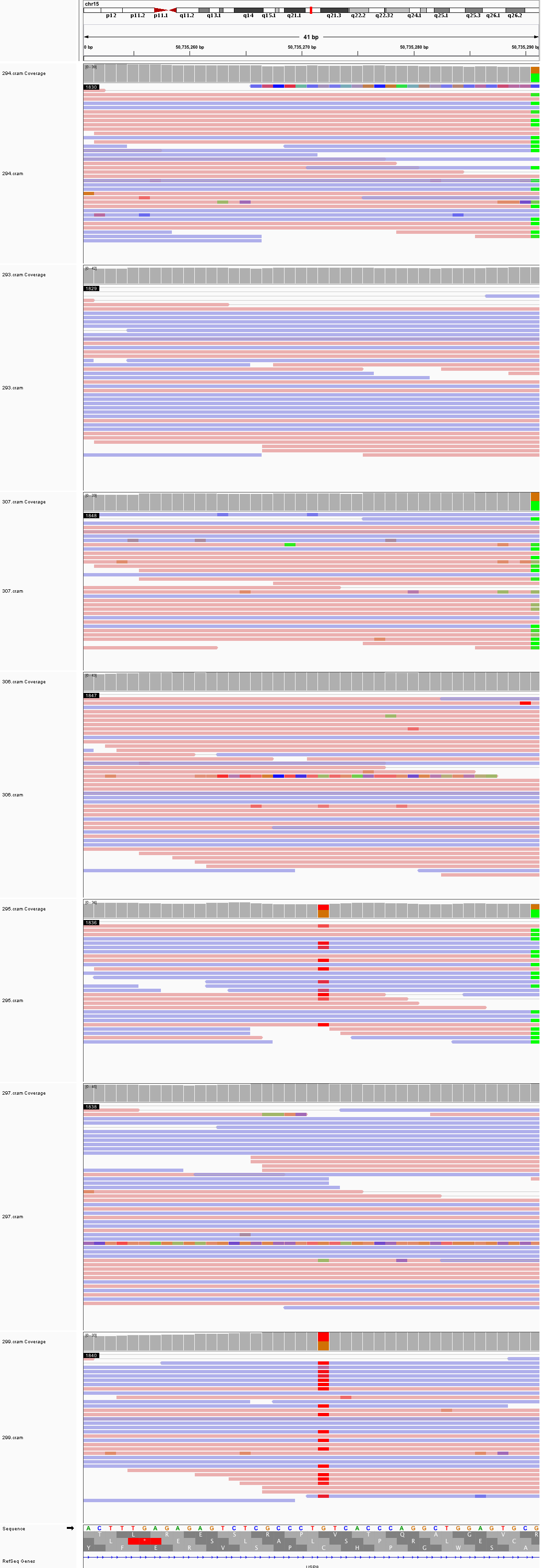

### chr15_62,497,018_62,497,058.png

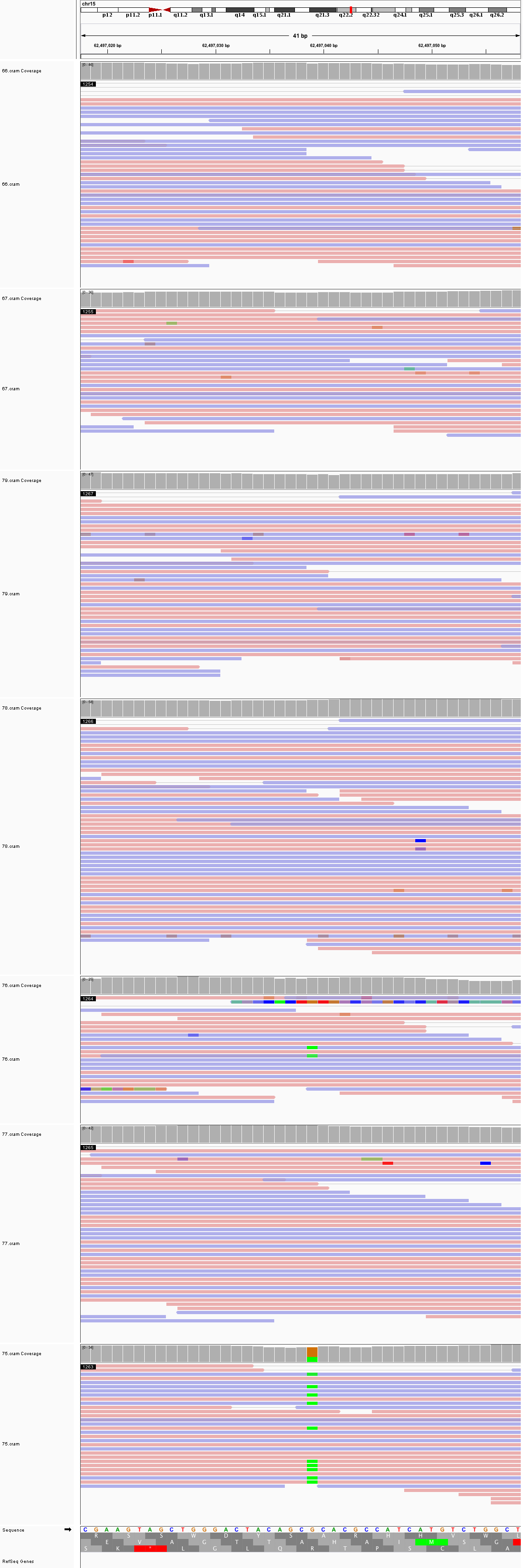

### chr15_84,622,257_84,622,297.png

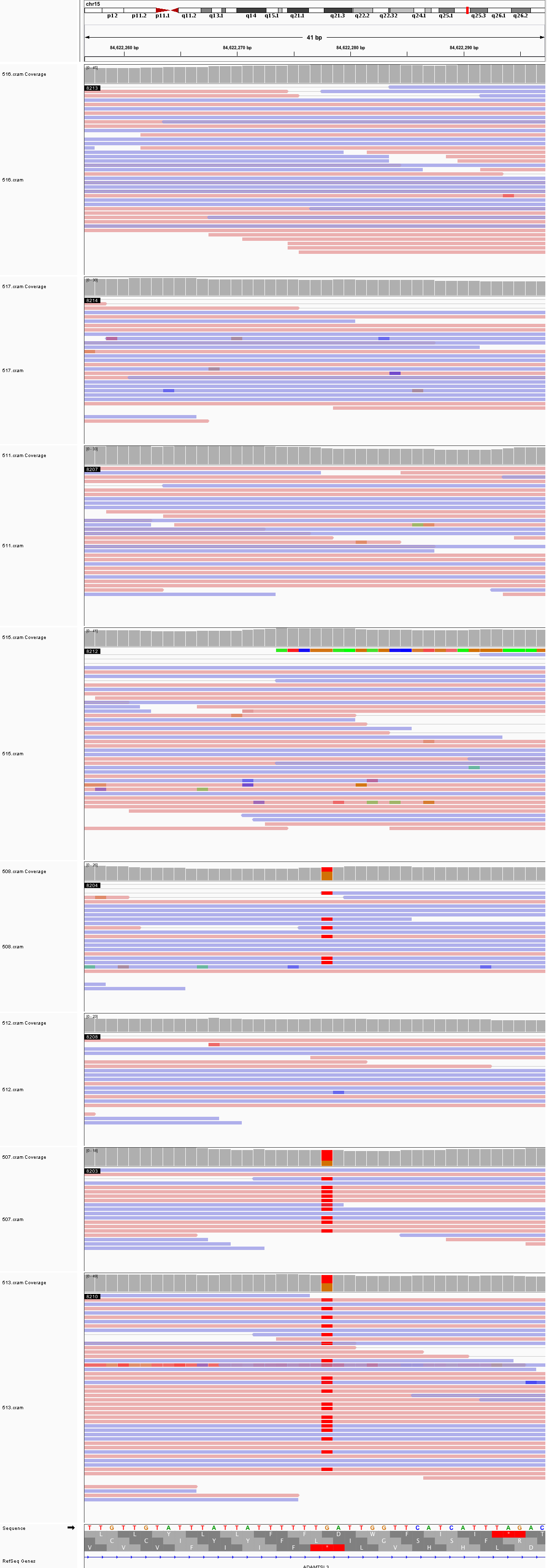

### chr15_96,799,375_96,799,415.png

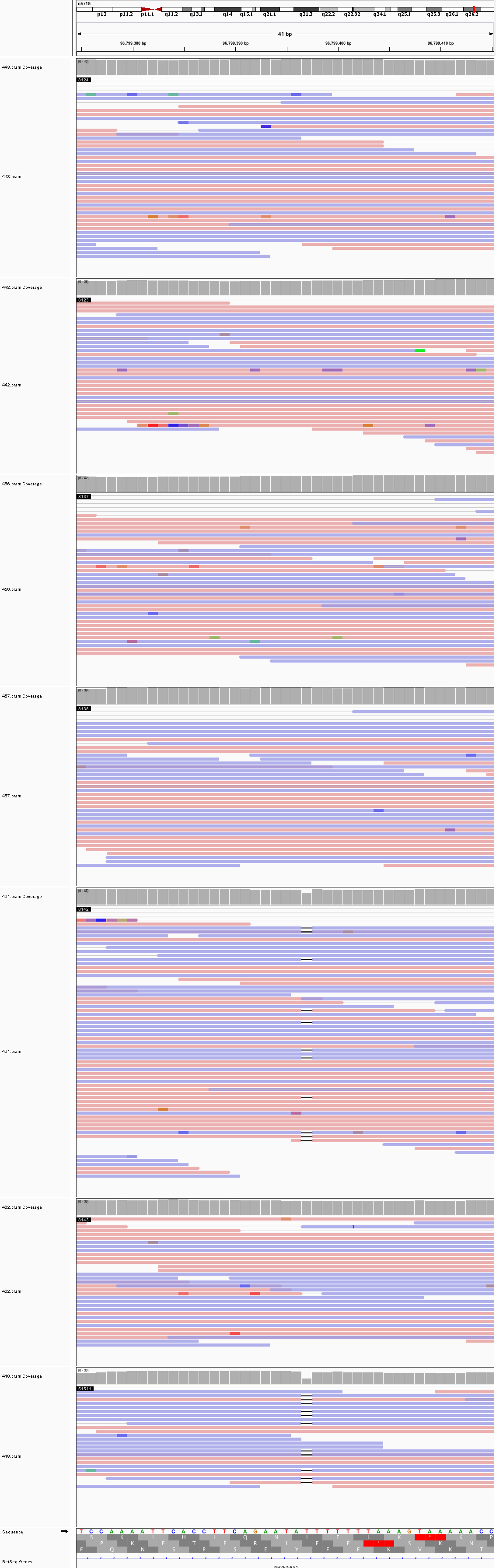

### chr16_2,645,692_2,645,732.png

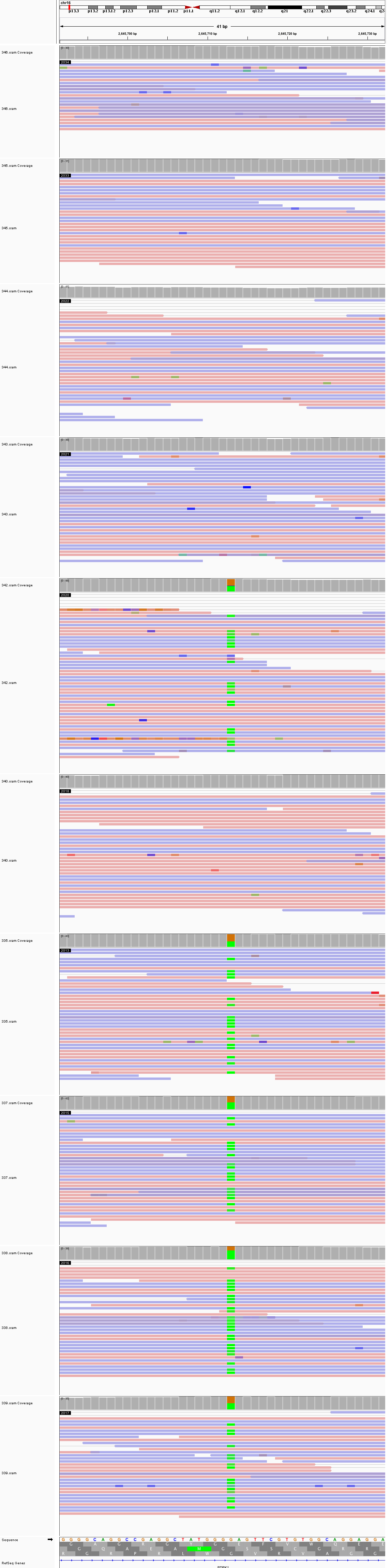

### chr16_75,007,082_75,007,122.png

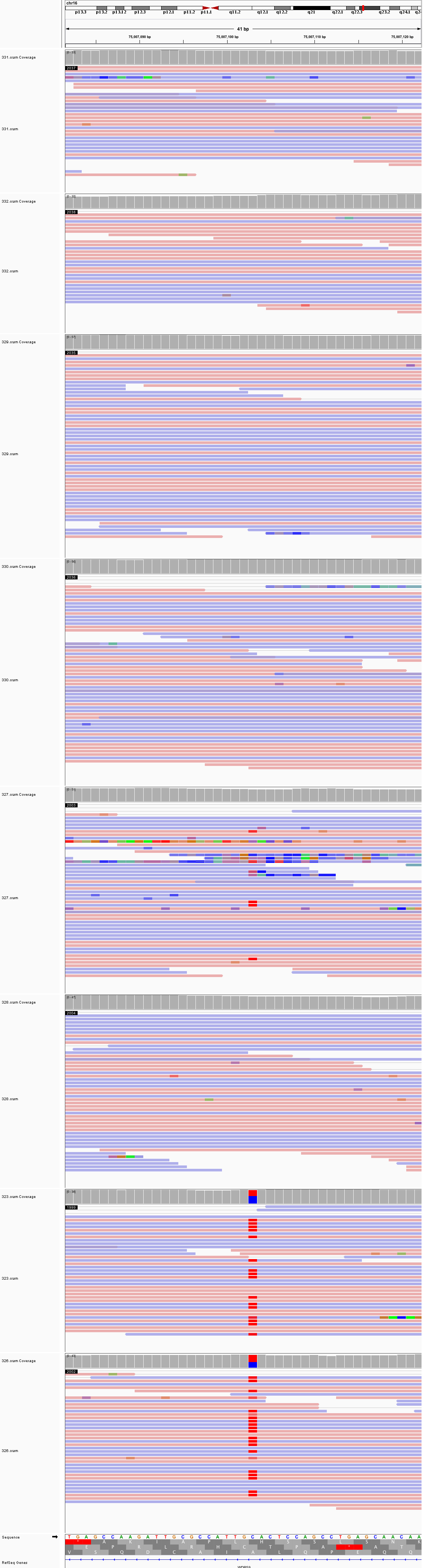

### chr17_74,941,746_74,941,786.png

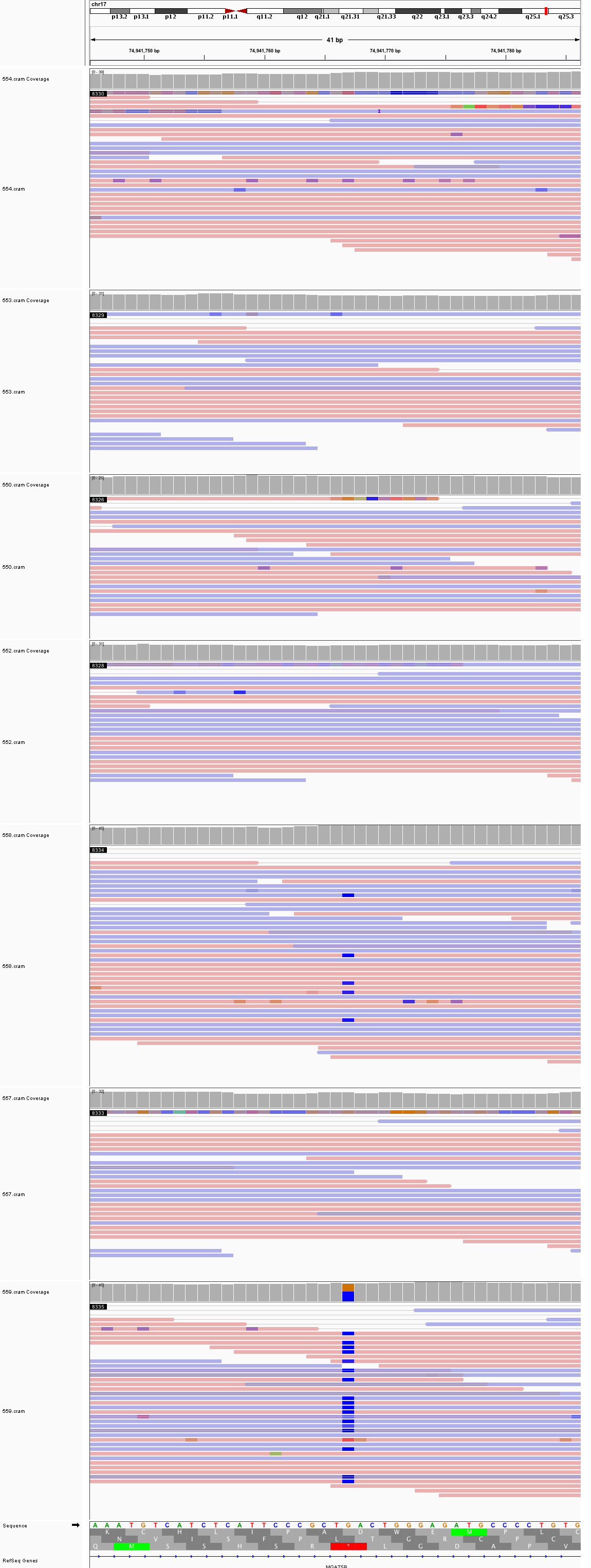

### chr18_11,075,806_11,075,846.png

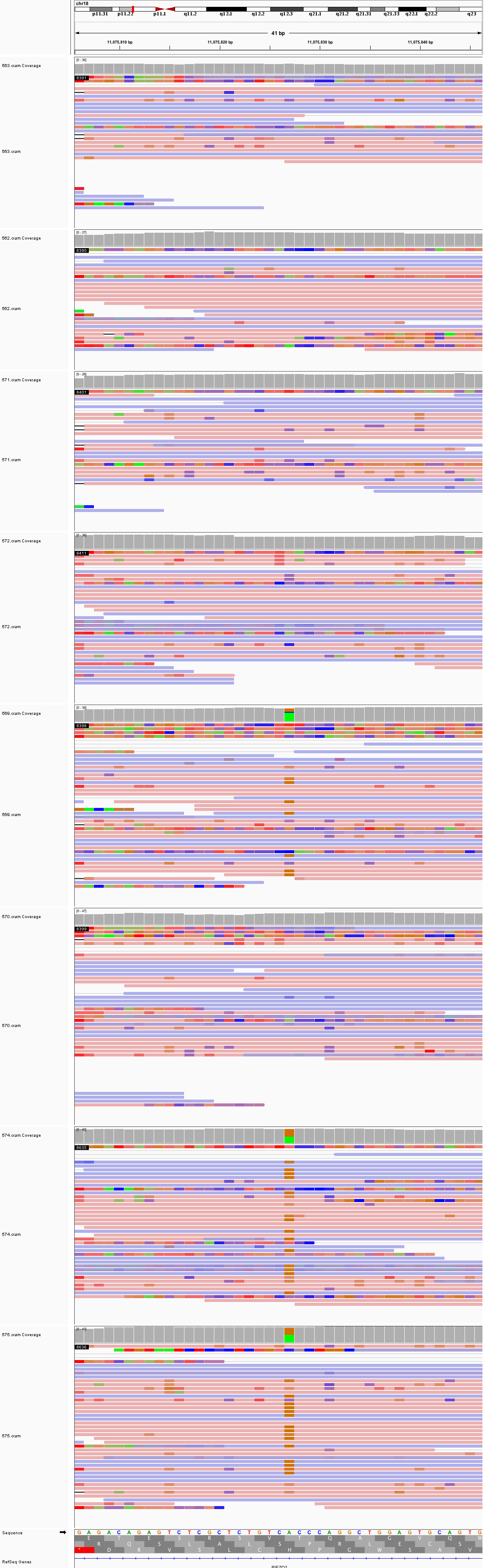

### chr18_38,144,262_38,144,302.png

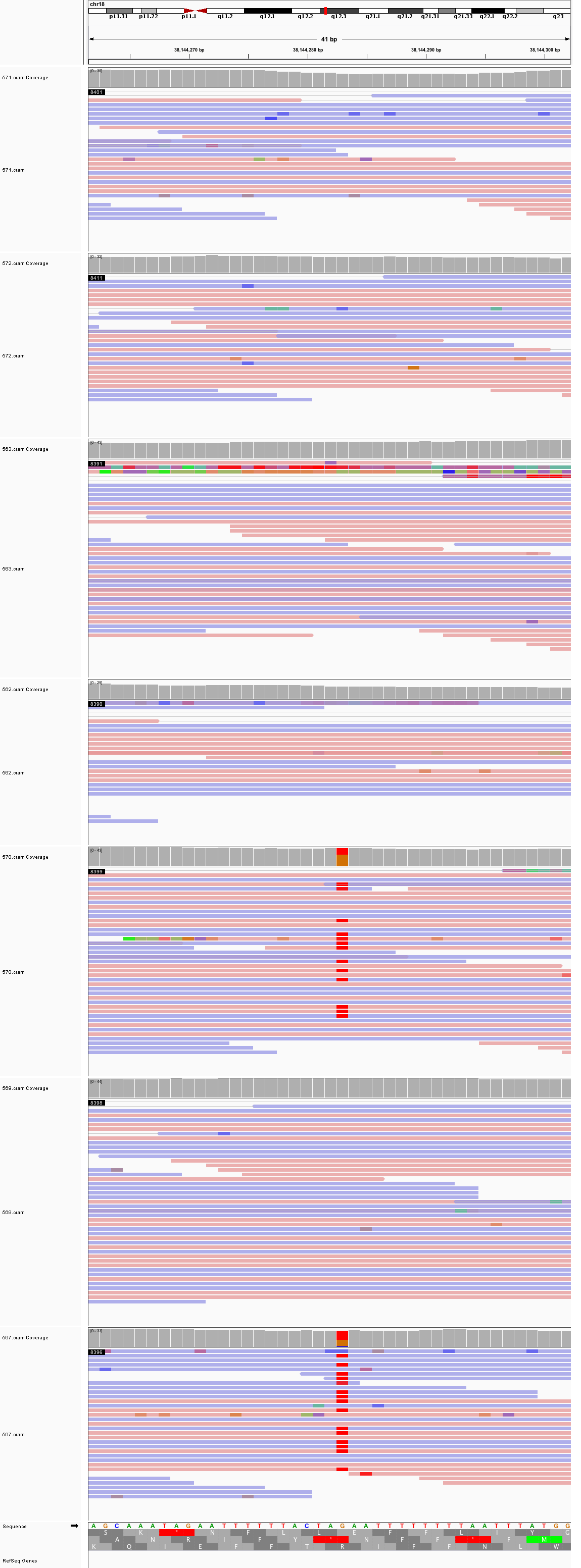

### chr18_49,438,608_49,438,648.png

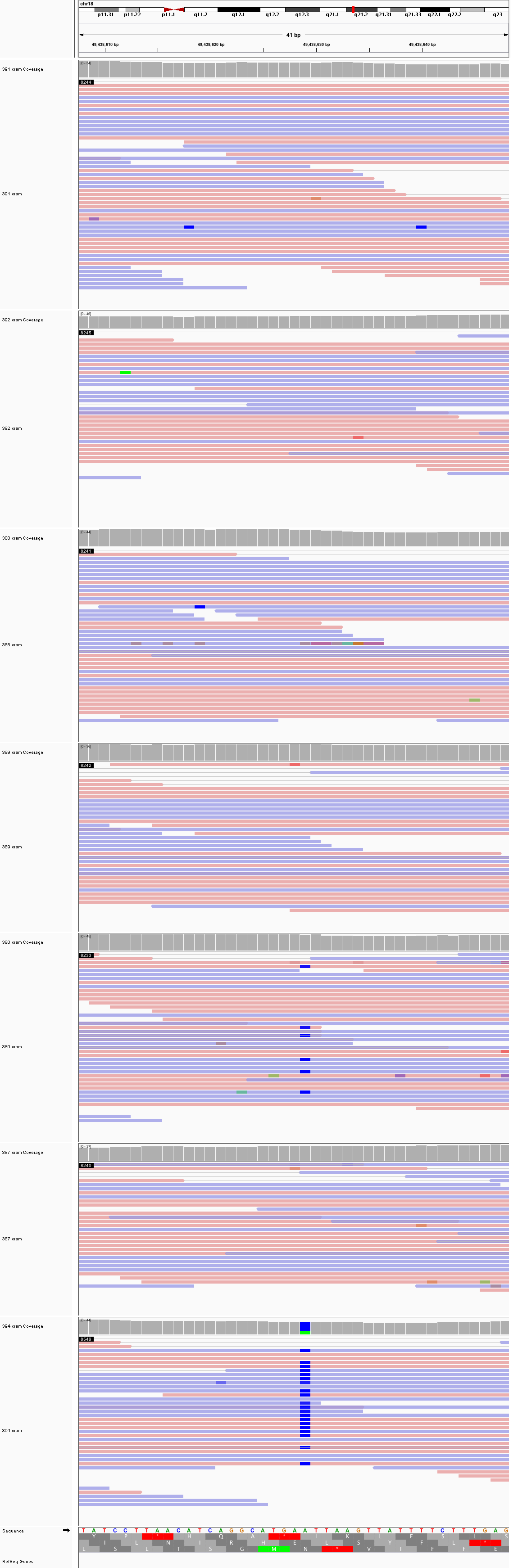

### chr18_77,088,981_77,089,021.png

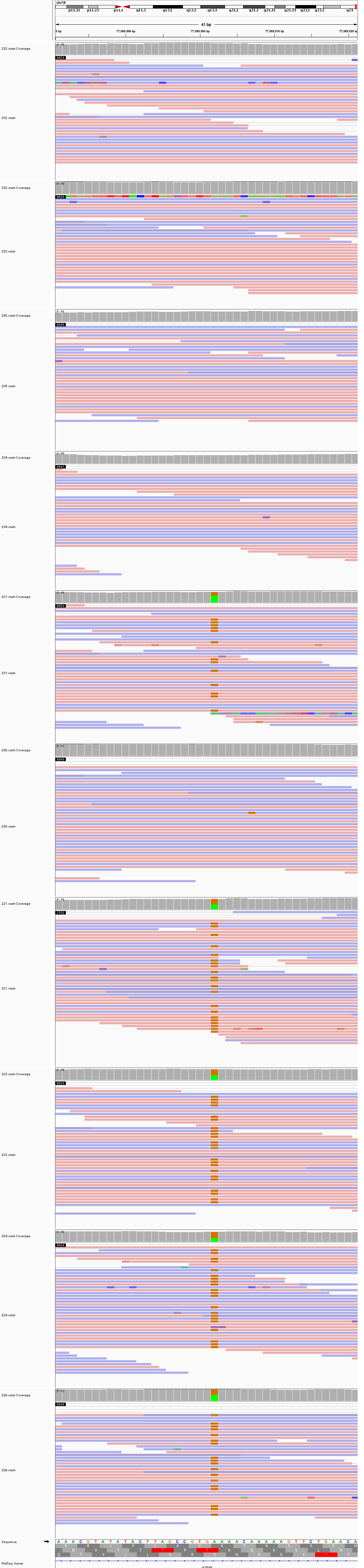

### chr19_9,154,731_9,154,771.png

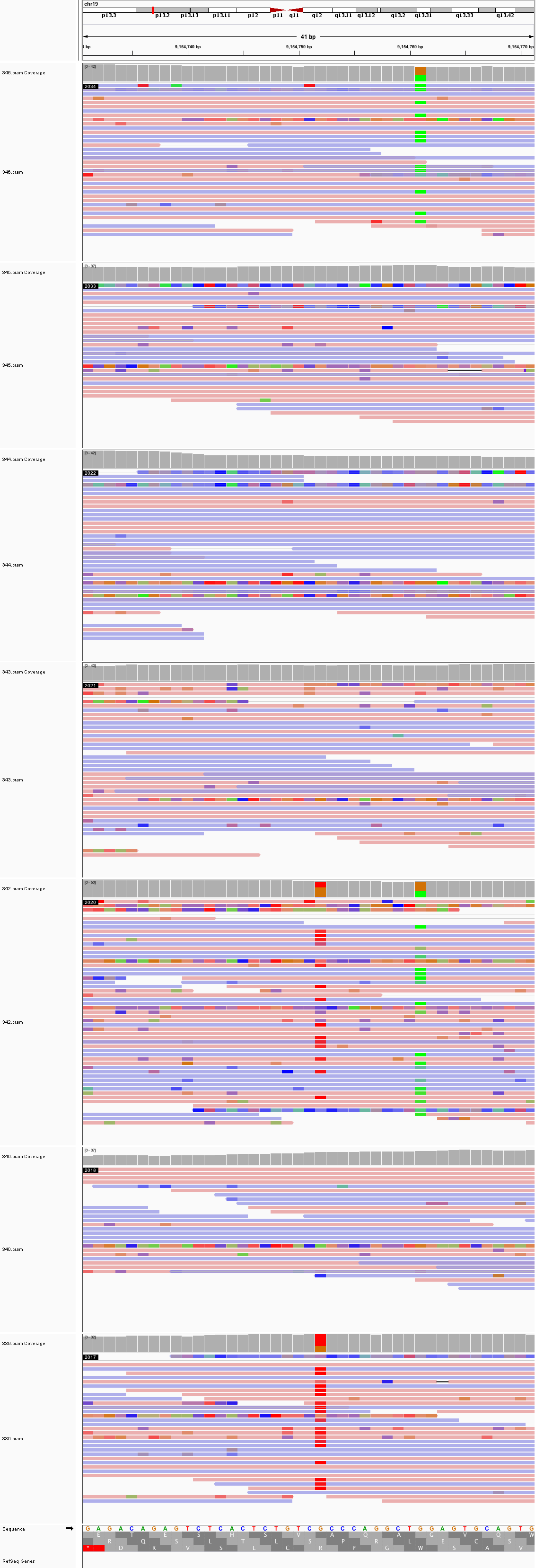

### chr19_40,907,429_40,907,469.png

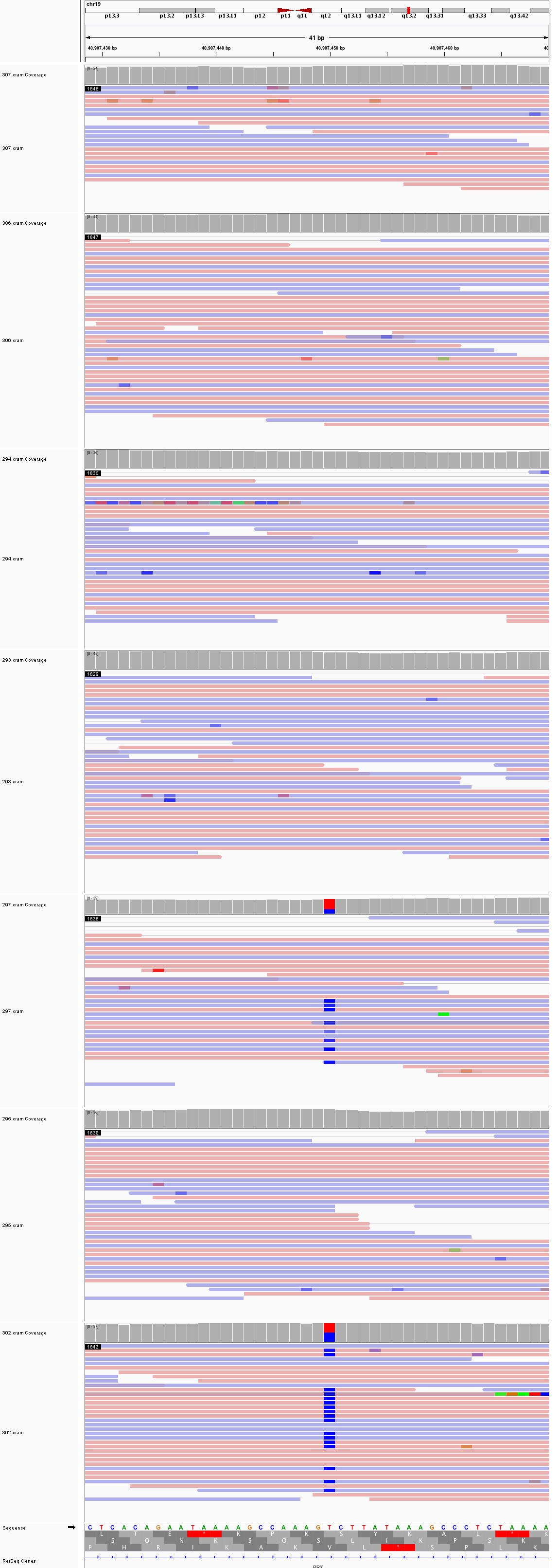

### chr20_31,092,479_31,092,519.png

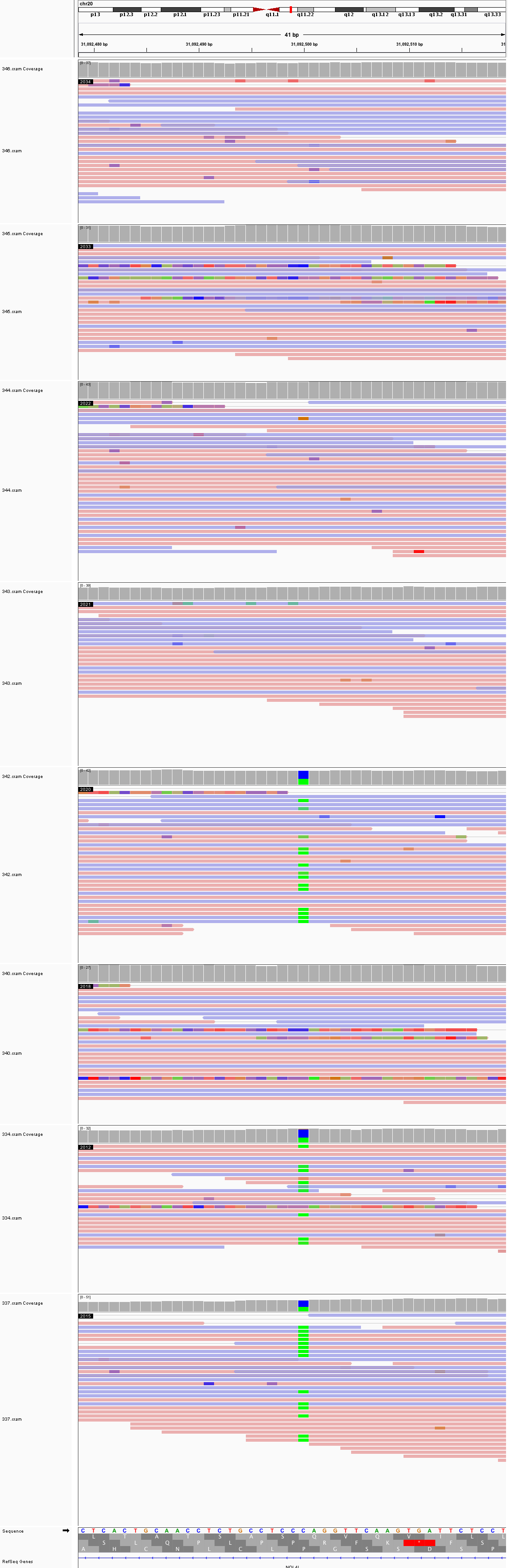

### chr20_55,778,553_55,778,593.png

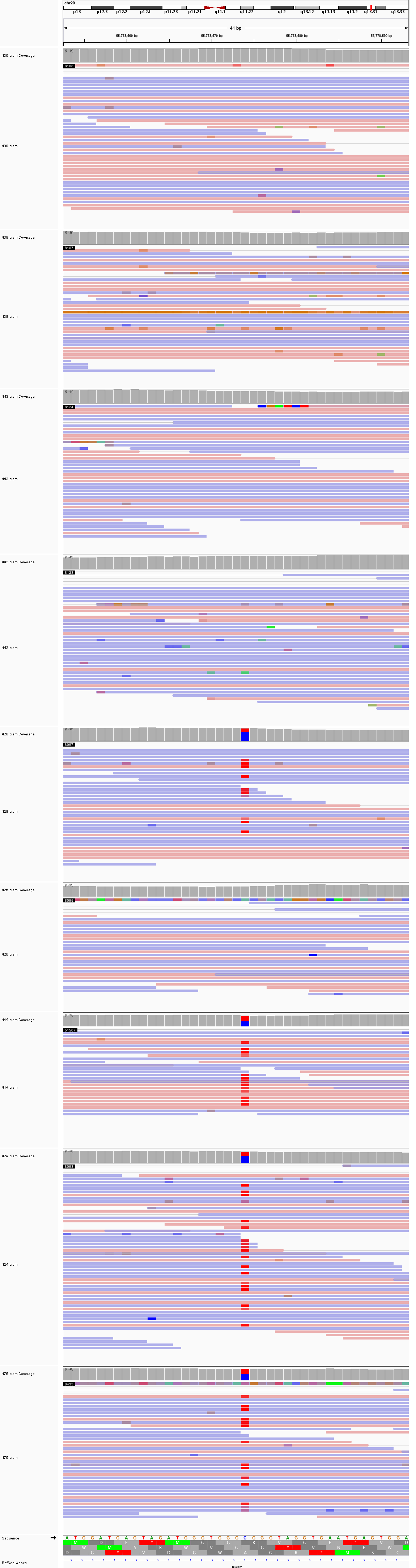

### chr20_757,737_757,777.png

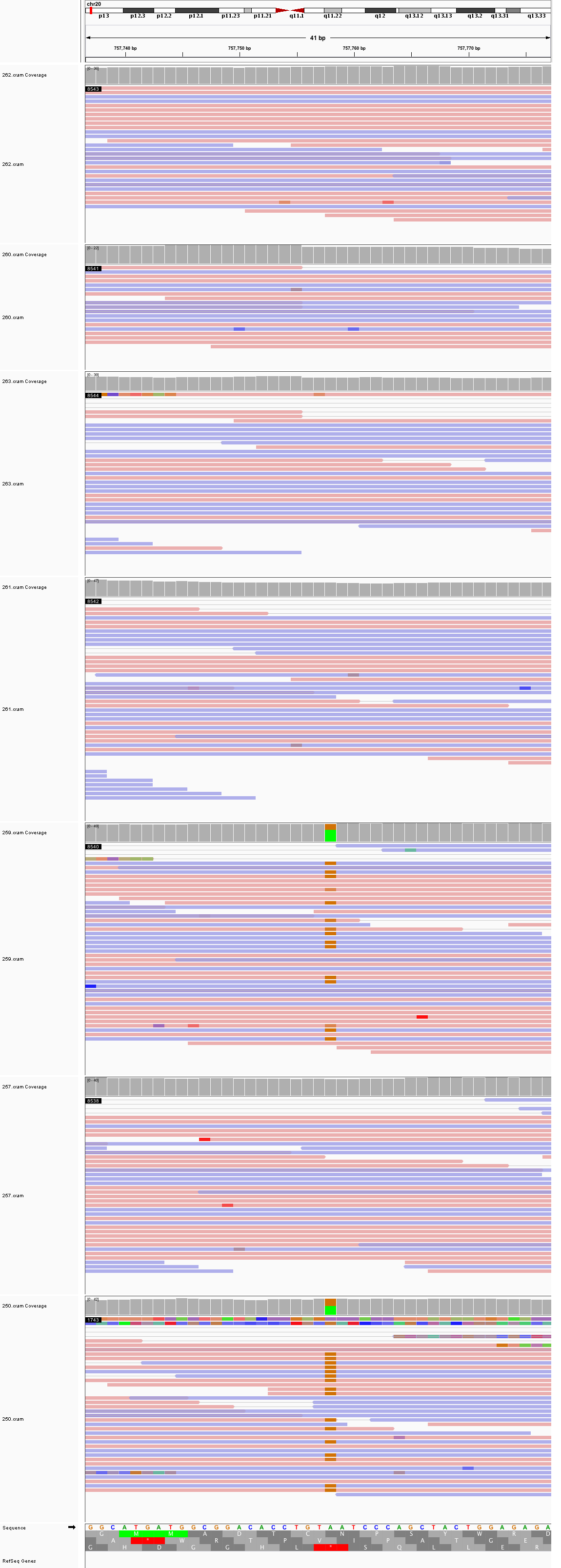
